# Selection for immune evasion in SARS-CoV-2 revealed by high-resolution epitope mapping combined with genome sequence analysis

**DOI:** 10.1101/2022.06.01.494373

**Authors:** Arnaud N’Guessan, Senthilkumar Kailasam, Fatima Mostefai, Raphael Poujol, Jean-Christophe Grenier, Paola Contini, Raffaele De Palma, Carsten Haber, Volker Stadler, Guillaume Bourque, Julie G. Hussin, B. Jesse Shapiro, Jörg H. Fritz, Ciriaco A. Piccirillo

## Abstract

A deeper understanding of the molecular determinants that drive humoral responses to coronaviruses, and in particular severe acute respiratory syndrome coronavirus 2 (SARS-CoV-2), is critical for improving and developing diagnostics, therapies and vaccines. Moreover, viral mutations can change key antigens in a manner that alters the ability of the immune system to detect and clear infections. In this study, we exploit a deep serological profiling strategy coupled with an integrated, computational framework for the analysis of SARS-CoV-2 humoral immune responses of asymptomatic or recovered COVID-19-positive patients relative to COVID-19-negative patients. We made use of a novel high-density peptide array (HDPA) spanning the entire proteomes of SARS-CoV-2 and endemic human coronaviruses to rapidly identify B cell epitopes recognized by distinct antibody isotypes in patients’ blood sera. Using our integrated computational pipeline, we then evaluated the fine immunological properties of detected SARS-CoV-2 epitopes and relate them to their evolutionary and structural properties. While some epitopes are common across all CoVs, others are private to specific hCoVs. We also highlight the existence of hotspots of pre-existing immunity and identify a subset of cross-reactive epitopes that contributes to increasing the overall humoral immune response to SARS-CoV-2. Using a public dataset of over 38,000 viral genomes from the early phase of the pandemic, capturing both inter- and within-host genetic viral diversity, we determined the evolutionary profile of epitopes and the differences across proteins, waves and SARS-CoV-2 variants, which have important implications for genomic surveillance and vaccine design. Lastly, we show that mutations in Spike and Nucleocapsid epitopes are under stronger selection between than within patients, suggesting that most of the selective pressure for immune evasion occurs upon transmission between hosts.

## INTRODUCTION

Coronaviruses constitute a large family of enveloped, positive-sense single-stranded RNA viruses that cause frequent diseases in birds and mammals. The *Coronaviridae* family includes four species that are endemic in the human population (hCoVs): the alpha-coronaviruses which include hCoV-229E and hCoV-NL63 and beta-coronavirus species which include hCoV-HKU1 and hCoV-OC43, and are usually associated with mild, self-limiting upper respiratory tract infections, although they can cause severe illness in immunocompromised patients (*1*). Three other beta-coronavirus species have recently emerged: Middle East respiratory syndrome-CoV (MERS-CoV), SARS-CoV-1, and SARS-CoV-2, all causing severe disease in humans (*2, 3*). Severe acute respiratory syndrome-coronavirus-2 (SARS-CoV-2) is a novel virus belonging to the *Coronaviridae* family that emerged in late 2019 and quickly spread throughout the world, causing a pandemic with morbidity, mortality, and economic disruption on a global scale with few precedents (*3*). The clinical course of COVID-19 is highly variable: some infected individuals are completely asymptomatic (*4*), while others experience a spectrum of clinical manifestations including fever, severe respiratory distress, pneumonia, diarrhea, blood clotting disorders, increased systemic cytokine release and, in <5% of cases, prolonged hospitalization and death (*5*). In addition to factors like viral exposure history, viral inoculum at infection, and the genetic background of the individual, the severity of COVID-19 and the response to treatment is also heavily influenced by other factors like sex, advanced age, ethnicity, and comorbidities such as cardiovascular disease, chronic lung disease, obesity, diabetes, and compromised immune function (*6–8*). An in-depth understanding of the immune response to SARS-CoV-2, particularly humoral, could improve our understanding of the diverse courses of disease and better guide the development of improved diagnostics and vaccines.

SARS-CoV-2 infection can elicit robust antibody responses in humans, and this response represents the primary focus of global efforts to develop accurate serology-based diagnostics and vaccination strategies against infection (*9, 10*). One prevailing view is that an underlying protective immune response directed towards endemic hCoVs is present in infected asymptomatic people consistent with the presence of cross-reactive antibodies between SARS-CoV-2 and hCoV antibodies in pre-pandemic sera from children and adults (*11–15*). Notably, structural proteins of SARS-CoV-2 show relatively high sequence identity (e.g. Spike: 29-35%, N: 28-50%) (*16*) with hCoVs and provide a molecular basis for this cross-reactivity (*16–18*). It is proposed that this response can partially inhibit viral replication and eliminate host infected cells with minimal pathology and inflammatory consequences. Considering the high propensity for SARS-CoV-2 to mutate viral proteins, notably in S protein, variants of concern (VOCs) and variants under investigation (VUIs) can acquire properties for increased transmissibility, disease severity, and/or immune evasion (*19*). Thus, promoting this cross-reactive, pre-existing memory immune response to common hCoVs may be an effective strategy against SARS-CoV-2 and future VOCs.

In order to better understand the molecular determinants underlying protective immunity to pathogens, including viruses, one must define the epitopes in various viral proteins, the minimal unit of an antigen that can be recognized by T and B cells and can elicit potent cellular and humoral immune responses, respectively. A recent study used VirScan technology, a high-throughput, programmable phage-display immunoprecipitation and sequencing (PhIP-Seq) method (*20*), to analyze epitopes of antiviral antibodies in sera of COVID-19 patients relative to pre–COVID-19 sera controls (*21*). However, the nature and dynamics of the peptide pools of VirScan/PHIP-seq may limit the resolution, sensitivity and breadth of specific epitope detection in infected individuals, in turn, providing a fragmented view of the complete footprint of epitope recognition by antibodies (*20, 22*).

In the current study, we provide a comprehensive analysis of SARS-CoV-2 humoral immune responses in a dataset of symptomatic or recovered COVID-19-positive and COVID-19-negative patients. We exploited a novel high-density peptide array (HDPA) by spotting overlapping 15-mer peptides derived from the entire SARS-CoV-2 and hCoVs proteomes to rapidly identify B cell epitopes recognized by distinct antibody isotypes in patients’ blood sera of individual patient groups. We then subjected our data to an integrated computational pipeline to evaluate the fine immunological properties of detected SARS-CoV-2 epitopes and relate them to their evolutionary and structural characteristics in relation to disease onset/susceptibility and clinical features. We show that while some epitopes are common (*public epitopes*) across all studied hCoVs (including SARS-CoV-2), others are unique (*private epitopes*) to a specific hCoV strain. Then, to highlight epitopes that have an important role for protecting against SARS-CoV-2 when an individual gets infected, we defined differential epitopes as epitope for which the response is at least two-times higher in COVID-19-positive than COVID-19-negative individuals. We also highlight hotspots of pre-existing immunity and a subset of cross-reactive epitopes that contributes to increasing the average humoral immune response to SARS-CoV-2. Finally, using a dataset of over 38,000 publicly available genome sequences, collected during the first two waves of the pandemic, we tracked single nucleotide variants (SNVs) within and between COVID-19 patients and found evidence for positive selection on nonsynonymous mutations in epitopes. Selection is stronger between than within patients, indicating that selection for immune evasion occurs mostly upon transmission between hosts. Overall, our results have implications for future genomic surveillance and vaccine design.

## RESULTS

### Antibody fingerprinting with high-density peptide arrays provides a high-resolution antibody epitope map across the SARS-CoV-2 proteome

Most previously reported high-resolution SARS-CoV-2 B cell epitope mapping strategies relied on VirScan/PHIP-seq methodology (*21, 23*). However, the nature and dynamics of the peptide pools of VirScan/PHIP-seq limit the resolution, sensitivity and breadth of specific epitope detection in infected individuals, providing a fragmented view of the complete footprint of epitope recognition by antibodies (*20, 22*). To assess the humoral immune response against SARS-CoV-2 at the epitope level, we used a novel high-density peptide array (HDPA) technology to define virus protein-specific B cell epitopes and potential antigenic hotspots for antibody reactivity. A high-resolution linear epitope map across the entire SARS-CoV-2 proteome was achieved using the PEPperCHIP^®^ SARS-CoV-2 proteome microarray technology (Fig. 1A) (*24*). We performed this assay on sera obtained from ten SARS-CoV-2-positive individuals (asymptomatic and recovered) and five SARS-CoV-2-negative, control subjects (SARS-CoV-2-negative) (Table S1). The degree of immune reactivity to spike protein (S), envelope protein (E), membrane glycoprotein (M), nucleocapsid phosphoprotein (N) and ORF1AB was measured in relative fluorescence units (RFU). Linear overlapping peptides of 15 amino acid length were used for each protein and a dual isotype analysis, determining IgG- and IgA-specific antibody responses, was performed (Fig. 1A). This was followed by a comprehensive analysis workflow to characterize the differential epitopes, their structural properties and utilize genome sequence analysis of arising SARS-CoV-2 variants to assess immune evasion potential.

**Fig. 1.**
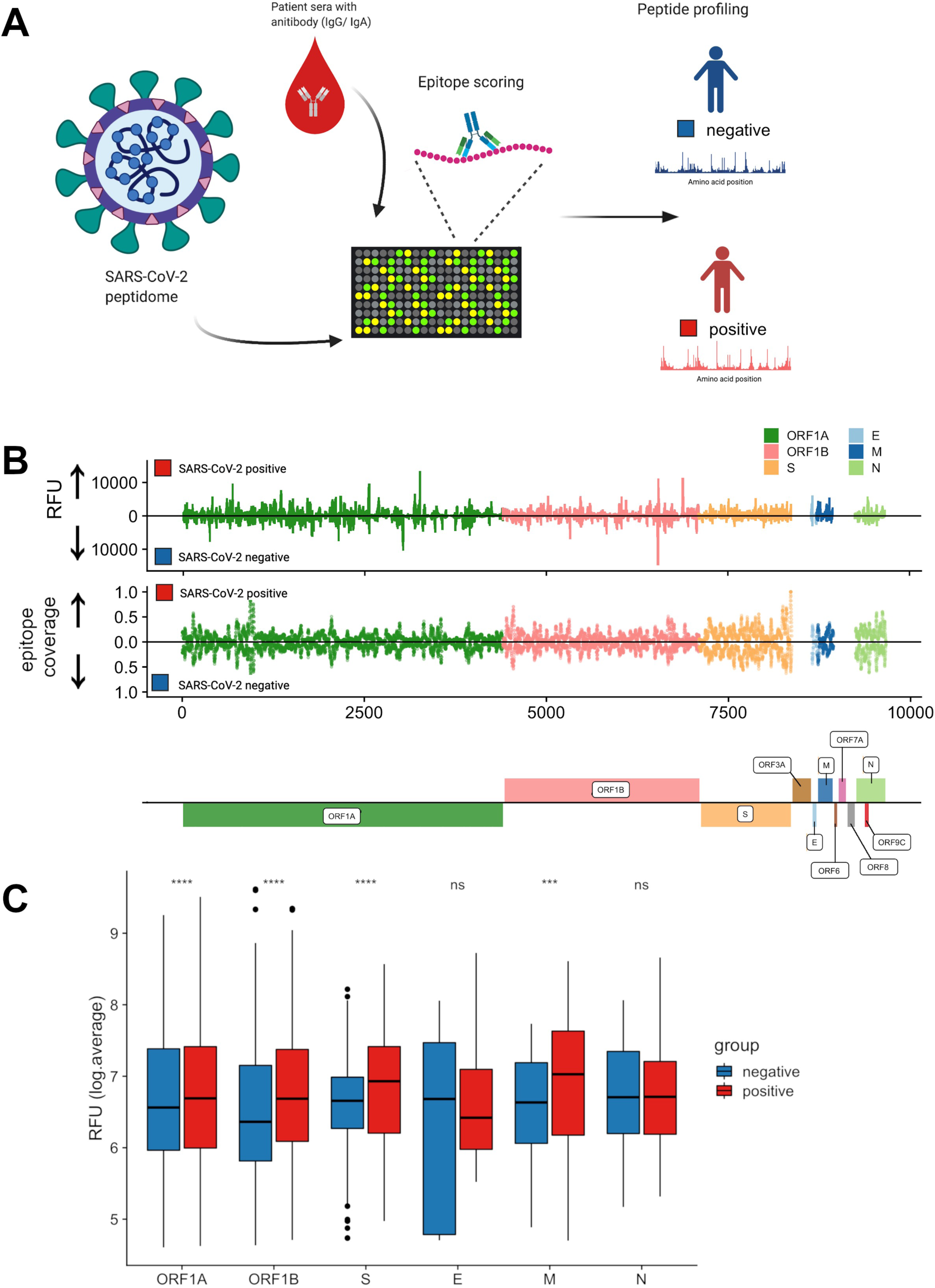
High-density peptide arrays (HDPA) provide high-resolution antibody epitope maps across the SARS-CoV-2 proteome. (A) Overview of analytical pipeline. The proteome of SARS-CoV-2 was translated into 15-mer overlapping peptides with a peptide-to-peptide overlap of 13 amino acids. The resulting individual peptides were printed in duplicates on the microarray. Sera from confirmed SARS-CoV-2-positive and -negative individuals were incubated on PEPperCHIP® the HDPA. Serum antibody binding was visualized using respective fluorescently labeled secondary antibodies (anti-human IgG and anti-human IgA). Image acquisition and data quantification resulted in epitope-specific antibody profiles for SARS-CoV-2. (B) Average relative fluorescent units (RFU) profiles and peptide coverages are plotted across the SARS-CoV-2 proteome (ORF1A, ORF1B, Spike (S) protein, Envelope (E) protein, Membrane (M) glycoprotein, Nucleocapsid (N) phosphoprotein). Antibody responses to each linear 15-mer peptide were mapped across the SARS-CoV-2 proteome and average RFU calculated for each amino acid residue. The normalized positional ‘epitope coverage’ at each protein residue location is defined as the ratio of total peptides mapped to each position by the total expected peptides (see Methods section). ‘Hotspots’ can be seen as spiked in the RFU or coverage distributions. (C) Comparison of mean RFU (log-scale) between SARS-CoV-2-positive and -negative sample groups for each viral protein. (ns: p > 0.05; *: p <= 0.05; **: p <= 0.01; ***: p <= 0.001; ****: p <= 0.0001).

Sera from SARS-CoV-2-positive individuals yielded strong immune reactivity (measured in RFU) in S and N proteins, as well as in select regions of ORF1AB (Fig. 1B). Antibody responses were also identified in samples from SARS-CoV-2-negative individuals (Fig. 1B), suggesting that HDPA technology is well suited to detect epitopes of pre-existing immune responses conferred through prior infections with hCoVs. Although sera from SARS-CoV-2-positive and -negative individuals had very similar epitope coverage per amino acid, some regions in the SARS-CoV-2 proteome were more immuno-dominant than others, exerting higher RFU values (Fig. 1B). Except for M protein, all proteins analyzed had stronger antibody responses to more unique peptides in the SARS-CoV-2-positive patient group (Fig. S1, Table 1, Table S2). In addition, the mean RFU values of SARS-CoV-2-positive sera were higher towards most regions of the SARS-CoV-2 proteome than in the SARS-CoV-2-negative group, demonstrating the elicitation of robust antibody responses to immuno-dominant epitopes upon SARS-CoV-2 infection (Fig. 1C). For further analysis of epitopes with greatest immuno-dominance, only peptide RFU values greater than or equal to 1000 were used in our further analysis. Taken together, our results demonstrate that the applied HDPA approach allows highly sensitive detection of a large pool of epitopes across the SARS-CoV-2 proteome.

**Table 1.** Numbers of SARS-CoV-2-specific epitope-defining peptides identified by HDPA. Numbers of SARS-CoV-2-specific epitope-defining peptides identified with high density peptide arrays (HDPA) in spike (S) protein, envelope (E) protein, membrane (M) glycoprotein, nucleocapsid (N) phosphoprotein, and ORF1ab. Numbers of unique peptides that showed a significant antibody response (RFU ≥ 1000) in SARS-CoV-2-negative and SARS-CoV-2-positive groups are depicted. Some peptides are present in both groups, referred to as overlap.

|  | **Spike** | **Nucleocapsid** | **Envelope** | **Membrane** | **ORF1ab** | **TOTAL** |
| --- | --- | --- | --- | --- | --- | --- |
| **SARS-CoV-2 negative** | 119 | 41 | 1 | 47 | 294 | 502 |
| **SARS-CoV-2 positive** | 195 | 69 | 6 | 29 | 549 | 848 |
| **Overlap** | 90 | 35 | 6 | 17 | 353 | 501 |
| **Total** | 404 | 145 | 13 | 93 | 1196 | 1851 |

### Structural features of identified epitopes and comparison with computationally predicted epitopes

An epitope is the minimal unit of an antigen that can be recognized by T and B cells and can elicit potent cellular and humoral immune responses, respectively. B cell epitopes can be divided into two major categories, namely linear and conformational epitopes. In a linear epitope, a stretch of continuous amino acids forms the antibody binding site, while amino acid residues that are brought together by protein folding form conformational epitopes. In our current study, antibody responses are detected using linear peptide arrays, and thus these epitopes are primarily linear in nature, although some linear epitopes contributing to conformational components of a protein may also be detected. Though there is a significant interest for short linear epitopes in vaccine design, most of the current SARS-CoV-2 vaccine immunogens are structural S proteins (*25*). Thus, we explored of whether the short peptide-based approach in the applied HDPA approach is able to reveal conformational epitope sites as well, as has been recently suggested (*21*). Using 3D structures and biophysical properties of the SARS-CoV-2 proteome, we applied the DiscoTope algorithm (*26*) to computationally predict conformational B cell epitopes as well as the Bepipred algorithm (*27*) to obtain linear B cell epitopes. We then compared these to the epitopes sites identified in our HDPA experiment. Apart from E and M proteins, we observed significant overlap of experimentally identified epitopes with predicted epitopes. (Fig. 2A), with approximately 38% of the total proteome being part of the amino acid residues contributing towards the B cell epitome of SARS-CoV-2 recognized in infected individuals. We observed overlap of mapped epitope sites with conformational epitopes predicted by Discotope, suggesting that the applied HDPA approach also identifies a considerable number of conformational epitopes.

It is important for epitopes to be solvent exposed to allow their amino acid side chains to interact with the antibody. In our study, there was no significant difference in the average normalized solvent accessibility (SASA) of epitopes compared to non-epitope regions (mean SASA = 5.3 Å^2^). However, residues in highly conserved epitope sites have lower solvent accessibility (Fig. S2A-C), suggesting that many identified epitopes might only get exposed through proteolysis or conformational changes throughout the infectious cycle or stage of anti-viral immune response. This implies that our linear peptide-based HDPA approach is able to capture more epitopes compared to those that use full-length antigens or protein domains to study immune profiling of antibody responses (*28, 29*). Using structural models we mapped the epitopes of S protein of SARS-CoV-2 identified by the HDPA approach, which revealed that many of the epitope sites identified in the S protein are located in the N terminal domain (NTD) and the receptor binding domain (RBD) (Fig. 2B, C). Interestingly, most epitope sites identified in the NTD and RBD have low conservation scores, while many other epitope sites identified in the S protein had high conservation scores (Fig. 3A, B). In addition, HDPA analysis revealed strong antibody immunoreactivity in a few epitope sites of E (Fig. 3C), M (Fig. 3D), N proteins (Fig. 3E), as well as ORF1A (Fig. S3) and ORF1B (Fig. S4). Taken together, our results demonstrate that the applied HDPA profiling strategy can identify a novel set of linear and conformational B cell epitopes unique in sera of SARS-CoV-2-infected individuals.

**Fig. 2.**
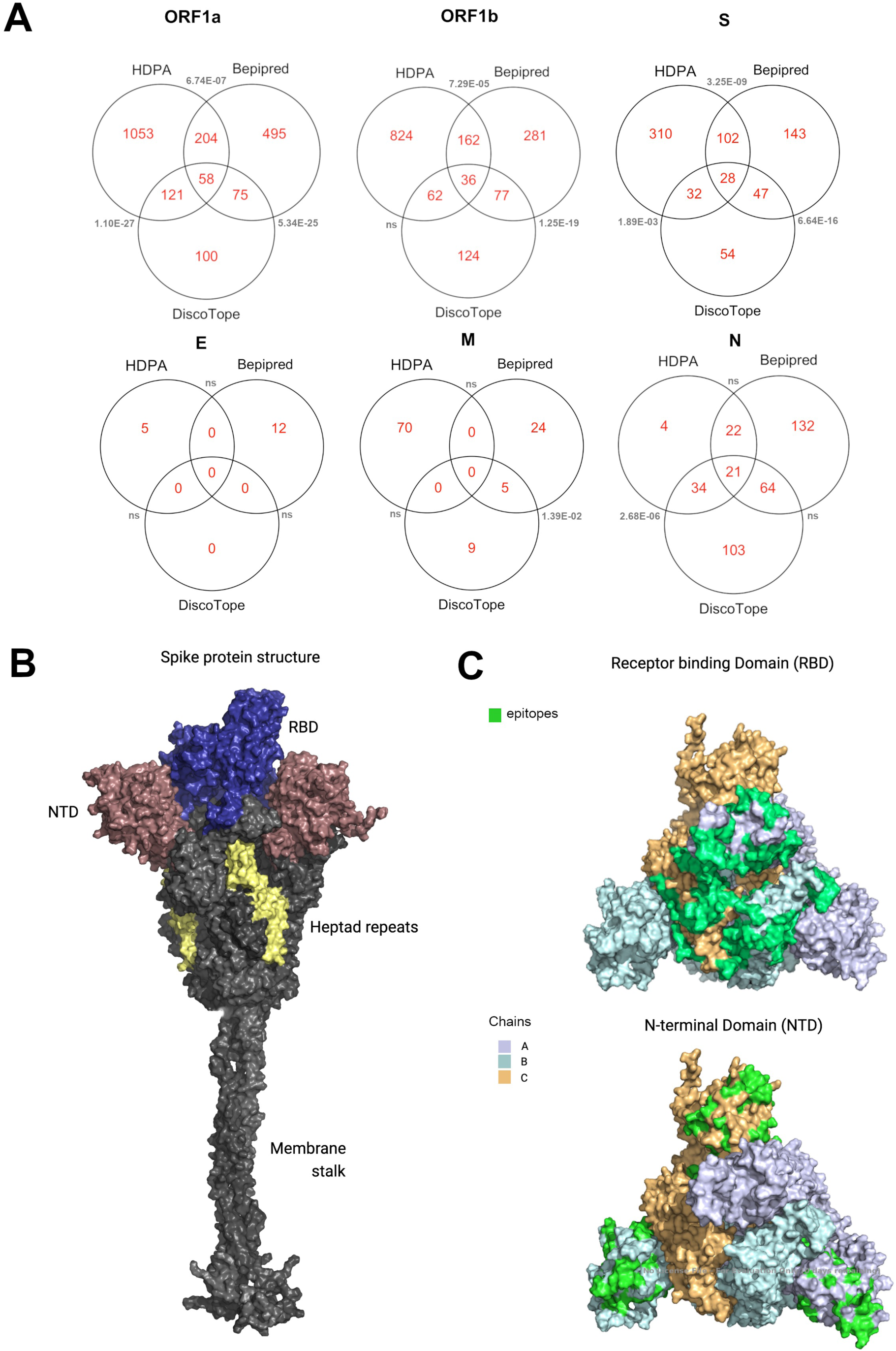
Structural features of identified epitopes and comparison with computationally predicted epitopes. (A) Venn diagrams comparing epitope sites identified by HDPA with computational approaches including Bepipred and Discotope. Overlap test for significance between epitope sites from HDPA and computational prediction methods; p-value (< 0.10) shown in grey. (B) Three-dimensional structural model of the full-length spike protein trimer in an open conformation with domains labelled; receptor binding domain (RBD); N-terminal domain (NTD). (C) Three-dimensional model of RBD and NTD highlighting epitope sites (green) identified by HDPA analysis on the surface of the RBD and NTD domains.

**Figure 3.**
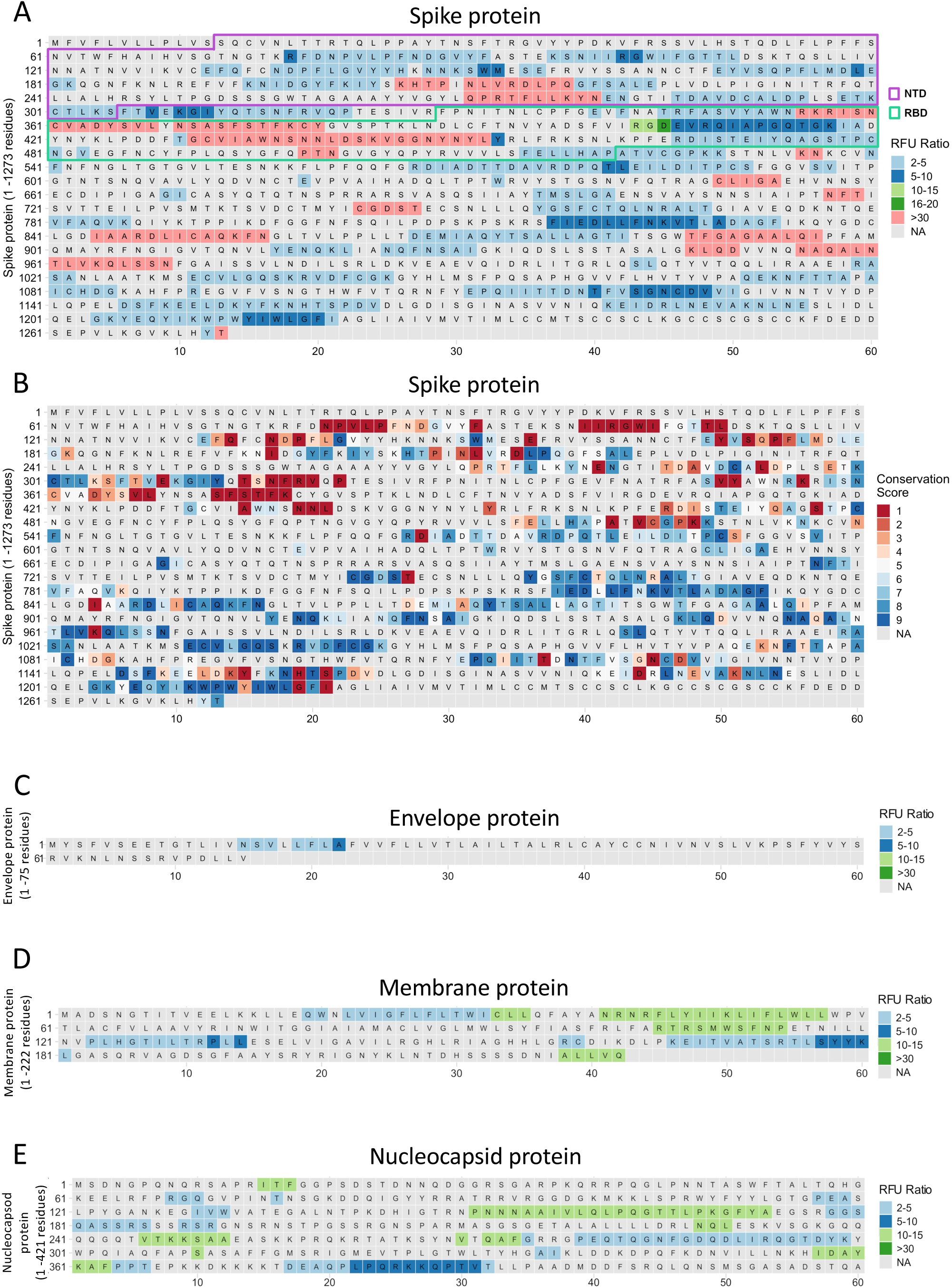
Identified differential epitope sites in structural proteins of SARS-CoV-2. Relative Fluorescence Unit (RFU) values of HDPA analysis were used to calculate ratio values to define differential epitope sites and are color coded on the distinct SARS-CoV-2 proteins. Residues that are not part of epitopes are marked in grey. (A) RFU values of differential epitope sites identified in SARS-CoV-2 Spike protein. N-terminal domain (NTD) and receptor-binding domain (RBD) are highlighted. (B) Conservation scores of physico-chemical properties of the SARS-CoV-2 Spike protein. (C) RFU values of differential epitope sites identified in SARS-CoV-2 Envelope protein. (D) RFU values of differential epitope sites identified in SARS-CoV-2 Membrane protein. (E) RFU values of differential epitope sites identified in SARS-CoV-2 Nucleocapsid protein.

### Cross-reactivity to endemic seasonal human coronaviruses is a significant driver of antibody responses to SARS-CoV-2 epitopes

In addition to the zoonotic pathogens SARS-CoV, MERS-CoV, and SARS-CoV-2, four other low-pathogenicity hCoVs are endemic and co-circulating in the human population (*30*): strains OC43 and HKU1 (beta-CoVs like SARS-CoV-2), and NL63 & 229E (alpha-CoVs), of which OC43 and 229E are the most common, accounting for 5-30% of common colds (*31*). Notably, structural proteins of SARS-CoV-2 show high amino acid sequence identity with hCoVs (*17, 18*). One prevailing view in our understanding of COVID-19 immunopathogenesis is that an underlying immune response towards endemic hCoVs is a hallmark feature of SARS-CoV-2-infected asymptomatic individuals. This pre-existing immunity is hypothesized to partially control viral replication and eliminate infected cells resulting in less severe pathology and inflammation (*11–15*).

Having established the link between structure accessibility and protein conservation (Fig. 3B; Fig. S2), we next asked how conservation is related to the humoral immune response. One way protein conservation could affect adaptive immunity is through cross-reactivity to related pathogens. To this end, we first analyzed the humoral immune response against the four hCoVs OC43, HKU1, NL63 and 229E at the epitope level using HDPA on sera from the same ten SARS-CoV-2-positive (asymptomatic or recovered) patients and five control subjects (SARS-CoV-2-negative; Table S1). HDPA yielded strong antibody reactivities to many distinct sites across the proteomes of all hCoVs (Table 2, Table S3-S6). We then defined cross-reactive epitopes based on the conservation of peptide residues across hCoVs and the presence of an immune response to epitopes in SARS-CoV-2 and at least of one of the endemic hCoVs (HKU1, NL63, OC43 or 229E; Fig. 4A). To evaluate the conservation of peptide sequences across hCoVs, we aligned the protein sequences of these viruses and calculated a conservation score, reflecting the conservation of physico-chemical properties in the alignment where identical residues score the highest (*32*). We also defined cross-reactivity at the level of epitope sites (single amino acids) to account for the possibility that a particular amino acid site within a 15-mers epitopes are associated with cross-reactive immunity. Epitope sites with a conservation score ≥ 6 and for which we obtained antibody reactivity for both SARS-CoV-2 and at least one of the hCoVs were considered as cross-reactive epitope sites. These cross-reactive epitope sites represent 27.2% of the pool of detected epitope sites by the applied HDPA assay (Table S7). We also carried out local alignment of the peptides from the HDPA (with RFU > = 1000) of all five viral strains to the SARS-CoV-2 proteome to evaluate the cross-reactivity profile of SARS-CoV-2 epitopes and identified hotspots of conserved epitopes (example for S protein in Fig. 4B).

**Fig. 4.**
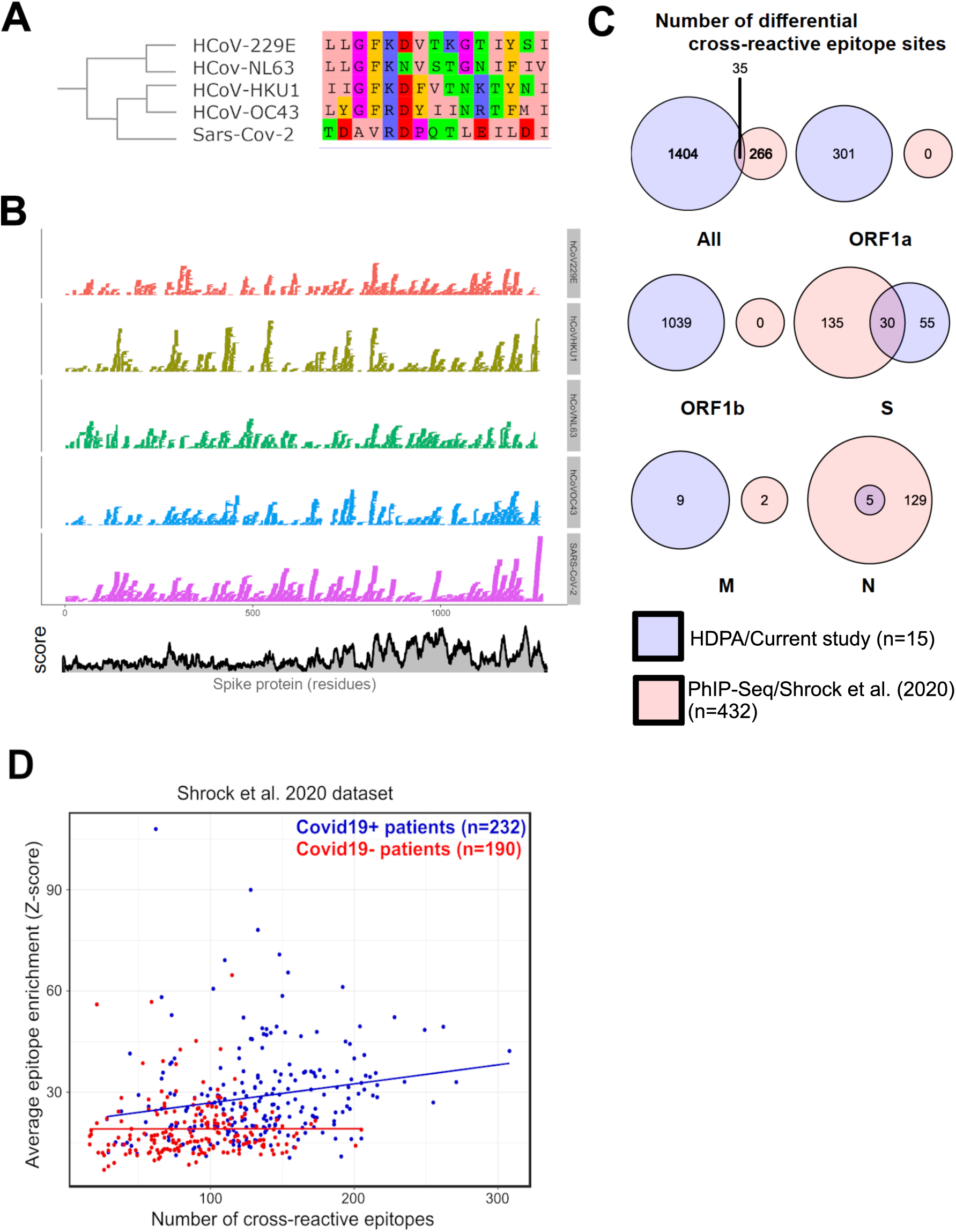
Cross-reactivity to endemic seasonal human coronaviruses is a significant driver of antibody responses to SARS-CoV-2 epitopes. (A) Schematic multiple sequence alignment of proteome sequences between hCoVs and SARS-CoV-2. We defined cross-reactive epitope sites based on peptide sequence conservation between human coronavirus strains (SARS-CoV-2, HKU1, OC43 and NL63) and the presence of an antibody response to the corresponding peptide in SARS-CoV-2 and at least one of the endemic human coronaviruses (229E, HKU1, OC43 and NL63). (B) Mapping of hCoVs epitope-defining peptides within the Spike (S) protein. The colors represent the 15-mer peptides to which an antibody response to the human coronavirus strains (SARS-CoV-2, 229E, HKU1, OC43 and NL63) has been detected. Conservation score (Cscore) was calculated based on this alignment. (C) Numbers of differential cross-reactive epitope sites across distinct viral proteins (ORF1A, ORF1B, Spike (S) protein, Envelope (E) protein, Membrane (M) glycoprotein, Nucleocapsid (N) phosphoprotein) detected in the current study using HDPA (blue) compared with a recently published PhiP-Seq study (red, n = 432; (*21*). (D) The average immune response to SARS-CoV-2 positively correlates with the number of cross-reactive epitopes. A linear regression between the average epitope Z-score per patient and the number of cross-reactive epitopes for both SARS-CoV-2-positive (blue; adjusted R^2^= 0.033; slope = 0.059; *p* = 0.003) and both SARS-CoV-2 negative (red; not significant; *P* > 0.5) patients was performed.

**Table 2.**
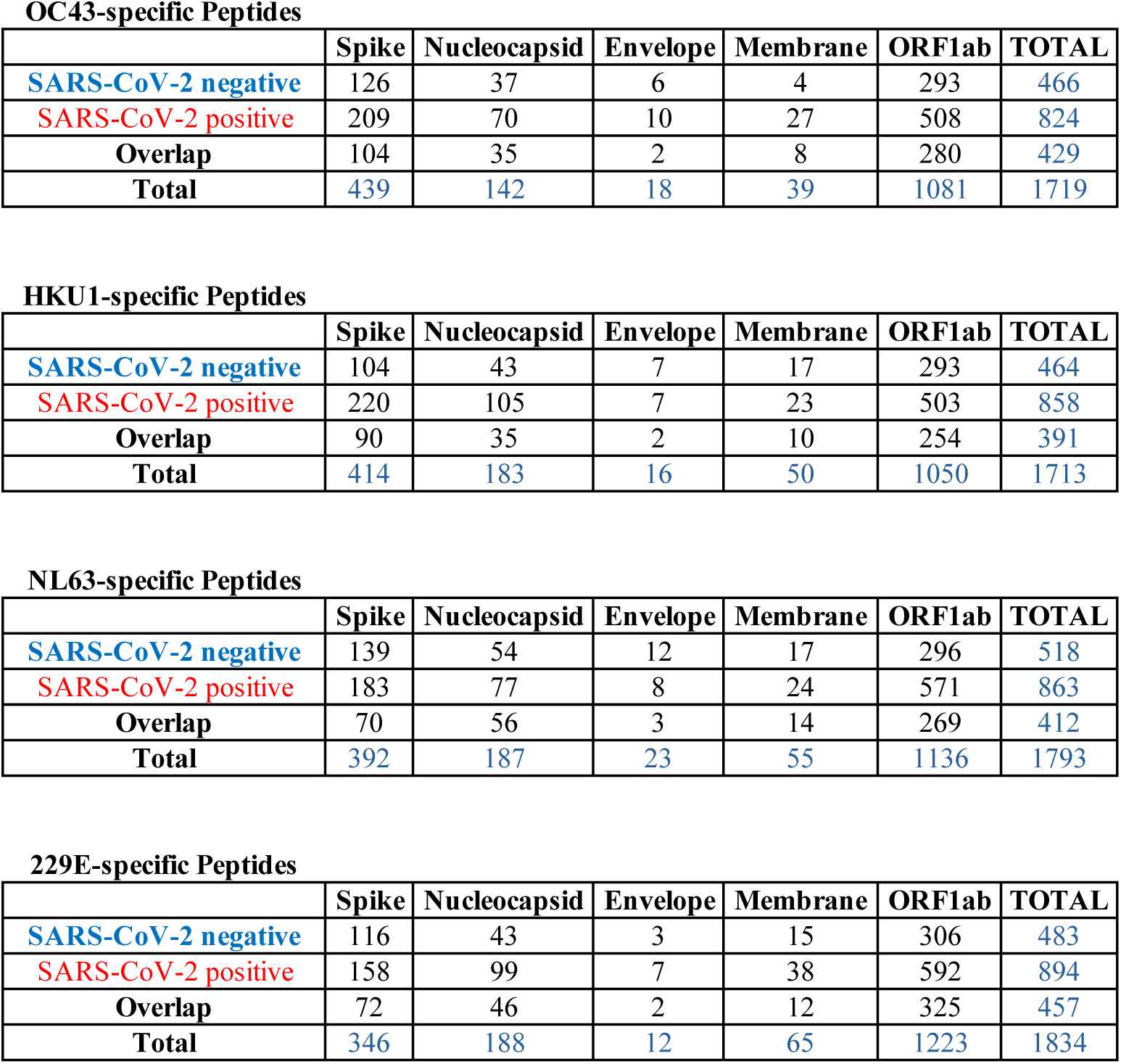
Numbers of seasonal endemic hCoV-specific epitope-defining peptides identified by HDPA. Numbers of seasonal endemic hCoV-specific (229E, HKU1, OC43 and NL63) epitope-defining peptides identified with high density peptide arrays (HDPA) in spike (S) protein, envelope (E) protein, membrane (M) glycoprotein, nucleocapsid (N) phosphoprotein, and ORF1ab. Numbers of unique peptides that showed a significant antibody response (RFU ≥ 1000) in SARS-CoV-2-negative and SARS-CoV-2-positive groups are depicted. Some peptides are present in both groups, referred to as overlap.

OC43-specific Peptides
|  | Spike | Nucleocapsid | Envelope | Membrane | ORF1ab | TOTAL |
| --- | --- | --- | --- | --- | --- | --- |
| **SARS-CoV-2 negative** | 126 | 37 | 6 | 4 | 293 | 466 |
| **SARS-CoV-2 positive** | 209 | 70 | 10 | 27 | 508 | 824 |
| **Overlap** | 104 | 35 | 2 | 8 | 280 | 429 |
| **Total** | 439 | 142 | 18 | 39 | 1081 | 1719 |

HKU1-specific Peptides
|  | Spike | Nucleocapsid | Envelope | Membrane | ORF1ab | TOTAL |
| --- | --- | --- | --- | --- | --- | --- |
| **SARS-CoV-2 negative** | 104 | 43 | 7 | 17 | 293 | 464 |
| **SARS-CoV-2 positive** | 220 | 105 | 7 | 23 | 503 | 858 |
| **Overlap** | 90 | 35 | 2 | 10 | 254 | 391 |
| **Total** | 414 | 183 | 16 | 50 | 1050 | 1713 |

NL63-specific Peptides
|  | Spike | Nucleocapsid | Envelope | Membrane | ORF1ab | TOTAL |
| --- | --- | --- | --- | --- | --- | --- |
| **SARS-CoV-2 negative** | 139 | 54 | 12 | 17 | 296 | 518 |
| **SARS-CoV-2 positive** | 183 | 77 | 8 | 24 | 571 | 863 |
| **Overlap** | 70 | 56 | 3 | 14 | 269 | 412 |
| **Total** | 392 | 187 | 23 | 55 | 1136 | 1793 |

Table 2
|  | Spike | Nucleocapsid | Envelope | Membrane | ORF1ab | TOTAL |
| --- | --- | --- | --- | --- | --- | --- |
| **SARS-CoV-2 negative** | 116 | 43 | 3 | 15 | 306 | 483 |
| **SARS-CoV-2 positive** | 158 | 99 | 7 | 38 | 592 | 894 |
| **Overlap** | 72 | 46 | 2 | 12 | 325 | 457 |
| **Total** | 346 | 188 | 12 | 65 | 1223 | 1834 |

Next, to be able to highlight cross-reactive epitope sites that are particularly important for the humoral immune response after exposure to SARS-CoV-2, we focused on differential cross-reactive epitope sites that give an antibody response signal in sera of SARS-CoV-2-positive over SARS-CoV-2-negative individuals. Although we analyzed sera from a smaller sized cohort compared to a recent study using PhIP-Seq (*21*), we were nonetheless able to sensitively detect more differential cross-reactive epitope sites discriminating virus exposed from non-exposed individuals (Fig. 4C). The increased sensitivity of HDPA over PhIP-Seq analysis for identifying cross-reactive epitope sites was further highlighted when performing a sensitivity analysis on the number of cross-reactive epitope sites that define a cross-reactive epitope (Fig. S5A).

We next analyzed if the humoral immune response to SARS-CoV-2 epitopes correlated with the number of cross-reactive epitopes identified. In other words, to what extent is the response to SARS-CoV-2 predictable based on cross-reactivity to other endemic hCoVs? We defined cross-reactive epitopes as peptide sequences with at least five cross-reactive epitope sites. We found a positive correlation between the average humoral immune response to SARS-CoV-2 epitopes and the number of cross-reactive epitopes per patient in a recently published PhIP-Seq dataset (*21*) (Fig. 4D) and in our HDPA dataset (Fig S5). We detected a stronger positive correlation between the average antibody response to SARS-CoV-2 epitopes and the number of cross-reactive epitopes in SARS-CoV-2-positive compared to negative patients (Fig. 4D; correlation coefficients: 4.42e-3 vs 1.54e-3; ANOVA *p* for *Covid*19*status* = 7.43e-10). This positive correlation is robust to the threshold number of cross-reactive epitope sites defining a cross-reactive epitope and is also replicated in our HDPA dataset (Fig. S5).

Leveraging the sample size in this same PhIP-Seq dataset, we aimed to identify the subset of cross-reactive epitopes with the greatest contribution in humoral immunity in SARS-CoV-2-positive patients using the IndVal test, which is a non-parametric test to identify significant associations from presence-absence data (*33*). This test allowed us to identify 75 epitopes that are significantly associated with SARS-CoV-2-positive over SARS-CoV-2-negative samples, with 16 out of these 75 epitopes (21.3%) being cross-reactive (Table S8.). These results again highlight the contribution of a subset of hCoV-cross-reactive epitopes for the humoral immune response to SARS-CoV-2.

### Point mutations and natural selection in epitopes occur at higher rates upon transmission than within patients

There is mounting evidence that mutations in SARS-CoV-2 enhance viral fitness, replication rate and transmissibility, and/or partially evade adaptive immunity that has been induced by prior infection or vaccination (*19*). Thus, it is essential to shed light on the interplay between SARS-CoV-2 mutations and the acquired immune response in infection. To this end, we tracked the evolution of SARS-CoV-2 B cell epitopes using single nucleotide variants (SNVs) identified in 38,685 SARS-CoV-2 genome sequences from the NCBI sequence read archive (Table S9). We selected SARS-CoV-2 samples from the first pandemic wave (defined as January 1 to July 31 2020) and the second wave (defined as August 1 to December 31 2020) sequenced using Illumina paired-end amplicons with a minimum average depth of coverage of 200x and fewer than 10,000 sites with a depth of coverage lower than 100x. Combined with additional filters to remove sequencing errors (see Methods for details), such deep coverage allowed us to identify SNVs that are polymorphic within patients, reflecting within-patient evolution (*34, 35*), as well as those that are shared between the consensus sequences of different patients. We refer to these within-patient SNVs as ’mutations’ and to between-patient SNVs (those present at >75% frequency within a sample and observed in three or more samples) as ’substitutions’ that have likely been transmitted across multiple patients. Our definitions of mutations and substitutions are not mutually exclusive: an SNV can be a mutation in one sample and a substitution in another. We counted the absolute number of substitutions relative to the Wuhan-1 reference genome, so the count does not reflect the unique number of substitution events along a phylogeny, but rather the prevalence of the substitutions in the database. As such, substitution counts are weighted to reflect their ’success’ in transmitting widely.

Using this dataset of mutations and substitutions, we first asked whether cross-reactive (public) epitopes evolved differently than epitopes private to SARS-CoV-2. We found that cross-reactive epitopes tend to evolve more slowly than SARS-CoV-2 private epitopes, accumulating fewer substitutions and having lower ratios of nonsynonymous to synonymous substitutions (Fig. S6A-D). The same trend of slower evolution in cross-reactive epitopes is also observed at the level of within-patient mutations, but the effect is much stronger at the level of substitutions between patients (Fig. S6A-D). This is consistent with the fact that these epitopes are conserved across multiple distinct hCoV strains and could be evolving under strong and longstanding functional constraints. However, purifying selection has less time to purge deleterious mutations within hosts, and is therefore more detectable over longer time scales spanning multiple transmission events.

Mutations in epitopes have the potential to evade or lessen the effectiveness of adaptive immunity conferred by infection or vaccination. A recent topic of debate has been the extent to which natural selection for immune evasion acts on SARS-CoV-2 during infection, or upon transmission (*19*). During influenza virus infection, most of the selective pressure for immune evasion occurs upon transmission, not within a patient (*36*). This is because viral loads often peak before the priming of adaptive immune responses. As such, peak viral transmission occurs before there is time for selection to act within a patient, and for immune evasion to occur. A similar ‘asynchrony’ transmission model has been proposed for SARS-CoV-2 (*37*), although data supporting such model has been lacking. To test the asynchrony model in SARS-CoV-2 we tracked SNVs within as well as between patients, within and outside epitope sites, and across the first two pandemic waves (Fig. 5, Fig. S7). Throughout both waves, we found consistently lower within-host mutation rates in epitopes sites when compared to non-epitope sites across most SARS-CoV-2 proteins (Fig. 5A, Fig. S7A). In contrast, the structural proteins S, M, and N had significantly higher rates of between-host substitution in epitope sites compared to non-epitope sites (Fig. 5B, Fig. S7B), suggesting stronger positive selection for epitope changes upon transmission than within hosts. To further assess the evidence for selection on epitopes, we used the ratio of nonsynonymous to synonymous SNVs both between patients (dN/dS) and within patients (pN/dS) calculated separately within and outside epitopes. Higher ratios indicate positive or relaxed purifying selection, whereas lower ratios indicate stronger purifying selection. We found that the structural proteins S and N have consistently higher nonsynonymous SNV rates in epitope sites, both within and between patients, and across both pandemic waves (Fig. 5C, D, Fig. S7C, D). While this result is consistent with positive selection of altered epitopes (immune evasion) occurring both within and between patients, dN/dS ratios (between patients) are consistently higher than pN/pS ratios (within patients). These observations indicate that nonsynonymous substitutions in S and N epitope sites accumulate most rapidly upon transmission, rather than within patients. Taken together these results support the notion that most of the selective pressure for immune evasion of SARS-CoV-2 occurs upon transmission between hosts, consistent with the asynchrony model (*36*).

**Fig. 5.**
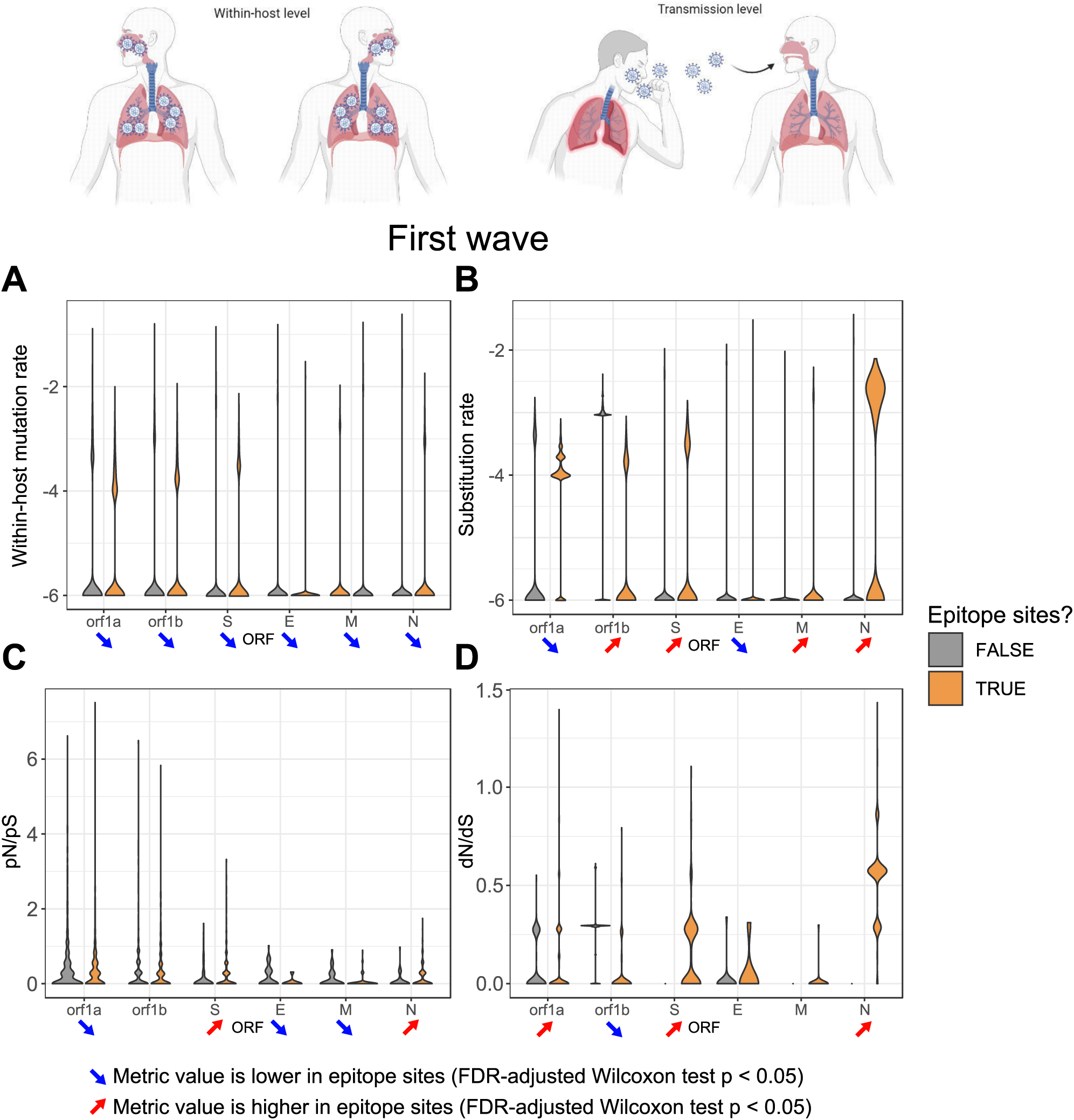
Evolutionary profiles of SARS-CoV-2 epitopes during the first pandemic wave. Mutations (within hosts) and substitutions (between hosts) in epitope sites (orange) vs non-epitope sites (grey) during the first pandemic SARS-CoV-2 wave (defined as January 1 to July 31, 2020) is depicted. For each metric, significantly lower values in epitope sites of a certain gene are represented by a blue arrow pointing down while significantly higher values in epitope sites of a certain gene are represented by a red arrow pointing up (FDR-adjusted Wilcoxon test p<0.05). (A) Distributions of sample mutation rates across proteins. (B) Distributions of sample substitution rates across proteins. (C) Distributions of pN/pS across proteins. (D) Distributions of dN/dS across proteins. Mutation and substitution rates are normalized by gene length and plotted on a log10 scale.

### Assessing the immune evasion potential of SARS-CoV-2 variants

The observation of similar pattern of mutations and selective pressures in epitopes across pandemic waves 1 and 2 (Fig. 5, Fig. S7) was surprising, given the expectation that increasing levels of immunity in the population would lead to increased selection for immune evasion over time. The second wave is characterized by the rise of variants of concern (VOCs) and variants under investigation (VUIs) with higher transmissibility and, in some cases, increased disease severity and acquired immune evasion phenotypes (*19*). The rise of VOCs has been suggested to be due to a shift in the SARS-CoV-2 fitness landscape, although the nature of such a shift is unclear (*38*). If part of this shift were due to rising population immunity from the first to second wave, one would expect increasing selection for immune evasion variants, resulting in higher frequencies of SNVs in epitopes in wave 2. Although we found a higher total number of nonsynonymous SNVs (including both mutations and substitutions) in epitope sites unique to wave 1 than unique to wave 2 (Fig. 6A), wave 2-specific SNVs reached higher frequencies across samples compared to wave 1-specific SNVs, consistent with increased selection for immune evasion over time (Fig. 6B). However, mutations common to both waves achieved the highest frequencies, indicating their early appearance and persistence over time (Fig. 6B).

**Fig. 6.**
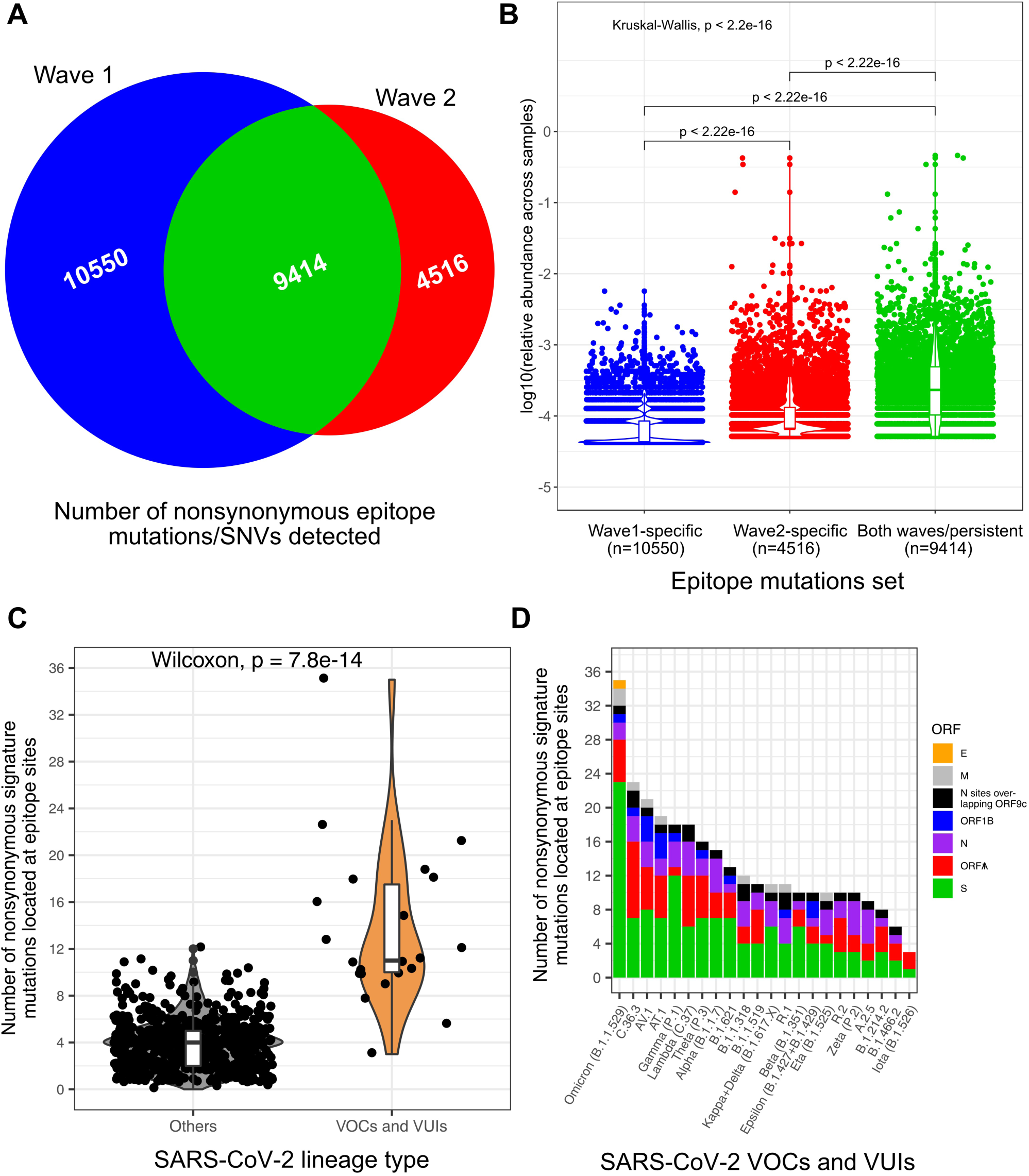
Assessing the immune evasion potential of SARS-CoV-2 variants. (A) Venn diagram showing the numbers of non-synonymous epitope mutations specific to SARS-CoV-2 pandemic wave 1 (blue; defined as January 1 to July 31, 2020), specific to wave 2 (red; defined as August 1 to December 31 2020), and shared between both waves (green). (B) Distribution of the relative abundance of non-synonymous mutations in epitopes across pandemic waves. For better visualization, we plotted the Y-axis on a log10 scale and represented the distributions with a jitter plot, a violin plot and a boxplot. The Wilcoxon test p-values are indicated above each pair of distributions. The Kruskal-Wallis test p-values are indicated at the top left to indicate the significance of the differences across all distributions. (C) Distribution of the numbers of signature mutations located at epitope sites across SARS-CoV-2 groups (grey for non-VOCs and non-VUIs, orange for VOCs and VUIs). The Wilcoxon test p-value is indicated at the top of the panel to show the significance of the differences between the two groups. (D) Distribution of the numbers of nonsynonymous signature mutations in epitopes of selected VOCs and VUIs. For each VOC or VUI we indicate the number of signature mutations in epitopes identified with HDPA across all analyzed ORFs: envelope (E) protein (orange), membrane (M) glycoprotein (grey), N sites overlapping ORF9c (black), ORF1b (blue), nucleocapsid (N) phosphoprotein (purple), ORF1A (red), spike (S) protein (green).

A likely driver of VOC evolution is selection for increased transmissibility. For example, the Delta VOC is estimated to be 76-117% more transmissible than non-VOCs and non-VUIs, while Gamma is 29-48% more transmissible and Alpha is 24-33% more transmissible than the original Wuhan SARS-CoV-2 strain (*39*). However, selection for immune evasion could also play a significant role for increased spread or VOCs and VUIs. To test this hypothesis, we defined signature mutations of each variant (see Methods; Table S10) as substitutions that are present in >=90% of sequences assigned to that lineage. We calculated the prevalence of substitutions in thousands of publicly available consensus sequences collected from NCBI during 2020 and added data from CoV-Spectrum about under-represented lineage in the database or lineages that emerged during 2021 (*40*). We focused on nonsynonymous signature mutations located in epitope sites and found that VOCs and VUIs contain significantly more signature mutations in epitopes compared to non-VOCs and non-VUIs (Fig. 6C) suggesting that evasion of the humoral immune response could be a significant driver of VOC/VUI evolution. We then ranked VOCs and VUIs based on their number of signature mutations in epitopes (Fig. 6D). We observed that Delta has an intermediate number of mutations in mapped epitopes (Fig. 6D). Many nonsynonymous epitope mutations were also found in the C.36.3 linage, which is thought to be highly transmissible (*41*). However, the most nonsynonymous epitope mutations were observed in Omicron (B.1.1.529, Fig. 6D), which is highly transmissible and immune evasive (*42*). Most epitope mutations in VOC/VUI occur in the S protein (Fig. 6D). However, normalizing by gene length revealed a relatively high density of epitope mutations in the N protein, especially at sites of the N protein that overlap with ORF9c (Fig. S8), a membrane-anchored protein of SARS-CoV-2 that can hinder interferon signaling, viral protein degradation and other stress response pathways when expressed in human lung epithelial cell lines (*43*). Finally, having established these general evolutionary patterns of mutation and selection on epitopes, we attempted to pinpoint specific epitope mutations that could hinder the immune response. For each epitope site, we extracted both the measured patient immune responses and the prevalence of nonsynonymous (missense) mutations from the NCBI dataset (Table S11). Among the most prevalent mutations identified are two mutations occurring at consecutive sites in the N protein (N:R203K and N:G204R) that overlap with ORF9c (encoded within the N gene). Taken together our observations show that high resolution epitope mapping combined with genome sequence analysis provides a powerful strategy to rapidly assess the immune evasion potential of emerging SARS-CoV-2 variants.

## DISCUSSION

An in-depth map of the breadth of the antigenic determinants of the immune response following infection with SARS-CoV-2 is key for a better understanding of the diagnostic markers, the identification of the correlates of protection and the monitoring of vaccine efficacy. We therefore set out to define the antigenic hotspots and epitope signatures of SARS-CoV-2-specific humoral immune responses in patients with COVID-19 and uninfected healthy controls using high-density peptide microarrays (HDPA) covering the entire proteomes of SARS-CoV-2 as well as of the four seasonal hCoVs (OC43, NL63, HKU1 and 229E). Our results demonstrate that the HDPA approach provides a sensitive, high-throughput antibody profiling strategy to identify linear and conformational B cell epitopes. Using structural models, we found that many of the epitope sites identified in the S protein are located in the NTD and RBD region of the S protein. Interestingly, most epitope sites identified in the NTD and RBD are poorly conserved across coronaviruses, while epitope elsewhere in the S protein were more highly conserved. In addition, HDPA analysis revealed strong and specific antibody immunoreactivity in select epitope sites of structural SARS-CoV-2 proteins (E, M, N proteins), as well as ORF1AB.

Antibody cross-reactivity with similar viral antigens affects the accuracy of serological tests, but also has the potential to elicit beneficial immunological memory responses that could affect the course of SARS-CoV-2 infections. Our results highlight a significant cross-reactivity between SARS-CoV-2 and hCoV B cell epitope sites in many viral proteins, demonstrating that HDPA allows to uncover a novel dimension of cross-reactive immunity relative to PhIP-Seq. The fact that more differential epitope sites in the S and N proteins were detected by a recent study using PhIP-Seq (*21*) probably reflects the lower sample size of our dataset. However, HDPA detected more cross-reactive epitope sites than PhIP-Seq with fewer patient samples analyzed, reflecting one of the benefits of our applied methodology. Sensitivity limitations of PhIP-seq to broadly detected polio epitopes have been previously reported (*20, 22*) and might contribute to the observed differences, similarly affecting detectability of CoV antigens. Such limitations are not observed in our HDPA approach (*44*) which typically yielded strong polio responsiveness in over 90% of sampled individuals (*45*).

Importantly, the cross-reactivity in identified B cell epitope sites positively relates to previous infections with seasonal common cold hCoVs. This suggests that immune memory conferred by previous seasonal hCoV infections positively influences SARS-CoV-2-specific antibody responses and may explain the large portion of SARS-CoV-2-infected individuals with mild and asymptomatic disease symptoms (*4*). Notably, there was little to no correlation between cross-reactivity and immune response in COVID-19 negative patients, suggesting that resistance to infection is not easily explained by cross reactivity. However, this molecular cross-reactivity can pose important complications in serological tests, particularly when studying asymptomatic patients. Cross-reactivity in immunodominant epitopes can be molecular determinants of strong immunity in individuals and therefore may serve as the basis for future pan-coronavirus vaccine design strategies. In turn, mutations in these cross-reactive epitopes can potentially breach pre-existing immune protection conferred by previous viral exposures, contributing to viral evolution, immune selection and immune evasion.

By combining our epitope dataset with publicly available SARS-CoV-2 genome sequences, we were able to study mutations that occur in epitopes and compare their rates of evolution and selective pressures to non-epitope sites. Ideally, we would have matched epitopes and viral mutations arising from the same patients to more directly infer selection for immune evasion. Although such matched data is currently rare, we were still able to make inferences about evolution within epitopes on a population-wide scale. First, we found that mutations in SARS-CoV-2 epitopes are under evolutionary constraints. SARS-CoV-2-specific epitopes that are cross-reactive with other endemic seasonal hCoVs tend to accumulate fewer substitutions and are under purifying selection against nonsynonymous changes. Second, epitopes in structural proteins S, M, and N accumulate more substitutions and are under stronger positive selection for nonsynonymous changes than non-epitopes. Natural selection favouring changes in epitope sites was therefore detectable during the first two pandemic waves. As population immunity accumulates over time, we would expect increasing selection for immune evasion. Consistent with this expectation, we observed that mutations in epitopes increased in frequency from the first to the second pandemic wave, and we expect this trend to continue.

Notably, we found much slower rates of evolution and weaker evidence for positive selection on epitopes within patients, indicating that most selection for immune evasion occurs upon transmission rather than within patients. This is consistent with asynchrony between peak viral loads (when selection is most efficient) and the adaptive immune response, as is the case for influenza (*36, 46*). Notable exceptions are chronic infections, in which significant adaptive evolution occurs within patients, likely including antibody evasion (*47, 48*). However, such infections likely represent a small minority of the sequences included in our dataset. While they may be important – particularly if a chronic infection is transmitted – they do not represent the vast majority of COVID-19 cases. Another non-exclusive explanation for the higher between-host mutation rate in epitope sites is the small transmission bottleneck (*49, 50*).

Consistent with a general trend of immune evasion, we observed that VOCs and VUIs contain significantly more signature mutations in epitopes than non-VOCs and non-VUIs demonstrating that evasion of the humoral immune response is a significant driver of VOC/VUI evolution. The most mutations in epitopes were found in the VOCs Delta, C.36.3, and especially Omicron (B.1.1.529). Most of these epitope mutations in VOCs are localized to the S protein, highlighting that polymorphism in the S protein critically impacts antigenicity in highly transmissible variants.

Much research rightly focuses on the S protein, but we also find mutation in the N protein epitopes that could be selected for immune evasion. The N protein had the highest dN/dS values during both pandemic waves analyzed, suggesting the presence of a subset of epitope substitutions under positive selection. After normalizing by gene length, we found the highest density of epitope mutations in the N protein, especially in regions of overlap with ORF9c. Orf9c is one of the four conserved overlapping genes (OLGs) of SARS-CoV-2 (*51*), wherein a single stretch of nucleotides encodes two distinct proteins in different reading frames. OLGs are ‘genes within genes’ that compress genomic information, thereby allowing genetic innovation via *overprinting* (*52, 53*). However, a single mutation in an OLG may alter two proteins at the same time, constraining evolution of the pre-existing open reading frame (ORF). Although, OLGs are known entities that contribute to the emergence and pathogenicity of new viruses (*54*), unfortunately, genome annotation methods typically miss OLGs, instead favoring one ORF per genomic region (*54*). Similarly, they remain inconsistently reported in viruses of the SARS *coronavirus* species (*55*). Importantly, annotations of *ORF9b* and *ORF9c* are conflicting or absent in the SARS-CoV-2 reference genome Wuhan-Hu-1 (NCBI: NC_045512.2) and genomic studies (*56, 57*). In addition, OLGs are often not displayed in genome browsers (*58*) and therefore such inconsistencies complicate research to decipher their role in infection and immunity.

The small protein encoded by the ORF9c OLG has recently been shown to constitute a membrane-associated protein to suppresses antiviral interferon and antigen-presentation responses and modify innate immune responses (*43, 59, 60*). Here, we found that N protein epitopes in the region overlap with ORF9c constitute an antigenic target of the humoral immune response and accumulate a high density of mutations in VOCs. It remains to be investigated if and to what extent ORF9c-specific immune responses contribute to host protection and if mutations could also affect these responses. Other OLG-derived proteins, including Orf3d, ORF8 and Orf9b, have been shown to elicit strong antibody responses in sera from COVID-19 patients (*61–63*), although their contribution to host protection remains unknown. Concerns have arisen that S-specific vaccine immunity conferred solely to S protein may fail to neutralize emerging variants of SARS-CoV-2 and contribute to selection of immune escape variants (*64–66*). Vaccination studies in rodent models using N protein as antigenic target have recently shown the establishment of protective immunity (*67*). Hence, expansion of viral antigenic targets in SARS-CoV-2 vaccines, including OLG proteins, to broaden epitope coverage and immune effector mechanisms should be a goal in the development of new COVID-19 vaccines.

## MATERIALS AND METHODS

### Serum Samples and Study Population

Recruitment of patients at the San Martino University Hospital (Genoa, Italy) was approved by the Institutional Review Board at Genoa University, approved by the Ethics Committee of Liguria Region (Comitato Etico Regione Liguria; N. CER Liguria 114/2020–ID 10420) and carried out in accordance with the principles of the Declaration of Helsinki. Positivity of SARS-CoV-2 infection was assessed both by PCR and measurement of specific antibodies (Cobas-Roche using Elecsys Anti-SARS-CoV-2 S). All patients gave their consent for participation in this study. Negative and asymptomatic individuals were health workers who were tested regularly in the hospital and classified according to serological and molecular tests for COVID-19 (Table S1). Recovered individuals (convalescent post-infection) were all patients previously admitted at the hospital due to lung pneumonia and were found to be positive to COVID-19, having severe (n = 2) and mild (n = 3) disease. Sera were collected according to standard procedures, by centrifugation.

### High-density peptide array (HDPA)

To analyze the antibody responses to SARS-CoV-2 at the epitope level we used a recently developed high-density peptide array (HDPA), the PEPperCHIP® Microarray (PEPperPRINT GmbH, Germany), covering the proteome of the SARS-CoV-2 isolate Wuhan-Hu-1 as well as the four seasonal hCoVs OC43, HKU1, NL63 and 229E (see Table S12 for accession numbers used). The protein sequences of ORF1A/B, Spike (S) protein, Envelope (E) protein, Membrane (M) glycoprotein, Nucleocapsid (N) phosphoprotein were translated into 15 amino acid peptides with a peptide overlap of 13 amino acids. This results in 27,540 individual peptides, which were printed in duplicates (55,080 spots). In addition, to ensure sensitivity controls of the PEPperCHIP® HDPA, positive controls were included to probe for antibody reactivity for influenza hemagglutinin (HA; YPYDVPDYAG, 360 spots) and polio virus (KEVPALTAVETGAT, 355 spots). These additional HA and polio peptides framing the microarrays were simultaneously stained as internal quality control to confirm assay performance and peptide microarray integrity. With this setup per chip, 15 samples (see Table S1) were analyzed.

At first, the peptide microarrays were incubated for 15 minutes in phosphate buffered saline supplemented with 0.05% Tween 20 (PBS-T, pH 7.4) and blocked for 30 minutes with Rockland Blocking Buffer (RL) (Rockland Immunochemicals) at room temperature. Prior to immunoassay, sera of patients were first heat-inactivated at 56°C for 30 minutes, and then the microarrays were incubated at serum dilutions of 1:500, 1:100 and 1:20 in 10% RL/PBS-T overnight at 4°C with orbital shaking. Microarrays were then washed (three times with PBS-T for 1 minute) and peptide binding was detected with isotype-specific secondary goat anti-human IgG (Fc) DyLight680 (ThermoFisher Scientific) and goat anti-human IgA (alpha chain) DyLight800 (Rockland Immunochemicals) antibodies at a final concentration of 0.1 μg/ml and 1 μg/ml, respectively (in 10% RL/PBS-T for 45 minutes at room temperature). Subsequent washing (three times with PBS-T for 1 minute) was followed by dipping the microarrays in 1mM TRIS pH 7.4 followed by drying with pressurized air. Acquisition of images was done using a LI-COR Odyssey CLx Infrared Imaging System (scanning offset 0.65 mm, resolution 21 μm). Data quantification and analysis was based on the assays at dilution 1:20. Using ImageJ software the resulting 32-bit gray-scale TIFF files were converted into 16-bit gray-scale TIFF files and then further analyzed using the PepSlide® Analyzer (SICASYS Software GmbH). The in house developed PEPperPRINT software algorithm was used to calculate median foreground intensities (background-corrected intensities) of each spot and spot-to-spot deviations of spot duplicates. A maximum spot-to-spot deviation of 40% was tolerated, otherwise, the corresponding intensity values were zeroed. To complement this analysis, acquired microarray scans were reassessed with respect to artifacts by visual inspection, and erroneous values were corrected manually. Based on averaged median foreground intensities, intensity maps were generated and interactions in the peptide maps highlighted by an intensity color code with red (IgG) or green (IgA) for high and white for low spot intensities. To identify the top IgG and IgA antibody responses of the human serum samples, the averaged intensity values were sorted by decreasing spot intensities. We further plotted averaged spot intensities of the assay against the peptide microarray content from the N-terminus of Spike (SARS-CoV-2) to the C-terminus of ORF1AB (HUK1) to visualize overall spot intensities and signal-to-noise ratios. The intensity plot was correlated with the peptide and intensity map as well as with visual inspection of the microarray scans to identify the main antibody responses of the human sera. In general, relative fluorescent units (RFU) of equal to or higher than 100 was considered a positive antibody response. However, as mentioned in the results sections, several sets of analysis were performed with and RFU cut-off of 1000 or higher.

### Defining cross-reactivity using protein conservation and immune response to endemic human coronaviruses

To find epitope sites associated with cross-reactivity, we first calculated the conservation of peptide sequences across endemic hCoVs (HKU1, NL63, OC43 or 229E). To do so, we aligned the reference sequences of the hCoVs (proteome) in Jalview and extracted the conservation score (CScore) (*68*). This conservation score reflects the conservation of physico-chemical properties in the alignment, where identical residues score the highest (*32*). Epitope sites with a conservation score ≥ 6 and for which we detected antibody responses for both SARS-CoV-2 and at least one of the endemic hCoVs were considered as cross-reactive epitope sites.

### Detecting epitopes that are significantly more prevalent in SARS-CoV-2 positive patients

First, antibody responses to each linear 15-mer peptide were mapped across the SARS-CoV-2 proteome and average RFU calculated for each amino acid residue.

Second, the normalized positional ‘epitope coverage’ at each amino acid residue within the proteins was defined as the ratio of total peptides mapped to each position by the total expected peptides, with values ranging between 0 to 1. A value of 1 in the SARS-CoV-2-positive group means that amino acid residues within the proteins were covered by peptides that showed immune response in all 10 SARS-CoV-2-positive patients and 14 peptides that overlap that position. (14 x 10 = 140 is the theoretical expected positional coverage to be 100%). Similarly, a value of 1 in SARS-CoV-2-negative group is 70 peptides with response (14 peptides x 5 SARS-CoV-2-negative patients = 70. i.e all 70 unique peptides that cover residue locations).

Third, to identify the epitopes that are particularly prevalent in SARS-CoV-2-positive subjects, we performed an indicator value analysis (*33*). This type of analysis is frequently used in ecology to determine whether species have significant associations with certain site groups. We applied this method to epitopes presence/absence data by replacing species with epitopes and site groups with patient groups defined by SARS-CoV-2 PCR status (positive or negative). The indicator value analysis measures the IndVal metric, which is the product of the specificity (e.g. the proportion of individuals within the whole dataset that exhibit a response to the epitope and belongs to a certain patient group) and the fidelity (e.g. the proportion of individuals within a certain patient group that exhibits a response) to the epitope. To control for the differences in sample size between patient groups, we used the group-equalized version of IndVal, IndValpag (*33*). The R function *multipatt* from the R package *indicspecies* allowed us to perform this analysis and evaluate the significance of the associations through permutation tests (*33*).

### Structural properties of B cell epitopes and B cell epitope prediction

SARS-CoV-2 protein sequences were obtained from Uniprot (*69*). Structure models of all 24 proteins in SARS-CoV-2 were obtained from I-TASSER (*70*). Solvent accessibility was calculated using freeSASA (*71*). Pymol was used for visualization. Using 3D structures and biophysical properties of the SARS-CoV-2 proteome, we applied the DiscoTope algorithm (*26*) to computationally predict conformational B cell epitopes with a significance threshold of -7.7 (75% specificity, 45% sensitivity). In addition, we used the Bepipred algorithm (*27*) to obtain linear B cell epitopes. Epitopes with minimum length of 7 amino acid residues and minimum score of 0.55 (80% specificity, 30% sensitivity) were used for the analysis.

### Epitope evolution profiling

To understand the evolution of SARS-CoV-2 epitopes in SARS-CoV2-positive patients, we made use of single nucleotide variants (SNVs) from 38,685 whole genome sequences from the NCBI sequence read archive (Table S9, see https://dataverse.harvard.edu/dataset.xhtml?persistentId=doi:10.7910/DVN/4ZXDW0). We selected SARS-CoV-2 samples from the first pandemic wave (defined as January 1 to July 31, 2020) and the second wave (defined as August 1 to December 31 2020) sequenced using Illumina paired-end amplicons with a minimum average depth of coverage of 200 x and fewer than 10,000 sites with a depth of coverage lower than 100x. We then retained single nucleotide variants present in both minus and plus strands at a minimum frequency of 2%, occurring at sites with a minimum depth of 100x, having a minimum within-sample frequency of 5% and located between sites 101 and 29778 of the genome to exclude sites at the extremities that are prone to sequencing errors and have been frequently masked (*72*). These additional filters allowed us to remove sequencing errors and provided deep coverage to identify SNVs that are polymorphic within patients, reflecting within-patient evolution (*34, 35*), as well as those that are shared between the consensus sequences of different patients.

Next, to compare epitope evolution from the evolution of non-epitope sites of the same protein, we measured the evolution rates at within-host (SNVs with a frequency <75% that are not transmitted for sure) and between-host/transmission level (SNVs with a frequency ≥75% that are observed in at least three samples). Because the number of SNVs observed will vary depending on sample coverage, which varies across samples, we estimated the evolution rates in each sample separately using the number of SNVs observed per site with adequate coverage. Such sites are defined as having a detection power of at least 80%, which is the probability of detecting five reads supporting the presence of a SNVs with a frequency of at least 5% in a site of coverage C, i.e. the minimum adequate coverage, under a binomial distribution. This approach has been used previously for similar purposes with the Lassa virus (*73*).

We also inferred selection in the proteins of interest (ORF1A/B, Spike (S) protein, Envelope (E) protein, Membrane (M) glycoprotein, Nucleocapsid (N) phosphoprotein) using dN/dS, the ratio of non-synonymous (dN) and synonymous substitutions rates (dS), which we calculated from the called SNVs in each sample (*74*).

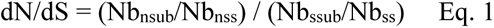

where Nb_nsub_ is the number of non-synonymous substitutions, Nb_nss_ is the number of non-synonymous sites, Nb_ssub_ is the number of synonymous substitutions, and Nb_ss_ is the number of synonymous sites. dN/dS can detect purifying selection (dN/dS<1), neutral evolution (dN/dS ≈ 1) and positive selection (dN/dS > 1). In each sample, we calculated dN/dS only if there were more than three SNVs including at least one synonymous SNV.

Finally, we inferred selection at the within-host level, using pN/pS, which we calculated from intrahost SNVs (iSNVs), i.e. SNVs that are not fixed (within-sample frequency <75%):

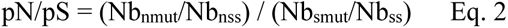

where Nb_nmut_ is the number of non-synonymous iSNVs, Nb_nss_ is the number of non-synonymous sites, Nb_smut_ is the number of synonymous iSNVs, and Nb_ss_ is the number of synonymous sites. These analyses have been implemented in R (https://github.com/arnaud00013/SARS-CoV-2-HPDA-evolutionary-analysis).

### Selection for immune escape in VOCs and VUIs genomes

To reveal VOCs and VUIs mutations possibly involved in selection for immune escape, we first defined the signature mutations of each variant (Table S10) as substitutions that are present in >=90% of sequences assigned to that lineage. We calculated the prevalence of substitutions in thousands of publicly available consensus sequences collected during 2020 and added data from CoV-Spectrum about under-represented lineage in the database or lineages that emerged during 2021 (*40*). Then, we only focused on nonsynonymous signature mutations in our database and asked if these signature mutations are located at epitope sites as these mutations can change the antibodies’ ability to recognize the epitopes. The signature mutation prevalence data were collected from our database of NCBI samples for the earlier lineages (PANGO v.2.1.7) and from GISAID data obtained from cov-spectrum for more recent lineages like Omicron. The database of lineage signature mutations is available on Github (https://github.com/arnaud00013/SARS-CoV-2-HPDA-evolutionary-analysis).

## SUPPLEMENTARY INFORMATION

**Figure S1.**
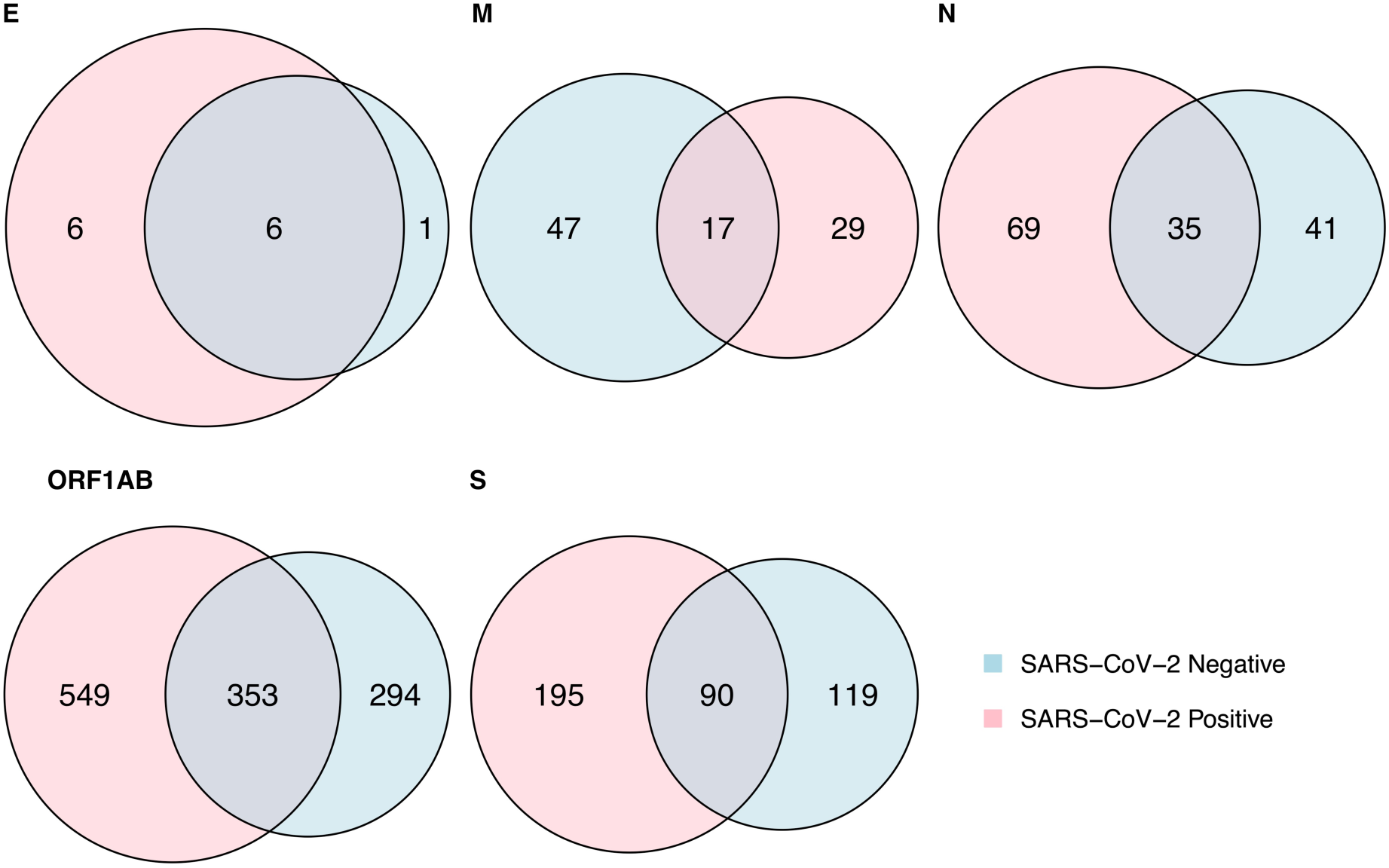
Numbers of SARS-CoV-2-specific epitope-defining peptides identified by HDPA. Numbers of SARS-CoV-2-specific epitopes identified with high density peptide arrays (HDPA) in proteins spike (S) protein, envelope (E) protein, membrane (M) glycoprotein, nucleocapsid (N) phosphoprotein, and ORF1AB. Numbers of unique peptides that showed a significant antibody response (RFU ≥ 1000) in SARS-CoV-2-negative (blue) and SARS-CoV-2-positive (red) groups are depicted. Some peptides are present in both groups, shown as overlap.

**Figure S2.**
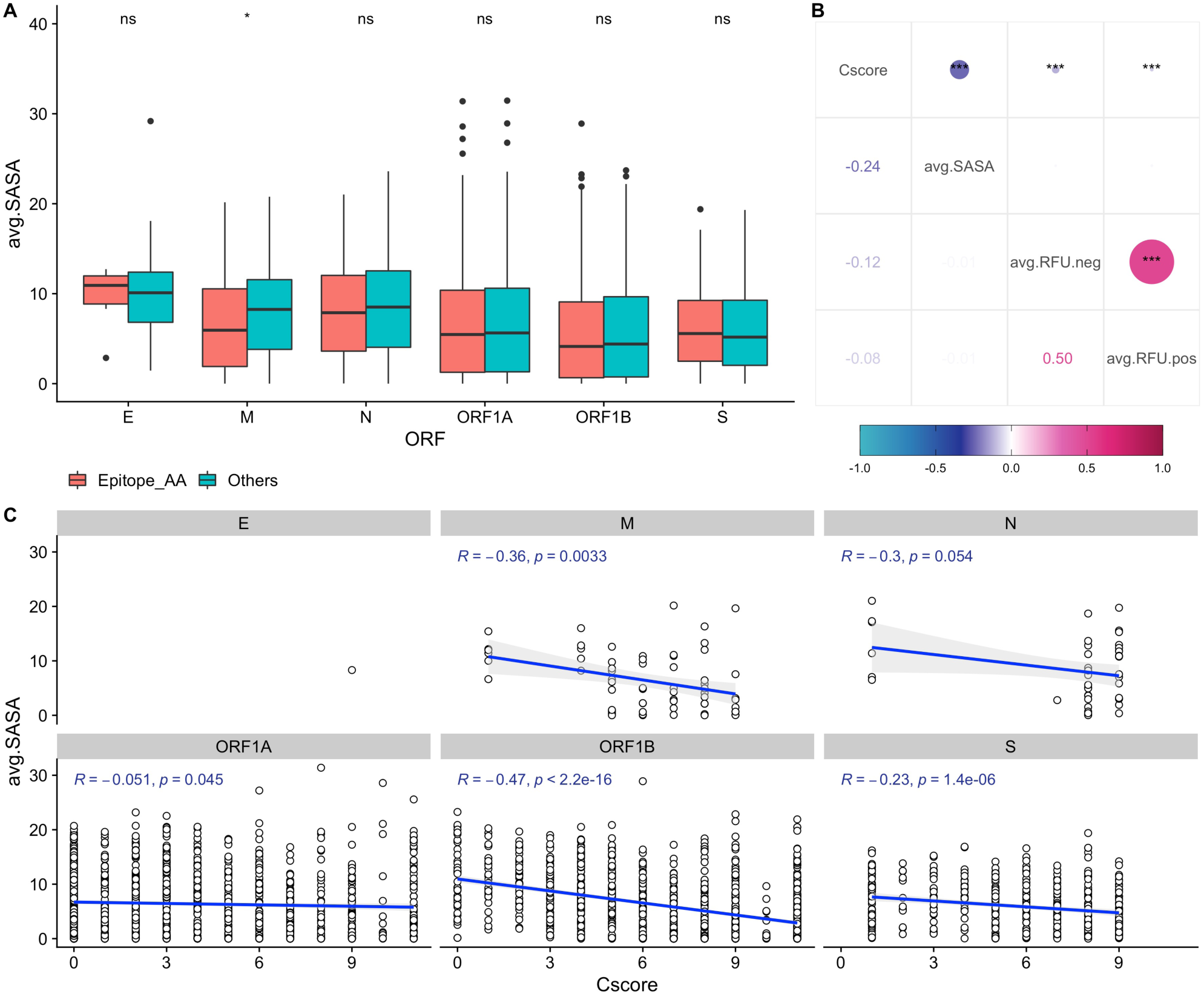
Relationship between structural properties of epitope sites residues. (A) Comparison of solvent accessibility (SASA) between residues in differential epitope sites and rest of the protein. The group “Epitope_AA” includes all residues that are part of differential epitopes and was compared with rest of the amino acids in the protein (“Others”). Student t-test was performed (ns = not significant, * = <0.05). (B) Correlation plot between RFU, SASA and, conservation score (Cscore) in differential epitope sites. Top triangle above the diagonal shows p-value between correlations (*** = > 0.001, **= > 0.01, * = > 0.05). Bottom triangle shows the correlation coefficients. (C) Correlation between conservation score and average solvent accessibility in differential epitope sites.

**Figure S3.**
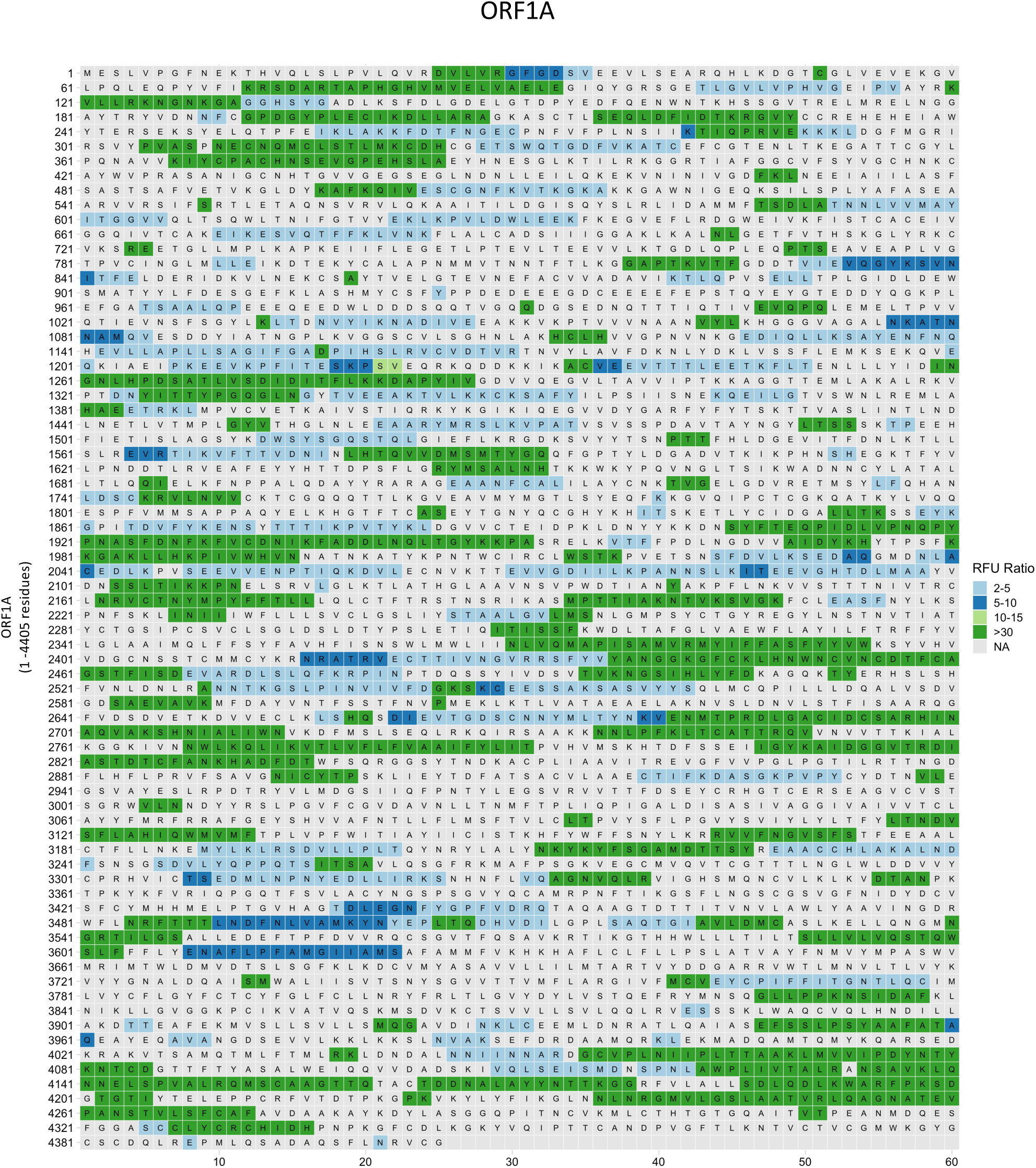
Identified differential epitope sites in ORF1A of SARS-CoV-2. Relative Fluorescence Unit (RFU) values of HDPA analysis were used to calculate ratio values to define differential epitope sites and are color coded. Residues that are not part of epitopes are marked in grey.

**Figure S4.**
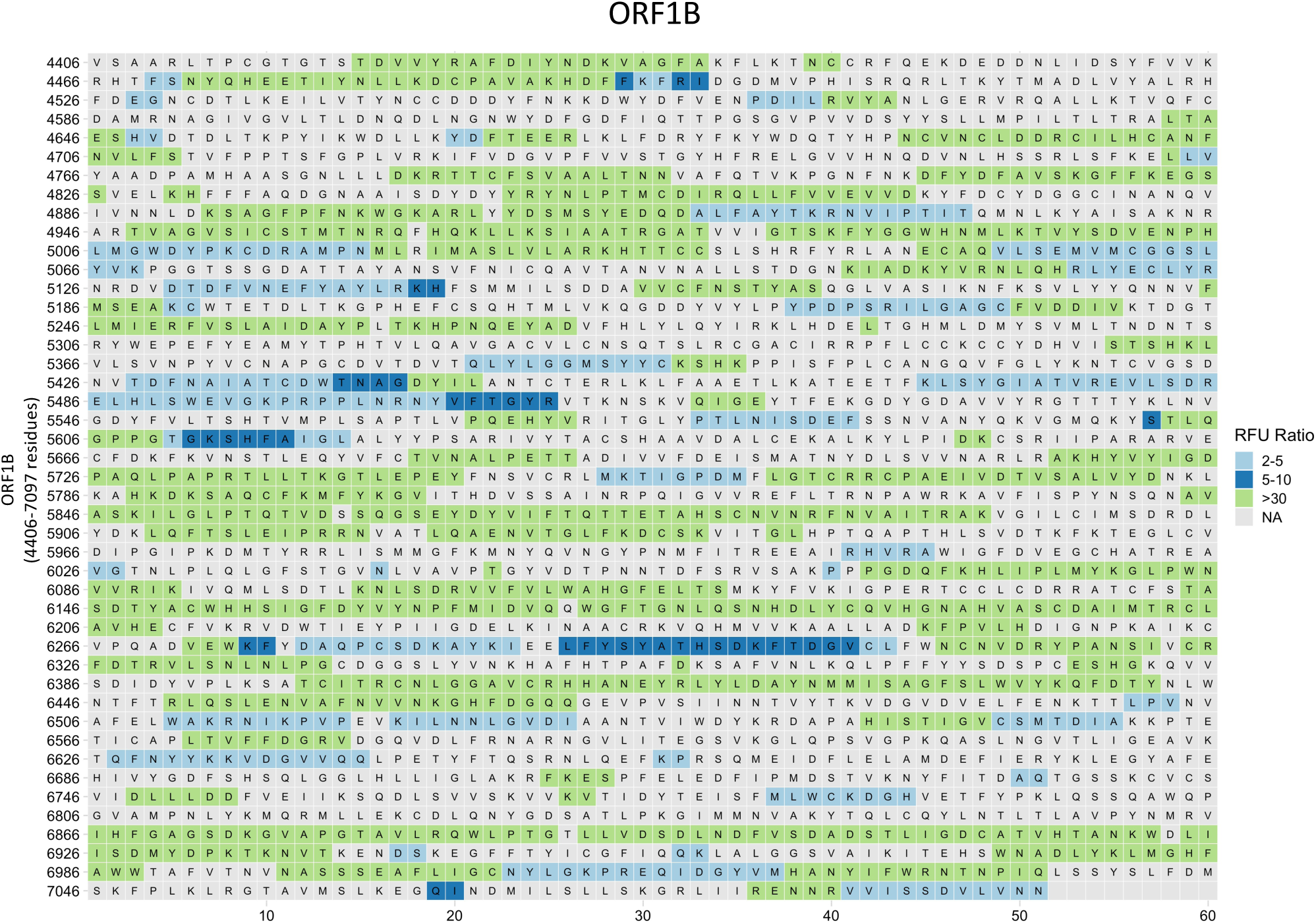
Identified differential epitope sites in ORF1B of SARS-CoV-2. Relative Fluorescence Unit (RFU) values of HDPA analysis were used to calculate ratio values to define differential epitope sites and are color coded. Residues that are not part of epitopes are marked in grey.

**Figure S5.**
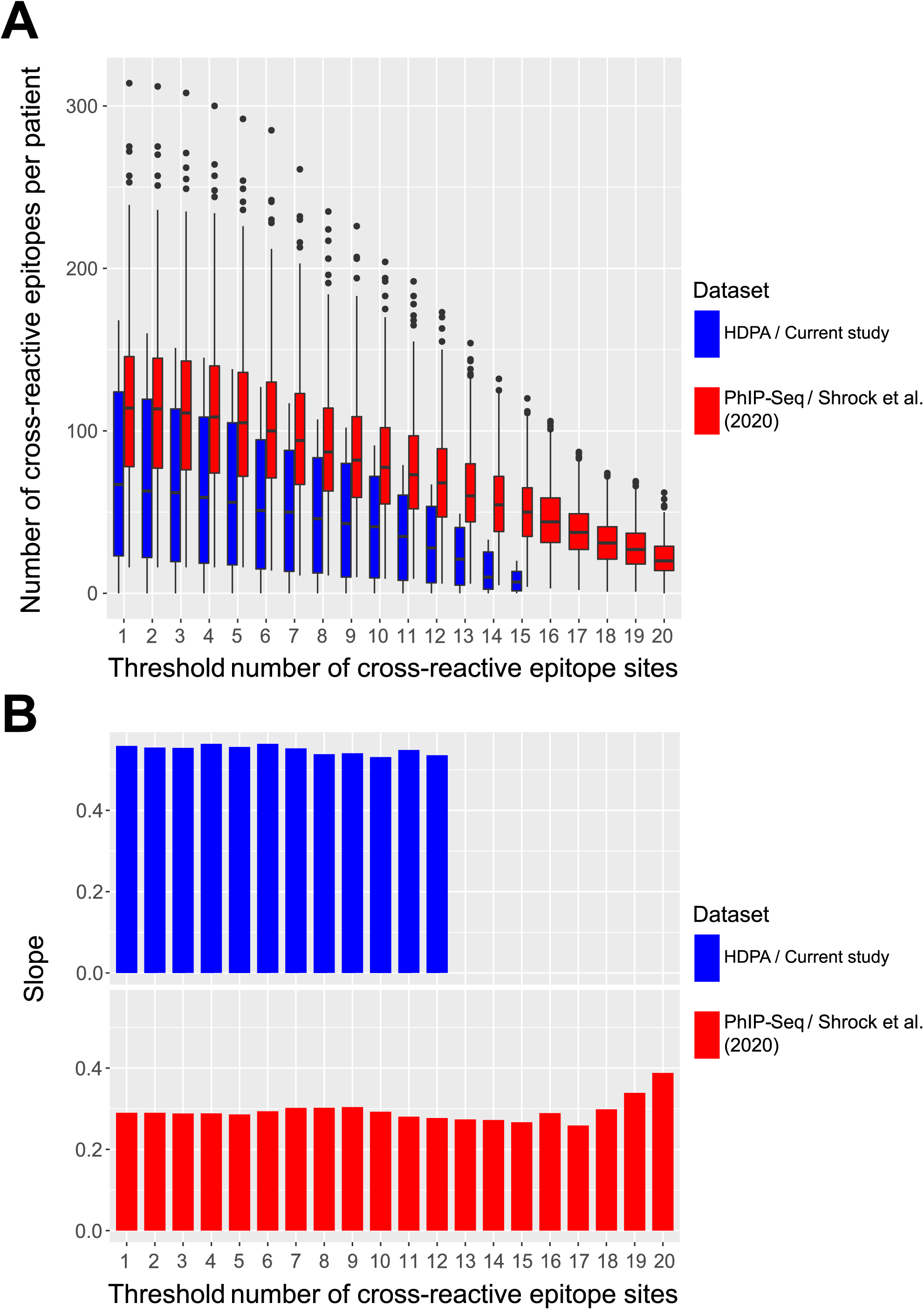
Sensitivity analysis of the number of cross-reactive epitope sites that define a cross-reactive epitope. (A) Numbers of cross-reactive epitopes per patient in relation to the number of cross-reactive epitope sites in HDPA (blue) and recently published PhiP-Seq study (red; (*21*)). (B) Slope of the correlation between the average antibody response and the number of cross-reactive epitopes in relation to the minimum number of cross-reactive epitope sites in HDPA (blue) and recently published PhiP-Seq study (red; (*21*)). Using HDPA, the antibody responses were measured as relative fluorescent units (RFU), while it was measured as the Z-score in a recently published PhiP-Seq study (*21*). Data have been normalized before performing the linear regressions to respect the assumption of normally distributed residues.

**Figure S6.**
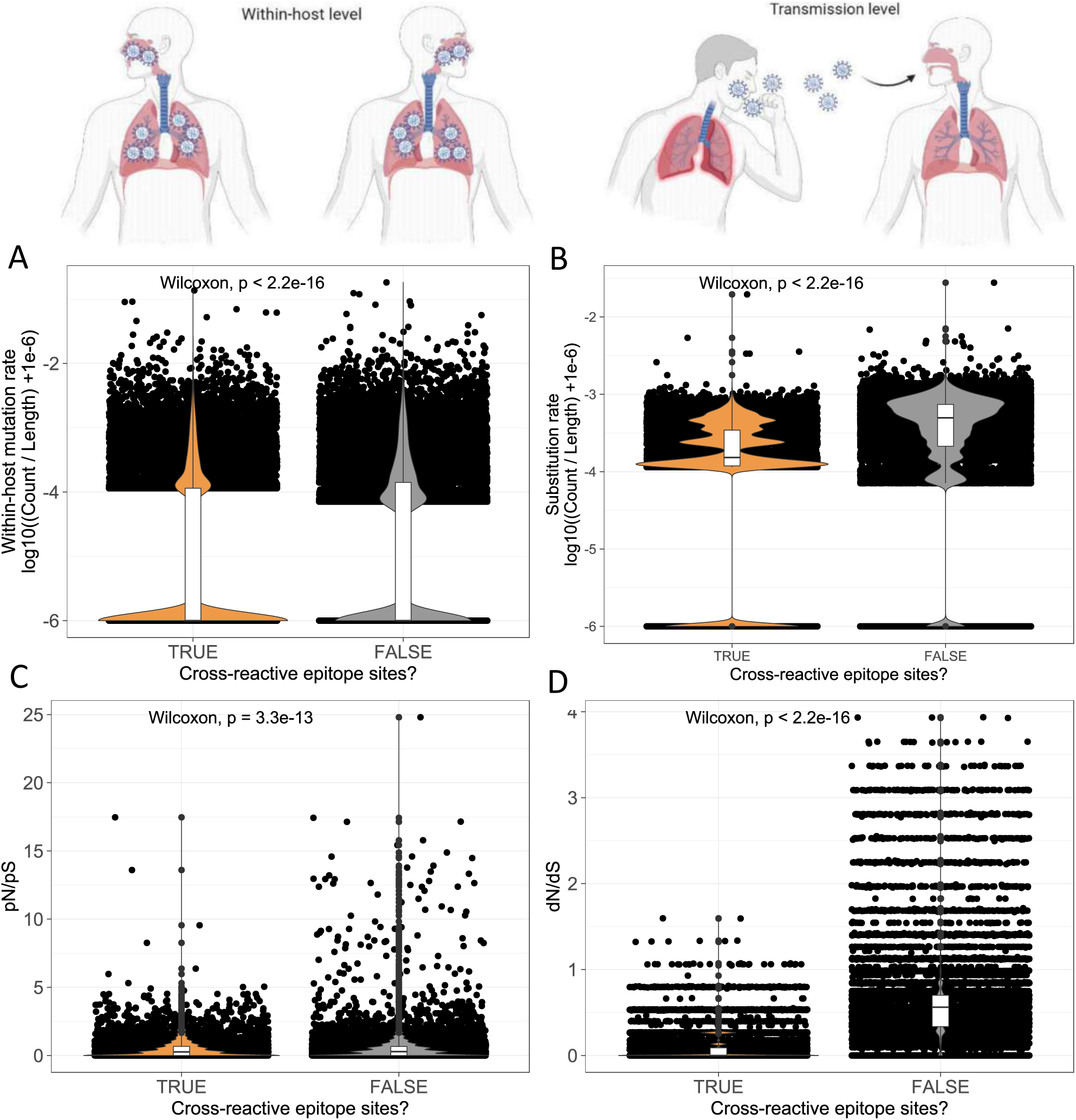
Evolution profile of cross-reactive epitope sites compared to the global epitope pool. (A) Distribution of the within-host mutation rates of cross-reactive epitope sites (orange) vs non-cross-reactive epitope sites (grey) across samples is shown. (B) Distribution of the substitution rate of cross-reactive epitope sites (orange) vs non-cross-reactive epitope sites (grey) across samples is shown. (C) Distribution of pN/pS of cross-reactive epitope sites (orange) vs non-cross-reactive epitope sites (grey) across samples is shown. (D) Distribution of dN/dS of cross-reactive epitope sites (orange) vs non-cross-reactive epitope sites (grey) across samples is shown.

**Figure S7.**
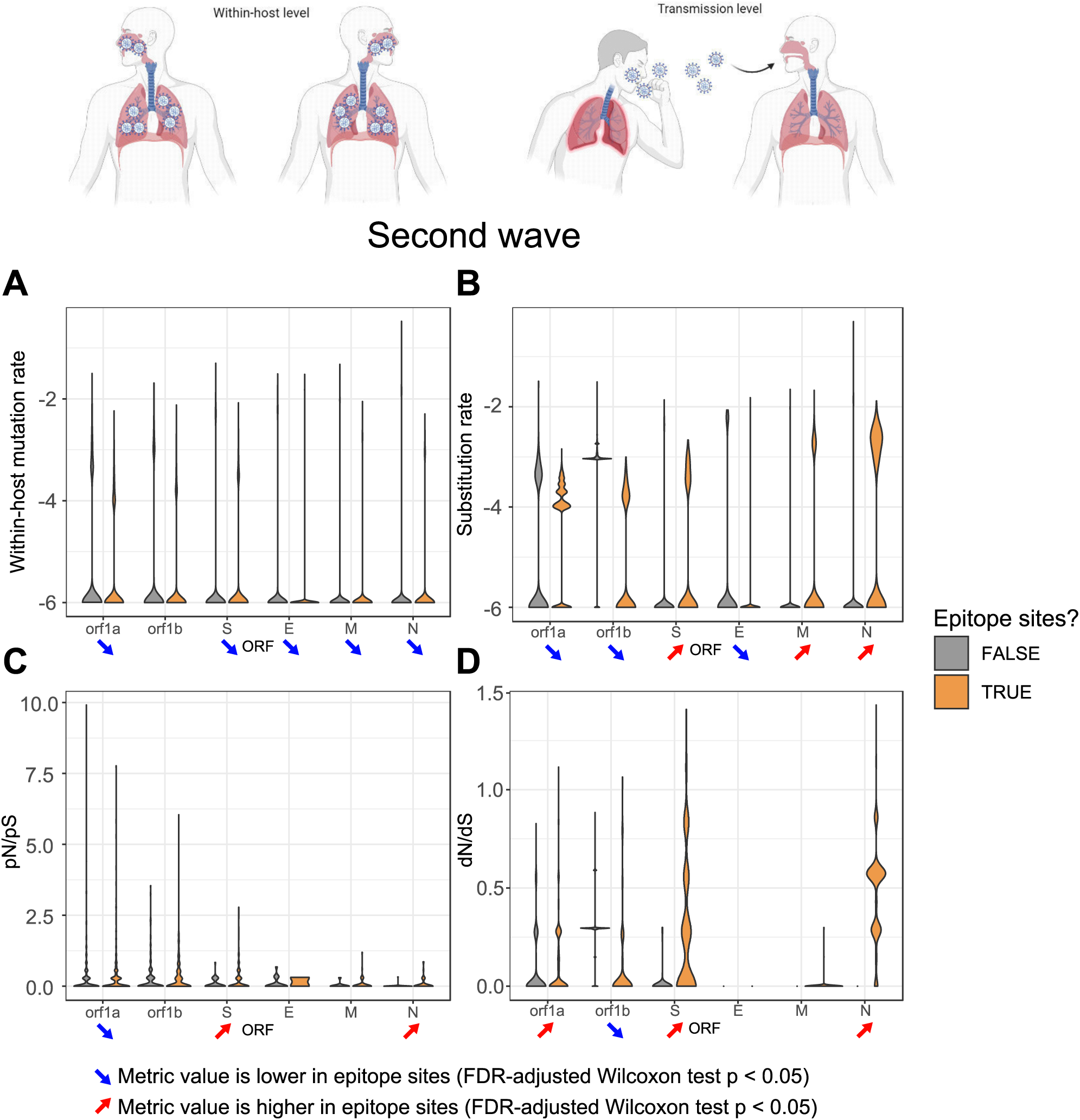
Evolutionary profile of SARS-CoV-2 epitopes during the second pandemic wave. The extent to which natural selection for immune evasion acts on SARS-CoV-2 during infection, or upon transmission is investigated. The distribution of evolutionary parameters in epitope sites (orange) vs non-epitope sites (grey) during the second pandemic SARS-CoV-2 wave (defined as August 1 to December 31 2020) is depicted. For each metric, significantly lower values in epitope sites of a certain gene are represented by a blue arrow pointing down while significantly higher values in epitope sites of a certain gene are represented by a red arrow pointing up (FDR-adjusted Wilcoxon test p<0.05). (A) Distributions of sample mutation rates (*log10(Count/gene length + 1e-6)*) across targeted proteins during were analyzed. (B) Distributions of sample substitution rate (*log10(Count/gene length + 1e-6)*) across targeted proteins were investigated. (C) Analysis of distributions of sample pN/pS across targeted proteins. (D) Distributions of sample dN/dS across targeted proteins were examined.

**Fig. S8.**
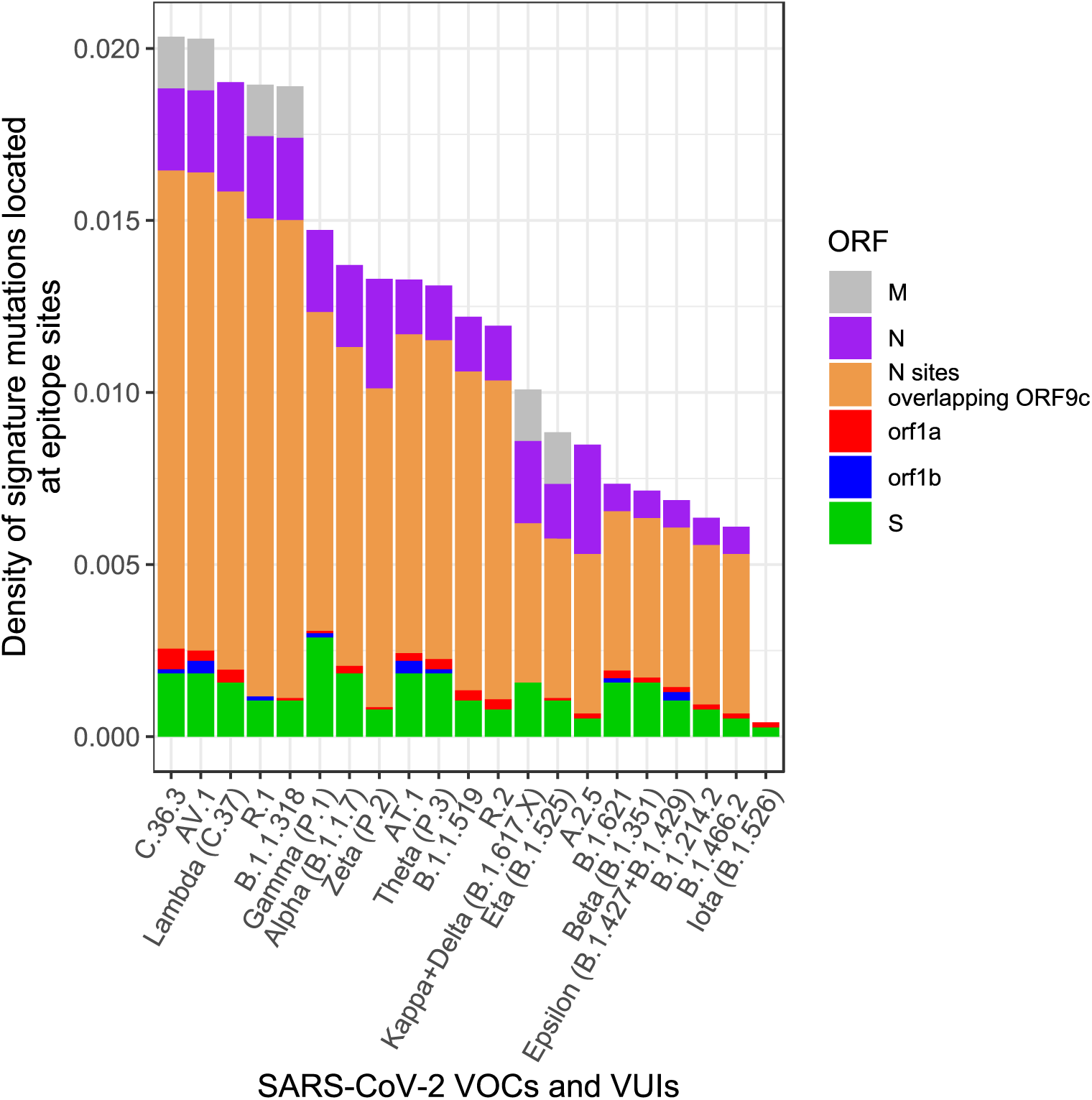
Density of nonsynonymous signature mutations of SARS-CoV-2 variants located at epitope sites. Density of nonsynonymous signature mutations in epitopes of selected VOCs and VUIs normalized by gene length. For each VOC/VUI we indicate the density of signature mutations in epitopes identified with HDPA across all analyzed ORFs: envelope (E) protein (orange), membrane (M) glycoprotein (grey), N sites overlapping ORF9c (black), ORF1b (blue), nucleocapsid (N) phosphoprotein (purple), ORF1A (red), spike (S) protein (green).

**Table S1. Serum Samples and Study Population.**

Positivity of SARS-CoV-2 infection was assessed both by PCR.

**Table S2. SARS-CoV-2-specific epitope information.**

Detailed information of SARS-CoV-2-specific epitopes identified with high density peptide arrays (HDPA) in structural proteins; spike (S) protein, envelope (E) protein, membrane (M) glycoprotein, nucleocapsid (N) phosphoprotein, and ORF1AB. Peptides that showed a significant antibody response (RFU ≥ 1000) in SARS-CoV-2-negative and SARS-CoV-2-positive groups are depicted. Some peptides are present in both groups, referred to as overlapping. Start and end of alignment amino acid (AA) positions of peptides with respect to their full-length protein was obtained by aligning the peptides to the protein sequence.

**Table S3. OC43-specific epitope information.**

Detailed information of OC43-specific epitopes identified with high density peptide arrays (HDPA) in structural proteins; spike (S) protein, envelope (E) protein, membrane (M) glycoprotein, nucleocapsid (N) phosphoprotein, and ORF1ab. Peptides that showed a significant antibody response (RFU ≥ 1000) in SARS-CoV-2-negative and SARS-CoV-2-positive groups are depicted. Some peptides are present in both groups, referred to as overlapping. Start and end of alignment amino acid (AA) positions of peptides with respect to their full-length protein was obtained by aligning the peptides to the protein sequence.

**Table S4. HKU1-specific epitope information.**

Detailed information of HKU1-specific epitopes identified with high density peptide arrays (HDPA) in structural proteins; spike (S) protein, envelope (E) protein, membrane (M) glycoprotein, nucleocapsid (N) phosphoprotein, and ORF1ab. Peptides that showed a significant antibody response (RFU ≥ 1000) in SARS-CoV-2-negative and SARS-CoV-2-positive groups are depicted. Some peptides are present in both groups, referred to as overlapping. Start and end of alignment amino acid (AA) positions of peptides with respect to their full-length protein was obtained by aligning the peptides to the protein sequence.

**Table S5. NL63-specific epitope information.**

Detailed information of NL63-specific epitopes identified with high density peptide arrays (HDPA) in structural proteins; spike (S) protein, envelope (E) protein, membrane (M) glycoprotein, nucleocapsid (N) phosphoprotein, and ORF1ab. Peptides that showed a significant antibody response (RFU ≥ 1000) in SARS-CoV-2-negative and SARS-CoV-2-positive groups are depicted. Some peptides are present in both groups, referred to as overlapping. Start and end of alignment amino acid (AA) positions of peptides with respect to their full-length protein was obtained by aligning the peptides to the protein sequence.

**Table S6. 229E-specific epitope information.**

Detailed information of 229E-specific epitopes identified with high density peptide arrays (HDPA) in structural proteins; spike (S) protein, envelope (E) protein, membrane (M) glycoprotein, nucleocapsid (N) phosphoprotein, and ORF1ab. Peptides that showed a significant antibody response (RFU ≥ 1000) in SARS-CoV-2-negative and SARS-CoV-2-positive groups are depicted. Some peptides are present in both groups, referred to as overlapping. Start and end of alignment amino acid (AA) positions of peptides with respect to their full-length protein was obtained by aligning the peptides to the protein sequence.

**Table S7. Cross-reactive epitope sites.**

The columns of the table indicate from left to right: the proteins of SARS-CoV-2, the amino acid position of the site in the protein, the average RFU of antibody responses detected in SARS-CoV-2-positive patients for epitopes mapping at the site, the average RFU of antibody responses detected in SARS-CoV-2-negative patients for epitopes mapping at the site, the presence of cross-reactivity, and the number of mutations observed in NCBI samples during 2020 (first and second wave).

**Table S8. Epitopes that are significantly associated to COVID-19-positive patients.**

Detailed information on epitopes from a recent PhIP-Seq study (*21*) dataset that are significant indicators or COVID-19-positive patients. The p-value was obtained from the IndVal test and are corrected for multiple testing using the Sidak method.

**Table S9. Features of the called SNVs.**

https://dataverse.harvard.edu/dataset.xhtml?persistentId=doi:10.7910/DVN/4ZXDW0

**Table S10. VOCs and VUIs nonsynonymous signature mutations in epitopes.**

**Table S11. Features of all the nonsynonymous mutations detected at epitope sites**.

**Table S12. Coronavirus taxonomy and sequence accession numbers for analyzed proteins.**

