## Supplementary Tables for "Selection for immune evasion in SARS-CoV-2 revealed by high-resolution epitope mapping combined with genome sequence analysis"

| Patient # | Clinical status | Age | Sex | Molecular diagnosis for SARS-CoV-2 (PCR) |
|---|---|---|---|---|
| | **Asymptomatics** | | | **SARS-CoV-2-positive** |
| 1 | | 56 | male | positive |
| 2 | | 53 | female | positive |
| 3 | | 28 | female | positive |
| 4 | | 35 | female | positive |
| 5 | | 31 | female | positive |
| | **Recovered** | | | **SARS-CoV-2-positive** |
| 6 | | 61 | male | positive |
| 7 | | 67 | male | positive |
| 8 | | 44 | female | positive |
| 9 | | 42 | male | positive |
| 10 | | 59 | female | positive |
| | **Negative** | | | **SARS-CoV-2-negative** |
| 11 | | 27 | male | negative |
| 12 | | 53 | female | negative |
| 13 | | 44 | male | negative |
| 14 | | 47 | male | negative |
| 15 | | 66 | female | negative |

Table S1

**Table S2. SARS-CoV2-specific Peptides**

| Lineage | Protein | HDPA peptide sequence | AA pos start | AA pos end | Group |
|---|---|---|---|---|---|
| SARS-CoV-2 | envelope | GSGMYSFVSEETGTL | 1 | 12 | Positive |
| SARS-CoV-2 | envelope | GSGSGMYSFVSEETG | 1 | 10 | Positive |
| SARS-CoV-2 | envelope | SGSGSGMYSFVSEET | 1 | 9 | Negative |
| SARS-CoV-2 | envelope | GMYSFVSEETGTLIV | 1 | 14 | Overlapping |
| SARS-CoV-2 | envelope | GSGSGSGMYSFVSEE | 1 | 8 | Overlapping |
| SARS-CoV-2 | envelope | FVSEETGTLIVNSVL | 4 | 18 | Positive |
| SARS-CoV-2 | envelope | VSEETGTLIVNSVLL | 5 | 19 | Positive |
| SARS-CoV-2 | envelope | SEETGTLIVNSVLLF | 6 | 20 | Positive |
| SARS-CoV-2 | envelope | EETGTLIVNSVLLFL | 7 | 21 | Overlapping |
| SARS-CoV-2 | envelope | ETGTLIVNSVLLFLA | 8 | 22 | Overlapping |
| SARS-CoV-2 | envelope | SSRVPDLLVGSGSGS | 67 | 75 | Positive |
| SARS-CoV-2 | envelope | SSRVPDLLVGSGSGS | 67 | 75 | Overlapping |
| SARS-CoV-2 | membrane | GSGSGMADSNGTITV | 1 | 10 | Positive |
| SARS-CoV-2 | membrane | GSGSGSGMADSNGTI | 1 | 8 | Positive |
| SARS-CoV-2 | membrane | GMADSNGTITVEELK | 1 | 14 | Negative |
| SARS-CoV-2 | membrane | GMSDNGPQNQRNAPR | 1 | 6 | Negative |
| SARS-CoV-2 | membrane | GSGMADSNGTITVEE | 1 | 12 | Negative |
| SARS-CoV-2 | membrane | GSGSGMADSNGTITV | 1 | 10 | Negative |
| SARS-CoV-2 | membrane | GSGSGSGMADSNGTI | 1 | 8 | Negative |
| SARS-CoV-2 | membrane | GSGMADSNGTITVEE | 1 | 12 | Overlapping |
| SARS-CoV-2 | membrane | GGPSDSTGSNQNGER | 2 | 7 | Negative |
| SARS-CoV-2 | membrane | PEAGLPYGANKDGII | 2 | 8 | Negative |
| SARS-CoV-2 | membrane | ADSNGTITVEELKKL | 2 | 16 | Overlapping |
| SARS-CoV-2 | membrane | SNGTITVEELKKLLE | 4 | 18 | Positive |
| SARS-CoV-2 | membrane | GTITVEELKKLLEQW | 6 | 20 | Negative |
| SARS-CoV-2 | membrane | TITVEELKKLLEQWN | 7 | 21 | Positive |
| SARS-CoV-2 | membrane | ITVEELKKLLEQWNL | 8 | 22 | Positive |
| SARS-CoV-2 | membrane | VEELKKLLEQWNLVI | 10 | 24 | Positive |
| SARS-CoV-2 | membrane | EELKKLLEQWNLVIG | 11 | 25 | Overlapping |
| SARS-CoV-2 | membrane | EQWNLVIGFLFLTWI | 18 | 32 | Overlapping |
| SARS-CoV-2 | membrane | NLVIGFLFLTWICLL | 21 | 35 | Positive |
| SARS-CoV-2 | membrane | GTDYKHWPQIAQFAP | 25 | 38 | Negative |
| SARS-CoV-2 | membrane | NRNRFLYIIKLIFLW | 41 | 55 | Positive |
| SARS-CoV-2 | membrane | NRFLYIIKLIFLWLL | 43 | 57 | Positive |
| SARS-CoV-2 | membrane | VLAAVYRINWITGGI | 66 | 80 | Negative |
| SARS-CoV-2 | membrane | LAAVYRINWITGGIA | 67 | 81 | Negative |
| SARS-CoV-2 | membrane | AAVYRINWITGGIAI | 68 | 82 | Negative |
| SARS-CoV-2 | membrane | NWITGGIAIAMACLV | 74 | 88 | Positive |
| SARS-CoV-2 | membrane | GLPNNTASWFTALTQ | 74 | 77 | Negative |
| SARS-CoV-2 | membrane | QGLPNNTASWFTALT | 74 | 77 | Negative |
| SARS-CoV-2 | membrane | GIAIAMACLVGLMWL | 79 | 93 | Negative |
| SARS-CoV-2 | membrane | GMEVTPSGTWLTYTG | 79 | 95 | Negative |
| SARS-CoV-2 | membrane | GMSRIGMEVTPSGTW | 79 | 92 | Negative |
| SARS-CoV-2 | membrane | EAGLPYGANKDGIIW | 89 | 92 | Negative |
| SARS-CoV-2 | membrane | GIIWVATEGALNTPK | 89 | 94 | Negative |
| SARS-CoV-2 | membrane | GLMWLSYFIASFRLF | 89 | 103 | Negative |

| | | | | | |
|---|---|---|---|---|---|
| SARS-CoV-2 | membrane | GLPYGANKDGIIWVA | 89 | 94 | Negative |
| SARS-CoV-2 | membrane | YGANKDGIIWVATEG | 89 | 94 | Negative |
| SARS-CoV-2 | membrane | RTRSMWSFNPETNIL | 105 | 119 | Positive |
| SARS-CoV-2 | membrane | SMWSFNPETNILLNV | 108 | 122 | Positive |
| SARS-CoV-2 | membrane | PETNILLNVPLHGTI | 114 | 128 | Positive |
| SARS-CoV-2 | membrane | ETNILLNVPLHGTIL | 115 | 129 | Overlapping |
| SARS-CoV-2 | membrane | NILLNVPLHGTILTR | 117 | 131 | Positive |
| SARS-CoV-2 | membrane | EGALNTPKDHIGTRN | 120 | 127 | Negative |
| SARS-CoV-2 | membrane | NVPLHGTILTRPLLE | 121 | 135 | Positive |
| SARS-CoV-2 | membrane | PLHGTILTRPLLESE | 123 | 137 | Overlapping |
| SARS-CoV-2 | membrane | GTGPEAGLPYGANKD | 130 | 142 | Negative |
| SARS-CoV-2 | membrane | TGPEAGLPYGANKDG | 130 | 142 | Negative |
| SARS-CoV-2 | membrane | GPEAGLPYGANKDGI | 132 | 142 | Negative |
| SARS-CoV-2 | membrane | PLLESELVIGAVILR | 132 | 146 | Overlapping |
| SARS-CoV-2 | membrane | LLESELVIGAVILRG | 133 | 147 | Positive |
| SARS-CoV-2 | membrane | LESELVIGAVILRGH | 134 | 148 | Overlapping |
| SARS-CoV-2 | membrane | ESELVIGAVILRGHL | 135 | 149 | Overlapping |
| SARS-CoV-2 | membrane | SELVIGAVILRGHLR | 136 | 150 | Positive |
| SARS-CoV-2 | membrane | ELVIGAVILRGHLRI | 137 | 151 | Negative |
| SARS-CoV-2 | membrane | IGAVILRGHLRIAGH | 140 | 154 | Positive |
| SARS-CoV-2 | membrane | GAVILRGHLRIAGHH | 141 | 155 | Positive |
| SARS-CoV-2 | membrane | GHLRIAGHHLGRCDI | 147 | 161 | Negative |
| SARS-CoV-2 | membrane | GHLRIAGHHLGRCDI | 147 | 161 | Overlapping |
| SARS-CoV-2 | membrane | QHGKEDLKFPRGQGV | 154 | 166 | Negative |
| SARS-CoV-2 | membrane | HLGRCDIKDLPKEIT | 155 | 169 | Positive |
| SARS-CoV-2 | membrane | HGKEDLKFPRGQGVP | 155 | 166 | Negative |
| SARS-CoV-2 | membrane | LGRCDIKDLPKEITV | 156 | 170 | Positive |
| SARS-CoV-2 | membrane | GKEDLKFPRGQGVPI | 157 | 166 | Negative |
| SARS-CoV-2 | membrane | GKMKDLSPRWYFYYL | 157 | 166 | Negative |
| SARS-CoV-2 | membrane | GRCDIKDLPKEITVA | 157 | 171 | Overlapping |
| SARS-CoV-2 | membrane | RCDIKDLPKEITVAT | 158 | 172 | Positive |
| SARS-CoV-2 | membrane | CDIKDLPKEITVATS | 159 | 173 | Overlapping |
| SARS-CoV-2 | membrane | DIKDLPKEITVATSR | 160 | 174 | Positive |
| SARS-CoV-2 | membrane | DETQALPQRQKKQQT | 160 | 166 | Negative |
| SARS-CoV-2 | membrane | EDLKFPRGQGVPINT | 160 | 166 | Negative |
| SARS-CoV-2 | membrane | IKDLPKEITVATSRT | 161 | 175 | Overlapping |
| SARS-CoV-2 | membrane | KDLPKEITVATSRTL | 162 | 176 | Positive |
| SARS-CoV-2 | membrane | KEITVATSRTLSYYK | 166 | 180 | Overlapping |
| SARS-CoV-2 | membrane | EITVATSRTLSYYKL | 167 | 181 | Overlapping |
| SARS-CoV-2 | membrane | EGSRGGSQASSRSSS | 171 | 177 | Negative |
| SARS-CoV-2 | membrane | YRRATRRIRGGDGKM | 178 | 192 | Negative |
| SARS-CoV-2 | membrane | ESKMSGKGQQQQGQT | 180 | 191 | Negative |
| SARS-CoV-2 | membrane | GRRGPEQTQGNFGDQ | 180 | 192 | Negative |
| SARS-CoV-2 | membrane | GPEQTQGNFGDQELI | 182 | 192 | Negative |
| SARS-CoV-2 | membrane | GTSPARMAGNGGDAA | 182 | 195 | Negative |
| SARS-CoV-2 | membrane | KFPRGQGVPINTNSS | 196 | 211 | Negative |
| SARS-CoV-2 | membrane | EQTQGNFGDQELIRQ | 202 | 204 | Negative |
| SARS-CoV-2 | membrane | GNFGDQELIRQGTDY | 202 | 204 | Negative |
| SARS-CoV-2 | membrane | GNYKLNTDHSSSSDN | 202 | 216 | Negative |

| | | | | | |
|---|---|---|---|---|---|
| SARS-CoV-2 | membrane | PEQTQGNFGDQELIR | 202 | 204 | Negative |
| SARS-CoV-2 | membrane | QGNFGDQELIRQGTD | 202 | 204 | Negative |
| SARS-CoV-2 | membrane | NYKLNTDHSSSSDNI | 203 | 217 | Overlapping |
| SARS-CoV-2 | membrane | KLNTDHSSSSDNIAL | 205 | 219 | Positive |
| SARS-CoV-2 | membrane | GVPINTNSSPDDQIG | 206 | 217 | Negative |
| SARS-CoV-2 | membrane | NTDHSSSSDNIALLV | 207 | 221 | Positive |
| SARS-CoV-2 | membrane | TDHSSSSDNIALLVQ | 208 | 222 | Positive |
| SARS-CoV-2 | membrane | DHSSSSDNIALLVQG | 209 | 222 | Positive |
| SARS-CoV-2 | membrane | SDNIALLVQGSGSGS | 214 | 222 | Positive |
| SARS-CoV-2 | nucleocapsid | GSGMSDNGPQNQRNA | 1 | 12 | Positive |
| SARS-CoV-2 | nucleocapsid | GSGSGMSDNGPQNQR | 1 | 10 | Positive |
| SARS-CoV-2 | nucleocapsid | GSGSGSGMSDNGPQN | 1 | 8 | Positive |
| SARS-CoV-2 | nucleocapsid | GSGMSDNGPQNQRNA | 1 | 12 | Negative |
| SARS-CoV-2 | nucleocapsid | GSGSGMSDNGPQNQR | 1 | 10 | Negative |
| SARS-CoV-2 | nucleocapsid | GSGSGSGMSDNGPQN | 1 | 8 | Negative |
| SARS-CoV-2 | nucleocapsid | GMSDNGPQNQRNAPR | 1 | 14 | Overlapping |
| SARS-CoV-2 | nucleocapsid | SDNGPQNQRNAPRIT | 2 | 16 | Positive |
| SARS-CoV-2 | nucleocapsid | NQRNAPRITFGGPSD | 8 | 22 | Positive |
| SARS-CoV-2 | nucleocapsid | NAPRITFGGPSDSTG | 11 | 25 | Positive |
| SARS-CoV-2 | nucleocapsid | GGPSDSTGSNQNGER | 18 | 32 | Negative |
| SARS-CoV-2 | nucleocapsid | GGPSDSTGSNQNGER | 18 | 32 | Overlapping |
| SARS-CoV-2 | nucleocapsid | PSDSTGSNQNGERSG | 20 | 34 | Positive |
| SARS-CoV-2 | nucleocapsid | DSTGSNQNGERSGAR | 22 | 36 | Negative |
| SARS-CoV-2 | nucleocapsid | ERSGARSKQRRPQGL | 31 | 45 | Overlapping |
| SARS-CoV-2 | nucleocapsid | QGLPNNTASWFTALT | 43 | 57 | Negative |
| SARS-CoV-2 | nucleocapsid | GLPNNTASWFTALTQ | 44 | 58 | Overlapping |
| SARS-CoV-2 | nucleocapsid | TASWFTALTQHGKED | 49 | 63 | Positive |
| SARS-CoV-2 | nucleocapsid | ASWFTALTQHGKEDL | 50 | 64 | Positive |
| SARS-CoV-2 | nucleocapsid | FTALTQHGKEDLKFP | 53 | 67 | Positive |
| SARS-CoV-2 | nucleocapsid | TALTQHGKEDLKFPR | 54 | 68 | Positive |
| SARS-CoV-2 | nucleocapsid | QHGKEDLKFPRGQGV | 58 | 72 | Overlapping |
| SARS-CoV-2 | nucleocapsid | HGKEDLKFPRGQGVP | 59 | 73 | Negative |
| SARS-CoV-2 | nucleocapsid | GKEDLKFPRGQGVPI | 60 | 74 | Negative |
| SARS-CoV-2 | nucleocapsid | KEDLKFPRGQGVPIN | 61 | 75 | Positive |
| SARS-CoV-2 | nucleocapsid | EDLKFPRGQGVPINT | 62 | 76 | Overlapping |
| SARS-CoV-2 | nucleocapsid | KFPRGQGVPINTNSS | 65 | 79 | Negative |
| SARS-CoV-2 | nucleocapsid | GVPINTNSSPDDQIG | 71 | 85 | Overlapping |
| SARS-CoV-2 | nucleocapsid | PINTNSSPDDQIGYY | 73 | 87 | Overlapping |
| SARS-CoV-2 | nucleocapsid | NSSPDDQIGYYRRAT | 77 | 91 | Positive |
| SARS-CoV-2 | nucleocapsid | SSPDDQIGYYRRATR | 78 | 92 | Negative |
| SARS-CoV-2 | nucleocapsid | SPDDQIGYYRRATRR | 79 | 93 | Overlapping |
| SARS-CoV-2 | nucleocapsid | PDDQIGYYRRATRRI | 80 | 94 | Negative |
| SARS-CoV-2 | nucleocapsid | DDQIGYYRRATRRIR | 81 | 95 | Overlapping |
| SARS-CoV-2 | nucleocapsid | YRRATRRIRGGDGKM | 87 | 101 | Negative |
| SARS-CoV-2 | nucleocapsid | DGKMKDLSPRWYFYY | 98 | 112 | Negative |
| SARS-CoV-2 | nucleocapsid | GKMKDLSPRWYFYYL | 99 | 113 | Negative |
| SARS-CoV-2 | nucleocapsid | GTGPEAGLPYGANKD | 114 | 128 | Negative |
| SARS-CoV-2 | nucleocapsid | TGPEAGLPYGANKDG | 115 | 129 | Overlapping |
| SARS-CoV-2 | nucleocapsid | GPEAGLPYGANKDGI | 116 | 130 | Overlapping |

| | | | | | |
|---|---|---|---|---|---|
| SARS-CoV-2 | nucleocapsid | PEAGLPYGANKDGII | 117 | 131 | Overlapping |
| SARS-CoV-2 | nucleocapsid | EAGLPYGANKDGIIW | 118 | 132 | Overlapping |
| SARS-CoV-2 | nucleocapsid | GLPYGANKDGIIWVA | 120 | 134 | Overlapping |
| SARS-CoV-2 | nucleocapsid | YGANKDGIIWVATEG | 123 | 137 | Negative |
| SARS-CoV-2 | nucleocapsid | NKDGIIWVATEGALN | 126 | 140 | Positive |
| SARS-CoV-2 | nucleocapsid | GIIWVATEGALNTPK | 129 | 143 | Negative |
| SARS-CoV-2 | nucleocapsid | VATEGALNTPKDHIG | 133 | 147 | Positive |
| SARS-CoV-2 | nucleocapsid | TEGALNTPKDHIGTR | 135 | 149 | Positive |
| SARS-CoV-2 | nucleocapsid | EGALNTPKDHIGTRN | 136 | 150 | Negative |
| SARS-CoV-2 | nucleocapsid | EGALNTPKDHIGTRN | 136 | 150 | Overlapping |
| SARS-CoV-2 | nucleocapsid | DHIGTRNPANNAAIV | 144 | 158 | Positive |
| SARS-CoV-2 | nucleocapsid | AAIVLQLPQGTTLPK | 155 | 169 | Positive |
| SARS-CoV-2 | nucleocapsid | AIVLQLPQGTTLPKG | 156 | 170 | Positive |
| SARS-CoV-2 | nucleocapsid | IVLQLPQGTTLPKGF | 157 | 171 | Positive |
| SARS-CoV-2 | nucleocapsid | LQLPQGTTLPKGFYA | 159 | 173 | Positive |
| SARS-CoV-2 | nucleocapsid | QLPQGTTLPKGFYAE | 160 | 174 | Positive |
| SARS-CoV-2 | nucleocapsid | LPQGTTLPKGFYAEG | 161 | 175 | Positive |
| SARS-CoV-2 | nucleocapsid | QGTTLPKGFYAEGSR | 163 | 177 | Positive |
| SARS-CoV-2 | nucleocapsid | AEGSRGGSQASSRSS | 173 | 187 | Positive |
| SARS-CoV-2 | nucleocapsid | EGSRGGSQASSRSSS | 174 | 188 | Negative |
| SARS-CoV-2 | nucleocapsid | EGSRGGSQASSRSSS | 174 | 188 | Overlapping |
| SARS-CoV-2 | nucleocapsid | GSRGGSQASSRSSSR | 175 | 189 | Overlapping |
| SARS-CoV-2 | nucleocapsid | SRGGSQASSRSSSRS | 176 | 190 | Negative |
| SARS-CoV-2 | nucleocapsid | RGGSQASSRSSSRSR | 177 | 191 | Positive |
| SARS-CoV-2 | nucleocapsid | GGSQASSRSSSRSRN | 178 | 192 | Positive |
| SARS-CoV-2 | nucleocapsid | GSQASSRSSSRSRNS | 179 | 193 | Positive |
| SARS-CoV-2 | nucleocapsid | SQASSRSSSRSRNSS | 180 | 194 | Positive |
| SARS-CoV-2 | nucleocapsid | QASSRSSSRSRNSSR | 181 | 195 | Positive |
| SARS-CoV-2 | nucleocapsid | ASSRSSSRSRNSSRN | 182 | 196 | Positive |
| SARS-CoV-2 | nucleocapsid | SSRSSSRSRNSSRNS | 183 | 197 | Overlapping |
| SARS-CoV-2 | nucleocapsid | SRSSSRSRNSSRNST | 184 | 198 | Overlapping |
| SARS-CoV-2 | nucleocapsid | RSSSRSRNSSRNSTP | 185 | 199 | Positive |
| SARS-CoV-2 | nucleocapsid | SSSRSRNSSRNSTPG | 186 | 200 | Overlapping |
| SARS-CoV-2 | nucleocapsid | SSRSRNSSRNSTPGS | 187 | 201 | Overlapping |
| SARS-CoV-2 | nucleocapsid | SRSRNSSRNSTPGSS | 188 | 202 | Negative |
| SARS-CoV-2 | nucleocapsid | RSRNSSRNSTPGSSR | 189 | 203 | Positive |
| SARS-CoV-2 | nucleocapsid | RNSSRNSTPGSSRGT | 191 | 205 | Positive |
| SARS-CoV-2 | nucleocapsid | NSSRNSTPGSSRGTS | 192 | 206 | Positive |
| SARS-CoV-2 | nucleocapsid | SSRNSTPGSSRGTSP | 193 | 207 | Negative |
| SARS-CoV-2 | nucleocapsid | RNSTPGSSRGTSPAR | 195 | 209 | Positive |
| SARS-CoV-2 | nucleocapsid | PGSSRGTSPARMAGN | 199 | 213 | Positive |
| SARS-CoV-2 | nucleocapsid | GSSRGTSPARMAGNG | 200 | 214 | Positive |
| SARS-CoV-2 | nucleocapsid | SSRGTSPARMAGNGG | 201 | 215 | Negative |
| SARS-CoV-2 | nucleocapsid | GTSPARMAGNGGDAA | 204 | 218 | Negative |
| SARS-CoV-2 | nucleocapsid | NQLESKMSGKGQQQQ | 228 | 242 | Positive |
| SARS-CoV-2 | nucleocapsid | ESKMSGKGQQQQGQT | 231 | 245 | Overlapping |
| SARS-CoV-2 | nucleocapsid | QGQTVTKKSAAEASK | 242 | 256 | Positive |
| SARS-CoV-2 | nucleocapsid | GQTVTKKSAAEASKK | 243 | 257 | Positive |
| SARS-CoV-2 | nucleocapsid | QTVTKKSAAEASKKP | 244 | 258 | Positive |

| | | | | | |
|---|---|---|---|---|---|
| SARS-CoV-2 | nucleocapsid | TVTKKSAAEASKKPR | 245 | 259 | Positive |
| SARS-CoV-2 | nucleocapsid | KKSAAEASKKPRQKR | 248 | 262 | Positive |
| SARS-CoV-2 | nucleocapsid | EASKKPRQKRTATKA | 253 | 267 | Overlapping |
| SARS-CoV-2 | nucleocapsid | SKKPRQKRTATKAYN | 255 | 269 | Negative |
| SARS-CoV-2 | nucleocapsid | KKPRQKRTATKAYNV | 256 | 270 | Positive |
| SARS-CoV-2 | nucleocapsid | GRRGPEQTQGNFGDQ | 275 | 289 | Overlapping |
| SARS-CoV-2 | nucleocapsid | RRGPEQTQGNFGDQE | 276 | 290 | Negative |
| SARS-CoV-2 | nucleocapsid | RGPEQTQGNFGDQEL | 277 | 291 | Negative |
| SARS-CoV-2 | nucleocapsid | GPEQTQGNFGDQELI | 278 | 292 | Overlapping |
| SARS-CoV-2 | nucleocapsid | PEQTQGNFGDQELIR | 279 | 293 | Overlapping |
| SARS-CoV-2 | nucleocapsid | EQTQGNFGDQELIRQ | 280 | 294 | Overlapping |
| SARS-CoV-2 | nucleocapsid | QGNFGDQELIRQGTD | 283 | 297 | Negative |
| SARS-CoV-2 | nucleocapsid | GNFGDQELIRQGTDY | 284 | 298 | Negative |
| SARS-CoV-2 | nucleocapsid | NFGDQELIRQGTDYK | 285 | 299 | Negative |
| SARS-CoV-2 | nucleocapsid | QELIRQGTDYKHWPQ | 289 | 303 | Positive |
| SARS-CoV-2 | nucleocapsid | ELIRQGTDYKHWPQI | 290 | 304 | Overlapping |
| SARS-CoV-2 | nucleocapsid | GTDYKHWPQIAQFAP | 295 | 309 | Negative |
| SARS-CoV-2 | nucleocapsid | TDYKHWPQIAQFAPS | 296 | 310 | Positive |
| SARS-CoV-2 | nucleocapsid | GMSRIGMEVTPSGTW | 316 | 330 | Overlapping |
| SARS-CoV-2 | nucleocapsid | GMEVTPSGTWLTYTG | 321 | 335 | Overlapping |
| SARS-CoV-2 | nucleocapsid | EVTPSGTWLTYTGAI | 323 | 337 | Overlapping |
| SARS-CoV-2 | nucleocapsid | TYTGAIKLDDKDPNF | 332 | 346 | Positive |
| SARS-CoV-2 | nucleocapsid | GAIKLDDKDPNFKDQ | 335 | 349 | Negative |
| SARS-CoV-2 | nucleocapsid | KLDDKDPNFKDQVIL | 338 | 352 | Positive |
| SARS-CoV-2 | nucleocapsid | LDDKDPNFKDQVILL | 339 | 353 | Positive |
| SARS-CoV-2 | nucleocapsid | KDPNFKDQVILLNKH | 342 | 356 | Negative |
| SARS-CoV-2 | nucleocapsid | NFKDQVILLNKHIDA | 345 | 359 | Positive |
| SARS-CoV-2 | nucleocapsid | NKHIDAYKTFPPTEP | 354 | 368 | Positive |
| SARS-CoV-2 | nucleocapsid | DAYKTFPPTEPKKDK | 358 | 372 | Positive |
| SARS-CoV-2 | nucleocapsid | YKTFPPTEPKKDKKK | 360 | 374 | Positive |
| SARS-CoV-2 | nucleocapsid | KTFPPTEPKKDKKKK | 361 | 375 | Positive |
| SARS-CoV-2 | nucleocapsid | TFPPTEPKKDKKKKA | 362 | 376 | Positive |
| SARS-CoV-2 | nucleocapsid | FPPTEPKKDKKKKAD | 363 | 377 | Positive |
| SARS-CoV-2 | nucleocapsid | PPTEPKKDKKKKADE | 364 | 378 | Negative |
| SARS-CoV-2 | nucleocapsid | PTEPKKDKKKKADET | 365 | 379 | Positive |
| SARS-CoV-2 | nucleocapsid | TEPKKDKKKKADETQ | 366 | 380 | Positive |
| SARS-CoV-2 | nucleocapsid | EPKKDKKKKADETQA | 367 | 381 | Overlapping |
| SARS-CoV-2 | nucleocapsid | ADETQALPQRQKKQQ | 376 | 390 | Positive |
| SARS-CoV-2 | nucleocapsid | DETQALPQRQKKQQT | 377 | 391 | Overlapping |
| SARS-CoV-2 | nucleocapsid | ETQALPQRQKKQQTV | 378 | 392 | Overlapping |
| SARS-CoV-2 | nucleocapsid | KQQTVTLLPAADLDD | 388 | 402 | Negative |
| SARS-CoV-2 | nucleocapsid | QQTVTLLPAADLDDF | 389 | 403 | Negative |
| SARS-CoV-2 | nucleocapsid | QTVTLLPAADLDDFS | 390 | 404 | Negative |
| SARS-CoV-2 | nucleocapsid | TVTLLPAADLDDFSK | 391 | 405 | Positive |
| SARS-CoV-2 | nucleocapsid | VTLLPAADLDDFSKQ | 392 | 406 | Negative |
| SARS-CoV-2 | nucleocapsid | TLLPAADLDDFSKQL | 393 | 407 | Positive |
| SARS-CoV-2 | nucleocapsid | LPAADLDDFSKQLQQ | 395 | 409 | Negative |
| SARS-CoV-2 | nucleocapsid | PAADLDDFSKQLQQS | 396 | 410 | Positive |
| SARS-CoV-2 | nucleocapsid | PAADLDDFSKQLQQS | 396 | 410 | Negative |

| | | | | | |
|---|---|---|---|---|---|
| SARS-CoV-2 | nucleocapsid | AADLDDFSKQLQQSM | 397 | 411 | Negative |
| SARS-CoV-2 | nucleocapsid | SMSSADSTQAGSGSG | 410 | 419 | Positive |
| SARS-CoV-2 | orf1ab | GSGMESLVPGFNEKT | 1 | 12 | Positive |
| SARS-CoV-2 | orf1ab | GSGSGSGMESLVPGF | 1 | 8 | Positive |
| SARS-CoV-2 | orf1ab | GSGSGMESLVPGFNE | 1 | 10 | Negative |
| SARS-CoV-2 | orf1ab | GSGSGSGMESLVPGF | 1 | 8 | Negative |
| SARS-CoV-2 | orf1ab | GMESLVPGFNEKTHV | 1 | 14 | Overlapping |
| SARS-CoV-2 | orf1ab | GSGSGMESLVPGFNE | 1 | 10 | Overlapping |
| SARS-CoV-2 | orf1ab | ESLVPGFNEKTHVQL | 2 | 16 | Overlapping |
| SARS-CoV-2 | orf1ab | EKTHVQLSLPVLQVR | 10 | 24 | Overlapping |
| SARS-CoV-2 | orf1ab | QVRDVLVRGFGDSVE | 22 | 36 | Positive |
| SARS-CoV-2 | orf1ab | RDVLVRGFGDSVEEV | 24 | 38 | Positive |
| SARS-CoV-2 | orf1ab | VRGFGDSVEEVLSEA | 28 | 42 | Positive |
| SARS-CoV-2 | orf1ab | GFGDSVEEVLSEARQ | 30 | 44 | Negative |
| SARS-CoV-2 | orf1ab | GDSVEEVLSEARQHL | 32 | 46 | Overlapping |
| SARS-CoV-2 | orf1ab | SVEEVLSEARQHLKD | 34 | 48 | Overlapping |
| SARS-CoV-2 | orf1ab | EEVLSEARQHLKDGT | 36 | 50 | Overlapping |
| SARS-CoV-2 | orf1ab | QHLKDGTCGLVEVEK | 44 | 58 | Positive |
| SARS-CoV-2 | orf1ab | GLVEVEKGVLPQLEQ | 52 | 66 | Overlapping |
| SARS-CoV-2 | orf1ab | KRSDARTAPHGHVMV | 72 | 86 | Positive |
| SARS-CoV-2 | orf1ab | SDARTAPHGHVMVEL | 74 | 88 | Positive |
| SARS-CoV-2 | orf1ab | ARTAPHGHVMVELVA | 76 | 90 | Positive |
| SARS-CoV-2 | orf1ab | TAPHGHVMVELVAEL | 78 | 92 | Positive |
| SARS-CoV-2 | orf1ab | PHGHVMVELVAELEG | 80 | 94 | Positive |
| SARS-CoV-2 | orf1ab | LVAELEGIQYGRSGE | 88 | 102 | Positive |
| SARS-CoV-2 | orf1ab | GIQYGRSGETLGVLV | 94 | 108 | Negative |
| SARS-CoV-2 | orf1ab | GRSGETLGVLVPHVG | 98 | 112 | Negative |
| SARS-CoV-2 | orf1ab | SGETLGVLVPHVGEI | 100 | 114 | Positive |
| SARS-CoV-2 | orf1ab | ETLGVLVPHVGEIPV | 102 | 116 | Overlapping |
| SARS-CoV-2 | orf1ab | KVLLRKNGNKGAGGH | 120 | 134 | Positive |
| SARS-CoV-2 | orf1ab | NKGAGGHSYGADLKS | 128 | 142 | Positive |
| SARS-CoV-2 | orf1ab | GAGGHSYGADLKSFD | 130 | 144 | Positive |
| SARS-CoV-2 | orf1ab | GGHSYGADLKSFDLG | 132 | 146 | Negative |
| SARS-CoV-2 | orf1ab | ADLKSFDLGDELGTD | 138 | 152 | Negative |
| SARS-CoV-2 | orf1ab | SFDLGDELGTDPYED | 142 | 156 | Overlapping |
| SARS-CoV-2 | orf1ab | DLGDELGTDPYEDFQ | 144 | 158 | Overlapping |
| SARS-CoV-2 | orf1ab | GDELGTDPYEDFQEN | 146 | 160 | Overlapping |
| SARS-CoV-2 | orf1ab | ELGTDPYEDFQENWN | 148 | 162 | Negative |
| SARS-CoV-2 | orf1ab | ELGTDPYEDFQENWN | 148 | 162 | Overlapping |
| SARS-CoV-2 | orf1ab | GTDPYEDFQENWNTK | 150 | 164 | Overlapping |
| SARS-CoV-2 | orf1ab | DPYEDFQENWNTKHS | 152 | 166 | Negative |
| SARS-CoV-2 | orf1ab | DFQENWNTKHSSGVT | 156 | 170 | Negative |
| SARS-CoV-2 | orf1ab | QENWNTKHSSGVTRE | 158 | 172 | Negative |
| SARS-CoV-2 | orf1ab | NWNTKHSSGVTRELM | 160 | 174 | Positive |
| SARS-CoV-2 | orf1ab | NTKHSSGVTRELMRE | 162 | 176 | Overlapping |
| SARS-CoV-2 | orf1ab | KHSSGVTRELMRELN | 164 | 178 | Negative |
| SARS-CoV-2 | orf1ab | GVTRELMRELNGGAY | 168 | 182 | Negative |
| SARS-CoV-2 | orf1ab | TRELMRELNGGAYTR | 170 | 184 | Negative |
| SARS-CoV-2 | orf1ab | ELMRELNGGAYTRYV | 172 | 186 | Negative |

| | | | | | |
|---|---|---|---|---|---|
| SARS-CoV-2 | orf1ab | ELMRELNGGAYTRYV | 172 | 186 | Overlapping |
| SARS-CoV-2 | orf1ab | MRELNGGAYTRYVDN | 174 | 188 | Positive |
| SARS-CoV-2 | orf1ab | ELNGGAYTRYVDNNF | 176 | 190 | Negative |
| SARS-CoV-2 | orf1ab | ELNGGAYTRYVDNNF | 176 | 190 | Overlapping |
| SARS-CoV-2 | orf1ab | GPDGYPLECIKDLLA | 192 | 206 | Positive |
| SARS-CoV-2 | orf1ab | DGYPLECIKDLLARA | 194 | 208 | Positive |
| SARS-CoV-2 | orf1ab | SEQLDFIDTKRGVYC | 216 | 230 | Positive |
| SARS-CoV-2 | orf1ab | IDTKRGVYCCREHEH | 222 | 236 | Positive |
| SARS-CoV-2 | orf1ab | CCREHEHEIAWYTER | 230 | 244 | Overlapping |
| SARS-CoV-2 | orf1ab | REHEHEIAWYTERSE | 232 | 246 | Overlapping |
| SARS-CoV-2 | orf1ab | HEHEIAWYTERSEKS | 234 | 248 | Positive |
| SARS-CoV-2 | orf1ab | TERSEKSYELQTPFE | 242 | 256 | Overlapping |
| SARS-CoV-2 | orf1ab | RSEKSYELQTPFEIK | 244 | 258 | Negative |
| SARS-CoV-2 | orf1ab | EKSYELQTPFEIKLA | 246 | 260 | Overlapping |
| SARS-CoV-2 | orf1ab | SYELQTPFEIKLAKK | 248 | 262 | Overlapping |
| SARS-CoV-2 | orf1ab | ELQTPFEIKLAKKFD | 250 | 264 | Negative |
| SARS-CoV-2 | orf1ab | ELQTPFEIKLAKKFD | 250 | 264 | Overlapping |
| SARS-CoV-2 | orf1ab | PFEIKLAKKFDTFNG | 254 | 268 | Positive |
| SARS-CoV-2 | orf1ab | EIKLAKKFDTFNGEC | 256 | 270 | Overlapping |
| SARS-CoV-2 | orf1ab | GECPNFVFPLNSIIK | 268 | 282 | Negative |
| SARS-CoV-2 | orf1ab | NFVFPLNSIIKTIQP | 272 | 286 | Positive |
| SARS-CoV-2 | orf1ab | KTIQPRVEKKKLDGF | 282 | 296 | Positive |
| SARS-CoV-2 | orf1ab | KKKLDGFMGRIRSVY | 290 | 304 | Negative |
| SARS-CoV-2 | orf1ab | DGFMGRIRSVYPVAS | 294 | 308 | Positive |
| SARS-CoV-2 | orf1ab | NECNQMCLSTLMKCD | 310 | 324 | Positive |
| SARS-CoV-2 | orf1ab | KCDHCGETSWQTGDF | 322 | 336 | Positive |
| SARS-CoV-2 | orf1ab | CGETSWQTGDFVKAT | 326 | 340 | Overlapping |
| SARS-CoV-2 | orf1ab | ETSWQTGDFVKATCE | 328 | 342 | Overlapping |
| SARS-CoV-2 | orf1ab | TCEFCGTENLTKEGA | 340 | 354 | Positive |
| SARS-CoV-2 | orf1ab | EFCGTENLTKEGATT | 342 | 356 | Overlapping |
| SARS-CoV-2 | orf1ab | NLTKEGATTCGYLPQ | 348 | 362 | Positive |
| SARS-CoV-2 | orf1ab | TKEGATTCGYLPQNA | 350 | 364 | Negative |
| SARS-CoV-2 | orf1ab | EGATTCGYLPQNAVV | 352 | 366 | Negative |
| SARS-CoV-2 | orf1ab | EGATTCGYLPQNAVV | 352 | 366 | Overlapping |
| SARS-CoV-2 | orf1ab | QNAVVKIYCPACHNS | 362 | 376 | Positive |
| SARS-CoV-2 | orf1ab | AVVKIYCPACHNSEV | 364 | 378 | Positive |
| SARS-CoV-2 | orf1ab | IYCPACHNSEVGPEH | 368 | 382 | Positive |
| SARS-CoV-2 | orf1ab | HNSEVGPEHSLAEYH | 374 | 388 | Positive |
| SARS-CoV-2 | orf1ab | SEVGPEHSLAEYHNE | 376 | 390 | Positive |
| SARS-CoV-2 | orf1ab | VGPEHSLAEYHNESG | 378 | 392 | Positive |
| SARS-CoV-2 | orf1ab | PEHSLAEYHNESGLK | 380 | 394 | Positive |
| SARS-CoV-2 | orf1ab | EYHNESGLKTILRKG | 386 | 400 | Overlapping |
| SARS-CoV-2 | orf1ab | ESGLKTILRKGGRTI | 390 | 404 | Overlapping |
| SARS-CoV-2 | orf1ab | NIGCNHTGVVGEGSE | 430 | 444 | Negative |
| SARS-CoV-2 | orf1ab | GCNHTGVVGEGSEGL | 432 | 446 | Negative |
| SARS-CoV-2 | orf1ab | NHTGVVGEGSEGLND | 434 | 448 | Overlapping |
| SARS-CoV-2 | orf1ab | TGVVGEGSEGLNDNL | 436 | 450 | Negative |
| SARS-CoV-2 | orf1ab | VVGEGSEGLNDNLLE | 438 | 452 | Overlapping |
| SARS-CoV-2 | orf1ab | GEGSEGLNDNLLEIL | 440 | 454 | Overlapping |

| | | | | | |
|---|---|---|---|---|---|
| SARS-CoV-2 | orf1ab | EGLNDNLLEILQKEK | 444 | 458 | Negative |
| SARS-CoV-2 | orf1ab | EGLNDNLLEILQKEK | 444 | 458 | Overlapping |
| SARS-CoV-2 | orf1ab | EILQKEKVNINIVGD | 452 | 466 | Negative |
| SARS-CoV-2 | orf1ab | EILQKEKVNINIVGD | 452 | 466 | Overlapping |
| SARS-CoV-2 | orf1ab | KVNINIVGDFKLNEE | 458 | 472 | Positive |
| SARS-CoV-2 | orf1ab | NINIVGDFKLNEEIA | 460 | 474 | Positive |
| SARS-CoV-2 | orf1ab | NIVGDFKLNEEIAII | 462 | 476 | Positive |
| SARS-CoV-2 | orf1ab | NEEIAIILASFSAST | 470 | 484 | Overlapping |
| SARS-CoV-2 | orf1ab | ASTSAFVETVKGLDY | 482 | 496 | Negative |
| SARS-CoV-2 | orf1ab | VETVKGLDYKAFKQI | 488 | 502 | Positive |
| SARS-CoV-2 | orf1ab | IVESCGNFKVTKGKA | 502 | 516 | Positive |
| SARS-CoV-2 | orf1ab | ESCGNFKVTKGKAKK | 504 | 518 | Negative |
| SARS-CoV-2 | orf1ab | YAFASEAARVVRSIF | 534 | 548 | Negative |
| SARS-CoV-2 | orf1ab | FASEAARVVRSIFSR | 536 | 550 | Positive |
| SARS-CoV-2 | orf1ab | RTLETAQNSVRVLQK | 550 | 564 | Negative |
| SARS-CoV-2 | orf1ab | LDGISQYSLRLIDAM | 570 | 584 | Negative |
| SARS-CoV-2 | orf1ab | GISQYSLRLIDAMMF | 572 | 586 | Negative |
| SARS-CoV-2 | orf1ab | DAMMFTSDLATNNLV | 582 | 596 | Positive |
| SARS-CoV-2 | orf1ab | SDLATNNLVVMAYIT | 588 | 602 | Positive |
| SARS-CoV-2 | orf1ab | TNNLVVMAYITGGVV | 592 | 606 | Negative |
| SARS-CoV-2 | orf1ab | NLVVMAYITGGVVQL | 594 | 608 | Positive |
| SARS-CoV-2 | orf1ab | VVMAYITGGVVQLTS | 596 | 610 | Positive |
| SARS-CoV-2 | orf1ab | GVVQLTSQWLTNIFG | 604 | 618 | Negative |
| SARS-CoV-2 | orf1ab | GTVYEKLKPVLDWLE | 618 | 632 | Negative |
| SARS-CoV-2 | orf1ab | VYEKLKPVLDWLEEK | 620 | 634 | Negative |
| SARS-CoV-2 | orf1ab | EKLKPVLDWLEEKFK | 622 | 636 | Overlapping |
| SARS-CoV-2 | orf1ab | PVLDWLEEKFKEGVE | 626 | 640 | Overlapping |
| SARS-CoV-2 | orf1ab | WLEEKFKEGVEFLRD | 630 | 644 | Overlapping |
| SARS-CoV-2 | orf1ab | EEKFKEGVEFLRDGW | 632 | 646 | Overlapping |
| SARS-CoV-2 | orf1ab | KEGVEFLRDGWEIVK | 636 | 650 | Overlapping |
| SARS-CoV-2 | orf1ab | GVEFLRDGWEIVKFI | 638 | 652 | Negative |
| SARS-CoV-2 | orf1ab | EFLRDGWEIVKFIST | 640 | 654 | Overlapping |
| SARS-CoV-2 | orf1ab | EIVGGQIVTCAKEIK | 658 | 672 | Negative |
| SARS-CoV-2 | orf1ab | EIKESVQTFFKLVNK | 670 | 684 | Overlapping |
| SARS-CoV-2 | orf1ab | NLGETFVTHSKGLYR | 704 | 718 | Positive |
| SARS-CoV-2 | orf1ab | GETFVTHSKGLYRKC | 706 | 720 | Negative |
| SARS-CoV-2 | orf1ab | REETGLLMPLKAPKE | 724 | 738 | Positive |
| SARS-CoV-2 | orf1ab | ETGLLMPLKAPKEII | 726 | 740 | Negative |
| SARS-CoV-2 | orf1ab | GLLMPLKAPKEIIFL | 728 | 742 | Negative |
| SARS-CoV-2 | orf1ab | LMPLKAPKEIIFLEG | 730 | 744 | Positive |
| SARS-CoV-2 | orf1ab | PLKAPKEIIFLEGET | 732 | 746 | Positive |
| SARS-CoV-2 | orf1ab | PKEIIFLEGETLPTE | 736 | 750 | Overlapping |
| SARS-CoV-2 | orf1ab | EIIFLEGETLPTEVL | 738 | 752 | Negative |
| SARS-CoV-2 | orf1ab | EIIFLEGETLPTEVL | 738 | 752 | Overlapping |
| SARS-CoV-2 | orf1ab | IFLEGETLPTEVLTE | 740 | 754 | Negative |
| SARS-CoV-2 | orf1ab | LEGETLPTEVLTEEV | 742 | 756 | Positive |
| SARS-CoV-2 | orf1ab | GETLPTEVLTEEVVL | 744 | 758 | Overlapping |
| SARS-CoV-2 | orf1ab | TLPTEVLTEEVVLKT | 746 | 760 | Negative |
| SARS-CoV-2 | orf1ab | TLPTEVLTEEVVLKT | 746 | 760 | Overlapping |

| | | | | | |
|---|---|---|---|---|---|
| SARS-CoV-2 | orf1ab | PTEVLTEEVVLKTGD | 748 | 762 | Positive |
| SARS-CoV-2 | orf1ab | PTEVLTEEVVLKTGD | 748 | 762 | Overlapping |
| SARS-CoV-2 | orf1ab | EVLTEEVVLKTGDLQ | 750 | 764 | Overlapping |
| SARS-CoV-2 | orf1ab | LTEEVVLKTGDLQPL | 752 | 766 | Overlapping |
| SARS-CoV-2 | orf1ab | EEVVLKTGDLQPLEQ | 754 | 768 | Overlapping |
| SARS-CoV-2 | orf1ab | TGDLQPLEQPTSEAV | 760 | 774 | Positive |
| SARS-CoV-2 | orf1ab | DLQPLEQPTSEAVEA | 762 | 776 | Positive |
| SARS-CoV-2 | orf1ab | QPLEQPTSEAVEAPL | 764 | 778 | Positive |
| SARS-CoV-2 | orf1ab | LEQPTSEAVEAPLVG | 766 | 780 | Positive |
| SARS-CoV-2 | orf1ab | QPTSEAVEAPLVGTP | 768 | 782 | Positive |
| SARS-CoV-2 | orf1ab | TSEAVEAPLVGTPVC | 770 | 784 | Positive |
| SARS-CoV-2 | orf1ab | EAVEAPLVGTPVCIN | 772 | 786 | Overlapping |
| SARS-CoV-2 | orf1ab | LVGTPVCINGLMLLE | 778 | 792 | Negative |
| SARS-CoV-2 | orf1ab | NGLMLLEIKDTEKYC | 786 | 800 | Positive |
| SARS-CoV-2 | orf1ab | LLEIKDTEKYCALAP | 790 | 804 | Positive |
| SARS-CoV-2 | orf1ab | EIKDTEKYCALAPNM | 792 | 806 | Overlapping |
| SARS-CoV-2 | orf1ab | TEKYCALAPNMMVTN | 796 | 810 | Positive |
| SARS-CoV-2 | orf1ab | KYCALAPNMMVTNNT | 798 | 812 | Negative |
| SARS-CoV-2 | orf1ab | GAPTKVTFGDDTVIE | 818 | 832 | Positive |
| SARS-CoV-2 | orf1ab | PTKVTFGDDTVIEVQ | 820 | 834 | Positive |
| SARS-CoV-2 | orf1ab | GDDTVIEVQGYKSVN | 826 | 840 | Negative |
| SARS-CoV-2 | orf1ab | VIEVQGYKSVNITFE | 830 | 844 | Positive |
| SARS-CoV-2 | orf1ab | EVQGYKSVNITFELD | 832 | 846 | Negative |
| SARS-CoV-2 | orf1ab | EVQGYKSVNITFELD | 832 | 846 | Overlapping |
| SARS-CoV-2 | orf1ab | NITFELDERIDKVLN | 840 | 854 | Positive |
| SARS-CoV-2 | orf1ab | TFELDERIDKVLNEK | 842 | 856 | Overlapping |
| SARS-CoV-2 | orf1ab | ELDERIDKVLNEKCS | 844 | 858 | Negative |
| SARS-CoV-2 | orf1ab | ELDERIDKVLNEKCS | 844 | 858 | Overlapping |
| SARS-CoV-2 | orf1ab | NEKCSAYTVELGTEV | 854 | 868 | Positive |
| SARS-CoV-2 | orf1ab | YTVELGTEVNEFACV | 860 | 874 | Negative |
| SARS-CoV-2 | orf1ab | TEVNEFACVVADAVI | 866 | 880 | Overlapping |
| SARS-CoV-2 | orf1ab | EFACVVADAVIKTLQ | 870 | 884 | Overlapping |
| SARS-CoV-2 | orf1ab | ADAVIKTLQPVSELL | 876 | 890 | Negative |
| SARS-CoV-2 | orf1ab | TLQPVSELLTPLGID | 882 | 896 | Negative |
| SARS-CoV-2 | orf1ab | ELLTPLGIDLDEWSM | 888 | 902 | Negative |
| SARS-CoV-2 | orf1ab | ELLTPLGIDLDEWSM | 888 | 902 | Overlapping |
| SARS-CoV-2 | orf1ab | PLGIDLDEWSMATYY | 892 | 906 | Overlapping |
| SARS-CoV-2 | orf1ab | GIDLDEWSMATYYLF | 894 | 908 | Negative |
| SARS-CoV-2 | orf1ab | DLDEWSMATYYLFDE | 896 | 910 | Positive |
| SARS-CoV-2 | orf1ab | DEWSMATYYLFDESG | 898 | 912 | Positive |
| SARS-CoV-2 | orf1ab | WSMATYYLFDESGEF | 900 | 914 | Positive |
| SARS-CoV-2 | orf1ab | FDESGEFKLASHMYC | 908 | 922 | Negative |
| SARS-CoV-2 | orf1ab | FDESGEFKLASHMYC | 908 | 922 | Overlapping |
| SARS-CoV-2 | orf1ab | ESGEFKLASHMYCSF | 910 | 924 | Overlapping |
| SARS-CoV-2 | orf1ab | SHMYCSFYPPDEDEE | 918 | 932 | Positive |
| SARS-CoV-2 | orf1ab | MYCSFYPPDEDEEEG | 920 | 934 | Positive |
| SARS-CoV-2 | orf1ab | CSFYPPDEDEEEGDC | 922 | 936 | Overlapping |
| SARS-CoV-2 | orf1ab | FYPPDEDEEEGDCEE | 924 | 938 | Overlapping |
| SARS-CoV-2 | orf1ab | PPDEDEEEGDCEEEE | 926 | 940 | Overlapping |

| | | | | | |
|---|---|---|---|---|---|
| SARS-CoV-2 | orf1ab | DEDEEEGDCEEEEFE | 928 | 942 | Positive |
| SARS-CoV-2 | orf1ab | DEDEEEGDCEEEEFE | 928 | 942 | Overlapping |
| SARS-CoV-2 | orf1ab | DEEEGDCEEEEFEPS | 930 | 944 | Overlapping |
| SARS-CoV-2 | orf1ab | EEGDCEEEEFEPSTQ | 932 | 946 | Overlapping |
| SARS-CoV-2 | orf1ab | GDCEEEEFEPSTQYE | 934 | 948 | Overlapping |
| SARS-CoV-2 | orf1ab | CEEEEFEPSTQYEYG | 936 | 950 | Overlapping |
| SARS-CoV-2 | orf1ab | EEEFEPSTQYEYGTE | 938 | 952 | Overlapping |
| SARS-CoV-2 | orf1ab | EFEPSTQYEYGTEDD | 940 | 954 | Overlapping |
| SARS-CoV-2 | orf1ab | EPSTQYEYGTEDDYQ | 942 | 956 | Positive |
| SARS-CoV-2 | orf1ab | EYGTEDDYQGKPLEF | 948 | 962 | Overlapping |
| SARS-CoV-2 | orf1ab | GTEDDYQGKPLEFGA | 950 | 964 | Overlapping |
| SARS-CoV-2 | orf1ab | EDDYQGKPLEFGATS | 952 | 966 | Positive |
| SARS-CoV-2 | orf1ab | EDDYQGKPLEFGATS | 952 | 966 | Overlapping |
| SARS-CoV-2 | orf1ab | FGATSAALQPEEEQE | 962 | 976 | Overlapping |
| SARS-CoV-2 | orf1ab | ATSAALQPEEEQEED | 964 | 978 | Positive |
| SARS-CoV-2 | orf1ab | ATSAALQPEEEQEED | 964 | 978 | Overlapping |
| SARS-CoV-2 | orf1ab | SAALQPEEEQEEDWL | 966 | 980 | Overlapping |
| SARS-CoV-2 | orf1ab | ALQPEEEQEEDWLDD | 968 | 982 | Overlapping |
| SARS-CoV-2 | orf1ab | QPEEEQEEDWLDDDS | 970 | 984 | Overlapping |
| SARS-CoV-2 | orf1ab | EEEQEEDWLDDDSQQ | 972 | 986 | Overlapping |
| SARS-CoV-2 | orf1ab | EQEEDWLDDDSQQTV | 974 | 988 | Positive |
| SARS-CoV-2 | orf1ab | EQEEDWLDDDSQQTV | 974 | 988 | Overlapping |
| SARS-CoV-2 | orf1ab | EEDWLDDDSQQTVGQ | 976 | 990 | Overlapping |
| SARS-CoV-2 | orf1ab | DWLDDDSQQTVGQQD | 978 | 992 | Positive |
| SARS-CoV-2 | orf1ab | DDSQQTVGQQDGSED | 982 | 996 | Positive |
| SARS-CoV-2 | orf1ab | VGQQDGSEDNQTTTI | 988 | 1002 | Positive |
| SARS-CoV-2 | orf1ab | DGSEDNQTTTIQTIV | 992 | 1006 | Overlapping |
| SARS-CoV-2 | orf1ab | SEDNQTTTIQTIVEV | 994 | 1008 | Positive |
| SARS-CoV-2 | orf1ab | IQTIVEVQPQLEMEL | 1002 | 1016 | Positive |
| SARS-CoV-2 | orf1ab | TIVEVQPQLEMELTP | 1004 | 1018 | Positive |
| SARS-CoV-2 | orf1ab | VQPQLEMELTPVVQT | 1008 | 1022 | Positive |
| SARS-CoV-2 | orf1ab | PQLEMELTPVVQTIE | 1010 | 1024 | Positive |
| SARS-CoV-2 | orf1ab | LEMELTPVVQTIEVN | 1012 | 1026 | Negative |
| SARS-CoV-2 | orf1ab | LTPVVQTIEVNSFSG | 1016 | 1030 | Positive |
| SARS-CoV-2 | orf1ab | PVVQTIEVNSFSGYL | 1018 | 1032 | Overlapping |
| SARS-CoV-2 | orf1ab | VQTIEVNSFSGYLKL | 1020 | 1034 | Positive |
| SARS-CoV-2 | orf1ab | TIEVNSFSGYLKLTD | 1022 | 1036 | Positive |
| SARS-CoV-2 | orf1ab | EVNSFSGYLKLTDNV | 1024 | 1038 | Positive |
| SARS-CoV-2 | orf1ab | LTDNVYIKNADIVEE | 1034 | 1048 | Overlapping |
| SARS-CoV-2 | orf1ab | DNVYIKNADIVEEAK | 1036 | 1050 | Positive |
| SARS-CoV-2 | orf1ab | VYIKNADIVEEAKKV | 1038 | 1052 | Positive |
| SARS-CoV-2 | orf1ab | NADIVEEAKKVKPTV | 1042 | 1056 | Overlapping |
| SARS-CoV-2 | orf1ab | EAKKVKPTVVVNAAN | 1048 | 1062 | Overlapping |
| SARS-CoV-2 | orf1ab | VKPTVVVNAANVYLK | 1052 | 1066 | Positive |
| SARS-CoV-2 | orf1ab | KHGGGVAGALNKATN | 1066 | 1080 | Negative |
| SARS-CoV-2 | orf1ab | GGGVAGALNKATNNA | 1068 | 1082 | Negative |
| SARS-CoV-2 | orf1ab | GVAGALNKATNNAMQ | 1070 | 1084 | Negative |
| SARS-CoV-2 | orf1ab | NKATNNAMQVESDDY | 1076 | 1090 | Positive |
| SARS-CoV-2 | orf1ab | NNAMQVESDDYIATN | 1080 | 1094 | Positive |

| | | | | | |
|---|---|---|---|---|---|
| SARS-CoV-2 | orf1ab | AMQVESDDYIATNGP | 1082 | 1096 | Negative |
| SARS-CoV-2 | orf1ab | QVESDDYIATNGPLK | 1084 | 1098 | Overlapping |
| SARS-CoV-2 | orf1ab | ESDDYIATNGPLKVG | 1086 | 1100 | Positive |
| SARS-CoV-2 | orf1ab | ESDDYIATNGPLKVG | 1086 | 1100 | Overlapping |
| SARS-CoV-2 | orf1ab | KVGGSCVLSGHNLAK | 1098 | 1112 | Negative |
| SARS-CoV-2 | orf1ab | GGSCVLSGHNLAKHC | 1100 | 1114 | Positive |
| SARS-CoV-2 | orf1ab | SCVLSGHNLAKHCLH | 1102 | 1116 | Positive |
| SARS-CoV-2 | orf1ab | KGEDIQLLKSAYENF | 1124 | 1138 | Overlapping |
| SARS-CoV-2 | orf1ab | EDIQLLKSAYENFNQ | 1126 | 1140 | Overlapping |
| SARS-CoV-2 | orf1ab | IQLLKSAYENFNQHE | 1128 | 1142 | Positive |
| SARS-CoV-2 | orf1ab | ENFNQHEVLLAPLLS | 1136 | 1150 | Overlapping |
| SARS-CoV-2 | orf1ab | QHEVLLAPLLSAGIF | 1140 | 1154 | Positive |
| SARS-CoV-2 | orf1ab | EVLLAPLLSAGIFGA | 1142 | 1156 | Negative |
| SARS-CoV-2 | orf1ab | EVLLAPLLSAGIFGA | 1142 | 1156 | Overlapping |
| SARS-CoV-2 | orf1ab | ADPIHSLRVCVDTVR | 1156 | 1170 | Positive |
| SARS-CoV-2 | orf1ab | PIHSLRVCVDTVRTN | 1158 | 1172 | Overlapping |
| SARS-CoV-2 | orf1ab | EMKSEKQVEQKIAEI | 1192 | 1206 | Overlapping |
| SARS-CoV-2 | orf1ab | KSEKQVEQKIAEIPK | 1194 | 1208 | Positive |
| SARS-CoV-2 | orf1ab | EKQVEQKIAEIPKEE | 1196 | 1210 | Overlapping |
| SARS-CoV-2 | orf1ab | EQKIAEIPKEEVKPF | 1200 | 1214 | Overlapping |
| SARS-CoV-2 | orf1ab | KIAEIPKEEVKPFIT | 1202 | 1216 | Positive |
| SARS-CoV-2 | orf1ab | AEIPKEEVKPFITES | 1204 | 1218 | Positive |
| SARS-CoV-2 | orf1ab | IPKEEVKPFITESKP | 1206 | 1220 | Positive |
| SARS-CoV-2 | orf1ab | IPKEEVKPFITESKP | 1206 | 1220 | Negative |
| SARS-CoV-2 | orf1ab | KEEVKPFITESKPSV | 1208 | 1222 | Overlapping |
| SARS-CoV-2 | orf1ab | EVKPFITESKPSVEQ | 1210 | 1224 | Negative |
| SARS-CoV-2 | orf1ab | SKPSVEQRKQDDKKI | 1218 | 1232 | Negative |
| SARS-CoV-2 | orf1ab | ACVEEVTTTLEETKF | 1234 | 1248 | Positive |
| SARS-CoV-2 | orf1ab | VEEVTTTLEETKFLT | 1236 | 1250 | Overlapping |
| SARS-CoV-2 | orf1ab | EVTTTLEETKFLTEN | 1238 | 1252 | Negative |
| SARS-CoV-2 | orf1ab | EVTTTLEETKFLTEN | 1238 | 1252 | Overlapping |
| SARS-CoV-2 | orf1ab | TLEETKFLTENLLLY | 1242 | 1256 | Positive |
| SARS-CoV-2 | orf1ab | EETKFLTENLLLYID | 1244 | 1258 | Overlapping |
| SARS-CoV-2 | orf1ab | TENLLLYIDINGNLH | 1250 | 1264 | Positive |
| SARS-CoV-2 | orf1ab | NLLLYIDINGNLHPD | 1252 | 1266 | Positive |
| SARS-CoV-2 | orf1ab | NGNLHPDSATLVSDI | 1260 | 1274 | Positive |
| SARS-CoV-2 | orf1ab | NLHPDSATLVSDIDI | 1262 | 1276 | Positive |
| SARS-CoV-2 | orf1ab | SDIDITFLKKDAPYI | 1272 | 1286 | Positive |
| SARS-CoV-2 | orf1ab | IVGDVVQEGVLTAVV | 1286 | 1300 | Positive |
| SARS-CoV-2 | orf1ab | GDVVQEGVLTAVVIP | 1288 | 1302 | Negative |
| SARS-CoV-2 | orf1ab | GVLTAVVIPTKKAGG | 1294 | 1308 | Negative |
| SARS-CoV-2 | orf1ab | GTTEMLAKALRKVPT | 1308 | 1322 | Negative |
| SARS-CoV-2 | orf1ab | TEMLAKALRKVPTDN | 1310 | 1324 | Overlapping |
| SARS-CoV-2 | orf1ab | TDNYITTYPGQGLNG | 1322 | 1336 | Positive |
| SARS-CoV-2 | orf1ab | PGQGLNGYTVEEAKT | 1330 | 1344 | Positive |
| SARS-CoV-2 | orf1ab | GYTVEEAKTVLKKCK | 1336 | 1350 | Negative |
| SARS-CoV-2 | orf1ab | TVEEAKTVLKKCKSA | 1338 | 1352 | Overlapping |
| SARS-CoV-2 | orf1ab | EEAKTVLKKCKSAFY | 1340 | 1354 | Overlapping |
| SARS-CoV-2 | orf1ab | AFYILPSIISNEKQE | 1352 | 1366 | Negative |

| | | | | | |
|---|---|---|---|---|---|
| SARS-CoV-2 | orf1ab | KQEILGTVSWNLREM | 1364 | 1378 | Positive |
| SARS-CoV-2 | orf1ab | EILGTVSWNLREMLA | 1366 | 1380 | Negative |
| SARS-CoV-2 | orf1ab | EILGTVSWNLREMLA | 1366 | 1380 | Overlapping |
| SARS-CoV-2 | orf1ab | TVSWNLREMLAHAEE | 1370 | 1384 | Positive |
| SARS-CoV-2 | orf1ab | NLREMLAHAEETRKL | 1374 | 1388 | Positive |
| SARS-CoV-2 | orf1ab | REMLAHAEETRKLMP | 1376 | 1390 | Positive |
| SARS-CoV-2 | orf1ab | AHAEETRKLMPVCVE | 1380 | 1394 | Positive |
| SARS-CoV-2 | orf1ab | ETRKLMPVCVETKAI | 1384 | 1398 | Overlapping |
| SARS-CoV-2 | orf1ab | PVCVETKAIVSTIQR | 1390 | 1404 | Overlapping |
| SARS-CoV-2 | orf1ab | ETKAIVSTIQRKYKG | 1394 | 1408 | Positive |
| SARS-CoV-2 | orf1ab | KAIVSTIQRKYKGIK | 1396 | 1410 | Positive |
| SARS-CoV-2 | orf1ab | IQRKYKGIKIQEGVV | 1402 | 1416 | Negative |
| SARS-CoV-2 | orf1ab | RKYKGIKIQEGVVDY | 1404 | 1418 | Negative |
| SARS-CoV-2 | orf1ab | YKGIKIQEGVVDYGA | 1406 | 1420 | Negative |
| SARS-CoV-2 | orf1ab | GIKIQEGVVDYGARF | 1408 | 1422 | Negative |
| SARS-CoV-2 | orf1ab | GVVDYGARFYFYTSK | 1414 | 1428 | Negative |
| SARS-CoV-2 | orf1ab | TVASLINTLNDLNET | 1430 | 1444 | Negative |
| SARS-CoV-2 | orf1ab | NTLNDLNETLVTMPL | 1436 | 1450 | Overlapping |
| SARS-CoV-2 | orf1ab | NETLVTMPLGYVTHG | 1442 | 1456 | Positive |
| SARS-CoV-2 | orf1ab | TLVTMPLGYVTHGLN | 1444 | 1458 | Positive |
| SARS-CoV-2 | orf1ab | VTMPLGYVTHGLNLE | 1446 | 1460 | Positive |
| SARS-CoV-2 | orf1ab | LGYVTHGLNLEEAAR | 1450 | 1464 | Positive |
| SARS-CoV-2 | orf1ab | THGLNLEEAARYMRS | 1454 | 1468 | Negative |
| SARS-CoV-2 | orf1ab | GLNLEEAARYMRSLK | 1456 | 1470 | Negative |
| SARS-CoV-2 | orf1ab | NLEEAARYMRSLKVP | 1458 | 1472 | Positive |
| SARS-CoV-2 | orf1ab | EEAARYMRSLKVPAT | 1460 | 1474 | Overlapping |
| SARS-CoV-2 | orf1ab | LTSSSKTPEEHFIET | 1490 | 1504 | Positive |
| SARS-CoV-2 | orf1ab | SSSKTPEEHFIETIS | 1492 | 1506 | Positive |
| SARS-CoV-2 | orf1ab | SKTPEEHFIETISLA | 1494 | 1508 | Negative |
| SARS-CoV-2 | orf1ab | TPEEHFIETISLAGS | 1496 | 1510 | Overlapping |
| SARS-CoV-2 | orf1ab | EEHFIETISLAGSYK | 1498 | 1512 | Overlapping |
| SARS-CoV-2 | orf1ab | IETISLAGSYKDWSY | 1502 | 1516 | Positive |
| SARS-CoV-2 | orf1ab | SYKDWSYSGQSTQLG | 1510 | 1524 | Positive |
| SARS-CoV-2 | orf1ab | KDWSYSGQSTQLGIE | 1512 | 1526 | Negative |
| SARS-CoV-2 | orf1ab | QLGIEFLKRGDKSVY | 1522 | 1536 | Negative |
| SARS-CoV-2 | orf1ab | GIEFLKRGDKSVYYT | 1524 | 1538 | Overlapping |
| SARS-CoV-2 | orf1ab | EFLKRGDKSVYYTSN | 1526 | 1540 | Overlapping |
| SARS-CoV-2 | orf1ab | NPTTFHLDGEVITFD | 1540 | 1554 | Positive |
| SARS-CoV-2 | orf1ab | FHLDGEVITFDNLKT | 1544 | 1558 | Negative |
| SARS-CoV-2 | orf1ab | GEVITFDNLKTLLSL | 1548 | 1562 | Negative |
| SARS-CoV-2 | orf1ab | TFDNLKTLLSLREVR | 1552 | 1566 | Negative |
| SARS-CoV-2 | orf1ab | EVRTIKVFTTVDNIN | 1564 | 1578 | Overlapping |
| SARS-CoV-2 | orf1ab | NINLHTQVVDMSMTY | 1576 | 1590 | Positive |
| SARS-CoV-2 | orf1ab | NLHTQVVDMSMTYGQ | 1578 | 1592 | Positive |
| SARS-CoV-2 | orf1ab | PTYLDGADVTKIKPH | 1596 | 1610 | Overlapping |
| SARS-CoV-2 | orf1ab | DGADVTKIKPHNSHE | 1600 | 1614 | Overlapping |
| SARS-CoV-2 | orf1ab | VTKIKPHNSHEGKTF | 1604 | 1618 | Positive |
| SARS-CoV-2 | orf1ab | KIKPHNSHEGKTFYV | 1606 | 1620 | Positive |
| SARS-CoV-2 | orf1ab | KPHNSHEGKTFYVLP | 1608 | 1622 | Positive |

| | | | | | |
|---|---|---|---|---|---|
| SARS-CoV-2 | orf1ab | HNSHEGKTFYVLPND | 1610 | 1624 | Positive |
| SARS-CoV-2 | orf1ab | SHEGKTFYVLPNDDT | 1612 | 1626 | Overlapping |
| SARS-CoV-2 | orf1ab | EGKTFYVLPNDDTLR | 1614 | 1628 | Negative |
| SARS-CoV-2 | orf1ab | EGKTFYVLPNDDTLR | 1614 | 1628 | Overlapping |
| SARS-CoV-2 | orf1ab | VLPNDDTLRVEAFEY | 1620 | 1634 | Positive |
| SARS-CoV-2 | orf1ab | DDTLRVEAFEYYHTT | 1624 | 1638 | Negative |
| SARS-CoV-2 | orf1ab | TLRVEAFEYYHTTDP | 1626 | 1640 | Positive |
| SARS-CoV-2 | orf1ab | RVEAFEYYHTTDPSF | 1628 | 1642 | Negative |
| SARS-CoV-2 | orf1ab | EAFEYYHTTDPSFLG | 1630 | 1644 | Overlapping |
| SARS-CoV-2 | orf1ab | TDPSFLGRYMSALNH | 1638 | 1652 | Positive |
| SARS-CoV-2 | orf1ab | ADNNCYLATALLTLQ | 1670 | 1684 | Negative |
| SARS-CoV-2 | orf1ab | NNCYLATALLTLQQI | 1672 | 1686 | Positive |
| SARS-CoV-2 | orf1ab | AGEAANFCALILAYC | 1704 | 1718 | Negative |
| SARS-CoV-2 | orf1ab | EAANFCALILAYCNK | 1706 | 1720 | Overlapping |
| SARS-CoV-2 | orf1ab | ILAYCNKTVGELGDV | 1714 | 1728 | Positive |
| SARS-CoV-2 | orf1ab | KTVGELGDVRETMSY | 1720 | 1734 | Positive |
| SARS-CoV-2 | orf1ab | VGELGDVRETMSYLF | 1722 | 1736 | Positive |
| SARS-CoV-2 | orf1ab | ELGDVRETMSYLFQH | 1724 | 1738 | Negative |
| SARS-CoV-2 | orf1ab | ELGDVRETMSYLFQH | 1724 | 1738 | Overlapping |
| SARS-CoV-2 | orf1ab | GDVRETMSYLFQHAN | 1726 | 1740 | Positive |
| SARS-CoV-2 | orf1ab | ETMSYLFQHANLDSC | 1730 | 1744 | Overlapping |
| SARS-CoV-2 | orf1ab | NLDSCKRVLNVVCKT | 1740 | 1754 | Positive |
| SARS-CoV-2 | orf1ab | CKTCGQQQTTLKGVE | 1752 | 1766 | Negative |
| SARS-CoV-2 | orf1ab | GQQQTTLKGVEAVMY | 1756 | 1770 | Negative |
| SARS-CoV-2 | orf1ab | GVEAVMYMGTLSYEQ | 1764 | 1778 | Negative |
| SARS-CoV-2 | orf1ab | EAVMYMGTLSYEQFK | 1766 | 1780 | Overlapping |
| SARS-CoV-2 | orf1ab | YMGTLSYEQFKKGVQ | 1770 | 1784 | Negative |
| SARS-CoV-2 | orf1ab | GTLSYEQFKKGVQIP | 1772 | 1786 | Negative |
| SARS-CoV-2 | orf1ab | KKGVQIPCTCGKQAT | 1780 | 1794 | Negative |
| SARS-CoV-2 | orf1ab | GVQIPCTCGKQATKY | 1782 | 1796 | Negative |
| SARS-CoV-2 | orf1ab | ASEYTGNYQCGHYKH | 1824 | 1838 | Positive |
| SARS-CoV-2 | orf1ab | EYTGNYQCGHYKHIT | 1826 | 1840 | Overlapping |
| SARS-CoV-2 | orf1ab | LLTKSSEYKGPITDV | 1852 | 1866 | Positive |
| SARS-CoV-2 | orf1ab | SSEYKGPITDVFYKE | 1856 | 1870 | Negative |
| SARS-CoV-2 | orf1ab | EYKGPITDVFYKENS | 1858 | 1872 | Overlapping |
| SARS-CoV-2 | orf1ab | KGPITDVFYKENSYT | 1860 | 1874 | Positive |
| SARS-CoV-2 | orf1ab | PITDVFYKENSYTTT | 1862 | 1876 | Overlapping |
| SARS-CoV-2 | orf1ab | TDVFYKENSYTTTIK | 1864 | 1878 | Positive |
| SARS-CoV-2 | orf1ab | YKENSYTTTIKPVTY | 1868 | 1882 | Positive |
| SARS-CoV-2 | orf1ab | ENSYTTTIKPVTYKL | 1870 | 1884 | Overlapping |
| SARS-CoV-2 | orf1ab | TTTIKPVTYKLDGVV | 1874 | 1888 | Negative |
| SARS-CoV-2 | orf1ab | VTYKLDGVVCTEIDP | 1880 | 1894 | Positive |
| SARS-CoV-2 | orf1ab | LDGVVCTEIDPKLDN | 1884 | 1898 | Negative |
| SARS-CoV-2 | orf1ab | GVVCTEIDPKLDNYY | 1886 | 1900 | Negative |
| SARS-CoV-2 | orf1ab | VCTEIDPKLDNYYKK | 1888 | 1902 | Positive |
| SARS-CoV-2 | orf1ab | TEIDPKLDNYYKKDN | 1890 | 1904 | Positive |
| SARS-CoV-2 | orf1ab | TEIDPKLDNYYKKDN | 1890 | 1904 | Negative |
| SARS-CoV-2 | orf1ab | NSYFTEQPIDLVPNQ | 1904 | 1918 | Positive |
| SARS-CoV-2 | orf1ab | TEQPIDLVPNQPYPN | 1908 | 1922 | Positive |

| | | | | | |
|---|---|---|---|---|---|
| SARS-CoV-2 | orf1ab | QPIDLVPNQPYPNAS | 1910 | 1924 | Positive |
| SARS-CoV-2 | orf1ab | NASFDNFKFVCDNIK | 1922 | 1936 | Positive |
| SARS-CoV-2 | orf1ab | KFADDLNQLTGYKKP | 1936 | 1950 | Positive |
| SARS-CoV-2 | orf1ab | ADDLNQLTGYKKPAS | 1938 | 1952 | Positive |
| SARS-CoV-2 | orf1ab | NQLTGYKKPASRELK | 1942 | 1956 | Positive |
| SARS-CoV-2 | orf1ab | SRELKVTFFPDLNGD | 1952 | 1966 | Negative |
| SARS-CoV-2 | orf1ab | ELKVTFFPDLNGDVV | 1954 | 1968 | Negative |
| SARS-CoV-2 | orf1ab | ELKVTFFPDLNGDVV | 1954 | 1968 | Overlapping |
| SARS-CoV-2 | orf1ab | FPDLNGDVVAIDYKH | 1960 | 1974 | Positive |
| SARS-CoV-2 | orf1ab | KKGAKLLHKPIVWHV | 1980 | 1994 | Positive |
| SARS-CoV-2 | orf1ab | KLLHKPIVWHVNNAT | 1984 | 1998 | Positive |
| SARS-CoV-2 | orf1ab | NATNKATYKPNTWCI | 1996 | 2010 | Positive |
| SARS-CoV-2 | orf1ab | NATNKATYKPNTWCI | 1996 | 2010 | Negative |
| SARS-CoV-2 | orf1ab | WSTKPVETSNSFDVL | 2014 | 2028 | Positive |
| SARS-CoV-2 | orf1ab | TKPVETSNSFDVLKS | 2016 | 2030 | Positive |
| SARS-CoV-2 | orf1ab | PVETSNSFDVLKSED | 2018 | 2032 | Positive |
| SARS-CoV-2 | orf1ab | PVETSNSFDVLKSED | 2018 | 2032 | Overlapping |
| SARS-CoV-2 | orf1ab | ETSNSFDVLKSEDAQ | 2020 | 2034 | Overlapping |
| SARS-CoV-2 | orf1ab | SNSFDVLKSEDAQGM | 2022 | 2036 | Negative |
| SARS-CoV-2 | orf1ab | SFDVLKSEDAQGMDN | 2024 | 2038 | Negative |
| SARS-CoV-2 | orf1ab | LKSEDAQGMDNLACE | 2028 | 2042 | Negative |
| SARS-CoV-2 | orf1ab | QGMDNLACEDLKPVS | 2034 | 2048 | Negative |
| SARS-CoV-2 | orf1ab | MDNLACEDLKPVSEE | 2036 | 2050 | Positive |
| SARS-CoV-2 | orf1ab | NLACEDLKPVSEEVV | 2038 | 2052 | Positive |
| SARS-CoV-2 | orf1ab | ACEDLKPVSEEVVEN | 2040 | 2054 | Positive |
| SARS-CoV-2 | orf1ab | EDLKPVSEEVVENPT | 2042 | 2056 | Overlapping |
| SARS-CoV-2 | orf1ab | LKPVSEEVVENPTIQ | 2044 | 2058 | Overlapping |
| SARS-CoV-2 | orf1ab | PVSEEVVENPTIQKD | 2046 | 2060 | Positive |
| SARS-CoV-2 | orf1ab | PVSEEVVENPTIQKD | 2046 | 2060 | Overlapping |
| SARS-CoV-2 | orf1ab | SEEVVENPTIQKDVL | 2048 | 2062 | Overlapping |
| SARS-CoV-2 | orf1ab | EVVENPTIQKDVLEC | 2050 | 2064 | Negative |
| SARS-CoV-2 | orf1ab | EVVENPTIQKDVLEC | 2050 | 2064 | Overlapping |
| SARS-CoV-2 | orf1ab | EVVGDIILKPANNSL | 2070 | 2084 | Negative |
| SARS-CoV-2 | orf1ab | EVVGDIILKPANNSL | 2070 | 2084 | Overlapping |
| SARS-CoV-2 | orf1ab | ILKPANNSLKITEEV | 2076 | 2090 | Negative |
| SARS-CoV-2 | orf1ab | NSLKITEEVGHTDLM | 2082 | 2096 | Positive |
| SARS-CoV-2 | orf1ab | LKITEEVGHTDLMAA | 2084 | 2098 | Negative |
| SARS-CoV-2 | orf1ab | ITEEVGHTDLMAAYV | 2086 | 2100 | Overlapping |
| SARS-CoV-2 | orf1ab | EEVGHTDLMAAYVDN | 2088 | 2102 | Overlapping |
| SARS-CoV-2 | orf1ab | HTDLMAAYVDNSSLT | 2092 | 2106 | Positive |
| SARS-CoV-2 | orf1ab | NSSLTIKKPNELSRV | 2102 | 2116 | Positive |
| SARS-CoV-2 | orf1ab | ELSRVLGLKTLATHG | 2112 | 2126 | Negative |
| SARS-CoV-2 | orf1ab | ELSRVLGLKTLATHG | 2112 | 2126 | Overlapping |
| SARS-CoV-2 | orf1ab | VLGLKTLATHGLAAV | 2116 | 2130 | Negative |
| SARS-CoV-2 | orf1ab | GLKTLATHGLAAVNS | 2118 | 2132 | Negative |
| SARS-CoV-2 | orf1ab | THGLAAVNSVPWDTI | 2124 | 2138 | Negative |
| SARS-CoV-2 | orf1ab | GLAAVNSVPWDTIAN | 2126 | 2140 | Negative |
| SARS-CoV-2 | orf1ab | SVPWDTIANYAKPFL | 2132 | 2146 | Positive |
| SARS-CoV-2 | orf1ab | NYAKPFLNKVVSTTT | 2140 | 2154 | Positive |

| | | | | | |
|---|---|---|---|---|---|
| SARS-CoV-2 | orf1ab | AKPFLNKVVSTTTNI | 2142 | 2156 | Negative |
| SARS-CoV-2 | orf1ab | NRVCTNYMPYFFTLL | 2162 | 2176 | Positive |
| SARS-CoV-2 | orf1ab | MPTTIAKNTVKSVGK | 2194 | 2208 | Positive |
| SARS-CoV-2 | orf1ab | EASFNYLKSPNFSKL | 2212 | 2226 | Overlapping |
| SARS-CoV-2 | orf1ab | NYLKSPNFSKLINII | 2216 | 2230 | Positive |
| SARS-CoV-2 | orf1ab | VCLGSLIYSTAALGV | 2238 | 2252 | Negative |
| SARS-CoV-2 | orf1ab | STAALGVLMSNLGMP | 2246 | 2260 | Positive |
| SARS-CoV-2 | orf1ab | AALGVLMSNLGMPSY | 2248 | 2262 | Positive |
| SARS-CoV-2 | orf1ab | NLGMPSYCTGYREGY | 2256 | 2270 | Overlapping |
| SARS-CoV-2 | orf1ab | GMPSYCTGYREGYLN | 2258 | 2272 | Overlapping |
| SARS-CoV-2 | orf1ab | YCTGYREGYLNSTNV | 2262 | 2276 | Overlapping |
| SARS-CoV-2 | orf1ab | TGYREGYLNSTNVTI | 2264 | 2278 | Overlapping |
| SARS-CoV-2 | orf1ab | YREGYLNSTNVTIAT | 2266 | 2280 | Negative |
| SARS-CoV-2 | orf1ab | EGYLNSTNVTIATYC | 2268 | 2282 | Negative |
| SARS-CoV-2 | orf1ab | EGYLNSTNVTIATYC | 2268 | 2282 | Overlapping |
| SARS-CoV-2 | orf1ab | CTGSIPCSVCLSGLD | 2282 | 2296 | Negative |
| SARS-CoV-2 | orf1ab | GLDSLDTYPSLETIQ | 2294 | 2308 | Negative |
| SARS-CoV-2 | orf1ab | DSLDTYPSLETIQIT | 2296 | 2310 | Positive |
| SARS-CoV-2 | orf1ab | TYPSLETIQITISSF | 2300 | 2314 | Positive |
| SARS-CoV-2 | orf1ab | GLVAEWFLAYILFTR | 2322 | 2336 | Negative |
| SARS-CoV-2 | orf1ab | VAEWFLAYILFTRFF | 2324 | 2338 | Positive |
| SARS-CoV-2 | orf1ab | EWFLAYILFTRFFYV | 2326 | 2340 | Overlapping |
| SARS-CoV-2 | orf1ab | VLGLAAIMQLFFSYF | 2340 | 2354 | Negative |
| SARS-CoV-2 | orf1ab | GLAAIMQLFFSYFAV | 2342 | 2356 | Negative |
| SARS-CoV-2 | orf1ab | NLVQMAPISAMVRMY | 2370 | 2384 | Positive |
| SARS-CoV-2 | orf1ab | PISAMVRMYIFFASF | 2376 | 2390 | Positive |
| SARS-CoV-2 | orf1ab | MVRMYIFFASFYYVW | 2380 | 2394 | Positive |
| SARS-CoV-2 | orf1ab | NRATRVECTTIVNGV | 2416 | 2430 | Overlapping |
| SARS-CoV-2 | orf1ab | RVECTTIVNGVRRSF | 2420 | 2434 | Overlapping |
| SARS-CoV-2 | orf1ab | ECTTIVNGVRRSFYV | 2422 | 2436 | Overlapping |
| SARS-CoV-2 | orf1ab | NGVRRSFYVYANGGK | 2428 | 2442 | Positive |
| SARS-CoV-2 | orf1ab | KGFCKLHNWNCVNCD | 2442 | 2456 | Positive |
| SARS-CoV-2 | orf1ab | NCDTFCAGSTFISDE | 2454 | 2468 | Positive |
| SARS-CoV-2 | orf1ab | DTFCAGSTFISDEVA | 2456 | 2470 | Positive |
| SARS-CoV-2 | orf1ab | SDEVARDLSLQFKRP | 2466 | 2480 | Positive |
| SARS-CoV-2 | orf1ab | EVARDLSLQFKRPIN | 2468 | 2482 | Overlapping |
| SARS-CoV-2 | orf1ab | PINPTDQSSYIVDSV | 2480 | 2494 | Overlapping |
| SARS-CoV-2 | orf1ab | NPTDQSSYIVDSVTV | 2482 | 2496 | Positive |
| SARS-CoV-2 | orf1ab | TDQSSYIVDSVTVKN | 2484 | 2498 | Positive |
| SARS-CoV-2 | orf1ab | DSVTVKNGSIHLYFD | 2492 | 2506 | Positive |
| SARS-CoV-2 | orf1ab | TYERHSLSHFVNLDN | 2512 | 2526 | Positive |
| SARS-CoV-2 | orf1ab | ERHSLSHFVNLDNLR | 2514 | 2528 | Overlapping |
| SARS-CoV-2 | orf1ab | NLRANNTKGSLPINV | 2526 | 2540 | Positive |
| SARS-CoV-2 | orf1ab | NNTKGSLPINVIVFD | 2530 | 2544 | Positive |
| SARS-CoV-2 | orf1ab | NNTKGSLPINVIVFD | 2530 | 2544 | Negative |
| SARS-CoV-2 | orf1ab | GSLPINVIVFDGKSK | 2534 | 2548 | Positive |
| SARS-CoV-2 | orf1ab | KCEESSAKSASVYYS | 2548 | 2562 | Overlapping |
| SARS-CoV-2 | orf1ab | EESSAKSASVYYSQL | 2550 | 2564 | Overlapping |
| SARS-CoV-2 | orf1ab | PILLLDQALVSDVGD | 2568 | 2582 | Overlapping |

| | | | | | |
|---|---|---|---|---|---|
| SARS-CoV-2 | orf1ab | LVSDVGDSAEVAVKM | 2576 | 2590 | Positive |
| SARS-CoV-2 | orf1ab | SDVGDSAEVAVKMFD | 2578 | 2592 | Positive |
| SARS-CoV-2 | orf1ab | VAVKMFDAYVNTFSS | 2586 | 2600 | Positive |
| SARS-CoV-2 | orf1ab | MFDAYVNTFSSTFNV | 2590 | 2604 | Negative |
| SARS-CoV-2 | orf1ab | VPMEKLKTLVATAEA | 2604 | 2618 | Positive |
| SARS-CoV-2 | orf1ab | MEKLKTLVATAEAEL | 2606 | 2620 | Negative |
| SARS-CoV-2 | orf1ab | ATAEAELAKNVSLDN | 2614 | 2628 | Negative |
| SARS-CoV-2 | orf1ab | AARQGFVDSDVETKD | 2636 | 2650 | Negative |
| SARS-CoV-2 | orf1ab | RQGFVDSDVETKDVV | 2638 | 2652 | Overlapping |
| SARS-CoV-2 | orf1ab | GFVDSDVETKDVVEC | 2640 | 2654 | Overlapping |
| SARS-CoV-2 | orf1ab | VDSDVETKDVVECLK | 2642 | 2656 | Positive |
| SARS-CoV-2 | orf1ab | VDSDVETKDVVECLK | 2642 | 2656 | Overlapping |
| SARS-CoV-2 | orf1ab | SDVETKDVVECLKLS | 2644 | 2658 | Overlapping |
| SARS-CoV-2 | orf1ab | VETKDVVECLKLSHQ | 2646 | 2660 | Positive |
| SARS-CoV-2 | orf1ab | DIEVTGDSCNNYMLT | 2662 | 2676 | Overlapping |
| SARS-CoV-2 | orf1ab | EVTGDSCNNYMLTYN | 2664 | 2678 | Negative |
| SARS-CoV-2 | orf1ab | EVTGDSCNNYMLTYN | 2664 | 2678 | Overlapping |
| SARS-CoV-2 | orf1ab | TGDSCNNYMLTYNKV | 2666 | 2680 | Overlapping |
| SARS-CoV-2 | orf1ab | NYMLTYNKVENMTPR | 2672 | 2686 | Positive |
| SARS-CoV-2 | orf1ab | NKVENMTPRDLGACI | 2678 | 2692 | Positive |
| SARS-CoV-2 | orf1ab | NMTPRDLGACIDCSA | 2682 | 2696 | Positive |
| SARS-CoV-2 | orf1ab | TPRDLGACIDCSARH | 2684 | 2698 | Positive |
| SARS-CoV-2 | orf1ab | RDLGACIDCSARHIN | 2686 | 2700 | Positive |
| SARS-CoV-2 | orf1ab | CSARHINAQVAKSHN | 2694 | 2708 | Positive |
| SARS-CoV-2 | orf1ab | NAQVAKSHNIALIWN | 2700 | 2714 | Positive |
| SARS-CoV-2 | orf1ab | NNLPFKLTCATTRQV | 2736 | 2750 | Positive |
| SARS-CoV-2 | orf1ab | NWLKQLIKVTLVFLF | 2768 | 2782 | Positive |
| SARS-CoV-2 | orf1ab | FLFVAAIFYLITPVH | 2780 | 2794 | Positive |
| SARS-CoV-2 | orf1ab | ITPVHVMSKHTDFSS | 2790 | 2804 | Positive |
| SARS-CoV-2 | orf1ab | PVHVMSKHTDFSSEI | 2792 | 2806 | Positive |
| SARS-CoV-2 | orf1ab | PVHVMSKHTDFSSEI | 2792 | 2806 | Overlapping |
| SARS-CoV-2 | orf1ab | TDFSSEIIGYKAIDG | 2800 | 2814 | Positive |
| SARS-CoV-2 | orf1ab | SEIIGYKAIDGGVTR | 2804 | 2818 | Positive |
| SARS-CoV-2 | orf1ab | RDIASTDTCFANKHA | 2818 | 2832 | Positive |
| SARS-CoV-2 | orf1ab | STDTCFANKHADFDT | 2822 | 2836 | Positive |
| SARS-CoV-2 | orf1ab | TNDKACPLIAAVITR | 2846 | 2860 | Negative |
| SARS-CoV-2 | orf1ab | VPGLPGTILRTTNGD | 2866 | 2880 | Negative |
| SARS-CoV-2 | orf1ab | GLPGTILRTTNGDFL | 2868 | 2882 | Overlapping |
| SARS-CoV-2 | orf1ab | NICYTPSKLIEYTDF | 2894 | 2908 | Positive |
| SARS-CoV-2 | orf1ab | SKLIEYTDFATSACV | 2900 | 2914 | Overlapping |
| SARS-CoV-2 | orf1ab | EYTDFATSACVLAAE | 2904 | 2918 | Overlapping |
| SARS-CoV-2 | orf1ab | TDFATSACVLAAECT | 2906 | 2920 | Positive |
| SARS-CoV-2 | orf1ab | ECTIFKDASGKPVPY | 2918 | 2932 | Overlapping |
| SARS-CoV-2 | orf1ab | VLEGSVAYESLRPDT | 2938 | 2952 | Positive |
| SARS-CoV-2 | orf1ab | EGSVAYESLRPDTRY | 2940 | 2954 | Overlapping |
| SARS-CoV-2 | orf1ab | AYESLRPDTRYVLMD | 2944 | 2958 | Positive |
| SARS-CoV-2 | orf1ab | ESLRPDTRYVLMDGS | 2946 | 2960 | Overlapping |
| SARS-CoV-2 | orf1ab | LRPDTRYVLMDGSII | 2948 | 2962 | Negative |
| SARS-CoV-2 | orf1ab | DGSIIQFPNTYLEGS | 2958 | 2972 | Overlapping |

| | | | | | |
|---|---|---|---|---|---|
| SARS-CoV-2 | orf1ab | YLEGSVRVVTTFDSE | 2968 | 2982 | Negative |
| SARS-CoV-2 | orf1ab | EGSVRVVTTFDSEYC | 2970 | 2984 | Negative |
| SARS-CoV-2 | orf1ab | EGSVRVVTTFDSEYC | 2970 | 2984 | Overlapping |
| SARS-CoV-2 | orf1ab | VTTFDSEYCRHGTCE | 2976 | 2990 | Positive |
| SARS-CoV-2 | orf1ab | TFDSEYCRHGTCERS | 2978 | 2992 | Positive |
| SARS-CoV-2 | orf1ab | DSEYCRHGTCERSEA | 2980 | 2994 | Overlapping |
| SARS-CoV-2 | orf1ab | EYCRHGTCERSEAGV | 2982 | 2996 | Overlapping |
| SARS-CoV-2 | orf1ab | ERSEAGVCVSTSGRW | 2990 | 3004 | Overlapping |
| SARS-CoV-2 | orf1ab | SEAGVCVSTSGRWVL | 2992 | 3006 | Positive |
| SARS-CoV-2 | orf1ab | TSGRWVLNNDYYRSL | 3000 | 3014 | Positive |
| SARS-CoV-2 | orf1ab | NDYYRSLPGVFCGVD | 3008 | 3022 | Negative |
| SARS-CoV-2 | orf1ab | YYRSLPGVFCGVDAV | 3010 | 3024 | Negative |
| SARS-CoV-2 | orf1ab | RSLPGVFCGVDAVNL | 3012 | 3026 | Positive |
| SARS-CoV-2 | orf1ab | RSLPGVFCGVDAVNL | 3012 | 3026 | Overlapping |
| SARS-CoV-2 | orf1ab | LPGVFCGVDAVNLLT | 3014 | 3028 | Negative |
| SARS-CoV-2 | orf1ab | GVFCGVDAVNLLTNM | 3016 | 3030 | Negative |
| SARS-CoV-2 | orf1ab | FCGVDAVNLLTNMFT | 3018 | 3032 | Negative |
| SARS-CoV-2 | orf1ab | GVDAVNLLTNMFTPL | 3020 | 3034 | Negative |
| SARS-CoV-2 | orf1ab | TNMFTPLIQPIGALD | 3028 | 3042 | Negative |
| SARS-CoV-2 | orf1ab | GEYSHVVAFNTLLFL | 3072 | 3086 | Negative |
| SARS-CoV-2 | orf1ab | LTPVYSFLPGVYSVI | 3094 | 3108 | Positive |
| SARS-CoV-2 | orf1ab | PVYSFLPGVYSVIYL | 3096 | 3110 | Negative |
| SARS-CoV-2 | orf1ab | LTNDVSFLAHIQWMV | 3116 | 3130 | Positive |
| SARS-CoV-2 | orf1ab | NDVSFLAHIQWMVMF | 3118 | 3132 | Positive |
| SARS-CoV-2 | orf1ab | RVVFNGVSFSTFEEA | 3164 | 3178 | Positive |
| SARS-CoV-2 | orf1ab | VFNGVSFSTFEEAAL | 3166 | 3180 | Positive |
| SARS-CoV-2 | orf1ab | NGVSFSTFEEAALCT | 3168 | 3182 | Positive |
| SARS-CoV-2 | orf1ab | TFEEAALCTFLLNKE | 3174 | 3188 | Overlapping |
| SARS-CoV-2 | orf1ab | EEAALCTFLLNKEMY | 3176 | 3190 | Overlapping |
| SARS-CoV-2 | orf1ab | NKEMYLKLRSDVLLP | 3186 | 3200 | Positive |
| SARS-CoV-2 | orf1ab | EMYLKLRSDVLLPLT | 3188 | 3202 | Overlapping |
| SARS-CoV-2 | orf1ab | NKYKYFSGAMDTTSY | 3212 | 3226 | Positive |
| SARS-CoV-2 | orf1ab | EAACCHLAKALNDFS | 3228 | 3242 | Overlapping |
| SARS-CoV-2 | orf1ab | SNSGSDVLYQPPQTS | 3242 | 3256 | Overlapping |
| SARS-CoV-2 | orf1ab | SDVLYQPPQTSITSA | 3246 | 3260 | Positive |
| SARS-CoV-2 | orf1ab | GKVEGCMVQVTCGTT | 3274 | 3288 | Negative |
| SARS-CoV-2 | orf1ab | VEGCMVQVTCGTTTL | 3276 | 3290 | Negative |
| SARS-CoV-2 | orf1ab | GCMVQVTCGTTTLNG | 3278 | 3292 | Negative |
| SARS-CoV-2 | orf1ab | LNGLWLDDVVYCPRH | 3290 | 3304 | Negative |
| SARS-CoV-2 | orf1ab | GLWLDDVVYCPRHVI | 3292 | 3306 | Negative |
| SARS-CoV-2 | orf1ab | DDVVYCPRHVICTSE | 3296 | 3310 | Positive |
| SARS-CoV-2 | orf1ab | HVICTSEDMLNPNYE | 3304 | 3318 | Overlapping |
| SARS-CoV-2 | orf1ab | TSEDMLNPNYEDLLI | 3308 | 3322 | Overlapping |
| SARS-CoV-2 | orf1ab | EDMLNPNYEDLLIRK | 3310 | 3324 | Overlapping |
| SARS-CoV-2 | orf1ab | NPNYEDLLIRKSNHN | 3314 | 3328 | Positive |
| SARS-CoV-2 | orf1ab | NYEDLLIRKSNHNFL | 3316 | 3330 | Positive |
| SARS-CoV-2 | orf1ab | EDLLIRKSNHNFLVQ | 3318 | 3332 | Overlapping |
| SARS-CoV-2 | orf1ab | NHNFLVQAGNVQLRV | 3326 | 3340 | Positive |
| SARS-CoV-2 | orf1ab | NFLVQAGNVQLRVIG | 3328 | 3342 | Positive |

| | | | | | |
|---|---|---|---|---|---|
| SARS-CoV-2 | orf1ab | VIGHSMQNCVLKLKV | 3340 | 3354 | Negative |
| SARS-CoV-2 | orf1ab | GHSMQNCVLKLKVDT | 3342 | 3356 | Positive |
| SARS-CoV-2 | orf1ab | SMQNCVLKLKVDTAN | 3344 | 3358 | Positive |
| SARS-CoV-2 | orf1ab | SVGFNIDYDCVSFCY | 3410 | 3424 | Negative |
| SARS-CoV-2 | orf1ab | GFNIDYDCVSFCYMH | 3412 | 3426 | Negative |
| SARS-CoV-2 | orf1ab | NIDYDCVSFCYMHHM | 3414 | 3428 | Positive |
| SARS-CoV-2 | orf1ab | DYDCVSFCYMHHMEL | 3416 | 3430 | Positive |
| SARS-CoV-2 | orf1ab | VSFCYMHHMELPTGV | 3420 | 3434 | Negative |
| SARS-CoV-2 | orf1ab | FCYMHHMELPTGVHA | 3422 | 3436 | Positive |
| SARS-CoV-2 | orf1ab | YMHHMELPTGVHAGT | 3424 | 3438 | Positive |
| SARS-CoV-2 | orf1ab | HHMELPTGVHAGTDL | 3426 | 3440 | Overlapping |
| SARS-CoV-2 | orf1ab | TGVHAGTDLEGNFYG | 3432 | 3446 | Overlapping |
| SARS-CoV-2 | orf1ab | TDLEGNFYGPFVDRQ | 3438 | 3452 | Positive |
| SARS-CoV-2 | orf1ab | FYGPFVDRQTAQAAG | 3444 | 3458 | Positive |
| SARS-CoV-2 | orf1ab | GPFVDRQTAQAAGTD | 3446 | 3460 | Overlapping |
| SARS-CoV-2 | orf1ab | GTDTTITVNVLAWLY | 3458 | 3472 | Negative |
| SARS-CoV-2 | orf1ab | NRFTTTLNDFNLVAM | 3484 | 3498 | Positive |
| SARS-CoV-2 | orf1ab | LNDFNLVAMKYNYEP | 3490 | 3504 | Negative |
| SARS-CoV-2 | orf1ab | NLVAMKYNYEPLTQD | 3494 | 3508 | Positive |
| SARS-CoV-2 | orf1ab | VAMKYNYEPLTQDHV | 3496 | 3510 | Positive |
| SARS-CoV-2 | orf1ab | MKYNYEPLTQDHVDI | 3498 | 3512 | Positive |
| SARS-CoV-2 | orf1ab | YEPLTQDHVDILGPL | 3502 | 3516 | Positive |
| SARS-CoV-2 | orf1ab | PLTQDHVDILGPLSA | 3504 | 3518 | Positive |
| SARS-CoV-2 | orf1ab | DHVDILGPLSAQTGI | 3508 | 3522 | Overlapping |
| SARS-CoV-2 | orf1ab | GPLSAQTGIAVLDMC | 3514 | 3528 | Positive |
| SARS-CoV-2 | orf1ab | NGRTILGSALLEDEF | 3540 | 3554 | Positive |
| SARS-CoV-2 | orf1ab | GSALLEDEFTPFDVV | 3546 | 3560 | Positive |
| SARS-CoV-2 | orf1ab | ALLEDEFTPFDVVRQ | 3548 | 3562 | Negative |
| SARS-CoV-2 | orf1ab | ALLEDEFTPFDVVRQ | 3548 | 3562 | Overlapping |
| SARS-CoV-2 | orf1ab | LEDEFTPFDVVRQCS | 3550 | 3564 | Positive |
| SARS-CoV-2 | orf1ab | DEFTPFDVVRQCSGV | 3552 | 3566 | Negative |
| SARS-CoV-2 | orf1ab | FTPFDVVRQCSGVTF | 3554 | 3568 | Negative |
| SARS-CoV-2 | orf1ab | SLLVLVQSTQWSLFF | 3590 | 3604 | Positive |
| SARS-CoV-2 | orf1ab | FFLYENAFLPFAMGI | 3604 | 3618 | Negative |
| SARS-CoV-2 | orf1ab | LYENAFLPFAMGIIA | 3606 | 3620 | Negative |
| SARS-CoV-2 | orf1ab | ENAFLPFAMGIIAMS | 3608 | 3622 | Overlapping |
| SARS-CoV-2 | orf1ab | SMWALIISVTSNYSG | 3732 | 3746 | Positive |
| SARS-CoV-2 | orf1ab | WALIISVTSNYSGVV | 3734 | 3748 | Negative |
| SARS-CoV-2 | orf1ab | YSGVVTTVMFLARGI | 3744 | 3758 | Negative |
| SARS-CoV-2 | orf1ab | GVVTTVMFLARGIVF | 3746 | 3760 | Negative |
| SARS-CoV-2 | orf1ab | VTTVMFLARGIVFMC | 3748 | 3762 | Positive |
| SARS-CoV-2 | orf1ab | CVEYCPIFFITGNTL | 3762 | 3776 | Positive |
| SARS-CoV-2 | orf1ab | EYCPIFFITGNTLQC | 3764 | 3778 | Overlapping |
| SARS-CoV-2 | orf1ab | DYLVSTQEFRYMNSQ | 3812 | 3826 | Overlapping |
| SARS-CoV-2 | orf1ab | YMNSQGLLPPKNSID | 3822 | 3836 | Positive |
| SARS-CoV-2 | orf1ab | NSQGLLPPKNSIDAF | 3824 | 3838 | Positive |
| SARS-CoV-2 | orf1ab | LLGVGGKPCIKVATV | 3844 | 3858 | Negative |
| SARS-CoV-2 | orf1ab | GVGGKPCIKVATVQS | 3846 | 3860 | Negative |
| SARS-CoV-2 | orf1ab | SKMSDVKCTSVVLLS | 3860 | 3874 | Negative |

| | | | | | |
|---|---|---|---|---|---|
| SARS-CoV-2 | orf1ab | RVESSSKLWAQCVQL | 3880 | 3894 | Negative |
| SARS-CoV-2 | orf1ab | ESSSKLWAQCVQLHN | 3882 | 3896 | Overlapping |
| SARS-CoV-2 | orf1ab | SSKLWAQCVQLHNDI | 3884 | 3898 | Overlapping |
| SARS-CoV-2 | orf1ab | KLWAQCVQLHNDILL | 3886 | 3900 | Positive |
| SARS-CoV-2 | orf1ab | VQLHNDILLAKDTTE | 3892 | 3906 | Negative |
| SARS-CoV-2 | orf1ab | LHNDILLAKDTTEAF | 3894 | 3908 | Positive |
| SARS-CoV-2 | orf1ab | NDILLAKDTTEAFEK | 3896 | 3910 | Positive |
| SARS-CoV-2 | orf1ab | ILLAKDTTEAFEKMV | 3898 | 3912 | Positive |
| SARS-CoV-2 | orf1ab | ILLAKDTTEAFEKMV | 3898 | 3912 | Negative |
| SARS-CoV-2 | orf1ab | LAKDTTEAFEKMVSL | 3900 | 3914 | Positive |
| SARS-CoV-2 | orf1ab | KDTTEAFEKMVSLLS | 3902 | 3916 | Positive |
| SARS-CoV-2 | orf1ab | TTEAFEKMVSLLSVL | 3904 | 3918 | Positive |
| SARS-CoV-2 | orf1ab | EAFEKMVSLLSVLLS | 3906 | 3920 | Overlapping |
| SARS-CoV-2 | orf1ab | SMQGAVDINKLCEEM | 3920 | 3934 | Positive |
| SARS-CoV-2 | orf1ab | QGAVDINKLCEEMLD | 3922 | 3936 | Positive |
| SARS-CoV-2 | orf1ab | AVDINKLCEEMLDNR | 3924 | 3938 | Overlapping |
| SARS-CoV-2 | orf1ab | DINKLCEEMLDNRAT | 3926 | 3940 | Positive |
| SARS-CoV-2 | orf1ab | NKLCEEMLDNRATLQ | 3928 | 3942 | Positive |
| SARS-CoV-2 | orf1ab | LCEEMLDNRATLQAI | 3930 | 3944 | Overlapping |
| SARS-CoV-2 | orf1ab | EEMLDNRATLQAIAS | 3932 | 3946 | Overlapping |
| SARS-CoV-2 | orf1ab | SEFSSLPSYAAFATA | 3946 | 3960 | Positive |
| SARS-CoV-2 | orf1ab | PSYAAFATAQEAYEQ | 3952 | 3966 | Positive |
| SARS-CoV-2 | orf1ab | AQEAYEQAVANGDSE | 3960 | 3974 | Overlapping |
| SARS-CoV-2 | orf1ab | EAYEQAVANGDSEVV | 3962 | 3976 | Overlapping |
| SARS-CoV-2 | orf1ab | YEQAVANGDSEVVLK | 3964 | 3978 | Negative |
| SARS-CoV-2 | orf1ab | QAVANGDSEVVLKKL | 3966 | 3980 | Negative |
| SARS-CoV-2 | orf1ab | NGDSEVVLKKLKKSL | 3970 | 3984 | Positive |
| SARS-CoV-2 | orf1ab | DSEVVLKKLKKSLNV | 3972 | 3986 | Overlapping |
| SARS-CoV-2 | orf1ab | EVVLKKLKKSLNVAK | 3974 | 3988 | Negative |
| SARS-CoV-2 | orf1ab | EVVLKKLKKSLNVAK | 3974 | 3988 | Overlapping |
| SARS-CoV-2 | orf1ab | KSEFDRDAAMQRKLE | 3988 | 4002 | Negative |
| SARS-CoV-2 | orf1ab | EFDRDAAMQRKLEKM | 3990 | 4004 | Overlapping |
| SARS-CoV-2 | orf1ab | KLEKMADQAMTQMYK | 4000 | 4014 | Overlapping |
| SARS-CoV-2 | orf1ab | EKMADQAMTQMYKQA | 4002 | 4016 | Overlapping |
| SARS-CoV-2 | orf1ab | RKLDNDALNNIINNA | 4038 | 4052 | Positive |
| SARS-CoV-2 | orf1ab | LDNDALNNIINNARD | 4040 | 4054 | Overlapping |
| SARS-CoV-2 | orf1ab | NDALNNIINNARDGC | 4042 | 4056 | Positive |
| SARS-CoV-2 | orf1ab | NNIINNARDGCVPLN | 4046 | 4060 | Positive |
| SARS-CoV-2 | orf1ab | NNARDGCVPLNIIPL | 4050 | 4064 | Positive |
| SARS-CoV-2 | orf1ab | DGCVPLNIIPLTTAA | 4054 | 4068 | Positive |
| SARS-CoV-2 | orf1ab | IPLTTAAKLMVVIPD | 4062 | 4076 | Positive |
| SARS-CoV-2 | orf1ab | VVIPDYNTYKNTCDG | 4072 | 4086 | Positive |
| SARS-CoV-2 | orf1ab | IPDYNTYKNTCDGTT | 4074 | 4088 | Positive |
| SARS-CoV-2 | orf1ab | GTTFTYASALWEIQQ | 4086 | 4100 | Negative |
| SARS-CoV-2 | orf1ab | ASALWEIQQVVDADS | 4092 | 4106 | Positive |
| SARS-CoV-2 | orf1ab | ALWEIQQVVDADSKI | 4094 | 4108 | Negative |
| SARS-CoV-2 | orf1ab | QVVDADSKIVQLSEI | 4100 | 4114 | Overlapping |
| SARS-CoV-2 | orf1ab | VDADSKIVQLSEISM | 4102 | 4116 | Positive |
| SARS-CoV-2 | orf1ab | ADSKIVQLSEISMDN | 4104 | 4118 | Negative |

| | | | | | |
|---|---|---|---|---|---|
| SARS-CoV-2 | orf1ab | IVQLSEISMDNSPNL | 4108 | 4122 | Negative |
| SARS-CoV-2 | orf1ab | SEISMDNSPNLAWPL | 4112 | 4126 | Positive |
| SARS-CoV-2 | orf1ab | NSPNLAWPLIVTALR | 4118 | 4132 | Positive |
| SARS-CoV-2 | orf1ab | NSAVKLQNNELSPVA | 4134 | 4148 | Positive |
| SARS-CoV-2 | orf1ab | NELSPVALRQMSCAA | 4142 | 4156 | Positive |
| SARS-CoV-2 | orf1ab | PVALRQMSCAAGTTQ | 4146 | 4160 | Positive |
| SARS-CoV-2 | orf1ab | TDDNALAYYNTTKGG | 4164 | 4178 | Positive |
| SARS-CoV-2 | orf1ab | SDLQDLKWARFPKSD | 4186 | 4200 | Positive |
| SARS-CoV-2 | orf1ab | TGTIYTELEPPCRFV | 4202 | 4216 | Positive |
| SARS-CoV-2 | orf1ab | TIYTELEPPCRFVTD | 4204 | 4218 | Positive |
| SARS-CoV-2 | orf1ab | YTELEPPCRFVTDTP | 4206 | 4220 | Negative |
| SARS-CoV-2 | orf1ab | ELEPPCRFVTDTPKG | 4208 | 4222 | Negative |
| SARS-CoV-2 | orf1ab | ELEPPCRFVTDTPKG | 4208 | 4222 | Overlapping |
| SARS-CoV-2 | orf1ab | EPPCRFVTDTPKGPK | 4210 | 4224 | Positive |
| SARS-CoV-2 | orf1ab | NLNRGMVLGSLAATV | 4236 | 4250 | Positive |
| SARS-CoV-2 | orf1ab | NRGMVLGSLAATVRL | 4238 | 4252 | Positive |
| SARS-CoV-2 | orf1ab | LQAGNATEVPANSTV | 4252 | 4266 | Positive |
| SARS-CoV-2 | orf1ab | NATEVPANSTVLSFC | 4256 | 4270 | Positive |
| SARS-CoV-2 | orf1ab | TEVPANSTVLSFCAF | 4258 | 4272 | Positive |
| SARS-CoV-2 | orf1ab | VTPEANMDQESFGGA | 4310 | 4324 | Positive |
| SARS-CoV-2 | orf1ab | PEANMDQESFGGASC | 4312 | 4326 | Negative |
| SARS-CoV-2 | orf1ab | GGASCCLYCRCHIDH | 4322 | 4336 | Positive |
| SARS-CoV-2 | orf1ab | PVGFTLKNTVCTVCG | 4360 | 4374 | Overlapping |
| SARS-CoV-2 | orf1ab | GMWKGYGCSCDQLRE | 4374 | 4388 | Negative |
| SARS-CoV-2 | orf1ab | LREPMLQSADAQSFL | 4386 | 4400 | Positive |
| SARS-CoV-2 | orf1ab | EPMLQSADAQSFLNR | 4388 | 4402 | Overlapping |
| SARS-CoV-2 | orf1ab | QSADAQSFLNRVCGV | 4392 | 4406 | Negative |
| SARS-CoV-2 | orf1ab | TGTSTDVVYRAFDIY | 4416 | 4430 | Positive |
| SARS-CoV-2 | orf1ab | TDVVYRAFDIYNDKV | 4420 | 4434 | Positive |
| SARS-CoV-2 | orf1ab | FLKTNCCRFQEKDED | 4440 | 4454 | Positive |
| SARS-CoV-2 | orf1ab | KTNCCRFQEKDEDDN | 4442 | 4456 | Overlapping |
| SARS-CoV-2 | orf1ab | NCCRFQEKDEDDNLI | 4444 | 4458 | Positive |
| SARS-CoV-2 | orf1ab | CRFQEKDEDDNLIDS | 4446 | 4460 | Positive |
| SARS-CoV-2 | orf1ab | FQEKDEDDNLIDSYF | 4448 | 4462 | Positive |
| SARS-CoV-2 | orf1ab | EKDEDDNLIDSYFVV | 4450 | 4464 | Positive |
| SARS-CoV-2 | orf1ab | EKDEDDNLIDSYFVV | 4450 | 4464 | Overlapping |
| SARS-CoV-2 | orf1ab | DEDDNLIDSYFVVKR | 4452 | 4466 | Overlapping |
| SARS-CoV-2 | orf1ab | NLIDSYFVVKRHTFS | 4456 | 4470 | Positive |
| SARS-CoV-2 | orf1ab | SYFVVKRHTFSNYQH | 4460 | 4474 | Positive |
| SARS-CoV-2 | orf1ab | FVVKRHTFSNYQHEE | 4462 | 4476 | Positive |
| SARS-CoV-2 | orf1ab | VKRHTFSNYQHEETI | 4464 | 4478 | Positive |
| SARS-CoV-2 | orf1ab | RHTFSNYQHEETIYN | 4466 | 4480 | Positive |
| SARS-CoV-2 | orf1ab | TFSNYQHEETIYNLL | 4468 | 4482 | Positive |
| SARS-CoV-2 | orf1ab | SNYQHEETIYNLLKD | 4470 | 4484 | Positive |
| SARS-CoV-2 | orf1ab | YQHEETIYNLLKDCP | 4472 | 4486 | Positive |
| SARS-CoV-2 | orf1ab | HEETIYNLLKDCPAV | 4474 | 4488 | Positive |
| SARS-CoV-2 | orf1ab | ETIYNLLKDCPAVAK | 4476 | 4490 | Positive |
| SARS-CoV-2 | orf1ab | IYNLLKDCPAVAKHD | 4478 | 4492 | Positive |
| SARS-CoV-2 | orf1ab | NLLKDCPAVAKHDFF | 4480 | 4494 | Positive |

| | | | | | |
|---|---|---|---|---|---|
| SARS-CoV-2 | orf1ab | KHDFFKFRIDGDMVP | 4490 | 4504 | Negative |
| SARS-CoV-2 | orf1ab | FRIDGDMVPHISRQR | 4496 | 4510 | Negative |
| SARS-CoV-2 | orf1ab | GDMVPHISRQRLTKY | 4500 | 4514 | Negative |
| SARS-CoV-2 | orf1ab | MVPHISRQRLTKYTM | 4502 | 4516 | Positive |
| SARS-CoV-2 | orf1ab | YTMADLVYALRHFDE | 4514 | 4528 | Negative |
| SARS-CoV-2 | orf1ab | DLVYALRHFDEGNCD | 4518 | 4532 | Positive |
| SARS-CoV-2 | orf1ab | ALRHFDEGNCDTLKE | 4522 | 4536 | Overlapping |
| SARS-CoV-2 | orf1ab | RHFDEGNCDTLKEIL | 4524 | 4538 | Positive |
| SARS-CoV-2 | orf1ab | FDEGNCDTLKEILVT | 4526 | 4540 | Negative |
| SARS-CoV-2 | orf1ab | FDEGNCDTLKEILVT | 4526 | 4540 | Overlapping |
| SARS-CoV-2 | orf1ab | EGNCDTLKEILVTYN | 4528 | 4542 | Negative |
| SARS-CoV-2 | orf1ab | EGNCDTLKEILVTYN | 4528 | 4542 | Overlapping |
| SARS-CoV-2 | orf1ab | NCDTLKEILVTYNCC | 4530 | 4544 | Positive |
| SARS-CoV-2 | orf1ab | LKEILVTYNCCDDDY | 4534 | 4548 | Positive |
| SARS-CoV-2 | orf1ab | EILVTYNCCDDDYFN | 4536 | 4550 | Negative |
| SARS-CoV-2 | orf1ab | EILVTYNCCDDDYFN | 4536 | 4550 | Overlapping |
| SARS-CoV-2 | orf1ab | TYNCCDDDYFNKKDW | 4540 | 4554 | Positive |
| SARS-CoV-2 | orf1ab | NCCDDDYFNKKDWYD | 4542 | 4556 | Overlapping |
| SARS-CoV-2 | orf1ab | CDDDYFNKKDWYDFV | 4544 | 4558 | Overlapping |
| SARS-CoV-2 | orf1ab | DDYFNKKDWYDFVEN | 4546 | 4560 | Negative |
| SARS-CoV-2 | orf1ab | YFNKKDWYDFVENPD | 4548 | 4562 | Positive |
| SARS-CoV-2 | orf1ab | NKKDWYDFVENPDIL | 4550 | 4564 | Positive |
| SARS-CoV-2 | orf1ab | ERVRQALLKTVQFCD | 4572 | 4586 | Overlapping |
| SARS-CoV-2 | orf1ab | VQFCDAMRNAGIVGV | 4582 | 4596 | Negative |
| SARS-CoV-2 | orf1ab | FCDAMRNAGIVGVLT | 4584 | 4598 | Negative |
| SARS-CoV-2 | orf1ab | DAMRNAGIVGVLTLD | 4586 | 4600 | Positive |
| SARS-CoV-2 | orf1ab | NAGIVGVLTLDNQDL | 4590 | 4604 | Positive |
| SARS-CoV-2 | orf1ab | GIVGVLTLDNQDLNG | 4592 | 4606 | Negative |
| SARS-CoV-2 | orf1ab | TLDNQDLNGNWYDFG | 4598 | 4612 | Positive |
| SARS-CoV-2 | orf1ab | DNQDLNGNWYDFGDF | 4600 | 4614 | Overlapping |
| SARS-CoV-2 | orf1ab | QDLNGNWYDFGDFIQ | 4602 | 4616 | Negative |
| SARS-CoV-2 | orf1ab | GSGVPVVDSYYSLLM | 4620 | 4634 | Negative |
| SARS-CoV-2 | orf1ab | GSGVPVVDSYYSLLM | 4620 | 4634 | Overlapping |
| SARS-CoV-2 | orf1ab | GVPVVDSYYSLLMPI | 4622 | 4636 | Overlapping |
| SARS-CoV-2 | orf1ab | PVVDSYYSLLMPILT | 4624 | 4638 | Positive |
| SARS-CoV-2 | orf1ab | PVVDSYYSLLMPILT | 4624 | 4638 | Overlapping |
| SARS-CoV-2 | orf1ab | ILTLTRALTAESHVD | 4636 | 4650 | Positive |
| SARS-CoV-2 | orf1ab | TAESHVDTDLTKPYI | 4644 | 4658 | Overlapping |
| SARS-CoV-2 | orf1ab | ESHVDTDLTKPYIKW | 4646 | 4660 | Overlapping |
| SARS-CoV-2 | orf1ab | HVDTDLTKPYIKWDL | 4648 | 4662 | Negative |
| SARS-CoV-2 | orf1ab | PYIKWDLLKYDFTEE | 4656 | 4670 | Positive |
| SARS-CoV-2 | orf1ab | WDLLKYDFTEERLKL | 4660 | 4674 | Positive |
| SARS-CoV-2 | orf1ab | TEERLKLFDRYFKYW | 4668 | 4682 | Overlapping |
| SARS-CoV-2 | orf1ab | ERLKLFDRYFKYWDQ | 4670 | 4684 | Overlapping |
| SARS-CoV-2 | orf1ab | QTYHPNCVNCLDDRC | 4684 | 4698 | Positive |
| SARS-CoV-2 | orf1ab | PNCVNCLDDRCILHC | 4688 | 4702 | Positive |
| SARS-CoV-2 | orf1ab | NCLDDRCILHCANFN | 4692 | 4706 | Positive |
| SARS-CoV-2 | orf1ab | TSFGPLVRKIFVDGV | 4716 | 4730 | Negative |
| SARS-CoV-2 | orf1ab | DGVPFVVSTGYHFRE | 4728 | 4742 | Overlapping |

| | | | | | |
|---|---|---|---|---|---|
| SARS-CoV-2 | orf1ab | FVVSTGYHFRELGVV | 4732 | 4746 | Negative |
| SARS-CoV-2 | orf1ab | VSTGYHFRELGVVHN | 4734 | 4748 | Positive |
| SARS-CoV-2 | orf1ab | TGYHFRELGVVHNQD | 4736 | 4750 | Positive |
| SARS-CoV-2 | orf1ab | YHFRELGVVHNQDVN | 4738 | 4752 | Positive |
| SARS-CoV-2 | orf1ab | FRELGVVHNQDVNLH | 4740 | 4754 | Overlapping |
| SARS-CoV-2 | orf1ab | ELGVVHNQDVNLHSS | 4742 | 4756 | Negative |
| SARS-CoV-2 | orf1ab | ELGVVHNQDVNLHSS | 4742 | 4756 | Overlapping |
| SARS-CoV-2 | orf1ab | GVVHNQDVNLHSSRL | 4744 | 4758 | Overlapping |
| SARS-CoV-2 | orf1ab | VHNQDVNLHSSRLSF | 4746 | 4760 | Positive |
| SARS-CoV-2 | orf1ab | NQDVNLHSSRLSFKE | 4748 | 4762 | Positive |
| SARS-CoV-2 | orf1ab | NLHSSRLSFKELLVY | 4752 | 4766 | Positive |
| SARS-CoV-2 | orf1ab | FKELLVYAADPAMHA | 4760 | 4774 | Overlapping |
| SARS-CoV-2 | orf1ab | ELLVYAADPAMHAAS | 4762 | 4776 | Negative |
| SARS-CoV-2 | orf1ab | ELLVYAADPAMHAAS | 4762 | 4776 | Overlapping |
| SARS-CoV-2 | orf1ab | NLLLDKRTTCFSVAA | 4778 | 4792 | Positive |
| SARS-CoV-2 | orf1ab | NFNKDFYDFAVSKGF | 4806 | 4820 | Positive |
| SARS-CoV-2 | orf1ab | NKDFYDFAVSKGFFK | 4808 | 4822 | Positive |
| SARS-CoV-2 | orf1ab | SVELKHFFFAQDGNA | 4826 | 4840 | Positive |
| SARS-CoV-2 | orf1ab | ELKHFFFAQDGNAAI | 4828 | 4842 | Negative |
| SARS-CoV-2 | orf1ab | ELKHFFFAQDGNAAI | 4828 | 4842 | Overlapping |
| SARS-CoV-2 | orf1ab | DYDYYRYNLPTMCDI | 4844 | 4858 | Positive |
| SARS-CoV-2 | orf1ab | TMCDIRQLLFVVEVV | 4854 | 4868 | Positive |
| SARS-CoV-2 | orf1ab | VVEVVDKYFDCYDGG | 4864 | 4878 | Positive |
| SARS-CoV-2 | orf1ab | EVVDKYFDCYDGGCI | 4866 | 4880 | Negative |
| SARS-CoV-2 | orf1ab | EVVDKYFDCYDGGCI | 4866 | 4880 | Overlapping |
| SARS-CoV-2 | orf1ab | NNLDKSAGFPFNKWG | 4888 | 4902 | Positive |
| SARS-CoV-2 | orf1ab | ARLYYDSMSYEDQDA | 4904 | 4918 | Positive |
| SARS-CoV-2 | orf1ab | LYYDSMSYEDQDALF | 4906 | 4920 | Positive |
| SARS-CoV-2 | orf1ab | SMSYEDQDALFAYTK | 4910 | 4924 | Positive |
| SARS-CoV-2 | orf1ab | SYEDQDALFAYTKRN | 4912 | 4926 | Positive |
| SARS-CoV-2 | orf1ab | EDQDALFAYTKRNVI | 4914 | 4928 | Overlapping |
| SARS-CoV-2 | orf1ab | NRARTVAGVSICSTM | 4944 | 4958 | Positive |
| SARS-CoV-2 | orf1ab | NRQFHQKLLKSIAAT | 4960 | 4974 | Positive |
| SARS-CoV-2 | orf1ab | TVVIGTSKFYGGWHN | 4978 | 4992 | Positive |
| SARS-CoV-2 | orf1ab | VIGTSKFYGGWHNML | 4980 | 4994 | Positive |
| SARS-CoV-2 | orf1ab | NMLKTVYSDVENPHL | 4992 | 5006 | Positive |
| SARS-CoV-2 | orf1ab | DVENPHLMGWDYPKC | 5000 | 5014 | Positive |
| SARS-CoV-2 | orf1ab | ENPHLMGWDYPKCDR | 5002 | 5016 | Overlapping |
| SARS-CoV-2 | orf1ab | PHLMGWDYPKCDRAM | 5004 | 5018 | Positive |
| SARS-CoV-2 | orf1ab | NMLRIMASLVLARKH | 5020 | 5034 | Positive |
| SARS-CoV-2 | orf1ab | RLANECAQVLSEMVM | 5046 | 5060 | Positive |
| SARS-CoV-2 | orf1ab | ECAQVLSEMVMCGGS | 5050 | 5064 | Overlapping |
| SARS-CoV-2 | orf1ab | TDGNKIADKYVRNLQ | 5102 | 5116 | Positive |
| SARS-CoV-2 | orf1ab | NLQHRLYECLYRNRD | 5114 | 5128 | Overlapping |
| SARS-CoV-2 | orf1ab | QHRLYECLYRNRDVD | 5116 | 5130 | Negative |
| SARS-CoV-2 | orf1ab | RLYECLYRNRDVDTD | 5118 | 5132 | Overlapping |
| SARS-CoV-2 | orf1ab | CLYRNRDVDTDFVNE | 5122 | 5136 | Overlapping |
| SARS-CoV-2 | orf1ab | YRNRDVDTDFVNEFY | 5124 | 5138 | Positive |
| SARS-CoV-2 | orf1ab | NRDVDTDFVNEFYAY | 5126 | 5140 | Positive |

| | | | | | |
|---|---|---|---|---|---|
| SARS-CoV-2 | orf1ab | EFYAYLRKHFSMMIL | 5136 | 5150 | Overlapping |
| SARS-CoV-2 | orf1ab | SMMILSDDAVVCFNS | 5146 | 5160 | Positive |
| SARS-CoV-2 | orf1ab | GLVASIKNFKSVLYY | 5166 | 5180 | Negative |
| SARS-CoV-2 | orf1ab | FKSVLYYQNNVFMSE | 5174 | 5188 | Positive |
| SARS-CoV-2 | orf1ab | YQNNVFMSEAKCWTE | 5180 | 5194 | Positive |
| SARS-CoV-2 | orf1ab | NNVFMSEAKCWTETD | 5182 | 5196 | Positive |
| SARS-CoV-2 | orf1ab | MSEAKCWTETDLTKG | 5186 | 5200 | Overlapping |
| SARS-CoV-2 | orf1ab | EAKCWTETDLTKGPH | 5188 | 5202 | Overlapping |
| SARS-CoV-2 | orf1ab | KCWTETDLTKGPHEF | 5190 | 5204 | Positive |
| SARS-CoV-2 | orf1ab | WTETDLTKGPHEFCS | 5192 | 5206 | Positive |
| SARS-CoV-2 | orf1ab | ETDLTKGPHEFCSQH | 5194 | 5208 | Positive |
| SARS-CoV-2 | orf1ab | ETDLTKGPHEFCSQH | 5194 | 5208 | Overlapping |
| SARS-CoV-2 | orf1ab | DLTKGPHEFCSQHTM | 5196 | 5210 | Positive |
| SARS-CoV-2 | orf1ab | TKGPHEFCSQHTMLV | 5198 | 5212 | Positive |
| SARS-CoV-2 | orf1ab | GPHEFCSQHTMLVKQ | 5200 | 5214 | Overlapping |
| SARS-CoV-2 | orf1ab | FCSQHTMLVKQGDDY | 5204 | 5218 | Positive |
| SARS-CoV-2 | orf1ab | QGDDYVYLPYPDPSR | 5214 | 5228 | Negative |
| SARS-CoV-2 | orf1ab | DDYVYLPYPDPSRIL | 5216 | 5230 | Overlapping |
| SARS-CoV-2 | orf1ab | PYPDPSRILGAGCFV | 5222 | 5236 | Positive |
| SARS-CoV-2 | orf1ab | TDGTLMIERFVSLAI | 5242 | 5256 | Positive |
| SARS-CoV-2 | orf1ab | AYPLTKHPNQEYADV | 5258 | 5272 | Positive |
| SARS-CoV-2 | orf1ab | PLTKHPNQEYADVFH | 5260 | 5274 | Positive |
| SARS-CoV-2 | orf1ab | NQEYADVFHLYLQYI | 5266 | 5280 | Positive |
| SARS-CoV-2 | orf1ab | EYADVFHLYLQYIRK | 5268 | 5282 | Overlapping |
| SARS-CoV-2 | orf1ab | HLYLQYIRKLHDELT | 5274 | 5288 | Positive |
| SARS-CoV-2 | orf1ab | YLQYIRKLHDELTGH | 5276 | 5290 | Positive |
| SARS-CoV-2 | orf1ab | QYIRKLHDELTGHML | 5278 | 5292 | Positive |
| SARS-CoV-2 | orf1ab | IRKLHDELTGHMLDM | 5280 | 5294 | Positive |
| SARS-CoV-2 | orf1ab | KLHDELTGHMLDMYS | 5282 | 5296 | Positive |
| SARS-CoV-2 | orf1ab | HDELTGHMLDMYSVM | 5284 | 5298 | Overlapping |
| SARS-CoV-2 | orf1ab | ELTGHMLDMYSVMLT | 5286 | 5300 | Negative |
| SARS-CoV-2 | orf1ab | ELTGHMLDMYSVMLT | 5286 | 5300 | Overlapping |
| SARS-CoV-2 | orf1ab | TGHMLDMYSVMLTND | 5288 | 5302 | Overlapping |
| SARS-CoV-2 | orf1ab | TNDNTSRYWEPEFYE | 5300 | 5314 | Overlapping |
| SARS-CoV-2 | orf1ab | RYWEPEFYEAMYTPH | 5306 | 5320 | Positive |
| SARS-CoV-2 | orf1ab | WEPEFYEAMYTPHTV | 5308 | 5322 | Overlapping |
| SARS-CoV-2 | orf1ab | PEFYEAMYTPHTVLQ | 5310 | 5324 | Overlapping |
| SARS-CoV-2 | orf1ab | FYEAMYTPHTVLQAV | 5312 | 5326 | Overlapping |
| SARS-CoV-2 | orf1ab | EAMYTPHTVLQAVGA | 5314 | 5328 | Overlapping |
| SARS-CoV-2 | orf1ab | DHVISTSHKLVLSVN | 5356 | 5370 | Positive |
| SARS-CoV-2 | orf1ab | SHKLVLSVNPYVCNA | 5362 | 5376 | Negative |
| SARS-CoV-2 | orf1ab | APGCDVTDVTQLYLG | 5376 | 5390 | Negative |
| SARS-CoV-2 | orf1ab | GCDVTDVTQLYLGGM | 5378 | 5392 | Negative |
| SARS-CoV-2 | orf1ab | TDVTQLYLGGMSYYC | 5382 | 5396 | Positive |
| SARS-CoV-2 | orf1ab | LYKNTCVGSDNVTDF | 5416 | 5430 | Negative |
| SARS-CoV-2 | orf1ab | KNTCVGSDNVTDFNA | 5418 | 5432 | Negative |
| SARS-CoV-2 | orf1ab | TCVGSDNVTDFNAIA | 5420 | 5434 | Negative |
| SARS-CoV-2 | orf1ab | VGSDNVTDFNAIATC | 5422 | 5436 | Negative |
| SARS-CoV-2 | orf1ab | SDNVTDFNAIATCDW | 5424 | 5438 | Overlapping |

| | | | | | |
|---|---|---|---|---|---|
| SARS-CoV-2 | orf1ab | TDFNAIATCDWTNAG | 5428 | 5442 | Positive |
| SARS-CoV-2 | orf1ab | CTERLKLFAAETLKA | 5450 | 5464 | Overlapping |
| SARS-CoV-2 | orf1ab | ERLKLFAAETLKATE | 5452 | 5466 | Overlapping |
| SARS-CoV-2 | orf1ab | LFAAETLKATEETFK | 5456 | 5470 | Positive |
| SARS-CoV-2 | orf1ab | AAETLKATEETFKLS | 5458 | 5472 | Overlapping |
| SARS-CoV-2 | orf1ab | ETLKATEETFKLSYG | 5460 | 5474 | Overlapping |
| SARS-CoV-2 | orf1ab | LKATEETFKLSYGIA | 5462 | 5476 | Negative |
| SARS-CoV-2 | orf1ab | ATEETFKLSYGIATV | 5464 | 5478 | Negative |
| SARS-CoV-2 | orf1ab | EETFKLSYGIATVRE | 5466 | 5480 | Overlapping |
| SARS-CoV-2 | orf1ab | SYGIATVREVLSDRE | 5472 | 5486 | Negative |
| SARS-CoV-2 | orf1ab | GIATVREVLSDRELH | 5474 | 5488 | Negative |
| SARS-CoV-2 | orf1ab | VREVLSDRELHLSWE | 5478 | 5492 | Overlapping |
| SARS-CoV-2 | orf1ab | EVLSDRELHLSWEVG | 5480 | 5494 | Negative |
| SARS-CoV-2 | orf1ab | EVLSDRELHLSWEVG | 5480 | 5494 | Overlapping |
| SARS-CoV-2 | orf1ab | DRELHLSWEVGKPRP | 5484 | 5498 | Overlapping |
| SARS-CoV-2 | orf1ab | ELHLSWEVGKPRPPL | 5486 | 5500 | Negative |
| SARS-CoV-2 | orf1ab | ELHLSWEVGKPRPPL | 5486 | 5500 | Overlapping |
| SARS-CoV-2 | orf1ab | EVGKPRPPLNRNYVF | 5492 | 5506 | Overlapping |
| SARS-CoV-2 | orf1ab | NSKVQIGEYTFEKGD | 5514 | 5528 | Positive |
| SARS-CoV-2 | orf1ab | QIGEYTFEKGDYGDA | 5518 | 5532 | Negative |
| SARS-CoV-2 | orf1ab | GEYTFEKGDYGDAVV | 5520 | 5534 | Negative |
| SARS-CoV-2 | orf1ab | NVGDYFVLTSHTVMP | 5544 | 5558 | Overlapping |
| SARS-CoV-2 | orf1ab | PTLVPQEHYVRITGL | 5562 | 5576 | Positive |
| SARS-CoV-2 | orf1ab | LVPQEHYVRITGLYP | 5564 | 5578 | Positive |
| SARS-CoV-2 | orf1ab | PQEHYVRITGLYPTL | 5566 | 5580 | Positive |
| SARS-CoV-2 | orf1ab | EHYVRITGLYPTLNI | 5568 | 5582 | Overlapping |
| SARS-CoV-2 | orf1ab | TGLYPTLNISDEFSS | 5574 | 5588 | Negative |
| SARS-CoV-2 | orf1ab | PTLNISDEFSSNVAN | 5578 | 5592 | Overlapping |
| SARS-CoV-2 | orf1ab | DEFSSNVANYQKVGM | 5584 | 5598 | Negative |
| SARS-CoV-2 | orf1ab | NYQKVGMQKYSTLQG | 5592 | 5606 | Positive |
| SARS-CoV-2 | orf1ab | QKVGMQKYSTLQGPP | 5594 | 5608 | Positive |
| SARS-CoV-2 | orf1ab | VGMQKYSTLQGPPGT | 5596 | 5610 | Positive |
| SARS-CoV-2 | orf1ab | MQKYSTLQGPPGTGK | 5598 | 5612 | Positive |
| SARS-CoV-2 | orf1ab | KYSTLQGPPGTGKSH | 5600 | 5614 | Positive |
| SARS-CoV-2 | orf1ab | GPPGTGKSHFAIGLA | 5606 | 5620 | Positive |
| SARS-CoV-2 | orf1ab | GPPGTGKSHFAIGLA | 5606 | 5620 | Overlapping |
| SARS-CoV-2 | orf1ab | AIGLALYYPSARIVY | 5616 | 5630 | Negative |
| SARS-CoV-2 | orf1ab | GLALYYPSARIVYTA | 5618 | 5632 | Negative |
| SARS-CoV-2 | orf1ab | YLPIDKCSRIIPARA | 5648 | 5662 | Positive |
| SARS-CoV-2 | orf1ab | PIDKCSRIIPARARV | 5650 | 5664 | Overlapping |
| SARS-CoV-2 | orf1ab | IPARARVECFDKFKV | 5658 | 5672 | Positive |
| SARS-CoV-2 | orf1ab | VECFDKFKVNSTLEQ | 5664 | 5678 | Negative |
| SARS-CoV-2 | orf1ab | STLEQYVFCTVNALP | 5674 | 5688 | Positive |
| SARS-CoV-2 | orf1ab | ALPETTADIVVFDEI | 5686 | 5700 | Positive |
| SARS-CoV-2 | orf1ab | PETTADIVVFDEISM | 5688 | 5702 | Overlapping |
| SARS-CoV-2 | orf1ab | ADIVVFDEISMATNY | 5692 | 5706 | Negative |
| SARS-CoV-2 | orf1ab | IVVFDEISMATNYDL | 5694 | 5708 | Positive |
| SARS-CoV-2 | orf1ab | VFDEISMATNYDLSV | 5696 | 5710 | Overlapping |
| SARS-CoV-2 | orf1ab | DEISMATNYDLSVVN | 5698 | 5712 | Overlapping |

| | | | | | |
|---|---|---|---|---|---|
| SARS-CoV-2 | orf1ab | NARLRAKHYVYIGDP | 5712 | 5726 | Positive |
| SARS-CoV-2 | orf1ab | GDPAQLPAPRTLLTK | 5724 | 5738 | Positive |
| SARS-CoV-2 | orf1ab | PAPRTLLTKGTLEPE | 5730 | 5744 | Positive |
| SARS-CoV-2 | orf1ab | PRTLLTKGTLEPEYF | 5732 | 5746 | Positive |
| SARS-CoV-2 | orf1ab | TLLTKGTLEPEYFNS | 5734 | 5748 | Positive |
| SARS-CoV-2 | orf1ab | LTKGTLEPEYFNSVC | 5736 | 5750 | Positive |
| SARS-CoV-2 | orf1ab | EPEYFNSVCRLMKTI | 5742 | 5756 | Overlapping |
| SARS-CoV-2 | orf1ab | PDMFLGTCRRCPAEI | 5758 | 5772 | Positive |
| SARS-CoV-2 | orf1ab | TCRRCPAEIVDTVSA | 5764 | 5778 | Positive |
| SARS-CoV-2 | orf1ab | KLKAHKDKSAQCFKM | 5784 | 5798 | Positive |
| SARS-CoV-2 | orf1ab | KAHKDKSAQCFKMFY | 5786 | 5800 | Positive |
| SARS-CoV-2 | orf1ab | QCFKMFYKGVITHDV | 5794 | 5808 | Positive |
| SARS-CoV-2 | orf1ab | FKMFYKGVITHDVSS | 5796 | 5810 | Positive |
| SARS-CoV-2 | orf1ab | YKGVITHDVSSAINR | 5800 | 5814 | Overlapping |
| SARS-CoV-2 | orf1ab | GVITHDVSSAINRPQ | 5802 | 5816 | Negative |
| SARS-CoV-2 | orf1ab | HDVSSAINRPQIGVV | 5806 | 5820 | Negative |
| SARS-CoV-2 | orf1ab | VSSAINRPQIGVVRE | 5808 | 5822 | Negative |
| SARS-CoV-2 | orf1ab | QIGVVREFLTRNPAW | 5816 | 5830 | Negative |
| SARS-CoV-2 | orf1ab | GVVREFLTRNPAWRK | 5818 | 5832 | Negative |
| SARS-CoV-2 | orf1ab | EFLTRNPAWRKAVFI | 5822 | 5836 | Negative |
| SARS-CoV-2 | orf1ab | NSQNAVASKILGLPT | 5840 | 5854 | Positive |
| SARS-CoV-2 | orf1ab | TVDSSQGSEYDYVIF | 5856 | 5870 | Positive |
| SARS-CoV-2 | orf1ab | EYDYVIFTQTTETAH | 5864 | 5878 | Positive |
| SARS-CoV-2 | orf1ab | TETAHSCNVNRFNVA | 5874 | 5888 | Positive |
| SARS-CoV-2 | orf1ab | NVAITRAKVGILCIM | 5886 | 5900 | Positive |
| SARS-CoV-2 | orf1ab | TRAKVGILCIMSDRD | 5890 | 5904 | Negative |
| SARS-CoV-2 | orf1ab | DRDLYDKLQFTSLEI | 5902 | 5916 | Positive |
| SARS-CoV-2 | orf1ab | NVATLQAENVTGLFK | 5920 | 5934 | Positive |
| SARS-CoV-2 | orf1ab | KVITGLHPTQAPTHL | 5938 | 5952 | Positive |
| SARS-CoV-2 | orf1ab | ITGLHPTQAPTHLSV | 5940 | 5954 | Negative |
| SARS-CoV-2 | orf1ab | GLHPTQAPTHLSVDT | 5942 | 5956 | Overlapping |
| SARS-CoV-2 | orf1ab | TEGLCVDIPGIPKDM | 5960 | 5974 | Negative |
| SARS-CoV-2 | orf1ab | GLCVDIPGIPKDMTY | 5962 | 5976 | Negative |
| SARS-CoV-2 | orf1ab | MMGFKMNYQVNGYPN | 5982 | 5996 | Negative |
| SARS-CoV-2 | orf1ab | NYQVNGYPNMFITRE | 5988 | 6002 | Overlapping |
| SARS-CoV-2 | orf1ab | QVNGYPNMFITREEA | 5990 | 6004 | Positive |
| SARS-CoV-2 | orf1ab | NGYPNMFITREEAIR | 5992 | 6006 | Negative |
| SARS-CoV-2 | orf1ab | NMFITREEAIRHVRA | 5996 | 6010 | Positive |
| SARS-CoV-2 | orf1ab | TREEAIRHVRAWIGF | 6000 | 6014 | Negative |
| SARS-CoV-2 | orf1ab | EEAIRHVRAWIGFDV | 6002 | 6016 | Overlapping |
| SARS-CoV-2 | orf1ab | RHVRAWIGFDVEGCH | 6006 | 6020 | Positive |
| SARS-CoV-2 | orf1ab | AWIGFDVEGCHATRE | 6010 | 6024 | Positive |
| SARS-CoV-2 | orf1ab | VEGCHATREAVGTNL | 6016 | 6030 | Negative |
| SARS-CoV-2 | orf1ab | TREAVGTNLPLQLGF | 6022 | 6036 | Positive |
| SARS-CoV-2 | orf1ab | EAVGTNLPLQLGFST | 6024 | 6038 | Overlapping |
| SARS-CoV-2 | orf1ab | VGTNLPLQLGFSTGV | 6026 | 6040 | Negative |
| SARS-CoV-2 | orf1ab | TNLPLQLGFSTGVNL | 6028 | 6042 | Negative |
| SARS-CoV-2 | orf1ab | TGVNLVAVPTGYVDT | 6038 | 6052 | Positive |
| SARS-CoV-2 | orf1ab | AVPTGYVDTPNNTDF | 6044 | 6058 | Overlapping |

| | | | | | |
|---|---|---|---|---|---|
| SARS-CoV-2 | orf1ab | PTGYVDTPNNTDFSR | 6046 | 6060 | Overlapping |
| SARS-CoV-2 | orf1ab | GYVDTPNNTDFSRVS | 6048 | 6062 | Overlapping |
| SARS-CoV-2 | orf1ab | NNTDFSRVSAKPPPG | 6054 | 6068 | Positive |
| SARS-CoV-2 | orf1ab | TDFSRVSAKPPPGDQ | 6056 | 6070 | Positive |
| SARS-CoV-2 | orf1ab | SAKPPPGDQFKHLIP | 6062 | 6076 | Positive |
| SARS-CoV-2 | orf1ab | KPPPGDQFKHLIPLM | 6064 | 6078 | Positive |
| SARS-CoV-2 | orf1ab | PPGDQFKHLIPLMYK | 6066 | 6080 | Positive |
| SARS-CoV-2 | orf1ab | KHLIPLMYKGLPWNV | 6072 | 6086 | Positive |
| SARS-CoV-2 | orf1ab | SDTLKNLSDRVVFVL | 6096 | 6110 | Positive |
| SARS-CoV-2 | orf1ab | FVLWAHGFELTSMKY | 6108 | 6122 | Positive |
| SARS-CoV-2 | orf1ab | AHGFELTSMKYFVKI | 6112 | 6126 | Positive |
| SARS-CoV-2 | orf1ab | ELTSMKYFVKIGPER | 6116 | 6130 | Negative |
| SARS-CoV-2 | orf1ab | VKIGPERTCCLCDRR | 6124 | 6138 | Negative |
| SARS-CoV-2 | orf1ab | TCFSTASDTYACWHH | 6140 | 6154 | Positive |
| SARS-CoV-2 | orf1ab | SDTYACWHHSIGFDY | 6146 | 6160 | Positive |
| SARS-CoV-2 | orf1ab | WHHSIGFDYVYNPFM | 6152 | 6166 | Positive |
| SARS-CoV-2 | orf1ab | DVQQWGFTGNLQSNH | 6168 | 6182 | Positive |
| SARS-CoV-2 | orf1ab | QQWGFTGNLQSNHDL | 6170 | 6184 | Positive |
| SARS-CoV-2 | orf1ab | WGFTGNLQSNHDLYC | 6172 | 6186 | Positive |
| SARS-CoV-2 | orf1ab | GNLQSNHDLYCQVHG | 6176 | 6190 | Positive |
| SARS-CoV-2 | orf1ab | LQSNHDLYCQVHGNA | 6178 | 6192 | Positive |
| SARS-CoV-2 | orf1ab | SNHDLYCQVHGNAHV | 6180 | 6194 | Positive |
| SARS-CoV-2 | orf1ab | HDLYCQVHGNAHVAS | 6182 | 6196 | Positive |
| SARS-CoV-2 | orf1ab | LYCQVHGNAHVASCD | 6184 | 6198 | Positive |
| SARS-CoV-2 | orf1ab | CQVHGNAHVASCDAI | 6186 | 6200 | Positive |
| SARS-CoV-2 | orf1ab | VHGNAHVASCDAIMT | 6188 | 6202 | Positive |
| SARS-CoV-2 | orf1ab | GNAHVASCDAIMTRC | 6190 | 6204 | Positive |
| SARS-CoV-2 | orf1ab | SCDAIMTRCLAVHEC | 6196 | 6210 | Positive |
| SARS-CoV-2 | orf1ab | AVHECFVKRVDWTIE | 6206 | 6220 | Negative |
| SARS-CoV-2 | orf1ab | RVDWTIEYPIIGDEL | 6214 | 6228 | Overlapping |
| SARS-CoV-2 | orf1ab | DWTIEYPIIGDELKI | 6216 | 6230 | Negative |
| SARS-CoV-2 | orf1ab | TIEYPIIGDELKINA | 6218 | 6232 | Positive |
| SARS-CoV-2 | orf1ab | EYPIIGDELKINAAC | 6220 | 6234 | Overlapping |
| SARS-CoV-2 | orf1ab | PIIGDELKINAACRK | 6222 | 6236 | Negative |
| SARS-CoV-2 | orf1ab | DELKINAACRKVQHM | 6226 | 6240 | Overlapping |
| SARS-CoV-2 | orf1ab | LLADKFPVLHDIGNP | 6246 | 6260 | Positive |
| SARS-CoV-2 | orf1ab | KFPVLHDIGNPKAIK | 6250 | 6264 | Positive |
| SARS-CoV-2 | orf1ab | PVLHDIGNPKAIKCV | 6252 | 6266 | Overlapping |
| SARS-CoV-2 | orf1ab | LHDIGNPKAIKCVPQ | 6254 | 6268 | Positive |
| SARS-CoV-2 | orf1ab | GNPKAIKCVPQADVE | 6258 | 6272 | Positive |
| SARS-CoV-2 | orf1ab | DVEWKFYDAQPCSDK | 6270 | 6284 | Overlapping |
| SARS-CoV-2 | orf1ab | EWKFYDAQPCSDKAY | 6272 | 6286 | Overlapping |
| SARS-CoV-2 | orf1ab | YDAQPCSDKAYKIEE | 6276 | 6290 | Negative |
| SARS-CoV-2 | orf1ab | PCSDKAYKIEELFYS | 6280 | 6294 | Positive |
| SARS-CoV-2 | orf1ab | SDKAYKIEELFYSYA | 6282 | 6296 | Positive |
| SARS-CoV-2 | orf1ab | KAYKIEELFYSYATH | 6284 | 6298 | Negative |
| SARS-CoV-2 | orf1ab | YKIEELFYSYATHSD | 6286 | 6300 | Positive |
| SARS-CoV-2 | orf1ab | IEELFYSYATHSDKF | 6288 | 6302 | Overlapping |
| SARS-CoV-2 | orf1ab | ELFYSYATHSDKFTD | 6290 | 6304 | Negative |

| | | | | | |
|---|---|---|---|---|---|
| SARS-CoV-2 | orf1ab | ELFYSYATHSDKFTD | 6290 | 6304 | Overlapping |
| SARS-CoV-2 | orf1ab | FYSYATHSDKFTDGV | 6292 | 6306 | Negative |
| SARS-CoV-2 | orf1ab | DGVCLFWNCNVDRYP | 6304 | 6318 | Positive |
| SARS-CoV-2 | orf1ab | NSIVCRFDTRVLSNL | 6320 | 6334 | Positive |
| SARS-CoV-2 | orf1ab | GCDGGSLYVNKHAFH | 6338 | 6352 | Overlapping |
| SARS-CoV-2 | orf1ab | DGGSLYVNKHAFHTP | 6340 | 6354 | Positive |
| SARS-CoV-2 | orf1ab | PAFDKSAFVNLKQLP | 6354 | 6368 | Negative |
| SARS-CoV-2 | orf1ab | DSPCESHGKQVVSDI | 6374 | 6388 | Positive |
| SARS-CoV-2 | orf1ab | PCESHGKQVVSDIDY | 6376 | 6390 | Positive |
| SARS-CoV-2 | orf1ab | ESHGKQVVSDIDYVP | 6378 | 6392 | Overlapping |
| SARS-CoV-2 | orf1ab | VVSDIDYVPLKSATC | 6384 | 6398 | Positive |
| SARS-CoV-2 | orf1ab | SDIDYVPLKSATCIT | 6386 | 6400 | Positive |
| SARS-CoV-2 | orf1ab | CITRCNLGGAVCRHH | 6398 | 6412 | Positive |
| SARS-CoV-2 | orf1ab | CNLGGAVCRHHANEY | 6402 | 6416 | Positive |
| SARS-CoV-2 | orf1ab | GAVCRHHANEYRLYL | 6406 | 6420 | Positive |
| SARS-CoV-2 | orf1ab | RHHANEYRLYLDAYN | 6410 | 6424 | Positive |
| SARS-CoV-2 | orf1ab | NEYRLYLDAYNMMIS | 6414 | 6428 | Positive |
| SARS-CoV-2 | orf1ab | NMMISAGFSLWVYKQ | 6424 | 6438 | Positive |
| SARS-CoV-2 | orf1ab | NTFTRLQSLENVAFN | 6446 | 6460 | Positive |
| SARS-CoV-2 | orf1ab | NVAFNVVNKGHFDGQ | 6456 | 6470 | Positive |
| SARS-CoV-2 | orf1ab | NVVNKGHFDGQQGEV | 6460 | 6474 | Positive |
| SARS-CoV-2 | orf1ab | DGQQGEVPVSIINNT | 6468 | 6482 | Overlapping |
| SARS-CoV-2 | orf1ab | VSIINNTVYTKVDGV | 6476 | 6490 | Negative |
| SARS-CoV-2 | orf1ab | IINNTVYTKVDGVDV | 6478 | 6492 | Overlapping |
| SARS-CoV-2 | orf1ab | NNTVYTKVDGVDVEL | 6480 | 6494 | Overlapping |
| SARS-CoV-2 | orf1ab | TVYTKVDGVDVELFE | 6482 | 6496 | Overlapping |
| SARS-CoV-2 | orf1ab | YTKVDGVDVELFENK | 6484 | 6498 | Negative |
| SARS-CoV-2 | orf1ab | KVDGVDVELFENKTT | 6486 | 6500 | Overlapping |
| SARS-CoV-2 | orf1ab | DGVDVELFENKTTLP | 6488 | 6502 | Positive |
| SARS-CoV-2 | orf1ab | DGVDVELFENKTTLP | 6488 | 6502 | Overlapping |
| SARS-CoV-2 | orf1ab | VDVELFENKTTLPVN | 6490 | 6504 | Positive |
| SARS-CoV-2 | orf1ab | VELFENKTTLPVNVA | 6492 | 6506 | Negative |
| SARS-CoV-2 | orf1ab | LFENKTTLPVNVAFE | 6494 | 6508 | Positive |
| SARS-CoV-2 | orf1ab | ENKTTLPVNVAFELW | 6496 | 6510 | Overlapping |
| SARS-CoV-2 | orf1ab | TLPVNVAFELWAKRN | 6500 | 6514 | Positive |
| SARS-CoV-2 | orf1ab | AFELWAKRNIKPVPE | 6506 | 6520 | Overlapping |
| SARS-CoV-2 | orf1ab | ELWAKRNIKPVPEVK | 6508 | 6522 | Negative |
| SARS-CoV-2 | orf1ab | ELWAKRNIKPVPEVK | 6508 | 6522 | Overlapping |
| SARS-CoV-2 | orf1ab | NIKPVPEVKILNNLG | 6514 | 6528 | Positive |
| SARS-CoV-2 | orf1ab | KPVPEVKILNNLGVD | 6516 | 6530 | Negative |
| SARS-CoV-2 | orf1ab | VPEVKILNNLGVDIA | 6518 | 6532 | Overlapping |
| SARS-CoV-2 | orf1ab | EVKILNNLGVDIAAN | 6520 | 6534 | Overlapping |
| SARS-CoV-2 | orf1ab | GVDIAANTVIWDYKR | 6528 | 6542 | Negative |
| SARS-CoV-2 | orf1ab | WDYKRDAPAHISTIG | 6538 | 6552 | Positive |
| SARS-CoV-2 | orf1ab | YKRDAPAHISTIGVC | 6540 | 6554 | Positive |
| SARS-CoV-2 | orf1ab | RDAPAHISTIGVCSM | 6542 | 6556 | Positive |
| SARS-CoV-2 | orf1ab | ISTIGVCSMTDIAKK | 6548 | 6562 | Positive |
| SARS-CoV-2 | orf1ab | TIGVCSMTDIAKKPT | 6550 | 6564 | Negative |
| SARS-CoV-2 | orf1ab | GVCSMTDIAKKPTET | 6552 | 6566 | Negative |

| | | | | | |
|---|---|---|---|---|---|
| SARS-CoV-2 | orf1ab | DIAKKPTETICAPLT | 6558 | 6572 | Positive |
| SARS-CoV-2 | orf1ab | PLTVFFDGRVDGQVD | 6570 | 6584 | Positive |
| SARS-CoV-2 | orf1ab | DGRVDGQVDLFRNAR | 6576 | 6590 | Positive |
| SARS-CoV-2 | orf1ab | DGRVDGQVDLFRNAR | 6576 | 6590 | Overlapping |
| SARS-CoV-2 | orf1ab | RVDGQVDLFRNARNG | 6578 | 6592 | Negative |
| SARS-CoV-2 | orf1ab | DGQVDLFRNARNGVL | 6580 | 6594 | Overlapping |
| SARS-CoV-2 | orf1ab | NARNGVLITEGSVKG | 6588 | 6602 | Positive |
| SARS-CoV-2 | orf1ab | GVLITEGSVKGLQPS | 6592 | 6606 | Negative |
| SARS-CoV-2 | orf1ab | VKGLQPSVGPKQASL | 6600 | 6614 | Negative |
| SARS-CoV-2 | orf1ab | GLQPSVGPKQASLNG | 6602 | 6616 | Negative |
| SARS-CoV-2 | orf1ab | QPSVGPKQASLNGVT | 6604 | 6618 | Negative |
| SARS-CoV-2 | orf1ab | GPKQASLNGVTLIGE | 6608 | 6622 | Positive |
| SARS-CoV-2 | orf1ab | LNGVTLIGEAVKTQF | 6614 | 6628 | Negative |
| SARS-CoV-2 | orf1ab | GVTLIGEAVKTQFNY | 6616 | 6630 | Negative |
| SARS-CoV-2 | orf1ab | IGEAVKTQFNYYKKV | 6620 | 6634 | Overlapping |
| SARS-CoV-2 | orf1ab | EAVKTQFNYYKKVDG | 6622 | 6636 | Overlapping |
| SARS-CoV-2 | orf1ab | VKTQFNYYKKVDGVV | 6624 | 6638 | Negative |
| SARS-CoV-2 | orf1ab | VDGVVQQLPETYFTQ | 6634 | 6648 | Negative |
| SARS-CoV-2 | orf1ab | GVVQQLPETYFTQSR | 6636 | 6650 | Negative |
| SARS-CoV-2 | orf1ab | PETYFTQSRNLQEFK | 6642 | 6656 | Negative |
| SARS-CoV-2 | orf1ab | LQEFKPRSQMEIDFL | 6652 | 6666 | Positive |
| SARS-CoV-2 | orf1ab | EFKPRSQMEIDFLEL | 6654 | 6668 | Overlapping |
| SARS-CoV-2 | orf1ab | RSQMEIDFLELAMDE | 6658 | 6672 | Overlapping |
| SARS-CoV-2 | orf1ab | QMEIDFLELAMDEFI | 6660 | 6674 | Overlapping |
| SARS-CoV-2 | orf1ab | EIDFLELAMDEFIER | 6662 | 6676 | Overlapping |
| SARS-CoV-2 | orf1ab | LELAMDEFIERYKLE | 6666 | 6680 | Overlapping |
| SARS-CoV-2 | orf1ab | MDEFIERYKLEGYAF | 6670 | 6684 | Positive |
| SARS-CoV-2 | orf1ab | EFIERYKLEGYAFEH | 6672 | 6686 | Overlapping |
| SARS-CoV-2 | orf1ab | KLEGYAFEHIVYGDF | 6678 | 6692 | Overlapping |
| SARS-CoV-2 | orf1ab | EGYAFEHIVYGDFSH | 6680 | 6694 | Negative |
| SARS-CoV-2 | orf1ab | EGYAFEHIVYGDFSH | 6680 | 6694 | Overlapping |
| SARS-CoV-2 | orf1ab | YAFEHIVYGDFSHSQ | 6682 | 6696 | Positive |
| SARS-CoV-2 | orf1ab | HIVYGDFSHSQLGGL | 6686 | 6700 | Negative |
| SARS-CoV-2 | orf1ab | LAKRFKESPFELEDF | 6706 | 6720 | Positive |
| SARS-CoV-2 | orf1ab | FKESPFELEDFIPMD | 6710 | 6724 | Overlapping |
| SARS-CoV-2 | orf1ab | ESPFELEDFIPMDST | 6712 | 6726 | Overlapping |
| SARS-CoV-2 | orf1ab | PFELEDFIPMDSTVK | 6714 | 6728 | Overlapping |
| SARS-CoV-2 | orf1ab | ELEDFIPMDSTVKNY | 6716 | 6730 | Negative |
| SARS-CoV-2 | orf1ab | ELEDFIPMDSTVKNY | 6716 | 6730 | Overlapping |
| SARS-CoV-2 | orf1ab | EDFIPMDSTVKNYFI | 6718 | 6732 | Overlapping |
| SARS-CoV-2 | orf1ab | FIPMDSTVKNYFITD | 6720 | 6734 | Negative |
| SARS-CoV-2 | orf1ab | PMDSTVKNYFITDAQ | 6722 | 6736 | Overlapping |
| SARS-CoV-2 | orf1ab | CSVIDLLLDDFVEII | 6744 | 6758 | Positive |
| SARS-CoV-2 | orf1ab | LLDDFVEIIKSQDLS | 6750 | 6764 | Overlapping |
| SARS-CoV-2 | orf1ab | DDFVEIIKSQDLSVV | 6752 | 6766 | Overlapping |
| SARS-CoV-2 | orf1ab | FVEIIKSQDLSVVSK | 6754 | 6768 | Positive |
| SARS-CoV-2 | orf1ab | EIIKSQDLSVVSKVV | 6756 | 6770 | Negative |
| SARS-CoV-2 | orf1ab | EIIKSQDLSVVSKVV | 6756 | 6770 | Overlapping |
| SARS-CoV-2 | orf1ab | EISFMLWCKDGHVET | 6778 | 6792 | Overlapping |

| | | | | | |
|---|---|---|---|---|---|
| SARS-CoV-2 | orf1ab | KDGHVETFYPKLQSS | 6786 | 6800 | Positive |
| SARS-CoV-2 | orf1ab | GHVETFYPKLQSSQA | 6788 | 6802 | Overlapping |
| SARS-CoV-2 | orf1ab | QPGVAMPNLYKMQRM | 6804 | 6818 | Negative |
| SARS-CoV-2 | orf1ab | GVAMPNLYKMQRMLL | 6806 | 6820 | Negative |
| SARS-CoV-2 | orf1ab | DLQNYGDSATLPKGI | 6824 | 6838 | Negative |
| SARS-CoV-2 | orf1ab | NMRVIHFGAGSDKGV | 6862 | 6876 | Positive |
| SARS-CoV-2 | orf1ab | SDKGVAPGTAVLRQW | 6872 | 6886 | Positive |
| SARS-CoV-2 | orf1ab | PTGTLLVDSDLNDFV | 6888 | 6902 | Positive |
| SARS-CoV-2 | orf1ab | GTLLVDSDLNDFVSD | 6890 | 6904 | Positive |
| SARS-CoV-2 | orf1ab | LLVDSDLNDFVSDAD | 6892 | 6906 | Positive |
| SARS-CoV-2 | orf1ab | VDSDLNDFVSDADST | 6894 | 6908 | Positive |
| SARS-CoV-2 | orf1ab | SDLNDFVSDADSTLI | 6896 | 6910 | Positive |
| SARS-CoV-2 | orf1ab | DFVSDADSTLIGDCA | 6900 | 6914 | Positive |
| SARS-CoV-2 | orf1ab | VSDADSTLIGDCATV | 6902 | 6916 | Positive |
| SARS-CoV-2 | orf1ab | DADSTLIGDCATVHT | 6904 | 6918 | Positive |
| SARS-CoV-2 | orf1ab | NKWDLIISDMYDPKT | 6920 | 6934 | Positive |
| SARS-CoV-2 | orf1ab | NVTKENDSKEGFFTY | 6936 | 6950 | Positive |
| SARS-CoV-2 | orf1ab | NVTKENDSKEGFFTY | 6936 | 6950 | Negative |
| SARS-CoV-2 | orf1ab | TKENDSKEGFFTYIC | 6938 | 6952 | Positive |
| SARS-CoV-2 | orf1ab | ENDSKEGFFTYICGF | 6940 | 6954 | Overlapping |
| SARS-CoV-2 | orf1ab | TEHSWNADLYKLMGH | 6970 | 6984 | Positive |
| SARS-CoV-2 | orf1ab | VTNVNASSSEAFLIG | 6992 | 7006 | Positive |
| SARS-CoV-2 | orf1ab | NVNASSSEAFLIGCN | 6994 | 7008 | Positive |
| SARS-CoV-2 | orf1ab | NASSSEAFLIGCNYL | 6996 | 7010 | Positive |
| SARS-CoV-2 | orf1ab | SEAFLIGCNYLGKPR | 7000 | 7014 | Positive |
| SARS-CoV-2 | orf1ab | LIGCNYLGKPREQID | 7004 | 7018 | Negative |
| SARS-CoV-2 | orf1ab | NYLGKPREQIDGYVM | 7008 | 7022 | Positive |
| SARS-CoV-2 | orf1ab | DGYVMHANYIFWRNT | 7018 | 7032 | Positive |
| SARS-CoV-2 | orf1ab | PIQLSSYSLFDMSKF | 7034 | 7048 | Negative |
| SARS-CoV-2 | orf1ab | LRGTAVMSLKEGQIN | 7052 | 7066 | Negative |
| SARS-CoV-2 | orf1ab | GTAVMSLKEGQINDM | 7054 | 7068 | Negative |
| SARS-CoV-2 | orf1ab | LKEGQINDMILSLLS | 7060 | 7074 | Overlapping |
| SARS-CoV-2 | orf1ab | EGQINDMILSLLSKG | 7062 | 7076 | Negative |
| SARS-CoV-2 | orf1ab | EGQINDMILSLLSKG | 7062 | 7076 | Overlapping |
| SARS-CoV-2 | orf1ab | LSLLSKGRLIIRENN | 7070 | 7084 | Positive |
| SARS-CoV-2 | orf1ab | LIIRENNRVVISSDV | 7078 | 7092 | Positive |
| SARS-CoV-2 | orf1ab | IRENNRVVISSDVLV | 7080 | 7094 | Positive |
| SARS-CoV-2 | orf1ab | ENNRVVISSDVLVNN | 7082 | 7096 | Overlapping |
| SARS-CoV-2 | orf1ab | NRVVISSDVLVNNGS | 7084 | 7096 | Positive |
| SARS-CoV-2 | orf1ab | SDVLVNNGSGSGSG | 7090 | 7096 | Positive |
| SARS-CoV-2 | spike | GSGMFVFLVLLPLVS | 1 | 12 | Positive |
| SARS-CoV-2 | spike | GSGSGMFVFLVLLPL | 1 | 10 | Positive |
| SARS-CoV-2 | spike | GSGSGSGMFVFLVLL | 1 | 8 | Positive |
| SARS-CoV-2 | spike | GSGSGMFVFLVLLPL | 1 | 10 | Negative |
| SARS-CoV-2 | spike | GSGSGSGMFVFLVLL | 1 | 8 | Negative |
| SARS-CoV-2 | spike | FLVLLPLVSSQCVNL | 4 | 18 | Negative |
| SARS-CoV-2 | spike | LPLVSSQCVNLTTRT | 8 | 22 | Negative |
| SARS-CoV-2 | spike | PLVSSQCVNLTTRTQ | 9 | 23 | Negative |
| SARS-CoV-2 | spike | NLTTRTQLPPAYTNS | 17 | 31 | Positive |

| | | | | | |
|---|---|---|---|---|---|
| SARS-CoV-2 | spike | TTRTQLPPAYTNSFT | 19 | 33 | Negative |
| SARS-CoV-2 | spike | GVYYPDKVFRSSVLH | 35 | 49 | Negative |
| SARS-CoV-2 | spike | SSVLHSTQDLFLPFF | 45 | 59 | Negative |
| SARS-CoV-2 | spike | GTNGTKRFDNPVLPF | 72 | 86 | Negative |
| SARS-CoV-2 | spike | RFDNPVLPFNDGVYF | 78 | 92 | Overlapping |
| SARS-CoV-2 | spike | FDNPVLPFNDGVYFA | 79 | 93 | Positive |
| SARS-CoV-2 | spike | PVLPFNDGVYFASTE | 82 | 96 | Overlapping |
| SARS-CoV-2 | spike | FNDGVYFASTEKSNI | 86 | 100 | Negative |
| SARS-CoV-2 | spike | NDGVYFASTEKSNII | 87 | 101 | Positive |
| SARS-CoV-2 | spike | GVYFASTEKSNIIRG | 89 | 103 | Negative |
| SARS-CoV-2 | spike | TEKSNIIRGWIFGTT | 95 | 109 | Negative |
| SARS-CoV-2 | spike | EKSNIIRGWIFGTTL | 96 | 110 | Overlapping |
| SARS-CoV-2 | spike | GTTLDSKTQSLLIVN | 107 | 121 | Negative |
| SARS-CoV-2 | spike | DSKTQSLLIVNNATN | 111 | 125 | Overlapping |
| SARS-CoV-2 | spike | TQSLLIVNNATNVVI | 114 | 128 | Negative |
| SARS-CoV-2 | spike | LIVNNATNVVIKVCE | 118 | 132 | Overlapping |
| SARS-CoV-2 | spike | NNATNVVIKVCEFQF | 121 | 135 | Positive |
| SARS-CoV-2 | spike | NATNVVIKVCEFQFC | 122 | 136 | Positive |
| SARS-CoV-2 | spike | NVVIKVCEFQFCNDP | 125 | 139 | Positive |
| SARS-CoV-2 | spike | VIKVCEFQFCNDPFL | 127 | 141 | Negative |
| SARS-CoV-2 | spike | EFQFCNDPFLGVYYH | 132 | 146 | Overlapping |
| SARS-CoV-2 | spike | FCNDPFLGVYYHKNN | 135 | 149 | Positive |
| SARS-CoV-2 | spike | NDPFLGVYYHKNNKS | 137 | 151 | Overlapping |
| SARS-CoV-2 | spike | GVYYHKNNKSWMESE | 142 | 156 | Overlapping |
| SARS-CoV-2 | spike | NNKSWMESEFRVYSS | 148 | 162 | Positive |
| SARS-CoV-2 | spike | NKSWMESEFRVYSSA | 149 | 163 | Positive |
| SARS-CoV-2 | spike | MESEFRVYSSANNCT | 153 | 167 | Overlapping |
| SARS-CoV-2 | spike | ESEFRVYSSANNCTF | 154 | 168 | Overlapping |
| SARS-CoV-2 | spike | SEFRVYSSANNCTFE | 155 | 169 | Positive |
| SARS-CoV-2 | spike | EFRVYSSANNCTFEY | 156 | 170 | Overlapping |
| SARS-CoV-2 | spike | NNCTFEYVSQPFLMD | 164 | 178 | Positive |
| SARS-CoV-2 | spike | NCTFEYVSQPFLMDL | 165 | 179 | Positive |
| SARS-CoV-2 | spike | FEYVSQPFLMDLEGK | 168 | 182 | Overlapping |
| SARS-CoV-2 | spike | EYVSQPFLMDLEGKQ | 169 | 183 | Overlapping |
| SARS-CoV-2 | spike | MDLEGKQGNFKNLRE | 177 | 191 | Negative |
| SARS-CoV-2 | spike | EGKQGNFKNLREFVF | 180 | 194 | Negative |
| SARS-CoV-2 | spike | EGKQGNFKNLREFVF | 180 | 194 | Overlapping |
| SARS-CoV-2 | spike | NFKNLREFVFKNIDG | 185 | 199 | Positive |
| SARS-CoV-2 | spike | NLREFVFKNIDGYFK | 188 | 202 | Positive |
| SARS-CoV-2 | spike | REFVFKNIDGYFKIY | 190 | 204 | Positive |
| SARS-CoV-2 | spike | EFVFKNIDGYFKIYS | 191 | 205 | Overlapping |
| SARS-CoV-2 | spike | KNIDGYFKIYSKHTP | 195 | 209 | Positive |
| SARS-CoV-2 | spike | NLVRDLPQGFSALEP | 211 | 225 | Positive |
| SARS-CoV-2 | spike | RDLPQGFSALEPLVD | 214 | 228 | Positive |
| SARS-CoV-2 | spike | GFSALEPLVDLPIGI | 219 | 233 | Negative |
| SARS-CoV-2 | spike | SALEPLVDLPIGINI | 221 | 235 | Positive |
| SARS-CoV-2 | spike | EPLVDLPIGINITRF | 224 | 238 | Overlapping |
| SARS-CoV-2 | spike | PLVDLPIGINITRFQ | 225 | 239 | Positive |
| SARS-CoV-2 | spike | PIGINITRFQTLLAL | 230 | 244 | Positive |

| | | | | | |
|---|---|---|---|---|---|
| SARS-CoV-2 | spike | GINITRFQTLLALHR | 232 | 246 | Negative |
| SARS-CoV-2 | spike | QPRTFLLKYNENGTI | 271 | 285 | Positive |
| SARS-CoV-2 | spike | PRTFLLKYNENGTIT | 272 | 286 | Positive |
| SARS-CoV-2 | spike | RTFLLKYNENGTITD | 273 | 287 | Positive |
| SARS-CoV-2 | spike | FLLKYNENGTITDAV | 275 | 289 | Positive |
| SARS-CoV-2 | spike | LLKYNENGTITDAVD | 276 | 290 | Positive |
| SARS-CoV-2 | spike | LKYNENGTITDAVDC | 277 | 291 | Positive |
| SARS-CoV-2 | spike | ENGTITDAVDCALDP | 281 | 295 | Overlapping |
| SARS-CoV-2 | spike | GTITDAVDCALDPLS | 283 | 297 | Negative |
| SARS-CoV-2 | spike | TITDAVDCALDPLSE | 284 | 298 | Positive |
| SARS-CoV-2 | spike | ITDAVDCALDPLSET | 285 | 299 | Overlapping |
| SARS-CoV-2 | spike | TDAVDCALDPLSETK | 286 | 300 | Positive |
| SARS-CoV-2 | spike | AVDCALDPLSETKCT | 288 | 302 | Positive |
| SARS-CoV-2 | spike | PLSETKCTLKSFTVE | 295 | 309 | Positive |
| SARS-CoV-2 | spike | ETKCTLKSFTVEKGI | 298 | 312 | Overlapping |
| SARS-CoV-2 | spike | VEKGIYQTSNFRVQP | 308 | 322 | Overlapping |
| SARS-CoV-2 | spike | EKGIYQTSNFRVQPT | 309 | 323 | Overlapping |
| SARS-CoV-2 | spike | YQTSNFRVQPTESIV | 313 | 327 | Positive |
| SARS-CoV-2 | spike | NFRVQPTESIVRFPN | 317 | 331 | Positive |
| SARS-CoV-2 | spike | VQPTESIVRFPNITN | 320 | 334 | Negative |
| SARS-CoV-2 | spike | PTESIVRFPNITNLC | 322 | 336 | Positive |
| SARS-CoV-2 | spike | ESIVRFPNITNLCPF | 324 | 338 | Overlapping |
| SARS-CoV-2 | spike | IVRFPNITNLCPFGE | 326 | 340 | Negative |
| SARS-CoV-2 | spike | VRFPNITNLCPFGEV | 327 | 341 | Positive |
| SARS-CoV-2 | spike | PNITNLCPFGEVFNA | 330 | 344 | Positive |
| SARS-CoV-2 | spike | GEVFNATRFASVYAW | 339 | 353 | Overlapping |
| SARS-CoV-2 | spike | EVFNATRFASVYAWN | 340 | 354 | Overlapping |
| SARS-CoV-2 | spike | NATRFASVYAWNRKR | 343 | 357 | Positive |
| SARS-CoV-2 | spike | NRKRISNCVADYSVL | 354 | 368 | Positive |
| SARS-CoV-2 | spike | NSASFSTFKCYGVSP | 370 | 384 | Positive |
| SARS-CoV-2 | spike | GVSPTKLNDLCFTNV | 381 | 395 | Negative |
| SARS-CoV-2 | spike | KLNDLCFTNVYADSF | 386 | 400 | Positive |
| SARS-CoV-2 | spike | NDLCFTNVYADSFVI | 388 | 402 | Positive |
| SARS-CoV-2 | spike | TNVYADSFVIRGDEV | 393 | 407 | Negative |
| SARS-CoV-2 | spike | NVYADSFVIRGDEVR | 394 | 408 | Negative |
| SARS-CoV-2 | spike | VYADSFVIRGDEVRQ | 395 | 409 | Negative |
| SARS-CoV-2 | spike | YADSFVIRGDEVRQI | 396 | 410 | Negative |
| SARS-CoV-2 | spike | VIRGDEVRQIAPGQT | 401 | 415 | Negative |
| SARS-CoV-2 | spike | RGDEVRQIAPGQTGK | 403 | 417 | Positive |
| SARS-CoV-2 | spike | DEVRQIAPGQTGKIA | 405 | 419 | Overlapping |
| SARS-CoV-2 | spike | EVRQIAPGQTGKIAD | 406 | 420 | Overlapping |
| SARS-CoV-2 | spike | RQIAPGQTGKIADYN | 408 | 422 | Positive |
| SARS-CoV-2 | spike | TGKIADYNYKLPDDF | 415 | 429 | Positive |
| SARS-CoV-2 | spike | GKIADYNYKLPDDFT | 416 | 430 | Negative |
| SARS-CoV-2 | spike | NYKLPDDFTGCVIAW | 422 | 436 | Positive |
| SARS-CoV-2 | spike | NSNNLDSKVGGNYNY | 437 | 451 | Positive |
| SARS-CoV-2 | spike | SNNLDSKVGGNYNYL | 438 | 452 | Positive |
| SARS-CoV-2 | spike | NNLDSKVGGNYNYLY | 439 | 453 | Positive |
| SARS-CoV-2 | spike | NLDSKVGGNYNYLYR | 440 | 454 | Positive |

| | | | | | |
|---|---|---|---|---|---|
| SARS-CoV-2 | spike | KVGGNYNYLYRLFRK | 444 | 458 | Positive |
| SARS-CoV-2 | spike | YRLFRKSNLKPFERD | 453 | 467 | Negative |
| SARS-CoV-2 | spike | SNLKPFERDISTEIY | 459 | 473 | Positive |
| SARS-CoV-2 | spike | NLKPFERDISTEIYQ | 460 | 474 | Positive |
| SARS-CoV-2 | spike | PFERDISTEIYQAGS | 463 | 477 | Negative |
| SARS-CoV-2 | spike | FERDISTEIYQAGST | 464 | 478 | Negative |
| SARS-CoV-2 | spike | ERDISTEIYQAGSTP | 465 | 479 | Overlapping |
| SARS-CoV-2 | spike | RDISTEIYQAGSTPC | 466 | 480 | Positive |
| SARS-CoV-2 | spike | TEIYQAGSTPCNGVE | 470 | 484 | Overlapping |
| SARS-CoV-2 | spike | EIYQAGSTPCNGVEG | 471 | 485 | Overlapping |
| SARS-CoV-2 | spike | IYQAGSTPCNGVEGF | 472 | 486 | Negative |
| SARS-CoV-2 | spike | GSTPCNGVEGFNCYF | 476 | 490 | Negative |
| SARS-CoV-2 | spike | GVEGFNCYFPLQSYG | 482 | 496 | Negative |
| SARS-CoV-2 | spike | VEGFNCYFPLQSYGF | 483 | 497 | Positive |
| SARS-CoV-2 | spike | EGFNCYFPLQSYGFQ | 484 | 498 | Negative |
| SARS-CoV-2 | spike | EGFNCYFPLQSYGFQ | 484 | 498 | Overlapping |
| SARS-CoV-2 | spike | PTNGVGYQPYRVVVL | 499 | 513 | Positive |
| SARS-CoV-2 | spike | GVGYQPYRVVVLSFE | 502 | 516 | Overlapping |
| SARS-CoV-2 | spike | FELLHAPATVCGPKK | 515 | 529 | Positive |
| SARS-CoV-2 | spike | ELLHAPATVCGPKKS | 516 | 530 | Negative |
| SARS-CoV-2 | spike | ELLHAPATVCGPKKS | 516 | 530 | Overlapping |
| SARS-CoV-2 | spike | APATVCGPKKSTNLV | 520 | 534 | Negative |
| SARS-CoV-2 | spike | CGPKKSTNLVKNKCV | 525 | 539 | Positive |
| SARS-CoV-2 | spike | NLVKNKCVNFNFNGL | 532 | 546 | Positive |
| SARS-CoV-2 | spike | NKCVNFNFNGLTGTG | 536 | 550 | Positive |
| SARS-CoV-2 | spike | KCVNFNFNGLTGTGV | 537 | 551 | Negative |
| SARS-CoV-2 | spike | CVNFNFNGLTGTGVL | 538 | 552 | Negative |
| SARS-CoV-2 | spike | NFNFNGLTGTGVLTE | 540 | 554 | Negative |
| SARS-CoV-2 | spike | NFNGLTGTGVLTESN | 542 | 556 | Positive |
| SARS-CoV-2 | spike | NGLTGTGVLTESNKK | 544 | 558 | Positive |
| SARS-CoV-2 | spike | GLTGTGVLTESNKKF | 545 | 559 | Negative |
| SARS-CoV-2 | spike | GTGVLTESNKKFLPF | 548 | 562 | Overlapping |
| SARS-CoV-2 | spike | TGVLTESNKKFLPFQ | 549 | 563 | Negative |
| SARS-CoV-2 | spike | GVLTESNKKFLPFQQ | 550 | 564 | Negative |
| SARS-CoV-2 | spike | LTESNKKFLPFQQFG | 552 | 566 | Negative |
| SARS-CoV-2 | spike | ESNKKFLPFQQFGRD | 554 | 568 | Overlapping |
| SARS-CoV-2 | spike | NKKFLPFQQFGRDIA | 556 | 570 | Positive |
| SARS-CoV-2 | spike | FQQFGRDIADTTDAV | 562 | 576 | Negative |
| SARS-CoV-2 | spike | QQFGRDIADTTDAVR | 563 | 577 | Negative |
| SARS-CoV-2 | spike | QFGRDIADTTDAVRD | 564 | 578 | Negative |
| SARS-CoV-2 | spike | GRDIADTTDAVRDPQ | 566 | 580 | Negative |
| SARS-CoV-2 | spike | RDIADTTDAVRDPQT | 567 | 581 | Positive |
| SARS-CoV-2 | spike | ADTTDAVRDPQTLEI | 570 | 584 | Positive |
| SARS-CoV-2 | spike | TDAVRDPQTLEILDI | 573 | 587 | Positive |
| SARS-CoV-2 | spike | DAVRDPQTLEILDIT | 574 | 588 | Positive |
| SARS-CoV-2 | spike | VRDPQTLEILDITPC | 576 | 590 | Negative |
| SARS-CoV-2 | spike | RDPQTLEILDITPCS | 577 | 591 | Overlapping |
| SARS-CoV-2 | spike | TLEILDITPCSFGGV | 581 | 595 | Overlapping |
| SARS-CoV-2 | spike | LEILDITPCSFGGVS | 582 | 596 | Overlapping |

| | | | | | |
|---|---|---|---|---|---|
| SARS-CoV-2 | spike | EILDITPCSFGGVSV | 583 | 597 | Negative |
| SARS-CoV-2 | spike | EILDITPCSFGGVSV | 583 | 597 | Overlapping |
| SARS-CoV-2 | spike | SFGGVSVITPGTNTS | 591 | 605 | Negative |
| SARS-CoV-2 | spike | FGGVSVITPGTNTSN | 592 | 606 | Negative |
| SARS-CoV-2 | spike | GGVSVITPGTNTSNQ | 593 | 607 | Negative |
| SARS-CoV-2 | spike | GVSVITPGTNTSNQV | 594 | 608 | Negative |
| SARS-CoV-2 | spike | VSVITPGTNTSNQVA | 595 | 609 | Negative |
| SARS-CoV-2 | spike | SVITPGTNTSNQVAV | 596 | 610 | Negative |
| SARS-CoV-2 | spike | ITPGTNTSNQVAVLY | 598 | 612 | Negative |
| SARS-CoV-2 | spike | GTNTSNQVAVLYQDV | 601 | 615 | Negative |
| SARS-CoV-2 | spike | TNTSNQVAVLYQDVN | 602 | 616 | Negative |
| SARS-CoV-2 | spike | NTSNQVAVLYQDVNC | 603 | 617 | Negative |
| SARS-CoV-2 | spike | SNQVAVLYQDVNCTE | 605 | 619 | Negative |
| SARS-CoV-2 | spike | NQVAVLYQDVNCTEV | 606 | 620 | Positive |
| SARS-CoV-2 | spike | NCTEVPVAIHADQLT | 616 | 630 | Positive |
| SARS-CoV-2 | spike | TEVPVAIHADQLTPT | 618 | 632 | Overlapping |
| SARS-CoV-2 | spike | EVPVAIHADQLTPTW | 619 | 633 | Overlapping |
| SARS-CoV-2 | spike | PVAIHADQLTPTWRV | 621 | 635 | Overlapping |
| SARS-CoV-2 | spike | CLIGAEHVNNSYECD | 649 | 663 | Positive |
| SARS-CoV-2 | spike | IGAEHVNNSYECDIP | 651 | 665 | Positive |
| SARS-CoV-2 | spike | EHVNNSYECDIPIGA | 654 | 668 | Overlapping |
| SARS-CoV-2 | spike | VNNSYECDIPIGAGI | 656 | 670 | Negative |
| SARS-CoV-2 | spike | NNSYECDIPIGAGIC | 657 | 671 | Positive |
| SARS-CoV-2 | spike | YECDIPIGAGICASY | 660 | 674 | Positive |
| SARS-CoV-2 | spike | ECDIPIGAGICASYQ | 661 | 675 | Overlapping |
| SARS-CoV-2 | spike | IPIGAGICASYQTQT | 664 | 678 | Positive |
| SARS-CoV-2 | spike | PIGAGICASYQTQTN | 665 | 679 | Positive |
| SARS-CoV-2 | spike | CASYQTQTNSPRRAR | 671 | 685 | Positive |
| SARS-CoV-2 | spike | TQTNSPRRARSVASQ | 676 | 690 | Negative |
| SARS-CoV-2 | spike | SPRRARSVASQSIIA | 680 | 694 | Negative |
| SARS-CoV-2 | spike | PRRARSVASQSIIAY | 681 | 695 | Negative |
| SARS-CoV-2 | spike | ASQSIIAYTMSLGAE | 688 | 702 | Negative |
| SARS-CoV-2 | spike | SQSIIAYTMSLGAEN | 689 | 703 | Positive |
| SARS-CoV-2 | spike | TMSLGAENSVAYSNN | 696 | 710 | Positive |
| SARS-CoV-2 | spike | GAENSVAYSNNSIAI | 700 | 714 | Positive |
| SARS-CoV-2 | spike | ENSVAYSNNSIAIPT | 702 | 716 | Overlapping |
| SARS-CoV-2 | spike | NSVAYSNNSIAIPTN | 703 | 717 | Positive |
| SARS-CoV-2 | spike | NNSIAIPTNFTISVT | 709 | 723 | Positive |
| SARS-CoV-2 | spike | IPTNFTISVTTEILP | 714 | 728 | Positive |
| SARS-CoV-2 | spike | PTNFTISVTTEILPV | 715 | 729 | Positive |
| SARS-CoV-2 | spike | TISVTTEILPVSMTK | 719 | 733 | Positive |
| SARS-CoV-2 | spike | ISVTTEILPVSMTKT | 720 | 734 | Negative |
| SARS-CoV-2 | spike | SVTTEILPVSMTKTS | 721 | 735 | Negative |
| SARS-CoV-2 | spike | VTTEILPVSMTKTSV | 722 | 736 | Positive |
| SARS-CoV-2 | spike | TEILPVSMTKTSVDC | 724 | 738 | Positive |
| SARS-CoV-2 | spike | EILPVSMTKTSVDCT | 725 | 739 | Negative |
| SARS-CoV-2 | spike | EILPVSMTKTSVDCT | 725 | 739 | Overlapping |
| SARS-CoV-2 | spike | ILPVSMTKTSVDCTM | 726 | 740 | Negative |
| SARS-CoV-2 | spike | PVSMTKTSVDCTMYI | 728 | 742 | Positive |

| | | | | | |
|---|---|---|---|---|---|
| SARS-CoV-2 | spike | PVSMTKTSVDCTMYI | 728 | 742 | Overlapping |
| SARS-CoV-2 | spike | SMTKTSVDCTMYICG | 730 | 744 | Positive |
| SARS-CoV-2 | spike | TMYICGDSTECSNLL | 739 | 753 | Positive |
| SARS-CoV-2 | spike | ECSNLLLQYGSFCTQ | 748 | 762 | Overlapping |
| SARS-CoV-2 | spike | NLLLQYGSFCTQLNR | 751 | 765 | Positive |
| SARS-CoV-2 | spike | YGSFCTQLNRALTGI | 756 | 770 | Negative |
| SARS-CoV-2 | spike | GSFCTQLNRALTGIA | 757 | 771 | Negative |
| SARS-CoV-2 | spike | FCTQLNRALTGIAVE | 759 | 773 | Negative |
| SARS-CoV-2 | spike | QLNRALTGIAVEQDK | 762 | 776 | Positive |
| SARS-CoV-2 | spike | LNRALTGIAVEQDKN | 763 | 777 | Positive |
| SARS-CoV-2 | spike | NRALTGIAVEQDKNT | 764 | 778 | Positive |
| SARS-CoV-2 | spike | RALTGIAVEQDKNTQ | 765 | 779 | Positive |
| SARS-CoV-2 | spike | ALTGIAVEQDKNTQE | 766 | 780 | Positive |
| SARS-CoV-2 | spike | LTGIAVEQDKNTQEV | 767 | 781 | Positive |
| SARS-CoV-2 | spike | TGIAVEQDKNTQEVF | 768 | 782 | Positive |
| SARS-CoV-2 | spike | GIAVEQDKNTQEVFA | 769 | 783 | Negative |
| SARS-CoV-2 | spike | IAVEQDKNTQEVFAQ | 770 | 784 | Positive |
| SARS-CoV-2 | spike | AVEQDKNTQEVFAQV | 771 | 785 | Overlapping |
| SARS-CoV-2 | spike | VEQDKNTQEVFAQVK | 772 | 786 | Overlapping |
| SARS-CoV-2 | spike | EQDKNTQEVFAQVKQ | 773 | 787 | Overlapping |
| SARS-CoV-2 | spike | QEVFAQVKQIYKTPP | 779 | 793 | Negative |
| SARS-CoV-2 | spike | ILPDPSKPSKRSFIE | 805 | 819 | Negative |
| SARS-CoV-2 | spike | PSKRSFIEDLLFNKV | 812 | 826 | Positive |
| SARS-CoV-2 | spike | SKRSFIEDLLFNKVT | 813 | 827 | Negative |
| SARS-CoV-2 | spike | FIEDLLFNKVTLADA | 817 | 831 | Positive |
| SARS-CoV-2 | spike | IEDLLFNKVTLADAG | 818 | 832 | Overlapping |
| SARS-CoV-2 | spike | EDLLFNKVTLADAGF | 819 | 833 | Overlapping |
| SARS-CoV-2 | spike | NKVTLADAGFIKQYG | 824 | 838 | Positive |
| SARS-CoV-2 | spike | ADAGFIKQYGDCLGD | 829 | 843 | Negative |
| SARS-CoV-2 | spike | DAGFIKQYGDCLGDI | 830 | 844 | Positive |
| SARS-CoV-2 | spike | AGFIKQYGDCLGDIA | 831 | 845 | Positive |
| SARS-CoV-2 | spike | ARDLICAQKFNGLTV | 846 | 860 | Positive |
| SARS-CoV-2 | spike | RDLICAQKFNGLTVL | 847 | 861 | Positive |
| SARS-CoV-2 | spike | NGLTVLPPLLTDEMI | 856 | 870 | Positive |
| SARS-CoV-2 | spike | GLTVLPPLLTDEMIA | 857 | 871 | Negative |
| SARS-CoV-2 | spike | PLLTDEMIAQYTSAL | 863 | 877 | Positive |
| SARS-CoV-2 | spike | LLTDEMIAQYTSALL | 864 | 878 | Positive |
| SARS-CoV-2 | spike | LTDEMIAQYTSALLA | 865 | 879 | Negative |
| SARS-CoV-2 | spike | TDEMIAQYTSALLAG | 866 | 880 | Positive |
| SARS-CoV-2 | spike | DEMIAQYTSALLAGT | 867 | 881 | Negative |
| SARS-CoV-2 | spike | EMIAQYTSALLAGTI | 868 | 882 | Overlapping |
| SARS-CoV-2 | spike | TFGAGAALQIPFAMQ | 887 | 901 | Positive |
| SARS-CoV-2 | spike | IPFAMQMAYRFNGIG | 896 | 910 | Positive |
| SARS-CoV-2 | spike | PFAMQMAYRFNGIGV | 897 | 911 | Negative |
| SARS-CoV-2 | spike | FAMQMAYRFNGIGVT | 898 | 912 | Negative |
| SARS-CoV-2 | spike | AMQMAYRFNGIGVTQ | 899 | 913 | Negative |
| SARS-CoV-2 | spike | MQMAYRFNGIGVTQN | 900 | 914 | Negative |
| SARS-CoV-2 | spike | GIGVTQNVLYENQKL | 908 | 922 | Negative |
| SARS-CoV-2 | spike | IGVTQNVLYENQKLI | 909 | 923 | Negative |

| | | | | | |
|---|---|---|---|---|---|
| SARS-CoV-2 | spike | GVTQNVLYENQKLIA | 910 | 924 | Negative |
| SARS-CoV-2 | spike | NVLYENQKLIANQFN | 914 | 928 | Positive |
| SARS-CoV-2 | spike | YENQKLIANQFNSAI | 917 | 931 | Positive |
| SARS-CoV-2 | spike | ENQKLIANQFNSAIG | 918 | 932 | Overlapping |
| SARS-CoV-2 | spike | NQKLIANQFNSAIGK | 919 | 933 | Positive |
| SARS-CoV-2 | spike | NSAIGKIQDSLSSTA | 928 | 942 | Positive |
| SARS-CoV-2 | spike | GKIQDSLSSTASALG | 932 | 946 | Negative |
| SARS-CoV-2 | spike | DSLSSTASALGKLQD | 936 | 950 | Positive |
| SARS-CoV-2 | spike | NAQALNTLVKQLSSN | 955 | 969 | Positive |
| SARS-CoV-2 | spike | SSVLNDILSRLDKVE | 974 | 988 | Negative |
| SARS-CoV-2 | spike | VLNDILSRLDKVEAE | 976 | 990 | Negative |
| SARS-CoV-2 | spike | LNDILSRLDKVEAEV | 977 | 991 | Positive |
| SARS-CoV-2 | spike | NDILSRLDKVEAEVQ | 978 | 992 | Positive |
| SARS-CoV-2 | spike | DILSRLDKVEAEVQI | 979 | 993 | Positive |
| SARS-CoV-2 | spike | ILSRLDKVEAEVQID | 980 | 994 | Positive |
| SARS-CoV-2 | spike | KVEAEVQIDRLITGR | 986 | 1000 | Negative |
| SARS-CoV-2 | spike | VEAEVQIDRLITGRL | 987 | 1001 | Positive |
| SARS-CoV-2 | spike | EAEVQIDRLITGRLQ | 988 | 1002 | Overlapping |
| SARS-CoV-2 | spike | AEVQIDRLITGRLQS | 989 | 1003 | Positive |
| SARS-CoV-2 | spike | EVQIDRLITGRLQSL | 990 | 1004 | Overlapping |
| SARS-CoV-2 | spike | LQTYVTQQLIRAAEI | 1004 | 1018 | Overlapping |
| SARS-CoV-2 | spike | QTYVTQQLIRAAEIR | 1005 | 1019 | Negative |
| SARS-CoV-2 | spike | EIRASANLAATKMSE | 1017 | 1031 | Overlapping |
| SARS-CoV-2 | spike | NLAATKMSECVLGQS | 1023 | 1037 | Positive |
| SARS-CoV-2 | spike | LAATKMSECVLGQSK | 1024 | 1038 | Positive |
| SARS-CoV-2 | spike | SECVLGQSKRVDFCG | 1030 | 1044 | Positive |
| SARS-CoV-2 | spike | ECVLGQSKRVDFCGK | 1031 | 1045 | Overlapping |
| SARS-CoV-2 | spike | RVDFCGKGYHLMSFP | 1039 | 1053 | Overlapping |
| SARS-CoV-2 | spike | GYHLMSFPQSAPHGV | 1046 | 1060 | Overlapping |
| SARS-CoV-2 | spike | YHLMSFPQSAPHGVV | 1047 | 1061 | Overlapping |
| SARS-CoV-2 | spike | HLMSFPQSAPHGVVF | 1048 | 1062 | Overlapping |
| SARS-CoV-2 | spike | MSFPQSAPHGVVFLH | 1050 | 1064 | Positive |
| SARS-CoV-2 | spike | PHGVVFLHVTYVPAQ | 1057 | 1071 | Positive |
| SARS-CoV-2 | spike | GVVFLHVTYVPAQEK | 1059 | 1073 | Negative |
| SARS-CoV-2 | spike | TYVPAQEKNFTTAPA | 1066 | 1080 | Positive |
| SARS-CoV-2 | spike | PAQEKNFTTAPAICH | 1069 | 1083 | Positive |
| SARS-CoV-2 | spike | AQEKNFTTAPAICHD | 1070 | 1084 | Positive |
| SARS-CoV-2 | spike | QEKNFTTAPAICHDG | 1071 | 1085 | Overlapping |
| SARS-CoV-2 | spike | EKNFTTAPAICHDGK | 1072 | 1086 | Overlapping |
| SARS-CoV-2 | spike | KNFTTAPAICHDGKA | 1073 | 1087 | Positive |
| SARS-CoV-2 | spike | NFTTAPAICHDGKAH | 1074 | 1088 | Positive |
| SARS-CoV-2 | spike | FTTAPAICHDGKAHF | 1075 | 1089 | Positive |
| SARS-CoV-2 | spike | TTAPAICHDGKAHFP | 1076 | 1090 | Positive |
| SARS-CoV-2 | spike | TAPAICHDGKAHFPR | 1077 | 1091 | Positive |
| SARS-CoV-2 | spike | APAICHDGKAHFPRE | 1078 | 1092 | Positive |
| SARS-CoV-2 | spike | PAICHDGKAHFPREG | 1079 | 1093 | Positive |
| SARS-CoV-2 | spike | AICHDGKAHFPREGV | 1080 | 1094 | Negative |
| SARS-CoV-2 | spike | ICHDGKAHFPREGVF | 1081 | 1095 | Positive |
| SARS-CoV-2 | spike | CHDGKAHFPREGVFV | 1082 | 1096 | Positive |

| | | | | | |
|---|---|---|---|---|---|
| SARS-CoV-2 | spike | HDGKAHFPREGVFVS | 1083 | 1097 | Positive |
| SARS-CoV-2 | spike | GKAHFPREGVFVSNG | 1085 | 1099 | Negative |
| SARS-CoV-2 | spike | AHFPREGVFVSNGTH | 1087 | 1101 | Positive |
| SARS-CoV-2 | spike | REGVFVSNGTHWFVT | 1091 | 1105 | Positive |
| SARS-CoV-2 | spike | EGVFVSNGTHWFVTQ | 1092 | 1106 | Overlapping |
| SARS-CoV-2 | spike | GVFVSNGTHWFVTQR | 1093 | 1107 | Negative |
| SARS-CoV-2 | spike | NGTHWFVTQRNFYEP | 1098 | 1112 | Overlapping |
| SARS-CoV-2 | spike | GTHWFVTQRNFYEPQ | 1099 | 1113 | Negative |
| SARS-CoV-2 | spike | TQRNFYEPQIITTDN | 1105 | 1119 | Negative |
| SARS-CoV-2 | spike | QRNFYEPQIITTDNT | 1106 | 1120 | Negative |
| SARS-CoV-2 | spike | NFYEPQIITTDNTFV | 1108 | 1122 | Positive |
| SARS-CoV-2 | spike | YEPQIITTDNTFVSG | 1110 | 1124 | Positive |
| SARS-CoV-2 | spike | EPQIITTDNTFVSGN | 1111 | 1125 | Overlapping |
| SARS-CoV-2 | spike | TDNTFVSGNCDVVIG | 1117 | 1131 | Positive |
| SARS-CoV-2 | spike | NTFVSGNCDVVIGIV | 1119 | 1133 | Negative |
| SARS-CoV-2 | spike | TFVSGNCDVVIGIVN | 1120 | 1134 | Negative |
| SARS-CoV-2 | spike | VIGIVNNTVYDPLQP | 1129 | 1143 | Positive |
| SARS-CoV-2 | spike | GIVNNTVYDPLQPEL | 1131 | 1145 | Overlapping |
| SARS-CoV-2 | spike | NNTVYDPLQPELDSF | 1134 | 1148 | Positive |
| SARS-CoV-2 | spike | VYDPLQPELDSFKEE | 1137 | 1151 | Positive |
| SARS-CoV-2 | spike | YDPLQPELDSFKEEL | 1138 | 1152 | Positive |
| SARS-CoV-2 | spike | PLQPELDSFKEELDK | 1140 | 1154 | Positive |
| SARS-CoV-2 | spike | LQPELDSFKEELDKY | 1141 | 1155 | Positive |
| SARS-CoV-2 | spike | PELDSFKEELDKYFK | 1143 | 1157 | Overlapping |
| SARS-CoV-2 | spike | ELDSFKEELDKYFKN | 1144 | 1158 | Negative |
| SARS-CoV-2 | spike | ELDSFKEELDKYFKN | 1144 | 1158 | Overlapping |
| SARS-CoV-2 | spike | DSFKEELDKYFKNHT | 1146 | 1160 | Positive |
| SARS-CoV-2 | spike | SFKEELDKYFKNHTS | 1147 | 1161 | Positive |
| SARS-CoV-2 | spike | KEELDKYFKNHTSPD | 1149 | 1163 | Positive |
| SARS-CoV-2 | spike | EELDKYFKNHTSPDV | 1150 | 1164 | Overlapping |
| SARS-CoV-2 | spike | ELDKYFKNHTSPDVD | 1151 | 1165 | Negative |
| SARS-CoV-2 | spike | ELDKYFKNHTSPDVD | 1151 | 1165 | Overlapping |
| SARS-CoV-2 | spike | LDKYFKNHTSPDVDL | 1152 | 1166 | Negative |
| SARS-CoV-2 | spike | YFKNHTSPDVDLGDI | 1155 | 1169 | Positive |
| SARS-CoV-2 | spike | FKNHTSPDVDLGDIS | 1156 | 1170 | Positive |
| SARS-CoV-2 | spike | KNHTSPDVDLGDISG | 1157 | 1171 | Positive |
| SARS-CoV-2 | spike | NHTSPDVDLGDISGI | 1158 | 1172 | Overlapping |
| SARS-CoV-2 | spike | HTSPDVDLGDISGIN | 1159 | 1173 | Negative |
| SARS-CoV-2 | spike | TSPDVDLGDISGINA | 1160 | 1174 | Overlapping |
| SARS-CoV-2 | spike | SPDVDLGDISGINAS | 1161 | 1175 | Negative |
| SARS-CoV-2 | spike | PDVDLGDISGINASV | 1162 | 1176 | Positive |
| SARS-CoV-2 | spike | DVDLGDISGINASVV | 1163 | 1177 | Positive |
| SARS-CoV-2 | spike | VDLGDISGINASVVN | 1164 | 1178 | Positive |
| SARS-CoV-2 | spike | GINASVVNIQKEIDR | 1171 | 1185 | Negative |
| SARS-CoV-2 | spike | NASVVNIQKEIDRLN | 1173 | 1187 | Positive |
| SARS-CoV-2 | spike | ASVVNIQKEIDRLNE | 1174 | 1188 | Overlapping |
| SARS-CoV-2 | spike | NIQKEIDRLNEVAKN | 1178 | 1192 | Positive |
| SARS-CoV-2 | spike | QKEIDRLNEVAKNLN | 1180 | 1194 | Positive |
| SARS-CoV-2 | spike | KEIDRLNEVAKNLNE | 1181 | 1195 | Overlapping |

| | | | | | |
|---|---|---|---|---|---|
| SARS-CoV-2 | spike | EIDRLNEVAKNLNES | 1182 | 1196 | Overlapping |
| SARS-CoV-2 | spike | NEVAKNLNESLIDLQ | 1187 | 1201 | Positive |
| SARS-CoV-2 | spike | NEVAKNLNESLIDLQ | 1187 | 1201 | Negative |
| SARS-CoV-2 | spike | EVAKNLNESLIDLQE | 1188 | 1202 | Overlapping |
| SARS-CoV-2 | spike | NLNESLIDLQELGKY | 1192 | 1206 | Positive |
| SARS-CoV-2 | spike | LNESLIDLQELGKYE | 1193 | 1207 | Overlapping |
| SARS-CoV-2 | spike | NESLIDLQELGKYEQ | 1194 | 1208 | Positive |
| SARS-CoV-2 | spike | ESLIDLQELGKYEQY | 1195 | 1209 | Overlapping |
| SARS-CoV-2 | spike | LQELGKYEQYIKWPW | 1200 | 1214 | Positive |
| SARS-CoV-2 | spike | QELGKYEQYIKWPWY | 1201 | 1215 | Negative |
| SARS-CoV-2 | spike | ELGKYEQYIKWPWYI | 1202 | 1216 | Negative |
| SARS-CoV-2 | spike | ELGKYEQYIKWPWYI | 1202 | 1216 | Overlapping |
| SARS-CoV-2 | spike | GKYEQYIKWPWYIWL | 1204 | 1218 | Negative |
| SARS-CoV-2 | spike | YEQYIKWPWYIWLGF | 1206 | 1220 | Overlapping |
| SARS-CoV-2 | spike | EQYIKWPWYIWLGFI | 1207 | 1221 | Overlapping |
| SARS-CoV-2 | spike | GLIAIVMVTIMLCCM | 1223 | 1237 | Negative |
| SARS-CoV-2 | spike | CCSCLKGCCSCGSCC | 1240 | 1254 | Positive |
| SARS-CoV-2 | spike | KGCCSCGSCCKFDED | 1245 | 1259 | Positive |
| SARS-CoV-2 | spike | GCCSCGSCCKFDEDD | 1246 | 1260 | Overlapping |
| SARS-CoV-2 | spike | CCSCGSCCKFDEDDS | 1247 | 1261 | Positive |
| SARS-CoV-2 | spike | CSCGSCCKFDEDDSE | 1248 | 1262 | Overlapping |
| SARS-CoV-2 | spike | SCGSCCKFDEDDSEP | 1249 | 1263 | Positive |
| SARS-CoV-2 | spike | CGSCCKFDEDDSEPV | 1250 | 1264 | Positive |
| SARS-CoV-2 | spike | GSCCKFDEDDSEPVL | 1251 | 1265 | Positive |
| SARS-CoV-2 | spike | SCCKFDEDDSEPVLK | 1252 | 1266 | Positive |
| SARS-CoV-2 | spike | CCKFDEDDSEPVLKG | 1253 | 1267 | Positive |
| SARS-CoV-2 | spike | CKFDEDDSEPVLKGV | 1254 | 1268 | Positive |
| SARS-CoV-2 | spike | KFDEDDSEPVLKGVK | 1255 | 1269 | Positive |
| SARS-CoV-2 | spike | KFDEDDSEPVLKGVK | 1255 | 1269 | Overlapping |
| SARS-CoV-2 | spike | FDEDDSEPVLKGVKL | 1256 | 1270 | Positive |
| SARS-CoV-2 | spike | FDEDDSEPVLKGVKL | 1256 | 1270 | Overlapping |
| SARS-CoV-2 | spike | DEDDSEPVLKGVKLH | 1257 | 1271 | Positive |
| SARS-CoV-2 | spike | EDDSEPVLKGVKLHY | 1258 | 1272 | Positive |
| SARS-CoV-2 | spike | EDDSEPVLKGVKLHY | 1258 | 1272 | Overlapping |
| SARS-CoV-2 | spike | DDSEPVLKGVKLHYT | 1259 | 1273 | Positive |
| SARS-CoV-2 | spike | SEPVLKGVKLHYTGS | 1261 | 1273 | Positive |
| SARS-CoV-2 | spike | EPVLKGVKLHYTGSG | 1262 | 1273 | Positive |

**Table S3. OC43-CoV2-specific Peptides**

| Lineage | Protein | HDPA peptide sequence | AA pos start | AA pos end | Group |
|---|---|---|---|---|---|
| hCoVOC43 | envelope | DAYLADTVWYVGQII | 5 | 19 | Positive |
| hCoVOC43 | envelope | FYNDVKPPVLDVDDV | 70 | 84 | Positive |
| hCoVOC43 | envelope | GSGMFMADAYLADTV | 1 | 12 | Positive |
| hCoVOC43 | envelope | GSGSGMFMADAYLAD | 1 | 10 | Positive |
| hCoVOC43 | envelope | GSGSGSGMFMADAYL | 1 | 8 | Positive |
| hCoVOC43 | envelope | NDVKPPVLDVDDVGS | 72 | 84 | Positive |
| hCoVOC43 | envelope | NRGRQFYEFYNDVKP | 62 | 76 | Positive |
| hCoVOC43 | envelope | PPVLDVDDVGSGSGS | 76 | 84 | Positive |
| hCoVOC43 | envelope | PVLDVDDVGSGSGSG | 77 | 84 | Positive |
| hCoVOC43 | envelope | YEFYNDVKPPVLDVD | 68 | 82 | Positive |
| hCoVOC43 | envelope | ADAYLADTVWYVGQI | 4 | 18 | Negative |
| hCoVOC43 | envelope | FYNDVKPPVLDVDDV | 70 | 84 | Negative |
| hCoVOC43 | envelope | GMCNTLVLSPSIYVF | 47 | 61 | Negative |
| hCoVOC43 | envelope | GMFMADAYLADTVWY | 1 | 14 | Negative |
| hCoVOC43 | envelope | GSGSGSGMFMADAYL | 1 | 8 | Negative |
| hCoVOC43 | envelope | SPSIYVFNRGRQFYE | 55 | 69 | Negative |
| hCoVOC43 | envelope | EFYNDVKPPVLDVDD | 69 | 83 | Overlapping |
| hCoVOC43 | envelope | PVLDVDDVGSGSGSG | 77 | 84 | Overlapping |
| hCoVOC43 | membrane | ADEAIKFLKEWNFSL | 15 | 29 | Positive |
| hCoVOC43 | membrane | DEAIKFLKEWNFSLG | 16 | 30 | Positive |
| hCoVOC43 | membrane | DMKGTMYVRPIIEDY | 128 | 142 | Positive |
| hCoVOC43 | membrane | GHLYIQGIKLGTGYS | 152 | 166 | Positive |
| hCoVOC43 | membrane | GSGMDTALLRNNIGS | 218 | 230 | Positive |
| hCoVOC43 | membrane | GSGMSSKTTPAPVYI | 1 | 12 | Positive |
| hCoVOC43 | membrane | GSGSGMSSKTTPAPV | 1 | 10 | Positive |
| hCoVOC43 | membrane | IRTGSFWSFNPETNN | 109 | 123 | Positive |
| hCoVOC43 | membrane | ISDTSGFAVYVKSKV | 193 | 207 | Positive |
| hCoVOC43 | membrane | KEWNFSLGIILLFIT | 23 | 37 | Positive |
| hCoVOC43 | membrane | KLGTGYSLADLPAYM | 160 | 174 | Positive |
| hCoVOC43 | membrane | KVGNYRLPSTQKGSG | 206 | 220 | Positive |
| hCoVOC43 | membrane | NLMCIDMKGTMYVRP | 123 | 137 | Positive |
| hCoVOC43 | membrane | NNLMCIDMKGTMYVR | 122 | 136 | Positive |
| hCoVOC43 | membrane | NNVYLGLSIVFTIVA | 79 | 93 | Positive |
| hCoVOC43 | membrane | NPETNNLMCIDMKGT | 118 | 132 | Positive |
| hCoVOC43 | membrane | NYRLPSTQKGSGMDT | 209 | 223 | Positive |
| hCoVOC43 | membrane | PETNNLMCIDMKGTM | 119 | 133 | Positive |
| hCoVOC43 | membrane | PIIEDYHTLTVTIIR | 137 | 151 | Positive |
| hCoVOC43 | membrane | RPIIEDYHTLTVTII | 136 | 150 | Positive |
| hCoVOC43 | membrane | RTGSFWSFNPETNNL | 110 | 124 | Positive |
| hCoVOC43 | membrane | SDTSGFAVYVKSKVG | 194 | 208 | Positive |
| hCoVOC43 | membrane | SFWSFNPETNNLMCI | 113 | 127 | Positive |
| hCoVOC43 | membrane | SGSGSGMSSKTTPAP | 1 | 9 | Positive |
| hCoVOC43 | membrane | SMFVYVIKMIILWLM | 48 | 62 | Positive |
| hCoVOC43 | membrane | TMYVRPIIEDYHTLT | 132 | 146 | Positive |
| hCoVOC43 | membrane | TNNLMCIDMKGTMYV | 121 | 135 | Positive |
| hCoVOC43 | membrane | GLSIVFTIVAIIMWI | 84 | 98 | Negative |
| hCoVOC43 | membrane | GSGMDTALLRNNIGS | 218 | 230 | Negative |

| | | | | | |
|---|---|---|---|---|---|
| hCoVOC43 | membrane | GSGSGSGMSSKTTPA | 1 | 8 | Negative |
| hCoVOC43 | membrane | MSSKTTPAPVYIWTA | 1 | 15 | Negative |
| hCoVOC43 | membrane | EAIKFLKEWNFSLGI | 17 | 31 | Overlapping |
| hCoVOC43 | membrane | EDYHTLTVTIIRGHL | 140 | 154 | Overlapping |
| hCoVOC43 | membrane | ETNNLMCIDMKGTMY | 120 | 134 | Overlapping |
| hCoVOC43 | membrane | EWNFSLGIILLFITI | 24 | 38 | Overlapping |
| hCoVOC43 | membrane | GMDTALLRNNIGSGS | 220 | 230 | Overlapping |
| hCoVOC43 | membrane | IEDYHTLTVTIIRGH | 139 | 153 | Overlapping |
| hCoVOC43 | membrane | PIIEDYHTLTVTIIR | 137 | 151 | Overlapping |
| hCoVOC43 | membrane | SSKTTPAPVYIWTAD | 2 | 16 | Overlapping |
| hCoVOC43 | nucleocapsid | AEDISLLKKMDEPYT | 428 | 442 | Positive |
| hCoVOC43 | nucleocapsid | AELAPTAGAFFFGSR | 314 | 328 | Positive |
| hCoVOC43 | nucleocapsid | DGMMNMSPKPQRQRG | 384 | 398 | Positive |
| hCoVOC43 | nucleocapsid | DISLLKKMDEPYTED | 430 | 444 | Positive |
| hCoVOC43 | nucleocapsid | DPSSDEAIPTRFPPG | 165 | 179 | Positive |
| hCoVOC43 | nucleocapsid | FDSTLSGFETIMKVL | 359 | 373 | Positive |
| hCoVOC43 | nucleocapsid | FGGGEMLKLGTSDPQ | 295 | 309 | Positive |
| hCoVOC43 | nucleocapsid | GGEMLKLGTSDPQFP | 297 | 311 | Positive |
| hCoVOC43 | nucleocapsid | GHKNGQGENDNISVA | 398 | 412 | Positive |
| hCoVOC43 | nucleocapsid | GKRGPNQNFGGGEML | 287 | 301 | Positive |
| hCoVOC43 | nucleocapsid | GMSFTPGKQSSSRAS | 1 | 14 | Positive |
| hCoVOC43 | nucleocapsid | GQGENDNISVAVPKS | 402 | 416 | Positive |
| hCoVOC43 | nucleocapsid | GRSAPNSRSTSRTSS | 192 | 206 | Positive |
| hCoVOC43 | nucleocapsid | GSGMSFTPGKQSSSR | 1 | 12 | Positive |
| hCoVOC43 | nucleocapsid | GSGRSAPNSRSTSRT | 190 | 204 | Positive |
| hCoVOC43 | nucleocapsid | GSGSGMSFTPGKQSS | 1 | 10 | Positive |
| hCoVOC43 | nucleocapsid | GSGSGSGMSFTPGKQ | 1 | 8 | Positive |
| hCoVOC43 | nucleocapsid | GSRSRANSGNRTPTS | 212 | 226 | Positive |
| hCoVOC43 | nucleocapsid | IEGSGRSAPNSRSTS | 188 | 202 | Positive |
| hCoVOC43 | nucleocapsid | KDATKPQQVTKHTAK | 246 | 260 | Positive |
| hCoVOC43 | nucleocapsid | KGYWYRHNRRSFKTA | 98 | 112 | Positive |
| hCoVOC43 | nucleocapsid | KHTAKEVRQKILNKP | 256 | 270 | Positive |
| hCoVOC43 | nucleocapsid | KLGKDATKPQQVTKH | 243 | 257 | Positive |
| hCoVOC43 | nucleocapsid | KMDEPYTEDTSEIGS | 436 | 448 | Positive |
| hCoVOC43 | nucleocapsid | KPQQVTKHTAKEVRQ | 250 | 264 | Positive |
| hCoVOC43 | nucleocapsid | LELAKVQNLSGNPDE | 329 | 343 | Positive |
| hCoVOC43 | nucleocapsid | LGTGPHAKDQYGTDI | 128 | 142 | Positive |
| hCoVOC43 | nucleocapsid | LKKMDEPYTEDTSEI | 434 | 448 | Positive |
| hCoVOC43 | nucleocapsid | NAYQQQDGMMNMSPK | 378 | 392 | Positive |
| hCoVOC43 | nucleocapsid | NENLNAYQQQDGMMN | 374 | 388 | Positive |
| hCoVOC43 | nucleocapsid | NKPRQKRSPNKQCTV | 268 | 282 | Positive |
| hCoVOC43 | nucleocapsid | NKSRELTAEDISLLK | 421 | 435 | Positive |
| hCoVOC43 | nucleocapsid | NLNAYQQQDGMMNMS | 376 | 390 | Positive |
| hCoVOC43 | nucleocapsid | NMSPKPQRQRGHKNG | 388 | 402 | Positive |
| hCoVOC43 | nucleocapsid | NPDEPQKDVYELRYN | 340 | 354 | Positive |
| hCoVOC43 | nucleocapsid | NQADVNTPADIVDRD | 151 | 165 | Positive |
| hCoVOC43 | nucleocapsid | NRRSFKTADGNQRQL | 105 | 119 | Positive |
| hCoVOC43 | nucleocapsid | NRSGNGILKWADQSD | 17 | 31 | Positive |
| hCoVOC43 | nucleocapsid | NRTPTSGVTPDMADQ | 221 | 235 | Positive |

| | | | | | |
|---|---|---|---|---|---|
| hCoVOC43 | nucleocapsid | NSGNRTPTSGVTPDM | 218 | 232 | Positive |
| hCoVOC43 | nucleocapsid | NSRSTSRTSSRASSA | 197 | 211 | Positive |
| hCoVOC43 | nucleocapsid | PGVPATEAKGYWYRH | 90 | 104 | Positive |
| hCoVOC43 | nucleocapsid | PHAKDQYGTDIDGVY | 132 | 146 | Positive |
| hCoVOC43 | nucleocapsid | PQRQRGHKNGQGEND | 393 | 407 | Positive |
| hCoVOC43 | nucleocapsid | PSSDEAIPTRFPPGT | 166 | 180 | Positive |
| hCoVOC43 | nucleocapsid | PTSGVTPDMADQIAS | 224 | 238 | Positive |
| hCoVOC43 | nucleocapsid | PYTEDTSEIGSGSGS | 440 | 448 | Positive |
| hCoVOC43 | nucleocapsid | QFQKGKEFEFVEGQG | 71 | 85 | Positive |
| hCoVOC43 | nucleocapsid | QNKSRELTAEDISLL | 420 | 434 | Positive |
| hCoVOC43 | nucleocapsid | QQNKSRELTAEDISL | 419 | 433 | Positive |
| hCoVOC43 | nucleocapsid | RDPSSDEAIPTRFPP | 164 | 178 | Positive |
| hCoVOC43 | nucleocapsid | RELTAEDISLLKKMD | 424 | 438 | Positive |
| hCoVOC43 | nucleocapsid | RLELAKVQNLSGNPD | 328 | 342 | Positive |
| hCoVOC43 | nucleocapsid | RSAPNSRSTSRTSSR | 193 | 207 | Positive |
| hCoVOC43 | nucleocapsid | RTSSRASSAGSRSRA | 203 | 217 | Positive |
| hCoVOC43 | nucleocapsid | SDEAIPTRFPPGTVL | 168 | 182 | Positive |
| hCoVOC43 | nucleocapsid | SDPQFPILAELAPTA | 306 | 320 | Positive |
| hCoVOC43 | nucleocapsid | SGITQFQKGKEFEFV | 67 | 81 | Positive |
| hCoVOC43 | nucleocapsid | SGNPDEPQKDVYELR | 338 | 352 | Positive |
| hCoVOC43 | nucleocapsid | SSDEAIPTRFPPGTV | 167 | 181 | Positive |
| hCoVOC43 | nucleocapsid | TAEDISLLKKMDEPY | 427 | 441 | Positive |
| hCoVOC43 | nucleocapsid | TDIDGVYWVASNQAD | 140 | 154 | Positive |
| hCoVOC43 | nucleocapsid | TEAKGYWYRHNRRSF | 95 | 109 | Positive |
| hCoVOC43 | nucleocapsid | TGPHAKDQYGTDIDG | 130 | 144 | Positive |
| hCoVOC43 | nucleocapsid | TPADIVDRDPSSDEA | 157 | 171 | Positive |
| hCoVOC43 | nucleocapsid | TPDMADQIASLVLAK | 229 | 243 | Positive |
| hCoVOC43 | nucleocapsid | TQFQKGKEFEFVEGQ | 70 | 84 | Positive |
| hCoVOC43 | nucleocapsid | VEGQGVPIAPGVPAT | 81 | 95 | Positive |
| hCoVOC43 | nucleocapsid | VTPDMADQIASLVLA | 228 | 242 | Positive |
| hCoVOC43 | nucleocapsid | DGVYWVASNQADVNT | 143 | 157 | Negative |
| hCoVOC43 | nucleocapsid | DQSDQFRNVQTRGRR | 28 | 42 | Negative |
| hCoVOC43 | nucleocapsid | ELAKVQNLSGNPDEP | 330 | 344 | Negative |
| hCoVOC43 | nucleocapsid | ELAPTAGAFFFGSRL | 315 | 329 | Negative |
| hCoVOC43 | nucleocapsid | ELRYNGAIRFDSTLS | 350 | 364 | Negative |
| hCoVOC43 | nucleocapsid | ELTAEDISLLKKMDE | 425 | 439 | Negative |
| hCoVOC43 | nucleocapsid | FQKGKEFEFVEGQGV | 72 | 85 | Negative |
| hCoVOC43 | nucleocapsid | FTPGKQSSSRASSGN | 3 | 17 | Negative |
| hCoVOC43 | nucleocapsid | GGGEMLKLGTSDPQF | 296 | 310 | Negative |
| hCoVOC43 | nucleocapsid | GGNVVPYYSWFSGIT | 56 | 70 | Negative |
| hCoVOC43 | nucleocapsid | GHKNGQGENDNISVA | 398 | 412 | Negative |
| hCoVOC43 | nucleocapsid | GKDATKPQQVTKHTA | 245 | 259 | Negative |
| hCoVOC43 | nucleocapsid | GKEFEFVEGQGVPIA | 75 | 89 | Negative |
| hCoVOC43 | nucleocapsid | GSGSGSGMSFTPGKQ | 1 | 8 | Negative |
| hCoVOC43 | nucleocapsid | GSRLELAKVQNLSGN | 326 | 340 | Negative |
| hCoVOC43 | nucleocapsid | GTGPHAKDQYGTDID | 129 | 143 | Negative |
| hCoVOC43 | nucleocapsid | GTSDPQFPILAELAP | 304 | 318 | Negative |
| hCoVOC43 | nucleocapsid | GTVLPQGYYIEGSGR | 179 | 193 | Negative |
| hCoVOC43 | nucleocapsid | GVTPDMADQIASLVL | 227 | 241 | Negative |

| | | | | | |
|---|---|---|---|---|---|
| hCoVOC43 | nucleocapsid | GVYWVASNQADVNTP | 144 | 158 | Negative |
| hCoVOC43 | nucleocapsid | ITQFQKGKEFEFVEG | 69 | 83 | Negative |
| hCoVOC43 | nucleocapsid | KEVRQKILNKPRQKR | 260 | 274 | Negative |
| hCoVOC43 | nucleocapsid | MSFTPGKQSSSRASS | 1 | 15 | Negative |
| hCoVOC43 | nucleocapsid | PADIVDRDPSSDEAI | 158 | 172 | Negative |
| hCoVOC43 | nucleocapsid | PATEAKGYWYRHNRR | 93 | 106 | Negative |
| hCoVOC43 | nucleocapsid | QGVPIAPGVPATEAK | 84 | 98 | Negative |
| hCoVOC43 | nucleocapsid | RFDSTLSGFETIMKV | 358 | 372 | Negative |
| hCoVOC43 | nucleocapsid | SFTPGKQSSSRASSG | 2 | 16 | Negative |
| hCoVOC43 | nucleocapsid | SGVTPDMADQIASLV | 226 | 240 | Negative |
| hCoVOC43 | nucleocapsid | SRTSSRASSAGSRSR | 202 | 216 | Negative |
| hCoVOC43 | nucleocapsid | SSRASSGNRSGNGIL | 10 | 24 | Negative |
| hCoVOC43 | nucleocapsid | SSSRASSGNRSGNGI | 9 | 23 | Negative |
| hCoVOC43 | nucleocapsid | TPGKQSSSRASSGNR | 4 | 18 | Negative |
| hCoVOC43 | nucleocapsid | TSRTSSRASSAGSRS | 201 | 215 | Negative |
| hCoVOC43 | nucleocapsid | VDRDPSSDEAIPTRF | 162 | 176 | Negative |
| hCoVOC43 | nucleocapsid | WFSGITQFQKGKEFE | 65 | 79 | Negative |
| hCoVOC43 | nucleocapsid | YLGTGPHAKDQYGTD | 127 | 141 | Negative |
| hCoVOC43 | nucleocapsid | DEAIPTRFPPGTVLP | 169 | 183 | Overlapping |
| hCoVOC43 | nucleocapsid | DEPYTEDTSEIGSGS | 438 | 448 | Overlapping |
| hCoVOC43 | nucleocapsid | EAIPTRFPPGTVLPQ | 170 | 184 | Overlapping |
| hCoVOC43 | nucleocapsid | EAKGYWYRHNRRSFK | 96 | 110 | Overlapping |
| hCoVOC43 | nucleocapsid | EDISLLKKMDEPYTE | 429 | 443 | Overlapping |
| hCoVOC43 | nucleocapsid | EFEFVEGQGVPIAPG | 77 | 91 | Overlapping |
| hCoVOC43 | nucleocapsid | EFVEGQGVPIAPGVP | 79 | 93 | Overlapping |
| hCoVOC43 | nucleocapsid | EGQGVPIAPGVPATE | 82 | 96 | Overlapping |
| hCoVOC43 | nucleocapsid | EGSGRSAPNSRSTSR | 189 | 203 | Overlapping |
| hCoVOC43 | nucleocapsid | ELAKVQNLSGNPDEP | 330 | 344 | Overlapping |
| hCoVOC43 | nucleocapsid | ELRYNGAIRFDSTLS | 350 | 364 | Overlapping |
| hCoVOC43 | nucleocapsid | ELTAEDISLLKKMDE | 425 | 439 | Overlapping |
| hCoVOC43 | nucleocapsid | EMLKLGTSDPQFPIL | 299 | 313 | Overlapping |
| hCoVOC43 | nucleocapsid | ENDNISVAVPKSRVQ | 405 | 419 | Overlapping |
| hCoVOC43 | nucleocapsid | ENLNAYQQQDGMMNM | 375 | 389 | Overlapping |
| hCoVOC43 | nucleocapsid | EPQKDVYELRYNGAI | 343 | 357 | Overlapping |
| hCoVOC43 | nucleocapsid | EPYTEDTSEIGSGSG | 439 | 448 | Overlapping |
| hCoVOC43 | nucleocapsid | ETIMKVLNENLNAYQ | 367 | 381 | Overlapping |
| hCoVOC43 | nucleocapsid | EVRQKILNKPRQKRS | 261 | 275 | Overlapping |
| hCoVOC43 | nucleocapsid | FEFVEGQGVPIAPGV | 78 | 92 | Overlapping |
| hCoVOC43 | nucleocapsid | GENDNISVAVPKSRV | 404 | 418 | Overlapping |
| hCoVOC43 | nucleocapsid | GGGEMLKLGTSDPQF | 296 | 310 | Overlapping |
| hCoVOC43 | nucleocapsid | GITQFQKGKEFEFVE | 68 | 82 | Overlapping |
| hCoVOC43 | nucleocapsid | GMMNMSPKPQRQRGH | 385 | 399 | Overlapping |
| hCoVOC43 | nucleocapsid | GNPDEPQKDVYELRY | 339 | 353 | Overlapping |
| hCoVOC43 | nucleocapsid | GPHAKDQYGTDIDGV | 131 | 145 | Overlapping |
| hCoVOC43 | nucleocapsid | GVPATEAKGYWYRHN | 91 | 105 | Overlapping |
| hCoVOC43 | nucleocapsid | GVPIAPGVPATEAKG | 85 | 99 | Overlapping |
| hCoVOC43 | nucleocapsid | KEFEFVEGQGVPIAP | 76 | 90 | Overlapping |
| hCoVOC43 | nucleocapsid | KSRELTAEDISLLKK | 422 | 436 | Overlapping |
| hCoVOC43 | nucleocapsid | KSRVQQNKSRELTAE | 415 | 429 | Overlapping |

hCoVOC43

| | | | | | |
|---|---|---|---|---|---|
| hCoVOC43 | nucleocapsid | NLSGNPDEPQKDVYE | 336 | 350 | Overlapping |
| hCoVOC43 | nucleocapsid | PILAELAPTAGAFFF | 311 | 325 | Overlapping |
| hCoVOC43 | nucleocapsid | SSRASSAGSRSRANS | 205 | 219 | Overlapping |
| hCoVOC43 | orf1ab | ADDSIILRQYNLVDI | 344 | 358 | Positive |
| hCoVOC43 | orf1ab | AFTAPEDDDFLEESD | 960 | 974 | Positive |
| hCoVOC43 | orf1ab | AGDKYYPVVVDDHVG | 826 | 840 | Positive |
| hCoVOC43 | orf1ab | AGEEVLAPKLYCAFT | 948 | 962 | Positive |
| hCoVOC43 | orf1ab | AGLELTDESCNNLVP | 2648 | 2662 | Positive |
| hCoVOC43 | orf1ab | AGQPCVASEQEESSE | 992 | 1006 | Positive |
| hCoVOC43 | orf1ab | AIHYLNSKIVDLAQH | 630 | 644 | Positive |
| hCoVOC43 | orf1ab | ALDDDARTFVQSNVD | 1488 | 1502 | Positive |
| hCoVOC43 | orf1ab | ALPEMVTDIVVVDEV | 5658 | 5672 | Positive |
| hCoVOC43 | orf1ab | ALYLDYVQHKCGNLH | 514 | 528 | Positive |
| hCoVOC43 | orf1ab | ANGHMAHGGGVAKAI | 1300 | 1314 | Positive |
| hCoVOC43 | orf1ab | ANKFEQFDQWYESTF | 2812 | 2826 | Positive |
| hCoVOC43 | orf1ab | ANMLPAHVFMRFYII | 2360 | 2374 | Positive |
| hCoVOC43 | orf1ab | ANTGTSVTETMFDVY | 2566 | 2580 | Positive |
| hCoVOC43 | orf1ab | APEFPWMFEDAEEKL | 14 | 28 | Positive |
| hCoVOC43 | orf1ab | APKLYCAFTAPEDDD | 954 | 968 | Positive |
| hCoVOC43 | orf1ab | ASEQEESSEVLEDTL | 998 | 1012 | Positive |
| hCoVOC43 | orf1ab | ATIRSVAYVANPTED | 314 | 328 | Positive |
| hCoVOC43 | orf1ab | AVPVNMRVLHLGAGS | 6856 | 6870 | Positive |
| hCoVOC43 | orf1ab | AVYDCFCNNINWNVE | 6180 | 6194 | Positive |
| hCoVOC43 | orf1ab | AWVGFDAEGAHATRD | 5984 | 5998 | Positive |
| hCoVOC43 | orf1ab | CKELEGVGAKVSAFL | 920 | 934 | Positive |
| hCoVOC43 | orf1ab | CLKHAEEYREYLESY | 6378 | 6392 | Positive |
| hCoVOC43 | orf1ab | CLQHLHVASNSKLWH | 3852 | 3866 | Positive |
| hCoVOC43 | orf1ab | CNIAKSAYERDRAVA | 3968 | 3982 | Positive |
| hCoVOC43 | orf1ab | CRVGLCEEADEGICF | 2970 | 2984 | Positive |
| hCoVOC43 | orf1ab | CSTLHNEILATSDLS | 3868 | 3882 | Positive |
| hCoVOC43 | orf1ab | CVASEQEESSEVLED | 996 | 1010 | Positive |
| hCoVOC43 | orf1ab | DALSISTIFTQSRVI | 6636 | 6650 | Positive |
| hCoVOC43 | orf1ab | DAMFFYGDVVSHICK | 1164 | 1178 | Positive |
| hCoVOC43 | orf1ab | DAVMHFGTLSREDLE | 1762 | 1776 | Positive |
| hCoVOC43 | orf1ab | DAYDQVHDEPKGKFS | 220 | 234 | Positive |
| hCoVOC43 | orf1ab | DCDSYKPGNTFITVE | 2442 | 2456 | Positive |
| hCoVOC43 | orf1ab | DDHRITSITSDKFDF | 1236 | 1250 | Positive |
| hCoVOC43 | orf1ab | DDHVGLLDQAWRVPC | 836 | 850 | Positive |
| hCoVOC43 | orf1ab | DFIALDQDVFIQKYG | 6662 | 6676 | Positive |
| hCoVOC43 | orf1ab | DFLEESDVEEDDVEG | 968 | 982 | Positive |
| hCoVOC43 | orf1ab | DFQLPVYASYKVLDN | 2784 | 2798 | Positive |
| hCoVOC43 | orf1ab | DGICPENHVMVDCRR | 50 | 64 | Positive |
| hCoVOC43 | orf1ab | DGLEAYADKTLQEMK | 266 | 280 | Positive |
| hCoVOC43 | orf1ab | DGLSCFKIGRRRICL | 724 | 738 | Positive |
| hCoVOC43 | orf1ab | DGMDAKQVDYVPLKS | 6350 | 6364 | Positive |
| hCoVOC43 | orf1ab | DGNVMHANYLFWRNS | 7016 | 7030 | Positive |
| hCoVOC43 | orf1ab | DGPSVETSDSQVEED | 1014 | 1028 | Positive |
| hCoVOC43 | orf1ab | DGTFEEMALTTFMIT | 3156 | 3170 | Positive |
| hCoVOC43 | orf1ab | DGYCKLFITKEEAVK | 5966 | 5980 | Positive |

| | | | | | |
|---|---|---|---|---|---|
| hCoVOC43 | orf1ab | DHGRGGFGHVYDFKV | 204 | 218 | Positive |
| hCoVOC43 | orf1ab | DIQQWGYIGSLSSNH | 6142 | 6156 | Positive |
| hCoVOC43 | orf1ab | DKVVAKIDKEDVVIF | 6450 | 6464 | Positive |
| hCoVOC43 | orf1ab | DLVQYDFTDYKLELF | 4634 | 4648 | Positive |
| hCoVOC43 | orf1ab | DMEKDFIALDQDVFI | 6658 | 6672 | Positive |
| hCoVOC43 | orf1ab | DMLDNYKAIDVVQYE | 2280 | 2294 | Positive |
| hCoVOC43 | orf1ab | DSLNAKLGFDSSKEF | 1932 | 1946 | Positive |
| hCoVOC43 | orf1ab | DSQVEEDVEMSDFVD | 1022 | 1036 | Positive |
| hCoVOC43 | orf1ab | DSSIHSYFITDEKSG | 6722 | 6736 | Positive |
| hCoVOC43 | orf1ab | DSVIVNDDTSETKYV | 2052 | 2066 | Positive |
| hCoVOC43 | orf1ab | DSVMSAVSAGLELTD | 2640 | 2654 | Positive |
| hCoVOC43 | orf1ab | DSVVEVVTTSLTPCG | 790 | 804 | Positive |
| hCoVOC43 | orf1ab | DWYDFVENPDIINVY | 4530 | 4544 | Positive |
| hCoVOC43 | orf1ab | DYVQHKCGNLHQREL | 518 | 532 | Positive |
| hCoVOC43 | orf1ab | EADRRAFVDYTGVLK | 2294 | 2308 | Positive |
| hCoVOC43 | orf1ab | EDAEEKLDNPSSSEV | 22 | 36 | Positive |
| hCoVOC43 | orf1ab | EDHKPQYSFKLVMNG | 5370 | 5384 | Positive |
| hCoVOC43 | orf1ab | EMSDFVDLESVIQDY | 1030 | 1044 | Positive |
| hCoVOC43 | orf1ab | EQDEIYAYTKRNVLP | 4888 | 4902 | Positive |
| hCoVOC43 | orf1ab | EQEESSEVLEDTLDD | 1000 | 1014 | Positive |
| hCoVOC43 | orf1ab | ESDVEEDDVEGEETD | 972 | 986 | Positive |
| hCoVOC43 | orf1ab | EYDYVIYSQTAETAH | 5836 | 5850 | Positive |
| hCoVOC43 | orf1ab | FAAETQKATEEAFKQ | 5430 | 5444 | Positive |
| hCoVOC43 | orf1ab | FDAEGAHATRDSIGT | 5988 | 6002 | Positive |
| hCoVOC43 | orf1ab | FDCQWDLIISDMYDP | 6916 | 6930 | Positive |
| hCoVOC43 | orf1ab | FEVDDTVDMEEFYAV | 894 | 908 | Positive |
| hCoVOC43 | orf1ab | FGHVYDFKVEDAYDQ | 210 | 224 | Positive |
| hCoVOC43 | orf1ab | FIDGLSCFKIGRRRI | 722 | 736 | Positive |
| hCoVOC43 | orf1ab | FINNTTYPTNVAVEL | 6464 | 6478 | Positive |
| hCoVOC43 | orf1ab | FKINDTTRKYVFTTI | 5642 | 5656 | Positive |
| hCoVOC43 | orf1ab | FKVEDAYDQVHDEPK | 216 | 230 | Positive |
| hCoVOC43 | orf1ab | FMPVQSRKFIVPWVM | 176 | 190 | Positive |
| hCoVOC43 | orf1ab | FMSESKCWVEHDINN | 5158 | 5172 | Positive |
| hCoVOC43 | orf1ab | FNRPSLVDDNKFDVL | 2000 | 2014 | Positive |
| hCoVOC43 | orf1ab | FSLWYKGSNLEEEIL | 3698 | 3712 | Positive |
| hCoVOC43 | orf1ab | FVDDLLKTDSVLLIE | 5208 | 5222 | Positive |
| hCoVOC43 | orf1ab | FVENPDIINVYKKLG | 4534 | 4548 | Positive |
| hCoVOC43 | orf1ab | FVSLAIDAYPLVYHE | 5224 | 5238 | Positive |
| hCoVOC43 | orf1ab | GCITFGKPVIWLSHE | 1976 | 1990 | Positive |
| hCoVOC43 | orf1ab | GCNPKGWTMGLFRRR | 120 | 134 | Positive |
| hCoVOC43 | orf1ab | GDPADSRDFLRVVFS | 1720 | 1734 | Positive |
| hCoVOC43 | orf1ab | GDPAQLPAPRVLLSK | 5696 | 5710 | Positive |
| hCoVOC43 | orf1ab | GDSLVNVIGSGSGSG | 7088 | 7095 | Positive |
| hCoVOC43 | orf1ab | GGAVCLKHAEEYREY | 6374 | 6388 | Positive |
| hCoVOC43 | orf1ab | GHMAHGGGVAKAIAV | 1302 | 1316 | Positive |
| hCoVOC43 | orf1ab | GHYTGQAGEMPCAII | 6434 | 6448 | Positive |
| hCoVOC43 | orf1ab | GNPKAIACVKDFDFK | 6232 | 6246 | Positive |
| hCoVOC43 | orf1ab | GPIFNRALVSATEFA | 4548 | 4562 | Positive |
| hCoVOC43 | orf1ab | GPPGTGKSHLAIGLA | 5578 | 5592 | Positive |

| | | | | | |
|---|---|---|---|---|---|
| hCoVOC43 | orf1ab | GRKIYEVERGLLHSS | 740 | 754 | Positive |
| hCoVOC43 | orf1ab | GRPARFVALVLAKGG | 1702 | 1716 | Positive |
| hCoVOC43 | orf1ab | GRRRICLSGRKIYEV | 732 | 746 | Positive |
| hCoVOC43 | orf1ab | GSCITPNVCFVKGDI | 1270 | 1284 | Positive |
| hCoVOC43 | orf1ab | GSEYDYVIYSQTAET | 5834 | 5848 | Positive |
| hCoVOC43 | orf1ab | GSGMSKINKYGLELH | 1 | 12 | Positive |
| hCoVOC43 | orf1ab | GSGSGSGMSKINKYG | 1 | 8 | Positive |
| hCoVOC43 | orf1ab | GSNLEEEILLMLASL | 3704 | 3718 | Positive |
| hCoVOC43 | orf1ab | GSPQPYCYTEGLMQN | 2912 | 2926 | Positive |
| hCoVOC43 | orf1ab | GYTVDCSCGKKLIHC | 1778 | 1792 | Positive |
| hCoVOC43 | orf1ab | HAFHTKPFARAAFEH | 6318 | 6332 | Positive |
| hCoVOC43 | orf1ab | HDFFTFDVEGSRVPH | 4468 | 4482 | Positive |
| hCoVOC43 | orf1ab | HDINNGPHEFCSQHT | 5168 | 5182 | Positive |
| hCoVOC43 | orf1ab | HHPELKLFRNLNIDV | 6486 | 6500 | Positive |
| hCoVOC43 | orf1ab | HLTFKIVQWQEAWLE | 1684 | 1698 | Positive |
| hCoVOC43 | orf1ab | HPNTVDCQDDRCIIH | 4660 | 4674 | Positive |
| hCoVOC43 | orf1ab | HRITSITSDKFDFII | 1238 | 1252 | Positive |
| hCoVOC43 | orf1ab | HVYDFKVEDAYDQVH | 212 | 226 | Positive |
| hCoVOC43 | orf1ab | HVYEVGDLMAQSSGV | 418 | 432 | Positive |
| hCoVOC43 | orf1ab | HYCSTLHNEILATSD | 3866 | 3880 | Positive |
| hCoVOC43 | orf1ab | HYLNSKIVDLAQHFS | 632 | 646 | Positive |
| hCoVOC43 | orf1ab | IEEVCDDYAKDNTVL | 3910 | 3924 | Positive |
| hCoVOC43 | orf1ab | IGMKRYCTVQGPPGT | 5568 | 5582 | Positive |
| hCoVOC43 | orf1ab | IKEPVHVYADDSIIL | 336 | 350 | Positive |
| hCoVOC43 | orf1ab | IMGSCSFEDELTPSD | 3524 | 3538 | Positive |
| hCoVOC43 | orf1ab | IMPMLTMCHALDCEL | 4610 | 4624 | Positive |
| hCoVOC43 | orf1ab | INDKVVAKIDKEDVV | 6448 | 6462 | Positive |
| hCoVOC43 | orf1ab | INNGPHEFCSQHTML | 5170 | 5184 | Positive |
| hCoVOC43 | orf1ab | IPIDLLNLREIKPAV | 2114 | 2128 | Positive |
| hCoVOC43 | orf1ab | IPSLAANTLNIIVPD | 4046 | 4060 | Positive |
| hCoVOC43 | orf1ab | IQTIQDSDGTNKQLN | 4082 | 4096 | Positive |
| hCoVOC43 | orf1ab | ISTTKVKSLSMIRGP | 6558 | 6572 | Positive |
| hCoVOC43 | orf1ab | ISVQELRYMNANGLR | 3792 | 3806 | Positive |
| hCoVOC43 | orf1ab | ITAVEGTKKLAARLS | 1432 | 1446 | Positive |
| hCoVOC43 | orf1ab | IVMNMDVDTHRYRLS | 4718 | 4732 | Positive |
| hCoVOC43 | orf1ab | IVVESDIYFVKNIPR | 684 | 698 | Positive |
| hCoVOC43 | orf1ab | IVVVDEVSMLTNYEL | 5666 | 5680 | Positive |
| hCoVOC43 | orf1ab | IYNREMKCYERVKDC | 4448 | 4462 | Positive |
| hCoVOC43 | orf1ab | KAEVVVNPANGHMAH | 1292 | 1306 | Positive |
| hCoVOC43 | orf1ab | KAMDTFTNNNGSDVL | 3218 | 3232 | Positive |
| hCoVOC43 | orf1ab | KAPPGEQFKHLIPLM | 6038 | 6052 | Positive |
| hCoVOC43 | orf1ab | KARLYYEALSFEEQD | 4876 | 4890 | Positive |
| hCoVOC43 | orf1ab | KAVAKAPPGEQFKHL | 6034 | 6048 | Positive |
| hCoVOC43 | orf1ab | KDCKFVAEHDFFTFD | 4460 | 4474 | Positive |
| hCoVOC43 | orf1ab | KDPVSYVLTHDVCRV | 4336 | 4350 | Positive |
| hCoVOC43 | orf1ab | KEARINDKKSKVVSA | 3998 | 4012 | Positive |
| hCoVOC43 | orf1ab | KEDVVIFINNTTYPT | 6458 | 6472 | Positive |
| hCoVOC43 | orf1ab | KEINIIKLSGVKKPF | 2034 | 2048 | Positive |
| hCoVOC43 | orf1ab | KEIVDTVSALVYENK | 5742 | 5756 | Positive |

| | | | | | |
|---|---|---|---|---|---|
| hCoVOC43 | orf1ab | KEQPTVKEIISMPKI | 858 | 872 | Positive |
| hCoVOC43 | orf1ab | KETTDMVKSKGVCAT | 1324 | 1338 | Positive |
| hCoVOC43 | orf1ab | KFGMTLVSIPIDLLN | 2106 | 2120 | Positive |
| hCoVOC43 | orf1ab | KGAYNKDHGRGGFGH | 198 | 212 | Positive |
| hCoVOC43 | orf1ab | KGSGSDFSLADSVVE | 780 | 794 | Positive |
| hCoVOC43 | orf1ab | KHAEEYREYLESYNT | 6380 | 6394 | Positive |
| hCoVOC43 | orf1ab | KIGFLPTFVGKIAQW | 2226 | 2240 | Positive |
| hCoVOC43 | orf1ab | KIGRRRICLSGRKIY | 730 | 744 | Positive |
| hCoVOC43 | orf1ab | KKSLNALIATAHSSI | 2592 | 2606 | Positive |
| hCoVOC43 | orf1ab | KLERMADLALTNMYK | 3984 | 3998 | Positive |
| hCoVOC43 | orf1ab | KLETDGICPENHVMV | 46 | 60 | Positive |
| hCoVOC43 | orf1ab | KLFSLFRHVAYGCSK | 2380 | 2394 | Positive |
| hCoVOC43 | orf1ab | KLKIQVVNSGPDQTC | 4130 | 4144 | Positive |
| hCoVOC43 | orf1ab | KLYCAFTAPEDDDFL | 956 | 970 | Positive |
| hCoVOC43 | orf1ab | KPPLNKNYVFTGYHF | 5470 | 5484 | Positive |
| hCoVOC43 | orf1ab | KQVVLVSNNQEDFDL | 1412 | 1426 | Positive |
| hCoVOC43 | orf1ab | KRPIQPTDVAYHTVT | 2466 | 2480 | Positive |
| hCoVOC43 | orf1ab | KSLSMIRGPPRAELN | 6564 | 6578 | Positive |
| hCoVOC43 | orf1ab | KTGHYTGQAGEMPCA | 6432 | 6446 | Positive |
| hCoVOC43 | orf1ab | KTLLYSFEAHKDSFK | 6258 | 6272 | Positive |
| hCoVOC43 | orf1ab | KVKSLSMIRGPPRAE | 6562 | 6576 | Positive |
| hCoVOC43 | orf1ab | KVLRYGYHVLHFITH | 2858 | 2872 | Positive |
| hCoVOC43 | orf1ab | KVPQAVETKVQCSTN | 5890 | 5904 | Positive |
| hCoVOC43 | orf1ab | KWQVVVNGKYFTFKQ | 1652 | 1666 | Positive |
| hCoVOC43 | orf1ab | KYTMLDLCYALRHFD | 4490 | 4504 | Positive |
| hCoVOC43 | orf1ab | LCCKCCYDHVMATDH | 5322 | 5336 | Positive |
| hCoVOC43 | orf1ab | LCFKVYYKGVTTHES | 5766 | 5780 | Positive |
| hCoVOC43 | orf1ab | LDKVDILLTVDGVNF | 1562 | 1576 | Positive |
| hCoVOC43 | orf1ab | LINDVAFVSTFNVLQ | 1464 | 1478 | Positive |
| hCoVOC43 | orf1ab | LISKCQITAVEGTKK | 1426 | 1440 | Positive |
| hCoVOC43 | orf1ab | LLEKGKLLIRDTNKE | 7070 | 7084 | Positive |
| hCoVOC43 | orf1ab | LLVGMVFVTLRSINH | 3668 | 3682 | Positive |
| hCoVOC43 | orf1ab | LPELFMLSTLHWSFR | 2340 | 2354 | Positive |
| hCoVOC43 | orf1ab | LREGLVRIVRTRSMS | 2954 | 2968 | Positive |
| hCoVOC43 | orf1ab | LVLAKGGFKFGDPAD | 1710 | 1724 | Positive |
| hCoVOC43 | orf1ab | LWHYCSTLHNEILAT | 3864 | 3878 | Positive |
| hCoVOC43 | orf1ab | LWYKGSNLEEEILLM | 3700 | 3714 | Positive |
| hCoVOC43 | orf1ab | MVFVTLRSINHDLFS | 3672 | 3686 | Positive |
| hCoVOC43 | orf1ab | NAAITDYNYYKYNLP | 4812 | 4826 | Positive |
| hCoVOC43 | orf1ab | NAAVFYAQSLFRPIL | 2542 | 2556 | Positive |
| hCoVOC43 | orf1ab | NAIPSLAANTLNIIV | 4044 | 4058 | Positive |
| hCoVOC43 | orf1ab | NALIATAHSSIKQGT | 2596 | 2610 | Positive |
| hCoVOC43 | orf1ab | NALSRAVNVPTIRKF | 2090 | 2104 | Positive |
| hCoVOC43 | orf1ab | NAPGCDVNDVTKLYL | 5348 | 5362 | Positive |
| hCoVOC43 | orf1ab | NARIRAKHYVYIGDP | 5684 | 5698 | Positive |
| hCoVOC43 | orf1ab | NASLYSSLVPHVRYN | 2926 | 2940 | Positive |
| hCoVOC43 | orf1ab | NASSSEGFLIGINYL | 6994 | 7008 | Positive |
| hCoVOC43 | orf1ab | NAYRLFDLVQYDFTD | 4628 | 4642 | Positive |
| hCoVOC43 | orf1ab | NCVKMLCDHAGTGMA | 4272 | 4286 | Positive |

| | | | | | |
|---|---|---|---|---|---|
| hCoVOC43 | orf1ab | NDCMLLCDILSIYAG | 4506 | 4520 | Positive |
| hCoVOC43 | orf1ab | NDEKVMTFYPRLQAA | 6784 | 6798 | Positive |
| hCoVOC43 | orf1ab | NDLGNQILDSYSVIL | 5258 | 5272 | Positive |
| hCoVOC43 | orf1ab | NDLYPFVSDSVATYF | 6894 | 6908 | Positive |
| hCoVOC43 | orf1ab | NDSHSMAVVDGKQID | 1222 | 1236 | Positive |
| hCoVOC43 | orf1ab | NDVAFVSTFNVLQDV | 1466 | 1480 | Positive |
| hCoVOC43 | orf1ab | NEISDDCNWPLVIIA | 4096 | 4110 | Positive |
| hCoVOC43 | orf1ab | NELMPAKLKIQVVNS | 4124 | 4138 | Positive |
| hCoVOC43 | orf1ab | NELSINTSCRVLQRV | 6200 | 6214 | Positive |
| hCoVOC43 | orf1ab | NEVSATVLQNNELMP | 4114 | 4128 | Positive |
| hCoVOC43 | orf1ab | NFDQKELLAYYNMLV | 1634 | 1648 | Positive |
| hCoVOC43 | orf1ab | NFKLIGHTVCDSLNA | 1922 | 1936 | Positive |
| hCoVOC43 | orf1ab | NFKLLGIGGVPIIEV | 3818 | 3832 | Positive |
| hCoVOC43 | orf1ab | NFLNRVRGTSVDARL | 4374 | 4388 | Positive |
| hCoVOC43 | orf1ab | NFMMLFVFQVYPILS | 3074 | 3088 | Positive |
| hCoVOC43 | orf1ab | NGPHEFCSQHTMLVK | 5172 | 5186 | Positive |
| hCoVOC43 | orf1ab | NGVIRDVSVEDVCFA | 2798 | 2812 | Positive |
| hCoVOC43 | orf1ab | NHVMVDCRRLLKQEC | 56 | 70 | Positive |
| hCoVOC43 | orf1ab | NIDYFNMRRAKFSLE | 556 | 570 | Positive |
| hCoVOC43 | orf1ab | NINWNVEYPIISNEL | 6188 | 6202 | Positive |
| hCoVOC43 | orf1ab | NKATVAQIKALFLDK | 1550 | 1564 | Positive |
| hCoVOC43 | orf1ab | NKDHGRGGFGHVYDF | 202 | 216 | Positive |
| hCoVOC43 | orf1ab | NKEVFVGDSLVNVIG | 7082 | 7095 | Positive |
| hCoVOC43 | orf1ab | NKFYQINGVRTVKYF | 1512 | 1526 | Positive |
| hCoVOC43 | orf1ab | NKHAFHTKPFARAAF | 6316 | 6330 | Positive |
| hCoVOC43 | orf1ab | NKIEDLSIRALQKRL | 5078 | 5092 | Positive |
| hCoVOC43 | orf1ab | NKIPANIVLPQGGYV | 1116 | 1130 | Positive |
| hCoVOC43 | orf1ab | NKLILINDVAFVSTF | 1460 | 1474 | Positive |
| hCoVOC43 | orf1ab | NKNYVFTGYHFTKNG | 5474 | 5488 | Positive |
| hCoVOC43 | orf1ab | NKQLNEISDDCNWPL | 4092 | 4106 | Positive |
| hCoVOC43 | orf1ab | NKQMANVSVLTTPFS | 2732 | 2746 | Positive |
| hCoVOC43 | orf1ab | NKVIYTTEIASKLTC | 2162 | 2176 | Positive |
| hCoVOC43 | orf1ab | NKYRYYSGKMDTAAY | 3194 | 3208 | Positive |
| hCoVOC43 | orf1ab | NLANAKGFIRFPEVL | 2940 | 2954 | Positive |
| hCoVOC43 | orf1ab | NLDEARFSGSANQQQ | 3946 | 3960 | Positive |
| hCoVOC43 | orf1ab | NLFKDCSKSYSGYHP | 5904 | 5918 | Positive |
| hCoVOC43 | orf1ab | NLGGAVCLKHAEEYR | 6372 | 6386 | Positive |
| hCoVOC43 | orf1ab | NLGSNGKPGNVGGND | 6622 | 6636 | Positive |
| hCoVOC43 | orf1ab | NLHQRELLGVSDVWH | 526 | 540 | Positive |
| hCoVOC43 | orf1ab | NLKQTFKSVLTTYYL | 1864 | 1878 | Positive |
| hCoVOC43 | orf1ab | NLLCRVTSSDFTVLF | 3302 | 3316 | Positive |
| hCoVOC43 | orf1ab | NLLHSKVKSVPNMHV | 2514 | 2528 | Positive |
| hCoVOC43 | orf1ab | NLNLPGCNGGSLYVN | 6302 | 6316 | Positive |
| hCoVOC43 | orf1ab | NLPGCNGGSLYVNKH | 6304 | 6318 | Positive |
| hCoVOC43 | orf1ab | NLPTMVDIKQLLFVL | 4824 | 4838 | Positive |
| hCoVOC43 | orf1ab | NLQDLWVLYKQQYSQ | 1094 | 1108 | Positive |
| hCoVOC43 | orf1ab | NLREIKPAVNVVKAV | 2120 | 2134 | Positive |
| hCoVOC43 | orf1ab | NLSSEDLKAVRSSFN | 1620 | 1634 | Positive |
| hCoVOC43 | orf1ab | NLVDIMSHFYMEADT | 354 | 368 | Positive |

| | | | | | |
|---|---|---|---|---|---|
| hCoVOC43 | orf1ab | NLVVQEFVSYDSSIH | 6712 | 6726 | Positive |
| hCoVOC43 | orf1ab | NLWNTFTKLQSLENV | 6412 | 6426 | Positive |
| hCoVOC43 | orf1ab | NMIDGFACTTCGHVY | 406 | 420 | Positive |
| hCoVOC43 | orf1ab | NMQLFEALQFTTLTL | 5874 | 5888 | Positive |
| hCoVOC43 | orf1ab | NMRVLHLGAGSEKGV | 6860 | 6874 | Positive |
| hCoVOC43 | orf1ab | NMTLNGLWLDDKVYC | 3270 | 3284 | Positive |
| hCoVOC43 | orf1ab | NMYLRSAVMQSVGAC | 5288 | 5302 | Positive |
| hCoVOC43 | orf1ab | NNCNWFVQSDKCSVE | 3460 | 3474 | Positive |
| hCoVOC43 | orf1ab | NNLVPTYLKSDNIVA | 2658 | 2672 | Positive |
| hCoVOC43 | orf1ab | NNNCFVNVSCLMLQS | 1668 | 1682 | Positive |
| hCoVOC43 | orf1ab | NNTTYPTNVAVELFA | 6466 | 6480 | Positive |
| hCoVOC43 | orf1ab | NNVVNYQHIGMKRYC | 5560 | 5574 | Positive |
| hCoVOC43 | orf1ab | NPAAVDSKCLTSIEE | 3898 | 3912 | Positive |
| hCoVOC43 | orf1ab | NPANGHMAHGGGVAK | 1298 | 1312 | Positive |
| hCoVOC43 | orf1ab | NPLFLFDEAGEEVLA | 940 | 954 | Positive |
| hCoVOC43 | orf1ab | NPSSSEVDMICSTTA | 30 | 44 | Positive |
| hCoVOC43 | orf1ab | NPTEDLCDGSVVIKE | 324 | 338 | Positive |
| hCoVOC43 | orf1ab | NPVLHTKSAAGYGGF | 436 | 450 | Positive |
| hCoVOC43 | orf1ab | NQALNSILDNAVKGC | 4026 | 4040 | Positive |
| hCoVOC43 | orf1ab | NQDLNGKWYDFGDYV | 4578 | 4592 | Positive |
| hCoVOC43 | orf1ab | NQFSSDFQHKLKKAC | 2706 | 2720 | Positive |
| hCoVOC43 | orf1ab | NQILDSYSVILSTCD | 5262 | 5276 | Positive |
| hCoVOC43 | orf1ab | NQSPQGNLGSNGKPG | 6616 | 6630 | Positive |
| hCoVOC43 | orf1ab | NRALVSATEFADKLV | 4552 | 4566 | Positive |
| hCoVOC43 | orf1ab | NREMKCYERVKDCKF | 4450 | 4464 | Positive |
| hCoVOC43 | orf1ab | NRFVPVGESFGKSLG | 1578 | 1592 | Positive |
| hCoVOC43 | orf1ab | NRGVYKPLLENIDYF | 546 | 560 | Positive |
| hCoVOC43 | orf1ab | NRIASCKWTDVDDYI | 5404 | 5418 | Positive |
| hCoVOC43 | orf1ab | NRSLRVKCSTIVGGM | 2404 | 2418 | Positive |
| hCoVOC43 | orf1ab | NSDYASKGYIANISA | 5132 | 5146 | Positive |
| hCoVOC43 | orf1ab | NSILDNAVKGCVPLN | 4030 | 4044 | Positive |
| hCoVOC43 | orf1ab | NTGRCTVNKHVAYQL | 138 | 152 | Positive |
| hCoVOC43 | orf1ab | NTVDCQDDRCIIHCA | 4662 | 4676 | Positive |
| hCoVOC43 | orf1ab | NVCALMSCNGNKIED | 5068 | 5082 | Positive |
| hCoVOC43 | orf1ab | NVDVVPEGWRVVNKF | 1500 | 1514 | Positive |
| hCoVOC43 | orf1ab | NVEYPIISNELSINT | 6192 | 6206 | Positive |
| hCoVOC43 | orf1ab | NVFMSESKCWVEHDI | 5156 | 5170 | Positive |
| hCoVOC43 | orf1ab | NVGRSIVYETDANKL | 1448 | 1462 | Positive |
| hCoVOC43 | orf1ab | NVMHANYLFWRNSTV | 7018 | 7032 | Positive |
| hCoVOC43 | orf1ab | NVTKHKCDINYKGKV | 1600 | 1614 | Positive |
| hCoVOC43 | orf1ab | NWNVEYPIISNELSI | 6190 | 6204 | Positive |
| hCoVOC43 | orf1ab | NYKAIDVVQYEADRR | 2284 | 2298 | Positive |
| hCoVOC43 | orf1ab | NYLCKPKVEIDGNVM | 7006 | 7020 | Positive |
| hCoVOC43 | orf1ab | NYQHIGMKRYCTVQG | 5564 | 5578 | Positive |
| hCoVOC43 | orf1ab | PAPRVLLSKGTLEPK | 5702 | 5716 | Positive |
| hCoVOC43 | orf1ab | PEMVTDIVVVDEVSM | 5660 | 5674 | Positive |
| hCoVOC43 | orf1ab | PGETFTVLAAYNGKP | 3354 | 3368 | Positive |
| hCoVOC43 | orf1ab | PHEFCSQHTMLVKMD | 5174 | 5188 | Positive |
| hCoVOC43 | orf1ab | PHSQISYSNFYASGC | 2884 | 2898 | Positive |

| | | | | | |
|---|---|---|---|---|---|
| hCoVOC43 | orf1ab | PLGMSLEACYVRGCN | 108 | 122 | Positive |
| hCoVOC43 | orf1ab | PLLENIDYFNMRRAK | 552 | 566 | Positive |
| hCoVOC43 | orf1ab | PMLTMCHALDCELYV | 4612 | 4626 | Positive |
| hCoVOC43 | orf1ab | PPGEQFKHLIPLMTR | 6040 | 6054 | Positive |
| hCoVOC43 | orf1ab | PQAVETKVQCSTNLF | 5892 | 5906 | Positive |
| hCoVOC43 | orf1ab | PSDVSLTYLLGTAKK | 1398 | 1412 | Positive |
| hCoVOC43 | orf1ab | PSDVYQQLAGIKLQS | 3536 | 3550 | Positive |
| hCoVOC43 | orf1ab | PSVETSDSQVEEDVE | 1016 | 1030 | Positive |
| hCoVOC43 | orf1ab | PTWSQELLFDVIVAW | 284 | 298 | Positive |
| hCoVOC43 | orf1ab | PVGESFGKSLGNVFC | 1582 | 1596 | Positive |
| hCoVOC43 | orf1ab | PVGIKDPVSYVLTHD | 4332 | 4346 | Positive |
| hCoVOC43 | orf1ab | PVNMRVLHLGAGSEK | 6858 | 6872 | Positive |
| hCoVOC43 | orf1ab | PVVVDDHVGLLDQAW | 832 | 846 | Positive |
| hCoVOC43 | orf1ab | PYHPNTVDCQDDRCI | 4658 | 4672 | Positive |
| hCoVOC43 | orf1ab | QDVLSLRHDIALDDD | 1478 | 1492 | Positive |
| hCoVOC43 | orf1ab | QGTQIYKVLDTFLSC | 2608 | 2622 | Positive |
| hCoVOC43 | orf1ab | QHIGMKRYCTVQGPP | 5566 | 5580 | Positive |
| hCoVOC43 | orf1ab | QNNVFMSESKCWVEH | 5154 | 5168 | Positive |
| hCoVOC43 | orf1ab | QPCVASEQEESSEVL | 994 | 1008 | Positive |
| hCoVOC43 | orf1ab | QQVLYYQNNVFMSES | 5148 | 5162 | Positive |
| hCoVOC43 | orf1ab | QQWGYIGSLSSNHDL | 6144 | 6158 | Positive |
| hCoVOC43 | orf1ab | QVPVGIKDPVSYVLT | 4330 | 4344 | Positive |
| hCoVOC43 | orf1ab | QYLNTTTLAVPVNMR | 6848 | 6862 | Positive |
| hCoVOC43 | orf1ab | RARTVAGVSILSTMT | 4918 | 4932 | Positive |
| hCoVOC43 | orf1ab | RCNLGGAVCLKHAEE | 6370 | 6384 | Positive |
| hCoVOC43 | orf1ab | RDFLRVVFSQVDLTG | 1726 | 1740 | Positive |
| hCoVOC43 | orf1ab | RDKLALGGSVAIKIT | 6954 | 6968 | Positive |
| hCoVOC43 | orf1ab | RDPRYVMRLQSAATI | 302 | 316 | Positive |
| hCoVOC43 | orf1ab | RDVSVEDVCFANKFE | 2802 | 2816 | Positive |
| hCoVOC43 | orf1ab | REDLEIGYTVDCSCG | 1772 | 1786 | Positive |
| hCoVOC43 | orf1ab | REIVMNASPYDLEVL | 78 | 92 | Positive |
| hCoVOC43 | orf1ab | REYLESYNTATTAGF | 6386 | 6400 | Positive |
| hCoVOC43 | orf1ab | RHDIALDDDARTFVQ | 1484 | 1498 | Positive |
| hCoVOC43 | orf1ab | RLKLFAAETQKATEE | 5426 | 5440 | Positive |
| hCoVOC43 | orf1ab | RLYYEALSFEEQDEI | 4878 | 4892 | Positive |
| hCoVOC43 | orf1ab | RSDGTFEEMALTTFM | 3154 | 3168 | Positive |
| hCoVOC43 | orf1ab | RSINHDLFSFIMFVG | 3678 | 3692 | Positive |
| hCoVOC43 | orf1ab | RTRLFKGTVCWIMAS | 3552 | 3566 | Positive |
| hCoVOC43 | orf1ab | RTVAGVSILSTMTGR | 4920 | 4934 | Positive |
| hCoVOC43 | orf1ab | RVKDCKFVAEHDFFT | 4458 | 4472 | Positive |
| hCoVOC43 | orf1ab | RYCTVQGPPGTGKSH | 5572 | 5586 | Positive |
| hCoVOC43 | orf1ab | SAKHVQGNVAKIAGV | 2682 | 2696 | Positive |
| hCoVOC43 | orf1ab | SDDGVVCYNSDYASK | 5124 | 5138 | Positive |
| hCoVOC43 | orf1ab | SDFGTSFVSKIVHFF | 646 | 660 | Positive |
| hCoVOC43 | orf1ab | SDFQHKLKKACCKTG | 2710 | 2724 | Positive |
| hCoVOC43 | orf1ab | SDFSLADSVVEVVTT | 784 | 798 | Positive |
| hCoVOC43 | orf1ab | SDFTVLFDRLSLTVM | 3310 | 3324 | Positive |
| hCoVOC43 | orf1ab | SDFVDLESVIQDYEN | 1032 | 1046 | Positive |
| hCoVOC43 | orf1ab | SDFYLPKIGFLPTFV | 2220 | 2234 | Positive |

| | | | | | |
|---|---|---|---|---|---|
| hCoVOC43 | orf1ab | SDGTNKQLNEISDDC | 4088 | 4102 | Positive |
| hCoVOC43 | orf1ab | SDIYFVKNIPRYASA | 688 | 702 | Positive |
| hCoVOC43 | orf1ab | SDKCSVEDFNVWALS | 3468 | 3482 | Positive |
| hCoVOC43 | orf1ab | SDLVIDALASMTGVS | 3490 | 3504 | Positive |
| hCoVOC43 | orf1ab | SDRFYRLANECAQVL | 5014 | 5028 | Positive |
| hCoVOC43 | orf1ab | SDTPCVYMDGMDAKQ | 6342 | 6356 | Positive |
| hCoVOC43 | orf1ab | SDVAFNRYLSLYNKY | 3182 | 3196 | Positive |
| hCoVOC43 | orf1ab | SDVDGLKYTKILKDD | 4166 | 4180 | Positive |
| hCoVOC43 | orf1ab | SDVWHKQLLLNRGVY | 536 | 550 | Positive |
| hCoVOC43 | orf1ab | SEISVIMHLQWLVMY | 3104 | 3118 | Positive |
| hCoVOC43 | orf1ab | SELTNGVYYRATTTY | 5500 | 5514 | Positive |
| hCoVOC43 | orf1ab | SEPPKVAAKICIVDN | 806 | 820 | Positive |
| hCoVOC43 | orf1ab | SERELILSWEIGKVK | 5456 | 5470 | Positive |
| hCoVOC43 | orf1ab | SESGAKETKEINIIK | 2026 | 2040 | Positive |
| hCoVOC43 | orf1ab | SESKCWVEHDINNGP | 5160 | 5174 | Positive |
| hCoVOC43 | orf1ab | SEVDMICSTTAQKLE | 34 | 48 | Positive |
| hCoVOC43 | orf1ab | SGSDFSLADSVVEVV | 782 | 796 | Positive |
| hCoVOC43 | orf1ab | SHEKASLNSLTYFNR | 1988 | 2002 | Positive |
| hCoVOC43 | orf1ab | SHSMAVVDGKQIDDH | 1224 | 1238 | Positive |
| hCoVOC43 | orf1ab | SIHSYFITDEKSGGS | 6724 | 6738 | Positive |
| hCoVOC43 | orf1ab | SKCWVEHDINNGPHE | 5162 | 5176 | Positive |
| hCoVOC43 | orf1ab | SKELLDVSLDSLGAA | 616 | 630 | Positive |
| hCoVOC43 | orf1ab | SKLWHYCSTLHNEIL | 3862 | 3876 | Positive |
| hCoVOC43 | orf1ab | SLAIDAYPLVYHENE | 5226 | 5240 | Positive |
| hCoVOC43 | orf1ab | SLEACYVRGCNPKGW | 112 | 126 | Positive |
| hCoVOC43 | orf1ab | SLETFTVCADGFMPF | 568 | 582 | Positive |
| hCoVOC43 | orf1ab | SLHLTFKIVQWQEAW | 1682 | 1696 | Positive |
| hCoVOC43 | orf1ab | SLTYFNRPSLVDDNK | 1996 | 2010 | Positive |
| hCoVOC43 | orf1ab | SMACPIVVAVIDQDF | 2834 | 2848 | Positive |
| hCoVOC43 | orf1ab | SMAVVDGKQIDDHRI | 1226 | 1240 | Positive |
| hCoVOC43 | orf1ab | SMFDVDKKSLNALIA | 2586 | 2600 | Positive |
| hCoVOC43 | orf1ab | SMPKIIKVFYELDND | 868 | 882 | Positive |
| hCoVOC43 | orf1ab | SMPVLYKYLNSPMER | 6806 | 6820 | Positive |
| hCoVOC43 | orf1ab | SMPYHPNTVDCQDDR | 4656 | 4670 | Positive |
| hCoVOC43 | orf1ab | SMVLIDVDVPFTAHF | 1182 | 1196 | Positive |
| hCoVOC43 | orf1ab | SNHAFWVFSYCRKLG | 3136 | 3150 | Positive |
| hCoVOC43 | orf1ab | SNHDLYCSVHKGAHV | 6154 | 6168 | Positive |
| hCoVOC43 | orf1ab | SNSKLWHYCSTLHNE | 3860 | 3874 | Positive |
| hCoVOC43 | orf1ab | SQLPLDVYDLTMPSQ | 754 | 768 | Positive |
| hCoVOC43 | orf1ab | STFNVLQDVLSLRHD | 1472 | 1486 | Positive |
| hCoVOC43 | orf1ab | SVPSDVSLTYLLGTA | 1396 | 1410 | Positive |
| hCoVOC43 | orf1ab | SVRSDGTFEEMALTT | 3152 | 3166 | Positive |
| hCoVOC43 | orf1ab | SYCRVGLCEEADEGI | 2968 | 2982 | Positive |
| hCoVOC43 | orf1ab | SYVLTHDVCRVCGFW | 4340 | 4354 | Positive |
| hCoVOC43 | orf1ab | TCRTDMEKDFIALDQ | 6654 | 6668 | Positive |
| hCoVOC43 | orf1ab | TCYRCPKEIVDTVSA | 5736 | 5750 | Positive |
| hCoVOC43 | orf1ab | TDCVFYFAVRKEGQD | 6588 | 6602 | Positive |
| hCoVOC43 | orf1ab | TDHKYVLSVSPYVCN | 5334 | 5348 | Positive |
| hCoVOC43 | orf1ab | TDIPQIKIVLLCYLF | 3748 | 3762 | Positive |

| | | | | | |
|---|---|---|---|---|---|
| hCoVOC43 | orf1ab | TDIVVVDEVSMLTNY | 5664 | 5678 | Positive |
| hCoVOC43 | orf1ab | TDLKFIDKLNVLFDG | 6526 | 6540 | Positive |
| hCoVOC43 | orf1ab | TDTTVQSKDTNFLNR | 4364 | 4378 | Positive |
| hCoVOC43 | orf1ab | TDVAYHTVTDVKQVG | 2472 | 2486 | Positive |
| hCoVOC43 | orf1ab | TDVDDYILANECTER | 5412 | 5426 | Positive |
| hCoVOC43 | orf1ab | TDVKCANVVLLNCLQ | 3840 | 3854 | Positive |
| hCoVOC43 | orf1ab | TDVKQVGCSMRLFYD | 2480 | 2494 | Positive |
| hCoVOC43 | orf1ab | TDVTGKLSDCLYLKN | 1850 | 1864 | Positive |
| hCoVOC43 | orf1ab | TEDLCDGSVVIKEPV | 326 | 340 | Positive |
| hCoVOC43 | orf1ab | TERLKLFAAETQKAT | 5424 | 5438 | Positive |
| hCoVOC43 | orf1ab | TEWPTATGDVVLATD | 1952 | 1966 | Positive |
| hCoVOC43 | orf1ab | TEYYEFLNKHFSMMI | 5108 | 5122 | Positive |
| hCoVOC43 | orf1ab | TFKEQPTVKEIISMP | 856 | 870 | Positive |
| hCoVOC43 | orf1ab | TGFCSKHQWNCIDCD | 2430 | 2444 | Positive |
| hCoVOC43 | orf1ab | TGYYGCWRHSVTCDY | 6120 | 6134 | Positive |
| hCoVOC43 | orf1ab | TIQDSDGTNKQLNEI | 4084 | 4098 | Positive |
| hCoVOC43 | orf1ab | TIQEIVSERELILSW | 5450 | 5464 | Positive |
| hCoVOC43 | orf1ab | TKEEAVKRVRAWVGF | 5974 | 5988 | Positive |
| hCoVOC43 | orf1ab | TLHNEILATSDLSVA | 3870 | 3884 | Positive |
| hCoVOC43 | orf1ab | TMADGSPQPYCYTEG | 2908 | 2922 | Positive |
| hCoVOC43 | orf1ab | TMLDLCYALRHFDRN | 4492 | 4506 | Positive |
| hCoVOC43 | orf1ab | TMLVKMDGDDVYLPY | 5182 | 5196 | Positive |
| hCoVOC43 | orf1ab | TMYITPVLFTLLYNN | 3616 | 3630 | Positive |
| hCoVOC43 | orf1ab | TNLICKALYLDYVQH | 508 | 522 | Positive |
| hCoVOC43 | orf1ab | TSDKFDFIIGHGMSF | 1244 | 1258 | Positive |
| hCoVOC43 | orf1ab | TSDSQVEEDVEMSDF | 1020 | 1034 | Positive |
| hCoVOC43 | orf1ab | TSSDFTVLFDRLSLT | 3308 | 3322 | Positive |
| hCoVOC43 | orf1ab | TTAGFTFWVYKTFDF | 6396 | 6410 | Positive |
| hCoVOC43 | orf1ab | TTHESSSAVNMQQIY | 5776 | 5790 | Positive |
| hCoVOC43 | orf1ab | TTKVKSLSMIRGPPR | 6560 | 6574 | Positive |
| hCoVOC43 | orf1ab | TVEAALDLSKELKRP | 2454 | 2468 | Positive |
| hCoVOC43 | orf1ab | TVKYFECTGGIDICS | 1522 | 1536 | Positive |
| hCoVOC43 | orf1ab | TVMSYQMRGCMLVLT | 3322 | 3336 | Positive |
| hCoVOC43 | orf1ab | TYFNRPSLVDDNKFD | 1998 | 2012 | Positive |
| hCoVOC43 | orf1ab | VANPTEDLCDGSVVI | 322 | 336 | Positive |
| hCoVOC43 | orf1ab | VASNSKLWHYCSTLH | 3858 | 3872 | Positive |
| hCoVOC43 | orf1ab | VAVELFAKRSVRHHP | 6474 | 6488 | Positive |
| hCoVOC43 | orf1ab | VDCQDDRCIIHCANF | 4664 | 4678 | Positive |
| hCoVOC43 | orf1ab | VDDTVDMEEFYAVVI | 896 | 910 | Positive |
| hCoVOC43 | orf1ab | VDDVDDGGDSSESGA | 2016 | 2030 | Positive |
| hCoVOC43 | orf1ab | VDIMSHFYMEADTVV | 357 | 370 | Positive |
| hCoVOC43 | orf1ab | VDMEEFYAVVIDAIE | 900 | 914 | Positive |
| hCoVOC43 | orf1ab | VDMICSTTAQKLETD | 36 | 50 | Positive |
| hCoVOC43 | orf1ab | VDNDLYPFVSDSVAT | 6892 | 6906 | Positive |
| hCoVOC43 | orf1ab | VDSKCLTSIEEVCDD | 3902 | 3916 | Positive |
| hCoVOC43 | orf1ab | VDVYDMWLTGCKYVV | 2072 | 2086 | Positive |
| hCoVOC43 | orf1ab | VEHDINNGPHEFCSQ | 5166 | 5180 | Positive |
| hCoVOC43 | orf1ab | VETSDSQVEEDVEMS | 1018 | 1032 | Positive |
| hCoVOC43 | orf1ab | VFIQKYGLEDYAFEH | 6670 | 6684 | Positive |

| | | | | | |
|---|---|---|---|---|---|
| hCoVOC43 | orf1ab | VGMVFVTLRSINHDL | 3670 | 3684 | Positive |
| hCoVOC43 | orf1ab | VIELIVSYALYTAWF | 2310 | 2324 | Positive |
| hCoVOC43 | orf1ab | VIYSQTAETAHSVNV | 5840 | 5854 | Positive |
| hCoVOC43 | orf1ab | VKSVPNMHVVVVEND | 2520 | 2534 | Positive |
| hCoVOC43 | orf1ab | VLARKHETCCSQSDR | 5002 | 5016 | Positive |
| hCoVOC43 | orf1ab | VLPVNPVLHTKSAAG | 432 | 446 | Positive |
| hCoVOC43 | orf1ab | VLQDVLSLRHDIALD | 1476 | 1490 | Positive |
| hCoVOC43 | orf1ab | VLSLRHDIALDDDAR | 1480 | 1494 | Positive |
| hCoVOC43 | orf1ab | VLYYQNNVFMSESKC | 5150 | 5164 | Positive |
| hCoVOC43 | orf1ab | VNDDTSETKYVKSLS | 2056 | 2070 | Positive |
| hCoVOC43 | orf1ab | VPDKSVYDQVVDNVY | 4058 | 4072 | Positive |
| hCoVOC43 | orf1ab | VPQENYSSIRFASVY | 5538 | 5552 | Positive |
| hCoVOC43 | orf1ab | VQGPPGTGKSHLAIG | 5576 | 5590 | Positive |
| hCoVOC43 | orf1ab | VRHHPELKLFRNLNI | 6484 | 6498 | Positive |
| hCoVOC43 | orf1ab | VSDSVATYFGDCITL | 6900 | 6914 | Positive |
| hCoVOC43 | orf1ab | VSVEDVCFANKFEQF | 2804 | 2818 | Positive |
| hCoVOC43 | orf1ab | VTETMFDVYVDTFLS | 2572 | 2586 | Positive |
| hCoVOC43 | orf1ab | VTFIDGLSCFKIGRR | 720 | 734 | Positive |
| hCoVOC43 | orf1ab | VVAKIDKEDVVIFIN | 6452 | 6466 | Positive |
| hCoVOC43 | orf1ab | VVDDHVGLLDQAWRV | 834 | 848 | Positive |
| hCoVOC43 | orf1ab | VVDEVSMLTNYELSV | 5668 | 5682 | Positive |
| hCoVOC43 | orf1ab | VVDVNDSHSMAVVDG | 1218 | 1232 | Positive |
| hCoVOC43 | orf1ab | VVENDADKANFLNAA | 2530 | 2544 | Positive |
| hCoVOC43 | orf1ab | VVEVVTTSLTPCGYS | 792 | 806 | Positive |
| hCoVOC43 | orf1ab | VVIDAIEEKLSPCKE | 908 | 922 | Positive |
| hCoVOC43 | orf1ab | VVIGTTKFYGGWDDM | 4952 | 4966 | Positive |
| hCoVOC43 | orf1ab | VVIKEPVHVYADDSI | 334 | 348 | Positive |
| hCoVOC43 | orf1ab | VVLELDPPCKFTVQD | 4184 | 4198 | Positive |
| hCoVOC43 | orf1ab | VVQYEADRRAFVDYT | 2290 | 2304 | Positive |
| hCoVOC43 | orf1ab | VVSHICKCGESMVLI | 1172 | 1186 | Positive |
| hCoVOC43 | orf1ab | VVSKSKELLDVSLDS | 612 | 626 | Positive |
| hCoVOC43 | orf1ab | VVVDKVGDTDCVFYF | 6580 | 6594 | Positive |
| hCoVOC43 | orf1ab | VVVVENDADKANFLN | 2528 | 2542 | Positive |
| hCoVOC43 | orf1ab | VYENKLKAKNESSSL | 5752 | 5766 | Positive |
| hCoVOC43 | orf1ab | VYETDANKLILINDV | 1454 | 1468 | Positive |
| hCoVOC43 | orf1ab | WGYIGSLSSNHDLYC | 6146 | 6160 | Positive |
| hCoVOC43 | orf1ab | WSVDAFNQFSSDFQH | 2700 | 2714 | Positive |
| hCoVOC43 | orf1ab | YCAFTAPEDDDFLEE | 958 | 972 | Positive |
| hCoVOC43 | orf1ab | YCEDHKPQYSFKLVM | 5368 | 5382 | Positive |
| hCoVOC43 | orf1ab | YDCFCNNINWNVEYP | 6182 | 6196 | Positive |
| hCoVOC43 | orf1ab | YDQVHDEPKGKFSKK | 222 | 236 | Positive |
| hCoVOC43 | orf1ab | YENVCFEFYTTEPEF | 1044 | 1058 | Positive |
| hCoVOC43 | orf1ab | YFEIYDGGCIPASQV | 4844 | 4858 | Positive |
| hCoVOC43 | orf1ab | YHENEEYQKVFRVYL | 5236 | 5250 | Positive |
| hCoVOC43 | orf1ab | YKGSNLEEEILLMLA | 3702 | 3716 | Positive |
| hCoVOC43 | orf1ab | YLDYVQHKCGNLHQR | 516 | 530 | Positive |
| hCoVOC43 | orf1ab | YPLVYHENEEYQKVF | 5232 | 5246 | Positive |
| hCoVOC43 | orf1ab | YVPSVEYTYTDEVIY | 3650 | 3664 | Positive |
| hCoVOC43 | orf1ab | YYEFLNKHFSMMILS | 5110 | 5124 | Positive |

| | | | | | |
|---|---|---|---|---|---|
| hCoVOC43 | orf1ab | YYPVVVDDHVGLLDQ | 830 | 844 | Positive |
| hCoVOC43 | orf1ab | ACKCGVKQEQRTGLD | 1748 | 1762 | Negative |
| hCoVOC43 | orf1ab | ACSQLAKAMDTFTNN | 3212 | 3226 | Negative |
| hCoVOC43 | orf1ab | ADGSPQPYCYTEGLM | 2910 | 2924 | Negative |
| hCoVOC43 | orf1ab | ADGVQCYTPHSQISY | 2876 | 2890 | Negative |
| hCoVOC43 | orf1ab | ADHLIDLSDCVVLVT | 6070 | 6084 | Negative |
| hCoVOC43 | orf1ab | ADNKVIYTTEIASKL | 2160 | 2174 | Negative |
| hCoVOC43 | orf1ab | AEEKLDNPSSSEVDM | 24 | 38 | Negative |
| hCoVOC43 | orf1ab | AEEYREYLESYNTAT | 6382 | 6396 | Negative |
| hCoVOC43 | orf1ab | AFVKWTMFMYVTTNM | 3576 | 3590 | Negative |
| hCoVOC43 | orf1ab | AGEMPCAIINDKVVA | 6440 | 6453 | Negative |
| hCoVOC43 | orf1ab | AIGLAVFYCTARVVY | 5588 | 5602 | Negative |
| hCoVOC43 | orf1ab | AIINDKVVAKIDKED | 6446 | 6460 | Negative |
| hCoVOC43 | orf1ab | AITDYNYYKYNLPTM | 4814 | 4828 | Negative |
| hCoVOC43 | orf1ab | AKETKEINIIKLSGV | 2030 | 2044 | Negative |
| hCoVOC43 | orf1ab | ALQSEFVNMASFVEY | 3926 | 3940 | Negative |
| hCoVOC43 | orf1ab | ANAKGFIRFPEVLRE | 2942 | 2956 | Negative |
| hCoVOC43 | orf1ab | ANECTERLKLFAAET | 5420 | 5434 | Negative |
| hCoVOC43 | orf1ab | ASGLSTDVQLRAFDI | 4392 | 4406 | Negative |
| hCoVOC43 | orf1ab | ATGLFADRDGYSFKK | 6020 | 6034 | Negative |
| hCoVOC43 | orf1ab | AVSANVCALMSCNGN | 5064 | 5078 | Negative |
| hCoVOC43 | orf1ab | CDGVNVTKHKCDINY | 1596 | 1610 | Negative |
| hCoVOC43 | orf1ab | CIWSVDAFNQFSSDF | 2698 | 2712 | Negative |
| hCoVOC43 | orf1ab | CKFVAEHDFFTFDVE | 4462 | 4476 | Negative |
| hCoVOC43 | orf1ab | CLGIGDSAVTYSRLI | 5940 | 5954 | Negative |
| hCoVOC43 | orf1ab | CMLLCDILSIYAGCE | 4508 | 4522 | Negative |
| hCoVOC43 | orf1ab | CSFEDELTPSDVYQQ | 3528 | 3542 | Negative |
| hCoVOC43 | orf1ab | DDCNWPLVIIANRYN | 4100 | 4114 | Negative |
| hCoVOC43 | orf1ab | DDDARTFVQSNVDVV | 1490 | 1504 | Negative |
| hCoVOC43 | orf1ab | DDKYKATGDLAVCLG | 5928 | 5942 | Negative |
| hCoVOC43 | orf1ab | DDLYVKRYERGCITF | 1966 | 1980 | Negative |
| hCoVOC43 | orf1ab | DDMLRRLIKDVDNPV | 4964 | 4978 | Negative |
| hCoVOC43 | orf1ab | DDTSETKYVKSLSIV | 2058 | 2072 | Negative |
| hCoVOC43 | orf1ab | DEKSGGSKSVCTVID | 6732 | 6746 | Negative |
| hCoVOC43 | orf1ab | DFEIACKCGVKQEQR | 1744 | 1758 | Negative |
| hCoVOC43 | orf1ab | DFNRIASCKWTDVDD | 5402 | 5416 | Negative |
| hCoVOC43 | orf1ab | DFNTILNTACGVFEV | 882 | 896 | Negative |
| hCoVOC43 | orf1ab | DGNAAITDYNYYKYN | 4810 | 4824 | Negative |
| hCoVOC43 | orf1ab | DGSCSCVSTDTTVQS | 4356 | 4370 | Negative |
| hCoVOC43 | orf1ab | DLEIGYTVDCSCGKK | 1774 | 1788 | Negative |
| hCoVOC43 | orf1ab | DLGVLIQNSAKHVQG | 2674 | 2688 | Negative |
| hCoVOC43 | orf1ab | DNDFNTILNTACGVF | 880 | 894 | Negative |
| hCoVOC43 | orf1ab | DNGALEAFKRSNNGV | 6542 | 6556 | Negative |
| hCoVOC43 | orf1ab | DNKFDVLKVDDVDDG | 2008 | 2023 | Negative |
| hCoVOC43 | orf1ab | DPKKTYLDFIQQGGT | 4254 | 4268 | Negative |
| hCoVOC43 | orf1ab | DSDVDTKCLADSVMS | 2630 | 2644 | Negative |
| hCoVOC43 | orf1ab | DVDDGGDSSESGAKE | 2018 | 2032 | Negative |
| hCoVOC43 | orf1ab | DVEEDDVEGEETDLT | 974 | 988 | Negative |
| hCoVOC43 | orf1ab | DVTKLYLGGMSYYCE | 5356 | 5370 | Negative |

| | | | | | |
|---|---|---|---|---|---|
| hCoVOC43 | orf1ab | DVVLATDDLYVKRYE | 1960 | 1974 | Negative |
| hCoVOC43 | orf1ab | DVYVDTFLSMFDVDK | 2578 | 2592 | Negative |
| hCoVOC43 | orf1ab | DYAKDNTVLQALQSE | 3916 | 3930 | Negative |
| hCoVOC43 | orf1ab | DYARESIFCSNTYGV | 6508 | 6522 | Negative |
| hCoVOC43 | orf1ab | DYVIAAPGCGVAIAD | 4590 | 4604 | Negative |
| hCoVOC43 | orf1ab | DYVIYSQTAETAHSV | 5838 | 5852 | Negative |
| hCoVOC43 | orf1ab | DYVPLKSATCITRCN | 6358 | 6372 | Negative |
| hCoVOC43 | orf1ab | EGWRVVNKFYQINGV | 1506 | 1520 | Negative |
| hCoVOC43 | orf1ab | EIDGNVMHANYLFWR | 7014 | 7028 | Negative |
| hCoVOC43 | orf1ab | EILATSDLSVAFEKL | 3874 | 3888 | Negative |
| hCoVOC43 | orf1ab | EILLMLASLFGTYTW | 3710 | 3724 | Negative |
| hCoVOC43 | orf1ab | EIVMCGGCYYVKPGG | 5030 | 5044 | Negative |
| hCoVOC43 | orf1ab | ELDNDFNTILNTACG | 878 | 892 | Negative |
| hCoVOC43 | orf1ab | ELEGVGAKVSAFLQK | 922 | 936 | Negative |
| hCoVOC43 | orf1ab | ELFMLSTLHWSFRLL | 2342 | 2356 | Negative |
| hCoVOC43 | orf1ab | ELFNKYFKHWSMPYH | 4646 | 4660 | Negative |
| hCoVOC43 | orf1ab | ELHWAPEFPWMFEDA | 10 | 24 | Negative |
| hCoVOC43 | orf1ab | ELIVSYALYTAWFYP | 2312 | 2326 | Negative |
| hCoVOC43 | orf1ab | ELKRPIQPTDVAYHT | 2464 | 2478 | Negative |
| hCoVOC43 | orf1ab | ELLDVSLDSLGAAIH | 618 | 632 | Negative |
| hCoVOC43 | orf1ab | ELNGVVVDKVGDTDC | 6576 | 6590 | Negative |
| hCoVOC43 | orf1ab | ELRYMNANGLRPPKN | 3796 | 3810 | Negative |
| hCoVOC43 | orf1ab | ELTCLRYFAKVGREI | 6090 | 6104 | Negative |
| hCoVOC43 | orf1ab | EVLLQDALQSREAVL | 90 | 104 | Negative |
| hCoVOC43 | orf1ab | EVLREGLVRIVRTRS | 2952 | 2966 | Negative |
| hCoVOC43 | orf1ab | EVVTTSLTPCGYSEP | 794 | 808 | Negative |
| hCoVOC43 | orf1ab | EVVVNPANGHMAHGG | 1294 | 1308 | Negative |
| hCoVOC43 | orf1ab | FCGRDVFDLIYQLFK | 3004 | 3018 | Negative |
| hCoVOC43 | orf1ab | FDCQWDLIISDMYDP | 6916 | 6930 | Negative |
| hCoVOC43 | orf1ab | FDGRDNGALEAFKRS | 6538 | 6552 | Negative |
| hCoVOC43 | orf1ab | FDVIVAWHVVRDPRY | 292 | 306 | Negative |
| hCoVOC43 | orf1ab | FEDELTPSDVYQQLA | 3530 | 3544 | Negative |
| hCoVOC43 | orf1ab | FHQKCLKSIAATRGV | 4936 | 4950 | Negative |
| hCoVOC43 | orf1ab | FLLDDLVPRAYYLAV | 582 | 596 | Negative |
| hCoVOC43 | orf1ab | FLNINDCTRIVPAKV | 5620 | 5634 | Negative |
| hCoVOC43 | orf1ab | FPEVLREGLVRIVRT | 2950 | 2964 | Negative |
| hCoVOC43 | orf1ab | FPLQLGFSTGIDFVV | 6004 | 6018 | Negative |
| hCoVOC43 | orf1ab | FSTGIDFVVEATGLF | 6010 | 6024 | Negative |
| hCoVOC43 | orf1ab | FTDESFYKNMYLRSA | 5280 | 5294 | Negative |
| hCoVOC43 | orf1ab | FTNRFVPVGESFGKS | 1576 | 1590 | Negative |
| hCoVOC43 | orf1ab | FVAEHDFFTFDVEGS | 4464 | 4478 | Negative |
| hCoVOC43 | orf1ab | FVEYKITEWPTATGD | 1946 | 1960 | Negative |
| hCoVOC43 | orf1ab | GALEAFKRSNNGVYI | 6544 | 6558 | Negative |
| hCoVOC43 | orf1ab | GCEQSYFTKKDWYDF | 4520 | 4534 | Negative |
| hCoVOC43 | orf1ab | GCHTGTDFNGDFYGP | 3416 | 3430 | Negative |
| hCoVOC43 | orf1ab | GDKLDQFFVVKRTDL | 4432 | 4446 | Negative |
| hCoVOC43 | orf1ab | GESMVLIDVDVPFTA | 1180 | 1194 | Negative |
| hCoVOC43 | orf1ab | GFDMLDNYKAIDVVQ | 2278 | 2292 | Negative |
| hCoVOC43 | orf1ab | GGIDICSQDKVFGYV | 1530 | 1544 | Negative |

| | | | | | |
|---|---|---|---|---|---|
| hCoVOC43 | orf1ab | GGMIRYYDVMANGGT | 2416 | 2430 | Negative |
| hCoVOC43 | orf1ab | GHYTGQAGEMPCAII | 6434 | 6448 | Negative |
| hCoVOC43 | orf1ab | GIGDSAVTYSRLISL | 5942 | 5956 | Negative |
| hCoVOC43 | orf1ab | GINYLCKPKVEIDGN | 7004 | 7018 | Negative |
| hCoVOC43 | orf1ab | GIVKMVNPTSKVEPC | 3248 | 3262 | Negative |
| hCoVOC43 | orf1ab | GKFVQVPVGIKDPVS | 4326 | 4340 | Negative |
| hCoVOC43 | orf1ab | GKIVYAILSDVDGLK | 4158 | 4172 | Negative |
| hCoVOC43 | orf1ab | GKLLIRDTNKEVFVG | 7074 | 7088 | Negative |
| hCoVOC43 | orf1ab | GKQIDDHRITSITSD | 1232 | 1246 | Negative |
| hCoVOC43 | orf1ab | GKSLGNVFCDGVNVT | 1588 | 1602 | Negative |
| hCoVOC43 | orf1ab | GKTVLGEYVFDKSEL | 5488 | 5502 | Negative |
| hCoVOC43 | orf1ab | GKVFFQFDNLSSEDL | 1612 | 1626 | Negative |
| hCoVOC43 | orf1ab | GLAVFYCTARVVYTA | 5590 | 5604 | Negative |
| hCoVOC43 | orf1ab | GLDAMFFYGDVVSHI | 1162 | 1176 | Negative |
| hCoVOC43 | orf1ab | GLDAVMHFGTLSRED | 1760 | 1774 | Negative |
| hCoVOC43 | orf1ab | GLFADRDGYSFKKAV | 6022 | 6036 | Negative |
| hCoVOC43 | orf1ab | GLFSLMNSLFRMPLG | 3772 | 3786 | Negative |
| hCoVOC43 | orf1ab | GLKYTKILKDDGNFV | 4170 | 4184 | Negative |
| hCoVOC43 | orf1ab | GLLDQAWRVPCAGRR | 840 | 854 | Negative |
| hCoVOC43 | orf1ab | GLLKEGSSVDLKHFF | 4792 | 4806 | Negative |
| hCoVOC43 | orf1ab | GLMQNASLYSSLVPH | 2922 | 2936 | Negative |
| hCoVOC43 | orf1ab | GLRPPKNSFEALMLN | 3804 | 3818 | Negative |
| hCoVOC43 | orf1ab | GLSTDVQLRAFDIYN | 4394 | 4408 | Negative |
| hCoVOC43 | orf1ab | GLVFGLYKQSCTGSP | 5384 | 5398 | Negative |
| hCoVOC43 | orf1ab | GLWCLMPTYTVHKSD | 2770 | 2784 | Negative |
| hCoVOC43 | orf1ab | GLYKQSCTGSPYIDD | 5388 | 5402 | Negative |
| hCoVOC43 | orf1ab | GMAITVKPDATTSQD | 4284 | 4298 | Negative |
| hCoVOC43 | orf1ab | GMSKINKYGLELHWA | 1 | 14 | Negative |
| hCoVOC43 | orf1ab | GQDVIFSQFDSLGVS | 6600 | 6614 | Negative |
| hCoVOC43 | orf1ab | GRDNGALEAFKRSNN | 6540 | 6554 | Negative |
| hCoVOC43 | orf1ab | GRDVFDLIYQLFKGL | 3006 | 3020 | Negative |
| hCoVOC43 | orf1ab | GRSIVYETDANKLIL | 1450 | 1464 | Negative |
| hCoVOC43 | orf1ab | GSCSFEDELTPSDVY | 3526 | 3540 | Negative |
| hCoVOC43 | orf1ab | GSGSGMSKINKYGLE | 1 | 10 | Negative |
| hCoVOC43 | orf1ab | GSGSGSGMSKINKYG | 1 | 8 | Negative |
| hCoVOC43 | orf1ab | GTAKKQVVLVSNNQE | 1408 | 1422 | Negative |
| hCoVOC43 | orf1ab | GTATEYASNSSILSL | 4234 | 4248 | Negative |
| hCoVOC43 | orf1ab | GTDFNGDFYGPYKDA | 3420 | 3434 | Negative |
| hCoVOC43 | orf1ab | GTGMAITVKPDATTS | 4282 | 4296 | Negative |
| hCoVOC43 | orf1ab | GTILVDNDLYPFVSD | 6888 | 6902 | Negative |
| hCoVOC43 | orf1ab | GTLSREDLEIGYTVD | 1768 | 1782 | Negative |
| hCoVOC43 | orf1ab | GVCATGDCYVSTGGK | 1334 | 1348 | Negative |
| hCoVOC43 | orf1ab | GVKPLLYVDQYGCDY | 246 | 260 | Negative |
| hCoVOC43 | orf1ab | GVKQEQRTGLDAVMH | 1752 | 1766 | Negative |
| hCoVOC43 | orf1ab | GVLIQNSAKHVQGNV | 2676 | 2690 | Negative |
| hCoVOC43 | orf1ab | GVLTLDNQDLNGKWY | 4572 | 4586 | Negative |
| hCoVOC43 | orf1ab | GVNVTKHKCDINYKG | 1598 | 1612 | Negative |
| hCoVOC43 | orf1ab | GVPFVVSIGYHYKEL | 4702 | 4716 | Negative |
| hCoVOC43 | orf1ab | GVQCYTPHSQISYSN | 2878 | 2892 | Negative |

| | | | | | |
|---|---|---|---|---|---|
| hCoVOC43 | orf1ab | GVSDVWHKQLLLNRG | 534 | 548 | Negative |
| hCoVOC43 | orf1ab | GVSILSTMTGRMFHQ | 4924 | 4938 | Negative |
| hCoVOC43 | orf1ab | GVSLETLLAAIKRLK | 3502 | 3516 | Negative |
| hCoVOC43 | orf1ab | GVSSNQSPQGNLGSN | 6612 | 6626 | Negative |
| hCoVOC43 | orf1ab | GVVKPGETFTVLAAY | 3350 | 3364 | Negative |
| hCoVOC43 | orf1ab | GVYNYKISVQELRYM | 3786 | 3800 | Negative |
| hCoVOC43 | orf1ab | GVYTNFKLIGHTVCD | 1918 | 1932 | Negative |
| hCoVOC43 | orf1ab | GWDDMLRRLIKDVDN | 4962 | 4976 | Negative |
| hCoVOC43 | orf1ab | GYVADFAYWFLTLCD | 1128 | 1142 | Negative |
| hCoVOC43 | orf1ab | HKPQYSFKLVMNGLV | 5372 | 5386 | Negative |
| hCoVOC43 | orf1ab | HRWDVVRPRIVQMFA | 6056 | 6070 | Negative |
| hCoVOC43 | orf1ab | HTVCDSLNAKLGFDS | 1928 | 1942 | Negative |
| hCoVOC43 | orf1ab | HVASASALYDLRTCC | 4746 | 4760 | Negative |
| hCoVOC43 | orf1ab | HVGLLDQAWRVPCAG | 838 | 852 | Negative |
| hCoVOC43 | orf1ab | ICSQDKVFGYVQQGI | 1534 | 1548 | Negative |
| hCoVOC43 | orf1ab | IDFVVEATGLFADRD | 6014 | 6028 | Negative |
| hCoVOC43 | orf1ab | IDVCWKHVIWDYARE | 6498 | 6512 | Negative |
| hCoVOC43 | orf1ab | IGGVPIIEVSQFQSK | 3824 | 3838 | Negative |
| hCoVOC43 | orf1ab | IGTNFPLQLGFSTGI | 6000 | 6014 | Negative |
| hCoVOC43 | orf1ab | IIVPDKSVYDQVVDN | 4056 | 4070 | Negative |
| hCoVOC43 | orf1ab | ILDSYSVILSTCDGQ | 5264 | 5278 | Negative |
| hCoVOC43 | orf1ab | ILGAGCFVDDLLKTD | 5202 | 5216 | Negative |
| hCoVOC43 | orf1ab | IMGSCSFEDELTPSD | 3524 | 3538 | Negative |
| hCoVOC43 | orf1ab | IMPMLTMCHALDCEL | 4610 | 4624 | Negative |
| hCoVOC43 | orf1ab | IRDTNKEVFVGDSLV | 7078 | 7092 | Negative |
| hCoVOC43 | orf1ab | ISADNKVIYTTEIAS | 2158 | 2172 | Negative |
| hCoVOC43 | orf1ab | ISDDCNWPLVIIANR | 4098 | 4112 | Negative |
| hCoVOC43 | orf1ab | IVGGMIRYYDVMANG | 2414 | 2428 | Negative |
| hCoVOC43 | orf1ab | IVMNMDVDTHRYRLS | 4718 | 4732 | Negative |
| hCoVOC43 | orf1ab | IVVVDEVSMLTNYEL | 5666 | 5680 | Negative |
| hCoVOC43 | orf1ab | KACNIAKSAYERDRA | 3966 | 3980 | Negative |
| hCoVOC43 | orf1ab | KAIDVVQYEADRRAF | 2286 | 2300 | Negative |
| hCoVOC43 | orf1ab | KATGDLAVCLGIGDS | 5932 | 5946 | Negative |
| hCoVOC43 | orf1ab | KCGVKQEQRTGLDAV | 1750 | 1764 | Negative |
| hCoVOC43 | orf1ab | KCIKCDLALKLKGLD | 1150 | 1164 | Negative |
| hCoVOC43 | orf1ab | KEFVEYKITEWPTAT | 1944 | 1958 | Negative |
| hCoVOC43 | orf1ab | KLELFNKYFKHWSMP | 4644 | 4658 | Negative |
| hCoVOC43 | orf1ab | KSSVKSYDSLVYTGV | 484 | 498 | Negative |
| hCoVOC43 | orf1ab | KVDILLTVDGVNFTN | 1564 | 1578 | Negative |
| hCoVOC43 | orf1ab | KVGDTDCVFYFAVRK | 6584 | 6598 | Negative |
| hCoVOC43 | orf1ab | KYGLEDYAFEHIVYG | 6674 | 6688 | Negative |
| hCoVOC43 | orf1ab | KYGLELHWAPEFPWM | 6 | 20 | Negative |
| hCoVOC43 | orf1ab | LDNGVIRDVSVEDVC | 2796 | 2810 | Negative |
| hCoVOC43 | orf1ab | LEKACNIAKSAYERD | 3964 | 3978 | Negative |
| hCoVOC43 | orf1ab | LGEYVFDKSELTNGV | 5492 | 5506 | Negative |
| hCoVOC43 | orf1ab | LGFSTGIDFVVEATG | 6008 | 6022 | Negative |
| hCoVOC43 | orf1ab | LKFIDKLNVLFDGRD | 6528 | 6542 | Negative |
| hCoVOC43 | orf1ab | LKGLDAMFFYGDVVS | 1160 | 1174 | Negative |
| hCoVOC43 | orf1ab | LLFDVIVAWHVVRDP | 290 | 304 | Negative |

| | | | | | |
|---|---|---|---|---|---|
| hCoVOC43 | orf1ab | LLRIVSSLVLARKHE | 4994 | 5008 | Negative |
| hCoVOC43 | orf1ab | LMGFKLDVTLDGYCK | 5956 | 5970 | Negative |
| hCoVOC43 | orf1ab | LRADQINDMVYSLLE | 7058 | 7072 | Negative |
| hCoVOC43 | orf1ab | LSMIRGPPRAELNGV | 6566 | 6580 | Negative |
| hCoVOC43 | orf1ab | LTDESCNNLVPTYLK | 2652 | 2666 | Negative |
| hCoVOC43 | orf1ab | LVGVLTLDNQDLNGK | 4570 | 4584 | Negative |
| hCoVOC43 | orf1ab | MDVDTHRYRLSLKDL | 4722 | 4736 | Negative |
| hCoVOC43 | orf1ab | MEADTVVNAFYGVAL | 364 | 378 | Negative |
| hCoVOC43 | orf1ab | MFADHLIDLSDCVVL | 6068 | 6082 | Negative |
| hCoVOC43 | orf1ab | MFDVYVDTFLSMFDV | 2576 | 2590 | Negative |
| hCoVOC43 | orf1ab | MICSTTAQKLETDGI | 38 | 52 | Negative |
| hCoVOC43 | orf1ab | MIRGPPRAELNGVVV | 6568 | 6582 | Negative |
| hCoVOC43 | orf1ab | MNGLVFGLYKQSCTG | 5382 | 5396 | Negative |
| hCoVOC43 | orf1ab | MTGVSLETLLAAIKR | 3500 | 3514 | Negative |
| hCoVOC43 | orf1ab | MVDIKQLLFVLEVVY | 4828 | 4842 | Negative |
| hCoVOC43 | orf1ab | NDYYRSLPGTFCGRD | 2994 | 3008 | Negative |
| hCoVOC43 | orf1ab | NGVRTVKYFECTGGI | 1518 | 1532 | Negative |
| hCoVOC43 | orf1ab | NPTSKVEPCVVSVTY | 3254 | 3268 | Negative |
| hCoVOC43 | orf1ab | NVTKHKCDINYKGKV | 1600 | 1614 | Negative |
| hCoVOC43 | orf1ab | PADSRDFLRVVFSQV | 1722 | 1736 | Negative |
| hCoVOC43 | orf1ab | PEFVKVLGLYVPKAT | 1056 | 1070 | Negative |
| hCoVOC43 | orf1ab | PFVSDSVATYFGDCI | 6898 | 6912 | Negative |
| hCoVOC43 | orf1ab | PKGVGSANIFIGDKV | 1812 | 1826 | Negative |
| hCoVOC43 | orf1ab | PLGVYNYKISVQELR | 3784 | 3798 | Negative |
| hCoVOC43 | orf1ab | PVLYKYLNSPMERVS | 6808 | 6822 | Negative |
| hCoVOC43 | orf1ab | PVQSRKFIVPWVMYL | 178 | 192 | Negative |
| hCoVOC43 | orf1ab | PVSYVLTHDVCRVCG | 4338 | 4352 | Negative |
| hCoVOC43 | orf1ab | PVYASYKVLDNGVIR | 2788 | 2802 | Negative |
| hCoVOC43 | orf1ab | QGGYVADFAYWFLTL | 1126 | 1140 | Negative |
| hCoVOC43 | orf1ab | QGRQIMGSCSFEDEL | 3520 | 3534 | Negative |
| hCoVOC43 | orf1ab | QIDDHRITSITSDKF | 1234 | 1248 | Negative |
| hCoVOC43 | orf1ab | QLPVYASYKVLDNGV | 2786 | 2800 | Negative |
| hCoVOC43 | orf1ab | QSRVISSFTCRTDME | 6646 | 6660 | Negative |
| hCoVOC43 | orf1ab | QTAETAHSVNVNRFN | 5844 | 5858 | Negative |
| hCoVOC43 | orf1ab | RAFVDYTGVLKIVIE | 2298 | 2312 | Negative |
| hCoVOC43 | orf1ab | RFDTRVLNNLNLPGC | 6294 | 6308 | Negative |
| hCoVOC43 | orf1ab | RLFYDRDGQRTYDDV | 2490 | 2504 | Negative |
| hCoVOC43 | orf1ab | RQIMGSCSFEDELTP | 3522 | 3536 | Negative |
| hCoVOC43 | orf1ab | RRICLSGRKIYEVER | 734 | 748 | Negative |
| hCoVOC43 | orf1ab | RSDKVDSTFVTEYYE | 5098 | 5112 | Negative |
| hCoVOC43 | orf1ab | RTDMEKDFIALDQDV | 6656 | 6670 | Negative |
| hCoVOC43 | orf1ab | SAKNRARTVAGVSIL | 4914 | 4928 | Negative |
| hCoVOC43 | orf1ab | SATIQEIVSERELIL | 5448 | 5462 | Negative |
| hCoVOC43 | orf1ab | SCGSVGYVIMGDCVK | 3390 | 3404 | Negative |
| hCoVOC43 | orf1ab | SCKWTDVDDYILANE | 5408 | 5422 | Negative |
| hCoVOC43 | orf1ab | SFGKSLGNVFCDGVN | 1586 | 1600 | Negative |
| hCoVOC43 | orf1ab | SFKDGLCMFWNCNVD | 6270 | 6284 | Negative |
| hCoVOC43 | orf1ab | SFVEYEVAKKNLDEA | 3936 | 3950 | Negative |
| hCoVOC43 | orf1ab | SGKMDTAAYREAACS | 3200 | 3214 | Negative |

| | | | | | |
|---|---|---|---|---|---|
| hCoVOC43 | orf1ab | SKGLLKEGSSVDLKH | 4790 | 4804 | Negative |
| hCoVOC43 | orf1ab | SKGVCATGDCYVSTG | 1332 | 1346 | Negative |
| hCoVOC43 | orf1ab | SKHQWNCIDCDSYKP | 2434 | 2448 | Negative |
| hCoVOC43 | orf1ab | SKKAYALIRGYRGVK | 234 | 248 | Negative |
| hCoVOC43 | orf1ab | SKRTRLFKGTVCWIM | 3550 | 3564 | Negative |
| hCoVOC43 | orf1ab | SLENVVYNLVKTGHY | 6422 | 6436 | Negative |
| hCoVOC43 | orf1ab | SLGNVFCDGVNVTKH | 1590 | 1604 | Negative |
| hCoVOC43 | orf1ab | SLGVSSNQSPQGNLG | 6610 | 6624 | Negative |
| hCoVOC43 | orf1ab | SRVPHIVRKDLTKYT | 4478 | 4492 | Negative |
| hCoVOC43 | orf1ab | TDLTIYNREMKCYER | 4444 | 4458 | Negative |
| hCoVOC43 | orf1ab | TFGVVKPGETFTVLA | 3348 | 3362 | Negative |
| hCoVOC43 | orf1ab | TLQNSRTPKYTFGVV | 3338 | 3352 | Negative |
| hCoVOC43 | orf1ab | TNFPLQLGFSTGIDF | 6002 | 6016 | Negative |
| hCoVOC43 | orf1ab | TNMFSITFCALCVIS | 3588 | 3602 | Negative |
| hCoVOC43 | orf1ab | TNMYKEARINDKKSK | 3994 | 4008 | Negative |
| hCoVOC43 | orf1ab | TSETKYVKSLSIVDV | 2060 | 2074 | Negative |
| hCoVOC43 | orf1ab | TTYYLDDVKKIEYKP | 1874 | 1888 | Negative |
| hCoVOC43 | orf1ab | VAGVSILSTMTGRMF | 4922 | 4936 | Negative |
| hCoVOC43 | orf1ab | VAKYTQLCQYLNTTT | 6840 | 6854 | Negative |
| hCoVOC43 | orf1ab | VDGLKYTKILKDDGN | 4168 | 4182 | Negative |
| hCoVOC43 | orf1ab | VDGVPFVVSIGYHYK | 4700 | 4714 | Negative |
| hCoVOC43 | orf1ab | VDGVYTNFKLIGHTV | 1916 | 1930 | Negative |
| hCoVOC43 | orf1ab | VDKNLITTANTGTSV | 2558 | 2572 | Negative |
| hCoVOC43 | orf1ab | VFGLYKQSCTGSPYI | 5386 | 5400 | Negative |
| hCoVOC43 | orf1ab | VILSTCDGQKFTDES | 5270 | 5284 | Negative |
| hCoVOC43 | orf1ab | VIVNDDTSETKYVKS | 2054 | 2068 | Negative |
| hCoVOC43 | orf1ab | VIVNNYDKSAGYPFN | 4858 | 4872 | Negative |
| hCoVOC43 | orf1ab | VKQVGCSMRLFYDRD | 2482 | 2496 | Negative |
| hCoVOC43 | orf1ab | VLQNNELMPAKLKIQ | 4120 | 4134 | Negative |
| hCoVOC43 | orf1ab | VPEGWRVVNKFYQIN | 1504 | 1518 | Negative |
| hCoVOC43 | orf1ab | VTGKLSDCLYLKNLK | 1852 | 1866 | Negative |
| hCoVOC43 | orf1ab | VVAVIDQDFGSTVFN | 2840 | 2854 | Negative |
| hCoVOC43 | orf1ab | VVNGKYFTFKQANNN | 1656 | 1670 | Negative |
| hCoVOC43 | orf1ab | VVSIGYHYKELGIVM | 4706 | 4720 | Negative |
| hCoVOC43 | orf1ab | WQEAWLEFRSGRPAR | 1692 | 1706 | Negative |
| hCoVOC43 | orf1ab | YASATIQEIVSEREL | 5446 | 5460 | Negative |
| hCoVOC43 | orf1ab | YCRARVEHPDVDGLC | 4308 | 4322 | Negative |
| hCoVOC43 | orf1ab | YKGVTTHESSSAVNM | 5772 | 5786 | Negative |
| hCoVOC43 | orf1ab | YKPLLENIDYFNMRR | 550 | 564 | Negative |
| hCoVOC43 | orf1ab | YNPLIVDIQQWGYIG | 6136 | 6150 | Negative |
| hCoVOC43 | orf1ab | YTGQAGEMPCAIIND | 6436 | 6450 | Negative |
| hCoVOC43 | orf1ab | YTNFKLIGHTVCDSL | 1920 | 1934 | Negative |
| hCoVOC43 | orf1ab | YTTEPEFVKVLGLYV | 1052 | 1066 | Negative |
| hCoVOC43 | orf1ab | YWGLFSLMNSLFRMP | 3770 | 3784 | Negative |
| hCoVOC43 | orf1ab | ACGVFEVDDTVDMEE | 890 | 904 | Overlapping |
| hCoVOC43 | orf1ab | AEHDFFTFDVEGSRV | 4466 | 4480 | Overlapping |
| hCoVOC43 | orf1ab | AIDAYPLVYHENEEY | 5228 | 5242 | Overlapping |
| hCoVOC43 | orf1ab | AIEEKLSPCKELEGV | 912 | 926 | Overlapping |
| hCoVOC43 | orf1ab | ARESIFCSNTYGVCM | 6510 | 6524 | Overlapping |

| | | | | | |
|---|---|---|---|---|---|
| hCoVOC43 | orf1ab | ATEFADKLVEVGLVG | 4558 | 4572 | Overlapping |
| hCoVOC43 | orf1ab | AVCLKHAEEYREYLE | 6376 | 6390 | Overlapping |
| hCoVOC43 | orf1ab | AVDDKYKATGDLAVC | 5926 | 5940 | Overlapping |
| hCoVOC43 | orf1ab | AYERDRAVAKKLERM | 3974 | 3988 | Overlapping |
| hCoVOC43 | orf1ab | CFEFYTTEPEFVKVL | 1048 | 1062 | Overlapping |
| hCoVOC43 | orf1ab | CWVEHDINNGPHEFC | 5164 | 5178 | Overlapping |
| hCoVOC43 | orf1ab | DDDFLEESDVEEDDV | 966 | 980 | Overlapping |
| hCoVOC43 | orf1ab | DELTPSDVYQQLAGI | 3532 | 3546 | Overlapping |
| hCoVOC43 | orf1ab | DESCNNLVPTYLKSD | 2654 | 2668 | Overlapping |
| hCoVOC43 | orf1ab | DEVSMLTNYELSVIN | 5670 | 5684 | Overlapping |
| hCoVOC43 | orf1ab | DGDDVYLPYPNPSRI | 5188 | 5202 | Overlapping |
| hCoVOC43 | orf1ab | DGGKYYTQRIIKAQF | 1896 | 1910 | Overlapping |
| hCoVOC43 | orf1ab | DGLEAYADKTLQEMK | 266 | 280 | Overlapping |
| hCoVOC43 | orf1ab | DGMDAKQVDYVPLKS | 6350 | 6364 | Overlapping |
| hCoVOC43 | orf1ab | DGQKFTDESFYKNMY | 5276 | 5290 | Overlapping |
| hCoVOC43 | orf1ab | DGQRTYDDVNASLFV | 2496 | 2510 | Overlapping |
| hCoVOC43 | orf1ab | DGTFEEMALTTFMIT | 3156 | 3170 | Overlapping |
| hCoVOC43 | orf1ab | DGVNFTNRFVPVGES | 1572 | 1586 | Overlapping |
| hCoVOC43 | orf1ab | DGVVCYNSDYASKGY | 5126 | 5140 | Overlapping |
| hCoVOC43 | orf1ab | DLESVIQDYENVCFE | 1036 | 1050 | Overlapping |
| hCoVOC43 | orf1ab | DTLDDGPSVETSDSQ | 1010 | 1024 | Overlapping |
| hCoVOC43 | orf1ab | DVEMSDFVDLESVIQ | 1028 | 1042 | Overlapping |
| hCoVOC43 | orf1ab | EAACSQLAKAMDTFT | 3210 | 3224 | Overlapping |
| hCoVOC43 | orf1ab | EAALDLSKELKRPIQ | 2456 | 2470 | Overlapping |
| hCoVOC43 | orf1ab | EACYVRGCNPKGWTM | 114 | 128 | Overlapping |
| hCoVOC43 | orf1ab | EADRRAFVDYTGVLK | 2294 | 2308 | Overlapping |
| hCoVOC43 | orf1ab | EAFKQSYASATIQEI | 5440 | 5454 | Overlapping |
| hCoVOC43 | orf1ab | EALSFEEQDEIYAYT | 4882 | 4896 | Overlapping |
| hCoVOC43 | orf1ab | EAWLEFRSGRPARFV | 1694 | 1708 | Overlapping |
| hCoVOC43 | orf1ab | ECTERLKLFAAETQK | 5422 | 5436 | Overlapping |
| hCoVOC43 | orf1ab | EDAEEKLDNPSSSEV | 22 | 36 | Overlapping |
| hCoVOC43 | orf1ab | EDFDLISKCQITAVE | 1422 | 1436 | Overlapping |
| hCoVOC43 | orf1ab | EDFNVWALSNGFSQV | 3474 | 3488 | Overlapping |
| hCoVOC43 | orf1ab | EDHKPQYSFKLVMNG | 5370 | 5384 | Overlapping |
| hCoVOC43 | orf1ab | EDLKAVRSSFNFDQK | 1624 | 1638 | Overlapping |
| hCoVOC43 | orf1ab | EDNPLFLFDEAGEEV | 938 | 952 | Overlapping |
| hCoVOC43 | orf1ab | EDYAFEHIVYGNFNQ | 6678 | 6692 | Overlapping |
| hCoVOC43 | orf1ab | EEADEGICFNFNGSW | 2976 | 2990 | Overlapping |
| hCoVOC43 | orf1ab | EEAVKRVRAWVGFDA | 5976 | 5990 | Overlapping |
| hCoVOC43 | orf1ab | EEDDVEGEETDLTVT | 976 | 990 | Overlapping |
| hCoVOC43 | orf1ab | EEDVEMSDFVDLESV | 1026 | 1040 | Overlapping |
| hCoVOC43 | orf1ab | EEEILLMLASLFGTY | 3708 | 3722 | Overlapping |
| hCoVOC43 | orf1ab | EEKLSPCKELEGVGA | 914 | 928 | Overlapping |
| hCoVOC43 | orf1ab | EEMALTTFMITKDSY | 3160 | 3174 | Overlapping |
| hCoVOC43 | orf1ab | EESSEVLEDTLDDGP | 1002 | 1016 | Overlapping |
| hCoVOC43 | orf1ab | EEVLAPKLYCAFTAP | 950 | 964 | Overlapping |
| hCoVOC43 | orf1ab | EEYQKVFRVYLAYIK | 5240 | 5254 | Overlapping |
| hCoVOC43 | orf1ab | EFADKLVEVGLVGVL | 4560 | 4574 | Overlapping |
| hCoVOC43 | orf1ab | EFCSQHTMLVKMDGD | 5176 | 5190 | Overlapping |

| | | | | | |
|---|---|---|---|---|---|
| hCoVOC43 | orf1ab | EFLNKHFSMMILSDD | 5112 | 5126 | Overlapping |
| hCoVOC43 | orf1ab | EFPWMFEDAEEKLDN | 16 | 30 | Overlapping |
| hCoVOC43 | orf1ab | EFRSGRPARFVALVL | 1698 | 1712 | Overlapping |
| hCoVOC43 | orf1ab | EFVNMASFVEYEVAK | 3930 | 3944 | Overlapping |
| hCoVOC43 | orf1ab | EFYAVVIDAIEEKLS | 904 | 918 | Overlapping |
| hCoVOC43 | orf1ab | EFYTTEPEFVKVLGL | 1050 | 1064 | Overlapping |
| hCoVOC43 | orf1ab | EGICFNFNGSWVLNN | 2980 | 2994 | Overlapping |
| hCoVOC43 | orf1ab | EGLVRIVRTRSMSYC | 2956 | 2970 | Overlapping |
| hCoVOC43 | orf1ab | EGSRVPHIVRKDLTK | 4476 | 4490 | Overlapping |
| hCoVOC43 | orf1ab | EGSSVDLKHFFFTQD | 4796 | 4810 | Overlapping |
| hCoVOC43 | orf1ab | EGTKKLAARLSFNVG | 1436 | 1450 | Overlapping |
| hCoVOC43 | orf1ab | EGVGAKVSAFLQKLE | 924 | 938 | Overlapping |
| hCoVOC43 | orf1ab | EGWRVVNKFYQINGV | 1506 | 1520 | Overlapping |
| hCoVOC43 | orf1ab | EHPDVDGLCKLRGKF | 4314 | 4328 | Overlapping |
| hCoVOC43 | orf1ab | EIACKCGVKQEQRTG | 1746 | 1760 | Overlapping |
| hCoVOC43 | orf1ab | EIAQLYGSCITPNVC | 1264 | 1278 | Overlapping |
| hCoVOC43 | orf1ab | EIDGNVMHANYLFWR | 7014 | 7028 | Overlapping |
| hCoVOC43 | orf1ab | EIGYTVDCSCGKKLI | 1776 | 1790 | Overlapping |
| hCoVOC43 | orf1ab | EILATSDLSVAFEKL | 3874 | 3888 | Overlapping |
| hCoVOC43 | orf1ab | EILLMLASLFGTYTW | 3710 | 3724 | Overlapping |
| hCoVOC43 | orf1ab | EIVMCGGCYYVKPGG | 5030 | 5044 | Overlapping |
| hCoVOC43 | orf1ab | EIYDGGCIPASQVIV | 4846 | 4860 | Overlapping |
| hCoVOC43 | orf1ab | EKASLNSLTYFNRPS | 1990 | 2003 | Overlapping |
| hCoVOC43 | orf1ab | EKDFIALDQDVFIQK | 6660 | 6674 | Overlapping |
| hCoVOC43 | orf1ab | EKGKLLIRDTNKEVF | 7072 | 7086 | Overlapping |
| hCoVOC43 | orf1ab | EKLAQLLIVLFANPA | 3886 | 3900 | Overlapping |
| hCoVOC43 | orf1ab | EKLDNPSSSEVDMIC | 26 | 40 | Overlapping |
| hCoVOC43 | orf1ab | EKVDGVYTNFKLIGH | 1914 | 1928 | Overlapping |
| hCoVOC43 | orf1ab | EKVMTFYPRLQAASD | 6786 | 6800 | Overlapping |
| hCoVOC43 | orf1ab | ELDNDFNTILNTACG | 878 | 892 | Overlapping |
| hCoVOC43 | orf1ab | ELEGVGAKVSAFLQK | 922 | 936 | Overlapping |
| hCoVOC43 | orf1ab | ELFMLSTLHWSFRLL | 2342 | 2356 | Overlapping |
| hCoVOC43 | orf1ab | ELFNKYFKHWSMPYH | 4646 | 4660 | Overlapping |
| hCoVOC43 | orf1ab | ELHWAPEFPWMFEDA | 10 | 24 | Overlapping |
| hCoVOC43 | orf1ab | ELIVSYALYTAWFYP | 2312 | 2326 | Overlapping |
| hCoVOC43 | orf1ab | ELKRPIQPTDVAYHT | 2464 | 2478 | Overlapping |
| hCoVOC43 | orf1ab | ELLDVSLDSLGAAIH | 618 | 632 | Overlapping |
| hCoVOC43 | orf1ab | ELNGVVVDKVGDTDC | 6576 | 6590 | Overlapping |
| hCoVOC43 | orf1ab | ELRYMNANGLRPPKN | 3796 | 3810 | Overlapping |
| hCoVOC43 | orf1ab | ELTCLRYFAKVGREI | 6090 | 6104 | Overlapping |
| hCoVOC43 | orf1ab | EMKALFPTWSQELLF | 278 | 292 | Overlapping |
| hCoVOC43 | orf1ab | EMKCYERVKDCKFVA | 4452 | 4466 | Overlapping |
| hCoVOC43 | orf1ab | EMPCAIINDKVVAKI | 6442 | 6456 | Overlapping |
| hCoVOC43 | orf1ab | EMSDFVDLESVIQDY | 1030 | 1044 | Overlapping |
| hCoVOC43 | orf1ab | ENDADKANFLNAAVF | 2532 | 2546 | Overlapping |
| hCoVOC43 | orf1ab | ENEEYQKVFRVYLAY | 5238 | 5252 | Overlapping |
| hCoVOC43 | orf1ab | ENGDKLDQFFVVKRT | 4430 | 4444 | Overlapping |
| hCoVOC43 | orf1ab | ENKLKAKNESSSLCF | 5754 | 5768 | Overlapping |
| hCoVOC43 | orf1ab | ENPDIINVYKKLGPI | 4536 | 4550 | Overlapping |

| | | | | | |
|---|---|---|---|---|---|
| hCoVOC43 | orf1ab | ENVVYNLVKTGHYTG | 6424 | 6438 | Overlapping |
| hCoVOC43 | orf1ab | EPCVVSVTYGNMTLN | 3260 | 3274 | Overlapping |
| hCoVOC43 | orf1ab | EPKGKFSKKAYALIR | 228 | 242 | Overlapping |
| hCoVOC43 | orf1ab | EPKYFNTVTKLMCCL | 5714 | 5728 | Overlapping |
| hCoVOC43 | orf1ab | EPVHVYADDSIILRQ | 338 | 352 | Overlapping |
| hCoVOC43 | orf1ab | EQDEIYAYTKRNVLP | 4888 | 4902 | Overlapping |
| hCoVOC43 | orf1ab | EQEESSEVLEDTLDD | 1000 | 1014 | Overlapping |
| hCoVOC43 | orf1ab | EQFDQWYESTFGLSY | 2816 | 2830 | Overlapping |
| hCoVOC43 | orf1ab | EQRTGLDAVMHFGTL | 1756 | 1770 | Overlapping |
| hCoVOC43 | orf1ab | EQSYFTKKDWYDFVE | 4522 | 4536 | Overlapping |
| hCoVOC43 | orf1ab | ERDRAVAKKLERMAD | 3976 | 3990 | Overlapping |
| hCoVOC43 | orf1ab | ERFVSLAIDAYPLVY | 5222 | 5236 | Overlapping |
| hCoVOC43 | orf1ab | ERGCITFGKPVIWLS | 1974 | 1988 | Overlapping |
| hCoVOC43 | orf1ab | ERMADLALTNMYKEA | 3986 | 4000 | Overlapping |
| hCoVOC43 | orf1ab | ESDVEEDDVEGEETD | 972 | 986 | Overlapping |
| hCoVOC43 | orf1ab | ESIFCSNTYGVCMYT | 6512 | 6526 | Overlapping |
| hCoVOC43 | orf1ab | ESSSLCFKVYYKGVT | 5762 | 5776 | Overlapping |
| hCoVOC43 | orf1ab | ESVIQDYENVCFEFY | 1038 | 1052 | Overlapping |
| hCoVOC43 | orf1ab | ESYNTATTAGFTFWV | 6390 | 6404 | Overlapping |
| hCoVOC43 | orf1ab | ETCCSQSDRFYRLAN | 5008 | 5022 | Overlapping |
| hCoVOC43 | orf1ab | ETDANKLILINDVAF | 1456 | 1470 | Overlapping |
| hCoVOC43 | orf1ab | ETDGICPENHVMVDC | 48 | 62 | Overlapping |
| hCoVOC43 | orf1ab | ETDLTVTSAGQPCVA | 984 | 998 | Overlapping |
| hCoVOC43 | orf1ab | ETFQNNVVNYQHIGM | 5556 | 5570 | Overlapping |
| hCoVOC43 | orf1ab | ETFTVCADGFMPFLL | 570 | 584 | Overlapping |
| hCoVOC43 | orf1ab | ETFTVLAAYNGKPQG | 3356 | 3370 | Overlapping |
| hCoVOC43 | orf1ab | ETKEINIIKLSGVKK | 2032 | 2046 | Overlapping |
| hCoVOC43 | orf1ab | ETKVQCSTNLFKDCS | 5896 | 5910 | Overlapping |
| hCoVOC43 | orf1ab | ETKYVKSLSIVDVYD | 2062 | 2076 | Overlapping |
| hCoVOC43 | orf1ab | ETLLAAIKRLKNGFQ | 3506 | 3520 | Overlapping |
| hCoVOC43 | orf1ab | ETMFDVYVDTFLSMF | 2574 | 2588 | Overlapping |
| hCoVOC43 | orf1ab | EVCDDYAKDNTVLQA | 3912 | 3926 | Overlapping |
| hCoVOC43 | orf1ab | EVFVGDSLVNVIGSG | 7084 | 7095 | Overlapping |
| hCoVOC43 | orf1ab | EVLEDTLDDGPSVET | 1006 | 1020 | Overlapping |
| hCoVOC43 | orf1ab | EVLLQDALQSREAVL | 90 | 104 | Overlapping |
| hCoVOC43 | orf1ab | EVLREGLVRIVRTRS | 2952 | 2966 | Overlapping |
| hCoVOC43 | orf1ab | EVVTTSLTPCGYSEP | 794 | 808 | Overlapping |
| hCoVOC43 | orf1ab | EVVVNPANGHMAHGG | 1294 | 1308 | Overlapping |
| hCoVOC43 | orf1ab | EYASNSSILSLCAFS | 4238 | 4252 | Overlapping |
| hCoVOC43 | orf1ab | EYDYVIYSQTAETAH | 5836 | 5850 | Overlapping |
| hCoVOC43 | orf1ab | EYKITEWPTATGDVV | 1948 | 1962 | Overlapping |
| hCoVOC43 | orf1ab | EYPIISNELSINTSC | 6194 | 6208 | Overlapping |
| hCoVOC43 | orf1ab | EYREYLESYNTATTA | 6384 | 6398 | Overlapping |
| hCoVOC43 | orf1ab | EYVFDKSELTNGVYY | 5494 | 5508 | Overlapping |
| hCoVOC43 | orf1ab | FDSSKEFVEYKITEW | 1940 | 1954 | Overlapping |
| hCoVOC43 | orf1ab | FEDELTPSDVYQQLA | 3530 | 3544 | Overlapping |
| hCoVOC43 | orf1ab | FEEQDEIYAYTKRNV | 4886 | 4900 | Overlapping |
| hCoVOC43 | orf1ab | FGKARLYYEALSFEE | 4874 | 4888 | Overlapping |
| hCoVOC43 | orf1ab | FVDDLLKTDSVLLIE | 5208 | 5222 | Overlapping |

| | | | | | |
|---|---|---|---|---|---|
| hCoVOC43 | orf1ab | FVPVGESFGKSLGNV | 1580 | 1594 | Overlapping |
| hCoVOC43 | orf1ab | GEETDLTVTSAGQPC | 982 | 996 | Overlapping |
| hCoVOC43 | orf1ab | GEKGAYNKDHGRGGF | 196 | 210 | Overlapping |
| hCoVOC43 | orf1ab | GEQFKHLIPLMTRGH | 6042 | 6056 | Overlapping |
| hCoVOC43 | orf1ab | GGDSSESGAKETKEI | 2022 | 2036 | Overlapping |
| hCoVOC43 | orf1ab | GGMIRYYDVMANGGT | 2416 | 2430 | Overlapping |
| hCoVOC43 | orf1ab | GIKDPVSYVLTHDVC | 4334 | 4348 | Overlapping |
| hCoVOC43 | orf1ab | GKPQGAFHVTMRSSY | 3366 | 3380 | Overlapping |
| hCoVOC43 | orf1ab | GLEDYAFEHIVYGNF | 6676 | 6690 | Overlapping |
| hCoVOC43 | orf1ab | GLELHWAPEFPWMFE | 8 | 22 | Overlapping |
| hCoVOC43 | orf1ab | GMSYYCEDHKPQYSF | 5364 | 5378 | Overlapping |
| hCoVOC43 | orf1ab | GMTLVSIPIDLLNLR | 2108 | 2122 | Overlapping |
| hCoVOC43 | orf1ab | GPDARTQGKQSYVLL | 1358 | 1372 | Overlapping |
| hCoVOC43 | orf1ab | GPLVRQIFVDGVPFV | 4692 | 4706 | Overlapping |
| hCoVOC43 | orf1ab | GPPGTGKSHLAIGLA | 5578 | 5592 | Overlapping |
| hCoVOC43 | orf1ab | GSCSFEDELTPSDVY | 3526 | 3540 | Overlapping |
| hCoVOC43 | orf1ab | GTNKQLNEISDDCNW | 4090 | 4104 | Overlapping |
| hCoVOC43 | orf1ab | GTSVRSDGTFEEMAL | 3150 | 3164 | Overlapping |
| hCoVOC43 | orf1ab | GVFEVDDTVDMEEFY | 892 | 906 | Overlapping |
| hCoVOC43 | orf1ab | GVGSANIFIGDKVGH | 1814 | 1828 | Overlapping |
| hCoVOC43 | orf1ab | GVPIIEVSQFQSKLT | 3826 | 3840 | Overlapping |
| hCoVOC43 | orf1ab | GVTTHESSSAVNMQQ | 5774 | 5788 | Overlapping |
| hCoVOC43 | orf1ab | HNEILATSDLSVAFE | 3872 | 3886 | Overlapping |
| hCoVOC43 | orf1ab | HWAPEFPWMFEDAEE | 12 | 26 | Overlapping |
| hCoVOC43 | orf1ab | ICKCGESMVLIDVDV | 1176 | 1190 | Overlapping |
| hCoVOC43 | orf1ab | IEEVCDDYAKDNTVL | 3910 | 3924 | Overlapping |
| hCoVOC43 | orf1ab | KEGQDVIFSQFDSLG | 6598 | 6612 | Overlapping |
| hCoVOC43 | orf1ab | KFEQFDQWYESTFGL | 2814 | 2828 | Overlapping |
| hCoVOC43 | orf1ab | KHETCCSQSDRFYRL | 5006 | 5020 | Overlapping |
| hCoVOC43 | orf1ab | KLEDNPLFLFDEAGE | 936 | 950 | Overlapping |
| hCoVOC43 | orf1ab | KLGTSVRSDGTFEEM | 3148 | 3162 | Overlapping |
| hCoVOC43 | orf1ab | KNESSSLCFKVYYKG | 5760 | 5774 | Overlapping |
| hCoVOC43 | orf1ab | KVEIDGNVMHANYLF | 7012 | 7026 | Overlapping |
| hCoVOC43 | orf1ab | KVPQAVETKVQCSTN | 5890 | 5904 | Overlapping |
| hCoVOC43 | orf1ab | LCEEADEGICFNFNG | 2974 | 2988 | Overlapping |
| hCoVOC43 | orf1ab | LDKVDILLTVDGVNF | 1562 | 1576 | Overlapping |
| hCoVOC43 | orf1ab | LEESDVEEDDVEGEE | 970 | 984 | Overlapping |
| hCoVOC43 | orf1ab | LIERFVSLAIDAYPL | 5220 | 5234 | Overlapping |
| hCoVOC43 | orf1ab | LIGINYLCKPKVEID | 7002 | 7016 | Overlapping |
| hCoVOC43 | orf1ab | LKVDDVDDGGDSSES | 2014 | 2028 | Overlapping |
| hCoVOC43 | orf1ab | LSFEEQDEIYAYTKR | 4884 | 4898 | Overlapping |
| hCoVOC43 | orf1ab | MEEFYAVVIDAIEEK | 902 | 916 | Overlapping |
| hCoVOC43 | orf1ab | MFEDAEEKLDNPSSS | 20 | 34 | Overlapping |
| hCoVOC43 | orf1ab | MVTDIVVVDEVSMLT | 5662 | 5676 | Overlapping |
| hCoVOC43 | orf1ab | NANASSSEGFLIGIN | 6992 | 7006 | Overlapping |
| hCoVOC43 | orf1ab | NFELTCLRYFAKVGR | 6088 | 6102 | Overlapping |
| hCoVOC43 | orf1ab | NGVVVDKVGDTDCVF | 6578 | 6592 | Overlapping |
| hCoVOC43 | orf1ab | NIGEYNVSKDGFFTY | 6934 | 6948 | Overlapping |
| hCoVOC43 | orf1ab | NLEEEILLMLASLFG | 3706 | 3720 | Overlapping |

| | | | | | |
|---|---|---|---|---|---|
| hCoVOC43 | orf1ab | NNYDCVVTTLISAGI | 1380 | 1394 | Overlapping |
| hCoVOC43 | orf1ab | NQEDFDLISKCQITA | 1420 | 1434 | Overlapping |
| hCoVOC43 | orf1ab | NSFEALMLNFKLLGI | 3810 | 3824 | Overlapping |
| hCoVOC43 | orf1ab | PEDDDFLEESDVEED | 964 | 978 | Overlapping |
| hCoVOC43 | orf1ab | PEMVTDIVVVDEVSM | 5660 | 5674 | Overlapping |
| hCoVOC43 | orf1ab | PIIEVSQFQSKLTDV | 3828 | 3842 | Overlapping |
| hCoVOC43 | orf1ab | PIISNELSINTSCRV | 6196 | 6210 | Overlapping |
| hCoVOC43 | orf1ab | PILMVDKNLITTANT | 2554 | 2568 | Overlapping |
| hCoVOC43 | orf1ab | PIQPTDVAYHTVTDV | 2468 | 2482 | Overlapping |
| hCoVOC43 | orf1ab | PIVVAVIDQDFGSTV | 2838 | 2852 | Overlapping |
| hCoVOC43 | orf1ab | PTASVSTSFLQSGIV | 3236 | 3250 | Overlapping |
| hCoVOC43 | orf1ab | PTMVDIKQLLFVLEV | 4826 | 4840 | Overlapping |
| hCoVOC43 | orf1ab | PTYTVHKSDFQLPVY | 2776 | 2790 | Overlapping |
| hCoVOC43 | orf1ab | PVGESFGKSLGNVFC | 1582 | 1596 | Overlapping |
| hCoVOC43 | orf1ab | PVGIKDPVSYVLTHD | 4332 | 4346 | Overlapping |
| hCoVOC43 | orf1ab | PVNPVLHTKSAAGYG | 434 | 448 | Overlapping |
| hCoVOC43 | orf1ab | PVVVDDHVGLLDQAW | 832 | 846 | Overlapping |
| hCoVOC43 | orf1ab | QEIVSERELILSWEI | 5452 | 5466 | Overlapping |
| hCoVOC43 | orf1ab | QGRQIMGSCSFEDEL | 3520 | 3534 | Overlapping |
| hCoVOC43 | orf1ab | QSEFVNMASFVEYEV | 3928 | 3942 | Overlapping |
| hCoVOC43 | orf1ab | QVEEDVEMSDFVDLE | 1024 | 1038 | Overlapping |
| hCoVOC43 | orf1ab | QVHDEPKGKFSKKAY | 224 | 238 | Overlapping |
| hCoVOC43 | orf1ab | QYEADRRAFVDYTGV | 2292 | 2306 | Overlapping |
| hCoVOC43 | orf1ab | RAELNGVVVDKVGDT | 6574 | 6588 | Overlapping |
| hCoVOC43 | orf1ab | RAFDIYNASVAGIGL | 4402 | 4416 | Overlapping |
| hCoVOC43 | orf1ab | RGPPRAELNGVVVDK | 6570 | 6584 | Overlapping |
| hCoVOC43 | orf1ab | RTGLDAVMHFGTLSR | 1758 | 1772 | Overlapping |
| hCoVOC43 | orf1ab | RVEHPDVDGLCKLRG | 4312 | 4326 | Overlapping |
| hCoVOC43 | orf1ab | SAKHVQGNVAKIAGV | 2682 | 2696 | Overlapping |
| hCoVOC43 | orf1ab | SIDSDVDTKCLADSV | 2628 | 2642 | Overlapping |
| hCoVOC43 | orf1ab | SKELKRPIQPTDVAY | 2462 | 2476 | Overlapping |
| hCoVOC43 | orf1ab | SMRLFYDRDGQRTYD | 2488 | 2502 | Overlapping |
| hCoVOC43 | orf1ab | SMSYCRVGLCEEADE | 2966 | 2980 | Overlapping |
| hCoVOC43 | orf1ab | SSKEFVEYKITEWPT | 1942 | 1956 | Overlapping |
| hCoVOC43 | orf1ab | SSVDLKHFFFTQDGN | 4798 | 4812 | Overlapping |
| hCoVOC43 | orf1ab | SVEDFNVWALSNGFS | 3472 | 3486 | Overlapping |
| hCoVOC43 | orf1ab | SVFSLWYKGSNLEEE | 3696 | 3710 | Overlapping |
| hCoVOC43 | orf1ab | SVPNMHVVVVENDAD | 2522 | 2536 | Overlapping |
| hCoVOC43 | orf1ab | TAPEDDDFLEESDVE | 962 | 976 | Overlapping |
| hCoVOC43 | orf1ab | TDYNYYKYNLPTMVD | 4816 | 4830 | Overlapping |
| hCoVOC43 | orf1ab | TEEAFKQSYASATIQ | 5438 | 5452 | Overlapping |
| hCoVOC43 | orf1ab | TEGLMQNASLYSSLV | 2920 | 2934 | Overlapping |
| hCoVOC43 | orf1ab | TEPEFVKVLGLYVPK | 1054 | 1068 | Overlapping |
| hCoVOC43 | orf1ab | TFEEMALTTFMITKD | 3158 | 3172 | Overlapping |
| hCoVOC43 | orf1ab | TGCMMNVAKYTQLCQ | 6834 | 6848 | Overlapping |
| hCoVOC43 | orf1ab | TGDCYVSTGGKLCKT | 1338 | 1352 | Overlapping |
| hCoVOC43 | orf1ab | TGDLAVCLGIGDSAV | 5934 | 5948 | Overlapping |
| hCoVOC43 | orf1ab | TGDVVLATDDLYVKR | 1958 | 1972 | Overlapping |
| hCoVOC43 | orf1ab | TGIDFVVEATGLFAD | 6012 | 6026 | Overlapping |

| | | | | | |
|---|---|---|---|---|---|
| hCoVOC43 | orf1ab | TGTSVTETMFDVYVD | 2568 | 2582 | Overlapping |
| hCoVOC43 | orf1ab | VDDVDDGGDSSESGA | 2016 | 2030 | Overlapping |
| hCoVOC43 | orf1ab | VDENGDKLDQFFVVK | 4428 | 4442 | Overlapping |
| hCoVOC43 | orf1ab | VDIMSHFYMEADTVV | 357 | 370 | Overlapping |
| hCoVOC43 | orf1ab | VEDAYDQVHDEPKGK | 218 | 232 | Overlapping |
| hCoVOC43 | orf1ab | VGLCEEADEGICFNF | 2972 | 2986 | Overlapping |
| hCoVOC43 | orf1ab | VIRDVSVEDVCFANK | 2800 | 2814 | Overlapping |
| hCoVOC43 | orf1ab | VLETFQNNVVNYQHI | 5554 | 5568 | Overlapping |
| hCoVOC43 | orf1ab | VMQFGYIDCEQDSCD | 384 | 398 | Overlapping |
| hCoVOC43 | orf1ab | VRTVKYFECTGGIDI | 1520 | 1534 | Overlapping |
| hCoVOC43 | orf1ab | VVNPANGHMAHGGGV | 1296 | 1310 | Overlapping |
| hCoVOC43 | orf1ab | VVQEFVSYDSSIHSY | 6714 | 6728 | Overlapping |
| hCoVOC43 | orf1ab | YDFKVEDAYDQVHDE | 214 | 228 | Overlapping |
| hCoVOC43 | orf1ab | YLESYNTATTAGFTF | 6388 | 6402 | Overlapping |
| hCoVOC43 | orf1ab | YTPHSQISYSNFYAS | 2882 | 2896 | Overlapping |
| hCoVOC43 | orf1ab | YYEALSFEEQDEIYA | 4880 | 4894 | Overlapping |
| hCoVOC43 | spike | AGVLTNHDVVYAQHC | 469 | 483 | Positive |
| hCoVOC43 | spike | AIEDLLFDKVKLSDV | 905 | 919 | Positive |
| hCoVOC43 | spike | AITTGYRFTNFEPFT | 759 | 773 | Positive |
| hCoVOC43 | spike | ANFILHDVNSGLTCS | 592 | 606 | Positive |
| hCoVOC43 | spike | CEYPQTICHPNLGNH | 169 | 183 | Positive |
| hCoVOC43 | spike | CGTSCFKKCGGCCDD | 1324 | 1338 | Positive |
| hCoVOC43 | spike | CHPNLGNHRKELWHL | 176 | 190 | Positive |
| hCoVOC43 | spike | DALEAEAQIDRLING | 1069 | 1083 | Positive |
| hCoVOC43 | spike | DFKEELDQWFKNQTS | 1232 | 1246 | Positive |
| hCoVOC43 | spike | DGIIFNAVDCMSDFM | 281 | 295 | Positive |
| hCoVOC43 | spike | DSLEPVGGLYEIQIP | 779 | 793 | Positive |
| hCoVOC43 | spike | EIKCKTQSIAPPTGV | 297 | 311 | Positive |
| hCoVOC43 | spike | EPFTVNSVNDSLEPV | 770 | 784 | Positive |
| hCoVOC43 | spike | EPITENNVVVMSTCA | 1198 | 1212 | Positive |
| hCoVOC43 | spike | FDATNSALVKIQAVV | 1022 | 1036 | Positive |
| hCoVOC43 | spike | FIEDSVFKPRPAGVL | 458 | 472 | Positive |
| hCoVOC43 | spike | FKPRPAGVLTNHDVV | 464 | 478 | Positive |
| hCoVOC43 | spike | FNPSTWNKRFGFIED | 447 | 461 | Positive |
| hCoVOC43 | spike | FNQDGIIFNAVDCMS | 278 | 292 | Positive |
| hCoVOC43 | spike | GAITTGYRFTNFEPF | 758 | 772 | Positive |
| hCoVOC43 | spike | GCCDDYTGYQELVIK | 1334 | 1348 | Positive |
| hCoVOC43 | spike | GDKCNIFANFILHDV | 585 | 599 | Positive |
| hCoVOC43 | spike | GIAPKSGYFVNVNNT | 1172 | 1186 | Positive |
| hCoVOC43 | spike | GPGKNNGIGTCPAGT | 503 | 517 | Positive |
| hCoVOC43 | spike | GPPPISTDTVDVTNG | 32 | 46 | Positive |
| hCoVOC43 | spike | GSGMFLILLISLPTA | 1 | 12 | Positive |
| hCoVOC43 | spike | GSGSGSGMFLILLIS | 1 | 8 | Positive |
| hCoVOC43 | spike | GSGYYYPEPITENNV | 1191 | 1205 | Positive |
| hCoVOC43 | spike | GYQELVIKTSHDDGS | 1341 | 1353 | Positive |
| hCoVOC43 | spike | GYRFTNFEPFTVNSV | 763 | 777 | Positive |
| hCoVOC43 | spike | GYYYPEPITENNVVV | 1193 | 1207 | Positive |
| hCoVOC43 | spike | ICHPNLGNHRKELWH | 175 | 189 | Positive |
| hCoVOC43 | spike | IDCAAFVCGDYAACK | 814 | 828 | Positive |

| | | | | | |
|---|---|---|---|---|---|
| hCoVOC43 | spike | IDTTATSCQLYYNLP | 424 | 438 | Positive |
| hCoVOC43 | spike | IGEHCSGLAVKSDYC | 548 | 562 | Positive |
| hCoVOC43 | spike | IPNLPDFKEELDQWF | 1227 | 1241 | Positive |
| hCoVOC43 | spike | IPSEFTIGNMVEFIQ | 792 | 806 | Positive |
| hCoVOC43 | spike | IQIPSEFTIGNMVEF | 790 | 804 | Positive |
| hCoVOC43 | spike | ITTGYRFTNFEPFTV | 760 | 774 | Positive |
| hCoVOC43 | spike | KCNIFANFILHDVNS | 587 | 601 | Positive |
| hCoVOC43 | spike | KDTGPPPISTDTVDV | 29 | 43 | Positive |
| hCoVOC43 | spike | KEELDQWFKNQTSVA | 1234 | 1248 | Positive |
| hCoVOC43 | spike | KLKDGVNFNVDDINF | 872 | 886 | Positive |
| hCoVOC43 | spike | KPRPAGVLTNHDVVY | 465 | 479 | Positive |
| hCoVOC43 | spike | KRFGFIEDSVFKPRP | 454 | 468 | Positive |
| hCoVOC43 | spike | KVKLSDVGFVEAYNN | 913 | 927 | Positive |
| hCoVOC43 | spike | LEVSVCQYNMCEYPQ | 159 | 173 | Positive |
| hCoVOC43 | spike | LEYWVTPLTSRQYLL | 262 | 276 | Positive |
| hCoVOC43 | spike | LHDVNSGLTCSTDLQ | 596 | 610 | Positive |
| hCoVOC43 | spike | LLAFNQDGIIFNAVD | 275 | 289 | Positive |
| hCoVOC43 | spike | LVIKTSHDDGSGSGS | 1345 | 1353 | Positive |
| hCoVOC43 | spike | LVKFSAAQAMEKVNE | 1101 | 1115 | Positive |
| hCoVOC43 | spike | MLNTSIPNLPDFKEE | 1222 | 1236 | Positive |
| hCoVOC43 | spike | MYTGSGYYYPEPITE | 1188 | 1202 | Positive |
| hCoVOC43 | spike | NADYLYFHFYQEGGT | 208 | 222 | Positive |
| hCoVOC43 | spike | NAEALNNLLQQLSNR | 1039 | 1053 | Positive |
| hCoVOC43 | spike | NAFNNALYAIQEGFD | 1009 | 1023 | Positive |
| hCoVOC43 | spike | NAILTEVNELLDTTQ | 842 | 856 | Positive |
| hCoVOC43 | spike | NALYAIQEGFDATNS | 1013 | 1027 | Positive |
| hCoVOC43 | spike | NANAEALNNLLQQLS | 1037 | 1051 | Positive |
| hCoVOC43 | spike | NATYYNSWQNLLYDS | 639 | 653 | Positive |
| hCoVOC43 | spike | NAVDCMSDFMSEIKC | 286 | 300 | Positive |
| hCoVOC43 | spike | NAYVSQQLSDSTLVK | 1089 | 1103 | Positive |
| hCoVOC43 | spike | NDSLEPVGGLYEIQI | 778 | 792 | Positive |
| hCoVOC43 | spike | NFEPFTVNSVNDSLE | 768 | 782 | Positive |
| hCoVOC43 | spike | NFILHDVNSGLTCST | 593 | 607 | Positive |
| hCoVOC43 | spike | NFNMSSLMSFIQADS | 361 | 375 | Positive |
| hCoVOC43 | spike | NFNVDDINFSPVLGC | 878 | 892 | Positive |
| hCoVOC43 | spike | NFTYDVNADYLYFHF | 202 | 216 | Positive |
| hCoVOC43 | spike | NGNLYGFRDYITNRT | 654 | 668 | Positive |
| hCoVOC43 | spike | NGRKVDLQLGNLGYL | 403 | 417 | Positive |
| hCoVOC43 | spike | NHDVVYAQHCFKAPK | 474 | 488 | Positive |
| hCoVOC43 | spike | NHIISLVQNAPYGLY | 1131 | 1145 | Positive |
| hCoVOC43 | spike | NIEAWLNDKSVPSPL | 336 | 350 | Positive |
| hCoVOC43 | spike | NINAILTEVNELLDT | 840 | 854 | Positive |
| hCoVOC43 | spike | NINDKDTGPPPISTD | 25 | 39 | Positive |
| hCoVOC43 | spike | NKLQGLLEVSVCQYN | 153 | 167 | Positive |
| hCoVOC43 | spike | NKRFGFIEDSVFKPR | 453 | 467 | Positive |
| hCoVOC43 | spike | NLCTPDPITFTGTYK | 524 | 538 | Positive |
| hCoVOC43 | spike | NLGNHRKELWHLDTG | 179 | 193 | Positive |
| hCoVOC43 | spike | NLGYLQSFNYRIDTT | 413 | 427 | Positive |
| hCoVOC43 | spike | NLKDIGTYEYYVKWP | 1283 | 1297 | Positive |

| | | | | | |
|---|---|---|---|---|---|
| hCoVOC43 | spike | NLLQQLSNRFGAISA | 1045 | 1059 | Positive |
| hCoVOC43 | spike | NLLYDSNGNLYGFRD | 648 | 662 | Positive |
| hCoVOC43 | spike | NLPAANVSVSRFNPS | 436 | 450 | Positive |
| hCoVOC43 | spike | NLPDFKEELDQWFKN | 1229 | 1243 | Positive |
| hCoVOC43 | spike | NLPNCNIEAWLNDKS | 331 | 345 | Positive |
| hCoVOC43 | spike | NLYGFRDYITNRTFM | 656 | 670 | Positive |
| hCoVOC43 | spike | NMALKGSVLLSRLWF | 77 | 91 | Positive |
| hCoVOC43 | spike | NMCEYPQTICHPNLG | 167 | 181 | Positive |
| hCoVOC43 | spike | NMSSLMSFIQADSFT | 363 | 377 | Positive |
| hCoVOC43 | spike | NMVEFIQTSSPKVTI | 800 | 814 | Positive |
| hCoVOC43 | spike | NNALYAIQEGFDATN | 1012 | 1026 | Positive |
| hCoVOC43 | spike | NNCTGGAEIRDLICV | 926 | 940 | Positive |
| hCoVOC43 | spike | NNGIGTCPAGTNYLT | 507 | 521 | Positive |
| hCoVOC43 | spike | NNIDAAKIYGMCFSS | 379 | 393 | Positive |
| hCoVOC43 | spike | NNLLQQLSNRFGAIS | 1044 | 1058 | Positive |
| hCoVOC43 | spike | NNSLTRQLQPINYFD | 704 | 718 | Positive |
| hCoVOC43 | spike | NNTWMYTGSGYYYPE | 1184 | 1198 | Positive |
| hCoVOC43 | spike | NNVVVMSTCAVNYTK | 1203 | 1217 | Positive |
| hCoVOC43 | spike | NPSTWNKRFGFIEDS | 448 | 462 | Positive |
| hCoVOC43 | spike | NQKLIANAFNNALYA | 1003 | 1017 | Positive |
| hCoVOC43 | spike | NQSYINLKDIGTYEY | 1278 | 1292 | Positive |
| hCoVOC43 | spike | NQTSVAPDLSLDYIN | 1243 | 1257 | Positive |
| hCoVOC43 | spike | NRFGAISASLQEILS | 1052 | 1066 | Positive |
| hCoVOC43 | spike | NRLQEAIKVLNQSYI | 1268 | 1282 | Positive |
| hCoVOC43 | spike | NRRSRGAITTGYRFT | 753 | 767 | Positive |
| hCoVOC43 | spike | NRTFMIRSCYSGRVS | 666 | 680 | Positive |
| hCoVOC43 | spike | NSALVKIQAVVNANA | 1026 | 1040 | Positive |
| hCoVOC43 | spike | NSKLTLEYWVTPLTS | 257 | 271 | Positive |
| hCoVOC43 | spike | NSLMNGVTLSTKLKD | 861 | 875 | Positive |
| hCoVOC43 | spike | NSLTRQLQPINYFDS | 705 | 719 | Positive |
| hCoVOC43 | spike | NSSEPALLFRNIKCN | 686 | 700 | Positive |
| hCoVOC43 | spike | NSTAISVQTCDLTVG | 729 | 743 | Positive |
| hCoVOC43 | spike | NSTQDGDNKLQGLLE | 146 | 160 | Positive |
| hCoVOC43 | spike | NSVNDSLEPVGGLYE | 775 | 789 | Positive |
| hCoVOC43 | spike | NVDDINFSPVLGCLG | 880 | 894 | Positive |
| hCoVOC43 | spike | NVSVSRFNPSTWNKR | 441 | 455 | Positive |
| hCoVOC43 | spike | NVYLGMALSHYYVMP | 239 | 253 | Positive |
| hCoVOC43 | spike | NWERKTFSNCNFNMS | 351 | 365 | Positive |
| hCoVOC43 | spike | NYLTCDNLCTPDPIT | 518 | 532 | Positive |
| hCoVOC43 | spike | NYRIDTTATSCQLYY | 421 | 435 | Positive |
| hCoVOC43 | spike | NYVFNNSLTRQLQPI | 700 | 714 | Positive |
| hCoVOC43 | spike | PAGVLTNHDVVYAQH | 468 | 482 | Positive |
| hCoVOC43 | spike | PEPITENNVVVMSTC | 1197 | 1211 | Positive |
| hCoVOC43 | spike | PITENNVVVMSTCAV | 1199 | 1213 | Positive |
| hCoVOC43 | spike | PLLSENQISGYTLAA | 951 | 965 | Positive |
| hCoVOC43 | spike | PNLPDFKEELDQWFK | 1228 | 1242 | Positive |
| hCoVOC43 | spike | PRPAGVLTNHDVVYA | 466 | 480 | Positive |
| hCoVOC43 | spike | PSEFTIGNMVEFIQT | 793 | 807 | Positive |
| hCoVOC43 | spike | PSTWNKRFGFIEDSV | 449 | 463 | Positive |

| | | | | | |
|---|---|---|---|---|---|
| hCoVOC43 | spike | PTSGSTYRNMALKGS | 69 | 83 | Positive |
| hCoVOC43 | spike | PVLGCLGSECSKASS | 888 | 902 | Positive |
| hCoVOC43 | spike | QEAIKVLNQSYINLK | 1271 | 1285 | Positive |
| hCoVOC43 | spike | QEGGTFYAYFTDTGV | 218 | 232 | Positive |
| hCoVOC43 | spike | QELVIKTSHDDGSGS | 1343 | 1353 | Positive |
| hCoVOC43 | spike | QIPSEFTIGNMVEFI | 791 | 805 | Positive |
| hCoVOC43 | spike | QNQKLIANAFNNALY | 1002 | 1016 | Positive |
| hCoVOC43 | spike | QSIAPPTGVYELNGY | 303 | 317 | Positive |
| hCoVOC43 | spike | RDYITNRTFMIRSCY | 661 | 675 | Positive |
| hCoVOC43 | spike | RFGFIEDSVFKPRPA | 455 | 469 | Positive |
| hCoVOC43 | spike | RFTNFEPFTVNSVND | 765 | 779 | Positive |
| hCoVOC43 | spike | SDFINGIFAKVKNTK | 97 | 111 | Positive |
| hCoVOC43 | spike | SDNINDKDTGPPPIS | 23 | 37 | Positive |
| hCoVOC43 | spike | SDSTLVKFSAAQAME | 1097 | 1111 | Positive |
| hCoVOC43 | spike | SDVGFVEAYNNCTGG | 917 | 931 | Positive |
| hCoVOC43 | spike | SDYCGGNSCTCRPQA | 559 | 573 | Positive |
| hCoVOC43 | spike | SECSKASSRSAIEDL | 895 | 909 | Positive |
| hCoVOC43 | spike | SEFTIGNMVEFIQTS | 794 | 808 | Positive |
| hCoVOC43 | spike | SEIKCKTQSIAPPTG | 296 | 310 | Positive |
| hCoVOC43 | spike | SENQISGYTLAATSA | 954 | 968 | Positive |
| hCoVOC43 | spike | SEPALLFRNIKCNYV | 688 | 702 | Positive |
| hCoVOC43 | spike | SGPGKNNGIGTCPAG | 502 | 516 | Positive |
| hCoVOC43 | spike | SGYYYPEPITENNVV | 1192 | 1206 | Positive |
| hCoVOC43 | spike | SIPNLPDFKEELDQW | 1226 | 1240 | Positive |
| hCoVOC43 | spike | SLDYINVTFLDLQVE | 1252 | 1266 | Positive |
| hCoVOC43 | spike | SLMSFIQADSFTCNN | 366 | 380 | Positive |
| hCoVOC43 | spike | SQLVEYGSFCDNINA | 829 | 843 | Positive |
| hCoVOC43 | spike | SRLDALEAEAQIDRL | 1066 | 1080 | Positive |
| hCoVOC43 | spike | STWNKRFGFIEDSVF | 450 | 464 | Positive |
| hCoVOC43 | spike | SVFKPRPAGVLTNHD | 462 | 476 | Positive |
| hCoVOC43 | spike | SYLGCVVNAYNSTAI | 719 | 733 | Positive |
| hCoVOC43 | spike | TDIILGVCVNYDLYG | 614 | 628 | Positive |
| hCoVOC43 | spike | TDLQKANTDIILGVC | 607 | 621 | Positive |
| hCoVOC43 | spike | TDTGVVTKFLFNVYL | 228 | 242 | Positive |
| hCoVOC43 | spike | TDTVDVTNGLGTYYV | 38 | 52 | Positive |
| hCoVOC43 | spike | TENNVVVMSTCAVNY | 1201 | 1215 | Positive |
| hCoVOC43 | spike | TGGAEIRDLICVQSY | 929 | 943 | Positive |
| hCoVOC43 | spike | TGPPPISTDTVDVTN | 31 | 45 | Positive |
| hCoVOC43 | spike | TGSGYYYPEPITENN | 1190 | 1204 | Positive |
| hCoVOC43 | spike | TGYQELVIKTSHDDG | 1340 | 1353 | Positive |
| hCoVOC43 | spike | TGYRFTNFEPFTVNS | 762 | 776 | Positive |
| hCoVOC43 | spike | TIDCAAFVCGDYAAC | 813 | 827 | Positive |
| hCoVOC43 | spike | TIGNMVEFIQTSSPK | 797 | 811 | Positive |
| hCoVOC43 | spike | TMDVLSQNQKLIANA | 996 | 1010 | Positive |
| hCoVOC43 | spike | TNHDVVYAQHCFKAP | 473 | 487 | Positive |
| hCoVOC43 | spike | TPDPITFTGTYKCPQ | 527 | 541 | Positive |
| hCoVOC43 | spike | TSDNINDKDTGPPPI | 22 | 36 | Positive |
| hCoVOC43 | spike | TSGSTYRNMALKGSV | 70 | 84 | Positive |
| hCoVOC43 | spike | TTGYRFTNFEPFTVN | 761 | 775 | Positive |

| | | | | | |
|---|---|---|---|---|---|
| hCoVOC43 | spike | TVDVTNGLGTYYVLD | 40 | 54 | Positive |
| hCoVOC43 | spike | VDLQLGNLGYLQSFN | 407 | 421 | Positive |
| hCoVOC43 | spike | VDVTNGLGTYYVLDR | 41 | 55 | Positive |
| hCoVOC43 | spike | VFKPRPAGVLTNHDV | 463 | 477 | Positive |
| hCoVOC43 | spike | VNSVNDSLEPVGGLY | 774 | 788 | Positive |
| hCoVOC43 | spike | VVMSTCAVNYTKAPY | 1206 | 1220 | Positive |
| hCoVOC43 | spike | VVVMSTCAVNYTKAP | 1205 | 1219 | Positive |
| hCoVOC43 | spike | WNKRFGFIEDSVFKP | 452 | 466 | Positive |
| hCoVOC43 | spike | YAIQEGFDATNSALV | 1016 | 1030 | Positive |
| hCoVOC43 | spike | YDLYGILGQGIFVEV | 624 | 638 | Positive |
| hCoVOC43 | spike | YEYYVKWPWYVWLLI | 1290 | 1304 | Positive |
| hCoVOC43 | spike | YFHFYQEGGTFYAYF | 213 | 227 | Positive |
| hCoVOC43 | spike | YPEPITENNVVVMST | 1196 | 1210 | Positive |
| hCoVOC43 | spike | YQELVIKTSHDDGSG | 1342 | 1353 | Positive |
| hCoVOC43 | spike | YRFTNFEPFTVNSVN | 764 | 778 | Positive |
| hCoVOC43 | spike | YSGRVSAAFHANSSE | 675 | 689 | Positive |
| hCoVOC43 | spike | YTGYQELVIKTSHDD | 1339 | 1353 | Positive |
| hCoVOC43 | spike | YYPEPITENNVVVMS | 1195 | 1209 | Positive |
| hCoVOC43 | spike | YYYPEPITENNVVVM | 1194 | 1208 | Positive |
| hCoVOC43 | spike | AAFHANSSEPALLFR | 681 | 695 | Negative |
| hCoVOC43 | spike | ADSCLQGDKCNIFAN | 579 | 593 | Negative |
| hCoVOC43 | spike | ADSFTCNNIDAAKIY | 373 | 387 | Negative |
| hCoVOC43 | spike | AFAVIGDLKCTSDNI | 12 | 26 | Negative |
| hCoVOC43 | spike | AILTEVNELLDTTQL | 843 | 857 | Negative |
| hCoVOC43 | spike | AISASLQEILSRLDA | 1056 | 1070 | Negative |
| hCoVOC43 | spike | ASLQEILSRLDALEA | 1059 | 1073 | Negative |
| hCoVOC43 | spike | ASSRSAIEDLLFDKV | 900 | 914 | Negative |
| hCoVOC43 | spike | AWLNDKSVPSPLNWE | 339 | 353 | Negative |
| hCoVOC43 | spike | DDINFSPVLGCLGSE | 882 | 896 | Negative |
| hCoVOC43 | spike | DGDNKLQGLLEVSVC | 150 | 164 | Negative |
| hCoVOC43 | spike | DIILGVCVNYDLYGI | 615 | 629 | Negative |
| hCoVOC43 | spike | DNINAILTEVNELLD | 839 | 853 | Negative |
| hCoVOC43 | spike | DVNADYLYFHFYQEG | 206 | 220 | Negative |
| hCoVOC43 | spike | DYLYFHFYQEGGTFY | 210 | 224 | Negative |
| hCoVOC43 | spike | ECSKASSRSAIEDLL | 896 | 910 | Negative |
| hCoVOC43 | spike | EGFDATNSALVKIQA | 1020 | 1034 | Negative |
| hCoVOC43 | spike | EIQIPSEFTIGNMVE | 789 | 803 | Negative |
| hCoVOC43 | spike | EIRDLICVQSYKGIK | 933 | 947 | Negative |
| hCoVOC43 | spike | ELDQWFKNQTSVAPD | 1236 | 1250 | Negative |
| hCoVOC43 | spike | ELLDTTQLQVANSLM | 850 | 864 | Negative |
| hCoVOC43 | spike | ELNGYTVQPIADVYR | 313 | 327 | Negative |
| hCoVOC43 | spike | ELVIKTSHDDGSGSG | 1344 | 1353 | Negative |
| hCoVOC43 | spike | ELWHLDTGVVSCLYK | 186 | 200 | Negative |
| hCoVOC43 | spike | FDKVKLSDVGFVEAY | 911 | 925 | Negative |
| hCoVOC43 | spike | FLDLQVEMNRLQEAI | 1260 | 1274 | Negative |
| hCoVOC43 | spike | FRDYITNRTFMIRSC | 660 | 674 | Negative |
| hCoVOC43 | spike | FTNFEPFTVNSVNDS | 766 | 780 | Negative |
| hCoVOC43 | spike | FVEAYNNCTGGAEIR | 921 | 935 | Negative |
| hCoVOC43 | spike | FYLNVQYRINGLGVT | 982 | 996 | Negative |

| | | | | | |
|---|---|---|---|---|---|
| hCoVOC43 | spike | GEHCSGLAVKSDYCG | 549 | 563 | Negative |
| hCoVOC43 | spike | GFIEDSVFKPRPAGV | 457 | 471 | Negative |
| hCoVOC43 | spike | GGCCDDYTGYQELVI | 1333 | 1347 | Negative |
| hCoVOC43 | spike | GGLYEIQIPSEFTIG | 785 | 799 | Negative |
| hCoVOC43 | spike | GGTFYAYFTDTGVVT | 220 | 234 | Negative |
| hCoVOC43 | spike | GIFVEVNATYYNSWQ | 633 | 647 | Negative |
| hCoVOC43 | spike | GIGEHCSGLAVKSDY | 547 | 561 | Negative |
| hCoVOC43 | spike | GIGTCPAGTNYLTCD | 509 | 523 | Negative |
| hCoVOC43 | spike | GIIFNAVDCMSDFMS | 282 | 296 | Negative |
| hCoVOC43 | spike | GILGQGIFVEVNATY | 628 | 642 | Negative |
| hCoVOC43 | spike | GKNNGIGTCPAGTNY | 505 | 519 | Negative |
| hCoVOC43 | spike | GLAVKSDYCGGNSCT | 554 | 568 | Negative |
| hCoVOC43 | spike | GLGTYYVLDRVYLNT | 46 | 60 | Negative |
| hCoVOC43 | spike | GLGVTMDVLSQNQKL | 992 | 1006 | Negative |
| hCoVOC43 | spike | GLTCSTDLQKANTDI | 602 | 616 | Negative |
| hCoVOC43 | spike | GLYEIQIPSEFTIGN | 786 | 800 | Negative |
| hCoVOC43 | spike | GMALSHYYVMPLTCN | 243 | 257 | Negative |
| hCoVOC43 | spike | GMCFSSITIDKFAIP | 388 | 402 | Negative |
| hCoVOC43 | spike | GMFLILLISLPTAFA | 1 | 14 | Negative |
| hCoVOC43 | spike | GNHRKELWHLDTGVV | 181 | 195 | Negative |
| hCoVOC43 | spike | GRKVDLQLGNLGYLQ | 404 | 418 | Negative |
| hCoVOC43 | spike | GSGSGMFLILLISLP | 1 | 10 | Negative |
| hCoVOC43 | spike | GSGSGSGMFLILLIS | 1 | 8 | Negative |
| hCoVOC43 | spike | GSGYYYPEPITENNV | 1191 | 1205 | Negative |
| hCoVOC43 | spike | GTCPAGTNYLTCDNL | 511 | 525 | Negative |
| hCoVOC43 | spike | GTNYLTCDNLCTPDP | 516 | 530 | Negative |
| hCoVOC43 | spike | GTYEYYVKWPWYVWL | 1288 | 1302 | Negative |
| hCoVOC43 | spike | GTYKCPQTKSLVGIG | 535 | 549 | Negative |
| hCoVOC43 | spike | GVAMLVLLFFICCCT | 1308 | 1322 | Negative |
| hCoVOC43 | spike | GVCVNYDLYGILGQG | 619 | 633 | Negative |
| hCoVOC43 | spike | GVNFNVDDINFSPVL | 876 | 890 | Negative |
| hCoVOC43 | spike | GVPFYLNVQYRINGL | 979 | 993 | Negative |
| hCoVOC43 | spike | GVTLSTKLKDGVNFN | 866 | 880 | Negative |
| hCoVOC43 | spike | GVTMDVLSQNQKLIA | 994 | 1008 | Negative |
| hCoVOC43 | spike | GVVSCLYKRNFTYDV | 193 | 207 | Negative |
| hCoVOC43 | spike | GVVTKFLFNVYLGMA | 231 | 245 | Negative |
| hCoVOC43 | spike | GVYELNGYTVQPIAD | 310 | 324 | Negative |
| hCoVOC43 | spike | GWSADSCLQGDKCNI | 576 | 590 | Negative |
| hCoVOC43 | spike | IEAWLNDKSVPSPLN | 337 | 351 | Negative |
| hCoVOC43 | spike | IGTYEYYVKWPWYVW | 1287 | 1301 | Negative |
| hCoVOC43 | spike | IILGVCVNYDLYGIL | 616 | 630 | Negative |
| hCoVOC43 | spike | KCKTQSIAPPTGVYE | 299 | 313 | Negative |
| hCoVOC43 | spike | KVDLQLGNLGYLQSF | 406 | 420 | Negative |
| hCoVOC43 | spike | LGNHRKELWHLDTGV | 180 | 194 | Negative |
| hCoVOC43 | spike | LGVCVNYDLYGILGQ | 618 | 632 | Negative |
| hCoVOC43 | spike | LGVTMDVLSQNQKLI | 993 | 1007 | Negative |
| hCoVOC43 | spike | LNTSIPNLPDFKEEL | 1223 | 1237 | Negative |
| hCoVOC43 | spike | LNVQYRINGLGVTMD | 984 | 998 | Negative |
| hCoVOC43 | spike | LQEILSRLDALEAEA | 1061 | 1075 | Negative |

| | | | | | |
|---|---|---|---|---|---|
| hCoVOC43 | spike | LTLEYWVTPLTSRQY | 260 | 274 | Negative |
| hCoVOC43 | spike | LWFKPPFLSDFINGI | 89 | 103 | Negative |
| hCoVOC43 | spike | MNGVTLSTKLKDGVN | 864 | 878 | Negative |
| hCoVOC43 | spike | NFCGNGNHIISLVQN | 1125 | 1139 | Negative |
| hCoVOC43 | spike | NGVTLSTKLKDGVNF | 865 | 879 | Negative |
| hCoVOC43 | spike | NHRKELWHLDTGVVS | 182 | 196 | Negative |
| hCoVOC43 | spike | NVQYRINGLGVTMDV | 985 | 999 | Negative |
| hCoVOC43 | spike | PFYLNVQYRINGLGV | 981 | 995 | Negative |
| hCoVOC43 | spike | RGAITTGYRFTNFEP | 757 | 771 | Negative |
| hCoVOC43 | spike | RKELWHLDTGVVSCL | 184 | 198 | Negative |
| hCoVOC43 | spike | RLDALEAEAQIDRLI | 1067 | 1081 | Negative |
| hCoVOC43 | spike | RVMYSEFPAITIGST | 116 | 130 | Negative |
| hCoVOC43 | spike | SAIEDLLFDKVKLSD | 904 | 918 | Negative |
| hCoVOC43 | spike | SFIQADSFTCNNIDA | 369 | 383 | Negative |
| hCoVOC43 | spike | SGSGSGMFLILLISL | 1 | 9 | Negative |
| hCoVOC43 | spike | SKLTLEYWVTPLTSR | 258 | 272 | Negative |
| hCoVOC43 | spike | SKNRRSRGAITTGYR | 751 | 765 | Negative |
| hCoVOC43 | spike | SRQYLLAFNQDGIIF | 271 | 285 | Negative |
| hCoVOC43 | spike | SSRINFCGNGNHIIS | 1121 | 1135 | Negative |
| hCoVOC43 | spike | SVAPDLSLDYINVTF | 1246 | 1260 | Negative |
| hCoVOC43 | spike | SVNDSLEPVGGLYEI | 776 | 790 | Negative |
| hCoVOC43 | spike | TAFAVIGDLKCTSDN | 11 | 25 | Negative |
| hCoVOC43 | spike | TFLDLQVEMNRLQEA | 1259 | 1273 | Negative |
| hCoVOC43 | spike | TGTYKCPQTKSLVGI | 534 | 548 | Negative |
| hCoVOC43 | spike | TGVVTKFLFNVYLGM | 230 | 244 | Negative |
| hCoVOC43 | spike | TINSTQDGDNKLQGL | 144 | 158 | Negative |
| hCoVOC43 | spike | TLEYWVTPLTSRQYL | 261 | 275 | Negative |
| hCoVOC43 | spike | TQLQVANSLMNGVTL | 855 | 869 | Negative |
| hCoVOC43 | spike | TSASLFPPWTAAAGV | 966 | 980 | Negative |
| hCoVOC43 | spike | TSIPNLPDFKEELDQ | 1225 | 1239 | Negative |
| hCoVOC43 | spike | TSRQYLLAFNQDGII | 270 | 284 | Negative |
| hCoVOC43 | spike | TTQLQVANSLMNGVT | 854 | 868 | Negative |
| hCoVOC43 | spike | TYKCPQTKSLVGIGE | 536 | 550 | Negative |
| hCoVOC43 | spike | VCVNYDLYGILGQGI | 620 | 634 | Negative |
| hCoVOC43 | spike | VGFVEAYNNCTGGAE | 919 | 933 | Negative |
| hCoVOC43 | spike | VKSQSSRINFCGNGN | 1117 | 1131 | Negative |
| hCoVOC43 | spike | VQYRINGLGVTMDVL | 986 | 1000 | Negative |
| hCoVOC43 | spike | VVQPRTINSTQDGDN | 139 | 153 | Negative |
| hCoVOC43 | spike | WFKPPFLSDFINGIF | 90 | 104 | Negative |
| hCoVOC43 | spike | YKCPQTKSLVGIGEH | 537 | 551 | Negative |
| hCoVOC43 | spike | YLNVQYRINGLGVTM | 983 | 997 | Negative |
| hCoVOC43 | spike | YRINGLGVTMDVLSQ | 988 | 1002 | Negative |
| hCoVOC43 | spike | YRNMALKGSVLLSRL | 75 | 89 | Negative |
| hCoVOC43 | spike | YRRKPNLPNCNIEAW | 326 | 340 | Negative |
| hCoVOC43 | spike | YSKNRRSRGAITTGY | 750 | 764 | Negative |
| hCoVOC43 | spike | YYNSWQNLLYDSNGN | 642 | 656 | Negative |
| hCoVOC43 | spike | AEALNNLLQQLSNRF | 1040 | 1054 | Overlapping |
| hCoVOC43 | spike | AEAQIDRLINGRLTA | 1073 | 1087 | Overlapping |
| hCoVOC43 | spike | AEIRDLICVQSYKGI | 932 | 946 | Overlapping |

| | | | | | |
|---|---|---|---|---|---|
| hCoVOC43 | spike | AVDCMSDFMSEIKCK | 287 | 301 | Overlapping |
| hCoVOC43 | spike | DDYTGYQELVIKTSH | 1337 | 1351 | Overlapping |
| hCoVOC43 | spike | DGIIFNAVDCMSDFM | 281 | 295 | Overlapping |
| hCoVOC43 | spike | DGVNFNVDDINFSPV | 875 | 889 | Overlapping |
| hCoVOC43 | spike | DYTGYQELVIKTSHD | 1338 | 1352 | Overlapping |
| hCoVOC43 | spike | EAEAQIDRLINGRLT | 1072 | 1086 | Overlapping |
| hCoVOC43 | spike | EAIKVLNQSYINLKD | 1272 | 1286 | Overlapping |
| hCoVOC43 | spike | EALNNLLQQLSNRFG | 1041 | 1055 | Overlapping |
| hCoVOC43 | spike | EAQIDRLINGRLTAL | 1074 | 1088 | Overlapping |
| hCoVOC43 | spike | EAWLNDKSVPSPLNW | 338 | 352 | Overlapping |
| hCoVOC43 | spike | EAYNNCTGGAEIRDL | 923 | 937 | Overlapping |
| hCoVOC43 | spike | ECVKSQSSRINFCGN | 1115 | 1129 | Overlapping |
| hCoVOC43 | spike | EDLLFDKVKLSDVGF | 907 | 921 | Overlapping |
| hCoVOC43 | spike | EDSVFKPRPAGVLTN | 460 | 474 | Overlapping |
| hCoVOC43 | spike | EELDQWFKNQTSVAP | 1235 | 1249 | Overlapping |
| hCoVOC43 | spike | EFIQTSSPKVTIDCA | 803 | 817 | Overlapping |
| hCoVOC43 | spike | EFPAITIGSTFVNTS | 121 | 135 | Overlapping |
| hCoVOC43 | spike | EFTIGNMVEFIQTSS | 795 | 809 | Overlapping |
| hCoVOC43 | spike | EGFDATNSALVKIQA | 1020 | 1034 | Overlapping |
| hCoVOC43 | spike | EGGTFYAYFTDTGVV | 219 | 233 | Overlapping |
| hCoVOC43 | spike | EHCSGLAVKSDYCGG | 550 | 564 | Overlapping |
| hCoVOC43 | spike | EILSRLDALEAEAQI | 1063 | 1077 | Overlapping |
| hCoVOC43 | spike | EIQIPSEFTIGNMVE | 789 | 803 | Overlapping |
| hCoVOC43 | spike | EIRDLICVQSYKGIK | 933 | 947 | Overlapping |
| hCoVOC43 | spike | EKVNECVKSQSSRIN | 1111 | 1125 | Overlapping |
| hCoVOC43 | spike | ELDQWFKNQTSVAPD | 1236 | 1250 | Overlapping |
| hCoVOC43 | spike | ELLDTTQLQVANSLM | 850 | 864 | Overlapping |
| hCoVOC43 | spike | ELNGYTVQPIADVYR | 313 | 327 | Overlapping |
| hCoVOC43 | spike | ELVIKTSHDDGSGSG | 1344 | 1353 | Overlapping |
| hCoVOC43 | spike | ELWHLDTGVVSCLYK | 186 | 200 | Overlapping |
| hCoVOC43 | spike | EMNRLQEAIKVLNQS | 1266 | 1280 | Overlapping |
| hCoVOC43 | spike | ENNVVVMSTCAVNYT | 1202 | 1216 | Overlapping |
| hCoVOC43 | spike | ENQISGYTLAATSAS | 955 | 969 | Overlapping |
| hCoVOC43 | spike | EPALLFRNIKCNYVF | 689 | 703 | Overlapping |
| hCoVOC43 | spike | EPVGGLYEIQIPSEF | 782 | 796 | Overlapping |
| hCoVOC43 | spike | ERKTFSNCNFNMSSL | 353 | 367 | Overlapping |
| hCoVOC43 | spike | EVNATYYNSWQNLLY | 637 | 651 | Overlapping |
| hCoVOC43 | spike | EVNELLDTTQLQVAN | 847 | 861 | Overlapping |
| hCoVOC43 | spike | EVSVCQYNMCEYPQT | 160 | 174 | Overlapping |
| hCoVOC43 | spike | EYGSFCDNINAILTE | 833 | 847 | Overlapping |
| hCoVOC43 | spike | EYPQTICHPNLGNHR | 170 | 184 | Overlapping |
| hCoVOC43 | spike | EYWVTPLTSRQYLLA | 263 | 277 | Overlapping |
| hCoVOC43 | spike | EYYVKWPWYVWLLIC | 1291 | 1305 | Overlapping |
| hCoVOC43 | spike | FGFIEDSVFKPRPAG | 456 | 470 | Overlapping |
| hCoVOC43 | spike | FNPSTWNKRFGFIED | 447 | 461 | Overlapping |
| hCoVOC43 | spike | GFDATNSALVKIQAV | 1021 | 1035 | Overlapping |
| hCoVOC43 | spike | GFVEAYNNCTGGAEI | 920 | 934 | Overlapping |
| hCoVOC43 | spike | GLLEVSVCQYNMCEY | 157 | 171 | Overlapping |
| hCoVOC43 | spike | GPPPISTDTVDVTNG | 32 | 46 | Overlapping |

| | | | | | |
|---|---|---|---|---|---|
| hCoVOC43 | spike | GTFYAYFTDTGVVTK | 221 | 235 | Overlapping |
| hCoVOC43 | spike | GVLTNHDVVYAQHCF | 470 | 484 | Overlapping |
| hCoVOC43 | spike | HRKELWHLDTGVVSC | 183 | 197 | Overlapping |
| hCoVOC43 | spike | IEDLLFDKVKLSDVG | 906 | 920 | Overlapping |
| hCoVOC43 | spike | IEDSVFKPRPAGVLT | 459 | 473 | Overlapping |
| hCoVOC43 | spike | IFNAVDCMSDFMSEI | 284 | 298 | Overlapping |
| hCoVOC43 | spike | ILTEVNELLDTTQLQ | 844 | 858 | Overlapping |
| hCoVOC43 | spike | KELWHLDTGVVSCLY | 185 | 199 | Overlapping |
| hCoVOC43 | spike | KKCGGCCDDYTGYQE | 1330 | 1344 | Overlapping |
| hCoVOC43 | spike | LTEVNELLDTTQLQV | 845 | 859 | Overlapping |
| hCoVOC43 | spike | MEKVNECVKSQSSRI | 1110 | 1124 | Overlapping |
| hCoVOC43 | spike | MVEFIQTSSPKVTID | 801 | 815 | Overlapping |
| hCoVOC43 | spike | NELLDTTQLQVANSL | 849 | 863 | Overlapping |
| hCoVOC43 | spike | NGRLTALNAYVSQQL | 1082 | 1096 | Overlapping |
| hCoVOC43 | spike | NGSCVGSGPGKNNGI | 496 | 510 | Overlapping |
| hCoVOC43 | spike | NTSIPNLPDFKEELD | 1224 | 1238 | Overlapping |
| hCoVOC43 | spike | NYDLYGILGQGIFVE | 623 | 637 | Overlapping |
| hCoVOC43 | spike | PIADVYRRKPNLPNC | 321 | 335 | Overlapping |
| hCoVOC43 | spike | PISTDTVDVTNGLGT | 35 | 49 | Overlapping |
| hCoVOC43 | spike | PITENNVVVMSTCAV | 1199 | 1213 | Overlapping |
| hCoVOC43 | spike | PTGVYELNGYTVQPI | 308 | 322 | Overlapping |
| hCoVOC43 | spike | PVGGLYEIQIPSEFT | 783 | 797 | Overlapping |
| hCoVOC43 | spike | QEGFDATNSALVKIQ | 1019 | 1033 | Overlapping |
| hCoVOC43 | spike | QEGGTFYAYFTDTGV | 218 | 232 | Overlapping |
| hCoVOC43 | spike | QEILSRLDALEAEAQ | 1062 | 1076 | Overlapping |
| hCoVOC43 | spike | QVEMNRLQEAIKVLN | 1264 | 1278 | Overlapping |
| hCoVOC43 | spike | RDLICVQSYKGIKVL | 935 | 949 | Overlapping |
| hCoVOC43 | spike | RIDTTATSCQLYYNL | 423 | 437 | Overlapping |
| hCoVOC43 | spike | SDFMSEIKCKTQSIA | 292 | 306 | Overlapping |
| hCoVOC43 | spike | SEFPAITIGSTFVNT | 120 | 134 | Overlapping |
| hCoVOC43 | spike | SFCDNINAILTEVNE | 836 | 850 | Overlapping |
| hCoVOC43 | spike | SLQEILSRLDALEAE | 1060 | 1074 | Overlapping |
| hCoVOC43 | spike | SSLMSFIQADSFTCN | 365 | 379 | Overlapping |
| hCoVOC43 | spike | SSRSAIEDLLFDKVK | 901 | 915 | Overlapping |
| hCoVOC43 | spike | STKLKDGVNFNVDDI | 870 | 884 | Overlapping |
| hCoVOC43 | spike | TEVNELLDTTQLQVA | 846 | 860 | Overlapping |
| hCoVOC43 | spike | TGVYELNGYTVQPIA | 309 | 323 | Overlapping |
| hCoVOC43 | spike | VEAYNNCTGGAEIRD | 922 | 936 | Overlapping |
| hCoVOC43 | spike | VEFIQTSSPKVTIDC | 802 | 816 | Overlapping |
| hCoVOC43 | spike | VEMNRLQEAIKVLNQ | 1265 | 1279 | Overlapping |
| hCoVOC43 | spike | VEVNATYYNSWQNLL | 636 | 650 | Overlapping |
| hCoVOC43 | spike | VEYGSFCDNINAILT | 832 | 846 | Overlapping |
| hCoVOC43 | spike | VIKTSHDDGSGSGSG | 1346 | 1353 | Overlapping |
| hCoVOC43 | spike | VLTNHDVVYAQHCFK | 471 | 485 | Overlapping |
| hCoVOC43 | spike | VTFLDLQVEMNRLQE | 1258 | 1272 | Overlapping |
| hCoVOC43 | spike | WERKTFSNCNFNMSS | 352 | 366 | Overlapping |
| hCoVOC43 | spike | YDVNADYLYFHFYQE | 205 | 219 | Overlapping |
| hCoVOC43 | spike | YELNGYTVQPIADVY | 312 | 326 | Overlapping |

**Table S4. HKU1-specific Peptides**

| Lineage | Protein | HDPA peptide sequence | AA pos start | AA pos end | Group |
|---|---|---|---|---|---|
| hCoVHKU1 | envelope | FFNDTAWYIGQILVL | 5 | 19 | Positive |
| hCoVHKU1 | envelope | GSGMVDLFFNDTAWY | 1 | 12 | Positive |
| hCoVHKU1 | envelope | GSGSGMVDLFFNDTA | 1 | 10 | Positive |
| hCoVHKU1 | envelope | GSGSGSGMVDLFFND | 1 | 8 | Positive |
| hCoVHKU1 | envelope | IPPTSDYLIGSGSGS | 74 | 82 | Positive |
| hCoVHKU1 | envelope | KLCMQLCGFCNFFII | 38 | 52 | Positive |
| hCoVHKU1 | envelope | SEQVIPPTSDYLIGS | 70 | 82 | Positive |
| hCoVHKU1 | envelope | FFNDTAWYIGQILVL | 5 | 19 | Negative |
| hCoVHKU1 | envelope | GMQLYKSYSEQVIPP | 62 | 76 | Negative |
| hCoVHKU1 | envelope | GSGMVDLFFNDTAWY | 1 | 12 | Negative |
| hCoVHKU1 | envelope | GSGSGMVDLFFNDTA | 1 | 10 | Negative |
| hCoVHKU1 | envelope | GSGSGSGMVDLFFND | 1 | 8 | Negative |
| hCoVHKU1 | envelope | MVDLFFNDTAWYIGQ | 1 | 15 | Negative |
| hCoVHKU1 | envelope | SGMVDLFFNDTAWYI | 1 | 13 | Negative |
| hCoVHKU1 | envelope | EQVIPPTSDYLIGSG | 71 | 82 | Overlapping |
| hCoVHKU1 | envelope | GMVDLFFNDTAWYIG | 1 | 14 | Overlapping |
| hCoVHKU1 | membrane | GHLYIQGVKLGTGYT | 148 | 162 | Positive |
| hCoVHKU1 | membrane | GSGMNKSFLPQFTSD | 1 | 12 | Positive |
| hCoVHKU1 | membrane | GSGSGMNKSFLPQFT | 1 | 10 | Positive |
| hCoVHKU1 | membrane | GSGSGSGMNKSFLPQ | 1 | 8 | Positive |
| hCoVHKU1 | membrane | GSWWSFNPETNNLMC | 108 | 122 | Positive |
| hCoVHKU1 | membrane | IRTGSWWSFNPETNN | 105 | 119 | Positive |
| hCoVHKU1 | membrane | NAFLAFSIVFTIISI | 76 | 90 | Positive |
| hCoVHKU1 | membrane | NKSFLPQFTSDQAVT | 2 | 16 | Positive |
| hCoVHKU1 | membrane | NLMCIDMKGKMFVRP | 119 | 133 | Positive |
| hCoVHKU1 | membrane | NNAFLAFSIVFTIIS | 75 | 89 | Positive |
| hCoVHKU1 | membrane | NNLMCIDMKGKMFVR | 118 | 132 | Positive |
| hCoVHKU1 | membrane | NYRLPSSKPSGMDTA | 205 | 219 | Positive |
| hCoVHKU1 | membrane | PETNNLMCIDMKGKM | 115 | 129 | Positive |
| hCoVHKU1 | membrane | PSGMDTALLRAGSGS | 213 | 223 | Positive |
| hCoVHKU1 | membrane | PVIEDYHTLTATVIR | 133 | 147 | Positive |
| hCoVHKU1 | membrane | RPVIEDYHTLTATVI | 132 | 146 | Positive |
| hCoVHKU1 | membrane | SDLPVYVTVAKVQVL | 164 | 178 | Positive |
| hCoVHKU1 | membrane | SDQAVTFLKEWNFSL | 11 | 25 | Positive |
| hCoVHKU1 | membrane | SMFVYLIKMIILWLM | 44 | 58 | Positive |
| hCoVHKU1 | membrane | SSKPSGMDTALLRAG | 210 | 223 | Positive |
| hCoVHKU1 | membrane | TLSDLPVYVTVAKVQ | 162 | 176 | Positive |
| hCoVHKU1 | membrane | TNNLMCIDMKGKMFV | 117 | 131 | Positive |
| hCoVHKU1 | membrane | TSDQAVTFLKEWNFS | 10 | 24 | Positive |
| hCoVHKU1 | membrane | AFLDKLDVNSGFAVF | 184 | 198 | Negative |
| hCoVHKU1 | membrane | AVTFLKEWNFSLGVI | 14 | 28 | Negative |
| hCoVHKU1 | membrane | GHLYIQGVKLGTGYT | 148 | 162 | Negative |
| hCoVHKU1 | membrane | GMDTALLRAGSGSGS | 215 | 223 | Negative |
| hCoVHKU1 | membrane | GMNKSFLPQFTSDQA | 1 | 14 | Negative |
| hCoVHKU1 | membrane | GSGSGMNKSFLPQFT | 1 | 10 | Negative |
| hCoVHKU1 | membrane | GSGSGSGMNKSFLPQ | 1 | 8 | Negative |
| hCoVHKU1 | membrane | GTGYTLSDLPVYVTV | 158 | 172 | Negative |

| | | | | | |
|---|---|---|---|---|---|
| hCoVHKU1 | membrane | GVILLFITIILQFGY | 26 | 40 | Negative |
| hCoVHKU1 | membrane | GVKLGTGYTLSDLPV | 154 | 168 | Negative |
| hCoVHKU1 | membrane | LDVNSGFAVFVKSKV | 189 | 203 | Negative |
| hCoVHKU1 | membrane | LGVILLFITIILQFG | 25 | 39 | Negative |
| hCoVHKU1 | membrane | LTATVIRGHLYIQGV | 141 | 155 | Negative |
| hCoVHKU1 | membrane | SKPSGMDTALLRAGS | 211 | 223 | Negative |
| hCoVHKU1 | membrane | VTFLKEWNFSLGVIL | 15 | 29 | Negative |
| hCoVHKU1 | membrane | YRLPSSKPSGMDTAL | 206 | 220 | Negative |
| hCoVHKU1 | membrane | YTLSDLPVYVTVAKV | 161 | 175 | Negative |
| hCoVHKU1 | membrane | EDYHTLTATVIRGHL | 136 | 150 | Overlapping |
| hCoVHKU1 | membrane | ETNNLMCIDMKGKMF | 116 | 130 | Overlapping |
| hCoVHKU1 | membrane | EWNFSLGVILLFITI | 20 | 34 | Overlapping |
| hCoVHKU1 | membrane | GMDTALLRAGSGSGS | 215 | 223 | Overlapping |
| hCoVHKU1 | membrane | IEDYHTLTATVIRGH | 135 | 149 | Overlapping |
| hCoVHKU1 | membrane | KEWNFSLGVILLFIT | 19 | 33 | Overlapping |
| hCoVHKU1 | membrane | PSSKPSGMDTALLRA | 209 | 223 | Overlapping |
| hCoVHKU1 | membrane | PVIEDYHTLTATVIR | 133 | 147 | Overlapping |
| hCoVHKU1 | membrane | SSKPSGMDTALLRAG | 210 | 223 | Overlapping |
| hCoVHKU1 | membrane | VIEDYHTLTATVIRG | 134 | 148 | Overlapping |
| hCoVHKU1 | nucleocapsid | AGTQHISNDFTPEDH | 411 | 425 | Positive |
| hCoVHKU1 | nucleocapsid | ASIPESKPLADDDSA | 228 | 234 | Positive |
| hCoVHKU1 | nucleocapsid | DDPYVEDSVAGSGSG | 432 | 441 | Positive |
| hCoVHKU1 | nucleocapsid | DFTPEDHSLLATLDD | 419 | 433 | Positive |
| hCoVHKU1 | nucleocapsid | DGVVWVAKEGAKTVN | 142 | 154 | Positive |
| hCoVHKU1 | nucleocapsid | DSEVSTDEVGDNVQI | 224 | 235 | Positive |
| hCoVHKU1 | nucleocapsid | EADSPVKDVFELHYS | 335 | 349 | Positive |
| hCoVHKU1 | nucleocapsid | EQIGYWNVQERWRMR | 95 | 105 | Positive |
| hCoVHKU1 | nucleocapsid | FDSTLPGFETIMKVL | 354 | 368 | Positive |
| hCoVHKU1 | nucleocapsid | FETIMKVLEENLNAY | 361 | 375 | Positive |
| hCoVHKU1 | nucleocapsid | FIEQISAFTKPSSIK | 88 | 97 | Positive |
| hCoVHKU1 | nucleocapsid | FWVANHQADTSTPSD | 145 | 159 | Positive |
| hCoVHKU1 | nucleocapsid | GHYAGSRSSSGNRSG | 6 | 20 | Positive |
| hCoVHKU1 | nucleocapsid | GRSASNSRPGSRSQS | 191 | 205 | Positive |
| hCoVHKU1 | nucleocapsid | GSGMSYTPGHYAGSR | 1 | 12 | Positive |
| hCoVHKU1 | nucleocapsid | GSGSGMSYTPGHYAG | 1 | 10 | Positive |
| hCoVHKU1 | nucleocapsid | GSGSGSGMSYTPGHY | 1 | 8 | Positive |
| hCoVHKU1 | nucleocapsid | GSRSSSGNRSGILKK | 10 | 24 | Positive |
| hCoVHKU1 | nucleocapsid | HQADTSTPSDVSSRD | 150 | 164 | Positive |
| hCoVHKU1 | nucleocapsid | HSDSIVKPDMADEIA | 222 | 236 | Positive |
| hCoVHKU1 | nucleocapsid | IPESKPLADDDSAII | 228 | 239 | Positive |
| hCoVHKU1 | nucleocapsid | IPHYSWFSGITQFQK | 59 | 73 | Positive |
| hCoVHKU1 | nucleocapsid | IRFDSTLPGFETIMK | 352 | 366 | Positive |
| hCoVHKU1 | nucleocapsid | ISNDFTPEDHSLLAT | 416 | 430 | Positive |
| hCoVHKU1 | nucleocapsid | IVKPDMADEIANLVL | 226 | 240 | Positive |
| hCoVHKU1 | nucleocapsid | KDEQIGYWNVQERWR | 94 | 105 | Positive |
| hCoVHKU1 | nucleocapsid | KDVFELHYSGSIRFD | 341 | 355 | Positive |
| hCoVHKU1 | nucleocapsid | KEMQSQSSHVAQNTV | 408 | 419 | Positive |
| hCoVHKU1 | nucleocapsid | KHCNVQQCFGKRGPS | 276 | 290 | Positive |
| hCoVHKU1 | nucleocapsid | KILTKPRQKRTPNKH | 263 | 277 | Positive |

| | | | | | |
|---|---|---|---|---|---|
| hCoVHKU1 | nucleocapsid | KLGKDSKPQQVTKQN | 242 | 256 | Positive |
| hCoVHKU1 | nucleocapsid | KPQQVTKQNAKEIRH | 248 | 262 | Positive |
| hCoVHKU1 | nucleocapsid | KQNAKEIRHKILTKP | 254 | 268 | Positive |
| hCoVHKU1 | nucleocapsid | KRDSEADSPVKDVFE | 331 | 345 | Positive |
| hCoVHKU1 | nucleocapsid | LADDDSAIIEIVNEV | 223 | 235 | Positive |
| hCoVHKU1 | nucleocapsid | LEENLNAYVNSNQNT | 368 | 382 | Positive |
| hCoVHKU1 | nucleocapsid | LNASIPESKPLADDD | 372 | 385 | Positive |
| hCoVHKU1 | nucleocapsid | LPEQFDSLNLSAGTQ | 400 | 414 | Positive |
| hCoVHKU1 | nucleocapsid | LSRSNSNFRHSDSIV | 213 | 227 | Positive |
| hCoVHKU1 | nucleocapsid | NAEMLKLGTNDPQFP | 295 | 309 | Positive |
| hCoVHKU1 | nucleocapsid | NAKEIRHKILTKPRQ | 256 | 270 | Positive |
| hCoVHKU1 | nucleocapsid | NASIPESKPLADDDS | 373 | 386 | Positive |
| hCoVHKU1 | nucleocapsid | NASYGESLEGVFWVA | 134 | 148 | Positive |
| hCoVHKU1 | nucleocapsid | NDFTPEDHSLLATLD | 418 | 432 | Positive |
| hCoVHKU1 | nucleocapsid | NFRHSDSIVKPDMAD | 219 | 233 | Positive |
| hCoVHKU1 | nucleocapsid | NKHCNVQQCFGKRGP | 275 | 289 | Positive |
| hCoVHKU1 | nucleocapsid | NLSAGTQHISNDFTP | 408 | 422 | Positive |
| hCoVHKU1 | nucleocapsid | NLVLAKLGKDSKPQQ | 237 | 251 | Positive |
| hCoVHKU1 | nucleocapsid | NNRSLSRSNSNFRHS | 209 | 223 | Positive |
| hCoVHKU1 | nucleocapsid | NQNTDSDSLSSKPQR | 379 | 393 | Positive |
| hCoVHKU1 | nucleocapsid | NRGRKTQPKFTVSTQ | 39 | 53 | Positive |
| hCoVHKU1 | nucleocapsid | NRSGILKKTSWADQS | 17 | 31 | Positive |
| hCoVHKU1 | nucleocapsid | NRSLSRSNSNFRHSD | 210 | 224 | Positive |
| hCoVHKU1 | nucleocapsid | NSNFRHSDSIVKPDM | 217 | 231 | Positive |
| hCoVHKU1 | nucleocapsid | NSNQNTDSDSLSSKP | 377 | 391 | Positive |
| hCoVHKU1 | nucleocapsid | NSRPGSRSQSRGPNN | 196 | 210 | Positive |
| hCoVHKU1 | nucleocapsid | PEQFDSLNLSAGTQH | 401 | 415 | Positive |
| hCoVHKU1 | nucleocapsid | PSDVSSRDPTTQEAI | 157 | 171 | Positive |
| hCoVHKU1 | nucleocapsid | PTPGAFFFGSKLDLV | 316 | 330 | Positive |
| hCoVHKU1 | nucleocapsid | PYVEDSVAGSGSGSG | 434 | 441 | Positive |
| hCoVHKU1 | nucleocapsid | QIGYWNVQERWRMRR | 98 | 105 | Positive |
| hCoVHKU1 | nucleocapsid | RDFKFSDGQGVPIAF | 75 | 89 | Positive |
| hCoVHKU1 | nucleocapsid | RDPTTQEAIPTRFPP | 163 | 177 | Positive |
| hCoVHKU1 | nucleocapsid | RDSEADSPVKDVFEL | 332 | 346 | Positive |
| hCoVHKU1 | nucleocapsid | REENVIQCFGPRDFN | 276 | 287 | Positive |
| hCoVHKU1 | nucleocapsid | RHSDSIVKPDMADEI | 221 | 235 | Positive |
| hCoVHKU1 | nucleocapsid | RSSSGNRSGILKKTS | 12 | 26 | Positive |
| hCoVHKU1 | nucleocapsid | SDGQGVPIAFGVPPS | 80 | 94 | Positive |
| hCoVHKU1 | nucleocapsid | SDSIVKPDMADEIAN | 223 | 237 | Positive |
| hCoVHKU1 | nucleocapsid | SDSLSSKPQRKRGVK | 384 | 398 | Positive |
| hCoVHKU1 | nucleocapsid | SDVSSRDPTTQEAIP | 158 | 172 | Positive |
| hCoVHKU1 | nucleocapsid | SEADSPVKDVFELHY | 334 | 348 | Positive |
| hCoVHKU1 | nucleocapsid | SEAKGYWYRHSRRSF | 94 | 108 | Positive |
| hCoVHKU1 | nucleocapsid | SERNYQTFNRGRKTQ | 31 | 45 | Positive |
| hCoVHKU1 | nucleocapsid | SIPESKPLADDDSAI | 422 | 436 | Positive |
| hCoVHKU1 | nucleocapsid | SLLATLDDPYVEDSV | 426 | 440 | Positive |
| hCoVHKU1 | nucleocapsid | SNDFTPEDHSLLATL | 417 | 431 | Positive |
| hCoVHKU1 | nucleocapsid | SNFRHSDSIVKPDMA | 218 | 232 | Positive |
| hCoVHKU1 | nucleocapsid | SSKPQRKRGVKQLPE | 388 | 402 | Positive |

| | | | | | |
|---|---|---|---|---|---|
| hCoVHKU1 | nucleocapsid | SSRASSRSSTRNNSR | 196 | 211 | Positive |
| hCoVHKU1 | nucleocapsid | SSRSSTRNNSRDSSR | 200 | 215 | Positive |
| hCoVHKU1 | nucleocapsid | STDEVGDNVQITYTY | 51 | 64 | Positive |
| hCoVHKU1 | nucleocapsid | STLPGFETIMKVLEE | 356 | 370 | Positive |
| hCoVHKU1 | nucleocapsid | SYGESLEGVFWVANH | 136 | 150 | Positive |
| hCoVHKU1 | nucleocapsid | TDEVGDNVQITYTYK | 80 | 89 | Positive |
| hCoVHKU1 | nucleocapsid | TDSDSLSSKPQRKRG | 382 | 396 | Positive |
| hCoVHKU1 | nucleocapsid | TGPHKDLKFRQRSDG | 75 | 82 | Positive |
| hCoVHKU1 | nucleocapsid | TLDDPYVEDSVAGSG | 430 | 441 | Positive |
| hCoVHKU1 | nucleocapsid | TPEDHSLLATLDDPY | 421 | 435 | Positive |
| hCoVHKU1 | nucleocapsid | TQEAIPTRFPPGTIL | 167 | 181 | Positive |
| hCoVHKU1 | nucleocapsid | TSTPSDVSSRDPTTQ | 154 | 168 | Positive |
| hCoVHKU1 | nucleocapsid | TVLNASIPESKPLAD | 239 | 249 | Positive |
| hCoVHKU1 | nucleocapsid | VANHQADTSTPSDVS | 147 | 161 | Positive |
| hCoVHKU1 | nucleocapsid | VDLPPKVHFYYLGTG | 117 | 130 | Positive |
| hCoVHKU1 | nucleocapsid | VLEENLNAYVNSNQN | 367 | 381 | Positive |
| hCoVHKU1 | nucleocapsid | VLNASIPESKPLADD | 416 | 433 | Positive |
| hCoVHKU1 | nucleocapsid | WNVQERWRMRRGQRV | 27 | 43 | Positive |
| hCoVHKU1 | nucleocapsid | WVANHQADTSTPSDV | 146 | 160 | Positive |
| hCoVHKU1 | nucleocapsid | YGESLEGVFWVANHQ | 137 | 151 | Positive |
| hCoVHKU1 | nucleocapsid | YWNVQERWRMRRGQR | 99 | 115 | Positive |
| hCoVHKU1 | nucleocapsid | ATLDDPYVEDSVAGS | 429 | 441 | Negative |
| hCoVHKU1 | nucleocapsid | CFGKRGPSQNFGNAE | 283 | 297 | Negative |
| hCoVHKU1 | nucleocapsid | DGQQKQLLPRWYFYY | 112 | 126 | Negative |
| hCoVHKU1 | nucleocapsid | EGVFWVANHQADTST | 142 | 156 | Negative |
| hCoVHKU1 | nucleocapsid | ELAPTPGAFFFGSKL | 313 | 327 | Negative |
| hCoVHKU1 | nucleocapsid | ELHYSGSIRFDSTLP | 345 | 359 | Negative |
| hCoVHKU1 | nucleocapsid | EMLKLGTNDPQFPIL | 297 | 311 | Negative |
| hCoVHKU1 | nucleocapsid | FELHYSGSIRFDSTL | 344 | 358 | Negative |
| hCoVHKU1 | nucleocapsid | FGVPPSEAKGYWYRH | 89 | 103 | Negative |
| hCoVHKU1 | nucleocapsid | FKFSDGQGVPIAFGV | 77 | 91 | Negative |
| hCoVHKU1 | nucleocapsid | FQKGRDFKFSDGQGV | 71 | 85 | Negative |
| hCoVHKU1 | nucleocapsid | GHYAGSRSSSGNRSG | 6 | 20 | Negative |
| hCoVHKU1 | nucleocapsid | GITQFQKGRDFKFSD | 67 | 81 | Negative |
| hCoVHKU1 | nucleocapsid | GKDSKPQQVTKQNAK | 244 | 258 | Negative |
| hCoVHKU1 | nucleocapsid | GQGVPIAFGVPPSEA | 82 | 96 | Negative |
| hCoVHKU1 | nucleocapsid | GRDFKFSDGQGVPIA | 74 | 88 | Negative |
| hCoVHKU1 | nucleocapsid | GSGMSYTPGHYAGSR | 1 | 12 | Negative |
| hCoVHKU1 | nucleocapsid | GSGSGMSYTPGHYAG | 1 | 10 | Negative |
| hCoVHKU1 | nucleocapsid | GSRSQSRGPNNRSLS | 200 | 214 | Negative |
| hCoVHKU1 | nucleocapsid | GTGPYANASYGESLE | 128 | 142 | Negative |
| hCoVHKU1 | nucleocapsid | GTILPQGYYVEGSGR | 178 | 192 | Negative |
| hCoVHKU1 | nucleocapsid | GTNDPQFPILAELAP | 302 | 316 | Negative |
| hCoVHKU1 | nucleocapsid | GVFWVANHQADTSTP | 143 | 157 | Negative |
| hCoVHKU1 | nucleocapsid | HSRRSFKTADGQQKQ | 103 | 117 | Negative |
| hCoVHKU1 | nucleocapsid | HYAGSRSSSGNRSGI | 7 | 21 | Negative |
| hCoVHKU1 | nucleocapsid | IMKVLEENLNAYVNS | 364 | 378 | Negative |
| hCoVHKU1 | nucleocapsid | IVKPDMADEIANLVL | 226 | 240 | Negative |
| hCoVHKU1 | nucleocapsid | KVLEENLNAYVNSNQ | 366 | 380 | Negative |

| | | | | | |
|---|---|---|---|---|---|
| hCoVHKU1 | nucleocapsid | LNAYVNSNQNTDSDS | 372 | 386 | Negative |
| hCoVHKU1 | nucleocapsid | NYQTFNRGRKTQPKF | 34 | 48 | Negative |
| hCoVHKU1 | nucleocapsid | PQGNTIPHYSWFSGI | 54 | 68 | Negative |
| hCoVHKU1 | nucleocapsid | QGVPIAFGVPPSEAK | 83 | 97 | Negative |
| hCoVHKU1 | nucleocapsid | QKGRDFKFSDGQGVP | 72 | 86 | Negative |
| hCoVHKU1 | nucleocapsid | RNYQTFNRGRKTQPK | 33 | 47 | Negative |
| hCoVHKU1 | nucleocapsid | SDSIVKPDMADEIAN | 223 | 237 | Negative |
| hCoVHKU1 | nucleocapsid | SFKTADGQQKQLLPR | 107 | 121 | Negative |
| hCoVHKU1 | nucleocapsid | SIRFDSTLPGFETIM | 351 | 365 | Negative |
| hCoVHKU1 | nucleocapsid | SRRSFKTADGQQKQL | 104 | 118 | Negative |
| hCoVHKU1 | nucleocapsid | TFNRGRKTQPKFTVS | 37 | 51 | Negative |
| hCoVHKU1 | nucleocapsid | VEGSGRSASNSRPGS | 187 | 201 | Negative |
| hCoVHKU1 | nucleocapsid | VSSRDPTTQEAIPTR | 160 | 174 | Negative |
| hCoVHKU1 | nucleocapsid | YRHSRRSFKTADGQQ | 101 | 115 | Negative |
| hCoVHKU1 | nucleocapsid | YTPGHYAGSRSSSGN | 3 | 17 | Negative |
| hCoVHKU1 | nucleocapsid | DSEADSPVKDVFELH | 333 | 347 | Overlapping |
| hCoVHKU1 | nucleocapsid | EADSPVKDVFELHYS | 335 | 349 | Overlapping |
| hCoVHKU1 | nucleocapsid | EAIPTRFPPGTILPQ | 169 | 183 | Overlapping |
| hCoVHKU1 | nucleocapsid | EAKGYWYRHSRRSFK | 95 | 109 | Overlapping |
| hCoVHKU1 | nucleocapsid | EDHSLLATLDDPYVE | 423 | 437 | Overlapping |
| hCoVHKU1 | nucleocapsid | EENLNAYVNSNQNTD | 369 | 383 | Overlapping |
| hCoVHKU1 | nucleocapsid | EGSGRSASNSRPGSR | 188 | 202 | Overlapping |
| hCoVHKU1 | nucleocapsid | EGVFWVANHQADTST | 142 | 156 | Overlapping |
| hCoVHKU1 | nucleocapsid | EIANLVLAKLGKDSK | 234 | 248 | Overlapping |
| hCoVHKU1 | nucleocapsid | EIRHKILTKPRQKRT | 259 | 273 | Overlapping |
| hCoVHKU1 | nucleocapsid | ELHYSGSIRFDSTLP | 345 | 359 | Overlapping |
| hCoVHKU1 | nucleocapsid | ENLNAYVNSNQNTDS | 370 | 384 | Overlapping |
| hCoVHKU1 | nucleocapsid | EQFDSLNLSAGTQHI | 402 | 416 | Overlapping |
| hCoVHKU1 | nucleocapsid | ERNYQTFNRGRKTQP | 32 | 46 | Overlapping |
| hCoVHKU1 | nucleocapsid | ESLEGVFWVANHQAD | 139 | 153 | Overlapping |
| hCoVHKU1 | nucleocapsid | ETIMKVLEENLNAYV | 362 | 376 | Overlapping |
| hCoVHKU1 | nucleocapsid | FTPEDHSLLATLDDP | 420 | 434 | Overlapping |
| hCoVHKU1 | nucleocapsid | GESLEGVFWVANHQA | 138 | 152 | Overlapping |
| hCoVHKU1 | nucleocapsid | GMSYTPGHYAGSRSS | 1 | 14 | Overlapping |
| hCoVHKU1 | nucleocapsid | GPYANASYGESLEGV | 130 | 144 | Overlapping |
| hCoVHKU1 | nucleocapsid | GTQHISNDFTPEDHS | 412 | 426 | Overlapping |
| hCoVHKU1 | nucleocapsid | GVPIAFGVPPSEAKG | 84 | 98 | Overlapping |
| hCoVHKU1 | nucleocapsid | GVPPSEAKGYWYRHS | 90 | 104 | Overlapping |
| hCoVHKU1 | nucleocapsid | KEIRHKILTKPRQKR | 258 | 272 | Overlapping |
| hCoVHKU1 | nucleocapsid | NAYVNSNQNTDSDSL | 373 | 387 | Overlapping |
| hCoVHKU1 | nucleocapsid | NLNAYVNSNQNTDSD | 371 | 385 | Overlapping |
| hCoVHKU1 | nucleocapsid | PEDHSLLATLDDPYV | 422 | 436 | Overlapping |
| hCoVHKU1 | nucleocapsid | PILAELAPTPGAFFF | 309 | 323 | Overlapping |
| hCoVHKU1 | nucleocapsid | PTTQEAIPTRFPPGT | 165 | 179 | Overlapping |
| hCoVHKU1 | nucleocapsid | PVKDVFELHYSGSIR | 339 | 353 | Overlapping |
| hCoVHKU1 | nucleocapsid | QEAIPTRFPPGTILP | 168 | 182 | Overlapping |
| hCoVHKU1 | nucleocapsid | RFDSTLPGFETIMKV | 353 | 367 | Overlapping |
| hCoVHKU1 | nucleocapsid | SSKPQRKRGVKQLPE | 388 | 402 | Overlapping |
| hCoVHKU1 | nucleocapsid | SSRDPTTQEAIPTRF | 161 | 175 | Overlapping |

| | | | | | |
|---|---|---|---|---|---|
| hCoVHKU1 | orf1ab | AADLPCTLHFSLFDD | 1274 | 1288 | Positive |
| hCoVHKU1 | orf1ab | ACSQLAKAMETFNHN | 3300 | 3314 | Positive |
| hCoVHKU1 | orf1ab | AEHDFFTFDIDGSRV | 4554 | 4568 | Positive |
| hCoVHKU1 | orf1ab | AESVIVEDDVIENVK | 740 | 754 | Positive |
| hCoVHKU1 | orf1ab | AETQKATEEAFKQSY | 5520 | 5534 | Positive |
| hCoVHKU1 | orf1ab | AIDAYPLVHHENEEY | 5316 | 5330 | Positive |
| hCoVHKU1 | orf1ab | AIFITMHSINHDIFS | 3760 | 3774 | Positive |
| hCoVHKU1 | orf1ab | ALPELVTDIVVVDEV | 5746 | 5760 | Positive |
| hCoVHKU1 | orf1ab | ALQTMLFSMVRKLDN | 4100 | 4114 | Positive |
| hCoVHKU1 | orf1ab | ANRYNEVANAVMQNN | 4198 | 4212 | Positive |
| hCoVHKU1 | orf1ab | APEFRWLLPDAAEEL | 14 | 28 | Positive |
| hCoVHKU1 | orf1ab | ARKHEFCCSHGDRFY | 5092 | 5106 | Positive |
| hCoVHKU1 | orf1ab | AVCSKHAEEYCNYLE | 6464 | 6478 | Positive |
| hCoVHKU1 | orf1ab | CEILCEYADCKESYF | 4600 | 4614 | Positive |
| hCoVHKU1 | orf1ab | CHDEASLNSLTYFNK | 2074 | 2088 | Positive |
| hCoVHKU1 | orf1ab | CKFSIQDVKGLKIKY | 4280 | 4294 | Positive |
| hCoVHKU1 | orf1ab | CLASIDEVSDDYVQD | 3994 | 4008 | Positive |
| hCoVHKU1 | orf1ab | CRVGACEYAEEGICF | 3058 | 3072 | Positive |
| hCoVHKU1 | orf1ab | CSEFEVEKGVTVDDF | 848 | 862 | Positive |
| hCoVHKU1 | orf1ab | CTGSPYIDDFNKIAS | 5482 | 5496 | Positive |
| hCoVHKU1 | orf1ab | CWVENDITNGPHEFC | 5252 | 5266 | Positive |
| hCoVHKU1 | orf1ab | DAIIVNDENSSIKVV | 2140 | 2154 | Positive |
| hCoVHKU1 | orf1ab | DALFVSYGSSVETET | 1130 | 1144 | Positive |
| hCoVHKU1 | orf1ab | DAVMHFGTLAKTDLF | 1848 | 1862 | Positive |
| hCoVHKU1 | orf1ab | DAYAEVHAEPKGKYS | 196 | 210 | Positive |
| hCoVHKU1 | orf1ab | DAYPLVHHENEEYQK | 5318 | 5332 | Positive |
| hCoVHKU1 | orf1ab | DCHWDLIISDMYDPL | 7004 | 7018 | Positive |
| hCoVHKU1 | orf1ab | DDEDVVTGDNDDEDV | 946 | 960 | Positive |
| hCoVHKU1 | orf1ab | DDEDVVTGDNNDEEI | 1076 | 1090 | Positive |
| hCoVHKU1 | orf1ab | DEPSDATDFIRVVLK | 1806 | 1820 | Positive |
| hCoVHKU1 | orf1ab | DFNKIASCKWTEVDD | 5490 | 5504 | Positive |
| hCoVHKU1 | orf1ab | DFSIKSIYNVDLCDC | 340 | 354 | Positive |
| hCoVHKU1 | orf1ab | DGLESKQVDYVPLRS | 6438 | 6452 | Positive |
| hCoVHKU1 | orf1ab | DGLKFTEESFYKNMY | 5364 | 5378 | Positive |
| hCoVHKU1 | orf1ab | DGQRVYDDVDASLFV | 2584 | 2598 | Positive |
| hCoVHKU1 | orf1ab | DGVVCYNSDYASKGY | 5214 | 5228 | Positive |
| hCoVHKU1 | orf1ab | DHIVYGSFNHKVIGG | 6770 | 6784 | Positive |
| hCoVHKU1 | orf1ab | DIQQWGYTGSLTSNH | 6230 | 6244 | Positive |
| hCoVHKU1 | orf1ab | DLFMLETMHWLIRFI | 2430 | 2444 | Positive |
| hCoVHKU1 | orf1ab | DLVQYDFTDYKLELF | 4722 | 4736 | Positive |
| hCoVHKU1 | orf1ab | DMDDNLFIAKYGLED | 6752 | 6766 | Positive |
| hCoVHKU1 | orf1ab | DNDDEDVVTGDNDDE | 954 | 968 | Positive |
| hCoVHKU1 | orf1ab | DNDDEDVVTGDNNDE | 1074 | 1088 | Positive |
| hCoVHKU1 | orf1ab | DNNDEEIVTGDNDDQ | 1084 | 1098 | Positive |
| hCoVHKU1 | orf1ab | DPAMHVASASALLDL | 4830 | 4844 | Positive |
| hCoVHKU1 | orf1ab | DRNDCSILCEILCEY | 4592 | 4606 | Positive |
| hCoVHKU1 | orf1ab | DSLFCSSTYGVCKYT | 6600 | 6614 | Positive |
| hCoVHKU1 | orf1ab | DSYYSYMMPMLTMCH | 4692 | 4706 | Positive |
| hCoVHKU1 | orf1ab | DTINDDEDVVTGDND | 942 | 956 | Positive |

| | | | | | |
|---|---|---|---|---|---|
| hCoVHKU1 | orf1ab | DTVNHDSFNLYGYSV | 418 | 432 | Positive |
| hCoVHKU1 | orf1ab | EDDVIENVKSSLSSY | 746 | 760 | Positive |
| hCoVHKU1 | orf1ab | EDVVTGDNDDEDVVT | 948 | 962 | Positive |
| hCoVHKU1 | orf1ab | EDVVTGDNNDEEIVT | 1078 | 1092 | Positive |
| hCoVHKU1 | orf1ab | EESFYKNMYLKSAVM | 5370 | 5384 | Positive |
| hCoVHKU1 | orf1ab | ELSTGCHTGTDFTGN | 3500 | 3514 | Positive |
| hCoVHKU1 | orf1ab | ENEEYQKVFRVYLEY | 5326 | 5340 | Positive |
| hCoVHKU1 | orf1ab | ENRYSVLSVDSVSEE | 2094 | 2108 | Positive |
| hCoVHKU1 | orf1ab | ESDADRANFLNAVVF | 2620 | 2634 | Positive |
| hCoVHKU1 | orf1ab | EYDYVIYSQTAETAH | 5924 | 5938 | Positive |
| hCoVHKU1 | orf1ab | FAAETQKATEEAFKQ | 5518 | 5532 | Positive |
| hCoVHKU1 | orf1ab | FCVKHQWNCFNCHSF | 2520 | 2534 | Positive |
| hCoVHKU1 | orf1ab | FDDILGKVCSEFEVE | 840 | 854 | Positive |
| hCoVHKU1 | orf1ab | FDMTKFSLKLAGTAV | 7128 | 7142 | Positive |
| hCoVHKU1 | orf1ab | FDQWYESTFGSVYYH | 2906 | 2920 | Positive |
| hCoVHKU1 | orf1ab | FETDAYKLFLSGDDC | 1542 | 1556 | Positive |
| hCoVHKU1 | orf1ab | FGEDFLGWIVPFGFM | 146 | 160 | Positive |
| hCoVHKU1 | orf1ab | FGKARLYYEALSFEE | 4962 | 4976 | Positive |
| hCoVHKU1 | orf1ab | FGKPVIWFCHDEASL | 2066 | 2080 | Positive |
| hCoVHKU1 | orf1ab | FIDMDDNLFIAKYGL | 6750 | 6764 | Positive |
| hCoVHKU1 | orf1ab | FITMHSINHDIFSLM | 3762 | 3776 | Positive |
| hCoVHKU1 | orf1ab | FKINDTTCKYVFTTI | 5730 | 5744 | Positive |
| hCoVHKU1 | orf1ab | FNDVYNDALFVSYGS | 1124 | 1138 | Positive |
| hCoVHKU1 | orf1ab | FTNDKTIFYTTEIAS | 2246 | 2260 | Positive |
| hCoVHKU1 | orf1ab | FVDDLLKTDSVLLIE | 5296 | 5310 | Positive |
| hCoVHKU1 | orf1ab | FVMQYGYVDCFNDNC | 356 | 370 | Positive |
| hCoVHKU1 | orf1ab | FVNEYYEFLCKHFSM | 5194 | 5208 | Positive |
| hCoVHKU1 | orf1ab | FVSLAIDAYPLVHHE | 5312 | 5326 | Positive |
| hCoVHKU1 | orf1ab | FWFCVTYVAMVIANH | 3212 | 3226 | Positive |
| hCoVHKU1 | orf1ab | GAGSDKEVAPGSAVL | 6954 | 6968 | Positive |
| hCoVHKU1 | orf1ab | GDEAMASRMYCTFAI | 906 | 920 | Positive |
| hCoVHKU1 | orf1ab | GDPAQLPAPRVLLSK | 5784 | 5798 | Positive |
| hCoVHKU1 | orf1ab | GFPKGYVMGLFRSYK | 114 | 128 | Positive |
| hCoVHKU1 | orf1ab | GGAVCSKHAEEYCNY | 6462 | 6476 | Positive |
| hCoVHKU1 | orf1ab | GGSLYVNKHAFHTNP | 6398 | 6412 | Positive |
| hCoVHKU1 | orf1ab | GHYDGRTGELPCAIM | 6522 | 6536 | Positive |
| hCoVHKU1 | orf1ab | GHYSLQDMRQKQSVW | 248 | 262 | Positive |
| hCoVHKU1 | orf1ab | GNPKGLACVKDYEFK | 6320 | 6334 | Positive |
| hCoVHKU1 | orf1ab | GPIFNRALLNTVIFA | 4636 | 4650 | Positive |
| hCoVHKU1 | orf1ab | GPPGTGKSHLAIGLA | 5666 | 5680 | Positive |
| hCoVHKU1 | orf1ab | GQTTHESSSAVNIQQ | 5862 | 5876 | Positive |
| hCoVHKU1 | orf1ab | GRPHRLVALVLAKGH | 1788 | 1802 | Positive |
| hCoVHKU1 | orf1ab | GRPQGAFHVTMRSSY | 3454 | 3468 | Positive |
| hCoVHKU1 | orf1ab | GSCTFEDELAPSDVY | 3614 | 3628 | Positive |
| hCoVHKU1 | orf1ab | GSGMIKTSKYGLGFK | 1 | 12 | Positive |
| hCoVHKU1 | orf1ab | GSGSGMIKTSKYGLG | 1 | 10 | Positive |
| hCoVHKU1 | orf1ab | GSGSGSGMIKTSKYG | 1 | 8 | Positive |
| hCoVHKU1 | orf1ab | GSNLEEDVLLFITAF | 3792 | 3806 | Positive |
| hCoVHKU1 | orf1ab | GSPYIDDFNKIASCK | 5484 | 5498 | Positive |

| | | | | | |
|---|---|---|---|---|---|
| hCoVHKU1 | orf1ab | GWVGFDVEGAHATRE | 6072 | 6086 | Positive |
| hCoVHKU1 | orf1ab | GYKIGCNCAGRIVHC | 1864 | 1878 | Positive |
| hCoVHKU1 | orf1ab | HAEPKGKYSQKAYAL | 202 | 216 | Positive |
| hCoVHKU1 | orf1ab | HAFHTNPFTRTVFEN | 6406 | 6420 | Positive |
| hCoVHKU1 | orf1ab | HDFFTFDIDGSRVPH | 4556 | 4570 | Positive |
| hCoVHKU1 | orf1ab | HHENEEYQKVFRVYL | 5324 | 5338 | Positive |
| hCoVHKU1 | orf1ab | HHPELKILRNLNIDI | 6574 | 6588 | Positive |
| hCoVHKU1 | orf1ab | HLSMTTSTTNFGEDF | 136 | 150 | Positive |
| hCoVHKU1 | orf1ab | HNEILSTSDLSVAFD | 3960 | 3974 | Positive |
| hCoVHKU1 | orf1ab | HNMTLIAADLPCTLH | 1268 | 1282 | Positive |
| hCoVHKU1 | orf1ab | HNNCYVNVACLMLQH | 1754 | 1768 | Positive |
| hCoVHKU1 | orf1ab | HPNTVDCDNDRCIIH | 4748 | 4762 | Positive |
| hCoVHKU1 | orf1ab | HSINHDIFSLMFLVG | 3766 | 3780 | Positive |
| hCoVHKU1 | orf1ab | HYTHLKCGSPYQHYD | 1914 | 1928 | Positive |
| hCoVHKU1 | orf1ab | IAVELFTKRSIRHHP | 6562 | 6576 | Positive |
| hCoVHKU1 | orf1ab | IDDFNKIASCKWTEV | 5488 | 5502 | Positive |
| hCoVHKU1 | orf1ab | IEDVEDVISSEAVED | 920 | 934 | Positive |
| hCoVHKU1 | orf1ab | IEVSQIQSKLTDVKC | 3918 | 3932 | Positive |
| hCoVHKU1 | orf1ab | IGMKRYCTVQGPPGT | 5656 | 5670 | Positive |
| hCoVHKU1 | orf1ab | ILFDGRDTGALEAFR | 6624 | 6638 | Positive |
| hCoVHKU1 | orf1ab | ILGSCTFEDELAPSD | 3612 | 3626 | Positive |
| hCoVHKU1 | orf1ab | INHDIFSLMFLVGRI | 3768 | 3782 | Positive |
| hCoVHKU1 | orf1ab | INNLLHSKVKVVPNL | 2600 | 2614 | Positive |
| hCoVHKU1 | orf1ab | IPALAANTLTIVIPD | 4134 | 4148 | Positive |
| hCoVHKU1 | orf1ab | IPDKQVFDKVVDNVY | 4146 | 4160 | Positive |
| hCoVHKU1 | orf1ab | IPDYAKILVNKFYTF | 664 | 678 | Positive |
| hCoVHKU1 | orf1ab | IPENPVLFTNSTDTV | 406 | 420 | Positive |
| hCoVHKU1 | orf1ab | IPIDLLCLRDDNQTL | 2202 | 2216 | Positive |
| hCoVHKU1 | orf1ab | IPSLMTVKVMFDLDS | 824 | 838 | Positive |
| hCoVHKU1 | orf1ab | IQEVLLLRHDIQLNN | 1564 | 1578 | Positive |
| hCoVHKU1 | orf1ab | IQTVQDADGINKQLT | 4170 | 4184 | Positive |
| hCoVHKU1 | orf1ab | IREPVHLLSADAIVL | 312 | 326 | Positive |
| hCoVHKU1 | orf1ab | ISMWYFGSNLEEDVL | 3786 | 3800 | Positive |
| hCoVHKU1 | orf1ab | ISTEKLSRLSMIKGP | 6646 | 6660 | Positive |
| hCoVHKU1 | orf1ab | ISTKEVKLKGVRKTV | 2122 | 2136 | Positive |
| hCoVHKU1 | orf1ab | ISVNDLCFANKFFQF | 2892 | 2906 | Positive |
| hCoVHKU1 | orf1ab | ISVQELRYMNANGLR | 3880 | 3894 | Positive |
| hCoVHKU1 | orf1ab | ITNGPHEFCSQHTML | 5258 | 5272 | Positive |
| hCoVHKU1 | orf1ab | IVEDDVIENVKSSLS | 744 | 758 | Positive |
| hCoVHKU1 | orf1ab | IVELVIGYSLYTVWF | 2398 | 2412 | Positive |
| hCoVHKU1 | orf1ab | IVVVDEVSMLTNYEL | 5754 | 5768 | Positive |
| hCoVHKU1 | orf1ab | IYDFDTYKALLVFND | 1112 | 1126 | Positive |
| hCoVHKU1 | orf1ab | KAKHYVYIGDPAQLP | 5776 | 5790 | Positive |
| hCoVHKU1 | orf1ab | KARLYYEALSFEEQN | 4964 | 4978 | Positive |
| hCoVHKU1 | orf1ab | KDPILYVLTHDVCQV | 4424 | 4438 | Positive |
| hCoVHKU1 | orf1ab | KEARINDKKSKVVSA | 4086 | 4100 | Positive |
| hCoVHKU1 | orf1ab | KEDFDIIQKCQITSV | 1508 | 1522 | Positive |
| hCoVHKU1 | orf1ab | KEHPVVGYQVRAFLN | 878 | 892 | Positive |
| hCoVHKU1 | orf1ab | KEIFVGDSLVNTCGS | 7170 | 7182 | Positive |

| | | | | | |
|---|---|---|---|---|---|
| hCoVHKU1 | orf1ab | KEIVETVSALVYDNK | 5830 | 5844 | Positive |
| hCoVHKU1 | orf1ab | KESYFSKKDWYDFVE | 4610 | 4624 | Positive |
| hCoVHKU1 | orf1ab | KFKHLIPLMSKGQKW | 6132 | 6146 | Positive |
| hCoVHKU1 | orf1ab | KFVYMHQLELSTGCH | 3492 | 3506 | Positive |
| hCoVHKU1 | orf1ab | KGHFKFDEPSDATDF | 1800 | 1814 | Positive |
| hCoVHKU1 | orf1ab | KHEFCCSHGDRFYRL | 5094 | 5108 | Positive |
| hCoVHKU1 | orf1ab | KHLIPLMSKGQKWDI | 6134 | 6148 | Positive |
| hCoVHKU1 | orf1ab | KHVQGNVAKAANISC | 2772 | 2786 | Positive |
| hCoVHKU1 | orf1ab | KIDGDYVYLPYPDPS | 5274 | 5288 | Positive |
| hCoVHKU1 | orf1ab | KIEPCIVSVTYGSMT | 3346 | 3360 | Positive |
| hCoVHKU1 | orf1ab | KLDKFFVVKRTNLEV | 4522 | 4536 | Positive |
| hCoVHKU1 | orf1ab | KLERMADLALTNMYK | 4072 | 4086 | Positive |
| hCoVHKU1 | orf1ab | KLVGHDICAQLNDKL | 2010 | 2024 | Positive |
| hCoVHKU1 | orf1ab | KPEATINQDSYGGAS | 4378 | 4392 | Positive |
| hCoVHKU1 | orf1ab | KPPLNKNYVFTGYHF | 5558 | 5572 | Positive |
| hCoVHKU1 | orf1ab | KPVIWFCHDEASLNS | 2068 | 2082 | Positive |
| hCoVHKU1 | orf1ab | KQLFYVYDVHKDNFK | 6346 | 6360 | Positive |
| hCoVHKU1 | orf1ab | KRPVNPTDASHYVVT | 2554 | 2568 | Positive |
| hCoVHKU1 | orf1ab | KSDIHLPAYASFKVI | 2870 | 2884 | Positive |
| hCoVHKU1 | orf1ab | KSPLHSLGGFPKGYV | 106 | 120 | Positive |
| hCoVHKU1 | orf1ab | KTIFYTTEIASKFTF | 2250 | 2264 | Positive |
| hCoVHKU1 | orf1ab | KTKRYVVHHLSMTTS | 128 | 142 | Positive |
| hCoVHKU1 | orf1ab | KTVAKAPPGEKFKHL | 6122 | 6136 | Positive |
| hCoVHKU1 | orf1ab | KVLRHGFHVLHFLTY | 2946 | 2960 | Positive |
| hCoVHKU1 | orf1ab | KYTMLDLCYALRHFD | 4578 | 4592 | Positive |
| hCoVHKU1 | orf1ab | LCCKCCYDHVMATNH | 5410 | 5424 | Positive |
| hCoVHKU1 | orf1ab | LCFKVYFKGQTTHES | 5854 | 5868 | Positive |
| hCoVHKU1 | orf1ab | LCVFAIFITMHSINH | 3756 | 3770 | Positive |
| hCoVHKU1 | orf1ab | LDALASMTGVSIETL | 3582 | 3596 | Positive |
| hCoVHKU1 | orf1ab | LDCELFVNDSYRQFD | 4708 | 4722 | Positive |
| hCoVHKU1 | orf1ab | LDDVEMVAYNPDLSQ | 1964 | 1978 | Positive |
| hCoVHKU1 | orf1ab | LDYVQCKCGNLEQNH | 486 | 500 | Positive |
| hCoVHKU1 | orf1ab | LFTNSTDTVNHDSFN | 412 | 426 | Positive |
| hCoVHKU1 | orf1ab | LFVNDSYRQFDLVQY | 4712 | 4726 | Positive |
| hCoVHKU1 | orf1ab | LGGMSYYCENHKPHY | 5450 | 5464 | Positive |
| hCoVHKU1 | orf1ab | LIERFVSLAIDAYPL | 5308 | 5322 | Positive |
| hCoVHKU1 | orf1ab | LIRDIFVHEDLHVVE | 76 | 90 | Positive |
| hCoVHKU1 | orf1ab | LLIQEFLQYDSSIHS | 6800 | 6814 | Positive |
| hCoVHKU1 | orf1ab | LLPDAAEELASPMKS | 20 | 34 | Positive |
| hCoVHKU1 | orf1ab | LREGVQLEKVLDTFV | 2694 | 2708 | Positive |
| hCoVHKU1 | orf1ab | LSFEEQNEIYAYTKR | 4972 | 4986 | Positive |
| hCoVHKU1 | orf1ab | LSTCDGLKFTEESFY | 5360 | 5374 | Positive |
| hCoVHKU1 | orf1ab | LTKNIGDYNVSKDGF | 7018 | 7032 | Positive |
| hCoVHKU1 | orf1ab | LTLDNQDLYGQWYDF | 4662 | 4676 | Positive |
| hCoVHKU1 | orf1ab | LTMCHVLDCELFVND | 4702 | 4716 | Positive |
| hCoVHKU1 | orf1ab | LVHHENEEYQKVFRV | 5322 | 5336 | Positive |
| hCoVHKU1 | orf1ab | LYVLTHDVCQVCGFW | 4428 | 4442 | Positive |
| hCoVHKU1 | orf1ab | MCHVLDCELFVNDSY | 4704 | 4718 | Positive |
| hCoVHKU1 | orf1ab | MDEDIGSTMFNVPTK | 2932 | 2946 | Positive |

| | | | | | |
|---|---|---|---|---|---|
| hCoVHKU1 | orf1ab | MISAVAAGLEFTDEN | 2730 | 2744 | Positive |
| hCoVHKU1 | orf1ab | MKRYCTVQGPPGTGK | 5658 | 5672 | Positive |
| hCoVHKU1 | orf1ab | MLCNRYNLCYDIGNP | 6308 | 6322 | Positive |
| hCoVHKU1 | orf1ab | MQYGYVDCFNDNCDF | 358 | 372 | Positive |
| hCoVHKU1 | orf1ab | MWYFGSNLEEDVLLF | 3788 | 3802 | Positive |
| hCoVHKU1 | orf1ab | NAAITDYNYYKYNLP | 4900 | 4914 | Positive |
| hCoVHKU1 | orf1ab | NADLYKLMSCFAFWT | 7060 | 7074 | Positive |
| hCoVHKU1 | orf1ab | NALNSCKEHPVVGYQ | 872 | 886 | Positive |
| hCoVHKU1 | orf1ab | NAPNCDVSDVTKLYL | 5436 | 5450 | Positive |
| hCoVHKU1 | orf1ab | NASLYTSLVPHTRYS | 3014 | 3028 | Positive |
| hCoVHKU1 | orf1ab | NAVMQNNELMPHKLK | 4206 | 4220 | Positive |
| hCoVHKU1 | orf1ab | NCHSFKPGNTFITVE | 2530 | 2544 | Positive |
| hCoVHKU1 | orf1ab | NCVKMLCDHAGTGMA | 4360 | 4374 | Positive |
| hCoVHKU1 | orf1ab | NCYRCPKEIVETVSA | 5824 | 5838 | Positive |
| hCoVHKU1 | orf1ab | NCYVNVACLMLQHIN | 1756 | 1770 | Positive |
| hCoVHKU1 | orf1ab | NDCSILCEILCEYAD | 4594 | 4608 | Positive |
| hCoVHKU1 | orf1ab | NDITNGPHEFCSQHT | 5256 | 5270 | Positive |
| hCoVHKU1 | orf1ab | NDLGTQILDSYSVIL | 5346 | 5360 | Positive |
| hCoVHKU1 | orf1ab | NDVCSTEDFNVWAMA | 3556 | 3570 | Positive |
| hCoVHKU1 | orf1ab | NDYYRSMPGTFCGRD | 3082 | 3096 | Positive |
| hCoVHKU1 | orf1ab | NELMPHKLKIQVVNS | 4212 | 4226 | Positive |
| hCoVHKU1 | orf1ab | NENLAVNLNICEPVL | 6022 | 6036 | Positive |
| hCoVHKU1 | orf1ab | NEVANAVMQNNELMP | 4202 | 4216 | Positive |
| hCoVHKU1 | orf1ab | NEVSINTSCRLLQRV | 6288 | 6302 | Positive |
| hCoVHKU1 | orf1ab | NFITLPLDKIQNQTL | 5970 | 5984 | Positive |
| hCoVHKU1 | orf1ab | NFKDGLCMFWNCNVD | 6358 | 6372 | Positive |
| hCoVHKU1 | orf1ab | NFKLVGHDICAQLND | 2008 | 2022 | Positive |
| hCoVHKU1 | orf1ab | NFLMLFVFQVYPICA | 3162 | 3176 | Positive |
| hCoVHKU1 | orf1ab | NFLNRVRGTSVNARL | 4462 | 4476 | Positive |
| hCoVHKU1 | orf1ab | NFNEKPVVMEIPSLM | 814 | 828 | Positive |
| hCoVHKU1 | orf1ab | NGHMLHGGGVAKAIA | 1388 | 1402 | Positive |
| hCoVHKU1 | orf1ab | NGISVTQTMFDVYVD | 2656 | 2670 | Positive |
| hCoVHKU1 | orf1ab | NGPFFSFEQSHNNCY | 1744 | 1758 | Positive |
| hCoVHKU1 | orf1ab | NGPHEFCSQHTMLVK | 5260 | 5274 | Positive |
| hCoVHKU1 | orf1ab | NHKVIGGLHLLIGLF | 6778 | 6792 | Positive |
| hCoVHKU1 | orf1ab | NHNNGNDVLYQPPTA | 3312 | 3326 | Positive |
| hCoVHKU1 | orf1ab | NHVKIDCRCILGQEW | 56 | 70 | Positive |
| hCoVHKU1 | orf1ab | NKEKTYYELTKSCGV | 4538 | 4552 | Positive |
| hCoVHKU1 | orf1ab | NKHAFHTNPFTRTVF | 6404 | 6418 | Positive |
| hCoVHKU1 | orf1ab | NKIASCKWTEVDDYV | 5492 | 5506 | Positive |
| hCoVHKU1 | orf1ab | NKLNENVVYLFDEAG | 892 | 906 | Positive |
| hCoVHKU1 | orf1ab | NKMVNWFNTMLDASA | 622 | 636 | Positive |
| hCoVHKU1 | orf1ab | NKNYVFTGYHFTSTG | 5562 | 5576 | Positive |
| hCoVHKU1 | orf1ab | NKQEASVPILTTPFS | 2820 | 2834 | Positive |
| hCoVHKU1 | orf1ab | NKQLTDISVDSNWPL | 4180 | 4194 | Positive |
| hCoVHKU1 | orf1ab | NKWQWQEAWYEFRAG | 1774 | 1788 | Positive |
| hCoVHKU1 | orf1ab | NKYRYYSGKMDTAAY | 3282 | 3296 | Positive |
| hCoVHKU1 | orf1ab | NLAVNLNICEPVLTY | 6024 | 6038 | Positive |
| hCoVHKU1 | orf1ab | NLEYPIISNEVSINT | 6280 | 6294 | Positive |

| | | | | | |
|---|---|---|---|---|---|
| hCoVHKU1 | orf1ab | NLFIAKYGLEDYAFD | 6756 | 6770 | Positive |
| hCoVHKU1 | orf1ab | NLFKDCSKSCLGYHP | 5992 | 6006 | Positive |
| hCoVHKU1 | orf1ab | NLGGAVCSKHAEEYC | 6460 | 6474 | Positive |
| hCoVHKU1 | orf1ab | NLGGNCAGNVIGNDA | 6710 | 6724 | Positive |
| hCoVHKU1 | orf1ab | NLKFNKWQWQEAWYE | 1770 | 1784 | Positive |
| hCoVHKU1 | orf1ab | NLKLLGIGGVPVIEV | 3906 | 3920 | Positive |
| hCoVHKU1 | orf1ab | NLKPMPFFYYSDTPC | 6420 | 6434 | Positive |
| hCoVHKU1 | orf1ab | NLLHSKVKVVPNLYV | 2602 | 2616 | Positive |
| hCoVHKU1 | orf1ab | NLNICEPVLTYSRLI | 6028 | 6042 | Positive |
| hCoVHKU1 | orf1ab | NLPGCNGGSLYVNKH | 6392 | 6406 | Positive |
| hCoVHKU1 | orf1ab | NLPTGCMMNVAKYTQ | 6918 | 6932 | Positive |
| hCoVHKU1 | orf1ab | NLPTMVDIKQLLFVL | 4912 | 4926 | Positive |
| hCoVHKU1 | orf1ab | NLRPDQLNDLVYSLI | 7144 | 7158 | Positive |
| hCoVHKU1 | orf1ab | NLTQTFTSMLTNYFL | 1950 | 1964 | Positive |
| hCoVHKU1 | orf1ab | NMLLKNIDLFVKRRA | 520 | 534 | Positive |
| hCoVHKU1 | orf1ab | NMNEPDYSALLCRVT | 3382 | 3396 | Positive |
| hCoVHKU1 | orf1ab | NMQLFESLNFITLPL | 5962 | 5976 | Positive |
| hCoVHKU1 | orf1ab | NMYLKSAVMQSVGAC | 5376 | 5390 | Positive |
| hCoVHKU1 | orf1ab | NNLVPTYLKSDNIVA | 2746 | 2760 | Positive |
| hCoVHKU1 | orf1ab | NNTSFPTNIAVELFT | 6554 | 6568 | Positive |
| hCoVHKU1 | orf1ab | NNVANYQHIGMKRYC | 5648 | 5662 | Positive |
| hCoVHKU1 | orf1ab | NQALNSILDNAVKGC | 4114 | 4128 | Positive |
| hCoVHKU1 | orf1ab | NQDLYGQWYDFGDFI | 4666 | 4680 | Positive |
| hCoVHKU1 | orf1ab | NQLTADLQHKLKKAC | 2794 | 2808 | Positive |
| hCoVHKU1 | orf1ab | NRAGIGLYYKVNCCR | 4498 | 4512 | Positive |
| hCoVHKU1 | orf1ab | NRALLNTVIFADTLV | 4640 | 4654 | Positive |
| hCoVHKU1 | orf1ab | NSDYASKGYIANISV | 5220 | 5234 | Positive |
| hCoVHKU1 | orf1ab | NSGSVNQQQIKQLEK | 4040 | 4054 | Positive |
| hCoVHKU1 | orf1ab | NSILDNAVKGCVPLS | 4118 | 4132 | Positive |
| hCoVHKU1 | orf1ab | NSSIKVVKSLSLVDV | 2148 | 2162 | Positive |
| hCoVHKU1 | orf1ab | NSSVIQEVLLLRHDI | 1560 | 1574 | Positive |
| hCoVHKU1 | orf1ab | NSTTWNGGAYSLFDM | 7116 | 7130 | Positive |
| hCoVHKU1 | orf1ab | NSWCRQLLLNRGDYN | 506 | 520 | Positive |
| hCoVHKU1 | orf1ab | NTNRAGIGLYYKVNC | 4496 | 4510 | Positive |
| hCoVHKU1 | orf1ab | NTPLSKDLPDDVVAA | 1890 | 1904 | Positive |
| hCoVHKU1 | orf1ab | NTTTLAVPVNMRVLH | 6938 | 6952 | Positive |
| hCoVHKU1 | orf1ab | NTVDCDNDRCIIHCA | 4750 | 4764 | Positive |
| hCoVHKU1 | orf1ab | NVACLMLQHINLKFN | 1760 | 1774 | Positive |
| hCoVHKU1 | orf1ab | NVCSLMACNGHKIED | 5156 | 5170 | Positive |
| hCoVHKU1 | orf1ab | NVDLCDCGFVMQYGY | 348 | 362 | Positive |
| hCoVHKU1 | orf1ab | NVFMSESKCWVENDI | 5244 | 5258 | Positive |
| hCoVHKU1 | orf1ab | NVGHYDGRTGELPCA | 6520 | 6534 | Positive |
| hCoVHKU1 | orf1ab | NVPLERVSLWNYGKP | 6902 | 6916 | Positive |
| hCoVHKU1 | orf1ab | NVPTKVLRHGFHVLH | 2942 | 2956 | Positive |
| hCoVHKU1 | orf1ab | NVSKDGFFTYICHLI | 7026 | 7040 | Positive |
| hCoVHKU1 | orf1ab | NVVTSVMESQISTKE | 2112 | 2126 | Positive |
| hCoVHKU1 | orf1ab | NWFNTMLDASAPATG | 626 | 640 | Positive |
| hCoVHKU1 | orf1ab | NWLSVNIFYFTDVPY | 3826 | 3840 | Positive |
| hCoVHKU1 | orf1ab | NWNLEYPIISNEVSI | 6278 | 6292 | Positive |

| | | | | | |
|---|---|---|---|---|---|
| hCoVHKU1 | orf1ab | NYGKPINLPTGCMMN | 6912 | 6926 | Positive |
| hCoVHKU1 | orf1ab | NYQHIGMKRYCTVQG | 5652 | 5666 | Positive |
| hCoVHKU1 | orf1ab | PDAAEELASPMKSDE | 22 | 36 | Positive |
| hCoVHKU1 | orf1ab | PFDCHWDLIISDMYD | 7002 | 7016 | Positive |
| hCoVHKU1 | orf1ab | PGETFTVLAAYNGRP | 3442 | 3456 | Positive |
| hCoVHKU1 | orf1ab | PHEFCSQHTMLVKID | 5262 | 5276 | Positive |
| hCoVHKU1 | orf1ab | PHIQISYNDFYASGC | 2972 | 2986 | Positive |
| hCoVHKU1 | orf1ab | PINLPTGCMMNVAKY | 6916 | 6930 | Positive |
| hCoVHKU1 | orf1ab | PLERVSLWNYGKPIN | 6904 | 6918 | Positive |
| hCoVHKU1 | orf1ab | PLGIKDPILYVLTHD | 4420 | 4434 | Positive |
| hCoVHKU1 | orf1ab | PLSKDLPDDVVAANM | 1892 | 1906 | Positive |
| hCoVHKU1 | orf1ab | PPGEKFKHLIPLMSK | 6128 | 6142 | Positive |
| hCoVHKU1 | orf1ab | PSDATDFIRVVLKQA | 1808 | 1822 | Positive |
| hCoVHKU1 | orf1ab | PSDVYQQLAGVKLQS | 3624 | 3638 | Positive |
| hCoVHKU1 | orf1ab | PTGCMMNVAKYTQLC | 6920 | 6934 | Positive |
| hCoVHKU1 | orf1ab | PTITHTNCWLRSVLL | 1154 | 1168 | Positive |
| hCoVHKU1 | orf1ab | PTLTQMNLKYAISAK | 4990 | 5004 | Positive |
| hCoVHKU1 | orf1ab | PTMVDIKQLLFVLEV | 4914 | 4928 | Positive |
| hCoVHKU1 | orf1ab | PVIEVSQIQSKLTDV | 3916 | 3930 | Positive |
| hCoVHKU1 | orf1ab | PVNPTDASHYVVTDI | 2556 | 2570 | Positive |
| hCoVHKU1 | orf1ab | PYIDDFNKIASCKWT | 5486 | 5500 | Positive |
| hCoVHKU1 | orf1ab | QDADGINKQLTDISV | 4174 | 4188 | Positive |
| hCoVHKU1 | orf1ab | QGRQILGSCTFEDEL | 3608 | 3622 | Positive |
| hCoVHKU1 | orf1ab | QHIGMKRYCTVQGPP | 5654 | 5668 | Positive |
| hCoVHKU1 | orf1ab | QNNVFMSESKCWVEN | 5242 | 5256 | Positive |
| hCoVHKU1 | orf1ab | QQVLYYQNNVFMSES | 5236 | 5250 | Positive |
| hCoVHKU1 | orf1ab | QQWGYTGSLTSNHDI | 6232 | 6246 | Positive |
| hCoVHKU1 | orf1ab | QRVYDDVDASLFVDI | 2586 | 2600 | Positive |
| hCoVHKU1 | orf1ab | QSCTGSPYIDDFNKI | 5480 | 5494 | Positive |
| hCoVHKU1 | orf1ab | QWYESTFGSVYYHNS | 2908 | 2922 | Positive |
| hCoVHKU1 | orf1ab | RCNLGGAVCSKHAEE | 6458 | 6472 | Positive |
| hCoVHKU1 | orf1ab | RDAQVVQLPVKDYVQ | 3520 | 3534 | Positive |
| hCoVHKU1 | orf1ab | RDDNQTLLVPKIFKA | 2210 | 2224 | Positive |
| hCoVHKU1 | orf1ab | RDFIDMDDNLFIAKY | 6748 | 6762 | Positive |
| hCoVHKU1 | orf1ab | RDIFVHEDLHVVEVL | 78 | 92 | Positive |
| hCoVHKU1 | orf1ab | RDISVNDLCFANKFF | 2890 | 2904 | Positive |
| hCoVHKU1 | orf1ab | RDSRFVMRLQTIATI | 278 | 292 | Positive |
| hCoVHKU1 | orf1ab | RDTRKEIFVGDSLVN | 7166 | 7180 | Positive |
| hCoVHKU1 | orf1ab | RKLDNQALNSILDNA | 4110 | 4124 | Positive |
| hCoVHKU1 | orf1ab | RLFYDRDGQRVYDDV | 2578 | 2592 | Positive |
| hCoVHKU1 | orf1ab | RLKLFAAETQKATEE | 5514 | 5528 | Positive |
| hCoVHKU1 | orf1ab | RLYYEALSFEEQNEI | 4966 | 4980 | Positive |
| hCoVHKU1 | orf1ab | RPDQLNDLVYSLIER | 7146 | 7160 | Positive |
| hCoVHKU1 | orf1ab | RPIDFFSLTASSIFG | 3110 | 3124 | Positive |
| hCoVHKU1 | orf1ab | RQILGSCTFEDELAP | 3610 | 3624 | Positive |
| hCoVHKU1 | orf1ab | RRADFACKFAVCGDG | 532 | 546 | Positive |
| hCoVHKU1 | orf1ab | RTVAGVSILSTMTGR | 5008 | 5022 | Positive |
| hCoVHKU1 | orf1ab | RYCTVQGPPGTGKSH | 5660 | 5674 | Positive |
| hCoVHKU1 | orf1ab | SDDGVVCYNSDYASK | 5212 | 5226 | Positive |

| | | | | | |
|---|---|---|---|---|---|
| hCoVHKU1 | orf1ab | SDDYVQDSTVLQALQ | 4002 | 4016 | Positive |
| hCoVHKU1 | orf1ab | SDFYADKILYQYENL | 1694 | 1708 | Positive |
| hCoVHKU1 | orf1ab | SDFYLPNISVFPIFV | 2308 | 2322 | Positive |
| hCoVHKU1 | orf1ab | SDLERDFIDMDDNLF | 6744 | 6758 | Positive |
| hCoVHKU1 | orf1ab | SDLIISNFKFDDYDF | 178 | 192 | Positive |
| hCoVHKU1 | orf1ab | SDMNCNIPTQCYYNN | 4228 | 4242 | Positive |
| hCoVHKU1 | orf1ab | SDMYDPLTKNIGDYN | 7012 | 7026 | Positive |
| hCoVHKU1 | orf1ab | SDREVILCWETGKVK | 5544 | 5558 | Positive |
| hCoVHKU1 | orf1ab | SDSLVTYFGDCMTLP | 6988 | 7002 | Positive |
| hCoVHKU1 | orf1ab | SDSVQCYTPHIQISY | 2964 | 2978 | Positive |
| hCoVHKU1 | orf1ab | SDTPCVYVDGLESKQ | 6430 | 6444 | Positive |
| hCoVHKU1 | orf1ab | SDVAYNRYLSLYNKY | 3270 | 3284 | Positive |
| hCoVHKU1 | orf1ab | SDYLLDLSDSVVFIT | 6158 | 6172 | Positive |
| hCoVHKU1 | orf1ab | SEESQGNVVTSVMES | 2106 | 2120 | Positive |
| hCoVHKU1 | orf1ab | SEISVIMHLQWIVMY | 3192 | 3206 | Positive |
| hCoVHKU1 | orf1ab | SELTNGVYYRATTTY | 5588 | 5602 | Positive |
| hCoVHKU1 | orf1ab | SEVKAQSSGVIPENP | 396 | 410 | Positive |
| hCoVHKU1 | orf1ab | SFVDYLLTTIPKAIV | 1196 | 1210 | Positive |
| hCoVHKU1 | orf1ab | SHFDVDRKSFNNFVN | 2674 | 2688 | Positive |
| hCoVHKU1 | orf1ab | SHGDRFYRLANECAQ | 5100 | 5114 | Positive |
| hCoVHKU1 | orf1ab | SIETLLAAIKRLYMG | 3592 | 3606 | Positive |
| hCoVHKU1 | orf1ab | SILVDNDLNPFVSDS | 6976 | 6990 | Positive |
| hCoVHKU1 | orf1ab | SKLWQYCSVLHNEIL | 3950 | 3964 | Positive |
| hCoVHKU1 | orf1ab | SLAIDAYPLVHHENE | 5314 | 5328 | Positive |
| hCoVHKU1 | orf1ab | SLEPRHFNSITKIMC | 5800 | 5814 | Positive |
| hCoVHKU1 | orf1ab | SLFDDNFCAFCTPKK | 1284 | 1298 | Positive |
| hCoVHKU1 | orf1ab | SLFDMTKFSLKLAGT | 7126 | 7140 | Positive |
| hCoVHKU1 | orf1ab | SLSSYEYCQPPKSVE | 756 | 770 | Positive |
| hCoVHKU1 | orf1ab | SLVDVWDMYLTGCDY | 2158 | 2172 | Positive |
| hCoVHKU1 | orf1ab | SMDCPIVVAVMDEDI | 2922 | 2936 | Positive |
| hCoVHKU1 | orf1ab | SMSSFELPQLYGLCI | 1346 | 1360 | Positive |
| hCoVHKU1 | orf1ab | SMTLNGLWLDDKVYC | 3358 | 3372 | Positive |
| hCoVHKU1 | orf1ab | SMTTSTTNFGEDFLG | 138 | 152 | Positive |
| hCoVHKU1 | orf1ab | SMTYCRVGACEYAEE | 3054 | 3068 | Positive |
| hCoVHKU1 | orf1ab | SNHDIICNVHKGAHV | 6242 | 6256 | Positive |
| hCoVHKU1 | orf1ab | SNLIRDIFVHEDLHV | 74 | 88 | Positive |
| hCoVHKU1 | orf1ab | SNLLIQEFLQYDSSI | 6798 | 6812 | Positive |
| hCoVHKU1 | orf1ab | SSFEPRSDLERDFID | 6738 | 6752 | Positive |
| hCoVHKU1 | orf1ab | SSIHSYFITDQECGS | 6810 | 6824 | Positive |
| hCoVHKU1 | orf1ab | STDTVNHDSFNLYGY | 416 | 430 | Positive |
| hCoVHKU1 | orf1ab | STDVQLRAFDICNTN | 4484 | 4498 | Positive |
| hCoVHKU1 | orf1ab | STEDFNVWAMANGFS | 3560 | 3574 | Positive |
| hCoVHKU1 | orf1ab | STFGSVYYHNSMDCP | 2912 | 2926 | Positive |
| hCoVHKU1 | orf1ab | SVEKICIIDNMYMGK | 768 | 782 | Positive |
| hCoVHKU1 | orf1ab | SVIVEDDVIENVKSS | 742 | 756 | Positive |
| hCoVHKU1 | orf1ab | SVLSVDSVSEESQGN | 2098 | 2112 | Positive |
| hCoVHKU1 | orf1ab | SVVVNGPFFSFEQSH | 1740 | 1754 | Positive |
| hCoVHKU1 | orf1ab | SVYSVPLVFQNNVAN | 5638 | 5652 | Positive |
| hCoVHKU1 | orf1ab | TCDGLKFTEESFYKN | 5362 | 5376 | Positive |

| | | | | | |
|---|---|---|---|---|---|
| hCoVHKU1 | orf1ab | TDASHYVVTDIKQVG | 2560 | 2574 | Positive |
| hCoVHKU1 | orf1ab | TDAYKLFLSGDDCFV | 1544 | 1558 | Positive |
| hCoVHKU1 | orf1ab | TDFIRVVLKQADLSG | 1812 | 1826 | Positive |
| hCoVHKU1 | orf1ab | TDIKQVGCMMRLFYD | 2568 | 2582 | Positive |
| hCoVHKU1 | orf1ab | TDISVDSNWPLVIIA | 4184 | 4198 | Positive |
| hCoVHKU1 | orf1ab | TDIVVVDEVSMLTNY | 5752 | 5766 | Positive |
| hCoVHKU1 | orf1ab | TDLKFIENLNILFDG | 6614 | 6628 | Positive |
| hCoVHKU1 | orf1ab | TDQECGSSKSVCTVI | 6818 | 6832 | Positive |
| hCoVHKU1 | orf1ab | TDSLCSSHYWSPQGN | 6696 | 6710 | Positive |
| hCoVHKU1 | orf1ab | TDVPYIKLILLSYLF | 3836 | 3850 | Positive |
| hCoVHKU1 | orf1ab | TEDVVDGDVVIREPV | 302 | 316 | Positive |
| hCoVHKU1 | orf1ab | TFEETSLTTFMITKD | 3246 | 3260 | Positive |
| hCoVHKU1 | orf1ab | TGCDYVVWVANELSR | 2168 | 2182 | Positive |
| hCoVHKU1 | orf1ab | TGDNDDEDVVTGDND | 952 | 966 | Positive |
| hCoVHKU1 | orf1ab | TGDNDDQIVVTGDDV | 1092 | 1106 | Positive |
| hCoVHKU1 | orf1ab | TGDNNDEEIVTGDND | 1082 | 1096 | Positive |
| hCoVHKU1 | orf1ab | TGFCVKHQWNCFNCH | 2518 | 2532 | Positive |
| hCoVHKU1 | orf1ab | TGKSHLAIGLAVYYY | 5670 | 5684 | Positive |
| hCoVHKU1 | orf1ab | TGYYGCWRHSYTCDY | 6208 | 6222 | Positive |
| hCoVHKU1 | orf1ab | THDVCQVCGFWRDGS | 4432 | 4446 | Positive |
| hCoVHKU1 | orf1ab | THLKCGSPYQHYDAC | 1916 | 1930 | Positive |
| hCoVHKU1 | orf1ab | THMDDFSIKSIYNVD | 336 | 350 | Positive |
| hCoVHKU1 | orf1ab | TIQEIVSDREVILCW | 5538 | 5552 | Positive |
| hCoVHKU1 | orf1ab | TKLYLGGMSYYCENH | 5446 | 5460 | Positive |
| hCoVHKU1 | orf1ab | TLISMWYFGSNLEED | 3784 | 3798 | Positive |
| hCoVHKU1 | orf1ab | TLPFDCHWDLIISDM | 7000 | 7014 | Positive |
| hCoVHKU1 | orf1ab | TMHSINHDIFSLMFL | 3764 | 3778 | Positive |
| hCoVHKU1 | orf1ab | TMHWLIRFIVFVANM | 2436 | 2450 | Positive |
| hCoVHKU1 | orf1ab | TMLDLCYALRHFDRN | 4580 | 4594 | Positive |
| hCoVHKU1 | orf1ab | TMLVKIDGDYVYLPY | 5270 | 5284 | Positive |
| hCoVHKU1 | orf1ab | TMYIIPVLCTLFYVN | 3704 | 3718 | Positive |
| hCoVHKU1 | orf1ab | TPKYTFGNVKPGETF | 3432 | 3446 | Positive |
| hCoVHKU1 | orf1ab | TSHFCNGSFICELCH | 2350 | 2364 | Positive |
| hCoVHKU1 | orf1ab | TTAGFTFWVYKNFDF | 6484 | 6498 | Positive |
| hCoVHKU1 | orf1ab | TTHESSSAVNIQQIY | 5864 | 5878 | Positive |
| hCoVHKU1 | orf1ab | TVEAAIELSKELKRP | 2542 | 2556 | Positive |
| hCoVHKU1 | orf1ab | TWLPDLFMLETMHWL | 2426 | 2440 | Positive |
| hCoVHKU1 | orf1ab | TYYELTKSCGVVAEH | 4542 | 4556 | Positive |
| hCoVHKU1 | orf1ab | VAAGLEFTDENYNNL | 2734 | 2748 | Positive |
| hCoVHKU1 | orf1ab | VADSYYSYMMPMLTM | 4690 | 4704 | Positive |
| hCoVHKU1 | orf1ab | VAQPTEDVVDGDVVI | 298 | 312 | Positive |
| hCoVHKU1 | orf1ab | VATGDVVLASDDLYV | 2042 | 2056 | Positive |
| hCoVHKU1 | orf1ab | VAVADSYYSYMMPML | 4688 | 4702 | Positive |
| hCoVHKU1 | orf1ab | VAVIGDEQIDGKFVT | 1314 | 1328 | Positive |
| hCoVHKU1 | orf1ab | VDCDNDRCIIHCANF | 4752 | 4766 | Positive |
| hCoVHKU1 | orf1ab | VDGDVVIREPVHLLS | 306 | 320 | Positive |
| hCoVHKU1 | orf1ab | VDGLKYTKIMKDDGN | 4256 | 4270 | Positive |
| hCoVHKU1 | orf1ab | VDGVNFKSISLTVGE | 1658 | 1672 | Positive |
| hCoVHKU1 | orf1ab | VDLPFVEYKVTVWPV | 2028 | 2042 | Positive |

| | | | | | |
|---|---|---|---|---|---|
| hCoVHKU1 | orf1ab | VDSVSEESQGNVVTS | 2102 | 2116 | Positive |
| hCoVHKU1 | orf1ab | VDVWDMYLTGCDYVV | 2160 | 2174 | Positive |
| hCoVHKU1 | orf1ab | VEDTINDDEDVVTGD | 940 | 954 | Positive |
| hCoVHKU1 | orf1ab | VELFTKRSIRHHPEL | 6564 | 6578 | Positive |
| hCoVHKU1 | orf1ab | VFAIFITMHSINHDI | 3758 | 3772 | Positive |
| hCoVHKU1 | orf1ab | VGACEYAEEGICFNF | 3060 | 3074 | Positive |
| hCoVHKU1 | orf1ab | VGHYTHLKCGSPYQH | 1912 | 1926 | Positive |
| hCoVHKU1 | orf1ab | VHEDLHVVEVLTKTA | 82 | 96 | Positive |
| hCoVHKU1 | orf1ab | VICSSSNMNEPDYSA | 3376 | 3390 | Positive |
| hCoVHKU1 | orf1ab | VIWFCHDEASLNSLT | 2070 | 2084 | Positive |
| hCoVHKU1 | orf1ab | VKETALITHALYLDY | 474 | 488 | Positive |
| hCoVHKU1 | orf1ab | VLARKHEFCCSHGDR | 5090 | 5104 | Positive |
| hCoVHKU1 | orf1ab | VLYYQNNVFMSESKC | 5238 | 5252 | Positive |
| hCoVHKU1 | orf1ab | VNHDSFNLYGYSVTP | 420 | 434 | Positive |
| hCoVHKU1 | orf1ab | VNSGSDMNCNIPTQC | 4224 | 4238 | Positive |
| hCoVHKU1 | orf1ab | VQSNLIRDIFVHEDL | 72 | 86 | Positive |
| hCoVHKU1 | orf1ab | VRGWVGFDVEGAHAT | 6070 | 6084 | Positive |
| hCoVHKU1 | orf1ab | VSNNKEDFDIIQKCQ | 1504 | 1518 | Positive |
| hCoVHKU1 | orf1ab | VSNSSVIQEVLLLRH | 1558 | 1572 | Positive |
| hCoVHKU1 | orf1ab | VTSVMESQISTKEVK | 2114 | 2128 | Positive |
| hCoVHKU1 | orf1ab | VVAEHDFFTFDIDGS | 4552 | 4566 | Positive |
| hCoVHKU1 | orf1ab | VVAVMDEDIGSTMFN | 2928 | 2942 | Positive |
| hCoVHKU1 | orf1ab | VVDEVSMLTNYELSV | 5756 | 5770 | Positive |
| hCoVHKU1 | orf1ab | VVESDADRANFLNAV | 2618 | 2632 | Positive |
| hCoVHKU1 | orf1ab | VVEVLTKTAVKSGTA | 88 | 102 | Positive |
| hCoVHKU1 | orf1ab | VVIGTTKFYGGWDDM | 5040 | 5054 | Positive |
| hCoVHKU1 | orf1ab | VVIREPVHLLSADAI | 310 | 324 | Positive |
| hCoVHKU1 | orf1ab | VVLELDPPCKFSIQD | 4272 | 4286 | Positive |
| hCoVHKU1 | orf1ab | VVNGPFFSFEQSHNN | 1742 | 1756 | Positive |
| hCoVHKU1 | orf1ab | VVRDSRFVMRLQTIA | 276 | 290 | Positive |
| hCoVHKU1 | orf1ab | VVTAMYKVVGFIRHI | 2462 | 2476 | Positive |
| hCoVHKU1 | orf1ab | VVTGDNDDEDVVTGD | 950 | 964 | Positive |
| hCoVHKU1 | orf1ab | VVTGDNNDEEIVTGD | 1080 | 1094 | Positive |
| hCoVHKU1 | orf1ab | VVVVESDADRANFLN | 2616 | 2630 | Positive |
| hCoVHKU1 | orf1ab | WFIDAFNQLTADLQH | 2788 | 2802 | Positive |
| hCoVHKU1 | orf1ab | WHVQSNLIRDIFVHE | 70 | 84 | Positive |
| hCoVHKU1 | orf1ab | YDSSIHSYFITDQEC | 6808 | 6822 | Positive |
| hCoVHKU1 | orf1ab | YELTKSCGVVAEHDF | 4544 | 4558 | Positive |
| hCoVHKU1 | orf1ab | YESTFGSVYYHNSMD | 2910 | 2924 | Positive |
| hCoVHKU1 | orf1ab | YFEIYDGGCIPASQV | 4932 | 4946 | Positive |
| hCoVHKU1 | orf1ab | YNSRTGYYGCWRHSY | 6204 | 6218 | Positive |
| hCoVHKU1 | orf1ab | YPLVHHENEEYQKVF | 5320 | 5334 | Positive |
| hCoVHKU1 | orf1ab | YSKLFITKDEAIKRV | 6056 | 6070 | Positive |
| hCoVHKU1 | orf1ab | YWSPRPGLWIPIIKS | 440 | 454 | Positive |
| hCoVHKU1 | orf1ab | YYEALSFEEQNEIYA | 4968 | 4982 | Positive |
| hCoVHKU1 | orf1ab | ADAKNSGSVNQQQIK | 4036 | 4050 | Negative |
| hCoVHKU1 | orf1ab | ADGINKQLTDISVDS | 4176 | 4190 | Negative |
| hCoVHKU1 | orf1ab | ADVSLTYLLGVVDKQ | 1486 | 1500 | Negative |
| hCoVHKU1 | orf1ab | AIELSKELKRPVNPT | 2546 | 2560 | Negative |

| | | | | | |
|---|---|---|---|---|---|
| hCoVHKU1 | orf1ab | AIYDCFCKSVNWNLE | 6268 | 6282 | Negative |
| hCoVHKU1 | orf1ab | AKYTQLCQYLNTTTL | 6928 | 6942 | Negative |
| hCoVHKU1 | orf1ab | ALLDLRTCCFSVAAI | 4840 | 4854 | Negative |
| hCoVHKU1 | orf1ab | ALQSEFVNMASFVEY | 4014 | 4028 | Negative |
| hCoVHKU1 | orf1ab | AMESVGFVYDNHVKI | 46 | 60 | Negative |
| hCoVHKU1 | orf1ab | ANECIERLKLFAAET | 5508 | 5522 | Negative |
| hCoVHKU1 | orf1ab | AVMDEDIGSTMFNVP | 2930 | 2944 | Negative |
| hCoVHKU1 | orf1ab | CLKCGFSFDLNGLDA | 1238 | 1252 | Negative |
| hCoVHKU1 | orf1ab | CSSSNMNEPDYSALL | 3378 | 3392 | Negative |
| hCoVHKU1 | orf1ab | CTFEDELAPSDVYQQ | 3616 | 3630 | Negative |
| hCoVHKU1 | orf1ab | DCFNDNCDFYGWVSG | 364 | 378 | Negative |
| hCoVHKU1 | orf1ab | DCGIKQESRVGVDAV | 1836 | 1850 | Negative |
| hCoVHKU1 | orf1ab | DDGNKLDKFFVVKRT | 4518 | 4532 | Negative |
| hCoVHKU1 | orf1ab | DDLYVKRYFKGCETF | 2052 | 2066 | Negative |
| hCoVHKU1 | orf1ab | DDMLRHLIKDVDNPV | 5052 | 5066 | Negative |
| hCoVHKU1 | orf1ab | DDNLFIAKYGLEDYA | 6754 | 6768 | Negative |
| hCoVHKU1 | orf1ab | DDVDASLFVDINNLL | 2590 | 2604 | Negative |
| hCoVHKU1 | orf1ab | DEAIKRVRGWVGFDV | 6064 | 6078 | Negative |
| hCoVHKU1 | orf1ab | DGDYVYLPYPDPSRI | 5276 | 5290 | Negative |
| hCoVHKU1 | orf1ab | DGNAAITDYNYYKYN | 4898 | 4912 | Negative |
| hCoVHKU1 | orf1ab | DIGNPKGLACVKDYE | 6318 | 6332 | Negative |
| hCoVHKU1 | orf1ab | DKQVILVSNNKEDFD | 1498 | 1512 | Negative |
| hCoVHKU1 | orf1ab | DLGVLIQNGAKHVQG | 2762 | 2776 | Negative |
| hCoVHKU1 | orf1ab | DNCDFYGWVSGNMMD | 368 | 382 | Negative |
| hCoVHKU1 | orf1ab | DNDLNPFVSDSLVTY | 6980 | 6994 | Negative |
| hCoVHKU1 | orf1ab | DPKKTYLDYIQQGGV | 4342 | 4356 | Negative |
| hCoVHKU1 | orf1ab | DSDVETRFITKSMIS | 2718 | 2732 | Negative |
| hCoVHKU1 | orf1ab | DSSEVKAQSSGVIPE | 394 | 408 | Negative |
| hCoVHKU1 | orf1ab | DTGALEAFRKARNGV | 6630 | 6644 | Negative |
| hCoVHKU1 | orf1ab | DVTKLYLGGMSYYCE | 5444 | 5458 | Negative |
| hCoVHKU1 | orf1ab | DVYVDTFMSHFDVDR | 2666 | 2680 | Negative |
| hCoVHKU1 | orf1ab | DYLLSKMTSLPKDWR | 1582 | 1596 | Negative |
| hCoVHKU1 | orf1ab | DYNVSKDGFFTYICH | 7024 | 7038 | Negative |
| hCoVHKU1 | orf1ab | DYVKDSLFCSSTYGV | 6596 | 6610 | Negative |
| hCoVHKU1 | orf1ab | DYVLANECIERLKLF | 5504 | 5518 | Negative |
| hCoVHKU1 | orf1ab | DYVPLRSATCITRCN | 6446 | 6460 | Negative |
| hCoVHKU1 | orf1ab | DYVSLVKLIVELVIG | 2390 | 2404 | Negative |
| hCoVHKU1 | orf1ab | DYVYLPYPDPSRILG | 5278 | 5292 | Negative |
| hCoVHKU1 | orf1ab | EAVEDTIDGVVEDTI | 930 | 944 | Negative |
| hCoVHKU1 | orf1ab | EFVNMASFVEYEVAK | 4018 | 4032 | Negative |
| hCoVHKU1 | orf1ab | EGFLIGINYLGKSSF | 7086 | 7100 | Negative |
| hCoVHKU1 | orf1ab | EGIVRIVRTRSMTYC | 3044 | 3058 | Negative |
| hCoVHKU1 | orf1ab | EGVQLEKVLDTFVGC | 2696 | 2710 | Negative |
| hCoVHKU1 | orf1ab | EILSTSDLSVAFDKL | 3962 | 3976 | Negative |
| hCoVHKU1 | orf1ab | EIVMCGGCYYVKPGG | 5118 | 5132 | Negative |
| hCoVHKU1 | orf1ab | ELELICDCGIKQESR | 1830 | 1844 | Negative |
| hCoVHKU1 | orf1ab | ELFNKYFKYWGMKYH | 4734 | 4748 | Negative |
| hCoVHKU1 | orf1ab | ELICDCGIKQESRVG | 1832 | 1846 | Negative |
| hCoVHKU1 | orf1ab | ELKRPVNPTDASHYV | 2552 | 2566 | Negative |

| | | | | | |
|---|---|---|---|---|---|
| hCoVHKU1 | orf1ab | ELPCAIMNDKVVVKI | 6530 | 6544 | Negative |
| hCoVHKU1 | orf1ab | ELRYMNANGLRPPRN | 3884 | 3898 | Negative |
| hCoVHKU1 | orf1ab | ELSKELKRPVNPTDA | 2548 | 2562 | Negative |
| hCoVHKU1 | orf1ab | ELTCLRYFAKLGREL | 6178 | 6192 | Negative |
| hCoVHKU1 | orf1ab | ELVIGYSLYTVWFYP | 2400 | 2414 | Negative |
| hCoVHKU1 | orf1ab | EMVAYNPDLSQYYCD | 1968 | 1982 | Negative |
| hCoVHKU1 | orf1ab | ENIGTNFPLQIGFST | 6086 | 6100 | Negative |
| hCoVHKU1 | orf1ab | EQIDGKFVTKFSGDK | 1320 | 1334 | Negative |
| hCoVHKU1 | orf1ab | EVEKGVTVDDFVAVV | 852 | 866 | Negative |
| hCoVHKU1 | orf1ab | EVHAEPKGKYSQKAY | 200 | 214 | Negative |
| hCoVHKU1 | orf1ab | EVLLLRHDIQLNNDV | 1566 | 1580 | Negative |
| hCoVHKU1 | orf1ab | EVLTKTAVKSGTAIL | 90 | 104 | Negative |
| hCoVHKU1 | orf1ab | EVSDDYVQDSTVLQA | 4000 | 4014 | Negative |
| hCoVHKU1 | orf1ab | EYAANSSILSLCAFS | 4326 | 4340 | Negative |
| hCoVHKU1 | orf1ab | FAAETQKATEEAFKQ | 5518 | 5532 | Negative |
| hCoVHKU1 | orf1ab | FAERDCYTFKKTVAK | 6112 | 6126 | Negative |
| hCoVHKU1 | orf1ab | FAIEDVEDVISSEAV | 918 | 932 | Negative |
| hCoVHKU1 | orf1ab | FCGRDLFDLFYQFFS | 3092 | 3106 | Negative |
| hCoVHKU1 | orf1ab | FDDILGKVCSEFEVE | 840 | 854 | Negative |
| hCoVHKU1 | orf1ab | FDDNFCAFCTPKKIF | 1286 | 1300 | Negative |
| hCoVHKU1 | orf1ab | FDEAGDEAMASRMYC | 902 | 916 | Negative |
| hCoVHKU1 | orf1ab | FDICNTNRAGIGLYY | 4492 | 4506 | Negative |
| hCoVHKU1 | orf1ab | FDLDSTFDDILGKVC | 834 | 848 | Negative |
| hCoVHKU1 | orf1ab | FDQQQLLAYYNFLTV | 1722 | 1736 | Negative |
| hCoVHKU1 | orf1ab | FDQWYESTFGSVYYH | 2906 | 2920 | Negative |
| hCoVHKU1 | orf1ab | FDVDRKSFNNFVNIA | 2676 | 2690 | Negative |
| hCoVHKU1 | orf1ab | FDVEGAHATRENIGT | 6076 | 6090 | Negative |
| hCoVHKU1 | orf1ab | FEDELAPSDVYQQLA | 3618 | 3632 | Negative |
| hCoVHKU1 | orf1ab | FHQKCLKSIAATRGV | 5024 | 5038 | Negative |
| hCoVHKU1 | orf1ab | FITVEAAIELSKELK | 2540 | 2554 | Negative |
| hCoVHKU1 | orf1ab | FLGTYTWTTILSLAI | 3806 | 3820 | Negative |
| hCoVHKU1 | orf1ab | FNDNCDFYGWVSGNM | 366 | 380 | Negative |
| hCoVHKU1 | orf1ab | FNDVYNDALFVSYGS | 1124 | 1138 | Negative |
| hCoVHKU1 | orf1ab | FPLQIGFSTGVDFVV | 6092 | 6106 | Negative |
| hCoVHKU1 | orf1ab | FQTVKPGNFNQDFYE | 4860 | 4874 | Negative |
| hCoVHKU1 | orf1ab | FSTGVDFVVEATGLF | 6098 | 6112 | Negative |
| hCoVHKU1 | orf1ab | FSVEDAYAEVHAEPK | 192 | 206 | Negative |
| hCoVHKU1 | orf1ab | FTDENYNNLVPTYLK | 2740 | 2754 | Negative |
| hCoVHKU1 | orf1ab | FTEESFYKNMYLKSA | 5368 | 5382 | Negative |
| hCoVHKU1 | orf1ab | GALEAFRKARNGVFI | 6632 | 6646 | Negative |
| hCoVHKU1 | orf1ab | GCFVDDLLKTDSVLL | 5294 | 5308 | Negative |
| hCoVHKU1 | orf1ab | GCHTGTDFTGNFYGP | 3504 | 3518 | Negative |
| hCoVHKU1 | orf1ab | GDGTPHPYCYSDGVM | 2998 | 3012 | Negative |
| hCoVHKU1 | orf1ab | GFIRFPDVISEGIVR | 3034 | 3048 | Negative |
| hCoVHKU1 | orf1ab | GFSFDLNGLDALFFY | 1242 | 1256 | Negative |
| hCoVHKU1 | orf1ab | GGVIRYYDITANGGT | 2504 | 2518 | Negative |
| hCoVHKU1 | orf1ab | GHYDGRTGELPCAIM | 6522 | 6536 | Negative |
| hCoVHKU1 | orf1ab | GIKPVLFVDQYGCDY | 222 | 236 | Negative |
| hCoVHKU1 | orf1ab | GIKQESRVGVDAVMH | 1838 | 1852 | Negative |

| | | | | | |
|---|---|---|---|---|---|
| hCoVHKU1 | orf1ab | GINKQLTDISVDSNW | 4178 | 4192 | Negative |
| hCoVHKU1 | orf1ab | GIVKMVSPTSKIEPC | 3336 | 3350 | Negative |
| hCoVHKU1 | orf1ab | GKFVQVPLGIKDPIL | 4414 | 4428 | Negative |
| hCoVHKU1 | orf1ab | GKLADCLQAYGHYSL | 238 | 252 | Negative |
| hCoVHKU1 | orf1ab | GKTVLGEYVFDKSEL | 5576 | 5590 | Negative |
| hCoVHKU1 | orf1ab | GLACVKDYEFKFYDA | 6324 | 6338 | Negative |
| hCoVHKU1 | orf1ab | GLAVYYYTARVVYTA | 5678 | 5692 | Negative |
| hCoVHKU1 | orf1ab | GLCPSTGQAMESVGF | 38 | 52 | Negative |
| hCoVHKU1 | orf1ab | GLFRRKKKSNLLIQE | 6790 | 6804 | Negative |
| hCoVHKU1 | orf1ab | GLGFKWAPEFRWLLP | 8 | 22 | Negative |
| hCoVHKU1 | orf1ab | GLIPRSYYLIQSGIF | 554 | 568 | Negative |
| hCoVHKU1 | orf1ab | GLKLKLTFNKQEASV | 2812 | 2826 | Negative |
| hCoVHKU1 | orf1ab | GLKYTKIMKDDGNCV | 4258 | 4272 | Negative |
| hCoVHKU1 | orf1ab | GLQLFTTWLPDLFML | 2420 | 2434 | Negative |
| hCoVHKU1 | orf1ab | GLRPPRNSFEAILLN | 3892 | 3906 | Negative |
| hCoVHKU1 | orf1ab | GLSTDVQLRAFDICN | 4482 | 4496 | Negative |
| hCoVHKU1 | orf1ab | GLVTICDFYSKLGVG | 2334 | 2348 | Negative |
| hCoVHKU1 | orf1ab | GLWIPIIKSSVKSYD | 446 | 460 | Negative |
| hCoVHKU1 | orf1ab | GLWSPTITHTNCWLR | 1150 | 1164 | Negative |
| hCoVHKU1 | orf1ab | GLYKQSCTGSPYIDD | 5476 | 5490 | Negative |
| hCoVHKU1 | orf1ab | GMIKTSKYGLGFKWA | 1 | 14 | Negative |
| hCoVHKU1 | orf1ab | GMSFSMSSFELPQLY | 1342 | 1356 | Negative |
| hCoVHKU1 | orf1ab | GRDLFDLFYQFFSSL | 3094 | 3108 | Negative |
| hCoVHKU1 | orf1ab | GRDTGALEAFRKARN | 6628 | 6642 | Negative |
| hCoVHKU1 | orf1ab | GSCTFEDELAPSDVY | 3614 | 3628 | Negative |
| hCoVHKU1 | orf1ab | GTDFTGNFYGPYRDA | 3508 | 3522 | Negative |
| hCoVHKU1 | orf1ab | GTGMAITIKPEATIN | 4370 | 4384 | Negative |
| hCoVHKU1 | orf1ab | GVAESVIVEDDVIEN | 738 | 752 | Negative |
| hCoVHKU1 | orf1ab | GVATEYAANSSILSL | 4322 | 4336 | Negative |
| hCoVHKU1 | orf1ab | GVDAVMHFGTLAKTD | 1846 | 1860 | Negative |
| hCoVHKU1 | orf1ab | GVGFTSHFCNGSFIC | 2346 | 2360 | Negative |
| hCoVHKU1 | orf1ab | GVKTVKYFECPNSIY | 1606 | 1620 | Negative |
| hCoVHKU1 | orf1ab | GVLIQNGAKHVQGNV | 2764 | 2778 | Negative |
| hCoVHKU1 | orf1ab | GVLTLDNQDLYGQWY | 4660 | 4674 | Negative |
| hCoVHKU1 | orf1ab | GVMKNASLYTSLVPH | 3010 | 3024 | Negative |
| hCoVHKU1 | orf1ab | GVNFKSISLTVGEVF | 1660 | 1674 | Negative |
| hCoVHKU1 | orf1ab | GVPFVVSIGYHYKEL | 4790 | 4804 | Negative |
| hCoVHKU1 | orf1ab | GVSIETLLAAIKRLY | 3590 | 3604 | Negative |
| hCoVHKU1 | orf1ab | GVSILSTMTGRMFHQ | 5012 | 5026 | Negative |
| hCoVHKU1 | orf1ab | GVTLCVLCFVSFMML | 3680 | 3694 | Negative |
| hCoVHKU1 | orf1ab | GVTVDDFVAVVCDAI | 856 | 870 | Negative |
| hCoVHKU1 | orf1ab | GVVGCKSIVKETALI | 466 | 480 | Negative |
| hCoVHKU1 | orf1ab | GVVLSNLLYILFFVS | 2838 | 2852 | Negative |
| hCoVHKU1 | orf1ab | GVYNYKISVQELRYM | 3874 | 3888 | Negative |
| hCoVHKU1 | orf1ab | GVYTNFKLVGHDICA | 2004 | 2018 | Negative |
| hCoVHKU1 | orf1ab | GWDDMLRHLIKDVDN | 5050 | 5064 | Negative |
| hCoVHKU1 | orf1ab | HDICAQLNDKLGFNV | 2014 | 2028 | Negative |
| hCoVHKU1 | orf1ab | HFKFDEPSDATDFIR | 1802 | 1816 | Negative |
| hCoVHKU1 | orf1ab | HLPAYASFKVIDNGV | 2874 | 2888 | Negative |

| | | | | | |
|---|---|---|---|---|---|
| hCoVHKU1 | orf1ab | HNSMDCPIVVAVMDE | 2920 | 2934 | Negative |
| hCoVHKU1 | orf1ab | HTGTDFTGNFYGPYR | 3506 | 3520 | Negative |
| hCoVHKU1 | orf1ab | HTRYSLANSNGFIRF | 3024 | 3038 | Negative |
| hCoVHKU1 | orf1ab | HVASASALLDLRTCC | 4834 | 4848 | Negative |
| hCoVHKU1 | orf1ab | HVLWLFSYCRKIGVN | 3226 | 3240 | Negative |
| hCoVHKU1 | orf1ab | HYVVTDIKQVGCMMR | 2564 | 2578 | Negative |
| hCoVHKU1 | orf1ab | ICDCGIKQESRVGVD | 1834 | 1848 | Negative |
| hCoVHKU1 | orf1ab | IEDLSIRNLQKRLYS | 5168 | 5182 | Negative |
| hCoVHKU1 | orf1ab | IFADTLVEVGLVGVL | 4648 | 4662 | Negative |
| hCoVHKU1 | orf1ab | IGFSTGVDFVVEATG | 6096 | 6110 | Negative |
| hCoVHKU1 | orf1ab | IGTNFPLQIGFSTGV | 6088 | 6102 | Negative |
| hCoVHKU1 | orf1ab | IIVNDENSSIKVVKS | 2142 | 2156 | Negative |
| hCoVHKU1 | orf1ab | IKSSVKSYDDLVYSG | 452 | 466 | Negative |
| hCoVHKU1 | orf1ab | ILGSCTFEDELAPSD | 3612 | 3626 | Negative |
| hCoVHKU1 | orf1ab | ILTTPFSLKGGVVLS | 2828 | 2842 | Negative |
| hCoVHKU1 | orf1ab | IMSGRMSLTVVSYQM | 3402 | 3416 | Negative |
| hCoVHKU1 | orf1ab | INGVKTVKYFECPNS | 1604 | 1618 | Negative |
| hCoVHKU1 | orf1ab | IVEDDVIENVKSSLS | 744 | 758 | Negative |
| hCoVHKU1 | orf1ab | IVGGVIRYYDITANG | 2502 | 2516 | Negative |
| hCoVHKU1 | orf1ab | IVPFGFMPSYVHKWF | 154 | 168 | Negative |
| hCoVHKU1 | orf1ab | IVTGDNDDQIVVTGD | 1090 | 1104 | Negative |
| hCoVHKU1 | orf1ab | KCGNLEQNHILGVNN | 492 | 506 | Negative |
| hCoVHKU1 | orf1ab | KCSDFYADKILYQYE | 1692 | 1706 | Negative |
| hCoVHKU1 | orf1ab | KFDEPSDATDFIRVV | 1804 | 1818 | Negative |
| hCoVHKU1 | orf1ab | KFSKETAAMVKSKGV | 1408 | 1422 | Negative |
| hCoVHKU1 | orf1ab | KIGVNVCSDSTFEET | 3236 | 3250 | Negative |
| hCoVHKU1 | orf1ab | KLELFNKYFKYWGMK | 4732 | 4746 | Negative |
| hCoVHKU1 | orf1ab | KLGVGFTSHFCNGSF | 2344 | 2358 | Negative |
| hCoVHKU1 | orf1ab | KQLLFVLEVVYKYFE | 4920 | 4934 | Negative |
| hCoVHKU1 | orf1ab | KVCSEFEVEKGVTVD | 846 | 860 | Negative |
| hCoVHKU1 | orf1ab | KVGELKVEFWFAMRK | 6672 | 6686 | Negative |
| hCoVHKU1 | orf1ab | KYGLGFKWAPEFRWL | 6 | 20 | Negative |
| hCoVHKU1 | orf1ab | LDGLIPRSYYLIQSG | 552 | 566 | Negative |
| hCoVHKU1 | orf1ab | LEFTDENYNNLVPTY | 2738 | 2752 | Negative |
| hCoVHKU1 | orf1ab | LIGLQLFTTWLPDLF | 2418 | 2432 | Negative |
| hCoVHKU1 | orf1ab | LKFIENLNILFDGRD | 6616 | 6630 | Negative |
| hCoVHKU1 | orf1ab | LLVDKKLITTACNGI | 2644 | 2658 | Negative |
| hCoVHKU1 | orf1ab | LMGFKLDLTLDGYSK | 6044 | 6058 | Negative |
| hCoVHKU1 | orf1ab | LQIGFSTGVDFVVEA | 6094 | 6108 | Negative |
| hCoVHKU1 | orf1ab | LRTCCFSVAAITSGI | 4844 | 4858 | Negative |
| hCoVHKU1 | orf1ab | LSMIKGPQRADLNGV | 6654 | 6668 | Negative |
| hCoVHKU1 | orf1ab | LTDCLYLKNLTQTFT | 1942 | 1956 | Negative |
| hCoVHKU1 | orf1ab | LTGDSVKFVYMHQLE | 3486 | 3500 | Negative |
| hCoVHKU1 | orf1ab | LVGVLTLDNQDLYGQ | 4658 | 4672 | Negative |
| hCoVHKU1 | orf1ab | MASFVEYEVAKKNLA | 4022 | 4036 | Negative |
| hCoVHKU1 | orf1ab | MDDFSIKSIYNVDLC | 338 | 352 | Negative |
| hCoVHKU1 | orf1ab | MFDVYYVDTFMSHFDV | 2664 | 2678 | Negative |
| hCoVHKU1 | orf1ab | MHQLELSTGCHTGTD | 3496 | 3510 | Negative |
| hCoVHKU1 | orf1ab | MIKGPQRADLNGVIV | 6656 | 6670 | Negative |

| | | | | | |
|---|---|---|---|---|---|
| hCoVHKU1 | orf1ab | MTGVSIETLLAAIKR | 3588 | 3602 | Negative |
| hCoVHKU1 | orf1ab | MVDIKQLLFVLEVVY | 4916 | 4930 | Negative |
| hCoVHKU1 | orf1ab | NCSVRVKCSTIVGGV | 2492 | 2506 | Negative |
| hCoVHKU1 | orf1ab | NEYYEFLCKHFSMMI | 5196 | 5210 | Negative |
| hCoVHKU1 | orf1ab | NNFVNIAHASLREGV | 2684 | 2698 | Negative |
| hCoVHKU1 | orf1ab | NSSIKVVKSLSLVDV | 2148 | 2162 | Negative |
| hCoVHKU1 | orf1ab | NTFITVEAAIELSKE | 2538 | 2552 | Negative |
| hCoVHKU1 | orf1ab | PAYASFKVIDNGVVR | 2876 | 2890 | Negative |
| hCoVHKU1 | orf1ab | PIIKSSVKSYDDLVY | 450 | 464 | Negative |
| hCoVHKU1 | orf1ab | PILLVDKKLITTACN | 2642 | 2656 | Negative |
| hCoVHKU1 | orf1ab | PKGLACVKDYEFKFY | 6322 | 6336 | Negative |
| hCoVHKU1 | orf1ab | PMGVYNYKISVQELR | 3872 | 3886 | Negative |
| hCoVHKU1 | orf1ab | PVAKSVKQLFYVYDV | 6340 | 6354 | Negative |
| hCoVHKU1 | orf1ab | PVWNSAVFISPYNSQ | 5888 | 5902 | Negative |
| hCoVHKU1 | orf1ab | PYNSQNYVAKRVLGV | 5898 | 5912 | Negative |
| hCoVHKU1 | orf1ab | PYTPKYTFGNVKPGE | 3430 | 3444 | Negative |
| hCoVHKU1 | orf1ab | QAGVATEYAANSSIL | 4320 | 4334 | Negative |
| hCoVHKU1 | orf1ab | QEIVSDREVILCWET | 5540 | 5554 | Negative |
| hCoVHKU1 | orf1ab | QGRQILGSCTFEDEL | 3608 | 3622 | Negative |
| hCoVHKU1 | orf1ab | QIVVTGDDVDDIESI | 1098 | 1112 | Negative |
| hCoVHKU1 | orf1ab | QPTEDVVDGDVVIRE | 300 | 314 | Negative |
| hCoVHKU1 | orf1ab | QVDYVPLRSATCITR | 6444 | 6458 | Negative |
| hCoVHKU1 | orf1ab | RAFDICNTNRAGIGL | 4490 | 4504 | Negative |
| hCoVHKU1 | orf1ab | RHVICSSSNMNEPDY | 3374 | 3388 | Negative |
| hCoVHKU1 | orf1ab | RQILGSCTFEDELAP | 3610 | 3624 | Negative |
| hCoVHKU1 | orf1ab | RVLFDYVSLVKLIVE | 2386 | 2400 | Negative |
| hCoVHKU1 | orf1ab | SAKNRARTVAGVSIL | 5002 | 5016 | Negative |
| hCoVHKU1 | orf1ab | SFKSENRYSVLSVDS | 2090 | 2104 | Negative |
| hCoVHKU1 | orf1ab | SFKVIDNGVVRDISV | 2880 | 2894 | Negative |
| hCoVHKU1 | orf1ab | SFVEYEVAKKNLADA | 4024 | 4038 | Negative |
| hCoVHKU1 | orf1ab | SGKMDTAAYREAACS | 3288 | 3302 | Negative |
| hCoVHKU1 | orf1ab | SIGYHYKELGVVMNL | 4796 | 4810 | Negative |
| hCoVHKU1 | orf1ab | SKELKRPVNPTDASH | 2550 | 2564 | Negative |
| hCoVHKU1 | orf1ab | SKGLFKEGSTVDLKH | 4878 | 4892 | Negative |
| hCoVHKU1 | orf1ab | SKMTSLPKDWRLINK | 1586 | 1600 | Negative |
| hCoVHKU1 | orf1ab | SKQVDYVPLRSATCI | 6442 | 6456 | Negative |
| hCoVHKU1 | orf1ab | SKTKRFIKETIYWIL | 3638 | 3652 | Negative |
| hCoVHKU1 | orf1ab | SLFDDNFCAFCTPKK | 1284 | 1298 | Negative |
| hCoVHKU1 | orf1ab | SNGFIRFPDVISEGI | 3032 | 3046 | Negative |
| hCoVHKU1 | orf1ab | SRVLSSFEPRSDLER | 6734 | 6748 | Negative |
| hCoVHKU1 | orf1ab | SSEGFLIGINYLGKS | 7084 | 7098 | Negative |
| hCoVHKU1 | orf1ab | SSVETETYFKVNGLW | 1138 | 1152 | Negative |
| hCoVHKU1 | orf1ab | SSVKSYDDLVYSGVV | 454 | 468 | Negative |
| hCoVHKU1 | orf1ab | STGCHTGTDFTGNFY | 3502 | 3516 | Negative |
| hCoVHKU1 | orf1ab | STGQAMESVGFVYDN | 42 | 56 | Negative |
| hCoVHKU1 | orf1ab | SVMESQISTKEVKLK | 2116 | 2130 | Negative |
| hCoVHKU1 | orf1ab | SVRVKCSTIVGGVIR | 2494 | 2508 | Negative |
| hCoVHKU1 | orf1ab | SVWHIQTVQDADGIN | 4166 | 4180 | Negative |
| hCoVHKU1 | orf1ab | SYKTKRYVVHHLSMT | 126 | 140 | Negative |

| | | | | | |
|---|---|---|---|---|---|
| hCoVHKU1 | orf1ab | TEVDDYVLANECIER | 5500 | 5514 | Negative |
| hCoVHKU1 | orf1ab | TFGLVTICDFYSKLG | 2332 | 2346 | Negative |
| hCoVHKU1 | orf1ab | TFTSMLTNYFLDDVE | 1954 | 1968 | Negative |
| hCoVHKU1 | orf1ab | TLVEVGLVGVLTLDN | 4652 | 4666 | Negative |
| hCoVHKU1 | orf1ab | TNFPLQIGFSTGVDF | 6090 | 6104 | Negative |
| hCoVHKU1 | orf1ab | TNMYKEARINDKKSK | 4082 | 4096 | Negative |
| hCoVHKU1 | orf1ab | TNVNASSSEGFLIGI | 7078 | 7092 | Negative |
| hCoVHKU1 | orf1ab | TYAAIDFVQYEVDRR | 2372 | 2386 | Negative |
| hCoVHKU1 | orf1ab | VDGLKYTKIMKDDGN | 4256 | 4270 | Negative |
| hCoVHKU1 | orf1ab | VDGVNFKSISLTVGE | 1658 | 1672 | Negative |
| hCoVHKU1 | orf1ab | VEDAYAEVHAEPKGK | 194 | 208 | Negative |
| hCoVHKU1 | orf1ab | VEGAHATRENIGTNF | 6078 | 6092 | Negative |
| hCoVHKU1 | orf1ab | VETETYFKVNGLWSP | 1140 | 1154 | Negative |
| hCoVHKU1 | orf1ab | VIANHVLWLFSYCRK | 3222 | 3236 | Negative |
| hCoVHKU1 | orf1ab | VILSTCDGLKFTEES | 5358 | 5372 | Negative |
| hCoVHKU1 | orf1ab | VIMHLQWIVMYGAIM | 3196 | 3210 | Negative |
| hCoVHKU1 | orf1ab | VIVNNYDKSAGYPFN | 4946 | 4960 | Negative |
| hCoVHKU1 | orf1ab | VIVNPANGHMLHGGG | 1382 | 1396 | Negative |
| hCoVHKU1 | orf1ab | VLHNEILSTSDLSVA | 3958 | 3972 | Negative |
| hCoVHKU1 | orf1ab | VNASSSEGFLIGINY | 7080 | 7094 | Negative |
| hCoVHKU1 | orf1ab | VNGLWSPTITHTNCW | 1148 | 1162 | Negative |
| hCoVHKU1 | orf1ab | VPILTTPFSLKGGVV | 2826 | 2840 | Negative |
| hCoVHKU1 | orf1ab | VTICDFYSKLGVGFT | 2336 | 2350 | Negative |
| hCoVHKU1 | orf1ab | VVSIGYHYKELGVVM | 4794 | 4808 | Negative |
| hCoVHKU1 | orf1ab | VVYLFDEAGDEAMAS | 898 | 912 | Negative |
| hCoVHKU1 | orf1ab | WIPIIKSSVKSYDDL | 448 | 462 | Negative |
| hCoVHKU1 | orf1ab | WTIFMYINTHMIGVT | 3668 | 3682 | Negative |
| hCoVHKU1 | orf1ab | YAFASDSVQCYTPHI | 2960 | 2974 | Negative |
| hCoVHKU1 | orf1ab | YASFKVIDNGVVRDI | 2878 | 2892 | Negative |
| hCoVHKU1 | orf1ab | YCRARVEHPDVDGIC | 4396 | 4410 | Negative |
| hCoVHKU1 | orf1ab | YGLEDYAFDHIVYGS | 6762 | 6776 | Negative |
| hCoVHKU1 | orf1ab | YGSMTLNGLWLDDKV | 3356 | 3370 | Negative |
| hCoVHKU1 | orf1ab | YLFDEAGDEAMASRM | 900 | 914 | Negative |
| hCoVHKU1 | orf1ab | YLGKSSFEIDGNVMH | 7094 | 7108 | Negative |
| hCoVHKU1 | orf1ab | YMGFQGRQILGSCTF | 3604 | 3618 | Negative |
| hCoVHKU1 | orf1ab | YNLVNVGHYDGRTGE | 6516 | 6530 | Negative |
| hCoVHKU1 | orf1ab | YREAACSQLAKAMET | 3296 | 3310 | Negative |
| hCoVHKU1 | orf1ab | YTNFKLVGHDICAQL | 2006 | 2020 | Negative |
| hCoVHKU1 | orf1ab | YTQLCQYLNTTTLAV | 6930 | 6944 | Negative |
| hCoVHKU1 | orf1ab | YTTEIASKFTFNLFC | 2254 | 2268 | Negative |
| hCoVHKU1 | orf1ab | YVVHHLSMTTSTTNF | 132 | 146 | Negative |
| hCoVHKU1 | orf1ab | AAEELASPMKSDEGG | 24 | 38 | Overlapping |
| hCoVHKU1 | orf1ab | AEEYCNYLESYNIVT | 6470 | 6484 | Overlapping |
| hCoVHKU1 | orf1ab | AVDDKYKVNENLAVN | 6014 | 6028 | Overlapping |
| hCoVHKU1 | orf1ab | CGVVAEHDFFTFDID | 4550 | 4564 | Overlapping |
| hCoVHKU1 | orf1ab | CTFAIEDVEDVISSE | 916 | 930 | Overlapping |
| hCoVHKU1 | orf1ab | DDEDVVTGDNDDEDV | 946 | 960 | Overlapping |
| hCoVHKU1 | orf1ab | DDEDVVTGDNNDEEI | 1076 | 1090 | Overlapping |
| hCoVHKU1 | orf1ab | DDLLKTDSVLLIERF | 5298 | 5312 | Overlapping |

| | | | | | |
|---|---|---|---|---|---|
| hCoVHKU1 | orf1ab | DDLVYSGVVGCKSIV | 460 | 474 | Overlapping |
| hCoVHKU1 | orf1ab | DDQIVVTGDDVDDIE | 1096 | 1110 | Overlapping |
| hCoVHKU1 | orf1ab | DDVDDIESIYDFDTY | 1104 | 1118 | Overlapping |
| hCoVHKU1 | orf1ab | DDYDFSVEDAYAEVH | 188 | 202 | Overlapping |
| hCoVHKU1 | orf1ab | DELAPSDVYQQLAGV | 3620 | 3634 | Overlapping |
| hCoVHKU1 | orf1ab | DENYNNLVPTYLKSD | 2742 | 2756 | Overlapping |
| hCoVHKU1 | orf1ab | DEVSMLTNYELSVIN | 5758 | 5772 | Overlapping |
| hCoVHKU1 | orf1ab | DGIDVTKLKCSDFYA | 1684 | 1698 | Overlapping |
| hCoVHKU1 | orf1ab | DGLESKQVDYVPLRS | 6438 | 6452 | Overlapping |
| hCoVHKU1 | orf1ab | DGLKFTEESFYKNMY | 5364 | 5378 | Overlapping |
| hCoVHKU1 | orf1ab | DGQRVYDDVDASLFV | 2584 | 2598 | Overlapping |
| hCoVHKU1 | orf1ab | DIESIYDFDTYKALL | 1108 | 1122 | Overlapping |
| hCoVHKU1 | orf1ab | DMLDTYAAIDFVQYE | 2368 | 2382 | Overlapping |
| hCoVHKU1 | orf1ab | DNDDEDVVTGDNDDE | 954 | 968 | Overlapping |
| hCoVHKU1 | orf1ab | DNDDEDVVTGDNNDE | 1074 | 1088 | Overlapping |
| hCoVHKU1 | orf1ab | DNDDQIVVTGDDVDD | 1094 | 1108 | Overlapping |
| hCoVHKU1 | orf1ab | DNNDEEIVTGDNDDQ | 1084 | 1098 | Overlapping |
| hCoVHKU1 | orf1ab | DTIDGVVEDTINDDE | 934 | 948 | Overlapping |
| hCoVHKU1 | orf1ab | DTINDDEDVVTGDND | 942 | 956 | Overlapping |
| hCoVHKU1 | orf1ab | DVEDVISSEAVEDTI | 922 | 936 | Overlapping |
| hCoVHKU1 | orf1ab | DVEMVAYNPDLSQYY | 1966 | 1980 | Overlapping |
| hCoVHKU1 | orf1ab | DWYDFVENPDIINIY | 4618 | 4632 | Overlapping |
| hCoVHKU1 | orf1ab | DYAFDHIVYGSFNHK | 6766 | 6780 | Overlapping |
| hCoVHKU1 | orf1ab | DYVDYTFVNEYYEFL | 5188 | 5202 | Overlapping |
| hCoVHKU1 | orf1ab | DYVIYSQTAETAHSV | 5926 | 5940 | Overlapping |
| hCoVHKU1 | orf1ab | DYVQDSTVLQALQSE | 4004 | 4018 | Overlapping |
| hCoVHKU1 | orf1ab | EAACSQLAKAMETFN | 3298 | 3312 | Overlapping |
| hCoVHKU1 | orf1ab | EAAIELSKELKRPVN | 2544 | 2558 | Overlapping |
| hCoVHKU1 | orf1ab | EAFKQSYASATIQEI | 5528 | 5542 | Overlapping |
| hCoVHKU1 | orf1ab | EAGDEAMASRMYCTF | 904 | 918 | Overlapping |
| hCoVHKU1 | orf1ab | EALSFEEQNEIYAYT | 4970 | 4984 | Overlapping |
| hCoVHKU1 | orf1ab | EAMASRMYCTFAIED | 908 | 922 | Overlapping |
| hCoVHKU1 | orf1ab | EATINQDSYGGASVC | 4380 | 4394 | Overlapping |
| hCoVHKU1 | orf1ab | EAVEDTIDGVVEDTI | 930 | 944 | Overlapping |
| hCoVHKU1 | orf1ab | EAWYEFRAGRPHRLV | 1780 | 1794 | Overlapping |
| hCoVHKU1 | orf1ab | ECIERLKLFAAETQK | 5510 | 5524 | Overlapping |
| hCoVHKU1 | orf1ab | ECPNSIYICSQGKDF | 1614 | 1628 | Overlapping |
| hCoVHKU1 | orf1ab | ECVTVDVLKDMPVLK | 684 | 698 | Overlapping |
| hCoVHKU1 | orf1ab | EDDVIENVKSSLSSY | 746 | 760 | Overlapping |
| hCoVHKU1 | orf1ab | EDFLGWIVPFGFMPS | 148 | 162 | Overlapping |
| hCoVHKU1 | orf1ab | EDFNVWAMANGFSQV | 3562 | 3576 | Overlapping |
| hCoVHKU1 | orf1ab | EDIGSTMFNVPTKVL | 2934 | 2948 | Overlapping |
| hCoVHKU1 | orf1ab | EDLHVVEVLTKTAVK | 84 | 98 | Overlapping |
| hCoVHKU1 | orf1ab | EDVISSEAVEDTIDG | 924 | 938 | Overlapping |
| hCoVHKU1 | orf1ab | EDVVTGDNDDEDVVT | 948 | 962 | Overlapping |
| hCoVHKU1 | orf1ab | EDVVTGDNNDEEIVT | 1078 | 1092 | Overlapping |
| hCoVHKU1 | orf1ab | EEDVLLFITAFLGTY | 3796 | 3810 | Overlapping |
| hCoVHKU1 | orf1ab | EEIVTGDNDDQIVVT | 1088 | 1102 | Overlapping |
| hCoVHKU1 | orf1ab | EELASPMKSDEGGLC | 26 | 40 | Overlapping |

| | | | | | |
|---|---|---|---|---|---|
| hCoVHKU1 | orf1ab | EESDLIISNFKFDDY | 176 | 190 | Overlapping |
| hCoVHKU1 | orf1ab | EESFYKNMYLKSAVM | 5370 | 5384 | Overlapping |
| hCoVHKU1 | orf1ab | EETSLTTFMITKDSY | 3248 | 3262 | Overlapping |
| hCoVHKU1 | orf1ab | EEYQKVFRVYLEYIK | 5328 | 5342 | Overlapping |
| hCoVHKU1 | orf1ab | EFCCSHGDRFYRLAN | 5096 | 5110 | Overlapping |
| hCoVHKU1 | orf1ab | EFCSQHTMLVKIDGD | 5264 | 5278 | Overlapping |
| hCoVHKU1 | orf1ab | EFEVEKGVTVDDFVA | 850 | 864 | Overlapping |
| hCoVHKU1 | orf1ab | EFKFYDAFPVAKSVK | 6332 | 6346 | Overlapping |
| hCoVHKU1 | orf1ab | EFLCKHFSMMILSDD | 5200 | 5214 | Overlapping |
| hCoVHKU1 | orf1ab | EFLQYDSSIHSYFIT | 6804 | 6818 | Overlapping |
| hCoVHKU1 | orf1ab | EFRAGRPHRLVALVL | 1784 | 1798 | Overlapping |
| hCoVHKU1 | orf1ab | EFRWLLPDAAEELAS | 16 | 30 | Overlapping |
| hCoVHKU1 | orf1ab | EFVKSKGLFKEGSTV | 4874 | 4888 | Overlapping |
| hCoVHKU1 | orf1ab | EFVNMASFVEYEVAK | 4018 | 4032 | Overlapping |
| hCoVHKU1 | orf1ab | EFYGFLKWLFIYVFS | 2228 | 2242 | Overlapping |
| hCoVHKU1 | orf1ab | EGFLIGINYLGKSSF | 7086 | 7100 | Overlapping |
| hCoVHKU1 | orf1ab | EGGLCPSTGQAMESV | 36 | 50 | Overlapping |
| hCoVHKU1 | orf1ab | EGICFNFNSSWVLNN | 3068 | 3082 | Overlapping |
| hCoVHKU1 | orf1ab | EGIVRIVRTRSMTYC | 3044 | 3058 | Overlapping |
| hCoVHKU1 | orf1ab | EGSTVDLKHFFFTQD | 4884 | 4898 | Overlapping |
| hCoVHKU1 | orf1ab | EGVQLEKVLDTFVGC | 2696 | 2710 | Overlapping |
| hCoVHKU1 | orf1ab | EHPDVDGICKLRGKF | 4402 | 4416 | Overlapping |
| hCoVHKU1 | orf1ab | EILSTSDLSVAFDKL | 3962 | 3976 | Overlapping |
| hCoVHKU1 | orf1ab | EIYDGGCIPASQVIV | 4934 | 4948 | Overlapping |
| hCoVHKU1 | orf1ab | EKGVTVDDFVAVVCD | 854 | 868 | Overlapping |
| hCoVHKU1 | orf1ab | EKICIIDNMYMGKCG | 770 | 784 | Overlapping |
| hCoVHKU1 | orf1ab | EKTYYELTKSCGVVA | 4540 | 4554 | Overlapping |
| hCoVHKU1 | orf1ab | ELELICDCGIKQESR | 1830 | 1844 | Overlapping |
| hCoVHKU1 | orf1ab | ELFNKYFKYWGMKYH | 4734 | 4748 | Overlapping |
| hCoVHKU1 | orf1ab | ELICDCGIKQESRVG | 1832 | 1846 | Overlapping |
| hCoVHKU1 | orf1ab | ELKRPVNPTDASHYV | 2552 | 2566 | Overlapping |
| hCoVHKU1 | orf1ab | ELPCAIMNDKVVVKI | 6530 | 6544 | Overlapping |
| hCoVHKU1 | orf1ab | ELRYMNANGLRPPRN | 3884 | 3898 | Overlapping |
| hCoVHKU1 | orf1ab | ELSKELKRPVNPTDA | 2548 | 2562 | Overlapping |
| hCoVHKU1 | orf1ab | ELTCLRYFAKLGREL | 6178 | 6192 | Overlapping |
| hCoVHKU1 | orf1ab | ELVIGYSLYTVWFYP | 2400 | 2414 | Overlapping |
| hCoVHKU1 | orf1ab | EMVAYNPDLSQYYCD | 1968 | 1982 | Overlapping |
| hCoVHKU1 | orf1ab | ENEEYQKVFRVYLEY | 5326 | 5340 | Overlapping |
| hCoVHKU1 | orf1ab | ENHKPHYSFKLVMNG | 5458 | 5472 | Overlapping |
| hCoVHKU1 | orf1ab | ENLNILFDGRDTGAL | 6620 | 6634 | Overlapping |
| hCoVHKU1 | orf1ab | ENLSLADISAVQSSF | 1706 | 1720 | Overlapping |
| hCoVHKU1 | orf1ab | ENPDIINIYKKLGPI | 4624 | 4638 | Overlapping |
| hCoVHKU1 | orf1ab | ENPVLFTNSTDTVNH | 408 | 422 | Overlapping |
| hCoVHKU1 | orf1ab | ENVIYNLVNVGHYDG | 6512 | 6526 | Overlapping |
| hCoVHKU1 | orf1ab | ENVVYLFDEAGDEAM | 896 | 910 | Overlapping |
| hCoVHKU1 | orf1ab | EPCIVSVTYGSMTLN | 3348 | 3362 | Overlapping |
| hCoVHKU1 | orf1ab | EPKGKYSQKAYALLR | 204 | 218 | Overlapping |
| hCoVHKU1 | orf1ab | EPRHFNSITKIMCCL | 5802 | 5816 | Overlapping |
| hCoVHKU1 | orf1ab | EPVHLLSADAIVLKL | 314 | 328 | Overlapping |

| | | | | | |
|---|---|---|---|---|---|
| hCoVHKU1 | orf1ab | EQNEIYAYTKRNVLP | 4976 | 4990 | Overlapping |
| hCoVHKU1 | orf1ab | EQQIYFFEGVAESVI | 730 | 744 | Overlapping |
| hCoVHKU1 | orf1ab | ERDCYTFKKTVAKAP | 6114 | 6128 | Overlapping |
| hCoVHKU1 | orf1ab | ERDKAVARKLERMAD | 4064 | 4078 | Overlapping |
| hCoVHKU1 | orf1ab | ERFVSLAIDAYPLVH | 5310 | 5324 | Overlapping |
| hCoVHKU1 | orf1ab | ERMADLALTNMYKEA | 4074 | 4088 | Overlapping |
| hCoVHKU1 | orf1ab | ERVSLWNYGKPINLP | 6906 | 6920 | Overlapping |
| hCoVHKU1 | orf1ab | ESDADRANFLNAVVF | 2620 | 2634 | Overlapping |
| hCoVHKU1 | orf1ab | ESIYDFDTYKALLVF | 1110 | 1124 | Overlapping |
| hCoVHKU1 | orf1ab | ESQGNVVTSVMESQI | 2108 | 2122 | Overlapping |
| hCoVHKU1 | orf1ab | ESRVGVDAVMHFGTL | 1842 | 1856 | Overlapping |
| hCoVHKU1 | orf1ab | ESVGFVYDNHVKIDC | 48 | 62 | Overlapping |
| hCoVHKU1 | orf1ab | ESYNIVTTAGFTFWV | 6478 | 6492 | Overlapping |
| hCoVHKU1 | orf1ab | ETAAMVKSKGVCQVG | 1412 | 1426 | Overlapping |
| hCoVHKU1 | orf1ab | ETALITHALYLDYVQ | 476 | 490 | Overlapping |
| hCoVHKU1 | orf1ab | ETFGKPVIWFCHDEA | 2064 | 2078 | Overlapping |
| hCoVHKU1 | orf1ab | ETFTVLAAYNGRPQG | 3444 | 3458 | Overlapping |
| hCoVHKU1 | orf1ab | ETIYWILISTFLFSC | 3646 | 3660 | Overlapping |
| hCoVHKU1 | orf1ab | ETLLAAIKRLYMGFQ | 3594 | 3608 | Overlapping |
| hCoVHKU1 | orf1ab | ETRFITKSMISAVAA | 2722 | 2736 | Overlapping |
| hCoVHKU1 | orf1ab | EVAPGSAVLRQWLPS | 6960 | 6974 | Overlapping |
| hCoVHKU1 | orf1ab | EVDRRVLFDYVSLVK | 2382 | 2396 | Overlapping |
| hCoVHKU1 | orf1ab | EVEKGVTVDDFVAVV | 852 | 866 | Overlapping |
| hCoVHKU1 | orf1ab | EVFGKILGNVFCDGI | 1672 | 1686 | Overlapping |
| hCoVHKU1 | orf1ab | EVHAEPKGKYSQKAY | 200 | 214 | Overlapping |
| hCoVHKU1 | orf1ab | EVKLKGVRKTVKIED | 2126 | 2140 | Overlapping |
| hCoVHKU1 | orf1ab | EVLLLRHDIQLNNDV | 1566 | 1580 | Overlapping |
| hCoVHKU1 | orf1ab | EVLTKTAVKSGTAIL | 90 | 104 | Overlapping |
| hCoVHKU1 | orf1ab | EVSDDYVQDSTVLQA | 4000 | 4014 | Overlapping |
| hCoVHKU1 | orf1ab | EYAEEGICFNFNSSW | 3064 | 3078 | Overlapping |
| hCoVHKU1 | orf1ab | EYCNYLESYNIVTTA | 6472 | 6486 | Overlapping |
| hCoVHKU1 | orf1ab | EYDYVIYSQTAETAH | 5924 | 5938 | Overlapping |
| hCoVHKU1 | orf1ab | EYKVTVWPVATGDVV | 2034 | 2048 | Overlapping |
| hCoVHKU1 | orf1ab | EYPIISNEVSINTSC | 6282 | 6296 | Overlapping |
| hCoVHKU1 | orf1ab | EYVFDKSELTNGVYY | 5582 | 5596 | Overlapping |
| hCoVHKU1 | orf1ab | FAIEDVEDVISSEAV | 918 | 932 | Overlapping |
| hCoVHKU1 | orf1ab | FEDELAPSDVYQQLA | 3618 | 3632 | Overlapping |
| hCoVHKU1 | orf1ab | FEEQNEIYAYTKRNV | 4974 | 4988 | Overlapping |
| hCoVHKU1 | orf1ab | FEPRSDLERDFIDMD | 6740 | 6754 | Overlapping |
| hCoVHKU1 | orf1ab | FGKILGNVFCDGIDV | 1674 | 1688 | Overlapping |
| hCoVHKU1 | orf1ab | FLNKLNENVVYLFDE | 890 | 904 | Overlapping |
| hCoVHKU1 | orf1ab | FVDDLLKTDSVLLIE | 5296 | 5310 | Overlapping |
| hCoVHKU1 | orf1ab | FYDRDGQRVYDDVDA | 2580 | 2594 | Overlapping |
| hCoVHKU1 | orf1ab | GCETFGKPVIWFCHD | 2062 | 2076 | Overlapping |
| hCoVHKU1 | orf1ab | GFDMLDTYAAIDFVQ | 2366 | 2380 | Overlapping |
| hCoVHKU1 | orf1ab | GIKDPILYVLTHDVC | 4422 | 4436 | Overlapping |
| hCoVHKU1 | orf1ab | GKSSFEIDGNVMHAN | 7096 | 7110 | Overlapping |
| hCoVHKU1 | orf1ab | GLFRSYKTKRYVVHH | 122 | 136 | Overlapping |
| hCoVHKU1 | orf1ab | GMAITIKPEATINQD | 4372 | 4386 | Overlapping |

| | | | | | |
|---|---|---|---|---|---|
| hCoVHKU1 | orf1ab | GMKYHPNTVDCDNDR | 4744 | 4758 | Overlapping |
| hCoVHKU1 | orf1ab | GMSYYCENHKPHYSF | 5452 | 5466 | Overlapping |
| hCoVHKU1 | orf1ab | GMVFGLYKQSCTGSP | 5472 | 5486 | Overlapping |
| hCoVHKU1 | orf1ab | GPLVRQIFVDGVPFV | 4780 | 4794 | Overlapping |
| hCoVHKU1 | orf1ab | GTPHPYCYSDGVMKN | 3000 | 3014 | Overlapping |
| hCoVHKU1 | orf1ab | GVIPENPVLFTNSTD | 404 | 418 | Overlapping |
| hCoVHKU1 | orf1ab | GVNVCSDSTFEETSL | 3238 | 3252 | Overlapping |
| hCoVHKU1 | orf1ab | GVPVIEVSQIQSKLT | 3914 | 3928 | Overlapping |
| hCoVHKU1 | orf1ab | GVVEDTINDDEDVVT | 938 | 952 | Overlapping |
| hCoVHKU1 | orf1ab | ICELELICDCGIKQE | 1828 | 1842 | Overlapping |
| hCoVHKU1 | orf1ab | IDGVVEDTINDDEDV | 936 | 950 | Overlapping |
| hCoVHKU1 | orf1ab | IEDAIIVNDENSSIK | 2138 | 2152 | Overlapping |
| hCoVHKU1 | orf1ab | IEDVEDVISSEAVED | 920 | 934 | Overlapping |
| hCoVHKU1 | orf1ab | INDDEDVVTGDNDDE | 944 | 958 | Overlapping |
| hCoVHKU1 | orf1ab | ISEGIVRIVRTRSMT | 3042 | 3056 | Overlapping |
| hCoVHKU1 | orf1ab | ISNFKFDDYDFSVED | 182 | 196 | Overlapping |
| hCoVHKU1 | orf1ab | KFDDYDFSVEDAYAE | 186 | 200 | Overlapping |
| hCoVHKU1 | orf1ab | LAKAMETFNHNNGND | 3304 | 3318 | Overlapping |
| hCoVHKU1 | orf1ab | LNENVVYLFDEAGDE | 894 | 908 | Overlapping |
| hCoVHKU1 | orf1ab | LVDNDLNPFVSDSLV | 6978 | 6992 | Overlapping |
| hCoVHKU1 | orf1ab | LVLDALASMTGVSIE | 3580 | 3594 | Overlapping |
| hCoVHKU1 | orf1ab | LVTDIVVVDEVSMLT | 5750 | 5764 | Overlapping |
| hCoVHKU1 | orf1ab | METFNHNNGNDVLYQ | 3308 | 3322 | Overlapping |
| hCoVHKU1 | orf1ab | NDEEIVTGDNDDQIV | 1086 | 1100 | Overlapping |
| hCoVHKU1 | orf1ab | NDVRDYLLSKMTSLP | 1578 | 1592 | Overlapping |
| hCoVHKU1 | orf1ab | NEPDYSALLCRVTLG | 3384 | 3398 | Overlapping |
| hCoVHKU1 | orf1ab | NFKFDDYDFSVEDAY | 184 | 198 | Overlapping |
| hCoVHKU1 | orf1ab | NLADAKNSGSVNQQQ | 4034 | 4048 | Overlapping |
| hCoVHKU1 | orf1ab | NLEEDVLLFITAFLG | 3794 | 3808 | Overlapping |
| hCoVHKU1 | orf1ab | NNGNDVLYQPPTASV | 3314 | 3328 | Overlapping |
| hCoVHKU1 | orf1ab | NNKEDFDIIQKCQIT | 1506 | 1520 | Overlapping |
| hCoVHKU1 | orf1ab | NSFEAILLNLKLLGI | 3898 | 3912 | Overlapping |
| hCoVHKU1 | orf1ab | PDDVVAANMFMGVGV | 1898 | 1912 | Overlapping |
| hCoVHKU1 | orf1ab | PELVTDIVVVDEVSM | 5748 | 5762 | Overlapping |
| hCoVHKU1 | orf1ab | PIISNEVSINTSCRL | 6284 | 6298 | Overlapping |
| hCoVHKU1 | orf1ab | PIVVAVMDEDIGSTM | 2926 | 2940 | Overlapping |
| hCoVHKU1 | orf1ab | PRSDLERDFIDMDDN | 6742 | 6756 | Overlapping |
| hCoVHKU1 | orf1ab | PTASVSTSFLQSGIV | 3324 | 3338 | Overlapping |
| hCoVHKU1 | orf1ab | PVIEVSQIQSKLTDV | 3916 | 3930 | Overlapping |
| hCoVHKU1 | orf1ab | PVKDYVQTVNVIAWL | 3528 | 3542 | Overlapping |
| hCoVHKU1 | orf1ab | PVLFTNSTDTVNHDS | 410 | 424 | Overlapping |
| hCoVHKU1 | orf1ab | PVNPTDASHYVVTDI | 2556 | 2570 | Overlapping |
| hCoVHKU1 | orf1ab | QSEFVNMASFVEYEV | 4016 | 4030 | Overlapping |
| hCoVHKU1 | orf1ab | QYENLSLADISAVQS | 1704 | 1718 | Overlapping |
| hCoVHKU1 | orf1ab | RTDYVDYTFVNEYYE | 5186 | 5200 | Overlapping |
| hCoVHKU1 | orf1ab | RVEHPDVDGICKLRG | 4400 | 4414 | Overlapping |
| hCoVHKU1 | orf1ab | SDEGGLCPSTGQAME | 34 | 48 | Overlapping |
| hCoVHKU1 | orf1ab | SDGVMKNASLYTSLV | 3008 | 3022 | Overlapping |
| hCoVHKU1 | orf1ab | SIDSDVETRFITKSM | 2716 | 2730 | Overlapping |

| | | | | | |
|---|---|---|---|---|---|
| hCoVHKU1 | orf1ab | SKCWVENDITNGPHE | 5250 | 5264 | Overlapping |
| hCoVHKU1 | orf1ab | SKETAAMVKSKGVCQ | 1410 | 1424 | Overlapping |
| hCoVHKU1 | orf1ab | SNSKLWQYCSVLHNE | 3948 | 3962 | Overlapping |
| hCoVHKU1 | orf1ab | SRMYCTFAIEDVEDV | 912 | 926 | Overlapping |
| hCoVHKU1 | orf1ab | SSEAVEDTIDGVVED | 928 | 942 | Overlapping |
| hCoVHKU1 | orf1ab | SSKSVCTVIDLLLDD | 6824 | 6838 | Overlapping |
| hCoVHKU1 | orf1ab | SVSEESQGNVVTSVM | 2104 | 2118 | Overlapping |
| hCoVHKU1 | orf1ab | TDYNYYKYNLPTMVD | 4904 | 4918 | Overlapping |
| hCoVHKU1 | orf1ab | TEEAFKQSYASATIQ | 5526 | 5540 | Overlapping |
| hCoVHKU1 | orf1ab | TEVDDYVLANECIER | 5500 | 5514 | Overlapping |
| hCoVHKU1 | orf1ab | TGDDVDDIESIYDFD | 1102 | 1116 | Overlapping |
| hCoVHKU1 | orf1ab | TGDNDDEDVVTGDND | 952 | 966 | Overlapping |
| hCoVHKU1 | orf1ab | TGDNDDEDVVTGDNN | 1072 | 1086 | Overlapping |
| hCoVHKU1 | orf1ab | TGDNDDQIVVTGDDV | 1092 | 1106 | Overlapping |
| hCoVHKU1 | orf1ab | TGDNNDEEIVTGDND | 1082 | 1096 | Overlapping |
| hCoVHKU1 | orf1ab | TGDVVLASDDLYVKR | 2044 | 2058 | Overlapping |
| hCoVHKU1 | orf1ab | TGELPCAIMNDKVVV | 6528 | 6542 | Overlapping |
| hCoVHKU1 | orf1ab | TGVDFVVEATGLFAE | 6100 | 6114 | Overlapping |
| hCoVHKU1 | orf1ab | TMLDASAPATGWLLY | 630 | 644 | Overlapping |
| hCoVHKU1 | orf1ab | TNLEVYNKEKTYYEL | 4532 | 4546 | Overlapping |
| hCoVHKU1 | orf1ab | TNSTDTVNHDSFNLY | 414 | 428 | Overlapping |
| hCoVHKU1 | orf1ab | TYCRVGACEYAEEGI | 3056 | 3070 | Overlapping |
| hCoVHKU1 | orf1ab | VDGVYTNFKLVGHDI | 2002 | 2016 | Overlapping |
| hCoVHKU1 | orf1ab | VEDTIDGVVEDTIND | 932 | 946 | Overlapping |
| hCoVHKU1 | orf1ab | VEDTINDDEDVVTGD | 940 | 954 | Overlapping |
| hCoVHKU1 | orf1ab | VHHLSMTTSTTNFGE | 134 | 148 | Overlapping |
| hCoVHKU1 | orf1ab | VISSEAVEDTIDGVV | 926 | 940 | Overlapping |
| hCoVHKU1 | orf1ab | VNPANGHMLHGGGVA | 1384 | 1398 | Overlapping |
| hCoVHKU1 | orf1ab | VPADVSLTYLLGVVD | 1484 | 1498 | Overlapping |
| hCoVHKU1 | orf1ab | VVDEVSMLTNYELSV | 5756 | 5770 | Overlapping |
| hCoVHKU1 | orf1ab | VVMEIPSLMTVKVMF | 820 | 834 | Overlapping |
| hCoVHKU1 | orf1ab | VVMNLDVDTHRYRLS | 4806 | 4820 | Overlapping |
| hCoVHKU1 | orf1ab | VVTGDDVDDIESIYD | 1100 | 1114 | Overlapping |
| hCoVHKU1 | orf1ab | VVTGDNDDEDVVTGD | 950 | 964 | Overlapping |
| hCoVHKU1 | orf1ab | VVTGDNNDEEIVTGD | 1080 | 1094 | Overlapping |
| hCoVHKU1 | orf1ab | VYDDVDASLFVDINN | 2588 | 2602 | Overlapping |
| hCoVHKU1 | orf1ab | VYERDKAVARKLERM | 4062 | 4076 | Overlapping |
| hCoVHKU1 | orf1ab | WQEAWYEFRAGRPHR | 1778 | 1792 | Overlapping |
| hCoVHKU1 | orf1ab | YAEVHAEPKGKYSQK | 198 | 212 | Overlapping |
| hCoVHKU1 | orf1ab | YDFSVEDAYAEVHAE | 190 | 204 | Overlapping |
| hCoVHKU1 | orf1ab | YDSSIHSYFITDQEC | 6808 | 6822 | Overlapping |
| hCoVHKU1 | orf1ab | YIEESDLIISNFKFD | 174 | 188 | Overlapping |
| hCoVHKU1 | orf1ab | YTPHIQISYNDFYAS | 2970 | 2984 | Overlapping |
| hCoVHKU1 | spike | ACHDLLSEYGTFCDN | 828 | 842 | Positive |
| hCoVHKU1 | spike | APKQGYFIKHNDHWM | 1175 | 1189 | Positive |
| hCoVHKU1 | spike | CCTGCGSACFSKCHN | 1323 | 1337 | Positive |
| hCoVHKU1 | spike | CDEYGGHHDFVIKTS | 1339 | 1353 | Positive |
| hCoVHKU1 | spike | CFSKCHNCCDEYGGH | 1331 | 1345 | Positive |
| hCoVHKU1 | spike | CHDLLSEYGTFCDNI | 829 | 843 | Positive |

hCoVHKU1

| | | | | | |
|---|---|---|---|---|---|
| hCoVHKU1 | spike | CHNCCDEYGGHHDFV | 1335 | 1349 | Positive |
| hCoVHKU1 | spike | CLPDPITAYDPRSCS | 520 | 534 | Positive |
| hCoVHKU1 | spike | CPIGTNYRSCESTTV | 495 | 509 | Positive |
| hCoVHKU1 | spike | DALEAQVQIDRLING | 1070 | 1084 | Positive |
| hCoVHKU1 | spike | DCSLFVCSNYAACHD | 817 | 831 | Positive |
| hCoVHKU1 | spike | DFVIKTSHDDGSGSG | 1347 | 1356 | Positive |
| hCoVHKU1 | spike | DHTDWCRCSCLPDPI | 511 | 525 | Positive |
| hCoVHKU1 | spike | DHWMFTGSSYYYPEP | 1186 | 1200 | Positive |
| hCoVHKU1 | spike | DLLQPNTEVFTDVCV | 608 | 622 | Positive |
| hCoVHKU1 | spike | DSIESVGGLYEIKIP | 781 | 795 | Positive |
| hCoVHKU1 | spike | DVGIAPKQGYFIKHN | 1171 | 1185 | Positive |
| hCoVHKU1 | spike | DYNSPSSSSSRRKRR | 746 | 760 | Positive |
| hCoVHKU1 | spike | EPISDKNVVFMNTCS | 1199 | 1213 | Positive |
| hCoVHKU1 | spike | FAINDLNTTVPRISE | 23 | 37 | Positive |
| hCoVHKU1 | spike | FDESKIYGSCFKSIV | 373 | 387 | Positive |
| hCoVHKU1 | spike | FDKSEPLCLFKKNFT | 176 | 190 | Positive |
| hCoVHKU1 | spike | FINNSYTIVVQPHNG | 130 | 144 | Positive |
| hCoVHKU1 | spike | FNVSFVNDSIESVGG | 774 | 788 | Positive |
| hCoVHKU1 | spike | FSKCHNCCDEYGGHH | 1332 | 1346 | Positive |
| hCoVHKU1 | spike | FVCSNYAACHDLLSE | 821 | 835 | Positive |
| hCoVHKU1 | spike | FVIKTSHDDGSGSGS | 1348 | 1356 | Positive |
| hCoVHKU1 | spike | FVNDSIESVGGLYEI | 778 | 792 | Positive |
| hCoVHKU1 | spike | FVTFEPFNVSFVNDS | 768 | 782 | Positive |
| hCoVHKU1 | spike | GHHDFVIKTSHDDGS | 1344 | 1356 | Positive |
| hCoVHKU1 | spike | GPHCGSSSRSFFEDL | 896 | 910 | Positive |
| hCoVHKU1 | spike | GQEEFIQTNSPKVTI | 802 | 816 | Positive |
| hCoVHKU1 | spike | GSACFSKCHNCCDEY | 1328 | 1342 | Positive |
| hCoVHKU1 | spike | GSGMLLIIFILPTTL | 1 | 12 | Positive |
| hCoVHKU1 | spike | GSGSGMLLIIFILPT | 1 | 10 | Positive |
| hCoVHKU1 | spike | GSGSGSGMLLIIFIL | 1 | 8 | Positive |
| hCoVHKU1 | spike | GTFYAYYADSGMPTT | 207 | 221 | Positive |
| hCoVHKU1 | spike | GYFIKHNDHWMFTGS | 1179 | 1193 | Positive |
| hCoVHKU1 | spike | HDFVIKTSHDDGSGS | 1346 | 1356 | Positive |
| hCoVHKU1 | spike | HHDFVIKTSHDDGSG | 1345 | 1356 | Positive |
| hCoVHKU1 | spike | HNCCDEYGGHHDFVI | 1336 | 1350 | Positive |
| hCoVHKU1 | spike | HNDHWMFTGSSYYYP | 1184 | 1198 | Positive |
| hCoVHKU1 | spike | HSVPKLSDFESELSH | 1226 | 1240 | Positive |
| hCoVHKU1 | spike | HTDSFSCNNFDESKI | 364 | 378 | Positive |
| hCoVHKU1 | spike | IAPKQGYFIKHNDHW | 1174 | 1188 | Positive |
| hCoVHKU1 | spike | IESVGGLYEIKIPTN | 783 | 797 | Positive |
| hCoVHKU1 | spike | IKHNDHWMFTGSSYY | 1182 | 1196 | Positive |
| hCoVHKU1 | spike | IKIPTNFTIVGQEEF | 792 | 806 | Positive |
| hCoVHKU1 | spike | ILDEVNGLLDTTQLH | 846 | 860 | Positive |
| hCoVHKU1 | spike | INATFLDLYYEMNLI | 1259 | 1273 | Positive |
| hCoVHKU1 | spike | INVTINNYNPSSWNR | 432 | 446 | Positive |
| hCoVHKU1 | spike | IPDLPDCDIDKWLNN | 321 | 335 | Positive |
| hCoVHKU1 | spike | IPNSRRSDLQLGSSG | 393 | 407 | Positive |
| hCoVHKU1 | spike | IPTNFTIVGQEEFIQ | 794 | 808 | Positive |
| hCoVHKU1 | spike | ISASYRFVTFEPFNV | 762 | 776 | Positive |

| | | | | | |
|---|---|---|---|---|---|
| hCoVHKU1 | spike | ISDKNVVFMNTCSVN | 1201 | 1215 | Positive |
| hCoVHKU1 | spike | IVGQEEFIQTNSPKV | 800 | 814 | Positive |
| hCoVHKU1 | spike | IVVQPHNGVLEITAC | 137 | 151 | Positive |
| hCoVHKU1 | spike | KCHNCCDEYGGHHDF | 1334 | 1348 | Positive |
| hCoVHKU1 | spike | KHNDHWMFTGSSYYY | 1183 | 1197 | Positive |
| hCoVHKU1 | spike | KNQTSIAPNLTLNLH | 1243 | 1257 | Positive |
| hCoVHKU1 | spike | KPSFASSCKSHKPPS | 478 | 492 | Positive |
| hCoVHKU1 | spike | KQGYFIKHNDHWMFT | 1177 | 1191 | Positive |
| hCoVHKU1 | spike | KSEPLCLFKKNFTYN | 178 | 192 | Positive |
| hCoVHKU1 | spike | KSHKPPSASCPIGTN | 486 | 500 | Positive |
| hCoVHKU1 | spike | LDALEAQVQIDRLIN | 1069 | 1083 | Positive |
| hCoVHKU1 | spike | LEITACQYTMCEYPH | 146 | 160 | Positive |
| hCoVHKU1 | spike | LLKFDNRGVITNAVD | 267 | 281 | Positive |
| hCoVHKU1 | spike | LNTTVPRISEYVVDV | 28 | 42 | Positive |
| hCoVHKU1 | spike | NADNLTDYSVSSCAL | 723 | 737 | Positive |
| hCoVHKU1 | spike | NAISSNTDNETLQYW | 243 | 257 | Positive |
| hCoVHKU1 | spike | NALLSIQNGFSATNS | 1014 | 1028 | Positive |
| hCoVHKU1 | spike | NAQALNSLLQQLFNK | 1040 | 1054 | Positive |
| hCoVHKU1 | spike | NATFLDLYYEMNLIQ | 1260 | 1274 | Positive |
| hCoVHKU1 | spike | NAVDCSSSFFSEIQC | 278 | 292 | Positive |
| hCoVHKU1 | spike | NAYVSQQLSDISLVK | 1090 | 1104 | Positive |
| hCoVHKU1 | spike | NCCDEYGGHHDFVIK | 1337 | 1351 | Positive |
| hCoVHKU1 | spike | NDHWMFTGSSYYYPE | 1185 | 1199 | Positive |
| hCoVHKU1 | spike | NDLLQPNTEVFTDVC | 607 | 621 | Positive |
| hCoVHKU1 | spike | NDLNTTVPRISEYVV | 26 | 40 | Positive |
| hCoVHKU1 | spike | NECVKSQSPRINFCG | 1115 | 1129 | Positive |
| hCoVHKU1 | spike | NESWHFDKSEPLCLF | 171 | 185 | Positive |
| hCoVHKU1 | spike | NFDESKIYGSCFKSI | 372 | 386 | Positive |
| hCoVHKU1 | spike | NFILNGINSGTTCSN | 593 | 607 | Positive |
| hCoVHKU1 | spike | NFNLSSHSVVYSRYC | 452 | 466 | Positive |
| hCoVHKU1 | spike | NFNVPSPLNWERKIF | 335 | 349 | Positive |
| hCoVHKU1 | spike | NFRDLSLKGTTYLST | 73 | 87 | Positive |
| hCoVHKU1 | spike | NFTIVGQEEFIQTNS | 797 | 811 | Positive |
| hCoVHKU1 | spike | NGLGVTMDVLNKNQK | 992 | 1006 | Positive |
| hCoVHKU1 | spike | NGLLDTTQLHVADTL | 851 | 865 | Positive |
| hCoVHKU1 | spike | NGRLTALNAYVSQQL | 1083 | 1097 | Positive |
| hCoVHKU1 | spike | NHILSLVQNAPYGLL | 1132 | 1146 | Positive |
| hCoVHKU1 | spike | NHSVPKLSDFESELS | 1225 | 1239 | Positive |
| hCoVHKU1 | spike | NINFKSLVGCLGPHC | 885 | 899 | Positive |
| hCoVHKU1 | spike | NINSILDEVNGLLDT | 842 | 856 | Positive |
| hCoVHKU1 | spike | NKFGAISSSLQEILS | 1053 | 1067 | Positive |
| hCoVHKU1 | spike | NKNQKLIATAFNNAL | 1002 | 1016 | Positive |
| hCoVHKU1 | spike | NKTLYSEFSTIVIGS | 114 | 128 | Positive |
| hCoVHKU1 | spike | NLHFDVDNINFKSLV | 878 | 892 | Positive |
| hCoVHKU1 | spike | NLHTINATFLDLYYE | 1255 | 1269 | Positive |
| hCoVHKU1 | spike | NLIQESIKSLNNSYI | 1271 | 1285 | Positive |
| hCoVHKU1 | spike | NLKCSYVLNNISLTT | 696 | 710 | Positive |
| hCoVHKU1 | spike | NLKDIGTYEMYVKWP | 1286 | 1300 | Positive |
| hCoVHKU1 | spike | NLLYDSNGNIIGFKD | 648 | 662 | Positive |

| | | | | | |
|---|---|---|---|---|---|
| hCoVHKU1 | spike | NLNTNLHFDVDNINF | 874 | 888 | Positive |
| hCoVHKU1 | spike | NLSSHSVVYSRYCFS | 454 | 468 | Positive |
| hCoVHKU1 | spike | NLSTLLRLVHTDSFS | 355 | 369 | Positive |
| hCoVHKU1 | spike | NLTLNLHTINATFLD | 1251 | 1265 | Positive |
| hCoVHKU1 | spike | NNCTGGSEIRDLLCV | 927 | 941 | Positive |
| hCoVHKU1 | spike | NNFDESKIYGSCFKS | 371 | 385 | Positive |
| hCoVHKU1 | spike | NNFNLSSHSVVYSRY | 451 | 465 | Positive |
| hCoVHKU1 | spike | NNISLTTQPYFDSYL | 704 | 718 | Positive |
| hCoVHKU1 | spike | NNRCNIFSNFILNGI | 585 | 599 | Positive |
| hCoVHKU1 | spike | NNSYINLKDIGTYEM | 1281 | 1295 | Positive |
| hCoVHKU1 | spike | NNTFCPCAKPSFASS | 470 | 484 | Positive |
| hCoVHKU1 | spike | NNYNPSSWNRRYGFN | 437 | 451 | Positive |
| hCoVHKU1 | spike | NPSSWNRRYGFNNFN | 440 | 454 | Positive |
| hCoVHKU1 | spike | NQTSIAPNLTLNLHT | 1244 | 1258 | Positive |
| hCoVHKU1 | spike | NRCNIFSNFILNGIN | 586 | 600 | Positive |
| hCoVHKU1 | spike | NRGVITNAVDCSSSF | 272 | 286 | Positive |
| hCoVHKU1 | spike | NRRYGFNNFNLSSHS | 445 | 459 | Positive |
| hCoVHKU1 | spike | NSALAKIQSVVNSNA | 1027 | 1041 | Positive |
| hCoVHKU1 | spike | NSILDEVNGLLDTTQ | 844 | 858 | Positive |
| hCoVHKU1 | spike | NSLLQQLFNKFGAIS | 1045 | 1059 | Positive |
| hCoVHKU1 | spike | NSPKVTIDCSLFVCS | 810 | 824 | Positive |
| hCoVHKU1 | spike | NSRRSDLQLGSSGFL | 395 | 409 | Positive |
| hCoVHKU1 | spike | NSYINLKDIGTYEMY | 1282 | 1296 | Positive |
| hCoVHKU1 | spike | NTDNETLQYWVTPLS | 248 | 262 | Positive |
| hCoVHKU1 | spike | NTEVFTDVCVDYDLY | 613 | 627 | Positive |
| hCoVHKU1 | spike | NTTVPRISEYVVDVS | 29 | 43 | Positive |
| hCoVHKU1 | spike | NVPSPLNWERKIFSN | 337 | 351 | Positive |
| hCoVHKU1 | spike | NVSFVNDSIESVGGL | 775 | 789 | Positive |
| hCoVHKU1 | spike | NVSTDWLYFHFYQER | 192 | 206 | Positive |
| hCoVHKU1 | spike | NVVFMNTCSVNFTKA | 1205 | 1219 | Positive |
| hCoVHKU1 | spike | NWERKIFSNCNFNLS | 343 | 357 | Positive |
| hCoVHKU1 | spike | NYAACHDLLSEYGTF | 825 | 839 | Positive |
| hCoVHKU1 | spike | NYNPSSWNRRYGFNN | 438 | 452 | Positive |
| hCoVHKU1 | spike | NYRSCESTTVLDHTD | 500 | 514 | Positive |
| hCoVHKU1 | spike | PEPISDKNVVFMNTC | 1198 | 1212 | Positive |
| hCoVHKU1 | spike | PFNVSFVNDSIESVG | 773 | 787 | Positive |
| hCoVHKU1 | spike | PHCGSSSRSFFEDLL | 897 | 911 | Positive |
| hCoVHKU1 | spike | PHNGVLEITACQYTM | 141 | 155 | Positive |
| hCoVHKU1 | spike | PIGTNYRSCESTTVL | 496 | 510 | Positive |
| hCoVHKU1 | spike | PISDKNVVFMNTCSV | 1200 | 1214 | Positive |
| hCoVHKU1 | spike | PKQGYFIKHNDHWMF | 1176 | 1190 | Positive |
| hCoVHKU1 | spike | PTNFTIVGQEEFIQT | 795 | 809 | Positive |
| hCoVHKU1 | spike | PTTLAVIGDFNCTNF | 9 | 23 | Positive |
| hCoVHKU1 | spike | QGYFIKHNDHWMFTG | 1178 | 1192 | Positive |
| hCoVHKU1 | spike | RCSCLPDPITAYDPR | 517 | 531 | Positive |
| hCoVHKU1 | spike | RDLLCVQSFNGIKVL | 936 | 950 | Positive |
| hCoVHKU1 | spike | RDLSLKGTTYLSTLW | 75 | 89 | Positive |
| hCoVHKU1 | spike | RFVTFEPFNVSFVND | 767 | 781 | Positive |
| hCoVHKU1 | spike | RGTFYAYYADSGMPT | 206 | 220 | Positive |

| | | | | | |
|---|---|---|---|---|---|
| hCoVHKU1 | spike | RLVHTDSFSCNNFDE | 361 | 375 | Positive |
| hCoVHKU1 | spike | SASYRFVTFEPFNVS | 763 | 777 | Positive |
| hCoVHKU1 | spike | SDFESELSHWFKNQT | 1232 | 1246 | Positive |
| hCoVHKU1 | spike | SDFNNGIFSRVKNTK | 96 | 110 | Positive |
| hCoVHKU1 | spike | SDISLVKFGAALAME | 1098 | 1112 | Positive |
| hCoVHKU1 | spike | SDKNVVFMNTCSVNF | 1202 | 1216 | Positive |
| hCoVHKU1 | spike | SDVGFVEAYNNCTGG | 918 | 932 | Positive |
| hCoVHKU1 | spike | SEFSTIVIGSVFINN | 119 | 133 | Positive |
| hCoVHKU1 | spike | SEIQCKTKSLLPNTG | 288 | 302 | Positive |
| hCoVHKU1 | spike | SELSHWFKNQTSIAP | 1236 | 1250 | Positive |
| hCoVHKU1 | spike | SEPLCLFKKNFTYNV | 179 | 193 | Positive |
| hCoVHKU1 | spike | SESQISGYTTAATVA | 955 | 969 | Positive |
| hCoVHKU1 | spike | SFVNDSIESVGGLYE | 777 | 791 | Positive |
| hCoVHKU1 | spike | SGDVGIAPKQGYFIK | 1169 | 1183 | Positive |
| hCoVHKU1 | spike | SILDEVNGLLDTTQL | 845 | 859 | Positive |
| hCoVHKU1 | spike | SISASYRFVTFEPFN | 761 | 775 | Positive |
| hCoVHKU1 | spike | SKCHNCCDEYGGHHD | 1333 | 1347 | Positive |
| hCoVHKU1 | spike | SLFVCSNYAACHDLL | 819 | 833 | Positive |
| hCoVHKU1 | spike | SNYAACHDLLSEYGT | 824 | 838 | Positive |
| hCoVHKU1 | spike | SSNLNTNLHFDVDNI | 872 | 886 | Positive |
| hCoVHKU1 | spike | STDAFLGWSYDTCVS | 570 | 584 | Positive |
| hCoVHKU1 | spike | STDWLYFHFYQERGT | 194 | 208 | Positive |
| hCoVHKU1 | spike | STIVIGSVFINNSYT | 122 | 136 | Positive |
| hCoVHKU1 | spike | SVFINNSYTIVVQPH | 128 | 142 | Positive |
| hCoVHKU1 | spike | SYYYPEPISDKNVVF | 1194 | 1208 | Positive |
| hCoVHKU1 | spike | TDAFLGWSYDTCVSN | 571 | 585 | Positive |
| hCoVHKU1 | spike | TDNETLQYWVTPLSK | 249 | 263 | Positive |
| hCoVHKU1 | spike | TDSFSCNNFDESKIY | 365 | 379 | Positive |
| hCoVHKU1 | spike | TDVCVDYDLYGITGQ | 618 | 632 | Positive |
| hCoVHKU1 | spike | TDWCRCSCLPDPITA | 513 | 527 | Positive |
| hCoVHKU1 | spike | TDWLYFHFYQERGTF | 195 | 209 | Positive |
| hCoVHKU1 | spike | TDYSVSSCALRMGSG | 728 | 742 | Positive |
| hCoVHKU1 | spike | TEVFTDVCVDYDLYG | 614 | 628 | Positive |
| hCoVHKU1 | spike | TFEPFNVSFVNDSIE | 770 | 784 | Positive |
| hCoVHKU1 | spike | TGGSEIRDLLCVQSF | 930 | 944 | Positive |
| hCoVHKU1 | spike | TGSSYYYPEPISDKN | 1191 | 1205 | Positive |
| hCoVHKU1 | spike | TIDCSLFVCSNYAAC | 815 | 829 | Positive |
| hCoVHKU1 | spike | TIVVQPHNGVLEITA | 136 | 150 | Positive |
| hCoVHKU1 | spike | TLYSEFSTIVIGSVF | 116 | 130 | Positive |
| hCoVHKU1 | spike | TMCEYPHTICKSKGS | 154 | 168 | Positive |
| hCoVHKU1 | spike | TMDVLNKNQKLIATA | 997 | 1011 | Positive |
| hCoVHKU1 | spike | TNFTIVGQEEFIQTN | 796 | 810 | Positive |
| hCoVHKU1 | spike | TVLDHTDWCRCSCLP | 508 | 522 | Positive |
| hCoVHKU1 | spike | VADTLMQGVTLSSNL | 861 | 875 | Positive |
| hCoVHKU1 | spike | VATVHRRIPDLPDCD | 314 | 328 | Positive |
| hCoVHKU1 | spike | VDEEKCGVLDGSYNV | 551 | 565 | Positive |
| hCoVHKU1 | spike | VFINNSYTIVVQPHN | 129 | 143 | Positive |
| hCoVHKU1 | spike | VSFVNDSIESVGGLY | 776 | 790 | Positive |
| hCoVHKU1 | spike | VVQPHNGVLEITACQ | 138 | 152 | Positive |

| | | | | | |
|---|---|---|---|---|---|
| hCoVHKU1 | spike | WNRRYGFNNFNLSSH | 444 | 458 | Positive |
| hCoVHKU1 | spike | YEMNLIQESIKSLNN | 1268 | 1282 | Positive |
| hCoVHKU1 | spike | YFIKHNDHWMFTGSS | 1180 | 1194 | Positive |
| hCoVHKU1 | spike | YGGHHDFVIKTSHDD | 1342 | 1356 | Positive |
| hCoVHKU1 | spike | YLNHSVPKLSDFESE | 1223 | 1237 | Positive |
| hCoVHKU1 | spike | YNSPSSSSSRRKRRS | 747 | 761 | Positive |
| hCoVHKU1 | spike | ADNLTDYSVSSCALR | 724 | 738 | Negative |
| hCoVHKU1 | spike | ADSGMPTTFLFSLYL | 214 | 228 | Negative |
| hCoVHKU1 | spike | AGFGVDEEKCGVLDG | 547 | 561 | Negative |
| hCoVHKU1 | spike | AVDCSSSFFSEIQCK | 279 | 293 | Negative |
| hCoVHKU1 | spike | AVIGDFNCTNFAIND | 13 | 27 | Negative |
| hCoVHKU1 | spike | CEYPHTICKSKGSSR | 156 | 170 | Negative |
| hCoVHKU1 | spike | CGVLDGSYNVSCLCS | 556 | 570 | Negative |
| hCoVHKU1 | spike | CNIFSNFILNGINSG | 588 | 602 | Negative |
| hCoVHKU1 | spike | CQLYYSLPAINVTIN | 423 | 437 | Negative |
| hCoVHKU1 | spike | CSQKKSLVGVGEHCA | 533 | 547 | Negative |
| hCoVHKU1 | spike | CVDYDLYGITGQGIF | 621 | 635 | Negative |
| hCoVHKU1 | spike | CVFNADNLTDYSVSS | 720 | 734 | Negative |
| hCoVHKU1 | spike | DFNCTNFAINDLNTT | 17 | 31 | Negative |
| hCoVHKU1 | spike | DGSYNVSCLCSTDAF | 560 | 574 | Negative |
| hCoVHKU1 | spike | DSYLGCVFNADNLTD | 715 | 729 | Negative |
| hCoVHKU1 | spike | EAQVQIDRLINGRLT | 1073 | 1087 | Negative |
| hCoVHKU1 | spike | ELSHWFKNQTSIAPN | 1237 | 1251 | Negative |
| hCoVHKU1 | spike | FFSEIQCKTKSLLPN | 286 | 300 | Negative |
| hCoVHKU1 | spike | FGVDEEKCGVLDGSY | 549 | 563 | Negative |
| hCoVHKU1 | spike | FNCTNFAINDLNTTV | 18 | 32 | Negative |
| hCoVHKU1 | spike | FRDLSLKGTTYLSTL | 74 | 88 | Negative |
| hCoVHKU1 | spike | FSLNVQYRINGLGVT | 983 | 997 | Negative |
| hCoVHKU1 | spike | GCVFNADNLTDYSVS | 719 | 733 | Negative |
| hCoVHKU1 | spike | GFGVDEEKCGVLDGS | 548 | 562 | Negative |
| hCoVHKU1 | spike | GFNNFNLSSHSVVYS | 449 | 463 | Negative |
| hCoVHKU1 | spike | GGHHDFVIKTSHDDG | 1343 | 1356 | Negative |
| hCoVHKU1 | spike | GGLYEIKIPTNFTIV | 787 | 801 | Negative |
| hCoVHKU1 | spike | GIFKEVSAVYYNSWQ | 633 | 647 | Negative |
| hCoVHKU1 | spike | GINSGTTCSNDLLQP | 598 | 612 | Negative |
| hCoVHKU1 | spike | GITGQGIFKEVSAVY | 628 | 642 | Negative |
| hCoVHKU1 | spike | GLCISGDVGIAPKQG | 1165 | 1179 | Negative |
| hCoVHKU1 | spike | GLGTYYILDRVYLNT | 45 | 59 | Negative |
| hCoVHKU1 | spike | GLGVTMDVLNKNQKL | 993 | 1007 | Negative |
| hCoVHKU1 | spike | GLLDTTQLHVADTLM | 852 | 866 | Negative |
| hCoVHKU1 | spike | GLLFMHFSYKPISFK | 1144 | 1158 | Negative |
| hCoVHKU1 | spike | GLYEIKIPTNFTIVG | 788 | 802 | Negative |
| hCoVHKU1 | spike | GMLLIIFILPTTLAV | 1 | 14 | Negative |
| hCoVHKU1 | spike | GSGSGMLLIIFILPT | 1 | 10 | Negative |
| hCoVHKU1 | spike | GSGSGSGMLLIIFIL | 1 | 8 | Negative |
| hCoVHKU1 | spike | GTNYRSCESTTVLDH | 498 | 512 | Negative |
| hCoVHKU1 | spike | GTTCSNDLLQPNTEV | 602 | 616 | Negative |
| hCoVHKU1 | spike | GTYEMYVKWPWYVWL | 1291 | 1305 | Negative |
| hCoVHKU1 | spike | GVITNAVDCSSSFFS | 274 | 288 | Negative |

| | | | | | |
|---|---|---|---|---|---|
| hCoVHKU1 | spike | GVLDGSYNVSCLCST | 557 | 571 | Negative |
| hCoVHKU1 | spike | GVLEITACQYTMCEY | 144 | 158 | Negative |
| hCoVHKU1 | spike | GVTLSSNLNTNLHFD | 868 | 882 | Negative |
| hCoVHKU1 | spike | GVTMDVLNKNQKLIA | 995 | 1009 | Negative |
| hCoVHKU1 | spike | GVYDLSGFTVKPVAT | 302 | 316 | Negative |
| hCoVHKU1 | spike | HCAGFGVDEEKCGVL | 545 | 559 | Negative |
| hCoVHKU1 | spike | HKPPSASCPIGTNYR | 488 | 502 | Negative |
| hCoVHKU1 | spike | IFSNFILNGINSGTT | 590 | 604 | Negative |
| hCoVHKU1 | spike | INNSYTIVVQPHNGV | 131 | 145 | Negative |
| hCoVHKU1 | spike | KSIVLDKFAIPNSRR | 384 | 398 | Negative |
| hCoVHKU1 | spike | KSLVGVGEHCAGFGV | 537 | 551 | Negative |
| hCoVHKU1 | spike | LLFDKVKLSDVGFVE | 910 | 924 | Negative |
| hCoVHKU1 | spike | LLQPNTEVFTDVCVD | 609 | 623 | Negative |
| hCoVHKU1 | spike | LNVQYRINGLGVTMD | 985 | 999 | Negative |
| hCoVHKU1 | spike | LPLTCNAISSNTDNE | 238 | 252 | Negative |
| hCoVHKU1 | spike | LPTTLAVIGDFNCTN | 8 | 22 | Negative |
| hCoVHKU1 | spike | LSKRQYLLKFDNRGV | 261 | 275 | Negative |
| hCoVHKU1 | spike | LTCNAISSNTDNETL | 240 | 254 | Negative |
| hCoVHKU1 | spike | LTDYSVSSCALRMGS | 727 | 741 | Negative |
| hCoVHKU1 | spike | LWYQKPFLSDFNNGI | 88 | 102 | Negative |
| hCoVHKU1 | spike | MEKVNECVKSQSPRI | 1111 | 1125 | Negative |
| hCoVHKU1 | spike | NCTNFAINDLNTTVP | 19 | 33 | Negative |
| hCoVHKU1 | spike | NGINSGTTCSNDLLQ | 597 | 611 | Negative |
| hCoVHKU1 | spike | NVTINNYNPSSWNRR | 433 | 447 | Negative |
| hCoVHKU1 | spike | PFSLNVQYRINGLGV | 982 | 996 | Negative |
| hCoVHKU1 | spike | PLTCNAISSNTDNET | 239 | 253 | Negative |
| hCoVHKU1 | spike | PNTGVYDLSGFTVKP | 299 | 313 | Negative |
| hCoVHKU1 | spike | PYFDSYLGCVFNADN | 712 | 726 | Negative |
| hCoVHKU1 | spike | QCKTKSLLPNTGVYD | 291 | 305 | Negative |
| hCoVHKU1 | spike | RCNIFSNFILNGINS | 587 | 601 | Negative |
| hCoVHKU1 | spike | RISEYVVDVSYGLGT | 34 | 48 | Negative |
| hCoVHKU1 | spike | RLDALEAQVQIDRLI | 1068 | 1082 | Negative |
| hCoVHKU1 | spike | SFFSEIQCKTKSLLP | 285 | 299 | Negative |
| hCoVHKU1 | spike | SKGSSRNESWHFDKS | 165 | 179 | Negative |
| hCoVHKU1 | spike | SKRQYLLKFDNRGVI | 262 | 276 | Negative |
| hCoVHKU1 | spike | SLNVQYRINGLGVTM | 984 | 998 | Negative |
| hCoVHKU1 | spike | SLVGVGEHCAGFGVD | 538 | 552 | Negative |
| hCoVHKU1 | spike | SRLDALEAQVQIDRL | 1067 | 1081 | Negative |
| hCoVHKU1 | spike | SRRSDLQLGSSGFLQ | 396 | 410 | Negative |
| hCoVHKU1 | spike | SSLQEILSRLDALEA | 1060 | 1074 | Negative |
| hCoVHKU1 | spike | SSRNESWHFDKSEPL | 168 | 182 | Negative |
| hCoVHKU1 | spike | SSRSFFEDLLFDKVK | 902 | 916 | Negative |
| hCoVHKU1 | spike | SSSRRKRRSISASYR | 753 | 767 | Negative |
| hCoVHKU1 | spike | SSSRSFFEDLLFDKV | 901 | 915 | Negative |
| hCoVHKU1 | spike | SSSSRRKRRSISASY | 752 | 766 | Negative |
| hCoVHKU1 | spike | SYTIVVQPHNGVLEI | 134 | 148 | Negative |
| hCoVHKU1 | spike | TQLHVADTLMQGVTL | 857 | 871 | Negative |
| hCoVHKU1 | spike | TSIAPNLTLNLHTIN | 1246 | 1260 | Negative |
| hCoVHKU1 | spike | TTQLHVADTLMQGVT | 856 | 870 | Negative |

| | | | | | |
|---|---|---|---|---|---|
| hCoVHKU1 | spike | TYYILDRVYLNTTIL | 48 | 62 | Negative |
| hCoVHKU1 | spike | VCVDYDLYGITGQGI | 620 | 634 | Negative |
| hCoVHKU1 | spike | VFNADNLTDYSVSSC | 721 | 735 | Negative |
| hCoVHKU1 | spike | VFTDVCVDYDLYGIT | 616 | 630 | Negative |
| hCoVHKU1 | spike | VITNAVDCSSSFFSE | 275 | 289 | Negative |
| hCoVHKU1 | spike | VKSQSPRINFCGNGN | 1118 | 1132 | Negative |
| hCoVHKU1 | spike | VLPLTCNAISSNTDN | 237 | 251 | Negative |
| hCoVHKU1 | spike | VLVSPGLCISGDVGI | 1160 | 1174 | Negative |
| hCoVHKU1 | spike | VQYRINGLGVTMDVL | 987 | 1001 | Negative |
| hCoVHKU1 | spike | YYNSWQNLLYDSNGN | 642 | 656 | Negative |
| hCoVHKU1 | spike | ATFLDLYYEMNLIQE | 1261 | 1275 | Overlapping |
| hCoVHKU1 | spike | CAGFGVDEEKCGVLD | 546 | 560 | Overlapping |
| hCoVHKU1 | spike | CESTTVLDHTDWCRC | 504 | 518 | Overlapping |
| hCoVHKU1 | spike | CGSACFSKCHNCCDE | 1327 | 1341 | Overlapping |
| hCoVHKU1 | spike | DAFLGWSYDTCVSNN | 572 | 586 | Overlapping |
| hCoVHKU1 | spike | DEEKCGVLDGSYNVS | 552 | 566 | Overlapping |
| hCoVHKU1 | spike | DESKIYGSCFKSIVL | 374 | 388 | Overlapping |
| hCoVHKU1 | spike | DEVNGLLDTTQLHVA | 848 | 862 | Overlapping |
| hCoVHKU1 | spike | DEYGGHHDFVIKTSH | 1340 | 1354 | Overlapping |
| hCoVHKU1 | spike | EAYNNCTGGSEIRDL | 924 | 938 | Overlapping |
| hCoVHKU1 | spike | ECVKSQSPRINFCGN | 1116 | 1130 | Overlapping |
| hCoVHKU1 | spike | EDLLFDKVKLSDVGF | 908 | 922 | Overlapping |
| hCoVHKU1 | spike | EEFIQTNSPKVTIDC | 804 | 818 | Overlapping |
| hCoVHKU1 | spike | EEKCGVLDGSYNVSC | 553 | 567 | Overlapping |
| hCoVHKU1 | spike | EFIQTNSPKVTIDCS | 805 | 819 | Overlapping |
| hCoVHKU1 | spike | EFSTIVIGSVFINNS | 120 | 134 | Overlapping |
| hCoVHKU1 | spike | EHCAGFGVDEEKCGV | 544 | 558 | Overlapping |
| hCoVHKU1 | spike | EIKIPTNFTIVGQEE | 791 | 805 | Overlapping |
| hCoVHKU1 | spike | EILSRLDALEAQVQI | 1064 | 1078 | Overlapping |
| hCoVHKU1 | spike | EIQCKTKSLLPNTGV | 289 | 303 | Overlapping |
| hCoVHKU1 | spike | EIRDLLCVQSFNGIK | 934 | 948 | Overlapping |
| hCoVHKU1 | spike | EITACQYTMCEYPHT | 147 | 161 | Overlapping |
| hCoVHKU1 | spike | EKCGVLDGSYNVSCL | 554 | 568 | Overlapping |
| hCoVHKU1 | spike | EKVNECVKSQSPRIN | 1112 | 1126 | Overlapping |
| hCoVHKU1 | spike | ELSHWFKNQTSIAPN | 1237 | 1251 | Overlapping |
| hCoVHKU1 | spike | EMNLIQESIKSLNNS | 1269 | 1283 | Overlapping |
| hCoVHKU1 | spike | EMYVKWPWYVWLLIS | 1294 | 1308 | Overlapping |
| hCoVHKU1 | spike | EPFNVSFVNDSIESV | 772 | 786 | Overlapping |
| hCoVHKU1 | spike | EPLCLFKKNFTYNVS | 180 | 194 | Overlapping |
| hCoVHKU1 | spike | ERGTFYAYYADSGMP | 205 | 219 | Overlapping |
| hCoVHKU1 | spike | ERKIFSNCNFNLSTL | 345 | 359 | Overlapping |
| hCoVHKU1 | spike | ESELSHWFKNQTSIA | 1235 | 1249 | Overlapping |
| hCoVHKU1 | spike | ESIKSLNNSYINLKD | 1275 | 1289 | Overlapping |
| hCoVHKU1 | spike | ESKIYGSCFKSIVLD | 375 | 389 | Overlapping |
| hCoVHKU1 | spike | ESQISGYTTAATVAA | 956 | 970 | Overlapping |
| hCoVHKU1 | spike | ESTTVLDHTDWCRCS | 505 | 519 | Overlapping |
| hCoVHKU1 | spike | ESVGGLYEIKIPTNF | 784 | 798 | Overlapping |
| hCoVHKU1 | spike | ESWHFDKSEPLCLFK | 172 | 186 | Overlapping |
| hCoVHKU1 | spike | ETLQYWVTPLSKRQY | 252 | 266 | Overlapping |

| | | | | | |
|---|---|---|---|---|---|
| hCoVHKU1 | spike | EVFTDVCVDYDLYGI | 615 | 629 | Overlapping |
| hCoVHKU1 | spike | EVNGLLDTTQLHVAD | 849 | 863 | Overlapping |
| hCoVHKU1 | spike | EVSAVYYNSWQNLLY | 637 | 651 | Overlapping |
| hCoVHKU1 | spike | EYGGHHDFVIKTSHD | 1341 | 1355 | Overlapping |
| hCoVHKU1 | spike | EYGTFCDNINSILDE | 835 | 849 | Overlapping |
| hCoVHKU1 | spike | EYPHTICKSKGSSRN | 157 | 171 | Overlapping |
| hCoVHKU1 | spike | EYVVDVSYGLGTYYI | 37 | 51 | Overlapping |
| hCoVHKU1 | spike | FEDLLFDKVKLSDVG | 907 | 921 | Overlapping |
| hCoVHKU1 | spike | FEPFNVSFVNDSIES | 771 | 785 | Overlapping |
| hCoVHKU1 | spike | FESELSHWFKNQTSI | 1234 | 1248 | Overlapping |
| hCoVHKU1 | spike | FVNDSIESVGGLYEI | 778 | 792 | Overlapping |
| hCoVHKU1 | spike | GEHCAGFGVDEEKCG | 543 | 557 | Overlapping |
| hCoVHKU1 | spike | GFVEAYNNCTGGSEI | 921 | 935 | Overlapping |
| hCoVHKU1 | spike | GGHHDFVIKTSHDDG | 1343 | 1356 | Overlapping |
| hCoVHKU1 | spike | GIAPKQGYFIKHNDH | 1173 | 1187 | Overlapping |
| hCoVHKU1 | spike | GIPFSLNVQYRINGL | 980 | 994 | Overlapping |
| hCoVHKU1 | spike | GMPTTFLFSLYLGTL | 217 | 231 | Overlapping |
| hCoVHKU1 | spike | GTFCDNINSILDEVN | 837 | 851 | Overlapping |
| hCoVHKU1 | spike | GVDEEKCGVLDGSYN | 550 | 564 | Overlapping |
| hCoVHKU1 | spike | GVGEHCAGFGVDEEK | 541 | 555 | Overlapping |
| hCoVHKU1 | spike | IESVGGLYEIKIPTN | 783 | 797 | Overlapping |
| hCoVHKU1 | spike | KEVSAVYYNSWQNLL | 636 | 650 | Overlapping |
| hCoVHKU1 | spike | KSQSPRINFCGNGNH | 1119 | 1133 | Overlapping |
| hCoVHKU1 | spike | LVGVGEHCAGFGVDE | 539 | 553 | Overlapping |
| hCoVHKU1 | spike | NDSIESVGGLYEIKI | 780 | 794 | Overlapping |
| hCoVHKU1 | spike | NNSYTIVVQPHNGVL | 132 | 146 | Overlapping |
| hCoVHKU1 | spike | NSRRSDLQLGSSGFL | 395 | 409 | Overlapping |
| hCoVHKU1 | spike | NSYTIVVQPHNGVLE | 133 | 147 | Overlapping |
| hCoVHKU1 | spike | PILSESQISGYTTAA | 952 | 966 | Overlapping |
| hCoVHKU1 | spike | PVATVHRRIPDLPDC | 313 | 327 | Overlapping |
| hCoVHKU1 | spike | QEEFIQTNSPKVTID | 803 | 817 | Overlapping |
| hCoVHKU1 | spike | QERGTFYAYYADSGM | 204 | 218 | Overlapping |
| hCoVHKU1 | spike | QESIKSLNNSYINLK | 1274 | 1288 | Overlapping |
| hCoVHKU1 | spike | RRSISASYRFVTFEP | 759 | 773 | Overlapping |
| hCoVHKU1 | spike | SEIRDLLCVQSFNGI | 933 | 947 | Overlapping |
| hCoVHKU1 | spike | SEYGTFCDNINSILD | 834 | 848 | Overlapping |
| hCoVHKU1 | spike | SEYVVDVSYGLGTYY | 36 | 50 | Overlapping |
| hCoVHKU1 | spike | SFVNDSIESVGGLYE | 777 | 791 | Overlapping |
| hCoVHKU1 | spike | SRRKRRSISASYRFV | 755 | 769 | Overlapping |
| hCoVHKU1 | spike | SSRRKRRSISASYRF | 754 | 768 | Overlapping |
| hCoVHKU1 | spike | TLMQGVTLSSNLNTN | 864 | 878 | Overlapping |
| hCoVHKU1 | spike | VEAYNNCTGGSEIRD | 923 | 937 | Overlapping |
| hCoVHKU1 | spike | VGEHCAGFGVDEEKC | 542 | 556 | Overlapping |
| hCoVHKU1 | spike | VGVGEHCAGFGVDEE | 540 | 554 | Overlapping |
| hCoVHKU1 | spike | VIKTSHDDGSGSGSG | 1349 | 1356 | Overlapping |
| hCoVHKU1 | spike | YEIKIPTNFTIVGQE | 790 | 804 | Overlapping |
| hCoVHKU1 | spike | YGGHHDFVIKTSHDD | 1342 | 1356 | Overlapping |
| hCoVHKU1 | spike | YGTFCDNINSILDEV | 836 | 850 | Overlapping |

**Table S5. NL63-specific Peptides**

| Lineage | Protein | HDPA peptide sequence | AA pos start | AA pos end | Group |
|---|---|---|---|---|---|
| hCoVNL63 | envelope | APVPAEVLNVGSGSG | 68 | 77 | Positive |
| hCoVNL63 | envelope | AYQDYMQIAPVPAEV | 60 | 74 | Positive |
| hCoVNL63 | envelope | GSGSGMFLRLIDDNG | 1 | 10 | Positive |
| hCoVNL63 | envelope | GSGSGSGMFLRLIDD | 1 | 8 | Positive |
| hCoVNL63 | envelope | PVPAEVLNVGSGSGS | 69 | 77 | Positive |
| hCoVNL63 | envelope | RLIDDNGIVLNSILW | 4 | 18 | Positive |
| hCoVNL63 | envelope | VPAEVLNVGSGSGSG | 70 | 77 | Positive |
| hCoVNL63 | envelope | APVPAEVLNVGSGSG | 68 | 77 | Negative |
| hCoVNL63 | envelope | FSRTLYQPVYKIFLA | 46 | 60 | Negative |
| hCoVNL63 | envelope | GIVLNSILWLLVMIF | 10 | 24 | Negative |
| hCoVNL63 | envelope | GSGMFLRLIDDNGIV | 1 | 12 | Negative |
| hCoVNL63 | envelope | GSGSGMFLRLIDDNG | 1 | 10 | Negative |
| hCoVNL63 | envelope | GSGSGSGMFLRLIDD | 1 | 8 | Negative |
| hCoVNL63 | envelope | IAPVPAEVLNVGSGS | 67 | 77 | Negative |
| hCoVNL63 | envelope | IDDNGIVLNSILWLL | 6 | 20 | Negative |
| hCoVNL63 | envelope | MFLRLIDDNGIVLNS | 1 | 15 | Negative |
| hCoVNL63 | envelope | PVYKIFLAYQDYMQI | 53 | 67 | Negative |
| hCoVNL63 | envelope | SGMFLRLIDDNGIVL | 1 | 13 | Negative |
| hCoVNL63 | envelope | SGSGMFLRLIDDNGI | 1 | 11 | Negative |
| hCoVNL63 | envelope | GMFLRLIDDNGIVLN | 1 | 14 | Overlapping |
| hCoVNL63 | envelope | GSGMFLRLIDDNGIV | 1 | 12 | Overlapping |
| hCoVNL63 | envelope | PVPAEVLNVGSGSGS | 69 | 77 | Overlapping |
| hCoVNL63 | membrane | DGHKIATRVQVGQLP | 152 | 166 | Positive |
| hCoVNL63 | membrane | GHKIATRVQVGQLPK | 153 | 167 | Positive |
| hCoVNL63 | membrane | GHYKYSRLLYGLKMS | 36 | 50 | Positive |
| hCoVNL63 | membrane | GSGMSNSSVPLLEVY | 1 | 12 | Positive |
| hCoVNL63 | membrane | GSGSGMSNSSVPLLE | 1 | 10 | Positive |
| hCoVNL63 | membrane | GSGSGSGMSNSSVPL | 1 | 8 | Positive |
| hCoVNL63 | membrane | GVASQEGVLSEREKL | 208 | 222 | Positive |
| hCoVNL63 | membrane | GWAFYVRAKHGDFSG | 194 | 208 | Positive |
| hCoVNL63 | membrane | LEVYVHLRNWNFSWN | 9 | 23 | Positive |
| hCoVNL63 | membrane | LVDGHKIATRVQVGQ | 150 | 164 | Positive |
| hCoVNL63 | membrane | NAIISLQVYGHNYYL | 117 | 131 | Positive |
| hCoVNL63 | membrane | NFNVDWVFFGFSILM | 70 | 84 | Positive |
| hCoVNL63 | membrane | NLILTLFIVVLQYGH | 23 | 37 | Positive |
| hCoVNL63 | membrane | NSSVPLLEVYVHLRN | 3 | 17 | Positive |
| hCoVNL63 | membrane | NWNFSWNLILTLFIV | 17 | 31 | Positive |
| hCoVNL63 | membrane | PLLEVYVHLRNWNFS | 7 | 21 | Positive |
| hCoVNL63 | membrane | PTGVTLTLLSGVLLV | 137 | 151 | Positive |
| hCoVNL63 | membrane | SGVASQEGVLSEREK | 207 | 221 | Positive |
| hCoVNL63 | membrane | SNSSVPLLEVYVHLR | 2 | 16 | Positive |
| hCoVNL63 | membrane | TGWAFYVRAKHGDFS | 193 | 207 | Positive |
| hCoVNL63 | membrane | VASQEGVLSEREKLL | 209 | 223 | Positive |
| hCoVNL63 | membrane | VDGHKIATRVQVGQL | 151 | 165 | Positive |
| hCoVNL63 | membrane | EGVLSEREKLLHLIG | 213 | 226 | Negative |
| hCoVNL63 | membrane | GDFSGVASQEGVLSE | 204 | 218 | Negative |
| hCoVNL63 | membrane | GRSVNETSQTGWAFY | 184 | 198 | Negative |

| | | | | | |
|---|---|---|---|---|---|
| hCoVNL63 | membrane | GSGMSNSSVPLLEVY | 1 | 12 | Negative |
| hCoVNL63 | membrane | GSGSGMSNSSVPLLE | 1 | 10 | Negative |
| hCoVNL63 | membrane | GSGSGSGMSNSSVPL | 1 | 8 | Negative |
| hCoVNL63 | membrane | GVASQEGVLSEREKL | 208 | 222 | Negative |
| hCoVNL63 | membrane | GVLLVDGHKIATRVQ | 147 | 161 | Negative |
| hCoVNL63 | membrane | GVLSEREKLLHLIGS | 214 | 226 | Negative |
| hCoVNL63 | membrane | GVTLTLLSGVLLVDG | 139 | 153 | Negative |
| hCoVNL63 | membrane | MAAPTGVTLTLLSGV | 134 | 148 | Negative |
| hCoVNL63 | membrane | NETSQTGWAFYVRAK | 188 | 202 | Negative |
| hCoVNL63 | membrane | NSFRLWRRVKTFWAF | 98 | 112 | Negative |
| hCoVNL63 | membrane | SGSGMSNSSVPLLEV | 1 | 11 | Negative |
| hCoVNL63 | membrane | SGSGSGMSNSSVPLL | 1 | 9 | Negative |
| hCoVNL63 | membrane | SSVPLLEVYVHLRNW | 4 | 18 | Negative |
| hCoVNL63 | membrane | WAFYVRAKHGDFSGV | 195 | 209 | Negative |
| hCoVNL63 | membrane | EGVLSEREKLLHLIG | 213 | 226 | Overlapping |
| hCoVNL63 | membrane | EREKLLHLIGSGSGS | 218 | 226 | Overlapping |
| hCoVNL63 | membrane | ETNAIISLQVYGHNY | 115 | 129 | Overlapping |
| hCoVNL63 | membrane | ETSQTGWAFYVRAKH | 189 | 203 | Overlapping |
| hCoVNL63 | membrane | EVYVHLRNWNFSWNL | 10 | 24 | Overlapping |
| hCoVNL63 | membrane | FSGVASQEGVLSERE | 206 | 220 | Overlapping |
| hCoVNL63 | membrane | GHNYYLPVMAAPTGV | 126 | 140 | Overlapping |
| hCoVNL63 | membrane | GMSNSSVPLLEVYVH | 1 | 14 | Overlapping |
| hCoVNL63 | membrane | NYYLPVMAAPTGVTL | 128 | 142 | Overlapping |
| hCoVNL63 | membrane | PETNAIISLQVYGHN | 114 | 128 | Overlapping |
| hCoVNL63 | membrane | PVMAAPTGVTLTLLS | 132 | 146 | Overlapping |
| hCoVNL63 | membrane | QEGVLSEREKLLHLI | 212 | 226 | Overlapping |
| hCoVNL63 | membrane | SEREKLLHLIGSGSG | 217 | 226 | Overlapping |
| hCoVNL63 | membrane | SNSSVPLLEVYVHLR | 2 | 16 | Overlapping |
| hCoVNL63 | nucleocapsid | AALFFDSEVSTDEVG | 285 | 299 | Positive |
| hCoVNL63 | nucleocapsid | ADDDSAIIEIVNEVL | 362 | 376 | Positive |
| hCoVNL63 | nucleocapsid | ADDRAARKKFPPPSF | 7 | 21 | Positive |
| hCoVNL63 | nucleocapsid | AQNTVLNASIPESKP | 346 | 360 | Positive |
| hCoVNL63 | nucleocapsid | ASIPESKPLADDDSA | 353 | 367 | Positive |
| hCoVNL63 | nucleocapsid | DEQIGYWNVQERWRM | 50 | 64 | Positive |
| hCoVNL63 | nucleocapsid | ENVIQCFGPRDFNHN | 244 | 258 | Positive |
| hCoVNL63 | nucleocapsid | FIEQISAFTKPSSIK | 322 | 336 | Positive |
| hCoVNL63 | nucleocapsid | FSIALPPELSVVEFE | 127 | 141 | Positive |
| hCoVNL63 | nucleocapsid | GPHKDLKFRQRSDGV | 84 | 98 | Positive |
| hCoVNL63 | nucleocapsid | GPRDFNHNMGDSDLV | 251 | 265 | Positive |
| hCoVNL63 | nucleocapsid | GSGMASVNWADDRAA | 1 | 12 | Positive |
| hCoVNL63 | nucleocapsid | GSGSGMASVNWADDR | 1 | 10 | Positive |
| hCoVNL63 | nucleocapsid | GSGSGSGMASVNWAD | 1 | 8 | Positive |
| hCoVNL63 | nucleocapsid | IEIVNEVLHGSGSGS | 369 | 377 | Positive |
| hCoVNL63 | nucleocapsid | IPESKPLADDDSAII | 355 | 369 | Positive |
| hCoVNL63 | nucleocapsid | IPNQAALFFDSEVST | 281 | 295 | Positive |
| hCoVNL63 | nucleocapsid | ISAFTKPSSIKEMQS | 326 | 340 | Positive |
| hCoVNL63 | nucleocapsid | KDEQIGYWNVQERWR | 49 | 63 | Positive |
| hCoVNL63 | nucleocapsid | KDNKNLPKFIEQISA | 314 | 328 | Positive |
| hCoVNL63 | nucleocapsid | KGFPQLAELIPNQAA | 272 | 286 | Positive |

| | | | | | |
|---|---|---|---|---|---|
| hCoVNL63 | nucleocapsid | KGNKDEQIGYWNVQE | 46 | 60 | Positive |
| hCoVNL63 | nucleocapsid | KKPNKPLSQPRADKP | 214 | 228 | Positive |
| hCoVNL63 | nucleocapsid | KPLADDDSAIIEIVN | 359 | 373 | Positive |
| hCoVNL63 | nucleocapsid | KPLEPKFSIALPPEL | 121 | 135 | Positive |
| hCoVNL63 | nucleocapsid | KVHFYYLGTGPHKDL | 75 | 89 | Positive |
| hCoVNL63 | nucleocapsid | LADDDSAIIEIVNEV | 361 | 375 | Positive |
| hCoVNL63 | nucleocapsid | LEPKFSIALPPELSV | 123 | 137 | Positive |
| hCoVNL63 | nucleocapsid | LNASIPESKPLADDD | 351 | 365 | Positive |
| hCoVNL63 | nucleocapsid | LPPELSVVEFEDRSN | 131 | 145 | Positive |
| hCoVNL63 | nucleocapsid | NASIPESKPLADDDS | 352 | 366 | Positive |
| hCoVNL63 | nucleocapsid | NKNLPKFIEQISAFT | 316 | 330 | Positive |
| hCoVNL63 | nucleocapsid | NKPLSQPRADKPSQL | 217 | 231 | Positive |
| hCoVNL63 | nucleocapsid | NLGFDNQSKSPSSSG | 195 | 209 | Positive |
| hCoVNL63 | nucleocapsid | NLPKFIEQISAFTKP | 318 | 332 | Positive |
| hCoVNL63 | nucleocapsid | NLVPIGKGNKDEQIG | 40 | 54 | Positive |
| hCoVNL63 | nucleocapsid | NNSSRASSRSSTRNN | 145 | 159 | Positive |
| hCoVNL63 | nucleocapsid | NQAALFFDSEVSTDE | 283 | 297 | Positive |
| hCoVNL63 | nucleocapsid | NQKPLEPKFSIALPP | 119 | 133 | Positive |
| hCoVNL63 | nucleocapsid | NQSKSPSSSGTSTPK | 200 | 214 | Positive |
| hCoVNL63 | nucleocapsid | NQSSSDLVAAVTLAL | 179 | 193 | Positive |
| hCoVNL63 | nucleocapsid | NRKRNQKPLEPKFSI | 115 | 129 | Positive |
| hCoVNL63 | nucleocapsid | NSRDSSRSTSRQQSR | 159 | 173 | Positive |
| hCoVNL63 | nucleocapsid | NSSRASSRSSTRNNS | 146 | 160 | Positive |
| hCoVNL63 | nucleocapsid | NVIQCFGPRDFNHNM | 245 | 259 | Positive |
| hCoVNL63 | nucleocapsid | PESKPLADDDSAIIE | 356 | 370 | Positive |
| hCoVNL63 | nucleocapsid | PKVHFYYLGTGPHKD | 74 | 88 | Positive |
| hCoVNL63 | nucleocapsid | PPELSVVEFEDRSNN | 132 | 146 | Positive |
| hCoVNL63 | nucleocapsid | PRDFNHNMGDSDLVQ | 252 | 266 | Positive |
| hCoVNL63 | nucleocapsid | QCFGPRDFNHNMGDS | 248 | 262 | Positive |
| hCoVNL63 | nucleocapsid | RNNSRDSSRSTSRQQ | 157 | 171 | Positive |
| hCoVNL63 | nucleocapsid | RSDGVVWVAKEGAKT | 94 | 108 | Positive |
| hCoVNL63 | nucleocapsid | SAFTKPSSIKEMQSQ | 327 | 341 | Positive |
| hCoVNL63 | nucleocapsid | SDGVVWVAKEGAKTV | 95 | 109 | Positive |
| hCoVNL63 | nucleocapsid | SDKAPYRVIPRNLVP | 29 | 43 | Positive |
| hCoVNL63 | nucleocapsid | SDLVAAVTLALKNLG | 183 | 197 | Positive |
| hCoVNL63 | nucleocapsid | SDSNQSSSDLVAAVT | 176 | 190 | Positive |
| hCoVNL63 | nucleocapsid | SIKEMQSQSSHVAQN | 334 | 348 | Positive |
| hCoVNL63 | nucleocapsid | SIPESKPLADDDSAI | 354 | 368 | Positive |
| hCoVNL63 | nucleocapsid | SKPLADDDSAIIEIV | 358 | 372 | Positive |
| hCoVNL63 | nucleocapsid | SSDKAPYRVIPRNLV | 28 | 42 | Positive |
| hCoVNL63 | nucleocapsid | SSDLVAAVTLALKNL | 182 | 196 | Positive |
| hCoVNL63 | nucleocapsid | STDEVGDNVQITYTY | 294 | 308 | Positive |
| hCoVNL63 | nucleocapsid | TGPHKDLKFRQRSDG | 83 | 97 | Positive |
| hCoVNL63 | nucleocapsid | TVLNASIPESKPLAD | 349 | 363 | Positive |
| hCoVNL63 | nucleocapsid | VAKEGAKTVNTSLGN | 101 | 115 | Positive |
| hCoVNL63 | nucleocapsid | VDLPPKVHFYYLGTG | 70 | 84 | Positive |
| hCoVNL63 | nucleocapsid | VLNASIPESKPLADD | 350 | 364 | Positive |
| hCoVNL63 | nucleocapsid | VVEFEDRSNNSSRAS | 137 | 151 | Positive |
| hCoVNL63 | nucleocapsid | AELIPNQAALFFDSE | 278 | 292 | Negative |

| | | | | | |
|---|---|---|---|---|---|
| hCoVNL63 | nucleocapsid | ALFFDSEVSTDEVGD | 286 | 300 | Negative |
| hCoVNL63 | nucleocapsid | ASSRSSTRNNSRDSS | 150 | 164 | Negative |
| hCoVNL63 | nucleocapsid | DDSAIIEIVNEVLHG | 364 | 377 | Negative |
| hCoVNL63 | nucleocapsid | DLVQNGVDAKGFPQL | 263 | 277 | Negative |
| hCoVNL63 | nucleocapsid | DSSRSTSRQQSRTRS | 162 | 176 | Negative |
| hCoVNL63 | nucleocapsid | EGAKTVNTSLGNRKR | 104 | 118 | Negative |
| hCoVNL63 | nucleocapsid | EIVNEVLHGSGSGSG | 370 | 377 | Negative |
| hCoVNL63 | nucleocapsid | ELIPNQAALFFDSEV | 279 | 293 | Negative |
| hCoVNL63 | nucleocapsid | ELSVVEFEDRSNNSS | 134 | 148 | Negative |
| hCoVNL63 | nucleocapsid | EMQSQSSHVAQNTVL | 337 | 351 | Negative |
| hCoVNL63 | nucleocapsid | EVGDNVQITYTYKML | 297 | 311 | Negative |
| hCoVNL63 | nucleocapsid | FDSEVSTDEVGDNVQ | 289 | 303 | Negative |
| hCoVNL63 | nucleocapsid | FFDSEVSTDEVGDNV | 288 | 302 | Negative |
| hCoVNL63 | nucleocapsid | FNHNMGDSDLVQNGV | 255 | 269 | Negative |
| hCoVNL63 | nucleocapsid | FRQRSDGVVWVAKEG | 91 | 105 | Negative |
| hCoVNL63 | nucleocapsid | FTKPSSIKEMQSQSS | 329 | 343 | Negative |
| hCoVNL63 | nucleocapsid | GDSDLVQNGVDAKGF | 260 | 274 | Negative |
| hCoVNL63 | nucleocapsid | GFDNQSKSPSSSGTS | 197 | 211 | Negative |
| hCoVNL63 | nucleocapsid | GFPQLAELIPNQAAL | 273 | 287 | Negative |
| hCoVNL63 | nucleocapsid | GKGNKDEQIGYWNVQ | 45 | 59 | Negative |
| hCoVNL63 | nucleocapsid | GMASVNWADDRAARK | 1 | 14 | Negative |
| hCoVNL63 | nucleocapsid | GSGMASVNWADDRAA | 1 | 12 | Negative |
| hCoVNL63 | nucleocapsid | GSGSGMASVNWADDR | 1 | 10 | Negative |
| hCoVNL63 | nucleocapsid | GSGSGSGMASVNWAD | 1 | 8 | Negative |
| hCoVNL63 | nucleocapsid | GTGPHKDLKFRQRSD | 82 | 96 | Negative |
| hCoVNL63 | nucleocapsid | GVDAKGFPQLAELIP | 268 | 282 | Negative |
| hCoVNL63 | nucleocapsid | GVVWVAKEGAKTVNT | 97 | 111 | Negative |
| hCoVNL63 | nucleocapsid | GYWNVQERWRMRRGQ | 54 | 68 | Negative |
| hCoVNL63 | nucleocapsid | HNMGDSDLVQNGVDA | 257 | 271 | Negative |
| hCoVNL63 | nucleocapsid | IGYWNVQERWRMRRG | 53 | 67 | Negative |
| hCoVNL63 | nucleocapsid | IIEIVNEVLHGSGSG | 368 | 377 | Negative |
| hCoVNL63 | nucleocapsid | KEGAKTVNTSLGNRK | 103 | 117 | Negative |
| hCoVNL63 | nucleocapsid | LFFDSEVSTDEVGDN | 287 | 301 | Negative |
| hCoVNL63 | nucleocapsid | LIPNQAALFFDSEVS | 280 | 294 | Negative |
| hCoVNL63 | nucleocapsid | LSVVEFEDRSNNSSR | 135 | 149 | Negative |
| hCoVNL63 | nucleocapsid | LVQNGVDAKGFPQLA | 264 | 278 | Negative |
| hCoVNL63 | nucleocapsid | PHKDLKFRQRSDGVV | 85 | 99 | Negative |
| hCoVNL63 | nucleocapsid | QNGVDAKGFPQLAEL | 266 | 280 | Negative |
| hCoVNL63 | nucleocapsid | QSRTRSDSNQSSSDL | 171 | 185 | Negative |
| hCoVNL63 | nucleocapsid | SKPLADDDSAIIEIV | 358 | 372 | Negative |
| hCoVNL63 | nucleocapsid | SKSPSSSGTSTPKKP | 202 | 216 | Negative |
| hCoVNL63 | nucleocapsid | SRASSRSSTRNNSRD | 148 | 162 | Negative |
| hCoVNL63 | nucleocapsid | SRDSSRSTSRQQSRT | 160 | 174 | Negative |
| hCoVNL63 | nucleocapsid | SRTRSDSNQSSSDLV | 172 | 186 | Negative |
| hCoVNL63 | nucleocapsid | SSIKEMQSQSSHVAQ | 333 | 347 | Negative |
| hCoVNL63 | nucleocapsid | SSRSTSRQQSRTRSD | 163 | 177 | Negative |
| hCoVNL63 | nucleocapsid | SSTRNNSRDSSRSTS | 154 | 168 | Negative |
| hCoVNL63 | nucleocapsid | STRNNSRDSSRSTSR | 155 | 169 | Negative |
| hCoVNL63 | nucleocapsid | SVVEFEDRSNNSSRA | 136 | 150 | Negative |

| | | | | | |
|---|---|---|---|---|---|
| hCoVNL63 | nucleocapsid | TSRQQSRTRSDSNQS | 167 | 181 | Negative |
| hCoVNL63 | nucleocapsid | VEFEDRSNNSSRASS | 138 | 152 | Negative |
| hCoVNL63 | nucleocapsid | VQNGVDAKGFPQLAE | 265 | 279 | Negative |
| hCoVNL63 | nucleocapsid | DDDSAIIEIVNEVLH | 363 | 377 | Overlapping |
| hCoVNL63 | nucleocapsid | DEVGDNVQITYTYKM | 296 | 310 | Overlapping |
| hCoVNL63 | nucleocapsid | DGVVWVAKEGAKTVN | 96 | 110 | Overlapping |
| hCoVNL63 | nucleocapsid | DSEVSTDEVGDNVQI | 290 | 304 | Overlapping |
| hCoVNL63 | nucleocapsid | EDRSNNSSRASSRSS | 141 | 155 | Overlapping |
| hCoVNL63 | nucleocapsid | EENVIQCFGPRDFNH | 243 | 257 | Overlapping |
| hCoVNL63 | nucleocapsid | EFEDRSNNSSRASSR | 139 | 153 | Overlapping |
| hCoVNL63 | nucleocapsid | EGAKTVNTSLGNRKR | 104 | 118 | Overlapping |
| hCoVNL63 | nucleocapsid | ELIPNQAALFFDSEV | 279 | 293 | Overlapping |
| hCoVNL63 | nucleocapsid | ELSVVEFEDRSNNSS | 134 | 148 | Overlapping |
| hCoVNL63 | nucleocapsid | EMQSQSSHVAQNTVL | 337 | 351 | Overlapping |
| hCoVNL63 | nucleocapsid | EPKFSIALPPELSVV | 124 | 138 | Overlapping |
| hCoVNL63 | nucleocapsid | EQIGYWNVQERWRMR | 51 | 65 | Overlapping |
| hCoVNL63 | nucleocapsid | EQISAFTKPSSIKEM | 324 | 338 | Overlapping |
| hCoVNL63 | nucleocapsid | ERWRMRRGQRVDLPP | 60 | 74 | Overlapping |
| hCoVNL63 | nucleocapsid | ESKPLADDDSAIIEI | 357 | 371 | Overlapping |
| hCoVNL63 | nucleocapsid | EVGDNVQITYTYKML | 297 | 311 | Overlapping |
| hCoVNL63 | nucleocapsid | EVSTDEVGDNVQITY | 292 | 306 | Overlapping |
| hCoVNL63 | nucleocapsid | FDSEVSTDEVGDNVQ | 289 | 303 | Overlapping |
| hCoVNL63 | nucleocapsid | FFDSEVSTDEVGDNV | 288 | 302 | Overlapping |
| hCoVNL63 | nucleocapsid | IEQISAFTKPSSIKE | 323 | 337 | Overlapping |
| hCoVNL63 | nucleocapsid | KEMQSQSSHVAQNTV | 336 | 350 | Overlapping |
| hCoVNL63 | nucleocapsid | NHNMGDSDLVQNGVD | 256 | 270 | Overlapping |
| hCoVNL63 | nucleocapsid | NKDEQIGYWNVQERW | 48 | 62 | Overlapping |
| hCoVNL63 | nucleocapsid | NMGDSDLVQNGVDAK | 258 | 272 | Overlapping |
| hCoVNL63 | nucleocapsid | NNSRDSSRSTSRQQS | 158 | 172 | Overlapping |
| hCoVNL63 | nucleocapsid | NNSSRASSRSSTRNN | 145 | 159 | Overlapping |
| hCoVNL63 | nucleocapsid | NSRDSSRSTSRQQSR | 159 | 173 | Overlapping |
| hCoVNL63 | nucleocapsid | NVQERWRMRRGQRVD | 57 | 71 | Overlapping |
| hCoVNL63 | nucleocapsid | PELSVVEFEDRSNNS | 133 | 147 | Overlapping |
| hCoVNL63 | nucleocapsid | PIGKGNKDEQIGYWN | 43 | 57 | Overlapping |
| hCoVNL63 | nucleocapsid | PLADDDSAIIEIVNE | 360 | 374 | Overlapping |
| hCoVNL63 | nucleocapsid | PLEPKFSIALPPELS | 122 | 136 | Overlapping |
| hCoVNL63 | nucleocapsid | PTREENVIQCFGPRD | 240 | 254 | Overlapping |
| hCoVNL63 | nucleocapsid | QERWRMRRGQRVDLP | 59 | 73 | Overlapping |
| hCoVNL63 | nucleocapsid | QIGYWNVQERWRMRR | 52 | 66 | Overlapping |
| hCoVNL63 | nucleocapsid | RDFNHNMGDSDLVQN | 253 | 267 | Overlapping |
| hCoVNL63 | nucleocapsid | RDSSRSTSRQQSRTR | 161 | 175 | Overlapping |
| hCoVNL63 | nucleocapsid | REENVIQCFGPRDFN | 242 | 256 | Overlapping |
| hCoVNL63 | nucleocapsid | RVDLPPKVHFYYLGT | 69 | 83 | Overlapping |
| hCoVNL63 | nucleocapsid | SDLVQNGVDAKGFPQ | 262 | 276 | Overlapping |
| hCoVNL63 | nucleocapsid | SDSNQSSSDLVAAVT | 176 | 190 | Overlapping |
| hCoVNL63 | nucleocapsid | SEVSTDEVGDNVQIT | 291 | 305 | Overlapping |
| hCoVNL63 | nucleocapsid | SNQSSSDLVAAVTLA | 178 | 192 | Overlapping |
| hCoVNL63 | nucleocapsid | SRSSTRNNSRDSSRS | 152 | 166 | Overlapping |
| hCoVNL63 | nucleocapsid | SRSTSRQQSRTRSDS | 164 | 178 | Overlapping |

| | | | | | |
|---|---|---|---|---|---|
| hCoVNL63 | nucleocapsid | SSRASSRSSTRNNSR | 147 | 161 | Overlapping |
| hCoVNL63 | nucleocapsid | SSRSSTRNNSRDSSR | 151 | 165 | Overlapping |
| hCoVNL63 | nucleocapsid | TDEVGDNVQITYTYK | 295 | 309 | Overlapping |
| hCoVNL63 | nucleocapsid | TREENVIQCFGPRDF | 241 | 255 | Overlapping |
| hCoVNL63 | nucleocapsid | TRNNSRDSSRSTSRQ | 156 | 170 | Overlapping |
| hCoVNL63 | nucleocapsid | VPTREENVIQCFGPR | 239 | 253 | Overlapping |
| hCoVNL63 | nucleocapsid | WNVQERWRMRRGQRV | 56 | 70 | Overlapping |
| hCoVNL63 | nucleocapsid | YWNVQERWRMRRGQR | 55 | 69 | Overlapping |
| hCoVNL63 | orf1ab | AAETIKAKEESVKSS | 5104 | 5118 | Positive |
| hCoVNL63 | orf1ab | AANSSEWAYCVDLHN | 3544 | 3558 | Positive |
| hCoVNL63 | orf1ab | ACEVVVTNLNKSAGW | 4528 | 4542 | Positive |
| hCoVNL63 | orf1ab | ACNVNRFNVAITRAK | 5526 | 5540 | Positive |
| hCoVNL63 | orf1ab | AGSLELTTMRYFVKI | 5754 | 5768 | Positive |
| hCoVNL63 | orf1ab | AIDILTEGQLQSLSK | 1302 | 1316 | Positive |
| hCoVNL63 | orf1ab | AKAMLDYAKDHNDML | 2912 | 2926 | Positive |
| hCoVNL63 | orf1ab | AKEESVKSSYAFATL | 5110 | 5124 | Positive |
| hCoVNL63 | orf1ab | AKRKMGLTPPLSILK | 6144 | 6158 | Positive |
| hCoVNL63 | orf1ab | ALKLYKPSVIHDIGN | 5886 | 5900 | Positive |
| hCoVNL63 | orf1ab | AMFTKVVVGPTTEVK | 518 | 532 | Positive |
| hCoVNL63 | orf1ab | ANEKFINGCGRNVQG | 5866 | 5880 | Positive |
| hCoVNL63 | orf1ab | ANMSMAECKATLGLT | 2328 | 2342 | Positive |
| hCoVNL63 | orf1ab | APHGPFDALWLSFKL | 3492 | 3506 | Positive |
| hCoVNL63 | orf1ab | ASEADYRCACYAHLA | 2898 | 2912 | Positive |
| hCoVNL63 | orf1ab | ASLYYESISYEEQDA | 4550 | 4564 | Positive |
| hCoVNL63 | orf1ab | AVDSLCAKAMTVYSI | 5284 | 5298 | Positive |
| hCoVNL63 | orf1ab | AVVNDKVFVRYGDVD | 6110 | 6124 | Positive |
| hCoVNL63 | orf1ab | CAPVVCPKGKIVVIA | 548 | 562 | Positive |
| hCoVNL63 | orf1ab | CAVTDAKWYCYDKQP | 5906 | 5920 | Positive |
| hCoVNL63 | orf1ab | CCMRAGSKVVNHNVL | 2388 | 2402 | Positive |
| hCoVNL63 | orf1ab | CEPIKLVNSELLSTL | 2288 | 2302 | Positive |
| hCoVNL63 | orf1ab | CKEVLKNCNVLENFI | 2250 | 2264 | Positive |
| hCoVNL63 | orf1ab | CLENNVCNVCGCWLG | 4018 | 4032 | Positive |
| hCoVNL63 | orf1ab | CNEFLATFIVCKIVL | 2108 | 2122 | Positive |
| hCoVNL63 | orf1ab | CPEILLSIDDGHLWN | 714 | 728 | Positive |
| hCoVNL63 | orf1ab | CSDCYDDMCVIHCAN | 4336 | 4350 | Positive |
| hCoVNL63 | orf1ab | CVYFSQLLCEPIKLV | 2280 | 2294 | Positive |
| hCoVNL63 | orf1ab | DADFSITGDCATVYL | 6534 | 6548 | Positive |
| hCoVNL63 | orf1ab | DAIIIDNDINDYVSD | 6520 | 6534 | Positive |
| hCoVNL63 | orf1ab | DAIWDAFVAAIKLVP | 408 | 422 | Positive |
| hCoVNL63 | orf1ab | DCYDDMCVIHCANFN | 4338 | 4352 | Positive |
| hCoVNL63 | orf1ab | DDGYICGDGLIDLLV | 2696 | 2710 | Positive |
| hCoVNL63 | orf1ab | DFSITGDCATVYLED | 6536 | 6550 | Positive |
| hCoVNL63 | orf1ab | DGELPVAVVNDKVFV | 6104 | 6118 | Positive |
| hCoVNL63 | orf1ab | DGENVSKDGFFTYLN | 6568 | 6582 | Positive |
| hCoVNL63 | orf1ab | DGLDLTPVIDDVDVV | 1418 | 1432 | Positive |
| hCoVNL63 | orf1ab | DGLIDSYFDNSSTLQ | 3590 | 3604 | Positive |
| hCoVNL63 | orf1ab | DGRVDDGYICGDGLI | 2692 | 2706 | Positive |
| hCoVNL63 | orf1ab | DGVPLVTTAGYHFKQ | 4374 | 4388 | Positive |
| hCoVNL63 | orf1ab | DGVVPLSVIPATSAS | 3716 | 3730 | Positive |

| | | | | | |
|---|---|---|---|---|---|
| hCoVNL63 | orf1ab | DGYLSSCCGTPAKKL | 258 | 272 | Positive |
| hCoVNL63 | orf1ab | DHYDGFYTQGRNLSD | 6270 | 6284 | Positive |
| hCoVNL63 | orf1ab | DIVVVDEVSMCTNYD | 5340 | 5354 | Positive |
| hCoVNL63 | orf1ab | DLTKGPHEFCSQHTM | 4842 | 4856 | Positive |
| hCoVNL63 | orf1ab | DMEYDFLNMDMGVFI | 6290 | 6304 | Positive |
| hCoVNL63 | orf1ab | DMLYSPPTISYNSTL | 2924 | 2938 | Positive |
| hCoVNL63 | orf1ab | DSFEGVLPINTDTVL | 1616 | 1630 | Positive |
| hCoVNL63 | orf1ab | DSHKGVCFGFDKWYV | 2676 | 2690 | Positive |
| hCoVNL63 | orf1ab | DSIVTLEQYSTCDIC | 1746 | 1760 | Positive |
| hCoVNL63 | orf1ab | DSKFQIGEFIFEKVE | 5160 | 5174 | Positive |
| hCoVNL63 | orf1ab | DSLCAKAMTVYSIDK | 5286 | 5300 | Positive |
| hCoVNL63 | orf1ab | DSVREEVDIIEQPFE | 1016 | 1030 | Positive |
| hCoVNL63 | orf1ab | DSYFDNSSTLQSVAS | 3594 | 3608 | Positive |
| hCoVNL63 | orf1ab | DTRTRSVFNLEGVNG | 5962 | 5976 | Positive |
| hCoVNL63 | orf1ab | DVIVHDVEPTHKVKL | 906 | 920 | Positive |
| hCoVNL63 | orf1ab | DVRDYKLANDVKDTL | 5086 | 5100 | Positive |
| hCoVNL63 | orf1ab | EADYRCACYAHLAKA | 2900 | 2914 | Positive |
| hCoVNL63 | orf1ab | ECEKFNVFNVVGPRT | 1334 | 1348 | Positive |
| hCoVNL63 | orf1ab | EFDHEISVQKKINRM | 3652 | 3666 | Positive |
| hCoVNL63 | orf1ab | EVKPFAVYKNVKFYL | 1256 | 1270 | Positive |
| hCoVNL63 | orf1ab | EYEGGCNTIELDSPC | 3854 | 3868 | Positive |
| hCoVNL63 | orf1ab | FAVYKNVKFYLGDIS | 1260 | 1274 | Positive |
| hCoVNL63 | orf1ab | FDGKPVLPKNMWEFR | 122 | 136 | Positive |
| hCoVNL63 | orf1ab | FDIYEGGCIKACEVV | 4518 | 4532 | Positive |
| hCoVNL63 | orf1ab | FDYHPNCSDCYDDMC | 4330 | 4344 | Positive |
| hCoVNL63 | orf1ab | FEDDVVTSLCKKSFG | 924 | 938 | Positive |
| hCoVNL63 | orf1ab | FEFEDDVVTSLCKKS | 922 | 936 | Positive |
| hCoVNL63 | orf1ab | FEMSCDCGKKFDDQV | 1130 | 1144 | Positive |
| hCoVNL63 | orf1ab | FFEIKHADLHSSQVC | 5558 | 5572 | Positive |
| hCoVNL63 | orf1ab | FIFEKVEYGSDTVTY | 5168 | 5182 | Positive |
| hCoVNL63 | orf1ab | FITMSSKGQKGDAEE | 1720 | 1734 | Positive |
| hCoVNL63 | orf1ab | FKYMVANGLHAPHGP | 3482 | 3496 | Positive |
| hCoVNL63 | orf1ab | FLELLPNVFKLKIST | 2828 | 2842 | Positive |
| hCoVNL63 | orf1ab | FLGASVTEDVKFAAS | 360 | 374 | Positive |
| hCoVNL63 | orf1ab | FMGFRFDISIPGSHS | 5624 | 5638 | Positive |
| hCoVNL63 | orf1ab | FNVFNVVGPRTGKHE | 1338 | 1352 | Positive |
| hCoVNL63 | orf1ab | FSPESVFSVFGDVKL | 3426 | 3440 | Positive |
| hCoVNL63 | orf1ab | FTKVVVGPTTEVKFS | 520 | 534 | Positive |
| hCoVNL63 | orf1ab | FTMFWAELFIYTNTI | 3260 | 3274 | Positive |
| hCoVNL63 | orf1ab | FVEKFNFVTDWLKTL | 730 | 744 | Positive |
| hCoVNL63 | orf1ab | FVKWEYEGGCNTIEL | 3850 | 3864 | Positive |
| hCoVNL63 | orf1ab | FVQDPAPIDIDAFPV | 1182 | 1196 | Positive |
| hCoVNL63 | orf1ab | GAGSVVFVDKYMCGF | 108 | 122 | Positive |
| hCoVNL63 | orf1ab | GAGVMLECEKFNVFN | 1328 | 1342 | Positive |
| hCoVNL63 | orf1ab | GASVTEDVKFAASTG | 362 | 376 | Positive |
| hCoVNL63 | orf1ab | GCACDRTTIQSVDIS | 4034 | 4048 | Positive |
| hCoVNL63 | orf1ab | GDAVMTRCLAVHDCF | 5838 | 5852 | Positive |
| hCoVNL63 | orf1ab | GFCPHRHKLRSRVKF | 1784 | 1798 | Positive |
| hCoVNL63 | orf1ab | GFGVFDIKNSSVNTV | 1226 | 1240 | Positive |

| | | | | | |
|---|---|---|---|---|---|
| hCoVNL63 | orf1ab | GGKISFSDDVIVHDV | 898 | 912 | Positive |
| hCoVNL63 | orf1ab | GHGAGSVVFVDKYMC | 106 | 120 | Positive |
| hCoVNL63 | orf1ab | GHGCACDRTTIQSVD | 4032 | 4046 | Positive |
| hCoVNL63 | orf1ab | GHTLKSGCKLTNAKP | 184 | 198 | Positive |
| hCoVNL63 | orf1ab | GHVVRAALKLYKPSV | 5880 | 5894 | Positive |
| hCoVNL63 | orf1ab | GKASLYYESISYEEQ | 4548 | 4562 | Positive |
| hCoVNL63 | orf1ab | GKPVLPKNMWEFRDY | 124 | 138 | Positive |
| hCoVNL63 | orf1ab | GKQKITTIQGPPGSG | 5244 | 5258 | Positive |
| hCoVNL63 | orf1ab | GMLNGMPVSSIKGDK | 6236 | 6250 | Positive |
| hCoVNL63 | orf1ab | GNFDDQPSLQVESAN | 3122 | 3136 | Positive |
| hCoVNL63 | orf1ab | GNGQAITTSVDANTN | 3958 | 3972 | Positive |
| hCoVNL63 | orf1ab | GPFIAGNTVHANYIF | 6646 | 6660 | Positive |
| hCoVNL63 | orf1ab | GPFVSFIYFITMSSK | 1712 | 1726 | Positive |
| hCoVNL63 | orf1ab | GPGNTFINGDIAREL | 2186 | 2200 | Positive |
| hCoVNL63 | orf1ab | GPHEFCSQHTMQIVD | 4846 | 4860 | Positive |
| hCoVNL63 | orf1ab | GPTTEVKFSVIELAT | 526 | 540 | Positive |
| hCoVNL63 | orf1ab | GSASEADYRCACYAH | 2896 | 2910 | Positive |
| hCoVNL63 | orf1ab | GSELTLKHFFFAQNG | 4470 | 4484 | Positive |
| hCoVNL63 | orf1ab | GSGMFYNQVTLAVAS | 1 | 12 | Positive |
| hCoVNL63 | orf1ab | GSGSGMFYNQVTLAV | 1 | 10 | Positive |
| hCoVNL63 | orf1ab | GSGSGSGMFYNQVTL | 1 | 8 | Positive |
| hCoVNL63 | orf1ab | GSPGYNVRNDGTVEF | 3084 | 3098 | Positive |
| hCoVNL63 | orf1ab | GWYDPVENEDIHRVY | 4198 | 4212 | Positive |
| hCoVNL63 | orf1ab | GYAMPQIYKLQRMCL | 6436 | 6450 | Positive |
| hCoVNL63 | orf1ab | HAYSTMRLHNFSVSH | 2994 | 3008 | Positive |
| hCoVNL63 | orf1ab | HDVEPTHKVKLIFEF | 910 | 924 | Positive |
| hCoVNL63 | orf1ab | HKENLFNKYFKHWSF | 4316 | 4330 | Positive |
| hCoVNL63 | orf1ab | HKVKLIFEFEDDVVT | 916 | 930 | Positive |
| hCoVNL63 | orf1ab | HLAKAMLDYAKDHND | 2910 | 2924 | Positive |
| hCoVNL63 | orf1ab | HLELKETVFVEPKDG | 848 | 862 | Positive |
| hCoVNL63 | orf1ab | HNDMLYSPPTISYNS | 2922 | 2936 | Positive |
| hCoVNL63 | orf1ab | HQKHLKSIVNTRNAT | 4610 | 4624 | Positive |
| hCoVNL63 | orf1ab | HQLECIVDDSVREEV | 1008 | 1022 | Positive |
| hCoVNL63 | orf1ab | HSDFGLDGLIDSYFD | 3584 | 3598 | Positive |
| hCoVNL63 | orf1ab | HSFDVYNLWQIFIET | 6068 | 6082 | Positive |
| hCoVNL63 | orf1ab | HVIAPSTTVLIDYDH | 2980 | 2994 | Positive |
| hCoVNL63 | orf1ab | IACCMRAGSKVVNHN | 2386 | 2400 | Positive |
| hCoVNL63 | orf1ab | IAKSEFDHEISVQKK | 3648 | 3662 | Positive |
| hCoVNL63 | orf1ab | IDGFGVFDIKNSSVN | 1224 | 1238 | Positive |
| hCoVNL63 | orf1ab | IDYDHAYSTMRLHNF | 2990 | 3004 | Positive |
| hCoVNL63 | orf1ab | IDYTTTVTSFHGYDF | 2488 | 2502 | Positive |
| hCoVNL63 | orf1ab | IETNLQSLENIAFNV | 6080 | 6094 | Positive |
| hCoVNL63 | orf1ab | IKNSSVNTVCFVDVD | 1232 | 1246 | Positive |
| hCoVNL63 | orf1ab | IKPVCAKSPPGEQFR | 5700 | 5714 | Positive |
| hCoVNL63 | orf1ab | INDDDYVIALTGTNQ | 56 | 70 | Positive |
| hCoVNL63 | orf1ab | INGMHKSFYVNANGG | 2154 | 2168 | Positive |
| hCoVNL63 | orf1ab | INKPLQVFVYSSNEE | 1396 | 1410 | Positive |
| hCoVNL63 | orf1ab | INRLLSVPSDSCNNV | 4742 | 4756 | Positive |
| hCoVNL63 | orf1ab | IPATSASKLTIVSPD | 3724 | 3738 | Positive |

| | | | | | |
|---|---|---|---|---|---|
| hCoVNL63 | orf1ab | IQQWGYVGSLSQNHH | 5810 | 5824 | Positive |
| hCoVNL63 | orf1ab | IQSVDISYLNRARGS | 4042 | 4056 | Positive |
| hCoVNL63 | orf1ab | ISAMHSLLFGMLRRL | 3688 | 3702 | Positive |
| hCoVNL63 | orf1ab | ITEDGINVKDVVVES | 1586 | 1600 | Positive |
| hCoVNL63 | orf1ab | ITTIQGPPGSGKSHC | 5248 | 5262 | Positive |
| hCoVNL63 | orf1ab | IVHDVEPTHKVKLIF | 908 | 922 | Positive |
| hCoVNL63 | orf1ab | IVSPDLESYSKIVCD | 3734 | 3748 | Positive |
| hCoVNL63 | orf1ab | IWVPHSFDVYNLWQI | 6064 | 6078 | Positive |
| hCoVNL63 | orf1ab | KCWVEEDLTKGPHEF | 4836 | 4850 | Positive |
| hCoVNL63 | orf1ab | KDGCDVGFCPHRHKL | 1778 | 1792 | Positive |
| hCoVNL63 | orf1ab | KDNDGRPVHVKEITK | 3764 | 3778 | Positive |
| hCoVNL63 | orf1ab | KELTANMSMAECKAT | 2324 | 2338 | Positive |
| hCoVNL63 | orf1ab | KESIPIVWGVKDFNT | 2404 | 2418 | Positive |
| hCoVNL63 | orf1ab | KETVFVEPKDGGQFF | 852 | 866 | Positive |
| hCoVNL63 | orf1ab | KEVLVLTGCCDNSYF | 4180 | 4194 | Positive |
| hCoVNL63 | orf1ab | KFGKASLYYESISYE | 4546 | 4560 | Positive |
| hCoVNL63 | orf1ab | KGQKGDAEEALSKLS | 1726 | 1740 | Positive |
| hCoVNL63 | orf1ab | KHEHSLLVEAYNSIL | 1350 | 1364 | Positive |
| hCoVNL63 | orf1ab | KHFFFAQNGDAAVKD | 4476 | 4490 | Positive |
| hCoVNL63 | orf1ab | KHLKSIVNTRNATVV | 4612 | 4626 | Positive |
| hCoVNL63 | orf1ab | KHVFKTLKPGDSFNI | 3038 | 3052 | Positive |
| hCoVNL63 | orf1ab | KINRMAEQAATQMYK | 3662 | 3676 | Positive |
| hCoVNL63 | orf1ab | KISFSDDVIVHDVEP | 900 | 914 | Positive |
| hCoVNL63 | orf1ab | KKYLVKTTKAKGLTF | 2424 | 2438 | Positive |
| hCoVNL63 | orf1ab | KLHPAFNVSDAYANL | 5222 | 5236 | Positive |
| hCoVNL63 | orf1ab | KLLDYDYATHGQLDG | 5926 | 5940 | Positive |
| hCoVNL63 | orf1ab | KLPQFYIYDEEGGYD | 966 | 980 | Positive |
| hCoVNL63 | orf1ab | KLSEYLISDSIVTLE | 1738 | 1752 | Positive |
| hCoVNL63 | orf1ab | KLTNAKPPKYSSKVV | 192 | 206 | Positive |
| hCoVNL63 | orf1ab | KLYKPSVIHDIGNPK | 5888 | 5902 | Positive |
| hCoVNL63 | orf1ab | KNMWEFRDYFNNNTD | 130 | 144 | Positive |
| hCoVNL63 | orf1ab | KPASKQCFKVFFKGN | 5436 | 5450 | Positive |
| hCoVNL63 | orf1ab | KPEDKLSVYDIACCM | 2376 | 2390 | Positive |
| hCoVNL63 | orf1ab | KPMPFFYFDDSDCDV | 6000 | 6014 | Positive |
| hCoVNL63 | orf1ab | KPNAIDVEHLELKET | 840 | 854 | Positive |
| hCoVNL63 | orf1ab | KQLEIVKLFLVKNPS | 5462 | 5476 | Positive |
| hCoVNL63 | orf1ab | KQPVNSNVKLLDYDY | 5918 | 5932 | Positive |
| hCoVNL63 | orf1ab | KRWLPPDAIIIDNDI | 6514 | 6528 | Positive |
| hCoVNL63 | orf1ab | KSVCKYTDFNEDVCV | 6184 | 6198 | Positive |
| hCoVNL63 | orf1ab | KTCVIANVPAIDVLK | 1538 | 1552 | Positive |
| hCoVNL63 | orf1ab | KVSQSNVHTPKHVFK | 3028 | 3042 | Positive |
| hCoVNL63 | orf1ab | KVVVGPTTEVKFSVI | 522 | 536 | Positive |
| hCoVNL63 | orf1ab | KWEYEGGCNTIELDS | 3852 | 3866 | Positive |
| hCoVNL63 | orf1ab | KYSSIYKLHPAFNVS | 5216 | 5230 | Positive |
| hCoVNL63 | orf1ab | LAFLGASVTEDVKFA | 358 | 372 | Positive |
| hCoVNL63 | orf1ab | LASIQHLHEGFGGKN | 3202 | 3216 | Positive |
| hCoVNL63 | orf1ab | LCQYLNSTTMCVPHN | 6478 | 6492 | Positive |
| hCoVNL63 | orf1ab | LDNQEDKFWCEDFYA | 4946 | 4960 | Positive |
| hCoVNL63 | orf1ab | LECIVDDSVREEVDI | 1010 | 1024 | Positive |

| | | | | | |
|---|---|---|---|---|---|
| hCoVNL63 | orf1ab | LFAAETIKAKEESVK | 5102 | 5116 | Positive |
| hCoVNL63 | orf1ab | LFENGIPLMPLLSCG | 1364 | 1378 | Positive |
| hCoVNL63 | orf1ab | LFYTIKFSGFKLDGF | 592 | 606 | Positive |
| hCoVNL63 | orf1ab | LGDGNALYNTEGGKT | 3822 | 3836 | Positive |
| hCoVNL63 | orf1ab | LHAPHGPFDALWLSF | 3490 | 3504 | Positive |
| hCoVNL63 | orf1ab | LHEVLTSAMNVIGQH | 950 | 964 | Positive |
| hCoVNL63 | orf1ab | LHGGGVARAIDILTE | 1294 | 1308 | Positive |
| hCoVNL63 | orf1ab | LKEKYTCSITFEMSC | 1120 | 1134 | Positive |
| hCoVNL63 | orf1ab | LKEVVGPKELLLSWE | 5124 | 5138 | Positive |
| hCoVNL63 | orf1ab | LLERFLPKCPEILLS | 706 | 720 | Positive |
| hCoVNL63 | orf1ab | LLLRNNGRFGGFSNH | 6710 | 6724 | Positive |
| hCoVNL63 | orf1ab | LLRTSIMKHDIKVIA | 1922 | 1936 | Positive |
| hCoVNL63 | orf1ab | LLYAILYFVFTRTVR | 2784 | 2798 | Positive |
| hCoVNL63 | orf1ab | LNELSSKVVCTYMDL | 6366 | 6380 | Positive |
| hCoVNL63 | orf1ab | LNGCRWWLCSTRVNV | 3152 | 3166 | Positive |
| hCoVNL63 | orf1ab | LPECNADIVVVDEVS | 5334 | 5348 | Positive |
| hCoVNL63 | orf1ab | LQDFDVVFGHGAGSV | 98 | 112 | Positive |
| hCoVNL63 | orf1ab | LQVFVYSSNEEQAVL | 1400 | 1414 | Positive |
| hCoVNL63 | orf1ab | LSKDYISSNGPLKVG | 1314 | 1328 | Positive |
| hCoVNL63 | orf1ab | LSVDFNGVLHKAYVD | 2302 | 2316 | Positive |
| hCoVNL63 | orf1ab | LSVPSDSCNNVNVRD | 4746 | 4760 | Positive |
| hCoVNL63 | orf1ab | LTPIKLNFGMLNGMP | 6228 | 6242 | Positive |
| hCoVNL63 | orf1ab | LTPVIDDVDVVKPFR | 1422 | 1436 | Positive |
| hCoVNL63 | orf1ab | LTSNGLLGNCAKRFR | 748 | 762 | Positive |
| hCoVNL63 | orf1ab | LTVKFVVESNVMDVN | 1560 | 1574 | Positive |
| hCoVNL63 | orf1ab | LVEAYNSILFENGIP | 1356 | 1370 | Positive |
| hCoVNL63 | orf1ab | LVTTAGYHFKQLGLV | 4378 | 4392 | Positive |
| hCoVNL63 | orf1ab | LWRVTAVHSDGMFVA | 314 | 328 | Positive |
| hCoVNL63 | orf1ab | LYYESISYEEQDALF | 4552 | 4566 | Positive |
| hCoVNL63 | orf1ab | MAEQAATQMYKEARS | 3666 | 3680 | Positive |
| hCoVNL63 | orf1ab | MYENSTILQAAGLCV | 4962 | 4976 | Positive |
| hCoVNL63 | orf1ab | NACVYFSQLLCEPIK | 2278 | 2292 | Positive |
| hCoVNL63 | orf1ab | NADIVVVDEVSMCTN | 5338 | 5352 | Positive |
| hCoVNL63 | orf1ab | NAFVKCNCGSESWSV | 240 | 254 | Positive |
| hCoVNL63 | orf1ab | NAHRYDVLLSDLSFN | 2354 | 2368 | Positive |
| hCoVNL63 | orf1ab | NAIDVEHLELKETVF | 842 | 856 | Positive |
| hCoVNL63 | orf1ab | NALPECNADIVVVDE | 5332 | 5346 | Positive |
| hCoVNL63 | orf1ab | NALYNTEGGKTFMYA | 3826 | 3840 | Positive |
| hCoVNL63 | orf1ab | NANGGTCFCNKHNFF | 2164 | 2178 | Positive |
| hCoVNL63 | orf1ab | NATVVIGTTKFYGGW | 4622 | 4636 | Positive |
| hCoVNL63 | orf1ab | NAVLFSTVVIKNLTP | 6216 | 6230 | Positive |
| hCoVNL63 | orf1ab | NAVYRAFGSPFITNG | 212 | 226 | Positive |
| hCoVNL63 | orf1ab | NCGEPWFLTDYNAIL | 674 | 688 | Positive |
| hCoVNL63 | orf1ab | NCGSESWSVGAWDGY | 246 | 260 | Positive |
| hCoVNL63 | orf1ab | NCLASECFVKSDIFG | 4286 | 4300 | Positive |
| hCoVNL63 | orf1ab | NCNVDMYPEFSIVCR | 5946 | 5960 | Positive |
| hCoVNL63 | orf1ab | NCNVLENFIVYNNSG | 2256 | 2270 | Positive |
| hCoVNL63 | orf1ab | NCYRLTSVEESFIDD | 4768 | 4782 | Positive |
| hCoVNL63 | orf1ab | NDGRPVHVKEITKEN | 3766 | 3780 | Positive |

| | | | | | |
|---|---|---|---|---|---|
| hCoVNL63 | orf1ab | NDGTVEFCYLHQIEL | 3092 | 3106 | Positive |
| hCoVNL63 | orf1ab | NDINDYVSDADFSIT | 6526 | 6540 | Positive |
| hCoVNL63 | orf1ab | NDLDHTSLVWKHIRD | 2034 | 2048 | Positive |
| hCoVNL63 | orf1ab | NDPVFTGDLFYTIKF | 584 | 598 | Positive |
| hCoVNL63 | orf1ab | NDSNGCVVEASTDFH | 994 | 1008 | Positive |
| hCoVNL63 | orf1ab | NDVKDTLRLFAAETI | 5094 | 5108 | Positive |
| hCoVNL63 | orf1ab | NEEFYNFLRLRGFFD | 4454 | 4468 | Positive |
| hCoVNL63 | orf1ab | NEEQAVLKFLDGLDL | 1408 | 1422 | Positive |
| hCoVNL63 | orf1ab | NENLLHGGGVARAID | 1290 | 1304 | Positive |
| hCoVNL63 | orf1ab | NEWAMANGYTSVSSV | 3170 | 3184 | Positive |
| hCoVNL63 | orf1ab | NEYCCFKHALGCDYV | 5788 | 5802 | Positive |
| hCoVNL63 | orf1ab | NFLFAAFITFLCFLV | 2732 | 2746 | Positive |
| hCoVNL63 | orf1ab | NFNQWHEAKFGVVTT | 2526 | 2540 | Positive |
| hCoVNL63 | orf1ab | NFNTLFATTIPGTAF | 4350 | 4364 | Positive |
| hCoVNL63 | orf1ab | NFSGALAEHDFFTWK | 4128 | 4142 | Positive |
| hCoVNL63 | orf1ab | NFYDFGDFVVSLPNM | 4252 | 4266 | Positive |
| hCoVNL63 | orf1ab | NGHYVVYDAANNAVY | 1834 | 1848 | Positive |
| hCoVNL63 | orf1ab | NGLHAPHGPFDALWL | 3488 | 3502 | Positive |
| hCoVNL63 | orf1ab | NGNFYDFGDFVVSLP | 4250 | 4264 | Positive |
| hCoVNL63 | orf1ab | NGRVVITNVGEPIIS | 1800 | 1814 | Positive |
| hCoVNL63 | orf1ab | NGVIREKLAIGGSVA | 6582 | 6596 | Positive |
| hCoVNL63 | orf1ab | NGYTSVSSVECYSIL | 3176 | 3190 | Positive |
| hCoVNL63 | orf1ab | NHHTFCNIHRNEHDA | 5822 | 5836 | Positive |
| hCoVNL63 | orf1ab | NHKPQLSFPLCSAGN | 5044 | 5058 | Positive |
| hCoVNL63 | orf1ab | NHNVLIKESIPIVWG | 2398 | 2412 | Positive |
| hCoVNL63 | orf1ab | NIAANSSEWAYCVDL | 3542 | 3556 | Positive |
| hCoVNL63 | orf1ab | NIGGAVCSKHANLYQ | 6034 | 6048 | Positive |
| hCoVNL63 | orf1ab | NIHRNEHDASGDAVM | 5828 | 5842 | Positive |
| hCoVNL63 | orf1ab | NILGYSSLCDEFTLA | 3216 | 3230 | Positive |
| hCoVNL63 | orf1ab | NKADLKFVKWEYEGG | 3844 | 3858 | Positive |
| hCoVNL63 | orf1ab | NKDIYCGNSMVCKMC | 2012 | 2026 | Positive |
| hCoVNL63 | orf1ab | NKDVNTHSVRLTITE | 4394 | 4408 | Positive |
| hCoVNL63 | orf1ab | NKFVPVKPASKQCFK | 5430 | 5444 | Positive |
| hCoVNL63 | orf1ab | NKFVTGDVGPFVSFI | 1704 | 1718 | Positive |
| hCoVNL63 | orf1ab | NKHAFHTPAYDKRAF | 5982 | 5996 | Positive |
| hCoVNL63 | orf1ab | NKINLCDDPEKAQSM | 3558 | 3572 | Positive |
| hCoVNL63 | orf1ab | NKNLNEGVLESFSVT | 4930 | 4944 | Positive |
| hCoVNL63 | orf1ab | NKNVSFLGKCLKMNC | 4080 | 4094 | Positive |
| hCoVNL63 | orf1ab | NKTTLPTNVAFELFA | 6130 | 6144 | Positive |
| hCoVNL63 | orf1ab | NKYFKHWSFDYHPNC | 4322 | 4336 | Positive |
| hCoVNL63 | orf1ab | NLARDGVVPLSVIPA | 3712 | 3726 | Positive |
| hCoVNL63 | orf1ab | NLEGVNGGSLYVNKH | 5970 | 5984 | Positive |
| hCoVNL63 | orf1ab | NLFVEKFNFVTDWLK | 728 | 742 | Positive |
| hCoVNL63 | orf1ab | NLGRCVRKLNRLKTC | 1526 | 1540 | Positive |
| hCoVNL63 | orf1ab | NLLHGGGVARAIDIL | 1292 | 1306 | Positive |
| hCoVNL63 | orf1ab | NLMLSDNVVAFLYAA | 3136 | 3150 | Positive |
| hCoVNL63 | orf1ab | NLNTLRRGAVLGFIG | 3888 | 3902 | Positive |
| hCoVNL63 | orf1ab | NLQSGKVIFGLKTMF | 3240 | 3254 | Positive |
| hCoVNL63 | orf1ab | NLRGWLIFSNSNYVL | 84 | 98 | Positive |

| | | | | | |
|---|---|---|---|---|---|
| hCoVNL63 | orf1ab | NLRLVDCAPVVCPKG | 542 | 556 | Positive |
| hCoVNL63 | orf1ab | NLYSFNKAIDGFGVF | 1216 | 1230 | Positive |
| hCoVNL63 | orf1ab | NMIRMISAMVLGSKH | 4666 | 4680 | Positive |
| hCoVNL63 | orf1ab | NMLRTLIDGVENPML | 4638 | 4652 | Positive |
| hCoVNL63 | orf1ab | NMRVLHYGAGSDKGV | 6492 | 6506 | Positive |
| hCoVNL63 | orf1ab | NNADCVACSKSARLK | 2132 | 2146 | Positive |
| hCoVNL63 | orf1ab | NNAVYDGARLFSSDL | 1844 | 1858 | Positive |
| hCoVNL63 | orf1ab | NNCWISTTLVQLQLT | 1060 | 1074 | Positive |
| hCoVNL63 | orf1ab | NNEIMPGKLKQKPMK | 3800 | 3814 | Positive |
| hCoVNL63 | orf1ab | NNFFISYAKPEDKLS | 2368 | 2382 | Positive |
| hCoVNL63 | orf1ab | NNGRFGGFSNHLVST | 6714 | 6728 | Positive |
| hCoVNL63 | orf1ab | NNISYVVTQNLFFML | 2770 | 2784 | Positive |
| hCoVNL63 | orf1ab | NNTSAQYIFSTVNAL | 5320 | 5334 | Positive |
| hCoVNL63 | orf1ab | NNVFMSTSKCWVEED | 4828 | 4842 | Positive |
| hCoVNL63 | orf1ab | NNVNVRDLQRRLYDN | 4754 | 4768 | Positive |
| hCoVNL63 | orf1ab | NQDLNGNFYDFGDFV | 4246 | 4260 | Positive |
| hCoVNL63 | orf1ab | NQDSYGGASICLYCR | 3972 | 3986 | Positive |
| hCoVNL63 | orf1ab | NQEDKFWCEDFYASM | 4948 | 4962 | Positive |
| hCoVNL63 | orf1ab | NQLRLAFLGASVTED | 354 | 368 | Positive |
| hCoVNL63 | orf1ab | NQVTLAVASDSEISG | 4 | 18 | Positive |
| hCoVNL63 | orf1ab | NRFNVAITRAKKGIF | 5530 | 5544 | Positive |
| hCoVNL63 | orf1ab | NRKQLEIVKLFLVKN | 5460 | 5474 | Positive |
| hCoVNL63 | orf1ab | NRKSKVISAMHSLLF | 3682 | 3696 | Positive |
| hCoVNL63 | orf1ab | NRMAEQAATQMYKEA | 3664 | 3678 | Positive |
| hCoVNL63 | orf1ab | NSATCYNSVSNEYCC | 5778 | 5792 | Positive |
| hCoVNL63 | orf1ab | NSEYRKVFYVLLDWV | 4912 | 4926 | Positive |
| hCoVNL63 | orf1ab | NSGLLTACAFSVDPA | 3918 | 3932 | Positive |
| hCoVNL63 | orf1ab | NSGSNITQIKNACVY | 2268 | 2282 | Positive |
| hCoVNL63 | orf1ab | NSIQGSYERFTLTTN | 6202 | 6216 | Positive |
| hCoVNL63 | orf1ab | NSLKALFSCDINKPL | 1386 | 1400 | Positive |
| hCoVNL63 | orf1ab | NSSEWAYCVDLHNKI | 3546 | 3560 | Positive |
| hCoVNL63 | orf1ab | NSSVNTVCFVDVDFH | 1234 | 1248 | Positive |
| hCoVNL63 | orf1ab | NSVLDLSKFECKHKA | 6672 | 6686 | Positive |
| hCoVNL63 | orf1ab | NSVSNEYCCFKHALG | 5784 | 5798 | Positive |
| hCoVNL63 | orf1ab | NTEGGKTFMYAYISN | 3830 | 3844 | Positive |
| hCoVNL63 | orf1ab | NTIELDSPCRFMVET | 3860 | 3874 | Positive |
| hCoVNL63 | orf1ab | NTVCFVDVDFHSVEI | 1238 | 1252 | Positive |
| hCoVNL63 | orf1ab | NVCTYEHVISFMGFR | 5614 | 5628 | Positive |
| hCoVNL63 | orf1ab | NVDMYPEFSIVCRFD | 5948 | 5962 | Positive |
| hCoVNL63 | orf1ab | NVFDNFNQWHEAKFG | 2522 | 2536 | Positive |
| hCoVNL63 | orf1ab | NVKLLDYDYATHGQL | 5924 | 5938 | Positive |
| hCoVNL63 | orf1ab | NVLAIESIHYLGTTG | 170 | 184 | Positive |
| hCoVNL63 | orf1ab | NVLENFIVYNNSGSN | 2258 | 2272 | Positive |
| hCoVNL63 | orf1ab | NVLPTMTQLNLKYAI | 4572 | 4586 | Positive |
| hCoVNL63 | orf1ab | NVNRFNVAITRAKKG | 5528 | 5542 | Positive |
| hCoVNL63 | orf1ab | NVPTIVSEKISVMDK | 1876 | 1890 | Positive |
| hCoVNL63 | orf1ab | NVQVDNGSSINRKQL | 5450 | 5464 | Positive |
| hCoVNL63 | orf1ab | NVSDAYANLVPYYQL | 5228 | 5242 | Positive |
| hCoVNL63 | orf1ab | NVSRHNLTKYTMMDL | 4150 | 4164 | Positive |

| | | | | | |
|---|---|---|---|---|---|
| hCoVNL63 | orf1ab | NVVPGDVIITSTSAG | 278 | 292 | Positive |
| hCoVNL63 | orf1ab | NVVPGVPTNVYLVGK | 2556 | 2570 | Positive |
| hCoVNL63 | orf1ab | NVYLVGKTLVFTLQA | 2564 | 2578 | Positive |
| hCoVNL63 | orf1ab | NYDLSVINQRLSYKH | 5352 | 5366 | Positive |
| hCoVNL63 | orf1ab | NYLGDFIQGPFIAGN | 6638 | 6652 | Positive |
| hCoVNL63 | orf1ab | NYVASRFLGLQIQTV | 5490 | 5504 | Positive |
| hCoVNL63 | orf1ab | NYVPLRASSCVTRCN | 6020 | 6034 | Positive |
| hCoVNL63 | orf1ab | NYVRFPEILARGFGL | 2644 | 2658 | Positive |
| hCoVNL63 | orf1ab | PESVFSVFGDVKLTL | 3428 | 3442 | Positive |
| hCoVNL63 | orf1ab | PHPSMDGYCKFKGKC | 3990 | 4004 | Positive |
| hCoVNL63 | orf1ab | PIDIDAFPVKPICSS | 1188 | 1202 | Positive |
| hCoVNL63 | orf1ab | PIKLNFGMLNGMPVS | 6230 | 6244 | Positive |
| hCoVNL63 | orf1ab | PKELLLSWESGKVKP | 5130 | 5144 | Positive |
| hCoVNL63 | orf1ab | PNCSDCYDDMCVIHC | 4334 | 4348 | Positive |
| hCoVNL63 | orf1ab | PPDAIIIDNDINDYV | 6518 | 6532 | Positive |
| hCoVNL63 | orf1ab | PRHVIAPSTTVLIDY | 2978 | 2992 | Positive |
| hCoVNL63 | orf1ab | PSMDGYCKFKGKCVQ | 3992 | 4006 | Positive |
| hCoVNL63 | orf1ab | PSYIAYENARQAYED | 3614 | 3628 | Positive |
| hCoVNL63 | orf1ab | PTAPAYVIIDKVDFV | 2210 | 2224 | Positive |
| hCoVNL63 | orf1ab | PTNVYLVGKTLVFTL | 2562 | 2576 | Positive |
| hCoVNL63 | orf1ab | PTTTGVLVRFVKSIA | 422 | 436 | Positive |
| hCoVNL63 | orf1ab | PVDYNVVTQRMCAIG | 5390 | 5404 | Positive |
| hCoVNL63 | orf1ab | PVENEDIHRVYASLG | 4202 | 4216 | Positive |
| hCoVNL63 | orf1ab | PVHVKEITKENVETL | 3770 | 3784 | Positive |
| hCoVNL63 | orf1ab | PVIDDVDVVKPFRVE | 1424 | 1438 | Positive |
| hCoVNL63 | orf1ab | PYPDPSRILSAGVFV | 4868 | 4882 | Positive |
| hCoVNL63 | orf1ab | PYVVISGFVSRVIRR | 798 | 812 | Positive |
| hCoVNL63 | orf1ab | QFRLSKMGVLKADDF | 6336 | 6350 | Positive |
| hCoVNL63 | orf1ab | QGPPGSGKSHCSIGL | 5252 | 5266 | Positive |
| hCoVNL63 | orf1ab | QGRNLSDFTPRSDME | 6278 | 6292 | Positive |
| hCoVNL63 | orf1ab | QHLHEGFGGKNILGY | 3206 | 3220 | Positive |
| hCoVNL63 | orf1ab | QIQTVDSSQGSEYDY | 5500 | 5514 | Positive |
| hCoVNL63 | orf1ab | QLLASIQHLHEGFGG | 3200 | 3214 | Positive |
| hCoVNL63 | orf1ab | QQAALDYLATVKTVP | 1482 | 1496 | Positive |
| hCoVNL63 | orf1ab | QVPIGCLDPIRFCLE | 4006 | 4020 | Positive |
| hCoVNL63 | orf1ab | QVVKPGHFNEEFYNF | 4446 | 4460 | Positive |
| hCoVNL63 | orf1ab | QWGYVGSLSQNHHTF | 5812 | 5826 | Positive |
| hCoVNL63 | orf1ab | RAALKLYKPSVIHDI | 5884 | 5898 | Positive |
| hCoVNL63 | orf1ab | RAFDIYNKNVSFLGK | 4074 | 4088 | Positive |
| hCoVNL63 | orf1ab | RCDMTFPCVSCVTFF | 814 | 828 | Positive |
| hCoVNL63 | orf1ab | RCLAVHDCFVKNVDW | 5844 | 5858 | Positive |
| hCoVNL63 | orf1ab | RCPAEIVNTVSELVY | 5414 | 5428 | Positive |
| hCoVNL63 | orf1ab | RDATFATHVCFKDCY | 650 | 664 | Positive |
| hCoVNL63 | orf1ab | RDYFNNNTDSIVIGG | 136 | 150 | Positive |
| hCoVNL63 | orf1ab | RDYKLANDVKDTLRL | 5088 | 5102 | Positive |
| hCoVNL63 | orf1ab | RERCDMTFPCVSCVT | 812 | 826 | Positive |
| hCoVNL63 | orf1ab | RGFFDEGSELTLKHF | 4464 | 4478 | Positive |
| hCoVNL63 | orf1ab | RLGDYVLTENALVRL | 488 | 502 | Positive |
| hCoVNL63 | orf1ab | RLYDNCYRLTSVEES | 4764 | 4778 | Positive |

| | | | | | |
|---|---|---|---|---|---|
| hCoVNL63 | orf1ab | RNLSDFTPRSDMEYD | 6280 | 6294 | Positive |
| hCoVNL63 | orf1ab | RNSTIMSLSYNSVLD | 6662 | 6676 | Positive |
| hCoVNL63 | orf1ab | RSDMEYDFLNMDMGV | 6288 | 6302 | Positive |
| hCoVNL63 | orf1ab | RTRSVFNLEGVNGGS | 5964 | 5978 | Positive |
| hCoVNL63 | orf1ab | RTSIMKHDIKVIAKA | 1924 | 1938 | Positive |
| hCoVNL63 | orf1ab | SAEWKCGYAMPQIYK | 6430 | 6444 | Positive |
| hCoVNL63 | orf1ab | SDAYANLVPYYQLIG | 5230 | 5244 | Positive |
| hCoVNL63 | orf1ab | SDCDVVQEQVNYVPL | 6010 | 6024 | Positive |
| hCoVNL63 | orf1ab | SDDDFVSAVANAHRY | 2344 | 2358 | Positive |
| hCoVNL63 | orf1ab | SDDVIVHDVEPTHKV | 904 | 918 | Positive |
| hCoVNL63 | orf1ab | SDFTGSVYGNFDDQP | 3114 | 3128 | Positive |
| hCoVNL63 | orf1ab | SDGMFVASSSYDALL | 322 | 336 | Positive |
| hCoVNL63 | orf1ab | SDIFGSDFKTFDLLK | 4296 | 4310 | Positive |
| hCoVNL63 | orf1ab | SDKCPIVVGVSERIN | 2542 | 2556 | Positive |
| hCoVNL63 | orf1ab | SDKGVAPGTTVLKRW | 6502 | 6516 | Positive |
| hCoVNL63 | orf1ab | SDLSTLAVTAIVVVG | 1856 | 1870 | Positive |
| hCoVNL63 | orf1ab | SDNNCWISTTLVQLQ | 1058 | 1072 | Positive |
| hCoVNL63 | orf1ab | SDNVVAFLYAALLNG | 3140 | 3154 | Positive |
| hCoVNL63 | orf1ab | SDSEISGFGFAIPSV | 12 | 26 | Positive |
| hCoVNL63 | orf1ab | SDTTLRCCTVTYLNE | 6354 | 6368 | Positive |
| hCoVNL63 | orf1ab | SDVKKLYLGGLNYYC | 5028 | 5042 | Positive |
| hCoVNL63 | orf1ab | SDWTDVRDYKLANDV | 5082 | 5096 | Positive |
| hCoVNL63 | orf1ab | SDWYIGAIVFRLSRL | 3408 | 3422 | Positive |
| hCoVNL63 | orf1ab | SDYLSNLSDILVFVL | 5738 | 5752 | Positive |
| hCoVNL63 | orf1ab | SEAFLIGINYLGDFI | 6630 | 6644 | Positive |
| hCoVNL63 | orf1ab | SECFVKSDIFGSDFK | 4290 | 4304 | Positive |
| hCoVNL63 | orf1ab | SEISGFGFAIPSVAV | 14 | 28 | Positive |
| hCoVNL63 | orf1ab | SEKISVMDKLDTGAQ | 1882 | 1896 | Positive |
| hCoVNL63 | orf1ab | SELVYENKFVPVKPA | 5424 | 5438 | Positive |
| hCoVNL63 | orf1ab | SERINVVPGVPTNVY | 2552 | 2566 | Positive |
| hCoVNL63 | orf1ab | SEYDYVIYAQTSDTA | 5510 | 5524 | Positive |
| hCoVNL63 | orf1ab | SEYLISDSIVTLEQY | 1740 | 1754 | Positive |
| hCoVNL63 | orf1ab | SFGPGNTFINGDIAR | 2184 | 2198 | Positive |
| hCoVNL63 | orf1ab | SFNNFFISYAKPEDK | 2366 | 2380 | Positive |
| hCoVNL63 | orf1ab | SGEWNAVYRAFGSPF | 208 | 222 | Positive |
| hCoVNL63 | orf1ab | SGKLLSELLKEKYTC | 1112 | 1126 | Positive |
| hCoVNL63 | orf1ab | SHNVQPLRAPTIANQ | 5200 | 5214 | Positive |
| hCoVNL63 | orf1ab | SIDDGHLWNLFVEKF | 720 | 734 | Positive |
| hCoVNL63 | orf1ab | SIEMQLFKVGKVDSI | 1080 | 1094 | Positive |
| hCoVNL63 | orf1ab | SIQHLHEGFGGKNIL | 3204 | 3218 | Positive |
| hCoVNL63 | orf1ab | SLDVKPYGYPNDFVG | 1652 | 1666 | Positive |
| hCoVNL63 | orf1ab | SLLVEAYNSILFENG | 1354 | 1368 | Positive |
| hCoVNL63 | orf1ab | SLSQNHHTFCNIHRN | 5818 | 5832 | Positive |
| hCoVNL63 | orf1ab | SMCTNYDLSVINQRL | 5348 | 5362 | Positive |
| hCoVNL63 | orf1ab | SMFSLLRTSIMKHDI | 1918 | 1932 | Positive |
| hCoVNL63 | orf1ab | SMMILSDDGVVCYNK | 4792 | 4806 | Positive |
| hCoVNL63 | orf1ab | SMNIAANSSEWAYCV | 3540 | 3554 | Positive |
| hCoVNL63 | orf1ab | SMPSYIAYENARQAY | 3612 | 3626 | Positive |
| hCoVNL63 | orf1ab | SMYNLLNFSGALAEH | 4122 | 4136 | Positive |

| | | | | | |
|---|---|---|---|---|---|
| hCoVNL63 | orf1ab | SNGLLGNCAKRFRRV | 750 | 764 | Positive |
| hCoVNL63 | orf1ab | SNHLVSTKGSGSGSG | 6722 | 6729 | Positive |
| hCoVNL63 | orf1ab | SNINRLLSVPSDSCN | 4740 | 4754 | Positive |
| hCoVNL63 | orf1ab | SQLLCEPIKLVNSEL | 2284 | 2298 | Positive |
| hCoVNL63 | orf1ab | SQNHHTFCNIHRNEH | 5820 | 5834 | Positive |
| hCoVNL63 | orf1ab | SSKGQKGDAEEALSK | 1724 | 1738 | Positive |
| hCoVNL63 | orf1ab | SSKSLGKQLGVVSDG | 1600 | 1614 | Positive |
| hCoVNL63 | orf1ab | SSNEEQAVLKFLDGL | 1406 | 1420 | Positive |
| hCoVNL63 | orf1ab | SSYDALLHRNSLDPF | 330 | 344 | Positive |
| hCoVNL63 | orf1ab | SVETVLNLARDGVVP | 3706 | 3720 | Positive |
| hCoVNL63 | orf1ab | SVTEDVKFAASTGVI | 364 | 378 | Positive |
| hCoVNL63 | orf1ab | SVYDIACCMRAGSKV | 2382 | 2396 | Positive |
| hCoVNL63 | orf1ab | SWESGKVKPPLNRNS | 5136 | 5150 | Positive |
| hCoVNL63 | orf1ab | SYEEQDALFALTKRN | 4558 | 4572 | Positive |
| hCoVNL63 | orf1ab | SYERLINTISPEKLK | 2868 | 2882 | Positive |
| hCoVNL63 | orf1ab | SYFDSKGWYDPVENE | 4192 | 4206 | Positive |
| hCoVNL63 | orf1ab | TCFGVSKPNAIDVEH | 834 | 848 | Positive |
| hCoVNL63 | orf1ab | TDAVVLLERYVSLAI | 4888 | 4902 | Positive |
| hCoVNL63 | orf1ab | TDLIEGSKPYSTLQP | 2620 | 2634 | Positive |
| hCoVNL63 | orf1ab | TDNNCWVNATCIILQ | 1674 | 1688 | Positive |
| hCoVNL63 | orf1ab | TDPSLIIASSPALVD | 4414 | 4428 | Positive |
| hCoVNL63 | orf1ab | TDTVLSVAPEVDWVA | 1626 | 1640 | Positive |
| hCoVNL63 | orf1ab | TDWLKTLKLTLTSNG | 738 | 752 | Positive |
| hCoVNL63 | orf1ab | TEDVKFAASTGVIDI | 366 | 380 | Positive |
| hCoVNL63 | orf1ab | TEHKENLFNKYFKHW | 4314 | 4328 | Positive |
| hCoVNL63 | orf1ab | TEYSWNKYLYELIQR | 6600 | 6614 | Positive |
| hCoVNL63 | orf1ab | TGDCATVYLEDKFDL | 6540 | 6554 | Positive |
| hCoVNL63 | orf1ab | TGKHEHSLLVEAYNS | 1348 | 1362 | Positive |
| hCoVNL63 | orf1ab | THGQLDGLCLFWNCN | 5934 | 5948 | Positive |
| hCoVNL63 | orf1ab | TKGPHEFCSQHTMQI | 4844 | 4858 | Positive |
| hCoVNL63 | orf1ab | TKLLDDSIEMQLFKV | 1074 | 1088 | Positive |
| hCoVNL63 | orf1ab | TKLVPGMIFVLTSHN | 5188 | 5202 | Positive |
| hCoVNL63 | orf1ab | TLEQYSTCDICKSTV | 1750 | 1764 | Positive |
| hCoVNL63 | orf1ab | TLIDGVENPMLMGWD | 4642 | 4656 | Positive |
| hCoVNL63 | orf1ab | TLLDNQEDKFWCEDF | 4944 | 4958 | Positive |
| hCoVNL63 | orf1ab | TMCVPHNMRVLHYGA | 6486 | 6500 | Positive |
| hCoVNL63 | orf1ab | TMFLFSVFFTMFWAE | 3252 | 3266 | Positive |
| hCoVNL63 | orf1ab | TMFLKHKFLFLQVFL | 3296 | 3310 | Positive |
| hCoVNL63 | orf1ab | TMRLHNFSVSHNGVF | 2998 | 3012 | Positive |
| hCoVNL63 | orf1ab | TMSSKGQKGDAEEAL | 1722 | 1736 | Positive |
| hCoVNL63 | orf1ab | TMTQLNLKYAISGKE | 4576 | 4590 | Positive |
| hCoVNL63 | orf1ab | TNAKPPKYSSKVVLS | 194 | 208 | Positive |
| hCoVNL63 | orf1ab | TNSDKCPIVVGVSER | 2540 | 2554 | Positive |
| hCoVNL63 | orf1ab | TPRSDMEYDFLNMDM | 6286 | 6300 | Positive |
| hCoVNL63 | orf1ab | TQLFEGDKFIGTFES | 2842 | 2856 | Positive |
| hCoVNL63 | orf1ab | TTAGYHFKQLGLVWN | 4380 | 4394 | Positive |
| hCoVNL63 | orf1ab | TTGDLAVQIGSNNVC | 5602 | 5616 | Positive |
| hCoVNL63 | orf1ab | TTSGDASTAYANSIF | 4718 | 4732 | Positive |
| hCoVNL63 | orf1ab | TVSDDDFVSAVANAH | 2342 | 2356 | Positive |

| | | | | | |
|---|---|---|---|---|---|
| hCoVNL63 | orf1ab | TWKDGRVIYGNVSRH | 4140 | 4154 | Positive |
| hCoVNL63 | orf1ab | TYMDLLLDDFVTILK | 6376 | 6390 | Positive |
| hCoVNL63 | orf1ab | VAITRAKKGIFCVMC | 5534 | 5548 | Positive |
| hCoVNL63 | orf1ab | VANGLHAPHGPFDAL | 3486 | 3500 | Positive |
| hCoVNL63 | orf1ab | VASDSEISGFGFAIP | 10 | 24 | Positive |
| hCoVNL63 | orf1ab | VASSFVSMPSYIAYE | 3606 | 3620 | Positive |
| hCoVNL63 | orf1ab | VCDGSVHYAGVVWTL | 3746 | 3760 | Positive |
| hCoVNL63 | orf1ab | VCFVDVDFHSVEIEA | 1240 | 1254 | Positive |
| hCoVNL63 | orf1ab | VCKMCLFSYQEFNDL | 2022 | 2036 | Positive |
| hCoVNL63 | orf1ab | VDDVVKTDAVVLLER | 4882 | 4896 | Positive |
| hCoVNL63 | orf1ab | VDEVSMCTNYDLSVI | 5344 | 5358 | Positive |
| hCoVNL63 | orf1ab | VDGFNEWAMANGYTS | 3166 | 3180 | Positive |
| hCoVNL63 | orf1ab | VDIIEQPFEEVEHVL | 1022 | 1036 | Positive |
| hCoVNL63 | orf1ab | VDLHNKINLCDDPEK | 3554 | 3568 | Positive |
| hCoVNL63 | orf1ab | VDSSQGSEYDYVIYA | 5504 | 5518 | Positive |
| hCoVNL63 | orf1ab | VESNVMDVNDCFKND | 1566 | 1580 | Positive |
| hCoVNL63 | orf1ab | VFEAPLHCVRNVFDN | 2512 | 2526 | Positive |
| hCoVNL63 | orf1ab | VFVDDVVKTDAVVLL | 4880 | 4894 | Positive |
| hCoVNL63 | orf1ab | VFVYSSNEEQAVLKF | 1402 | 1416 | Positive |
| hCoVNL63 | orf1ab | VHDCFVKNVDWTVTY | 5848 | 5862 | Positive |
| hCoVNL63 | orf1ab | VHEVIIDNKPYRWML | 6400 | 6414 | Positive |
| hCoVNL63 | orf1ab | VHSDGMFVASSSYDA | 320 | 334 | Positive |
| hCoVNL63 | orf1ab | VKPGHFNEEFYNFLR | 4448 | 4462 | Positive |
| hCoVNL63 | orf1ab | VLKDGCDVGFCPHRH | 1776 | 1790 | Positive |
| hCoVNL63 | orf1ab | VMDVNDCFKNDNVVL | 1570 | 1584 | Positive |
| hCoVNL63 | orf1ab | VNAANENLLHGGGVA | 1286 | 1300 | Positive |
| hCoVNL63 | orf1ab | VNTRNATVVIGTTKF | 4618 | 4632 | Positive |
| hCoVNL63 | orf1ab | VNTVSELVYENKFVP | 5420 | 5434 | Positive |
| hCoVNL63 | orf1ab | VPGDVIITSTSAGCG | 280 | 294 | Positive |
| hCoVNL63 | orf1ab | VRNVFDNFNQWHEAK | 2520 | 2534 | Positive |
| hCoVNL63 | orf1ab | VRRDATFATHVCFKD | 648 | 662 | Positive |
| hCoVNL63 | orf1ab | VSDADFSITGDCATV | 6532 | 6546 | Positive |
| hCoVNL63 | orf1ab | VSLWRVTAVHSDGMF | 312 | 326 | Positive |
| hCoVNL63 | orf1ab | VSVEQLLASIQHLHE | 3196 | 3210 | Positive |
| hCoVNL63 | orf1ab | VTDAKWYCYDKQPVN | 5908 | 5922 | Positive |
| hCoVNL63 | orf1ab | VVAMSGHMLFNFLFA | 2722 | 2736 | Positive |
| hCoVNL63 | orf1ab | VVDGCSVIVRRDATF | 640 | 654 | Positive |
| hCoVNL63 | orf1ab | VVEASTDFHQLECIV | 1000 | 1014 | Positive |
| hCoVNL63 | orf1ab | VVEVKSAIVCASVLK | 1764 | 1778 | Positive |
| hCoVNL63 | orf1ab | VVGVSERINVVPGVP | 2548 | 2562 | Positive |
| hCoVNL63 | orf1ab | VVLKITEDGINVKDV | 1582 | 1596 | Positive |
| hCoVNL63 | orf1ab | VVSDGVDSFEGVLPI | 1610 | 1624 | Positive |
| hCoVNL63 | orf1ab | VVVDEVSMCTNYDLS | 5342 | 5356 | Positive |
| hCoVNL63 | orf1ab | VVVESSKSLGKQLGV | 1596 | 1610 | Positive |
| hCoVNL63 | orf1ab | VVVTLKDSDVNDMVL | 6688 | 6702 | Positive |
| hCoVNL63 | orf1ab | VWEQFCIDNCGEPWF | 666 | 680 | Positive |
| hCoVNL63 | orf1ab | VYDFKVSAAEFKYMV | 3472 | 3486 | Positive |
| hCoVNL63 | orf1ab | VYLEDKFDLLISDMY | 6546 | 6560 | Positive |
| hCoVNL63 | orf1ab | WCEDFYASMYENSTI | 4954 | 4968 | Positive |

| | | | | | |
|---|---|---|---|---|---|
| hCoVNL63 | orf1ab | WLGHGCACDRTTIQS | 4030 | 4044 | Positive |
| hCoVNL63 | orf1ab | WLPPDAIIIDNDIND | 6516 | 6530 | Positive |
| hCoVNL63 | orf1ab | WNNMLRTLIDGVENP | 4636 | 4650 | Positive |
| hCoVNL63 | orf1ab | WSFDYHPNCSDCYDD | 4328 | 4342 | Positive |
| hCoVNL63 | orf1ab | YDDMCVIHCANFNTL | 4340 | 4354 | Positive |
| hCoVNL63 | orf1ab | YDNCYRLTSVEESFI | 4766 | 4780 | Positive |
| hCoVNL63 | orf1ab | YDPVENEDIHRVYAS | 4200 | 4214 | Positive |
| hCoVNL63 | orf1ab | YDYATHGQLDGLCLF | 5930 | 5944 | Positive |
| hCoVNL63 | orf1ab | YESISYEEQDALFAL | 4554 | 4568 | Positive |
| hCoVNL63 | orf1ab | YFSQLLCEPIKLVNS | 2282 | 2296 | Positive |
| hCoVNL63 | orf1ab | YHPNCSDCYDDMCVI | 4332 | 4346 | Positive |
| hCoVNL63 | orf1ab | YMVANGLHAPHGPFD | 3484 | 3498 | Positive |
| hCoVNL63 | orf1ab | YNKYKYYSGSASEAD | 2888 | 2902 | Positive |
| hCoVNL63 | orf1ab | YNLLNFSGALAEHDF | 4124 | 4138 | Positive |
| hCoVNL63 | orf1ab | YNSILFENGIPLMPL | 1360 | 1374 | Positive |
| hCoVNL63 | orf1ab | YNTDLIEGSKPYSTL | 2618 | 2632 | Positive |
| hCoVNL63 | orf1ab | YQEFNDLDHTSLVWK | 2030 | 2044 | Positive |
| hCoVNL63 | orf1ab | YQNNVFMSTSKCWVE | 4826 | 4840 | Positive |
| hCoVNL63 | orf1ab | YRLTSVEESFIDDYY | 4770 | 4784 | Positive |
| hCoVNL63 | orf1ab | YRLYSGDTFWRYDFD | 2228 | 2242 | Positive |
| hCoVNL63 | orf1ab | YSIDKCTRIIPARAR | 5296 | 5310 | Positive |
| hCoVNL63 | orf1ab | YSTMRLHNFSVSHNG | 2996 | 3010 | Positive |
| hCoVNL63 | orf1ab | YTIKFSGFKLDGFNH | 594 | 608 | Positive |
| hCoVNL63 | orf1ab | YVEAYNTFTQAGFNI | 6050 | 6064 | Positive |
| hCoVNL63 | orf1ab | YVNANGGTCFCNKHN | 2162 | 2176 | Positive |
| hCoVNL63 | orf1ab | AAFGNTGVCYDFDGV | 2578 | 2592 | Negative |
| hCoVNL63 | orf1ab | ACYEGIASGVFGVNL | 3054 | 3068 | Negative |
| hCoVNL63 | orf1ab | AFFLSKHSDFGLDGL | 3578 | 3592 | Negative |
| hCoVNL63 | orf1ab | AFGLQDCVTGINDDD | 46 | 60 | Negative |
| hCoVNL63 | orf1ab | AITPYVCNASGCGVS | 5014 | 5028 | Negative |
| hCoVNL63 | orf1ab | ALAEHDFFTWKDGRV | 4132 | 4146 | Negative |
| hCoVNL63 | orf1ab | ALCDAMVAKGVVGVL | 4228 | 4242 | Negative |
| hCoVNL63 | orf1ab | ALDGDIYLLFTNSIL | 1450 | 1464 | Negative |
| hCoVNL63 | orf1ab | ALLHRNSLDPFCFDV | 334 | 348 | Negative |
| hCoVNL63 | orf1ab | ALVRLTTEVVRGVRD | 498 | 512 | Negative |
| hCoVNL63 | orf1ab | ATVYLEDKFDLLISD | 6544 | 6558 | Negative |
| hCoVNL63 | orf1ab | CNGTDIDKCVRAFDI | 4064 | 4078 | Negative |
| hCoVNL63 | orf1ab | CRFVAFGLQDCVTGI | 42 | 56 | Negative |
| hCoVNL63 | orf1ab | CVALCDAMVAKGVVG | 4226 | 4240 | Negative |
| hCoVNL63 | orf1ab | CVTFFYEFLDTCFGV | 824 | 838 | Negative |
| hCoVNL63 | orf1ab | DALWLSFKLLGIGGD | 3498 | 3512 | Negative |
| hCoVNL63 | orf1ab | DDFVSAVANAHRYDV | 2346 | 2360 | Negative |
| hCoVNL63 | orf1ab | DDFVTASDTTLRCCT | 6348 | 6362 | Negative |
| hCoVNL63 | orf1ab | DDVVTSLCKKSFGKS | 926 | 940 | Negative |
| hCoVNL63 | orf1ab | DELGVRVLDQSDNNC | 1048 | 1062 | Negative |
| hCoVNL63 | orf1ab | DEQNCDVLKEVLVLT | 4172 | 4186 | Negative |
| hCoVNL63 | orf1ab | DFDITESKYSCKEVL | 2240 | 2254 | Negative |
| hCoVNL63 | orf1ab | DFHQLECIVDDSVRE | 1006 | 1020 | Negative |
| hCoVNL63 | orf1ab | DFHSVEIEAGEVKPF | 1246 | 1260 | Negative |

| | | | | | |
|---|---|---|---|---|---|
| hCoVNL63 | orf1ab | DFVNGFYRLYSGDTF | 2222 | 2236 | Negative |
| hCoVNL63 | orf1ab | DHVFGTDHKFILAIT | 5002 | 5016 | Negative |
| hCoVNL63 | orf1ab | DNHLSTFYPQLQSAE | 6418 | 6432 | Negative |
| hCoVNL63 | orf1ab | DSDVNDMVLSLIKSG | 6694 | 6708 | Negative |
| hCoVNL63 | orf1ab | DTFWRYDFDITESKY | 2234 | 2248 | Negative |
| hCoVNL63 | orf1ab | DVGPFVSFIYFITMS | 1710 | 1724 | Negative |
| hCoVNL63 | orf1ab | DYVIALTGTNQLCAK | 60 | 74 | Negative |
| hCoVNL63 | orf1ab | EGFGGKNILGYSSLC | 3210 | 3224 | Negative |
| hCoVNL63 | orf1ab | EGLGGDNVYCYNTDL | 2608 | 2622 | Negative |
| hCoVNL63 | orf1ab | EGLHEVLTSAMNVIG | 948 | 962 | Negative |
| hCoVNL63 | orf1ab | EGNFSFFDCGVNALD | 1438 | 1452 | Negative |
| hCoVNL63 | orf1ab | EGQLQSLSKDYISSN | 1308 | 1322 | Negative |
| hCoVNL63 | orf1ab | EGSKPYSTLQPNAYY | 2624 | 2638 | Negative |
| hCoVNL63 | orf1ab | EGVNGGSLYVNKHAF | 5972 | 5986 | Negative |
| hCoVNL63 | orf1ab | EILLSIDDGHLWNLF | 716 | 730 | Negative |
| hCoVNL63 | orf1ab | EIMPGKLKQKPMKAE | 3802 | 3816 | Negative |
| hCoVNL63 | orf1ab | EIVNTVSELVYENKF | 5418 | 5432 | Negative |
| hCoVNL63 | orf1ab | ELAQVLTEVVYSNGG | 4696 | 4710 | Negative |
| hCoVNL63 | orf1ab | ELAVNSGLLTACAFS | 3914 | 3928 | Negative |
| hCoVNL63 | orf1ab | ELFIYTNTIWINPVI | 3266 | 3280 | Negative |
| hCoVNL63 | orf1ab | ELGYIADISAFKATL | 4810 | 4824 | Negative |
| hCoVNL63 | orf1ab | ELIQRFAFWTLFCTS | 6610 | 6624 | Negative |
| hCoVNL63 | orf1ab | ELKETVFVEPKDGGQ | 850 | 864 | Negative |
| hCoVNL63 | orf1ab | ELLKEKYTCSITFEM | 1118 | 1132 | Negative |
| hCoVNL63 | orf1ab | ELLLLDFKTAVFVYT | 624 | 638 | Negative |
| hCoVNL63 | orf1ab | ELLLSWESGKVKPPL | 5132 | 5146 | Negative |
| hCoVNL63 | orf1ab | ELLPNVFKLKISTQL | 2830 | 2844 | Negative |
| hCoVNL63 | orf1ab | ELLQFVTDPSLIIAS | 4408 | 4422 | Negative |
| hCoVNL63 | orf1ab | ELPVAVVNDKVFVRY | 6106 | 6120 | Negative |
| hCoVNL63 | orf1ab | ELSHLISGSLGDSGK | 1100 | 1114 | Negative |
| hCoVNL63 | orf1ab | ELSSKVVCTYMDLLL | 6368 | 6382 | Negative |
| hCoVNL63 | orf1ab | ELTLKHFFFAQNGDA | 4472 | 4486 | Negative |
| hCoVNL63 | orf1ab | ELTTMRYFVKIGPIK | 5758 | 5772 | Negative |
| hCoVNL63 | orf1ab | EVIIDNKPYRWMLWC | 6402 | 6416 | Negative |
| hCoVNL63 | orf1ab | EVLKNCNVLENFIVY | 2252 | 2266 | Negative |
| hCoVNL63 | orf1ab | EVLTSAMNVIGQHIK | 952 | 966 | Negative |
| hCoVNL63 | orf1ab | EVSMCTNYDLSVINQ | 5346 | 5360 | Negative |
| hCoVNL63 | orf1ab | EVVGPKELLLSWESG | 5126 | 5140 | Negative |
| hCoVNL63 | orf1ab | EVVVTNLNKSAGWPL | 4530 | 4544 | Negative |
| hCoVNL63 | orf1ab | FDDQPSLQVESANLM | 3124 | 3138 | Negative |
| hCoVNL63 | orf1ab | FDDQVGCLFWIMPYT | 1140 | 1154 | Negative |
| hCoVNL63 | orf1ab | FDIYEGGCIKACEVV | 4518 | 4532 | Negative |
| hCoVNL63 | orf1ab | FDNFNQWHEAKFGVV | 2524 | 2538 | Negative |
| hCoVNL63 | orf1ab | FDNSIQGSYERFTLT | 6200 | 6214 | Negative |
| hCoVNL63 | orf1ab | FDSKGWYDPVENEDI | 4194 | 4208 | Negative |
| hCoVNL63 | orf1ab | FDVYNLWQIFIETNL | 6070 | 6084 | Negative |
| hCoVNL63 | orf1ab | FDYHPNCSDCYDDMC | 4330 | 4344 | Negative |
| hCoVNL63 | orf1ab | FEKVEYGSDTVTYKS | 5170 | 5184 | Negative |
| hCoVNL63 | orf1ab | FFDEGSELTLKHFFF | 4466 | 4480 | Negative |

| | | | | | |
|---|---|---|---|---|---|
| hCoVNL63 | orf1ab | FFNAFVKCNCGSESW | 238 | 252 | Negative |
| hCoVNL63 | orf1ab | FGNTGVCYDFDGVTT | 2580 | 2594 | Negative |
| hCoVNL63 | orf1ab | FLDGLDLTPVIDDVD | 1416 | 1430 | Negative |
| hCoVNL63 | orf1ab | FLGVVGVTMHGSVLR | 3012 | 3026 | Negative |
| hCoVNL63 | orf1ab | FLSKHSDFGLDGLID | 3580 | 3594 | Negative |
| hCoVNL63 | orf1ab | FNDNQAITQVPATSI | 2442 | 2456 | Negative |
| hCoVNL63 | orf1ab | FNSACTRLEGLGGDN | 2600 | 2614 | Negative |
| hCoVNL63 | orf1ab | FNVVKKGCFTGVDGE | 6092 | 6106 | Negative |
| hCoVNL63 | orf1ab | FSVSHNGVFLGVVGV | 3004 | 3018 | Negative |
| hCoVNL63 | orf1ab | FTGVDGELPVAVVND | 6100 | 6114 | Negative |
| hCoVNL63 | orf1ab | FVAFGLQDCVTGIND | 44 | 58 | Negative |
| hCoVNL63 | orf1ab | GCFTGVDGELPVAVV | 6098 | 6112 | Negative |
| hCoVNL63 | orf1ab | GCGVSDVKKLYLGGL | 5024 | 5038 | Negative |
| hCoVNL63 | orf1ab | GCVVEASTDFHQLEC | 998 | 1012 | Negative |
| hCoVNL63 | orf1ab | GDGLIDLLVNVLSIF | 2702 | 2716 | Negative |
| hCoVNL63 | orf1ab | GDNVYCYNTDLIEGS | 2612 | 2626 | Negative |
| hCoVNL63 | orf1ab | GECRDSHKGVCFGFD | 2672 | 2686 | Negative |
| hCoVNL63 | orf1ab | GFDKWYVNDGRVDDG | 2684 | 2698 | Negative |
| hCoVNL63 | orf1ab | GFLETVCSVAYTAGV | 774 | 788 | Negative |
| hCoVNL63 | orf1ab | GFNEWAMANGYTSVS | 3168 | 3182 | Negative |
| hCoVNL63 | orf1ab | GFVSRVIRRERCDMT | 804 | 818 | Negative |
| hCoVNL63 | orf1ab | GGGVARAIDILTEGQ | 1296 | 1310 | Negative |
| hCoVNL63 | orf1ab | GGWNNMLRTLIDGVE | 4634 | 4648 | Negative |
| hCoVNL63 | orf1ab | GHGAGSVVFVDKYMC | 106 | 120 | Negative |
| hCoVNL63 | orf1ab | GHGCACDRTTIQSVD | 4032 | 4046 | Negative |
| hCoVNL63 | orf1ab | GIASGVFGVNLRTNF | 3058 | 3072 | Negative |
| hCoVNL63 | orf1ab | GIAYTTFSGSFDNGH | 1822 | 1836 | Negative |
| hCoVNL63 | orf1ab | GIGGDRCIKISTVQS | 3508 | 3522 | Negative |
| hCoVNL63 | orf1ab | GINVKDVVVESSKSL | 1590 | 1604 | Negative |
| hCoVNL63 | orf1ab | GINYLGDFIQGPFIA | 6636 | 6650 | Negative |
| hCoVNL63 | orf1ab | GKIVVIAGQAFFYSG | 556 | 570 | Negative |
| hCoVNL63 | orf1ab | GKNILGYSSLCDEFT | 3214 | 3228 | Negative |
| hCoVNL63 | orf1ab | GKQTELAVNSGLLTA | 3910 | 3924 | Negative |
| hCoVNL63 | orf1ab | GLCLFWNCNVDMYPE | 5940 | 5954 | Negative |
| hCoVNL63 | orf1ab | GLDGLIDSYFDNSST | 3588 | 3602 | Negative |
| hCoVNL63 | orf1ab | GLIDLLVNVLSIFSS | 2704 | 2718 | Negative |
| hCoVNL63 | orf1ab | GLLTACAFSVDPATT | 3920 | 3934 | Negative |
| hCoVNL63 | orf1ab | GLNVLWNKFVTGDVG | 1698 | 1712 | Negative |
| hCoVNL63 | orf1ab | GLQDCVTGINDDDYV | 48 | 62 | Negative |
| hCoVNL63 | orf1ab | GLQIQTVDSSQGSEY | 5498 | 5512 | Negative |
| hCoVNL63 | orf1ab | GLTVSDDDFVSAVAN | 2340 | 2354 | Negative |
| hCoVNL63 | orf1ab | GLVVKHITNITGVSL | 300 | 314 | Negative |
| hCoVNL63 | orf1ab | GLVWNKDVNTHSVRL | 4390 | 4404 | Negative |
| hCoVNL63 | orf1ab | GLYDDILTNNKPWFV | 386 | 400 | Negative |
| hCoVNL63 | orf1ab | GMFYNQVTLAVASDS | 1 | 14 | Negative |
| hCoVNL63 | orf1ab | GSESWSVGAWDGYLS | 248 | 262 | Negative |
| hCoVNL63 | orf1ab | GSFINGACGSPGYNV | 3076 | 3090 | Negative |
| hCoVNL63 | orf1ab | GSGSGMFYNQVTLAV | 1 | 10 | Negative |
| hCoVNL63 | orf1ab | GSGSGSGMFYNQVTL | 1 | 8 | Negative |

| | | | | | |
|---|---|---|---|---|---|
| hCoVNL63 | orf1ab | GTDHKFILAITPYVC | 5006 | 5020 | Negative |
| hCoVNL63 | orf1ab | GTDIDKCVRAFDIYN | 4066 | 4080 | Negative |
| hCoVNL63 | orf1ab | GTTDNNCWVNATCII | 1672 | 1686 | Negative |
| hCoVNL63 | orf1ab | GTVEFCYLHQIELGS | 3094 | 3108 | Negative |
| hCoVNL63 | orf1ab | GVARAIDILTEGQLQ | 1298 | 1312 | Negative |
| hCoVNL63 | orf1ab | GVCFGFDKWYVNDGR | 2680 | 2694 | Negative |
| hCoVNL63 | orf1ab | GVCYDFDGVTTSDKC | 2584 | 2598 | Negative |
| hCoVNL63 | orf1ab | GVDGELPVAVVNDKV | 6102 | 6116 | Negative |
| hCoVNL63 | orf1ab | GVFDIKNSSVNTVCF | 1228 | 1242 | Negative |
| hCoVNL63 | orf1ab | GVFGVNLRTNFTIKG | 3062 | 3076 | Negative |
| hCoVNL63 | orf1ab | GVFLGVVGVTMHGSV | 3010 | 3024 | Negative |
| hCoVNL63 | orf1ab | GVFVQDPAPIDIDAF | 1180 | 1194 | Negative |
| hCoVNL63 | orf1ab | GVIDISAGMFGLYDD | 376 | 390 | Negative |
| hCoVNL63 | orf1ab | GVKDFNTLSQEGKKY | 2412 | 2426 | Negative |
| hCoVNL63 | orf1ab | GVKGSGHYQTNLYSF | 1206 | 1220 | Negative |
| hCoVNL63 | orf1ab | GVLGDGNALYNTEGG | 3820 | 3834 | Negative |
| hCoVNL63 | orf1ab | GVLPINTDTVLSVAP | 1620 | 1634 | Negative |
| hCoVNL63 | orf1ab | GVLTLDNQDLNGNFY | 4240 | 4254 | Negative |
| hCoVNL63 | orf1ab | GVLVRFVKSIASTVL | 426 | 440 | Negative |
| hCoVNL63 | orf1ab | GVMEPVDYNVVTQRM | 5386 | 5400 | Negative |
| hCoVNL63 | orf1ab | GVMLECEKFNVFNVV | 1330 | 1344 | Negative |
| hCoVNL63 | orf1ab | GVNLQSGKVIFGLKT | 3238 | 3252 | Negative |
| hCoVNL63 | orf1ab | GVSDVKKLYLGGLNY | 5026 | 5040 | Negative |
| hCoVNL63 | orf1ab | GVSERINVVPGVPTN | 2550 | 2564 | Negative |
| hCoVNL63 | orf1ab | GVSFIDYTTTVTSFH | 2484 | 2498 | Negative |
| hCoVNL63 | orf1ab | GVTYQLAWDVIRKDL | 150 | 164 | Negative |
| hCoVNL63 | orf1ab | GVVCYNKDYAELGYI | 4800 | 4814 | Negative |
| hCoVNL63 | orf1ab | GVVGVTMHGSVLRIK | 3014 | 3028 | Negative |
| hCoVNL63 | orf1ab | GVVTTNSDKCPIVVG | 2536 | 2550 | Negative |
| hCoVNL63 | orf1ab | GVWLGDTVTCPRHVI | 2968 | 2982 | Negative |
| hCoVNL63 | orf1ab | GWLGMDVESAHVCGD | 5652 | 5666 | Negative |
| hCoVNL63 | orf1ab | GYICGDGLIDLLVNV | 2698 | 2712 | Negative |
| hCoVNL63 | orf1ab | HDIKVIAKAPKRTGV | 1930 | 1944 | Negative |
| hCoVNL63 | orf1ab | HFNEEFYNFLRLRGF | 4452 | 4466 | Negative |
| hCoVNL63 | orf1ab | HFSMMILSDDGVVCY | 4790 | 4804 | Negative |
| hCoVNL63 | orf1ab | HKGVCFGFDKWYVND | 2678 | 2692 | Negative |
| hCoVNL63 | orf1ab | HKLRSRVKFVNGRVV | 1790 | 1804 | Negative |
| hCoVNL63 | orf1ab | HNFSVSHNGVFLGVV | 3002 | 3016 | Negative |
| hCoVNL63 | orf1ab | HNGVFLGVVGVTMHG | 3008 | 3022 | Negative |
| hCoVNL63 | orf1ab | IAFNVVKKGCFTGVD | 6090 | 6104 | Negative |
| hCoVNL63 | orf1ab | IDGVENPMLMGWDYP | 4644 | 4658 | Negative |
| hCoVNL63 | orf1ab | IDNDINDYVSDADFS | 6524 | 6538 | Negative |
| hCoVNL63 | orf1ab | IELDSPCRFMVETPN | 3862 | 3876 | Negative |
| hCoVNL63 | orf1ab | IGTNVPLQVGFSNGV | 5668 | 5682 | Negative |
| hCoVNL63 | orf1ab | IKQPFSFSFRDELGV | 1038 | 1052 | Negative |
| hCoVNL63 | orf1ab | ILACYEGIASGVFGV | 3052 | 3066 | Negative |
| hCoVNL63 | orf1ab | ILAITPYVCNASGCG | 5012 | 5026 | Negative |
| hCoVNL63 | orf1ab | IRMISAMVLGSKHVN | 4668 | 4682 | Negative |
| hCoVNL63 | orf1ab | IRRERCDMTFPCVSC | 810 | 824 | Negative |

| | | | | | |
|---|---|---|---|---|---|
| hCoVNL63 | orf1ab | ISGFVSRVIRRERCD | 802 | 816 | Negative |
| hCoVNL63 | orf1ab | ITKENVETLTWPLIL | 3776 | 3790 | Negative |
| hCoVNL63 | orf1ab | IYTGDWEGLHEVLTS | 942 | 956 | Negative |
| hCoVNL63 | orf1ab | KAYVDVLCNSFFKEL | 2312 | 2326 | Negative |
| hCoVNL63 | orf1ab | KITEYSWNKYLYELI | 6598 | 6612 | Negative |
| hCoVNL63 | orf1ab | KLVNSELLSTLSVDF | 2292 | 2306 | Negative |
| hCoVNL63 | orf1ab | KNDNVVLKITEDGIN | 1578 | 1592 | Negative |
| hCoVNL63 | orf1ab | KQLGVVSDGVDSFEG | 1606 | 1620 | Negative |
| hCoVNL63 | orf1ab | KSLGKQLGVVSDGVD | 1602 | 1616 | Negative |
| hCoVNL63 | orf1ab | KTLFDSLKFFEIKHA | 5550 | 5564 | Negative |
| hCoVNL63 | orf1ab | KVDFVNGFYRLYSGD | 2220 | 2234 | Negative |
| hCoVNL63 | orf1ab | KVDSIVQKCYELSHL | 1090 | 1104 | Negative |
| hCoVNL63 | orf1ab | KVEYGSDTVTYKSTV | 5172 | 5186 | Negative |
| hCoVNL63 | orf1ab | LDDFVTILKSLDLGV | 6382 | 6396 | Negative |
| hCoVNL63 | orf1ab | LDKQGQLLDTKLNGI | 1466 | 1480 | Negative |
| hCoVNL63 | orf1ab | LDNQDLNGNFYDFGD | 4244 | 4258 | Negative |
| hCoVNL63 | orf1ab | LFEGDKFIGTFESAA | 2844 | 2858 | Negative |
| hCoVNL63 | orf1ab | LGKQLGVVSDGVDSF | 1604 | 1618 | Negative |
| hCoVNL63 | orf1ab | LGMDVESAHVCGDNI | 5654 | 5668 | Negative |
| hCoVNL63 | orf1ab | LGVVSDGVDSFEGVL | 1608 | 1622 | Negative |
| hCoVNL63 | orf1ab | LICGYLVCTYWGILY | 3446 | 3460 | Negative |
| hCoVNL63 | orf1ab | LIDSYFDNSSTLQSV | 3592 | 3606 | Negative |
| hCoVNL63 | orf1ab | LIEGSKPYSTLQPNA | 2622 | 2636 | Negative |
| hCoVNL63 | orf1ab | LIFEFEDDVVTSLCK | 920 | 934 | Negative |
| hCoVNL63 | orf1ab | LILNCERVVKLQNNE | 3788 | 3802 | Negative |
| hCoVNL63 | orf1ab | LLAFFLSKHSDFGLD | 3576 | 3590 | Negative |
| hCoVNL63 | orf1ab | LLLDDFVTILKSLDL | 6380 | 6394 | Negative |
| hCoVNL63 | orf1ab | LNGIAYTTFSGSFDN | 1820 | 1834 | Negative |
| hCoVNL63 | orf1ab | LSDQFKTTGDLAVQI | 5596 | 5610 | Negative |
| hCoVNL63 | orf1ab | LTEGQLQSLSKDYIS | 1306 | 1320 | Negative |
| hCoVNL63 | orf1ab | LTLDNQDLNGNFYDF | 4242 | 4256 | Negative |
| hCoVNL63 | orf1ab | LVTTAGYHFKQLGLV | 4378 | 4392 | Negative |
| hCoVNL63 | orf1ab | MCLEPCNLYNYGAGI | 6448 | 6462 | Negative |
| hCoVNL63 | orf1ab | MDVESAHVCGDNIGT | 5656 | 5670 | Negative |
| hCoVNL63 | orf1ab | MGLTPPLSILKNLGV | 6148 | 6162 | Negative |
| hCoVNL63 | orf1ab | MKGTGVFVQDPAPID | 1176 | 1190 | Negative |
| hCoVNL63 | orf1ab | MLCTKCAYDHVFGTD | 4994 | 5008 | Negative |
| hCoVNL63 | orf1ab | MSSVETVLNLARDGV | 3704 | 3718 | Negative |
| hCoVNL63 | orf1ab | MYGVNLQSGKVIFGL | 3236 | 3250 | Negative |
| hCoVNL63 | orf1ab | NDFVGGFRVLGTTDN | 1662 | 1676 | Negative |
| hCoVNL63 | orf1ab | NLYNYGAGIKLPSGI | 6454 | 6468 | Negative |
| hCoVNL63 | orf1ab | PEVDWVAFYGFEKAA | 1634 | 1648 | Negative |
| hCoVNL63 | orf1ab | PFRVEGNFSFFDCGV | 1434 | 1448 | Negative |
| hCoVNL63 | orf1ab | PGDSFNILACYEGIA | 3046 | 3060 | Negative |
| hCoVNL63 | orf1ab | PIKLVNSELLSTLSV | 2290 | 2304 | Negative |
| hCoVNL63 | orf1ab | PIVVGVSERINVVPG | 2546 | 2560 | Negative |
| hCoVNL63 | orf1ab | PSINDLSFDKNLGRC | 1516 | 1530 | Negative |
| hCoVNL63 | orf1ab | PSLQVESANLMLSDN | 3128 | 3142 | Negative |
| hCoVNL63 | orf1ab | PVCAKSPPGEQFRHL | 5702 | 5716 | Negative |

| | | | | | |
|---|---|---|---|---|---|
| hCoVNL63 | orf1ab | PVFFNAFVKCNCGSE | 236 | 250 | Negative |
| hCoVNL63 | orf1ab | PVFTGDLFYTIKFSG | 586 | 600 | Negative |
| hCoVNL63 | orf1ab | QIGEFIFEKVEYGSD | 5164 | 5178 | Negative |
| hCoVNL63 | orf1ab | QLGLVWNKDVNTHSV | 4388 | 4402 | Negative |
| hCoVNL63 | orf1ab | RFDISIPGSHSLFCT | 5628 | 5642 | Negative |
| hCoVNL63 | orf1ab | RFDTRTRSVFNLEGV | 5960 | 5974 | Negative |
| hCoVNL63 | orf1ab | RGSSAARLEPCNGTD | 4054 | 4068 | Negative |
| hCoVNL63 | orf1ab | RGVRDARIKKAMFTK | 508 | 522 | Negative |
| hCoVNL63 | orf1ab | RKHFSMMILSDDGVV | 4788 | 4802 | Negative |
| hCoVNL63 | orf1ab | RTLATRYCRVGECRD | 2662 | 2676 | Negative |
| hCoVNL63 | orf1ab | RVECYSGFKPNNTSA | 5310 | 5324 | Negative |
| hCoVNL63 | orf1ab | RVGECRDSHKGVCFG | 2670 | 2684 | Negative |
| hCoVNL63 | orf1ab | RVMITKGVMEPVDYN | 5380 | 5394 | Negative |
| hCoVNL63 | orf1ab | RVTAVHSDGMFVASS | 316 | 330 | Negative |
| hCoVNL63 | orf1ab | SAAAGTFVLDMRSYE | 2856 | 2870 | Negative |
| hCoVNL63 | orf1ab | SAFKATLYYQNNVFM | 4818 | 4832 | Negative |
| hCoVNL63 | orf1ab | SARLKRVPLQTIING | 2142 | 2156 | Negative |
| hCoVNL63 | orf1ab | SASKLTIVSPDLESY | 3728 | 3742 | Negative |
| hCoVNL63 | orf1ab | SDGVDSFEGVLPINT | 1612 | 1626 | Negative |
| hCoVNL63 | orf1ab | SEISGFGFAIPSVAV | 14 | 28 | Negative |
| hCoVNL63 | orf1ab | SIASTVLTVSNGVII | 434 | 448 | Negative |
| hCoVNL63 | orf1ab | SKGLNVLWNKFVTGD | 1696 | 1710 | Negative |
| hCoVNL63 | orf1ab | SKHVNCCTATDRFYR | 4678 | 4692 | Negative |
| hCoVNL63 | orf1ab | SKLTDLKCTNVVLLG | 3522 | 3536 | Negative |
| hCoVNL63 | orf1ab | SKSARLKRVPLQTII | 2140 | 2154 | Negative |
| hCoVNL63 | orf1ab | SKVHEVIIDNKPYRW | 6398 | 6412 | Negative |
| hCoVNL63 | orf1ab | SLDPFCFDVNTLLSN | 340 | 354 | Negative |
| hCoVNL63 | orf1ab | SSATDAIIAVELLLL | 614 | 628 | Negative |
| hCoVNL63 | orf1ab | SSIKGDKGVEKLVNW | 6244 | 6258 | Negative |
| hCoVNL63 | orf1ab | SSKVVCTYMDLLLDD | 6370 | 6384 | Negative |
| hCoVNL63 | orf1ab | SSVECYSILAAKTGV | 3182 | 3196 | Negative |
| hCoVNL63 | orf1ab | SVNTSSSEAFLIGIN | 6624 | 6638 | Negative |
| hCoVNL63 | orf1ab | TAFGPLCRKVFIDGV | 4362 | 4376 | Negative |
| hCoVNL63 | orf1ab | TGVSVEQLLASIQHL | 3194 | 3208 | Negative |
| hCoVNL63 | orf1ab | TNVPLQVGFSNGVNF | 5670 | 5684 | Negative |
| hCoVNL63 | orf1ab | TPYVCNASGCGVSDV | 5016 | 5030 | Negative |
| hCoVNL63 | orf1ab | TVGGVSLLSTMTTRQ | 4594 | 4608 | Negative |
| hCoVNL63 | orf1ab | VDFNGVLHKAYVDVL | 2304 | 2318 | Negative |
| hCoVNL63 | orf1ab | VEHLELKETVFVEPK | 846 | 860 | Negative |
| hCoVNL63 | orf1ab | VESSKSLGKQLGVVS | 1598 | 1612 | Negative |
| hCoVNL63 | orf1ab | VIIDKVDFVNGFYRL | 2216 | 2230 | Negative |
| hCoVNL63 | orf1ab | VIREKLAIGGSVAIK | 6584 | 6598 | Negative |
| hCoVNL63 | orf1ab | VITNVGEPIISQPSK | 1804 | 1818 | Negative |
| hCoVNL63 | orf1ab | VKSIASTVLTVSNGV | 432 | 446 | Negative |
| hCoVNL63 | orf1ab | VLDLSKFECKHKATV | 6674 | 6688 | Negative |
| hCoVNL63 | orf1ab | VLDQSDNNCWISTTL | 1054 | 1068 | Negative |
| hCoVNL63 | orf1ab | VLKADDFVTASDTTL | 6344 | 6358 | Negative |
| hCoVNL63 | orf1ab | VLKRWLPPDAIIIDN | 6512 | 6526 | Negative |
| hCoVNL63 | orf1ab | VLNDPVFTGDLFYTI | 582 | 596 | Negative |

| | | | | | |
|---|---|---|---|---|---|
| hCoVNL63 | orf1ab | VRDARIKKAMFTKVV | 510 | 524 | Negative |
| hCoVNL63 | orf1ab | VRLTTEVVRGVRDAR | 500 | 514 | Negative |
| hCoVNL63 | orf1ab | VSHNGVFLGVVGVTM | 3006 | 3020 | Negative |
| hCoVNL63 | orf1ab | VTKFKRVFGDLSYGV | 2746 | 2760 | Negative |
| hCoVNL63 | orf1ab | VTLKDSDVNDMVLSL | 6690 | 6704 | Negative |
| hCoVNL63 | orf1ab | VTTNSDKCPIVVGVS | 2538 | 2552 | Negative |
| hCoVNL63 | orf1ab | VVATYKFVLWDYEAE | 6162 | 6176 | Negative |
| hCoVNL63 | orf1ab | VVGGCVTSNVPTIVS | 1868 | 1882 | Negative |
| hCoVNL63 | orf1ab | VVGVLTLDNQDLNGN | 4238 | 4252 | Negative |
| hCoVNL63 | orf1ab | VVISGFVSRVIRRER | 800 | 814 | Negative |
| hCoVNL63 | orf1ab | VYLICGYLVCTYWGI | 3444 | 3458 | Negative |
| hCoVNL63 | orf1ab | WLGDTVTCPRHVIAP | 2970 | 2984 | Negative |
| hCoVNL63 | orf1ab | WTLNDVKDNDGRPVH | 3758 | 3772 | Negative |
| hCoVNL63 | orf1ab | WYVCLFVVALFIGVS | 2472 | 2486 | Negative |
| hCoVNL63 | orf1ab | YDFGDFVVSLPNMGV | 4254 | 4268 | Negative |
| hCoVNL63 | orf1ab | YKPSVIHDIGNPKGV | 5890 | 5904 | Negative |
| hCoVNL63 | orf1ab | YLGVKGSGHYQTNLY | 1204 | 1218 | Negative |
| hCoVNL63 | orf1ab | YLISDSIVTLEQYST | 1742 | 1756 | Negative |
| hCoVNL63 | orf1ab | YLPYPDPSRILSAGV | 4866 | 4880 | Negative |
| hCoVNL63 | orf1ab | YSKIVCDGSVHYAGV | 3742 | 3756 | Negative |
| hCoVNL63 | orf1ab | YVLTENALVRLTTEV | 492 | 506 | Negative |
| hCoVNL63 | orf1ab | YVVDDIYYPASCNGV | 874 | 888 | Negative |
| hCoVNL63 | orf1ab | YWFNRFFKCTMGVYD | 3460 | 3474 | Negative |
| hCoVNL63 | orf1ab | AEEALSKLSEYLISD | 1732 | 1746 | Overlapping |
| hCoVNL63 | orf1ab | AEHDFFTWKDGRVIY | 4134 | 4148 | Overlapping |
| hCoVNL63 | orf1ab | AVELLLLDFKTAVFV | 622 | 636 | Overlapping |
| hCoVNL63 | orf1ab | CIVDDSVREEVDIIE | 1012 | 1026 | Overlapping |
| hCoVNL63 | orf1ab | DDDYVIALTGTNQLC | 58 | 72 | Overlapping |
| hCoVNL63 | orf1ab | DDSDCDVVQEQVNYV | 6008 | 6022 | Overlapping |
| hCoVNL63 | orf1ab | DEEGGYDVSKPVMIS | 974 | 988 | Overlapping |
| hCoVNL63 | orf1ab | DGCSVIVRRDATFAT | 642 | 656 | Overlapping |
| hCoVNL63 | orf1ab | DGDIYLLFTNSILML | 1452 | 1466 | Overlapping |
| hCoVNL63 | orf1ab | DGELPVAVVNDKVFV | 6104 | 6118 | Overlapping |
| hCoVNL63 | orf1ab | DGENVSKDGFFTYLN | 6568 | 6582 | Overlapping |
| hCoVNL63 | orf1ab | DGFNHQFVTASSATD | 604 | 618 | Overlapping |
| hCoVNL63 | orf1ab | DGGVLGDGNALYNTE | 3818 | 3832 | Overlapping |
| hCoVNL63 | orf1ab | DGLDLTPVIDDVDVV | 1418 | 1432 | Overlapping |
| hCoVNL63 | orf1ab | DGLIDSYFDNSSTLQ | 3590 | 3604 | Overlapping |
| hCoVNL63 | orf1ab | DGNALYNTEGGKTFM | 3824 | 3838 | Overlapping |
| hCoVNL63 | orf1ab | DGRVDDGYICGDGLI | 2692 | 2706 | Overlapping |
| hCoVNL63 | orf1ab | DGSVHYAGVVWTLND | 3748 | 3762 | Overlapping |
| hCoVNL63 | orf1ab | DGVTTSDKCIFNSAC | 2590 | 2604 | Overlapping |
| hCoVNL63 | orf1ab | DGVVPLSVIPATSAS | 3716 | 3730 | Overlapping |
| hCoVNL63 | orf1ab | DHEISVQKKINRMAE | 3654 | 3668 | Overlapping |
| hCoVNL63 | orf1ab | DNSSTLQSVASSFVS | 3598 | 3612 | Overlapping |
| hCoVNL63 | orf1ab | DSIVIGGVTYQLAWD | 144 | 158 | Overlapping |
| hCoVNL63 | orf1ab | DSVREEVDIIEQPFE | 1016 | 1030 | Overlapping |
| hCoVNL63 | orf1ab | DVEVFNRLATSDWTD | 5072 | 5086 | Overlapping |
| hCoVNL63 | orf1ab | DWEGLHEVLTSAMNV | 946 | 960 | Overlapping |

| | | | | | |
|---|---|---|---|---|---|
| hCoVNL63 | orf1ab | DYEAERPFTSYTKSV | 6172 | 6186 | Overlapping |
| hCoVNL63 | orf1ab | DYVSDADFSITGDCA | 6530 | 6544 | Overlapping |
| hCoVNL63 | orf1ab | EADYRCACYAHLAKA | 2900 | 2914 | Overlapping |
| hCoVNL63 | orf1ab | EAERPFTSYTKSVCK | 6174 | 6188 | Overlapping |
| hCoVNL63 | orf1ab | EAKFGVVTTNSDKCP | 2532 | 2546 | Overlapping |
| hCoVNL63 | orf1ab | EALSKLSEYLISDSI | 1734 | 1748 | Overlapping |
| hCoVNL63 | orf1ab | EAPLHCVRNVFDNFN | 2514 | 2528 | Overlapping |
| hCoVNL63 | orf1ab | EASTDFHQLECIVDD | 1002 | 1016 | Overlapping |
| hCoVNL63 | orf1ab | EAYNSILFENGIPLM | 1358 | 1372 | Overlapping |
| hCoVNL63 | orf1ab | EAYNTFTQAGFNIWV | 6052 | 6066 | Overlapping |
| hCoVNL63 | orf1ab | ECKATLGLTVSDDDF | 2334 | 2348 | Overlapping |
| hCoVNL63 | orf1ab | ECNADIVVVDEVSMC | 5336 | 5350 | Overlapping |
| hCoVNL63 | orf1ab | ECYSGFKPNNTSAQY | 5312 | 5326 | Overlapping |
| hCoVNL63 | orf1ab | EDFNFEHVVYGDVSK | 6310 | 6324 | Overlapping |
| hCoVNL63 | orf1ab | EDFYASMYENSTILQ | 4956 | 4970 | Overlapping |
| hCoVNL63 | orf1ab | EDGINVKDVVVESSK | 1588 | 1602 | Overlapping |
| hCoVNL63 | orf1ab | EDIHRVYASLGKIVA | 4206 | 4220 | Overlapping |
| hCoVNL63 | orf1ab | EDKFWCEDFYASMYE | 4950 | 4964 | Overlapping |
| hCoVNL63 | orf1ab | EDKLSVYDIACCMRA | 2378 | 2392 | Overlapping |
| hCoVNL63 | orf1ab | EDVCVCFDNSIQGSY | 6194 | 6208 | Overlapping |
| hCoVNL63 | orf1ab | EEDLTKGPHEFCSQH | 4840 | 4854 | Overlapping |
| hCoVNL63 | orf1ab | EEQDALFALTKRNVL | 4560 | 4574 | Overlapping |
| hCoVNL63 | orf1ab | EESFIDDYYGYLRKH | 4776 | 4790 | Overlapping |
| hCoVNL63 | orf1ab | EESVKSSYAFATLKE | 5112 | 5126 | Overlapping |
| hCoVNL63 | orf1ab | EEVDIIEQPFEEVEH | 1020 | 1034 | Overlapping |
| hCoVNL63 | orf1ab | EEVEHVLSIKQPFSF | 1030 | 1044 | Overlapping |
| hCoVNL63 | orf1ab | EFDHEISVQKKINRM | 3652 | 3666 | Overlapping |
| hCoVNL63 | orf1ab | EFLATFIVCKIVLFV | 2110 | 2124 | Overlapping |
| hCoVNL63 | orf1ab | EFNDLDHTSLVWKHI | 2032 | 2046 | Overlapping |
| hCoVNL63 | orf1ab | EFRDYFNNNTDSIVI | 134 | 148 | Overlapping |
| hCoVNL63 | orf1ab | EFSIVCRFDTRTRSV | 5954 | 5968 | Overlapping |
| hCoVNL63 | orf1ab | EFTLAEVVKQMYGVN | 3226 | 3240 | Overlapping |
| hCoVNL63 | orf1ab | EGDGGVLGDGNALYN | 3816 | 3830 | Overlapping |
| hCoVNL63 | orf1ab | EGDKFIGTFESAAAG | 2846 | 2860 | Overlapping |
| hCoVNL63 | orf1ab | EGFGGKNILGYSSLC | 3210 | 3224 | Overlapping |
| hCoVNL63 | orf1ab | EGGCIKACEVVVTNL | 4522 | 4536 | Overlapping |
| hCoVNL63 | orf1ab | EGGCNTIELDSPCRF | 3856 | 3870 | Overlapping |
| hCoVNL63 | orf1ab | EGGKTFMYAYISNKA | 3832 | 3846 | Overlapping |
| hCoVNL63 | orf1ab | EGGYDVSKPVMISQW | 976 | 990 | Overlapping |
| hCoVNL63 | orf1ab | EGKKYLVKTTKAKGL | 2422 | 2436 | Overlapping |
| hCoVNL63 | orf1ab | EGLGGDNVYCYNTDL | 2608 | 2622 | Overlapping |
| hCoVNL63 | orf1ab | EGLHEVLTSAMNVIG | 948 | 962 | Overlapping |
| hCoVNL63 | orf1ab | EGNFSFFDCGVNALD | 1438 | 1452 | Overlapping |
| hCoVNL63 | orf1ab | EGQLQSLSKDYISSN | 1308 | 1322 | Overlapping |
| hCoVNL63 | orf1ab | EGSKPYSTLQPNAYY | 2624 | 2638 | Overlapping |
| hCoVNL63 | orf1ab | EGVNGGSLYVNKHAF | 5972 | 5986 | Overlapping |
| hCoVNL63 | orf1ab | EHEQSMYNLLNFSGA | 4118 | 4132 | Overlapping |
| hCoVNL63 | orf1ab | EHSLLVEAYNSILFE | 1352 | 1366 | Overlapping |
| hCoVNL63 | orf1ab | EIKHADLHSSQVCGL | 5560 | 5574 | Overlapping |

| | | | | | |
|---|---|---|---|---|---|
| hCoVNL63 | orf1ab | EILARGFGLRTIRTL | 2650 | 2664 | Overlapping |
| hCoVNL63 | orf1ab | EILLSIDDGHLWNLF | 716 | 730 | Overlapping |
| hCoVNL63 | orf1ab | EIMPGKLKQKPMKAE | 3802 | 3816 | Overlapping |
| hCoVNL63 | orf1ab | EISVQKKINRMAEQA | 3656 | 3670 | Overlapping |
| hCoVNL63 | orf1ab | EIVNTVSELVYENKF | 5418 | 5432 | Overlapping |
| hCoVNL63 | orf1ab | EKFINGCGRNVQGHV | 5868 | 5882 | Overlapping |
| hCoVNL63 | orf1ab | EKFNFVTDWLKTLKL | 732 | 746 | Overlapping |
| hCoVNL63 | orf1ab | EKYTCSITFEMSCDC | 1122 | 1136 | Overlapping |
| hCoVNL63 | orf1ab | ELAQVLTEVVYSNGG | 4696 | 4710 | Overlapping |
| hCoVNL63 | orf1ab | ELAVNSGLLTACAFS | 3914 | 3928 | Overlapping |
| hCoVNL63 | orf1ab | ELFIYTNTIWINPVI | 3266 | 3280 | Overlapping |
| hCoVNL63 | orf1ab | ELGYIADISAFKATL | 4810 | 4824 | Overlapping |
| hCoVNL63 | orf1ab | ELIQRFAFWTLFCTS | 6610 | 6624 | Overlapping |
| hCoVNL63 | orf1ab | ELKETVFVEPKDGGQ | 850 | 864 | Overlapping |
| hCoVNL63 | orf1ab | ELLKEKYTCSITFEM | 1118 | 1132 | Overlapping |
| hCoVNL63 | orf1ab | ELLLLDFKTAVFVYT | 624 | 638 | Overlapping |
| hCoVNL63 | orf1ab | ELLLSWESGKVKPPL | 5132 | 5146 | Overlapping |
| hCoVNL63 | orf1ab | ELLPNVFKLKISTQL | 2830 | 2844 | Overlapping |
| hCoVNL63 | orf1ab | ELLQFVTDPSLIIAS | 4408 | 4422 | Overlapping |
| hCoVNL63 | orf1ab | ELPVAVVNDKVFVRY | 6106 | 6120 | Overlapping |
| hCoVNL63 | orf1ab | ELSSKVVCTYMDLLL | 6368 | 6382 | Overlapping |
| hCoVNL63 | orf1ab | ELTLKHFFFAQNGDA | 4472 | 4486 | Overlapping |
| hCoVNL63 | orf1ab | ELTTMRYFVKIGPIK | 5758 | 5772 | Overlapping |
| hCoVNL63 | orf1ab | EMQLFKVGKVDSIVQ | 1082 | 1096 | Overlapping |
| hCoVNL63 | orf1ab | ENALVRLTTEVVRGV | 496 | 510 | Overlapping |
| hCoVNL63 | orf1ab | ENARQAYEDAIANGS | 3620 | 3634 | Overlapping |
| hCoVNL63 | orf1ab | ENEDIHRVYASLGKI | 4204 | 4218 | Overlapping |
| hCoVNL63 | orf1ab | ENGIPLMPLLSCGIF | 1366 | 1380 | Overlapping |
| hCoVNL63 | orf1ab | ENGQLKVFEAPLHCV | 2506 | 2520 | Overlapping |
| hCoVNL63 | orf1ab | ENIAFNVVKKGCFTG | 6088 | 6102 | Overlapping |
| hCoVNL63 | orf1ab | ENLFNKYFKHWSFDY | 4318 | 4332 | Overlapping |
| hCoVNL63 | orf1ab | ENNVCNVCGCWLGHG | 4020 | 4034 | Overlapping |
| hCoVNL63 | orf1ab | ENPMLMGWDYPKCDR | 4648 | 4662 | Overlapping |
| hCoVNL63 | orf1ab | ENSTILQAAGLCVVC | 4964 | 4978 | Overlapping |
| hCoVNL63 | orf1ab | ENVSKDGFFTYLNGV | 6570 | 6584 | Overlapping |
| hCoVNL63 | orf1ab | EPCNGTDIDKCVRAF | 4062 | 4076 | Overlapping |
| hCoVNL63 | orf1ab | EPIISQPSKLLNGIA | 1810 | 1824 | Overlapping |
| hCoVNL63 | orf1ab | EPKDGGQFFVSGDYL | 858 | 872 | Overlapping |
| hCoVNL63 | orf1ab | EQAATQMYKEARSVN | 3668 | 3682 | Overlapping |
| hCoVNL63 | orf1ab | EQAVLKFLDGLDLTP | 1410 | 1424 | Overlapping |
| hCoVNL63 | orf1ab | EQFCIDNCGEPWFLT | 668 | 682 | Overlapping |
| hCoVNL63 | orf1ab | EQPFEEVEHVLSIKQ | 1026 | 1040 | Overlapping |
| hCoVNL63 | orf1ab | EQSMYNLLNFSGALA | 4120 | 4134 | Overlapping |
| hCoVNL63 | orf1ab | EQYSTCDICKSTVVE | 1752 | 1766 | Overlapping |
| hCoVNL63 | orf1ab | ERARTVGGVSLLSTM | 4590 | 4604 | Overlapping |
| hCoVNL63 | orf1ab | ERFLPKCPEILLSID | 708 | 722 | Overlapping |
| hCoVNL63 | orf1ab | ERFTHWFTYVCSLIA | 3370 | 3384 | Overlapping |
| hCoVNL63 | orf1ab | ERLINTISPEKLKNY | 2870 | 2884 | Overlapping |
| hCoVNL63 | orf1ab | ERPFTSYTKSVCKYT | 6176 | 6190 | Overlapping |

| | | | | | |
|---|---|---|---|---|---|
| hCoVNL63 | orf1ab | ESCTIYMCVVPSIND | 1506 | 1520 | Overlapping |
| hCoVNL63 | orf1ab | ESGKVKPPLNRNSVF | 5138 | 5152 | Overlapping |
| hCoVNL63 | orf1ab | ESKVLLERFLPKCPE | 702 | 716 | Overlapping |
| hCoVNL63 | orf1ab | ESWSVGAWDGYLSSC | 250 | 264 | Overlapping |
| hCoVNL63 | orf1ab | ESYSKIVCDGSVHYA | 3740 | 3754 | Overlapping |
| hCoVNL63 | orf1ab | ETIKAKEESVKSSYA | 5106 | 5120 | Overlapping |
| hCoVNL63 | orf1ab | ETLTWPLILNCERVV | 3782 | 3796 | Overlapping |
| hCoVNL63 | orf1ab | ETVLNLARDGVVPLS | 3708 | 3722 | Overlapping |
| hCoVNL63 | orf1ab | EVFNRLATSDWTDVR | 5074 | 5088 | Overlapping |
| hCoVNL63 | orf1ab | EVIIDNKPYRWMLWC | 6402 | 6416 | Overlapping |
| hCoVNL63 | orf1ab | EVKFSVIELATVNLR | 530 | 544 | Overlapping |
| hCoVNL63 | orf1ab | EVKSAIVCASVLKDG | 1766 | 1780 | Overlapping |
| hCoVNL63 | orf1ab | EVLKNCNVLENFIVY | 2252 | 2266 | Overlapping |
| hCoVNL63 | orf1ab | EVLTSAMNVIGQHIK | 952 | 966 | Overlapping |
| hCoVNL63 | orf1ab | EVSMCTNYDLSVINQ | 5346 | 5360 | Overlapping |
| hCoVNL63 | orf1ab | EVVGPKELLLSWESG | 5126 | 5140 | Overlapping |
| hCoVNL63 | orf1ab | EVVVTNLNKSAGWPL | 4530 | 4544 | Overlapping |
| hCoVNL63 | orf1ab | EWKCGYAMPQIYKLQ | 6432 | 6446 | Overlapping |
| hCoVNL63 | orf1ab | EWNAVYRAFGSPFIT | 210 | 224 | Overlapping |
| hCoVNL63 | orf1ab | EYDFLNMDMGVFINK | 6292 | 6306 | Overlapping |
| hCoVNL63 | orf1ab | EYEGGCNTIELDSPC | 3854 | 3868 | Overlapping |
| hCoVNL63 | orf1ab | EYGSDTVTYKSTVTT | 5174 | 5188 | Overlapping |
| hCoVNL63 | orf1ab | EYRKVFYVLLDWVKH | 4914 | 4928 | Overlapping |
| hCoVNL63 | orf1ab | FFYFDDSDCDVVQEQ | 6004 | 6018 | Overlapping |
| hCoVNL63 | orf1ab | FNEDVCVCFDNSIQG | 6192 | 6206 | Overlapping |
| hCoVNL63 | orf1ab | FNRLATSDWTDVRDY | 5076 | 5090 | Overlapping |
| hCoVNL63 | orf1ab | FRDELGVRVLDQSDN | 1046 | 1060 | Overlapping |
| hCoVNL63 | orf1ab | FVDVDFHSVEIEAGE | 1242 | 1256 | Overlapping |
| hCoVNL63 | orf1ab | FYNQVTLAVASDSEI | 2 | 16 | Overlapping |
| hCoVNL63 | orf1ab | GDAEEALSKLSEYLI | 1730 | 1744 | Overlapping |
| hCoVNL63 | orf1ab | GEPWFLTDYNAILQS | 676 | 690 | Overlapping |
| hCoVNL63 | orf1ab | GIPLMPLLSCGIFGV | 1368 | 1382 | Overlapping |
| hCoVNL63 | orf1ab | GKSIIYTGDWEGLHE | 938 | 952 | Overlapping |
| hCoVNL63 | orf1ab | GLEDFNFEHVVYGDV | 6308 | 6322 | Overlapping |
| hCoVNL63 | orf1ab | GLTNQVVKPGHFNEE | 4442 | 4456 | Overlapping |
| hCoVNL63 | orf1ab | GMFVASSSYDALLHR | 324 | 338 | Overlapping |
| hCoVNL63 | orf1ab | GMHKSFYVNANGGTC | 2156 | 2170 | Overlapping |
| hCoVNL63 | orf1ab | GMPVSSIKGDKGVEK | 6240 | 6254 | Overlapping |
| hCoVNL63 | orf1ab | GMSLLDIIVKPVFFN | 226 | 240 | Overlapping |
| hCoVNL63 | orf1ab | GPDVFLHKCYRCPAE | 5404 | 5418 | Overlapping |
| hCoVNL63 | orf1ab | GQHIKLPQFYIYDEE | 962 | 976 | Overlapping |
| hCoVNL63 | orf1ab | GTGVFVQDPAPIDID | 1178 | 1192 | Overlapping |
| hCoVNL63 | orf1ab | GVDSFEGVLPINTDT | 1614 | 1628 | Overlapping |
| hCoVNL63 | orf1ab | GVENPMLMGWDYPKC | 4646 | 4660 | Overlapping |
| hCoVNL63 | orf1ab | GVLESFSVTLLDNQE | 4936 | 4950 | Overlapping |
| hCoVNL63 | orf1ab | HCVRNVFDNFNQWHE | 2518 | 2532 | Overlapping |
| hCoVNL63 | orf1ab | HRHKLRSRVKFVNGR | 1788 | 1802 | Overlapping |
| hCoVNL63 | orf1ab | HRNEHDASGDAVMTR | 5830 | 5844 | Overlapping |
| hCoVNL63 | orf1ab | HVPHPSMDGYCKFKG | 3988 | 4002 | Overlapping |

| | | | | | |
|---|---|---|---|---|---|
| hCoVNL63 | orf1ab | IDVEHLELKETVFVE | 844 | 858 | Overlapping |
| hCoVNL63 | orf1ab | IIEQPFEEVEHVLSI | 1024 | 1038 | Overlapping |
| hCoVNL63 | orf1ab | IIIDNDINDYVSDAD | 6522 | 6536 | Overlapping |
| hCoVNL63 | orf1ab | ISDMYDGRIKFCDGE | 6556 | 6570 | Overlapping |
| hCoVNL63 | orf1ab | KDHNDMLYSPPTISY | 2920 | 2934 | Overlapping |
| hCoVNL63 | orf1ab | KENVETLTWPLILNC | 3778 | 3792 | Overlapping |
| hCoVNL63 | orf1ab | KSEFDHEISVQKKIN | 3650 | 3664 | Overlapping |
| hCoVNL63 | orf1ab | LHEGFGGKNILGYSS | 3208 | 3222 | Overlapping |
| hCoVNL63 | orf1ab | LTENALVRLTTEVVR | 494 | 508 | Overlapping |
| hCoVNL63 | orf1ab | LTPVIDDVDVVKPFR | 1422 | 1436 | Overlapping |
| hCoVNL63 | orf1ab | MEPVDYNVVTQRMCA | 5388 | 5402 | Overlapping |
| hCoVNL63 | orf1ab | MVDPTTVLNDPVFTG | 576 | 590 | Overlapping |
| hCoVNL63 | orf1ab | NEGVLESFSVTLLDN | 4934 | 4948 | Overlapping |
| hCoVNL63 | orf1ab | NEHDASGDAVMTRCL | 5832 | 5846 | Overlapping |
| hCoVNL63 | orf1ab | NFDEQNCDVLKEVLV | 4170 | 4184 | Overlapping |
| hCoVNL63 | orf1ab | NLNEGVLESFSVTLL | 4932 | 4946 | Overlapping |
| hCoVNL63 | orf1ab | NNNTDSIVIGGVTYQ | 140 | 154 | Overlapping |
| hCoVNL63 | orf1ab | NQWHEAKFGVVTTNS | 2528 | 2542 | Overlapping |
| hCoVNL63 | orf1ab | NTGVCYDFDGVTTSD | 2582 | 2596 | Overlapping |
| hCoVNL63 | orf1ab | NVETLTWPLILNCER | 3780 | 3794 | Overlapping |
| hCoVNL63 | orf1ab | PAEIVNTVSELVYEN | 5416 | 5430 | Overlapping |
| hCoVNL63 | orf1ab | PAPIDIDAFPVKPIC | 1186 | 1200 | Overlapping |
| hCoVNL63 | orf1ab | PESVFSVFGDVKLTL | 3428 | 3442 | Overlapping |
| hCoVNL63 | orf1ab | PFEEVEHVLSIKQPF | 1028 | 1042 | Overlapping |
| hCoVNL63 | orf1ab | PIDIDAFPVKPICSS | 1188 | 1202 | Overlapping |
| hCoVNL63 | orf1ab | PIGCLDPIRFCLENN | 4008 | 4022 | Overlapping |
| hCoVNL63 | orf1ab | PIMGLTNCLASECFV | 4280 | 4294 | Overlapping |
| hCoVNL63 | orf1ab | PQFYIYDEEGGYDVS | 968 | 982 | Overlapping |
| hCoVNL63 | orf1ab | PTHKVKLIFEFEDDV | 914 | 928 | Overlapping |
| hCoVNL63 | orf1ab | PTIVSEKISVMDKLD | 1878 | 1892 | Overlapping |
| hCoVNL63 | orf1ab | PVAVVNDKVFVRYGD | 6108 | 6122 | Overlapping |
| hCoVNL63 | orf1ab | PVDYNVVTQRMCAIG | 5390 | 5404 | Overlapping |
| hCoVNL63 | orf1ab | PVENEDIHRVYASLG | 4202 | 4216 | Overlapping |
| hCoVNL63 | orf1ab | PVHVKEITKENVETL | 3770 | 3784 | Overlapping |
| hCoVNL63 | orf1ab | PVIDDVDVVKPFRVE | 1424 | 1438 | Overlapping |
| hCoVNL63 | orf1ab | PVLPKNMWEFRDYFN | 126 | 140 | Overlapping |
| hCoVNL63 | orf1ab | PVMISQWPISNDSNG | 984 | 998 | Overlapping |
| hCoVNL63 | orf1ab | PVNSNVKLLDYDYAT | 5920 | 5934 | Overlapping |
| hCoVNL63 | orf1ab | PVSSIKGDKGVEKLV | 6242 | 6256 | Overlapping |
| hCoVNL63 | orf1ab | RLEGLGGDNVYCYNT | 2606 | 2620 | Overlapping |
| hCoVNL63 | orf1ab | RLEPCNGTDIDKCVR | 4060 | 4074 | Overlapping |
| hCoVNL63 | orf1ab | RVDDGYICGDGLIDL | 2694 | 2708 | Overlapping |
| hCoVNL63 | orf1ab | RVEGNFSFFDCGVNA | 1436 | 1450 | Overlapping |
| hCoVNL63 | orf1ab | RYDFDITESKYSCKE | 2238 | 2252 | Overlapping |
| hCoVNL63 | orf1ab | SDSCNNVNVRDLQRR | 4750 | 4764 | Overlapping |
| hCoVNL63 | orf1ab | SFSDDVIVHDVEPTH | 902 | 916 | Overlapping |
| hCoVNL63 | orf1ab | SKPVMISQWPISNDS | 982 | 996 | Overlapping |
| hCoVNL63 | orf1ab | SSKGQKGDAEEALSK | 1724 | 1738 | Overlapping |
| hCoVNL63 | orf1ab | SSKSLGKQLGVVSDG | 1600 | 1614 | Overlapping |

| | | | | | |
|---|---|---|---|---|---|
| hCoVNL63 | orf1ab | SSKVVCTYMDLLLDD | 6370 | 6384 | Overlapping |
| hCoVNL63 | orf1ab | SSKVVLSGEWNAVYR | 202 | 216 | Overlapping |
| hCoVNL63 | orf1ab | SVEESFIDDYYGYLR | 4774 | 4788 | Overlapping |
| hCoVNL63 | orf1ab | SVTEDVKFAASTGVI | 364 | 378 | Overlapping |
| hCoVNL63 | orf1ab | SVTLLDNQEDKFWCE | 4942 | 4956 | Overlapping |
| hCoVNL63 | orf1ab | TDFNEDVCVCFDNSI | 6190 | 6204 | Overlapping |
| hCoVNL63 | orf1ab | TDYNAILQSNNPQCA | 682 | 696 | Overlapping |
| hCoVNL63 | orf1ab | TEDVKFAASTGVIDI | 366 | 380 | Overlapping |
| hCoVNL63 | orf1ab | TGCCDNSYFDSKGWY | 4186 | 4200 | Overlapping |
| hCoVNL63 | orf1ab | TGDVGPFVSFIYFIT | 1708 | 1722 | Overlapping |
| hCoVNL63 | orf1ab | TGDWEGLHEVLTSAM | 944 | 958 | Overlapping |
| hCoVNL63 | orf1ab | TGINDDDYVIALTGT | 54 | 68 | Overlapping |
| hCoVNL63 | orf1ab | TGSLDVEVFNRLATS | 5068 | 5082 | Overlapping |
| hCoVNL63 | orf1ab | TMMDLVYAMRNFDEQ | 4160 | 4174 | Overlapping |
| hCoVNL63 | orf1ab | TTEVKFSVIELATVN | 528 | 542 | Overlapping |
| hCoVNL63 | orf1ab | VDDSVREEVDIIEQP | 1014 | 1028 | Overlapping |
| hCoVNL63 | orf1ab | VDIIEQPFEEVEHVL | 1022 | 1036 | Overlapping |
| hCoVNL63 | orf1ab | VEPTHKVKLIFEFED | 912 | 926 | Overlapping |
| hCoVNL63 | orf1ab | VEQLLASIQHLHEGF | 3198 | 3212 | Overlapping |
| hCoVNL63 | orf1ab | VLIDYDHAYSTMRLH | 2988 | 3002 | Overlapping |
| hCoVNL63 | orf1ab | VMDVNDCFKNDNVVL | 1570 | 1584 | Overlapping |
| hCoVNL63 | orf1ab | VMEHEQSMYNLLNFS | 4116 | 4130 | Overlapping |
| hCoVNL63 | orf1ab | VPSDSCNNVNVRDLQ | 4748 | 4762 | Overlapping |
| hCoVNL63 | orf1ab | VREEVDIIEQPFEEV | 1018 | 1032 | Overlapping |
| hCoVNL63 | orf1ab | VVGPTTEVKFSVIEL | 524 | 538 | Overlapping |
| hCoVNL63 | orf1ab | VYSSNEEQAVLKFLD | 1404 | 1418 | Overlapping |
| hCoVNL63 | orf1ab | WHEAKFGVVTTNSDK | 2530 | 2544 | Overlapping |
| hCoVNL63 | orf1ab | WVEEDLTKGPHEFCS | 4838 | 4852 | Overlapping |
| hCoVNL63 | orf1ab | YAISGKERARTVGGV | 4584 | 4598 | Overlapping |
| hCoVNL63 | orf1ab | YDPVENEDIHRVYAS | 4200 | 4214 | Overlapping |
| hCoVNL63 | orf1ab | YFDDSDCDVVQEQVN | 6006 | 6020 | Overlapping |
| hCoVNL63 | orf1ab | YFNNNTDSIVIGGVT | 138 | 152 | Overlapping |
| hCoVNL63 | orf1ab | YPEFSIVCRFDTRTR | 5952 | 5966 | Overlapping |
| hCoVNL63 | spike | ALNAFVSQVLNKYTE | 1069 | 1083 | Positive |
| hCoVNL63 | spike | ANFLDDNVLPETYVA | 461 | 475 | Positive |
| hCoVNL63 | spike | ANSLDNLKSGVIVFK | 304 | 318 | Positive |
| hCoVNL63 | spike | ATGSDVNCNGYQHNS | 277 | 291 | Positive |
| hCoVNL63 | spike | ATVTVNVTTHNGRVV | 188 | 202 | Positive |
| hCoVNL63 | spike | CCDCGSTKLPYYEFE | 1337 | 1351 | Positive |
| hCoVNL63 | spike | CCNCLTSSMRGCCDC | 1326 | 1340 | Positive |
| hCoVNL63 | spike | CKTIEDALRLSAHLE | 811 | 825 | Positive |
| hCoVNL63 | spike | CNGYQHNSVVDVMRY | 284 | 298 | Positive |
| hCoVNL63 | spike | DAERMAMYTGSLIGG | 922 | 936 | Positive |
| hCoVNL63 | spike | DCGSTKLPYYEFEKV | 1339 | 1353 | Positive |
| hCoVNL63 | spike | DGFYSANFLDDNVLP | 456 | 470 | Positive |
| hCoVNL63 | spike | DGLLFLHTVLLPTDY | 1124 | 1138 | Positive |
| hCoVNL63 | spike | DGRIPNGFPFNNWFL | 223 | 237 | Positive |
| hCoVNL63 | spike | DVNCNGYQHNSVVDV | 281 | 295 | Positive |
| hCoVNL63 | spike | FDLTPFNLTYLNLSS | 1236 | 1250 | Positive |

| | | | | | |
|---|---|---|---|---|---|
| hCoVNL63 | spike | FDSNAFSLANVTSFG | 835 | 849 | Positive |
| hCoVNL63 | spike | FGICADGSLIPVRPR | 734 | 748 | Positive |
| hCoVNL63 | spike | FINSTINTTHVSTFV | 352 | 366 | Positive |
| hCoVNL63 | spike | FNATGSDVNCNGYQH | 275 | 289 | Positive |
| hCoVNL63 | spike | GNPRCKNLLKQYTSA | 796 | 810 | Positive |
| hCoVNL63 | spike | GPSSQPYYCFINSTI | 343 | 357 | Positive |
| hCoVNL63 | spike | GSDVNCNGYQHNSVV | 279 | 293 | Positive |
| hCoVNL63 | spike | GSGMKLFLILLVLPL | 1 | 12 | Positive |
| hCoVNL63 | spike | GSGSGSGMKLFLILL | 1 | 8 | Positive |
| hCoVNL63 | spike | GSPGDSSWHIYLKSG | 534 | 548 | Positive |
| hCoVNL63 | spike | HTDINFTATASFGGS | 482 | 496 | Positive |
| hCoVNL63 | spike | IEAVNFNVTTASATD | 397 | 411 | Positive |
| hCoVNL63 | spike | IFSVQQDGRIPNGFP | 217 | 231 | Positive |
| hCoVNL63 | spike | IPDYVDVNKTLQEFA | 1211 | 1225 | Positive |
| hCoVNL63 | spike | IPSNWTTSVQVEYLQ | 765 | 779 | Positive |
| hCoVNL63 | spike | IPVRPRNSSDNGISA | 743 | 757 | Positive |
| hCoVNL63 | spike | ISRVELHTVIPDYVD | 1202 | 1216 | Positive |
| hCoVNL63 | spike | IYDRLDSIQADQQVD | 1046 | 1060 | Positive |
| hCoVNL63 | spike | KIQDVVNQQGSALNH | 1013 | 1027 | Positive |
| hCoVNL63 | spike | KLPYYEFEKVHVQGS | 1344 | 1356 | Positive |
| hCoVNL63 | spike | KLQCEHLQFGLQDGF | 444 | 458 | Positive |
| hCoVNL63 | spike | KSGSPGDSSWHIYLK | 532 | 546 | Positive |
| hCoVNL63 | spike | LEATWHYTSYTIVGA | 581 | 595 | Positive |
| hCoVNL63 | spike | LETNDVSSMLTFDSN | 824 | 838 | Positive |
| hCoVNL63 | spike | LFSKVVTSGLGTVDV | 877 | 891 | Positive |
| hCoVNL63 | spike | LGGLTSAAAIPFSLA | 939 | 953 | Positive |
| hCoVNL63 | spike | LNGNTSVCVRTSHFS | 509 | 523 | Positive |
| hCoVNL63 | spike | LPTDYKNVKAWSGIC | 1134 | 1148 | Positive |
| hCoVNL63 | spike | LPVLSDFVQIYNCNV | 1183 | 1197 | Positive |
| hCoVNL63 | spike | LQCEHLQFGLQDGFY | 445 | 459 | Positive |
| hCoVNL63 | spike | NAFSLANVTSFGDYN | 838 | 852 | Positive |
| hCoVNL63 | spike | NAFVSQVLNKYTEVR | 1071 | 1085 | Positive |
| hCoVNL63 | spike | NANLSMLQLGVPDNS | 22 | 36 | Positive |
| hCoVNL63 | spike | NASSSFDCIVNLLFT | 119 | 133 | Positive |
| hCoVNL63 | spike | NASVTLKICKFSRNT | 98 | 112 | Positive |
| hCoVNL63 | spike | NATGSDVNCNGYQHN | 276 | 290 | Positive |
| hCoVNL63 | spike | NATVTVNVTTHNGRV | 187 | 201 | Positive |
| hCoVNL63 | spike | NCLTSSMRGCCDCGS | 1328 | 1342 | Positive |
| hCoVNL63 | spike | NCNGYQHNSVVDVMR | 283 | 297 | Positive |
| hCoVNL63 | spike | NCNVTFVNISRVELH | 1194 | 1208 | Positive |
| hCoVNL63 | spike | NDVSSMLTFDSNAFS | 827 | 841 | Positive |
| hCoVNL63 | spike | NESRYGLQNLLQLPN | 699 | 713 | Positive |
| hCoVNL63 | spike | NFDLTPFNLTYLNLS | 1235 | 1249 | Positive |
| hCoVNL63 | spike | NFLDDNVLPETYVAL | 462 | 476 | Positive |
| hCoVNL63 | spike | NFNVTTASATDFWTV | 401 | 415 | Positive |
| hCoVNL63 | spike | NFPLEATWHYTSYTI | 578 | 592 | Positive |
| hCoVNL63 | spike | NGFKYFDLGFIEAVN | 387 | 401 | Positive |
| hCoVNL63 | spike | NGRVVNYTVCDDCNG | 198 | 212 | Positive |
| hCoVNL63 | spike | NHLTSQLRHNFQAIS | 1026 | 1040 | Positive |

| | | | | | |
|---|---|---|---|---|---|
| hCoVNL63 | spike | NHRSAFALHTGYYDA | 70 | 84 | Positive |
| hCoVNL63 | spike | NIRSSRIAGRSALED | 861 | 875 | Positive |
| hCoVNL63 | spike | NISRVELHTVIPDYV | 1201 | 1215 | Positive |
| hCoVNL63 | spike | NKAINNIVASFSSVN | 980 | 994 | Positive |
| hCoVNL63 | spike | NKIQDVVNQQGSALN | 1012 | 1026 | Positive |
| hCoVNL63 | spike | NKTLQEFAQNLPKYV | 1218 | 1232 | Positive |
| hCoVNL63 | spike | NKYTEVRGSRRLAQQ | 1079 | 1093 | Positive |
| hCoVNL63 | spike | NLKSGVIVFKTLQYD | 309 | 323 | Positive |
| hCoVNL63 | spike | NLLGFKNVSTGNIFI | 660 | 674 | Positive |
| hCoVNL63 | spike | NLLKQYTSACKTIED | 802 | 816 | Positive |
| hCoVNL63 | spike | NLLQLPNFYYVSNGG | 707 | 721 | Positive |
| hCoVNL63 | spike | NLLYCDSPFEKLQCE | 434 | 448 | Positive |
| hCoVNL63 | spike | NLNFSANSLDNLKSG | 299 | 313 | Positive |
| hCoVNL63 | spike | NLPKYVKPNFDLTPF | 1227 | 1241 | Positive |
| hCoVNL63 | spike | NLSIPSNWTTSVQVE | 762 | 776 | Positive |
| hCoVNL63 | spike | NLSMLQLGVPDNSST | 24 | 38 | Positive |
| hCoVNL63 | spike | NLSSELKQLEAKTAS | 1247 | 1261 | Positive |
| hCoVNL63 | spike | NLSSVLPQRNIRSSR | 852 | 866 | Positive |
| hCoVNL63 | spike | NLTYLNLSSELKQLE | 1242 | 1256 | Positive |
| hCoVNL63 | spike | NLVLNNCTKYNIYDY | 621 | 635 | Positive |
| hCoVNL63 | spike | NNCTKYNIYDYVGTG | 625 | 639 | Positive |
| hCoVNL63 | spike | NNFQKFKTICFSTVE | 558 | 572 | Positive |
| hCoVNL63 | spike | NNIVASFSSVNDAIT | 984 | 998 | Positive |
| hCoVNL63 | spike | NNWFLLTNGSTLVDG | 233 | 247 | Positive |
| hCoVNL63 | spike | NQKILAASFNKAINN | 971 | 985 | Positive |
| hCoVNL63 | spike | NQPDQVAVYQQSIIG | 679 | 693 | Positive |
| hCoVNL63 | spike | NQSLAGGITYVSNSG | 645 | 659 | Positive |
| hCoVNL63 | spike | NRFENYIKWPWWVWL | 1287 | 1301 | Positive |
| hCoVNL63 | spike | NRVKSGSPGDSSWHI | 529 | 543 | Positive |
| hCoVNL63 | spike | NRYGFCGNGTHIFSI | 1104 | 1118 | Positive |
| hCoVNL63 | spike | NSGNLLGFKNVSTGN | 657 | 671 | Positive |
| hCoVNL63 | spike | NSLDNLKSGVIVFKT | 305 | 319 | Positive |
| hCoVNL63 | spike | NSSDNGISAIITANL | 749 | 763 | Positive |
| hCoVNL63 | spike | NSSSGVLDTTIPFGP | 330 | 344 | Positive |
| hCoVNL63 | spike | NSSTIVTGLLPTHWF | 35 | 49 | Positive |
| hCoVNL63 | spike | NSTYVDLKLLNRFEN | 1277 | 1291 | Positive |
| hCoVNL63 | spike | NSVVDVMRYNLNFSA | 290 | 304 | Positive |
| hCoVNL63 | spike | NVLPETYVALPIYYQ | 467 | 481 | Positive |
| hCoVNL63 | spike | NVSATNIQNLLYCDS | 426 | 440 | Positive |
| hCoVNL63 | spike | NVSTGNIFIVTPCNQ | 666 | 680 | Positive |
| hCoVNL63 | spike | NVTFVNISRVELHTV | 1196 | 1210 | Positive |
| hCoVNL63 | spike | NVTTHNGRVVNYTVC | 193 | 207 | Positive |
| hCoVNL63 | spike | NWTTSVQVEYLQITS | 768 | 782 | Positive |
| hCoVNL63 | spike | NYVALQTDVLQENQK | 959 | 973 | Positive |
| hCoVNL63 | spike | PLEATWHYTSYTIVG | 580 | 594 | Positive |
| hCoVNL63 | spike | PLGITISGETVRLHL | 139 | 153 | Positive |
| hCoVNL63 | spike | PRNSSDNGISAIITA | 747 | 761 | Positive |
| hCoVNL63 | spike | PTDYKNVKAWSGICV | 1135 | 1149 | Positive |
| hCoVNL63 | spike | PTHWFCANQSTSVYS | 45 | 59 | Positive |

| | | | | | |
|---|---|---|---|---|---|
| hCoVNL63 | spike | PVPGLKSSTGFVYFN | 262 | 276 | Positive |
| hCoVNL63 | spike | PVRPRNSSDNGISAI | 744 | 758 | Positive |
| hCoVNL63 | spike | PYPVSGIREFSNLVL | 610 | 624 | Positive |
| hCoVNL63 | spike | QEFAQNLPKYVKPNF | 1222 | 1236 | Positive |
| hCoVNL63 | spike | QGSALNHLTSQLRHN | 1021 | 1035 | Positive |
| hCoVNL63 | spike | QHNSVVDVMRYNLNF | 288 | 302 | Positive |
| hCoVNL63 | spike | QNLLYCDSPFEKLQC | 433 | 447 | Positive |
| hCoVNL63 | spike | QQGSALNHLTSQLRH | 1020 | 1034 | Positive |
| hCoVNL63 | spike | REFSNLVLNNCTKYN | 617 | 631 | Positive |
| hCoVNL63 | spike | RFENYIKWPWWVWLI | 1288 | 1302 | Positive |
| hCoVNL63 | spike | RNSSDNGISAIITAN | 748 | 762 | Positive |
| hCoVNL63 | spike | RPRNSSDNGISAIIT | 746 | 760 | Positive |
| hCoVNL63 | spike | RTSHFSIRYIYNRVK | 518 | 532 | Positive |
| hCoVNL63 | spike | SAIITANLSIPSNWT | 756 | 770 | Positive |
| hCoVNL63 | spike | SALEDLLFSKVVTSG | 871 | 885 | Positive |
| hCoVNL63 | spike | SCYVCKPHQVNISLN | 496 | 510 | Positive |
| hCoVNL63 | spike | SDFVQIYNCNVTFVN | 1187 | 1201 | Positive |
| hCoVNL63 | spike | SDNGISAIITANLSI | 751 | 765 | Positive |
| hCoVNL63 | spike | SDNGVFRVTSRVMFQ | 1166 | 1180 | Positive |
| hCoVNL63 | spike | SDVNCNGYQHNSVVD | 280 | 294 | Positive |
| hCoVNL63 | spike | SELKQLEAKTASLFQ | 1250 | 1264 | Positive |
| hCoVNL63 | spike | SGSGMKLFLILLVLP | 1 | 11 | Positive |
| hCoVNL63 | spike | SMLQLGVPDNSSTIV | 26 | 40 | Positive |
| hCoVNL63 | spike | SMLTFDSNAFSLANV | 831 | 845 | Positive |
| hCoVNL63 | spike | SRVELHTVIPDYVDV | 1203 | 1217 | Positive |
| hCoVNL63 | spike | SSSGVLDTTIPFGPS | 331 | 345 | Positive |
| hCoVNL63 | spike | SSTIVTGLLPTHWFC | 36 | 50 | Positive |
| hCoVNL63 | spike | STKLPYYEFEKVHVQ | 1342 | 1356 | Positive |
| hCoVNL63 | spike | STVEVPGSCNFPLEA | 569 | 583 | Positive |
| hCoVNL63 | spike | TAEAIHTVTIALNKI | 1000 | 1014 | Positive |
| hCoVNL63 | spike | TDFWTVAFATFVDVL | 410 | 424 | Positive |
| hCoVNL63 | spike | TDNIFSVQQDGRIPN | 214 | 228 | Positive |
| hCoVNL63 | spike | TDVLQENQKILAASF | 965 | 979 | Positive |
| hCoVNL63 | spike | TDYKNVKAWSGICVD | 1136 | 1150 | Positive |
| hCoVNL63 | spike | TEQLGAPLGITISGE | 133 | 147 | Positive |
| hCoVNL63 | spike | TEVRGSRRLAQQKIN | 1082 | 1096 | Positive |
| hCoVNL63 | spike | TFDSNAFSLANVTSF | 834 | 848 | Positive |
| hCoVNL63 | spike | TGVPYPVSGIREFSN | 607 | 621 | Positive |
| hCoVNL63 | spike | TGYYDANQYYIYVTN | 79 | 93 | Positive |
| hCoVNL63 | spike | TIEDALRLSAHLETN | 813 | 827 | Positive |
| hCoVNL63 | spike | TKLPYYEFEKVHVQG | 1343 | 1356 | Positive |
| hCoVNL63 | spike | TKLSVKCYFNYSCVF | 169 | 183 | Positive |
| hCoVNL63 | spike | TSGLGTVDVDYKSCT | 883 | 897 | Positive |
| hCoVNL63 | spike | TTHNGRVVNYTVCDD | 195 | 209 | Positive |
| hCoVNL63 | spike | TTHVSTFVGILPPTV | 359 | 373 | Positive |
| hCoVNL63 | spike | TVELQGLIDQINSTY | 1266 | 1280 | Positive |
| hCoVNL63 | spike | TVEVPGSCNFPLEAT | 570 | 584 | Positive |
| hCoVNL63 | spike | TYVALPIYYQHTDIN | 472 | 486 | Positive |
| hCoVNL63 | spike | VADAERMAMYTGSLI | 920 | 934 | Positive |

| | | | | | |
|---|---|---|---|---|---|
| hCoVNL63 | spike | VDGVSRLYQPLRLTC | 245 | 259 | Positive |
| hCoVNL63 | spike | VDVDYKSCTKGLSIA | 889 | 903 | Positive |
| hCoVNL63 | spike | VELHTVIPDYVDVNK | 1205 | 1219 | Positive |
| hCoVNL63 | spike | VEYLQITSTPIVVDC | 775 | 789 | Positive |
| hCoVNL63 | spike | VFCCLSTGCCGCCNC | 1315 | 1329 | Positive |
| hCoVNL63 | spike | VLPETYVALPIYYQH | 468 | 482 | Positive |
| hCoVNL63 | spike | VNCNGYQHNSVVDVM | 282 | 296 | Positive |
| hCoVNL63 | spike | VPDNSSTIVTGLLPT | 32 | 46 | Positive |
| hCoVNL63 | spike | VTNEIGLNASVTLKI | 91 | 105 | Positive |
| hCoVNL63 | spike | VVDVMRYNLNFSANS | 292 | 306 | Positive |
| hCoVNL63 | spike | VVTSGLGTVDVDYKS | 881 | 895 | Positive |
| hCoVNL63 | spike | WPVPGLKSSTGFVYF | 261 | 275 | Positive |
| hCoVNL63 | spike | YEFEKVHVQGSGSGS | 1348 | 1356 | Positive |
| hCoVNL63 | spike | YVALPIYYQHTDINF | 473 | 487 | Positive |
| hCoVNL63 | spike | YVTNEIGLNASVTLK | 90 | 104 | Positive |
| hCoVNL63 | spike | ACAQYYNGIMVLPGV | 906 | 920 | Negative |
| hCoVNL63 | spike | ADAERMAMYTGSLIG | 921 | 935 | Negative |
| hCoVNL63 | spike | ADGSLIPVRPRNSSD | 738 | 752 | Negative |
| hCoVNL63 | spike | AERMAMYTGSLIGGM | 923 | 937 | Negative |
| hCoVNL63 | spike | AGRSALEDLLFSKVV | 868 | 882 | Negative |
| hCoVNL63 | spike | ALEDLLFSKVVTSGL | 872 | 886 | Negative |
| hCoVNL63 | spike | ALYVTWSEGNSITGV | 595 | 609 | Negative |
| hCoVNL63 | spike | AQYYNGIMVLPGVAD | 908 | 922 | Negative |
| hCoVNL63 | spike | ASLFQTTVELQGLID | 1260 | 1274 | Negative |
| hCoVNL63 | spike | ATFVDVLVNVSATNI | 418 | 432 | Negative |
| hCoVNL63 | spike | ATNIQNLLYCDSPFE | 429 | 443 | Negative |
| hCoVNL63 | spike | AWSGICVDGIYGYVL | 1143 | 1157 | Negative |
| hCoVNL63 | spike | CTKYNIYDYVGTGII | 627 | 641 | Negative |
| hCoVNL63 | spike | DDCNGYTDNIFSVQQ | 208 | 222 | Negative |
| hCoVNL63 | spike | DDNVLPETYVALPIY | 465 | 479 | Negative |
| hCoVNL63 | spike | DGRIPNGFPFNNWFL | 223 | 237 | Negative |
| hCoVNL63 | spike | DVLFYCSNSSSGVLD | 323 | 337 | Negative |
| hCoVNL63 | spike | DYVDVNKTLQEFAQN | 1213 | 1227 | Negative |
| hCoVNL63 | spike | EGNSITGVPYPVSGI | 602 | 616 | Negative |
| hCoVNL63 | spike | EIVVARTGQFYINGF | 375 | 389 | Negative |
| hCoVNL63 | spike | ELHTVIPDYVDVNKT | 1206 | 1220 | Negative |
| hCoVNL63 | spike | ELKQLEAKTASLFQT | 1251 | 1265 | Negative |
| hCoVNL63 | spike | ELQGLIDQINSTYVD | 1268 | 1282 | Negative |
| hCoVNL63 | spike | FEKLQCEHLQFGLQD | 442 | 456 | Negative |
| hCoVNL63 | spike | FENYIKWPWWVWLII | 1289 | 1303 | Negative |
| hCoVNL63 | spike | FGDYNLSSVLPQRNI | 848 | 862 | Negative |
| hCoVNL63 | spike | FSANSLDNLKSGVIV | 302 | 316 | Negative |
| hCoVNL63 | spike | FSIRYIYNRVKSGSP | 522 | 536 | Negative |
| hCoVNL63 | spike | FSVVNATVTVNVTTH | 183 | 197 | Negative |
| hCoVNL63 | spike | FVSQVLNKYTEVRGS | 1073 | 1087 | Negative |
| hCoVNL63 | spike | FYINGFKYFDLGFIE | 384 | 398 | Negative |
| hCoVNL63 | spike | GAMTAVNESRYGLQN | 693 | 707 | Negative |
| hCoVNL63 | spike | GETVRLHLYNVTRTF | 146 | 160 | Negative |
| hCoVNL63 | spike | GICADGSLIPVRPRN | 735 | 749 | Negative |

| | | | | | |
|---|---|---|---|---|---|
| hCoVNL63 | spike | GICVDGIYGYVLRQP | 1146 | 1160 | Negative |
| hCoVNL63 | spike | GIIRSSNQSLAGGIT | 639 | 653 | Negative |
| hCoVNL63 | spike | GILPPTVREIVVART | 367 | 381 | Negative |
| hCoVNL63 | spike | GIMVLPGVADAERMA | 913 | 927 | Negative |
| hCoVNL63 | spike | GISAIITANLSIPSN | 754 | 768 | Negative |
| hCoVNL63 | spike | GITISGETVRLHLYN | 141 | 155 | Negative |
| hCoVNL63 | spike | GITYVSNSGNLLGFK | 651 | 665 | Negative |
| hCoVNL63 | spike | GLGTVDVDYKSCTKG | 885 | 899 | Negative |
| hCoVNL63 | spike | GLIDQINSTYVDLKL | 1271 | 1285 | Negative |
| hCoVNL63 | spike | GLKSSTGFVYFNATG | 265 | 279 | Negative |
| hCoVNL63 | spike | GLLPTHWFCANQSTS | 42 | 56 | Negative |
| hCoVNL63 | spike | GLNASVTLKICKFSR | 96 | 110 | Negative |
| hCoVNL63 | spike | GLQDGFYSANFLDDN | 453 | 467 | Negative |
| hCoVNL63 | spike | GLQNLLQLPNFYYVS | 704 | 718 | Negative |
| hCoVNL63 | spike | GLSIADLACAQYYNG | 899 | 913 | Negative |
| hCoVNL63 | spike | GLTSAAAIPFSLALQ | 941 | 955 | Negative |
| hCoVNL63 | spike | GMKLFLILLVLPLAS | 1 | 14 | Negative |
| hCoVNL63 | spike | GMVLGGLTSAAAIPF | 936 | 950 | Negative |
| hCoVNL63 | spike | GNSITGVPYPVSGIR | 603 | 617 | Negative |
| hCoVNL63 | spike | GQFYINGFKYFDLGF | 382 | 396 | Negative |
| hCoVNL63 | spike | GRIPNGFPFNNWFLL | 224 | 238 | Negative |
| hCoVNL63 | spike | GRSALEDLLFSKVVT | 869 | 883 | Negative |
| hCoVNL63 | spike | GRVVNYTVCDDCNGY | 199 | 213 | Negative |
| hCoVNL63 | spike | GSGSGMKLFLILLVL | 1 | 10 | Negative |
| hCoVNL63 | spike | GSGSGSGMKLFLILL | 1 | 8 | Negative |
| hCoVNL63 | spike | GTGIIRSSNQSLAGG | 637 | 651 | Negative |
| hCoVNL63 | spike | GTHIFSIVNSAPDGL | 1112 | 1126 | Negative |
| hCoVNL63 | spike | GTVDVDYKSCTKGLS | 887 | 901 | Negative |
| hCoVNL63 | spike | GVFRVTSRVMFQPRL | 1169 | 1183 | Negative |
| hCoVNL63 | spike | GVIVFKTLQYDVLFY | 313 | 327 | Negative |
| hCoVNL63 | spike | GVLDTTIPFGPSSQP | 334 | 348 | Negative |
| hCoVNL63 | spike | GVSRLYQPLRLTCLW | 247 | 261 | Negative |
| hCoVNL63 | spike | GYTDNIFSVQQDGRI | 212 | 226 | Negative |
| hCoVNL63 | spike | GYVLRQPNLVLYSDN | 1154 | 1168 | Negative |
| hCoVNL63 | spike | HIFSIVNSAPDGLLF | 1114 | 1128 | Negative |
| hCoVNL63 | spike | IAGRSALEDLLFSKV | 867 | 881 | Negative |
| hCoVNL63 | spike | IIRSSNQSLAGGITY | 640 | 654 | Negative |
| hCoVNL63 | spike | ILPPTVREIVVARTG | 368 | 382 | Negative |
| hCoVNL63 | spike | IMVLPGVADAERMAM | 914 | 928 | Negative |
| hCoVNL63 | spike | IRSSNQSLAGGITYV | 641 | 655 | Negative |
| hCoVNL63 | spike | IYVTNEIGLNASVTL | 89 | 103 | Negative |
| hCoVNL63 | spike | KGLSIADLACAQYYN | 898 | 912 | Negative |
| hCoVNL63 | spike | KTIEDALRLSAHLET | 812 | 826 | Negative |
| hCoVNL63 | spike | LEAKTASLFQTTVEL | 1255 | 1269 | Negative |
| hCoVNL63 | spike | LEDLLFSKVVTSGLG | 873 | 887 | Negative |
| hCoVNL63 | spike | LGTVDVDYKSCTKGL | 886 | 900 | Negative |
| hCoVNL63 | spike | LIPVRPRNSSDNGIS | 742 | 756 | Negative |
| hCoVNL63 | spike | LLFSKVVTSGLGTVD | 876 | 890 | Negative |
| hCoVNL63 | spike | LLFTEQLGAPLGITI | 130 | 144 | Negative |

| | | | | | |
|---|---|---|---|---|---|
| hCoVNL63 | spike | LLPTDYKNVKAWSGI | 1133 | 1147 | Negative |
| hCoVNL63 | spike | LLQLPNFYYVSNGGN | 708 | 722 | Negative |
| hCoVNL63 | spike | LNFSANSLDNLKSGV | 300 | 314 | Negative |
| hCoVNL63 | spike | LPNFYYVSNGGNNCT | 711 | 725 | Negative |
| hCoVNL63 | spike | LSIADLACAQYYNGI | 900 | 914 | Negative |
| hCoVNL63 | spike | LTFDSNAFSLANVTS | 833 | 847 | Negative |
| hCoVNL63 | spike | MYTGSLIGGMVLGGL | 928 | 942 | Negative |
| hCoVNL63 | spike | NCTKYNIYDYVGTGI | 626 | 640 | Negative |
| hCoVNL63 | spike | NFYYVSNGGNNCTTA | 713 | 727 | Negative |
| hCoVNL63 | spike | NNFQKFKTICFSTVE | 558 | 572 | Negative |
| hCoVNL63 | spike | PGLKSSTGFVYFNAT | 264 | 278 | Negative |
| hCoVNL63 | spike | PIYYQHTDINFTATA | 477 | 491 | Negative |
| hCoVNL63 | spike | PNFYYVSNGGNNCTT | 712 | 726 | Negative |
| hCoVNL63 | spike | PYYCFINSTINTTHV | 348 | 362 | Negative |
| hCoVNL63 | spike | QGLIDQINSTYVDLK | 1270 | 1284 | Negative |
| hCoVNL63 | spike | QLEAKTASLFQTTVE | 1254 | 1268 | Negative |
| hCoVNL63 | spike | QLPNFYYVSNGGNNC | 710 | 724 | Negative |
| hCoVNL63 | spike | QPYYCFINSTINTTH | 347 | 361 | Negative |
| hCoVNL63 | spike | REIVVARTGQFYING | 374 | 388 | Negative |
| hCoVNL63 | spike | RSALEDLLFSKVVTS | 870 | 884 | Negative |
| hCoVNL63 | spike | RSSNQSLAGGITYVS | 642 | 656 | Negative |
| hCoVNL63 | spike | RVMFQPRLPVLSDFV | 1176 | 1190 | Negative |
| hCoVNL63 | spike | SAHLETNDVSSMLTF | 821 | 835 | Negative |
| hCoVNL63 | spike | SALEDLLFSKVVTSG | 871 | 885 | Negative |
| hCoVNL63 | spike | SANSLDNLKSGVIVF | 303 | 317 | Negative |
| hCoVNL63 | spike | SGICVDGIYGYVLRQ | 1145 | 1159 | Negative |
| hCoVNL63 | spike | SLFQTTVELQGLIDQ | 1261 | 1275 | Negative |
| hCoVNL63 | spike | SLIPVRPRNSSDNGI | 741 | 755 | Negative |
| hCoVNL63 | spike | SNGGNNCTTAVMTYS | 718 | 732 | Negative |
| hCoVNL63 | spike | SQVLNKYTEVRGSRR | 1075 | 1089 | Negative |
| hCoVNL63 | spike | SRRLAQQKINECVKS | 1087 | 1101 | Negative |
| hCoVNL63 | spike | SSDNGISAIITANLS | 750 | 764 | Negative |
| hCoVNL63 | spike | SSMLTFDSNAFSLAN | 830 | 844 | Negative |
| hCoVNL63 | spike | SSRIAGRSALEDLLF | 864 | 878 | Negative |
| hCoVNL63 | spike | SSVLPQRNIRSSRIA | 854 | 868 | Negative |
| hCoVNL63 | spike | SSVNDAITQTAEAIH | 991 | 1005 | Negative |
| hCoVNL63 | spike | STLVDGVSRLYQPLR | 242 | 256 | Negative |
| hCoVNL63 | spike | TCNSNANLSMLQLGV | 18 | 32 | Negative |
| hCoVNL63 | spike | TFDFLSNASSSFDCI | 113 | 127 | Negative |
| hCoVNL63 | spike | TGIIRSSNQSLAGGI | 638 | 652 | Negative |
| hCoVNL63 | spike | TKYNIYDYVGTGIIR | 628 | 642 | Negative |
| hCoVNL63 | spike | TLVDGVSRLYQPLRL | 243 | 257 | Negative |
| hCoVNL63 | spike | TSVQVEYLQITSTPI | 771 | 785 | Negative |
| hCoVNL63 | spike | VDVNKTLQEFAQNLP | 1215 | 1229 | Negative |
| hCoVNL63 | spike | VELQGLIDQINSTYV | 1267 | 1281 | Negative |
| hCoVNL63 | spike | VLFYCSNSSSGVLDT | 324 | 338 | Negative |
| hCoVNL63 | spike | VLRQPNLVLYSDNGV | 1156 | 1170 | Negative |
| hCoVNL63 | spike | VNLLFTEQLGAPLGI | 128 | 142 | Negative |
| hCoVNL63 | spike | VNYTVCDDCNGYTDN | 202 | 216 | Negative |

| | | | | | |
|---|---|---|---|---|---|
| hCoVNL63 | spike | VSQVLNKYTEVRGSR | 1074 | 1088 | Negative |
| hCoVNL63 | spike | WFLLTNGSTLVDGVS | 235 | 249 | Negative |
| hCoVNL63 | spike | YCFINSTINTTHVST | 350 | 364 | Negative |
| hCoVNL63 | spike | YKNVKAWSGICVDGI | 1138 | 1152 | Negative |
| hCoVNL63 | spike | YTGSLIGGMVLGGLT | 929 | 943 | Negative |
| hCoVNL63 | spike | YTVCDDCNGYTDNIF | 204 | 218 | Negative |
| hCoVNL63 | spike | AEAIHTVTIALNKIQ | 1001 | 1015 | Overlapping |
| hCoVNL63 | spike | CEHLQFGLQDGFYSA | 447 | 461 | Overlapping |
| hCoVNL63 | spike | DGSLIPVRPRNSSDN | 739 | 753 | Overlapping |
| hCoVNL63 | spike | EAIHTVTIALNKIQD | 1002 | 1016 | Overlapping |
| hCoVNL63 | spike | EAKTASLFQTTVELQ | 1256 | 1270 | Overlapping |
| hCoVNL63 | spike | EATWHYTSYTIVGAL | 582 | 596 | Overlapping |
| hCoVNL63 | spike | EAVNFNVTTASATDF | 398 | 412 | Overlapping |
| hCoVNL63 | spike | EDALRLSAHLETNDV | 815 | 829 | Overlapping |
| hCoVNL63 | spike | EDLLFSKVVTSGLGT | 874 | 888 | Overlapping |
| hCoVNL63 | spike | EFEKVHVQGSGSGSG | 1349 | 1356 | Overlapping |
| hCoVNL63 | spike | EFSNLVLNNCTKYNI | 618 | 632 | Overlapping |
| hCoVNL63 | spike | EGNSITGVPYPVSGI | 602 | 616 | Overlapping |
| hCoVNL63 | spike | EHLQFGLQDGFYSAN | 448 | 462 | Overlapping |
| hCoVNL63 | spike | EIGLNASVTLKICKF | 94 | 108 | Overlapping |
| hCoVNL63 | spike | EIVVARTGQFYINGF | 375 | 389 | Overlapping |
| hCoVNL63 | spike | EKLQCEHLQFGLQDG | 443 | 457 | Overlapping |
| hCoVNL63 | spike | ELHTVIPDYVDVNKT | 1206 | 1220 | Overlapping |
| hCoVNL63 | spike | ELKQLEAKTASLFQT | 1251 | 1265 | Overlapping |
| hCoVNL63 | spike | ELQGLIDQINSTYVD | 1268 | 1282 | Overlapping |
| hCoVNL63 | spike | ENQKILAASFNKAIN | 970 | 984 | Overlapping |
| hCoVNL63 | spike | ENYIKWPWWVWLIIS | 1290 | 1304 | Overlapping |
| hCoVNL63 | spike | EQLGAPLGITISGET | 134 | 148 | Overlapping |
| hCoVNL63 | spike | ERMAMYTGSLIGGMV | 924 | 938 | Overlapping |
| hCoVNL63 | spike | ESRYGLQNLLQLPNF | 700 | 714 | Overlapping |
| hCoVNL63 | spike | ETNDVSSMLTFDSNA | 825 | 839 | Overlapping |
| hCoVNL63 | spike | ETVRLHLYNVTRTFY | 147 | 161 | Overlapping |
| hCoVNL63 | spike | ETYVALPIYYQHTDI | 471 | 485 | Overlapping |
| hCoVNL63 | spike | EVPGSCNFPLEATWH | 572 | 586 | Overlapping |
| hCoVNL63 | spike | EVRGSRRLAQQKINE | 1083 | 1097 | Overlapping |
| hCoVNL63 | spike | EYLQITSTPIVVDCA | 776 | 790 | Overlapping |
| hCoVNL63 | spike | GAPLGITISGETVRL | 137 | 151 | Overlapping |
| hCoVNL63 | spike | GFKYFDLGFIEAVNF | 388 | 402 | Overlapping |
| hCoVNL63 | spike | GVADAERMAMYTGSL | 919 | 933 | Overlapping |
| hCoVNL63 | spike | GVPDNSSTIVTGLLP | 31 | 45 | Overlapping |
| hCoVNL63 | spike | GVPYPVSGIREFSNL | 608 | 622 | Overlapping |
| hCoVNL63 | spike | IEAVNFNVTTASATD | 397 | 411 | Overlapping |
| hCoVNL63 | spike | IEDALRLSAHLETND | 814 | 828 | Overlapping |
| hCoVNL63 | spike | LDDNVLPETYVALPI | 464 | 478 | Overlapping |
| hCoVNL63 | spike | LQLPNFYYVSNGGNN | 709 | 723 | Overlapping |
| hCoVNL63 | spike | NCTTAVMTYSNFGIC | 723 | 737 | Overlapping |
| hCoVNL63 | spike | NEIGLNASVTLKICK | 93 | 107 | Overlapping |
| hCoVNL63 | spike | NFSANSLDNLKSGVI | 301 | 315 | Overlapping |
| hCoVNL63 | spike | NGGNNCTTAVMTYSN | 719 | 733 | Overlapping |

| | | | | | |
|---|---|---|---|---|---|
| hCoVNL63 | spike | NLLFTEQLGAPLGIT | 129 | 143 | Overlapping |
| hCoVNL63 | spike | NNCTTAVMTYSNFGI | 722 | 736 | Overlapping |
| hCoVNL63 | spike | NSNANLSMLQLGVPD | 20 | 34 | Overlapping |
| hCoVNL63 | spike | NSTINTTHVSTFVGI | 354 | 368 | Overlapping |
| hCoVNL63 | spike | NWFLLTNGSTLVDGV | 234 | 248 | Overlapping |
| hCoVNL63 | spike | NYTVCDDCNGYTDNI | 203 | 217 | Overlapping |
| hCoVNL63 | spike | PETYVALPIYYQHTD | 470 | 484 | Overlapping |
| hCoVNL63 | spike | PIVVDCATYVCNGNP | 784 | 798 | Overlapping |
| hCoVNL63 | spike | PTVREIVVARTGQFY | 371 | 385 | Overlapping |
| hCoVNL63 | spike | PVLSDFVQIYNCNVT | 1184 | 1198 | Overlapping |
| hCoVNL63 | spike | PVPGLKSSTGFVYFN | 262 | 276 | Overlapping |
| hCoVNL63 | spike | PVSGIREFSNLVLNN | 612 | 626 | Overlapping |
| hCoVNL63 | spike | PYYEFEKVHVQGSGS | 1346 | 1356 | Overlapping |
| hCoVNL63 | spike | QENQKILAASFNKAI | 969 | 983 | Overlapping |
| hCoVNL63 | spike | QVEYLQITSTPIVVD | 774 | 788 | Overlapping |
| hCoVNL63 | spike | RVELHTVIPDYVDVN | 1204 | 1218 | Overlapping |
| hCoVNL63 | spike | SEGNSITGVPYPVSG | 601 | 615 | Overlapping |
| hCoVNL63 | spike | SMRGCCDCGSTKLPY | 1333 | 1347 | Overlapping |
| hCoVNL63 | spike | SNSSSGVLDTTIPFG | 329 | 343 | Overlapping |
| hCoVNL63 | spike | SQPYYCFINSTINTT | 346 | 360 | Overlapping |
| hCoVNL63 | spike | STKLPYYEFEKVHVQ | 1342 | 1356 | Overlapping |
| hCoVNL63 | spike | SVVNATVTVNVTTHN | 184 | 198 | Overlapping |
| hCoVNL63 | spike | TDINFTATASFGGSC | 483 | 497 | Overlapping |
| hCoVNL63 | spike | TGSDVNCNGYQHNSV | 278 | 292 | Overlapping |
| hCoVNL63 | spike | TNEIGLNASVTLKIC | 92 | 106 | Overlapping |
| hCoVNL63 | spike | VEVPGSCNFPLEATW | 571 | 585 | Overlapping |

**Table S6. 229E-specific Peptides**

| Lineage | Protein | HDPA peptide sequence | AA pos start | AA pos end | Group |
|---|---|---|---|---|---|
| hCoV229E | envelope | DHALIVNVLLWCVVL | 8 | 22 | Positive |
| hCoV229E | envelope | GPIKNVYHIYQSYMH | 52 | 66 | Positive |
| hCoV229E | envelope | GSGMFLKLVDDHALI | 1 | 12 | Positive |
| hCoV229E | envelope | GSGSGMFLKLVDDHA | 1 | 10 | Positive |
| hCoV229E | envelope | GSGSGSGMFLKLVDD | 1 | 8 | Positive |
| hCoV229E | envelope | HMFCNRTVYGPIKNV | 43 | 57 | Positive |
| hCoV229E | envelope | NRTVYGPIKNVYHIY | 47 | 61 | Positive |
| hCoV229E | envelope | GMFLKLVDDHALIVN | 1 | 14 | Negative |
| hCoV229E | envelope | GSGSGMFLKLVDDHA | 1 | 10 | Negative |
| hCoV229E | envelope | GSGSGSGMFLKLVDD | 1 | 8 | Negative |
| hCoV229E | envelope | DDHALIVNVLLWCVV | 7 | 21 | Overlapping |
| hCoV229E | envelope | LVDDHALIVNVLLWC | 5 | 19 | Overlapping |
| hCoV229E | membrane | ARTFWAWNPEVNAIT | 105 | 119 | Positive |
| hCoV229E | membrane | ASGVQVHNLPEYMTV | 156 | 170 | Positive |
| hCoV229E | membrane | AWNPEVNAITVTTVL | 110 | 124 | Positive |
| hCoV229E | membrane | DFSAVSSPMSNMTEN | 204 | 218 | Positive |
| hCoV229E | membrane | DGHRLASGVQVHNLP | 151 | 165 | Positive |
| hCoV229E | membrane | GHRLASGVQVHNLPE | 152 | 166 | Positive |
| hCoV229E | membrane | GHYKYSRLFYGLKML | 35 | 49 | Positive |
| hCoV229E | membrane | GSGSGSGMSNDNCTG | 1 | 8 | Positive |
| hCoV229E | membrane | GWVFYVRVKHGDFSA | 193 | 207 | Positive |
| hCoV229E | membrane | HRLASGVQVHNLPEY | 153 | 167 | Positive |
| hCoV229E | membrane | IYSRVGRSVNSQNST | 178 | 192 | Positive |
| hCoV229E | membrane | LPEYMTVAVPSTTII | 164 | 178 | Positive |
| hCoV229E | membrane | LYVDGHRLASGVQVH | 148 | 162 | Positive |
| hCoV229E | membrane | MYFANSFRLFRRART | 93 | 107 | Positive |
| hCoV229E | membrane | NAITVTTVLGQTYYQ | 116 | 130 | Positive |
| hCoV229E | membrane | NDNCTGDIVTHLKNW | 3 | 17 | Positive |
| hCoV229E | membrane | NERLLHFFGSGSGSG | 218 | 225 | Positive |
| hCoV229E | membrane | NLPEYMTVAVPSTTI | 163 | 177 | Positive |
| hCoV229E | membrane | NMTENERLLHFFGSG | 214 | 225 | Positive |
| hCoV229E | membrane | NPEVNAITVTTVLGQ | 112 | 126 | Positive |
| hCoV229E | membrane | NWDSNWAFVAFSFLM | 69 | 83 | Positive |
| hCoV229E | membrane | NWNFGWNVILTIFIV | 16 | 30 | Positive |
| hCoV229E | membrane | PMSNMTENERLLHFF | 211 | 225 | Positive |
| hCoV229E | membrane | PTGITVTLLSGVLYV | 136 | 150 | Positive |
| hCoV229E | membrane | QVHNLPEYMTVAVPS | 160 | 174 | Positive |
| hCoV229E | membrane | RRARTFWAWNPEVNA | 103 | 117 | Positive |
| hCoV229E | membrane | SGVQVHNLPEYMTVA | 157 | 171 | Positive |
| hCoV229E | membrane | SNMTENERLLHFFGS | 213 | 225 | Positive |
| hCoV229E | membrane | TFWAWNPEVNAITVT | 107 | 121 | Positive |
| hCoV229E | membrane | TGDIVTHLKNWNFGW | 7 | 21 | Positive |
| hCoV229E | membrane | TGWVFYVRVKHGDFS | 192 | 206 | Positive |
| hCoV229E | membrane | VHNLPEYMTVAVPST | 161 | 175 | Positive |
| hCoV229E | membrane | VMYFANSFRLFRRAR | 92 | 106 | Positive |
| hCoV229E | membrane | WAWNPEVNAITVTTV | 109 | 123 | Positive |
| hCoV229E | membrane | WNPEVNAITVTTVLG | 111 | 125 | Positive |

| | | | | | |
|---|---|---|---|---|---|
| hCoV229E | membrane | WVMYFANSFRLFRRA | 91 | 105 | Positive |
| hCoV229E | membrane | YSRVGRSVNSQNSTG | 179 | 193 | Positive |
| hCoV229E | membrane | YVDGHRLASGVQVHN | 149 | 163 | Positive |
| hCoV229E | membrane | AVSSPMSNMTENERL | 207 | 221 | Negative |
| hCoV229E | membrane | FDTWANWDSNWAFVA | 64 | 78 | Negative |
| hCoV229E | membrane | FSAVSSPMSNMTENE | 205 | 219 | Negative |
| hCoV229E | membrane | GDIVTHLKNWNFGWN | 8 | 22 | Negative |
| hCoV229E | membrane | GITVTLLSGVLYVDG | 138 | 152 | Negative |
| hCoV229E | membrane | GQTYYQPIQQAPTGI | 125 | 139 | Negative |
| hCoV229E | membrane | GSGMSNDNCTGDIVT | 1 | 12 | Negative |
| hCoV229E | membrane | GSGSGMSNDNCTGDI | 1 | 10 | Negative |
| hCoV229E | membrane | GSGSGSGMSNDNCTG | 1 | 8 | Negative |
| hCoV229E | membrane | GVLYVDGHRLASGVQ | 146 | 160 | Negative |
| hCoV229E | membrane | GVQVHNLPEYMTVAV | 158 | 172 | Negative |
| hCoV229E | membrane | SAVSSPMSNMTENER | 206 | 220 | Negative |
| hCoV229E | membrane | SGVLYVDGHRLASGV | 145 | 159 | Negative |
| hCoV229E | membrane | SSPMSNMTENERLLH | 209 | 223 | Negative |
| hCoV229E | membrane | VSSPMSNMTENERLL | 208 | 222 | Negative |
| hCoV229E | membrane | DFSAVSSPMSNMTEN | 204 | 218 | Overlapping |
| hCoV229E | membrane | ENERLLHFFGSGSGS | 217 | 225 | Overlapping |
| hCoV229E | membrane | EVNAITVTTVLGQTY | 114 | 128 | Overlapping |
| hCoV229E | membrane | EYMTVAVPSTTIIYS | 166 | 180 | Overlapping |
| hCoV229E | membrane | FWAWNPEVNAITVTT | 108 | 122 | Overlapping |
| hCoV229E | membrane | GMSNDNCTGDIVTHL | 1 | 14 | Overlapping |
| hCoV229E | membrane | GSGMSNDNCTGDIVT | 1 | 12 | Overlapping |
| hCoV229E | membrane | GSGSGMSNDNCTGDI | 1 | 10 | Overlapping |
| hCoV229E | membrane | PEVNAITVTTVLGQT | 113 | 127 | Overlapping |
| hCoV229E | membrane | PEYMTVAVPSTTIIY | 165 | 179 | Overlapping |
| hCoV229E | membrane | RTFWAWNPEVNAITV | 106 | 120 | Overlapping |
| hCoV229E | membrane | TENERLLHFFGSGSG | 216 | 225 | Overlapping |
| hCoV229E | nucleocapsid | AAALKSLGFDKPQEK | 187 | 201 | Positive |
| hCoV229E | nucleocapsid | AELVPSTAAMLFDSH | 293 | 307 | Positive |
| hCoV229E | nucleocapsid | ALKSLGFDKPQEKDK | 189 | 203 | Positive |
| hCoV229E | nucleocapsid | APSRSQSRSQSRSRV | 147 | 160 | Positive |
| hCoV229E | nucleocapsid | DAKFRERVEGVVWVA | 90 | 104 | Positive |
| hCoV229E | nucleocapsid | DHNFGSAGVVANGVK | 271 | 285 | Positive |
| hCoV229E | nucleocapsid | DHPHLGKFLEELNAF | 330 | 344 | Positive |
| hCoV229E | nucleocapsid | DKPQEKDKKSAKTGT | 196 | 210 | Positive |
| hCoV229E | nucleocapsid | DSHIVSKESGNTVVL | 305 | 319 | Positive |
| hCoV229E | nucleocapsid | DSRAPSRSQSRSQSR | 144 | 158 | Positive |
| hCoV229E | nucleocapsid | EPVRDEVSIETDIID | 372 | 386 | Positive |
| hCoV229E | nucleocapsid | ETDIIDEVNGSGSGS | 381 | 389 | Positive |
| hCoV229E | nucleocapsid | EVSIETDIIDEVNGS | 377 | 389 | Positive |
| hCoV229E | nucleocapsid | FDKPQEKDKKSAKTG | 195 | 209 | Positive |
| hCoV229E | nucleocapsid | FDSHIVSKESGNTVV | 304 | 318 | Positive |
| hCoV229E | nucleocapsid | FLEELNAFTREMQQQ | 337 | 351 | Positive |
| hCoV229E | nucleocapsid | GSGMATVKWADASEP | 1 | 12 | Positive |
| hCoV229E | nucleocapsid | GSGSGMATVKWADAS | 1 | 10 | Positive |
| hCoV229E | nucleocapsid | GSGSGSGMATVKWAD | 1 | 8 | Positive |

| | | | | | |
|---|---|---|---|---|---|
| hCoV229E | nucleocapsid | GTPKPSRNQSPASSQ | 209 | 223 | Positive |
| hCoV229E | nucleocapsid | GYPQFAELVPSTAAM | 288 | 302 | Positive |
| hCoV229E | nucleocapsid | GYWNVQKRFRTRKGK | 56 | 70 | Positive |
| hCoV229E | nucleocapsid | HKDAKFRERVEGVVW | 88 | 102 | Positive |
| hCoV229E | nucleocapsid | IPHFNQKLPNGVTVA | 126 | 139 | Positive |
| hCoV229E | nucleocapsid | IPRNLVPVNKKDKNK | 39 | 53 | Positive |
| hCoV229E | nucleocapsid | IVSKESGNTVVLTFT | 308 | 322 | Positive |
| hCoV229E | nucleocapsid | KDHPHLGKFLEELNA | 329 | 343 | Positive |
| hCoV229E | nucleocapsid | KHEMQKPRWKRQPND | 241 | 255 | Positive |
| hCoV229E | nucleocapsid | KKDKNKLIGYWNVQK | 48 | 62 | Positive |
| hCoV229E | nucleocapsid | KLIGYWNVQKRFRTR | 53 | 67 | Positive |
| hCoV229E | nucleocapsid | KLPNGVTVAEEPDSR | 132 | 146 | Positive |
| hCoV229E | nucleocapsid | KNKLIGYWNVQKRFR | 51 | 65 | Positive |
| hCoV229E | nucleocapsid | KPRWKRQPNDDVTSN | 246 | 260 | Positive |
| hCoV229E | nucleocapsid | KSLGFDKPQEKDKKS | 191 | 205 | Positive |
| hCoV229E | nucleocapsid | KSQSRNPSSDRNHNS | 164 | 178 | Positive |
| hCoV229E | nucleocapsid | KVIPRNLVPVNKKDK | 37 | 51 | Positive |
| hCoV229E | nucleocapsid | LARSQSSETKEQKHE | 229 | 243 | Positive |
| hCoV229E | nucleocapsid | LEFNPSQTSPATVEP | 359 | 373 | Positive |
| hCoV229E | nucleocapsid | LGFDKPQEKDKKSAK | 193 | 207 | Positive |
| hCoV229E | nucleocapsid | LKSLGFDKPQEKDKK | 190 | 204 | Positive |
| hCoV229E | nucleocapsid | LVDSEQPWKVIPRNL | 29 | 43 | Positive |
| hCoV229E | nucleocapsid | NAFTREMQQQPLLNP | 342 | 356 | Positive |
| hCoV229E | nucleocapsid | NDDVTSNVTQCFGPR | 254 | 268 | Positive |
| hCoV229E | nucleocapsid | NHNSQDDIMKAVAAA | 175 | 189 | Positive |
| hCoV229E | nucleocapsid | NKKDKNKLIGYWNVQ | 47 | 61 | Positive |
| hCoV229E | nucleocapsid | NKLIGYWNVQKRFRT | 52 | 66 | Positive |
| hCoV229E | nucleocapsid | NLVPVNKKDKNKLIG | 42 | 56 | Positive |
| hCoV229E | nucleocapsid | NPSSDRNHNSQDDIM | 169 | 183 | Positive |
| hCoV229E | nucleocapsid | NSEPEIPHFNQKLPN | 121 | 135 | Positive |
| hCoV229E | nucleocapsid | NSQDDIMKAVAAALK | 177 | 191 | Positive |
| hCoV229E | nucleocapsid | PATVEPVRDEVSIET | 368 | 382 | Positive |
| hCoV229E | nucleocapsid | PHKDAKFRERVEGVV | 87 | 101 | Positive |
| hCoV229E | nucleocapsid | PKDHPHLGKFLEELN | 328 | 342 | Positive |
| hCoV229E | nucleocapsid | PLLNPSALEFNPSQT | 352 | 366 | Positive |
| hCoV229E | nucleocapsid | PLLVDSEQPWKVIPR | 27 | 41 | Positive |
| hCoV229E | nucleocapsid | PNGVTVAEEPDSRAP | 134 | 148 | Positive |
| hCoV229E | nucleocapsid | PRWKRQPNDDVTSNV | 247 | 261 | Positive |
| hCoV229E | nucleocapsid | PSALEFNPSQTSPAT | 356 | 370 | Positive |
| hCoV229E | nucleocapsid | PSSDRNHNSQDDIMK | 170 | 184 | Positive |
| hCoV229E | nucleocapsid | PTGYGVRRKNSEPEI | 112 | 126 | Positive |
| hCoV229E | nucleocapsid | PVRDEVSIETDIIDE | 373 | 387 | Positive |
| hCoV229E | nucleocapsid | RAPSRSQSRSQSRSR | 146 | 160 | Positive |
| hCoV229E | nucleocapsid | RDEVSIETDIIDEVN | 375 | 389 | Positive |
| hCoV229E | nucleocapsid | RFRTRKGKRVDLSPK | 63 | 77 | Positive |
| hCoV229E | nucleocapsid | RNHNSQDDIMKAVAA | 174 | 188 | Positive |
| hCoV229E | nucleocapsid | RNPSSDRNHNSQDDI | 168 | 182 | Positive |
| hCoV229E | nucleocapsid | RTRKGKRVDLSPKLH | 65 | 79 | Positive |
| hCoV229E | nucleocapsid | RWKRQPNDDVTSNVT | 248 | 262 | Positive |

| | | | | | |
|---|---|---|---|---|---|
| hCoV229E | nucleocapsid | SAKTGTPKPSRNQSP | 205 | 219 | Positive |
| hCoV229E | nucleocapsid | SDRNHNSQDDIMKAV | 172 | 186 | Positive |
| hCoV229E | nucleocapsid | SEQPWKVIPRNLVPV | 32 | 46 | Positive |
| hCoV229E | nucleocapsid | SIETDIIDEVNGSGS | 379 | 389 | Positive |
| hCoV229E | nucleocapsid | SKSQSRNPSSDRNHN | 163 | 177 | Positive |
| hCoV229E | nucleocapsid | SLGFDKPQEKDKKSA | 192 | 206 | Positive |
| hCoV229E | nucleocapsid | SNVTQCFGPRDLDHN | 259 | 273 | Positive |
| hCoV229E | nucleocapsid | SPLLVDSEQPWKVIP | 26 | 40 | Positive |
| hCoV229E | nucleocapsid | SQTSPATVEPVRDEV | 364 | 378 | Positive |
| hCoV229E | nucleocapsid | SRNPSSDRNHNSQDD | 167 | 181 | Positive |
| hCoV229E | nucleocapsid | SRSQSRSQSRSRVES | 149 | 163 | Positive |
| hCoV229E | nucleocapsid | SSDRNHNSQDDIMKA | 171 | 185 | Positive |
| hCoV229E | nucleocapsid | TDIIDEVNGSGSGSG | 382 | 389 | Positive |
| hCoV229E | nucleocapsid | TEPTGYGVRRKNSEP | 110 | 124 | Positive |
| hCoV229E | nucleocapsid | TGPHKDAKFRERVEG | 85 | 99 | Positive |
| hCoV229E | nucleocapsid | TVAEEPDSRAPSRSQ | 138 | 152 | Positive |
| hCoV229E | nucleocapsid | TVEPVRDEVSIETDI | 370 | 384 | Positive |
| hCoV229E | nucleocapsid | TVPKDHPHLGKFLEE | 326 | 340 | Positive |
| hCoV229E | nucleocapsid | VAEEPDSRAPSRSQS | 139 | 153 | Positive |
| hCoV229E | nucleocapsid | VDLSPKLHFYYLGTG | 72 | 86 | Positive |
| hCoV229E | nucleocapsid | VEPVRDEVSIETDII | 371 | 385 | Positive |
| hCoV229E | nucleocapsid | VESKSQSRNPSSDRN | 162 | 175 | Positive |
| hCoV229E | nucleocapsid | VRRKNSEPEIPHFNQ | 117 | 131 | Positive |
| hCoV229E | nucleocapsid | VTVAEEPDSRAPSRS | 137 | 151 | Positive |
| hCoV229E | nucleocapsid | WKRQPNDDVTSNVTQ | 249 | 263 | Positive |
| hCoV229E | nucleocapsid | ADASEPQRGRQGRIP | 7 | 21 | Negative |
| hCoV229E | nucleocapsid | AVDGAKTEPTGYGVR | 104 | 118 | Negative |
| hCoV229E | nucleocapsid | DDIMKAVAAALKSLG | 180 | 194 | Negative |
| hCoV229E | nucleocapsid | EGVVWVAVDGAKTEP | 98 | 112 | Negative |
| hCoV229E | nucleocapsid | ELNAFTREMQQQPLL | 340 | 354 | Negative |
| hCoV229E | nucleocapsid | ELVPSTAAMLFDSHI | 294 | 308 | Negative |
| hCoV229E | nucleocapsid | FGPRDLDHNFGSAGV | 265 | 279 | Negative |
| hCoV229E | nucleocapsid | FRERVEGVVWVAVDG | 93 | 107 | Negative |
| hCoV229E | nucleocapsid | FTREMQQQPLLNPSA | 344 | 358 | Negative |
| hCoV229E | nucleocapsid | GKFLEELNAFTREMQ | 335 | 349 | Negative |
| hCoV229E | nucleocapsid | GKRVDLSPKLHFYYL | 69 | 83 | Negative |
| hCoV229E | nucleocapsid | GMATVKWADASEPQR | 1 | 14 | Negative |
| hCoV229E | nucleocapsid | GSGSGMATVKWADAS | 1 | 10 | Negative |
| hCoV229E | nucleocapsid | GSGSGSGMATVKWAD | 1 | 8 | Negative |
| hCoV229E | nucleocapsid | GTGPHKDAKFRERVE | 84 | 98 | Negative |
| hCoV229E | nucleocapsid | GVKAKGYPQFAELVP | 283 | 297 | Negative |
| hCoV229E | nucleocapsid | GVRRKNSEPEIPHFN | 116 | 130 | Negative |
| hCoV229E | nucleocapsid | GVTVAEEPDSRAPSR | 136 | 150 | Negative |
| hCoV229E | nucleocapsid | GVVANGVKAKGYPQF | 278 | 292 | Negative |
| hCoV229E | nucleocapsid | GVVWVAVDGAKTEPT | 99 | 113 | Negative |
| hCoV229E | nucleocapsid | KFLEELNAFTREMQQ | 336 | 350 | Negative |
| hCoV229E | nucleocapsid | KWADASEPQRGRQGR | 5 | 19 | Negative |
| hCoV229E | nucleocapsid | LDHNFGSAGVVANGV | 270 | 284 | Negative |
| hCoV229E | nucleocapsid | LPNGVTVAEEPDSRA | 133 | 147 | Negative |

| | | | | | |
|---|---|---|---|---|---|
| hCoV229E | nucleocapsid | NGVKAKGYPQFAELV | 282 | 296 | Negative |
| hCoV229E | nucleocapsid | NGVTVAEEPDSRAPS | 135 | 149 | Negative |
| hCoV229E | nucleocapsid | QKLPNGVTVAEEPDS | 131 | 145 | Negative |
| hCoV229E | nucleocapsid | QSRSQSRSRVESKSQ | 152 | 166 | Negative |
| hCoV229E | nucleocapsid | QSRSRVESKSQSRNP | 156 | 170 | Negative |
| hCoV229E | nucleocapsid | RNQSPASSQSAAKIL | 215 | 229 | Negative |
| hCoV229E | nucleocapsid | RSRVESKSQSRNPSS | 158 | 172 | Negative |
| hCoV229E | nucleocapsid | RVEGVVWVAVDGAKT | 96 | 110 | Negative |
| hCoV229E | nucleocapsid | RVESKSQSRNPSSDR | 160 | 174 | Negative |
| hCoV229E | nucleocapsid | SKESGNTVVLTFTTR | 310 | 324 | Negative |
| hCoV229E | nucleocapsid | SPATVEPVRDEVSIE | 367 | 381 | Negative |
| hCoV229E | nucleocapsid | SQSRSQSRSRVESKS | 151 | 164 | Negative |
| hCoV229E | nucleocapsid | SQSRSRVESKSQSRN | 155 | 169 | Negative |
| hCoV229E | nucleocapsid | SRSRVESKSQSRNPS | 157 | 171 | Negative |
| hCoV229E | nucleocapsid | TDIIDEVNGSGSGSG | 382 | 389 | Negative |
| hCoV229E | nucleocapsid | TSPATVEPVRDEVSI | 366 | 380 | Negative |
| hCoV229E | nucleocapsid | VRDEVSIETDIIDEV | 374 | 388 | Negative |
| hCoV229E | nucleocapsid | YLGTGPHKDAKFRER | 82 | 96 | Negative |
| hCoV229E | nucleocapsid | AEEPDSRAPSRSQSR | 140 | 154 | Overlapping |
| hCoV229E | nucleocapsid | AFTREMQQQPLLNPS | 343 | 357 | Overlapping |
| hCoV229E | nucleocapsid | DEVSIETDIIDEVNG | 376 | 389 | Overlapping |
| hCoV229E | nucleocapsid | DSRAPSRSQSRSQSR | 144 | 158 | Overlapping |
| hCoV229E | nucleocapsid | EELNAFTREMQQQPL | 339 | 353 | Overlapping |
| hCoV229E | nucleocapsid | EEPDSRAPSRSQSRS | 141 | 155 | Overlapping |
| hCoV229E | nucleocapsid | EFNPSQTSPATVEPV | 360 | 374 | Overlapping |
| hCoV229E | nucleocapsid | EGVVWVAVDGAKTEP | 98 | 112 | Overlapping |
| hCoV229E | nucleocapsid | EIPHFNQKLPNGVTV | 125 | 139 | Overlapping |
| hCoV229E | nucleocapsid | EKDKKSAKTGTPKPS | 200 | 214 | Overlapping |
| hCoV229E | nucleocapsid | ELNAFTREMQQQPLL | 340 | 354 | Overlapping |
| hCoV229E | nucleocapsid | ELVPSTAAMLFDSHI | 294 | 308 | Overlapping |
| hCoV229E | nucleocapsid | EMQKPRWKRQPNDDV | 243 | 257 | Overlapping |
| hCoV229E | nucleocapsid | EMQQQPLLNPSALEF | 347 | 361 | Overlapping |
| hCoV229E | nucleocapsid | EPDSRAPSRSQSRSQ | 142 | 156 | Overlapping |
| hCoV229E | nucleocapsid | EPEIPHFNQKLPNGV | 123 | 137 | Overlapping |
| hCoV229E | nucleocapsid | EPQRGRQGRIPYSLY | 11 | 25 | Overlapping |
| hCoV229E | nucleocapsid | EPTGYGVRRKNSEPE | 111 | 125 | Overlapping |
| hCoV229E | nucleocapsid | EPVRDEVSIETDIID | 372 | 386 | Overlapping |
| hCoV229E | nucleocapsid | EQKHEMQKPRWKRQP | 239 | 253 | Overlapping |
| hCoV229E | nucleocapsid | EQPWKVIPRNLVPVN | 33 | 47 | Overlapping |
| hCoV229E | nucleocapsid | ERVEGVVWVAVDGAK | 95 | 109 | Overlapping |
| hCoV229E | nucleocapsid | ESGNTVVLTFTTRVT | 312 | 326 | Overlapping |
| hCoV229E | nucleocapsid | ESKSQSRNPSSDRNH | 162 | 176 | Overlapping |
| hCoV229E | nucleocapsid | ETDIIDEVNGSGSGS | 381 | 389 | Overlapping |
| hCoV229E | nucleocapsid | ETKEQKHEMQKPRWK | 236 | 250 | Overlapping |
| hCoV229E | nucleocapsid | EVSIETDIIDEVNGS | 377 | 389 | Overlapping |
| hCoV229E | nucleocapsid | FTREMQQQPLLNPSA | 344 | 358 | Overlapping |
| hCoV229E | nucleocapsid | GFDKPQEKDKKSAKT | 194 | 208 | Overlapping |
| hCoV229E | nucleocapsid | GPHKDAKFRERVEGV | 86 | 100 | Overlapping |
| hCoV229E | nucleocapsid | GPRDLDHNFGSAGVV | 266 | 280 | Overlapping |

| | | | | | |
|---|---|---|---|---|---|
| hCoV229E | nucleocapsid | HEMQKPRWKRQPNDD | 242 | 256 | Overlapping |
| hCoV229E | nucleocapsid | HFNQKLPNGVTVAEE | 128 | 142 | Overlapping |
| hCoV229E | nucleocapsid | HLGKFLEELNAFTRE | 333 | 347 | Overlapping |
| hCoV229E | nucleocapsid | IETDIIDEVNGSGSG | 380 | 389 | Overlapping |
| hCoV229E | nucleocapsid | KEQKHEMQKPRWKRQ | 238 | 252 | Overlapping |
| hCoV229E | nucleocapsid | LEELNAFTREMQQQP | 338 | 352 | Overlapping |
| hCoV229E | nucleocapsid | MQQQPLLNPSALEFN | 348 | 362 | Overlapping |
| hCoV229E | nucleocapsid | NQKLPNGVTVAEEPD | 130 | 144 | Overlapping |
| hCoV229E | nucleocapsid | PATVEPVRDEVSIET | 368 | 382 | Overlapping |
| hCoV229E | nucleocapsid | PVRDEVSIETDIIDE | 373 | 387 | Overlapping |
| hCoV229E | nucleocapsid | PYSLYSPLLVDSEQP | 21 | 35 | Overlapping |
| hCoV229E | nucleocapsid | QEKDKKSAKTGTPKP | 199 | 213 | Overlapping |
| hCoV229E | nucleocapsid | QSRNPSSDRNHNSQD | 166 | 180 | Overlapping |
| hCoV229E | nucleocapsid | RDLDHNFGSAGVVAN | 268 | 282 | Overlapping |
| hCoV229E | nucleocapsid | REMQQQPLLNPSALE | 346 | 360 | Overlapping |
| hCoV229E | nucleocapsid | RVDLSPKLHFYYLGT | 71 | 85 | Overlapping |
| hCoV229E | nucleocapsid | SEPEIPHFNQKLPNG | 122 | 136 | Overlapping |
| hCoV229E | nucleocapsid | SETKEQKHEMQKPRW | 235 | 249 | Overlapping |
| hCoV229E | nucleocapsid | SRAPSRSQSRSQSRS | 145 | 158 | Overlapping |
| hCoV229E | nucleocapsid | TREMQQQPLLNPSAL | 345 | 359 | Overlapping |
| hCoV229E | nucleocapsid | VAVDGAKTEPTGYGV | 103 | 117 | Overlapping |
| hCoV229E | nucleocapsid | VEGVVWVAVDGAKTE | 97 | 111 | Overlapping |
| hCoV229E | nucleocapsid | VRDEVSIETDIIDEV | 374 | 388 | Overlapping |
| hCoV229E | orf1ab | AEAEWIELFPHNDRI | 810 | 824 | Positive |
| hCoV229E | orf1ab | AEDTLNKLSKYLVNE | 1755 | 1769 | Positive |
| hCoV229E | orf1ab | AGLYYESISYEEQDA | 4575 | 4589 | Positive |
| hCoV229E | orf1ab | AGVCFGFDKWFVNDG | 2705 | 2719 | Positive |
| hCoV229E | orf1ab | AHGGGLAKALDVYTK | 1309 | 1323 | Positive |
| hCoV229E | orf1ab | AHPTMDGFCQYKGKW | 4015 | 4029 | Positive |
| hCoV229E | orf1ab | AIDGENVSKEGFFTY | 6595 | 6609 | Positive |
| hCoV229E | orf1ab | AIKLYNPKAIHDIGN | 5911 | 5925 | Positive |
| hCoV229E | orf1ab | AISGDKVFVRDGNTD | 6135 | 6149 | Positive |
| hCoV229E | orf1ab | AKDEISVTPIKNTVD | 1245 | 1259 | Positive |
| hCoV229E | orf1ab | AKEESVKSSYAYATL | 5135 | 5149 | Positive |
| hCoV229E | orf1ab | AKERCTHWCTYLFSL | 3393 | 3407 | Positive |
| hCoV229E | orf1ab | AKESLRLFAAETVKA | 5121 | 5135 | Positive |
| hCoV229E | orf1ab | AKHDFFTWHEGRTIY | 4159 | 4173 | Positive |
| hCoV229E | orf1ab | AKPEEKLSAYDLACC | 2397 | 2411 | Positive |
| hCoV229E | orf1ab | ALGCDYVYNPYVIDI | 5821 | 5835 | Positive |
| hCoV229E | orf1ab | ANENAINKGGRTVQS | 5891 | 5905 | Positive |
| hCoV229E | orf1ab | APLKCVRNVFENFED | 2541 | 2555 | Positive |
| hCoV229E | orf1ab | AQGPFIDGNIIHANY | 6673 | 6687 | Positive |
| hCoV229E | orf1ab | ATSAARLVVVVPDHD | 3751 | 3765 | Positive |
| hCoV229E | orf1ab | AVAKHDFFTWHEGRT | 4157 | 4171 | Positive |
| hCoV229E | orf1ab | AVETCTIAGKAFDKV | 476 | 490 | Positive |
| hCoV229E | orf1ab | AYETARQEYENAVAN | 3643 | 3657 | Positive |
| hCoV229E | orf1ab | CDEMVLKGVVGVLTL | 4255 | 4269 | Positive |
| hCoV229E | orf1ab | CEVVHDQVNYVPLRA | 6037 | 6051 | Positive |
| hCoV229E | orf1ab | CFVASSTFVEEEHVN | 326 | 340 | Positive |

| | | | | | |
|---|---|---|---|---|---|
| hCoV229E | orf1ab | CGSPGYNLKNGEVEF | 3109 | 3123 | Positive |
| hCoV229E | orf1ab | CKVCGCWLNHGCTCD | 4049 | 4063 | Positive |
| hCoV229E | orf1ab | CNAVAKHDFFTWHEG | 4155 | 4169 | Positive |
| hCoV229E | orf1ab | CRQYSNESWFEDDYK | 668 | 682 | Positive |
| hCoV229E | orf1ab | CVDCDSYGYGSTFIT | 2201 | 2215 | Positive |
| hCoV229E | orf1ab | CVDCHDEMCILHCSN | 4361 | 4375 | Positive |
| hCoV229E | orf1ab | CVPHNMRVLHLGAGS | 6517 | 6531 | Positive |
| hCoV229E | orf1ab | CVRFKNVDKDDAFYI | 4119 | 4133 | Positive |
| hCoV229E | orf1ab | CVRNVFENFEDWHYA | 2545 | 2559 | Positive |
| hCoV229E | orf1ab | CYVNLPTYYIYDEEG | 962 | 976 | Positive |
| hCoV229E | orf1ab | DADFSVTGDCATVYL | 6563 | 6577 | Positive |
| hCoV229E | orf1ab | DAIVVDNDVVDYVSD | 6549 | 6563 | Positive |
| hCoV229E | orf1ab | DALDKLYKVFGSPVM | 210 | 224 | Positive |
| hCoV229E | orf1ab | DALFLSFKLMGIGGP | 3523 | 3537 | Positive |
| hCoV229E | orf1ab | DALKFFEVTMTDLQS | 5579 | 5593 | Positive |
| hCoV229E | orf1ab | DCDSYGYGSTFITPE | 2203 | 2217 | Positive |
| hCoV229E | orf1ab | DCHDEMCILHCSNFN | 4363 | 4377 | Positive |
| hCoV229E | orf1ab | DEHVFYKKDGVYYPS | 868 | 882 | Positive |
| hCoV229E | orf1ab | DFLPRSTMEEDFLNM | 6313 | 6327 | Positive |
| hCoV229E | orf1ab | DFLVHNFVIFFTWLL | 1927 | 1941 | Positive |
| hCoV229E | orf1ab | DGELPVAISGDKVFV | 6129 | 6143 | Positive |
| hCoV229E | orf1ab | DGENVSKEGFFTYIN | 6597 | 6611 | Positive |
| hCoV229E | orf1ab | DGFVHYAGVVWTLQE | 3773 | 3787 | Positive |
| hCoV229E | orf1ab | DGNIIHANYVFWRNS | 6679 | 6693 | Positive |
| hCoV229E | orf1ab | DGVKTVLLFYMLLGF | 3461 | 3475 | Positive |
| hCoV229E | orf1ab | DGVPVVATAGYHFKQ | 4399 | 4413 | Positive |
| hCoV229E | orf1ab | DGVYYPSNGTNILPV | 876 | 890 | Positive |
| hCoV229E | orf1ab | DHDSFVKMMVDGFVH | 3763 | 3777 | Positive |
| hCoV229E | orf1ab | DHEQSMYNLLKGCNA | 4143 | 4157 | Positive |
| hCoV229E | orf1ab | DHFGENEEIVINGRT | 138 | 152 | Positive |
| hCoV229E | orf1ab | DHKPHLSFPLCSAGN | 5069 | 5083 | Positive |
| hCoV229E | orf1ab | DHNDILYTPPTVSYG | 2947 | 2961 | Positive |
| hCoV229E | orf1ab | DHRFILAITPYVCNT | 5033 | 5047 | Positive |
| hCoV229E | orf1ab | DIEPVYRVKLCFEFE | 910 | 924 | Positive |
| hCoV229E | orf1ab | DIRDYKLANDAKESL | 5111 | 5125 | Positive |
| hCoV229E | orf1ab | DLSIGPHEFCSQHTM | 4867 | 4881 | Positive |
| hCoV229E | orf1ab | DMYDGRTKAIDGENV | 6587 | 6601 | Positive |
| hCoV229E | orf1ab | DNAVATFYPQLQSAE | 6447 | 6461 | Positive |
| hCoV229E | orf1ab | DQVEEVNSIFDIETV | 1011 | 1025 | Positive |
| hCoV229E | orf1ab | DSFVKMMVDGFVHYA | 3765 | 3779 | Positive |
| hCoV229E | orf1ab | DSILQSVASSFVGMP | 3625 | 3639 | Positive |
| hCoV229E | orf1ab | DSLCAKAVTAYSVDK | 5311 | 5325 | Positive |
| hCoV229E | orf1ab | DSYFENDSILQSVAS | 3619 | 3633 | Positive |
| hCoV229E | orf1ab | DTRTRSTLNLEGVNG | 5987 | 6001 | Positive |
| hCoV229E | orf1ab | DYVYNPYVIDIQQWG | 5825 | 5839 | Positive |
| hCoV229E | orf1ab | EDDYKAFISVLDITD | 678 | 692 | Positive |
| hCoV229E | orf1ab | EFDRESSVQKKINRM | 3677 | 3691 | Positive |
| hCoV229E | orf1ab | EHDSGVVTVELEPPC | 3879 | 3893 | Positive |
| hCoV229E | orf1ab | EHESKFWDESFYASM | 4973 | 4987 | Positive |

| | | | | | |
|---|---|---|---|---|---|
| hCoV229E | orf1ab | EPCNGTDIDYCVRAF | 4087 | 4101 | Positive |
| hCoV229E | orf1ab | ETVNGYAKSNFVKDD | 2023 | 2037 | Positive |
| hCoV229E | orf1ab | EWPLFESDKEEDVQQ | 988 | 999 | Positive |
| hCoV229E | orf1ab | FAEDVCTCYDNSIQG | 6217 | 6231 | Positive |
| hCoV229E | orf1ab | FATVDHSAFAYESAV | 1673 | 1687 | Positive |
| hCoV229E | orf1ab | FCVRFGPFEFCSETV | 2011 | 2025 | Positive |
| hCoV229E | orf1ab | FDITESKYSCKEVFK | 2263 | 2277 | Positive |
| hCoV229E | orf1ab | FEDQPNLQVESANQM | 3149 | 3163 | Positive |
| hCoV229E | orf1ab | FEDWHYAKFGFTPLN | 2553 | 2567 | Positive |
| hCoV229E | orf1ab | FEKVDYGSDTVTYKS | 5195 | 5209 | Positive |
| hCoV229E | orf1ab | FFEVTMTDLQSESSC | 5583 | 5597 | Positive |
| hCoV229E | orf1ab | FFYYDDGSCEVVHDQ | 6029 | 6043 | Positive |
| hCoV229E | orf1ab | FKLMGIGGPRTIKVS | 3529 | 3543 | Positive |
| hCoV229E | orf1ab | FLNAFDVFVTTIQTV | 458 | 472 | Positive |
| hCoV229E | orf1ab | FLPVEYVSYFDGVKT | 3451 | 3465 | Positive |
| hCoV229E | orf1ab | FMPYFMHDVVSSFEG | 2511 | 2525 | Positive |
| hCoV229E | orf1ab | FNSTTLCVPHNMRVL | 6511 | 6525 | Positive |
| hCoV229E | orf1ab | FPHNDRIKSFSTFES | 818 | 832 | Positive |
| hCoV229E | orf1ab | FQLDYSIDVIDNEII | 630 | 644 | Positive |
| hCoV229E | orf1ab | FRRMFGDLSVGVCTV | 2775 | 2789 | Positive |
| hCoV229E | orf1ab | FVDHFGENEEIVING | 136 | 150 | Positive |
| hCoV229E | orf1ab | FVMFWAELFVYTTTI | 3285 | 3299 | Positive |
| hCoV229E | orf1ab | FVNDGRVGNGYVCGT | 2715 | 2729 | Positive |
| hCoV229E | orf1ab | FVNDKITEFQLDYSI | 622 | 636 | Positive |
| hCoV229E | orf1ab | GAGDAGHSLTWLWLL | 2483 | 2497 | Positive |
| hCoV229E | orf1ab | GAVPSDLNTSELLTK | 1641 | 1655 | Positive |
| hCoV229E | orf1ab | GCWLNHGCTCDRTAI | 4053 | 4067 | Positive |
| hCoV229E | orf1ab | GDLAVQIGNNNVCTY | 5629 | 5643 | Positive |
| hCoV229E | orf1ab | GFRACRFVSLDLQDC | 38 | 52 | Positive |
| hCoV229E | orf1ab | GGFEDQPNLQVESAN | 3147 | 3161 | Positive |
| hCoV229E | orf1ab | GGLNYYCVDHKPHLS | 5061 | 5075 | Positive |
| hCoV229E | orf1ab | GGPRTIKVSTVQSKL | 3535 | 3549 | Positive |
| hCoV229E | orf1ab | GGVSLLATMTTRQFH | 4621 | 4635 | Positive |
| hCoV229E | orf1ab | GKAGLYYESISYEEQ | 4573 | 4587 | Positive |
| hCoV229E | orf1ab | GKERARTVGGVSLLA | 4613 | 4627 | Positive |
| hCoV229E | orf1ab | GKQRITTIQGPPGSG | 5269 | 5283 | Positive |
| hCoV229E | orf1ab | GLGDLVDSYFENDSI | 3613 | 3627 | Positive |
| hCoV229E | orf1ab | GNAIATVKSEDGNIK | 6269 | 6283 | Positive |
| hCoV229E | orf1ab | GNITKTNVQPTGPAY | 2223 | 2237 | Positive |
| hCoV229E | orf1ab | GNVVFYQGDVDTVVN | 1280 | 1293 | Positive |
| hCoV229E | orf1ab | GPFIDGNIIHANYVF | 6675 | 6689 | Positive |
| hCoV229E | orf1ab | GPHEFCSQHTMQIVD | 4871 | 4885 | Positive |
| hCoV229E | orf1ab | GQAITSTIDSNTTQD | 3985 | 3999 | Positive |
| hCoV229E | orf1ab | GSCEVVHDQVNYVPL | 6035 | 6049 | Positive |
| hCoV229E | orf1ab | GSELTLKHFFFTQKG | 4495 | 4509 | Positive |
| hCoV229E | orf1ab | GSGKSHCSIGIGVYY | 5281 | 5295 | Positive |
| hCoV229E | orf1ab | GSGMACNRVTLAVAS | 1 | 12 | Positive |
| hCoV229E | orf1ab | GSGQAITSTIDSNTT | 3983 | 3997 | Positive |
| hCoV229E | orf1ab | GSGSGMACNRVTLAV | 1 | 10 | Positive |

| | | | | | |
|---|---|---|---|---|---|
| hCoV229E | orf1ab | GSGSGQAITSTIDSN | 3981 | 3995 | Positive |
| hCoV229E | orf1ab | GSGSGSGMACNRVTL | 1 | 8 | Positive |
| hCoV229E | orf1ab | GSLPYSSIQANAYYK | 2651 | 2665 | Positive |
| hCoV229E | orf1ab | GSPVMTNGSNILEAF | 220 | 234 | Positive |
| hCoV229E | orf1ab | GYVGSLSNNHHAICN | 5839 | 5853 | Positive |
| hCoV229E | orf1ab | HAICNVHRNEHVASG | 5849 | 5863 | Positive |
| hCoV229E | orf1ab | HDEMCILHCSNFNTL | 4365 | 4379 | Positive |
| hCoV229E | orf1ab | HDFFTWHEGRTIYGN | 4161 | 4175 | Positive |
| hCoV229E | orf1ab | HGVTLKIKVSQTNMH | 3047 | 3061 | Positive |
| hCoV229E | orf1ab | HKEVLFNKYFKYWGQ | 4341 | 4355 | Positive |
| hCoV229E | orf1ab | HNDRIKSFSTFESAY | 820 | 834 | Positive |
| hCoV229E | orf1ab | HSDFGLGDLVDSYFE | 3609 | 3623 | Positive |
| hCoV229E | orf1ab | HVNRMDTFCFNVRNS | 338 | 352 | Positive |
| hCoV229E | orf1ab | IADDTYVMGLHGNQT | 56 | 70 | Positive |
| hCoV229E | orf1ab | IEEIEYVHGDALHTL | 174 | 188 | Positive |
| hCoV229E | orf1ab | IFDIETVDVKHEMSP | 1019 | 1033 | Positive |
| hCoV229E | orf1ab | IKLVDSELLSTLSVD | 2313 | 2327 | Positive |
| hCoV229E | orf1ab | ILEAFTKPVFISALV | 230 | 244 | Positive |
| hCoV229E | orf1ab | ILQSVASSFVGMPSF | 3627 | 3641 | Positive |
| hCoV229E | orf1ab | INGACGSPGYNLKNG | 3105 | 3119 | Positive |
| hCoV229E | orf1ab | IPATSAARLVVVVPD | 3749 | 3763 | Positive |
| hCoV229E | orf1ab | IPKEEFVVKEKLNAF | 1262 | 1275 | Positive |
| hCoV229E | orf1ab | IPSNVYLVGKTLIFT | 2587 | 2601 | Positive |
| hCoV229E | orf1ab | IQQWGYVGSLSNNHH | 5835 | 5849 | Positive |
| hCoV229E | orf1ab | ISDHEFTSAISNAHR | 2365 | 2379 | Positive |
| hCoV229E | orf1ab | ITDAAVKAAESKAFV | 690 | 704 | Positive |
| hCoV229E | orf1ab | ITELLQFVTDPTLIV | 4431 | 4445 | Positive |
| hCoV229E | orf1ab | ITESKYSCKEVFKNC | 2265 | 2279 | Positive |
| hCoV229E | orf1ab | ITTIQGPPGSGKSHC | 5273 | 5287 | Positive |
| hCoV229E | orf1ab | IVCKVENKAEAEWIE | 802 | 816 | Positive |
| hCoV229E | orf1ab | KAAGGKVSFSDDVEV | 894 | 908 | Positive |
| hCoV229E | orf1ab | KAEEFVAASDITLKC | 6375 | 6389 | Positive |
| hCoV229E | orf1ab | KAMLDFSRDHNDILY | 2939 | 2953 | Positive |
| hCoV229E | orf1ab | KAPQRTGVVLKRSLK | 1961 | 1975 | Positive |
| hCoV229E | orf1ab | KCSCTSGRLEESGAV | 1125 | 1139 | Positive |
| hCoV229E | orf1ab | KCWTEEDLSIGPHEF | 4861 | 4875 | Positive |
| hCoV229E | orf1ab | KDADYNAKDEISVTP | 1239 | 1253 | Positive |
| hCoV229E | orf1ab | KDCVDINKAIPSGNL | 1513 | 1527 | Positive |
| hCoV229E | orf1ab | KDDAFYIVKRCIKSV | 4127 | 4141 | Positive |
| hCoV229E | orf1ab | KDDYCDGSLGCKMCL | 2035 | 2049 | Positive |
| hCoV229E | orf1ab | KDNDGKNVHLKDVTK | 3789 | 3803 | Positive |
| hCoV229E | orf1ab | KDVTKENQEILVWPL | 3799 | 3813 | Positive |
| hCoV229E | orf1ab | KEASKQCFKIFERGS | 5461 | 5475 | Positive |
| hCoV229E | orf1ab | KEEFVVKEKLNAFLV | 1263 | 1277 | Positive |
| hCoV229E | orf1ab | KEHIGLAGKVRVGTG | 1331 | 1345 | Positive |
| hCoV229E | orf1ab | KEKLNAFLVHGNVVF | 1269 | 1283 | Positive |
| hCoV229E | orf1ab | KETNWFLHFIPFDVI | 2115 | 2129 | Positive |
| hCoV229E | orf1ab | KEVKTSSKVVLSDAL | 198 | 212 | Positive |
| hCoV229E | orf1ab | KFGKAGLYYESISYE | 4571 | 4585 | Positive |

| | | | | | |
|---|---|---|---|---|---|
| hCoV229E | orf1ab | KGCNAVAKHDFFTWH | 4153 | 4167 | Positive |
| hCoV229E | orf1ab | KGDKGDAEDTLNKLS | 1749 | 1763 | Positive |
| hCoV229E | orf1ab | KGDKLSVEHYNEWAQ | 3185 | 3199 | Positive |
| hCoV229E | orf1ab | KGHYTVYDTAKKSMY | 1861 | 1875 | Positive |
| hCoV229E | orf1ab | KHERDLLVKAYNTIN | 1365 | 1379 | Positive |
| hCoV229E | orf1ab | KHFFFTQKGDAAIKD | 4501 | 4515 | Positive |
| hCoV229E | orf1ab | KIEQKPVSLIKVAPK | 1445 | 1459 | Positive |
| hCoV229E | orf1ab | KIMAKAPQRTGVVLK | 1957 | 1971 | Positive |
| hCoV229E | orf1ab | KLYNPKAIHDIGNPK | 5913 | 5927 | Positive |
| hCoV229E | orf1ab | KNINWFVYVRKDGKH | 6283 | 6296 | Positive |
| hCoV229E | orf1ab | KNPINSNVKTLEYDY | 5943 | 5957 | Positive |
| hCoV229E | orf1ab | KPLDYKRQNNLAIEE | 162 | 176 | Positive |
| hCoV229E | orf1ab | KQCFKIFERGSVQVD | 5465 | 5479 | Positive |
| hCoV229E | orf1ab | KRWLPHDAIVVDNDV | 6543 | 6557 | Positive |
| hCoV229E | orf1ab | KVITIKVTEDGVNVH | 1601 | 1615 | Positive |
| hCoV229E | orf1ab | KVKYATVVVGSTEEV | 518 | 532 | Positive |
| hCoV229E | orf1ab | KYATVVVGSTEEVKS | 520 | 534 | Positive |
| hCoV229E | orf1ab | KYWGQDYHPDCVDCH | 4351 | 4365 | Positive |
| hCoV229E | orf1ab | LAEKFDYNVSVMQMD | 3357 | 3371 | Positive |
| hCoV229E | orf1ab | LAIEEIEYVHGDALH | 172 | 186 | Positive |
| hCoV229E | orf1ab | LAKAMLDFSRDHNDI | 2937 | 2951 | Positive |
| hCoV229E | orf1ab | LAVIDEHVFYKKDGV | 864 | 878 | Positive |
| hCoV229E | orf1ab | LAVQIGNNNVCTYEH | 5631 | 5645 | Positive |
| hCoV229E | orf1ab | LCQYFNSTTLCVPHN | 6507 | 6521 | Positive |
| hCoV229E | orf1ab | LCRPIKLVDSELLST | 2309 | 2323 | Positive |
| hCoV229E | orf1ab | LDAVKQGAKPVGNCV | 3961 | 3975 | Positive |
| hCoV229E | orf1ab | LDEHESKFWDESFYA | 4971 | 4985 | Positive |
| hCoV229E | orf1ab | LDENAQKFFDFGDFL | 1915 | 1929 | Positive |
| hCoV229E | orf1ab | LEEFDVVFGKRGGGN | 98 | 112 | Positive |
| hCoV229E | orf1ab | LEYDYMTHGQMEGLC | 5953 | 5967 | Positive |
| hCoV229E | orf1ab | LFAAETVKAKEESVK | 5127 | 5141 | Positive |
| hCoV229E | orf1ab | LGLSISDHEFTSAIS | 2361 | 2375 | Positive |
| hCoV229E | orf1ab | LGYSSLNDEFSINEV | 3243 | 3257 | Positive |
| hCoV229E | orf1ab | LILTCERVVKLQNNE | 3813 | 3827 | Positive |
| hCoV229E | orf1ab | LKCVRNVFENFEDWH | 2543 | 2557 | Positive |
| hCoV229E | orf1ab | LKEIVGPKELLLLWE | 5149 | 5163 | Positive |
| hCoV229E | orf1ab | LKIKVSQTNMHTPRH | 3051 | 3065 | Positive |
| hCoV229E | orf1ab | LKPHFISQGLDAAWN | 1714 | 1727 | Positive |
| hCoV229E | orf1ab | LLKSLDLTVVSKVHE | 6417 | 6431 | Positive |
| hCoV229E | orf1ab | LLVRGNGKCLSFSNH | 6739 | 6753 | Positive |
| hCoV229E | orf1ab | LNRLMCDIVCTIPAD | 1561 | 1575 | Positive |
| hCoV229E | orf1ab | LPEVNADIVVVDEVS | 5359 | 5373 | Positive |
| hCoV229E | orf1ab | LPTYYIYDEEGGNDL | 966 | 980 | Positive |
| hCoV229E | orf1ab | LQDFLPRSTMEEDFL | 6311 | 6325 | Positive |
| hCoV229E | orf1ab | LQEVKDNDGKNVHLK | 3785 | 3799 | Positive |
| hCoV229E | orf1ab | LSDDGVVCYNKTYAE | 4821 | 4834 | Positive |
| hCoV229E | orf1ab | LSGVAYTAFSGPVDK | 1847 | 1861 | Positive |
| hCoV229E | orf1ab | LVDSYFENDSILQSV | 3617 | 3631 | Positive |
| hCoV229E | orf1ab | LWESGKAKPPLNRNS | 5161 | 5175 | Positive |

| | | | | | |
|---|---|---|---|---|---|
| hCoV229E | orf1ab | LYELVQRFSFWTMFC | 6637 | 6651 | Positive |
| hCoV229E | orf1ab | LYYESISYEEQDAIF | 4577 | 4591 | Positive |
| hCoV229E | orf1ab | MAEQAAAAMYKEARA | 3691 | 3705 | Positive |
| hCoV229E | orf1ab | MEPIDYNVVTQRMCA | 5413 | 5427 | Positive |
| hCoV229E | orf1ab | MLDFSRDHNDILYTP | 2941 | 2955 | Positive |
| hCoV229E | orf1ab | MSPFEMPFEELNGLK | 1031 | 1045 | Positive |
| hCoV229E | orf1ab | MYAYVTTKPGMKYVK | 3863 | 3877 | Positive |
| hCoV229E | orf1ab | MYEKSTVLQAAGLCV | 4987 | 5001 | Positive |
| hCoV229E | orf1ab | NADIVVVDEVSMCTN | 5363 | 5377 | Positive |
| hCoV229E | orf1ab | NAFDVFVTTIQTVFD | 460 | 474 | Positive |
| hCoV229E | orf1ab | NAFLVHGNVVFYQGD | 1273 | 1287 | Positive |
| hCoV229E | orf1ab | NALPEVNADIVVVDE | 5357 | 5371 | Positive |
| hCoV229E | orf1ab | NALVKLVTTKLKGVR | 498 | 512 | Positive |
| hCoV229E | orf1ab | NALYNNEGGRAFMYA | 3851 | 3865 | Positive |
| hCoV229E | orf1ab | NANEADYRCACYAYL | 2923 | 2937 | Positive |
| hCoV229E | orf1ab | NANMSLADCKSALGL | 2349 | 2363 | Positive |
| hCoV229E | orf1ab | NANRFNVAITRAKKG | 5553 | 5567 | Positive |
| hCoV229E | orf1ab | NATVVIGTTKFYGGW | 4647 | 4661 | Positive |
| hCoV229E | orf1ab | NAVANGSSPQIIKQL | 3653 | 3667 | Positive |
| hCoV229E | orf1ab | NAVLFSATAVKTGGK | 6241 | 6255 | Positive |
| hCoV229E | orf1ab | NCLASECFMKSDIFG | 4311 | 4325 | Positive |
| hCoV229E | orf1ab | NCNNFNVKKLQRQLY | 4777 | 4791 | Positive |
| hCoV229E | orf1ab | NCNVDMYPEFSIVCR | 5971 | 5985 | Positive |
| hCoV229E | orf1ab | NCVKMLTNGSGSGQA | 3973 | 3987 | Positive |
| hCoV229E | orf1ab | NCYRNSNVDESFVDD | 4793 | 4807 | Positive |
| hCoV229E | orf1ab | NDGKNVHLKDVTKEN | 3791 | 3805 | Positive |
| hCoV229E | orf1ab | NDKITEFQLDYSIDV | 624 | 638 | Positive |
| hCoV229E | orf1ab | NDLSLPVMISEWPLF | 978 | 991 | Positive |
| hCoV229E | orf1ab | NDMVLSLVRSGKLLV | 6727 | 6741 | Positive |
| hCoV229E | orf1ab | NDVVDYVSDADFSVT | 6555 | 6569 | Positive |
| hCoV229E | orf1ab | NEADYRCACYAYLAK | 2925 | 2939 | Positive |
| hCoV229E | orf1ab | NEHVASGDAIMTRCL | 5857 | 5871 | Positive |
| hCoV229E | orf1ab | NENLAHGGGLAKALD | 1305 | 1319 | Positive |
| hCoV229E | orf1ab | NEVVKQMFGVNLQSG | 3255 | 3269 | Positive |
| hCoV229E | orf1ab | NFGMLNGNAIATVKS | 6263 | 6277 | Positive |
| hCoV229E | orf1ab | NFNTLFATTIPNTAF | 4375 | 4389 | Positive |
| hCoV229E | orf1ab | NFVSSYAKPEEKLSA | 2391 | 2405 | Positive |
| hCoV229E | orf1ab | NGFICEKLAIGGSIA | 6611 | 6625 | Positive |
| hCoV229E | orf1ab | NGIRVLKTSDNNCWV | 1689 | 1703 | Positive |
| hCoV229E | orf1ab | NGLNAPNGPFDALFL | 3513 | 3527 | Positive |
| hCoV229E | orf1ab | NGSVLEMAKEVKTSS | 190 | 204 | Positive |
| hCoV229E | orf1ab | NGVFMSTAKCWTEED | 4853 | 4867 | Positive |
| hCoV229E | orf1ab | NGVLHKAYIDVLRNS | 2329 | 2343 | Positive |
| hCoV229E | orf1ab | NGVQRLFYVNANGGS | 2177 | 2191 | Positive |
| hCoV229E | orf1ab | NGYVCGTGLWNLVFN | 2723 | 2737 | Positive |
| hCoV229E | orf1ab | NHGCTCDRTAIQSFD | 4057 | 4071 | Positive |
| hCoV229E | orf1ab | NHHAICNVHRNEHVA | 5847 | 5861 | Positive |
| hCoV229E | orf1ab | NIASNSKEWAYCVEM | 3567 | 3581 | Positive |
| hCoV229E | orf1ab | NIIHANYVFWRNSTV | 6681 | 6695 | Positive |

| | | | | | |
|---|---|---|---|---|---|
| hCoV229E | orf1ab | NILACYDGCAQGVFG | 3077 | 3091 | Positive |
| hCoV229E | orf1ab | NILPTMTQLNLKYAI | 4597 | 4611 | Positive |
| hCoV229E | orf1ab | NILSMFSSSFSVAAM | 2737 | 2751 | Positive |
| hCoV229E | orf1ab | NKAEAEWIELFPHND | 808 | 822 | Positive |
| hCoV229E | orf1ab | NKAIPSGNLIKFDID | 1519 | 1532 | Positive |
| hCoV229E | orf1ab | NKDASFIGKNLKSNC | 4105 | 4119 | Positive |
| hCoV229E | orf1ab | NKDVNTHSTRLTITE | 4419 | 4433 | Positive |
| hCoV229E | orf1ab | NKEFYDFLRSQGFFD | 4479 | 4493 | Positive |
| hCoV229E | orf1ab | NKFVLGDVEIFVAFV | 1727 | 1741 | Positive |
| hCoV229E | orf1ab | NKFVPVKEASKQCFK | 5455 | 5469 | Positive |
| hCoV229E | orf1ab | NKGGRTVQSHIMRAA | 5897 | 5911 | Positive |
| hCoV229E | orf1ab | NKGSFVGADGELPVA | 6121 | 6135 | Positive |
| hCoV229E | orf1ab | NKINLCDDPETAQEL | 3583 | 3597 | Positive |
| hCoV229E | orf1ab | NKIQPWTNDALNTIC | 1223 | 1237 | Positive |
| hCoV229E | orf1ab | NKTLNEGVLESFSVT | 4955 | 4969 | Positive |
| hCoV229E | orf1ab | NKYFKYWGQDYHPDC | 4347 | 4361 | Positive |
| hCoV229E | orf1ab | NLAHGGGLAKALDVY | 1307 | 1321 | Positive |
| hCoV229E | orf1ab | NLEGVNGGSLYVNNH | 5995 | 6009 | Positive |
| hCoV229E | orf1ab | NLFEGDKFVGTFESA | 2869 | 2883 | Positive |
| hCoV229E | orf1ab | NLKNGEVEFVYMHQI | 3115 | 3129 | Positive |
| hCoV229E | orf1ab | NLNNLRRGAVLGYIG | 3913 | 3927 | Positive |
| hCoV229E | orf1ab | NLQSGKTTSMFKSIS | 3265 | 3279 | Positive |
| hCoV229E | orf1ab | NLTQQGLLGTCAKRF | 744 | 758 | Positive |
| hCoV229E | orf1ab | NLVFNILSMFSSSFS | 2733 | 2747 | Positive |
| hCoV229E | orf1ab | NLVTMIAYAILYFFA | 2805 | 2819 | Positive |
| hCoV229E | orf1ab | NMARNGVVPLSVIPA | 3737 | 3751 | Positive |
| hCoV229E | orf1ab | NMHTPRHSFRTLKSG | 3059 | 3073 | Positive |
| hCoV229E | orf1ab | NMNIASNSKEWAYCV | 3565 | 3579 | Positive |
| hCoV229E | orf1ab | NMRTNWTIRGSFING | 3093 | 3107 | Positive |
| hCoV229E | orf1ab | NMSLADCKSALGLSI | 2351 | 2365 | Positive |
| hCoV229E | orf1ab | NNCWVNAVCIALQYL | 1699 | 1712 | Positive |
| hCoV229E | orf1ab | NNEGGRAFMYAYVTT | 3855 | 3869 | Positive |
| hCoV229E | orf1ab | NNEIMPGKMKVKATK | 3825 | 3839 | Positive |
| hCoV229E | orf1ab | NNEQGTPLTPILSCG | 1379 | 1393 | Positive |
| hCoV229E | orf1ab | NNHAFHTPAYDKRAM | 6007 | 6021 | Positive |
| hCoV229E | orf1ab | NNLAIEEIEYVHGDA | 170 | 184 | Positive |
| hCoV229E | orf1ab | NNLRRGAVLGYIGAT | 3915 | 3929 | Positive |
| hCoV229E | orf1ab | NNNGTNVTQVKNASV | 2289 | 2303 | Positive |
| hCoV229E | orf1ab | NNNVCTYEHVISYMG | 5637 | 5651 | Positive |
| hCoV229E | orf1ab | NNTSAQYVFSTVNAL | 5345 | 5359 | Positive |
| hCoV229E | orf1ab | NNVYCYNTELMEGSL | 2639 | 2653 | Positive |
| hCoV229E | orf1ab | NQDLNGNFYDFGDFV | 4271 | 4285 | Positive |
| hCoV229E | orf1ab | NQEILVWPLILTCER | 3805 | 3819 | Positive |
| hCoV229E | orf1ab | NQMLTVNVVAFLYAA | 3161 | 3175 | Positive |
| hCoV229E | orf1ab | NQTLFCNIMKFSDRP | 68 | 82 | Positive |
| hCoV229E | orf1ab | NRFNVAITRAKKGIF | 5555 | 5569 | Positive |
| hCoV229E | orf1ab | NRKSKVVSAMHSLLF | 3707 | 3721 | Positive |
| hCoV229E | orf1ab | NRMAEQAAAAMYKEA | 3689 | 3703 | Positive |
| hCoV229E | orf1ab | NRMDTFCFNVRNSVT | 340 | 354 | Positive |

| | | | | | |
|---|---|---|---|---|---|
| hCoV229E | orf1ab | NRRQLDVVKRFIHKN | 5485 | 5499 | Positive |
| hCoV229E | orf1ab | NRVTLAVASDSEISA | 4 | 18 | Positive |
| hCoV229E | orf1ab | NRYKYYSGNANEADY | 2915 | 2929 | Positive |
| hCoV229E | orf1ab | NSIFDIETVDVKHEM | 1017 | 1031 | Positive |
| hCoV229E | orf1ab | NSIQGSYERFTLSTN | 6227 | 6241 | Positive |
| hCoV229E | orf1ab | NSKEWAYCVEMHNKI | 3571 | 3585 | Positive |
| hCoV229E | orf1ab | NSLSAEGRKYIVKTS | 2439 | 2453 | Positive |
| hCoV229E | orf1ab | NSNVDESFVDDFYGY | 4797 | 4811 | Positive |
| hCoV229E | orf1ab | NSNVKTLEYDYMTHG | 5947 | 5961 | Positive |
| hCoV229E | orf1ab | NSTWSKAVFISPYNS | 5499 | 5513 | Positive |
| hCoV229E | orf1ab | NSVLDLSKFNCKHKA | 6701 | 6715 | Positive |
| hCoV229E | orf1ab | NSVMLQIQLTGILDG | 1057 | 1071 | Positive |
| hCoV229E | orf1ab | NSVSNDYCCFKHALG | 5809 | 5823 | Positive |
| hCoV229E | orf1ab | NSVTSINSAIVCASV | 1789 | 1803 | Positive |
| hCoV229E | orf1ab | NTDYFEMKNWFDPIE | 4215 | 4229 | Positive |
| hCoV229E | orf1ab | NTHSTRLTITELLQF | 4423 | 4437 | Positive |
| hCoV229E | orf1ab | NTSSSEAFVVGINYL | 6655 | 6669 | Positive |
| hCoV229E | orf1ab | NTTSAIDYDHEYSIM | 3011 | 3025 | Positive |
| hCoV229E | orf1ab | NVDESFVDDFYGYLQ | 4799 | 4813 | Positive |
| hCoV229E | orf1ab | NVDMYPEFSIVCRFD | 5973 | 5987 | Positive |
| hCoV229E | orf1ab | NVHDVTVTTDKSFEQ | 1613 | 1627 | Positive |
| hCoV229E | orf1ab | NVHLKDVTKENQEIL | 3795 | 3809 | Positive |
| hCoV229E | orf1ab | NVHRNEHVASGDAIM | 5853 | 5867 | Positive |
| hCoV229E | orf1ab | NVKTLEYDYMTHGQM | 5949 | 5963 | Positive |
| hCoV229E | orf1ab | NVQRCTRKLNRLMCD | 1553 | 1567 | Positive |
| hCoV229E | orf1ab | NVSDAYANLVPYYQL | 5253 | 5267 | Positive |
| hCoV229E | orf1ab | NVSRQDLTKYTMMDL | 4175 | 4189 | Positive |
| hCoV229E | orf1ab | NVSYIVTQNLVTMIA | 2797 | 2811 | Positive |
| hCoV229E | orf1ab | NWFDPIENEDIHRVY | 4223 | 4237 | Positive |
| hCoV229E | orf1ab | NWTIRGSFINGACGS | 3097 | 3111 | Positive |
| hCoV229E | orf1ab | NYDLSVINQRISYKH | 5377 | 5391 | Positive |
| hCoV229E | orf1ab | NYLGDFAQGPFIDGN | 6667 | 6681 | Positive |
| hCoV229E | orf1ab | NYLRCFLLYFVAQMI | 2091 | 2105 | Positive |
| hCoV229E | orf1ab | NYVAARLLGLQTQTV | 5515 | 5529 | Positive |
| hCoV229E | orf1ab | NYVPLRATNCITKCN | 6045 | 6059 | Positive |
| hCoV229E | orf1ab | NYVRDWLKSLKLNLT | 732 | 746 | Positive |
| hCoV229E | orf1ab | PDCVDCHDEMCILHC | 4359 | 4373 | Positive |
| hCoV229E | orf1ab | PGMLFILTSHNVAPL | 5217 | 5231 | Positive |
| hCoV229E | orf1ab | PGTAVLKRWLPHDAI | 6537 | 6551 | Positive |
| hCoV229E | orf1ab | PHDAIVVDNDVVDYV | 6547 | 6561 | Positive |
| hCoV229E | orf1ab | PHNMRVLHLGAGSDY | 6519 | 6533 | Positive |
| hCoV229E | orf1ab | PIDLPPSHATTYLSL | 5607 | 5621 | Positive |
| hCoV229E | orf1ab | PIENEDIHRVYAALG | 4227 | 4241 | Positive |
| hCoV229E | orf1ab | PINSNVKTLEYDYMT | 5945 | 5959 | Positive |
| hCoV229E | orf1ab | PKELLLLWESGKAKP | 5155 | 5169 | Positive |
| hCoV229E | orf1ab | PPGEQFTHIVPLLRK | 5733 | 5747 | Positive |
| hCoV229E | orf1ab | PQRTGVVLKRSLKYN | 1963 | 1977 | Positive |
| hCoV229E | orf1ab | PSEKDKHLDNNVQRC | 1543 | 1557 | Positive |
| hCoV229E | orf1ab | PTTVTCENLESAVLF | 608 | 622 | Positive |

| | | | | | |
|---|---|---|---|---|---|
| hCoV229E | orf1ab | PVEYVSYFDGVKTVL | 3453 | 3467 | Positive |
| hCoV229E | orf1ab | PVRARAPPGEQFTHI | 5727 | 5741 | Positive |
| hCoV229E | orf1ab | PYPDPSRIISAGVFV | 4893 | 4907 | Positive |
| hCoV229E | orf1ab | QDYHPDCVDCHDEMC | 4355 | 4369 | Positive |
| hCoV229E | orf1ab | QGPPGSGKSHCSIGI | 5277 | 5291 | Positive |
| hCoV229E | orf1ab | QLEPCVQSRLLSGVA | 1837 | 1851 | Positive |
| hCoV229E | orf1ab | QPTGPAYVMIDKVEF | 2231 | 2245 | Positive |
| hCoV229E | orf1ab | QQEATHIAEDVVDQV | 997 | 1013 | Positive |
| hCoV229E | orf1ab | QRITTIQGPPGSGKS | 5271 | 5285 | Positive |
| hCoV229E | orf1ab | QRQLYDNCYRNSNVD | 4787 | 4801 | Positive |
| hCoV229E | orf1ab | QTEFVSNSHLLTHCA | 3937 | 3951 | Positive |
| hCoV229E | orf1ab | QTVDSAQGSEYDYVI | 5527 | 5541 | Positive |
| hCoV229E | orf1ab | QVPIGTNDPIRFCLE | 4031 | 4045 | Positive |
| hCoV229E | orf1ab | QWGYVGSLSNNHHAI | 5837 | 5851 | Positive |
| hCoV229E | orf1ab | QYFNSTTLCVPHNMR | 6509 | 6523 | Positive |
| hCoV229E | orf1ab | QYSNESWFEDDYKAF | 670 | 684 | Positive |
| hCoV229E | orf1ab | RAAIKLYNPKAIHDI | 5909 | 5923 | Positive |
| hCoV229E | orf1ab | RCDVLLSDLSFNNFV | 2379 | 2393 | Positive |
| hCoV229E | orf1ab | RCPAEIVNTVSELVY | 5439 | 5453 | Positive |
| hCoV229E | orf1ab | RDGVQVGYCVHGIKY | 1805 | 1819 | Positive |
| hCoV229E | orf1ab | RDIVCKVENKAEAEW | 800 | 814 | Positive |
| hCoV229E | orf1ab | RDYKLANDAKESLRL | 5113 | 5127 | Positive |
| hCoV229E | orf1ab | RDYVDKWDDFCRQYS | 658 | 672 | Positive |
| hCoV229E | orf1ab | RGRAIIVSVEQLEPC | 1827 | 1841 | Positive |
| hCoV229E | orf1ab | RICRFGVAFLPVEYV | 3443 | 3457 | Positive |
| hCoV229E | orf1ab | RKMAQPSGFVEKCVV | 2969 | 2983 | Positive |
| hCoV229E | orf1ab | RLEPCNGTDIDYCVR | 4085 | 4099 | Positive |
| hCoV229E | orf1ab | RLMCDIVCTIPADYI | 1563 | 1577 | Positive |
| hCoV229E | orf1ab | RLSKMGILKAEEFVA | 6367 | 6381 | Positive |
| hCoV229E | orf1ab | RPIKLVDSELLSTLS | 2311 | 2325 | Positive |
| hCoV229E | orf1ab | RQNNLAIEEIEYVHG | 168 | 182 | Positive |
| hCoV229E | orf1ab | RTRSTLNLEGVNGGS | 5989 | 6003 | Positive |
| hCoV229E | orf1ab | RYFDCYEGGCITSRE | 4541 | 4555 | Positive |
| hCoV229E | orf1ab | SAARLVVVVPDHDSF | 3753 | 3767 | Positive |
| hCoV229E | orf1ab | SAEWKCGYSMPGIYK | 6459 | 6473 | Positive |
| hCoV229E | orf1ab | SAVLFVNDKITEFQL | 618 | 632 | Positive |
| hCoV229E | orf1ab | SDAYANLVPYYQLIG | 5255 | 5269 | Positive |
| hCoV229E | orf1ab | SDEGYTVVIGDVAYF | 555 | 568 | Positive |
| hCoV229E | orf1ab | SDGYFRLMASPNSVL | 570 | 584 | Positive |
| hCoV229E | orf1ab | SDIFGQDFKTFDLLK | 4321 | 4335 | Positive |
| hCoV229E | orf1ab | SDITLKCCTVTYLND | 6383 | 6397 | Positive |
| hCoV229E | orf1ab | SDLNTSELLTKAIDV | 1645 | 1659 | Positive |
| hCoV229E | orf1ab | SDLSFNNFVSSYAKP | 2385 | 2399 | Positive |
| hCoV229E | orf1ab | SDNNCWVNAVCIALQ | 1697 | 1711 | Positive |
| hCoV229E | orf1ab | SDRTLFDALKFFEVT | 5573 | 5587 | Positive |
| hCoV229E | orf1ab | SDSEISAYGCSTIAQ | 12 | 26 | Positive |
| hCoV229E | orf1ab | SDWSDIRDYKLANDA | 5107 | 5121 | Positive |
| hCoV229E | orf1ab | SDYGVAPGTAVLKRW | 6531 | 6545 | Positive |
| hCoV229E | orf1ab | SEISAYGCSTIAQAV | 14 | 28 | Positive |

| | | | | | |
|---|---|---|---|---|---|
| hCoV229E | orf1ab | SEYDYVIFAQTSDTA | 5535 | 5549 | Positive |
| hCoV229E | orf1ab | SHSLFCTRDFAMRHV | 5661 | 5675 | Positive |
| hCoV229E | orf1ab | SIGPHEFCSQHTMQI | 4869 | 4883 | Positive |
| hCoV229E | orf1ab | SKHPKPEYRKVFYAL | 4933 | 4947 | Positive |
| hCoV229E | orf1ab | SLADCKSALGLSISD | 2353 | 2367 | Positive |
| hCoV229E | orf1ab | SLSNNHHAICNVHRN | 5843 | 5857 | Positive |
| hCoV229E | orf1ab | SLYVNNHAFHTPAYD | 6003 | 6017 | Positive |
| hCoV229E | orf1ab | SMCTNYDLSVINQRI | 5373 | 5387 | Positive |
| hCoV229E | orf1ab | SMDIDVFNKLSTSDW | 5095 | 5109 | Positive |
| hCoV229E | orf1ab | SMFKSISLFAGFFVM | 3273 | 3287 | Positive |
| hCoV229E | orf1ab | SMLTLDDRGLALDNA | 1489 | 1503 | Positive |
| hCoV229E | orf1ab | SMMILSDDGVVCYNK | 4817 | 4831 | Positive |
| hCoV229E | orf1ab | SMPGSHSLFCTRDFA | 5657 | 5671 | Positive |
| hCoV229E | orf1ab | SMYDGDRFVKHDLSL | 1873 | 1887 | Positive |
| hCoV229E | orf1ab | SMYNLLKGCNAVAKH | 4147 | 4161 | Positive |
| hCoV229E | orf1ab | SNCVRFKNVDKDDAF | 4117 | 4131 | Positive |
| hCoV229E | orf1ab | SNNHHAICNVHRNEH | 5845 | 5859 | Positive |
| hCoV229E | orf1ab | SNVRRQVASGVIDIS | 368 | 382 | Positive |
| hCoV229E | orf1ab | SPEKLKSYAASYNRY | 2903 | 2917 | Positive |
| hCoV229E | orf1ab | SQLDENAQKFFDFGD | 1913 | 1927 | Positive |
| hCoV229E | orf1ab | STEEVKSSRVERSNA | 528 | 542 | Positive |
| hCoV229E | orf1ab | STFESAYMPIADPTH | 828 | 842 | Positive |
| hCoV229E | orf1ab | STFITPEVSRELGNI | 2211 | 2225 | Positive |
| hCoV229E | orf1ab | STNLFEGDKFVGTFE | 2867 | 2881 | Positive |
| hCoV229E | orf1ab | SVTLLDEHESKFWDE | 4967 | 4981 | Positive |
| hCoV229E | orf1ab | SYEEQDAIFLLTKRN | 4583 | 4597 | Positive |
| hCoV229E | orf1ab | TCENLESAVLFVNDK | 612 | 626 | Positive |
| hCoV229E | orf1ab | TDAVILLERYVSLAI | 4913 | 4927 | Positive |
| hCoV229E | orf1ab | TDECRLAMLGAEMTS | 354 | 368 | Positive |
| hCoV229E | orf1ab | TDFAEDVCTCYDNSI | 6215 | 6229 | Positive |
| hCoV229E | orf1ab | TDKSFEQQVGVIADK | 1621 | 1635 | Positive |
| hCoV229E | orf1ab | TDLQSESSCGLFKDC | 5589 | 5603 | Positive |
| hCoV229E | orf1ab | TDPLFSNIQPFIVMV | 2069 | 2083 | Positive |
| hCoV229E | orf1ab | TDPTLIVASSPALVD | 4439 | 4453 | Positive |
| hCoV229E | orf1ab | TDQYLCGADGKPVMS | 116 | 130 | Positive |
| hCoV229E | orf1ab | TDSMLTLDDRGLALD | 1487 | 1501 | Positive |
| hCoV229E | orf1ab | TDTEVCKVKDFVSGL | 1421 | 1435 | Positive |
| hCoV229E | orf1ab | TDVTKLYLGGLNYYC | 5053 | 5067 | Positive |
| hCoV229E | orf1ab | TEDLLCVADDKPIVL | 1471 | 1485 | Positive |
| hCoV229E | orf1ab | TEHKEVLFNKYFKYW | 4339 | 4353 | Positive |
| hCoV229E | orf1ab | TFESAAAGTFVIDIR | 2879 | 2893 | Positive |
| hCoV229E | orf1ab | TFVEEEHVNRMDTFC | 332 | 346 | Positive |
| hCoV229E | orf1ab | TGCCNTDYFEMKNWF | 4211 | 4225 | Positive |
| hCoV229E | orf1ab | TGELVRFVKSICNSA | 426 | 440 | Positive |
| hCoV229E | orf1ab | TGFMVMDYKCSCTSG | 1117 | 1131 | Positive |
| hCoV229E | orf1ab | TGVCVERLLHAIQVL | 3219 | 3233 | Positive |
| hCoV229E | orf1ab | THGQMEGLCLFWNCN | 5959 | 5973 | Positive |
| hCoV229E | orf1ab | THIAEDVVDQVEEVN | 1005 | 1017 | Positive |
| hCoV229E | orf1ab | TILDIGQARVAYQVA | 4525 | 4539 | Positive |

| | | | | | |
|---|---|---|---|---|---|
| hCoV229E | orf1ab | TINENQAVTQIPATS | 2463 | 2477 | Positive |
| hCoV229E | orf1ab | TIQTVFDCAVETCTI | 468 | 482 | Positive |
| hCoV229E | orf1ab | TKLVPGMLFILTSHN | 5213 | 5227 | Positive |
| hCoV229E | orf1ab | TKTDAVILLERYVSL | 4911 | 4925 | Positive |
| hCoV229E | orf1ab | TLLDEHESKFWDESF | 4969 | 4983 | Positive |
| hCoV229E | orf1ab | TLNEGVLESFSVTLL | 4957 | 4971 | Positive |
| hCoV229E | orf1ab | TMANQEKYSTIYKLH | 5235 | 5249 | Positive |
| hCoV229E | orf1ab | TMHGVTLKIKVSQTN | 3045 | 3059 | Positive |
| hCoV229E | orf1ab | TMTDLQSESSCGLFK | 5587 | 5601 | Positive |
| hCoV229E | orf1ab | TMTQLNLKYAISGKE | 4601 | 4615 | Positive |
| hCoV229E | orf1ab | TNIAFELFAKRKVGL | 6161 | 6175 | Positive |
| hCoV229E | orf1ab | TPEVSRELGNITKTN | 2215 | 2229 | Positive |
| hCoV229E | orf1ab | TSAIDYDHEYSIMRL | 3013 | 3027 | Positive |
| hCoV229E | orf1ab | TSGDLAVQIGNNNVC | 5627 | 5641 | Positive |
| hCoV229E | orf1ab | TSGRLEESGAVLFCT | 1129 | 1143 | Positive |
| hCoV229E | orf1ab | TTFDCYNLWQTFTDV | 6093 | 6107 | Positive |
| hCoV229E | orf1ab | TTSGDATTAYANSVF | 4743 | 4757 | Positive |
| hCoV229E | orf1ab | TVAGIPSNVYLVGKT | 2583 | 2597 | Positive |
| hCoV229E | orf1ab | TVDHSAFAYESAVVN | 1675 | 1689 | Positive |
| hCoV229E | orf1ab | TVNALPEVNADIVVV | 5355 | 5369 | Positive |
| hCoV229E | orf1ab | TYMDLLLDDFVSLLK | 6405 | 6419 | Positive |
| hCoV229E | orf1ab | TYYIYDEEGGNDLSL | 968 | 982 | Positive |
| hCoV229E | orf1ab | VADDKPIVLFTDSML | 1477 | 1491 | Positive |
| hCoV229E | orf1ab | VAITRAKKGIFCIMS | 5559 | 5573 | Positive |
| hCoV229E | orf1ab | VAPGTAVLKRWLPHD | 6535 | 6549 | Positive |
| hCoV229E | orf1ab | VAQMISTVGVFLGYK | 2101 | 2115 | Positive |
| hCoV229E | orf1ab | VASDSEISAYGCSTI | 10 | 24 | Positive |
| hCoV229E | orf1ab | VASGVIDISTGWFDV | 374 | 388 | Positive |
| hCoV229E | orf1ab | VASSFVGMPSFVAYE | 3631 | 3645 | Positive |
| hCoV229E | orf1ab | VASSTFVEEEHVNRM | 328 | 342 | Positive |
| hCoV229E | orf1ab | VDCFVASSTFVEEEH | 324 | 338 | Positive |
| hCoV229E | orf1ab | VDDFYGYLQKHFSMM | 4805 | 4819 | Positive |
| hCoV229E | orf1ab | VDEVSMCTNYDLSVI | 5369 | 5383 | Positive |
| hCoV229E | orf1ab | VDGFGVNKIQPWTND | 1217 | 1231 | Positive |
| hCoV229E | orf1ab | VDNDVVDYVSDADFS | 6553 | 6567 | Positive |
| hCoV229E | orf1ab | VDSAQGSEYDYVIFA | 5529 | 5543 | Positive |
| hCoV229E | orf1ab | VDSNAGVCFGFDKWF | 2701 | 2715 | Positive |
| hCoV229E | orf1ab | VDTIVPPCPSILKVI | 704 | 718 | Positive |
| hCoV229E | orf1ab | VDVKHEMSPFEMPFE | 1025 | 1039 | Positive |
| hCoV229E | orf1ab | VEEVNSIFDIETVDV | 1013 | 1027 | Positive |
| hCoV229E | orf1ab | VFGSPVMTNGSNILE | 218 | 232 | Positive |
| hCoV229E | orf1ab | VFSNSNYLLEEFDVV | 90 | 104 | Positive |
| hCoV229E | orf1ab | VFYALLDWVKHLNKT | 4943 | 4957 | Positive |
| hCoV229E | orf1ab | VGIADDTYVMGLHGN | 54 | 68 | Positive |
| hCoV229E | orf1ab | VGSLSNNHHAICNVH | 5841 | 5855 | Positive |
| hCoV229E | orf1ab | VHEVIIDNKPWRWML | 6429 | 6443 | Positive |
| hCoV229E | orf1ab | VHGDALHTLRNGSVL | 180 | 194 | Positive |
| hCoV229E | orf1ab | VHGNVVFYQGDVDTV | 1277 | 1291 | Positive |
| hCoV229E | orf1ab | VIDEHVFYKKDGVYY | 866 | 880 | Positive |

| | | | | | |
|---|---|---|---|---|---|
| hCoV229E | orf1ab | VIDISTGWFDVYDDI | 378 | 392 | Positive |
| hCoV229E | orf1ab | VLFVNDKITEFQLDY | 620 | 634 | Positive |
| hCoV229E | orf1ab | VMISEWPLFESDKEE | 984 | 999 | Positive |
| hCoV229E | orf1ab | VMSEDLWQFVDHFGE | 128 | 142 | Positive |
| hCoV229E | orf1ab | VNEAQVQLEHYSSCV | 1767 | 1781 | Positive |
| hCoV229E | orf1ab | VNGGSLYVNNHAFHT | 5999 | 6013 | Positive |
| hCoV229E | orf1ab | VNLPTYYIYDEEGGN | 964 | 978 | Positive |
| hCoV229E | orf1ab | VNTVSELVYENKFVP | 5445 | 5459 | Positive |
| hCoV229E | orf1ab | VPEKFLNAFDVFVTT | 454 | 467 | Positive |
| hCoV229E | orf1ab | VPSDLNTSELLTKAI | 1643 | 1657 | Positive |
| hCoV229E | orf1ab | VRKDGKHVDHYDGFY | 6291 | 6305 | Positive |
| hCoV229E | orf1ab | VSDADFSVTGDCATV | 6561 | 6575 | Positive |
| hCoV229E | orf1ab | VSEIVNTVAGIPSNV | 2577 | 2591 | Positive |
| hCoV229E | orf1ab | VSFSDDVEVKDIEPV | 900 | 914 | Positive |
| hCoV229E | orf1ab | VSTVKPKPVISQLDE | 1903 | 1917 | Positive |
| hCoV229E | orf1ab | VSVMQMDIQGFVNIF | 3365 | 3379 | Positive |
| hCoV229E | orf1ab | VTDAKWYCYDKNPIN | 5933 | 5947 | Positive |
| hCoV229E | orf1ab | VTEDGVNVHDVTVTT | 1607 | 1621 | Positive |
| hCoV229E | orf1ab | VTPIKNTVDTIPKEE | 1251 | 1265 | Positive |
| hCoV229E | orf1ab | VTTGELVRFVKSICN | 424 | 438 | Positive |
| hCoV229E | orf1ab | VTVELEPPCRFVIDT | 3885 | 3899 | Positive |
| hCoV229E | orf1ab | VTVTTDKSFEQQVGV | 1617 | 1631 | Positive |
| hCoV229E | orf1ab | VVDQVEEVNSIFDIE | 1009 | 1023 | Positive |
| hCoV229E | orf1ab | VVDYVSDADFSVTGD | 6557 | 6571 | Positive |
| hCoV229E | orf1ab | VVHDQVNYVPLRATN | 6039 | 6053 | Positive |
| hCoV229E | orf1ab | VVKEKLNAFLVHGNV | 1267 | 1281 | Positive |
| hCoV229E | orf1ab | VVLMGILSNMNIASN | 3557 | 3571 | Positive |
| hCoV229E | orf1ab | VVMVGDYVAPVSTVK | 1893 | 1907 | Positive |
| hCoV229E | orf1ab | VVNGIRVLKTSDNNC | 1687 | 1701 | Positive |
| hCoV229E | orf1ab | VVPSEKDKHLDNNVQ | 1541 | 1555 | Positive |
| hCoV229E | orf1ab | VVSKVHEVIIDNKPW | 6425 | 6439 | Positive |
| hCoV229E | orf1ab | VVSSFEGYDFKYIEN | 2519 | 2533 | Positive |
| hCoV229E | orf1ab | VVVDEVSMCTNYDLS | 5367 | 5381 | Positive |
| hCoV229E | orf1ab | VVVGSTEEVKSSRVE | 524 | 538 | Positive |
| hCoV229E | orf1ab | VVVPDHDSFVKMMVD | 3759 | 3773 | Positive |
| hCoV229E | orf1ab | VVVQLKDSDINDMVL | 6717 | 6731 | Positive |
| hCoV229E | orf1ab | VYCPRHVIASNTTSA | 3001 | 3015 | Positive |
| hCoV229E | orf1ab | VYFSQLLCRPIKLVD | 2303 | 2317 | Positive |
| hCoV229E | orf1ab | VYTDTEVCKVKDFVS | 1419 | 1433 | Positive |
| hCoV229E | orf1ab | WDESFYASMYEKSTV | 4979 | 4993 | Positive |
| hCoV229E | orf1ab | WDYHITKVLAEKFDY | 3349 | 3363 | Positive |
| hCoV229E | orf1ab | WIELFPHNDRIKSFS | 814 | 828 | Positive |
| hCoV229E | orf1ab | WLVFSNSNYLLEEFD | 88 | 102 | Positive |
| hCoV229E | orf1ab | WQFVDHFGENEEIVI | 134 | 148 | Positive |
| hCoV229E | orf1ab | YAELGYIADISAFKA | 4833 | 4847 | Positive |
| hCoV229E | orf1ab | YDEEGGNDLSLPVMI | 972 | 986 | Positive |
| hCoV229E | orf1ab | YDYMTHGQMEGLCLF | 5955 | 5969 | Positive |
| hCoV229E | orf1ab | YESISYEEQDAIFLL | 4579 | 4593 | Positive |
| hCoV229E | orf1ab | YFENDSILQSVASSF | 3621 | 3635 | Positive |

| | | | | | |
|---|---|---|---|---|---|
| hCoV229E | orf1ab | YHPDCVDCHDEMCIL | 4357 | 4371 | Positive |
| hCoV229E | orf1ab | YKGKWVQVPIGTNDP | 4025 | 4039 | Positive |
| hCoV229E | orf1ab | YKRQNNLAIEEIEYV | 166 | 180 | Positive |
| hCoV229E | orf1ab | YKYYSGNANEADYRC | 2917 | 2931 | Positive |
| hCoV229E | orf1ab | YRNSNVDESFVDDFY | 4795 | 4809 | Positive |
| hCoV229E | orf1ab | YSGETFWRYNFDITE | 2253 | 2267 | Positive |
| hCoV229E | orf1ab | YSVDKCTRIIPARAR | 5321 | 5335 | Positive |
| hCoV229E | orf1ab | YYDDGSCEVVHDQVN | 6031 | 6045 | Positive |
| hCoV229E | orf1ab | ADYNAKDEISVTPIK | 1241 | 1255 | Negative |
| hCoV229E | orf1ab | AFCDEMVLKGVVGVL | 4253 | 4267 | Negative |
| hCoV229E | orf1ab | AFGNAGVCYDIFGVT | 2605 | 2619 | Negative |
| hCoV229E | orf1ab | AGGLELTTMRYFVKI | 5779 | 5793 | Negative |
| hCoV229E | orf1ab | ALGKVVANAMLKCVA | 4239 | 4253 | Negative |
| hCoV229E | orf1ab | ALQYLKPHFISQGLD | 1709 | 1723 | Negative |
| hCoV229E | orf1ab | APLRAPTMANQEKYS | 5229 | 5243 | Negative |
| hCoV229E | orf1ab | ARLKRFPVNTIVNGV | 2165 | 2179 | Negative |
| hCoV229E | orf1ab | ATVKSEDGNIKNINW | 6273 | 6287 | Negative |
| hCoV229E | orf1ab | AWNKFVLGDVEIFVA | 1725 | 1739 | Negative |
| hCoV229E | orf1ab | CGGILAVIDEHVFYK | 860 | 874 | Negative |
| hCoV229E | orf1ab | CGMTLKFVANIEGVS | 300 | 314 | Negative |
| hCoV229E | orf1ab | CGSQTVLRCGDCLRR | 5003 | 5017 | Negative |
| hCoV229E | orf1ab | CVHGIKYYSRVKSVR | 1813 | 1827 | Negative |
| hCoV229E | orf1ab | CYDGCAQGVFGVNMR | 3081 | 3095 | Negative |
| hCoV229E | orf1ab | DAAWNKFVLGDVEIF | 1723 | 1737 | Negative |
| hCoV229E | orf1ab | DDGVVCYNKTYAELG | 4823 | 4837 | Negative |
| hCoV229E | orf1ab | DDTYVMGLHGNQTLF | 58 | 72 | Negative |
| hCoV229E | orf1ab | DDVEVKDIEPVYRVK | 904 | 918 | Negative |
| hCoV229E | orf1ab | DELFVTVIVIKVISF | 2131 | 2145 | Negative |
| hCoV229E | orf1ab | DFKYIENGQLKNFEA | 2527 | 2541 | Negative |
| hCoV229E | orf1ab | DGCAQGVFGVNMRTN | 3083 | 3097 | Negative |
| hCoV229E | orf1ab | DISAFKATLYYQNGV | 4841 | 4855 | Negative |
| hCoV229E | orf1ab | DKGDAEDTLNKLSKY | 1751 | 1765 | Negative |
| hCoV229E | orf1ab | DMYPEFSIVCRFDTR | 5975 | 5989 | Negative |
| hCoV229E | orf1ab | DNEIIVKPNISLCVP | 640 | 654 | Negative |
| hCoV229E | orf1ab | DRGLALDNALSGVLS | 1495 | 1509 | Negative |
| hCoV229E | orf1ab | DSDINDMVLSLVRSG | 6723 | 6737 | Negative |
| hCoV229E | orf1ab | DSELLSTLSVDFNGV | 2317 | 2331 | Negative |
| hCoV229E | orf1ab | DSGVVTVELEPPCRF | 3881 | 3895 | Negative |
| hCoV229E | orf1ab | DTVVNGVDFDFIVNA | 1289 | 1303 | Negative |
| hCoV229E | orf1ab | DVDTVVNGVDFDFIV | 1287 | 1301 | Negative |
| hCoV229E | orf1ab | DVGLCMYRLLKDLHT | 1103 | 1117 | Negative |
| hCoV229E | orf1ab | DVKIMAKAPQRTGVV | 1955 | 1969 | Negative |
| hCoV229E | orf1ab | DYVLLDNALVKLVTT | 492 | 506 | Negative |
| hCoV229E | orf1ab | DYVSLLVMLLCAISN | 3419 | 3433 | Negative |
| hCoV229E | orf1ab | ECVDSNAGVCFGFDK | 2699 | 2713 | Negative |
| hCoV229E | orf1ab | EFVVKEKLNAFLVHG | 1265 | 1278 | Negative |
| hCoV229E | orf1ab | EFYDFLRSQGFFDEG | 4481 | 4495 | Negative |
| hCoV229E | orf1ab | EGNALYNNEGGRAFM | 3849 | 3863 | Negative |
| hCoV229E | orf1ab | EGVNGGSLYVNNHAF | 5997 | 6011 | Negative |

| | | | | | |
|---|---|---|---|---|---|
| hCoV229E | orf1ab | EGYTVVIGDVAYFAS | 556 | 570 | Negative |
| hCoV229E | orf1ab | EIEYVHGDALHTLRN | 176 | 190 | Negative |
| hCoV229E | orf1ab | EILVWPLILTCERVV | 3807 | 3821 | Negative |
| hCoV229E | orf1ab | EIVGPKELLLLWESG | 5151 | 5165 | Negative |
| hCoV229E | orf1ab | EIVNTVAGIPSNVYL | 2579 | 2593 | Negative |
| hCoV229E | orf1ab | EIVNTVSELVYENKF | 5443 | 5457 | Negative |
| hCoV229E | orf1ab | ELAQVLTEVVYSNGG | 4721 | 4735 | Negative |
| hCoV229E | orf1ab | ELFPHNDRIKSFSTF | 816 | 830 | Negative |
| hCoV229E | orf1ab | ELFVYTTTIWVNPGF | 3291 | 3305 | Negative |
| hCoV229E | orf1ab | ELGNITKTNVQPTGP | 2221 | 2235 | Negative |
| hCoV229E | orf1ab | ELGYIADISAFKATL | 4836 | 4849 | Negative |
| hCoV229E | orf1ab | ELLLLWESGKAKPPL | 5157 | 5171 | Negative |
| hCoV229E | orf1ab | ELLQFVTDPTLIVAS | 4433 | 4447 | Negative |
| hCoV229E | orf1ab | ELLSTLSVDFNGVLH | 2319 | 2333 | Negative |
| hCoV229E | orf1ab | ELLTKAIDVDWVEFY | 1651 | 1665 | Negative |
| hCoV229E | orf1ab | ELMEGSLPYSSIQAN | 2648 | 2661 | Negative |
| hCoV229E | orf1ab | ELPVAISGDKVFVRD | 6131 | 6145 | Negative |
| hCoV229E | orf1ab | ELTLKHFFFTQKGDA | 4497 | 4511 | Negative |
| hCoV229E | orf1ab | ELTTMRYFVKIGAVK | 5783 | 5797 | Negative |
| hCoV229E | orf1ab | ELVQRFSFWTMFCTS | 6639 | 6653 | Negative |
| hCoV229E | orf1ab | ELVRFVKSICNSAVA | 428 | 442 | Negative |
| hCoV229E | orf1ab | EMVLKGVVGVLTLDN | 4257 | 4271 | Negative |
| hCoV229E | orf1ab | EVIIDNKPWRWMLWC | 6431 | 6445 | Negative |
| hCoV229E | orf1ab | EVLFNKYFKYWGQDY | 4343 | 4357 | Negative |
| hCoV229E | orf1ab | EVSMCTNYDLSVINQ | 5371 | 5385 | Negative |
| hCoV229E | orf1ab | EVSRELGNITKTNVQ | 2217 | 2231 | Negative |
| hCoV229E | orf1ab | EVTMTDLQSESSCGL | 5585 | 5599 | Negative |
| hCoV229E | orf1ab | EVVVTNLNKSAGWPL | 4555 | 4569 | Negative |
| hCoV229E | orf1ab | FDCYEGGCITSREVV | 4543 | 4557 | Negative |
| hCoV229E | orf1ab | FDFGDFLVHNFVIFF | 1923 | 1937 | Negative |
| hCoV229E | orf1ab | FDITESKYSCKEVFK | 2263 | 2277 | Negative |
| hCoV229E | orf1ab | FDPIENEDIHRVYAA | 4225 | 4239 | Negative |
| hCoV229E | orf1ab | FDVFVTTIQTVFDCA | 462 | 476 | Negative |
| hCoV229E | orf1ab | FDVYDDIFAESKPWF | 386 | 400 | Negative |
| hCoV229E | orf1ab | FDYNVSVMQMDIQGF | 3361 | 3375 | Negative |
| hCoV229E | orf1ab | FEGDKFVGTFESAAA | 2871 | 2885 | Negative |
| hCoV229E | orf1ab | FEGYDFKYIENGQLK | 2523 | 2537 | Negative |
| hCoV229E | orf1ab | FERGSVQVDNGSSIN | 5471 | 5485 | Negative |
| hCoV229E | orf1ab | FFDEGSELTLKHFFF | 4491 | 4505 | Negative |
| hCoV229E | orf1ab | FGENEEIVINGRTYV | 140 | 154 | Negative |
| hCoV229E | orf1ab | FGMLRRLDMSSVDNI | 3721 | 3735 | Negative |
| hCoV229E | orf1ab | FLNAFDVFVTTIQTV | 458 | 472 | Negative |
| hCoV229E | orf1ab | FRGAVSCGHYQTNIY | 1197 | 1211 | Negative |
| hCoV229E | orf1ab | FSDDVEVKDIEPVYR | 902 | 916 | Negative |
| hCoV229E | orf1ab | FSGPVDKGHYTVYDT | 1855 | 1869 | Negative |
| hCoV229E | orf1ab | FSNGVDFVAQPEGCV | 5703 | 5717 | Negative |
| hCoV229E | orf1ab | FTPLNKQSCPIVVGV | 2563 | 2577 | Negative |
| hCoV229E | orf1ab | FVKMMVDGFVHYAGV | 3767 | 3781 | Negative |
| hCoV229E | orf1ab | FVRHVLFGCENPDCI | 2145 | 2159 | Negative |

| | | | | | |
|---|---|---|---|---|---|
| hCoV229E | orf1ab | FVSLDLQDCIVGIAD | 44 | 58 | Negative |
| hCoV229E | orf1ab | GCDYVYNPYVIDIQQ | 5823 | 5837 | Negative |
| hCoV229E | orf1ab | GCNVTDVTKLYLGGL | 5049 | 5063 | Negative |
| hCoV229E | orf1ab | GDAEDTLNKLSKYLV | 1753 | 1766 | Negative |
| hCoV229E | orf1ab | GEGFNILACYDGCAQ | 3073 | 3087 | Negative |
| hCoV229E | orf1ab | GEQFTHIVPLLRKGQ | 5735 | 5749 | Negative |
| hCoV229E | orf1ab | GETFWRYNFDITESK | 2256 | 2269 | Negative |
| hCoV229E | orf1ab | GFNILACYDGCAQGV | 3075 | 3089 | Negative |
| hCoV229E | orf1ab | GGNVTYTDQYLCGAD | 110 | 124 | Negative |
| hCoV229E | orf1ab | GIFGVKLETSLEVLL | 1393 | 1407 | Negative |
| hCoV229E | orf1ab | GIGGPRTIKVSTVQS | 3533 | 3547 | Negative |
| hCoV229E | orf1ab | GIISEGNALYNNEGG | 3845 | 3859 | Negative |
| hCoV229E | orf1ab | GILAVIDEHVFYKKD | 862 | 876 | Negative |
| hCoV229E | orf1ab | GILDGDYAMQFFKMG | 1067 | 1081 | Negative |
| hCoV229E | orf1ab | GILSNMNIASNSKEW | 3561 | 3575 | Negative |
| hCoV229E | orf1ab | GINYLGDFAQGPFID | 6665 | 6679 | Negative |
| hCoV229E | orf1ab | GKAFDKVFDYVLLDN | 484 | 498 | Negative |
| hCoV229E | orf1ab | GKDLNANMSLADCKS | 2345 | 2358 | Negative |
| hCoV229E | orf1ab | GKHVDHYDGFYTQGR | 6295 | 6309 | Negative |
| hCoV229E | orf1ab | GKNVHLKDVTKENQE | 3793 | 3807 | Negative |
| hCoV229E | orf1ab | GKQILGYSSLNDEFS | 3239 | 3253 | Negative |
| hCoV229E | orf1ab | GKQTEFVSNSHLLTH | 3935 | 3949 | Negative |
| hCoV229E | orf1ab | GKTTSMFKSISLFAG | 3269 | 3283 | Negative |
| hCoV229E | orf1ab | GKVSFSDDVEVKDIE | 898 | 912 | Negative |
| hCoV229E | orf1ab | GKVVANAMLKCVAFC | 4241 | 4255 | Negative |
| hCoV229E | orf1ab | GLAKALDVYTKGKLQ | 1313 | 1327 | Negative |
| hCoV229E | orf1ab | GLALDNALSGVLSAA | 1497 | 1511 | Negative |
| hCoV229E | orf1ab | GLCLFWNCNVDMYPE | 5965 | 5979 | Negative |
| hCoV229E | orf1ab | GLCMYRLLKDLHTGF | 1105 | 1119 | Negative |
| hCoV229E | orf1ab | GLGDLVDSYFENDSI | 3613 | 3627 | Negative |
| hCoV229E | orf1ab | GLGGNNVYCYNTELM | 2635 | 2649 | Negative |
| hCoV229E | orf1ab | GLHGNQTLFCNIMKF | 64 | 78 | Negative |
| hCoV229E | orf1ab | GLKILKQLDNNCWVN | 1043 | 1057 | Negative |
| hCoV229E | orf1ab | GLTFKTYAFDKTYIV | 784 | 798 | Negative |
| hCoV229E | orf1ab | GLTFLLTINENQAVT | 2457 | 2471 | Negative |
| hCoV229E | orf1ab | GLTSQTVKPGHFNKE | 4467 | 4481 | Negative |
| hCoV229E | orf1ab | GLVCLIQFYLCFFMP | 2499 | 2513 | Negative |
| hCoV229E | orf1ab | GMACNRVTLAVASDS | 1 | 14 | Negative |
| hCoV229E | orf1ab | GMLNGNAIATVKSED | 6265 | 6279 | Negative |
| hCoV229E | orf1ab | GMPSFVAYETARQEY | 3637 | 3651 | Negative |
| hCoV229E | orf1ab | GNAGVCYDIFGVTTP | 2607 | 2621 | Negative |
| hCoV229E | orf1ab | GRVGNGYVCGTGLWN | 2719 | 2733 | Negative |
| hCoV229E | orf1ab | GSFVGADGELPVAIS | 6123 | 6137 | Negative |
| hCoV229E | orf1ab | GSGKSHCSIGIGVYY | 5281 | 5295 | Negative |
| hCoV229E | orf1ab | GSGSGQAITSTIDSN | 3981 | 3995 | Negative |
| hCoV229E | orf1ab | GTAFLGVVGATMHGV | 3035 | 3049 | Negative |
| hCoV229E | orf1ab | GTCAKRFKRWLGILL | 752 | 766 | Negative |
| hCoV229E | orf1ab | GTDHRFILAITPYVC | 5031 | 5045 | Negative |
| hCoV229E | orf1ab | GTDIDYCVRAFDVYN | 4091 | 4105 | Negative |

| | | | | | |
|---|---|---|---|---|---|
| hCoV229E | orf1ab | GTNDPIRFCLENTVC | 4035 | 4049 | Negative |
| hCoV229E | orf1ab | GVFLGYKETNWFLHF | 2109 | 2123 | Negative |
| hCoV229E | orf1ab | GVIKNVNYVRDWLKS | 726 | 740 | Negative |
| hCoV229E | orf1ab | GVLTLDNQDLNGNFY | 4265 | 4279 | Negative |
| hCoV229E | orf1ab | GVMEPIDYNVVTQRM | 5411 | 5425 | Negative |
| hCoV229E | orf1ab | GVMVECDSLRIFNVV | 1345 | 1359 | Negative |
| hCoV229E | orf1ab | GVMYGGFEDQPNLQV | 3143 | 3157 | Negative |
| hCoV229E | orf1ab | GVNKIQPWTNDALNT | 1221 | 1235 | Negative |
| hCoV229E | orf1ab | GVNLQSGKTTSMFKS | 3263 | 3277 | Negative |
| hCoV229E | orf1ab | GVNMRTNWTIRGSFI | 3091 | 3105 | Negative |
| hCoV229E | orf1ab | GVNVHDVTVTTDKSF | 1611 | 1625 | Negative |
| hCoV229E | orf1ab | GVQVGYCVHGIKYYS | 1807 | 1821 | Negative |
| hCoV229E | orf1ab | GVREGGLNKVKYATV | 510 | 524 | Negative |
| hCoV229E | orf1ab | GVSVWRVIALQSVDC | 312 | 326 | Negative |
| hCoV229E | orf1ab | GVTTPEKCIFTSACT | 2617 | 2631 | Negative |
| hCoV229E | orf1ab | GVVCYNKTYAELGYI | 4825 | 4839 | Negative |
| hCoV229E | orf1ab | GVVLKRSLKYNLKAS | 1967 | 1981 | Negative |
| hCoV229E | orf1ab | GVVTVELEPPCRFVI | 3883 | 3897 | Negative |
| hCoV229E | orf1ab | GWDYPKCDRAMPSMI | 4679 | 4693 | Negative |
| hCoV229E | orf1ab | GWLGMDVEGAHVTGD | 5677 | 5691 | Negative |
| hCoV229E | orf1ab | GYNLKNGEVEFVYMH | 3113 | 3127 | Negative |
| hCoV229E | orf1ab | HFSMMILSDDGVVCY | 4815 | 4829 | Negative |
| hCoV229E | orf1ab | HGIKYYSRVKSVRGR | 1815 | 1829 | Negative |
| hCoV229E | orf1ab | HIGLAGKVRVGTGVM | 1333 | 1347 | Negative |
| hCoV229E | orf1ab | HKAYIDVLRNSFGKD | 2333 | 2347 | Negative |
| hCoV229E | orf1ab | HLDNNVQRCTRKLNR | 1549 | 1563 | Negative |
| hCoV229E | orf1ab | HLSFPLCSAGNVFGL | 5073 | 5087 | Negative |
| hCoV229E | orf1ab | HSFRTLKSGEGFNIL | 3065 | 3079 | Negative |
| hCoV229E | orf1ab | HTPRHSFRTLKSGEG | 3061 | 3075 | Negative |
| hCoV229E | orf1ab | HYAKFGFTPLNKQSC | 2557 | 2571 | Negative |
| hCoV229E | orf1ab | IANNYSKLSDEGYTV | 546 | 560 | Negative |
| hCoV229E | orf1ab | IGAIIFRICRFGVAF | 3437 | 3451 | Negative |
| hCoV229E | orf1ab | ILKAEEFVAASDITL | 6373 | 6387 | Negative |
| hCoV229E | orf1ab | ILKVIDGGKIWNGVI | 714 | 728 | Negative |
| hCoV229E | orf1ab | IMRAAIKLYNPKAIH | 5907 | 5921 | Negative |
| hCoV229E | orf1ab | IRMLSAMILGSKHVT | 4693 | 4707 | Negative |
| hCoV229E | orf1ab | ISDMYDGRTKAIDGE | 6585 | 6599 | Negative |
| hCoV229E | orf1ab | IVVDNDVVDYVSDAD | 6551 | 6565 | Negative |
| hCoV229E | orf1ab | IVVGVSEIVNTVAGI | 2573 | 2587 | Negative |
| hCoV229E | orf1ab | KEGFFTYINGFICEK | 6603 | 6617 | Negative |
| hCoV229E | orf1ab | KLCFEFEDEKLVDVC | 918 | 932 | Negative |
| hCoV229E | orf1ab | KSGEGFNILACYDGC | 3071 | 3085 | Negative |
| hCoV229E | orf1ab | KVTEYSWNKKLYELV | 6627 | 6641 | Negative |
| hCoV229E | orf1ab | KYGLEDFNFEHVVYG | 6335 | 6349 | Negative |
| hCoV229E | orf1ab | LACYDGCAQGVFGVN | 3079 | 3093 | Negative |
| hCoV229E | orf1ab | LCFALRNFDEKDCGV | 4189 | 4203 | Negative |
| hCoV229E | orf1ab | LDDRGLALDNALSGV | 1493 | 1507 | Negative |
| hCoV229E | orf1ab | LDITDAAVKAAESKA | 688 | 702 | Negative |
| hCoV229E | orf1ab | LDNQDLNGNFYDFGD | 4269 | 4283 | Negative |

| | | | | | |
|---|---|---|---|---|---|
| hCoV229E | orf1ab | LDYSIDVIDNEIIVK | 632 | 646 | Negative |
| hCoV229E | orf1ab | LEMAKEVKTSSKVVL | 194 | 208 | Negative |
| hCoV229E | orf1ab | LGMDVEGAHVTGDNV | 5679 | 5693 | Negative |
| hCoV229E | orf1ab | LHLLISQVRLSKMGI | 6359 | 6373 | Negative |
| hCoV229E | orf1ab | LHNFSIISGTAFLGV | 3027 | 3041 | Negative |
| hCoV229E | orf1ab | LKCVAFCDEMVLKGV | 4249 | 4263 | Negative |
| hCoV229E | orf1ab | LKKAMNVAKAEFDRE | 3667 | 3681 | Negative |
| hCoV229E | orf1ab | LLDAEFVEPGCGGIL | 850 | 864 | Negative |
| hCoV229E | orf1ab | LLGTCAKRFKRWLGI | 750 | 764 | Negative |
| hCoV229E | orf1ab | LLYWINRFCKCTLGV | 3483 | 3497 | Negative |
| hCoV229E | orf1ab | LNLKYAISGKERART | 4605 | 4619 | Negative |
| hCoV229E | orf1ab | LNMDIGVFIQKYGLE | 6325 | 6339 | Negative |
| hCoV229E | orf1ab | LQGLENIAFNVVNKG | 6109 | 6123 | Negative |
| hCoV229E | orf1ab | LQSAEWKCGYSMPGI | 6457 | 6471 | Negative |
| hCoV229E | orf1ab | LTDLKCTNVVLMGIL | 3549 | 3563 | Negative |
| hCoV229E | orf1ab | LTGILDGDYAMQFFK | 1065 | 1079 | Negative |
| hCoV229E | orf1ab | LTIANNYSKLSDEGY | 544 | 558 | Negative |
| hCoV229E | orf1ab | LTLDDRGLALDNALS | 1491 | 1505 | Negative |
| hCoV229E | orf1ab | LTLDNQDLNGNFYDF | 4267 | 4281 | Negative |
| hCoV229E | orf1ab | LVGKTLIFTLQAAFG | 2593 | 2607 | Negative |
| hCoV229E | orf1ab | MADVDDPKLMGWDYP | 4669 | 4683 | Negative |
| hCoV229E | orf1ab | MAKEVKTSSKVVLSD | 196 | 210 | Negative |
| hCoV229E | orf1ab | MDIGVFIQKYGLEDF | 6327 | 6341 | Negative |
| hCoV229E | orf1ab | MDVEGAHVTGDNVGT | 5681 | 5695 | Negative |
| hCoV229E | orf1ab | MIDKVEFENGFYRLY | 2239 | 2253 | Negative |
| hCoV229E | orf1ab | MKVKATKGEGDGGII | 3833 | 3846 | Negative |
| hCoV229E | orf1ab | MLCTKCAYDHVFGTD | 5019 | 5033 | Negative |
| hCoV229E | orf1ab | MSSVDNILNMARNGV | 3729 | 3743 | Negative |
| hCoV229E | orf1ab | MTSNVRRQVASGVID | 366 | 380 | Negative |
| hCoV229E | orf1ab | MVDGFVHYAGVVWTL | 3771 | 3785 | Negative |
| hCoV229E | orf1ab | MVGDYVAPVSTVKPK | 1895 | 1909 | Negative |
| hCoV229E | orf1ab | NENQAVTQIPATSIV | 2465 | 2479 | Negative |
| hCoV229E | orf1ab | NEVVKQMFGVNLQSG | 3255 | 3269 | Negative |
| hCoV229E | orf1ab | NIYSQNLCVDGFGVN | 1209 | 1223 | Negative |
| hCoV229E | orf1ab | NNYSKLSDEGYTVVI | 548 | 562 | Negative |
| hCoV229E | orf1ab | PAYVMIDKVEFENGF | 2235 | 2249 | Negative |
| hCoV229E | orf1ab | PGDAVITTQQAGAGV | 282 | 296 | Negative |
| hCoV229E | orf1ab | PIKNTVDTIPKEEFV | 1253 | 1267 | Negative |
| hCoV229E | orf1ab | PPGSGKSHCSIGIGV | 5279 | 5293 | Negative |
| hCoV229E | orf1ab | PVCSSIFRGAVSCGH | 1191 | 1205 | Negative |
| hCoV229E | orf1ab | QGDVDTVVNGVDFDF | 1285 | 1299 | Negative |
| hCoV229E | orf1ab | QKHFSMMILSDDGVV | 4813 | 4827 | Negative |
| hCoV229E | orf1ab | QLGLVWNKDVNTHST | 4413 | 4427 | Negative |
| hCoV229E | orf1ab | QPFIVMVLLLIFGDN | 2077 | 2091 | Negative |
| hCoV229E | orf1ab | QTVFDCAVETCTIAG | 470 | 484 | Negative |
| hCoV229E | orf1ab | QVGYCVHGIKYYSRV | 1809 | 1823 | Negative |
| hCoV229E | orf1ab | QYLKPHFISQGLDAA | 1711 | 1725 | Negative |
| hCoV229E | orf1ab | RAFDVYNKDASFIGK | 4099 | 4113 | Negative |
| hCoV229E | orf1ab | RAYVESYNIFTQAGF | 6073 | 6087 | Negative |

| | | | | | |
|---|---|---|---|---|---|
| hCoV229E | orf1ab | RFDTRTRSTLNLEGV | 5985 | 5999 | Negative |
| hCoV229E | orf1ab | RFDVSMPGSHSLFCT | 5653 | 5667 | Negative |
| hCoV229E | orf1ab | RGSSAARLEPCNGTD | 4079 | 4093 | Negative |
| hCoV229E | orf1ab | RSTMEEDFLNMDIGV | 6317 | 6331 | Negative |
| hCoV229E | orf1ab | RTLFDALKFFEVTMT | 5575 | 5589 | Negative |
| hCoV229E | orf1ab | RVKLCFEFEDEKLVD | 916 | 930 | Negative |
| hCoV229E | orf1ab | SAFKATLYYQNGVFM | 4843 | 4857 | Negative |
| hCoV229E | orf1ab | SCGIFGVKLETSLEV | 1391 | 1405 | Negative |
| hCoV229E | orf1ab | SFDSSYLNRVRGSSA | 4069 | 4083 | Negative |
| hCoV229E | orf1ab | SFGKDLNANMSLADC | 2343 | 2357 | Negative |
| hCoV229E | orf1ab | SFNVSDAYANLVPYY | 5251 | 5265 | Negative |
| hCoV229E | orf1ab | SFSTFESAYMPIADP | 826 | 840 | Negative |
| hCoV229E | orf1ab | SKGVMEPIDYNVVTQ | 5409 | 5423 | Negative |
| hCoV229E | orf1ab | SKHPKPEYRKVFYAL | 4933 | 4947 | Negative |
| hCoV229E | orf1ab | SKLTDLKCTNVVLMG | 3547 | 3561 | Negative |
| hCoV229E | orf1ab | SLNDEFSINEVVKQM | 3247 | 3261 | Negative |
| hCoV229E | orf1ab | SSDVLVFVLWAGGLE | 5769 | 5783 | Negative |
| hCoV229E | orf1ab | SSKTVCTYMDLLLDD | 6399 | 6413 | Negative |
| hCoV229E | orf1ab | STGLTSQTVKPGHFN | 4465 | 4479 | Negative |
| hCoV229E | orf1ab | STGWFDVYDDIFAES | 382 | 396 | Negative |
| hCoV229E | orf1ab | SVNTSSSEAFVVGIN | 6653 | 6667 | Negative |
| hCoV229E | orf1ab | TAFGPLCRKVFIDGV | 4387 | 4401 | Negative |
| hCoV229E | orf1ab | TNVPLQVGFSNGVDF | 5695 | 5709 | Negative |
| hCoV229E | orf1ab | TQLNLKYAISGKERA | 4603 | 4617 | Negative |
| hCoV229E | orf1ab | TSELLTKAIDVDWVE | 1649 | 1663 | Negative |
| hCoV229E | orf1ab | TSTIDSNTTQDTYGG | 3989 | 4003 | Negative |
| hCoV229E | orf1ab | VAFLPVEYVSYFDGV | 3449 | 3463 | Negative |
| hCoV229E | orf1ab | VCTCYDNSIQGSYER | 6221 | 6235 | Negative |
| hCoV229E | orf1ab | VDSNAGVCFGFDKWF | 2701 | 2715 | Negative |
| hCoV229E | orf1ab | VEGAHVTGDNVGTNV | 5683 | 5697 | Negative |
| hCoV229E | orf1ab | VEHYNEWAQANGFTA | 3191 | 3205 | Negative |
| hCoV229E | orf1ab | VELLDAEFVEPGCGG | 848 | 862 | Negative |
| hCoV229E | orf1ab | VEMHNKINLCDDPET | 3579 | 3593 | Negative |
| hCoV229E | orf1ab | VFVDDVTKTDAVILL | 4905 | 4919 | Negative |
| hCoV229E | orf1ab | VGFSNGVDFVAQPEG | 5701 | 5715 | Negative |
| hCoV229E | orf1ab | VGTNVPLQVGFSNGV | 5693 | 5707 | Negative |
| hCoV229E | orf1ab | VGVSEIVNTVAGIPS | 2575 | 2589 | Negative |
| hCoV229E | orf1ab | VIASNTTSAIDYDHE | 3007 | 3021 | Negative |
| hCoV229E | orf1ab | VKATKGEGDGGIISE | 3835 | 3849 | Negative |
| hCoV229E | orf1ab | VKDIEPVYRVKLCFE | 908 | 922 | Negative |
| hCoV229E | orf1ab | VKHLNKTLNEGVLES | 4951 | 4965 | Negative |
| hCoV229E | orf1ab | VKSSRVERSNAVLTI | 532 | 546 | Negative |
| hCoV229E | orf1ab | VLDDFIVFNNNGTNV | 2281 | 2295 | Negative |
| hCoV229E | orf1ab | VLRNSFGKDLNANMS | 2339 | 2353 | Negative |
| hCoV229E | orf1ab | VMGLHGNQTLFCNIM | 62 | 76 | Negative |
| hCoV229E | orf1ab | VPLQVGFSNGVDFVA | 5697 | 5711 | Negative |
| hCoV229E | orf1ab | VSCYVNLPTYYIYDE | 960 | 974 | Negative |
| hCoV229E | orf1ab | VTKENQEILVWPLIL | 3801 | 3815 | Negative |
| hCoV229E | orf1ab | VVATYKFVLWDYEAE | 6187 | 6201 | Negative |

| | | | | | |
|---|---|---|---|---|---|
| hCoV229E | orf1ab | VVFYQGDVDTVVNGV | 1281 | 1295 | Negative |
| hCoV229E | orf1ab | VVGINYLGDFAQGPF | 6663 | 6677 | Negative |
| hCoV229E | orf1ab | VVGVLTLDNQDLNGN | 4263 | 4277 | Negative |
| hCoV229E | orf1ab | VVNGVDFDFIVNAAN | 1291 | 1305 | Negative |
| hCoV229E | orf1ab | VYRVKLCFEFEDEKL | 914 | 928 | Negative |
| hCoV229E | orf1ab | WAELFVYTTTIWVNP | 3289 | 3303 | Negative |
| hCoV229E | orf1ab | WYIGAIIFRICRFGV | 3435 | 3449 | Negative |
| hCoV229E | orf1ab | YAISGKERARTVGGV | 4609 | 4623 | Negative |
| hCoV229E | orf1ab | YDFGDFVLCPPGMGI | 4279 | 4293 | Negative |
| hCoV229E | orf1ab | YLPYPDPSRIISAGV | 4891 | 4905 | Negative |
| hCoV229E | orf1ab | YSKLSDEGYTVVIGD | 550 | 564 | Negative |
| hCoV229E | orf1ab | YTAEQCIRGAMGDVG | 1091 | 1105 | Negative |
| hCoV229E | orf1ab | YVMIDKVEFENGFYR | 2237 | 2251 | Negative |
| hCoV229E | orf1ab | YWINRFCKCTLGVYD | 3485 | 3499 | Negative |
| hCoV229E | orf1ab | AAEAKVITIKVTEDG | 1597 | 1611 | Overlapping |
| hCoV229E | orf1ab | AAETVKAKEESVKSS | 5129 | 5143 | Overlapping |
| hCoV229E | orf1ab | ADPTHFDIEEVELLD | 838 | 852 | Overlapping |
| hCoV229E | orf1ab | AGHSLTWLWLLCGLV | 2487 | 2501 | Overlapping |
| hCoV229E | orf1ab | AIDYDHEYSIMRLHN | 3015 | 3029 | Overlapping |
| hCoV229E | orf1ab | AITSTIDSNTTQDTY | 3987 | 4001 | Overlapping |
| hCoV229E | orf1ab | AVDPAAAYLDAVKQG | 3953 | 3967 | Overlapping |
| hCoV229E | orf1ab | AVDSLCAKAVTAYSV | 5309 | 5323 | Overlapping |
| hCoV229E | orf1ab | AVLTIANNYSKLSDE | 542 | 556 | Overlapping |
| hCoV229E | orf1ab | CFEFEDEKLVDVCEK | 920 | 934 | Overlapping |
| hCoV229E | orf1ab | CYEGGCITSREVVVT | 4545 | 4559 | Overlapping |
| hCoV229E | orf1ab | DAEFVEPGCGGILAV | 852 | 866 | Overlapping |
| hCoV229E | orf1ab | DAIVVDNDVVDYVSD | 6549 | 6563 | Overlapping |
| hCoV229E | orf1ab | DDFIVFNNNGTNVTQ | 2283 | 2297 | Overlapping |
| hCoV229E | orf1ab | DDGSCEVVHDQVNYV | 6033 | 6047 | Overlapping |
| hCoV229E | orf1ab | DEISVTPIKNTVDTI | 1248 | 1260 | Overlapping |
| hCoV229E | orf1ab | DFCRQYSNESWFEDD | 666 | 680 | Overlapping |
| hCoV229E | orf1ab | DFGLGDLVDSYFEND | 3611 | 3625 | Overlapping |
| hCoV229E | orf1ab | DGELPVAISGDKVFV | 6129 | 6143 | Overlapping |
| hCoV229E | orf1ab | DGENVSKEGFFTYIN | 6597 | 6611 | Overlapping |
| hCoV229E | orf1ab | DGFVHYAGVVWTLQE | 3773 | 3787 | Overlapping |
| hCoV229E | orf1ab | DGGIISEGNALYNNE | 3843 | 3857 | Overlapping |
| hCoV229E | orf1ab | DGKPVMSEDLWQFVD | 124 | 138 | Overlapping |
| hCoV229E | orf1ab | DHEFTSAISNAHRCD | 2367 | 2381 | Overlapping |
| hCoV229E | orf1ab | DHEYSIMRLHNFSII | 3019 | 3033 | Overlapping |
| hCoV229E | orf1ab | DHSAFAYESAVVNGI | 1677 | 1691 | Overlapping |
| hCoV229E | orf1ab | DHVFGTDHRFILAIT | 5027 | 5041 | Overlapping |
| hCoV229E | orf1ab | DHYDGFYTQGRNLQD | 6299 | 6313 | Overlapping |
| hCoV229E | orf1ab | DIEEVELLDAEFVEP | 844 | 858 | Overlapping |
| hCoV229E | orf1ab | DIETVDVKHEMSPFE | 1021 | 1035 | Overlapping |
| hCoV229E | orf1ab | DLWQFVDHFGENEEI | 132 | 146 | Overlapping |
| hCoV229E | orf1ab | DNEIIVKPNISLCVP | 640 | 654 | Overlapping |
| hCoV229E | orf1ab | DRESSVQKKINRMAE | 3679 | 3693 | Overlapping |
| hCoV229E | orf1ab | DVEIFVAFVYYVARL | 1733 | 1747 | Overlapping |
| hCoV229E | orf1ab | DYDHEYSIMRLHNFS | 3017 | 3031 | Overlapping |

| | | | | | |
|---|---|---|---|---|---|
| hCoV229E | orf1ab | DYFEMKNWFDPIENE | 4217 | 4231 | Overlapping |
| hCoV229E | orf1ab | DYVSDADFSVTGDCA | 6559 | 6573 | Overlapping |
| hCoV229E | orf1ab | EAERPLTSFTKSVCG | 6199 | 6213 | Overlapping |
| hCoV229E | orf1ab | EAFTKPVFISALVQC | 232 | 246 | Overlapping |
| hCoV229E | orf1ab | EAKVITIKVTEDGVN | 1599 | 1613 | Overlapping |
| hCoV229E | orf1ab | EAQVQLEHYSSCVEC | 1769 | 1783 | Overlapping |
| hCoV229E | orf1ab | EATHIAEDVVDQVEE | 999 | 1015 | Overlapping |
| hCoV229E | orf1ab | ECDSLRIFNVVGPRK | 1349 | 1363 | Overlapping |
| hCoV229E | orf1ab | ECRLAMLGAEMTSNV | 356 | 370 | Overlapping |
| hCoV229E | orf1ab | ECVDSNAGVCFGFDK | 2699 | 2713 | Overlapping |
| hCoV229E | orf1ab | ECYSGFKPNNTSAQY | 5337 | 5351 | Overlapping |
| hCoV229E | orf1ab | EDAFSILAAKTGVCV | 3209 | 3223 | Overlapping |
| hCoV229E | orf1ab | EDDYKAFISVLDITD | 678 | 692 | Overlapping |
| hCoV229E | orf1ab | EDEKLVDVCEKAIGK | 924 | 938 | Overlapping |
| hCoV229E | orf1ab | EDFNFEHVVYGDVSK | 6339 | 6353 | Overlapping |
| hCoV229E | orf1ab | EDGVNVHDVTVTTDK | 1609 | 1623 | Overlapping |
| hCoV229E | orf1ab | EDIHRVYAALGKVVA | 4231 | 4245 | Overlapping |
| hCoV229E | orf1ab | EDVCTCYDNSIQGSY | 6219 | 6233 | Overlapping |
| hCoV229E | orf1ab | EDVQQAQQEATHIAE | 992 | 1007 | Overlapping |
| hCoV229E | orf1ab | EDVVDQVEEVNSIFD | 1007 | 1021 | Overlapping |
| hCoV229E | orf1ab | EEDFLNMDIGVFIQK | 6321 | 6335 | Overlapping |
| hCoV229E | orf1ab | EEDLSIGPHEFCSQH | 4865 | 4879 | Overlapping |
| hCoV229E | orf1ab | EEFVAASDITLKCCT | 6377 | 6391 | Overlapping |
| hCoV229E | orf1ab | EEGGNDLSLPVMISE | 974 | 988 | Overlapping |
| hCoV229E | orf1ab | EEHVNRMDTFCFNVR | 336 | 350 | Overlapping |
| hCoV229E | orf1ab | EEIVINGRTYVCAWL | 144 | 158 | Overlapping |
| hCoV229E | orf1ab | EELNGLKILKQLDNN | 1039 | 1053 | Overlapping |
| hCoV229E | orf1ab | EEQDAIFLLTKRNIL | 4585 | 4599 | Overlapping |
| hCoV229E | orf1ab | EESVKSSYAYATLKE | 5137 | 5151 | Overlapping |
| hCoV229E | orf1ab | EEVELLDAEFVEPGC | 846 | 860 | Overlapping |
| hCoV229E | orf1ab | EEVKSSRVERSNAVL | 530 | 544 | Overlapping |
| hCoV229E | orf1ab | EFDRESSVQKKINRM | 3677 | 3691 | Overlapping |
| hCoV229E | orf1ab | EFDVVFGKRGGGNVT | 100 | 114 | Overlapping |
| hCoV229E | orf1ab | EFEDEKLVDVCEKAI | 922 | 936 | Overlapping |
| hCoV229E | orf1ab | EFSINEVVKQMFGVN | 3251 | 3265 | Overlapping |
| hCoV229E | orf1ab | EFSIVCRFDTRTRST | 5979 | 5993 | Overlapping |
| hCoV229E | orf1ab | EFTSAISNAHRCDVL | 2369 | 2383 | Overlapping |
| hCoV229E | orf1ab | EFVEPGCGGILAVID | 854 | 868 | Overlapping |
| hCoV229E | orf1ab | EFVSNSHLLTHCAFA | 3939 | 3953 | Overlapping |
| hCoV229E | orf1ab | EFYGFKDAVTFATVD | 1663 | 1677 | Overlapping |
| hCoV229E | orf1ab | EGDGGIISEGNALYN | 3841 | 3855 | Overlapping |
| hCoV229E | orf1ab | EGGCITSREVVVTNL | 4547 | 4561 | Overlapping |
| hCoV229E | orf1ab | EGGRAFMYAYVTTKP | 3857 | 3871 | Overlapping |
| hCoV229E | orf1ab | EGNALYNNEGGRAFM | 3849 | 3863 | Overlapping |
| hCoV229E | orf1ab | EGVNGGSLYVNNHAF | 5997 | 6011 | Overlapping |
| hCoV229E | orf1ab | EGYTVVIGDVAYFAS | 556 | 570 | Overlapping |
| hCoV229E | orf1ab | EHDSGVVTVELEPPC | 3879 | 3893 | Overlapping |
| hCoV229E | orf1ab | EHESKFWDESFYASM | 4973 | 4987 | Overlapping |
| hCoV229E | orf1ab | EHYSSCVECDAKFKN | 1775 | 1789 | Overlapping |

| | | | | | |
|---|---|---|---|---|---|
| hCoV229E | orf1ab | EIEYVHGDALHTLRN | 176 | 190 | Overlapping |
| hCoV229E | orf1ab | EIFVAFVYYYVARLVK | 1735 | 1749 | Overlapping |
| hCoV229E | orf1ab | EIIVKPNISLCVPLY | 642 | 656 | Overlapping |
| hCoV229E | orf1ab | EILVWPLILTCERVV | 3807 | 3821 | Overlapping |
| hCoV229E | orf1ab | EIMPGKMKVKATKGE | 3827 | 3841 | Overlapping |
| hCoV229E | orf1ab | EIVGPKELLLLWESG | 5151 | 5165 | Overlapping |
| hCoV229E | orf1ab | EIVNTVAGIPSNVYL | 2579 | 2593 | Overlapping |
| hCoV229E | orf1ab | EIVNTVSELVYENKF | 5443 | 5457 | Overlapping |
| hCoV229E | orf1ab | EKCVVRVCYGNTVLN | 2979 | 2993 | Overlapping |
| hCoV229E | orf1ab | EKDKHLDNNVQRCTR | 1545 | 1559 | Overlapping |
| hCoV229E | orf1ab | EKFDYNVSVMQMDIQ | 3359 | 3373 | Overlapping |
| hCoV229E | orf1ab | EKFLNAFDVFVTTIQ | 456 | 470 | Overlapping |
| hCoV229E | orf1ab | EKLKSYAASYNRYKY | 2905 | 2919 | Overlapping |
| hCoV229E | orf1ab | EKLSAYDLACCMRAG | 2401 | 2415 | Overlapping |
| hCoV229E | orf1ab | EKSTVLQAAGLCVVC | 4989 | 5003 | Overlapping |
| hCoV229E | orf1ab | ELFPHNDRIKSFSTF | 816 | 830 | Overlapping |
| hCoV229E | orf1ab | ELFVYTTTIWVNPGF | 3291 | 3305 | Overlapping |
| hCoV229E | orf1ab | ELGNITKTNVQPTGP | 2221 | 2235 | Overlapping |
| hCoV229E | orf1ab | ELGYIADISAFKATL | 4836 | 4849 | Overlapping |
| hCoV229E | orf1ab | ELLLLWESGKAKPPL | 5157 | 5171 | Overlapping |
| hCoV229E | orf1ab | ELLQFVTDPTLIVAS | 4433 | 4447 | Overlapping |
| hCoV229E | orf1ab | ELLSTLSVDFNGVLH | 2319 | 2333 | Overlapping |
| hCoV229E | orf1ab | ELLTKAIDVDWVEFY | 1651 | 1665 | Overlapping |
| hCoV229E | orf1ab | ELMEGSLPYSSIQAN | 2648 | 2661 | Overlapping |
| hCoV229E | orf1ab | ELPVAISGDKVFVRD | 6131 | 6145 | Overlapping |
| hCoV229E | orf1ab | ELTLKHFFFTQKGDA | 4497 | 4511 | Overlapping |
| hCoV229E | orf1ab | ELTTMRYFVKIGAVK | 5783 | 5797 | Overlapping |
| hCoV229E | orf1ab | ELVQRFSFWTMFCTS | 6639 | 6653 | Overlapping |
| hCoV229E | orf1ab | ELVRFVKSICNSAVA | 428 | 442 | Overlapping |
| hCoV229E | orf1ab | EMCILHCSNFNTLFA | 4367 | 4381 | Overlapping |
| hCoV229E | orf1ab | EMPFEELNGLKILKQ | 1035 | 1049 | Overlapping |
| hCoV229E | orf1ab | EMVLKGVVGVLTLDN | 4257 | 4271 | Overlapping |
| hCoV229E | orf1ab | ENAINKGGRTVQSHI | 5893 | 5907 | Overlapping |
| hCoV229E | orf1ab | ENAQKFFDFGDFLVH | 1917 | 1931 | Overlapping |
| hCoV229E | orf1ab | ENDSILQSVASSFVG | 3623 | 3637 | Overlapping |
| hCoV229E | orf1ab | ENEDIHRVYAALGKV | 4229 | 4243 | Overlapping |
| hCoV229E | orf1ab | ENEEIVINGRTYVCA | 142 | 156 | Overlapping |
| hCoV229E | orf1ab | ENFEDWHYAKFGFTP | 2551 | 2565 | Overlapping |
| hCoV229E | orf1ab | ENIAFNVVNKGSFVG | 6113 | 6127 | Overlapping |
| hCoV229E | orf1ab | ENLESAVLFVNDKIT | 614 | 628 | Overlapping |
| hCoV229E | orf1ab | ENTVCKVCGCWLNHG | 4045 | 4059 | Overlapping |
| hCoV229E | orf1ab | ENVSKEGFFTYINGF | 6599 | 6613 | Overlapping |
| hCoV229E | orf1ab | EPCVQSRLLSGVAYT | 1839 | 1853 | Overlapping |
| hCoV229E | orf1ab | EPVYRVKLCFEFEDE | 912 | 926 | Overlapping |
| hCoV229E | orf1ab | EQAAAAMYKEARAVN | 3693 | 3707 | Overlapping |
| hCoV229E | orf1ab | EQGTPLTPILSCGIF | 1381 | 1395 | Overlapping |
| hCoV229E | orf1ab | EQKPVSLIKVAPKPY | 1447 | 1461 | Overlapping |
| hCoV229E | orf1ab | EQSMYNLLKGCNAVA | 4145 | 4159 | Overlapping |
| hCoV229E | orf1ab | ERARTVGGVSLLATM | 4615 | 4629 | Overlapping |

| | | | | | |
|---|---|---|---|---|---|
| hCoV229E | orf1ab | ERCTHWCTYLFSLIA | 3395 | 3409 | Overlapping |
| hCoV229E | orf1ab | ERDLLVKAYNTINNE | 1367 | 1381 | Overlapping |
| hCoV229E | orf1ab | ERPLTSFTKSVCGYT | 6201 | 6215 | Overlapping |
| hCoV229E | orf1ab | ERSNAVLTIANNYSK | 538 | 552 | Overlapping |
| hCoV229E | orf1ab | ESAAAGTFVIDIRSY | 2881 | 2895 | Overlapping |
| hCoV229E | orf1ab | ESFYASMYEKSTVLQ | 4981 | 4995 | Overlapping |
| hCoV229E | orf1ab | ESGAVLFCTPTKKAF | 1135 | 1149 | Overlapping |
| hCoV229E | orf1ab | ESGKAKPPLNRNSVF | 5163 | 5177 | Overlapping |
| hCoV229E | orf1ab | ESKFWDESFYASMYE | 4975 | 4989 | Overlapping |
| hCoV229E | orf1ab | ESKYSCKEVFKNCNV | 2267 | 2281 | Overlapping |
| hCoV229E | orf1ab | ESLRLFAAETVKAKE | 5123 | 5137 | Overlapping |
| hCoV229E | orf1ab | ESSVQKKINRMAEQA | 3681 | 3695 | Overlapping |
| hCoV229E | orf1ab | ESWFEDDYKAFISVL | 674 | 688 | Overlapping |
| hCoV229E | orf1ab | ESYNIFTQAGFNIWV | 6077 | 6091 | Overlapping |
| hCoV229E | orf1ab | ETARQEYENAVANGS | 3645 | 3659 | Overlapping |
| hCoV229E | orf1ab | ETCTIAGKAFDKVFD | 478 | 492 | Overlapping |
| hCoV229E | orf1ab | ETVDVKHEMSPFEMP | 1023 | 1037 | Overlapping |
| hCoV229E | orf1ab | ETVKAKEESVKSSYA | 5131 | 5145 | Overlapping |
| hCoV229E | orf1ab | EVIIDNKPWRWMLWC | 6431 | 6445 | Overlapping |
| hCoV229E | orf1ab | EVKDNDGKNVHLKDV | 3787 | 3801 | Overlapping |
| hCoV229E | orf1ab | EVLFNKYFKYWGQDY | 4343 | 4357 | Overlapping |
| hCoV229E | orf1ab | EVNADIVVVDEVSMC | 5361 | 5375 | Overlapping |
| hCoV229E | orf1ab | EVNSIFDIETVDVKH | 1015 | 1029 | Overlapping |
| hCoV229E | orf1ab | EVSMCTNYDLSVINQ | 5371 | 5385 | Overlapping |
| hCoV229E | orf1ab | EVSRELGNITKTNVQ | 2217 | 2231 | Overlapping |
| hCoV229E | orf1ab | EVTMTDLQSESSCGL | 5585 | 5599 | Overlapping |
| hCoV229E | orf1ab | EVVVTNLNKSAGWPL | 4555 | 4569 | Overlapping |
| hCoV229E | orf1ab | EWKCGYSMPGIYKTQ | 6461 | 6475 | Overlapping |
| hCoV229E | orf1ab | EWPLFESDKEEDVQQ | 988 | 999 | Overlapping |
| hCoV229E | orf1ab | EYRKVFYALLDWVKH | 4939 | 4953 | Overlapping |
| hCoV229E | orf1ab | EYSIMRLHNFSIISG | 3021 | 3035 | Overlapping |
| hCoV229E | orf1ab | EYVHGDALHTLRNGS | 178 | 192 | Overlapping |
| hCoV229E | orf1ab | EYVSYFDGVKTVLLF | 3455 | 3469 | Overlapping |
| hCoV229E | orf1ab | FAEDVCTCYDNSIQG | 6217 | 6231 | Overlapping |
| hCoV229E | orf1ab | FDCYNLWQTFTDVNL | 6095 | 6109 | Overlapping |
| hCoV229E | orf1ab | FDGVMYGGFEDQPNL | 3141 | 3155 | Overlapping |
| hCoV229E | orf1ab | FDPIENEDIHRVYAA | 4225 | 4239 | Overlapping |
| hCoV229E | orf1ab | FEMKNWFDPIENEDI | 4219 | 4233 | Overlapping |
| hCoV229E | orf1ab | FESDKEEDVQQAQQE | 987 | 999 | Overlapping |
| hCoV229E | orf1ab | FGGKQILGYSSLNDE | 3237 | 3251 | Overlapping |
| hCoV229E | orf1ab | FLPVEYVSYFDGVKT | 3451 | 3465 | Overlapping |
| hCoV229E | orf1ab | FNKLSTSDWSDIRDY | 5101 | 5115 | Overlapping |
| hCoV229E | orf1ab | FVGELKAAEAKVITI | 1591 | 1605 | Overlapping |
| hCoV229E | orf1ab | FVNDKITEFQLDYSI | 622 | 636 | Overlapping |
| hCoV229E | orf1ab | FVPVKEASKQCFKIF | 5457 | 5471 | Overlapping |
| hCoV229E | orf1ab | GEFVFEKVDYGSDTV | 5191 | 5205 | Overlapping |
| hCoV229E | orf1ab | GEVEFVYMHQIELGS | 3119 | 3133 | Overlapping |
| hCoV229E | orf1ab | GLAGKVRVGTGVMVE | 1335 | 1349 | Overlapping |
| hCoV229E | orf1ab | GLDAAWNKFVLGDVE | 1721 | 1735 | Overlapping |

| | | | | | |
|---|---|---|---|---|---|
| hCoV229E | orf1ab | GLEDFNFEHVVYGDV | 6337 | 6351 | Overlapping |
| hCoV229E | orf1ab | GLELTTMRYFVKIGA | 5781 | 5795 | Overlapping |
| hCoV229E | orf1ab | GLENIAFNVVNKGSF | 6111 | 6125 | Overlapping |
| hCoV229E | orf1ab | GLQTQTVDSAQGSEY | 5523 | 5537 | Overlapping |
| hCoV229E | orf1ab | GLVWNKDVNTHSTRL | 4415 | 4429 | Overlapping |
| hCoV229E | orf1ab | GMPSFVAYETARQEY | 3637 | 3651 | Overlapping |
| hCoV229E | orf1ab | GPDVFLHKCYRCPAE | 5429 | 5443 | Overlapping |
| hCoV229E | orf1ab | GPVDKGHYTVYDTAK | 1857 | 1871 | Overlapping |
| hCoV229E | orf1ab | GTGVMVECDSLRIFN | 1343 | 1357 | Overlapping |
| hCoV229E | orf1ab | GTPLTPILSCGIFGV | 1383 | 1397 | Overlapping |
| hCoV229E | orf1ab | GVAYTAFSGPVDKGH | 1849 | 1863 | Overlapping |
| hCoV229E | orf1ab | GVLESFSVTLLDEHE | 4961 | 4975 | Overlapping |
| hCoV229E | orf1ab | GYTDFAEDVCTCYDN | 6213 | 6227 | Overlapping |
| hCoV229E | orf1ab | HEMSPFEMPFEELNG | 1029 | 1043 | Overlapping |
| hCoV229E | orf1ab | HFDIEEVELLDAEFV | 842 | 856 | Overlapping |
| hCoV229E | orf1ab | HRNEHVASGDAIMTR | 5855 | 5869 | Overlapping |
| hCoV229E | orf1ab | HVDHYDGFYTQGRNL | 6298 | 6311 | Overlapping |
| hCoV229E | orf1ab | IAEDVVDQVEEVNSI | 1005 | 1019 | Overlapping |
| hCoV229E | orf1ab | IDVIDNEIIVKPNIS | 636 | 650 | Overlapping |
| hCoV229E | orf1ab | IEEIEYVHGDALHTL | 174 | 188 | Overlapping |
| hCoV229E | orf1ab | ISEGNALYNNEGGRA | 3848 | 3861 | Overlapping |
| hCoV229E | orf1ab | ISEWPLFESDKEEDV | 986 | 999 | Overlapping |
| hCoV229E | orf1ab | IVVDNDVVDYVSDAD | 6551 | 6565 | Overlapping |
| hCoV229E | orf1ab | KAEEFVAASDITLKC | 6375 | 6389 | Overlapping |
| hCoV229E | orf1ab | KAEFDRESSVQKKIN | 3675 | 3689 | Overlapping |
| hCoV229E | orf1ab | KDGKHVDHYDGFYTQ | 6293 | 6307 | Overlapping |
| hCoV229E | orf1ab | KEEDVQQAQQEATHI | 992 | 1001 | Overlapping |
| hCoV229E | orf1ab | KEEFVVKEKLNAFLV | 1263 | 1277 | Overlapping |
| hCoV229E | orf1ab | KENQEILVWPLILTC | 3803 | 3817 | Overlapping |
| hCoV229E | orf1ab | KGEGDGGIISEGNAL | 3839 | 3853 | Overlapping |
| hCoV229E | orf1ab | KLSDEGYTVVIGDVA | 552 | 566 | Overlapping |
| hCoV229E | orf1ab | KNCNVLDDFIVFNNN | 2277 | 2291 | Overlapping |
| hCoV229E | orf1ab | KNGEVEFVYMHQIEL | 3117 | 3131 | Overlapping |
| hCoV229E | orf1ab | KTLEYDYMTHGQMEG | 5951 | 5965 | Overlapping |
| hCoV229E | orf1ab | KVDYGSDTVTYKSTA | 5197 | 5211 | Overlapping |
| hCoV229E | orf1ab | KWEHDSGVVTVELEP | 3877 | 3891 | Overlapping |
| hCoV229E | orf1ab | LDYKRQNNLAIEEIE | 164 | 178 | Overlapping |
| hCoV229E | orf1ab | LEESGAVLFCTPTKK | 1133 | 1147 | Overlapping |
| hCoV229E | orf1ab | LTCERVVKLQNNEIM | 3815 | 3829 | Overlapping |
| hCoV229E | orf1ab | LVDSELLSTLSVDFN | 2315 | 2329 | Overlapping |
| hCoV229E | orf1ab | MVECDSLRIFNVVGP | 1347 | 1361 | Overlapping |
| hCoV229E | orf1ab | NDEFSINEVVKQMFG | 3249 | 3263 | Overlapping |
| hCoV229E | orf1ab | NDGRVGNGYVCGTGL | 2717 | 2731 | Overlapping |
| hCoV229E | orf1ab | NEGVLESFSVTLLDE | 4959 | 4973 | Overlapping |
| hCoV229E | orf1ab | NEWAQANGFTAMNGE | 3195 | 3209 | Overlapping |
| hCoV229E | orf1ab | NFDEKDCGVFKEILV | 4195 | 4209 | Overlapping |
| hCoV229E | orf1ab | NGEDAFSILAAKTGV | 3207 | 3221 | Overlapping |
| hCoV229E | orf1ab | NGVDFDFIVNAANEN | 1293 | 1307 | Overlapping |
| hCoV229E | orf1ab | NGVDFVAQPEGCVLT | 5705 | 5719 | Overlapping |

| | | | | | |
|---|---|---|---|---|---|
| hCoV229E | orf1ab | NKQSCPIVVGVSEIV | 2567 | 2581 | Overlapping |
| hCoV229E | orf1ab | NLMADVDDPKLMGWD | 4667 | 4681 | Overlapping |
| hCoV229E | orf1ab | NLYNYGAGLKLPSGI | 6483 | 6497 | Overlapping |
| hCoV229E | orf1ab | NMLKNLMADVDDPKL | 4663 | 4677 | Overlapping |
| hCoV229E | orf1ab | NMRVLHLGAGSDYGV | 6521 | 6535 | Overlapping |
| hCoV229E | orf1ab | NNFNVKKLQRQLYDN | 4779 | 4793 | Overlapping |
| hCoV229E | orf1ab | NNYSKLSDEGYTVVI | 548 | 562 | Overlapping |
| hCoV229E | orf1ab | NTELMEGSLPYSSIQ | 2645 | 2659 | Overlapping |
| hCoV229E | orf1ab | PAEIVNTVSELVYEN | 5441 | 5455 | Overlapping |
| hCoV229E | orf1ab | PEEKLSAYDLACCMR | 2399 | 2413 | Overlapping |
| hCoV229E | orf1ab | PEPVNPVSFVVKPVC | 1179 | 1193 | Overlapping |
| hCoV229E | orf1ab | PETAQELLLALLAFF | 3591 | 3605 | Overlapping |
| hCoV229E | orf1ab | PFDVICDELFVTVIV | 2125 | 2139 | Overlapping |
| hCoV229E | orf1ab | PFEELNGLKILKQLD | 1037 | 1051 | Overlapping |
| hCoV229E | orf1ab | PFEMPFEELNGLKIL | 1033 | 1047 | Overlapping |
| hCoV229E | orf1ab | PGMKYVKWEHDSGVV | 3871 | 3885 | Overlapping |
| hCoV229E | orf1ab | PIADPTHFDIEEVEL | 836 | 850 | Overlapping |
| hCoV229E | orf1ab | PIDLPPSHATTYLSL | 5607 | 5621 | Overlapping |
| hCoV229E | orf1ab | PIDYNVVTQRMCAIG | 5415 | 5429 | Overlapping |
| hCoV229E | orf1ab | PIENEDIHRVYAALG | 4227 | 4241 | Overlapping |
| hCoV229E | orf1ab | PIGTNDPIRFCLENT | 4033 | 4047 | Overlapping |
| hCoV229E | orf1ab | PLFESDKEEDVQQAQ | 983 | 997 | Overlapping |
| hCoV229E | orf1ab | PLNKQSCPIVVGVSE | 2565 | 2579 | Overlapping |
| hCoV229E | orf1ab | PSFVAYETARQEYEN | 3639 | 3653 | Overlapping |
| hCoV229E | orf1ab | PTHFDIEEVELLDAE | 840 | 854 | Overlapping |
| hCoV229E | orf1ab | PTMDGFCQYKGKWVQ | 4017 | 4031 | Overlapping |
| hCoV229E | orf1ab | PVAISGDKVFVRDGN | 6133 | 6147 | Overlapping |
| hCoV229E | orf1ab | PVEYVSYFDGVKTVL | 3453 | 3467 | Overlapping |
| hCoV229E | orf1ab | PVKEASKQCFKIFER | 5459 | 5473 | Overlapping |
| hCoV229E | orf1ab | PVMGMTNCLASECFM | 4305 | 4319 | Overlapping |
| hCoV229E | orf1ab | PVMTNGSNILEAFTK | 222 | 236 | Overlapping |
| hCoV229E | orf1ab | PVNPVSFVVKPVCSS | 1181 | 1195 | Overlapping |
| hCoV229E | orf1ab | QLEHYSSCVECDAKF | 1773 | 1787 | Overlapping |
| hCoV229E | orf1ab | RVECYSGFKPNNTSA | 5335 | 5349 | Overlapping |
| hCoV229E | orf1ab | RVERSNAVLTIANNY | 536 | 550 | Overlapping |
| hCoV229E | orf1ab | SAVVNGIRVLKTSDN | 1685 | 1699 | Overlapping |
| hCoV229E | orf1ab | SDKEEDVQQAQQEAT | 987 | 1001 | Overlapping |
| hCoV229E | orf1ab | SEDLWQFVDHFGENE | 130 | 144 | Overlapping |
| hCoV229E | orf1ab | SRELGNITKTNVQPT | 2219 | 2233 | Overlapping |
| hCoV229E | orf1ab | SSKTVCTYMDLLLDD | 6399 | 6413 | Overlapping |
| hCoV229E | orf1ab | SSRVERSNAVLTIAN | 534 | 548 | Overlapping |
| hCoV229E | orf1ab | SSTFVEEEHVNRMDT | 330 | 344 | Overlapping |
| hCoV229E | orf1ab | SVLEMAKEVKTSSKV | 192 | 206 | Overlapping |
| hCoV229E | orf1ab | TDVNLQGLENIAFNV | 6105 | 6119 | Overlapping |
| hCoV229E | orf1ab | TEFQLDYSIDVIDNE | 628 | 642 | Overlapping |
| hCoV229E | orf1ab | TFVEEEHVNRMDTFC | 332 | 346 | Overlapping |
| hCoV229E | orf1ab | TGDCATVYLEDKFDL | 6569 | 6583 | Overlapping |
| hCoV229E | orf1ab | TGDNVGTNVPLQVGF | 5689 | 5703 | Overlapping |
| hCoV229E | orf1ab | TGPAYVMIDKVEFEN | 2233 | 2247 | Overlapping |

| | | | | | |
|---|---|---|---|---|---|
| hCoV229E | orf1ab | TMEEDFLNMDIGVFI | 6319 | 6333 | Overlapping |
| hCoV229E | orf1ab | TMMDLCFALRNFDEK | 4185 | 4199 | Overlapping |
| hCoV229E | orf1ab | VDNDVVDYVSDADFS | 6553 | 6567 | Overlapping |
| hCoV229E | orf1ab | VEEEHVNRMDTFCFN | 334 | 348 | Overlapping |
| hCoV229E | orf1ab | VFENFEDWHYAKFGF | 2549 | 2563 | Overlapping |
| hCoV229E | orf1ab | VGECVDSNAGVCFGF | 2697 | 2711 | Overlapping |
| hCoV229E | orf1ab | VIDNEIIVKPNISLC | 638 | 652 | Overlapping |
| hCoV229E | orf1ab | VIRDIVCKVENKAEA | 798 | 812 | Overlapping |
| hCoV229E | orf1ab | VKHEMSPFEMPFEEL | 1027 | 1041 | Overlapping |
| hCoV229E | orf1ab | VKRDGVQVGYCVHGI | 1803 | 1817 | Overlapping |
| hCoV229E | orf1ab | VLKTSDNNCWVNAVC | 1693 | 1707 | Overlapping |
| hCoV229E | orf1ab | VMDYKCSCTSGRLEE | 1121 | 1135 | Overlapping |
| hCoV229E | orf1ab | VMISEWPLFESDKEE | 984 | 999 | Overlapping |
| hCoV229E | orf1ab | VPDHDSFVKMMVDGF | 3761 | 3775 | Overlapping |
| hCoV229E | orf1ab | WDDFCRQYSNESWFE | 664 | 678 | Overlapping |
| hCoV229E | orf1ab | WDNMLKNLMADVDDP | 4661 | 4675 | Overlapping |
| hCoV229E | orf1ab | WFEDDYKAFISVLDI | 676 | 690 | Overlapping |
| hCoV229E | orf1ab | WGQDYHPDCVDCHDE | 4353 | 4367 | Overlapping |
| hCoV229E | orf1ab | WTEEDLSIGPHEFCS | 4863 | 4877 | Overlapping |
| hCoV229E | orf1ab | YDFLRSQGFFDEGSE | 4483 | 4497 | Overlapping |
| hCoV229E | orf1ab | YIYDEEGGNDLSLPV | 970 | 984 | Overlapping |
| hCoV229E | orf1ab | YLGGLNYYCVDHKPH | 5059 | 5073 | Overlapping |
| hCoV229E | orf1ab | YMPIADPTHFDIEEV | 834 | 848 | Overlapping |
| hCoV229E | orf1ab | YNPKAIHDIGNPKGI | 5915 | 5929 | Overlapping |
| hCoV229E | orf1ab | YPEFSIVCRFDTRTR | 5977 | 5991 | Overlapping |
| hCoV229E | orf1ab | YVESYNIFTQAGFNI | 6075 | 6089 | Overlapping |
| hCoV229E | spike | ADVSEMLTFDKKAFT | 646 | 660 | Positive |
| hCoV229E | spike | AELNYTVQKLQTLID | 1079 | 1093 | Positive |
| hCoV229E | spike | AIEDILFSKLVTSGL | 691 | 705 | Positive |
| hCoV229E | spike | AMPIMANLVNHKSHN | 397 | 408 | Positive |
| hCoV229E | spike | ANVSSFGDYNLSSVI | 662 | 676 | Positive |
| hCoV229E | spike | ASSDVIRYNINFEEN | 114 | 126 | Positive |
| hCoV229E | spike | CAMPIMANLVNHKSH | 396 | 408 | Positive |
| hCoV229E | spike | CDQLSFDVPDGFYST | 264 | 278 | Positive |
| hCoV229E | spike | DADYKKCTKGLSIAD | 709 | 723 | Positive |
| hCoV229E | spike | DAERMAMYTGSLIGG | 741 | 755 | Positive |
| hCoV229E | spike | DAFTGVNDAITQTSQ | 807 | 821 | Positive |
| hCoV229E | spike | DAHIPSGTVLGNFYC | 154 | 168 | Positive |
| hCoV229E | spike | DGFYSTSPIQPVELP | 273 | 287 | Positive |
| hCoV229E | spike | DLVVEQYNQTILNLT | 1054 | 1068 | Positive |
| hCoV229E | spike | DQLSFDVPDGFYSTS | 265 | 279 | Positive |
| hCoV229E | spike | EISTLENKSAELNYT | 1070 | 1084 | Positive |
| hCoV229E | spike | FDVPDGFYSTSPIQP | 269 | 282 | Positive |
| hCoV229E | spike | FGDYNLSSVIPSLPR | 667 | 680 | Positive |
| hCoV229E | spike | FNNWFLLTNTSSVVD | 54 | 68 | Positive |
| hCoV229E | spike | GFYSTSPIQPVELPM | 274 | 288 | Positive |
| hCoV229E | spike | GHFYINGYRYFSLGD | 199 | 213 | Positive |
| hCoV229E | spike | GPGKCYNCRPSVINI | 313 | 327 | Positive |
| hCoV229E | spike | GSGMFVLLVAYALLH | 1 | 12 | Positive |

| | | | | | |
|---|---|---|---|---|---|
| hCoV229E | spike | GSGSGMFVLLVAYAL | 1 | 10 | Positive |
| hCoV229E | spike | GSGSGSGMFVLLVAY | 1 | 8 | Positive |
| hCoV229E | spike | HIFSLVNAAPEGLVF | 933 | 947 | Positive |
| hCoV229E | spike | HSVCNGCVGHSENVF | 26 | 40 | Positive |
| hCoV229E | spike | IAVQPRNVSYDSVSA | 562 | 576 | Positive |
| hCoV229E | spike | IDVNKTLQELSYKLP | 1034 | 1048 | Positive |
| hCoV229E | spike | IENCNVTFVNISRSE | 1011 | 1025 | Positive |
| hCoV229E | spike | ILAASFNKAMTNIVD | 793 | 807 | Positive |
| hCoV229E | spike | INRLRCDQLSFDVPD | 259 | 273 | Positive |
| hCoV229E | spike | IPGGCAMPIMANLVN | 392 | 403 | Positive |
| hCoV229E | spike | IPSGTVLGNFYCFVN | 157 | 171 | Positive |
| hCoV229E | spike | IPSLPRSGSRVAGRS | 676 | 690 | Positive |
| hCoV229E | spike | IPSNWTTSVQVEYLQ | 584 | 598 | Positive |
| hCoV229E | spike | IPTIADFVQIENCNV | 1002 | 1016 | Positive |
| hCoV229E | spike | IRYNINFEENLRRGT | 117 | 131 | Positive |
| hCoV229E | spike | ISTLENKSAELNYTV | 1071 | 1085 | Positive |
| hCoV229E | spike | ITGVPKPVEGVSSFM | 425 | 439 | Positive |
| hCoV229E | spike | IYDVSGVGVIRISND | 451 | 465 | Positive |
| hCoV229E | spike | KFEHQRGPGKCYNCR | 308 | 320 | Positive |
| hCoV229E | spike | KIQDVVNQQGNSLNH | 832 | 846 | Positive |
| hCoV229E | spike | KPVEGVSSFMNVTLN | 430 | 444 | Positive |
| hCoV229E | spike | LENKSAELNYTVQKL | 1074 | 1088 | Positive |
| hCoV229E | spike | LNLTSEISTLENKSA | 1065 | 1079 | Positive |
| hCoV229E | spike | MANLVNHKSHNIGSL | 401 | 415 | Positive |
| hCoV229E | spike | MPIMANLVNHKSHNI | 398 | 412 | Positive |
| hCoV229E | spike | NAAPEGLVFLHTVLL | 939 | 953 | Positive |
| hCoV229E | spike | NAATIVCTVALASYA | 223 | 238 | Positive |
| hCoV229E | spike | NCLWSVSGSQFTTGF | 80 | 94 | Positive |
| hCoV229E | spike | NCNVTFVNISRSELQ | 1013 | 1027 | Positive |
| hCoV229E | spike | NCRPSVINITLANFN | 319 | 333 | Positive |
| hCoV229E | spike | NCTDAVLTYSSFGVC | 542 | 556 | Positive |
| hCoV229E | spike | NETKGPLCVDTSHFT | 333 | 347 | Positive |
| hCoV229E | spike | NFEENLRRGTILFKT | 122 | 136 | Positive |
| hCoV229E | spike | NFNVTNAATIVCTVA | 218 | 233 | Positive |
| hCoV229E | spike | NHKSHNIGSLYVSWS | 408 | 420 | Positive |
| hCoV229E | spike | NHLTSQLRQNFQAIS | 845 | 859 | Positive |
| hCoV229E | spike | NIGSLYVSWSDGDVI | 411 | 425 | Positive |
| hCoV229E | spike | NINFEENLRRGTILF | 120 | 134 | Positive |
| hCoV229E | spike | NISRSELQTIVPEYI | 1020 | 1034 | Positive |
| hCoV229E | spike | NKCTKYNIYDVSGVG | 444 | 458 | Positive |
| hCoV229E | spike | NKIQDVVNQQGNSLN | 831 | 845 | Positive |
| hCoV229E | spike | NKSAELNYTVQKLQT | 1076 | 1090 | Positive |
| hCoV229E | spike | NKTLQELSYKLPNYT | 1037 | 1051 | Positive |
| hCoV229E | spike | NLALYKEGNYYRITS | 980 | 994 | Positive |
| hCoV229E | spike | NLLGFKDVTNGTIYS | 479 | 493 | Positive |
| hCoV229E | spike | NLRRGTILFKTSYGA | 126 | 139 | Positive |
| hCoV229E | spike | NLSIPSNWTTSVQVE | 581 | 595 | Positive |
| hCoV229E | spike | NLTSEISTLENKSAE | 1066 | 1080 | Positive |
| hCoV229E | spike | NLVNHKSHNIGSLYV | 403 | 417 | Positive |

| | | | | | |
|---|---|---|---|---|---|
| hCoV229E | spike | NNTLVSGDAHIPSGT | 147 | 161 | Positive |
| hCoV229E | spike | NNWFLLTNTSSVVDG | 55 | 69 | Positive |
| hCoV229E | spike | NQKILAASFNKAMTN | 790 | 804 | Positive |
| hCoV229E | spike | NQQGNSLNHLTSQLR | 838 | 852 | Positive |
| hCoV229E | spike | NRLRCDQLSFDVPDG | 260 | 274 | Positive |
| hCoV229E | spike | NRVETYIKWPWWVWL | 1106 | 1120 | Positive |
| hCoV229E | spike | NSAMLESADVSEMLT | 639 | 653 | Positive |
| hCoV229E | spike | NSLNHLTSQLRQNFQ | 842 | 856 | Positive |
| hCoV229E | spike | NSTLVDLKWLNRVET | 1096 | 1110 | Positive |
| hCoV229E | spike | NSVINRLRCDQLSFD | 256 | 270 | Positive |
| hCoV229E | spike | NTLVSGDAHIPSGTV | 148 | 162 | Positive |
| hCoV229E | spike | NTSHSVCNGCVGHSE | 23 | 37 | Positive |
| hCoV229E | spike | NVFAVESGGYIPSNF | 38 | 52 | Positive |
| hCoV229E | spike | NVKLARWSASINTGN | 356 | 368 | Positive |
| hCoV229E | spike | NVRCVELLKQYTSAC | 616 | 630 | Positive |
| hCoV229E | spike | NVSYDSVSAIVTANL | 568 | 582 | Positive |
| hCoV229E | spike | NVVEMPKFFYASNGT | 526 | 540 | Positive |
| hCoV229E | spike | NWTTSVQVEYLQITS | 587 | 601 | Positive |
| hCoV229E | spike | NYTVPDLVVEQYNQT | 1049 | 1063 | Positive |
| hCoV229E | spike | NYTVQKLQTLIDNIN | 1082 | 1096 | Positive |
| hCoV229E | spike | NYVALQTDVLQENQK | 778 | 792 | Positive |
| hCoV229E | spike | PEGLVFLHTVLLPTQ | 942 | 956 | Positive |
| hCoV229E | spike | PGGCAMPIMANLVNH | 393 | 403 | Positive |
| hCoV229E | spike | PIMANLVNHKSHNIG | 399 | 413 | Positive |
| hCoV229E | spike | PKPVEGVSSFMNVTL | 429 | 443 | Positive |
| hCoV229E | spike | PLCVDTSHFTTQFVD | 338 | 351 | Positive |
| hCoV229E | spike | PNYTVPDLVVEQYNQ | 1048 | 1062 | Positive |
| hCoV229E | spike | PVEGVSSFMNVTLNK | 431 | 445 | Positive |
| hCoV229E | spike | PVELPMSIVSLPVYH | 284 | 297 | Positive |
| hCoV229E | spike | PYYDVEKIHIQGSGS | 1163 | 1173 | Positive |
| hCoV229E | spike | QELSYKLPNYTVPDL | 1041 | 1055 | Positive |
| hCoV229E | spike | QPVELPMSIVSLPVY | 282 | 296 | Positive |
| hCoV229E | spike | RCDQLSFDVPDGFYS | 263 | 277 | Positive |
| hCoV229E | spike | RCVELLKQYTSACKT | 618 | 632 | Positive |
| hCoV229E | spike | REFVISRTGHFYING | 191 | 205 | Positive |
| hCoV229E | spike | RTGHFYINGYRYFSL | 197 | 211 | Positive |
| hCoV229E | spike | SAIVTANLSIPSNWT | 575 | 589 | Positive |
| hCoV229E | spike | SEISTLENKSAELNY | 1069 | 1083 | Positive |
| hCoV229E | spike | SELQTIVPEYIDVNK | 1024 | 1038 | Positive |
| hCoV229E | spike | SEMLTFDKKAFTLAN | 649 | 663 | Positive |
| hCoV229E | spike | SENVFAVESGGYIPS | 36 | 50 | Positive |
| hCoV229E | spike | SHSVCNGCVGHSENV | 25 | 39 | Positive |
| hCoV229E | spike | SMLLLCCCSTGCCGF | 1131 | 1145 | Positive |
| hCoV229E | spike | SNASSDVIRYNINFE | 110 | 124 | Positive |
| hCoV229E | spike | SRTGHFYINGYRYFS | 196 | 210 | Positive |
| hCoV229E | spike | SSDVIRYNINFEENL | 114 | 127 | Positive |
| hCoV229E | spike | STLENKSAELNYTVQ | 1072 | 1086 | Positive |
| hCoV229E | spike | TDAVLTYSSFGVCAD | 544 | 558 | Positive |
| hCoV229E | spike | TDVLQENQKILAASF | 784 | 798 | Positive |

| | | | | | |
|---|---|---|---|---|---|
| hCoV229E | spike | TGHFYINGYRYFSLG | 198 | 212 | Positive |
| hCoV229E | spike | TIEDALRNSAMLESA | 632 | 646 | Positive |
| hCoV229E | spike | TLENKSAELNYTVQK | 1073 | 1087 | Positive |
| hCoV229E | spike | TLIDNINSTLVDLKW | 1090 | 1104 | Positive |
| hCoV229E | spike | TLQELSYKLPNYTVP | 1039 | 1053 | Positive |
| hCoV229E | spike | TNGTIYSITPCNPPD | 487 | 501 | Positive |
| hCoV229E | spike | TSEISTLENKSAELN | 1068 | 1082 | Positive |
| hCoV229E | spike | TSGLGTVDADYKKCT | 702 | 716 | Positive |
| hCoV229E | spike | TSGNLLGFKDVTNGT | 476 | 490 | Positive |
| hCoV229E | spike | TSRIMFEPRIPTIAD | 993 | 1007 | Positive |
| hCoV229E | spike | TVPDLVVEQYNQTIL | 1051 | 1065 | Positive |
| hCoV229E | spike | TVREFVISRTGHFYI | 189 | 203 | Positive |
| hCoV229E | spike | VADAERMAMYTGSLI | 739 | 753 | Positive |
| hCoV229E | spike | VDADYKKCTKGLSIA | 708 | 722 | Positive |
| hCoV229E | spike | VDAFTGVNDAITQTS | 806 | 820 | Positive |
| hCoV229E | spike | VDTSHFTTQFVDNVK | 341 | 351 | Positive |
| hCoV229E | spike | VEAVNFNVTNAATIV | 214 | 226 | Positive |
| hCoV229E | spike | VELPMSIVSLPVYHK | 284 | 298 | Positive |
| hCoV229E | spike | VEQYNQTILNLTSEI | 1057 | 1071 | Positive |
| hCoV229E | spike | VESGGYIPSNFSFNN | 42 | 56 | Positive |
| hCoV229E | spike | VEYLQITSTPIVVDC | 594 | 608 | Positive |
| hCoV229E | spike | VFAVESGGYIPSNFS | 39 | 53 | Positive |
| hCoV229E | spike | VNFNVTNAATIVCTV | 217 | 232 | Positive |
| hCoV229E | spike | VPDGFYSTSPIQPVE | 271 | 285 | Positive |
| hCoV229E | spike | VPDLVVEQYNQTILN | 1052 | 1066 | Positive |
| hCoV229E | spike | VPEYIDVNKTLQELS | 1030 | 1044 | Positive |
| hCoV229E | spike | VSEMLTFDKKAFTLA | 648 | 662 | Positive |
| hCoV229E | spike | VVEMPKFFYASNGTY | 527 | 541 | Positive |
| hCoV229E | spike | VVGAMLSENFTSYGF | 510 | 524 | Positive |
| hCoV229E | spike | YDVEKIHIQGSGSGS | 1165 | 1173 | Positive |
| hCoV229E | spike | YHKHTFIVLHVKFEH | 296 | 309 | Positive |
| hCoV229E | spike | YVALQTDVLQENQKI | 779 | 793 | Positive |
| hCoV229E | spike | YYDVEKIHIQGSGSG | 1164 | 1173 | Positive |
| hCoV229E | spike | ACAQYYNGIMVLPGV | 725 | 739 | Negative |
| hCoV229E | spike | ADAERMAMYTGSLIG | 740 | 754 | Negative |
| hCoV229E | spike | ADGSIIAVQPRNVSY | 557 | 571 | Negative |
| hCoV229E | spike | AGCQTTNGTNTSHSV | 14 | 28 | Negative |
| hCoV229E | spike | AMTNIVDAFTGVNDA | 801 | 815 | Negative |
| hCoV229E | spike | AQYYNGIMVLPGVAD | 727 | 741 | Negative |
| hCoV229E | spike | AWSGLCVDGINGYVL | 962 | 976 | Negative |
| hCoV229E | spike | CQTTNGTNTSHSVCN | 16 | 30 | Negative |
| hCoV229E | spike | CRPSVINITLANFNE | 320 | 334 | Negative |
| hCoV229E | spike | CTKYNIYDVSGVGVI | 446 | 460 | Negative |
| hCoV229E | spike | DVEAVNFNVTNAATI | 213 | 226 | Negative |
| hCoV229E | spike | DVEAWSGLCVDGING | 959 | 973 | Negative |
| hCoV229E | spike | DVEKIHIQGSGSGSG | 1166 | 1173 | Negative |
| hCoV229E | spike | EGLVFLHTVLLPTQY | 943 | 957 | Negative |
| hCoV229E | spike | EGNYYRITSRIMFEP | 986 | 1000 | Negative |
| hCoV229E | spike | EGVSSFMNVTLNKCT | 433 | 447 | Negative |

| | | | | | |
|---|---|---|---|---|---|
| hCoV229E | spike | EISTLENKSAELNYT | 1070 | 1084 | Negative |
| hCoV229E | spike | ELLKQYTSACKTIED | 621 | 635 | Negative |
| hCoV229E | spike | ELNYTVQKLQTLIDN | 1080 | 1094 | Negative |
| hCoV229E | spike | ELPMSIVSLPVYHKH | 285 | 299 | Negative |
| hCoV229E | spike | ELQTIVPEYIDVNKT | 1025 | 1039 | Negative |
| hCoV229E | spike | ELSYKLPNYTVPDLV | 1042 | 1056 | Negative |
| hCoV229E | spike | EMLTFDKKAFTLANV | 650 | 664 | Negative |
| hCoV229E | spike | EYIDVNKTLQELSYK | 1032 | 1046 | Negative |
| hCoV229E | spike | FEHQRGPGKCYNCRP | 308 | 322 | Negative |
| hCoV229E | spike | FLLTNTSSVVDGVVR | 58 | 72 | Negative |
| hCoV229E | spike | FNETKGPLCVDTSHF | 332 | 346 | Negative |
| hCoV229E | spike | FSLGDVEAVNFNVTN | 209 | 222 | Negative |
| hCoV229E | spike | FSNVVEMPKFFYASN | 524 | 538 | Negative |
| hCoV229E | spike | FTGVNDAITQTSQAL | 809 | 823 | Negative |
| hCoV229E | spike | GCQTTNGTNTSHSVC | 15 | 29 | Negative |
| hCoV229E | spike | GDVEAVNFNVTNAAT | 212 | 226 | Negative |
| hCoV229E | spike | GFSNVVEMPKFFYAS | 523 | 537 | Negative |
| hCoV229E | spike | GHFYINGYRYFSLGD | 199 | 213 | Negative |
| hCoV229E | spike | GHSENVFAVESGGYI | 34 | 48 | Negative |
| hCoV229E | spike | GIALGGLTSAASIPF | 755 | 769 | Negative |
| hCoV229E | spike | GIMVLPGVADAERMA | 732 | 746 | Negative |
| hCoV229E | spike | GINGYVLRQPNLALY | 970 | 984 | Negative |
| hCoV229E | spike | GITYTSTSGNLLGFK | 470 | 484 | Negative |
| hCoV229E | spike | GKVNNFVKFGSVCFS | 374 | 388 | Negative |
| hCoV229E | spike | GLCVDGINGYVLRQP | 965 | 979 | Negative |
| hCoV229E | spike | GLGTVDADYKKCTKG | 704 | 718 | Negative |
| hCoV229E | spike | GLSIADLACAQYYNG | 718 | 732 | Negative |
| hCoV229E | spike | GLTSAASIPFSLAIQ | 760 | 774 | Negative |
| hCoV229E | spike | GMFVLLVAYALLHIA | 1 | 14 | Negative |
| hCoV229E | spike | GRSAIEDILFSKLVT | 688 | 702 | Negative |
| hCoV229E | spike | GSGMFVLLVAYALLH | 1 | 12 | Negative |
| hCoV229E | spike | GSGSGMFVLLVAYAL | 1 | 10 | Negative |
| hCoV229E | spike | GSGSGSGMFVLLVAY | 1 | 8 | Negative |
| hCoV229E | spike | GTHIFSLVNAAPEGL | 931 | 945 | Negative |
| hCoV229E | spike | GTNTSHSVCNGCVGH | 21 | 35 | Negative |
| hCoV229E | spike | GTVDADYKKCTKGLS | 706 | 720 | Negative |
| hCoV229E | spike | GVADAERMAMYTGSL | 738 | 752 | Negative |
| hCoV229E | spike | GVCADGSIIAVQPRN | 554 | 568 | Negative |
| hCoV229E | spike | GVGVIRISNDTFLNG | 456 | 470 | Negative |
| hCoV229E | spike | GVIRISNDTFLNGIT | 458 | 472 | Negative |
| hCoV229E | spike | GVNDAITQTSQALQT | 811 | 825 | Negative |
| hCoV229E | spike | GVSSFMNVTLNKCTK | 434 | 448 | Negative |
| hCoV229E | spike | GVVRSFQPLLLNCLW | 69 | 83 | Negative |
| hCoV229E | spike | HIAGCQTTNGTNTSH | 12 | 26 | Negative |
| hCoV229E | spike | IAGCQTTNGTNTSHS | 13 | 27 | Negative |
| hCoV229E | spike | IDVNKTLQELSYKLP | 1034 | 1048 | Negative |
| hCoV229E | spike | IMANLVNHKSHNIGS | 400 | 414 | Negative |
| hCoV229E | spike | IMFEPRIPTIADFVQ | 996 | 1010 | Negative |
| hCoV229E | spike | IMVLPGVADAERMAM | 733 | 747 | Negative |

| | | | | | |
|---|---|---|---|---|---|
| hCoV229E | spike | IRISNDTFLNGITYT | 460 | 474 | Negative |
| hCoV229E | spike | ISRSELQTIVPEYID | 1021 | 1035 | Negative |
| hCoV229E | spike | IVPEYIDVNKTLQEL | 1029 | 1043 | Negative |
| hCoV229E | spike | IYDRLDIIQADQQVD | 865 | 879 | Negative |
| hCoV229E | spike | KAMTNIVDAFTGVND | 800 | 814 | Negative |
| hCoV229E | spike | KCTKYNIYDVSGVGV | 445 | 459 | Negative |
| hCoV229E | spike | KDVEAWSGLCVDGIN | 958 | 972 | Negative |
| hCoV229E | spike | KGLSIADLACAQYYN | 717 | 731 | Negative |
| hCoV229E | spike | KLPNYTVPDLVVEQY | 1046 | 1060 | Negative |
| hCoV229E | spike | KTIEDALRNSAMLES | 631 | 645 | Negative |
| hCoV229E | spike | LGTVDADYKKCTKGL | 705 | 719 | Negative |
| hCoV229E | spike | LHIAGCQTTNGTNTS | 11 | 25 | Negative |
| hCoV229E | spike | LQELSYKLPNYTVPD | 1040 | 1054 | Negative |
| hCoV229E | spike | LSIADLACAQYYNGI | 719 | 733 | Negative |
| hCoV229E | spike | LTSEISTLENKSAEL | 1067 | 1081 | Negative |
| hCoV229E | spike | MLSENFTSYGFSNVV | 514 | 528 | Negative |
| hCoV229E | spike | NETKGPLCVDTSHFT | 333 | 347 | Negative |
| hCoV229E | spike | NFSFNNWFLLTNTSS | 51 | 65 | Negative |
| hCoV229E | spike | PIVVDCSTYVCNGNV | 603 | 617 | Negative |
| hCoV229E | spike | PTIADFVQIENCNVT | 1003 | 1017 | Negative |
| hCoV229E | spike | QSRLNYVALQTDVLQ | 774 | 788 | Negative |
| hCoV229E | spike | QTLIDNINSTLVDLK | 1089 | 1103 | Negative |
| hCoV229E | spike | RMAMYTGSLIGGIAL | 744 | 758 | Negative |
| hCoV229E | spike | RSAIEDILFSKLVTS | 689 | 703 | Negative |
| hCoV229E | spike | RVETYIKWPWWVWLC | 1107 | 1121 | Negative |
| hCoV229E | spike | SAIEDILFSKLVTSG | 690 | 704 | Negative |
| hCoV229E | spike | SKRYGFCGNGTHIFS | 922 | 936 | Negative |
| hCoV229E | spike | SLGDVEAVNFNVTNA | 210 | 224 | Negative |
| hCoV229E | spike | SSIKGCCESTKLPYY | 1151 | 1165 | Negative |
| hCoV229E | spike | SSVIPSLPRSGSRVA | 673 | 687 | Negative |
| hCoV229E | spike | SSVVDGVVRSFQPLL | 64 | 78 | Negative |
| hCoV229E | spike | STKLPYYDVEKIHIQ | 1159 | 1173 | Negative |
| hCoV229E | spike | STLVDLKWLNRVETY | 1097 | 1111 | Negative |
| hCoV229E | spike | SYKLPNYTVPDLVVE | 1044 | 1058 | Negative |
| hCoV229E | spike | TKYNIYDVSGVGVIR | 447 | 461 | Negative |
| hCoV229E | spike | TLVDLKWLNRVETYI | 1098 | 1112 | Negative |
| hCoV229E | spike | TNTSHSVCNGCVGHS | 23 | 36 | Negative |
| hCoV229E | spike | TNTSSVVDGVVRSFQ | 61 | 75 | Negative |
| hCoV229E | spike | TPIVVDCSTYVCNGN | 602 | 616 | Negative |
| hCoV229E | spike | VEAVNFNVTNAATIV | 214 | 226 | Negative |
| hCoV229E | spike | VELLKQYTSACKTIE | 620 | 634 | Negative |
| hCoV229E | spike | VGVIRISNDTFLNGI | 457 | 471 | Negative |
| hCoV229E | spike | VIRISNDTFLNGITY | 459 | 473 | Negative |
| hCoV229E | spike | WFLLTNTSSVVDGVV | 57 | 71 | Negative |
| hCoV229E | spike | YKDVEAWSGLCVDGI | 957 | 970 | Negative |
| hCoV229E | spike | YNCTDAVLTYSSFGV | 541 | 555 | Negative |
| hCoV229E | spike | AERMAMYTGSLIGGI | 742 | 756 | Overlapping |
| hCoV229E | spike | DGDVITGVPKPVEGV | 421 | 435 | Overlapping |
| hCoV229E | spike | DGSIIAVQPRNVSYD | 558 | 572 | Overlapping |

| | | | | | |
|---|---|---|---|---|---|
| hCoV229E | spike | DVIRYNINFEENLRR | 115 | 129 | Overlapping |
| hCoV229E | spike | EAVNFNVTNAATIVC | 215 | 230 | Overlapping |
| hCoV229E | spike | EAWSGLCVDGINGYV | 961 | 975 | Overlapping |
| hCoV229E | spike | EDALRNSAMLESADV | 634 | 648 | Overlapping |
| hCoV229E | spike | EDILFSKLVTSGLGT | 693 | 707 | Overlapping |
| hCoV229E | spike | EENLRRGTILFKTSY | 124 | 138 | Overlapping |
| hCoV229E | spike | EFVISRTGHFYINGY | 192 | 206 | Overlapping |
| hCoV229E | spike | EGLVFLHTVLLPTQY | 943 | 957 | Overlapping |
| hCoV229E | spike | EGNYYRITSRIMFEP | 986 | 1000 | Overlapping |
| hCoV229E | spike | EGVSSFMNVTLNKCT | 433 | 447 | Overlapping |
| hCoV229E | spike | EHQRGPGKCYNCRPS | 311 | 323 | Overlapping |
| hCoV229E | spike | ELLKQYTSACKTIED | 621 | 635 | Overlapping |
| hCoV229E | spike | ELNYTVQKLQTLIDN | 1080 | 1094 | Overlapping |
| hCoV229E | spike | ELPMSIVSLPVYHKH | 285 | 299 | Overlapping |
| hCoV229E | spike | ELQTIVPEYIDVNKT | 1025 | 1039 | Overlapping |
| hCoV229E | spike | ELSYKLPNYTVPDLV | 1042 | 1056 | Overlapping |
| hCoV229E | spike | EMLTFDKKAFTLANV | 650 | 664 | Overlapping |
| hCoV229E | spike | EMPKFFYASNGTYNC | 529 | 543 | Overlapping |
| hCoV229E | spike | ENCNVTFVNISRSEL | 1012 | 1026 | Overlapping |
| hCoV229E | spike | ENFTSYGFSNVVEMP | 517 | 531 | Overlapping |
| hCoV229E | spike | ENKSAELNYTVQKLQ | 1075 | 1089 | Overlapping |
| hCoV229E | spike | ENLRRGTILFKTSYG | 125 | 139 | Overlapping |
| hCoV229E | spike | ENQKILAASFNKAMT | 789 | 803 | Overlapping |
| hCoV229E | spike | ENVFAVESGGYIPSN | 37 | 51 | Overlapping |
| hCoV229E | spike | EPRIPTIADFVQIEN | 999 | 1013 | Overlapping |
| hCoV229E | spike | EQYNQTILNLTSEIS | 1058 | 1072 | Overlapping |
| hCoV229E | spike | ERMAMYTGSLIGGIA | 743 | 757 | Overlapping |
| hCoV229E | spike | ESADVSEMLTFDKKA | 644 | 658 | Overlapping |
| hCoV229E | spike | ESGGYIPSNFSFNNW | 43 | 57 | Overlapping |
| hCoV229E | spike | ESTKLPYYDVEKIHI | 1158 | 1172 | Overlapping |
| hCoV229E | spike | ETKGPLCVDTSHFTT | 334 | 348 | Overlapping |
| hCoV229E | spike | ETTSAFVGALPKTVR | 177 | 191 | Overlapping |
| hCoV229E | spike | ETYIKWPWWVWLCIS | 1109 | 1123 | Overlapping |
| hCoV229E | spike | EVRASRQLAQQKVNE | 902 | 916 | Overlapping |
| hCoV229E | spike | EYLQITSTPIVVDCS | 595 | 609 | Overlapping |
| hCoV229E | spike | FEENLRRGTILFKTS | 123 | 137 | Overlapping |
| hCoV229E | spike | FEPRIPTIADFVQIE | 998 | 1012 | Overlapping |
| hCoV229E | spike | FYINGYRYFSLGDVE | 201 | 215 | Overlapping |
| hCoV229E | spike | GDVITGVPKPVEGVS | 422 | 436 | Overlapping |
| hCoV229E | spike | GHSENVFAVESGGYI | 34 | 48 | Overlapping |
| hCoV229E | spike | GPLCVDTSHFTTQFV | 337 | 351 | Overlapping |
| hCoV229E | spike | GTNTSHSVCNGCVGH | 21 | 35 | Overlapping |
| hCoV229E | spike | GVPKPVEGVSSFMNV | 427 | 441 | Overlapping |
| hCoV229E | spike | IEDALRNSAMLESAD | 633 | 647 | Overlapping |
| hCoV229E | spike | IEDILFSKLVTSGLG | 692 | 706 | Overlapping |
| hCoV229E | spike | KEGNYYRITSRIMFE | 985 | 999 | Overlapping |
| hCoV229E | spike | NASSDVIRYNINFEE | 114 | 125 | Overlapping |
| hCoV229E | spike | NFNETKGPLCVDTSH | 331 | 345 | Overlapping |
| hCoV229E | spike | NKAMTNIVDAFTGVN | 799 | 813 | Overlapping |

| | | | | | |
|---|---|---|---|---|---|
| hCoV229E | spike | NQTILNLTSEISTLE | 1061 | 1075 | Overlapping |
| hCoV229E | spike | NWFLLTNTSSVVDGV | 56 | 70 | Overlapping |
| hCoV229E | spike | PEYIDVNKTLQELSY | 1031 | 1045 | Overlapping |
| hCoV229E | spike | PIQPVELPMSIVSLP | 280 | 294 | Overlapping |
| hCoV229E | spike | PVEGVSSFMNVTLNK | 431 | 445 | Overlapping |
| hCoV229E | spike | PVELPMSIVSLPVYH | 284 | 297 | Overlapping |
| hCoV229E | spike | QENQKILAASFNKAM | 788 | 802 | Overlapping |
| hCoV229E | spike | QVEYLQITSTPIVVD | 593 | 607 | Overlapping |
| hCoV229E | spike | SDGDVITGVPKPVEG | 420 | 434 | Overlapping |
| hCoV229E | spike | SDVIRYNINFEENLR | 114 | 128 | Overlapping |
| hCoV229E | spike | SENFTSYGFSNVVEM | 516 | 530 | Overlapping |
| hCoV229E | spike | SFDVPDGFYSTSPIQ | 268 | 282 | Overlapping |
| hCoV229E | spike | TEVRASRQLAQQKVN | 901 | 915 | Overlapping |
| hCoV229E | spike | TGVNDAITQTSQALQ | 810 | 824 | Overlapping |
| hCoV229E | spike | VEAWSGLCVDGINGY | 960 | 974 | Overlapping |
| hCoV229E | spike | VEGVSSFMNVTLNKC | 432 | 446 | Overlapping |
| hCoV229E | spike | VEMPKFFYASNGTYN | 528 | 542 | Overlapping |
| hCoV229E | spike | VETYIKWPWWVWLCI | 1108 | 1122 | Overlapping |
| hCoV229E | spike | YINGYRYFSLGDVEA | 202 | 216 | Overlapping |

### Table S7

| Protein | Amino Acid Position in Protein | Average RFU of Antibody responses in SARS-CoV-2-positive patients | Average RFU of Antibody responsee in SARS-CoV-2-negative patients | Cross-reactive epitope site | # of unique mutations observed in public database |
|---|---|---|---|---|---|
| orf1a | 1 | 2084 | 1532 | FALSE | 2 |
| orf1a | 2 | 3408 | 3833 | FALSE | 3 |
| orf1a | 3 | 3408 | 3833 | FALSE | 3 |
| orf1a | 4 | 3408 | 3833 | FALSE | 3 |
| orf1a | 5 | 3408 | 3833 | FALSE | 4 |
| orf1a | 6 | 3408 | 3833 | FALSE | 3 |
| orf1a | 7 | 3408 | 3833 | FALSE | 3 |
| orf1a | 8 | 3408 | 3833 | FALSE | 3 |
| orf1a | 9 | 3408 | 3833 | FALSE | 2 |
| orf1a | 10 | 4024 | 4615 | FALSE | 4 |
| orf1a | 11 | 4620 | 6845 | FALSE | 2 |
| orf1a | 12 | 4620 | 6845 | FALSE | 2 |
| orf1a | 13 | 4977 | 6845 | FALSE | 4 |
| orf1a | 14 | 4977 | 6845 | FALSE | 3 |
| orf1a | 15 | 5964 | 9239 | FALSE | 1 |
| orf1a | 16 | 5964 | 9239 | FALSE | 2 |
| orf1a | 17 | 5874 | 7741 | FALSE | 3 |
| orf1a | 18 | 5874 | 7741 | FALSE | 4 |
| orf1a | 19 | 5874 | 7741 | FALSE | 4 |
| orf1a | 20 | 5874 | 7741 | FALSE | 4 |
| orf1a | 21 | 5874 | 7741 | FALSE | 2 |
| orf1a | 22 | 4989 | 7741 | FALSE | 3 |
| orf1a | 23 | 4989 | 7741 | FALSE | 2 |
| orf1a | 24 | 9288 | 7741 | FALSE | 4 |
| orf1a | 25 | 12702 | 0 | FALSE | 2 |
| orf1a | 26 | 12702 | 0 | FALSE | 2 |
| orf1a | 27 | 12702 | 0 | FALSE | 4 |
| orf1a | 28 | 8996 | 0 | FALSE | 4 |
| orf1a | 29 | 8996 | 0 | FALSE | 3 |
| orf1a | 30 | 8996 | 1400 | FALSE | 4 |
| orf1a | 31 | 8996 | 1400 | FALSE | 1 |
| orf1a | 32 | 7032 | 1322 | FALSE | 1 |
| orf1a | 33 | 7032 | 1322 | FALSE | 4 |
| orf1a | 34 | 5237 | 1337 | FALSE | 3 |
| orf1a | 35 | 5237 | 1337 | FALSE | 5 |
| orf1a | 36 | 8968 | 4813 | FALSE | 4 |
| orf1a | 37 | 9543 | 4813 | FALSE | 4 |
| orf1a | 38 | 9543 | 4813 | FALSE | 5 |
| orf1a | 39 | 8138 | 4813 | FALSE | 3 |
| orf1a | 40 | 8138 | 4813 | FALSE | 3 |
| orf1a | 41 | 8138 | 4813 | FALSE | 3 |
| orf1a | 42 | 8138 | 4813 | FALSE | 2 |
| orf1a | 43 | 8957 | 4813 | FALSE | 3 |
| orf1a | 44 | 8178 | 4813 | FALSE | 2 |
| orf1a | 45 | 8178 | 5382 | FALSE | 2 |
| orf1a | 46 | 8178 | 5382 | FALSE | 3 |
| orf1a | 47 | 9058 | 6210 | FALSE | 4 |
| orf1a | 48 | 9058 | 6210 | FALSE | 4 |
| orf1a | 49 | 11528 | 7420 | FALSE | 4 |
| orf1a | 50 | 11528 | 7420 | FALSE | 1 |
| orf1a | 51 | 1941 | 0 | FALSE | 3 |
| orf1a | 52 | 1557 | 14588 | FALSE | 4 |
| orf1a | 53 | 1557 | 14588 | FALSE | 1 |
| orf1a | 54 | 1557 | 14588 | FALSE | 1 |
| orf1a | 55 | 1557 | 14588 | FALSE | 3 |
| orf1a | 56 | 1557 | 14588 | FALSE | 2 |
| orf1a | 57 | 1557 | 14588 | FALSE | 1 |
| orf1a | 58 | 1557 | 14588 | FALSE | 2 |
| orf1a | 59 | 1173 | 14588 | FALSE | 6 |
| orf1a | 60 | 1173 | 14588 | FALSE | 2 |
| orf1a | 61 | 1173 | 14588 | FALSE | 5 |

| | | | | | |
|---|---|---|---|---|---|
| orf1a | 62 | 1173 | 14588 | FALSE | 2 |
| orf1a | 63 | 1173 | 14588 | FALSE | 1 |
| orf1a | 64 | 1173 | 14588 | FALSE | 2 |
| orf1a | 65 | 1173 | 14588 | FALSE | 1 |
| orf1a | 66 | 1173 | 14588 | FALSE | 5 |
| orf1a | 72 | 1424 | 0 | FALSE | 1 |
| orf1a | 73 | 1424 | 0 | FALSE | 5 |
| orf1a | 74 | 3489 | 0 | FALSE | 3 |
| orf1a | 75 | 3489 | 0 | FALSE | 4 |
| orf1a | 76 | 3373 | 0 | FALSE | 3 |
| orf1a | 77 | 3373 | 0 | FALSE | 3 |
| orf1a | 78 | 3011 | 0 | FALSE | 2 |
| orf1a | 79 | 3011 | 0 | FALSE | 4 |
| orf1a | 80 | 3148 | 0 | FALSE | 2 |
| orf1a | 81 | 3148 | 0 | FALSE | 2 |
| orf1a | 82 | 3148 | 0 | FALSE | 4 |
| orf1a | 83 | 3148 | 0 | FALSE | 2 |
| orf1a | 84 | 3148 | 0 | FALSE | 3 |
| orf1a | 85 | 3148 | 0 | FALSE | 6 |
| orf1a | 86 | 3148 | 0 | FALSE | 2 |
| orf1a | 87 | 3492 | 0 | FALSE | 2 |
| orf1a | 88 | 3079 | 0 | FALSE | 3 |
| orf1a | 89 | 2358 | 0 | FALSE | 2 |
| orf1a | 90 | 2358 | 0 | FALSE | 3 |
| orf1a | 91 | 2135 | 0 | FALSE | 2 |
| orf1a | 92 | 2135 | 0 | FALSE | 4 |
| orf1a | 93 | 2422 | 0 | FALSE | 5 |
| orf1a | 94 | 2422 | 3092 | FALSE | 4 |
| orf1a | 95 | 1012 | 3092 | FALSE | 3 |
| orf1a | 96 | 1012 | 3092 | FALSE | 5 |
| orf1a | 97 | 1012 | 3092 | FALSE | 3 |
| orf1a | 98 | 1012 | 2646 | FALSE | 3 |
| orf1a | 99 | 1012 | 2646 | FALSE | 1 |
| orf1a | 100 | 2905 | 2646 | FALSE | 4 |
| orf1a | 101 | 2905 | 2646 | FALSE | 1 |
| orf1a | 102 | 5353 | 3132 | FALSE | 2 |
| orf1a | 103 | 6800 | 3132 | FALSE | 3 |
| orf1a | 104 | 6800 | 3132 | FALSE | 2 |
| orf1a | 105 | 6800 | 3132 | FALSE | 4 |
| orf1a | 106 | 6800 | 3132 | FALSE | 4 |
| orf1a | 107 | 6800 | 3132 | FALSE | 3 |
| orf1a | 108 | 6800 | 3132 | FALSE | 2 |
| orf1a | 109 | 6800 | 3152 | FALSE | 2 |
| orf1a | 110 | 6800 | 3152 | FALSE | 4 |
| orf1a | 111 | 6800 | 3152 | FALSE | 2 |
| orf1a | 112 | 6800 | 3152 | FALSE | 4 |
| orf1a | 113 | 6800 | 4102 | FALSE | 2 |
| orf1a | 114 | 6800 | 4102 | FALSE | 2 |
| orf1a | 115 | 12697 | 4102 | FALSE | 2 |
| orf1a | 116 | 12697 | 4102 | FALSE | 3 |
| orf1a | 120 | 1797 | 0 | FALSE | 4 |
| orf1a | 121 | 1797 | 0 | FALSE | 2 |
| orf1a | 122 | 1797 | 0 | FALSE | 4 |
| orf1a | 123 | 1797 | 0 | FALSE | 2 |
| orf1a | 124 | 1797 | 0 | FALSE | 3 |
| orf1a | 125 | 1797 | 0 | FALSE | 2 |
| orf1a | 126 | 1797 | 0 | FALSE | 2 |
| orf1a | 127 | 1797 | 0 | FALSE | 3 |
| orf1a | 128 | 5193 | 0 | FALSE | 1 |
| orf1a | 129 | 5193 | 0 | FALSE | 3 |
| orf1a | 130 | 3822 | 0 | FALSE | 3 |
| orf1a | 131 | 3822 | 0 | FALSE | 4 |

| | | | | | |
|---|---|---|---|---|---|
| orf1a | 132 | 3822 | 1767 | FALSE | 4 |
| orf1a | 133 | 3822 | 1767 | FALSE | 4 |
| orf1a | 134 | 3822 | 1767 | FALSE | 2 |
| orf1a | 135 | 4835 | 1767 | FALSE | 3 |
| orf1a | 136 | 4835 | 1767 | FALSE | 4 |
| orf1a | 137 | 4835 | 1767 | FALSE | 7 |
| orf1a | 138 | 4835 | 2438 | FALSE | 2 |
| orf1a | 139 | 4835 | 2438 | FALSE | 3 |
| orf1a | 140 | 4835 | 2438 | FALSE | 5 |
| orf1a | 141 | 4835 | 2438 | FALSE | 4 |
| orf1a | 142 | 3033 | 1925 | FALSE | 2 |
| orf1a | 143 | 1644 | 1925 | FALSE | 5 |
| orf1a | 144 | 1672 | 2070 | FALSE | 4 |
| orf1a | 145 | 1771 | 2070 | FALSE | 4 |
| orf1a | 146 | 1799 | 2316 | FALSE | 5 |
| orf1a | 147 | 1799 | 2408 | FALSE | 1 |
| orf1a | 148 | 3257 | 3137 | FALSE | 4 |
| orf1a | 149 | 3257 | 3137 | FALSE | 3 |
| orf1a | 150 | 3187 | 4391 | FALSE | 5 |
| orf1a | 151 | 3187 | 4391 | FALSE | 3 |
| orf1a | 152 | 3187 | 4227 | FALSE | 5 |
| orf1a | 153 | 3187 | 4329 | FALSE | 3 |
| orf1a | 154 | 3187 | 4329 | FALSE | 4 |
| orf1a | 155 | 3187 | 4329 | FALSE | 6 |
| orf1a | 156 | 3187 | 4062 | FALSE | 6 |
| orf1a | 157 | 3477 | 4592 | FALSE | 2 |
| orf1a | 158 | 3477 | 4062 | FALSE | 2 |
| orf1a | 159 | 3960 | 4190 | FALSE | 4 |
| orf1a | 160 | 4008 | 4190 | FALSE | 1 |
| orf1a | 161 | 5091 | 4470 | FALSE | 8 |
| orf1a | 162 | 4703 | 4326 | FALSE | 4 |
| orf1a | 163 | 2733 | 4391 | FALSE | 2 |
| orf1a | 164 | 2733 | 3953 | FALSE | 2 |
| orf1a | 165 | 3069 | 1788 | FALSE | 2 |
| orf1a | 166 | 3069 | 1788 | FALSE | 5 |
| orf1a | 167 | 3069 | 1661 | FALSE | 1 |
| orf1a | 168 | 3069 | 3997 | FALSE | 3 |
| orf1a | 169 | 3069 | 3997 | FALSE | 3 |
| orf1a | 170 | 3069 | 3411 | FALSE | 3 |
| orf1a | 171 | 3069 | 3739 | FALSE | 3 |
| orf1a | 172 | 6227 | 4985 | FALSE | 5 |
| orf1a | 173 | 6227 | 5877 | FALSE | 5 |
| orf1a | 174 | 5285 | 5877 | FALSE | 3 |
| orf1a | 175 | 5473 | 5877 | FALSE | 4 |
| orf1a | 176 | 6493 | 5527 | FALSE | 6 |
| orf1a | 177 | 7716 | 5804 | FALSE | 2 |
| orf1a | 178 | 7716 | 5804 | FALSE | 2 |
| orf1a | 179 | 7716 | 6364 | FALSE | 4 |
| orf1a | 180 | 7716 | 6364 | FALSE | 7 |
| orf1a | 181 | 7716 | 6364 | FALSE | 3 |
| orf1a | 182 | 7716 | 6364 | FALSE | 3 |
| orf1a | 183 | 7716 | 5034 | FALSE | 1 |
| orf1a | 184 | 7716 | 5034 | FALSE | 3 |
| orf1a | 185 | 7716 | 6386 | FALSE | 2 |
| orf1a | 186 | 7716 | 6386 | FALSE | 3 |
| orf1a | 187 | 6048 | 4127 | FALSE | 1 |
| orf1a | 188 | 6048 | 4127 | FALSE | 4 |
| orf1a | 189 | 10576 | 4127 | FALSE | 3 |
| orf1a | 190 | 10576 | 4127 | FALSE | 3 |
| orf1a | 192 | 2011 | 0 | FALSE | 7 |
| orf1a | 193 | 2011 | 0 | FALSE | 5 |
| orf1a | 194 | 1744 | 0 | FALSE | 7 |

| | | | | | |
|---|---|---|---|---|---|
| orf1a | 195 | 1744 | 0 | FALSE | 3 |
| orf1a | 196 | 1744 | 0 | FALSE | 4 |
| orf1a | 197 | 1744 | 0 | FALSE | 4 |
| orf1a | 198 | 1744 | 0 | FALSE | 4 |
| orf1a | 199 | 1744 | 0 | FALSE | 6 |
| orf1a | 200 | 1744 | 0 | FALSE | 3 |
| orf1a | 201 | 1744 | 0 | FALSE | 0 |
| orf1a | 202 | 1744 | 0 | FALSE | 3 |
| orf1a | 203 | 1744 | 0 | FALSE | 2 |
| orf1a | 204 | 1744 | 0 | FALSE | 3 |
| orf1a | 205 | 1744 | 0 | FALSE | 3 |
| orf1a | 206 | 1744 | 0 | FALSE | 5 |
| orf1a | 207 | 1210 | 0 | FALSE | 6 |
| orf1a | 208 | 1210 | 0 | FALSE | 3 |
| orf1a | 216 | 1349 | 0 | FALSE | 4 |
| orf1a | 217 | 1349 | 0 | FALSE | 3 |
| orf1a | 218 | 1349 | 0 | FALSE | 2 |
| orf1a | 219 | 1349 | 0 | FALSE | 5 |
| orf1a | 220 | 1349 | 0 | FALSE | 6 |
| orf1a | 221 | 1349 | 0 | FALSE | 2 |
| orf1a | 222 | 1452 | 0 | FALSE | 3 |
| orf1a | 223 | 1452 | 0 | FALSE | 5 |
| orf1a | 224 | 1452 | 0 | FALSE | 3 |
| orf1a | 225 | 1452 | 0 | FALSE | 4 |
| orf1a | 226 | 1452 | 0 | FALSE | 3 |
| orf1a | 227 | 1452 | 0 | FALSE | 3 |
| orf1a | 228 | 1452 | 0 | FALSE | 3 |
| orf1a | 229 | 1452 | 0 | FALSE | 4 |
| orf1a | 230 | 1466 | 1156 | FALSE | 3 |
| orf1a | 231 | 1486 | 1156 | FALSE | 4 |
| orf1a | 232 | 2693 | 1681 | FALSE | 3 |
| orf1a | 233 | 2693 | 1681 | FALSE | 4 |
| orf1a | 234 | 2481 | 1681 | FALSE | 3 |
| orf1a | 235 | 2481 | 1681 | FALSE | 4 |
| orf1a | 236 | 2481 | 1681 | FALSE | 1 |
| orf1a | 237 | 2552 | 1681 | FALSE | 3 |
| orf1a | 238 | 2552 | 1681 | FALSE | 3 |
| orf1a | 239 | 2552 | 1681 | FALSE | 2 |
| orf1a | 240 | 2552 | 1681 | FALSE | 1 |
| orf1a | 241 | 2552 | 1681 | FALSE | 3 |
| orf1a | 242 | 2873 | 1581 | FALSE | 3 |
| orf1a | 243 | 2873 | 1581 | FALSE | 0 |
| orf1a | 244 | 2873 | 2235 | FALSE | 3 |
| orf1a | 245 | 3573 | 2667 | FALSE | 5 |
| orf1a | 246 | 4927 | 4042 | FALSE | 3 |
| orf1a | 247 | 6159 | 6052 | FALSE | 6 |
| orf1a | 248 | 6608 | 4881 | FALSE | 4 |
| orf1a | 249 | 9308 | 4881 | FALSE | 3 |
| orf1a | 250 | 8975 | 4634 | FALSE | 4 |
| orf1a | 251 | 8975 | 4634 | FALSE | 2 |
| orf1a | 252 | 8975 | 4634 | FALSE | 3 |
| orf1a | 253 | 8975 | 4634 | FALSE | 3 |
| orf1a | 254 | 8566 | 4634 | FALSE | 4 |
| orf1a | 255 | 8566 | 4634 | FALSE | 0 |
| orf1a | 256 | 8898 | 4477 | FALSE | 2 |
| orf1a | 257 | 10475 | 5043 | FALSE | 4 |
| orf1a | 258 | 10475 | 5043 | FALSE | 2 |
| orf1a | 259 | 10475 | 4821 | FALSE | 4 |
| orf1a | 260 | 10475 | 4821 | FALSE | 3 |
| orf1a | 261 | 8476 | 3297 | FALSE | 2 |
| orf1a | 262 | 8476 | 3297 | FALSE | 1 |
| orf1a | 263 | 8351 | 3940 | FALSE | 2 |

| | | | | | |
|---|---|---|---|---|---|
| orf1a | 264 | 8351 | 3940 | FALSE | 2 |
| orf1a | 265 | 8706 | 3537 | FALSE | 3 |
| orf1a | 266 | 8706 | 3537 | FALSE | 4 |
| orf1a | 267 | 8706 | 3537 | FALSE | 3 |
| orf1a | 268 | 8706 | 2278 | FALSE | 5 |
| orf1a | 269 | 10890 | 2278 | FALSE | 2 |
| orf1a | 270 | 10890 | 2278 | FALSE | 1 |
| orf1a | 271 | 0 | 1020 | FALSE | 3 |
| orf1a | 272 | 1497 | 1020 | FALSE | 4 |
| orf1a | 273 | 1497 | 1020 | FALSE | 2 |
| orf1a | 274 | 1497 | 1020 | FALSE | 4 |
| orf1a | 275 | 1497 | 1020 | FALSE | 3 |
| orf1a | 276 | 1497 | 1020 | FALSE | 3 |
| orf1a | 277 | 1497 | 1020 | FALSE | 2 |
| orf1a | 278 | 1497 | 1020 | FALSE | 2 |
| orf1a | 279 | 1497 | 1020 | FALSE | 3 |
| orf1a | 280 | 1497 | 1020 | FALSE | 2 |
| orf1a | 281 | 1497 | 1020 | FALSE | 4 |
| orf1a | 282 | 5608 | 1020 | FALSE | 1 |
| orf1a | 283 | 5608 | 0 | FALSE | 2 |
| orf1a | 284 | 5608 | 0 | FALSE | 3 |
| orf1a | 285 | 5608 | 0 | FALSE | 2 |
| orf1a | 286 | 5608 | 0 | FALSE | 3 |
| orf1a | 287 | 9720 | 0 | FALSE | 6 |
| orf1a | 288 | 9720 | 0 | FALSE | 3 |
| orf1a | 289 | 9720 | 0 | FALSE | 3 |
| orf1a | 290 | 9720 | 4368 | FALSE | 5 |
| orf1a | 291 | 9720 | 4368 | FALSE | 3 |
| orf1a | 292 | 9720 | 4368 | FALSE | 1 |
| orf1a | 293 | 9720 | 4368 | FALSE | 1 |
| orf1a | 294 | 5602 | 4368 | FALSE | 2 |
| orf1a | 295 | 5602 | 4368 | FALSE | 5 |
| orf1a | 296 | 5602 | 4368 | FALSE | 3 |
| orf1a | 297 | 1483 | 4368 | FALSE | 1 |
| orf1a | 298 | 1483 | 4368 | FALSE | 4 |
| orf1a | 299 | 1483 | 4368 | FALSE | 1 |
| orf1a | 300 | 1483 | 4368 | FALSE | 3 |
| orf1a | 301 | 1483 | 4368 | FALSE | 6 |
| orf1a | 302 | 1483 | 4368 | FALSE | 2 |
| orf1a | 303 | 1483 | 4368 | FALSE | 3 |
| orf1a | 304 | 1483 | 4368 | FALSE | 1 |
| orf1a | 305 | 1483 | 0 | FALSE | 2 |
| orf1a | 306 | 1483 | 0 | FALSE | 1 |
| orf1a | 307 | 1483 | 0 | FALSE | 7 |
| orf1a | 308 | 1483 | 0 | FALSE | 3 |
| orf1a | 310 | 1079 | 0 | FALSE | 3 |
| orf1a | 311 | 1079 | 0 | FALSE | 2 |
| orf1a | 312 | 1079 | 0 | FALSE | 4 |
| orf1a | 313 | 1079 | 0 | FALSE | 2 |
| orf1a | 314 | 1079 | 0 | FALSE | 4 |
| orf1a | 315 | 1079 | 0 | FALSE | 2 |
| orf1a | 316 | 1079 | 0 | FALSE | 3 |
| orf1a | 317 | 1079 | 0 | FALSE | 3 |
| orf1a | 318 | 1079 | 0 | FALSE | 2 |
| orf1a | 319 | 1079 | 0 | FALSE | 2 |
| orf1a | 320 | 1079 | 0 | FALSE | 2 |
| orf1a | 321 | 1079 | 0 | FALSE | 3 |
| orf1a | 322 | 1275 | 0 | FALSE | 3 |
| orf1a | 323 | 1275 | 0 | FALSE | 2 |
| orf1a | 324 | 1275 | 0 | FALSE | 3 |
| orf1a | 325 | 1470 | 0 | FALSE | 5 |
| orf1a | 326 | 2079 | 1459 | FALSE | 4 |

| | | | | | |
|---|---|---|---|---|---|
| orf1a | 327 | 2079 | 1459 | FALSE | 3 |
| orf1a | 328 | 5491 | 1963 | FALSE | 1 |
| orf1a | 329 | 5491 | 1963 | FALSE | 3 |
| orf1a | 330 | 5491 | 1963 | FALSE | 4 |
| orf1a | 331 | 5491 | 1963 | FALSE | 4 |
| orf1a | 332 | 5491 | 1963 | FALSE | 2 |
| orf1a | 333 | 5491 | 1963 | FALSE | 3 |
| orf1a | 334 | 5491 | 1963 | FALSE | 4 |
| orf1a | 335 | 5491 | 1963 | FALSE | 4 |
| orf1a | 336 | 5491 | 1963 | FALSE | 1 |
| orf1a | 337 | 6497 | 1963 | FALSE | 2 |
| orf1a | 338 | 6497 | 1963 | FALSE | 3 |
| orf1a | 339 | 6497 | 1963 | FALSE | 3 |
| orf1a | 340 | 5402 | 1963 | FALSE | 1 |
| orf1a | 341 | 6081 | 2468 | FALSE | 2 |
| orf1a | 342 | 6446 | 3670 | FALSE | 6 |
| orf1a | 343 | 4465 | 6075 | FALSE | 4 |
| orf1a | 344 | 4465 | 6075 | FALSE | 2 |
| orf1a | 345 | 4465 | 6075 | FALSE | 3 |
| orf1a | 346 | 4465 | 6075 | FALSE | 1 |
| orf1a | 347 | 4465 | 6075 | FALSE | 5 |
| orf1a | 348 | 4798 | 6075 | FALSE | 6 |
| orf1a | 349 | 4798 | 6075 | FALSE | 3 |
| orf1a | 350 | 4798 | 4280 | FALSE | 2 |
| orf1a | 351 | 4798 | 4280 | FALSE | 0 |
| orf1a | 352 | 4275 | 7827 | FALSE | 5 |
| orf1a | 353 | 4275 | 7827 | FALSE | 3 |
| orf1a | 354 | 4275 | 7827 | FALSE | 2 |
| orf1a | 355 | 4926 | 7827 | FALSE | 2 |
| orf1a | 356 | 4926 | 7827 | FALSE | 3 |
| orf1a | 357 | 4181 | 8411 | FALSE | 3 |
| orf1a | 358 | 4181 | 8411 | FALSE | 4 |
| orf1a | 359 | 4181 | 8411 | FALSE | 1 |
| orf1a | 360 | 4181 | 8411 | FALSE | 1 |
| orf1a | 361 | 4181 | 8411 | FALSE | 3 |
| orf1a | 362 | 3667 | 8411 | FALSE | 4 |
| orf1a | 363 | 3218 | 8411 | FALSE | 3 |
| orf1a | 364 | 2959 | 8411 | FALSE | 4 |
| orf1a | 365 | 2959 | 11375 | FALSE | 3 |
| orf1a | 366 | 2959 | 11375 | FALSE | 4 |
| orf1a | 367 | 1768 | 0 | FALSE | 4 |
| orf1a | 368 | 1952 | 0 | FALSE | 3 |
| orf1a | 369 | 1952 | 0 | FALSE | 1 |
| orf1a | 370 | 1952 | 0 | FALSE | 3 |
| orf1a | 371 | 1952 | 0 | FALSE | 2 |
| orf1a | 372 | 1952 | 0 | FALSE | 2 |
| orf1a | 373 | 1952 | 0 | FALSE | 1 |
| orf1a | 374 | 1756 | 0 | FALSE | 2 |
| orf1a | 375 | 1756 | 0 | FALSE | 2 |
| orf1a | 376 | 1756 | 0 | FALSE | 3 |
| orf1a | 377 | 1784 | 0 | FALSE | 3 |
| orf1a | 378 | 1840 | 0 | FALSE | 1 |
| orf1a | 379 | 1823 | 0 | FALSE | 6 |
| orf1a | 380 | 1786 | 0 | FALSE | 6 |
| orf1a | 381 | 1786 | 0 | FALSE | 4 |
| orf1a | 382 | 1786 | 0 | FALSE | 2 |
| orf1a | 383 | 1679 | 0 | FALSE | 2 |
| orf1a | 384 | 1679 | 0 | FALSE | 3 |
| orf1a | 385 | 1679 | 0 | FALSE | 4 |
| orf1a | 386 | 3661 | 7707 | FALSE | 6 |
| orf1a | 387 | 3661 | 7707 | FALSE | 1 |
| orf1a | 388 | 3661 | 7707 | FALSE | 2 |

| | | | | | |
|---|---|---|---|---|---|
| orf1a | 389 | 4160 | 7707 | FALSE | 3 |
| orf1a | 390 | 6265 | 8146 | FALSE | 4 |
| orf1a | 391 | 8519 | 8146 | FALSE | 2 |
| orf1a | 392 | 8519 | 8146 | FALSE | 5 |
| orf1a | 393 | 10653 | 8146 | FALSE | 4 |
| orf1a | 394 | 10653 | 8146 | FALSE | 1 |
| orf1a | 395 | 15179 | 8146 | FALSE | 4 |
| orf1a | 396 | 15179 | 8146 | FALSE | 1 |
| orf1a | 397 | 15179 | 8146 | FALSE | 3 |
| orf1a | 398 | 15179 | 8146 | FALSE | 4 |
| orf1a | 399 | 15179 | 8146 | FALSE | 5 |
| orf1a | 400 | 15179 | 8146 | FALSE | 1 |
| orf1a | 401 | 16787 | 8586 | FALSE | 3 |
| orf1a | 402 | 16787 | 8586 | FALSE | 3 |
| orf1a | 403 | 16787 | 8586 | FALSE | 3 |
| orf1a | 404 | 16787 | 8586 | FALSE | 1 |
| orf1a | 430 | 0 | 1485 | FALSE | 1 |
| orf1a | 431 | 0 | 1485 | FALSE | 1 |
| orf1a | 432 | 0 | 2540 | FALSE | 3 |
| orf1a | 433 | 0 | 2540 | FALSE | 4 |
| orf1a | 434 | 1874 | 2730 | FALSE | 1 |
| orf1a | 435 | 1874 | 2730 | FALSE | 2 |
| orf1a | 436 | 1874 | 2082 | FALSE | 4 |
| orf1a | 437 | 1874 | 2082 | FALSE | 4 |
| orf1a | 438 | 1956 | 2107 | FALSE | 4 |
| orf1a | 439 | 1956 | 2107 | FALSE | 2 |
| orf1a | 440 | 1813 | 2178 | FALSE | 5 |
| orf1a | 441 | 1813 | 2178 | FALSE | 2 |
| orf1a | 442 | 1813 | 2178 | FALSE | 7 |
| orf1a | 443 | 1813 | 2178 | FALSE | 4 |
| orf1a | 444 | 3917 | 4326 | FALSE | 6 |
| orf1a | 445 | 3917 | 4610 | FALSE | 5 |
| orf1a | 446 | 3917 | 4610 | FALSE | 5 |
| orf1a | 447 | 3917 | 4722 | FALSE | 5 |
| orf1a | 448 | 3917 | 4722 | FALSE | 6 |
| orf1a | 449 | 4734 | 4924 | FALSE | 2 |
| orf1a | 450 | 4734 | 4924 | FALSE | 4 |
| orf1a | 451 | 4734 | 6195 | FALSE | 3 |
| orf1a | 452 | 6287 | 5869 | FALSE | 3 |
| orf1a | 453 | 7120 | 7103 | FALSE | 3 |
| orf1a | 454 | 7120 | 7103 | FALSE | 4 |
| orf1a | 455 | 8554 | 7988 | FALSE | 4 |
| orf1a | 456 | 8554 | 7988 | FALSE | 3 |
| orf1a | 457 | 8554 | 7988 | FALSE | 4 |
| orf1a | 458 | 7673 | 7988 | FALSE | 3 |
| orf1a | 459 | 9098 | 4892 | FALSE | 2 |
| orf1a | 460 | 6604 | 4892 | FALSE | 3 |
| orf1a | 461 | 6604 | 4892 | FALSE | 4 |
| orf1a | 462 | 5376 | 4892 | FALSE | 3 |
| orf1a | 463 | 5376 | 4892 | FALSE | 3 |
| orf1a | 464 | 5376 | 4892 | FALSE | 4 |
| orf1a | 465 | 5376 | 4892 | FALSE | 4 |
| orf1a | 466 | 5376 | 4892 | FALSE | 1 |
| orf1a | 467 | 3208 | 0 | FALSE | 3 |
| orf1a | 468 | 3208 | 0 | FALSE | 3 |
| orf1a | 469 | 3208 | 0 | FALSE | 3 |
| orf1a | 470 | 2814 | 2480 | FALSE | 2 |
| orf1a | 471 | 2814 | 2480 | FALSE | 4 |
| orf1a | 472 | 2814 | 2480 | FALSE | 2 |
| orf1a | 473 | 2480 | 2480 | FALSE | 1 |
| orf1a | 474 | 2480 | 2480 | FALSE | 3 |
| orf1a | 475 | 2768 | 2480 | FALSE | 1 |

| | | | | | |
|---|---|---|---|---|---|
| orf1a | 476 | 2768 | 2480 | FALSE | 1 |
| orf1a | 477 | 1235 | 2480 | FALSE | 2 |
| orf1a | 478 | 1235 | 2480 | FALSE | 2 |
| orf1a | 479 | 1235 | 2480 | FALSE | 3 |
| orf1a | 480 | 1235 | 2480 | FALSE | 4 |
| orf1a | 481 | 1235 | 2480 | FALSE | 1 |
| orf1a | 482 | 1235 | 1741 | FALSE | 2 |
| orf1a | 483 | 1235 | 1741 | FALSE | 3 |
| orf1a | 484 | 1235 | 1741 | FALSE | 3 |
| orf1a | 485 | 0 | 1001 | FALSE | 4 |
| orf1a | 486 | 0 | 1001 | FALSE | 5 |
| orf1a | 487 | 0 | 1001 | FALSE | 3 |
| orf1a | 488 | 1880 | 1001 | FALSE | 2 |
| orf1a | 489 | 1880 | 1001 | FALSE | 2 |
| orf1a | 490 | 1880 | 1001 | FALSE | 2 |
| orf1a | 491 | 1880 | 1001 | FALSE | 4 |
| orf1a | 492 | 1880 | 1001 | FALSE | 2 |
| orf1a | 493 | 1880 | 1001 | FALSE | 5 |
| orf1a | 494 | 1880 | 1001 | FALSE | 2 |
| orf1a | 495 | 1880 | 1001 | FALSE | 2 |
| orf1a | 496 | 1880 | 1001 | FALSE | 3 |
| orf1a | 497 | 1880 | 0 | FALSE | 4 |
| orf1a | 498 | 1880 | 0 | FALSE | 5 |
| orf1a | 499 | 1880 | 0 | FALSE | 3 |
| orf1a | 500 | 1880 | 0 | FALSE | 0 |
| orf1a | 501 | 1880 | 0 | FALSE | 1 |
| orf1a | 502 | 6843 | 0 | FALSE | 2 |
| orf1a | 503 | 11807 | 0 | FALSE | 1 |
| orf1a | 504 | 11807 | 5826 | FALSE | 3 |
| orf1a | 505 | 11807 | 5826 | FALSE | 2 |
| orf1a | 506 | 11807 | 5826 | FALSE | 4 |
| orf1a | 507 | 11807 | 5826 | FALSE | 4 |
| orf1a | 508 | 11807 | 5826 | FALSE | 2 |
| orf1a | 509 | 11807 | 5826 | FALSE | 2 |
| orf1a | 510 | 11807 | 5826 | FALSE | 2 |
| orf1a | 511 | 11807 | 5826 | FALSE | 4 |
| orf1a | 512 | 11807 | 5826 | FALSE | 4 |
| orf1a | 513 | 11807 | 5826 | FALSE | 1 |
| orf1a | 514 | 11807 | 5826 | FALSE | 5 |
| orf1a | 515 | 11807 | 5826 | FALSE | 1 |
| orf1a | 516 | 11807 | 5826 | FALSE | 6 |
| orf1a | 517 | 0 | 5826 | FALSE | 2 |
| orf1a | 518 | 0 | 5826 | FALSE | 3 |
| orf1a | 534 | 0 | 1108 | FALSE | 2 |
| orf1a | 535 | 0 | 1108 | FALSE | 2 |
| orf1a | 536 | 1142 | 1108 | FALSE | 1 |
| orf1a | 537 | 1142 | 1108 | FALSE | 3 |
| orf1a | 538 | 1142 | 1108 | FALSE | 2 |
| orf1a | 539 | 1142 | 1108 | FALSE | 4 |
| orf1a | 540 | 1142 | 1108 | FALSE | 3 |
| orf1a | 541 | 1142 | 1108 | FALSE | 5 |
| orf1a | 542 | 1142 | 1108 | FALSE | 2 |
| orf1a | 543 | 1142 | 1108 | FALSE | 1 |
| orf1a | 544 | 1142 | 1108 | FALSE | 4 |
| orf1a | 545 | 1142 | 1108 | FALSE | 5 |
| orf1a | 546 | 1142 | 1108 | FALSE | 3 |
| orf1a | 547 | 1142 | 1108 | FALSE | 4 |
| orf1a | 548 | 1142 | 1108 | FALSE | 1 |
| orf1a | 549 | 1142 | 0 | FALSE | 4 |
| orf1a | 550 | 1142 | 1985 | FALSE | 2 |
| orf1a | 551 | 0 | 1985 | FALSE | 2 |
| orf1a | 552 | 0 | 1985 | FALSE | 3 |

| | | | | | |
|---|---|---|---|---|---|
| orf1a | 553 | 0 | 1985 | FALSE | 2 |
| orf1a | 554 | 0 | 1985 | FALSE | 5 |
| orf1a | 555 | 0 | 1985 | FALSE | 5 |
| orf1a | 556 | 0 | 1985 | FALSE | 5 |
| orf1a | 557 | 0 | 1985 | FALSE | 4 |
| orf1a | 558 | 0 | 1985 | FALSE | 5 |
| orf1a | 559 | 0 | 1985 | FALSE | 2 |
| orf1a | 560 | 0 | 1985 | FALSE | 2 |
| orf1a | 561 | 0 | 1985 | FALSE | 2 |
| orf1a | 562 | 0 | 1985 | FALSE | 0 |
| orf1a | 563 | 0 | 1985 | FALSE | 1 |
| orf1a | 564 | 0 | 1985 | FALSE | 2 |
| orf1a | 570 | 0 | 1523 | FALSE | 2 |
| orf1a | 571 | 0 | 1523 | FALSE | 1 |
| orf1a | 572 | 0 | 2442 | FALSE | 3 |
| orf1a | 573 | 0 | 2442 | FALSE | 3 |
| orf1a | 574 | 0 | 2442 | FALSE | 2 |
| orf1a | 575 | 0 | 2442 | FALSE | 3 |
| orf1a | 576 | 0 | 2442 | FALSE | 2 |
| orf1a | 577 | 0 | 2442 | FALSE | 3 |
| orf1a | 578 | 0 | 2442 | FALSE | 3 |
| orf1a | 579 | 0 | 2442 | FALSE | 4 |
| orf1a | 580 | 0 | 2442 | FALSE | 3 |
| orf1a | 581 | 0 | 2442 | FALSE | 4 |
| orf1a | 582 | 3451 | 2442 | FALSE | 1 |
| orf1a | 583 | 3451 | 2442 | FALSE | 3 |
| orf1a | 584 | 3451 | 2442 | FALSE | 2 |
| orf1a | 585 | 3451 | 3361 | FALSE | 1 |
| orf1a | 586 | 3451 | 3361 | FALSE | 4 |
| orf1a | 587 | 3451 | 0 | FALSE | 2 |
| orf1a | 588 | 4585 | 0 | FALSE | 1 |
| orf1a | 589 | 4585 | 0 | FALSE | 6 |
| orf1a | 590 | 4585 | 0 | FALSE | 4 |
| orf1a | 591 | 4585 | 0 | FALSE | 3 |
| orf1a | 592 | 4585 | 2052 | FALSE | 2 |
| orf1a | 593 | 4585 | 2052 | FALSE | 3 |
| orf1a | 594 | 7773 | 2052 | FALSE | 2 |
| orf1a | 595 | 7773 | 2052 | FALSE | 1 |
| orf1a | 596 | 6566 | 2052 | FALSE | 0 |
| orf1a | 597 | 8643 | 2052 | FALSE | 2 |
| orf1a | 598 | 8643 | 2052 | FALSE | 1 |
| orf1a | 599 | 8643 | 2052 | FALSE | 3 |
| orf1a | 600 | 8643 | 2052 | FALSE | 3 |
| orf1a | 601 | 8643 | 2052 | FALSE | 2 |
| orf1a | 602 | 8643 | 2052 | FALSE | 1 |
| orf1a | 603 | 9539 | 2052 | FALSE | 3 |
| orf1a | 604 | 9539 | 4044 | FALSE | 5 |
| orf1a | 605 | 9539 | 4044 | FALSE | 4 |
| orf1a | 606 | 9539 | 4044 | FALSE | 2 |
| orf1a | 607 | 9539 | 6036 | FALSE | 2 |
| orf1a | 608 | 9539 | 6036 | FALSE | 3 |
| orf1a | 609 | 1740 | 6036 | FALSE | 3 |
| orf1a | 610 | 1740 | 6036 | FALSE | 2 |
| orf1a | 611 | 0 | 6036 | FALSE | 3 |
| orf1a | 612 | 0 | 6036 | FALSE | 4 |
| orf1a | 613 | 0 | 6036 | FALSE | 2 |
| orf1a | 614 | 0 | 6036 | FALSE | 3 |
| orf1a | 615 | 0 | 6036 | FALSE | 1 |
| orf1a | 616 | 0 | 6036 | FALSE | 4 |
| orf1a | 617 | 0 | 6036 | FALSE | 0 |
| orf1a | 618 | 0 | 4409 | FALSE | 5 |
| orf1a | 619 | 0 | 2781 | FALSE | 3 |

| | | | | | |
|---|---|---|---|---|---|
| orf1a | 620 | 0 | 1902 | FALSE | 3 |
| orf1a | 621 | 0 | 1902 | FALSE | 3 |
| orf1a | 622 | 11608 | 3204 | FALSE | 4 |
| orf1a | 623 | 11608 | 3204 | FALSE | 1 |
| orf1a | 624 | 11608 | 3204 | FALSE | 3 |
| orf1a | 625 | 11608 | 3204 | FALSE | 0 |
| orf1a | 626 | 5102 | 2497 | FALSE | 5 |
| orf1a | 627 | 5102 | 2497 | FALSE | 3 |
| orf1a | 628 | 5102 | 2497 | FALSE | 1 |
| orf1a | 629 | 5102 | 2497 | FALSE | 2 |
| orf1a | 630 | 6892 | 2465 | FALSE | 3 |
| orf1a | 631 | 6892 | 2465 | FALSE | 2 |
| orf1a | 632 | 10026 | 4539 | FALSE | 4 |
| orf1a | 633 | 10026 | 4734 | FALSE | 4 |
| orf1a | 634 | 10026 | 4734 | FALSE | 2 |
| orf1a | 635 | 10026 | 5198 | FALSE | 3 |
| orf1a | 636 | 9223 | 4978 | FALSE | 3 |
| orf1a | 637 | 8984 | 4875 | FALSE | 1 |
| orf1a | 638 | 8984 | 5815 | FALSE | 2 |
| orf1a | 639 | 8984 | 5815 | FALSE | 3 |
| orf1a | 640 | 8861 | 5776 | FALSE | 4 |
| orf1a | 641 | 11939 | 7484 | FALSE | 1 |
| orf1a | 642 | 11939 | 7484 | FALSE | 1 |
| orf1a | 643 | 11939 | 7484 | FALSE | 1 |
| orf1a | 644 | 11939 | 7484 | FALSE | 3 |
| orf1a | 645 | 11289 | 8352 | FALSE | 1 |
| orf1a | 646 | 11289 | 8352 | FALSE | 1 |
| orf1a | 647 | 4410 | 7328 | FALSE | 3 |
| orf1a | 648 | 4410 | 7328 | FALSE | 5 |
| orf1a | 649 | 4410 | 7328 | FALSE | 5 |
| orf1a | 650 | 4410 | 7328 | FALSE | 5 |
| orf1a | 651 | 7633 | 9380 | FALSE | 2 |
| orf1a | 652 | 7633 | 9380 | FALSE | 5 |
| orf1a | 653 | 7633 | 5423 | FALSE | 2 |
| orf1a | 654 | 7633 | 5423 | FALSE | 4 |
| orf1a | 658 | 0 | 1571 | FALSE | 3 |
| orf1a | 659 | 0 | 1571 | FALSE | 2 |
| orf1a | 660 | 0 | 1571 | FALSE | 3 |
| orf1a | 661 | 0 | 1571 | FALSE | 6 |
| orf1a | 662 | 0 | 1571 | FALSE | 3 |
| orf1a | 663 | 0 | 1571 | FALSE | 3 |
| orf1a | 664 | 0 | 1571 | FALSE | 0 |
| orf1a | 665 | 0 | 1571 | FALSE | 4 |
| orf1a | 666 | 0 | 1571 | FALSE | 2 |
| orf1a | 667 | 0 | 1571 | FALSE | 2 |
| orf1a | 668 | 0 | 1571 | FALSE | 4 |
| orf1a | 669 | 0 | 1571 | FALSE | 4 |
| orf1a | 670 | 9709 | 2985 | FALSE | 5 |
| orf1a | 671 | 9709 | 2985 | FALSE | 4 |
| orf1a | 672 | 9709 | 2985 | FALSE | 3 |
| orf1a | 673 | 9709 | 4399 | FALSE | 3 |
| orf1a | 674 | 9709 | 4399 | FALSE | 4 |
| orf1a | 675 | 9709 | 4399 | FALSE | 3 |
| orf1a | 676 | 9709 | 4399 | FALSE | 5 |
| orf1a | 677 | 9709 | 4399 | FALSE | 3 |
| orf1a | 678 | 9709 | 4399 | FALSE | 1 |
| orf1a | 679 | 9709 | 4399 | FALSE | 2 |
| orf1a | 680 | 9709 | 4399 | FALSE | 5 |
| orf1a | 681 | 9709 | 4399 | FALSE | 2 |
| orf1a | 682 | 9709 | 4399 | FALSE | 1 |
| orf1a | 683 | 9709 | 4399 | FALSE | 1 |
| orf1a | 684 | 9709 | 4399 | FALSE | 3 |

| | | | | | |
|---|---|---|---|---|---|
| orf1a | 704 | 1937 | 0 | FALSE | 2 |
| orf1a | 705 | 1937 | 0 | FALSE | 2 |
| orf1a | 706 | 1937 | 1217 | FALSE | 7 |
| orf1a | 707 | 1937 | 1217 | FALSE | 6 |
| orf1a | 708 | 1937 | 1217 | FALSE | 5 |
| orf1a | 709 | 1937 | 1217 | FALSE | 1 |
| orf1a | 710 | 1937 | 1217 | FALSE | 4 |
| orf1a | 711 | 1937 | 1217 | FALSE | 4 |
| orf1a | 712 | 1937 | 1217 | FALSE | 4 |
| orf1a | 713 | 1937 | 1217 | FALSE | 7 |
| orf1a | 714 | 1937 | 1217 | FALSE | 6 |
| orf1a | 715 | 1937 | 1217 | FALSE | 3 |
| orf1a | 716 | 1937 | 1217 | FALSE | 3 |
| orf1a | 717 | 1937 | 1217 | FALSE | 3 |
| orf1a | 718 | 1937 | 1217 | FALSE | 2 |
| orf1a | 719 | 0 | 1217 | FALSE | 3 |
| orf1a | 720 | 0 | 1217 | FALSE | 3 |
| orf1a | 724 | 2706 | 0 | FALSE | 7 |
| orf1a | 725 | 2706 | 0 | FALSE | 6 |
| orf1a | 726 | 2706 | 2185 | FALSE | 3 |
| orf1a | 727 | 2706 | 2185 | FALSE | 3 |
| orf1a | 728 | 2706 | 2392 | FALSE | 3 |
| orf1a | 729 | 2706 | 2392 | FALSE | 1 |
| orf1a | 730 | 1752 | 2392 | FALSE | 4 |
| orf1a | 731 | 1752 | 2392 | FALSE | 3 |
| orf1a | 732 | 1467 | 2392 | FALSE | 4 |
| orf1a | 733 | 1467 | 2392 | FALSE | 2 |
| orf1a | 734 | 1467 | 2392 | FALSE | 4 |
| orf1a | 735 | 1467 | 2392 | FALSE | 5 |
| orf1a | 736 | 2150 | 1763 | FALSE | 6 |
| orf1a | 737 | 2150 | 1763 | FALSE | 2 |
| orf1a | 738 | 3296 | 2799 | FALSE | 4 |
| orf1a | 739 | 3361 | 2799 | FALSE | 2 |
| orf1a | 740 | 3361 | 2666 | FALSE | 3 |
| orf1a | 741 | 3361 | 2735 | FALSE | 2 |
| orf1a | 742 | 3262 | 2735 | FALSE | 2 |
| orf1a | 743 | 3262 | 2757 | FALSE | 2 |
| orf1a | 744 | 2845 | 2790 | FALSE | 1 |
| orf1a | 745 | 3130 | 2790 | FALSE | 3 |
| orf1a | 746 | 3133 | 2490 | FALSE | 1 |
| orf1a | 747 | 3482 | 2490 | FALSE | 1 |
| orf1a | 748 | 4394 | 2515 | FALSE | 4 |
| orf1a | 749 | 4394 | 2515 | FALSE | 4 |
| orf1a | 750 | 5333 | 4067 | FALSE | 2 |
| orf1a | 751 | 5322 | 4486 | FALSE | 3 |
| orf1a | 752 | 6134 | 4651 | FALSE | 2 |
| orf1a | 753 | 6383 | 4759 | FALSE | 4 |
| orf1a | 754 | 7755 | 5199 | FALSE | 4 |
| orf1a | 755 | 7755 | 5415 | FALSE | 3 |
| orf1a | 756 | 7755 | 5415 | FALSE | 2 |
| orf1a | 757 | 8012 | 5415 | FALSE | 4 |
| orf1a | 758 | 8012 | 5415 | FALSE | 4 |
| orf1a | 759 | 9104 | 5577 | FALSE | 1 |
| orf1a | 760 | 8124 | 5577 | FALSE | 3 |
| orf1a | 761 | 8911 | 6890 | FALSE | 0 |
| orf1a | 762 | 8123 | 6890 | FALSE | 4 |
| orf1a | 763 | 8837 | 7832 | FALSE | 3 |
| orf1a | 764 | 7758 | 7832 | FALSE | 2 |
| orf1a | 765 | 7278 | 6351 | FALSE | 2 |
| orf1a | 766 | 6635 | 6351 | FALSE | 3 |
| orf1a | 767 | 5601 | 6626 | FALSE | 3 |
| orf1a | 768 | 5221 | 6626 | FALSE | 2 |

| | | | | | |
|---|---|---|---|---|---|
| orf1a | 769 | 2733 | 0 | FALSE | 2 |
| orf1a | 770 | 2720 | 0 | FALSE | 2 |
| orf1a | 771 | 2720 | 0 | FALSE | 3 |
| orf1a | 772 | 3661 | 12251 | FALSE | 1 |
| orf1a | 773 | 3661 | 12251 | FALSE | 1 |
| orf1a | 774 | 3661 | 12251 | FALSE | 3 |
| orf1a | 775 | 3645 | 12251 | FALSE | 2 |
| orf1a | 776 | 3645 | 12251 | FALSE | 3 |
| orf1a | 777 | 3799 | 12251 | FALSE | 3 |
| orf1a | 778 | 3799 | 8533 | FALSE | 3 |
| orf1a | 779 | 4568 | 8533 | FALSE | 1 |
| orf1a | 780 | 4568 | 8533 | FALSE | 1 |
| orf1a | 781 | 5669 | 8533 | FALSE | 1 |
| orf1a | 782 | 5669 | 8533 | FALSE | 3 |
| orf1a | 783 | 7993 | 8533 | FALSE | 2 |
| orf1a | 784 | 7993 | 8533 | FALSE | 1 |
| orf1a | 785 | 10717 | 8533 | FALSE | 2 |
| orf1a | 786 | 7741 | 8533 | FALSE | 2 |
| orf1a | 787 | 1788 | 1097 | FALSE | 2 |
| orf1a | 788 | 1788 | 1097 | FALSE | 4 |
| orf1a | 789 | 1788 | 1097 | FALSE | 2 |
| orf1a | 790 | 4395 | 1097 | FALSE | 2 |
| orf1a | 791 | 4395 | 1097 | FALSE | 1 |
| orf1a | 792 | 7546 | 3490 | FALSE | 1 |
| orf1a | 793 | 7546 | 5884 | FALSE | 1 |
| orf1a | 794 | 7546 | 5884 | FALSE | 2 |
| orf1a | 795 | 7546 | 5884 | FALSE | 2 |
| orf1a | 796 | 5942 | 5884 | FALSE | 1 |
| orf1a | 797 | 5942 | 5884 | FALSE | 3 |
| orf1a | 798 | 5942 | 3758 | FALSE | 3 |
| orf1a | 799 | 5942 | 3758 | FALSE | 2 |
| orf1a | 800 | 5942 | 3758 | FALSE | 1 |
| orf1a | 801 | 7327 | 3758 | FALSE | 2 |
| orf1a | 802 | 7327 | 3758 | FALSE | 4 |
| orf1a | 803 | 7327 | 3758 | FALSE | 3 |
| orf1a | 804 | 7327 | 3758 | FALSE | 4 |
| orf1a | 805 | 7490 | 3758 | FALSE | 0 |
| orf1a | 806 | 7490 | 3758 | FALSE | 1 |
| orf1a | 807 | 1132 | 1633 | FALSE | 1 |
| orf1a | 808 | 1132 | 1633 | FALSE | 2 |
| orf1a | 809 | 1132 | 1633 | FALSE | 2 |
| orf1a | 810 | 1132 | 1633 | FALSE | 2 |
| orf1a | 811 | 0 | 1633 | FALSE | 1 |
| orf1a | 812 | 0 | 1633 | FALSE | 3 |
| orf1a | 818 | 1481 | 0 | FALSE | 1 |
| orf1a | 819 | 1481 | 0 | FALSE | 0 |
| orf1a | 820 | 1576 | 0 | FALSE | 3 |
| orf1a | 821 | 1576 | 0 | FALSE | 1 |
| orf1a | 822 | 1576 | 0 | FALSE | 2 |
| orf1a | 823 | 1576 | 0 | FALSE | 4 |
| orf1a | 824 | 1576 | 0 | FALSE | 3 |
| orf1a | 825 | 1576 | 0 | FALSE | 1 |
| orf1a | 826 | 1576 | 1167 | FALSE | 3 |
| orf1a | 827 | 1576 | 1167 | FALSE | 2 |
| orf1a | 828 | 1576 | 1167 | FALSE | 2 |
| orf1a | 829 | 1576 | 1167 | FALSE | 1 |
| orf1a | 830 | 4321 | 1167 | FALSE | 3 |
| orf1a | 831 | 4321 | 1167 | FALSE | 2 |
| orf1a | 832 | 5966 | 1272 | FALSE | 1 |
| orf1a | 833 | 8208 | 1272 | FALSE | 2 |
| orf1a | 834 | 8208 | 1272 | FALSE | 2 |
| orf1a | 835 | 10355 | 1272 | FALSE | 2 |

| | | | | | |
|---|---|---|---|---|---|
| orf1a | 836 | 10355 | 1272 | FALSE | 2 |
| orf1a | 837 | 10355 | 1272 | FALSE | 1 |
| orf1a | 838 | 10355 | 1272 | FALSE | 1 |
| orf1a | 839 | 10355 | 1272 | FALSE | 2 |
| orf1a | 840 | 8179 | 1272 | FALSE | 2 |
| orf1a | 841 | 8179 | 1325 | FALSE | 2 |
| orf1a | 842 | 5989 | 1431 | FALSE | 3 |
| orf1a | 843 | 5989 | 1431 | FALSE | 0 |
| orf1a | 844 | 6653 | 3227 | FALSE | 1 |
| orf1a | 845 | 6058 | 3227 | FALSE | 1 |
| orf1a | 846 | 6058 | 3227 | FALSE | 2 |
| orf1a | 847 | 4432 | 4178 | FALSE | 1 |
| orf1a | 848 | 4432 | 4178 | FALSE | 3 |
| orf1a | 849 | 4432 | 4178 | FALSE | 2 |
| orf1a | 850 | 4432 | 4178 | FALSE | 1 |
| orf1a | 851 | 4432 | 4178 | FALSE | 3 |
| orf1a | 852 | 4432 | 4178 | FALSE | 0 |
| orf1a | 853 | 4432 | 4178 | FALSE | 2 |
| orf1a | 854 | 3579 | 4178 | FALSE | 3 |
| orf1a | 855 | 3900 | 4178 | FALSE | 2 |
| orf1a | 856 | 3900 | 4178 | FALSE | 3 |
| orf1a | 857 | 5045 | 5023 | FALSE | 3 |
| orf1a | 858 | 5045 | 5023 | FALSE | 1 |
| orf1a | 859 | 1446 | 0 | FALSE | 2 |
| orf1a | 860 | 1446 | 2832 | FALSE | 1 |
| orf1a | 861 | 1446 | 2832 | FALSE | 1 |
| orf1a | 862 | 1446 | 2832 | FALSE | 0 |
| orf1a | 863 | 1446 | 2832 | FALSE | 2 |
| orf1a | 864 | 1446 | 2832 | FALSE | 3 |
| orf1a | 865 | 1446 | 2832 | FALSE | 5 |
| orf1a | 866 | 1218 | 2017 | FALSE | 3 |
| orf1a | 867 | 1218 | 2017 | FALSE | 2 |
| orf1a | 868 | 1218 | 2017 | FALSE | 5 |
| orf1a | 869 | 1066 | 2017 | FALSE | 4 |
| orf1a | 870 | 2930 | 2225 | FALSE | 2 |
| orf1a | 871 | 2930 | 2225 | FALSE | 1 |
| orf1a | 872 | 2930 | 2225 | FALSE | 4 |
| orf1a | 873 | 2930 | 2225 | FALSE | 1 |
| orf1a | 874 | 2930 | 2225 | FALSE | 3 |
| orf1a | 875 | 2930 | 1921 | FALSE | 4 |
| orf1a | 876 | 2930 | 1622 | FALSE | 2 |
| orf1a | 877 | 2930 | 1622 | FALSE | 0 |
| orf1a | 878 | 2930 | 1622 | FALSE | 4 |
| orf1a | 879 | 2930 | 1622 | FALSE | 3 |
| orf1a | 880 | 2930 | 1622 | FALSE | 1 |
| orf1a | 881 | 8523 | 1831 | FALSE | 1 |
| orf1a | 882 | 8523 | 2203 | FALSE | 2 |
| orf1a | 883 | 8523 | 2203 | FALSE | 3 |
| orf1a | 884 | 8523 | 2203 | FALSE | 1 |
| orf1a | 885 | 0 | 1984 | FALSE | 4 |
| orf1a | 886 | 0 | 1984 | FALSE | 2 |
| orf1a | 887 | 0 | 1984 | FALSE | 3 |
| orf1a | 888 | 10084 | 4602 | FALSE | 2 |
| orf1a | 889 | 10084 | 4602 | FALSE | 1 |
| orf1a | 890 | 10084 | 4602 | FALSE | 1 |
| orf1a | 891 | 10084 | 5795 | FALSE | 2 |
| orf1a | 892 | 5454 | 4760 | FALSE | 3 |
| orf1a | 893 | 5454 | 4760 | FALSE | 4 |
| orf1a | 894 | 5454 | 10582 | FALSE | 5 |
| orf1a | 895 | 5454 | 10582 | FALSE | 1 |
| orf1a | 896 | 3501 | 10582 | FALSE | 2 |
| orf1a | 897 | 3501 | 12492 | FALSE | 0 |

| | | | | | |
|---|---|---|---|---|---|
| orf1a | 898 | 3153 | 12492 | FALSE | 2 |
| orf1a | 899 | 3153 | 12492 | FALSE | 3 |
| orf1a | 900 | 2927 | 12492 | FALSE | 4 |
| orf1a | 901 | 2927 | 12492 | FALSE | 3 |
| orf1a | 902 | 2927 | 12492 | FALSE | 1 |
| orf1a | 903 | 1905 | 17763 | FALSE | 2 |
| orf1a | 904 | 1905 | 17763 | FALSE | 2 |
| orf1a | 905 | 1905 | 17763 | FALSE | 0 |
| orf1a | 906 | 1905 | 17763 | FALSE | 2 |
| orf1a | 907 | 1411 | 33874 | FALSE | 1 |
| orf1a | 908 | 1447 | 14209 | FALSE | 0 |
| orf1a | 909 | 1447 | 4377 | TRUE | 3 |
| orf1a | 910 | 3265 | 6502 | TRUE | 1 |
| orf1a | 911 | 4001 | 6502 | FALSE | 4 |
| orf1a | 912 | 4001 | 6502 | TRUE | 1 |
| orf1a | 913 | 4490 | 6502 | FALSE | 2 |
| orf1a | 914 | 4490 | 6502 | FALSE | 3 |
| orf1a | 915 | 5119 | 6502 | FALSE | 4 |
| orf1a | 916 | 5119 | 6502 | FALSE | 1 |
| orf1a | 917 | 5119 | 6502 | FALSE | 2 |
| orf1a | 918 | 4837 | 6502 | FALSE | 3 |
| orf1a | 919 | 4837 | 6502 | FALSE | 3 |
| orf1a | 920 | 3861 | 6502 | TRUE | 2 |
| orf1a | 921 | 3861 | 6502 | TRUE | 2 |
| orf1a | 922 | 3702 | 5407 | TRUE | 3 |
| orf1a | 923 | 3890 | 6093 | FALSE | 3 |
| orf1a | 924 | 3559 | 5021 | FALSE | 2 |
| orf1a | 925 | 2864 | 1415 | FALSE | 1 |
| orf1a | 926 | 3165 | 2045 | FALSE | 5 |
| orf1a | 927 | 3165 | 2045 | FALSE | 4 |
| orf1a | 928 | 3507 | 2227 | FALSE | 4 |
| orf1a | 929 | 3507 | 2227 | FALSE | 1 |
| orf1a | 930 | 4084 | 2561 | FALSE | 6 |
| orf1a | 931 | 4084 | 2561 | FALSE | 3 |
| orf1a | 932 | 5013 | 3665 | FALSE | 4 |
| orf1a | 933 | 5049 | 3665 | FALSE | 3 |
| orf1a | 934 | 4768 | 3747 | FALSE | 2 |
| orf1a | 935 | 4954 | 3747 | FALSE | 1 |
| orf1a | 936 | 5319 | 3951 | FALSE | 4 |
| orf1a | 937 | 5450 | 4176 | FALSE | 2 |
| orf1a | 938 | 5879 | 4383 | FALSE | 2 |
| orf1a | 939 | 6285 | 4535 | FALSE | 4 |
| orf1a | 940 | 6227 | 4409 | FALSE | 4 |
| orf1a | 941 | 6619 | 4626 | FALSE | 1 |
| orf1a | 942 | 6456 | 4626 | FALSE | 3 |
| orf1a | 943 | 6964 | 4916 | FALSE | 2 |
| orf1a | 944 | 6964 | 4916 | FALSE | 1 |
| orf1a | 945 | 7057 | 4943 | FALSE | 2 |
| orf1a | 946 | 7057 | 4943 | FALSE | 2 |
| orf1a | 947 | 6272 | 4777 | FALSE | 1 |
| orf1a | 948 | 6253 | 4110 | FALSE | 6 |
| orf1a | 949 | 7297 | 4041 | FALSE | 2 |
| orf1a | 950 | 7013 | 4876 | FALSE | 3 |
| orf1a | 951 | 6457 | 4685 | FALSE | 3 |
| orf1a | 952 | 7217 | 5213 | FALSE | 3 |
| orf1a | 953 | 6393 | 5504 | FALSE | 6 |
| orf1a | 954 | 6393 | 5504 | FALSE | 4 |
| orf1a | 955 | 6711 | 6312 | FALSE | 2 |
| orf1a | 956 | 6711 | 6312 | FALSE | 3 |
| orf1a | 957 | 7845 | 6312 | FALSE | 3 |
| orf1a | 958 | 7845 | 6312 | FALSE | 2 |
| orf1a | 959 | 7845 | 6312 | FALSE | 3 |

| | | | | | |
|---|---|---|---|---|---|
| orf1a | 960 | 7845 | 6312 | FALSE | 4 |
| orf1a | 961 | 7845 | 6312 | FALSE | 3 |
| orf1a | 962 | 7066 | 4614 | FALSE | 1 |
| orf1a | 963 | 7245 | 6198 | FALSE | 5 |
| orf1a | 964 | 6373 | 4747 | FALSE | 2 |
| orf1a | 965 | 6849 | 3217 | FALSE | 4 |
| orf1a | 966 | 6174 | 2927 | FALSE | 3 |
| orf1a | 967 | 3800 | 1519 | FALSE | 3 |
| orf1a | 968 | 3497 | 1503 | FALSE | 3 |
| orf1a | 969 | 3497 | 1503 | FALSE | 5 |
| orf1a | 970 | 4145 | 1609 | FALSE | 1 |
| orf1a | 971 | 4145 | 1609 | FALSE | 3 |
| orf1a | 972 | 4560 | 3170 | FALSE | 5 |
| orf1a | 973 | 4560 | 3170 | FALSE | 5 |
| orf1a | 974 | 4628 | 3630 | FALSE | 5 |
| orf1a | 975 | 4628 | 3630 | FALSE | 4 |
| orf1a | 976 | 4903 | 4065 | FALSE | 5 |
| orf1a | 977 | 4990 | 4381 | FALSE | 3 |
| orf1a | 978 | 4892 | 4381 | FALSE | 4 |
| orf1a | 979 | 4897 | 4698 | FALSE | 5 |
| orf1a | 980 | 4897 | 4698 | FALSE | 4 |
| orf1a | 981 | 5025 | 4913 | FALSE | 4 |
| orf1a | 982 | 4915 | 4913 | FALSE | 3 |
| orf1a | 983 | 5121 | 5444 | FALSE | 3 |
| orf1a | 984 | 5121 | 5444 | FALSE | 4 |
| orf1a | 985 | 5026 | 6521 | FALSE | 5 |
| orf1a | 986 | 5026 | 6521 | FALSE | 2 |
| orf1a | 987 | 4982 | 6153 | FALSE | 1 |
| orf1a | 988 | 4756 | 6153 | FALSE | 0 |
| orf1a | 989 | 4672 | 5804 | FALSE | 4 |
| orf1a | 990 | 4672 | 5804 | FALSE | 4 |
| orf1a | 991 | 1211 | 0 | FALSE | 3 |
| orf1a | 992 | 1413 | 1289 | FALSE | 4 |
| orf1a | 993 | 1469 | 1289 | FALSE | 4 |
| orf1a | 994 | 1528 | 1289 | FALSE | 2 |
| orf1a | 995 | 1528 | 1289 | FALSE | 4 |
| orf1a | 996 | 1528 | 1289 | FALSE | 4 |
| orf1a | 997 | 1572 | 1289 | FALSE | 3 |
| orf1a | 998 | 1572 | 1289 | FALSE | 5 |
| orf1a | 999 | 1572 | 1289 | FALSE | 2 |
| orf1a | 1000 | 1572 | 1289 | FALSE | 3 |
| orf1a | 1001 | 1572 | 1289 | FALSE | 3 |
| orf1a | 1002 | 1563 | 1289 | FALSE | 2 |
| orf1a | 1003 | 1648 | 1289 | FALSE | 3 |
| orf1a | 1004 | 1611 | 1289 | FALSE | 1 |
| orf1a | 1005 | 1611 | 1289 | FALSE | 0 |
| orf1a | 1006 | 1611 | 1289 | FALSE | 3 |
| orf1a | 1007 | 1558 | 0 | FALSE | 4 |
| orf1a | 1008 | 1561 | 0 | FALSE | 4 |
| orf1a | 1009 | 1504 | 0 | FALSE | 1 |
| orf1a | 1010 | 1808 | 0 | FALSE | 3 |
| orf1a | 1011 | 1808 | 0 | FALSE | 3 |
| orf1a | 1012 | 1808 | 1162 | FALSE | 2 |
| orf1a | 1013 | 1808 | 1162 | FALSE | 4 |
| orf1a | 1014 | 1808 | 1162 | FALSE | 3 |
| orf1a | 1015 | 1808 | 1162 | FALSE | 2 |
| orf1a | 1016 | 1701 | 1162 | FALSE | 2 |
| orf1a | 1017 | 1739 | 1162 | FALSE | 2 |
| orf1a | 1018 | 1660 | 1469 | FALSE | 2 |
| orf1a | 1019 | 1700 | 1469 | FALSE | 3 |
| orf1a | 1020 | 1899 | 1469 | FALSE | 2 |
| orf1a | 1021 | 1899 | 1469 | FALSE | 5 |

| | | | | | |
|---|---|---|---|---|---|
| orf1a | 1022 | 1829 | 1469 | FALSE | 3 |
| orf1a | 1023 | 1866 | 1469 | FALSE | 0 |
| orf1a | 1024 | 2111 | 1469 | FALSE | 5 |
| orf1a | 1025 | 2067 | 1469 | FALSE | 4 |
| orf1a | 1026 | 2067 | 1469 | FALSE | 1 |
| orf1a | 1027 | 2067 | 1776 | FALSE | 3 |
| orf1a | 1028 | 2067 | 1776 | FALSE | 1 |
| orf1a | 1029 | 2067 | 1776 | FALSE | 2 |
| orf1a | 1030 | 2067 | 1776 | FALSE | 5 |
| orf1a | 1031 | 2217 | 1776 | FALSE | 2 |
| orf1a | 1032 | 2217 | 1776 | FALSE | 2 |
| orf1a | 1033 | 2593 | 0 | FALSE | 3 |
| orf1a | 1034 | 2504 | 1169 | FALSE | 1 |
| orf1a | 1035 | 2358 | 1169 | FALSE | 4 |
| orf1a | 1036 | 2235 | 1169 | FALSE | 5 |
| orf1a | 1037 | 2457 | 1169 | FALSE | 3 |
| orf1a | 1038 | 2760 | 1169 | FALSE | 2 |
| orf1a | 1039 | 2621 | 1169 | FALSE | 3 |
| orf1a | 1040 | 2621 | 1169 | FALSE | 3 |
| orf1a | 1041 | 2621 | 1169 | FALSE | 4 |
| orf1a | 1042 | 5411 | 1150 | FALSE | 2 |
| orf1a | 1043 | 5411 | 1150 | FALSE | 4 |
| orf1a | 1044 | 5411 | 1150 | FALSE | 3 |
| orf1a | 1045 | 5411 | 1150 | FALSE | 1 |
| orf1a | 1046 | 5411 | 1150 | FALSE | 4 |
| orf1a | 1047 | 5411 | 1150 | FALSE | 4 |
| orf1a | 1048 | 6766 | 3690 | FALSE | 5 |
| orf1a | 1049 | 7920 | 6212 | FALSE | 7 |
| orf1a | 1050 | 7920 | 6212 | FALSE | 2 |
| orf1a | 1051 | 9979 | 6212 | FALSE | 3 |
| orf1a | 1052 | 7750 | 6212 | FALSE | 2 |
| orf1a | 1053 | 9009 | 6212 | FALSE | 0 |
| orf1a | 1054 | 9009 | 6212 | FALSE | 5 |
| orf1a | 1055 | 9009 | 6212 | FALSE | 1 |
| orf1a | 1056 | 9009 | 6212 | FALSE | 4 |
| orf1a | 1057 | 6623 | 11313 | FALSE | 1 |
| orf1a | 1058 | 6623 | 11313 | TRUE | 0 |
| orf1a | 1059 | 6623 | 11313 | TRUE | 1 |
| orf1a | 1060 | 6623 | 11313 | FALSE | 3 |
| orf1a | 1061 | 6623 | 11313 | FALSE | 6 |
| orf1a | 1062 | 6623 | 11313 | FALSE | 2 |
| orf1a | 1063 | 1062 | 0 | FALSE | 1 |
| orf1a | 1064 | 1062 | 0 | FALSE | 2 |
| orf1a | 1065 | 1062 | 0 | TRUE | 7 |
| orf1a | 1066 | 1062 | 1460 | TRUE | 3 |
| orf1a | 1067 | 0 | 1460 | FALSE | 6 |
| orf1a | 1068 | 0 | 1460 | TRUE | 3 |
| orf1a | 1069 | 0 | 1460 | FALSE | 3 |
| orf1a | 1070 | 0 | 2385 | FALSE | 4 |
| orf1a | 1071 | 0 | 2385 | TRUE | 3 |
| orf1a | 1072 | 0 | 2385 | FALSE | 4 |
| orf1a | 1073 | 0 | 2385 | FALSE | 3 |
| orf1a | 1074 | 0 | 2385 | TRUE | 4 |
| orf1a | 1075 | 0 | 2385 | TRUE | 2 |
| orf1a | 1076 | 14683 | 2385 | FALSE | 1 |
| orf1a | 1077 | 14683 | 2385 | FALSE | 5 |
| orf1a | 1078 | 14683 | 2385 | FALSE | 1 |
| orf1a | 1079 | 14683 | 2385 | FALSE | 4 |
| orf1a | 1080 | 16921 | 2385 | FALSE | 5 |
| orf1a | 1081 | 16921 | 3311 | FALSE | 3 |
| orf1a | 1082 | 16921 | 2880 | FALSE | 5 |
| orf1a | 1083 | 16921 | 2880 | FALSE | 2 |

| | | | | | |
|---|---|---|---|---|---|
| orf1a | 1084 | 17060 | 3644 | FALSE | 3 |
| orf1a | 1085 | 17060 | 3728 | FALSE | 1 |
| orf1a | 1086 | 10366 | 5819 | FALSE | 4 |
| orf1a | 1087 | 10366 | 5819 | TRUE | 2 |
| orf1a | 1088 | 10366 | 5819 | FALSE | 3 |
| orf1a | 1089 | 10366 | 5819 | FALSE | 2 |
| orf1a | 1090 | 10366 | 5819 | FALSE | 2 |
| orf1a | 1091 | 9887 | 5819 | FALSE | 0 |
| orf1a | 1092 | 9887 | 5819 | FALSE | 2 |
| orf1a | 1093 | 9887 | 5819 | FALSE | 2 |
| orf1a | 1094 | 9887 | 5819 | FALSE | 3 |
| orf1a | 1095 | 8727 | 5819 | FALSE | 4 |
| orf1a | 1096 | 8727 | 5819 | TRUE | 4 |
| orf1a | 1097 | 8727 | 6380 | TRUE | 2 |
| orf1a | 1098 | 8727 | 5708 | FALSE | 2 |
| orf1a | 1099 | 7497 | 6874 | FALSE | 4 |
| orf1a | 1100 | 6774 | 6874 | FALSE | 4 |
| orf1a | 1101 | 1712 | 1676 | FALSE | 2 |
| orf1a | 1102 | 1640 | 1676 | FALSE | 2 |
| orf1a | 1103 | 1640 | 1676 | FALSE | 2 |
| orf1a | 1104 | 1640 | 1676 | FALSE | 2 |
| orf1a | 1105 | 1640 | 1676 | FALSE | 1 |
| orf1a | 1106 | 1640 | 1676 | FALSE | 4 |
| orf1a | 1107 | 1640 | 1676 | FALSE | 4 |
| orf1a | 1108 | 1640 | 1676 | FALSE | 2 |
| orf1a | 1109 | 1640 | 1676 | FALSE | 1 |
| orf1a | 1110 | 1640 | 1676 | FALSE | 2 |
| orf1a | 1111 | 1640 | 1676 | FALSE | 4 |
| orf1a | 1112 | 1640 | 1676 | FALSE | 0 |
| orf1a | 1113 | 1640 | 0 | FALSE | 2 |
| orf1a | 1114 | 1640 | 0 | FALSE | 3 |
| orf1a | 1115 | 1568 | 0 | FALSE | 2 |
| orf1a | 1116 | 1568 | 0 | FALSE | 3 |
| orf1a | 1124 | 5607 | 3264 | FALSE | 3 |
| orf1a | 1125 | 5607 | 3264 | FALSE | 6 |
| orf1a | 1126 | 9339 | 4084 | FALSE | 2 |
| orf1a | 1127 | 9339 | 4084 | FALSE | 5 |
| orf1a | 1128 | 8188 | 4084 | FALSE | 2 |
| orf1a | 1129 | 8188 | 4084 | FALSE | 4 |
| orf1a | 1130 | 8188 | 4084 | FALSE | 2 |
| orf1a | 1131 | 8188 | 4084 | FALSE | 3 |
| orf1a | 1132 | 8188 | 4084 | FALSE | 3 |
| orf1a | 1133 | 8188 | 4084 | FALSE | 4 |
| orf1a | 1134 | 8188 | 4084 | FALSE | 4 |
| orf1a | 1135 | 8188 | 4084 | FALSE | 3 |
| orf1a | 1136 | 9480 | 4305 | FALSE | 4 |
| orf1a | 1137 | 9480 | 4305 | FALSE | 4 |
| orf1a | 1138 | 9480 | 4305 | FALSE | 3 |
| orf1a | 1139 | 11030 | 4652 | FALSE | 2 |
| orf1a | 1140 | 9422 | 4652 | FALSE | 3 |
| orf1a | 1141 | 7016 | 4967 | FALSE | 2 |
| orf1a | 1142 | 6881 | 2671 | FALSE | 3 |
| orf1a | 1143 | 8366 | 2671 | FALSE | 3 |
| orf1a | 1144 | 8366 | 2671 | FALSE | 3 |
| orf1a | 1145 | 8366 | 2671 | FALSE | 1 |
| orf1a | 1146 | 8366 | 2671 | FALSE | 2 |
| orf1a | 1147 | 8366 | 2671 | FALSE | 2 |
| orf1a | 1148 | 8366 | 2671 | FALSE | 4 |
| orf1a | 1149 | 8366 | 2671 | FALSE | 1 |
| orf1a | 1150 | 8366 | 2671 | FALSE | 3 |
| orf1a | 1151 | 3930 | 1523 | FALSE | 4 |
| orf1a | 1152 | 3930 | 1523 | FALSE | 5 |

| | | | | | |
|---|---|---|---|---|---|
| orf1a | 1153 | 3930 | 1523 | FALSE | 4 |
| orf1a | 1154 | 3930 | 1523 | FALSE | 2 |
| orf1a | 1155 | 6479 | 1523 | FALSE | 4 |
| orf1a | 1156 | 5836 | 1523 | FALSE | 4 |
| orf1a | 1157 | 5194 | 0 | FALSE | 1 |
| orf1a | 1158 | 3422 | 1394 | FALSE | 4 |
| orf1a | 1159 | 3422 | 1394 | FALSE | 3 |
| orf1a | 1160 | 3422 | 1394 | FALSE | 3 |
| orf1a | 1161 | 3422 | 1394 | FALSE | 1 |
| orf1a | 1162 | 3422 | 1394 | FALSE | 1 |
| orf1a | 1163 | 3422 | 1394 | FALSE | 2 |
| orf1a | 1164 | 3422 | 1394 | FALSE | 1 |
| orf1a | 1165 | 3422 | 1394 | FALSE | 2 |
| orf1a | 1166 | 3422 | 1394 | FALSE | 3 |
| orf1a | 1167 | 3422 | 1394 | FALSE | 6 |
| orf1a | 1168 | 3422 | 1394 | FALSE | 1 |
| orf1a | 1169 | 3422 | 1394 | FALSE | 2 |
| orf1a | 1170 | 3422 | 1394 | FALSE | 4 |
| orf1a | 1171 | 2536 | 1394 | FALSE | 3 |
| orf1a | 1172 | 2536 | 1394 | FALSE | 0 |
| orf1a | 1192 | 7346 | 10336 | FALSE | 3 |
| orf1a | 1193 | 7346 | 10336 | FALSE | 3 |
| orf1a | 1194 | 8728 | 10336 | FALSE | 2 |
| orf1a | 1195 | 8728 | 10336 | FALSE | 0 |
| orf1a | 1196 | 9434 | 5598 | FALSE | 3 |
| orf1a | 1197 | 9434 | 5598 | FALSE | 3 |
| orf1a | 1198 | 9434 | 5598 | FALSE | 1 |
| orf1a | 1199 | 9434 | 5598 | FALSE | 3 |
| orf1a | 1200 | 9911 | 4946 | FALSE | 4 |
| orf1a | 1201 | 9911 | 4946 | FALSE | 4 |
| orf1a | 1202 | 8646 | 4946 | FALSE | 4 |
| orf1a | 1203 | 8646 | 4946 | FALSE | 4 |
| orf1a | 1204 | 7730 | 4946 | FALSE | 5 |
| orf1a | 1205 | 7730 | 4946 | FALSE | 4 |
| orf1a | 1206 | 6768 | 4224 | FALSE | 3 |
| orf1a | 1207 | 6704 | 2696 | FALSE | 3 |
| orf1a | 1208 | 7903 | 2494 | FALSE | 1 |
| orf1a | 1209 | 7566 | 2494 | FALSE | 8 |
| orf1a | 1210 | 7566 | 2281 | FALSE | 4 |
| orf1a | 1211 | 6730 | 1808 | FALSE | 2 |
| orf1a | 1212 | 6730 | 1808 | FALSE | 3 |
| orf1a | 1213 | 6730 | 1808 | FALSE | 3 |
| orf1a | 1214 | 6730 | 1808 | FALSE | 0 |
| orf1a | 1215 | 5802 | 1413 | FALSE | 2 |
| orf1a | 1216 | 5802 | 1413 | FALSE | 3 |
| orf1a | 1217 | 6751 | 1413 | FALSE | 1 |
| orf1a | 1218 | 6751 | 1315 | FALSE | 4 |
| orf1a | 1219 | 8110 | 1315 | FALSE | 4 |
| orf1a | 1220 | 8110 | 1315 | FALSE | 2 |
| orf1a | 1221 | 13298 | 1308 | FALSE | 2 |
| orf1a | 1222 | 13298 | 1308 | FALSE | 2 |
| orf1a | 1223 | 0 | 1121 | FALSE | 1 |
| orf1a | 1224 | 0 | 1121 | FALSE | 4 |
| orf1a | 1225 | 0 | 1021 | FALSE | 6 |
| orf1a | 1226 | 0 | 1021 | FALSE | 2 |
| orf1a | 1227 | 0 | 1021 | FALSE | 3 |
| orf1a | 1228 | 0 | 1021 | FALSE | 2 |
| orf1a | 1229 | 0 | 1021 | FALSE | 4 |
| orf1a | 1230 | 0 | 1021 | FALSE | 5 |
| orf1a | 1231 | 0 | 1021 | FALSE | 3 |
| orf1a | 1232 | 0 | 1021 | FALSE | 6 |
| orf1a | 1234 | 1553 | 0 | FALSE | 4 |

| | | | | | |
|---|---|---|---|---|---|
| orf1a | 1235 | 1553 | 0 | FALSE | 2 |
| orf1a | 1236 | 12163 | 1852 | FALSE | 5 |
| orf1a | 1237 | 12163 | 1852 | FALSE | 3 |
| orf1a | 1238 | 10414 | 3197 | FALSE | 1 |
| orf1a | 1239 | 10414 | 3197 | FALSE | 4 |
| orf1a | 1240 | 10414 | 3197 | FALSE | 1 |
| orf1a | 1241 | 10414 | 3197 | FALSE | 1 |
| orf1a | 1242 | 8108 | 3197 | FALSE | 3 |
| orf1a | 1243 | 8108 | 3197 | FALSE | 4 |
| orf1a | 1244 | 11228 | 4426 | FALSE | 5 |
| orf1a | 1245 | 11228 | 4426 | FALSE | 1 |
| orf1a | 1246 | 11228 | 4426 | FALSE | 2 |
| orf1a | 1247 | 11228 | 4426 | FALSE | 7 |
| orf1a | 1248 | 11228 | 4426 | FALSE | 2 |
| orf1a | 1249 | 12107 | 4426 | FALSE | 5 |
| orf1a | 1250 | 10424 | 4426 | FALSE | 4 |
| orf1a | 1251 | 9394 | 5161 | FALSE | 3 |
| orf1a | 1252 | 9483 | 5161 | FALSE | 2 |
| orf1a | 1253 | 9553 | 5962 | FALSE | 0 |
| orf1a | 1254 | 9553 | 5962 | FALSE | 3 |
| orf1a | 1255 | 9553 | 5962 | FALSE | 2 |
| orf1a | 1256 | 9553 | 5962 | FALSE | 0 |
| orf1a | 1257 | 11820 | 5962 | FALSE | 1 |
| orf1a | 1258 | 11820 | 5962 | FALSE | 2 |
| orf1a | 1259 | 4290 | 0 | FALSE | 3 |
| orf1a | 1260 | 5893 | 0 | FALSE | 2 |
| orf1a | 1261 | 5893 | 0 | FALSE | 4 |
| orf1a | 1262 | 7591 | 0 | FALSE | 2 |
| orf1a | 1263 | 7591 | 0 | FALSE | 3 |
| orf1a | 1264 | 7591 | 0 | FALSE | 1 |
| orf1a | 1265 | 10804 | 0 | FALSE | 2 |
| orf1a | 1266 | 10804 | 0 | FALSE | 1 |
| orf1a | 1267 | 10892 | 0 | FALSE | 3 |
| orf1a | 1268 | 10892 | 0 | FALSE | 2 |
| orf1a | 1269 | 10892 | 0 | FALSE | 2 |
| orf1a | 1270 | 10892 | 0 | FALSE | 2 |
| orf1a | 1271 | 10892 | 0 | FALSE | 2 |
| orf1a | 1272 | 10755 | 0 | FALSE | 5 |
| orf1a | 1273 | 10755 | 0 | FALSE | 4 |
| orf1a | 1274 | 10755 | 0 | FALSE | 4 |
| orf1a | 1275 | 10772 | 0 | FALSE | 5 |
| orf1a | 1276 | 10772 | 0 | FALSE | 3 |
| orf1a | 1277 | 10345 | 0 | FALSE | 2 |
| orf1a | 1278 | 10345 | 0 | FALSE | 1 |
| orf1a | 1279 | 10345 | 0 | FALSE | 4 |
| orf1a | 1280 | 10345 | 0 | FALSE | 2 |
| orf1a | 1281 | 10345 | 0 | FALSE | 1 |
| orf1a | 1282 | 10345 | 0 | FALSE | 4 |
| orf1a | 1283 | 10345 | 0 | FALSE | 4 |
| orf1a | 1284 | 10345 | 0 | FALSE | 3 |
| orf1a | 1285 | 10345 | 0 | FALSE | 5 |
| orf1a | 1286 | 5818 | 0 | FALSE | 0 |
| orf1a | 1287 | 1291 | 0 | FALSE | 4 |
| orf1a | 1288 | 1291 | 2058 | FALSE | 2 |
| orf1a | 1289 | 1291 | 2058 | FALSE | 2 |
| orf1a | 1290 | 1291 | 2058 | FALSE | 2 |
| orf1a | 1291 | 1291 | 2058 | FALSE | 2 |
| orf1a | 1292 | 1291 | 2058 | FALSE | 3 |
| orf1a | 1293 | 1291 | 2058 | FALSE | 5 |
| orf1a | 1294 | 1291 | 2266 | FALSE | 6 |
| orf1a | 1295 | 1291 | 2266 | FALSE | 2 |
| orf1a | 1296 | 1291 | 2266 | FALSE | 2 |

| | | | | | |
|---|---|---|---|---|---|
| orf1a | 1297 | 1291 | 2266 | FALSE | 3 |
| orf1a | 1298 | 1291 | 2266 | FALSE | 3 |
| orf1a | 1299 | 1291 | 2266 | FALSE | 2 |
| orf1a | 1300 | 1291 | 2266 | FALSE | 2 |
| orf1a | 1301 | 0 | 2266 | FALSE | 0 |
| orf1a | 1302 | 0 | 2266 | FALSE | 4 |
| orf1a | 1303 | 0 | 2475 | FALSE | 2 |
| orf1a | 1304 | 0 | 2475 | FALSE | 5 |
| orf1a | 1305 | 0 | 2475 | FALSE | 4 |
| orf1a | 1306 | 0 | 2475 | FALSE | 4 |
| orf1a | 1307 | 0 | 2475 | FALSE | 3 |
| orf1a | 1308 | 0 | 3794 | FALSE | 2 |
| orf1a | 1309 | 0 | 5114 | FALSE | 2 |
| orf1a | 1310 | 2244 | 3063 | FALSE | 3 |
| orf1a | 1311 | 2244 | 3063 | FALSE | 3 |
| orf1a | 1312 | 2244 | 3063 | FALSE | 4 |
| orf1a | 1313 | 2244 | 3063 | FALSE | 3 |
| orf1a | 1314 | 2244 | 3063 | FALSE | 5 |
| orf1a | 1315 | 2244 | 3063 | FALSE | 2 |
| orf1a | 1316 | 2244 | 3063 | FALSE | 6 |
| orf1a | 1317 | 2244 | 3063 | FALSE | 4 |
| orf1a | 1318 | 2244 | 3063 | FALSE | 3 |
| orf1a | 1319 | 2244 | 3063 | FALSE | 4 |
| orf1a | 1320 | 2244 | 3063 | FALSE | 5 |
| orf1a | 1321 | 2244 | 3063 | FALSE | 3 |
| orf1a | 1322 | 3809 | 3063 | FALSE | 3 |
| orf1a | 1323 | 3809 | 1011 | FALSE | 6 |
| orf1a | 1324 | 3809 | 1011 | FALSE | 1 |
| orf1a | 1325 | 10070 | 0 | FALSE | 1 |
| orf1a | 1326 | 10070 | 0 | FALSE | 0 |
| orf1a | 1327 | 10070 | 0 | FALSE | 2 |
| orf1a | 1328 | 10070 | 0 | FALSE | 3 |
| orf1a | 1329 | 10070 | 0 | FALSE | 4 |
| orf1a | 1330 | 6081 | 0 | FALSE | 4 |
| orf1a | 1331 | 6081 | 0 | FALSE | 4 |
| orf1a | 1332 | 6081 | 0 | FALSE | 5 |
| orf1a | 1333 | 6081 | 0 | FALSE | 4 |
| orf1a | 1334 | 6081 | 0 | FALSE | 2 |
| orf1a | 1335 | 6081 | 0 | FALSE | 1 |
| orf1a | 1336 | 6081 | 3025 | FALSE | 3 |
| orf1a | 1337 | 2092 | 3025 | FALSE | 3 |
| orf1a | 1338 | 10771 | 2254 | FALSE | 2 |
| orf1a | 1339 | 10771 | 2254 | FALSE | 3 |
| orf1a | 1340 | 13291 | 4535 | FALSE | 4 |
| orf1a | 1341 | 13291 | 4535 | FALSE | 6 |
| orf1a | 1342 | 13291 | 4535 | FALSE | 4 |
| orf1a | 1343 | 13291 | 4535 | FALSE | 1 |
| orf1a | 1344 | 13291 | 4535 | FALSE | 3 |
| orf1a | 1345 | 15158 | 4535 | FALSE | 5 |
| orf1a | 1346 | 15158 | 4535 | FALSE | 5 |
| orf1a | 1347 | 15158 | 4535 | FALSE | 1 |
| orf1a | 1348 | 15158 | 4535 | FALSE | 5 |
| orf1a | 1349 | 15158 | 4535 | FALSE | 2 |
| orf1a | 1350 | 15158 | 4535 | FALSE | 0 |
| orf1a | 1351 | 15158 | 4837 | FALSE | 3 |
| orf1a | 1352 | 15158 | 4227 | FALSE | 6 |
| orf1a | 1353 | 15181 | 4776 | FALSE | 1 |
| orf1a | 1354 | 15181 | 4776 | FALSE | 4 |
| orf1a | 1355 | 0 | 1181 | FALSE | 3 |
| orf1a | 1356 | 0 | 1181 | FALSE | 3 |
| orf1a | 1357 | 0 | 1181 | FALSE | 7 |
| orf1a | 1358 | 0 | 1181 | FALSE | 4 |

| | | | | | |
|---|---|---|---|---|---|
| orf1a | 1359 | 0 | 1181 | FALSE | 3 |
| orf1a | 1360 | 0 | 1181 | FALSE | 4 |
| orf1a | 1361 | 0 | 1181 | FALSE | 4 |
| orf1a | 1362 | 0 | 1181 | FALSE | 2 |
| orf1a | 1363 | 0 | 1181 | FALSE | 5 |
| orf1a | 1364 | 4224 | 1181 | FALSE | 2 |
| orf1a | 1365 | 4224 | 1181 | FALSE | 1 |
| orf1a | 1366 | 7709 | 2822 | FALSE | 3 |
| orf1a | 1367 | 7709 | 3643 | FALSE | 1 |
| orf1a | 1368 | 7709 | 3643 | FALSE | 2 |
| orf1a | 1369 | 7709 | 3643 | FALSE | 3 |
| orf1a | 1370 | 5975 | 3643 | FALSE | 2 |
| orf1a | 1371 | 5975 | 3643 | FALSE | 2 |
| orf1a | 1372 | 5975 | 3643 | FALSE | 2 |
| orf1a | 1373 | 5975 | 3643 | FALSE | 3 |
| orf1a | 1374 | 6696 | 3643 | FALSE | 1 |
| orf1a | 1375 | 6696 | 3643 | FALSE | 2 |
| orf1a | 1376 | 5878 | 3643 | FALSE | 4 |
| orf1a | 1377 | 5878 | 3643 | FALSE | 6 |
| orf1a | 1378 | 5878 | 3643 | FALSE | 3 |
| orf1a | 1379 | 6209 | 3643 | FALSE | 4 |
| orf1a | 1380 | 5343 | 3643 | FALSE | 1 |
| orf1a | 1381 | 4173 | 0 | FALSE | 3 |
| orf1a | 1382 | 4173 | 0 | FALSE | 6 |
| orf1a | 1383 | 4173 | 0 | FALSE | 4 |
| orf1a | 1384 | 5104 | 2244 | FALSE | 1 |
| orf1a | 1385 | 5624 | 2244 | FALSE | 2 |
| orf1a | 1386 | 5624 | 2244 | FALSE | 5 |
| orf1a | 1387 | 5624 | 2244 | FALSE | 2 |
| orf1a | 1388 | 5624 | 2244 | FALSE | 1 |
| orf1a | 1389 | 4188 | 2244 | FALSE | 0 |
| orf1a | 1390 | 3374 | 2495 | FALSE | 3 |
| orf1a | 1391 | 3638 | 2495 | FALSE | 2 |
| orf1a | 1392 | 3638 | 2495 | FALSE | 2 |
| orf1a | 1393 | 3638 | 2495 | FALSE | 6 |
| orf1a | 1394 | 3360 | 2495 | FALSE | 4 |
| orf1a | 1395 | 3751 | 2495 | FALSE | 4 |
| orf1a | 1396 | 3423 | 2495 | FALSE | 3 |
| orf1a | 1397 | 3423 | 2495 | FALSE | 5 |
| orf1a | 1398 | 3423 | 2495 | FALSE | 0 |
| orf1a | 1399 | 2366 | 2621 | FALSE | 2 |
| orf1a | 1400 | 2366 | 2621 | FALSE | 1 |
| orf1a | 1401 | 2366 | 2621 | FALSE | 2 |
| orf1a | 1402 | 2366 | 2193 | FALSE | 1 |
| orf1a | 1403 | 2366 | 2193 | FALSE | 2 |
| orf1a | 1404 | 2366 | 2774 | FALSE | 4 |
| orf1a | 1405 | 1572 | 2927 | FALSE | 1 |
| orf1a | 1406 | 1572 | 2387 | FALSE | 2 |
| orf1a | 1407 | 1572 | 2387 | FALSE | 4 |
| orf1a | 1408 | 1572 | 2920 | FALSE | 3 |
| orf1a | 1409 | 1456 | 2920 | FALSE | 1 |
| orf1a | 1410 | 1456 | 2920 | FALSE | 1 |
| orf1a | 1411 | 0 | 2920 | FALSE | 2 |
| orf1a | 1412 | 0 | 2920 | FALSE | 4 |
| orf1a | 1413 | 0 | 2920 | FALSE | 4 |
| orf1a | 1414 | 0 | 5056 | FALSE | 3 |
| orf1a | 1415 | 0 | 5056 | FALSE | 3 |
| orf1a | 1416 | 0 | 5056 | FALSE | 2 |
| orf1a | 1417 | 0 | 5986 | FALSE | 1 |
| orf1a | 1418 | 0 | 5986 | FALSE | 3 |
| orf1a | 1419 | 0 | 6476 | FALSE | 2 |
| orf1a | 1420 | 0 | 6476 | FALSE | 1 |

| | | | | | |
|---|---|---|---|---|---|
| orf1a | 1421 | 0 | 9061 | FALSE | 4 |
| orf1a | 1422 | 0 | 9061 | FALSE | 2 |
| orf1a | 1423 | 0 | 13603 | FALSE | 1 |
| orf1a | 1424 | 0 | 13603 | FALSE | 2 |
| orf1a | 1425 | 0 | 13603 | FALSE | 1 |
| orf1a | 1426 | 0 | 13603 | FALSE | 5 |
| orf1a | 1427 | 0 | 13603 | FALSE | 2 |
| orf1a | 1428 | 0 | 13603 | FALSE | 3 |
| orf1a | 1430 | 0 | 3867 | FALSE | 3 |
| orf1a | 1431 | 0 | 3867 | FALSE | 0 |
| orf1a | 1432 | 0 | 3867 | FALSE | 3 |
| orf1a | 1433 | 0 | 3867 | FALSE | 1 |
| orf1a | 1434 | 0 | 3867 | FALSE | 2 |
| orf1a | 1435 | 0 | 3867 | FALSE | 2 |
| orf1a | 1436 | 1478 | 3893 | FALSE | 3 |
| orf1a | 1437 | 1478 | 3893 | FALSE | 2 |
| orf1a | 1438 | 1478 | 3893 | FALSE | 2 |
| orf1a | 1439 | 1478 | 3893 | FALSE | 3 |
| orf1a | 1440 | 1478 | 3893 | FALSE | 1 |
| orf1a | 1441 | 1478 | 3893 | FALSE | 3 |
| orf1a | 1442 | 2084 | 3893 | FALSE | 3 |
| orf1a | 1443 | 2084 | 3893 | FALSE | 1 |
| orf1a | 1444 | 1727 | 3893 | FALSE | 2 |
| orf1a | 1445 | 1727 | 3918 | FALSE | 1 |
| orf1a | 1446 | 1617 | 3918 | FALSE | 3 |
| orf1a | 1447 | 1617 | 3918 | FALSE | 3 |
| orf1a | 1448 | 1617 | 3918 | FALSE | 2 |
| orf1a | 1449 | 1617 | 3918 | FALSE | 4 |
| orf1a | 1450 | 1546 | 3918 | FALSE | 4 |
| orf1a | 1451 | 1563 | 0 | FALSE | 5 |
| orf1a | 1452 | 1563 | 0 | FALSE | 3 |
| orf1a | 1453 | 1563 | 0 | FALSE | 1 |
| orf1a | 1454 | 1563 | 2950 | FALSE | 3 |
| orf1a | 1455 | 1563 | 2950 | FALSE | 2 |
| orf1a | 1456 | 1563 | 6263 | FALSE | 2 |
| orf1a | 1457 | 1187 | 6263 | FALSE | 1 |
| orf1a | 1458 | 2838 | 6263 | FALSE | 4 |
| orf1a | 1459 | 3447 | 6263 | FALSE | 0 |
| orf1a | 1460 | 8491 | 5028 | FALSE | 3 |
| orf1a | 1461 | 10292 | 5028 | FALSE | 0 |
| orf1a | 1462 | 10292 | 5028 | FALSE | 2 |
| orf1a | 1463 | 10292 | 5028 | FALSE | 3 |
| orf1a | 1464 | 10292 | 5028 | FALSE | 7 |
| orf1a | 1465 | 13302 | 5028 | FALSE | 2 |
| orf1a | 1466 | 13302 | 5028 | FALSE | 4 |
| orf1a | 1467 | 13302 | 5028 | FALSE | 4 |
| orf1a | 1468 | 13302 | 5028 | FALSE | 2 |
| orf1a | 1469 | 13302 | 5443 | FALSE | 3 |
| orf1a | 1470 | 13302 | 5443 | FALSE | 5 |
| orf1a | 1471 | 13302 | 4410 | FALSE | 5 |
| orf1a | 1472 | 13302 | 4410 | FALSE | 3 |
| orf1a | 1473 | 16057 | 4410 | FALSE | 3 |
| orf1a | 1474 | 16057 | 4410 | FALSE | 1 |
| orf1a | 1490 | 4190 | 0 | FALSE | 1 |
| orf1a | 1491 | 4190 | 0 | FALSE | 3 |
| orf1a | 1492 | 2612 | 0 | FALSE | 4 |
| orf1a | 1493 | 2612 | 0 | FALSE | 3 |
| orf1a | 1494 | 2612 | 1369 | FALSE | 2 |
| orf1a | 1495 | 2612 | 1369 | FALSE | 1 |
| orf1a | 1496 | 4401 | 1304 | FALSE | 1 |
| orf1a | 1497 | 4401 | 1304 | FALSE | 3 |
| orf1a | 1498 | 7824 | 5655 | FALSE | 1 |

| | | | | | |
|---|---|---|---|---|---|
| orf1a | 1499 | 7824 | 5655 | FALSE | 1 |
| orf1a | 1500 | 7824 | 5655 | FALSE | 2 |
| orf1a | 1501 | 7824 | 5655 | FALSE | 3 |
| orf1a | 1502 | 7607 | 5655 | FALSE | 2 |
| orf1a | 1503 | 7607 | 5655 | FALSE | 1 |
| orf1a | 1504 | 7607 | 5655 | FALSE | 4 |
| orf1a | 1505 | 7851 | 5655 | FALSE | 3 |
| orf1a | 1506 | 7851 | 5655 | FALSE | 3 |
| orf1a | 1507 | 8376 | 5655 | FALSE | 3 |
| orf1a | 1508 | 8376 | 5655 | FALSE | 4 |
| orf1a | 1509 | 8376 | 6512 | FALSE | 1 |
| orf1a | 1510 | 8307 | 6512 | FALSE | 2 |
| orf1a | 1511 | 10080 | 7830 | FALSE | 2 |
| orf1a | 1512 | 10080 | 6642 | FALSE | 3 |
| orf1a | 1513 | 5993 | 1888 | FALSE | 2 |
| orf1a | 1514 | 5993 | 1888 | FALSE | 4 |
| orf1a | 1515 | 5993 | 1888 | FALSE | 3 |
| orf1a | 1516 | 5993 | 1888 | FALSE | 2 |
| orf1a | 1517 | 7422 | 1888 | FALSE | 3 |
| orf1a | 1518 | 7422 | 1888 | FALSE | 3 |
| orf1a | 1519 | 7422 | 1888 | FALSE | 2 |
| orf1a | 1520 | 7422 | 1888 | FALSE | 3 |
| orf1a | 1521 | 7422 | 1888 | FALSE | 4 |
| orf1a | 1522 | 7422 | 1889 | FALSE | 4 |
| orf1a | 1523 | 7422 | 1889 | FALSE | 4 |
| orf1a | 1524 | 4267 | 4348 | FALSE | 6 |
| orf1a | 1525 | 1112 | 4348 | FALSE | 2 |
| orf1a | 1526 | 3000 | 4009 | FALSE | 5 |
| orf1a | 1527 | 3000 | 4716 | FALSE | 2 |
| orf1a | 1528 | 3000 | 4716 | FALSE | 2 |
| orf1a | 1529 | 3000 | 4716 | FALSE | 4 |
| orf1a | 1530 | 3000 | 4716 | FALSE | 2 |
| orf1a | 1531 | 3000 | 4716 | FALSE | 4 |
| orf1a | 1532 | 3000 | 4716 | FALSE | 2 |
| orf1a | 1533 | 3000 | 4716 | FALSE | 4 |
| orf1a | 1534 | 3000 | 4716 | FALSE | 5 |
| orf1a | 1535 | 3000 | 4716 | FALSE | 4 |
| orf1a | 1536 | 3000 | 4716 | FALSE | 5 |
| orf1a | 1537 | 3000 | 6129 | FALSE | 2 |
| orf1a | 1538 | 3000 | 6129 | FALSE | 1 |
| orf1a | 1539 | 4889 | 2992 | FALSE | 3 |
| orf1a | 1540 | 3037 | 2992 | FALSE | 3 |
| orf1a | 1541 | 1185 | 0 | FALSE | 5 |
| orf1a | 1542 | 1185 | 0 | FALSE | 2 |
| orf1a | 1543 | 1185 | 0 | FALSE | 2 |
| orf1a | 1544 | 1185 | 1362 | FALSE | 1 |
| orf1a | 1545 | 1185 | 1362 | FALSE | 3 |
| orf1a | 1546 | 1185 | 1362 | FALSE | 1 |
| orf1a | 1547 | 1185 | 1362 | FALSE | 2 |
| orf1a | 1548 | 1185 | 1812 | FALSE | 3 |
| orf1a | 1549 | 1185 | 1812 | FALSE | 1 |
| orf1a | 1550 | 1185 | 1812 | FALSE | 3 |
| orf1a | 1551 | 1185 | 1812 | FALSE | 3 |
| orf1a | 1552 | 1185 | 1655 | FALSE | 2 |
| orf1a | 1553 | 1185 | 1655 | FALSE | 2 |
| orf1a | 1554 | 1185 | 1655 | FALSE | 5 |
| orf1a | 1555 | 0 | 1655 | FALSE | 2 |
| orf1a | 1556 | 0 | 1655 | FALSE | 2 |
| orf1a | 1557 | 0 | 1655 | FALSE | 2 |
| orf1a | 1558 | 0 | 1655 | FALSE | 1 |
| orf1a | 1559 | 0 | 1802 | FALSE | 2 |
| orf1a | 1560 | 0 | 1802 | FALSE | 1 |

| | | | | | |
|---|---|---|---|---|---|
| orf1a | 1561 | 0 | 1802 | FALSE | 3 |
| orf1a | 1562 | 0 | 1802 | FALSE | 5 |
| orf1a | 1563 | 0 | 1342 | FALSE | 2 |
| orf1a | 1564 | 18082 | 3350 | FALSE | 1 |
| orf1a | 1565 | 18082 | 3350 | FALSE | 2 |
| orf1a | 1566 | 18082 | 3350 | FALSE | 5 |
| orf1a | 1567 | 18082 | 5359 | FALSE | 5 |
| orf1a | 1568 | 18082 | 5359 | FALSE | 1 |
| orf1a | 1569 | 18082 | 5359 | FALSE | 4 |
| orf1a | 1570 | 18082 | 5359 | FALSE | 4 |
| orf1a | 1571 | 18082 | 5359 | FALSE | 1 |
| orf1a | 1572 | 18082 | 5359 | FALSE | 3 |
| orf1a | 1573 | 18082 | 5359 | FALSE | 4 |
| orf1a | 1574 | 18082 | 5359 | FALSE | 4 |
| orf1a | 1575 | 18082 | 5359 | FALSE | 3 |
| orf1a | 1576 | 10996 | 5359 | FALSE | 4 |
| orf1a | 1577 | 10996 | 5359 | FALSE | 4 |
| orf1a | 1578 | 9936 | 5359 | FALSE | 2 |
| orf1a | 1579 | 5864 | 0 | FALSE | 5 |
| orf1a | 1580 | 5864 | 0 | FALSE | 2 |
| orf1a | 1581 | 5864 | 0 | FALSE | 5 |
| orf1a | 1582 | 5864 | 0 | FALSE | 5 |
| orf1a | 1583 | 5864 | 0 | FALSE | 4 |
| orf1a | 1584 | 5864 | 0 | FALSE | 4 |
| orf1a | 1585 | 5864 | 0 | FALSE | 4 |
| orf1a | 1586 | 5864 | 0 | FALSE | 3 |
| orf1a | 1587 | 5864 | 0 | FALSE | 3 |
| orf1a | 1588 | 5864 | 0 | FALSE | 3 |
| orf1a | 1589 | 5864 | 0 | FALSE | 3 |
| orf1a | 1590 | 5864 | 0 | FALSE | 2 |
| orf1a | 1591 | 7818 | 0 | FALSE | 3 |
| orf1a | 1592 | 7818 | 0 | FALSE | 2 |
| orf1a | 1596 | 4131 | 6897 | FALSE | 2 |
| orf1a | 1597 | 4131 | 6897 | FALSE | 2 |
| orf1a | 1598 | 4131 | 6897 | FALSE | 3 |
| orf1a | 1599 | 4131 | 6897 | FALSE | 2 |
| orf1a | 1600 | 3311 | 4425 | FALSE | 1 |
| orf1a | 1601 | 3311 | 4425 | FALSE | 1 |
| orf1a | 1602 | 3311 | 4425 | FALSE | 1 |
| orf1a | 1603 | 3311 | 4425 | FALSE | 1 |
| orf1a | 1604 | 3388 | 4425 | FALSE | 1 |
| orf1a | 1605 | 3388 | 4425 | FALSE | 4 |
| orf1a | 1606 | 2915 | 4425 | FALSE | 1 |
| orf1a | 1607 | 2915 | 4425 | FALSE | 2 |
| orf1a | 1608 | 3029 | 4425 | FALSE | 3 |
| orf1a | 1609 | 3029 | 4425 | FALSE | 3 |
| orf1a | 1610 | 2862 | 4425 | FALSE | 2 |
| orf1a | 1611 | 2681 | 1953 | FALSE | 3 |
| orf1a | 1612 | 3330 | 1502 | FALSE | 1 |
| orf1a | 1613 | 3330 | 1502 | FALSE | 2 |
| orf1a | 1614 | 4177 | 5409 | FALSE | 4 |
| orf1a | 1615 | 4460 | 6273 | FALSE | 2 |
| orf1a | 1616 | 4460 | 6273 | FALSE | 1 |
| orf1a | 1617 | 4460 | 6273 | FALSE | 1 |
| orf1a | 1618 | 4460 | 6273 | FALSE | 2 |
| orf1a | 1619 | 4565 | 6273 | FALSE | 2 |
| orf1a | 1620 | 4333 | 6273 | FALSE | 1 |
| orf1a | 1621 | 4924 | 6273 | FALSE | 3 |
| orf1a | 1622 | 4924 | 6273 | FALSE | 2 |
| orf1a | 1623 | 5097 | 6273 | FALSE | 1 |
| orf1a | 1624 | 5097 | 5278 | FALSE | 1 |
| orf1a | 1625 | 5664 | 5278 | FALSE | 2 |

| | | | | | |
|---|---|---|---|---|---|
| orf1a | 1626 | 5261 | 5278 | FALSE | 5 |
| orf1a | 1627 | 5125 | 6334 | FALSE | 3 |
| orf1a | 1628 | 5125 | 5693 | FALSE | 6 |
| orf1a | 1629 | 3217 | 3372 | FALSE | 4 |
| orf1a | 1630 | 6890 | 6730 | FALSE | 4 |
| orf1a | 1631 | 6890 | 6730 | FALSE | 4 |
| orf1a | 1632 | 6890 | 6730 | FALSE | 0 |
| orf1a | 1633 | 6890 | 6730 | FALSE | 4 |
| orf1a | 1634 | 6890 | 6730 | FALSE | 2 |
| orf1a | 1635 | 8634 | 6730 | FALSE | 2 |
| orf1a | 1636 | 8634 | 6730 | FALSE | 2 |
| orf1a | 1637 | 8634 | 6730 | FALSE | 1 |
| orf1a | 1638 | 8560 | 6730 | FALSE | 2 |
| orf1a | 1639 | 8560 | 8089 | FALSE | 3 |
| orf1a | 1640 | 8560 | 8089 | FALSE | 3 |
| orf1a | 1641 | 9989 | 8089 | FALSE | 1 |
| orf1a | 1642 | 9989 | 8089 | FALSE | 4 |
| orf1a | 1643 | 9989 | 11769 | FALSE | 3 |
| orf1a | 1644 | 9989 | 11769 | FALSE | 2 |
| orf1a | 1645 | 8265 | 0 | FALSE | 5 |
| orf1a | 1646 | 8265 | 0 | FALSE | 3 |
| orf1a | 1647 | 8265 | 0 | FALSE | 5 |
| orf1a | 1648 | 8265 | 0 | FALSE | 2 |
| orf1a | 1649 | 8265 | 0 | FALSE | 5 |
| orf1a | 1650 | 8265 | 0 | FALSE | 0 |
| orf1a | 1651 | 8265 | 0 | FALSE | 2 |
| orf1a | 1652 | 8265 | 0 | FALSE | 2 |
| orf1a | 1670 | 0 | 3704 | TRUE | 4 |
| orf1a | 1671 | 0 | 3704 | FALSE | 0 |
| orf1a | 1672 | 1186 | 3704 | TRUE | 2 |
| orf1a | 1673 | 1186 | 3704 | TRUE | 3 |
| orf1a | 1674 | 1186 | 3704 | TRUE | 3 |
| orf1a | 1675 | 1186 | 3704 | TRUE | 3 |
| orf1a | 1676 | 1186 | 3704 | TRUE | 2 |
| orf1a | 1677 | 1186 | 3704 | TRUE | 5 |
| orf1a | 1678 | 1186 | 3704 | TRUE | 1 |
| orf1a | 1679 | 1186 | 3704 | TRUE | 4 |
| orf1a | 1680 | 1186 | 3704 | TRUE | 4 |
| orf1a | 1681 | 1186 | 3704 | TRUE | 3 |
| orf1a | 1682 | 1186 | 3704 | TRUE | 3 |
| orf1a | 1683 | 1186 | 3704 | TRUE | 3 |
| orf1a | 1684 | 1186 | 3704 | TRUE | 1 |
| orf1a | 1685 | 1186 | 0 | FALSE | 0 |
| orf1a | 1686 | 1186 | 0 | FALSE | 2 |
| orf1a | 1704 | 0 | 1438 | FALSE | 5 |
| orf1a | 1705 | 0 | 1438 | FALSE | 2 |
| orf1a | 1706 | 14732 | 4238 | FALSE | 4 |
| orf1a | 1707 | 14732 | 4238 | FALSE | 4 |
| orf1a | 1708 | 14732 | 4238 | FALSE | 2 |
| orf1a | 1709 | 14732 | 4238 | FALSE | 1 |
| orf1a | 1710 | 14732 | 4238 | FALSE | 2 |
| orf1a | 1711 | 14732 | 4238 | FALSE | 3 |
| orf1a | 1712 | 14732 | 4238 | FALSE | 2 |
| orf1a | 1713 | 14732 | 4238 | FALSE | 1 |
| orf1a | 1714 | 8146 | 4238 | FALSE | 2 |
| orf1a | 1715 | 8146 | 4238 | FALSE | 2 |
| orf1a | 1716 | 8146 | 4238 | FALSE | 3 |
| orf1a | 1717 | 8146 | 4238 | FALSE | 2 |
| orf1a | 1718 | 8146 | 4238 | FALSE | 1 |
| orf1a | 1719 | 8146 | 7038 | FALSE | 4 |
| orf1a | 1720 | 5888 | 7038 | FALSE | 3 |
| orf1a | 1721 | 1466 | 0 | FALSE | 1 |

| | | | | | |
|---|---|---|---|---|---|
| orf1a | 1722 | 5315 | 0 | FALSE | 4 |
| orf1a | 1723 | 5315 | 0 | FALSE | 1 |
| orf1a | 1724 | 7866 | 4691 | FALSE | 3 |
| orf1a | 1725 | 7866 | 4691 | FALSE | 3 |
| orf1a | 1726 | 6932 | 4691 | FALSE | 2 |
| orf1a | 1727 | 6932 | 4691 | FALSE | 2 |
| orf1a | 1728 | 6932 | 4691 | FALSE | 2 |
| orf1a | 1729 | 7827 | 4691 | FALSE | 3 |
| orf1a | 1730 | 8378 | 4433 | FALSE | 4 |
| orf1a | 1731 | 8378 | 4433 | FALSE | 1 |
| orf1a | 1732 | 8378 | 4433 | FALSE | 3 |
| orf1a | 1733 | 8378 | 4433 | FALSE | 4 |
| orf1a | 1734 | 8378 | 4433 | FALSE | 5 |
| orf1a | 1735 | 9253 | 4433 | FALSE | 1 |
| orf1a | 1736 | 9253 | 4433 | FALSE | 1 |
| orf1a | 1737 | 8717 | 4433 | FALSE | 3 |
| orf1a | 1738 | 8717 | 4433 | FALSE | 2 |
| orf1a | 1739 | 7441 | 3916 | FALSE | 6 |
| orf1a | 1740 | 6691 | 3916 | FALSE | 2 |
| orf1a | 1741 | 8032 | 3916 | FALSE | 1 |
| orf1a | 1742 | 8032 | 3916 | FALSE | 3 |
| orf1a | 1743 | 8032 | 3916 | FALSE | 2 |
| orf1a | 1744 | 8032 | 3916 | FALSE | 6 |
| orf1a | 1745 | 3690 | 0 | FALSE | 1 |
| orf1a | 1746 | 3690 | 0 | FALSE | 4 |
| orf1a | 1747 | 3690 | 0 | FALSE | 2 |
| orf1a | 1748 | 3690 | 0 | FALSE | 1 |
| orf1a | 1749 | 3690 | 0 | FALSE | 3 |
| orf1a | 1750 | 3690 | 0 | FALSE | 4 |
| orf1a | 1751 | 3690 | 0 | FALSE | 1 |
| orf1a | 1752 | 3690 | 3056 | FALSE | 0 |
| orf1a | 1753 | 3690 | 3056 | FALSE | 1 |
| orf1a | 1754 | 3690 | 3056 | FALSE | 1 |
| orf1a | 1755 | 0 | 3056 | FALSE | 1 |
| orf1a | 1756 | 0 | 2148 | FALSE | 6 |
| orf1a | 1757 | 0 | 2148 | FALSE | 2 |
| orf1a | 1758 | 0 | 2148 | FALSE | 3 |
| orf1a | 1759 | 0 | 2148 | FALSE | 3 |
| orf1a | 1760 | 0 | 2148 | FALSE | 2 |
| orf1a | 1761 | 0 | 2148 | FALSE | 3 |
| orf1a | 1762 | 0 | 2148 | FALSE | 2 |
| orf1a | 1763 | 0 | 2148 | FALSE | 2 |
| orf1a | 1764 | 0 | 9985 | FALSE | 2 |
| orf1a | 1765 | 0 | 9985 | FALSE | 4 |
| orf1a | 1766 | 10829 | 9867 | FALSE | 4 |
| orf1a | 1767 | 10829 | 11570 | FALSE | 2 |
| orf1a | 1768 | 10829 | 11570 | FALSE | 1 |
| orf1a | 1769 | 10829 | 11570 | FALSE | 2 |
| orf1a | 1770 | 10829 | 9713 | FALSE | 3 |
| orf1a | 1771 | 10829 | 11831 | FALSE | 2 |
| orf1a | 1772 | 10829 | 9889 | FALSE | 3 |
| orf1a | 1773 | 10829 | 9889 | FALSE | 2 |
| orf1a | 1774 | 10829 | 9889 | FALSE | 2 |
| orf1a | 1775 | 10829 | 9889 | FALSE | 2 |
| orf1a | 1776 | 10829 | 9889 | FALSE | 3 |
| orf1a | 1777 | 10829 | 9889 | FALSE | 4 |
| orf1a | 1778 | 10829 | 9889 | FALSE | 7 |
| orf1a | 1779 | 10829 | 5946 | FALSE | 2 |
| orf1a | 1780 | 10829 | 5385 | FALSE | 3 |
| orf1a | 1781 | 0 | 2514 | FALSE | 2 |
| orf1a | 1782 | 0 | 3063 | FALSE | 3 |
| orf1a | 1783 | 0 | 3063 | FALSE | 2 |

| | | | | | |
|---|---|---|---|---|---|
| orf1a | 1784 | 0 | 3063 | FALSE | 4 |
| orf1a | 1785 | 0 | 3323 | FALSE | 1 |
| orf1a | 1786 | 0 | 3323 | FALSE | 2 |
| orf1a | 1787 | 0 | 3923 | FALSE | 1 |
| orf1a | 1788 | 0 | 3923 | FALSE | 3 |
| orf1a | 1789 | 0 | 3923 | FALSE | 2 |
| orf1a | 1790 | 0 | 3923 | FALSE | 4 |
| orf1a | 1791 | 0 | 3923 | FALSE | 1 |
| orf1a | 1792 | 0 | 3923 | FALSE | 1 |
| orf1a | 1793 | 0 | 3923 | FALSE | 5 |
| orf1a | 1794 | 0 | 3923 | FALSE | 1 |
| orf1a | 1795 | 0 | 4709 | FALSE | 2 |
| orf1a | 1796 | 0 | 4709 | FALSE | 2 |
| orf1a | 1824 | 1800 | 0 | FALSE | 2 |
| orf1a | 1825 | 1800 | 0 | FALSE | 2 |
| orf1a | 1826 | 4123 | 3130 | FALSE | 2 |
| orf1a | 1827 | 4123 | 3130 | FALSE | 2 |
| orf1a | 1828 | 4123 | 3130 | FALSE | 2 |
| orf1a | 1829 | 4123 | 3130 | FALSE | 1 |
| orf1a | 1830 | 4123 | 3130 | FALSE | 3 |
| orf1a | 1831 | 4123 | 3130 | FALSE | 2 |
| orf1a | 1832 | 4123 | 3130 | FALSE | 0 |
| orf1a | 1833 | 4123 | 3130 | FALSE | 1 |
| orf1a | 1834 | 4123 | 3130 | FALSE | 2 |
| orf1a | 1835 | 4123 | 3130 | FALSE | 1 |
| orf1a | 1836 | 4123 | 3130 | FALSE | 2 |
| orf1a | 1837 | 4123 | 3130 | FALSE | 2 |
| orf1a | 1838 | 4123 | 3130 | FALSE | 3 |
| orf1a | 1839 | 8770 | 3130 | FALSE | 3 |
| orf1a | 1840 | 8770 | 3130 | FALSE | 1 |
| orf1a | 1852 | 1958 | 0 | FALSE | 3 |
| orf1a | 1853 | 1958 | 0 | FALSE | 2 |
| orf1a | 1854 | 1958 | 0 | FALSE | 2 |
| orf1a | 1855 | 1958 | 0 | FALSE | 1 |
| orf1a | 1856 | 1958 | 1322 | FALSE | 5 |
| orf1a | 1857 | 1958 | 1322 | FALSE | 2 |
| orf1a | 1858 | 7047 | 2528 | FALSE | 3 |
| orf1a | 1859 | 7047 | 2528 | FALSE | 3 |
| orf1a | 1860 | 5185 | 2528 | FALSE | 1 |
| orf1a | 1861 | 5185 | 2528 | FALSE | 4 |
| orf1a | 1862 | 4739 | 2856 | FALSE | 2 |
| orf1a | 1863 | 4739 | 2856 | FALSE | 1 |
| orf1a | 1864 | 5955 | 2856 | FALSE | 3 |
| orf1a | 1865 | 5955 | 2856 | FALSE | 1 |
| orf1a | 1866 | 5955 | 2856 | FALSE | 3 |
| orf1a | 1867 | 6755 | 2856 | FALSE | 3 |
| orf1a | 1868 | 6169 | 2856 | FALSE | 2 |
| orf1a | 1869 | 6169 | 2856 | FALSE | 2 |
| orf1a | 1870 | 7469 | 3023 | FALSE | 3 |
| orf1a | 1871 | 7469 | 3307 | FALSE | 1 |
| orf1a | 1872 | 7469 | 3307 | FALSE | 2 |
| orf1a | 1873 | 6691 | 3395 | FALSE | 2 |
| orf1a | 1874 | 6691 | 2600 | FALSE | 1 |
| orf1a | 1875 | 7737 | 2600 | FALSE | 2 |
| orf1a | 1876 | 7737 | 2600 | FALSE | 2 |
| orf1a | 1877 | 10182 | 2226 | FALSE | 2 |
| orf1a | 1878 | 10182 | 2226 | FALSE | 6 |
| orf1a | 1879 | 9255 | 2226 | FALSE | 3 |
| orf1a | 1880 | 6842 | 2226 | FALSE | 1 |
| orf1a | 1881 | 6842 | 2226 | FALSE | 5 |
| orf1a | 1882 | 6842 | 2226 | FALSE | 3 |
| orf1a | 1883 | 8644 | 2226 | FALSE | 3 |

| | | | | | |
|---|---|---|---|---|---|
| orf1a | 1884 | 8644 | 2666 | FALSE | 2 |
| orf1a | 1885 | 2017 | 2150 | FALSE | 5 |
| orf1a | 1886 | 2017 | 6352 | FALSE | 4 |
| orf1a | 1887 | 2017 | 6352 | FALSE | 1 |
| orf1a | 1888 | 1592 | 6352 | FALSE | 4 |
| orf1a | 1889 | 1592 | 11693 | FALSE | 3 |
| orf1a | 1890 | 1481 | 8143 | FALSE | 2 |
| orf1a | 1891 | 1481 | 8143 | FALSE | 4 |
| orf1a | 1892 | 1481 | 8143 | FALSE | 1 |
| orf1a | 1893 | 1481 | 8143 | FALSE | 2 |
| orf1a | 1894 | 1481 | 8143 | FALSE | 3 |
| orf1a | 1895 | 1302 | 8143 | FALSE | 2 |
| orf1a | 1896 | 1302 | 8143 | FALSE | 3 |
| orf1a | 1897 | 1302 | 8143 | FALSE | 2 |
| orf1a | 1898 | 1302 | 8143 | FALSE | 3 |
| orf1a | 1899 | 1302 | 10002 | FALSE | 1 |
| orf1a | 1900 | 1302 | 10002 | FALSE | 3 |
| orf1a | 1901 | 1302 | 1044 | FALSE | 5 |
| orf1a | 1902 | 1302 | 1044 | FALSE | 3 |
| orf1a | 1903 | 1370 | 1044 | FALSE | 3 |
| orf1a | 1904 | 1451 | 1044 | FALSE | 3 |
| orf1a | 1905 | 1612 | 0 | FALSE | 2 |
| orf1a | 1906 | 1612 | 0 | FALSE | 3 |
| orf1a | 1907 | 1612 | 0 | FALSE | 3 |
| orf1a | 1908 | 1593 | 0 | FALSE | 2 |
| orf1a | 1909 | 1593 | 0 | FALSE | 2 |
| orf1a | 1910 | 1543 | 0 | FALSE | 3 |
| orf1a | 1911 | 1543 | 0 | FALSE | 2 |
| orf1a | 1912 | 1543 | 0 | FALSE | 1 |
| orf1a | 1913 | 1543 | 0 | FALSE | 2 |
| orf1a | 1914 | 1543 | 0 | FALSE | 3 |
| orf1a | 1915 | 1543 | 0 | FALSE | 3 |
| orf1a | 1916 | 1543 | 0 | FALSE | 3 |
| orf1a | 1917 | 1543 | 0 | FALSE | 6 |
| orf1a | 1918 | 1543 | 0 | FALSE | 3 |
| orf1a | 1919 | 1529 | 0 | FALSE | 4 |
| orf1a | 1920 | 1529 | 0 | FALSE | 3 |
| orf1a | 1921 | 1529 | 0 | FALSE | 6 |
| orf1a | 1922 | 1546 | 0 | FALSE | 5 |
| orf1a | 1923 | 1461 | 0 | FALSE | 3 |
| orf1a | 1924 | 1461 | 0 | FALSE | 2 |
| orf1a | 1925 | 1629 | 0 | FALSE | 2 |
| orf1a | 1926 | 1629 | 0 | FALSE | 3 |
| orf1a | 1927 | 1629 | 0 | FALSE | 2 |
| orf1a | 1928 | 1629 | 0 | FALSE | 2 |
| orf1a | 1929 | 1629 | 0 | FALSE | 1 |
| orf1a | 1930 | 1629 | 0 | FALSE | 2 |
| orf1a | 1931 | 1629 | 0 | FALSE | 2 |
| orf1a | 1932 | 1629 | 0 | FALSE | 4 |
| orf1a | 1933 | 1629 | 0 | FALSE | 3 |
| orf1a | 1934 | 1629 | 0 | FALSE | 2 |
| orf1a | 1935 | 1629 | 0 | FALSE | 3 |
| orf1a | 1936 | 1324 | 0 | FALSE | 3 |
| orf1a | 1937 | 1019 | 0 | FALSE | 5 |
| orf1a | 1938 | 1272 | 0 | FALSE | 4 |
| orf1a | 1939 | 1272 | 0 | FALSE | 3 |
| orf1a | 1940 | 1272 | 0 | FALSE | 2 |
| orf1a | 1941 | 1272 | 0 | FALSE | 2 |
| orf1a | 1942 | 1298 | 0 | FALSE | 2 |
| orf1a | 1943 | 1298 | 0 | FALSE | 4 |
| orf1a | 1944 | 1298 | 0 | FALSE | 3 |
| orf1a | 1945 | 1298 | 0 | FALSE | 3 |

| | | | | | |
|---|---|---|---|---|---|
| orf1a | 1946 | 1298 | 0 | FALSE | 3 |
| orf1a | 1947 | 1298 | 0 | FALSE | 3 |
| orf1a | 1948 | 1298 | 0 | FALSE | 2 |
| orf1a | 1949 | 1298 | 0 | FALSE | 2 |
| orf1a | 1950 | 1298 | 0 | FALSE | 4 |
| orf1a | 1951 | 1437 | 0 | FALSE | 4 |
| orf1a | 1952 | 1437 | 2330 | FALSE | 3 |
| orf1a | 1953 | 1350 | 2330 | FALSE | 3 |
| orf1a | 1954 | 4412 | 2529 | FALSE | 4 |
| orf1a | 1955 | 4412 | 2529 | FALSE | 3 |
| orf1a | 1956 | 4412 | 2529 | FALSE | 4 |
| orf1a | 1957 | 5433 | 2529 | FALSE | 1 |
| orf1a | 1958 | 5433 | 2529 | FALSE | 4 |
| orf1a | 1959 | 5433 | 2529 | FALSE | 4 |
| orf1a | 1960 | 4437 | 2529 | FALSE | 2 |
| orf1a | 1961 | 4437 | 2529 | FALSE | 3 |
| orf1a | 1962 | 4437 | 2529 | FALSE | 5 |
| orf1a | 1963 | 4437 | 2529 | FALSE | 2 |
| orf1a | 1964 | 4437 | 2529 | FALSE | 3 |
| orf1a | 1965 | 4437 | 2529 | FALSE | 2 |
| orf1a | 1966 | 4437 | 2529 | FALSE | 3 |
| orf1a | 1967 | 4437 | 2596 | FALSE | 2 |
| orf1a | 1968 | 4437 | 2596 | FALSE | 2 |
| orf1a | 1969 | 1447 | 0 | FALSE | 4 |
| orf1a | 1970 | 1447 | 0 | FALSE | 1 |
| orf1a | 1971 | 1447 | 0 | FALSE | 2 |
| orf1a | 1972 | 1447 | 0 | FALSE | 1 |
| orf1a | 1973 | 1447 | 0 | FALSE | 1 |
| orf1a | 1974 | 1447 | 0 | FALSE | 2 |
| orf1a | 1980 | 2463 | 0 | FALSE | 4 |
| orf1a | 1981 | 2463 | 0 | FALSE | 2 |
| orf1a | 1982 | 2463 | 0 | FALSE | 4 |
| orf1a | 1983 | 2463 | 0 | FALSE | 4 |
| orf1a | 1984 | 1785 | 0 | FALSE | 2 |
| orf1a | 1985 | 1785 | 0 | FALSE | 3 |
| orf1a | 1986 | 1785 | 0 | FALSE | 2 |
| orf1a | 1987 | 1785 | 0 | FALSE | 3 |
| orf1a | 1988 | 1785 | 0 | FALSE | 2 |
| orf1a | 1989 | 1785 | 0 | FALSE | 4 |
| orf1a | 1990 | 1785 | 0 | FALSE | 4 |
| orf1a | 1991 | 1785 | 0 | FALSE | 4 |
| orf1a | 1992 | 1785 | 0 | FALSE | 3 |
| orf1a | 1993 | 1785 | 0 | FALSE | 5 |
| orf1a | 1994 | 1785 | 0 | FALSE | 4 |
| orf1a | 1995 | 1107 | 0 | FALSE | 3 |
| orf1a | 1996 | 1474 | 1117 | FALSE | 1 |
| orf1a | 1997 | 1474 | 1117 | FALSE | 3 |
| orf1a | 1998 | 1474 | 1117 | FALSE | 2 |
| orf1a | 1999 | 1840 | 1117 | FALSE | 0 |
| orf1a | 2000 | 1840 | 1117 | FALSE | 2 |
| orf1a | 2001 | 1840 | 1117 | FALSE | 4 |
| orf1a | 2002 | 1840 | 1117 | FALSE | 2 |
| orf1a | 2003 | 1840 | 1117 | FALSE | 1 |
| orf1a | 2004 | 1840 | 1117 | FALSE | 2 |
| orf1a | 2005 | 1840 | 1117 | FALSE | 2 |
| orf1a | 2006 | 1840 | 1117 | FALSE | 3 |
| orf1a | 2007 | 1840 | 1117 | FALSE | 4 |
| orf1a | 2008 | 1840 | 1117 | FALSE | 0 |
| orf1a | 2009 | 1840 | 1117 | FALSE | 1 |
| orf1a | 2010 | 1840 | 1117 | FALSE | 3 |
| orf1a | 2014 | 2253 | 0 | FALSE | 2 |
| orf1a | 2015 | 2253 | 0 | FALSE | 5 |

| | | | | | |
|---|---|---|---|---|---|
| orf1a | 2016 | 4638 | 0 | FALSE | 4 |
| orf1a | 2017 | 4638 | 0 | FALSE | 2 |
| orf1a | 2018 | 4803 | 3715 | FALSE | 4 |
| orf1a | 2019 | 4803 | 3715 | FALSE | 2 |
| orf1a | 2020 | 5439 | 3272 | FALSE | 3 |
| orf1a | 2021 | 5439 | 3272 | FALSE | 2 |
| orf1a | 2022 | 5439 | 2852 | FALSE | 3 |
| orf1a | 2023 | 5439 | 2852 | FALSE | 3 |
| orf1a | 2024 | 5439 | 2506 | FALSE | 3 |
| orf1a | 2025 | 5439 | 2506 | FALSE | 2 |
| orf1a | 2026 | 5439 | 2506 | FALSE | 2 |
| orf1a | 2027 | 5439 | 2506 | FALSE | 2 |
| orf1a | 2028 | 5439 | 2328 | FALSE | 4 |
| orf1a | 2029 | 5758 | 2328 | FALSE | 3 |
| orf1a | 2030 | 5758 | 2328 | FALSE | 2 |
| orf1a | 2031 | 5897 | 2328 | FALSE | 4 |
| orf1a | 2032 | 5897 | 2328 | FALSE | 3 |
| orf1a | 2033 | 11802 | 1866 | FALSE | 4 |
| orf1a | 2034 | 11802 | 1785 | FALSE | 3 |
| orf1a | 2035 | 0 | 1367 | FALSE | 6 |
| orf1a | 2036 | 2146 | 1367 | FALSE | 4 |
| orf1a | 2037 | 2146 | 1416 | FALSE | 4 |
| orf1a | 2038 | 4823 | 1416 | FALSE | 2 |
| orf1a | 2039 | 4823 | 1189 | FALSE | 3 |
| orf1a | 2040 | 6929 | 1189 | FALSE | 4 |
| orf1a | 2041 | 6929 | 1189 | FALSE | 2 |
| orf1a | 2042 | 10885 | 3098 | FALSE | 1 |
| orf1a | 2043 | 10885 | 3771 | FALSE | 5 |
| orf1a | 2044 | 9784 | 3724 | FALSE | 2 |
| orf1a | 2045 | 9784 | 3724 | FALSE | 3 |
| orf1a | 2046 | 10127 | 5068 | FALSE | 2 |
| orf1a | 2047 | 10127 | 5068 | FALSE | 3 |
| orf1a | 2048 | 10774 | 4273 | FALSE | 3 |
| orf1a | 2049 | 10774 | 4698 | FALSE | 3 |
| orf1a | 2050 | 10576 | 4872 | FALSE | 3 |
| orf1a | 2051 | 11250 | 4872 | FALSE | 4 |
| orf1a | 2052 | 11250 | 4872 | FALSE | 3 |
| orf1a | 2053 | 11295 | 4872 | FALSE | 3 |
| orf1a | 2054 | 11295 | 4872 | FALSE | 2 |
| orf1a | 2055 | 11405 | 4872 | FALSE | 2 |
| orf1a | 2056 | 11405 | 4872 | FALSE | 2 |
| orf1a | 2057 | 10446 | 4838 | FALSE | 1 |
| orf1a | 2058 | 10446 | 4838 | FALSE | 3 |
| orf1a | 2059 | 11311 | 5018 | FALSE | 3 |
| orf1a | 2060 | 11311 | 5018 | FALSE | 2 |
| orf1a | 2061 | 12146 | 3923 | FALSE | 2 |
| orf1a | 2062 | 12146 | 3923 | FALSE | 2 |
| orf1a | 2063 | 8995 | 5279 | FALSE | 3 |
| orf1a | 2064 | 8995 | 5279 | FALSE | 1 |
| orf1a | 2070 | 8497 | 2433 | FALSE | 1 |
| orf1a | 2071 | 8497 | 2433 | FALSE | 3 |
| orf1a | 2072 | 8497 | 2433 | FALSE | 2 |
| orf1a | 2073 | 8497 | 2433 | FALSE | 4 |
| orf1a | 2074 | 8497 | 2433 | FALSE | 3 |
| orf1a | 2075 | 8497 | 2433 | FALSE | 0 |
| orf1a | 2076 | 8497 | 2326 | FALSE | 2 |
| orf1a | 2077 | 8497 | 2326 | FALSE | 4 |
| orf1a | 2078 | 8497 | 2326 | FALSE | 3 |
| orf1a | 2079 | 8497 | 2326 | FALSE | 3 |
| orf1a | 2080 | 8497 | 2326 | FALSE | 4 |
| orf1a | 2081 | 8497 | 2326 | FALSE | 5 |
| orf1a | 2082 | 8457 | 2326 | FALSE | 4 |

| | | | | | |
|---|---|---|---|---|---|
| orf1a | 2083 | 8457 | 2326 | FALSE | 3 |
| orf1a | 2084 | 8457 | 2157 | FALSE | 2 |
| orf1a | 2085 | 8378 | 1743 | FALSE | 3 |
| orf1a | 2086 | 13733 | 2543 | FALSE | 3 |
| orf1a | 2087 | 13733 | 2543 | FALSE | 3 |
| orf1a | 2088 | 14470 | 5569 | FALSE | 3 |
| orf1a | 2089 | 14470 | 5569 | FALSE | 5 |
| orf1a | 2090 | 14470 | 5569 | FALSE | 1 |
| orf1a | 2091 | 14470 | 6163 | FALSE | 4 |
| orf1a | 2092 | 13069 | 6163 | FALSE | 2 |
| orf1a | 2093 | 13069 | 6163 | FALSE | 2 |
| orf1a | 2094 | 13069 | 6163 | FALSE | 2 |
| orf1a | 2095 | 13069 | 6163 | FALSE | 2 |
| orf1a | 2096 | 13069 | 6163 | FALSE | 2 |
| orf1a | 2097 | 13655 | 6163 | FALSE | 2 |
| orf1a | 2098 | 13655 | 6163 | FALSE | 1 |
| orf1a | 2099 | 13655 | 7099 | FALSE | 1 |
| orf1a | 2100 | 13655 | 7099 | FALSE | 3 |
| orf1a | 2101 | 12736 | 7838 | FALSE | 2 |
| orf1a | 2102 | 11201 | 7838 | FALSE | 4 |
| orf1a | 2103 | 1926 | 0 | FALSE | 4 |
| orf1a | 2104 | 1926 | 0 | FALSE | 2 |
| orf1a | 2105 | 1926 | 0 | FALSE | 2 |
| orf1a | 2106 | 1926 | 0 | FALSE | 1 |
| orf1a | 2107 | 1991 | 0 | FALSE | 4 |
| orf1a | 2108 | 1991 | 0 | FALSE | 3 |
| orf1a | 2109 | 1991 | 0 | FALSE | 2 |
| orf1a | 2110 | 1991 | 0 | FALSE | 3 |
| orf1a | 2111 | 1991 | 0 | FALSE | 2 |
| orf1a | 2112 | 3174 | 2395 | FALSE | 2 |
| orf1a | 2113 | 3174 | 2395 | FALSE | 2 |
| orf1a | 2114 | 3174 | 2395 | FALSE | 3 |
| orf1a | 2115 | 3174 | 2395 | FALSE | 4 |
| orf1a | 2116 | 3174 | 2054 | FALSE | 5 |
| orf1a | 2117 | 4357 | 2054 | FALSE | 2 |
| orf1a | 2118 | 4357 | 2382 | FALSE | 1 |
| orf1a | 2119 | 4357 | 2382 | FALSE | 4 |
| orf1a | 2120 | 4357 | 2382 | FALSE | 1 |
| orf1a | 2121 | 4357 | 2382 | FALSE | 3 |
| orf1a | 2122 | 4357 | 2382 | FALSE | 3 |
| orf1a | 2123 | 4357 | 2382 | FALSE | 4 |
| orf1a | 2124 | 4357 | 3477 | FALSE | 3 |
| orf1a | 2125 | 4357 | 3477 | FALSE | 2 |
| orf1a | 2126 | 4357 | 4162 | FALSE | 4 |
| orf1a | 2127 | 0 | 5045 | FALSE | 4 |
| orf1a | 2128 | 0 | 5045 | FALSE | 5 |
| orf1a | 2129 | 0 | 5045 | FALSE | 4 |
| orf1a | 2130 | 0 | 5045 | FALSE | 3 |
| orf1a | 2131 | 0 | 6269 | FALSE | 1 |
| orf1a | 2132 | 1424 | 6269 | FALSE | 1 |
| orf1a | 2133 | 1424 | 7721 | FALSE | 2 |
| orf1a | 2134 | 1424 | 7721 | FALSE | 3 |
| orf1a | 2135 | 1424 | 7721 | FALSE | 5 |
| orf1a | 2136 | 1424 | 7721 | FALSE | 3 |
| orf1a | 2137 | 1424 | 7721 | FALSE | 2 |
| orf1a | 2138 | 1424 | 7721 | FALSE | 2 |
| orf1a | 2139 | 1424 | 7585 | FALSE | 4 |
| orf1a | 2140 | 1364 | 7585 | FALSE | 3 |
| orf1a | 2141 | 1364 | 0 | FALSE | 0 |
| orf1a | 2142 | 1364 | 2713 | FALSE | 1 |
| orf1a | 2143 | 1364 | 2713 | FALSE | 1 |
| orf1a | 2144 | 1364 | 2713 | FALSE | 4 |

| | | | | | |
|---|---|---|---|---|---|
| orf1a | 2145 | 1364 | 2713 | FALSE | 3 |
| orf1a | 2146 | 1364 | 2713 | FALSE | 1 |
| orf1a | 2147 | 1243 | 2713 | FALSE | 2 |
| orf1a | 2148 | 1243 | 2713 | FALSE | 3 |
| orf1a | 2149 | 1243 | 2713 | FALSE | 5 |
| orf1a | 2150 | 1243 | 2713 | FALSE | 2 |
| orf1a | 2151 | 1243 | 2713 | FALSE | 3 |
| orf1a | 2152 | 1243 | 2713 | FALSE | 2 |
| orf1a | 2153 | 1243 | 2713 | FALSE | 2 |
| orf1a | 2154 | 1243 | 2713 | FALSE | 2 |
| orf1a | 2155 | 0 | 2713 | FALSE | 3 |
| orf1a | 2156 | 0 | 2713 | FALSE | 2 |
| orf1a | 2162 | 11957 | 0 | FALSE | 1 |
| orf1a | 2163 | 11957 | 0 | FALSE | 2 |
| orf1a | 2164 | 11957 | 0 | FALSE | 2 |
| orf1a | 2165 | 11957 | 0 | FALSE | 2 |
| orf1a | 2166 | 11957 | 0 | FALSE | 2 |
| orf1a | 2167 | 11957 | 0 | FALSE | 1 |
| orf1a | 2168 | 11957 | 0 | FALSE | 2 |
| orf1a | 2169 | 11957 | 0 | FALSE | 2 |
| orf1a | 2170 | 11957 | 0 | FALSE | 4 |
| orf1a | 2171 | 11957 | 0 | FALSE | 1 |
| orf1a | 2172 | 11957 | 0 | FALSE | 3 |
| orf1a | 2173 | 11957 | 0 | FALSE | 2 |
| orf1a | 2174 | 11957 | 0 | FALSE | 3 |
| orf1a | 2175 | 11957 | 0 | FALSE | 0 |
| orf1a | 2176 | 11957 | 0 | FALSE | 2 |
| orf1a | 2194 | 2239 | 0 | FALSE | 3 |
| orf1a | 2195 | 2239 | 0 | FALSE | 2 |
| orf1a | 2196 | 2239 | 0 | FALSE | 2 |
| orf1a | 2197 | 2239 | 0 | FALSE | 2 |
| orf1a | 2198 | 2239 | 0 | FALSE | 2 |
| orf1a | 2199 | 2239 | 0 | FALSE | 2 |
| orf1a | 2200 | 2239 | 0 | FALSE | 3 |
| orf1a | 2201 | 2239 | 0 | FALSE | 2 |
| orf1a | 2202 | 2239 | 0 | FALSE | 3 |
| orf1a | 2203 | 2239 | 0 | FALSE | 3 |
| orf1a | 2204 | 2239 | 0 | FALSE | 2 |
| orf1a | 2205 | 2239 | 0 | FALSE | 1 |
| orf1a | 2206 | 2239 | 0 | FALSE | 1 |
| orf1a | 2207 | 2239 | 0 | FALSE | 2 |
| orf1a | 2208 | 2239 | 0 | FALSE | 0 |
| orf1a | 2212 | 19011 | 8381 | FALSE | 2 |
| orf1a | 2213 | 19011 | 8381 | FALSE | 2 |
| orf1a | 2214 | 19011 | 8381 | FALSE | 4 |
| orf1a | 2215 | 19011 | 8381 | FALSE | 0 |
| orf1a | 2216 | 10588 | 8381 | FALSE | 3 |
| orf1a | 2217 | 10588 | 8381 | FALSE | 1 |
| orf1a | 2218 | 10588 | 8381 | FALSE | 0 |
| orf1a | 2219 | 10588 | 8381 | FALSE | 2 |
| orf1a | 2220 | 10588 | 8381 | FALSE | 2 |
| orf1a | 2221 | 10588 | 8381 | FALSE | 2 |
| orf1a | 2222 | 10588 | 8381 | FALSE | 2 |
| orf1a | 2223 | 10588 | 8381 | FALSE | 0 |
| orf1a | 2224 | 10588 | 8381 | FALSE | 2 |
| orf1a | 2225 | 10588 | 8381 | FALSE | 1 |
| orf1a | 2226 | 10588 | 8381 | FALSE | 2 |
| orf1a | 2227 | 2166 | 0 | FALSE | 3 |
| orf1a | 2228 | 2166 | 0 | FALSE | 1 |
| orf1a | 2229 | 2166 | 0 | FALSE | 2 |
| orf1a | 2230 | 2166 | 0 | FALSE | 2 |
| orf1a | 2238 | 0 | 1370 | FALSE | 2 |

| | | | | | |
|---|---|---|---|---|---|
| orf1a | 2239 | 0 | 1370 | FALSE | 4 |
| orf1a | 2240 | 0 | 1370 | FALSE | 2 |
| orf1a | 2241 | 0 | 1370 | FALSE | 1 |
| orf1a | 2242 | 0 | 1370 | FALSE | 2 |
| orf1a | 2243 | 0 | 1370 | FALSE | 2 |
| orf1a | 2244 | 0 | 1370 | FALSE | 2 |
| orf1a | 2245 | 0 | 1370 | FALSE | 1 |
| orf1a | 2246 | 5291 | 1370 | FALSE | 3 |
| orf1a | 2247 | 5291 | 1370 | FALSE | 3 |
| orf1a | 2248 | 3473 | 1370 | FALSE | 3 |
| orf1a | 2249 | 3473 | 1370 | FALSE | 6 |
| orf1a | 2250 | 3473 | 1370 | FALSE | 1 |
| orf1a | 2251 | 3473 | 1370 | FALSE | 5 |
| orf1a | 2252 | 3473 | 1370 | FALSE | 2 |
| orf1a | 2253 | 3473 | 0 | FALSE | 2 |
| orf1a | 2254 | 3473 | 0 | FALSE | 3 |
| orf1a | 2255 | 3473 | 0 | FALSE | 3 |
| orf1a | 2256 | 4695 | 5556 | FALSE | 4 |
| orf1a | 2257 | 4695 | 5556 | FALSE | 1 |
| orf1a | 2258 | 4477 | 6903 | FALSE | 3 |
| orf1a | 2259 | 4477 | 6903 | FALSE | 1 |
| orf1a | 2260 | 4477 | 6903 | FALSE | 2 |
| orf1a | 2261 | 4341 | 6903 | FALSE | 2 |
| orf1a | 2262 | 4212 | 5902 | FALSE | 1 |
| orf1a | 2263 | 4638 | 5902 | FALSE | 0 |
| orf1a | 2264 | 4247 | 4538 | FALSE | 1 |
| orf1a | 2265 | 4247 | 4538 | FALSE | 3 |
| orf1a | 2266 | 4247 | 3897 | FALSE | 3 |
| orf1a | 2267 | 4247 | 3897 | FALSE | 8 |
| orf1a | 2268 | 4514 | 4873 | FALSE | 6 |
| orf1a | 2269 | 4514 | 4873 | FALSE | 6 |
| orf1a | 2270 | 4514 | 4873 | FALSE | 3 |
| orf1a | 2271 | 4114 | 4804 | FALSE | 2 |
| orf1a | 2272 | 4114 | 4804 | FALSE | 4 |
| orf1a | 2273 | 4059 | 4422 | FALSE | 3 |
| orf1a | 2274 | 4059 | 4422 | FALSE | 4 |
| orf1a | 2275 | 4059 | 4422 | FALSE | 1 |
| orf1a | 2276 | 4059 | 4422 | FALSE | 5 |
| orf1a | 2277 | 4266 | 4487 | FALSE | 1 |
| orf1a | 2278 | 4266 | 4487 | FALSE | 1 |
| orf1a | 2279 | 5450 | 5152 | FALSE | 3 |
| orf1a | 2280 | 5450 | 5152 | FALSE | 4 |
| orf1a | 2281 | 5450 | 6580 | FALSE | 2 |
| orf1a | 2282 | 5450 | 5502 | FALSE | 1 |
| orf1a | 2283 | 0 | 1193 | FALSE | 1 |
| orf1a | 2284 | 0 | 1193 | FALSE | 1 |
| orf1a | 2285 | 0 | 1193 | FALSE | 2 |
| orf1a | 2286 | 0 | 1193 | FALSE | 1 |
| orf1a | 2287 | 0 | 1193 | FALSE | 2 |
| orf1a | 2288 | 0 | 1193 | FALSE | 4 |
| orf1a | 2289 | 0 | 1193 | FALSE | 1 |
| orf1a | 2290 | 0 | 1193 | FALSE | 1 |
| orf1a | 2291 | 0 | 1193 | FALSE | 0 |
| orf1a | 2292 | 0 | 1193 | FALSE | 2 |
| orf1a | 2293 | 0 | 1193 | FALSE | 1 |
| orf1a | 2294 | 0 | 13717 | FALSE | 5 |
| orf1a | 2295 | 0 | 13717 | FALSE | 1 |
| orf1a | 2296 | 1876 | 13717 | FALSE | 2 |
| orf1a | 2297 | 1876 | 26241 | FALSE | 2 |
| orf1a | 2298 | 1876 | 26241 | FALSE | 1 |
| orf1a | 2299 | 1876 | 26241 | FALSE | 6 |
| orf1a | 2300 | 3148 | 26241 | FALSE | 4 |

| | | | | | |
|---|---|---|---|---|---|
| orf1a | 2301 | 3148 | 26241 | FALSE | 2 |
| orf1a | 2302 | 3148 | 26241 | FALSE | 3 |
| orf1a | 2303 | 3148 | 26241 | FALSE | 2 |
| orf1a | 2304 | 3148 | 26241 | FALSE | 2 |
| orf1a | 2305 | 3148 | 26241 | FALSE | 2 |
| orf1a | 2306 | 3148 | 26241 | FALSE | 4 |
| orf1a | 2307 | 3148 | 26241 | FALSE | 2 |
| orf1a | 2308 | 3148 | 26241 | FALSE | 1 |
| orf1a | 2309 | 3148 | 0 | FALSE | 5 |
| orf1a | 2310 | 3148 | 0 | FALSE | 2 |
| orf1a | 2311 | 6964 | 0 | FALSE | 3 |
| orf1a | 2312 | 6964 | 0 | FALSE | 3 |
| orf1a | 2313 | 6964 | 0 | FALSE | 2 |
| orf1a | 2314 | 6964 | 0 | FALSE | 1 |
| orf1a | 2322 | 0 | 8015 | FALSE | 6 |
| orf1a | 2323 | 0 | 8015 | FALSE | 0 |
| orf1a | 2324 | 6205 | 8015 | FALSE | 1 |
| orf1a | 2325 | 6205 | 8015 | FALSE | 3 |
| orf1a | 2326 | 5650 | 6606 | FALSE | 5 |
| orf1a | 2327 | 5650 | 6606 | FALSE | 3 |
| orf1a | 2328 | 5650 | 6606 | FALSE | 3 |
| orf1a | 2329 | 5650 | 6606 | FALSE | 3 |
| orf1a | 2330 | 5650 | 6606 | FALSE | 1 |
| orf1a | 2331 | 5650 | 6606 | FALSE | 1 |
| orf1a | 2332 | 5650 | 6606 | FALSE | 1 |
| orf1a | 2333 | 5650 | 6606 | FALSE | 4 |
| orf1a | 2334 | 5650 | 6606 | FALSE | 2 |
| orf1a | 2335 | 5650 | 6606 | FALSE | 2 |
| orf1a | 2336 | 5650 | 6606 | FALSE | 4 |
| orf1a | 2337 | 5650 | 5198 | FALSE | 1 |
| orf1a | 2338 | 5650 | 5198 | FALSE | 3 |
| orf1a | 2339 | 5095 | 5198 | FALSE | 2 |
| orf1a | 2340 | 5095 | 4842 | FALSE | 0 |
| orf1a | 2341 | 0 | 4485 | FALSE | 3 |
| orf1a | 2342 | 0 | 3918 | FALSE | 0 |
| orf1a | 2343 | 0 | 3918 | FALSE | 3 |
| orf1a | 2344 | 0 | 3918 | FALSE | 3 |
| orf1a | 2345 | 0 | 3918 | FALSE | 4 |
| orf1a | 2346 | 0 | 3918 | FALSE | 2 |
| orf1a | 2347 | 0 | 3918 | FALSE | 3 |
| orf1a | 2348 | 0 | 3918 | FALSE | 3 |
| orf1a | 2349 | 0 | 3918 | FALSE | 3 |
| orf1a | 2350 | 0 | 3918 | FALSE | 1 |
| orf1a | 2351 | 0 | 3918 | FALSE | 2 |
| orf1a | 2352 | 0 | 3918 | FALSE | 3 |
| orf1a | 2353 | 0 | 3918 | FALSE | 2 |
| orf1a | 2354 | 0 | 3918 | FALSE | 0 |
| orf1a | 2355 | 0 | 3351 | FALSE | 6 |
| orf1a | 2356 | 0 | 3351 | FALSE | 3 |
| orf1a | 2370 | 12993 | 0 | FALSE | 1 |
| orf1a | 2371 | 12993 | 0 | FALSE | 2 |
| orf1a | 2372 | 12993 | 0 | FALSE | 1 |
| orf1a | 2373 | 12993 | 0 | FALSE | 3 |
| orf1a | 2374 | 12993 | 0 | FALSE | 3 |
| orf1a | 2375 | 12993 | 0 | FALSE | 3 |
| orf1a | 2376 | 7038 | 0 | FALSE | 4 |
| orf1a | 2377 | 7038 | 0 | FALSE | 3 |
| orf1a | 2378 | 7038 | 0 | FALSE | 3 |
| orf1a | 2379 | 7038 | 0 | FALSE | 3 |
| orf1a | 2380 | 5098 | 0 | FALSE | 1 |
| orf1a | 2381 | 5098 | 0 | FALSE | 3 |
| orf1a | 2382 | 5098 | 0 | FALSE | 1 |

| | | | | | |
|---|---|---|---|---|---|
| orf1a | 2383 | 5098 | 0 | FALSE | 0 |
| orf1a | 2384 | 5098 | 0 | FALSE | 1 |
| orf1a | 2385 | 1151 | 0 | FALSE | 3 |
| orf1a | 2386 | 1151 | 0 | FALSE | 2 |
| orf1a | 2387 | 1151 | 0 | FALSE | 2 |
| orf1a | 2388 | 1151 | 0 | FALSE | 2 |
| orf1a | 2389 | 1151 | 0 | FALSE | 1 |
| orf1a | 2390 | 1151 | 0 | FALSE | 1 |
| orf1a | 2391 | 1218 | 0 | FALSE | 4 |
| orf1a | 2392 | 1218 | 0 | FALSE | 3 |
| orf1a | 2393 | 1218 | 0 | FALSE | 3 |
| orf1a | 2394 | 1218 | 0 | FALSE | 1 |
| orf1a | 2416 | 14774 | 1793 | TRUE | 2 |
| orf1a | 2417 | 14774 | 1793 | FALSE | 4 |
| orf1a | 2418 | 14774 | 1793 | FALSE | 5 |
| orf1a | 2419 | 14774 | 1793 | FALSE | 1 |
| orf1a | 2420 | 10348 | 2004 | TRUE | 6 |
| orf1a | 2421 | 10348 | 2004 | TRUE | 2 |
| orf1a | 2422 | 8167 | 2104 | FALSE | 4 |
| orf1a | 2423 | 8167 | 2104 | TRUE | 3 |
| orf1a | 2424 | 8167 | 2104 | TRUE | 4 |
| orf1a | 2425 | 8167 | 2104 | TRUE | 2 |
| orf1a | 2426 | 8167 | 2104 | TRUE | 1 |
| orf1a | 2427 | 8167 | 2104 | TRUE | 2 |
| orf1a | 2428 | 7089 | 2104 | TRUE | 4 |
| orf1a | 2429 | 7089 | 2104 | TRUE | 4 |
| orf1a | 2430 | 7089 | 2104 | TRUE | 4 |
| orf1a | 2431 | 5168 | 2208 | FALSE | 4 |
| orf1a | 2432 | 5168 | 2208 | TRUE | 4 |
| orf1a | 2433 | 5168 | 2208 | TRUE | 3 |
| orf1a | 2434 | 5168 | 2208 | TRUE | 3 |
| orf1a | 2435 | 4917 | 2205 | FALSE | 1 |
| orf1a | 2436 | 4917 | 2205 | TRUE | 4 |
| orf1a | 2437 | 2777 | 0 | TRUE | 2 |
| orf1a | 2438 | 2777 | 0 | TRUE | 3 |
| orf1a | 2439 | 2777 | 0 | TRUE | 1 |
| orf1a | 2440 | 2777 | 0 | TRUE | 3 |
| orf1a | 2441 | 2777 | 0 | TRUE | 4 |
| orf1a | 2442 | 1894 | 0 | FALSE | 4 |
| orf1a | 2443 | 1011 | 0 | FALSE | 5 |
| orf1a | 2444 | 1011 | 0 | TRUE | 3 |
| orf1a | 2445 | 1011 | 0 | TRUE | 3 |
| orf1a | 2446 | 1011 | 0 | FALSE | 1 |
| orf1a | 2447 | 1011 | 0 | TRUE | 4 |
| orf1a | 2448 | 1011 | 0 | TRUE | 2 |
| orf1a | 2449 | 1011 | 0 | TRUE | 1 |
| orf1a | 2450 | 1011 | 0 | TRUE | 3 |
| orf1a | 2451 | 1011 | 0 | TRUE | 2 |
| orf1a | 2452 | 1011 | 0 | TRUE | 2 |
| orf1a | 2453 | 1011 | 0 | TRUE | 0 |
| orf1a | 2454 | 1264 | 0 | TRUE | 1 |
| orf1a | 2455 | 1264 | 0 | TRUE | 3 |
| orf1a | 2456 | 1191 | 0 | FALSE | 4 |
| orf1a | 2457 | 1281 | 0 | TRUE | 4 |
| orf1a | 2458 | 1281 | 0 | TRUE | 4 |
| orf1a | 2459 | 1281 | 0 | FALSE | 1 |
| orf1a | 2460 | 1281 | 0 | FALSE | 5 |
| orf1a | 2461 | 1281 | 0 | TRUE | 1 |
| orf1a | 2462 | 1281 | 0 | TRUE | 1 |
| orf1a | 2463 | 1281 | 0 | TRUE | 2 |
| orf1a | 2464 | 1281 | 0 | TRUE | 3 |
| orf1a | 2465 | 1281 | 0 | TRUE | 0 |

| | | | | | |
|---|---|---|---|---|---|
| orf1a | 2466 | 3220 | 0 | TRUE | 5 |
| orf1a | 2467 | 3220 | 0 | FALSE | 3 |
| orf1a | 2468 | 7558 | 3265 | FALSE | 4 |
| orf1a | 2469 | 9068 | 3265 | FALSE | 1 |
| orf1a | 2470 | 9068 | 3265 | TRUE | 5 |
| orf1a | 2471 | 11742 | 3265 | FALSE | 4 |
| orf1a | 2472 | 11742 | 3265 | TRUE | 1 |
| orf1a | 2473 | 11742 | 3265 | TRUE | 3 |
| orf1a | 2474 | 11742 | 3265 | TRUE | 1 |
| orf1a | 2475 | 11742 | 3265 | FALSE | 2 |
| orf1a | 2476 | 11742 | 3265 | FALSE | 3 |
| orf1a | 2477 | 11742 | 3265 | TRUE | 2 |
| orf1a | 2478 | 11742 | 3265 | TRUE | 3 |
| orf1a | 2479 | 11742 | 3265 | TRUE | 1 |
| orf1a | 2480 | 7382 | 3005 | TRUE | 5 |
| orf1a | 2481 | 8272 | 3005 | TRUE | 1 |
| orf1a | 2482 | 7079 | 3005 | TRUE | 1 |
| orf1a | 2483 | 3513 | 2832 | TRUE | 2 |
| orf1a | 2484 | 5192 | 2832 | TRUE | 2 |
| orf1a | 2485 | 5192 | 2832 | FALSE | 2 |
| orf1a | 2486 | 5192 | 2832 | FALSE | 2 |
| orf1a | 2487 | 5192 | 2832 | TRUE | 2 |
| orf1a | 2488 | 5192 | 2832 | FALSE | 3 |
| orf1a | 2489 | 5192 | 2832 | FALSE | 1 |
| orf1a | 2490 | 5192 | 2832 | TRUE | 4 |
| orf1a | 2491 | 5192 | 2832 | TRUE | 3 |
| orf1a | 2492 | 4661 | 2832 | TRUE | 2 |
| orf1a | 2493 | 4661 | 2832 | FALSE | 4 |
| orf1a | 2494 | 4661 | 2832 | TRUE | 2 |
| orf1a | 2495 | 5393 | 0 | FALSE | 6 |
| orf1a | 2496 | 5393 | 0 | FALSE | 3 |
| orf1a | 2497 | 7530 | 0 | FALSE | 2 |
| orf1a | 2498 | 7530 | 0 | FALSE | 4 |
| orf1a | 2499 | 1474 | 0 | FALSE | 2 |
| orf1a | 2500 | 1474 | 0 | FALSE | 3 |
| orf1a | 2501 | 1474 | 0 | TRUE | 2 |
| orf1a | 2502 | 1474 | 0 | TRUE | 1 |
| orf1a | 2503 | 1474 | 0 | TRUE | 2 |
| orf1a | 2504 | 1474 | 0 | TRUE | 1 |
| orf1a | 2505 | 1474 | 0 | FALSE | 3 |
| orf1a | 2506 | 1474 | 0 | FALSE | 1 |
| orf1a | 2512 | 2865 | 0 | TRUE | 3 |
| orf1a | 2513 | 2865 | 0 | FALSE | 0 |
| orf1a | 2514 | 6020 | 4548 | FALSE | 4 |
| orf1a | 2515 | 6020 | 4548 | FALSE | 3 |
| orf1a | 2516 | 6020 | 4548 | FALSE | 1 |
| orf1a | 2517 | 6020 | 4548 | FALSE | 4 |
| orf1a | 2518 | 6020 | 4548 | FALSE | 4 |
| orf1a | 2519 | 6020 | 4548 | FALSE | 1 |
| orf1a | 2520 | 6020 | 4548 | FALSE | 5 |
| orf1a | 2521 | 6020 | 4548 | FALSE | 0 |
| orf1a | 2522 | 6020 | 4548 | FALSE | 2 |
| orf1a | 2523 | 6020 | 4548 | FALSE | 2 |
| orf1a | 2524 | 6020 | 4548 | FALSE | 0 |
| orf1a | 2525 | 6020 | 4548 | FALSE | 3 |
| orf1a | 2526 | 5109 | 4548 | FALSE | 2 |
| orf1a | 2527 | 5857 | 4548 | FALSE | 2 |
| orf1a | 2528 | 5857 | 4548 | FALSE | 3 |
| orf1a | 2529 | 2377 | 0 | FALSE | 5 |
| orf1a | 2530 | 3424 | 1281 | FALSE | 0 |
| orf1a | 2531 | 3424 | 1281 | FALSE | 2 |
| orf1a | 2532 | 3424 | 1281 | FALSE | 2 |

| | | | | | |
|---|---|---|---|---|---|
| orf1a | 2533 | 3424 | 1281 | FALSE | 0 |
| orf1a | 2534 | 3124 | 1281 | FALSE | 3 |
| orf1a | 2535 | 3124 | 1281 | FALSE | 4 |
| orf1a | 2536 | 3124 | 1281 | FALSE | 1 |
| orf1a | 2537 | 3124 | 1281 | FALSE | 3 |
| orf1a | 2538 | 3124 | 1281 | FALSE | 1 |
| orf1a | 2539 | 3124 | 1281 | FALSE | 2 |
| orf1a | 2540 | 3124 | 1281 | FALSE | 0 |
| orf1a | 2541 | 3498 | 1281 | FALSE | 1 |
| orf1a | 2542 | 3498 | 1281 | FALSE | 2 |
| orf1a | 2543 | 3498 | 1281 | FALSE | 4 |
| orf1a | 2544 | 3498 | 1281 | FALSE | 2 |
| orf1a | 2545 | 2526 | 0 | FALSE | 5 |
| orf1a | 2546 | 2526 | 0 | FALSE | 1 |
| orf1a | 2547 | 2526 | 0 | FALSE | 2 |
| orf1a | 2548 | 21770 | 3173 | FALSE | 2 |
| orf1a | 2549 | 31392 | 3173 | FALSE | 3 |
| orf1a | 2550 | 17958 | 7096 | FALSE | 5 |
| orf1a | 2551 | 17958 | 7096 | FALSE | 3 |
| orf1a | 2552 | 17958 | 7096 | FALSE | 2 |
| orf1a | 2553 | 17958 | 7096 | FALSE | 3 |
| orf1a | 2554 | 17958 | 7096 | FALSE | 2 |
| orf1a | 2555 | 17958 | 7096 | FALSE | 1 |
| orf1a | 2556 | 17958 | 7096 | TRUE | 4 |
| orf1a | 2557 | 17958 | 7096 | TRUE | 5 |
| orf1a | 2558 | 17958 | 7096 | TRUE | 2 |
| orf1a | 2559 | 17958 | 7096 | TRUE | 3 |
| orf1a | 2560 | 17958 | 7096 | TRUE | 4 |
| orf1a | 2561 | 17958 | 7096 | TRUE | 1 |
| orf1a | 2562 | 17958 | 7096 | TRUE | 2 |
| orf1a | 2563 | 13480 | 8077 | TRUE | 4 |
| orf1a | 2564 | 13480 | 8077 | FALSE | 2 |
| orf1a | 2568 | 1592 | 2747 | TRUE | 5 |
| orf1a | 2569 | 1592 | 2747 | TRUE | 1 |
| orf1a | 2570 | 1592 | 2747 | TRUE | 4 |
| orf1a | 2571 | 1592 | 2747 | TRUE | 1 |
| orf1a | 2572 | 1592 | 2747 | TRUE | 3 |
| orf1a | 2573 | 1592 | 2747 | TRUE | 2 |
| orf1a | 2574 | 1592 | 2747 | FALSE | 3 |
| orf1a | 2575 | 1592 | 2747 | FALSE | 4 |
| orf1a | 2576 | 1500 | 2747 | FALSE | 3 |
| orf1a | 2577 | 1500 | 2747 | FALSE | 2 |
| orf1a | 2578 | 2226 | 2747 | FALSE | 2 |
| orf1a | 2579 | 2226 | 2747 | FALSE | 2 |
| orf1a | 2580 | 2226 | 2747 | FALSE | 3 |
| orf1a | 2581 | 2226 | 2747 | FALSE | 2 |
| orf1a | 2582 | 2226 | 2747 | FALSE | 0 |
| orf1a | 2583 | 2543 | 0 | FALSE | 2 |
| orf1a | 2584 | 2543 | 0 | FALSE | 7 |
| orf1a | 2585 | 2543 | 0 | FALSE | 2 |
| orf1a | 2586 | 2148 | 0 | FALSE | 3 |
| orf1a | 2587 | 2148 | 0 | FALSE | 2 |
| orf1a | 2588 | 2148 | 0 | FALSE | 2 |
| orf1a | 2589 | 2148 | 0 | FALSE | 2 |
| orf1a | 2590 | 2148 | 1486 | FALSE | 1 |
| orf1a | 2591 | 2517 | 1486 | FALSE | 3 |
| orf1a | 2592 | 2517 | 1486 | FALSE | 1 |
| orf1a | 2593 | 1357 | 1486 | FALSE | 1 |
| orf1a | 2594 | 1357 | 1486 | FALSE | 1 |
| orf1a | 2595 | 1357 | 1486 | FALSE | 4 |
| orf1a | 2596 | 1357 | 1486 | FALSE | 2 |
| orf1a | 2597 | 1357 | 1486 | FALSE | 4 |

| | | | | | |
|---|---|---|---|---|---|
| orf1a | 2598 | 1357 | 1486 | FALSE | 1 |
| orf1a | 2599 | 1357 | 1486 | FALSE | 2 |
| orf1a | 2600 | 1357 | 1486 | FALSE | 2 |
| orf1a | 2601 | 0 | 1486 | FALSE | 3 |
| orf1a | 2602 | 0 | 1486 | FALSE | 1 |
| orf1a | 2603 | 0 | 1486 | FALSE | 3 |
| orf1a | 2604 | 1251 | 1486 | FALSE | 5 |
| orf1a | 2605 | 1251 | 0 | FALSE | 4 |
| orf1a | 2606 | 1251 | 1255 | FALSE | 4 |
| orf1a | 2607 | 1251 | 1255 | FALSE | 5 |
| orf1a | 2608 | 1251 | 1255 | FALSE | 3 |
| orf1a | 2609 | 1251 | 1255 | FALSE | 4 |
| orf1a | 2610 | 1251 | 1255 | FALSE | 0 |
| orf1a | 2611 | 1251 | 1255 | FALSE | 1 |
| orf1a | 2612 | 1251 | 1255 | FALSE | 2 |
| orf1a | 2613 | 1251 | 1255 | FALSE | 2 |
| orf1a | 2614 | 1251 | 2277 | FALSE | 3 |
| orf1a | 2615 | 1251 | 2277 | FALSE | 2 |
| orf1a | 2616 | 1251 | 2277 | FALSE | 3 |
| orf1a | 2617 | 1251 | 2277 | FALSE | 4 |
| orf1a | 2618 | 1251 | 2277 | FALSE | 2 |
| orf1a | 2619 | 0 | 2277 | FALSE | 2 |
| orf1a | 2620 | 0 | 2277 | FALSE | 2 |
| orf1a | 2621 | 0 | 3300 | FALSE | 3 |
| orf1a | 2622 | 0 | 3300 | FALSE | 6 |
| orf1a | 2623 | 0 | 3300 | FALSE | 2 |
| orf1a | 2624 | 0 | 3300 | FALSE | 5 |
| orf1a | 2625 | 0 | 3300 | FALSE | 3 |
| orf1a | 2626 | 0 | 3300 | FALSE | 4 |
| orf1a | 2627 | 0 | 3300 | FALSE | 2 |
| orf1a | 2628 | 0 | 3300 | FALSE | 4 |
| orf1a | 2636 | 0 | 5019 | FALSE | 4 |
| orf1a | 2637 | 0 | 5019 | FALSE | 2 |
| orf1a | 2638 | 1665 | 5976 | FALSE | 5 |
| orf1a | 2639 | 1665 | 5976 | FALSE | 3 |
| orf1a | 2640 | 1654 | 5339 | FALSE | 5 |
| orf1a | 2641 | 1654 | 5339 | FALSE | 3 |
| orf1a | 2642 | 1843 | 4898 | FALSE | 1 |
| orf1a | 2643 | 1843 | 4898 | FALSE | 0 |
| orf1a | 2644 | 2141 | 4274 | FALSE | 3 |
| orf1a | 2645 | 2141 | 4274 | FALSE | 1 |
| orf1a | 2646 | 1978 | 4274 | FALSE | 1 |
| orf1a | 2647 | 1978 | 4274 | FALSE | 3 |
| orf1a | 2648 | 1978 | 4274 | FALSE | 3 |
| orf1a | 2649 | 1978 | 4274 | FALSE | 2 |
| orf1a | 2650 | 1978 | 4274 | FALSE | 2 |
| orf1a | 2651 | 1978 | 4125 | FALSE | 3 |
| orf1a | 2652 | 1978 | 4125 | FALSE | 1 |
| orf1a | 2653 | 2031 | 3423 | FALSE | 3 |
| orf1a | 2654 | 2031 | 3423 | FALSE | 2 |
| orf1a | 2655 | 2221 | 2145 | FALSE | 5 |
| orf1a | 2656 | 2221 | 2145 | FALSE | 3 |
| orf1a | 2657 | 2316 | 1154 | FALSE | 1 |
| orf1a | 2658 | 2316 | 1154 | FALSE | 2 |
| orf1a | 2659 | 1005 | 0 | FALSE | 3 |
| orf1a | 2660 | 1005 | 0 | FALSE | 3 |
| orf1a | 2662 | 9060 | 1710 | FALSE | 2 |
| orf1a | 2663 | 9060 | 1710 | FALSE | 2 |
| orf1a | 2664 | 10791 | 2537 | FALSE | 6 |
| orf1a | 2665 | 10791 | 2537 | FALSE | 1 |
| orf1a | 2666 | 9235 | 2318 | FALSE | 1 |
| orf1a | 2667 | 9235 | 2318 | FALSE | 4 |

| | | | | | |
|---|---|---|---|---|---|
| orf1a | 2668 | 9235 | 2318 | FALSE | 2 |
| orf1a | 2669 | 9235 | 2318 | FALSE | 3 |
| orf1a | 2670 | 9235 | 2318 | FALSE | 2 |
| orf1a | 2671 | 9235 | 2318 | FALSE | 1 |
| orf1a | 2672 | 8070 | 2318 | FALSE | 2 |
| orf1a | 2673 | 8070 | 2318 | FALSE | 3 |
| orf1a | 2674 | 8070 | 2318 | FALSE | 1 |
| orf1a | 2675 | 8070 | 2318 | FALSE | 3 |
| orf1a | 2676 | 8070 | 2318 | FALSE | 4 |
| orf1a | 2677 | 7822 | 2440 | FALSE | 3 |
| orf1a | 2678 | 9947 | 2440 | FALSE | 5 |
| orf1a | 2679 | 8808 | 1222 | FALSE | 2 |
| orf1a | 2680 | 8808 | 1222 | FALSE | 3 |
| orf1a | 2681 | 10929 | 0 | FALSE | 3 |
| orf1a | 2682 | 8910 | 0 | FALSE | 2 |
| orf1a | 2683 | 8910 | 0 | FALSE | 3 |
| orf1a | 2684 | 6702 | 0 | FALSE | 3 |
| orf1a | 2685 | 6702 | 0 | FALSE | 5 |
| orf1a | 2686 | 7885 | 0 | FALSE | 5 |
| orf1a | 2687 | 8780 | 0 | FALSE | 3 |
| orf1a | 2688 | 8780 | 0 | FALSE | 4 |
| orf1a | 2689 | 8780 | 0 | FALSE | 2 |
| orf1a | 2690 | 8780 | 0 | FALSE | 4 |
| orf1a | 2691 | 8780 | 0 | FALSE | 2 |
| orf1a | 2692 | 8780 | 0 | FALSE | 0 |
| orf1a | 2693 | 6363 | 0 | FALSE | 4 |
| orf1a | 2694 | 5464 | 0 | FALSE | 3 |
| orf1a | 2695 | 5464 | 0 | FALSE | 3 |
| orf1a | 2696 | 5464 | 0 | FALSE | 4 |
| orf1a | 2697 | 5612 | 0 | FALSE | 2 |
| orf1a | 2698 | 5612 | 0 | FALSE | 3 |
| orf1a | 2699 | 7834 | 0 | FALSE | 5 |
| orf1a | 2700 | 5824 | 0 | FALSE | 4 |
| orf1a | 2701 | 1838 | 0 | FALSE | 3 |
| orf1a | 2702 | 1838 | 0 | FALSE | 3 |
| orf1a | 2703 | 1838 | 0 | FALSE | 2 |
| orf1a | 2704 | 1838 | 0 | FALSE | 4 |
| orf1a | 2705 | 1838 | 0 | FALSE | 2 |
| orf1a | 2706 | 1838 | 0 | FALSE | 3 |
| orf1a | 2707 | 1838 | 0 | FALSE | 3 |
| orf1a | 2708 | 1838 | 0 | FALSE | 2 |
| orf1a | 2709 | 1806 | 0 | FALSE | 1 |
| orf1a | 2710 | 1806 | 0 | FALSE | 5 |
| orf1a | 2711 | 1806 | 0 | FALSE | 1 |
| orf1a | 2712 | 1806 | 0 | FALSE | 1 |
| orf1a | 2713 | 1806 | 0 | FALSE | 4 |
| orf1a | 2714 | 1806 | 0 | FALSE | 2 |
| orf1a | 2736 | 5081 | 0 | FALSE | 1 |
| orf1a | 2737 | 5081 | 0 | FALSE | 1 |
| orf1a | 2738 | 5081 | 0 | FALSE | 0 |
| orf1a | 2739 | 5081 | 0 | FALSE | 4 |
| orf1a | 2740 | 5081 | 0 | FALSE | 4 |
| orf1a | 2741 | 5081 | 0 | FALSE | 3 |
| orf1a | 2742 | 5081 | 0 | FALSE | 1 |
| orf1a | 2743 | 5081 | 0 | FALSE | 3 |
| orf1a | 2744 | 5081 | 0 | FALSE | 5 |
| orf1a | 2745 | 5081 | 0 | FALSE | 2 |
| orf1a | 2746 | 5081 | 0 | FALSE | 4 |
| orf1a | 2747 | 5081 | 0 | FALSE | 3 |
| orf1a | 2748 | 5081 | 0 | FALSE | 1 |
| orf1a | 2749 | 5081 | 0 | FALSE | 2 |
| orf1a | 2750 | 5081 | 0 | FALSE | 2 |

| | | | | | |
|---|---|---|---|---|---|
| orf1a | 2768 | 2047 | 0 | FALSE | 2 |
| orf1a | 2769 | 2047 | 0 | FALSE | 4 |
| orf1a | 2770 | 2047 | 0 | FALSE | 1 |
| orf1a | 2771 | 2047 | 0 | FALSE | 1 |
| orf1a | 2772 | 2047 | 0 | FALSE | 3 |
| orf1a | 2773 | 2047 | 0 | FALSE | 2 |
| orf1a | 2774 | 2047 | 0 | FALSE | 3 |
| orf1a | 2775 | 2047 | 0 | FALSE | 2 |
| orf1a | 2776 | 2047 | 0 | FALSE | 3 |
| orf1a | 2777 | 2047 | 0 | FALSE | 2 |
| orf1a | 2778 | 2047 | 0 | FALSE | 2 |
| orf1a | 2779 | 2047 | 0 | FALSE | 3 |
| orf1a | 2780 | 1618 | 0 | FALSE | 2 |
| orf1a | 2781 | 1618 | 0 | FALSE | 3 |
| orf1a | 2782 | 1618 | 0 | FALSE | 0 |
| orf1a | 2783 | 1190 | 0 | FALSE | 2 |
| orf1a | 2784 | 1190 | 0 | FALSE | 1 |
| orf1a | 2785 | 1190 | 0 | FALSE | 2 |
| orf1a | 2786 | 1190 | 0 | FALSE | 4 |
| orf1a | 2787 | 1190 | 0 | FALSE | 3 |
| orf1a | 2788 | 1190 | 0 | FALSE | 4 |
| orf1a | 2789 | 1190 | 0 | FALSE | 2 |
| orf1a | 2790 | 2922 | 0 | FALSE | 3 |
| orf1a | 2791 | 2922 | 0 | FALSE | 2 |
| orf1a | 2792 | 3253 | 3626 | FALSE | 2 |
| orf1a | 2793 | 3253 | 3626 | FALSE | 2 |
| orf1a | 2794 | 3253 | 3626 | FALSE | 3 |
| orf1a | 2795 | 3459 | 3626 | FALSE | 3 |
| orf1a | 2796 | 3459 | 3626 | FALSE | 4 |
| orf1a | 2797 | 3459 | 3626 | FALSE | 1 |
| orf1a | 2798 | 3459 | 3626 | FALSE | 1 |
| orf1a | 2799 | 3459 | 3626 | FALSE | 2 |
| orf1a | 2800 | 4518 | 3626 | FALSE | 1 |
| orf1a | 2801 | 4518 | 3626 | FALSE | 3 |
| orf1a | 2802 | 4518 | 3626 | FALSE | 2 |
| orf1a | 2803 | 4518 | 3626 | FALSE | 2 |
| orf1a | 2804 | 4439 | 3626 | FALSE | 1 |
| orf1a | 2805 | 4872 | 3626 | FALSE | 2 |
| orf1a | 2806 | 4872 | 3626 | FALSE | 3 |
| orf1a | 2807 | 6887 | 0 | FALSE | 1 |
| orf1a | 2808 | 6887 | 0 | FALSE | 3 |
| orf1a | 2809 | 6887 | 0 | FALSE | 3 |
| orf1a | 2810 | 6887 | 0 | FALSE | 2 |
| orf1a | 2811 | 6887 | 0 | FALSE | 2 |
| orf1a | 2812 | 6887 | 0 | FALSE | 2 |
| orf1a | 2813 | 6887 | 0 | FALSE | 3 |
| orf1a | 2814 | 6887 | 0 | FALSE | 2 |
| orf1a | 2815 | 3408 | 0 | FALSE | 2 |
| orf1a | 2816 | 3408 | 0 | TRUE | 2 |
| orf1a | 2817 | 3408 | 0 | TRUE | 2 |
| orf1a | 2818 | 13406 | 0 | TRUE | 6 |
| orf1a | 2819 | 23405 | 0 | FALSE | 1 |
| orf1a | 2820 | 23405 | 0 | TRUE | 2 |
| orf1a | 2821 | 23405 | 0 | FALSE | 4 |
| orf1a | 2822 | 12309 | 0 | TRUE | 1 |
| orf1a | 2823 | 12309 | 0 | FALSE | 2 |
| orf1a | 2824 | 12309 | 0 | FALSE | 3 |
| orf1a | 2825 | 12309 | 0 | FALSE | 3 |
| orf1a | 2826 | 12309 | 0 | TRUE | 1 |
| orf1a | 2827 | 12309 | 0 | TRUE | 1 |
| orf1a | 2828 | 12309 | 0 | FALSE | 3 |
| orf1a | 2829 | 12309 | 0 | TRUE | 5 |

| | | | | | |
|---|---|---|---|---|---|
| orf1a | 2830 | 12309 | 0 | FALSE | 1 |
| orf1a | 2831 | 12309 | 0 | FALSE | 4 |
| orf1a | 2832 | 12309 | 0 | FALSE | 3 |
| orf1a | 2833 | 1213 | 0 | FALSE | 3 |
| orf1a | 2834 | 1213 | 0 | FALSE | 2 |
| orf1a | 2835 | 1213 | 0 | FALSE | 3 |
| orf1a | 2836 | 1213 | 0 | FALSE | 2 |
| orf1a | 2846 | 0 | 1153 | FALSE | 2 |
| orf1a | 2847 | 0 | 1153 | FALSE | 1 |
| orf1a | 2848 | 0 | 1153 | FALSE | 2 |
| orf1a | 2849 | 0 | 1153 | FALSE | 0 |
| orf1a | 2850 | 0 | 1153 | FALSE | 5 |
| orf1a | 2851 | 0 | 1153 | FALSE | 6 |
| orf1a | 2852 | 0 | 1153 | FALSE | 4 |
| orf1a | 2853 | 0 | 1153 | FALSE | 3 |
| orf1a | 2854 | 0 | 1153 | FALSE | 2 |
| orf1a | 2855 | 0 | 1153 | FALSE | 4 |
| orf1a | 2856 | 0 | 1153 | FALSE | 3 |
| orf1a | 2857 | 0 | 1153 | FALSE | 3 |
| orf1a | 2858 | 0 | 1153 | FALSE | 0 |
| orf1a | 2859 | 0 | 1153 | FALSE | 1 |
| orf1a | 2860 | 0 | 1153 | FALSE | 4 |
| orf1a | 2866 | 0 | 3545 | FALSE | 1 |
| orf1a | 2867 | 0 | 3545 | FALSE | 1 |
| orf1a | 2868 | 2004 | 6088 | FALSE | 3 |
| orf1a | 2869 | 2004 | 6088 | FALSE | 1 |
| orf1a | 2870 | 2004 | 6088 | FALSE | 2 |
| orf1a | 2871 | 2004 | 6088 | FALSE | 0 |
| orf1a | 2872 | 2004 | 6088 | FALSE | 1 |
| orf1a | 2873 | 2004 | 6088 | FALSE | 3 |
| orf1a | 2874 | 2004 | 6088 | FALSE | 1 |
| orf1a | 2875 | 2004 | 6088 | FALSE | 2 |
| orf1a | 2876 | 2004 | 6088 | FALSE | 1 |
| orf1a | 2877 | 2004 | 6088 | FALSE | 2 |
| orf1a | 2878 | 2004 | 6088 | FALSE | 1 |
| orf1a | 2879 | 2004 | 6088 | FALSE | 2 |
| orf1a | 2880 | 2004 | 6088 | FALSE | 2 |
| orf1a | 2881 | 2004 | 8632 | FALSE | 2 |
| orf1a | 2882 | 2004 | 8632 | FALSE | 0 |
| orf1a | 2894 | 1003 | 0 | FALSE | 2 |
| orf1a | 2895 | 1003 | 0 | FALSE | 3 |
| orf1a | 2896 | 1003 | 0 | FALSE | 1 |
| orf1a | 2897 | 1003 | 0 | FALSE | 2 |
| orf1a | 2898 | 1003 | 0 | FALSE | 3 |
| orf1a | 2899 | 1003 | 0 | FALSE | 3 |
| orf1a | 2900 | 1059 | 2323 | FALSE | 3 |
| orf1a | 2901 | 1059 | 2323 | FALSE | 2 |
| orf1a | 2902 | 1059 | 2323 | FALSE | 4 |
| orf1a | 2903 | 1059 | 2323 | FALSE | 2 |
| orf1a | 2904 | 4554 | 4923 | FALSE | 2 |
| orf1a | 2905 | 4554 | 4923 | FALSE | 3 |
| orf1a | 2906 | 4535 | 4923 | FALSE | 2 |
| orf1a | 2907 | 4535 | 4923 | FALSE | 5 |
| orf1a | 2908 | 4535 | 4923 | FALSE | 3 |
| orf1a | 2909 | 5241 | 4923 | FALSE | 4 |
| orf1a | 2910 | 5241 | 4923 | FALSE | 2 |
| orf1a | 2911 | 5241 | 4923 | FALSE | 4 |
| orf1a | 2912 | 5241 | 4923 | FALSE | 3 |
| orf1a | 2913 | 5241 | 4923 | FALSE | 4 |
| orf1a | 2914 | 5241 | 4923 | FALSE | 4 |
| orf1a | 2915 | 6273 | 7523 | FALSE | 4 |
| orf1a | 2916 | 6273 | 7523 | FALSE | 2 |

| | | | | | |
|---|---|---|---|---|---|
| orf1a | 2917 | 6273 | 7523 | FALSE | 4 |
| orf1a | 2918 | 7189 | 5007 | FALSE | 2 |
| orf1a | 2919 | 7646 | 2492 | FALSE | 3 |
| orf1a | 2920 | 7646 | 2492 | FALSE | 4 |
| orf1a | 2921 | 10853 | 2492 | FALSE | 2 |
| orf1a | 2922 | 10853 | 2492 | FALSE | 2 |
| orf1a | 2923 | 10853 | 2492 | FALSE | 4 |
| orf1a | 2924 | 10853 | 2492 | FALSE | 4 |
| orf1a | 2925 | 10853 | 2492 | FALSE | 4 |
| orf1a | 2926 | 10853 | 2492 | FALSE | 3 |
| orf1a | 2927 | 10853 | 2492 | FALSE | 1 |
| orf1a | 2928 | 10853 | 2492 | FALSE | 3 |
| orf1a | 2929 | 10853 | 2492 | FALSE | 6 |
| orf1a | 2930 | 10853 | 2492 | FALSE | 5 |
| orf1a | 2931 | 10853 | 2492 | FALSE | 2 |
| orf1a | 2932 | 10853 | 2492 | FALSE | 2 |
| orf1a | 2938 | 2531 | 0 | FALSE | 3 |
| orf1a | 2939 | 2531 | 0 | FALSE | 3 |
| orf1a | 2940 | 5891 | 10721 | FALSE | 5 |
| orf1a | 2941 | 5891 | 10721 | FALSE | 3 |
| orf1a | 2942 | 5891 | 10721 | FALSE | 2 |
| orf1a | 2943 | 5891 | 10721 | FALSE | 5 |
| orf1a | 2944 | 5359 | 10721 | FALSE | 5 |
| orf1a | 2945 | 5359 | 10721 | FALSE | 1 |
| orf1a | 2946 | 6360 | 10737 | FALSE | 2 |
| orf1a | 2947 | 6360 | 10737 | FALSE | 1 |
| orf1a | 2948 | 6360 | 9424 | FALSE | 5 |
| orf1a | 2949 | 6360 | 9424 | FALSE | 6 |
| orf1a | 2950 | 6360 | 9424 | FALSE | 3 |
| orf1a | 2951 | 6360 | 9424 | FALSE | 3 |
| orf1a | 2952 | 6360 | 9424 | FALSE | 1 |
| orf1a | 2953 | 6786 | 9424 | FALSE | 3 |
| orf1a | 2954 | 6786 | 9424 | FALSE | 2 |
| orf1a | 2955 | 7065 | 7696 | FALSE | 5 |
| orf1a | 2956 | 7065 | 7696 | FALSE | 3 |
| orf1a | 2957 | 7065 | 7696 | FALSE | 2 |
| orf1a | 2958 | 5917 | 6315 | FALSE | 3 |
| orf1a | 2959 | 6855 | 6315 | FALSE | 5 |
| orf1a | 2960 | 6855 | 6315 | FALSE | 3 |
| orf1a | 2961 | 1327 | 1861 | FALSE | 2 |
| orf1a | 2962 | 1327 | 1861 | FALSE | 1 |
| orf1a | 2963 | 1327 | 2172 | FALSE | 2 |
| orf1a | 2964 | 1327 | 2172 | FALSE | 2 |
| orf1a | 2965 | 1327 | 2172 | FALSE | 2 |
| orf1a | 2966 | 1327 | 2172 | FALSE | 3 |
| orf1a | 2967 | 1327 | 2172 | FALSE | 2 |
| orf1a | 2968 | 1327 | 1588 | FALSE | 2 |
| orf1a | 2969 | 1327 | 1588 | FALSE | 2 |
| orf1a | 2970 | 5677 | 6552 | FALSE | 4 |
| orf1a | 2971 | 5677 | 6552 | FALSE | 5 |
| orf1a | 2972 | 5677 | 6552 | FALSE | 3 |
| orf1a | 2973 | 7128 | 7647 | FALSE | 3 |
| orf1a | 2974 | 7128 | 7647 | FALSE | 0 |
| orf1a | 2975 | 7128 | 7647 | FALSE | 6 |
| orf1a | 2976 | 5922 | 7647 | FALSE | 5 |
| orf1a | 2977 | 5922 | 7647 | FALSE | 2 |
| orf1a | 2978 | 5160 | 7647 | FALSE | 4 |
| orf1a | 2979 | 5160 | 7647 | FALSE | 1 |
| orf1a | 2980 | 5839 | 6334 | FALSE | 5 |
| orf1a | 2981 | 5839 | 6334 | FALSE | 3 |
| orf1a | 2982 | 5302 | 5670 | FALSE | 4 |
| orf1a | 2983 | 5302 | 6447 | TRUE | 2 |

| | | | | | |
|---|---|---|---|---|---|
| orf1a | 2984 | 5302 | 6447 | TRUE | 2 |
| orf1a | 2985 | 4520 | 3034 | TRUE | 5 |
| orf1a | 2986 | 4520 | 3034 | FALSE | 4 |
| orf1a | 2987 | 4520 | 3034 | TRUE | 2 |
| orf1a | 2988 | 4520 | 3034 | FALSE | 3 |
| orf1a | 2989 | 4520 | 3034 | TRUE | 3 |
| orf1a | 2990 | 6001 | 5428 | FALSE | 4 |
| orf1a | 2991 | 6463 | 5428 | FALSE | 3 |
| orf1a | 2992 | 5872 | 5428 | TRUE | 3 |
| orf1a | 2993 | 6342 | 5428 | TRUE | 3 |
| orf1a | 2994 | 6342 | 5428 | FALSE | 3 |
| orf1a | 2995 | 5944 | 6514 | TRUE | 4 |
| orf1a | 2996 | 5944 | 6514 | TRUE | 2 |
| orf1a | 2997 | 7838 | 9018 | TRUE | 3 |
| orf1a | 2998 | 7838 | 9018 | TRUE | 0 |
| orf1a | 2999 | 7838 | 9018 | TRUE | 2 |
| orf1a | 3000 | 6237 | 9018 | TRUE | 2 |
| orf1a | 3001 | 6237 | 9018 | TRUE | 3 |
| orf1a | 3002 | 6237 | 9018 | FALSE | 6 |
| orf1a | 3003 | 6237 | 9018 | FALSE | 4 |
| orf1a | 3004 | 6237 | 9018 | TRUE | 2 |
| orf1a | 3005 | 1290 | 0 | TRUE | 4 |
| orf1a | 3006 | 1290 | 0 | TRUE | 4 |
| orf1a | 3007 | 1433 | 0 | TRUE | 2 |
| orf1a | 3008 | 1433 | 9637 | TRUE | 1 |
| orf1a | 3009 | 1433 | 9637 | FALSE | 2 |
| orf1a | 3010 | 1433 | 8007 | FALSE | 0 |
| orf1a | 3011 | 1433 | 8007 | FALSE | 1 |
| orf1a | 3012 | 1524 | 6140 | TRUE | 1 |
| orf1a | 3013 | 1524 | 6140 | TRUE | 2 |
| orf1a | 3014 | 1524 | 5172 | FALSE | 2 |
| orf1a | 3015 | 1570 | 5172 | TRUE | 3 |
| orf1a | 3016 | 1570 | 6248 | TRUE | 3 |
| orf1a | 3017 | 1570 | 6248 | TRUE | 2 |
| orf1a | 3018 | 1570 | 5547 | TRUE | 1 |
| orf1a | 3019 | 1570 | 5547 | TRUE | 5 |
| orf1a | 3020 | 1570 | 10058 | TRUE | 3 |
| orf1a | 3021 | 1570 | 10058 | FALSE | 2 |
| orf1a | 3022 | 1570 | 10058 | FALSE | 2 |
| orf1a | 3023 | 1570 | 10128 | FALSE | 4 |
| orf1a | 3024 | 1570 | 10128 | FALSE | 3 |
| orf1a | 3025 | 1570 | 10878 | FALSE | 4 |
| orf1a | 3026 | 1570 | 10878 | FALSE | 1 |
| orf1a | 3027 | 0 | 12997 | FALSE | 2 |
| orf1a | 3028 | 0 | 10619 | FALSE | 3 |
| orf1a | 3029 | 0 | 12706 | FALSE | 0 |
| orf1a | 3030 | 0 | 12706 | FALSE | 3 |
| orf1a | 3031 | 0 | 13424 | FALSE | 1 |
| orf1a | 3032 | 0 | 13424 | FALSE | 3 |
| orf1a | 3033 | 0 | 19116 | FALSE | 1 |
| orf1a | 3034 | 0 | 19116 | FALSE | 2 |
| orf1a | 3035 | 0 | 1108 | FALSE | 4 |
| orf1a | 3036 | 0 | 1108 | FALSE | 3 |
| orf1a | 3037 | 0 | 1108 | FALSE | 3 |
| orf1a | 3038 | 0 | 1108 | FALSE | 1 |
| orf1a | 3039 | 0 | 1108 | FALSE | 1 |
| orf1a | 3040 | 0 | 1108 | FALSE | 3 |
| orf1a | 3041 | 0 | 1108 | FALSE | 3 |
| orf1a | 3042 | 0 | 1108 | FALSE | 3 |
| orf1a | 3072 | 0 | 1926 | FALSE | 5 |
| orf1a | 3073 | 0 | 1926 | FALSE | 1 |
| orf1a | 3074 | 0 | 1926 | FALSE | 4 |

| | | | | | |
|---|---|---|---|---|---|
| orf1a | 3075 | 0 | 1926 | FALSE | 1 |
| orf1a | 3076 | 0 | 1926 | FALSE | 3 |
| orf1a | 3077 | 0 | 1926 | FALSE | 1 |
| orf1a | 3078 | 0 | 1926 | FALSE | 3 |
| orf1a | 3079 | 0 | 1926 | FALSE | 3 |
| orf1a | 3080 | 0 | 1926 | FALSE | 2 |
| orf1a | 3081 | 0 | 1926 | FALSE | 2 |
| orf1a | 3082 | 0 | 1926 | FALSE | 3 |
| orf1a | 3083 | 0 | 1926 | FALSE | 3 |
| orf1a | 3084 | 0 | 1926 | FALSE | 1 |
| orf1a | 3085 | 0 | 1926 | FALSE | 4 |
| orf1a | 3086 | 0 | 1926 | FALSE | 1 |
| orf1a | 3094 | 1416 | 0 | FALSE | 0 |
| orf1a | 3095 | 1416 | 0 | FALSE | 1 |
| orf1a | 3096 | 1416 | 1221 | FALSE | 2 |
| orf1a | 3097 | 1416 | 1221 | FALSE | 1 |
| orf1a | 3098 | 1416 | 1221 | FALSE | 1 |
| orf1a | 3099 | 1416 | 1221 | FALSE | 2 |
| orf1a | 3100 | 1416 | 1221 | FALSE | 1 |
| orf1a | 3101 | 1416 | 1221 | FALSE | 0 |
| orf1a | 3102 | 1416 | 1221 | FALSE | 3 |
| orf1a | 3103 | 1416 | 1221 | FALSE | 1 |
| orf1a | 3104 | 1416 | 1221 | FALSE | 2 |
| orf1a | 3105 | 1416 | 1221 | FALSE | 3 |
| orf1a | 3106 | 1416 | 1221 | FALSE | 2 |
| orf1a | 3107 | 1416 | 1221 | FALSE | 0 |
| orf1a | 3108 | 1416 | 1221 | FALSE | 3 |
| orf1a | 3109 | 0 | 1221 | FALSE | 1 |
| orf1a | 3110 | 0 | 1221 | FALSE | 1 |
| orf1a | 3116 | 1018 | 0 | FALSE | 2 |
| orf1a | 3117 | 1018 | 0 | FALSE | 4 |
| orf1a | 3118 | 1267 | 0 | FALSE | 2 |
| orf1a | 3119 | 1267 | 0 | FALSE | 4 |
| orf1a | 3120 | 1267 | 0 | FALSE | 2 |
| orf1a | 3121 | 1267 | 0 | FALSE | 3 |
| orf1a | 3122 | 1267 | 0 | FALSE | 2 |
| orf1a | 3123 | 1267 | 0 | FALSE | 4 |
| orf1a | 3124 | 1267 | 0 | FALSE | 4 |
| orf1a | 3125 | 1267 | 0 | FALSE | 3 |
| orf1a | 3126 | 1267 | 0 | FALSE | 6 |
| orf1a | 3127 | 1267 | 0 | FALSE | 3 |
| orf1a | 3128 | 1267 | 0 | FALSE | 5 |
| orf1a | 3129 | 1267 | 0 | FALSE | 2 |
| orf1a | 3130 | 1267 | 0 | FALSE | 3 |
| orf1a | 3131 | 1516 | 0 | FALSE | 3 |
| orf1a | 3132 | 1516 | 0 | FALSE | 2 |
| orf1a | 3164 | 1225 | 0 | FALSE | 6 |
| orf1a | 3165 | 1225 | 0 | FALSE | 3 |
| orf1a | 3166 | 1188 | 0 | FALSE | 3 |
| orf1a | 3167 | 1188 | 0 | FALSE | 0 |
| orf1a | 3168 | 1209 | 0 | FALSE | 3 |
| orf1a | 3169 | 1209 | 0 | FALSE | 2 |
| orf1a | 3170 | 1209 | 0 | FALSE | 0 |
| orf1a | 3171 | 1209 | 0 | FALSE | 3 |
| orf1a | 3172 | 1209 | 0 | FALSE | 1 |
| orf1a | 3173 | 1209 | 0 | FALSE | 2 |
| orf1a | 3174 | 1245 | 2168 | FALSE | 3 |
| orf1a | 3175 | 1245 | 2168 | FALSE | 0 |
| orf1a | 3176 | 6708 | 5601 | FALSE | 2 |
| orf1a | 3177 | 6708 | 5601 | FALSE | 1 |
| orf1a | 3178 | 6708 | 5601 | FALSE | 2 |
| orf1a | 3179 | 7317 | 5601 | FALSE | 2 |

| | | | | | |
|---|---|---|---|---|---|
| orf1a | 3180 | 7317 | 5601 | FALSE | 5 |
| orf1a | 3181 | 8088 | 5601 | FALSE | 2 |
| orf1a | 3182 | 8088 | 5601 | FALSE | 3 |
| orf1a | 3183 | 10374 | 5601 | FALSE | 4 |
| orf1a | 3184 | 10374 | 5601 | FALSE | 3 |
| orf1a | 3185 | 10374 | 5601 | FALSE | 0 |
| orf1a | 3186 | 11092 | 5601 | FALSE | 4 |
| orf1a | 3187 | 11092 | 5601 | FALSE | 2 |
| orf1a | 3188 | 11590 | 5850 | FALSE | 2 |
| orf1a | 3189 | 15014 | 6464 | FALSE | 4 |
| orf1a | 3190 | 15014 | 6464 | FALSE | 3 |
| orf1a | 3191 | 15239 | 6472 | FALSE | 2 |
| orf1a | 3192 | 15239 | 6472 | FALSE | 2 |
| orf1a | 3193 | 15239 | 6472 | FALSE | 2 |
| orf1a | 3194 | 15239 | 6472 | FALSE | 4 |
| orf1a | 3195 | 15239 | 6472 | FALSE | 4 |
| orf1a | 3196 | 15239 | 6472 | FALSE | 2 |
| orf1a | 3197 | 15239 | 6472 | FALSE | 3 |
| orf1a | 3198 | 15239 | 6472 | FALSE | 4 |
| orf1a | 3199 | 15239 | 6472 | FALSE | 1 |
| orf1a | 3200 | 15239 | 6472 | FALSE | 2 |
| orf1a | 3201 | 15078 | 6472 | FALSE | 1 |
| orf1a | 3202 | 15078 | 6472 | FALSE | 2 |
| orf1a | 3212 | 2492 | 0 | TRUE | 1 |
| orf1a | 3213 | 2492 | 0 | TRUE | 2 |
| orf1a | 3214 | 2492 | 0 | TRUE | 2 |
| orf1a | 3215 | 2492 | 0 | TRUE | 1 |
| orf1a | 3216 | 2492 | 0 | TRUE | 3 |
| orf1a | 3217 | 2492 | 0 | TRUE | 0 |
| orf1a | 3218 | 2492 | 0 | TRUE | 4 |
| orf1a | 3219 | 2492 | 0 | TRUE | 5 |
| orf1a | 3220 | 2492 | 0 | FALSE | 2 |
| orf1a | 3221 | 2492 | 0 | TRUE | 3 |
| orf1a | 3222 | 2492 | 0 | TRUE | 5 |
| orf1a | 3223 | 2492 | 0 | TRUE | 1 |
| orf1a | 3224 | 2492 | 0 | TRUE | 4 |
| orf1a | 3225 | 2492 | 0 | FALSE | 3 |
| orf1a | 3226 | 2492 | 0 | TRUE | 3 |
| orf1a | 3228 | 21472 | 9671 | FALSE | 3 |
| orf1a | 3229 | 21472 | 9671 | TRUE | 3 |
| orf1a | 3230 | 21472 | 9671 | TRUE | 2 |
| orf1a | 3231 | 21472 | 9671 | TRUE | 1 |
| orf1a | 3232 | 21472 | 9671 | TRUE | 1 |
| orf1a | 3233 | 21472 | 9671 | TRUE | 2 |
| orf1a | 3234 | 21472 | 9671 | TRUE | 3 |
| orf1a | 3235 | 21472 | 9671 | TRUE | 3 |
| orf1a | 3236 | 21472 | 9671 | TRUE | 4 |
| orf1a | 3237 | 21472 | 9671 | TRUE | 2 |
| orf1a | 3238 | 21472 | 9671 | TRUE | 4 |
| orf1a | 3239 | 21472 | 9671 | FALSE | 2 |
| orf1a | 3240 | 21472 | 9671 | TRUE | 5 |
| orf1a | 3241 | 21472 | 9671 | TRUE | 5 |
| orf1a | 3242 | 11284 | 6067 | TRUE | 2 |
| orf1a | 3243 | 1095 | 2463 | TRUE | 3 |
| orf1a | 3244 | 1095 | 2463 | TRUE | 1 |
| orf1a | 3245 | 1095 | 2463 | FALSE | 4 |
| orf1a | 3246 | 7286 | 2463 | TRUE | 3 |
| orf1a | 3247 | 7286 | 2463 | TRUE | 2 |
| orf1a | 3248 | 7286 | 2463 | TRUE | 4 |
| orf1a | 3249 | 7286 | 2463 | TRUE | 3 |
| orf1a | 3250 | 7286 | 2463 | TRUE | 2 |
| orf1a | 3251 | 7286 | 2463 | TRUE | 3 |

| | | | | | |
|---|---|---|---|---|---|
| orf1a | 3252 | 7286 | 2463 | TRUE | 3 |
| orf1a | 3253 | 7286 | 2463 | TRUE | 5 |
| orf1a | 3254 | 7286 | 2463 | TRUE | 7 |
| orf1a | 3255 | 7286 | 2463 | TRUE | 5 |
| orf1a | 3256 | 7286 | 2463 | TRUE | 1 |
| orf1a | 3257 | 13476 | 0 | TRUE | 2 |
| orf1a | 3258 | 13476 | 0 | FALSE | 4 |
| orf1a | 3259 | 13476 | 0 | FALSE | 3 |
| orf1a | 3260 | 13476 | 0 | FALSE | 4 |
| orf1a | 3274 | 0 | 4721 | TRUE | 5 |
| orf1a | 3275 | 0 | 4721 | FALSE | 1 |
| orf1a | 3276 | 0 | 2935 | TRUE | 2 |
| orf1a | 3277 | 0 | 2935 | TRUE | 2 |
| orf1a | 3278 | 0 | 2573 | FALSE | 4 |
| orf1a | 3279 | 0 | 2573 | TRUE | 2 |
| orf1a | 3280 | 0 | 2573 | TRUE | 5 |
| orf1a | 3281 | 0 | 2573 | TRUE | 3 |
| orf1a | 3282 | 0 | 2573 | FALSE | 2 |
| orf1a | 3283 | 0 | 2573 | FALSE | 3 |
| orf1a | 3284 | 0 | 2573 | FALSE | 3 |
| orf1a | 3285 | 0 | 2573 | FALSE | 2 |
| orf1a | 3286 | 0 | 2573 | TRUE | 6 |
| orf1a | 3287 | 0 | 2573 | TRUE | 3 |
| orf1a | 3288 | 0 | 2573 | TRUE | 2 |
| orf1a | 3289 | 0 | 1500 | TRUE | 2 |
| orf1a | 3290 | 0 | 4161 | TRUE | 3 |
| orf1a | 3291 | 0 | 5668 | TRUE | 2 |
| orf1a | 3292 | 0 | 6761 | TRUE | 2 |
| orf1a | 3293 | 0 | 9216 | TRUE | 1 |
| orf1a | 3294 | 0 | 9216 | TRUE | 5 |
| orf1a | 3295 | 0 | 9216 | TRUE | 2 |
| orf1a | 3296 | 1476 | 9216 | FALSE | 4 |
| orf1a | 3297 | 1476 | 9216 | TRUE | 3 |
| orf1a | 3298 | 1476 | 9216 | FALSE | 2 |
| orf1a | 3299 | 1476 | 9216 | TRUE | 3 |
| orf1a | 3300 | 1476 | 9216 | TRUE | 1 |
| orf1a | 3301 | 1476 | 9216 | TRUE | 2 |
| orf1a | 3302 | 1476 | 9216 | TRUE | 4 |
| orf1a | 3303 | 1476 | 9216 | TRUE | 2 |
| orf1a | 3304 | 1532 | 5156 | TRUE | 3 |
| orf1a | 3305 | 1532 | 3713 | TRUE | 5 |
| orf1a | 3306 | 1532 | 3713 | TRUE | 5 |
| orf1a | 3307 | 1532 | 1097 | TRUE | 2 |
| orf1a | 3308 | 6635 | 1066 | TRUE | 3 |
| orf1a | 3309 | 6635 | 1066 | TRUE | 3 |
| orf1a | 3310 | 11221 | 2713 | FALSE | 3 |
| orf1a | 3311 | 12845 | 2713 | TRUE | 2 |
| orf1a | 3312 | 12845 | 2713 | FALSE | 3 |
| orf1a | 3313 | 12845 | 2713 | FALSE | 4 |
| orf1a | 3314 | 10318 | 2713 | FALSE | 2 |
| orf1a | 3315 | 10318 | 2713 | TRUE | 3 |
| orf1a | 3316 | 9503 | 2713 | TRUE | 2 |
| orf1a | 3317 | 9503 | 2713 | TRUE | 1 |
| orf1a | 3318 | 11092 | 3245 | FALSE | 1 |
| orf1a | 3319 | 12042 | 3859 | FALSE | 1 |
| orf1a | 3320 | 12042 | 3859 | FALSE | 2 |
| orf1a | 3321 | 12042 | 3859 | FALSE | 2 |
| orf1a | 3322 | 12042 | 3859 | FALSE | 3 |
| orf1a | 3323 | 12118 | 4335 | FALSE | 6 |
| orf1a | 3324 | 12118 | 4335 | FALSE | 0 |
| orf1a | 3325 | 8988 | 4310 | FALSE | 2 |
| orf1a | 3326 | 7837 | 4310 | FALSE | 3 |

| | | | | | |
|---|---|---|---|---|---|
| orf1a | 3327 | 7837 | 4310 | FALSE | 4 |
| orf1a | 3328 | 6906 | 4310 | FALSE | 3 |
| orf1a | 3329 | 8573 | 4310 | FALSE | 1 |
| orf1a | 3330 | 8573 | 4310 | FALSE | 2 |
| orf1a | 3331 | 9971 | 4310 | FALSE | 2 |
| orf1a | 3332 | 9971 | 4310 | FALSE | 4 |
| orf1a | 3333 | 1703 | 0 | FALSE | 3 |
| orf1a | 3334 | 1703 | 0 | FALSE | 4 |
| orf1a | 3335 | 1703 | 0 | FALSE | 1 |
| orf1a | 3336 | 1703 | 0 | FALSE | 2 |
| orf1a | 3337 | 1703 | 0 | FALSE | 0 |
| orf1a | 3338 | 1703 | 0 | FALSE | 3 |
| orf1a | 3339 | 1703 | 0 | FALSE | 5 |
| orf1a | 3340 | 1703 | 1063 | FALSE | 2 |
| orf1a | 3341 | 1324 | 1063 | FALSE | 2 |
| orf1a | 3342 | 1245 | 1063 | FALSE | 1 |
| orf1a | 3343 | 1167 | 1063 | FALSE | 2 |
| orf1a | 3344 | 1255 | 1063 | FALSE | 2 |
| orf1a | 3345 | 1255 | 1063 | FALSE | 4 |
| orf1a | 3346 | 1255 | 1063 | FALSE | 2 |
| orf1a | 3347 | 1255 | 1063 | FALSE | 2 |
| orf1a | 3348 | 1255 | 1063 | FALSE | 2 |
| orf1a | 3349 | 1255 | 1063 | FALSE | 3 |
| orf1a | 3350 | 1255 | 1063 | FALSE | 2 |
| orf1a | 3351 | 1255 | 1063 | FALSE | 5 |
| orf1a | 3352 | 1255 | 1063 | FALSE | 2 |
| orf1a | 3353 | 1255 | 1063 | FALSE | 5 |
| orf1a | 3354 | 1255 | 1063 | FALSE | 2 |
| orf1a | 3355 | 1255 | 0 | FALSE | 1 |
| orf1a | 3356 | 1255 | 0 | FALSE | 5 |
| orf1a | 3357 | 1342 | 0 | FALSE | 4 |
| orf1a | 3358 | 1342 | 0 | FALSE | 1 |
| orf1a | 3410 | 0 | 4694 | TRUE | 4 |
| orf1a | 3411 | 0 | 4694 | TRUE | 2 |
| orf1a | 3412 | 0 | 5081 | TRUE | 4 |
| orf1a | 3413 | 0 | 5081 | TRUE | 3 |
| orf1a | 3414 | 3051 | 5081 | TRUE | 4 |
| orf1a | 3415 | 3051 | 5081 | TRUE | 2 |
| orf1a | 3416 | 2374 | 5081 | FALSE | 2 |
| orf1a | 3417 | 2374 | 5081 | FALSE | 1 |
| orf1a | 3418 | 2374 | 5081 | FALSE | 1 |
| orf1a | 3419 | 2374 | 5081 | FALSE | 1 |
| orf1a | 3420 | 2374 | 3952 | TRUE | 4 |
| orf1a | 3421 | 2374 | 3952 | FALSE | 3 |
| orf1a | 3422 | 2402 | 3952 | TRUE | 3 |
| orf1a | 3423 | 2402 | 3952 | TRUE | 4 |
| orf1a | 3424 | 2175 | 3952 | TRUE | 3 |
| orf1a | 3425 | 2175 | 3580 | TRUE | 2 |
| orf1a | 3426 | 2621 | 2867 | TRUE | 5 |
| orf1a | 3427 | 2621 | 1566 | TRUE | 4 |
| orf1a | 3428 | 2621 | 1566 | TRUE | 6 |
| orf1a | 3429 | 2406 | 1566 | TRUE | 3 |
| orf1a | 3430 | 2406 | 1566 | TRUE | 2 |
| orf1a | 3431 | 2868 | 1566 | TRUE | 2 |
| orf1a | 3432 | 2561 | 1499 | TRUE | 3 |
| orf1a | 3433 | 2561 | 1499 | TRUE | 3 |
| orf1a | 3434 | 2561 | 1499 | TRUE | 2 |
| orf1a | 3435 | 2561 | 1402 | TRUE | 2 |
| orf1a | 3436 | 2561 | 1402 | TRUE | 4 |
| orf1a | 3437 | 2586 | 1402 | TRUE | 5 |
| orf1a | 3438 | 5874 | 1402 | TRUE | 3 |
| orf1a | 3439 | 7025 | 1402 | TRUE | 4 |

| | | | | | |
|---|---|---|---|---|---|
| orf1a | 3440 | 7025 | 1402 | TRUE | 3 |
| orf1a | 3441 | 7751 | 1365 | TRUE | 3 |
| orf1a | 3442 | 7751 | 1365 | TRUE | 4 |
| orf1a | 3443 | 7751 | 1365 | FALSE | 2 |
| orf1a | 3444 | 6076 | 1365 | TRUE | 1 |
| orf1a | 3445 | 6076 | 1365 | TRUE | 3 |
| orf1a | 3446 | 5172 | 1486 | TRUE | 2 |
| orf1a | 3447 | 6055 | 1606 | TRUE | 3 |
| orf1a | 3448 | 6055 | 1606 | TRUE | 2 |
| orf1a | 3449 | 6055 | 1606 | FALSE | 3 |
| orf1a | 3450 | 6055 | 1606 | TRUE | 4 |
| orf1a | 3451 | 6055 | 1606 | FALSE | 4 |
| orf1a | 3452 | 6055 | 1606 | FALSE | 2 |
| orf1a | 3453 | 1304 | 1606 | FALSE | 2 |
| orf1a | 3454 | 1304 | 1606 | FALSE | 5 |
| orf1a | 3455 | 1304 | 1606 | FALSE | 3 |
| orf1a | 3456 | 1304 | 1606 | FALSE | 4 |
| orf1a | 3457 | 1304 | 1606 | FALSE | 4 |
| orf1a | 3458 | 1304 | 10329 | FALSE | 2 |
| orf1a | 3459 | 1558 | 10329 | FALSE | 4 |
| orf1a | 3460 | 1558 | 10329 | FALSE | 3 |
| orf1a | 3461 | 0 | 19053 | FALSE | 1 |
| orf1a | 3462 | 0 | 19053 | FALSE | 4 |
| orf1a | 3463 | 0 | 19053 | FALSE | 3 |
| orf1a | 3464 | 0 | 19053 | FALSE | 1 |
| orf1a | 3465 | 0 | 19053 | FALSE | 4 |
| orf1a | 3466 | 0 | 19053 | FALSE | 3 |
| orf1a | 3467 | 0 | 19053 | FALSE | 2 |
| orf1a | 3468 | 0 | 19053 | FALSE | 0 |
| orf1a | 3469 | 0 | 19053 | FALSE | 2 |
| orf1a | 3470 | 0 | 19053 | FALSE | 1 |
| orf1a | 3471 | 0 | 19053 | FALSE | 2 |
| orf1a | 3472 | 0 | 19053 | FALSE | 2 |
| orf1a | 3484 | 8911 | 0 | FALSE | 3 |
| orf1a | 3485 | 8911 | 0 | FALSE | 5 |
| orf1a | 3486 | 8911 | 0 | FALSE | 3 |
| orf1a | 3487 | 8911 | 0 | FALSE | 3 |
| orf1a | 3488 | 8911 | 0 | FALSE | 4 |
| orf1a | 3489 | 8911 | 0 | FALSE | 2 |
| orf1a | 3490 | 8911 | 1346 | FALSE | 4 |
| orf1a | 3491 | 8911 | 1346 | FALSE | 1 |
| orf1a | 3492 | 8911 | 1346 | FALSE | 3 |
| orf1a | 3493 | 8911 | 1346 | FALSE | 1 |
| orf1a | 3494 | 12066 | 1346 | FALSE | 2 |
| orf1a | 3495 | 12066 | 1346 | FALSE | 1 |
| orf1a | 3496 | 8847 | 1346 | FALSE | 3 |
| orf1a | 3497 | 8847 | 1346 | FALSE | 4 |
| orf1a | 3498 | 7655 | 1346 | FALSE | 4 |
| orf1a | 3499 | 7236 | 1346 | FALSE | 2 |
| orf1a | 3500 | 7236 | 1346 | FALSE | 3 |
| orf1a | 3501 | 7236 | 1346 | FALSE | 2 |
| orf1a | 3502 | 5839 | 1346 | FALSE | 2 |
| orf1a | 3503 | 5839 | 1346 | FALSE | 3 |
| orf1a | 3504 | 3799 | 1346 | FALSE | 5 |
| orf1a | 3505 | 3799 | 0 | FALSE | 2 |
| orf1a | 3506 | 3799 | 0 | FALSE | 2 |
| orf1a | 3507 | 3799 | 0 | FALSE | 4 |
| orf1a | 3508 | 3690 | 1089 | FALSE | 4 |
| orf1a | 3509 | 2249 | 1089 | FALSE | 3 |
| orf1a | 3510 | 2249 | 1089 | FALSE | 5 |
| orf1a | 3511 | 2226 | 1089 | FALSE | 4 |
| orf1a | 3512 | 2226 | 1089 | FALSE | 1 |

| | | | | | |
|---|---|---|---|---|---|
| orf1a | 3513 | 1917 | 1089 | FALSE | 2 |
| orf1a | 3514 | 2154 | 1089 | FALSE | 3 |
| orf1a | 3515 | 2154 | 1089 | FALSE | 3 |
| orf1a | 3516 | 2154 | 1089 | FALSE | 3 |
| orf1a | 3517 | 2218 | 1089 | FALSE | 5 |
| orf1a | 3518 | 2218 | 1089 | FALSE | 3 |
| orf1a | 3519 | 2677 | 1089 | FALSE | 3 |
| orf1a | 3520 | 2677 | 1089 | FALSE | 2 |
| orf1a | 3521 | 2677 | 1089 | FALSE | 1 |
| orf1a | 3522 | 2677 | 1089 | FALSE | 2 |
| orf1a | 3523 | 2628 | 0 | FALSE | 4 |
| orf1a | 3524 | 2628 | 0 | FALSE | 7 |
| orf1a | 3525 | 2628 | 0 | FALSE | 0 |
| orf1a | 3526 | 2628 | 0 | FALSE | 5 |
| orf1a | 3527 | 2628 | 0 | FALSE | 5 |
| orf1a | 3528 | 2628 | 0 | FALSE | 3 |
| orf1a | 3540 | 1890 | 0 | FALSE | 4 |
| orf1a | 3541 | 1890 | 0 | FALSE | 5 |
| orf1a | 3542 | 1890 | 0 | FALSE | 5 |
| orf1a | 3543 | 1890 | 0 | FALSE | 4 |
| orf1a | 3544 | 1890 | 0 | FALSE | 2 |
| orf1a | 3545 | 1890 | 0 | FALSE | 3 |
| orf1a | 3546 | 1458 | 0 | FALSE | 3 |
| orf1a | 3547 | 1458 | 0 | FALSE | 2 |
| orf1a | 3548 | 1651 | 3395 | FALSE | 4 |
| orf1a | 3549 | 1651 | 3395 | FALSE | 1 |
| orf1a | 3550 | 1571 | 3395 | FALSE | 2 |
| orf1a | 3551 | 1571 | 3395 | FALSE | 5 |
| orf1a | 3552 | 1571 | 2965 | FALSE | 3 |
| orf1a | 3553 | 1571 | 2965 | FALSE | 1 |
| orf1a | 3554 | 1571 | 5209 | FALSE | 1 |
| orf1a | 3555 | 1465 | 5209 | FALSE | 2 |
| orf1a | 3556 | 1465 | 5209 | FALSE | 3 |
| orf1a | 3557 | 1465 | 5209 | FALSE | 2 |
| orf1a | 3558 | 1465 | 5209 | FALSE | 3 |
| orf1a | 3559 | 1465 | 5209 | FALSE | 4 |
| orf1a | 3560 | 1465 | 5209 | FALSE | 3 |
| orf1a | 3561 | 1684 | 5209 | FALSE | 3 |
| orf1a | 3562 | 1684 | 5209 | FALSE | 1 |
| orf1a | 3563 | 1013 | 8837 | FALSE | 1 |
| orf1a | 3564 | 1013 | 8837 | FALSE | 2 |
| orf1a | 3565 | 0 | 8837 | FALSE | 3 |
| orf1a | 3566 | 0 | 8837 | FALSE | 1 |
| orf1a | 3567 | 0 | 16432 | FALSE | 3 |
| orf1a | 3568 | 0 | 16432 | FALSE | 4 |
| orf1a | 3590 | 1121 | 0 | FALSE | 2 |
| orf1a | 3591 | 1121 | 0 | FALSE | 3 |
| orf1a | 3592 | 1121 | 0 | FALSE | 4 |
| orf1a | 3593 | 1121 | 0 | FALSE | 4 |
| orf1a | 3594 | 1121 | 0 | FALSE | 2 |
| orf1a | 3595 | 1121 | 0 | FALSE | 4 |
| orf1a | 3596 | 1121 | 0 | FALSE | 3 |
| orf1a | 3597 | 1121 | 0 | FALSE | 6 |
| orf1a | 3598 | 1121 | 0 | FALSE | 2 |
| orf1a | 3599 | 1121 | 0 | FALSE | 5 |
| orf1a | 3600 | 1121 | 0 | FALSE | 5 |
| orf1a | 3601 | 1121 | 0 | FALSE | 3 |
| orf1a | 3602 | 1121 | 0 | FALSE | 4 |
| orf1a | 3603 | 1121 | 0 | FALSE | 2 |
| orf1a | 3604 | 1121 | 1575 | FALSE | 3 |
| orf1a | 3605 | 0 | 1575 | FALSE | 1 |
| orf1a | 3606 | 0 | 1340 | FALSE | 3 |

| | | | | | |
|---|---|---|---|---|---|
| orf1a | 3607 | 0 | 1340 | FALSE | 3 |
| orf1a | 3608 | 10998 | 1376 | FALSE | 3 |
| orf1a | 3609 | 10998 | 1376 | FALSE | 0 |
| orf1a | 3610 | 10998 | 1376 | FALSE | 5 |
| orf1a | 3611 | 10998 | 1376 | FALSE | 4 |
| orf1a | 3612 | 10998 | 1376 | FALSE | 1 |
| orf1a | 3613 | 10998 | 1376 | FALSE | 3 |
| orf1a | 3614 | 10998 | 1376 | FALSE | 2 |
| orf1a | 3615 | 10998 | 1376 | FALSE | 5 |
| orf1a | 3616 | 10998 | 1376 | FALSE | 2 |
| orf1a | 3617 | 10998 | 1376 | FALSE | 3 |
| orf1a | 3618 | 10998 | 1376 | FALSE | 2 |
| orf1a | 3619 | 10998 | 1310 | FALSE | 4 |
| orf1a | 3620 | 10998 | 1310 | FALSE | 4 |
| orf1a | 3621 | 10998 | 1413 | FALSE | 3 |
| orf1a | 3622 | 10998 | 1413 | FALSE | 4 |
| orf1a | 3732 | 1917 | 0 | FALSE | 2 |
| orf1a | 3733 | 1917 | 0 | FALSE | 5 |
| orf1a | 3734 | 1917 | 3064 | FALSE | 3 |
| orf1a | 3735 | 1917 | 3064 | FALSE | 2 |
| orf1a | 3736 | 1917 | 3064 | FALSE | 3 |
| orf1a | 3737 | 1917 | 3064 | FALSE | 2 |
| orf1a | 3738 | 1917 | 3064 | FALSE | 2 |
| orf1a | 3739 | 1917 | 3064 | FALSE | 3 |
| orf1a | 3740 | 1917 | 3064 | FALSE | 4 |
| orf1a | 3741 | 1917 | 3064 | FALSE | 3 |
| orf1a | 3742 | 1917 | 3064 | FALSE | 1 |
| orf1a | 3743 | 1917 | 3064 | FALSE | 1 |
| orf1a | 3744 | 1917 | 2792 | FALSE | 2 |
| orf1a | 3745 | 1917 | 2792 | FALSE | 2 |
| orf1a | 3746 | 1917 | 7440 | FALSE | 4 |
| orf1a | 3747 | 0 | 7440 | FALSE | 2 |
| orf1a | 3748 | 2116 | 7440 | FALSE | 3 |
| orf1a | 3749 | 2116 | 9628 | FALSE | 1 |
| orf1a | 3750 | 2116 | 9628 | FALSE | 3 |
| orf1a | 3751 | 2116 | 9628 | FALSE | 5 |
| orf1a | 3752 | 2116 | 9628 | FALSE | 2 |
| orf1a | 3753 | 2116 | 9628 | FALSE | 3 |
| orf1a | 3754 | 2116 | 9628 | FALSE | 2 |
| orf1a | 3755 | 2116 | 9628 | FALSE | 3 |
| orf1a | 3756 | 2116 | 9628 | FALSE | 2 |
| orf1a | 3757 | 2116 | 9628 | FALSE | 6 |
| orf1a | 3758 | 2116 | 9628 | FALSE | 5 |
| orf1a | 3759 | 2116 | 16736 | FALSE | 5 |
| orf1a | 3760 | 2116 | 16736 | FALSE | 2 |
| orf1a | 3761 | 2116 | 0 | FALSE | 4 |
| orf1a | 3762 | 7287 | 0 | FALSE | 3 |
| orf1a | 3763 | 12457 | 0 | FALSE | 3 |
| orf1a | 3764 | 13249 | 3154 | FALSE | 3 |
| orf1a | 3765 | 13249 | 3154 | FALSE | 1 |
| orf1a | 3766 | 13249 | 3154 | FALSE | 4 |
| orf1a | 3767 | 13249 | 3154 | FALSE | 3 |
| orf1a | 3768 | 13249 | 3154 | FALSE | 3 |
| orf1a | 3769 | 13249 | 3154 | FALSE | 2 |
| orf1a | 3770 | 13249 | 3154 | FALSE | 1 |
| orf1a | 3771 | 13249 | 3154 | FALSE | 1 |
| orf1a | 3772 | 13249 | 3154 | FALSE | 2 |
| orf1a | 3773 | 13249 | 3154 | FALSE | 1 |
| orf1a | 3774 | 13249 | 3154 | FALSE | 3 |
| orf1a | 3775 | 13249 | 3154 | FALSE | 4 |
| orf1a | 3776 | 13249 | 3154 | FALSE | 3 |
| orf1a | 3777 | 14040 | 3154 | FALSE | 6 |

| | | | | | |
|---|---|---|---|---|---|
| orf1a | 3778 | 14040 | 3154 | FALSE | 2 |
| orf1a | 3812 | 2065 | 1339 | TRUE | 4 |
| orf1a | 3813 | 2065 | 1339 | TRUE | 4 |
| orf1a | 3814 | 2065 | 1339 | FALSE | 4 |
| orf1a | 3815 | 2065 | 1339 | TRUE | 2 |
| orf1a | 3816 | 2065 | 1339 | TRUE | 3 |
| orf1a | 3817 | 2065 | 1339 | FALSE | 2 |
| orf1a | 3818 | 2065 | 1339 | TRUE | 2 |
| orf1a | 3819 | 2065 | 1339 | TRUE | 3 |
| orf1a | 3820 | 2065 | 1339 | TRUE | 1 |
| orf1a | 3821 | 2065 | 1339 | TRUE | 3 |
| orf1a | 3822 | 1978 | 1339 | TRUE | 2 |
| orf1a | 3823 | 1978 | 1339 | TRUE | 5 |
| orf1a | 3824 | 1738 | 1339 | TRUE | 5 |
| orf1a | 3825 | 1738 | 1339 | TRUE | 2 |
| orf1a | 3826 | 1738 | 1339 | TRUE | 3 |
| orf1a | 3827 | 1575 | 0 | TRUE | 4 |
| orf1a | 3828 | 1575 | 0 | TRUE | 3 |
| orf1a | 3829 | 1575 | 0 | FALSE | 4 |
| orf1a | 3830 | 1575 | 0 | TRUE | 6 |
| orf1a | 3831 | 1575 | 0 | TRUE | 6 |
| orf1a | 3832 | 1575 | 0 | FALSE | 7 |
| orf1a | 3833 | 1575 | 0 | TRUE | 1 |
| orf1a | 3834 | 1575 | 0 | FALSE | 3 |
| orf1a | 3835 | 1575 | 0 | FALSE | 1 |
| orf1a | 3836 | 1575 | 0 | FALSE | 2 |
| orf1a | 3837 | 1258 | 0 | FALSE | 3 |
| orf1a | 3838 | 1258 | 0 | FALSE | 1 |
| orf1a | 3844 | 0 | 3866 | TRUE | 2 |
| orf1a | 3845 | 0 | 3866 | TRUE | 3 |
| orf1a | 3846 | 0 | 4039 | TRUE | 2 |
| orf1a | 3847 | 0 | 4039 | TRUE | 1 |
| orf1a | 3848 | 0 | 4039 | TRUE | 1 |
| orf1a | 3849 | 0 | 4039 | TRUE | 4 |
| orf1a | 3850 | 0 | 4039 | FALSE | 1 |
| orf1a | 3851 | 0 | 4039 | FALSE | 3 |
| orf1a | 3852 | 0 | 4039 | TRUE | 6 |
| orf1a | 3853 | 0 | 4039 | TRUE | 3 |
| orf1a | 3854 | 0 | 4039 | TRUE | 2 |
| orf1a | 3855 | 0 | 4039 | TRUE | 1 |
| orf1a | 3856 | 0 | 4039 | TRUE | 6 |
| orf1a | 3857 | 0 | 4039 | TRUE | 3 |
| orf1a | 3858 | 0 | 4039 | TRUE | 2 |
| orf1a | 3859 | 0 | 4213 | TRUE | 3 |
| orf1a | 3860 | 0 | 3934 | TRUE | 4 |
| orf1a | 3861 | 0 | 3842 | TRUE | 1 |
| orf1a | 3862 | 0 | 3842 | TRUE | 2 |
| orf1a | 3863 | 0 | 3842 | TRUE | 3 |
| orf1a | 3864 | 0 | 3842 | TRUE | 2 |
| orf1a | 3865 | 0 | 3842 | TRUE | 1 |
| orf1a | 3866 | 0 | 3842 | TRUE | 3 |
| orf1a | 3867 | 0 | 3842 | TRUE | 6 |
| orf1a | 3868 | 0 | 3842 | TRUE | 3 |
| orf1a | 3869 | 0 | 3842 | TRUE | 3 |
| orf1a | 3870 | 0 | 3842 | TRUE | 0 |
| orf1a | 3871 | 0 | 3842 | TRUE | 4 |
| orf1a | 3872 | 0 | 3842 | TRUE | 0 |
| orf1a | 3873 | 0 | 3842 | TRUE | 2 |
| orf1a | 3874 | 0 | 3842 | TRUE | 2 |
| orf1a | 3880 | 0 | 2627 | FALSE | 3 |
| orf1a | 3881 | 0 | 2627 | FALSE | 3 |
| orf1a | 3882 | 9588 | 4736 | FALSE | 4 |

| | | | | | |
|---|---|---|---|---|---|
| orf1a | 3883 | 9588 | 4736 | TRUE | 3 |
| orf1a | 3884 | 7944 | 7430 | TRUE | 2 |
| orf1a | 3885 | 7944 | 7430 | TRUE | 2 |
| orf1a | 3886 | 6915 | 7430 | FALSE | 0 |
| orf1a | 3887 | 6915 | 7430 | FALSE | 6 |
| orf1a | 3888 | 6915 | 7430 | TRUE | 4 |
| orf1a | 3889 | 6915 | 7430 | FALSE | 3 |
| orf1a | 3890 | 6915 | 7430 | TRUE | 2 |
| orf1a | 3891 | 6915 | 7430 | TRUE | 3 |
| orf1a | 3892 | 6915 | 6533 | TRUE | 5 |
| orf1a | 3893 | 6915 | 6533 | FALSE | 2 |
| orf1a | 3894 | 6102 | 6533 | TRUE | 2 |
| orf1a | 3895 | 6102 | 8096 | TRUE | 3 |
| orf1a | 3896 | 5656 | 8096 | TRUE | 0 |
| orf1a | 3897 | 2708 | 11027 | TRUE | 2 |
| orf1a | 3898 | 2423 | 7788 | TRUE | 3 |
| orf1a | 3899 | 2276 | 1230 | TRUE | 2 |
| orf1a | 3900 | 2090 | 1230 | FALSE | 4 |
| orf1a | 3901 | 1912 | 1230 | FALSE | 3 |
| orf1a | 3902 | 1915 | 1230 | FALSE | 3 |
| orf1a | 3903 | 1915 | 1230 | FALSE | 5 |
| orf1a | 3904 | 2828 | 1230 | FALSE | 3 |
| orf1a | 3905 | 2828 | 1230 | FALSE | 2 |
| orf1a | 3906 | 4694 | 6872 | FALSE | 4 |
| orf1a | 3907 | 4694 | 8779 | FALSE | 3 |
| orf1a | 3908 | 4694 | 8779 | FALSE | 3 |
| orf1a | 3909 | 4989 | 8779 | FALSE | 4 |
| orf1a | 3910 | 4989 | 8779 | FALSE | 1 |
| orf1a | 3911 | 5240 | 8779 | FALSE | 3 |
| orf1a | 3912 | 5240 | 8779 | FALSE | 3 |
| orf1a | 3913 | 5805 | 12513 | FALSE | 1 |
| orf1a | 3914 | 5805 | 12513 | FALSE | 2 |
| orf1a | 3915 | 6548 | 12513 | FALSE | 5 |
| orf1a | 3916 | 6548 | 12513 | FALSE | 2 |
| orf1a | 3917 | 8862 | 12513 | FALSE | 3 |
| orf1a | 3918 | 8862 | 12513 | FALSE | 2 |
| orf1a | 3919 | 12157 | 12513 | FALSE | 3 |
| orf1a | 3920 | 6911 | 12513 | FALSE | 2 |
| orf1a | 3921 | 1665 | 0 | FALSE | 5 |
| orf1a | 3922 | 2063 | 0 | FALSE | 3 |
| orf1a | 3923 | 2063 | 0 | FALSE | 4 |
| orf1a | 3924 | 2434 | 1754 | FALSE | 3 |
| orf1a | 3925 | 2434 | 1754 | FALSE | 3 |
| orf1a | 3926 | 2520 | 1754 | FALSE | 5 |
| orf1a | 3927 | 2520 | 1754 | FALSE | 1 |
| orf1a | 3928 | 4287 | 1754 | FALSE | 4 |
| orf1a | 3929 | 4287 | 1754 | FALSE | 1 |
| orf1a | 3930 | 5108 | 1447 | FALSE | 4 |
| orf1a | 3931 | 5108 | 1447 | FALSE | 4 |
| orf1a | 3932 | 8399 | 8250 | FALSE | 2 |
| orf1a | 3933 | 8399 | 8250 | FALSE | 2 |
| orf1a | 3934 | 8399 | 8250 | FALSE | 5 |
| orf1a | 3935 | 9435 | 8250 | FALSE | 3 |
| orf1a | 3936 | 9435 | 8250 | FALSE | 3 |
| orf1a | 3937 | 10703 | 8250 | FALSE | 3 |
| orf1a | 3938 | 10703 | 8250 | FALSE | 5 |
| orf1a | 3939 | 11382 | 9550 | FALSE | 4 |
| orf1a | 3940 | 11382 | 9550 | FALSE | 2 |
| orf1a | 3941 | 12318 | 9550 | FALSE | 3 |
| orf1a | 3942 | 12318 | 9550 | FALSE | 2 |
| orf1a | 3943 | 13099 | 9550 | FALSE | 3 |
| orf1a | 3944 | 13099 | 9550 | FALSE | 4 |

| | | | | | |
|---|---|---|---|---|---|
| orf1a | 3945 | 13336 | 11652 | FALSE | 5 |
| orf1a | 3946 | 12006 | 11652 | FALSE | 3 |
| orf1a | 3947 | 4028 | 0 | FALSE | 3 |
| orf1a | 3948 | 4028 | 0 | FALSE | 1 |
| orf1a | 3949 | 4028 | 0 | FALSE | 4 |
| orf1a | 3950 | 4028 | 0 | FALSE | 3 |
| orf1a | 3951 | 4028 | 0 | FALSE | 2 |
| orf1a | 3952 | 2767 | 0 | FALSE | 3 |
| orf1a | 3953 | 2767 | 0 | FALSE | 2 |
| orf1a | 3954 | 2767 | 0 | FALSE | 0 |
| orf1a | 3955 | 2767 | 0 | FALSE | 2 |
| orf1a | 3956 | 2767 | 0 | FALSE | 5 |
| orf1a | 3957 | 2767 | 0 | FALSE | 2 |
| orf1a | 3958 | 2767 | 0 | FALSE | 1 |
| orf1a | 3959 | 2767 | 0 | FALSE | 2 |
| orf1a | 3960 | 7511 | 1228 | FALSE | 2 |
| orf1a | 3961 | 9253 | 1228 | FALSE | 3 |
| orf1a | 3962 | 12537 | 8489 | FALSE | 2 |
| orf1a | 3963 | 12537 | 8489 | FALSE | 3 |
| orf1a | 3964 | 12537 | 7420 | FALSE | 2 |
| orf1a | 3965 | 12537 | 7420 | FALSE | 2 |
| orf1a | 3966 | 12537 | 6592 | FALSE | 4 |
| orf1a | 3967 | 15294 | 6592 | FALSE | 3 |
| orf1a | 3968 | 15294 | 6592 | FALSE | 2 |
| orf1a | 3969 | 15294 | 6592 | FALSE | 5 |
| orf1a | 3970 | 12505 | 6592 | FALSE | 1 |
| orf1a | 3971 | 12505 | 6592 | FALSE | 5 |
| orf1a | 3972 | 11094 | 6143 | FALSE | 2 |
| orf1a | 3973 | 11094 | 6143 | FALSE | 3 |
| orf1a | 3974 | 10576 | 5351 | FALSE | 4 |
| orf1a | 3975 | 9505 | 6381 | FALSE | 1 |
| orf1a | 3976 | 9505 | 6381 | FALSE | 3 |
| orf1a | 3977 | 4285 | 2213 | FALSE | 1 |
| orf1a | 3978 | 4285 | 2213 | FALSE | 1 |
| orf1a | 3979 | 4285 | 2247 | FALSE | 3 |
| orf1a | 3980 | 4285 | 2247 | TRUE | 3 |
| orf1a | 3981 | 4285 | 2454 | TRUE | 2 |
| orf1a | 3982 | 4285 | 2454 | TRUE | 3 |
| orf1a | 3983 | 4285 | 2454 | TRUE | 3 |
| orf1a | 3984 | 4285 | 2454 | TRUE | 1 |
| orf1a | 3985 | 5755 | 2454 | TRUE | 3 |
| orf1a | 3986 | 5755 | 2454 | TRUE | 1 |
| orf1a | 3987 | 7467 | 2180 | TRUE | 2 |
| orf1a | 3988 | 7467 | 1927 | TRUE | 1 |
| orf1a | 3989 | 0 | 1420 | TRUE | 5 |
| orf1a | 3990 | 6112 | 3080 | FALSE | 3 |
| orf1a | 3991 | 6112 | 3080 | TRUE | 0 |
| orf1a | 3992 | 6112 | 3080 | TRUE | 3 |
| orf1a | 3993 | 6112 | 3080 | TRUE | 5 |
| orf1a | 3994 | 6112 | 3080 | TRUE | 0 |
| orf1a | 3995 | 6112 | 3080 | FALSE | 3 |
| orf1a | 3996 | 6112 | 3080 | TRUE | 3 |
| orf1a | 3997 | 6112 | 3080 | TRUE | 3 |
| orf1a | 3998 | 6112 | 3080 | TRUE | 3 |
| orf1a | 3999 | 6112 | 3080 | TRUE | 4 |
| orf1a | 4000 | 4976 | 2362 | TRUE | 1 |
| orf1a | 4001 | 4976 | 2362 | TRUE | 5 |
| orf1a | 4002 | 5774 | 3578 | TRUE | 5 |
| orf1a | 4003 | 5774 | 3937 | TRUE | 4 |
| orf1a | 4004 | 5774 | 3937 | TRUE | 0 |
| orf1a | 4005 | 5605 | 3951 | TRUE | 3 |
| orf1a | 4006 | 5605 | 3951 | TRUE | 1 |

| | | | | | |
|---|---|---|---|---|---|
| orf1a | 4007 | 5605 | 3951 | TRUE | 1 |
| orf1a | 4008 | 5605 | 3951 | TRUE | 4 |
| orf1a | 4009 | 5605 | 3951 | TRUE | 4 |
| orf1a | 4010 | 5605 | 3951 | TRUE | 3 |
| orf1a | 4011 | 5605 | 3951 | TRUE | 3 |
| orf1a | 4012 | 5605 | 3951 | TRUE | 3 |
| orf1a | 4013 | 5605 | 3951 | TRUE | 2 |
| orf1a | 4014 | 5605 | 3951 | TRUE | 3 |
| orf1a | 4015 | 7369 | 6617 | TRUE | 3 |
| orf1a | 4016 | 7369 | 6617 | TRUE | 5 |
| orf1a | 4038 | 4346 | 0 | TRUE | 5 |
| orf1a | 4039 | 4346 | 0 | TRUE | 2 |
| orf1a | 4040 | 3688 | 1875 | TRUE | 2 |
| orf1a | 4041 | 3688 | 1875 | TRUE | 1 |
| orf1a | 4042 | 3455 | 1875 | TRUE | 1 |
| orf1a | 4043 | 3455 | 1875 | TRUE | 2 |
| orf1a | 4044 | 3455 | 1875 | TRUE | 3 |
| orf1a | 4045 | 3455 | 1875 | TRUE | 3 |
| orf1a | 4046 | 4072 | 1875 | TRUE | 2 |
| orf1a | 4047 | 4072 | 1875 | TRUE | 2 |
| orf1a | 4048 | 4072 | 1875 | TRUE | 0 |
| orf1a | 4049 | 4072 | 1875 | TRUE | 5 |
| orf1a | 4050 | 4104 | 1875 | TRUE | 3 |
| orf1a | 4051 | 4104 | 1875 | TRUE | 3 |
| orf1a | 4052 | 4104 | 1875 | TRUE | 5 |
| orf1a | 4053 | 3983 | 1875 | FALSE | 0 |
| orf1a | 4054 | 3611 | 1875 | FALSE | 1 |
| orf1a | 4055 | 3921 | 0 | TRUE | 4 |
| orf1a | 4056 | 3921 | 0 | TRUE | 1 |
| orf1a | 4057 | 4309 | 0 | TRUE | 2 |
| orf1a | 4058 | 4309 | 0 | TRUE | 4 |
| orf1a | 4059 | 4309 | 0 | TRUE | 5 |
| orf1a | 4060 | 4309 | 0 | TRUE | 2 |
| orf1a | 4061 | 3193 | 0 | TRUE | 3 |
| orf1a | 4062 | 2737 | 0 | TRUE | 3 |
| orf1a | 4063 | 2737 | 0 | TRUE | 2 |
| orf1a | 4064 | 2737 | 0 | FALSE | 3 |
| orf1a | 4065 | 1974 | 0 | FALSE | 3 |
| orf1a | 4066 | 1974 | 0 | FALSE | 1 |
| orf1a | 4067 | 1974 | 0 | FALSE | 2 |
| orf1a | 4068 | 1974 | 0 | FALSE | 4 |
| orf1a | 4069 | 1825 | 0 | FALSE | 3 |
| orf1a | 4070 | 1825 | 0 | FALSE | 4 |
| orf1a | 4071 | 1825 | 0 | FALSE | 3 |
| orf1a | 4072 | 1988 | 0 | FALSE | 2 |
| orf1a | 4073 | 1988 | 0 | FALSE | 2 |
| orf1a | 4074 | 2245 | 0 | FALSE | 1 |
| orf1a | 4075 | 2245 | 0 | FALSE | 2 |
| orf1a | 4076 | 2245 | 0 | FALSE | 2 |
| orf1a | 4077 | 2351 | 0 | FALSE | 0 |
| orf1a | 4078 | 2351 | 0 | FALSE | 1 |
| orf1a | 4079 | 2351 | 0 | FALSE | 2 |
| orf1a | 4080 | 2351 | 0 | FALSE | 2 |
| orf1a | 4081 | 2351 | 0 | FALSE | 4 |
| orf1a | 4082 | 2351 | 0 | FALSE | 2 |
| orf1a | 4083 | 2351 | 0 | FALSE | 4 |
| orf1a | 4084 | 2351 | 0 | FALSE | 2 |
| orf1a | 4085 | 2351 | 0 | FALSE | 1 |
| orf1a | 4086 | 2351 | 1652 | FALSE | 3 |
| orf1a | 4087 | 2417 | 1652 | FALSE | 2 |
| orf1a | 4088 | 2417 | 1652 | FALSE | 4 |
| orf1a | 4089 | 0 | 1652 | FALSE | 0 |

| | | | | | |
|---|---|---|---|---|---|
| orf1a | 4090 | 0 | 1652 | FALSE | 2 |
| orf1a | 4091 | 0 | 1652 | FALSE | 2 |
| orf1a | 4092 | 1389 | 1652 | FALSE | 2 |
| orf1a | 4093 | 1389 | 1652 | FALSE | 1 |
| orf1a | 4094 | 1389 | 1708 | FALSE | 3 |
| orf1a | 4095 | 1389 | 1708 | FALSE | 2 |
| orf1a | 4096 | 1389 | 1708 | FALSE | 3 |
| orf1a | 4097 | 1389 | 1708 | FALSE | 2 |
| orf1a | 4098 | 1389 | 1708 | FALSE | 1 |
| orf1a | 4099 | 1389 | 1708 | FALSE | 3 |
| orf1a | 4100 | 1635 | 1559 | FALSE | 2 |
| orf1a | 4101 | 1635 | 1513 | FALSE | 3 |
| orf1a | 4102 | 2037 | 1513 | FALSE | 4 |
| orf1a | 4103 | 2037 | 1513 | FALSE | 3 |
| orf1a | 4104 | 2037 | 1420 | FALSE | 4 |
| orf1a | 4105 | 2037 | 1420 | FALSE | 5 |
| orf1a | 4106 | 2037 | 1420 | FALSE | 3 |
| orf1a | 4107 | 2361 | 1420 | FALSE | 1 |
| orf1a | 4108 | 2361 | 1319 | FALSE | 0 |
| orf1a | 4109 | 2361 | 1171 | FALSE | 1 |
| orf1a | 4110 | 2361 | 1171 | FALSE | 2 |
| orf1a | 4111 | 2361 | 1171 | FALSE | 2 |
| orf1a | 4112 | 2543 | 1171 | FALSE | 2 |
| orf1a | 4113 | 2543 | 1171 | FALSE | 1 |
| orf1a | 4114 | 2543 | 1171 | FALSE | 0 |
| orf1a | 4115 | 2874 | 1126 | FALSE | 1 |
| orf1a | 4116 | 2874 | 1126 | FALSE | 3 |
| orf1a | 4117 | 2908 | 1126 | FALSE | 3 |
| orf1a | 4118 | 2247 | 1126 | FALSE | 1 |
| orf1a | 4119 | 2247 | 1017 | FALSE | 3 |
| orf1a | 4120 | 2247 | 1017 | FALSE | 3 |
| orf1a | 4121 | 2247 | 1017 | FALSE | 5 |
| orf1a | 4122 | 2247 | 1017 | FALSE | 0 |
| orf1a | 4123 | 2247 | 0 | FALSE | 3 |
| orf1a | 4124 | 2247 | 0 | FALSE | 2 |
| orf1a | 4125 | 2247 | 0 | FALSE | 4 |
| orf1a | 4126 | 2247 | 0 | FALSE | 1 |
| orf1a | 4127 | 1586 | 0 | FALSE | 0 |
| orf1a | 4128 | 1586 | 0 | FALSE | 5 |
| orf1a | 4129 | 1586 | 0 | FALSE | 2 |
| orf1a | 4130 | 1586 | 0 | FALSE | 4 |
| orf1a | 4131 | 1586 | 0 | FALSE | 0 |
| orf1a | 4132 | 1586 | 0 | FALSE | 5 |
| orf1a | 4134 | 1095 | 0 | FALSE | 5 |
| orf1a | 4135 | 1095 | 0 | FALSE | 2 |
| orf1a | 4136 | 1095 | 0 | FALSE | 3 |
| orf1a | 4137 | 1095 | 0 | FALSE | 4 |
| orf1a | 4138 | 1095 | 0 | FALSE | 3 |
| orf1a | 4139 | 1095 | 0 | FALSE | 0 |
| orf1a | 4140 | 1095 | 0 | FALSE | 5 |
| orf1a | 4141 | 1095 | 0 | FALSE | 2 |
| orf1a | 4142 | 1533 | 0 | FALSE | 4 |
| orf1a | 4143 | 1533 | 0 | FALSE | 2 |
| orf1a | 4144 | 1533 | 0 | FALSE | 3 |
| orf1a | 4145 | 1533 | 0 | FALSE | 0 |
| orf1a | 4146 | 1430 | 0 | FALSE | 2 |
| orf1a | 4147 | 1430 | 0 | FALSE | 4 |
| orf1a | 4148 | 1430 | 0 | FALSE | 3 |
| orf1a | 4149 | 1598 | 0 | FALSE | 2 |
| orf1a | 4150 | 1598 | 0 | FALSE | 5 |
| orf1a | 4151 | 1598 | 0 | FALSE | 3 |
| orf1a | 4152 | 1598 | 0 | FALSE | 4 |

| | | | | | |
|---|---|---|---|---|---|
| orf1a | 4153 | 1598 | 0 | FALSE | 2 |
| orf1a | 4154 | 1598 | 0 | FALSE | 1 |
| orf1a | 4155 | 1598 | 0 | FALSE | 2 |
| orf1a | 4156 | 1598 | 0 | FALSE | 3 |
| orf1a | 4157 | 1226 | 0 | FALSE | 5 |
| orf1a | 4158 | 1226 | 0 | FALSE | 3 |
| orf1a | 4159 | 1226 | 0 | FALSE | 3 |
| orf1a | 4160 | 1226 | 0 | FALSE | 1 |
| orf1a | 4164 | 7235 | 0 | FALSE | 1 |
| orf1a | 4165 | 7235 | 0 | FALSE | 5 |
| orf1a | 4166 | 7235 | 0 | FALSE | 3 |
| orf1a | 4167 | 7235 | 0 | FALSE | 0 |
| orf1a | 4168 | 7235 | 0 | FALSE | 4 |
| orf1a | 4169 | 7235 | 0 | FALSE | 2 |
| orf1a | 4170 | 7235 | 0 | FALSE | 2 |
| orf1a | 4171 | 7235 | 0 | FALSE | 3 |
| orf1a | 4172 | 7235 | 0 | FALSE | 2 |
| orf1a | 4173 | 7235 | 0 | FALSE | 3 |
| orf1a | 4174 | 7235 | 0 | FALSE | 4 |
| orf1a | 4175 | 7235 | 0 | FALSE | 4 |
| orf1a | 4176 | 7235 | 0 | FALSE | 5 |
| orf1a | 4177 | 7235 | 0 | FALSE | 5 |
| orf1a | 4178 | 7235 | 0 | FALSE | 6 |
| orf1a | 4186 | 5107 | 0 | FALSE | 3 |
| orf1a | 4187 | 5107 | 0 | FALSE | 3 |
| orf1a | 4188 | 5107 | 0 | FALSE | 2 |
| orf1a | 4189 | 5107 | 0 | FALSE | 3 |
| orf1a | 4190 | 5107 | 0 | FALSE | 4 |
| orf1a | 4191 | 5107 | 0 | FALSE | 3 |
| orf1a | 4192 | 5107 | 0 | FALSE | 2 |
| orf1a | 4193 | 5107 | 0 | FALSE | 2 |
| orf1a | 4194 | 5107 | 0 | FALSE | 5 |
| orf1a | 4195 | 5107 | 0 | FALSE | 5 |
| orf1a | 4196 | 5107 | 0 | FALSE | 3 |
| orf1a | 4197 | 5107 | 0 | FALSE | 2 |
| orf1a | 4198 | 5107 | 0 | FALSE | 3 |
| orf1a | 4199 | 5107 | 0 | FALSE | 3 |
| orf1a | 4200 | 5107 | 0 | FALSE | 1 |
| orf1a | 4202 | 1204 | 0 | TRUE | 2 |
| orf1a | 4203 | 1204 | 0 | FALSE | 3 |
| orf1a | 4204 | 1172 | 0 | TRUE | 1 |
| orf1a | 4205 | 1172 | 0 | TRUE | 3 |
| orf1a | 4206 | 1172 | 5617 | FALSE | 2 |
| orf1a | 4207 | 1172 | 5617 | FALSE | 3 |
| orf1a | 4208 | 4297 | 5790 | TRUE | 2 |
| orf1a | 4209 | 4297 | 5790 | TRUE | 4 |
| orf1a | 4210 | 4316 | 5790 | TRUE | 3 |
| orf1a | 4211 | 4316 | 5790 | TRUE | 4 |
| orf1a | 4212 | 4316 | 5790 | TRUE | 3 |
| orf1a | 4213 | 4316 | 5790 | TRUE | 2 |
| orf1a | 4214 | 4316 | 5790 | TRUE | 4 |
| orf1a | 4215 | 4316 | 5790 | TRUE | 0 |
| orf1a | 4216 | 4316 | 5790 | FALSE | 1 |
| orf1a | 4217 | 5094 | 5790 | TRUE | 2 |
| orf1a | 4218 | 5094 | 5790 | TRUE | 5 |
| orf1a | 4219 | 6412 | 5790 | TRUE | 3 |
| orf1a | 4220 | 6412 | 5790 | TRUE | 5 |
| orf1a | 4221 | 6412 | 5848 | FALSE | 2 |
| orf1a | 4222 | 6412 | 5848 | FALSE | 4 |
| orf1a | 4223 | 4391 | 0 | FALSE | 5 |
| orf1a | 4224 | 4391 | 0 | FALSE | 3 |
| orf1a | 4236 | 4031 | 0 | TRUE | 2 |

| | | | | | |
|---|---|---|---|---|---|
| orf1a | 4237 | 4031 | 0 | TRUE | 1 |
| orf1a | 4238 | 6452 | 0 | FALSE | 2 |
| orf1a | 4239 | 6452 | 0 | TRUE | 3 |
| orf1a | 4240 | 6452 | 0 | TRUE | 5 |
| orf1a | 4241 | 6452 | 0 | TRUE | 3 |
| orf1a | 4242 | 6452 | 0 | TRUE | 1 |
| orf1a | 4243 | 6452 | 0 | TRUE | 2 |
| orf1a | 4244 | 6452 | 0 | TRUE | 2 |
| orf1a | 4245 | 6452 | 0 | FALSE | 3 |
| orf1a | 4246 | 6452 | 0 | TRUE | 2 |
| orf1a | 4247 | 6452 | 0 | TRUE | 4 |
| orf1a | 4248 | 6452 | 0 | TRUE | 3 |
| orf1a | 4249 | 6452 | 0 | TRUE | 1 |
| orf1a | 4250 | 6452 | 0 | FALSE | 3 |
| orf1a | 4251 | 8873 | 0 | FALSE | 3 |
| orf1a | 4252 | 5501 | 0 | FALSE | 1 |
| orf1a | 4253 | 2128 | 0 | FALSE | 4 |
| orf1a | 4254 | 2128 | 0 | FALSE | 5 |
| orf1a | 4255 | 2128 | 0 | FALSE | 3 |
| orf1a | 4256 | 2900 | 0 | FALSE | 2 |
| orf1a | 4257 | 2900 | 0 | FALSE | 3 |
| orf1a | 4258 | 2271 | 0 | FALSE | 4 |
| orf1a | 4259 | 2271 | 0 | FALSE | 2 |
| orf1a | 4260 | 2271 | 0 | FALSE | 4 |
| orf1a | 4261 | 2271 | 0 | FALSE | 2 |
| orf1a | 4262 | 2271 | 0 | FALSE | 2 |
| orf1a | 4263 | 2271 | 0 | FALSE | 1 |
| orf1a | 4264 | 2271 | 0 | FALSE | 1 |
| orf1a | 4265 | 2271 | 0 | FALSE | 2 |
| orf1a | 4266 | 2271 | 0 | FALSE | 1 |
| orf1a | 4267 | 2343 | 0 | FALSE | 1 |
| orf1a | 4268 | 2343 | 0 | FALSE | 2 |
| orf1a | 4269 | 2343 | 0 | FALSE | 2 |
| orf1a | 4270 | 2343 | 0 | FALSE | 2 |
| orf1a | 4271 | 1014 | 0 | FALSE | 3 |
| orf1a | 4272 | 1014 | 0 | FALSE | 1 |
| orf1a | 4310 | 1374 | 0 | FALSE | 3 |
| orf1a | 4311 | 1374 | 0 | FALSE | 1 |
| orf1a | 4312 | 1374 | 1357 | TRUE | 4 |
| orf1a | 4313 | 1374 | 1357 | TRUE | 2 |
| orf1a | 4314 | 1374 | 1357 | TRUE | 6 |
| orf1a | 4315 | 1374 | 1357 | TRUE | 2 |
| orf1a | 4316 | 1374 | 1357 | TRUE | 3 |
| orf1a | 4317 | 1374 | 1357 | TRUE | 3 |
| orf1a | 4318 | 1374 | 1357 | TRUE | 3 |
| orf1a | 4319 | 1374 | 1357 | TRUE | 4 |
| orf1a | 4320 | 1374 | 1357 | TRUE | 2 |
| orf1a | 4321 | 1374 | 1357 | TRUE | 3 |
| orf1a | 4322 | 2060 | 1357 | TRUE | 1 |
| orf1a | 4323 | 2060 | 1357 | TRUE | 2 |
| orf1a | 4324 | 2060 | 1357 | TRUE | 2 |
| orf1a | 4325 | 2746 | 1357 | TRUE | 3 |
| orf1a | 4326 | 2746 | 1357 | TRUE | 2 |
| orf1a | 4327 | 2746 | 0 | TRUE | 4 |
| orf1a | 4328 | 2746 | 0 | TRUE | 4 |
| orf1a | 4329 | 2746 | 0 | TRUE | 2 |
| orf1a | 4330 | 2746 | 0 | TRUE | 5 |
| orf1a | 4331 | 2746 | 0 | TRUE | 3 |
| orf1a | 4332 | 2746 | 0 | TRUE | 5 |
| orf1a | 4333 | 2746 | 0 | TRUE | 3 |
| orf1a | 4334 | 2746 | 0 | TRUE | 2 |
| orf1a | 4335 | 2746 | 0 | FALSE | 2 |

| | | | | | |
|---|---|---|---|---|---|
| orf1a | 4336 | 2746 | 0 | TRUE | 3 |
| orf1a | 4360 | 2154 | 1671 | FALSE | 4 |
| orf1a | 4361 | 2154 | 1671 | FALSE | 3 |
| orf1a | 4362 | 2154 | 1671 | FALSE | 2 |
| orf1a | 4363 | 2154 | 1671 | FALSE | 2 |
| orf1a | 4364 | 2154 | 1671 | FALSE | 2 |
| orf1a | 4365 | 2154 | 1671 | FALSE | 3 |
| orf1a | 4366 | 2154 | 1671 | FALSE | 4 |
| orf1a | 4367 | 2154 | 1671 | FALSE | 2 |
| orf1a | 4368 | 2154 | 1671 | TRUE | 1 |
| orf1a | 4369 | 2154 | 1671 | TRUE | 2 |
| orf1a | 4370 | 2154 | 1671 | TRUE | 3 |
| orf1a | 4371 | 2154 | 1671 | TRUE | 3 |
| orf1a | 4372 | 2154 | 1671 | TRUE | 3 |
| orf1a | 4373 | 2154 | 1671 | TRUE | 2 |
| orf1a | 4374 | 2154 | 1883 | TRUE | 1 |
| orf1a | 4375 | 0 | 2308 | TRUE | 0 |
| orf1a | 4376 | 0 | 2308 | TRUE | 3 |
| orf1a | 4377 | 0 | 2308 | FALSE | 1 |
| orf1a | 4378 | 0 | 2308 | FALSE | 4 |
| orf1a | 4379 | 0 | 2308 | FALSE | 1 |
| orf1a | 4380 | 0 | 2308 | TRUE | 2 |
| orf1a | 4381 | 0 | 2308 | TRUE | 0 |
| orf1a | 4382 | 0 | 2308 | TRUE | 2 |
| orf1a | 4383 | 0 | 2308 | TRUE | 1 |
| orf1a | 4384 | 0 | 2308 | FALSE | 2 |
| orf1a | 4385 | 0 | 2308 | FALSE | 3 |
| orf1a | 4386 | 2389 | 2308 | FALSE | 3 |
| orf1a | 4387 | 2389 | 2308 | FALSE | 5 |
| orf1a | 4388 | 6421 | 3186 | FALSE | 2 |
| orf1a | 4389 | 6421 | 3626 | FALSE | 3 |
| orf1a | 4390 | 6421 | 3626 | FALSE | 1 |
| orf1a | 4391 | 6421 | 3626 | FALSE | 3 |
| orf1a | 4392 | 6421 | 3626 | FALSE | 3 |
| orf1a | 4393 | 6421 | 3626 | FALSE | 2 |
| orf1a | 4394 | 6421 | 3626 | FALSE | 5 |
| orf1a | 4395 | 6421 | 3626 | FALSE | 1 |
| orf1a | 4396 | 6421 | 3626 | FALSE | 3 |
| orf1a | 4397 | 6421 | 3626 | FALSE | 1 |
| orf1a | 4398 | 6421 | 3626 | FALSE | 3 |
| orf1a | 4399 | 6421 | 3626 | FALSE | 3 |
| orf1a | 4400 | 6421 | 3626 | FALSE | 3 |
| orf1a | 4401 | 7765 | 3626 | FALSE | 2 |
| orf1b | 15 | 1899 | 0 | FALSE | 3 |
| orf1b | 16 | 1899 | 0 | TRUE | 4 |
| orf1b | 17 | 1899 | 0 | TRUE | 2 |
| orf1b | 18 | 1899 | 0 | TRUE | 4 |
| orf1b | 19 | 4396 | 0 | FALSE | 3 |
| orf1b | 20 | 4396 | 0 | TRUE | 3 |
| orf1b | 21 | 4396 | 0 | TRUE | 1 |
| orf1b | 22 | 4396 | 0 | FALSE | 2 |
| orf1b | 23 | 4396 | 0 | TRUE | 2 |
| orf1b | 24 | 4396 | 0 | TRUE | 2 |
| orf1b | 25 | 4396 | 0 | TRUE | 5 |
| orf1b | 26 | 4396 | 0 | TRUE | 1 |
| orf1b | 27 | 4396 | 0 | TRUE | 3 |
| orf1b | 28 | 4396 | 0 | TRUE | 2 |
| orf1b | 29 | 4396 | 0 | FALSE | 1 |
| orf1b | 30 | 6894 | 0 | TRUE | 3 |
| orf1b | 31 | 6894 | 0 | TRUE | 2 |
| orf1b | 32 | 6894 | 0 | TRUE | 3 |
| orf1b | 33 | 6894 | 0 | TRUE | 3 |

| | | | | | |
|---|---|---|---|---|---|
| orf1b | 39 | 1460 | 0 | FALSE | 2 |
| orf1b | 40 | 1460 | 0 | FALSE | 3 |
| orf1b | 41 | 1401 | 1077 | FALSE | 1 |
| orf1b | 42 | 1401 | 1077 | FALSE | 2 |
| orf1b | 43 | 1624 | 1077 | FALSE | 2 |
| orf1b | 44 | 1624 | 1077 | FALSE | 1 |
| orf1b | 45 | 1531 | 1077 | FALSE | 2 |
| orf1b | 46 | 1531 | 1077 | FALSE | 2 |
| orf1b | 47 | 1479 | 1077 | FALSE | 1 |
| orf1b | 48 | 1479 | 1077 | FALSE | 2 |
| orf1b | 49 | 4473 | 6964 | FALSE | 4 |
| orf1b | 50 | 4473 | 6964 | FALSE | 4 |
| orf1b | 51 | 4347 | 5528 | FALSE | 3 |
| orf1b | 52 | 4347 | 5528 | FALSE | 1 |
| orf1b | 53 | 4347 | 5528 | FALSE | 4 |
| orf1b | 54 | 4829 | 5528 | FALSE | 6 |
| orf1b | 55 | 5099 | 5528 | FALSE | 3 |
| orf1b | 56 | 5417 | 7011 | FALSE | 4 |
| orf1b | 57 | 5417 | 7011 | FALSE | 4 |
| orf1b | 58 | 6109 | 7011 | FALSE | 5 |
| orf1b | 59 | 5702 | 7011 | FALSE | 3 |
| orf1b | 60 | 6165 | 7011 | FALSE | 4 |
| orf1b | 61 | 5845 | 7011 | FALSE | 4 |
| orf1b | 62 | 6313 | 7011 | FALSE | 3 |
| orf1b | 63 | 5947 | 7011 | FALSE | 3 |
| orf1b | 64 | 3525 | 1219 | FALSE | 5 |
| orf1b | 65 | 3934 | 1219 | FALSE | 4 |
| orf1b | 66 | 4138 | 0 | FALSE | 3 |
| orf1b | 67 | 3684 | 0 | FALSE | 2 |
| orf1b | 68 | 3684 | 0 | FALSE | 3 |
| orf1b | 69 | 3441 | 0 | FALSE | 3 |
| orf1b | 70 | 2828 | 0 | FALSE | 1 |
| orf1b | 71 | 2756 | 0 | FALSE | 3 |
| orf1b | 72 | 2756 | 0 | FALSE | 3 |
| orf1b | 73 | 2791 | 0 | TRUE | 4 |
| orf1b | 74 | 2920 | 0 | FALSE | 4 |
| orf1b | 75 | 3611 | 0 | FALSE | 2 |
| orf1b | 76 | 3718 | 0 | FALSE | 1 |
| orf1b | 77 | 3524 | 0 | FALSE | 4 |
| orf1b | 78 | 3661 | 0 | TRUE | 2 |
| orf1b | 79 | 4190 | 0 | TRUE | 2 |
| orf1b | 80 | 3999 | 0 | TRUE | 2 |
| orf1b | 81 | 3999 | 0 | TRUE | 4 |
| orf1b | 82 | 4558 | 0 | TRUE | 6 |
| orf1b | 83 | 4558 | 0 | FALSE | 4 |
| orf1b | 84 | 5170 | 0 | FALSE | 2 |
| orf1b | 85 | 5170 | 0 | TRUE | 6 |
| orf1b | 86 | 5917 | 0 | TRUE | 3 |
| orf1b | 87 | 5917 | 0 | TRUE | 2 |
| orf1b | 88 | 6852 | 0 | FALSE | 5 |
| orf1b | 89 | 6852 | 1280 | TRUE | 2 |
| orf1b | 90 | 5368 | 1280 | TRUE | 2 |
| orf1b | 91 | 5368 | 1280 | FALSE | 2 |
| orf1b | 92 | 8954 | 1280 | TRUE | 2 |
| orf1b | 93 | 8954 | 1280 | FALSE | 1 |
| orf1b | 94 | 0 | 1280 | TRUE | 3 |
| orf1b | 95 | 0 | 1373 | TRUE | 2 |
| orf1b | 96 | 0 | 1373 | TRUE | 4 |
| orf1b | 97 | 0 | 1373 | TRUE | 2 |
| orf1b | 98 | 0 | 1373 | TRUE | 3 |
| orf1b | 99 | 0 | 1361 | TRUE | 4 |
| orf1b | 100 | 0 | 1361 | TRUE | 2 |

| | | | | | |
|---|---|---|---|---|---|
| orf1b | 101 | 1440 | 1361 | TRUE | 3 |
| orf1b | 102 | 1440 | 1361 | TRUE | 4 |
| orf1b | 103 | 1440 | 1361 | TRUE | 2 |
| orf1b | 104 | 1440 | 1401 | FALSE | 3 |
| orf1b | 105 | 1440 | 1401 | FALSE | 3 |
| orf1b | 106 | 1440 | 1401 | FALSE | 3 |
| orf1b | 107 | 1440 | 1401 | FALSE | 3 |
| orf1b | 108 | 1440 | 1401 | FALSE | 3 |
| orf1b | 109 | 1440 | 1401 | FALSE | 2 |
| orf1b | 110 | 1440 | 1337 | TRUE | 2 |
| orf1b | 111 | 1440 | 1337 | TRUE | 3 |
| orf1b | 112 | 1440 | 1337 | FALSE | 3 |
| orf1b | 113 | 1440 | 1171 | TRUE | 3 |
| orf1b | 114 | 1440 | 1006 | TRUE | 4 |
| orf1b | 115 | 1440 | 1006 | TRUE | 5 |
| orf1b | 116 | 0 | 1006 | TRUE | 4 |
| orf1b | 117 | 1085 | 1006 | TRUE | 4 |
| orf1b | 118 | 1085 | 1006 | TRUE | 2 |
| orf1b | 119 | 1085 | 1006 | TRUE | 3 |
| orf1b | 120 | 1085 | 1006 | TRUE | 0 |
| orf1b | 121 | 1058 | 1233 | TRUE | 3 |
| orf1b | 122 | 1058 | 1233 | TRUE | 1 |
| orf1b | 123 | 6011 | 1233 | TRUE | 3 |
| orf1b | 124 | 6011 | 1233 | TRUE | 2 |
| orf1b | 125 | 5637 | 3890 | TRUE | 1 |
| orf1b | 126 | 5637 | 3890 | TRUE | 1 |
| orf1b | 127 | 6660 | 6220 | TRUE | 3 |
| orf1b | 128 | 6660 | 6965 | TRUE | 4 |
| orf1b | 129 | 6048 | 6965 | TRUE | 2 |
| orf1b | 130 | 6048 | 6965 | TRUE | 3 |
| orf1b | 131 | 6048 | 6965 | TRUE | 3 |
| orf1b | 132 | 6668 | 6965 | FALSE | 4 |
| orf1b | 133 | 5390 | 6965 | TRUE | 3 |
| orf1b | 134 | 5390 | 6965 | TRUE | 3 |
| orf1b | 135 | 4957 | 5603 | TRUE | 2 |
| orf1b | 136 | 5188 | 6063 | FALSE | 1 |
| orf1b | 137 | 5188 | 6063 | TRUE | 2 |
| orf1b | 138 | 4518 | 6063 | TRUE | 3 |
| orf1b | 139 | 4315 | 6063 | FALSE | 2 |
| orf1b | 140 | 4302 | 5924 | TRUE | 3 |
| orf1b | 141 | 3809 | 5367 | TRUE | 5 |
| orf1b | 142 | 2776 | 2183 | TRUE | 3 |
| orf1b | 143 | 2669 | 2027 | TRUE | 2 |
| orf1b | 144 | 2771 | 2027 | TRUE | 5 |
| orf1b | 145 | 2771 | 1775 | TRUE | 4 |
| orf1b | 146 | 2771 | 1775 | TRUE | 3 |
| orf1b | 147 | 2667 | 1775 | FALSE | 1 |
| orf1b | 148 | 2612 | 1775 | TRUE | 1 |
| orf1b | 149 | 3037 | 1775 | TRUE | 3 |
| orf1b | 150 | 2132 | 1288 | FALSE | 4 |
| orf1b | 151 | 2132 | 1288 | TRUE | 1 |
| orf1b | 152 | 2132 | 1288 | FALSE | 6 |
| orf1b | 153 | 2132 | 1288 | TRUE | 1 |
| orf1b | 154 | 2309 | 1288 | TRUE | 4 |
| orf1b | 155 | 2309 | 1288 | TRUE | 3 |
| orf1b | 156 | 3438 | 1230 | TRUE | 0 |
| orf1b | 157 | 3438 | 1230 | FALSE | 2 |
| orf1b | 158 | 4624 | 1144 | FALSE | 6 |
| orf1b | 159 | 4624 | 1144 | TRUE | 4 |
| orf1b | 160 | 4624 | 0 | FALSE | 5 |
| orf1b | 161 | 4624 | 0 | TRUE | 5 |
| orf1b | 162 | 8133 | 0 | TRUE | 4 |

| | | | | | |
|---|---|---|---|---|---|
| orf1b | 163 | 8133 | 0 | TRUE | 4 |
| orf1b | 171 | 7144 | 5827 | FALSE | 4 |
| orf1b | 172 | 7144 | 5827 | FALSE | 1 |
| orf1b | 173 | 7144 | 5827 | FALSE | 1 |
| orf1b | 174 | 7144 | 5827 | FALSE | 5 |
| orf1b | 175 | 7144 | 5827 | FALSE | 3 |
| orf1b | 176 | 7144 | 5827 | FALSE | 5 |
| orf1b | 177 | 7144 | 5827 | FALSE | 4 |
| orf1b | 178 | 7144 | 5827 | FALSE | 1 |
| orf1b | 179 | 7144 | 5827 | FALSE | 2 |
| orf1b | 180 | 7144 | 5827 | FALSE | 2 |
| orf1b | 181 | 7144 | 5017 | FALSE | 2 |
| orf1b | 182 | 7144 | 5017 | FALSE | 1 |
| orf1b | 183 | 7144 | 3848 | FALSE | 3 |
| orf1b | 184 | 7144 | 3848 | TRUE | 2 |
| orf1b | 185 | 4455 | 3848 | TRUE | 1 |
| orf1b | 186 | 3111 | 2858 | TRUE | 4 |
| orf1b | 187 | 3111 | 2858 | FALSE | 3 |
| orf1b | 188 | 3111 | 2858 | TRUE | 3 |
| orf1b | 189 | 2816 | 2858 | TRUE | 2 |
| orf1b | 190 | 2816 | 2858 | FALSE | 3 |
| orf1b | 191 | 2816 | 4094 | TRUE | 2 |
| orf1b | 192 | 2816 | 4094 | TRUE | 0 |
| orf1b | 193 | 2816 | 4094 | TRUE | 1 |
| orf1b | 194 | 2816 | 4094 | FALSE | 4 |
| orf1b | 195 | 2816 | 4094 | TRUE | 2 |
| orf1b | 196 | 2816 | 4037 | FALSE | 3 |
| orf1b | 197 | 2385 | 4037 | FALSE | 3 |
| orf1b | 198 | 2385 | 6567 | FALSE | 3 |
| orf1b | 199 | 2151 | 3467 | TRUE | 1 |
| orf1b | 200 | 1511 | 3467 | TRUE | 1 |
| orf1b | 201 | 1511 | 3102 | TRUE | 4 |
| orf1b | 202 | 1511 | 3102 | TRUE | 3 |
| orf1b | 203 | 1511 | 3102 | TRUE | 3 |
| orf1b | 204 | 1154 | 3102 | TRUE | 1 |
| orf1b | 205 | 1154 | 3102 | TRUE | 3 |
| orf1b | 206 | 1154 | 1948 | TRUE | 1 |
| orf1b | 207 | 1154 | 1948 | TRUE | 5 |
| orf1b | 208 | 1154 | 1948 | TRUE | 2 |
| orf1b | 209 | 1154 | 1948 | TRUE | 2 |
| orf1b | 210 | 1154 | 1948 | TRUE | 3 |
| orf1b | 211 | 1154 | 1948 | TRUE | 3 |
| orf1b | 212 | 1215 | 1948 | TRUE | 1 |
| orf1b | 213 | 1215 | 1948 | TRUE | 3 |
| orf1b | 214 | 0 | 2009 | TRUE | 1 |
| orf1b | 215 | 0 | 2009 | TRUE | 2 |
| orf1b | 219 | 1315 | 1137 | TRUE | 4 |
| orf1b | 220 | 1315 | 1137 | TRUE | 3 |
| orf1b | 221 | 2271 | 4759 | TRUE | 3 |
| orf1b | 222 | 2271 | 4759 | TRUE | 3 |
| orf1b | 223 | 2320 | 3838 | TRUE | 3 |
| orf1b | 224 | 2320 | 3838 | TRUE | 2 |
| orf1b | 225 | 2320 | 3838 | FALSE | 4 |
| orf1b | 226 | 2320 | 3838 | TRUE | 2 |
| orf1b | 227 | 2320 | 3838 | TRUE | 2 |
| orf1b | 228 | 2320 | 3838 | FALSE | 3 |
| orf1b | 229 | 2320 | 3838 | TRUE | 1 |
| orf1b | 230 | 2320 | 3838 | TRUE | 3 |
| orf1b | 231 | 2320 | 3838 | TRUE | 2 |
| orf1b | 232 | 2320 | 3838 | TRUE | 0 |
| orf1b | 233 | 2320 | 3838 | TRUE | 3 |
| orf1b | 234 | 2432 | 5638 | TRUE | 3 |

| | | | | | |
|---|---|---|---|---|---|
| orf1b | 235 | 2320 | 5638 | TRUE | 1 |
| orf1b | 236 | 2193 | 2455 | TRUE | 1 |
| orf1b | 237 | 2193 | 2455 | TRUE | 4 |
| orf1b | 238 | 1314 | 0 | TRUE | 1 |
| orf1b | 239 | 1314 | 0 | FALSE | 4 |
| orf1b | 240 | 1314 | 0 | FALSE | 5 |
| orf1b | 241 | 1314 | 0 | FALSE | 3 |
| orf1b | 242 | 1314 | 0 | FALSE | 1 |
| orf1b | 243 | 3657 | 1571 | FALSE | 4 |
| orf1b | 244 | 3657 | 1571 | FALSE | 4 |
| orf1b | 245 | 6309 | 7955 | FALSE | 4 |
| orf1b | 246 | 6309 | 7955 | FALSE | 3 |
| orf1b | 247 | 6309 | 6605 | FALSE | 3 |
| orf1b | 248 | 6309 | 6605 | FALSE | 2 |
| orf1b | 249 | 6309 | 6605 | FALSE | 4 |
| orf1b | 250 | 7022 | 6605 | FALSE | 3 |
| orf1b | 251 | 7022 | 6605 | FALSE | 4 |
| orf1b | 252 | 7022 | 6605 | FALSE | 1 |
| orf1b | 253 | 7022 | 6605 | FALSE | 5 |
| orf1b | 254 | 7022 | 6605 | FALSE | 3 |
| orf1b | 255 | 6383 | 6605 | FALSE | 2 |
| orf1b | 256 | 6383 | 6605 | FALSE | 3 |
| orf1b | 257 | 6383 | 6605 | FALSE | 2 |
| orf1b | 258 | 7549 | 7864 | FALSE | 4 |
| orf1b | 259 | 6900 | 7864 | FALSE | 4 |
| orf1b | 260 | 2779 | 1206 | FALSE | 2 |
| orf1b | 261 | 2779 | 1206 | FALSE | 0 |
| orf1b | 262 | 2779 | 0 | FALSE | 2 |
| orf1b | 263 | 2779 | 0 | FALSE | 0 |
| orf1b | 264 | 2779 | 0 | FALSE | 2 |
| orf1b | 265 | 2779 | 0 | FALSE | 3 |
| orf1b | 266 | 2779 | 0 | FALSE | 2 |
| orf1b | 267 | 2276 | 1139 | TRUE | 5 |
| orf1b | 268 | 2276 | 1139 | TRUE | 2 |
| orf1b | 269 | 3707 | 4000 | TRUE | 5 |
| orf1b | 270 | 4068 | 4000 | TRUE | 2 |
| orf1b | 271 | 4068 | 4000 | TRUE | 2 |
| orf1b | 272 | 4068 | 4000 | TRUE | 0 |
| orf1b | 273 | 4068 | 4000 | TRUE | 2 |
| orf1b | 274 | 4171 | 4000 | TRUE | 1 |
| orf1b | 275 | 4171 | 4000 | TRUE | 2 |
| orf1b | 276 | 4171 | 4000 | TRUE | 4 |
| orf1b | 277 | 4171 | 4000 | FALSE | 1 |
| orf1b | 278 | 4171 | 4000 | TRUE | 0 |
| orf1b | 279 | 4171 | 4000 | FALSE | 4 |
| orf1b | 280 | 4171 | 4000 | FALSE | 2 |
| orf1b | 281 | 4171 | 4000 | TRUE | 4 |
| orf1b | 282 | 10865 | 6861 | TRUE | 3 |
| orf1b | 283 | 6197 | 6861 | TRUE | 4 |
| orf1b | 284 | 1530 | 0 | TRUE | 2 |
| orf1b | 285 | 1530 | 0 | TRUE | 3 |
| orf1b | 286 | 1530 | 0 | TRUE | 2 |
| orf1b | 287 | 1432 | 0 | TRUE | 4 |
| orf1b | 288 | 1432 | 0 | TRUE | 2 |
| orf1b | 289 | 1432 | 0 | TRUE | 3 |
| orf1b | 290 | 1432 | 0 | FALSE | 1 |
| orf1b | 291 | 1325 | 0 | TRUE | 4 |
| orf1b | 292 | 1325 | 0 | FALSE | 1 |
| orf1b | 293 | 1325 | 0 | TRUE | 4 |
| orf1b | 294 | 1325 | 0 | TRUE | 3 |
| orf1b | 295 | 1325 | 0 | TRUE | 2 |
| orf1b | 296 | 1325 | 0 | TRUE | 1 |

| | | | | | |
|---|---|---|---|---|---|
| orf1b | 297 | 1325 | 0 | TRUE | 2 |
| orf1b | 298 | 1257 | 0 | TRUE | 4 |
| orf1b | 299 | 1257 | 0 | TRUE | 2 |
| orf1b | 300 | 1257 | 0 | TRUE | 3 |
| orf1b | 301 | 1257 | 0 | FALSE | 3 |
| orf1b | 302 | 1005 | 0 | TRUE | 4 |
| orf1b | 303 | 1005 | 0 | TRUE | 2 |
| orf1b | 304 | 1005 | 0 | TRUE | 3 |
| orf1b | 305 | 1005 | 0 | TRUE | 2 |
| orf1b | 315 | 0 | 1646 | TRUE | 4 |
| orf1b | 316 | 0 | 1646 | TRUE | 3 |
| orf1b | 317 | 0 | 1646 | TRUE | 3 |
| orf1b | 318 | 0 | 1646 | TRUE | 4 |
| orf1b | 319 | 0 | 1646 | TRUE | 2 |
| orf1b | 320 | 0 | 1646 | TRUE | 4 |
| orf1b | 321 | 0 | 1646 | TRUE | 5 |
| orf1b | 322 | 0 | 1646 | TRUE | 2 |
| orf1b | 323 | 0 | 1646 | TRUE | 2 |
| orf1b | 324 | 0 | 1646 | TRUE | 3 |
| orf1b | 325 | 0 | 1646 | TRUE | 1 |
| orf1b | 326 | 0 | 1646 | TRUE | 1 |
| orf1b | 327 | 1638 | 1432 | TRUE | 2 |
| orf1b | 328 | 1638 | 1432 | TRUE | 3 |
| orf1b | 329 | 1638 | 1432 | TRUE | 4 |
| orf1b | 330 | 1638 | 1217 | TRUE | 3 |
| orf1b | 331 | 1638 | 1960 | TRUE | 2 |
| orf1b | 332 | 1638 | 1960 | TRUE | 2 |
| orf1b | 333 | 1800 | 1960 | TRUE | 5 |
| orf1b | 334 | 1800 | 1960 | TRUE | 4 |
| orf1b | 335 | 1623 | 1960 | TRUE | 2 |
| orf1b | 336 | 1623 | 1960 | TRUE | 1 |
| orf1b | 337 | 1725 | 1960 | TRUE | 1 |
| orf1b | 338 | 1725 | 1960 | TRUE | 2 |
| orf1b | 339 | 1783 | 2052 | TRUE | 2 |
| orf1b | 340 | 1783 | 2052 | TRUE | 3 |
| orf1b | 341 | 3501 | 3226 | TRUE | 3 |
| orf1b | 342 | 3967 | 3627 | TRUE | 1 |
| orf1b | 343 | 3667 | 6178 | TRUE | 3 |
| orf1b | 344 | 3667 | 6178 | TRUE | 2 |
| orf1b | 345 | 3440 | 6178 | TRUE | 4 |
| orf1b | 346 | 3440 | 6873 | TRUE | 1 |
| orf1b | 347 | 3300 | 6873 | TRUE | 1 |
| orf1b | 348 | 3418 | 6873 | TRUE | 3 |
| orf1b | 349 | 3418 | 6873 | TRUE | 4 |
| orf1b | 350 | 3934 | 6873 | TRUE | 2 |
| orf1b | 351 | 4001 | 6873 | TRUE | 2 |
| orf1b | 352 | 4222 | 6873 | TRUE | 2 |
| orf1b | 353 | 4222 | 6873 | TRUE | 2 |
| orf1b | 354 | 4521 | 8033 | TRUE | 2 |
| orf1b | 355 | 4521 | 8033 | TRUE | 2 |
| orf1b | 356 | 2279 | 18933 | FALSE | 2 |
| orf1b | 357 | 2279 | 18933 | TRUE | 2 |
| orf1b | 358 | 2614 | 0 | TRUE | 3 |
| orf1b | 359 | 2593 | 1183 | TRUE | 3 |
| orf1b | 360 | 2992 | 1183 | TRUE | 3 |
| orf1b | 361 | 4359 | 3587 | TRUE | 4 |
| orf1b | 362 | 4973 | 3587 | FALSE | 2 |
| orf1b | 363 | 4973 | 3587 | FALSE | 5 |
| orf1b | 364 | 4973 | 3587 | FALSE | 3 |
| orf1b | 365 | 4973 | 3587 | FALSE | 2 |
| orf1b | 366 | 5116 | 3587 | FALSE | 5 |
| orf1b | 367 | 5116 | 3587 | FALSE | 1 |

| | | | | | |
|---|---|---|---|---|---|
| orf1b | 368 | 5116 | 3587 | TRUE | 2 |
| orf1b | 369 | 5116 | 3587 | TRUE | 5 |
| orf1b | 370 | 5116 | 3587 | TRUE | 4 |
| orf1b | 371 | 5116 | 3587 | TRUE | 3 |
| orf1b | 372 | 5116 | 3587 | TRUE | 2 |
| orf1b | 373 | 5116 | 3587 | TRUE | 4 |
| orf1b | 374 | 6410 | 4388 | TRUE | 6 |
| orf1b | 375 | 6410 | 4388 | TRUE | 2 |
| orf1b | 377 | 6748 | 0 | TRUE | 0 |
| orf1b | 378 | 6748 | 0 | TRUE | 4 |
| orf1b | 379 | 6748 | 0 | TRUE | 2 |
| orf1b | 380 | 6748 | 0 | FALSE | 3 |
| orf1b | 381 | 6748 | 0 | TRUE | 3 |
| orf1b | 382 | 6748 | 0 | TRUE | 2 |
| orf1b | 383 | 6748 | 0 | TRUE | 4 |
| orf1b | 384 | 6748 | 0 | TRUE | 2 |
| orf1b | 385 | 6748 | 0 | TRUE | 1 |
| orf1b | 386 | 6748 | 0 | TRUE | 2 |
| orf1b | 387 | 6748 | 0 | TRUE | 2 |
| orf1b | 388 | 6748 | 0 | TRUE | 2 |
| orf1b | 389 | 6748 | 0 | TRUE | 0 |
| orf1b | 390 | 6748 | 0 | FALSE | 3 |
| orf1b | 391 | 6748 | 0 | TRUE | 2 |
| orf1b | 405 | 1355 | 0 | FALSE | 3 |
| orf1b | 406 | 1355 | 0 | FALSE | 3 |
| orf1b | 407 | 4347 | 0 | TRUE | 3 |
| orf1b | 408 | 4347 | 0 | TRUE | 1 |
| orf1b | 409 | 4347 | 0 | TRUE | 3 |
| orf1b | 410 | 4347 | 0 | TRUE | 3 |
| orf1b | 411 | 4347 | 0 | TRUE | 5 |
| orf1b | 412 | 4347 | 0 | TRUE | 4 |
| orf1b | 413 | 4347 | 0 | FALSE | 0 |
| orf1b | 414 | 4347 | 0 | TRUE | 5 |
| orf1b | 415 | 4347 | 0 | TRUE | 6 |
| orf1b | 416 | 4347 | 0 | TRUE | 4 |
| orf1b | 417 | 4347 | 0 | TRUE | 1 |
| orf1b | 418 | 4347 | 0 | TRUE | 5 |
| orf1b | 419 | 4347 | 0 | TRUE | 2 |
| orf1b | 420 | 7340 | 0 | TRUE | 0 |
| orf1b | 421 | 7340 | 0 | TRUE | 5 |
| orf1b | 425 | 9990 | 0 | TRUE | 2 |
| orf1b | 426 | 9990 | 0 | TRUE | 2 |
| orf1b | 427 | 6108 | 3325 | TRUE | 6 |
| orf1b | 428 | 6108 | 3325 | TRUE | 4 |
| orf1b | 429 | 6108 | 3325 | TRUE | 2 |
| orf1b | 430 | 6108 | 3325 | TRUE | 3 |
| orf1b | 431 | 6108 | 3325 | TRUE | 3 |
| orf1b | 432 | 6108 | 3325 | TRUE | 2 |
| orf1b | 433 | 6108 | 3325 | FALSE | 0 |
| orf1b | 434 | 6108 | 3325 | TRUE | 2 |
| orf1b | 435 | 6108 | 3325 | TRUE | 4 |
| orf1b | 436 | 6108 | 3325 | TRUE | 5 |
| orf1b | 437 | 6108 | 3325 | TRUE | 3 |
| orf1b | 438 | 6108 | 3325 | TRUE | 1 |
| orf1b | 439 | 6108 | 3325 | TRUE | 4 |
| orf1b | 440 | 4814 | 3325 | TRUE | 4 |
| orf1b | 441 | 4814 | 3325 | TRUE | 1 |
| orf1b | 443 | 1514 | 0 | TRUE | 1 |
| orf1b | 444 | 1514 | 0 | FALSE | 3 |
| orf1b | 445 | 1514 | 0 | TRUE | 1 |
| orf1b | 446 | 1514 | 0 | TRUE | 2 |
| orf1b | 447 | 1514 | 0 | TRUE | 2 |

| | | | | | |
|---|---|---|---|---|---|
| orf1b | 448 | 1514 | 0 | TRUE | 4 |
| orf1b | 449 | 1514 | 0 | TRUE | 2 |
| orf1b | 450 | 1514 | 0 | FALSE | 1 |
| orf1b | 451 | 1514 | 0 | TRUE | 3 |
| orf1b | 452 | 1514 | 0 | TRUE | 5 |
| orf1b | 453 | 1828 | 0 | TRUE | 2 |
| orf1b | 454 | 1828 | 0 | TRUE | 4 |
| orf1b | 455 | 1828 | 0 | TRUE | 4 |
| orf1b | 456 | 1828 | 0 | TRUE | 3 |
| orf1b | 457 | 1828 | 0 | TRUE | 5 |
| orf1b | 458 | 2142 | 0 | TRUE | 4 |
| orf1b | 459 | 2142 | 0 | FALSE | 1 |
| orf1b | 460 | 2142 | 0 | TRUE | 1 |
| orf1b | 461 | 2142 | 0 | TRUE | 4 |
| orf1b | 462 | 2142 | 0 | TRUE | 6 |
| orf1b | 463 | 6797 | 0 | TRUE | 2 |
| orf1b | 464 | 6797 | 0 | TRUE | 2 |
| orf1b | 465 | 7361 | 4578 | TRUE | 3 |
| orf1b | 466 | 7361 | 4578 | FALSE | 2 |
| orf1b | 467 | 7361 | 4578 | TRUE | 1 |
| orf1b | 468 | 8405 | 4578 | TRUE | 4 |
| orf1b | 469 | 8405 | 4578 | FALSE | 5 |
| orf1b | 470 | 8405 | 4578 | TRUE | 1 |
| orf1b | 471 | 8405 | 4578 | TRUE | 3 |
| orf1b | 472 | 8405 | 4578 | TRUE | 4 |
| orf1b | 473 | 8405 | 4578 | TRUE | 2 |
| orf1b | 474 | 8405 | 4578 | TRUE | 2 |
| orf1b | 475 | 8405 | 4578 | TRUE | 1 |
| orf1b | 476 | 8405 | 4578 | FALSE | 4 |
| orf1b | 477 | 8405 | 4578 | TRUE | 3 |
| orf1b | 478 | 7926 | 4578 | TRUE | 2 |
| orf1b | 479 | 7926 | 4578 | TRUE | 2 |
| orf1b | 487 | 11301 | 0 | TRUE | 2 |
| orf1b | 488 | 11301 | 0 | FALSE | 4 |
| orf1b | 489 | 11301 | 0 | TRUE | 4 |
| orf1b | 490 | 11301 | 0 | FALSE | 5 |
| orf1b | 491 | 11301 | 0 | TRUE | 1 |
| orf1b | 492 | 11301 | 0 | TRUE | 1 |
| orf1b | 493 | 11301 | 0 | TRUE | 4 |
| orf1b | 494 | 11301 | 0 | TRUE | 1 |
| orf1b | 495 | 11301 | 0 | TRUE | 3 |
| orf1b | 496 | 11301 | 0 | TRUE | 4 |
| orf1b | 497 | 11301 | 0 | TRUE | 1 |
| orf1b | 498 | 11301 | 0 | TRUE | 5 |
| orf1b | 499 | 11301 | 0 | TRUE | 2 |
| orf1b | 500 | 11301 | 0 | TRUE | 1 |
| orf1b | 501 | 11301 | 0 | TRUE | 4 |
| orf1b | 503 | 1879 | 0 | TRUE | 2 |
| orf1b | 504 | 1879 | 0 | TRUE | 2 |
| orf1b | 505 | 1739 | 0 | TRUE | 3 |
| orf1b | 506 | 1739 | 0 | TRUE | 1 |
| orf1b | 507 | 1739 | 0 | TRUE | 1 |
| orf1b | 508 | 1739 | 0 | TRUE | 3 |
| orf1b | 509 | 1860 | 0 | TRUE | 2 |
| orf1b | 510 | 1860 | 0 | TRUE | 1 |
| orf1b | 511 | 5740 | 0 | TRUE | 2 |
| orf1b | 512 | 5740 | 0 | FALSE | 1 |
| orf1b | 513 | 9463 | 4727 | TRUE | 2 |
| orf1b | 514 | 9463 | 4727 | TRUE | 3 |
| orf1b | 515 | 9463 | 4727 | TRUE | 3 |
| orf1b | 516 | 9463 | 4727 | TRUE | 1 |
| orf1b | 517 | 9463 | 4727 | TRUE | 3 |

| | | | | | |
|---|---|---|---|---|---|
| orf1b | 518 | 10727 | 4727 | TRUE | 5 |
| orf1b | 519 | 10727 | 4727 | TRUE | 2 |
| orf1b | 520 | 12553 | 4727 | TRUE | 3 |
| orf1b | 521 | 12553 | 4727 | TRUE | 4 |
| orf1b | 522 | 12553 | 4727 | TRUE | 4 |
| orf1b | 523 | 12553 | 4727 | TRUE | 2 |
| orf1b | 524 | 15166 | 4727 | TRUE | 3 |
| orf1b | 525 | 15166 | 4727 | FALSE | 3 |
| orf1b | 526 | 14428 | 4727 | TRUE | 2 |
| orf1b | 527 | 14428 | 4727 | TRUE | 1 |
| orf1b | 543 | 12448 | 0 | TRUE | 2 |
| orf1b | 544 | 12448 | 0 | TRUE | 2 |
| orf1b | 545 | 12448 | 0 | TRUE | 4 |
| orf1b | 546 | 12448 | 0 | TRUE | 5 |
| orf1b | 547 | 12448 | 0 | TRUE | 3 |
| orf1b | 548 | 12448 | 0 | TRUE | 0 |
| orf1b | 549 | 12448 | 0 | TRUE | 8 |
| orf1b | 550 | 12448 | 0 | TRUE | 2 |
| orf1b | 551 | 12448 | 0 | TRUE | 4 |
| orf1b | 552 | 12448 | 0 | TRUE | 3 |
| orf1b | 553 | 12448 | 0 | TRUE | 1 |
| orf1b | 554 | 12448 | 0 | TRUE | 1 |
| orf1b | 555 | 12448 | 0 | TRUE | 2 |
| orf1b | 556 | 12448 | 0 | TRUE | 2 |
| orf1b | 557 | 12448 | 0 | TRUE | 4 |
| orf1b | 559 | 7566 | 0 | TRUE | 2 |
| orf1b | 560 | 7566 | 0 | TRUE | 3 |
| orf1b | 561 | 7566 | 0 | TRUE | 3 |
| orf1b | 562 | 7566 | 0 | TRUE | 2 |
| orf1b | 563 | 7566 | 0 | TRUE | 1 |
| orf1b | 564 | 7566 | 0 | TRUE | 4 |
| orf1b | 565 | 7566 | 0 | TRUE | 2 |
| orf1b | 566 | 7566 | 0 | TRUE | 1 |
| orf1b | 567 | 7566 | 0 | TRUE | 2 |
| orf1b | 568 | 7566 | 0 | TRUE | 1 |
| orf1b | 569 | 7566 | 0 | TRUE | 2 |
| orf1b | 570 | 7566 | 0 | TRUE | 3 |
| orf1b | 571 | 7566 | 0 | TRUE | 3 |
| orf1b | 572 | 7566 | 0 | TRUE | 3 |
| orf1b | 573 | 7566 | 0 | TRUE | 2 |
| orf1b | 577 | 1912 | 0 | TRUE | 3 |
| orf1b | 578 | 1912 | 0 | TRUE | 3 |
| orf1b | 579 | 1643 | 0 | TRUE | 5 |
| orf1b | 580 | 1643 | 0 | TRUE | 4 |
| orf1b | 581 | 1643 | 0 | TRUE | 5 |
| orf1b | 582 | 1643 | 0 | TRUE | 3 |
| orf1b | 583 | 1643 | 0 | TRUE | 6 |
| orf1b | 584 | 1643 | 0 | TRUE | 1 |
| orf1b | 585 | 1643 | 0 | TRUE | 4 |
| orf1b | 586 | 1643 | 0 | TRUE | 2 |
| orf1b | 587 | 1643 | 0 | TRUE | 2 |
| orf1b | 588 | 1643 | 0 | TRUE | 3 |
| orf1b | 589 | 1643 | 0 | TRUE | 4 |
| orf1b | 590 | 1643 | 0 | TRUE | 3 |
| orf1b | 591 | 2817 | 0 | TRUE | 3 |
| orf1b | 592 | 3269 | 0 | TRUE | 4 |
| orf1b | 593 | 3269 | 0 | TRUE | 3 |
| orf1b | 594 | 5166 | 0 | TRUE | 4 |
| orf1b | 595 | 5166 | 0 | TRUE | 2 |
| orf1b | 596 | 5166 | 0 | TRUE | 3 |
| orf1b | 597 | 5166 | 0 | TRUE | 2 |
| orf1b | 598 | 5166 | 0 | FALSE | 1 |

| | | | | | |
|---|---|---|---|---|---|
| orf1b | 599 | 6185 | 0 | TRUE | 3 |
| orf1b | 600 | 6185 | 0 | TRUE | 4 |
| orf1b | 601 | 7176 | 3002 | TRUE | 3 |
| orf1b | 602 | 7176 | 3002 | TRUE | 2 |
| orf1b | 603 | 6231 | 3002 | FALSE | 3 |
| orf1b | 604 | 6231 | 3002 | TRUE | 3 |
| orf1b | 605 | 6231 | 3002 | TRUE | 3 |
| orf1b | 606 | 6444 | 3002 | FALSE | 5 |
| orf1b | 607 | 6444 | 3002 | FALSE | 6 |
| orf1b | 608 | 6444 | 3002 | TRUE | 1 |
| orf1b | 609 | 6444 | 3002 | TRUE | 4 |
| orf1b | 610 | 6444 | 3002 | TRUE | 1 |
| orf1b | 611 | 6444 | 3002 | TRUE | 2 |
| orf1b | 612 | 6444 | 3002 | FALSE | 2 |
| orf1b | 613 | 6444 | 3002 | TRUE | 2 |
| orf1b | 614 | 6254 | 3002 | TRUE | 1 |
| orf1b | 615 | 6254 | 3002 | TRUE | 3 |
| orf1b | 616 | 1509 | 0 | TRUE | 2 |
| orf1b | 617 | 1509 | 0 | TRUE | 3 |
| orf1b | 619 | 5998 | 0 | TRUE | 1 |
| orf1b | 620 | 5998 | 0 | TRUE | 4 |
| orf1b | 621 | 5998 | 0 | TRUE | 3 |
| orf1b | 622 | 5998 | 0 | TRUE | 3 |
| orf1b | 623 | 5998 | 0 | TRUE | 1 |
| orf1b | 624 | 5998 | 0 | TRUE | 4 |
| orf1b | 625 | 5998 | 0 | TRUE | 4 |
| orf1b | 626 | 5998 | 0 | TRUE | 1 |
| orf1b | 627 | 5998 | 0 | TRUE | 2 |
| orf1b | 628 | 5998 | 0 | TRUE | 4 |
| orf1b | 629 | 5998 | 0 | TRUE | 2 |
| orf1b | 630 | 5998 | 0 | TRUE | 3 |
| orf1b | 631 | 5998 | 0 | TRUE | 4 |
| orf1b | 632 | 5998 | 0 | TRUE | 2 |
| orf1b | 633 | 5998 | 0 | TRUE | 1 |
| orf1b | 645 | 1512 | 0 | TRUE | 3 |
| orf1b | 646 | 1512 | 0 | TRUE | 1 |
| orf1b | 647 | 1512 | 0 | FALSE | 6 |
| orf1b | 648 | 1512 | 0 | TRUE | 2 |
| orf1b | 649 | 7060 | 2871 | FALSE | 5 |
| orf1b | 650 | 7060 | 2871 | TRUE | 3 |
| orf1b | 651 | 7060 | 2871 | TRUE | 2 |
| orf1b | 652 | 7060 | 2871 | TRUE | 3 |
| orf1b | 653 | 7060 | 2871 | TRUE | 4 |
| orf1b | 654 | 7060 | 2871 | TRUE | 3 |
| orf1b | 655 | 7060 | 2871 | FALSE | 2 |
| orf1b | 656 | 7060 | 2871 | TRUE | 1 |
| orf1b | 657 | 7060 | 2871 | TRUE | 3 |
| orf1b | 658 | 7060 | 2871 | TRUE | 5 |
| orf1b | 659 | 7060 | 2871 | TRUE | 4 |
| orf1b | 660 | 12607 | 2871 | TRUE | 3 |
| orf1b | 661 | 12607 | 2871 | TRUE | 3 |
| orf1b | 662 | 12607 | 2871 | TRUE | 3 |
| orf1b | 663 | 12607 | 2871 | TRUE | 1 |
| orf1b | 701 | 7762 | 0 | FALSE | 2 |
| orf1b | 702 | 7762 | 0 | FALSE | 2 |
| orf1b | 703 | 7762 | 0 | FALSE | 2 |
| orf1b | 704 | 7762 | 0 | FALSE | 2 |
| orf1b | 705 | 7762 | 0 | FALSE | 3 |
| orf1b | 706 | 7762 | 0 | FALSE | 1 |
| orf1b | 707 | 7762 | 0 | FALSE | 4 |
| orf1b | 708 | 7762 | 0 | FALSE | 3 |
| orf1b | 709 | 7762 | 0 | FALSE | 3 |

| | | | | | |
|---|---|---|---|---|---|
| orf1b | 710 | 7762 | 0 | FALSE | 1 |
| orf1b | 711 | 7762 | 0 | FALSE | 2 |
| orf1b | 712 | 7762 | 0 | FALSE | 3 |
| orf1b | 713 | 7772 | 2969 | FALSE | 1 |
| orf1b | 714 | 7772 | 2969 | FALSE | 2 |
| orf1b | 715 | 7772 | 2098 | FALSE | 1 |
| orf1b | 716 | 7782 | 2098 | FALSE | 2 |
| orf1b | 717 | 4425 | 2175 | FALSE | 1 |
| orf1b | 718 | 4425 | 2175 | FALSE | 1 |
| orf1b | 719 | 4425 | 2175 | FALSE | 3 |
| orf1b | 720 | 4425 | 2175 | FALSE | 2 |
| orf1b | 721 | 3436 | 2388 | FALSE | 3 |
| orf1b | 722 | 3436 | 2388 | FALSE | 2 |
| orf1b | 723 | 2572 | 2388 | TRUE | 1 |
| orf1b | 724 | 2572 | 2388 | TRUE | 3 |
| orf1b | 725 | 7399 | 2388 | TRUE | 3 |
| orf1b | 726 | 7399 | 2388 | TRUE | 3 |
| orf1b | 727 | 7399 | 2388 | TRUE | 3 |
| orf1b | 728 | 7336 | 2194 | TRUE | 4 |
| orf1b | 729 | 7336 | 2194 | FALSE | 5 |
| orf1b | 730 | 7336 | 2677 | TRUE | 2 |
| orf1b | 731 | 7336 | 2677 | TRUE | 2 |
| orf1b | 732 | 8589 | 3028 | TRUE | 2 |
| orf1b | 733 | 8589 | 3028 | FALSE | 4 |
| orf1b | 734 | 8589 | 3028 | FALSE | 1 |
| orf1b | 735 | 7846 | 2632 | FALSE | 4 |
| orf1b | 736 | 9124 | 2237 | TRUE | 2 |
| orf1b | 737 | 9124 | 2237 | TRUE | 2 |
| orf1b | 738 | 14357 | 2237 | FALSE | 6 |
| orf1b | 739 | 14357 | 2237 | TRUE | 1 |
| orf1b | 740 | 4132 | 2237 | TRUE | 0 |
| orf1b | 741 | 4132 | 2237 | TRUE | 4 |
| orf1b | 742 | 4132 | 2237 | TRUE | 2 |
| orf1b | 743 | 4132 | 2237 | TRUE | 3 |
| orf1b | 744 | 4132 | 2237 | TRUE | 3 |
| orf1b | 745 | 3225 | 2237 | TRUE | 3 |
| orf1b | 746 | 3225 | 2237 | FALSE | 1 |
| orf1b | 747 | 3225 | 2237 | TRUE | 5 |
| orf1b | 748 | 3225 | 2237 | TRUE | 2 |
| orf1b | 749 | 3225 | 2237 | FALSE | 3 |
| orf1b | 750 | 2319 | 0 | TRUE | 2 |
| orf1b | 751 | 2319 | 0 | TRUE | 3 |
| orf1b | 752 | 2319 | 0 | TRUE | 1 |
| orf1b | 753 | 2319 | 0 | TRUE | 2 |
| orf1b | 754 | 2319 | 0 | TRUE | 2 |
| orf1b | 755 | 2319 | 0 | TRUE | 4 |
| orf1b | 756 | 2319 | 0 | TRUE | 2 |
| orf1b | 757 | 2319 | 0 | TRUE | 2 |
| orf1b | 758 | 2319 | 0 | TRUE | 4 |
| orf1b | 759 | 2319 | 0 | TRUE | 3 |
| orf1b | 765 | 0 | 2675 | TRUE | 3 |
| orf1b | 766 | 0 | 2675 | FALSE | 3 |
| orf1b | 767 | 0 | 2675 | FALSE | 3 |
| orf1b | 768 | 0 | 2675 | FALSE | 3 |
| orf1b | 769 | 0 | 2675 | FALSE | 2 |
| orf1b | 770 | 0 | 2675 | FALSE | 2 |
| orf1b | 771 | 0 | 2675 | FALSE | 4 |
| orf1b | 772 | 0 | 2675 | FALSE | 1 |
| orf1b | 773 | 1233 | 2675 | TRUE | 3 |
| orf1b | 774 | 1233 | 2675 | TRUE | 3 |
| orf1b | 775 | 1233 | 2675 | TRUE | 1 |
| orf1b | 776 | 1233 | 2675 | TRUE | 1 |

| | | | | | |
|---|---|---|---|---|---|
| orf1b | 777 | 1233 | 2675 | TRUE | 2 |
| orf1b | 778 | 1233 | 2675 | TRUE | 1 |
| orf1b | 779 | 1482 | 2675 | FALSE | 2 |
| orf1b | 780 | 1482 | 0 | TRUE | 4 |
| orf1b | 781 | 7026 | 0 | TRUE | 2 |
| orf1b | 782 | 7026 | 0 | TRUE | 2 |
| orf1b | 783 | 7026 | 0 | TRUE | 3 |
| orf1b | 784 | 7026 | 0 | TRUE | 3 |
| orf1b | 785 | 6278 | 2419 | TRUE | 4 |
| orf1b | 786 | 6278 | 2419 | TRUE | 3 |
| orf1b | 787 | 8425 | 6679 | TRUE | 2 |
| orf1b | 788 | 9863 | 6679 | TRUE | 3 |
| orf1b | 789 | 7358 | 6679 | TRUE | 3 |
| orf1b | 790 | 7358 | 6679 | TRUE | 2 |
| orf1b | 791 | 8370 | 6679 | TRUE | 5 |
| orf1b | 792 | 8370 | 6679 | TRUE | 4 |
| orf1b | 793 | 7995 | 5127 | TRUE | 3 |
| orf1b | 794 | 8442 | 5127 | TRUE | 3 |
| orf1b | 795 | 8011 | 5127 | TRUE | 5 |
| orf1b | 796 | 7290 | 5127 | TRUE | 2 |
| orf1b | 797 | 7065 | 5127 | TRUE | 5 |
| orf1b | 798 | 7065 | 5127 | TRUE | 5 |
| orf1b | 799 | 6494 | 4560 | TRUE | 5 |
| orf1b | 800 | 6648 | 4917 | TRUE | 4 |
| orf1b | 801 | 6648 | 4917 | TRUE | 1 |
| orf1b | 802 | 5781 | 2971 | FALSE | 4 |
| orf1b | 803 | 5220 | 2971 | TRUE | 2 |
| orf1b | 804 | 5809 | 2971 | TRUE | 4 |
| orf1b | 805 | 5809 | 2971 | TRUE | 2 |
| orf1b | 806 | 4792 | 2971 | FALSE | 4 |
| orf1b | 807 | 4792 | 2971 | TRUE | 1 |
| orf1b | 808 | 2151 | 1156 | TRUE | 2 |
| orf1b | 809 | 2151 | 1156 | TRUE | 0 |
| orf1b | 810 | 2185 | 1156 | TRUE | 2 |
| orf1b | 811 | 2185 | 1156 | TRUE | 3 |
| orf1b | 812 | 1752 | 1156 | TRUE | 4 |
| orf1b | 813 | 1752 | 1159 | TRUE | 5 |
| orf1b | 814 | 1292 | 1163 | TRUE | 5 |
| orf1b | 815 | 1867 | 1383 | TRUE | 2 |
| orf1b | 816 | 1867 | 1383 | TRUE | 2 |
| orf1b | 817 | 1867 | 1383 | TRUE | 2 |
| orf1b | 818 | 3018 | 1383 | TRUE | 1 |
| orf1b | 819 | 3018 | 1383 | TRUE | 1 |
| orf1b | 820 | 3018 | 1383 | FALSE | 1 |
| orf1b | 821 | 4063 | 1383 | FALSE | 2 |
| orf1b | 822 | 4063 | 1383 | FALSE | 3 |
| orf1b | 823 | 4063 | 1383 | FALSE | 4 |
| orf1b | 824 | 4063 | 1383 | FALSE | 2 |
| orf1b | 825 | 4063 | 1383 | FALSE | 3 |
| orf1b | 826 | 4063 | 1383 | TRUE | 3 |
| orf1b | 827 | 4063 | 1383 | TRUE | 4 |
| orf1b | 828 | 4063 | 1603 | TRUE | 3 |
| orf1b | 829 | 4063 | 1603 | TRUE | 2 |
| orf1b | 830 | 5108 | 0 | TRUE | 5 |
| orf1b | 831 | 5108 | 0 | TRUE | 6 |
| orf1b | 832 | 5108 | 0 | TRUE | 3 |
| orf1b | 833 | 5108 | 0 | TRUE | 1 |
| orf1b | 834 | 5108 | 0 | TRUE | 3 |
| orf1b | 835 | 5108 | 0 | TRUE | 4 |
| orf1b | 841 | 6872 | 0 | TRUE | 2 |
| orf1b | 842 | 6872 | 0 | TRUE | 2 |
| orf1b | 843 | 6872 | 0 | TRUE | 1 |

| | | | | | |
|---|---|---|---|---|---|
| orf1b | 844 | 6872 | 0 | TRUE | 2 |
| orf1b | 845 | 6872 | 0 | TRUE | 1 |
| orf1b | 846 | 6872 | 0 | TRUE | 3 |
| orf1b | 847 | 6872 | 0 | TRUE | 4 |
| orf1b | 848 | 6872 | 0 | TRUE | 4 |
| orf1b | 849 | 6872 | 0 | TRUE | 5 |
| orf1b | 850 | 6872 | 0 | TRUE | 3 |
| orf1b | 851 | 6872 | 0 | TRUE | 3 |
| orf1b | 852 | 6872 | 0 | TRUE | 3 |
| orf1b | 853 | 6872 | 0 | TRUE | 3 |
| orf1b | 854 | 6872 | 0 | TRUE | 5 |
| orf1b | 855 | 6872 | 0 | TRUE | 2 |
| orf1b | 857 | 1418 | 0 | TRUE | 2 |
| orf1b | 858 | 1418 | 0 | TRUE | 1 |
| orf1b | 859 | 1553 | 0 | TRUE | 3 |
| orf1b | 860 | 1553 | 0 | TRUE | 1 |
| orf1b | 861 | 1553 | 0 | TRUE | 5 |
| orf1b | 862 | 1553 | 0 | TRUE | 0 |
| orf1b | 863 | 1553 | 0 | TRUE | 2 |
| orf1b | 864 | 1553 | 0 | TRUE | 3 |
| orf1b | 865 | 1450 | 0 | TRUE | 1 |
| orf1b | 866 | 1450 | 0 | TRUE | 4 |
| orf1b | 867 | 3905 | 4323 | TRUE | 3 |
| orf1b | 868 | 3905 | 4323 | TRUE | 0 |
| orf1b | 869 | 3905 | 4323 | TRUE | 5 |
| orf1b | 870 | 3905 | 4323 | TRUE | 4 |
| orf1b | 871 | 3905 | 4323 | TRUE | 2 |
| orf1b | 872 | 4527 | 4323 | FALSE | 2 |
| orf1b | 873 | 3909 | 4323 | TRUE | 2 |
| orf1b | 874 | 5435 | 4323 | FALSE | 2 |
| orf1b | 875 | 4557 | 4323 | TRUE | 1 |
| orf1b | 876 | 4557 | 4323 | TRUE | 2 |
| orf1b | 877 | 3945 | 4323 | FALSE | 3 |
| orf1b | 878 | 3945 | 4323 | TRUE | 1 |
| orf1b | 879 | 3547 | 4323 | TRUE | 4 |
| orf1b | 880 | 4028 | 4323 | TRUE | 5 |
| orf1b | 881 | 3673 | 4323 | FALSE | 3 |
| orf1b | 882 | 1662 | 0 | FALSE | 2 |
| orf1b | 883 | 2636 | 2211 | FALSE | 1 |
| orf1b | 884 | 2636 | 2211 | FALSE | 2 |
| orf1b | 885 | 3735 | 4507 | FALSE | 0 |
| orf1b | 886 | 3735 | 4507 | FALSE | 2 |
| orf1b | 887 | 3392 | 3908 | FALSE | 2 |
| orf1b | 888 | 3569 | 3908 | FALSE | 2 |
| orf1b | 889 | 3569 | 3908 | FALSE | 2 |
| orf1b | 890 | 3734 | 3908 | FALSE | 2 |
| orf1b | 891 | 3734 | 3908 | FALSE | 0 |
| orf1b | 892 | 3982 | 3908 | FALSE | 2 |
| orf1b | 893 | 3982 | 3908 | FALSE | 0 |
| orf1b | 894 | 4286 | 3908 | FALSE | 1 |
| orf1b | 895 | 4286 | 3908 | FALSE | 1 |
| orf1b | 896 | 4627 | 3908 | FALSE | 0 |
| orf1b | 897 | 4627 | 3908 | FALSE | 3 |
| orf1b | 898 | 4449 | 4332 | FALSE | 2 |
| orf1b | 899 | 3618 | 3678 | FALSE | 3 |
| orf1b | 900 | 2032 | 1285 | FALSE | 5 |
| orf1b | 901 | 2032 | 1285 | FALSE | 4 |
| orf1b | 902 | 1539 | 1059 | FALSE | 1 |
| orf1b | 903 | 1539 | 1059 | FALSE | 2 |
| orf1b | 904 | 1539 | 1059 | FALSE | 5 |
| orf1b | 905 | 1670 | 1059 | FALSE | 4 |
| orf1b | 906 | 1670 | 1059 | FALSE | 4 |

| | | | | | |
|---|---|---|---|---|---|
| orf1b | 907 | 2016 | 1139 | FALSE | 4 |
| orf1b | 908 | 2016 | 1139 | FALSE | 4 |
| orf1b | 909 | 2022 | 1589 | FALSE | 2 |
| orf1b | 910 | 2022 | 1589 | FALSE | 2 |
| orf1b | 911 | 2884 | 1511 | FALSE | 0 |
| orf1b | 912 | 2884 | 1511 | FALSE | 1 |
| orf1b | 913 | 3972 | 5456 | FALSE | 4 |
| orf1b | 914 | 4458 | 6336 | FALSE | 3 |
| orf1b | 915 | 4458 | 6336 | FALSE | 2 |
| orf1b | 916 | 4458 | 6336 | FALSE | 2 |
| orf1b | 917 | 4458 | 6336 | FALSE | 2 |
| orf1b | 918 | 4458 | 6336 | FALSE | 4 |
| orf1b | 919 | 4458 | 6336 | FALSE | 3 |
| orf1b | 920 | 5123 | 6336 | FALSE | 2 |
| orf1b | 921 | 5123 | 6336 | FALSE | 2 |
| orf1b | 922 | 7883 | 7615 | FALSE | 2 |
| orf1b | 923 | 7883 | 7615 | FALSE | 4 |
| orf1b | 924 | 9820 | 9324 | FALSE | 2 |
| orf1b | 925 | 9820 | 9324 | FALSE | 2 |
| orf1b | 926 | 9412 | 13347 | FALSE | 4 |
| orf1b | 927 | 9412 | 13347 | FALSE | 3 |
| orf1b | 955 | 2132 | 0 | TRUE | 2 |
| orf1b | 956 | 2132 | 0 | TRUE | 5 |
| orf1b | 957 | 2132 | 0 | TRUE | 5 |
| orf1b | 958 | 2132 | 0 | TRUE | 3 |
| orf1b | 959 | 2132 | 0 | TRUE | 3 |
| orf1b | 960 | 2132 | 0 | TRUE | 3 |
| orf1b | 961 | 2132 | 1230 | TRUE | 3 |
| orf1b | 962 | 2132 | 1230 | TRUE | 3 |
| orf1b | 963 | 2132 | 1230 | TRUE | 1 |
| orf1b | 964 | 2132 | 1230 | TRUE | 0 |
| orf1b | 965 | 2132 | 1230 | TRUE | 3 |
| orf1b | 966 | 2132 | 1230 | TRUE | 2 |
| orf1b | 967 | 2132 | 1230 | TRUE | 2 |
| orf1b | 968 | 2132 | 1230 | TRUE | 3 |
| orf1b | 969 | 2132 | 1230 | TRUE | 1 |
| orf1b | 970 | 0 | 1230 | TRUE | 3 |
| orf1b | 971 | 0 | 1230 | TRUE | 2 |
| orf1b | 972 | 0 | 1230 | TRUE | 3 |
| orf1b | 973 | 0 | 1230 | TRUE | 3 |
| orf1b | 974 | 0 | 1230 | TRUE | 6 |
| orf1b | 975 | 0 | 1717 | TRUE | 2 |
| orf1b | 976 | 0 | 2203 | TRUE | 4 |
| orf1b | 977 | 0 | 3119 | TRUE | 3 |
| orf1b | 978 | 0 | 3119 | TRUE | 1 |
| orf1b | 979 | 0 | 3119 | TRUE | 3 |
| orf1b | 980 | 0 | 3119 | TRUE | 3 |
| orf1b | 981 | 8692 | 3119 | TRUE | 4 |
| orf1b | 982 | 8692 | 3119 | TRUE | 2 |
| orf1b | 983 | 8692 | 3119 | TRUE | 3 |
| orf1b | 984 | 8692 | 3119 | TRUE | 2 |
| orf1b | 985 | 8692 | 3119 | TRUE | 2 |
| orf1b | 986 | 8692 | 3119 | TRUE | 3 |
| orf1b | 987 | 8692 | 3119 | FALSE | 1 |
| orf1b | 988 | 8692 | 3119 | TRUE | 0 |
| orf1b | 989 | 8692 | 3119 | TRUE | 2 |
| orf1b | 990 | 8692 | 4034 | TRUE | 3 |
| orf1b | 991 | 8692 | 4034 | TRUE | 1 |
| orf1b | 992 | 8692 | 0 | TRUE | 2 |
| orf1b | 993 | 8692 | 0 | TRUE | 1 |
| orf1b | 994 | 8692 | 0 | TRUE | 4 |
| orf1b | 995 | 8692 | 0 | TRUE | 3 |

| | | | | | |
|---|---|---|---|---|---|
| orf1b | 1015 | 0 | 2572 | TRUE | 3 |
| orf1b | 1016 | 0 | 2572 | TRUE | 2 |
| orf1b | 1017 | 0 | 2670 | TRUE | 2 |
| orf1b | 1018 | 0 | 2670 | TRUE | 1 |
| orf1b | 1019 | 0 | 2844 | TRUE | 3 |
| orf1b | 1020 | 0 | 2844 | TRUE | 2 |
| orf1b | 1021 | 0 | 2594 | TRUE | 3 |
| orf1b | 1022 | 0 | 2594 | TRUE | 2 |
| orf1b | 1023 | 9628 | 2309 | TRUE | 3 |
| orf1b | 1024 | 9628 | 2309 | FALSE | 1 |
| orf1b | 1025 | 9628 | 2309 | FALSE | 3 |
| orf1b | 1026 | 9628 | 2309 | TRUE | 2 |
| orf1b | 1027 | 8007 | 2309 | TRUE | 1 |
| orf1b | 1028 | 8007 | 2309 | TRUE | 5 |
| orf1b | 1029 | 8007 | 2309 | TRUE | 3 |
| orf1b | 1030 | 8007 | 2243 | TRUE | 2 |
| orf1b | 1031 | 8007 | 2243 | TRUE | 2 |
| orf1b | 1032 | 8007 | 2068 | FALSE | 3 |
| orf1b | 1033 | 8007 | 2068 | FALSE | 2 |
| orf1b | 1034 | 8007 | 1507 | TRUE | 3 |
| orf1b | 1035 | 8007 | 1507 | TRUE | 3 |
| orf1b | 1036 | 8007 | 1168 | FALSE | 3 |
| orf1b | 1037 | 8007 | 1168 | TRUE | 4 |
| orf1b | 1038 | 6386 | 0 | TRUE | 2 |
| orf1b | 1039 | 6386 | 0 | FALSE | 3 |
| orf1b | 1040 | 6386 | 0 | TRUE | 3 |
| orf1b | 1041 | 6386 | 0 | TRUE | 3 |
| orf1b | 1049 | 1842 | 1534 | FALSE | 2 |
| orf1b | 1050 | 1842 | 1534 | TRUE | 3 |
| orf1b | 1051 | 3626 | 4662 | TRUE | 3 |
| orf1b | 1052 | 3626 | 4662 | FALSE | 6 |
| orf1b | 1053 | 3626 | 4662 | TRUE | 3 |
| orf1b | 1054 | 3626 | 4662 | TRUE | 1 |
| orf1b | 1055 | 3176 | 4662 | TRUE | 4 |
| orf1b | 1056 | 3176 | 4662 | TRUE | 1 |
| orf1b | 1057 | 4629 | 3408 | TRUE | 3 |
| orf1b | 1058 | 4629 | 3408 | FALSE | 4 |
| orf1b | 1059 | 6285 | 3669 | TRUE | 3 |
| orf1b | 1060 | 6285 | 3669 | FALSE | 4 |
| orf1b | 1061 | 6285 | 3398 | TRUE | 4 |
| orf1b | 1062 | 6285 | 3398 | TRUE | 3 |
| orf1b | 1063 | 6285 | 4364 | TRUE | 3 |
| orf1b | 1064 | 7766 | 4718 | TRUE | 2 |
| orf1b | 1065 | 12590 | 5992 | TRUE | 5 |
| orf1b | 1066 | 14385 | 5945 | TRUE | 3 |
| orf1b | 1067 | 14385 | 5945 | TRUE | 3 |
| orf1b | 1068 | 14385 | 5945 | TRUE | 2 |
| orf1b | 1069 | 14385 | 5945 | TRUE | 3 |
| orf1b | 1070 | 16244 | 5945 | TRUE | 2 |
| orf1b | 1071 | 16244 | 5730 | TRUE | 2 |
| orf1b | 1072 | 16968 | 6664 | TRUE | 5 |
| orf1b | 1073 | 16968 | 6182 | FALSE | 2 |
| orf1b | 1074 | 19825 | 6648 | FALSE | 3 |
| orf1b | 1075 | 19825 | 6648 | FALSE | 4 |
| orf1b | 1076 | 19825 | 7382 | TRUE | 4 |
| orf1b | 1077 | 18500 | 6097 | TRUE | 2 |
| orf1b | 1078 | 18500 | 5347 | FALSE | 4 |
| orf1b | 1079 | 16931 | 5403 | TRUE | 3 |
| orf1b | 1080 | 13072 | 3836 | TRUE | 5 |
| orf1b | 1081 | 13072 | 3836 | TRUE | 3 |
| orf1b | 1082 | 13072 | 3836 | FALSE | 3 |
| orf1b | 1083 | 11032 | 3522 | TRUE | 5 |

| | | | | | |
|---|---|---|---|---|---|
| orf1b | 1084 | 11032 | 3522 | TRUE | 4 |
| orf1b | 1085 | 10290 | 3736 | TRUE | 2 |
| orf1b | 1086 | 10290 | 3752 | TRUE | 2 |
| orf1b | 1087 | 10290 | 3752 | TRUE | 2 |
| orf1b | 1088 | 10290 | 3964 | TRUE | 3 |
| orf1b | 1089 | 10290 | 3964 | TRUE | 3 |
| orf1b | 1090 | 10290 | 3964 | TRUE | 5 |
| orf1b | 1091 | 10458 | 3761 | FALSE | 4 |
| orf1b | 1092 | 10001 | 4302 | TRUE | 4 |
| orf1b | 1093 | 10001 | 4302 | TRUE | 2 |
| orf1b | 1094 | 8498 | 3087 | TRUE | 1 |
| orf1b | 1095 | 8498 | 3087 | FALSE | 2 |
| orf1b | 1096 | 8498 | 3087 | TRUE | 2 |
| orf1b | 1097 | 8498 | 3087 | FALSE | 4 |
| orf1b | 1098 | 9693 | 3780 | TRUE | 2 |
| orf1b | 1099 | 9693 | 3780 | TRUE | 2 |
| orf1b | 1100 | 11469 | 1934 | FALSE | 2 |
| orf1b | 1101 | 11469 | 1934 | TRUE | 2 |
| orf1b | 1102 | 11469 | 1934 | TRUE | 4 |
| orf1b | 1103 | 11469 | 1934 | TRUE | 2 |
| orf1b | 1104 | 11469 | 1934 | TRUE | 4 |
| orf1b | 1105 | 11469 | 1934 | TRUE | 3 |
| orf1b | 1113 | 1134 | 0 | TRUE | 2 |
| orf1b | 1114 | 1134 | 0 | TRUE | 2 |
| orf1b | 1115 | 1134 | 0 | TRUE | 1 |
| orf1b | 1116 | 1134 | 0 | TRUE | 3 |
| orf1b | 1117 | 1134 | 3170 | TRUE | 2 |
| orf1b | 1118 | 1134 | 3170 | TRUE | 3 |
| orf1b | 1119 | 1134 | 3180 | TRUE | 4 |
| orf1b | 1120 | 1134 | 3180 | FALSE | 2 |
| orf1b | 1121 | 1134 | 3180 | TRUE | 2 |
| orf1b | 1122 | 1134 | 3180 | TRUE | 2 |
| orf1b | 1123 | 1134 | 3180 | TRUE | 3 |
| orf1b | 1124 | 1134 | 3180 | TRUE | 1 |
| orf1b | 1125 | 1134 | 3180 | TRUE | 2 |
| orf1b | 1126 | 1134 | 3180 | TRUE | 3 |
| orf1b | 1127 | 1134 | 3180 | TRUE | 3 |
| orf1b | 1128 | 0 | 3180 | TRUE | 0 |
| orf1b | 1129 | 0 | 3180 | TRUE | 6 |
| orf1b | 1130 | 0 | 3180 | TRUE | 5 |
| orf1b | 1131 | 0 | 3180 | TRUE | 5 |
| orf1b | 1132 | 0 | 3190 | TRUE | 5 |
| orf1b | 1133 | 0 | 3190 | TRUE | 1 |
| orf1b | 1143 | 1610 | 1410 | FALSE | 0 |
| orf1b | 1144 | 1610 | 1410 | FALSE | 3 |
| orf1b | 1145 | 1610 | 1410 | FALSE | 4 |
| orf1b | 1146 | 1610 | 1410 | FALSE | 4 |
| orf1b | 1147 | 1610 | 1410 | FALSE | 3 |
| orf1b | 1148 | 1610 | 1410 | FALSE | 0 |
| orf1b | 1149 | 1610 | 1410 | FALSE | 3 |
| orf1b | 1150 | 1610 | 1410 | FALSE | 2 |
| orf1b | 1151 | 1610 | 1410 | FALSE | 2 |
| orf1b | 1152 | 1610 | 1410 | FALSE | 2 |
| orf1b | 1153 | 1610 | 1410 | FALSE | 2 |
| orf1b | 1154 | 1610 | 1410 | FALSE | 3 |
| orf1b | 1155 | 1610 | 1410 | FALSE | 2 |
| orf1b | 1156 | 1610 | 1410 | FALSE | 3 |
| orf1b | 1157 | 1610 | 1410 | FALSE | 3 |
| orf1b | 1161 | 1611 | 0 | FALSE | 2 |
| orf1b | 1162 | 1611 | 0 | FALSE | 2 |
| orf1b | 1163 | 2855 | 0 | FALSE | 3 |
| orf1b | 1164 | 2855 | 0 | FALSE | 3 |

| | | | | | |
|---|---|---|---|---|---|
| orf1b | 1165 | 4677 | 0 | FALSE | 2 |
| orf1b | 1166 | 4677 | 0 | FALSE | 2 |
| orf1b | 1167 | 7223 | 4039 | FALSE | 4 |
| orf1b | 1168 | 7223 | 4039 | FALSE | 4 |
| orf1b | 1169 | 7223 | 4039 | FALSE | 1 |
| orf1b | 1170 | 7223 | 4039 | FALSE | 2 |
| orf1b | 1171 | 7223 | 4039 | FALSE | 2 |
| orf1b | 1172 | 7223 | 4039 | FALSE | 2 |
| orf1b | 1173 | 7223 | 3026 | FALSE | 1 |
| orf1b | 1174 | 7223 | 3026 | FALSE | 3 |
| orf1b | 1175 | 7223 | 3026 | FALSE | 2 |
| orf1b | 1176 | 8346 | 3026 | FALSE | 2 |
| orf1b | 1177 | 6385 | 2579 | FALSE | 3 |
| orf1b | 1178 | 7548 | 2579 | FALSE | 1 |
| orf1b | 1179 | 7548 | 2579 | FALSE | 3 |
| orf1b | 1180 | 7640 | 2579 | FALSE | 1 |
| orf1b | 1181 | 7640 | 2579 | FALSE | 3 |
| orf1b | 1182 | 1482 | 1119 | FALSE | 2 |
| orf1b | 1183 | 1482 | 1157 | TRUE | 5 |
| orf1b | 1184 | 1482 | 1157 | FALSE | 5 |
| orf1b | 1185 | 1482 | 1157 | TRUE | 2 |
| orf1b | 1186 | 1482 | 1157 | TRUE | 2 |
| orf1b | 1187 | 1482 | 1157 | TRUE | 2 |
| orf1b | 1188 | 1482 | 1235 | TRUE | 0 |
| orf1b | 1189 | 1482 | 1235 | TRUE | 2 |
| orf1b | 1190 | 1482 | 1235 | FALSE | 2 |
| orf1b | 1191 | 1368 | 1235 | FALSE | 3 |
| orf1b | 1192 | 1140 | 1233 | FALSE | 0 |
| orf1b | 1193 | 1200 | 1233 | TRUE | 1 |
| orf1b | 1194 | 1200 | 1233 | TRUE | 5 |
| orf1b | 1195 | 1662 | 1233 | TRUE | 3 |
| orf1b | 1196 | 1662 | 1233 | TRUE | 1 |
| orf1b | 1197 | 7446 | 1233 | TRUE | 6 |
| orf1b | 1198 | 7446 | 0 | TRUE | 1 |
| orf1b | 1199 | 6322 | 0 | TRUE | 2 |
| orf1b | 1200 | 6322 | 0 | TRUE | 3 |
| orf1b | 1201 | 6322 | 0 | TRUE | 3 |
| orf1b | 1202 | 6322 | 0 | FALSE | 2 |
| orf1b | 1203 | 6322 | 0 | TRUE | 2 |
| orf1b | 1204 | 6322 | 0 | TRUE | 4 |
| orf1b | 1205 | 5294 | 1172 | TRUE | 2 |
| orf1b | 1206 | 5887 | 1172 | TRUE | 3 |
| orf1b | 1207 | 5887 | 1172 | TRUE | 3 |
| orf1b | 1208 | 6658 | 1172 | TRUE | 3 |
| orf1b | 1209 | 6658 | 1172 | TRUE | 2 |
| orf1b | 1210 | 7473 | 1172 | TRUE | 4 |
| orf1b | 1211 | 7473 | 1172 | TRUE | 3 |
| orf1b | 1212 | 3142 | 1172 | TRUE | 2 |
| orf1b | 1213 | 3142 | 1172 | TRUE | 2 |
| orf1b | 1214 | 3580 | 1172 | TRUE | 1 |
| orf1b | 1215 | 3580 | 1940 | TRUE | 2 |
| orf1b | 1216 | 3580 | 1940 | TRUE | 1 |
| orf1b | 1217 | 3580 | 2055 | TRUE | 2 |
| orf1b | 1218 | 3580 | 2055 | TRUE | 4 |
| orf1b | 1219 | 3580 | 2055 | TRUE | 3 |
| orf1b | 1220 | 0 | 2497 | TRUE | 3 |
| orf1b | 1221 | 0 | 2497 | TRUE | 1 |
| orf1b | 1222 | 0 | 2497 | TRUE | 0 |
| orf1b | 1223 | 0 | 2497 | TRUE | 3 |
| orf1b | 1224 | 0 | 2497 | TRUE | 3 |
| orf1b | 1225 | 0 | 2497 | TRUE | 4 |
| orf1b | 1226 | 0 | 2497 | TRUE | 4 |

| | | | | | |
|---|---|---|---|---|---|
| orf1b | 1227 | 0 | 2497 | TRUE | 1 |
| orf1b | 1228 | 0 | 2497 | TRUE | 2 |
| orf1b | 1229 | 0 | 2497 | TRUE | 2 |
| orf1b | 1230 | 0 | 2284 | TRUE | 1 |
| orf1b | 1231 | 0 | 2284 | FALSE | 2 |
| orf1b | 1247 | 1915 | 0 | FALSE | 2 |
| orf1b | 1248 | 1915 | 0 | FALSE | 2 |
| orf1b | 1249 | 3317 | 3470 | FALSE | 3 |
| orf1b | 1250 | 3317 | 3470 | FALSE | 3 |
| orf1b | 1251 | 3317 | 3470 | TRUE | 0 |
| orf1b | 1252 | 3317 | 3470 | TRUE | 0 |
| orf1b | 1253 | 3317 | 3470 | TRUE | 3 |
| orf1b | 1254 | 3317 | 3470 | TRUE | 4 |
| orf1b | 1255 | 3317 | 3470 | FALSE | 1 |
| orf1b | 1256 | 3317 | 3470 | TRUE | 0 |
| orf1b | 1257 | 2874 | 3470 | TRUE | 1 |
| orf1b | 1258 | 2874 | 3470 | TRUE | 2 |
| orf1b | 1259 | 2874 | 3470 | TRUE | 2 |
| orf1b | 1260 | 2874 | 3470 | TRUE | 3 |
| orf1b | 1261 | 2874 | 3470 | TRUE | 2 |
| orf1b | 1262 | 3514 | 3470 | TRUE | 3 |
| orf1b | 1263 | 3514 | 2411 | FALSE | 4 |
| orf1b | 1264 | 1103 | 1352 | FALSE | 4 |
| orf1b | 1265 | 1103 | 1352 | FALSE | 2 |
| orf1b | 1266 | 1103 | 1352 | FALSE | 1 |
| orf1b | 1267 | 1103 | 1352 | FALSE | 1 |
| orf1b | 1268 | 1103 | 1352 | FALSE | 1 |
| orf1b | 1269 | 1103 | 1352 | FALSE | 3 |
| orf1b | 1270 | 1103 | 1352 | FALSE | 2 |
| orf1b | 1271 | 1103 | 1352 | FALSE | 2 |
| orf1b | 1272 | 0 | 1352 | TRUE | 2 |
| orf1b | 1273 | 1459 | 1352 | TRUE | 3 |
| orf1b | 1274 | 1459 | 1352 | TRUE | 2 |
| orf1b | 1275 | 1459 | 1352 | TRUE | 0 |
| orf1b | 1276 | 1459 | 1352 | FALSE | 4 |
| orf1b | 1277 | 1459 | 1352 | TRUE | 3 |
| orf1b | 1278 | 1459 | 0 | TRUE | 1 |
| orf1b | 1279 | 1459 | 0 | TRUE | 5 |
| orf1b | 1280 | 1459 | 0 | TRUE | 0 |
| orf1b | 1281 | 1459 | 0 | TRUE | 2 |
| orf1b | 1282 | 1459 | 0 | TRUE | 3 |
| orf1b | 1283 | 1459 | 0 | FALSE | 0 |
| orf1b | 1284 | 1459 | 0 | FALSE | 0 |
| orf1b | 1285 | 1360 | 0 | TRUE | 3 |
| orf1b | 1286 | 1360 | 0 | TRUE | 2 |
| orf1b | 1287 | 1658 | 2760 | TRUE | 2 |
| orf1b | 1288 | 1691 | 2760 | TRUE | 5 |
| orf1b | 1289 | 1691 | 2760 | TRUE | 2 |
| orf1b | 1290 | 1691 | 2760 | TRUE | 1 |
| orf1b | 1291 | 1691 | 4192 | TRUE | 3 |
| orf1b | 1292 | 1691 | 4192 | TRUE | 2 |
| orf1b | 1293 | 1737 | 4192 | TRUE | 2 |
| orf1b | 1294 | 1737 | 4192 | TRUE | 2 |
| orf1b | 1295 | 1669 | 3243 | TRUE | 3 |
| orf1b | 1296 | 1669 | 3243 | TRUE | 2 |
| orf1b | 1297 | 1799 | 3067 | TRUE | 2 |
| orf1b | 1298 | 1799 | 3067 | TRUE | 2 |
| orf1b | 1299 | 1799 | 3067 | TRUE | 3 |
| orf1b | 1300 | 2063 | 3067 | TRUE | 2 |
| orf1b | 1301 | 2063 | 3067 | TRUE | 4 |
| orf1b | 1302 | 1928 | 3144 | TRUE | 3 |
| orf1b | 1303 | 1928 | 3144 | TRUE | 2 |

| | | | | | |
|---|---|---|---|---|---|
| orf1b | 1304 | 1928 | 3144 | TRUE | 2 |
| orf1b | 1305 | 1928 | 3144 | TRUE | 2 |
| orf1b | 1306 | 1928 | 2317 | TRUE | 3 |
| orf1b | 1307 | 1928 | 2317 | TRUE | 0 |
| orf1b | 1308 | 1906 | 2317 | TRUE | 2 |
| orf1b | 1309 | 1906 | 2317 | TRUE | 1 |
| orf1b | 1310 | 2383 | 2804 | TRUE | 3 |
| orf1b | 1311 | 2068 | 2804 | TRUE | 2 |
| orf1b | 1312 | 1439 | 0 | TRUE | 3 |
| orf1b | 1313 | 1439 | 0 | TRUE | 0 |
| orf1b | 1314 | 1439 | 0 | TRUE | 1 |
| orf1b | 1315 | 1439 | 0 | TRUE | 4 |
| orf1b | 1316 | 1439 | 0 | TRUE | 3 |
| orf1b | 1317 | 1439 | 0 | TRUE | 4 |
| orf1b | 1318 | 1439 | 0 | TRUE | 3 |
| orf1b | 1319 | 1439 | 0 | TRUE | 0 |
| orf1b | 1320 | 1439 | 0 | TRUE | 4 |
| orf1b | 1321 | 1439 | 0 | TRUE | 1 |
| orf1b | 1322 | 1439 | 0 | TRUE | 0 |
| orf1b | 1323 | 2160 | 0 | FALSE | 4 |
| orf1b | 1324 | 2160 | 0 | TRUE | 2 |
| orf1b | 1325 | 2160 | 0 | TRUE | 5 |
| orf1b | 1326 | 2881 | 0 | TRUE | 6 |
| orf1b | 1327 | 2881 | 0 | TRUE | 2 |
| orf1b | 1328 | 2881 | 0 | TRUE | 2 |
| orf1b | 1329 | 2937 | 0 | TRUE | 4 |
| orf1b | 1330 | 2937 | 0 | TRUE | 3 |
| orf1b | 1331 | 2558 | 0 | TRUE | 3 |
| orf1b | 1332 | 2558 | 0 | TRUE | 6 |
| orf1b | 1333 | 2274 | 0 | TRUE | 2 |
| orf1b | 1334 | 2274 | 0 | TRUE | 3 |
| orf1b | 1335 | 2036 | 0 | TRUE | 1 |
| orf1b | 1336 | 2036 | 0 | TRUE | 4 |
| orf1b | 1337 | 2036 | 0 | TRUE | 2 |
| orf1b | 1338 | 1825 | 0 | TRUE | 5 |
| orf1b | 1339 | 1825 | 0 | TRUE | 2 |
| orf1b | 1340 | 1825 | 0 | TRUE | 2 |
| orf1b | 1341 | 3474 | 1999 | TRUE | 6 |
| orf1b | 1342 | 3474 | 1999 | TRUE | 2 |
| orf1b | 1343 | 3474 | 1999 | TRUE | 3 |
| orf1b | 1344 | 3554 | 1999 | FALSE | 2 |
| orf1b | 1345 | 3554 | 1999 | FALSE | 5 |
| orf1b | 1346 | 3905 | 1999 | FALSE | 1 |
| orf1b | 1347 | 3905 | 1999 | FALSE | 1 |
| orf1b | 1348 | 4525 | 1999 | FALSE | 4 |
| orf1b | 1349 | 4525 | 1999 | FALSE | 2 |
| orf1b | 1350 | 5672 | 1999 | FALSE | 5 |
| orf1b | 1351 | 5672 | 1999 | FALSE | 2 |
| orf1b | 1352 | 5672 | 1999 | FALSE | 4 |
| orf1b | 1353 | 5672 | 1999 | FALSE | 2 |
| orf1b | 1354 | 5672 | 1999 | FALSE | 2 |
| orf1b | 1355 | 5672 | 1999 | FALSE | 0 |
| orf1b | 1357 | 1273 | 0 | TRUE | 4 |
| orf1b | 1358 | 1273 | 0 | TRUE | 2 |
| orf1b | 1359 | 1273 | 0 | FALSE | 5 |
| orf1b | 1360 | 1273 | 0 | TRUE | 2 |
| orf1b | 1361 | 1273 | 0 | TRUE | 7 |
| orf1b | 1362 | 1273 | 0 | TRUE | 3 |
| orf1b | 1363 | 2586 | 0 | TRUE | 4 |
| orf1b | 1364 | 2586 | 0 | TRUE | 3 |
| orf1b | 1365 | 2586 | 0 | TRUE | 4 |
| orf1b | 1366 | 2586 | 0 | TRUE | 2 |

| | | | | | |
|---|---|---|---|---|---|
| orf1b | 1367 | 2586 | 0 | TRUE | 3 |
| orf1b | 1368 | 2586 | 0 | TRUE | 4 |
| orf1b | 1369 | 2586 | 0 | TRUE | 3 |
| orf1b | 1370 | 2586 | 0 | FALSE | 3 |
| orf1b | 1371 | 2586 | 0 | TRUE | 1 |
| orf1b | 1372 | 3242 | 0 | TRUE | 3 |
| orf1b | 1373 | 3242 | 0 | TRUE | 3 |
| orf1b | 1374 | 3242 | 0 | TRUE | 2 |
| orf1b | 1375 | 3242 | 0 | TRUE | 3 |
| orf1b | 1376 | 3242 | 0 | TRUE | 1 |
| orf1b | 1377 | 3242 | 0 | FALSE | 5 |
| orf1b | 1383 | 2042 | 0 | TRUE | 2 |
| orf1b | 1384 | 2042 | 0 | TRUE | 3 |
| orf1b | 1385 | 2508 | 0 | FALSE | 1 |
| orf1b | 1386 | 2508 | 0 | FALSE | 4 |
| orf1b | 1387 | 2508 | 0 | FALSE | 3 |
| orf1b | 1388 | 2508 | 0 | TRUE | 5 |
| orf1b | 1389 | 2508 | 0 | TRUE | 4 |
| orf1b | 1390 | 2508 | 0 | TRUE | 2 |
| orf1b | 1391 | 2508 | 0 | TRUE | 2 |
| orf1b | 1392 | 2508 | 0 | TRUE | 5 |
| orf1b | 1393 | 2520 | 0 | FALSE | 2 |
| orf1b | 1394 | 2520 | 0 | TRUE | 2 |
| orf1b | 1395 | 2305 | 0 | FALSE | 0 |
| orf1b | 1396 | 2305 | 0 | TRUE | 2 |
| orf1b | 1397 | 2305 | 0 | FALSE | 3 |
| orf1b | 1398 | 2393 | 0 | FALSE | 2 |
| orf1b | 1399 | 2068 | 2727 | TRUE | 2 |
| orf1b | 1400 | 1766 | 2727 | FALSE | 4 |
| orf1b | 1401 | 1766 | 9301 | TRUE | 4 |
| orf1b | 1402 | 1766 | 9301 | TRUE | 3 |
| orf1b | 1403 | 1766 | 9301 | TRUE | 3 |
| orf1b | 1404 | 1766 | 9301 | TRUE | 4 |
| orf1b | 1405 | 1766 | 7493 | TRUE | 3 |
| orf1b | 1406 | 1766 | 7493 | TRUE | 4 |
| orf1b | 1407 | 1766 | 6085 | FALSE | 3 |
| orf1b | 1408 | 1377 | 6085 | TRUE | 2 |
| orf1b | 1409 | 1377 | 6085 | TRUE | 4 |
| orf1b | 1410 | 1094 | 6085 | FALSE | 6 |
| orf1b | 1411 | 1094 | 6085 | TRUE | 3 |
| orf1b | 1412 | 1094 | 6085 | TRUE | 2 |
| orf1b | 1413 | 1094 | 6085 | FALSE | 4 |
| orf1b | 1414 | 0 | 7204 | FALSE | 2 |
| orf1b | 1415 | 0 | 5980 | FALSE | 2 |
| orf1b | 1416 | 0 | 2682 | FALSE | 2 |
| orf1b | 1417 | 0 | 4333 | FALSE | 4 |
| orf1b | 1418 | 0 | 4333 | FALSE | 3 |
| orf1b | 1419 | 0 | 4333 | FALSE | 2 |
| orf1b | 1420 | 0 | 4484 | FALSE | 1 |
| orf1b | 1421 | 0 | 3814 | FALSE | 4 |
| orf1b | 1422 | 0 | 4466 | FALSE | 3 |
| orf1b | 1423 | 0 | 4466 | FALSE | 3 |
| orf1b | 1424 | 0 | 4466 | FALSE | 4 |
| orf1b | 1425 | 0 | 4466 | FALSE | 4 |
| orf1b | 1426 | 0 | 4466 | TRUE | 3 |
| orf1b | 1427 | 0 | 4466 | TRUE | 3 |
| orf1b | 1428 | 0 | 4466 | FALSE | 3 |
| orf1b | 1429 | 0 | 4466 | FALSE | 4 |
| orf1b | 1430 | 0 | 5544 | TRUE | 3 |
| orf1b | 1431 | 0 | 5544 | TRUE | 3 |
| orf1b | 1432 | 0 | 1804 | FALSE | 3 |
| orf1b | 1433 | 0 | 1804 | FALSE | 3 |

| | | | | | |
|---|---|---|---|---|---|
| orf1b | 1434 | 0 | 1804 | TRUE | 4 |
| orf1b | 1435 | 0 | 1804 | TRUE | 1 |
| orf1b | 1439 | 1327 | 0 | FALSE | 2 |
| orf1b | 1440 | 1327 | 0 | TRUE | 1 |
| orf1b | 1441 | 1327 | 0 | TRUE | 4 |
| orf1b | 1442 | 1327 | 0 | TRUE | 2 |
| orf1b | 1443 | 1327 | 0 | TRUE | 1 |
| orf1b | 1444 | 1327 | 0 | TRUE | 3 |
| orf1b | 1445 | 1327 | 0 | TRUE | 4 |
| orf1b | 1446 | 1327 | 0 | TRUE | 2 |
| orf1b | 1447 | 1327 | 0 | TRUE | 4 |
| orf1b | 1448 | 1327 | 0 | TRUE | 2 |
| orf1b | 1449 | 1327 | 0 | TRUE | 2 |
| orf1b | 1450 | 1327 | 0 | TRUE | 3 |
| orf1b | 1451 | 1327 | 0 | TRUE | 3 |
| orf1b | 1452 | 1327 | 0 | TRUE | 3 |
| orf1b | 1453 | 1327 | 0 | TRUE | 2 |
| orf1b | 1455 | 1286 | 0 | TRUE | 5 |
| orf1b | 1456 | 1286 | 0 | TRUE | 1 |
| orf1b | 1457 | 1286 | 0 | TRUE | 4 |
| orf1b | 1458 | 1286 | 0 | TRUE | 1 |
| orf1b | 1459 | 1286 | 0 | TRUE | 2 |
| orf1b | 1460 | 1286 | 0 | TRUE | 5 |
| orf1b | 1461 | 1286 | 0 | TRUE | 5 |
| orf1b | 1462 | 1286 | 0 | TRUE | 2 |
| orf1b | 1463 | 3442 | 0 | TRUE | 2 |
| orf1b | 1464 | 3442 | 0 | TRUE | 2 |
| orf1b | 1465 | 3442 | 0 | TRUE | 3 |
| orf1b | 1466 | 3442 | 0 | TRUE | 5 |
| orf1b | 1467 | 3442 | 0 | TRUE | 2 |
| orf1b | 1468 | 3442 | 0 | TRUE | 2 |
| orf1b | 1469 | 3442 | 0 | TRUE | 3 |
| orf1b | 1470 | 5598 | 0 | TRUE | 3 |
| orf1b | 1471 | 5598 | 0 | TRUE | 2 |
| orf1b | 1472 | 5598 | 0 | TRUE | 3 |
| orf1b | 1473 | 3659 | 0 | TRUE | 4 |
| orf1b | 1474 | 3659 | 0 | TRUE | 4 |
| orf1b | 1475 | 3659 | 0 | TRUE | 2 |
| orf1b | 1476 | 3659 | 0 | TRUE | 4 |
| orf1b | 1477 | 3659 | 0 | TRUE | 2 |
| orf1b | 1478 | 1720 | 0 | TRUE | 2 |
| orf1b | 1479 | 1720 | 0 | TRUE | 1 |
| orf1b | 1480 | 1720 | 0 | TRUE | 4 |
| orf1b | 1481 | 1720 | 0 | TRUE | 4 |
| orf1b | 1482 | 1720 | 0 | TRUE | 2 |
| orf1b | 1483 | 1720 | 0 | TRUE | 4 |
| orf1b | 1484 | 1720 | 0 | TRUE | 2 |
| orf1b | 1485 | 1499 | 0 | TRUE | 3 |
| orf1b | 1486 | 1499 | 0 | TRUE | 0 |
| orf1b | 1487 | 1499 | 0 | TRUE | 6 |
| orf1b | 1488 | 1057 | 0 | TRUE | 3 |
| orf1b | 1489 | 1057 | 1146 | TRUE | 4 |
| orf1b | 1490 | 1057 | 1146 | TRUE | 2 |
| orf1b | 1491 | 1057 | 1146 | TRUE | 5 |
| orf1b | 1492 | 1057 | 1146 | TRUE | 2 |
| orf1b | 1493 | 1057 | 1146 | TRUE | 3 |
| orf1b | 1494 | 1057 | 1146 | TRUE | 4 |
| orf1b | 1495 | 1057 | 1146 | TRUE | 2 |
| orf1b | 1496 | 1057 | 1146 | TRUE | 3 |
| orf1b | 1497 | 1057 | 1146 | TRUE | 4 |
| orf1b | 1498 | 1057 | 1146 | TRUE | 2 |
| orf1b | 1499 | 1057 | 1146 | TRUE | 3 |

| | | | | | |
|---|---|---|---|---|---|
| orf1b | 1500 | 0 | 1146 | TRUE | 3 |
| orf1b | 1501 | 1174 | 1146 | FALSE | 4 |
| orf1b | 1502 | 1174 | 1146 | FALSE | 5 |
| orf1b | 1503 | 1174 | 1146 | FALSE | 1 |
| orf1b | 1504 | 1174 | 0 | FALSE | 3 |
| orf1b | 1505 | 1174 | 0 | FALSE | 3 |
| orf1b | 1506 | 1174 | 0 | FALSE | 2 |
| orf1b | 1507 | 1174 | 0 | FALSE | 1 |
| orf1b | 1508 | 1174 | 0 | FALSE | 0 |
| orf1b | 1509 | 1174 | 0 | FALSE | 4 |
| orf1b | 1510 | 1174 | 0 | FALSE | 3 |
| orf1b | 1511 | 1174 | 0 | FALSE | 2 |
| orf1b | 1512 | 1174 | 0 | FALSE | 4 |
| orf1b | 1513 | 1174 | 0 | FALSE | 4 |
| orf1b | 1514 | 1174 | 0 | FALSE | 5 |
| orf1b | 1515 | 1174 | 0 | FALSE | 3 |
| orf1b | 1519 | 1593 | 0 | FALSE | 4 |
| orf1b | 1520 | 1593 | 0 | FALSE | 2 |
| orf1b | 1521 | 1593 | 0 | FALSE | 2 |
| orf1b | 1522 | 1593 | 0 | FALSE | 2 |
| orf1b | 1523 | 1593 | 0 | FALSE | 3 |
| orf1b | 1524 | 1593 | 0 | FALSE | 5 |
| orf1b | 1525 | 1593 | 0 | TRUE | 5 |
| orf1b | 1526 | 1593 | 0 | FALSE | 1 |
| orf1b | 1527 | 1593 | 0 | FALSE | 3 |
| orf1b | 1528 | 1593 | 0 | FALSE | 3 |
| orf1b | 1529 | 1593 | 0 | FALSE | 3 |
| orf1b | 1530 | 1593 | 0 | FALSE | 2 |
| orf1b | 1531 | 1593 | 0 | FALSE | 3 |
| orf1b | 1532 | 1593 | 0 | TRUE | 3 |
| orf1b | 1533 | 1593 | 0 | FALSE | 3 |
| orf1b | 1537 | 1340 | 0 | TRUE | 6 |
| orf1b | 1538 | 1340 | 0 | TRUE | 2 |
| orf1b | 1539 | 1340 | 2031 | TRUE | 4 |
| orf1b | 1540 | 1340 | 2031 | TRUE | 2 |
| orf1b | 1541 | 1271 | 3236 | TRUE | 4 |
| orf1b | 1542 | 1271 | 3236 | TRUE | 1 |
| orf1b | 1543 | 1271 | 3236 | TRUE | 5 |
| orf1b | 1544 | 1271 | 3236 | TRUE | 5 |
| orf1b | 1545 | 1271 | 3236 | FALSE | 1 |
| orf1b | 1546 | 1271 | 3236 | FALSE | 3 |
| orf1b | 1547 | 1271 | 3236 | FALSE | 5 |
| orf1b | 1548 | 1271 | 3236 | FALSE | 3 |
| orf1b | 1549 | 1271 | 3236 | FALSE | 2 |
| orf1b | 1550 | 1271 | 3236 | FALSE | 3 |
| orf1b | 1551 | 1271 | 3236 | FALSE | 2 |
| orf1b | 1552 | 1201 | 3236 | FALSE | 3 |
| orf1b | 1553 | 1201 | 3236 | FALSE | 4 |
| orf1b | 1554 | 1201 | 4440 | FALSE | 4 |
| orf1b | 1555 | 1201 | 4440 | FALSE | 2 |
| orf1b | 1559 | 0 | 2869 | FALSE | 3 |
| orf1b | 1560 | 0 | 2869 | FALSE | 3 |
| orf1b | 1561 | 0 | 5578 | FALSE | 4 |
| orf1b | 1562 | 0 | 5578 | FALSE | 1 |
| orf1b | 1563 | 0 | 5578 | FALSE | 3 |
| orf1b | 1564 | 0 | 5578 | FALSE | 4 |
| orf1b | 1565 | 0 | 5578 | FALSE | 2 |
| orf1b | 1566 | 0 | 5578 | FALSE | 4 |
| orf1b | 1567 | 0 | 5578 | FALSE | 2 |
| orf1b | 1568 | 0 | 5578 | FALSE | 4 |
| orf1b | 1569 | 0 | 5578 | FALSE | 3 |
| orf1b | 1570 | 0 | 5578 | FALSE | 3 |

| | | | | | |
|---|---|---|---|---|---|
| orf1b | 1571 | 0 | 5578 | FALSE | 5 |
| orf1b | 1572 | 0 | 5578 | FALSE | 3 |
| orf1b | 1573 | 0 | 5578 | FALSE | 4 |
| orf1b | 1574 | 0 | 8287 | FALSE | 3 |
| orf1b | 1575 | 0 | 8287 | FALSE | 2 |
| orf1b | 1581 | 0 | 1282 | FALSE | 2 |
| orf1b | 1582 | 0 | 1282 | FALSE | 4 |
| orf1b | 1583 | 0 | 1282 | FALSE | 4 |
| orf1b | 1584 | 0 | 1282 | FALSE | 1 |
| orf1b | 1585 | 0 | 1282 | FALSE | 2 |
| orf1b | 1586 | 0 | 1282 | FALSE | 2 |
| orf1b | 1587 | 1106 | 1176 | FALSE | 2 |
| orf1b | 1588 | 1106 | 1176 | FALSE | 1 |
| orf1b | 1589 | 1511 | 1176 | FALSE | 4 |
| orf1b | 1590 | 1511 | 1176 | FALSE | 3 |
| orf1b | 1591 | 1511 | 1385 | TRUE | 2 |
| orf1b | 1592 | 1511 | 1385 | TRUE | 1 |
| orf1b | 1593 | 1511 | 1385 | TRUE | 1 |
| orf1b | 1594 | 1511 | 1385 | TRUE | 4 |
| orf1b | 1595 | 2071 | 1385 | TRUE | 3 |
| orf1b | 1596 | 2071 | 1437 | TRUE | 6 |
| orf1b | 1597 | 2071 | 1437 | TRUE | 1 |
| orf1b | 1598 | 2071 | 1437 | TRUE | 3 |
| orf1b | 1599 | 2071 | 1666 | TRUE | 4 |
| orf1b | 1600 | 2071 | 1666 | TRUE | 5 |
| orf1b | 1601 | 9999 | 4623 | TRUE | 5 |
| orf1b | 1602 | 11269 | 5216 | FALSE | 1 |
| orf1b | 1603 | 11269 | 5216 | FALSE | 4 |
| orf1b | 1604 | 12828 | 5216 | TRUE | 4 |
| orf1b | 1605 | 11150 | 5216 | TRUE | 5 |
| orf1b | 1606 | 11150 | 5898 | TRUE | 2 |
| orf1b | 1607 | 11150 | 5898 | TRUE | 4 |
| orf1b | 1608 | 11150 | 5898 | TRUE | 5 |
| orf1b | 1609 | 9898 | 5898 | TRUE | 4 |
| orf1b | 1610 | 10856 | 5898 | FALSE | 2 |
| orf1b | 1611 | 10856 | 5898 | TRUE | 3 |
| orf1b | 1612 | 10856 | 5898 | TRUE | 3 |
| orf1b | 1613 | 10856 | 5898 | TRUE | 2 |
| orf1b | 1614 | 10856 | 6842 | FALSE | 3 |
| orf1b | 1615 | 10856 | 5745 | TRUE | 3 |
| orf1b | 1616 | 1107 | 1358 | TRUE | 7 |
| orf1b | 1617 | 1107 | 1358 | TRUE | 4 |
| orf1b | 1618 | 1107 | 1358 | TRUE | 2 |
| orf1b | 1619 | 1107 | 1358 | TRUE | 3 |
| orf1b | 1620 | 1135 | 1358 | TRUE | 4 |
| orf1b | 1621 | 3074 | 1358 | TRUE | 4 |
| orf1b | 1622 | 3074 | 1358 | TRUE | 3 |
| orf1b | 1623 | 7574 | 8034 | TRUE | 3 |
| orf1b | 1624 | 10794 | 8034 | TRUE | 3 |
| orf1b | 1625 | 10794 | 6370 | TRUE | 3 |
| orf1b | 1626 | 10794 | 6370 | TRUE | 3 |
| orf1b | 1627 | 10794 | 5691 | TRUE | 4 |
| orf1b | 1628 | 10794 | 5691 | TRUE | 3 |
| orf1b | 1629 | 10794 | 5691 | TRUE | 3 |
| orf1b | 1630 | 10794 | 6774 | FALSE | 4 |
| orf1b | 1631 | 10794 | 6774 | TRUE | 3 |
| orf1b | 1632 | 10794 | 6774 | TRUE | 3 |
| orf1b | 1633 | 10794 | 6774 | TRUE | 2 |
| orf1b | 1634 | 10794 | 6774 | TRUE | 4 |
| orf1b | 1635 | 10794 | 6774 | TRUE | 3 |
| orf1b | 1636 | 16574 | 6774 | TRUE | 3 |
| orf1b | 1637 | 8788 | 6774 | TRUE | 2 |

| | | | | | |
|---|---|---|---|---|---|
| orf1b | 1638 | 1003 | 2176 | TRUE | 3 |
| orf1b | 1639 | 1003 | 2176 | TRUE | 4 |
| orf1b | 1640 | 1003 | 2973 | TRUE | 2 |
| orf1b | 1641 | 1003 | 2973 | TRUE | 2 |
| orf1b | 1642 | 1003 | 0 | TRUE | 5 |
| orf1b | 1643 | 1887 | 1443 | TRUE | 5 |
| orf1b | 1644 | 1887 | 1443 | FALSE | 4 |
| orf1b | 1645 | 3215 | 2361 | FALSE | 2 |
| orf1b | 1646 | 3215 | 2361 | FALSE | 5 |
| orf1b | 1647 | 2872 | 2471 | FALSE | 2 |
| orf1b | 1648 | 2872 | 2471 | FALSE | 1 |
| orf1b | 1649 | 2872 | 2471 | FALSE | 3 |
| orf1b | 1650 | 2872 | 2471 | FALSE | 3 |
| orf1b | 1651 | 2872 | 2471 | FALSE | 1 |
| orf1b | 1652 | 3245 | 2471 | FALSE | 5 |
| orf1b | 1653 | 3203 | 2471 | FALSE | 1 |
| orf1b | 1654 | 3203 | 2471 | FALSE | 2 |
| orf1b | 1655 | 4641 | 2471 | FALSE | 2 |
| orf1b | 1656 | 4641 | 2471 | FALSE | 5 |
| orf1b | 1657 | 4641 | 2471 | FALSE | 3 |
| orf1b | 1658 | 4953 | 2813 | FALSE | 3 |
| orf1b | 1659 | 4953 | 2813 | TRUE | 4 |
| orf1b | 1660 | 5805 | 2800 | FALSE | 2 |
| orf1b | 1661 | 4383 | 2800 | FALSE | 5 |
| orf1b | 1662 | 5191 | 0 | TRUE | 2 |
| orf1b | 1663 | 4385 | 0 | FALSE | 2 |
| orf1b | 1664 | 4385 | 0 | FALSE | 2 |
| orf1b | 1665 | 3878 | 0 | TRUE | 4 |
| orf1b | 1666 | 3878 | 0 | FALSE | 5 |
| orf1b | 1667 | 3878 | 0 | FALSE | 3 |
| orf1b | 1668 | 4056 | 0 | TRUE | 4 |
| orf1b | 1669 | 4056 | 0 | FALSE | 3 |
| orf1b | 1670 | 1752 | 0 | TRUE | 3 |
| orf1b | 1671 | 1679 | 0 | TRUE | 2 |
| orf1b | 1672 | 1679 | 0 | TRUE | 3 |
| orf1b | 1673 | 1679 | 0 | TRUE | 2 |
| orf1b | 1674 | 1679 | 0 | TRUE | 2 |
| orf1b | 1675 | 1679 | 0 | TRUE | 5 |
| orf1b | 1676 | 1298 | 0 | TRUE | 4 |
| orf1b | 1677 | 1298 | 0 | TRUE | 3 |
| orf1b | 1678 | 1366 | 0 | TRUE | 3 |
| orf1b | 1679 | 1366 | 0 | TRUE | 2 |
| orf1b | 1680 | 1387 | 0 | TRUE | 5 |
| orf1b | 1681 | 1387 | 0 | TRUE | 2 |
| orf1b | 1682 | 1387 | 0 | TRUE | 6 |
| orf1b | 1683 | 1387 | 0 | TRUE | 2 |
| orf1b | 1684 | 1387 | 0 | TRUE | 2 |
| orf1b | 1685 | 1387 | 0 | TRUE | 1 |
| orf1b | 1695 | 9410 | 0 | TRUE | 3 |
| orf1b | 1696 | 9410 | 0 | FALSE | 3 |
| orf1b | 1697 | 9410 | 0 | TRUE | 2 |
| orf1b | 1698 | 9410 | 0 | TRUE | 3 |
| orf1b | 1699 | 9410 | 0 | TRUE | 3 |
| orf1b | 1700 | 9410 | 0 | TRUE | 1 |
| orf1b | 1701 | 9410 | 0 | TRUE | 4 |
| orf1b | 1702 | 9410 | 0 | TRUE | 3 |
| orf1b | 1703 | 9410 | 0 | TRUE | 2 |
| orf1b | 1704 | 9410 | 0 | TRUE | 3 |
| orf1b | 1705 | 9410 | 0 | FALSE | 4 |
| orf1b | 1706 | 9410 | 0 | TRUE | 6 |
| orf1b | 1707 | 5435 | 0 | FALSE | 1 |
| orf1b | 1708 | 5435 | 0 | FALSE | 3 |

| | | | | | |
|---|---|---|---|---|---|
| orf1b | 1709 | 5435 | 0 | FALSE | 3 |
| orf1b | 1710 | 1460 | 0 | TRUE | 3 |
| orf1b | 1711 | 1874 | 0 | TRUE | 3 |
| orf1b | 1712 | 1874 | 0 | FALSE | 4 |
| orf1b | 1713 | 1874 | 0 | TRUE | 3 |
| orf1b | 1714 | 1874 | 0 | TRUE | 1 |
| orf1b | 1715 | 1874 | 3416 | TRUE | 2 |
| orf1b | 1716 | 1874 | 3416 | TRUE | 2 |
| orf1b | 1717 | 1874 | 3416 | TRUE | 2 |
| orf1b | 1718 | 1874 | 3416 | TRUE | 2 |
| orf1b | 1719 | 1874 | 3416 | TRUE | 4 |
| orf1b | 1720 | 1874 | 3416 | FALSE | 1 |
| orf1b | 1721 | 1874 | 3416 | TRUE | 2 |
| orf1b | 1722 | 2287 | 3416 | TRUE | 4 |
| orf1b | 1723 | 2287 | 2654 | TRUE | 6 |
| orf1b | 1724 | 2287 | 2654 | TRUE | 3 |
| orf1b | 1725 | 2287 | 2654 | TRUE | 4 |
| orf1b | 1726 | 0 | 2654 | TRUE | 5 |
| orf1b | 1727 | 0 | 2654 | TRUE | 3 |
| orf1b | 1728 | 0 | 2654 | FALSE | 7 |
| orf1b | 1729 | 0 | 2654 | FALSE | 6 |
| orf1b | 1730 | 0 | 1131 | FALSE | 3 |
| orf1b | 1731 | 0 | 1131 | FALSE | 4 |
| orf1b | 1732 | 0 | 1131 | FALSE | 4 |
| orf1b | 1733 | 0 | 1131 | FALSE | 1 |
| orf1b | 1734 | 0 | 1131 | FALSE | 3 |
| orf1b | 1735 | 0 | 1131 | FALSE | 3 |
| orf1b | 1736 | 0 | 1131 | FALSE | 4 |
| orf1b | 1737 | 0 | 1131 | FALSE | 4 |
| orf1b | 1739 | 3180 | 0 | FALSE | 2 |
| orf1b | 1740 | 3180 | 0 | FALSE | 6 |
| orf1b | 1741 | 3180 | 0 | FALSE | 2 |
| orf1b | 1742 | 3180 | 0 | FALSE | 4 |
| orf1b | 1743 | 3180 | 0 | FALSE | 3 |
| orf1b | 1744 | 3180 | 0 | FALSE | 4 |
| orf1b | 1745 | 7067 | 0 | FALSE | 2 |
| orf1b | 1746 | 7067 | 0 | FALSE | 3 |
| orf1b | 1747 | 7067 | 0 | FALSE | 2 |
| orf1b | 1748 | 7067 | 0 | FALSE | 3 |
| orf1b | 1749 | 7067 | 0 | FALSE | 5 |
| orf1b | 1750 | 7067 | 0 | FALSE | 2 |
| orf1b | 1751 | 5275 | 0 | FALSE | 3 |
| orf1b | 1752 | 5275 | 0 | FALSE | 2 |
| orf1b | 1753 | 5275 | 0 | FALSE | 2 |
| orf1b | 1754 | 6323 | 0 | FALSE | 2 |
| orf1b | 1755 | 6323 | 0 | FALSE | 2 |
| orf1b | 1756 | 6323 | 0 | TRUE | 2 |
| orf1b | 1757 | 6323 | 0 | TRUE | 1 |
| orf1b | 1758 | 6323 | 0 | TRUE | 5 |
| orf1b | 1759 | 6323 | 0 | TRUE | 2 |
| orf1b | 1760 | 1691 | 0 | TRUE | 5 |
| orf1b | 1761 | 1691 | 0 | TRUE | 2 |
| orf1b | 1762 | 1691 | 0 | TRUE | 3 |
| orf1b | 1763 | 1691 | 0 | TRUE | 3 |
| orf1b | 1764 | 1691 | 0 | TRUE | 2 |
| orf1b | 1765 | 1691 | 0 | TRUE | 2 |
| orf1b | 1767 | 1347 | 0 | TRUE | 3 |
| orf1b | 1768 | 1347 | 0 | TRUE | 1 |
| orf1b | 1769 | 1496 | 0 | TRUE | 1 |
| orf1b | 1770 | 1496 | 0 | TRUE | 3 |
| orf1b | 1771 | 1375 | 0 | TRUE | 2 |
| orf1b | 1772 | 1375 | 0 | TRUE | 2 |

| | | | | | |
|---|---|---|---|---|---|
| orf1b | 1773 | 1375 | 0 | TRUE | 3 |
| orf1b | 1774 | 1375 | 0 | TRUE | 6 |
| orf1b | 1775 | 1700 | 0 | TRUE | 3 |
| orf1b | 1776 | 1700 | 0 | TRUE | 1 |
| orf1b | 1777 | 2131 | 0 | TRUE | 2 |
| orf1b | 1778 | 2131 | 0 | TRUE | 4 |
| orf1b | 1779 | 2594 | 0 | TRUE | 4 |
| orf1b | 1780 | 2594 | 0 | TRUE | 2 |
| orf1b | 1781 | 2701 | 0 | TRUE | 2 |
| orf1b | 1782 | 2927 | 0 | TRUE | 4 |
| orf1b | 1783 | 3035 | 0 | TRUE | 7 |
| orf1b | 1784 | 3267 | 0 | TRUE | 2 |
| orf1b | 1785 | 2979 | 0 | TRUE | 4 |
| orf1b | 1786 | 3287 | 0 | TRUE | 3 |
| orf1b | 1787 | 3214 | 0 | TRUE | 4 |
| orf1b | 1788 | 3214 | 0 | TRUE | 2 |
| orf1b | 1789 | 2948 | 0 | TRUE | 2 |
| orf1b | 1790 | 2987 | 0 | FALSE | 0 |
| orf1b | 1791 | 2987 | 0 | TRUE | 2 |
| orf1b | 1792 | 2843 | 0 | TRUE | 3 |
| orf1b | 1793 | 2843 | 0 | TRUE | 3 |
| orf1b | 1794 | 2429 | 0 | TRUE | 1 |
| orf1b | 1795 | 2331 | 0 | TRUE | 1 |
| orf1b | 1796 | 2130 | 0 | TRUE | 3 |
| orf1b | 1797 | 2130 | 0 | FALSE | 1 |
| orf1b | 1798 | 1740 | 0 | TRUE | 3 |
| orf1b | 1799 | 1740 | 0 | FALSE | 2 |
| orf1b | 1800 | 1903 | 0 | TRUE | 1 |
| orf1b | 1801 | 1903 | 0 | FALSE | 1 |
| orf1b | 1802 | 1465 | 0 | TRUE | 3 |
| orf1b | 1803 | 1465 | 0 | TRUE | 3 |
| orf1b | 1804 | 1844 | 0 | TRUE | 1 |
| orf1b | 1805 | 1844 | 1661 | TRUE | 3 |
| orf1b | 1806 | 1844 | 1661 | TRUE | 4 |
| orf1b | 1807 | 1844 | 1661 | TRUE | 2 |
| orf1b | 1808 | 1844 | 1661 | TRUE | 4 |
| orf1b | 1809 | 1844 | 1661 | TRUE | 3 |
| orf1b | 1810 | 0 | 1661 | TRUE | 3 |
| orf1b | 1811 | 0 | 1661 | TRUE | 2 |
| orf1b | 1812 | 0 | 1661 | TRUE | 2 |
| orf1b | 1813 | 1491 | 3208 | TRUE | 1 |
| orf1b | 1814 | 1491 | 3208 | TRUE | 2 |
| orf1b | 1815 | 1491 | 2690 | TRUE | 5 |
| orf1b | 1816 | 1491 | 2690 | TRUE | 3 |
| orf1b | 1817 | 1603 | 2690 | TRUE | 3 |
| orf1b | 1818 | 1603 | 2690 | TRUE | 4 |
| orf1b | 1819 | 2517 | 2803 | TRUE | 3 |
| orf1b | 1820 | 2517 | 3565 | TRUE | 1 |
| orf1b | 1821 | 2517 | 3427 | TRUE | 3 |
| orf1b | 1822 | 2517 | 3427 | TRUE | 2 |
| orf1b | 1823 | 2517 | 3427 | TRUE | 4 |
| orf1b | 1824 | 2517 | 3427 | TRUE | 4 |
| orf1b | 1825 | 2317 | 2978 | TRUE | 4 |
| orf1b | 1826 | 2317 | 2978 | TRUE | 3 |
| orf1b | 1827 | 2317 | 2978 | TRUE | 4 |
| orf1b | 1828 | 2435 | 2146 | TRUE | 6 |
| orf1b | 1829 | 2435 | 2146 | TRUE | 2 |
| orf1b | 1830 | 2435 | 2483 | TRUE | 2 |
| orf1b | 1831 | 2435 | 2483 | TRUE | 4 |
| orf1b | 1832 | 2745 | 2483 | TRUE | 3 |
| orf1b | 1833 | 2745 | 2483 | TRUE | 3 |
| orf1b | 1834 | 1715 | 2097 | TRUE | 3 |

| | | | | | |
|---|---|---|---|---|---|
| orf1b | 1835 | 1715 | 2097 | FALSE | 3 |
| orf1b | 1836 | 1715 | 1181 | FALSE | 5 |
| orf1b | 1837 | 1715 | 1181 | FALSE | 2 |
| orf1b | 1838 | 1715 | 1181 | FALSE | 2 |
| orf1b | 1839 | 1715 | 1181 | FALSE | 4 |
| orf1b | 1845 | 1746 | 0 | FALSE | 1 |
| orf1b | 1846 | 1746 | 0 | FALSE | 2 |
| orf1b | 1847 | 1746 | 0 | FALSE | 7 |
| orf1b | 1848 | 1746 | 0 | FALSE | 6 |
| orf1b | 1849 | 1490 | 0 | TRUE | 4 |
| orf1b | 1850 | 1490 | 0 | TRUE | 5 |
| orf1b | 1851 | 1554 | 1041 | TRUE | 2 |
| orf1b | 1852 | 1554 | 1041 | TRUE | 1 |
| orf1b | 1853 | 1599 | 1041 | TRUE | 5 |
| orf1b | 1854 | 1599 | 1041 | TRUE | 3 |
| orf1b | 1855 | 1599 | 1041 | TRUE | 4 |
| orf1b | 1856 | 1599 | 1041 | FALSE | 2 |
| orf1b | 1857 | 1539 | 1041 | TRUE | 4 |
| orf1b | 1858 | 1539 | 1041 | FALSE | 3 |
| orf1b | 1859 | 1539 | 1041 | FALSE | 4 |
| orf1b | 1860 | 1513 | 1041 | TRUE | 2 |
| orf1b | 1861 | 1513 | 1041 | TRUE | 2 |
| orf1b | 1862 | 1513 | 1041 | TRUE | 1 |
| orf1b | 1863 | 1513 | 1041 | TRUE | 4 |
| orf1b | 1864 | 1564 | 1041 | TRUE | 2 |
| orf1b | 1865 | 1564 | 1041 | TRUE | 3 |
| orf1b | 1866 | 1488 | 0 | FALSE | 5 |
| orf1b | 1867 | 1488 | 0 | FALSE | 2 |
| orf1b | 1868 | 1064 | 0 | FALSE | 5 |
| orf1b | 1869 | 9393 | 1123 | FALSE | 4 |
| orf1b | 1870 | 9393 | 1123 | FALSE | 2 |
| orf1b | 1871 | 10101 | 5607 | FALSE | 4 |
| orf1b | 1872 | 14620 | 5607 | FALSE | 5 |
| orf1b | 1873 | 14620 | 5607 | FALSE | 4 |
| orf1b | 1874 | 14620 | 5607 | FALSE | 2 |
| orf1b | 1875 | 14620 | 4462 | FALSE | 0 |
| orf1b | 1876 | 14620 | 4462 | FALSE | 2 |
| orf1b | 1877 | 14620 | 4462 | FALSE | 5 |
| orf1b | 1878 | 14620 | 4462 | FALSE | 3 |
| orf1b | 1879 | 10370 | 4462 | FALSE | 4 |
| orf1b | 1880 | 10370 | 4462 | FALSE | 3 |
| orf1b | 1881 | 10826 | 4462 | FALSE | 4 |
| orf1b | 1882 | 10826 | 4462 | FALSE | 4 |
| orf1b | 1883 | 10826 | 3770 | FALSE | 1 |
| orf1b | 1884 | 8528 | 5535 | FALSE | 4 |
| orf1b | 1885 | 6983 | 5535 | FALSE | 2 |
| orf1b | 1886 | 5471 | 1016 | FALSE | 3 |
| orf1b | 1887 | 12461 | 1438 | FALSE | 0 |
| orf1b | 1888 | 12461 | 1438 | FALSE | 2 |
| orf1b | 1889 | 12503 | 1982 | FALSE | 2 |
| orf1b | 1890 | 12503 | 2173 | FALSE | 1 |
| orf1b | 1891 | 12503 | 2068 | FALSE | 3 |
| orf1b | 1892 | 12503 | 2068 | FALSE | 1 |
| orf1b | 1893 | 12503 | 2068 | FALSE | 3 |
| orf1b | 1894 | 14630 | 2068 | FALSE | 3 |
| orf1b | 1895 | 14630 | 2068 | FALSE | 7 |
| orf1b | 1896 | 15238 | 2068 | FALSE | 5 |
| orf1b | 1897 | 15238 | 2068 | FALSE | 3 |
| orf1b | 1898 | 15238 | 2281 | FALSE | 3 |
| orf1b | 1899 | 15238 | 2281 | FALSE | 4 |
| orf1b | 1900 | 19535 | 2281 | FALSE | 1 |
| orf1b | 1901 | 19535 | 2281 | FALSE | 3 |

| | | | | | |
|---|---|---|---|---|---|
| orf1b | 1902 | 12713 | 2561 | TRUE | 2 |
| orf1b | 1903 | 7031 | 2561 | FALSE | 4 |
| orf1b | 1904 | 1350 | 1546 | TRUE | 4 |
| orf1b | 1905 | 1350 | 1546 | TRUE | 4 |
| orf1b | 1906 | 1350 | 0 | FALSE | 5 |
| orf1b | 1907 | 1350 | 0 | TRUE | 3 |
| orf1b | 1908 | 1350 | 0 | FALSE | 2 |
| orf1b | 1909 | 1350 | 0 | FALSE | 2 |
| orf1b | 1910 | 1350 | 0 | FALSE | 0 |
| orf1b | 1911 | 1350 | 0 | TRUE | 5 |
| orf1b | 1912 | 1350 | 0 | TRUE | 1 |
| orf1b | 1913 | 1350 | 0 | TRUE | 2 |
| orf1b | 1914 | 1350 | 0 | TRUE | 6 |
| orf1b | 1915 | 1350 | 0 | TRUE | 2 |
| orf1b | 1916 | 1350 | 0 | TRUE | 2 |
| orf1b | 1917 | 1350 | 0 | TRUE | 6 |
| orf1b | 1919 | 1058 | 0 | TRUE | 0 |
| orf1b | 1920 | 1058 | 0 | TRUE | 2 |
| orf1b | 1921 | 1058 | 0 | TRUE | 3 |
| orf1b | 1922 | 1058 | 0 | TRUE | 4 |
| orf1b | 1923 | 1058 | 0 | TRUE | 5 |
| orf1b | 1924 | 1058 | 0 | TRUE | 2 |
| orf1b | 1925 | 1058 | 0 | TRUE | 2 |
| orf1b | 1926 | 1058 | 0 | FALSE | 5 |
| orf1b | 1927 | 1058 | 0 | TRUE | 2 |
| orf1b | 1928 | 1058 | 0 | TRUE | 4 |
| orf1b | 1929 | 1058 | 0 | TRUE | 3 |
| orf1b | 1930 | 1058 | 0 | TRUE | 2 |
| orf1b | 1931 | 1058 | 0 | TRUE | 4 |
| orf1b | 1932 | 1058 | 0 | TRUE | 3 |
| orf1b | 1933 | 1058 | 0 | TRUE | 2 |
| orf1b | 1937 | 1293 | 1520 | TRUE | 0 |
| orf1b | 1938 | 1293 | 1520 | FALSE | 1 |
| orf1b | 1939 | 1180 | 1520 | TRUE | 4 |
| orf1b | 1940 | 1180 | 1520 | FALSE | 1 |
| orf1b | 1941 | 1180 | 1520 | TRUE | 3 |
| orf1b | 1942 | 1180 | 1520 | TRUE | 2 |
| orf1b | 1943 | 1180 | 1520 | TRUE | 2 |
| orf1b | 1944 | 1180 | 1520 | FALSE | 0 |
| orf1b | 1945 | 1180 | 1520 | TRUE | 3 |
| orf1b | 1946 | 1180 | 1520 | TRUE | 1 |
| orf1b | 1947 | 1180 | 1520 | TRUE | 1 |
| orf1b | 1948 | 1180 | 1520 | TRUE | 1 |
| orf1b | 1949 | 1180 | 1520 | TRUE | 3 |
| orf1b | 1950 | 1180 | 1520 | TRUE | 2 |
| orf1b | 1951 | 1180 | 1520 | TRUE | 2 |
| orf1b | 1952 | 1068 | 0 | TRUE | 1 |
| orf1b | 1953 | 1068 | 1075 | TRUE | 2 |
| orf1b | 1954 | 0 | 1075 | TRUE | 3 |
| orf1b | 1955 | 0 | 1075 | TRUE | 2 |
| orf1b | 1956 | 0 | 1075 | TRUE | 2 |
| orf1b | 1957 | 0 | 1075 | TRUE | 3 |
| orf1b | 1958 | 0 | 1075 | TRUE | 2 |
| orf1b | 1959 | 0 | 1075 | TRUE | 3 |
| orf1b | 1960 | 0 | 1075 | TRUE | 3 |
| orf1b | 1961 | 0 | 1075 | FALSE | 3 |
| orf1b | 1962 | 0 | 1075 | TRUE | 2 |
| orf1b | 1963 | 0 | 1075 | TRUE | 4 |
| orf1b | 1964 | 0 | 1075 | FALSE | 0 |
| orf1b | 1965 | 0 | 1075 | TRUE | 3 |
| orf1b | 1966 | 0 | 1075 | FALSE | 0 |
| orf1b | 1967 | 0 | 1075 | TRUE | 4 |

| | | | | | |
|---|---|---|---|---|---|
| orf1b | 1973 | 1354 | 0 | TRUE | 2 |
| orf1b | 1974 | 1354 | 0 | TRUE | 2 |
| orf1b | 1975 | 5755 | 0 | TRUE | 3 |
| orf1b | 1976 | 5755 | 0 | TRUE | 2 |
| orf1b | 1977 | 7578 | 10600 | TRUE | 4 |
| orf1b | 1978 | 7578 | 10600 | FALSE | 3 |
| orf1b | 1979 | 7578 | 10600 | FALSE | 3 |
| orf1b | 1980 | 7578 | 10600 | FALSE | 1 |
| orf1b | 1981 | 7578 | 10600 | FALSE | 0 |
| orf1b | 1982 | 7578 | 10600 | FALSE | 2 |
| orf1b | 1983 | 6880 | 10600 | FALSE | 1 |
| orf1b | 1984 | 6880 | 10600 | FALSE | 3 |
| orf1b | 1985 | 6955 | 10600 | FALSE | 3 |
| orf1b | 1986 | 6955 | 10600 | FALSE | 0 |
| orf1b | 1987 | 6955 | 10600 | FALSE | 0 |
| orf1b | 1988 | 7578 | 10600 | FALSE | 2 |
| orf1b | 1989 | 7578 | 10600 | FALSE | 0 |
| orf1b | 1990 | 7756 | 10600 | TRUE | 0 |
| orf1b | 1991 | 7756 | 10600 | FALSE | 3 |
| orf1b | 1992 | 4465 | 0 | FALSE | 2 |
| orf1b | 1993 | 4465 | 0 | FALSE | 4 |
| orf1b | 1994 | 4465 | 0 | TRUE | 3 |
| orf1b | 1995 | 4465 | 0 | TRUE | 5 |
| orf1b | 1996 | 4465 | 0 | TRUE | 1 |
| orf1b | 1997 | 3536 | 0 | TRUE | 3 |
| orf1b | 1998 | 4656 | 0 | TRUE | 3 |
| orf1b | 1999 | 4656 | 0 | TRUE | 3 |
| orf1b | 2000 | 1679 | 0 | TRUE | 2 |
| orf1b | 2001 | 2075 | 0 | TRUE | 4 |
| orf1b | 2002 | 2075 | 0 | TRUE | 2 |
| orf1b | 2003 | 2075 | 0 | TRUE | 2 |
| orf1b | 2004 | 2075 | 0 | TRUE | 2 |
| orf1b | 2005 | 1750 | 0 | TRUE | 2 |
| orf1b | 2006 | 1750 | 0 | TRUE | 3 |
| orf1b | 2007 | 1750 | 0 | TRUE | 3 |
| orf1b | 2008 | 1750 | 0 | TRUE | 1 |
| orf1b | 2009 | 1775 | 0 | TRUE | 2 |
| orf1b | 2010 | 1775 | 0 | TRUE | 0 |
| orf1b | 2011 | 1775 | 0 | TRUE | 1 |
| orf1b | 2012 | 1808 | 0 | TRUE | 3 |
| orf1b | 2013 | 2786 | 0 | TRUE | 1 |
| orf1b | 2014 | 2786 | 0 | TRUE | 1 |
| orf1b | 2015 | 2786 | 0 | TRUE | 1 |
| orf1b | 2016 | 2891 | 0 | TRUE | 1 |
| orf1b | 2017 | 2891 | 0 | FALSE | 0 |
| orf1b | 2018 | 2891 | 0 | TRUE | 2 |
| orf1b | 2019 | 2891 | 0 | TRUE | 2 |
| orf1b | 2020 | 3786 | 0 | TRUE | 3 |
| orf1b | 2021 | 3786 | 0 | TRUE | 2 |
| orf1b | 2022 | 3786 | 0 | FALSE | 0 |
| orf1b | 2023 | 3732 | 0 | TRUE | 2 |
| orf1b | 2024 | 4672 | 0 | FALSE | 0 |
| orf1b | 2025 | 4672 | 0 | FALSE | 2 |
| orf1b | 2026 | 4672 | 0 | TRUE | 2 |
| orf1b | 2027 | 4672 | 0 | TRUE | 1 |
| orf1b | 2028 | 3623 | 0 | TRUE | 2 |
| orf1b | 2029 | 3623 | 0 | TRUE | 2 |
| orf1b | 2030 | 3623 | 0 | TRUE | 1 |
| orf1b | 2031 | 3623 | 0 | TRUE | 1 |
| orf1b | 2032 | 3623 | 0 | TRUE | 1 |
| orf1b | 2033 | 3623 | 0 | TRUE | 1 |
| orf1b | 2034 | 3623 | 0 | TRUE | 1 |

| | | | | | |
|---|---|---|---|---|---|
| orf1b | 2035 | 3623 | 0 | TRUE | 4 |
| orf1b | 2036 | 3623 | 0 | TRUE | 2 |
| orf1b | 2037 | 3623 | 0 | TRUE | 3 |
| orf1b | 2045 | 1177 | 0 | TRUE | 1 |
| orf1b | 2046 | 1177 | 0 | TRUE | 3 |
| orf1b | 2047 | 1177 | 0 | TRUE | 4 |
| orf1b | 2048 | 1177 | 0 | TRUE | 2 |
| orf1b | 2049 | 1177 | 0 | TRUE | 5 |
| orf1b | 2050 | 1177 | 0 | TRUE | 1 |
| orf1b | 2051 | 1177 | 0 | TRUE | 3 |
| orf1b | 2052 | 1177 | 0 | TRUE | 3 |
| orf1b | 2053 | 1177 | 0 | TRUE | 2 |
| orf1b | 2054 | 1177 | 0 | TRUE | 4 |
| orf1b | 2055 | 1116 | 0 | TRUE | 0 |
| orf1b | 2056 | 1116 | 0 | TRUE | 3 |
| orf1b | 2057 | 1116 | 0 | TRUE | 5 |
| orf1b | 2058 | 1116 | 0 | TRUE | 4 |
| orf1b | 2059 | 1338 | 0 | TRUE | 1 |
| orf1b | 2060 | 1419 | 0 | TRUE | 3 |
| orf1b | 2061 | 1419 | 0 | TRUE | 3 |
| orf1b | 2062 | 1419 | 0 | TRUE | 2 |
| orf1b | 2063 | 1419 | 0 | TRUE | 4 |
| orf1b | 2064 | 1419 | 0 | TRUE | 3 |
| orf1b | 2065 | 1419 | 0 | TRUE | 3 |
| orf1b | 2066 | 1419 | 0 | TRUE | 4 |
| orf1b | 2067 | 1614 | 1007 | TRUE | 3 |
| orf1b | 2068 | 1614 | 1007 | TRUE | 5 |
| orf1b | 2069 | 1614 | 1007 | TRUE | 4 |
| orf1b | 2070 | 1893 | 1007 | TRUE | 3 |
| orf1b | 2071 | 1893 | 1007 | FALSE | 5 |
| orf1b | 2072 | 1893 | 1007 | TRUE | 3 |
| orf1b | 2073 | 1893 | 1007 | FALSE | 2 |
| orf1b | 2074 | 2004 | 1007 | TRUE | 5 |
| orf1b | 2075 | 2004 | 1485 | FALSE | 3 |
| orf1b | 2076 | 2004 | 1485 | TRUE | 2 |
| orf1b | 2077 | 2140 | 4798 | FALSE | 1 |
| orf1b | 2078 | 2140 | 4798 | FALSE | 3 |
| orf1b | 2079 | 4127 | 4236 | TRUE | 1 |
| orf1b | 2080 | 4127 | 4236 | TRUE | 2 |
| orf1b | 2081 | 3549 | 3403 | FALSE | 2 |
| orf1b | 2082 | 3936 | 3882 | FALSE | 1 |
| orf1b | 2083 | 3936 | 3588 | FALSE | 1 |
| orf1b | 2084 | 3936 | 3588 | FALSE | 2 |
| orf1b | 2085 | 3361 | 3255 | FALSE | 2 |
| orf1b | 2086 | 3361 | 3255 | FALSE | 3 |
| orf1b | 2087 | 3521 | 2882 | FALSE | 4 |
| orf1b | 2088 | 3521 | 2882 | FALSE | 2 |
| orf1b | 2089 | 3315 | 2882 | FALSE | 3 |
| orf1b | 2090 | 3315 | 2966 | FALSE | 4 |
| orf1b | 2091 | 3315 | 3305 | FALSE | 4 |
| orf1b | 2092 | 3464 | 2628 | FALSE | 3 |
| orf1b | 2093 | 4193 | 2628 | FALSE | 2 |
| orf1b | 2094 | 3552 | 2635 | FALSE | 1 |
| orf1b | 2095 | 5205 | 2672 | FALSE | 4 |
| orf1b | 2096 | 5771 | 2842 | FALSE | 3 |
| orf1b | 2097 | 5771 | 2842 | FALSE | 3 |
| orf1b | 2098 | 5771 | 2840 | TRUE | 2 |
| orf1b | 2099 | 5230 | 2840 | TRUE | 2 |
| orf1b | 2100 | 5825 | 3054 | FALSE | 2 |
| orf1b | 2101 | 5825 | 3054 | TRUE | 2 |
| orf1b | 2102 | 6588 | 4022 | TRUE | 1 |
| orf1b | 2103 | 6588 | 4022 | FALSE | 2 |

| | | | | | |
|---|---|---|---|---|---|
| orf1b | 2104 | 7765 | 4022 | TRUE | 1 |
| orf1b | 2105 | 7024 | 3453 | TRUE | 5 |
| orf1b | 2106 | 7024 | 2310 | TRUE | 2 |
| orf1b | 2107 | 7411 | 2854 | TRUE | 2 |
| orf1b | 2108 | 7202 | 2854 | TRUE | 2 |
| orf1b | 2109 | 7202 | 2854 | TRUE | 6 |
| orf1b | 2110 | 6357 | 2839 | TRUE | 3 |
| orf1b | 2111 | 6357 | 2839 | FALSE | 4 |
| orf1b | 2112 | 6357 | 2839 | FALSE | 5 |
| orf1b | 2113 | 5789 | 2839 | FALSE | 2 |
| orf1b | 2114 | 6410 | 2839 | FALSE | 0 |
| orf1b | 2115 | 6410 | 3214 | FALSE | 1 |
| orf1b | 2116 | 6410 | 3214 | FALSE | 3 |
| orf1b | 2117 | 7530 | 2851 | FALSE | 5 |
| orf1b | 2118 | 7530 | 2851 | TRUE | 4 |
| orf1b | 2119 | 7620 | 2657 | TRUE | 4 |
| orf1b | 2120 | 8299 | 2903 | TRUE | 3 |
| orf1b | 2121 | 8299 | 2903 | FALSE | 2 |
| orf1b | 2122 | 8286 | 2520 | TRUE | 2 |
| orf1b | 2123 | 8286 | 2520 | FALSE | 2 |
| orf1b | 2124 | 8286 | 2520 | FALSE | 2 |
| orf1b | 2125 | 8286 | 2520 | FALSE | 0 |
| orf1b | 2126 | 8286 | 2520 | TRUE | 2 |
| orf1b | 2127 | 8286 | 10809 | FALSE | 4 |
| orf1b | 2128 | 10442 | 10809 | TRUE | 3 |
| orf1b | 2129 | 10442 | 10809 | FALSE | 2 |
| orf1b | 2130 | 10442 | 12427 | TRUE | 2 |
| orf1b | 2131 | 10442 | 12427 | FALSE | 5 |
| orf1b | 2132 | 7979 | 16103 | FALSE | 4 |
| orf1b | 2133 | 7979 | 16103 | FALSE | 4 |
| orf1b | 2134 | 0 | 43964 | FALSE | 2 |
| orf1b | 2135 | 0 | 43964 | FALSE | 3 |
| orf1b | 2136 | 0 | 43964 | FALSE | 3 |
| orf1b | 2137 | 2139 | 43964 | FALSE | 4 |
| orf1b | 2138 | 2139 | 43964 | FALSE | 4 |
| orf1b | 2139 | 2941 | 43964 | FALSE | 3 |
| orf1b | 2140 | 2941 | 43964 | FALSE | 3 |
| orf1b | 2141 | 11222 | 43964 | FALSE | 5 |
| orf1b | 2142 | 11222 | 0 | FALSE | 4 |
| orf1b | 2143 | 11222 | 0 | FALSE | 3 |
| orf1b | 2144 | 11222 | 0 | FALSE | 2 |
| orf1b | 2145 | 11222 | 0 | FALSE | 2 |
| orf1b | 2146 | 11222 | 0 | FALSE | 3 |
| orf1b | 2147 | 10228 | 0 | FALSE | 3 |
| orf1b | 2148 | 10228 | 0 | FALSE | 5 |
| orf1b | 2149 | 10228 | 3475 | FALSE | 2 |
| orf1b | 2150 | 10228 | 3475 | FALSE | 3 |
| orf1b | 2151 | 10228 | 4988 | FALSE | 2 |
| orf1b | 2152 | 12925 | 4988 | FALSE | 1 |
| orf1b | 2153 | 12925 | 4988 | FALSE | 4 |
| orf1b | 2154 | 17516 | 4988 | FALSE | 1 |
| orf1b | 2155 | 17516 | 4988 | FALSE | 1 |
| orf1b | 2156 | 7247 | 4988 | FALSE | 1 |
| orf1b | 2157 | 4190 | 4988 | FALSE | 3 |
| orf1b | 2158 | 4190 | 4988 | FALSE | 2 |
| orf1b | 2159 | 4190 | 4988 | FALSE | 6 |
| orf1b | 2160 | 4190 | 4988 | FALSE | 3 |
| orf1b | 2161 | 4190 | 4988 | FALSE | 0 |
| orf1b | 2162 | 1132 | 4988 | FALSE | 5 |
| orf1b | 2163 | 1132 | 4988 | FALSE | 3 |
| orf1b | 2164 | 1132 | 6501 | FALSE | 1 |
| orf1b | 2165 | 1132 | 6501 | FALSE | 5 |

| | | | | | |
|---|---|---|---|---|---|
| orf1b | 2166 | 1132 | 0 | FALSE | 3 |
| orf1b | 2167 | 1132 | 0 | FALSE | 2 |
| orf1b | 2168 | 1132 | 0 | FALSE | 4 |
| orf1b | 2169 | 1305 | 0 | FALSE | 2 |
| orf1b | 2170 | 1305 | 0 | FALSE | 4 |
| orf1b | 2171 | 1305 | 0 | FALSE | 2 |
| orf1b | 2172 | 1391 | 0 | FALSE | 4 |
| orf1b | 2173 | 1391 | 0 | FALSE | 3 |
| orf1b | 2174 | 1391 | 0 | FALSE | 2 |
| orf1b | 2175 | 1297 | 1100 | FALSE | 2 |
| orf1b | 2176 | 1297 | 1100 | FALSE | 2 |
| orf1b | 2177 | 1297 | 1630 | FALSE | 4 |
| orf1b | 2178 | 1297 | 1630 | FALSE | 3 |
| orf1b | 2179 | 1413 | 1468 | FALSE | 3 |
| orf1b | 2180 | 1413 | 1468 | FALSE | 5 |
| orf1b | 2181 | 1413 | 1468 | FALSE | 2 |
| orf1b | 2182 | 1413 | 1468 | FALSE | 3 |
| orf1b | 2183 | 1413 | 1468 | FALSE | 3 |
| orf1b | 2184 | 1428 | 1468 | FALSE | 2 |
| orf1b | 2185 | 1428 | 1468 | FALSE | 0 |
| orf1b | 2186 | 1428 | 1468 | FALSE | 3 |
| orf1b | 2187 | 1801 | 1468 | FALSE | 3 |
| orf1b | 2188 | 1801 | 1468 | FALSE | 6 |
| orf1b | 2189 | 1801 | 1468 | FALSE | 4 |
| orf1b | 2190 | 2399 | 1713 | FALSE | 4 |
| orf1b | 2191 | 2399 | 3509 | FALSE | 2 |
| orf1b | 2192 | 2399 | 3783 | FALSE | 1 |
| orf1b | 2193 | 2399 | 3783 | FALSE | 2 |
| orf1b | 2194 | 2922 | 8899 | FALSE | 1 |
| orf1b | 2195 | 2922 | 8899 | FALSE | 2 |
| orf1b | 2196 | 2922 | 8899 | FALSE | 2 |
| orf1b | 2197 | 2922 | 8899 | FALSE | 3 |
| orf1b | 2198 | 2922 | 8899 | FALSE | 5 |
| orf1b | 2199 | 2922 | 6264 | FALSE | 3 |
| orf1b | 2200 | 2922 | 6264 | FALSE | 2 |
| orf1b | 2201 | 2922 | 5671 | FALSE | 3 |
| orf1b | 2202 | 0 | 5671 | FALSE | 2 |
| orf1b | 2203 | 0 | 4789 | FALSE | 3 |
| orf1b | 2204 | 0 | 4789 | FALSE | 3 |
| orf1b | 2205 | 0 | 4789 | FALSE | 3 |
| orf1b | 2206 | 0 | 3419 | FALSE | 1 |
| orf1b | 2207 | 1317 | 3419 | FALSE | 6 |
| orf1b | 2208 | 1317 | 3419 | FALSE | 4 |
| orf1b | 2209 | 1317 | 3419 | FALSE | 3 |
| orf1b | 2210 | 1317 | 3419 | FALSE | 4 |
| orf1b | 2211 | 1317 | 3419 | FALSE | 3 |
| orf1b | 2212 | 1317 | 3419 | FALSE | 3 |
| orf1b | 2213 | 1317 | 4040 | FALSE | 3 |
| orf1b | 2214 | 1317 | 4176 | FALSE | 2 |
| orf1b | 2215 | 1317 | 6922 | FALSE | 3 |
| orf1b | 2216 | 1317 | 7734 | FALSE | 4 |
| orf1b | 2217 | 1317 | 7734 | FALSE | 1 |
| orf1b | 2218 | 1317 | 10531 | FALSE | 3 |
| orf1b | 2219 | 6363 | 7537 | FALSE | 5 |
| orf1b | 2220 | 6363 | 7537 | FALSE | 2 |
| orf1b | 2221 | 9944 | 7820 | FALSE | 2 |
| orf1b | 2222 | 18570 | 7820 | FALSE | 4 |
| orf1b | 2223 | 18570 | 6889 | FALSE | 5 |
| orf1b | 2224 | 18570 | 6889 | FALSE | 3 |
| orf1b | 2225 | 18570 | 6889 | FALSE | 2 |
| orf1b | 2226 | 18570 | 6889 | FALSE | 4 |
| orf1b | 2227 | 18570 | 6889 | FALSE | 4 |

| | | | | | |
|---|---|---|---|---|---|
| orf1b | 2228 | 18570 | 7086 | FALSE | 1 |
| orf1b | 2229 | 18570 | 7086 | FALSE | 3 |
| orf1b | 2230 | 18570 | 5068 | FALSE | 0 |
| orf1b | 2231 | 18570 | 5068 | FALSE | 1 |
| orf1b | 2232 | 18570 | 5068 | FALSE | 2 |
| orf1b | 2233 | 18570 | 4796 | FALSE | 1 |
| orf1b | 2234 | 20685 | 5607 | FALSE | 1 |
| orf1b | 2235 | 20685 | 6890 | FALSE | 0 |
| orf1b | 2236 | 0 | 5986 | FALSE | 3 |
| orf1b | 2237 | 0 | 5986 | FALSE | 3 |
| orf1b | 2238 | 0 | 7862 | FALSE | 4 |
| orf1b | 2239 | 0 | 7862 | FALSE | 1 |
| orf1b | 2240 | 0 | 7862 | FALSE | 1 |
| orf1b | 2241 | 0 | 5745 | FALSE | 3 |
| orf1b | 2242 | 0 | 5745 | FALSE | 1 |
| orf1b | 2243 | 0 | 5745 | FALSE | 1 |
| orf1b | 2244 | 0 | 5745 | FALSE | 1 |
| orf1b | 2245 | 0 | 5745 | FALSE | 2 |
| orf1b | 2246 | 0 | 5745 | FALSE | 3 |
| orf1b | 2247 | 0 | 5745 | FALSE | 3 |
| orf1b | 2248 | 0 | 6765 | FALSE | 4 |
| orf1b | 2249 | 0 | 6765 | FALSE | 3 |
| orf1b | 2250 | 0 | 1511 | FALSE | 3 |
| orf1b | 2251 | 4656 | 1511 | FALSE | 1 |
| orf1b | 2252 | 4656 | 1511 | TRUE | 2 |
| orf1b | 2253 | 4829 | 3057 | TRUE | 2 |
| orf1b | 2254 | 4829 | 3057 | TRUE | 4 |
| orf1b | 2255 | 4829 | 3057 | TRUE | 2 |
| orf1b | 2256 | 4829 | 3830 | TRUE | 4 |
| orf1b | 2257 | 4111 | 2889 | TRUE | 4 |
| orf1b | 2258 | 4111 | 2889 | TRUE | 3 |
| orf1b | 2259 | 4885 | 3184 | TRUE | 4 |
| orf1b | 2260 | 4885 | 3184 | TRUE | 4 |
| orf1b | 2261 | 4607 | 3370 | TRUE | 2 |
| orf1b | 2262 | 4607 | 3370 | TRUE | 3 |
| orf1b | 2263 | 4607 | 3370 | FALSE | 2 |
| orf1b | 2264 | 4607 | 3370 | TRUE | 1 |
| orf1b | 2265 | 4269 | 3112 | TRUE | 1 |
| orf1b | 2266 | 4246 | 3112 | FALSE | 1 |
| orf1b | 2267 | 4246 | 3112 | FALSE | 2 |
| orf1b | 2268 | 4109 | 2873 | FALSE | 3 |
| orf1b | 2269 | 4258 | 2873 | FALSE | 1 |
| orf1b | 2270 | 4258 | 2873 | FALSE | 3 |
| orf1b | 2271 | 4353 | 3443 | FALSE | 3 |
| orf1b | 2272 | 4526 | 3748 | FALSE | 1 |
| orf1b | 2273 | 4526 | 3748 | FALSE | 3 |
| orf1b | 2274 | 4196 | 3701 | FALSE | 4 |
| orf1b | 2275 | 4196 | 3701 | TRUE | 2 |
| orf1b | 2276 | 4044 | 3765 | FALSE | 1 |
| orf1b | 2277 | 4252 | 4265 | FALSE | 1 |
| orf1b | 2278 | 4252 | 4265 | FALSE | 2 |
| orf1b | 2279 | 4139 | 5680 | FALSE | 1 |
| orf1b | 2280 | 4482 | 6226 | TRUE | 4 |
| orf1b | 2281 | 4305 | 6226 | TRUE | 0 |
| orf1b | 2282 | 4305 | 6226 | FALSE | 4 |
| orf1b | 2283 | 4305 | 6226 | TRUE | 0 |
| orf1b | 2284 | 4169 | 6226 | TRUE | 1 |
| orf1b | 2285 | 4169 | 5674 | FALSE | 2 |
| orf1b | 2286 | 3973 | 6219 | TRUE | 2 |
| orf1b | 2287 | 3973 | 6219 | TRUE | 1 |
| orf1b | 2288 | 3973 | 6219 | FALSE | 1 |
| orf1b | 2289 | 3973 | 6219 | TRUE | 1 |

| | | | | | |
|---|---|---|---|---|---|
| orf1b | 2290 | 3973 | 6219 | TRUE | 3 |
| orf1b | 2291 | 3973 | 6219 | TRUE | 1 |
| orf1b | 2292 | 3750 | 6209 | TRUE | 3 |
| orf1b | 2293 | 3750 | 6209 | TRUE | 1 |
| orf1b | 2294 | 1648 | 1253 | FALSE | 2 |
| orf1b | 2295 | 1648 | 1253 | FALSE | 3 |
| orf1b | 2296 | 0 | 1253 | FALSE | 1 |
| orf1b | 2297 | 0 | 1253 | FALSE | 2 |
| orf1b | 2298 | 0 | 1253 | FALSE | 2 |
| orf1b | 2299 | 0 | 1253 | FALSE | 2 |
| orf1b | 2305 | 1137 | 0 | FALSE | 2 |
| orf1b | 2306 | 1137 | 0 | FALSE | 4 |
| orf1b | 2307 | 1137 | 0 | FALSE | 4 |
| orf1b | 2308 | 1137 | 0 | FALSE | 2 |
| orf1b | 2309 | 2038 | 4340 | FALSE | 1 |
| orf1b | 2310 | 2038 | 4340 | FALSE | 2 |
| orf1b | 2311 | 5790 | 7585 | FALSE | 2 |
| orf1b | 2312 | 5790 | 7585 | FALSE | 1 |
| orf1b | 2313 | 4550 | 6658 | FALSE | 3 |
| orf1b | 2314 | 4550 | 6658 | FALSE | 0 |
| orf1b | 2315 | 6341 | 7325 | FALSE | 3 |
| orf1b | 2316 | 6341 | 7325 | FALSE | 0 |
| orf1b | 2317 | 7909 | 7060 | FALSE | 1 |
| orf1b | 2318 | 7909 | 7060 | FALSE | 3 |
| orf1b | 2319 | 7909 | 6746 | FALSE | 0 |
| orf1b | 2320 | 8361 | 6746 | FALSE | 2 |
| orf1b | 2321 | 7619 | 6251 | FALSE | 4 |
| orf1b | 2322 | 7619 | 6251 | FALSE | 3 |
| orf1b | 2323 | 7619 | 6251 | FALSE | 2 |
| orf1b | 2324 | 7912 | 6443 | TRUE | 2 |
| orf1b | 2325 | 7912 | 6443 | TRUE | 2 |
| orf1b | 2326 | 8194 | 5751 | TRUE | 2 |
| orf1b | 2327 | 8194 | 5751 | TRUE | 3 |
| orf1b | 2328 | 10646 | 6019 | TRUE | 2 |
| orf1b | 2329 | 10646 | 6019 | FALSE | 2 |
| orf1b | 2330 | 9646 | 4373 | FALSE | 3 |
| orf1b | 2331 | 9646 | 4373 | TRUE | 1 |
| orf1b | 2332 | 2060 | 2611 | FALSE | 2 |
| orf1b | 2333 | 2060 | 2611 | TRUE | 2 |
| orf1b | 2334 | 2060 | 1306 | TRUE | 2 |
| orf1b | 2335 | 2060 | 1306 | TRUE | 1 |
| orf1b | 2343 | 1752 | 0 | FALSE | 1 |
| orf1b | 2344 | 1752 | 0 | FALSE | 3 |
| orf1b | 2345 | 1752 | 0 | FALSE | 3 |
| orf1b | 2346 | 1752 | 0 | TRUE | 2 |
| orf1b | 2347 | 1752 | 0 | TRUE | 2 |
| orf1b | 2348 | 1752 | 0 | TRUE | 2 |
| orf1b | 2349 | 1411 | 1457 | TRUE | 2 |
| orf1b | 2350 | 1411 | 1457 | TRUE | 2 |
| orf1b | 2351 | 1276 | 1762 | TRUE | 3 |
| orf1b | 2352 | 1276 | 1762 | TRUE | 4 |
| orf1b | 2353 | 2346 | 1762 | TRUE | 2 |
| orf1b | 2354 | 2346 | 1762 | TRUE | 2 |
| orf1b | 2355 | 2346 | 1762 | TRUE | 2 |
| orf1b | 2356 | 2346 | 1762 | TRUE | 2 |
| orf1b | 2357 | 2346 | 1762 | TRUE | 3 |
| orf1b | 2358 | 2544 | 1762 | TRUE | 1 |
| orf1b | 2359 | 2544 | 1762 | TRUE | 3 |
| orf1b | 2360 | 2544 | 1762 | TRUE | 2 |
| orf1b | 2361 | 2544 | 1762 | TRUE | 2 |
| orf1b | 2362 | 2544 | 1762 | TRUE | 2 |
| orf1b | 2363 | 2544 | 1762 | FALSE | 2 |

| | | | | | |
|---|---|---|---|---|---|
| orf1b | 2364 | 3281 | 2068 | FALSE | 4 |
| orf1b | 2365 | 3281 | 2068 | FALSE | 1 |
| orf1b | 2366 | 5557 | 0 | FALSE | 4 |
| orf1b | 2367 | 5557 | 0 | FALSE | 4 |
| orf1b | 2377 | 11289 | 4150 | FALSE | 3 |
| orf1b | 2378 | 11289 | 4150 | FALSE | 3 |
| orf1b | 2379 | 11289 | 4150 | FALSE | 3 |
| orf1b | 2380 | 11289 | 4150 | FALSE | 2 |
| orf1b | 2381 | 11289 | 4150 | FALSE | 5 |
| orf1b | 2382 | 11289 | 4150 | FALSE | 3 |
| orf1b | 2383 | 11289 | 4150 | FALSE | 4 |
| orf1b | 2384 | 11289 | 4150 | FALSE | 4 |
| orf1b | 2385 | 6353 | 4150 | FALSE | 3 |
| orf1b | 2386 | 6353 | 4150 | TRUE | 3 |
| orf1b | 2387 | 4196 | 2671 | TRUE | 7 |
| orf1b | 2388 | 4196 | 2671 | TRUE | 6 |
| orf1b | 2389 | 4196 | 2671 | TRUE | 5 |
| orf1b | 2390 | 4196 | 2671 | TRUE | 3 |
| orf1b | 2391 | 4196 | 2671 | TRUE | 3 |
| orf1b | 2392 | 1832 | 1192 | TRUE | 3 |
| orf1b | 2393 | 1832 | 1192 | TRUE | 2 |
| orf1b | 2394 | 1832 | 1192 | TRUE | 6 |
| orf1b | 2395 | 1832 | 1192 | FALSE | 0 |
| orf1b | 2396 | 1832 | 1192 | FALSE | 3 |
| orf1b | 2397 | 1832 | 1192 | TRUE | 4 |
| orf1b | 2398 | 1832 | 1192 | FALSE | 3 |
| orf1b | 2399 | 1832 | 1192 | TRUE | 4 |
| orf1b | 2400 | 2040 | 1192 | TRUE | 3 |
| orf1b | 2401 | 2040 | 1192 | FALSE | 5 |
| orf1b | 2403 | 0 | 1950 | TRUE | 4 |
| orf1b | 2404 | 0 | 1950 | TRUE | 6 |
| orf1b | 2405 | 0 | 3132 | TRUE | 6 |
| orf1b | 2406 | 0 | 3132 | TRUE | 2 |
| orf1b | 2407 | 0 | 3132 | TRUE | 2 |
| orf1b | 2408 | 0 | 3132 | FALSE | 1 |
| orf1b | 2409 | 0 | 3132 | FALSE | 2 |
| orf1b | 2410 | 0 | 3132 | TRUE | 3 |
| orf1b | 2411 | 0 | 3132 | TRUE | 2 |
| orf1b | 2412 | 0 | 3132 | TRUE | 2 |
| orf1b | 2413 | 0 | 3132 | TRUE | 2 |
| orf1b | 2414 | 0 | 3132 | TRUE | 4 |
| orf1b | 2415 | 0 | 3132 | TRUE | 1 |
| orf1b | 2416 | 0 | 3132 | TRUE | 4 |
| orf1b | 2417 | 0 | 3132 | TRUE | 3 |
| orf1b | 2418 | 0 | 4313 | FALSE | 2 |
| orf1b | 2419 | 0 | 4313 | TRUE | 1 |
| orf1b | 2423 | 0 | 1322 | FALSE | 1 |
| orf1b | 2424 | 0 | 1322 | FALSE | 2 |
| orf1b | 2425 | 0 | 1322 | FALSE | 3 |
| orf1b | 2426 | 0 | 1322 | TRUE | 2 |
| orf1b | 2427 | 0 | 1322 | TRUE | 1 |
| orf1b | 2428 | 0 | 1322 | TRUE | 2 |
| orf1b | 2429 | 0 | 1322 | FALSE | 3 |
| orf1b | 2430 | 0 | 1322 | TRUE | 2 |
| orf1b | 2431 | 0 | 1322 | TRUE | 5 |
| orf1b | 2432 | 0 | 1322 | TRUE | 3 |
| orf1b | 2433 | 0 | 1322 | TRUE | 3 |
| orf1b | 2434 | 0 | 1322 | TRUE | 5 |
| orf1b | 2435 | 0 | 1322 | TRUE | 1 |
| orf1b | 2436 | 0 | 1322 | TRUE | 4 |
| orf1b | 2437 | 0 | 1322 | FALSE | 3 |
| orf1b | 2461 | 4405 | 0 | TRUE | 1 |

| | | | | | |
|---|---|---|---|---|---|
| orf1b | 2462 | 4405 | 0 | TRUE | 3 |
| orf1b | 2463 | 4405 | 0 | TRUE | 3 |
| orf1b | 2464 | 4405 | 0 | TRUE | 2 |
| orf1b | 2465 | 4405 | 0 | TRUE | 1 |
| orf1b | 2466 | 4405 | 0 | TRUE | 4 |
| orf1b | 2467 | 4405 | 0 | TRUE | 0 |
| orf1b | 2468 | 4405 | 0 | FALSE | 3 |
| orf1b | 2469 | 4405 | 0 | TRUE | 3 |
| orf1b | 2470 | 4405 | 0 | TRUE | 2 |
| orf1b | 2471 | 7918 | 0 | TRUE | 4 |
| orf1b | 2472 | 7918 | 0 | TRUE | 4 |
| orf1b | 2473 | 7918 | 0 | TRUE | 3 |
| orf1b | 2474 | 7918 | 0 | TRUE | 2 |
| orf1b | 2475 | 7918 | 0 | TRUE | 4 |
| orf1b | 2476 | 11432 | 0 | TRUE | 2 |
| orf1b | 2477 | 11432 | 0 | TRUE | 3 |
| orf1b | 2478 | 11432 | 0 | TRUE | 1 |
| orf1b | 2479 | 11432 | 0 | TRUE | 1 |
| orf1b | 2480 | 11432 | 0 | TRUE | 2 |
| orf1b | 2481 | 11432 | 0 | TRUE | 4 |
| orf1b | 2482 | 11432 | 0 | FALSE | 2 |
| orf1b | 2483 | 11432 | 0 | TRUE | 2 |
| orf1b | 2484 | 11432 | 0 | TRUE | 3 |
| orf1b | 2485 | 11432 | 0 | TRUE | 4 |
| orf1b | 2487 | 12629 | 0 | TRUE | 4 |
| orf1b | 2488 | 12629 | 0 | TRUE | 3 |
| orf1b | 2489 | 7022 | 0 | TRUE | 1 |
| orf1b | 2490 | 7022 | 0 | TRUE | 4 |
| orf1b | 2491 | 5497 | 0 | TRUE | 3 |
| orf1b | 2492 | 5497 | 0 | TRUE | 3 |
| orf1b | 2493 | 4174 | 0 | TRUE | 3 |
| orf1b | 2494 | 4174 | 0 | TRUE | 2 |
| orf1b | 2495 | 4616 | 0 | TRUE | 4 |
| orf1b | 2496 | 4616 | 0 | FALSE | 3 |
| orf1b | 2497 | 4616 | 0 | FALSE | 2 |
| orf1b | 2498 | 4616 | 0 | TRUE | 1 |
| orf1b | 2499 | 4179 | 0 | TRUE | 4 |
| orf1b | 2500 | 4179 | 0 | TRUE | 2 |
| orf1b | 2501 | 3896 | 0 | TRUE | 4 |
| orf1b | 2502 | 2804 | 0 | TRUE | 2 |
| orf1b | 2503 | 2668 | 0 | TRUE | 3 |
| orf1b | 2504 | 2825 | 0 | TRUE | 2 |
| orf1b | 2505 | 2825 | 0 | TRUE | 2 |
| orf1b | 2506 | 2879 | 0 | FALSE | 3 |
| orf1b | 2507 | 2879 | 0 | TRUE | 3 |
| orf1b | 2508 | 3154 | 0 | TRUE | 4 |
| orf1b | 2509 | 3154 | 0 | TRUE | 5 |
| orf1b | 2510 | 1442 | 0 | TRUE | 5 |
| orf1b | 2511 | 1442 | 0 | TRUE | 4 |
| orf1b | 2512 | 1442 | 0 | TRUE | 2 |
| orf1b | 2513 | 1442 | 0 | FALSE | 6 |
| orf1b | 2514 | 1603 | 0 | FALSE | 5 |
| orf1b | 2515 | 1603 | 0 | TRUE | 1 |
| orf1b | 2516 | 1576 | 0 | TRUE | 2 |
| orf1b | 2517 | 1576 | 0 | TRUE | 3 |
| orf1b | 2519 | 3073 | 0 | TRUE | 3 |
| orf1b | 2520 | 3073 | 0 | TRUE | 2 |
| orf1b | 2521 | 3073 | 0 | TRUE | 3 |
| orf1b | 2522 | 3073 | 0 | TRUE | 4 |
| orf1b | 2523 | 3073 | 0 | TRUE | 4 |
| orf1b | 2524 | 3073 | 0 | FALSE | 3 |
| orf1b | 2525 | 3073 | 0 | TRUE | 2 |

| | | | | | |
|---|---|---|---|---|---|
| orf1b | 2526 | 3073 | 0 | FALSE | 3 |
| orf1b | 2527 | 3073 | 0 | TRUE | 4 |
| orf1b | 2528 | 3073 | 0 | TRUE | 1 |
| orf1b | 2529 | 3073 | 0 | TRUE | 2 |
| orf1b | 2530 | 3073 | 0 | TRUE | 5 |
| orf1b | 2531 | 3073 | 0 | TRUE | 3 |
| orf1b | 2532 | 3073 | 0 | TRUE | 3 |
| orf1b | 2533 | 3073 | 0 | TRUE | 3 |
| orf1b | 2535 | 2915 | 1872 | TRUE | 3 |
| orf1b | 2536 | 2915 | 1872 | TRUE | 0 |
| orf1b | 2537 | 4572 | 1872 | TRUE | 2 |
| orf1b | 2538 | 4572 | 1872 | TRUE | 3 |
| orf1b | 2539 | 8540 | 4662 | TRUE | 2 |
| orf1b | 2540 | 8540 | 4662 | FALSE | 0 |
| orf1b | 2541 | 8540 | 4662 | TRUE | 3 |
| orf1b | 2542 | 8540 | 4662 | TRUE | 3 |
| orf1b | 2543 | 8540 | 4662 | FALSE | 4 |
| orf1b | 2544 | 8540 | 4662 | TRUE | 6 |
| orf1b | 2545 | 8540 | 4662 | FALSE | 4 |
| orf1b | 2546 | 8540 | 4662 | FALSE | 2 |
| orf1b | 2547 | 8540 | 4662 | TRUE | 2 |
| orf1b | 2548 | 8540 | 4662 | TRUE | 3 |
| orf1b | 2549 | 8540 | 4662 | FALSE | 1 |
| orf1b | 2550 | 9946 | 5591 | TRUE | 3 |
| orf1b | 2551 | 9946 | 5591 | TRUE | 6 |
| orf1b | 2552 | 11185 | 5591 | TRUE | 4 |
| orf1b | 2553 | 11185 | 5591 | TRUE | 3 |
| orf1b | 2569 | 4969 | 0 | TRUE | 1 |
| orf1b | 2570 | 4969 | 0 | TRUE | 1 |
| orf1b | 2571 | 4969 | 0 | TRUE | 2 |
| orf1b | 2572 | 4969 | 0 | TRUE | 1 |
| orf1b | 2573 | 4969 | 0 | TRUE | 4 |
| orf1b | 2574 | 4969 | 0 | TRUE | 0 |
| orf1b | 2575 | 4969 | 0 | TRUE | 2 |
| orf1b | 2576 | 4969 | 0 | TRUE | 0 |
| orf1b | 2577 | 4969 | 0 | TRUE | 2 |
| orf1b | 2578 | 4969 | 0 | TRUE | 0 |
| orf1b | 2579 | 4969 | 0 | TRUE | 1 |
| orf1b | 2580 | 4969 | 0 | TRUE | 3 |
| orf1b | 2581 | 4969 | 0 | TRUE | 1 |
| orf1b | 2582 | 4969 | 0 | TRUE | 2 |
| orf1b | 2583 | 4969 | 0 | FALSE | 2 |
| orf1b | 2591 | 3706 | 0 | FALSE | 2 |
| orf1b | 2592 | 3706 | 0 | TRUE | 1 |
| orf1b | 2593 | 3055 | 0 | TRUE | 0 |
| orf1b | 2594 | 3055 | 0 | TRUE | 4 |
| orf1b | 2595 | 4145 | 0 | TRUE | 0 |
| orf1b | 2596 | 4145 | 0 | TRUE | 2 |
| orf1b | 2597 | 4145 | 0 | TRUE | 2 |
| orf1b | 2598 | 4145 | 0 | TRUE | 1 |
| orf1b | 2599 | 3630 | 0 | TRUE | 2 |
| orf1b | 2600 | 3630 | 0 | TRUE | 2 |
| orf1b | 2601 | 3630 | 0 | TRUE | 2 |
| orf1b | 2602 | 3630 | 0 | TRUE | 0 |
| orf1b | 2603 | 3630 | 1294 | TRUE | 0 |
| orf1b | 2604 | 3630 | 1294 | TRUE | 0 |
| orf1b | 2605 | 3630 | 1294 | TRUE | 2 |
| orf1b | 2606 | 3605 | 1294 | TRUE | 1 |
| orf1b | 2607 | 3636 | 1294 | TRUE | 1 |
| orf1b | 2608 | 4047 | 1294 | TRUE | 2 |
| orf1b | 2609 | 4047 | 1294 | TRUE | 1 |
| orf1b | 2610 | 2908 | 1294 | TRUE | 4 |

| | | | | | |
|---|---|---|---|---|---|
| orf1b | 2611 | 2908 | 1294 | TRUE | 0 |
| orf1b | 2612 | 2908 | 1294 | TRUE | 3 |
| orf1b | 2613 | 2908 | 1294 | TRUE | 6 |
| orf1b | 2614 | 3730 | 1294 | TRUE | 2 |
| orf1b | 2615 | 3730 | 1294 | TRUE | 2 |
| orf1b | 2616 | 3730 | 1294 | TRUE | 2 |
| orf1b | 2617 | 2814 | 1294 | TRUE | 1 |
| orf1b | 2618 | 2814 | 0 | TRUE | 1 |
| orf1b | 2619 | 2814 | 0 | TRUE | 0 |
| orf1b | 2620 | 2814 | 0 | FALSE | 1 |
| orf1b | 2621 | 2814 | 0 | FALSE | 1 |
| orf1b | 2622 | 2357 | 0 | FALSE | 2 |
| orf1b | 2623 | 2357 | 0 | FALSE | 2 |
| orf1b | 2624 | 2357 | 0 | TRUE | 0 |
| orf1b | 2625 | 2357 | 0 | FALSE | 2 |
| orf1b | 2626 | 2357 | 0 | TRUE | 0 |
| orf1b | 2627 | 2357 | 0 | TRUE | 0 |
| orf1b | 2628 | 2357 | 0 | TRUE | 0 |
| orf1b | 2629 | 2357 | 0 | TRUE | 1 |
| orf1b | 2630 | 2357 | 0 | TRUE | 0 |
| orf1b | 2631 | 2357 | 0 | TRUE | 1 |
| orf1b | 2633 | 0 | 1151 | FALSE | 2 |
| orf1b | 2634 | 0 | 1151 | FALSE | 1 |
| orf1b | 2635 | 0 | 1151 | FALSE | 2 |
| orf1b | 2636 | 0 | 1151 | FALSE | 0 |
| orf1b | 2637 | 0 | 1151 | FALSE | 1 |
| orf1b | 2638 | 0 | 1151 | FALSE | 0 |
| orf1b | 2639 | 0 | 1151 | FALSE | 0 |
| orf1b | 2640 | 0 | 1151 | TRUE | 2 |
| orf1b | 2641 | 0 | 1151 | FALSE | 2 |
| orf1b | 2642 | 0 | 1151 | TRUE | 1 |
| orf1b | 2643 | 0 | 1151 | TRUE | 3 |
| orf1b | 2644 | 0 | 1151 | TRUE | 1 |
| orf1b | 2645 | 0 | 1151 | TRUE | 1 |
| orf1b | 2646 | 0 | 1151 | FALSE | 2 |
| orf1b | 2647 | 0 | 1151 | TRUE | 3 |
| orf1b | 2651 | 0 | 1394 | TRUE | 0 |
| orf1b | 2652 | 0 | 1394 | TRUE | 1 |
| orf1b | 2653 | 0 | 1344 | TRUE | 3 |
| orf1b | 2654 | 0 | 1344 | TRUE | 1 |
| orf1b | 2655 | 0 | 1344 | TRUE | 4 |
| orf1b | 2656 | 0 | 1344 | FALSE | 1 |
| orf1b | 2657 | 0 | 1344 | FALSE | 1 |
| orf1b | 2658 | 0 | 1344 | FALSE | 1 |
| orf1b | 2659 | 11773 | 1674 | FALSE | 2 |
| orf1b | 2660 | 11773 | 1674 | FALSE | 0 |
| orf1b | 2661 | 9213 | 5649 | FALSE | 1 |
| orf1b | 2662 | 9213 | 5649 | FALSE | 5 |
| orf1b | 2663 | 9213 | 5649 | FALSE | 2 |
| orf1b | 2664 | 9213 | 5649 | FALSE | 3 |
| orf1b | 2665 | 9213 | 5649 | TRUE | 1 |
| orf1b | 2666 | 9213 | 6359 | TRUE | 3 |
| orf1b | 2667 | 9213 | 6359 | TRUE | 4 |
| orf1b | 2668 | 9213 | 7372 | TRUE | 2 |
| orf1b | 2669 | 8885 | 7372 | FALSE | 0 |
| orf1b | 2670 | 8885 | 7372 | FALSE | 1 |
| orf1b | 2671 | 8885 | 7372 | TRUE | 1 |
| orf1b | 2672 | 8885 | 7372 | TRUE | 2 |
| orf1b | 2673 | 8885 | 7372 | TRUE | 2 |
| orf1b | 2674 | 7923 | 8631 | TRUE | 3 |
| orf1b | 2675 | 7923 | 8631 | TRUE | 3 |
| orf1b | 2676 | 7902 | 0 | TRUE | 3 |

| | | | | | |
|---|---|---|---|---|---|
| orf1b | 2677 | 4493 | 0 | TRUE | 1 |
| orf1b | 2678 | 4493 | 0 | TRUE | 0 |
| orf1b | 2679 | 7323 | 0 | TRUE | 1 |
| orf1b | 2680 | 7323 | 0 | FALSE | 0 |
| orf1b | 2681 | 8333 | 2870 | FALSE | 1 |
| orf1b | 2682 | 8333 | 2870 | FALSE | 1 |
| orf1b | 2683 | 8333 | 2870 | FALSE | 2 |
| orf1b | 2684 | 8419 | 2870 | FALSE | 2 |
| orf1b | 2685 | 8419 | 2870 | FALSE | 1 |
| orf1b | 2686 | 8419 | 2870 | FALSE | 0 |
| orf1b | 2687 | 8419 | 2870 | FALSE | 3 |
| orf1b | 2688 | 8419 | 2870 | FALSE | 1 |
| orf1b | 2689 | 8419 | 2870 | FALSE | 0 |
| orf1b | 2690 | 8419 | 2870 | FALSE | 2 |
| orf1b | 2691 | 8419 | 2870 | FALSE | 1 |
| orf1b | 2692 | 10252 | 2870 | FALSE | 3 |
| orf1b | 2693 | 10252 | 2870 | FALSE | 2 |
| orf1b | 2694 | 9342 | 2870 | FALSE | 2 |
| orf1b | 2695 | 9342 | 2870 | FALSE | 1 |
| S | 1 | 1420 | 1720 | FALSE | 2 |
| S | 2 | 1420 | 1720 | FALSE | 2 |
| S | 3 | 1420 | 1720 | FALSE | 2 |
| S | 4 | 1420 | 1512 | FALSE | 3 |
| S | 5 | 1420 | 1512 | FALSE | 1 |
| S | 6 | 1420 | 1512 | FALSE | 3 |
| S | 7 | 1420 | 1512 | FALSE | 3 |
| S | 8 | 1420 | 1563 | FALSE | 3 |
| S | 9 | 1420 | 1554 | FALSE | 2 |
| S | 10 | 1420 | 1554 | FALSE | 2 |
| S | 11 | 1132 | 1498 | FALSE | 4 |
| S | 12 | 1132 | 1498 | FALSE | 2 |
| S | 13 | 0 | 1498 | FALSE | 4 |
| S | 14 | 0 | 1498 | FALSE | 4 |
| S | 15 | 0 | 1498 | FALSE | 3 |
| S | 16 | 0 | 1498 | FALSE | 1 |
| S | 17 | 1534 | 1498 | FALSE | 2 |
| S | 18 | 1534 | 1498 | FALSE | 2 |
| S | 19 | 1534 | 1975 | FALSE | 2 |
| S | 20 | 1534 | 1975 | FALSE | 4 |
| S | 21 | 1534 | 1975 | FALSE | 4 |
| S | 22 | 1534 | 1975 | FALSE | 2 |
| S | 23 | 1534 | 2129 | FALSE | 4 |
| S | 24 | 1534 | 2735 | FALSE | 3 |
| S | 25 | 1534 | 2735 | FALSE | 5 |
| S | 26 | 1534 | 2735 | FALSE | 6 |
| S | 27 | 1534 | 2735 | FALSE | 4 |
| S | 28 | 1534 | 2735 | FALSE | 3 |
| S | 29 | 1534 | 2735 | FALSE | 1 |
| S | 30 | 1534 | 2735 | FALSE | 3 |
| S | 31 | 1534 | 2735 | FALSE | 1 |
| S | 32 | 0 | 2735 | FALSE | 1 |
| S | 33 | 0 | 2735 | FALSE | 1 |
| S | 35 | 0 | 2328 | FALSE | 3 |
| S | 36 | 0 | 2328 | FALSE | 1 |
| S | 37 | 0 | 2328 | FALSE | 2 |
| S | 38 | 0 | 2328 | FALSE | 3 |
| S | 39 | 0 | 2328 | FALSE | 4 |
| S | 40 | 0 | 2328 | FALSE | 1 |
| S | 41 | 0 | 2328 | FALSE | 4 |
| S | 42 | 0 | 2328 | FALSE | 4 |
| S | 43 | 0 | 2328 | FALSE | 3 |
| S | 44 | 0 | 2328 | FALSE | 2 |

| | | | | | |
|---|---|---|---|---|---|
| S | 45 | 0 | 3180 | FALSE | 4 |
| S | 46 | 0 | 3180 | FALSE | 2 |
| S | 47 | 0 | 3180 | FALSE | 2 |
| S | 48 | 0 | 3180 | FALSE | 1 |
| S | 49 | 0 | 3180 | FALSE | 3 |
| S | 50 | 0 | 4031 | FALSE | 3 |
| S | 51 | 0 | 4031 | FALSE | 3 |
| S | 52 | 0 | 4031 | FALSE | 4 |
| S | 53 | 0 | 4031 | FALSE | 5 |
| S | 54 | 0 | 4031 | FALSE | 3 |
| S | 55 | 0 | 4031 | FALSE | 2 |
| S | 56 | 0 | 4031 | FALSE | 3 |
| S | 57 | 0 | 4031 | FALSE | 3 |
| S | 58 | 0 | 4031 | FALSE | 4 |
| S | 59 | 0 | 4031 | FALSE | 3 |
| S | 72 | 0 | 1442 | FALSE | 4 |
| S | 73 | 0 | 1442 | FALSE | 4 |
| S | 74 | 0 | 1442 | FALSE | 3 |
| S | 75 | 0 | 1442 | FALSE | 4 |
| S | 76 | 0 | 1442 | FALSE | 2 |
| S | 77 | 0 | 1442 | FALSE | 3 |
| S | 78 | 7490 | 1409 | FALSE | 7 |
| S | 79 | 5277 | 1409 | FALSE | 1 |
| S | 80 | 5277 | 1409 | FALSE | 2 |
| S | 81 | 5277 | 1409 | FALSE | 1 |
| S | 82 | 3092 | 1408 | FALSE | 3 |
| S | 83 | 3092 | 1408 | FALSE | 4 |
| S | 84 | 3092 | 1408 | FALSE | 4 |
| S | 85 | 3092 | 1408 | FALSE | 4 |
| S | 86 | 3092 | 1361 | FALSE | 0 |
| S | 87 | 2800 | 1334 | FALSE | 3 |
| S | 88 | 2800 | 1334 | FALSE | 2 |
| S | 89 | 2800 | 1331 | FALSE | 3 |
| S | 90 | 2800 | 1331 | FALSE | 3 |
| S | 91 | 2800 | 1331 | FALSE | 2 |
| S | 92 | 2800 | 1331 | FALSE | 1 |
| S | 93 | 1862 | 1316 | FALSE | 4 |
| S | 94 | 1562 | 1316 | FALSE | 3 |
| S | 95 | 1562 | 1242 | FALSE | 2 |
| S | 96 | 3635 | 1892 | FALSE | 6 |
| S | 97 | 6635 | 2014 | FALSE | 2 |
| S | 98 | 6635 | 2014 | FALSE | 4 |
| S | 99 | 6635 | 2014 | FALSE | 2 |
| S | 100 | 6635 | 2014 | FALSE | 3 |
| S | 101 | 6635 | 2278 | FALSE | 2 |
| S | 102 | 11930 | 2278 | FALSE | 7 |
| S | 103 | 11930 | 2278 | FALSE | 5 |
| S | 104 | 11930 | 2756 | FALSE | 3 |
| S | 105 | 11930 | 2756 | FALSE | 1 |
| S | 106 | 11930 | 2756 | FALSE | 2 |
| S | 107 | 11930 | 2707 | FALSE | 4 |
| S | 108 | 11930 | 2707 | FALSE | 3 |
| S | 109 | 11930 | 2707 | FALSE | 3 |
| S | 110 | 11930 | 3550 | FALSE | 3 |
| S | 111 | 1308 | 2180 | FALSE | 3 |
| S | 112 | 1308 | 2180 | FALSE | 2 |
| S | 113 | 1308 | 2180 | FALSE | 1 |
| S | 114 | 1308 | 2087 | FALSE | 3 |
| S | 115 | 1308 | 2087 | FALSE | 2 |
| S | 116 | 1308 | 2087 | FALSE | 3 |
| S | 117 | 1308 | 2087 | FALSE | 2 |
| S | 118 | 1449 | 1994 | FALSE | 2 |

| | | | | | |
|---|---|---|---|---|---|
| S | 119 | 1449 | 1994 | FALSE | 2 |
| S | 120 | 1449 | 1994 | FALSE | 1 |
| S | 121 | 2371 | 1994 | FALSE | 3 |
| S | 122 | 2411 | 1789 | FALSE | 1 |
| S | 123 | 2411 | 1789 | FALSE | 4 |
| S | 124 | 2411 | 1789 | FALSE | 3 |
| S | 125 | 2287 | 1789 | FALSE | 1 |
| S | 126 | 2532 | 1808 | FALSE | 3 |
| S | 127 | 2532 | 1543 | FALSE | 1 |
| S | 128 | 2532 | 1543 | FALSE | 3 |
| S | 129 | 2532 | 1364 | FALSE | 5 |
| S | 130 | 2532 | 1364 | FALSE | 2 |
| S | 131 | 2532 | 1364 | FALSE | 3 |
| S | 132 | 4568 | 2247 | FALSE | 4 |
| S | 133 | 5163 | 2513 | FALSE | 2 |
| S | 134 | 5163 | 2513 | FALSE | 3 |
| S | 135 | 4677 | 2513 | FALSE | 0 |
| S | 136 | 4769 | 2513 | FALSE | 2 |
| S | 137 | 4623 | 2028 | FALSE | 2 |
| S | 138 | 4623 | 2028 | FALSE | 2 |
| S | 139 | 4623 | 2028 | FALSE | 4 |
| S | 140 | 5331 | 2028 | FALSE | 4 |
| S | 141 | 5331 | 2028 | FALSE | 4 |
| S | 142 | 4533 | 1915 | FALSE | 4 |
| S | 143 | 4533 | 1915 | FALSE | 2 |
| S | 144 | 4533 | 1915 | FALSE | 5 |
| S | 145 | 4533 | 1915 | FALSE | 4 |
| S | 146 | 4533 | 1915 | FALSE | 5 |
| S | 147 | 1796 | 1216 | FALSE | 3 |
| S | 148 | 2652 | 1216 | FALSE | 4 |
| S | 149 | 5205 | 1216 | FALSE | 4 |
| S | 150 | 5797 | 1216 | FALSE | 6 |
| S | 151 | 5797 | 1216 | FALSE | 5 |
| S | 152 | 6797 | 1296 | FALSE | 3 |
| S | 153 | 11076 | 2136 | FALSE | 2 |
| S | 154 | 12180 | 6226 | FALSE | 3 |
| S | 155 | 14019 | 6226 | FALSE | 3 |
| S | 156 | 13158 | 5229 | FALSE | 4 |
| S | 157 | 14340 | 7851 | FALSE | 2 |
| S | 158 | 14340 | 7851 | FALSE | 0 |
| S | 159 | 14340 | 7851 | FALSE | 1 |
| S | 160 | 14340 | 7851 | FALSE | 1 |
| S | 161 | 14340 | 7851 | FALSE | 2 |
| S | 162 | 14340 | 7851 | FALSE | 2 |
| S | 163 | 15353 | 7851 | FALSE | 4 |
| S | 164 | 15129 | 7851 | FALSE | 2 |
| S | 165 | 13586 | 7851 | FALSE | 2 |
| S | 166 | 13586 | 7851 | FALSE | 5 |
| S | 167 | 13586 | 7851 | FALSE | 2 |
| S | 168 | 13205 | 7661 | FALSE | 2 |
| S | 169 | 12970 | 2697 | FALSE | 1 |
| S | 170 | 9608 | 2697 | FALSE | 1 |
| S | 171 | 10872 | 3425 | FALSE | 1 |
| S | 172 | 10872 | 3425 | FALSE | 4 |
| S | 173 | 10872 | 3425 | FALSE | 4 |
| S | 174 | 10872 | 3425 | FALSE | 5 |
| S | 175 | 10872 | 3425 | FALSE | 3 |
| S | 176 | 10872 | 3425 | FALSE | 3 |
| S | 177 | 10872 | 2632 | FALSE | 4 |
| S | 178 | 10872 | 2632 | FALSE | 3 |
| S | 179 | 13337 | 2632 | FALSE | 3 |
| S | 180 | 13275 | 6202 | FALSE | 5 |

| | | | | | |
|---|---|---|---|---|---|
| S | 181 | 13275 | 6202 | FALSE | 5 |
| S | 182 | 13275 | 6202 | FALSE | 0 |
| S | 183 | 9448 | 6941 | FALSE | 4 |
| S | 184 | 7168 | 8053 | FALSE | 5 |
| S | 185 | 5751 | 8053 | FALSE | 2 |
| S | 186 | 5751 | 8053 | FALSE | 3 |
| S | 187 | 5751 | 8053 | FALSE | 2 |
| S | 188 | 6461 | 8053 | FALSE | 0 |
| S | 189 | 6461 | 8053 | FALSE | 1 |
| S | 190 | 8153 | 8053 | FALSE | 3 |
| S | 191 | 8425 | 6948 | FALSE | 2 |
| S | 192 | 8425 | 8916 | FALSE | 4 |
| S | 193 | 8425 | 8916 | FALSE | 2 |
| S | 194 | 8425 | 8916 | FALSE | 2 |
| S | 195 | 7501 | 3632 | FALSE | 2 |
| S | 196 | 7501 | 3632 | FALSE | 5 |
| S | 197 | 7501 | 3632 | FALSE | 4 |
| S | 198 | 7501 | 3632 | FALSE | 5 |
| S | 199 | 7501 | 3632 | FALSE | 2 |
| S | 200 | 8647 | 3632 | FALSE | 1 |
| S | 201 | 8647 | 3632 | FALSE | 1 |
| S | 202 | 8647 | 3632 | FALSE | 3 |
| S | 203 | 8665 | 3632 | FALSE | 3 |
| S | 204 | 8665 | 3632 | FALSE | 1 |
| S | 205 | 5538 | 3632 | FALSE | 2 |
| S | 206 | 1289 | 0 | FALSE | 0 |
| S | 207 | 1289 | 0 | FALSE | 3 |
| S | 208 | 1289 | 0 | FALSE | 1 |
| S | 209 | 1289 | 0 | FALSE | 2 |
| S | 211 | 5911 | 0 | FALSE | 1 |
| S | 212 | 5911 | 0 | FALSE | 1 |
| S | 213 | 5911 | 0 | FALSE | 4 |
| S | 214 | 10607 | 0 | FALSE | 4 |
| S | 215 | 10607 | 0 | FALSE | 4 |
| S | 216 | 10607 | 0 | FALSE | 1 |
| S | 217 | 10607 | 0 | FALSE | 3 |
| S | 218 | 10607 | 0 | FALSE | 4 |
| S | 219 | 10607 | 3692 | FALSE | 3 |
| S | 220 | 10607 | 3692 | FALSE | 0 |
| S | 221 | 7561 | 3692 | FALSE | 4 |
| S | 222 | 7561 | 3692 | FALSE | 3 |
| S | 223 | 7561 | 3692 | FALSE | 2 |
| S | 224 | 9990 | 3885 | FALSE | 0 |
| S | 225 | 6795 | 3885 | FALSE | 2 |
| S | 226 | 6921 | 3885 | FALSE | 1 |
| S | 227 | 6921 | 3885 | FALSE | 2 |
| S | 228 | 6921 | 3885 | FALSE | 4 |
| S | 229 | 5524 | 3885 | FALSE | 0 |
| S | 230 | 4469 | 3885 | FALSE | 3 |
| S | 231 | 4469 | 3885 | FALSE | 1 |
| S | 232 | 4469 | 3016 | FALSE | 5 |
| S | 233 | 4469 | 3016 | FALSE | 0 |
| S | 234 | 4469 | 2677 | FALSE | 1 |
| S | 235 | 4469 | 2677 | FALSE | 2 |
| S | 236 | 4898 | 2677 | FALSE | 1 |
| S | 237 | 4898 | 2677 | FALSE | 4 |
| S | 238 | 4898 | 2677 | FALSE | 1 |
| S | 239 | 1403 | 1277 | FALSE | 2 |
| S | 240 | 1303 | 1277 | FALSE | 2 |
| S | 241 | 1303 | 1277 | FALSE | 0 |
| S | 242 | 1303 | 1277 | FALSE | 1 |
| S | 243 | 1303 | 1277 | FALSE | 4 |

| | | | | | |
|---|---|---|---|---|---|
| S | 244 | 1303 | 1277 | FALSE | 2 |
| S | 245 | 0 | 1277 | FALSE | 5 |
| S | 246 | 0 | 1277 | FALSE | 2 |
| S | 271 | 1716 | 0 | FALSE | 0 |
| S | 272 | 1627 | 0 | FALSE | 1 |
| S | 273 | 1705 | 0 | FALSE | 2 |
| S | 274 | 1705 | 0 | FALSE | 2 |
| S | 275 | 1550 | 0 | FALSE | 1 |
| S | 276 | 1490 | 0 | FALSE | 1 |
| S | 277 | 1412 | 0 | FALSE | 0 |
| S | 278 | 1412 | 0 | FALSE | 1 |
| S | 279 | 1412 | 0 | FALSE | 1 |
| S | 280 | 1412 | 0 | FALSE | 0 |
| S | 281 | 4118 | 1723 | FALSE | 2 |
| S | 282 | 4118 | 1723 | FALSE | 0 |
| S | 283 | 4118 | 2255 | FALSE | 3 |
| S | 284 | 3765 | 2255 | FALSE | 0 |
| S | 285 | 3460 | 1904 | FALSE | 0 |
| S | 286 | 4290 | 1904 | FALSE | 1 |
| S | 287 | 4634 | 1904 | FALSE | 1 |
| S | 288 | 4640 | 1904 | FALSE | 2 |
| S | 289 | 4640 | 1904 | FALSE | 1 |
| S | 290 | 5148 | 1904 | FALSE | 2 |
| S | 291 | 5798 | 1904 | FALSE | 2 |
| S | 292 | 6754 | 1904 | FALSE | 1 |
| S | 293 | 6754 | 1904 | FALSE | 3 |
| S | 294 | 6754 | 1904 | FALSE | 3 |
| S | 295 | 5873 | 1904 | FALSE | 2 |
| S | 296 | 3803 | 1995 | FALSE | 1 |
| S | 297 | 3803 | 1995 | FALSE | 2 |
| S | 298 | 4902 | 1715 | FALSE | 1 |
| S | 299 | 5418 | 1715 | FALSE | 4 |
| S | 300 | 6151 | 1970 | FALSE | 1 |
| S | 301 | 5543 | 1970 | FALSE | 2 |
| S | 302 | 5543 | 1970 | FALSE | 4 |
| S | 303 | 6452 | 1970 | FALSE | 1 |
| S | 304 | 6452 | 1970 | FALSE | 0 |
| S | 305 | 6452 | 1970 | FALSE | 0 |
| S | 306 | 6452 | 1970 | FALSE | 2 |
| S | 307 | 6452 | 1970 | FALSE | 1 |
| S | 308 | 14042 | 2031 | FALSE | 1 |
| S | 309 | 12968 | 2907 | FALSE | 3 |
| S | 310 | 21532 | 2907 | FALSE | 0 |
| S | 311 | 21532 | 2907 | FALSE | 2 |
| S | 312 | 21532 | 2907 | FALSE | 1 |
| S | 313 | 17719 | 3843 | FALSE | 0 |
| S | 314 | 17719 | 3843 | FALSE | 1 |
| S | 315 | 17719 | 3843 | FALSE | 3 |
| S | 316 | 17719 | 3843 | FALSE | 1 |
| S | 317 | 14160 | 3843 | FALSE | 1 |
| S | 318 | 14160 | 3843 | FALSE | 3 |
| S | 319 | 14160 | 3843 | FALSE | 0 |
| S | 320 | 14160 | 3533 | FALSE | 2 |
| S | 321 | 14160 | 3533 | FALSE | 1 |
| S | 322 | 11124 | 3533 | FALSE | 1 |
| S | 323 | 4469 | 4224 | FALSE | 1 |
| S | 324 | 7238 | 5630 | FALSE | 1 |
| S | 325 | 7238 | 5630 | FALSE | 1 |
| S | 326 | 7238 | 4129 | FALSE | 0 |
| S | 327 | 6314 | 4129 | FALSE | 2 |
| S | 328 | 7345 | 4129 | FALSE | 3 |
| S | 329 | 7345 | 4129 | FALSE | 4 |

| | | | | | |
|---|---|---|---|---|---|
| S | 330 | 6401 | 4129 | FALSE | 2 |
| S | 331 | 6401 | 4129 | FALSE | 2 |
| S | 332 | 6984 | 4129 | FALSE | 2 |
| S | 333 | 6984 | 4129 | FALSE | 6 |
| S | 334 | 6984 | 4129 | FALSE | 2 |
| S | 335 | 6984 | 4736 | FALSE | 4 |
| S | 336 | 6984 | 4736 | FALSE | 2 |
| S | 337 | 8272 | 4736 | FALSE | 3 |
| S | 338 | 8272 | 4736 | FALSE | 4 |
| S | 339 | 2534 | 2791 | FALSE | 4 |
| S | 340 | 6858 | 2832 | FALSE | 0 |
| S | 341 | 6858 | 3684 | FALSE | 1 |
| S | 342 | 8580 | 3684 | FALSE | 3 |
| S | 343 | 6751 | 3684 | FALSE | 1 |
| S | 344 | 6751 | 3684 | FALSE | 4 |
| S | 345 | 8442 | 3684 | FALSE | 2 |
| S | 346 | 8442 | 3684 | FALSE | 3 |
| S | 347 | 8442 | 3684 | FALSE | 1 |
| S | 348 | 8442 | 3684 | FALSE | 4 |
| S | 349 | 8442 | 3684 | FALSE | 1 |
| S | 350 | 8442 | 3684 | FALSE | 0 |
| S | 351 | 8442 | 3684 | FALSE | 2 |
| S | 352 | 8442 | 3684 | FALSE | 3 |
| S | 353 | 8442 | 3684 | FALSE | 3 |
| S | 354 | 10000 | 2914 | FALSE | 4 |
| S | 355 | 5083 | 0 | FALSE | 2 |
| S | 356 | 5083 | 0 | FALSE | 4 |
| S | 357 | 5083 | 0 | FALSE | 2 |
| S | 358 | 8903 | 0 | FALSE | 1 |
| S | 359 | 8903 | 0 | FALSE | 3 |
| S | 360 | 8903 | 0 | FALSE | 1 |
| S | 361 | 8903 | 0 | FALSE | 3 |
| S | 362 | 8903 | 0 | FALSE | 3 |
| S | 363 | 8903 | 0 | FALSE | 1 |
| S | 364 | 8903 | 0 | FALSE | 2 |
| S | 365 | 8903 | 0 | FALSE | 2 |
| S | 366 | 8903 | 0 | FALSE | 3 |
| S | 367 | 8903 | 0 | FALSE | 3 |
| S | 368 | 8903 | 0 | FALSE | 1 |
| S | 370 | 1060 | 0 | FALSE | 3 |
| S | 371 | 1060 | 0 | FALSE | 4 |
| S | 372 | 1060 | 0 | FALSE | 4 |
| S | 373 | 1060 | 0 | FALSE | 3 |
| S | 374 | 1060 | 0 | FALSE | 1 |
| S | 375 | 1060 | 0 | FALSE | 3 |
| S | 376 | 1060 | 0 | FALSE | 4 |
| S | 377 | 1060 | 0 | FALSE | 1 |
| S | 378 | 1060 | 0 | FALSE | 1 |
| S | 379 | 1060 | 0 | FALSE | 3 |
| S | 380 | 1060 | 0 | FALSE | 1 |
| S | 381 | 1060 | 3466 | FALSE | 2 |
| S | 382 | 1060 | 3466 | FALSE | 2 |
| S | 383 | 1060 | 3466 | FALSE | 3 |
| S | 384 | 1060 | 3466 | FALSE | 5 |
| S | 385 | 0 | 3466 | FALSE | 3 |
| S | 386 | 1274 | 3466 | FALSE | 1 |
| S | 387 | 1274 | 3466 | FALSE | 2 |
| S | 388 | 2122 | 3466 | FALSE | 0 |
| S | 389 | 2122 | 3466 | FALSE | 1 |
| S | 390 | 2122 | 3466 | FALSE | 3 |
| S | 391 | 2122 | 3466 | FALSE | 2 |
| S | 392 | 2122 | 3466 | FALSE | 3 |

| | | | | | |
|---|---|---|---|---|---|
| S | 393 | 2122 | 2945 | FALSE | 3 |
| S | 394 | 2122 | 3168 | FALSE | 3 |
| S | 395 | 2122 | 2645 | FALSE | 5 |
| S | 396 | 2122 | 2335 | FALSE | 1 |
| S | 397 | 2122 | 2335 | FALSE | 4 |
| S | 398 | 2122 | 2335 | FALSE | 2 |
| S | 399 | 2122 | 2335 | FALSE | 2 |
| S | 400 | 2122 | 2335 | FALSE | 1 |
| S | 401 | 2970 | 2076 | FALSE | 4 |
| S | 402 | 2970 | 2076 | FALSE | 1 |
| S | 403 | 25344 | 2076 | FALSE | 4 |
| S | 404 | 25344 | 2076 | FALSE | 5 |
| S | 405 | 29448 | 1926 | FALSE | 4 |
| S | 406 | 20947 | 2138 | FALSE | 4 |
| S | 407 | 20947 | 2138 | FALSE | 3 |
| S | 408 | 16149 | 2091 | FALSE | 4 |
| S | 409 | 16149 | 1786 | FALSE | 2 |
| S | 410 | 16149 | 1963 | FALSE | 2 |
| S | 411 | 16149 | 1877 | FALSE | 2 |
| S | 412 | 16149 | 1877 | FALSE | 3 |
| S | 413 | 16149 | 1877 | FALSE | 6 |
| S | 414 | 16149 | 1877 | FALSE | 4 |
| S | 415 | 11158 | 1877 | FALSE | 2 |
| S | 416 | 11158 | 1959 | FALSE | 2 |
| S | 417 | 11158 | 1959 | FALSE | 4 |
| S | 418 | 8320 | 1959 | FALSE | 1 |
| S | 419 | 8320 | 1959 | FALSE | 2 |
| S | 420 | 2013 | 2350 | FALSE | 1 |
| S | 421 | 1368 | 1288 | FALSE | 1 |
| S | 422 | 2087 | 1288 | FALSE | 2 |
| S | 423 | 2199 | 1288 | FALSE | 2 |
| S | 424 | 2199 | 1288 | FALSE | 4 |
| S | 425 | 2199 | 1288 | FALSE | 1 |
| S | 426 | 2199 | 1288 | FALSE | 3 |
| S | 427 | 2199 | 1288 | FALSE | 4 |
| S | 428 | 2199 | 1288 | FALSE | 1 |
| S | 429 | 2199 | 1288 | FALSE | 2 |
| S | 430 | 4246 | 1288 | FALSE | 3 |
| S | 431 | 4246 | 0 | FALSE | 1 |
| S | 432 | 4246 | 0 | FALSE | 3 |
| S | 433 | 4246 | 0 | FALSE | 2 |
| S | 434 | 4246 | 0 | FALSE | 3 |
| S | 435 | 4246 | 0 | FALSE | 4 |
| S | 436 | 4246 | 0 | FALSE | 2 |
| S | 437 | 2976 | 0 | FALSE | 1 |
| S | 438 | 2099 | 0 | FALSE | 3 |
| S | 439 | 8497 | 0 | FALSE | 2 |
| S | 440 | 7232 | 0 | FALSE | 1 |
| S | 441 | 7232 | 0 | FALSE | 2 |
| S | 442 | 7232 | 0 | FALSE | 3 |
| S | 443 | 7232 | 0 | FALSE | 1 |
| S | 444 | 6011 | 0 | FALSE | 2 |
| S | 445 | 6011 | 0 | FALSE | 1 |
| S | 446 | 6011 | 0 | FALSE | 4 |
| S | 447 | 6011 | 0 | FALSE | 3 |
| S | 448 | 6011 | 0 | FALSE | 0 |
| S | 449 | 6011 | 0 | FALSE | 1 |
| S | 450 | 6011 | 0 | FALSE | 3 |
| S | 451 | 6011 | 0 | FALSE | 2 |
| S | 452 | 6770 | 0 | FALSE | 3 |
| S | 453 | 8619 | 1148 | FALSE | 2 |
| S | 454 | 2282 | 1148 | FALSE | 2 |

| | | | | | |
|---|---|---|---|---|---|
| S | 455 | 1126 | 1148 | FALSE | 1 |
| S | 456 | 1126 | 1148 | FALSE | 0 |
| S | 457 | 1126 | 1148 | FALSE | 6 |
| S | 458 | 1126 | 1148 | FALSE | 3 |
| S | 459 | 1057 | 1148 | FALSE | 1 |
| S | 460 | 2235 | 1148 | FALSE | 3 |
| S | 461 | 2235 | 1148 | FALSE | 2 |
| S | 462 | 2235 | 1148 | FALSE | 2 |
| S | 463 | 2235 | 1295 | FALSE | 4 |
| S | 464 | 2235 | 1479 | FALSE | 1 |
| S | 465 | 6410 | 3664 | FALSE | 2 |
| S | 466 | 9789 | 3664 | FALSE | 2 |
| S | 467 | 9789 | 3664 | FALSE | 1 |
| S | 468 | 9789 | 4502 | FALSE | 2 |
| S | 469 | 9789 | 4502 | FALSE | 1 |
| S | 470 | 7664 | 3379 | FALSE | 1 |
| S | 471 | 8102 | 3390 | FALSE | 2 |
| S | 472 | 8102 | 3107 | FALSE | 1 |
| S | 473 | 8102 | 3107 | FALSE | 0 |
| S | 474 | 8884 | 3107 | FALSE | 4 |
| S | 475 | 9568 | 3107 | FALSE | 4 |
| S | 476 | 9568 | 2899 | FALSE | 4 |
| S | 477 | 9568 | 2899 | FALSE | 4 |
| S | 478 | 9568 | 3081 | FALSE | 3 |
| S | 479 | 9568 | 3257 | FALSE | 3 |
| S | 480 | 9794 | 2097 | FALSE | 2 |
| S | 481 | 5570 | 2097 | FALSE | 1 |
| S | 482 | 5570 | 4359 | FALSE | 4 |
| S | 483 | 8985 | 4359 | FALSE | 2 |
| S | 484 | 8049 | 5268 | FALSE | 5 |
| S | 485 | 10302 | 6161 | FALSE | 3 |
| S | 486 | 10528 | 7076 | FALSE | 0 |
| S | 487 | 10528 | 8266 | FALSE | 1 |
| S | 488 | 10528 | 8266 | FALSE | 3 |
| S | 489 | 10528 | 8266 | FALSE | 2 |
| S | 490 | 10528 | 8266 | FALSE | 3 |
| S | 491 | 10528 | 10023 | FALSE | 3 |
| S | 492 | 10528 | 10023 | FALSE | 1 |
| S | 493 | 10528 | 10023 | FALSE | 1 |
| S | 494 | 10528 | 10023 | FALSE | 3 |
| S | 495 | 10528 | 10023 | FALSE | 4 |
| S | 496 | 10528 | 10023 | FALSE | 3 |
| S | 497 | 10528 | 7387 | FALSE | 2 |
| S | 498 | 6490 | 7387 | FALSE | 1 |
| S | 499 | 1934 | 0 | FALSE | 3 |
| S | 500 | 1934 | 0 | FALSE | 2 |
| S | 501 | 1934 | 0 | FALSE | 3 |
| S | 502 | 1520 | 2933 | FALSE | 3 |
| S | 503 | 1520 | 2933 | FALSE | 0 |
| S | 504 | 1520 | 2933 | FALSE | 1 |
| S | 505 | 1520 | 2933 | FALSE | 1 |
| S | 506 | 1520 | 2933 | FALSE | 1 |
| S | 507 | 1520 | 2933 | FALSE | 3 |
| S | 508 | 1520 | 2933 | FALSE | 1 |
| S | 509 | 1520 | 2933 | FALSE | 2 |
| S | 510 | 1520 | 2933 | FALSE | 2 |
| S | 511 | 1520 | 2933 | FALSE | 4 |
| S | 512 | 1520 | 2933 | FALSE | 4 |
| S | 513 | 1520 | 2933 | FALSE | 3 |
| S | 514 | 1107 | 2933 | FALSE | 3 |
| S | 515 | 11566 | 2933 | FALSE | 1 |
| S | 516 | 11114 | 4244 | FALSE | 1 |

| | | | | | |
|---|---|---|---|---|---|
| S | 517 | 16117 | 4900 | FALSE | 3 |
| S | 518 | 16117 | 4900 | FALSE | 1 |
| S | 519 | 16117 | 4900 | FALSE | 2 |
| S | 520 | 16117 | 3867 | FALSE | 5 |
| S | 521 | 16117 | 3867 | FALSE | 2 |
| S | 522 | 16117 | 3867 | FALSE | 5 |
| S | 523 | 16117 | 3867 | FALSE | 3 |
| S | 524 | 16117 | 3867 | FALSE | 3 |
| S | 525 | 11281 | 3867 | FALSE | 1 |
| S | 526 | 11281 | 3867 | FALSE | 1 |
| S | 527 | 11281 | 3867 | FALSE | 2 |
| S | 528 | 11281 | 3867 | FALSE | 6 |
| S | 529 | 11281 | 3867 | FALSE | 7 |
| S | 530 | 5910 | 3867 | FALSE | 2 |
| S | 531 | 1610 | 1801 | FALSE | 2 |
| S | 532 | 2890 | 1801 | FALSE | 2 |
| S | 533 | 2890 | 1801 | FALSE | 4 |
| S | 534 | 2890 | 1801 | FALSE | 3 |
| S | 535 | 2890 | 0 | FALSE | 2 |
| S | 536 | 2784 | 0 | FALSE | 4 |
| S | 537 | 2784 | 1710 | FALSE | 2 |
| S | 538 | 2784 | 1741 | FALSE | 3 |
| S | 539 | 2784 | 1741 | FALSE | 2 |
| S | 540 | 3372 | 1548 | FALSE | 1 |
| S | 541 | 3372 | 1548 | FALSE | 4 |
| S | 542 | 2871 | 1548 | FALSE | 3 |
| S | 543 | 2871 | 1548 | FALSE | 4 |
| S | 544 | 2485 | 1548 | FALSE | 3 |
| S | 545 | 2485 | 3132 | FALSE | 4 |
| S | 546 | 2485 | 3132 | FALSE | 1 |
| S | 547 | 1924 | 3132 | FALSE | 3 |
| S | 548 | 1943 | 2628 | FALSE | 5 |
| S | 549 | 1943 | 2400 | FALSE | 1 |
| S | 550 | 1943 | 3222 | FALSE | 5 |
| S | 551 | 1733 | 3222 | FALSE | 3 |
| S | 552 | 1733 | 3190 | FALSE | 3 |
| S | 553 | 1733 | 3393 | FALSE | 1 |
| S | 554 | 3611 | 3615 | FALSE | 4 |
| S | 555 | 3611 | 3921 | FALSE | 4 |
| S | 556 | 3506 | 3921 | FALSE | 4 |
| S | 557 | 3779 | 3921 | FALSE | 1 |
| S | 558 | 3779 | 3921 | FALSE | 4 |
| S | 559 | 4269 | 3921 | FALSE | 2 |
| S | 560 | 4269 | 3355 | FALSE | 4 |
| S | 561 | 4269 | 3355 | FALSE | 3 |
| S | 562 | 4269 | 3109 | FALSE | 5 |
| S | 563 | 4836 | 3262 | FALSE | 2 |
| S | 564 | 4836 | 3399 | FALSE | 5 |
| S | 565 | 4836 | 2468 | FALSE | 2 |
| S | 566 | 4836 | 3095 | FALSE | 4 |
| S | 567 | 8959 | 3369 | FALSE | 2 |
| S | 568 | 8959 | 3369 | FALSE | 2 |
| S | 569 | 14164 | 2857 | FALSE | 2 |
| S | 570 | 9780 | 2857 | FALSE | 3 |
| S | 571 | 13233 | 2857 | FALSE | 3 |
| S | 572 | 13233 | 2857 | FALSE | 1 |
| S | 573 | 14089 | 2857 | FALSE | 3 |
| S | 574 | 10889 | 2857 | FALSE | 4 |
| S | 575 | 10889 | 2857 | FALSE | 7 |
| S | 576 | 10889 | 2753 | FALSE | 5 |
| S | 577 | 12630 | 2680 | FALSE | 3 |
| S | 578 | 12630 | 3050 | FALSE | 3 |

| | | | | | |
|---|---|---|---|---|---|
| S | 579 | 12630 | 3403 | FALSE | 5 |
| S | 580 | 12630 | 3403 | FALSE | 6 |
| S | 581 | 10778 | 1530 | FALSE | 2 |
| S | 582 | 7602 | 1482 | FALSE | 4 |
| S | 583 | 8207 | 2439 | FALSE | 4 |
| S | 584 | 8207 | 2439 | FALSE | 1 |
| S | 585 | 9406 | 2439 | FALSE | 3 |
| S | 586 | 9406 | 2439 | FALSE | 6 |
| S | 587 | 9406 | 2439 | FALSE | 3 |
| S | 588 | 8127 | 2439 | FALSE | 1 |
| S | 589 | 9837 | 2439 | FALSE | 2 |
| S | 590 | 9837 | 2439 | FALSE | 1 |
| S | 591 | 9837 | 3240 | FALSE | 4 |
| S | 592 | 6584 | 3983 | FALSE | 1 |
| S | 593 | 6584 | 3614 | FALSE | 2 |
| S | 594 | 6584 | 3652 | FALSE | 2 |
| S | 595 | 6584 | 3392 | FALSE | 2 |
| S | 596 | 9118 | 3380 | FALSE | 6 |
| S | 597 | 11833 | 3507 | FALSE | 3 |
| S | 598 | 0 | 3284 | FALSE | 2 |
| S | 599 | 0 | 3284 | FALSE | 3 |
| S | 600 | 0 | 3284 | FALSE | 3 |
| S | 601 | 0 | 3209 | FALSE | 3 |
| S | 602 | 0 | 3118 | FALSE | 3 |
| S | 603 | 0 | 3074 | FALSE | 2 |
| S | 604 | 0 | 3074 | FALSE | 2 |
| S | 605 | 0 | 2970 | FALSE | 2 |
| S | 606 | 1512 | 2694 | FALSE | 2 |
| S | 607 | 1512 | 2393 | FALSE | 2 |
| S | 608 | 1512 | 2232 | FALSE | 1 |
| S | 609 | 1512 | 1709 | FALSE | 2 |
| S | 610 | 1512 | 1783 | FALSE | 3 |
| S | 611 | 1512 | 1912 | FALSE | 3 |
| S | 612 | 1512 | 1912 | FALSE | 4 |
| S | 613 | 1512 | 2091 | FALSE | 5 |
| S | 614 | 1512 | 2091 | FALSE | 4 |
| S | 615 | 1512 | 2091 | FALSE | 3 |
| S | 616 | 1606 | 1698 | FALSE | 3 |
| S | 617 | 1606 | 1720 | FALSE | 3 |
| S | 618 | 1516 | 1601 | FALSE | 5 |
| S | 619 | 4826 | 2334 | FALSE | 1 |
| S | 620 | 4826 | 2947 | FALSE | 3 |
| S | 621 | 4065 | 2478 | FALSE | 5 |
| S | 622 | 4065 | 2478 | FALSE | 5 |
| S | 623 | 4065 | 2478 | FALSE | 4 |
| S | 624 | 4065 | 2478 | FALSE | 2 |
| S | 625 | 4065 | 2478 | FALSE | 3 |
| S | 626 | 4065 | 2478 | FALSE | 2 |
| S | 627 | 4065 | 2478 | FALSE | 2 |
| S | 628 | 4065 | 2478 | FALSE | 2 |
| S | 629 | 4065 | 2478 | FALSE | 4 |
| S | 630 | 4065 | 2478 | FALSE | 2 |
| S | 631 | 4402 | 2478 | FALSE | 4 |
| S | 632 | 4402 | 2478 | FALSE | 3 |
| S | 633 | 5593 | 2670 | FALSE | 1 |
| S | 634 | 1691 | 1540 | FALSE | 2 |
| S | 635 | 1691 | 1540 | FALSE | 1 |
| S | 649 | 1227 | 0 | FALSE | 3 |
| S | 650 | 1227 | 0 | FALSE | 2 |
| S | 651 | 1237 | 0 | FALSE | 0 |
| S | 652 | 1237 | 0 | FALSE | 3 |
| S | 653 | 1237 | 0 | FALSE | 4 |

| | | | | | |
|---|---|---|---|---|---|
| S | 654 | 1497 | 6420 | FALSE | 3 |
| S | 655 | 1497 | 6420 | FALSE | 4 |
| S | 656 | 1497 | 4224 | FALSE | 4 |
| S | 657 | 1717 | 4224 | FALSE | 3 |
| S | 658 | 1717 | 4224 | FALSE | 4 |
| S | 659 | 1717 | 4224 | FALSE | 1 |
| S | 660 | 5515 | 4224 | FALSE | 4 |
| S | 661 | 6380 | 4104 | FALSE | 4 |
| S | 662 | 6380 | 4104 | FALSE | 4 |
| S | 663 | 6380 | 4104 | FALSE | 3 |
| S | 664 | 6378 | 4104 | FALSE | 2 |
| S | 665 | 5354 | 4104 | FALSE | 6 |
| S | 666 | 5811 | 4104 | FALSE | 1 |
| S | 667 | 5811 | 4104 | FALSE | 4 |
| S | 668 | 5811 | 4104 | FALSE | 3 |
| S | 669 | 6285 | 2945 | FALSE | 2 |
| S | 670 | 6285 | 2945 | FALSE | 3 |
| S | 671 | 5813 | 3862 | FALSE | 3 |
| S | 672 | 6242 | 3862 | FALSE | 4 |
| S | 673 | 6242 | 3862 | FALSE | 3 |
| S | 674 | 6242 | 3862 | FALSE | 2 |
| S | 675 | 4176 | 3862 | FALSE | 5 |
| S | 676 | 1442 | 1760 | FALSE | 3 |
| S | 677 | 1442 | 1760 | FALSE | 6 |
| S | 678 | 1442 | 1760 | FALSE | 2 |
| S | 679 | 1518 | 1760 | FALSE | 1 |
| S | 680 | 2037 | 1568 | FALSE | 4 |
| S | 681 | 2037 | 1637 | FALSE | 4 |
| S | 682 | 2037 | 1637 | FALSE | 6 |
| S | 683 | 2037 | 1637 | FALSE | 6 |
| S | 684 | 2037 | 1637 | FALSE | 4 |
| S | 685 | 2037 | 1637 | FALSE | 2 |
| S | 686 | 0 | 1637 | FALSE | 3 |
| S | 687 | 0 | 1637 | FALSE | 3 |
| S | 688 | 0 | 1653 | FALSE | 4 |
| S | 689 | 2630 | 1653 | FALSE | 5 |
| S | 690 | 2630 | 1653 | FALSE | 2 |
| S | 691 | 2630 | 1618 | FALSE | 3 |
| S | 692 | 2630 | 1618 | FALSE | 2 |
| S | 693 | 2630 | 1618 | FALSE | 5 |
| S | 694 | 2630 | 1618 | FALSE | 5 |
| S | 695 | 2630 | 1739 | FALSE | 2 |
| S | 696 | 7478 | 1701 | FALSE | 1 |
| S | 697 | 7478 | 1701 | FALSE | 2 |
| S | 698 | 7478 | 1701 | FALSE | 1 |
| S | 699 | 7478 | 1701 | FALSE | 2 |
| S | 700 | 7342 | 1701 | FALSE | 5 |
| S | 701 | 7342 | 1701 | FALSE | 5 |
| S | 702 | 7354 | 3738 | FALSE | 4 |
| S | 703 | 6607 | 5775 | FALSE | 2 |
| S | 704 | 7270 | 5775 | FALSE | 4 |
| S | 705 | 7270 | 5775 | FALSE | 3 |
| S | 706 | 7270 | 5775 | FALSE | 5 |
| S | 707 | 7270 | 5775 | FALSE | 6 |
| S | 708 | 7270 | 5775 | FALSE | 3 |
| S | 709 | 6455 | 5775 | FALSE | 3 |
| S | 710 | 6455 | 5775 | FALSE | 2 |
| S | 711 | 5477 | 5775 | FALSE | 3 |
| S | 712 | 5477 | 5775 | FALSE | 3 |
| S | 713 | 5477 | 5775 | FALSE | 5 |
| S | 714 | 4876 | 5775 | FALSE | 2 |
| S | 715 | 4528 | 5775 | FALSE | 3 |

| | | | | | |
|---|---|---|---|---|---|
| S | 716 | 4528 | 5775 | FALSE | 5 |
| S | 717 | 2400 | 0 | FALSE | 1 |
| S | 718 | 2490 | 0 | FALSE | 2 |
| S | 719 | 2133 | 0 | FALSE | 1 |
| S | 720 | 2133 | 1101 | FALSE | 2 |
| S | 721 | 2133 | 1829 | FALSE | 5 |
| S | 722 | 1914 | 1829 | FALSE | 2 |
| S | 723 | 1914 | 1829 | FALSE | 3 |
| S | 724 | 2611 | 1829 | FALSE | 3 |
| S | 725 | 3448 | 3176 | FALSE | 4 |
| S | 726 | 3448 | 2980 | FALSE | 2 |
| S | 727 | 3448 | 2980 | FALSE | 4 |
| S | 728 | 3266 | 2967 | FALSE | 4 |
| S | 729 | 3420 | 2967 | FALSE | 5 |
| S | 730 | 3157 | 2967 | FALSE | 3 |
| S | 731 | 3157 | 2967 | FALSE | 5 |
| S | 732 | 3157 | 2967 | FALSE | 5 |
| S | 733 | 3157 | 2967 | FALSE | 2 |
| S | 734 | 3332 | 2967 | FALSE | 3 |
| S | 735 | 3332 | 3340 | TRUE | 4 |
| S | 736 | 3332 | 3536 | TRUE | 1 |
| S | 737 | 3540 | 3536 | TRUE | 5 |
| S | 738 | 3540 | 3536 | TRUE | 3 |
| S | 739 | 4116 | 3536 | TRUE | 5 |
| S | 740 | 3470 | 2549 | TRUE | 5 |
| S | 741 | 3470 | 2902 | TRUE | 2 |
| S | 742 | 3470 | 2902 | TRUE | 2 |
| S | 743 | 4799 | 0 | TRUE | 3 |
| S | 744 | 4799 | 0 | TRUE | 2 |
| S | 745 | 8372 | 0 | FALSE | 2 |
| S | 746 | 8372 | 0 | TRUE | 3 |
| S | 747 | 8372 | 0 | FALSE | 3 |
| S | 748 | 6526 | 4182 | FALSE | 5 |
| S | 749 | 6526 | 4182 | TRUE | 3 |
| S | 750 | 6526 | 4182 | FALSE | 2 |
| S | 751 | 7356 | 4182 | FALSE | 0 |
| S | 752 | 7356 | 4182 | FALSE | 4 |
| S | 753 | 7356 | 4182 | FALSE | 4 |
| S | 754 | 7017 | 4182 | FALSE | 4 |
| S | 755 | 7017 | 4182 | FALSE | 2 |
| S | 756 | 7017 | 2843 | FALSE | 2 |
| S | 757 | 7017 | 2364 | FALSE | 6 |
| S | 758 | 7017 | 2364 | FALSE | 3 |
| S | 759 | 7017 | 2248 | FALSE | 2 |
| S | 760 | 7017 | 2248 | FALSE | 4 |
| S | 761 | 7017 | 2248 | FALSE | 2 |
| S | 762 | 5306 | 2248 | FALSE | 7 |
| S | 763 | 4573 | 1604 | FALSE | 3 |
| S | 764 | 6421 | 1604 | FALSE | 3 |
| S | 765 | 5879 | 1604 | FALSE | 7 |
| S | 766 | 4317 | 1604 | FALSE | 5 |
| S | 767 | 3963 | 1604 | FALSE | 2 |
| S | 768 | 3770 | 1604 | FALSE | 1 |
| S | 769 | 3770 | 3446 | FALSE | 4 |
| S | 770 | 3460 | 3446 | FALSE | 2 |
| S | 771 | 3291 | 3358 | FALSE | 5 |
| S | 772 | 7067 | 3717 | FALSE | 5 |
| S | 773 | 7564 | 4266 | FALSE | 3 |
| S | 774 | 7564 | 4857 | FALSE | 2 |
| S | 775 | 7564 | 4857 | FALSE | 5 |
| S | 776 | 7564 | 4857 | FALSE | 2 |
| S | 777 | 8167 | 4857 | FALSE | 5 |

| | | | | | |
|---|---|---|---|---|---|
| S | 778 | 8514 | 4857 | FALSE | 3 |
| S | 779 | 8267 | 4717 | FALSE | 5 |
| S | 780 | 8737 | 4717 | FALSE | 4 |
| S | 781 | 10162 | 4717 | FALSE | 2 |
| S | 782 | 11165 | 4717 | FALSE | 3 |
| S | 783 | 12307 | 4717 | FALSE | 4 |
| S | 784 | 12307 | 3652 | FALSE | 4 |
| S | 785 | 15771 | 3652 | FALSE | 5 |
| S | 786 | 18673 | 4486 | FALSE | 4 |
| S | 787 | 10049 | 5307 | FALSE | 3 |
| S | 788 | 0 | 4155 | FALSE | 3 |
| S | 789 | 0 | 4155 | FALSE | 4 |
| S | 790 | 0 | 4155 | FALSE | 1 |
| S | 791 | 0 | 4155 | FALSE | 2 |
| S | 792 | 0 | 4155 | FALSE | 3 |
| S | 793 | 0 | 4155 | FALSE | 3 |
| S | 805 | 0 | 2222 | FALSE | 1 |
| S | 806 | 0 | 2222 | FALSE | 1 |
| S | 807 | 0 | 2222 | FALSE | 2 |
| S | 808 | 0 | 2222 | FALSE | 6 |
| S | 809 | 0 | 2222 | FALSE | 4 |
| S | 810 | 0 | 2222 | FALSE | 3 |
| S | 811 | 0 | 2222 | FALSE | 3 |
| S | 812 | 2663 | 2222 | FALSE | 4 |
| S | 813 | 2663 | 1638 | TRUE | 3 |
| S | 814 | 2663 | 1638 | FALSE | 2 |
| S | 815 | 2663 | 1638 | TRUE | 3 |
| S | 816 | 2663 | 1638 | TRUE | 1 |
| S | 817 | 14810 | 1638 | TRUE | 2 |
| S | 818 | 13438 | 2140 | TRUE | 3 |
| S | 819 | 14745 | 2499 | TRUE | 1 |
| S | 820 | 14745 | 2569 | TRUE | 2 |
| S | 821 | 14745 | 2569 | TRUE | 3 |
| S | 822 | 14745 | 2569 | TRUE | 3 |
| S | 823 | 14745 | 2569 | TRUE | 2 |
| S | 824 | 13467 | 2569 | TRUE | 1 |
| S | 825 | 13467 | 2569 | TRUE | 2 |
| S | 826 | 13467 | 2569 | TRUE | 4 |
| S | 827 | 15267 | 2569 | FALSE | 2 |
| S | 828 | 15267 | 3074 | TRUE | 4 |
| S | 829 | 15267 | 2827 | TRUE | 3 |
| S | 830 | 13759 | 2827 | FALSE | 3 |
| S | 831 | 12164 | 2827 | TRUE | 3 |
| S | 832 | 9257 | 2827 | TRUE | 3 |
| S | 833 | 9245 | 2720 | TRUE | 2 |
| S | 834 | 3835 | 2085 | TRUE | 4 |
| S | 835 | 3835 | 2085 | TRUE | 3 |
| S | 836 | 3835 | 2085 | FALSE | 3 |
| S | 837 | 3835 | 2085 | FALSE | 3 |
| S | 838 | 3835 | 2085 | FALSE | 1 |
| S | 839 | 2853 | 2085 | FALSE | 3 |
| S | 840 | 2853 | 2085 | FALSE | 2 |
| S | 841 | 2853 | 2085 | FALSE | 3 |
| S | 842 | 2853 | 2085 | FALSE | 3 |
| S | 843 | 2853 | 2085 | FALSE | 0 |
| S | 844 | 2853 | 0 | FALSE | 2 |
| S | 845 | 1001 | 0 | FALSE | 2 |
| S | 846 | 2162 | 0 | FALSE | 3 |
| S | 847 | 9089 | 0 | FALSE | 5 |
| S | 848 | 9089 | 0 | TRUE | 4 |
| S | 849 | 9089 | 0 | TRUE | 4 |
| S | 850 | 9089 | 0 | TRUE | 4 |

| | | | | | |
|---|---|---|---|---|---|
| S | 851 | 9089 | 0 | TRUE | 5 |
| S | 852 | 9089 | 0 | TRUE | 3 |
| S | 853 | 9089 | 0 | TRUE | 4 |
| S | 854 | 9089 | 0 | FALSE | 5 |
| S | 855 | 9089 | 0 | TRUE | 2 |
| S | 856 | 6777 | 0 | TRUE | 4 |
| S | 857 | 6777 | 4686 | TRUE | 5 |
| S | 858 | 6777 | 4686 | TRUE | 3 |
| S | 859 | 6777 | 4686 | TRUE | 3 |
| S | 860 | 6777 | 4686 | TRUE | 4 |
| S | 861 | 9084 | 4686 | TRUE | 3 |
| S | 862 | 2151 | 4686 | FALSE | 3 |
| S | 863 | 3915 | 4686 | FALSE | 3 |
| S | 864 | 3838 | 4686 | FALSE | 4 |
| S | 865 | 3838 | 2524 | FALSE | 3 |
| S | 866 | 4831 | 2524 | FALSE | 1 |
| S | 867 | 4831 | 2132 | FALSE | 3 |
| S | 868 | 5679 | 2711 | FALSE | 2 |
| S | 869 | 5679 | 2711 | FALSE | 3 |
| S | 870 | 5679 | 2711 | FALSE | 3 |
| S | 871 | 6071 | 2711 | FALSE | 5 |
| S | 872 | 6071 | 2382 | FALSE | 4 |
| S | 873 | 6071 | 2382 | FALSE | 4 |
| S | 874 | 6071 | 2382 | FALSE | 4 |
| S | 875 | 6071 | 2382 | FALSE | 5 |
| S | 876 | 6071 | 2382 | FALSE | 2 |
| S | 877 | 6071 | 2382 | FALSE | 4 |
| S | 878 | 8325 | 2382 | FALSE | 2 |
| S | 879 | 9975 | 2382 | FALSE | 7 |
| S | 880 | 9975 | 2851 | FALSE | 3 |
| S | 881 | 9071 | 2851 | FALSE | 4 |
| S | 882 | 9071 | 4159 | FALSE | 3 |
| S | 887 | 1032 | 0 | FALSE | 2 |
| S | 888 | 1032 | 0 | FALSE | 2 |
| S | 889 | 1032 | 0 | FALSE | 2 |
| S | 890 | 1032 | 0 | FALSE | 4 |
| S | 891 | 1032 | 0 | FALSE | 2 |
| S | 892 | 1032 | 0 | FALSE | 5 |
| S | 893 | 1032 | 0 | FALSE | 4 |
| S | 894 | 1032 | 0 | FALSE | 3 |
| S | 895 | 1032 | 0 | FALSE | 4 |
| S | 896 | 1452 | 0 | FALSE | 5 |
| S | 897 | 1452 | 3703 | TRUE | 3 |
| S | 898 | 1452 | 3095 | TRUE | 2 |
| S | 899 | 1452 | 2402 | FALSE | 4 |
| S | 900 | 1452 | 2188 | TRUE | 5 |
| S | 901 | 1452 | 2188 | TRUE | 1 |
| S | 902 | 1872 | 2188 | TRUE | 2 |
| S | 903 | 1872 | 2188 | TRUE | 5 |
| S | 904 | 1872 | 2188 | FALSE | 1 |
| S | 905 | 1872 | 2188 | TRUE | 5 |
| S | 906 | 1872 | 2188 | TRUE | 4 |
| S | 907 | 1872 | 2188 | TRUE | 2 |
| S | 908 | 1872 | 2286 | TRUE | 1 |
| S | 909 | 1872 | 2192 | TRUE | 5 |
| S | 910 | 1872 | 3633 | TRUE | 1 |
| S | 911 | 0 | 3633 | TRUE | 4 |
| S | 912 | 0 | 3622 | TRUE | 1 |
| S | 913 | 0 | 3848 | TRUE | 4 |
| S | 914 | 3158 | 4557 | TRUE | 4 |
| S | 915 | 3158 | 5560 | TRUE | 5 |
| S | 916 | 3158 | 5560 | TRUE | 4 |

| | | | | | |
|---|---|---|---|---|---|
| S | 917 | 14947 | 5560 | TRUE | 4 |
| S | 918 | 13816 | 4889 | TRUE | 3 |
| S | 919 | 10642 | 4889 | TRUE | 1 |
| S | 920 | 10642 | 4889 | TRUE | 6 |
| S | 921 | 10642 | 4889 | TRUE | 3 |
| S | 922 | 10642 | 4889 | TRUE | 5 |
| S | 923 | 10642 | 5625 | TRUE | 4 |
| S | 924 | 10642 | 7576 | TRUE | 6 |
| S | 925 | 10642 | 2874 | TRUE | 2 |
| S | 926 | 10642 | 2874 | TRUE | 5 |
| S | 927 | 10642 | 2874 | TRUE | 4 |
| S | 928 | 7698 | 2874 | TRUE | 3 |
| S | 929 | 8606 | 2874 | FALSE | 2 |
| S | 930 | 8606 | 2874 | TRUE | 5 |
| S | 931 | 8606 | 2874 | TRUE | 3 |
| S | 932 | 4073 | 3070 | FALSE | 4 |
| S | 933 | 1580 | 3267 | FALSE | 2 |
| S | 934 | 1810 | 3267 | FALSE | 2 |
| S | 935 | 1810 | 3267 | FALSE | 5 |
| S | 936 | 1728 | 3267 | FALSE | 5 |
| S | 937 | 1728 | 3267 | FALSE | 1 |
| S | 938 | 1728 | 3267 | FALSE | 3 |
| S | 939 | 1728 | 3267 | FALSE | 3 |
| S | 940 | 1728 | 3267 | FALSE | 3 |
| S | 941 | 1728 | 3267 | FALSE | 5 |
| S | 942 | 1728 | 3267 | FALSE | 3 |
| S | 943 | 1647 | 3267 | FALSE | 2 |
| S | 944 | 1647 | 3267 | FALSE | 3 |
| S | 945 | 1647 | 3267 | FALSE | 1 |
| S | 946 | 1647 | 3267 | FALSE | 5 |
| S | 947 | 1647 | 0 | FALSE | 1 |
| S | 948 | 1647 | 0 | FALSE | 3 |
| S | 949 | 1647 | 0 | FALSE | 4 |
| S | 950 | 1647 | 0 | FALSE | 5 |
| S | 955 | 1123 | 0 | FALSE | 3 |
| S | 956 | 1123 | 0 | FALSE | 1 |
| S | 957 | 1123 | 0 | FALSE | 3 |
| S | 958 | 1123 | 0 | FALSE | 3 |
| S | 959 | 1123 | 0 | FALSE | 5 |
| S | 960 | 1123 | 0 | TRUE | 4 |
| S | 961 | 1123 | 0 | FALSE | 5 |
| S | 962 | 1123 | 0 | TRUE | 1 |
| S | 963 | 1123 | 0 | TRUE | 3 |
| S | 964 | 1123 | 0 | FALSE | 3 |
| S | 965 | 1123 | 0 | TRUE | 1 |
| S | 966 | 1123 | 0 | TRUE | 1 |
| S | 967 | 1123 | 0 | FALSE | 5 |
| S | 968 | 1123 | 0 | FALSE | 4 |
| S | 969 | 1123 | 0 | TRUE | 3 |
| S | 974 | 0 | 6996 | TRUE | 1 |
| S | 975 | 0 | 6996 | FALSE | 3 |
| S | 976 | 0 | 4827 | FALSE | 2 |
| S | 977 | 1862 | 4827 | TRUE | 1 |
| S | 978 | 1574 | 4827 | TRUE | 3 |
| S | 979 | 1711 | 4827 | FALSE | 2 |
| S | 980 | 1662 | 4827 | TRUE | 2 |
| S | 981 | 1662 | 4827 | TRUE | 3 |
| S | 982 | 1662 | 4827 | TRUE | 2 |
| S | 983 | 1662 | 4827 | TRUE | 3 |
| S | 984 | 1662 | 4827 | TRUE | 1 |
| S | 985 | 1662 | 4827 | TRUE | 1 |
| S | 986 | 1662 | 4150 | FALSE | 3 |

| | | | | | |
|---|---|---|---|---|---|
| S | 987 | 4508 | 4150 | TRUE | 3 |
| S | 988 | 5971 | 6508 | TRUE | 2 |
| S | 989 | 6037 | 6386 | TRUE | 4 |
| S | 990 | 6735 | 5679 | TRUE | 4 |
| S | 991 | 6735 | 6435 | FALSE | 2 |
| S | 992 | 7178 | 6435 | TRUE | 4 |
| S | 993 | 7767 | 6435 | TRUE | 3 |
| S | 994 | 8409 | 6435 | TRUE | 5 |
| S | 995 | 10358 | 6435 | TRUE | 3 |
| S | 996 | 10358 | 6435 | TRUE | 1 |
| S | 997 | 10358 | 6435 | TRUE | 2 |
| S | 998 | 10358 | 6435 | TRUE | 3 |
| S | 999 | 10358 | 6435 | TRUE | 5 |
| S | 1000 | 10358 | 6435 | TRUE | 3 |
| S | 1001 | 10358 | 7648 | TRUE | 3 |
| S | 1002 | 8962 | 7648 | TRUE | 3 |
| S | 1003 | 10555 | 2853 | TRUE | 3 |
| S | 1004 | 7717 | 3073 | TRUE | 4 |
| S | 1005 | 1026 | 2500 | FALSE | 4 |
| S | 1006 | 1026 | 2500 | FALSE | 2 |
| S | 1007 | 1026 | 2500 | FALSE | 3 |
| S | 1008 | 1026 | 2500 | FALSE | 3 |
| S | 1009 | 1026 | 2500 | FALSE | 3 |
| S | 1010 | 1026 | 2500 | FALSE | 3 |
| S | 1011 | 1026 | 2500 | FALSE | 2 |
| S | 1012 | 1026 | 2500 | FALSE | 3 |
| S | 1013 | 1026 | 2500 | FALSE | 2 |
| S | 1014 | 1026 | 2500 | FALSE | 1 |
| S | 1015 | 1026 | 2500 | FALSE | 6 |
| S | 1016 | 1026 | 2500 | FALSE | 4 |
| S | 1017 | 3776 | 2725 | FALSE | 2 |
| S | 1018 | 3776 | 2725 | FALSE | 3 |
| S | 1019 | 6526 | 2442 | FALSE | 2 |
| S | 1020 | 6526 | 3177 | FALSE | 4 |
| S | 1021 | 6526 | 3177 | FALSE | 2 |
| S | 1022 | 6526 | 3177 | FALSE | 4 |
| S | 1023 | 4240 | 3177 | FALSE | 0 |
| S | 1024 | 3183 | 3177 | FALSE | 3 |
| S | 1025 | 3183 | 3177 | FALSE | 4 |
| S | 1026 | 3183 | 3177 | FALSE | 2 |
| S | 1027 | 3183 | 3177 | FALSE | 5 |
| S | 1028 | 3183 | 3177 | FALSE | 0 |
| S | 1029 | 3183 | 3177 | FALSE | 3 |
| S | 1030 | 5436 | 3177 | TRUE | 3 |
| S | 1031 | 4822 | 2243 | TRUE | 2 |
| S | 1032 | 4481 | 1309 | TRUE | 3 |
| S | 1033 | 4481 | 1309 | TRUE | 4 |
| S | 1034 | 4481 | 1309 | TRUE | 1 |
| S | 1035 | 4481 | 1309 | TRUE | 4 |
| S | 1036 | 4481 | 1309 | TRUE | 1 |
| S | 1037 | 4481 | 1309 | TRUE | 2 |
| S | 1038 | 5113 | 1309 | FALSE | 3 |
| S | 1039 | 5219 | 1342 | TRUE | 4 |
| S | 1040 | 5219 | 1342 | TRUE | 2 |
| S | 1041 | 5219 | 1342 | FALSE | 3 |
| S | 1042 | 5219 | 1342 | TRUE | 1 |
| S | 1043 | 5219 | 1342 | TRUE | 1 |
| S | 1044 | 5219 | 1342 | TRUE | 5 |
| S | 1045 | 2893 | 1342 | TRUE | 2 |
| S | 1046 | 1289 | 2781 | TRUE | 1 |
| S | 1047 | 1419 | 3165 | TRUE | 1 |
| S | 1048 | 1439 | 2981 | TRUE | 2 |

| | | | | | |
|---|---|---|---|---|---|
| S | 1049 | 1439 | 2981 | TRUE | 4 |
| S | 1050 | 1402 | 2981 | TRUE | 3 |
| S | 1051 | 1402 | 2981 | TRUE | 2 |
| S | 1052 | 1402 | 2981 | FALSE | 2 |
| S | 1053 | 1402 | 2981 | FALSE | 2 |
| S | 1054 | 1379 | 3516 | FALSE | 5 |
| S | 1055 | 1379 | 3516 | FALSE | 3 |
| S | 1056 | 1379 | 3516 | FALSE | 3 |
| S | 1057 | 1891 | 3516 | TRUE | 6 |
| S | 1058 | 1891 | 3516 | FALSE | 2 |
| S | 1059 | 1891 | 3475 | TRUE | 3 |
| S | 1060 | 1891 | 3475 | TRUE | 3 |
| S | 1061 | 2092 | 3237 | TRUE | 2 |
| S | 1062 | 2229 | 2889 | TRUE | 2 |
| S | 1063 | 2594 | 3351 | TRUE | 3 |
| S | 1064 | 2594 | 3351 | TRUE | 1 |
| S | 1065 | 3938 | 3351 | TRUE | 6 |
| S | 1066 | 2938 | 3351 | TRUE | 5 |
| S | 1067 | 2938 | 3351 | TRUE | 0 |
| S | 1068 | 2938 | 3351 | FALSE | 3 |
| S | 1069 | 2554 | 3351 | TRUE | 2 |
| S | 1070 | 2310 | 3351 | TRUE | 2 |
| S | 1071 | 9441 | 3239 | FALSE | 3 |
| S | 1072 | 9914 | 3040 | FALSE | 4 |
| S | 1073 | 8573 | 3040 | FALSE | 4 |
| S | 1074 | 7995 | 2937 | FALSE | 4 |
| S | 1075 | 7262 | 2937 | FALSE | 2 |
| S | 1076 | 6611 | 2937 | FALSE | 2 |
| S | 1077 | 6213 | 2937 | FALSE | 1 |
| S | 1078 | 6014 | 2937 | FALSE | 4 |
| S | 1079 | 5477 | 2937 | FALSE | 4 |
| S | 1080 | 5477 | 2453 | FALSE | 3 |
| S | 1081 | 5478 | 2453 | FALSE | 3 |
| S | 1082 | 5444 | 2453 | FALSE | 2 |
| S | 1083 | 5153 | 2453 | FALSE | 1 |
| S | 1084 | 5394 | 2453 | FALSE | 8 |
| S | 1085 | 5687 | 2358 | FALSE | 6 |
| S | 1086 | 2997 | 1808 | FALSE | 4 |
| S | 1087 | 2671 | 1491 | FALSE | 4 |
| S | 1088 | 2744 | 1491 | FALSE | 2 |
| S | 1089 | 2566 | 1491 | FALSE | 3 |
| S | 1090 | 2614 | 1491 | FALSE | 3 |
| S | 1091 | 3781 | 1491 | FALSE | 5 |
| S | 1092 | 4406 | 4328 | FALSE | 0 |
| S | 1093 | 4449 | 4219 | FALSE | 2 |
| S | 1094 | 5000 | 4219 | FALSE | 3 |
| S | 1095 | 5000 | 5024 | FALSE | 5 |
| S | 1096 | 5507 | 5024 | FALSE | 2 |
| S | 1097 | 5608 | 5024 | FALSE | 4 |
| S | 1098 | 5623 | 4437 | FALSE | 1 |
| S | 1099 | 5623 | 3963 | FALSE | 6 |
| S | 1100 | 5623 | 4360 | FALSE | 2 |
| S | 1101 | 5623 | 4360 | FALSE | 2 |
| S | 1102 | 6432 | 4360 | FALSE | 5 |
| S | 1103 | 6432 | 4360 | FALSE | 3 |
| S | 1104 | 6432 | 4360 | FALSE | 4 |
| S | 1105 | 6432 | 4192 | FALSE | 5 |
| S | 1106 | 4605 | 3739 | FALSE | 2 |
| S | 1107 | 1154 | 2368 | FALSE | 6 |
| S | 1108 | 1094 | 2014 | FALSE | 4 |
| S | 1109 | 1094 | 2014 | FALSE | 2 |
| S | 1110 | 3635 | 2014 | FALSE | 2 |

| | | | | | |
|---|---|---|---|---|---|
| S | 1111 | 6178 | 1845 | FALSE | 4 |
| S | 1112 | 6178 | 1845 | FALSE | 2 |
| S | 1113 | 7434 | 1796 | FALSE | 5 |
| S | 1114 | 7434 | 1846 | FALSE | 3 |
| S | 1115 | 7434 | 1846 | FALSE | 2 |
| S | 1116 | 7434 | 1846 | FALSE | 5 |
| S | 1117 | 7592 | 1846 | FALSE | 3 |
| S | 1118 | 7592 | 1846 | FALSE | 3 |
| S | 1119 | 7592 | 1863 | FALSE | 3 |
| S | 1120 | 7592 | 1435 | FALSE | 2 |
| S | 1121 | 7592 | 1538 | FALSE | 3 |
| S | 1122 | 7592 | 1538 | FALSE | 4 |
| S | 1123 | 9232 | 1538 | FALSE | 4 |
| S | 1124 | 9232 | 1538 | FALSE | 5 |
| S | 1125 | 9404 | 1538 | FALSE | 1 |
| S | 1126 | 8223 | 1571 | FALSE | 0 |
| S | 1127 | 8223 | 1571 | FALSE | 2 |
| S | 1128 | 8223 | 1571 | FALSE | 2 |
| S | 1129 | 4869 | 1571 | FALSE | 5 |
| S | 1130 | 4869 | 1571 | FALSE | 3 |
| S | 1131 | 3149 | 3579 | FALSE | 3 |
| S | 1132 | 1458 | 3579 | FALSE | 2 |
| S | 1133 | 1458 | 3579 | FALSE | 5 |
| S | 1134 | 4053 | 4402 | FALSE | 2 |
| S | 1135 | 4053 | 7595 | FALSE | 3 |
| S | 1136 | 4053 | 7595 | FALSE | 2 |
| S | 1137 | 3845 | 7595 | FALSE | 2 |
| S | 1138 | 3865 | 7595 | FALSE | 2 |
| S | 1139 | 3865 | 7595 | FALSE | 3 |
| S | 1140 | 3758 | 7595 | FALSE | 4 |
| S | 1141 | 3705 | 7595 | FALSE | 3 |
| S | 1142 | 3705 | 7595 | FALSE | 2 |
| S | 1143 | 5874 | 4407 | TRUE | 4 |
| S | 1144 | 6752 | 3670 | FALSE | 2 |
| S | 1145 | 6752 | 3670 | TRUE | 4 |
| S | 1146 | 6917 | 2689 | FALSE | 4 |
| S | 1147 | 6554 | 2689 | TRUE | 2 |
| S | 1148 | 6554 | 2689 | TRUE | 2 |
| S | 1149 | 8823 | 2689 | FALSE | 6 |
| S | 1150 | 9538 | 3935 | FALSE | 5 |
| S | 1151 | 9308 | 3327 | FALSE | 3 |
| S | 1152 | 9570 | 3145 | TRUE | 3 |
| S | 1153 | 10085 | 3145 | FALSE | 5 |
| S | 1154 | 10085 | 3145 | FALSE | 4 |
| S | 1155 | 11197 | 3145 | TRUE | 5 |
| S | 1156 | 11190 | 3145 | TRUE | 1 |
| S | 1157 | 10693 | 3145 | FALSE | 2 |
| S | 1158 | 9413 | 3290 | TRUE | 2 |
| S | 1159 | 9311 | 3477 | FALSE | 3 |
| S | 1160 | 8879 | 3324 | TRUE | 4 |
| S | 1161 | 9655 | 3146 | FALSE | 1 |
| S | 1162 | 9653 | 3146 | FALSE | 8 |
| S | 1163 | 9192 | 3146 | FALSE | 4 |
| S | 1164 | 6384 | 3146 | FALSE | 1 |
| S | 1165 | 3277 | 2128 | FALSE | 4 |
| S | 1166 | 2431 | 2381 | FALSE | 1 |
| S | 1167 | 2431 | 2693 | FALSE | 4 |
| S | 1168 | 2431 | 2693 | FALSE | 4 |
| S | 1169 | 2431 | 2693 | FALSE | 3 |
| S | 1170 | 2040 | 2693 | FALSE | 2 |
| S | 1171 | 1881 | 3723 | FALSE | 5 |
| S | 1172 | 2008 | 3723 | FALSE | 1 |

| | | | | | |
|---|---|---|---|---|---|
| S | 1173 | 1736 | 3913 | FALSE | 3 |
| S | 1174 | 1736 | 2955 | FALSE | 2 |
| S | 1175 | 1778 | 3345 | FALSE | 2 |
| S | 1176 | 1778 | 4422 | FALSE | 3 |
| S | 1177 | 1953 | 4422 | FALSE | 3 |
| S | 1178 | 1804 | 4422 | FALSE | 4 |
| S | 1179 | 1455 | 4422 | FALSE | 1 |
| S | 1180 | 1843 | 4422 | FALSE | 2 |
| S | 1181 | 8304 | 3322 | FALSE | 4 |
| S | 1182 | 7943 | 3407 | FALSE | 3 |
| S | 1183 | 7943 | 3407 | FALSE | 3 |
| S | 1184 | 7943 | 3407 | FALSE | 4 |
| S | 1185 | 7943 | 3407 | FALSE | 4 |
| S | 1186 | 7943 | 2298 | FALSE | 2 |
| S | 1187 | 7305 | 2213 | FALSE | 3 |
| S | 1188 | 8055 | 2971 | FALSE | 3 |
| S | 1189 | 8582 | 3300 | FALSE | 4 |
| S | 1190 | 8582 | 3300 | FALSE | 5 |
| S | 1191 | 8582 | 3300 | FALSE | 4 |
| S | 1192 | 8508 | 3300 | FALSE | 4 |
| S | 1193 | 8737 | 3057 | FALSE | 3 |
| S | 1194 | 8694 | 3057 | FALSE | 1 |
| S | 1195 | 9073 | 3905 | FALSE | 3 |
| S | 1196 | 7452 | 4253 | FALSE | 2 |
| S | 1197 | 7510 | 4492 | FALSE | 3 |
| S | 1198 | 7510 | 4492 | FALSE | 1 |
| S | 1199 | 7510 | 4492 | FALSE | 2 |
| S | 1200 | 7024 | 4492 | FALSE | 4 |
| S | 1201 | 7024 | 4205 | FALSE | 3 |
| S | 1202 | 7868 | 4187 | FALSE | 4 |
| S | 1203 | 7628 | 3993 | TRUE | 2 |
| S | 1204 | 7628 | 3711 | TRUE | 4 |
| S | 1205 | 7628 | 3711 | TRUE | 3 |
| S | 1206 | 9855 | 3713 | TRUE | 2 |
| S | 1207 | 10189 | 3641 | TRUE | 4 |
| S | 1208 | 10855 | 3867 | TRUE | 3 |
| S | 1209 | 11552 | 3867 | TRUE | 1 |
| S | 1210 | 13508 | 3008 | TRUE | 1 |
| S | 1211 | 13508 | 3008 | TRUE | 4 |
| S | 1212 | 13508 | 3008 | TRUE | 3 |
| S | 1213 | 13508 | 3008 | TRUE | 2 |
| S | 1214 | 13508 | 3008 | TRUE | 4 |
| S | 1215 | 17613 | 3008 | TRUE | 4 |
| S | 1216 | 17613 | 3096 | TRUE | 7 |
| S | 1217 | 20429 | 2818 | TRUE | 6 |
| S | 1218 | 20429 | 2818 | TRUE | 2 |
| S | 1219 | 20429 | 3358 | FALSE | 6 |
| S | 1220 | 20429 | 3358 | FALSE | 1 |
| S | 1221 | 10957 | 2991 | FALSE | 1 |
| S | 1223 | 0 | 1975 | FALSE | 3 |
| S | 1224 | 0 | 1975 | FALSE | 1 |
| S | 1225 | 0 | 1975 | FALSE | 2 |
| S | 1226 | 0 | 1975 | FALSE | 4 |
| S | 1227 | 0 | 1975 | FALSE | 2 |
| S | 1228 | 0 | 1975 | FALSE | 2 |
| S | 1229 | 0 | 1975 | FALSE | 5 |
| S | 1230 | 0 | 1975 | FALSE | 4 |
| S | 1231 | 0 | 1975 | FALSE | 1 |
| S | 1232 | 0 | 1975 | FALSE | 2 |
| S | 1233 | 0 | 1975 | FALSE | 4 |
| S | 1234 | 0 | 1975 | FALSE | 3 |
| S | 1235 | 0 | 1975 | FALSE | 3 |

| | | | | | |
|---|---|---|---|---|---|
| S | 1236 | 0 | 1975 | FALSE | 2 |
| S | 1237 | 0 | 1975 | FALSE | 6 |
| S | 1246 | 1862 | 1088 | FALSE | 2 |
| S | 1247 | 1493 | 1088 | FALSE | 2 |
| S | 1248 | 1652 | 1159 | FALSE | 3 |
| S | 1249 | 1677 | 1159 | FALSE | 3 |
| S | 1250 | 1762 | 1159 | FALSE | 2 |
| S | 1251 | 1795 | 1159 | FALSE | 2 |
| S | 1252 | 1807 | 1159 | FALSE | 3 |
| S | 1253 | 1769 | 1159 | FALSE | 2 |
| S | 1254 | 1742 | 1159 | FALSE | 2 |
| S | 1255 | 1747 | 1223 | FALSE | 2 |
| S | 1256 | 1725 | 1719 | FALSE | 0 |
| S | 1257 | 1724 | 1719 | FALSE | 4 |
| S | 1258 | 2709 | 2597 | FALSE | 2 |
| S | 1259 | 2680 | 2597 | FALSE | 3 |
| S | 1260 | 2680 | 2597 | FALSE | 3 |
| S | 1261 | 3015 | 2813 | FALSE | 3 |
| S | 1262 | 3228 | 2813 | FALSE | 4 |
| S | 1263 | 3370 | 3460 | FALSE | 4 |
| S | 1264 | 3640 | 3460 | FALSE | 6 |
| S | 1265 | 3849 | 3460 | FALSE | 4 |
| S | 1266 | 4138 | 3460 | FALSE | 2 |
| S | 1267 | 4278 | 3460 | FALSE | 3 |
| S | 1268 | 4587 | 3460 | FALSE | 3 |
| S | 1269 | 5096 | 3460 | FALSE | 2 |
| S | 1270 | 5567 | 3972 | FALSE | 3 |
| S | 1271 | 7159 | 4061 | FALSE | 3 |
| S | 1272 | 9142 | 4061 | FALSE | 3 |
| S | 1273 | 8329 | 0 | FALSE | 2 |
| E | 1 | 2447 | 6688 | FALSE | 2 |
| E | 2 | 2447 | 6688 | FALSE | 2 |
| E | 3 | 2447 | 4598 | FALSE | 1 |
| E | 4 | 2309 | 8945 | FALSE | 1 |
| E | 5 | 2126 | 8945 | FALSE | 2 |
| E | 6 | 5923 | 8945 | FALSE | 5 |
| E | 7 | 7607 | 7218 | FALSE | 2 |
| E | 8 | 7910 | 5721 | FALSE | 2 |
| E | 9 | 7910 | 5721 | FALSE | 3 |
| E | 10 | 7910 | 5721 | FALSE | 3 |
| E | 11 | 8977 | 5721 | FALSE | 2 |
| E | 12 | 8977 | 5721 | FALSE | 3 |
| E | 13 | 9908 | 5721 | FALSE | 3 |
| E | 14 | 9908 | 5721 | FALSE | 3 |
| E | 15 | 11387 | 5624 | FALSE | 1 |
| E | 16 | 11387 | 5624 | FALSE | 3 |
| E | 17 | 11387 | 5624 | FALSE | 3 |
| E | 18 | 11387 | 6142 | FALSE | 3 |
| E | 19 | 12391 | 3879 | FALSE | 4 |
| E | 20 | 15140 | 3879 | FALSE | 0 |
| E | 21 | 11032 | 3879 | FALSE | 2 |
| E | 22 | 13057 | 1730 | FALSE | 4 |
| M | 1 | 2892 | 1973 | FALSE | 3 |
| M | 2 | 2784 | 2035 | FALSE | 4 |
| M | 3 | 2784 | 2035 | FALSE | 7 |
| M | 4 | 2686 | 2035 | FALSE | 2 |
| M | 5 | 2686 | 2035 | FALSE | 1 |
| M | 6 | 2686 | 2795 | FALSE | 4 |
| M | 7 | 2564 | 2795 | FALSE | 3 |
| M | 8 | 2518 | 2795 | FALSE | 6 |
| M | 9 | 2518 | 2795 | FALSE | 3 |
| M | 10 | 3493 | 2795 | FALSE | 5 |

| | | | | | |
|---|---|---|---|---|---|
| M | 11 | 7062 | 4227 | FALSE | 3 |
| M | 12 | 7062 | 4227 | FALSE | 2 |
| M | 13 | 9188 | 5228 | FALSE | 3 |
| M | 14 | 9188 | 5228 | FALSE | 2 |
| M | 15 | 9188 | 5556 | FALSE | 5 |
| M | 16 | 9188 | 5556 | FALSE | 3 |
| M | 17 | 10226 | 6358 | FALSE | 3 |
| M | 18 | 10273 | 5612 | FALSE | 2 |
| M | 19 | 11482 | 5612 | FALSE | 4 |
| M | 20 | 11482 | 5612 | FALSE | 6 |
| M | 21 | 10333 | 5177 | FALSE | 3 |
| M | 22 | 11617 | 5177 | FALSE | 2 |
| M | 23 | 13219 | 5177 | FALSE | 5 |
| M | 24 | 13219 | 5177 | FALSE | 3 |
| M | 25 | 12824 | 5177 | FALSE | 3 |
| M | 26 | 6443 | 2628 | FALSE | 0 |
| M | 27 | 6443 | 2628 | FALSE | 2 |
| M | 28 | 6443 | 2628 | FALSE | 2 |
| M | 29 | 6443 | 2628 | FALSE | 3 |
| M | 30 | 6443 | 2628 | FALSE | 4 |
| M | 31 | 6443 | 2628 | FALSE | 2 |
| M | 32 | 6443 | 2628 | FALSE | 3 |
| M | 33 | 2286 | 0 | FALSE | 3 |
| M | 34 | 2286 | 0 | FALSE | 2 |
| M | 35 | 2286 | 0 | FALSE | 2 |
| M | 41 | 6356 | 0 | FALSE | 2 |
| M | 42 | 6356 | 0 | FALSE | 3 |
| M | 43 | 4280 | 0 | FALSE | 2 |
| M | 44 | 4280 | 0 | FALSE | 3 |
| M | 45 | 4280 | 0 | FALSE | 1 |
| M | 46 | 4280 | 0 | FALSE | 3 |
| M | 47 | 4280 | 0 | FALSE | 1 |
| M | 48 | 4280 | 0 | FALSE | 0 |
| M | 49 | 4280 | 0 | FALSE | 2 |
| M | 50 | 4280 | 0 | FALSE | 1 |
| M | 51 | 4280 | 0 | FALSE | 1 |
| M | 52 | 4280 | 0 | FALSE | 1 |
| M | 53 | 4280 | 0 | FALSE | 1 |
| M | 54 | 4280 | 0 | FALSE | 2 |
| M | 55 | 4280 | 0 | FALSE | 1 |
| M | 56 | 2205 | 0 | FALSE | 2 |
| M | 57 | 2205 | 0 | FALSE | 4 |
| M | 66 | 0 | 4345 | FALSE | 6 |
| M | 67 | 0 | 3349 | FALSE | 2 |
| M | 68 | 0 | 2938 | FALSE | 4 |
| M | 69 | 0 | 2938 | FALSE | 4 |
| M | 70 | 0 | 2938 | FALSE | 4 |
| M | 71 | 0 | 2938 | FALSE | 4 |
| M | 72 | 0 | 2938 | FALSE | 3 |
| M | 73 | 0 | 2938 | FALSE | 2 |
| M | 74 | 4151 | 2938 | FALSE | 1 |
| M | 75 | 4151 | 2938 | FALSE | 2 |
| M | 76 | 4151 | 2938 | FALSE | 6 |
| M | 77 | 4151 | 2938 | FALSE | 3 |
| M | 78 | 4151 | 2938 | FALSE | 4 |
| M | 79 | 4151 | 2981 | FALSE | 3 |
| M | 80 | 4151 | 2981 | FALSE | 1 |
| M | 81 | 4151 | 2526 | FALSE | 5 |
| M | 82 | 4151 | 2613 | FALSE | 2 |
| M | 83 | 4151 | 3109 | FALSE | 4 |
| M | 84 | 4151 | 3109 | FALSE | 3 |
| M | 85 | 4151 | 3109 | FALSE | 3 |

| | | | | | |
|---|---|---|---|---|---|
| M | 86 | 4151 | 3109 | FALSE | 1 |
| M | 87 | 4151 | 3109 | FALSE | 1 |
| M | 88 | 4151 | 3109 | FALSE | 4 |
| M | 89 | 0 | 2296 | FALSE | 4 |
| M | 90 | 0 | 2296 | FALSE | 2 |
| M | 91 | 0 | 2296 | FALSE | 3 |
| M | 92 | 0 | 2296 | FALSE | 0 |
| M | 93 | 0 | 2296 | FALSE | 2 |
| M | 94 | 0 | 1483 | FALSE | 4 |
| M | 95 | 0 | 1483 | FALSE | 3 |
| M | 96 | 0 | 1483 | FALSE | 1 |
| M | 97 | 0 | 1483 | FALSE | 1 |
| M | 98 | 0 | 1483 | FALSE | 5 |
| M | 99 | 0 | 1483 | FALSE | 2 |
| M | 100 | 0 | 1483 | FALSE | 4 |
| M | 101 | 0 | 1483 | FALSE | 0 |
| M | 102 | 0 | 1483 | FALSE | 3 |
| M | 103 | 0 | 1483 | FALSE | 0 |
| M | 105 | 1138 | 0 | TRUE | 3 |
| M | 106 | 1138 | 0 | TRUE | 3 |
| M | 107 | 1138 | 0 | FALSE | 4 |
| M | 108 | 1640 | 0 | TRUE | 2 |
| M | 109 | 1640 | 0 | TRUE | 3 |
| M | 110 | 1640 | 0 | TRUE | 3 |
| M | 111 | 1640 | 0 | TRUE | 2 |
| M | 112 | 1640 | 0 | TRUE | 3 |
| M | 113 | 1640 | 0 | TRUE | 3 |
| M | 114 | 1590 | 0 | TRUE | 6 |
| M | 115 | 3075 | 2346 | TRUE | 4 |
| M | 116 | 3075 | 2346 | TRUE | 2 |
| M | 117 | 2884 | 2346 | TRUE | 4 |
| M | 118 | 2884 | 2346 | FALSE | 1 |
| M | 119 | 2884 | 2346 | TRUE | 2 |
| M | 120 | 3466 | 2346 | TRUE | 2 |
| M | 121 | 3191 | 2346 | FALSE | 2 |
| M | 122 | 3191 | 2346 | FALSE | 4 |
| M | 123 | 3576 | 1775 | FALSE | 2 |
| M | 124 | 3576 | 1775 | FALSE | 2 |
| M | 125 | 3576 | 1775 | FALSE | 3 |
| M | 126 | 3576 | 1775 | FALSE | 4 |
| M | 127 | 3576 | 1775 | FALSE | 2 |
| M | 128 | 3576 | 1775 | FALSE | 3 |
| M | 129 | 4165 | 1775 | FALSE | 3 |
| M | 130 | 3117 | 1203 | FALSE | 5 |
| M | 131 | 3117 | 1203 | FALSE | 2 |
| M | 132 | 5867 | 1143 | FALSE | 6 |
| M | 133 | 5417 | 1143 | FALSE | 3 |
| M | 134 | 8261 | 1112 | FALSE | 1 |
| M | 135 | 8435 | 5046 | FALSE | 2 |
| M | 136 | 7981 | 5046 | FALSE | 4 |
| M | 137 | 7981 | 4444 | FALSE | 1 |
| M | 138 | 8936 | 5092 | FALSE | 4 |
| M | 139 | 8936 | 5092 | FALSE | 2 |
| M | 140 | 8333 | 5092 | FALSE | 1 |
| M | 141 | 7908 | 5092 | FALSE | 3 |
| M | 142 | 7908 | 5092 | FALSE | 4 |
| M | 143 | 7908 | 5092 | FALSE | 3 |
| M | 144 | 7908 | 5092 | FALSE | 4 |
| M | 145 | 7908 | 5092 | FALSE | 2 |
| M | 146 | 7908 | 5092 | FALSE | 3 |
| M | 147 | 6689 | 4494 | FALSE | 3 |
| M | 148 | 7132 | 4494 | FALSE | 2 |

| | | | | | |
|---|---|---|---|---|---|
| M | 149 | 4459 | 5183 | FALSE | 1 |
| M | 150 | 1520 | 1340 | FALSE | 2 |
| M | 151 | 1485 | 1340 | FALSE | 0 |
| M | 152 | 1485 | 1294 | FALSE | 4 |
| M | 153 | 1485 | 1294 | FALSE | 3 |
| M | 154 | 1485 | 1294 | FALSE | 4 |
| M | 155 | 1685 | 1294 | FALSE | 2 |
| M | 156 | 1728 | 1294 | FALSE | 2 |
| M | 157 | 1638 | 1418 | FALSE | 1 |
| M | 158 | 2990 | 1418 | FALSE | 4 |
| M | 159 | 2985 | 1445 | FALSE | 1 |
| M | 160 | 2864 | 1445 | FALSE | 4 |
| M | 161 | 2875 | 1916 | FALSE | 2 |
| M | 162 | 3213 | 2331 | FALSE | 4 |
| M | 163 | 3213 | 2331 | FALSE | 4 |
| M | 164 | 3213 | 2331 | FALSE | 6 |
| M | 165 | 3213 | 2331 | FALSE | 4 |
| M | 166 | 5541 | 2228 | FALSE | 4 |
| M | 167 | 6119 | 2569 | FALSE | 4 |
| M | 168 | 6119 | 2569 | FALSE | 4 |
| M | 169 | 6119 | 2569 | FALSE | 2 |
| M | 170 | 6443 | 2569 | FALSE | 5 |
| M | 171 | 6767 | 2569 | FALSE | 3 |
| M | 172 | 7266 | 2795 | FALSE | 3 |
| M | 173 | 8744 | 2795 | FALSE | 5 |
| M | 174 | 9711 | 3218 | FALSE | 3 |
| M | 175 | 11321 | 3218 | FALSE | 4 |
| M | 176 | 13403 | 2925 | FALSE | 3 |
| M | 177 | 23553 | 2925 | FALSE | 3 |
| M | 178 | 23553 | 2925 | FALSE | 3 |
| M | 179 | 23553 | 2925 | FALSE | 1 |
| M | 180 | 23553 | 2925 | FALSE | 2 |
| M | 181 | 13626 | 3933 | FALSE | 1 |
| M | 202 | 0 | 4987 | FALSE | 4 |
| M | 203 | 2111 | 4259 | FALSE | 3 |
| M | 204 | 2111 | 4259 | FALSE | 2 |
| M | 205 | 1683 | 4259 | FALSE | 1 |
| M | 206 | 1683 | 4259 | FALSE | 3 |
| M | 207 | 1731 | 4259 | FALSE | 2 |
| M | 208 | 6730 | 4259 | FALSE | 3 |
| M | 209 | 5350 | 4259 | FALSE | 3 |
| M | 210 | 5350 | 4259 | FALSE | 3 |
| M | 211 | 5350 | 4259 | FALSE | 3 |
| M | 212 | 5350 | 4259 | FALSE | 2 |
| M | 213 | 5350 | 4259 | FALSE | 3 |
| M | 214 | 5350 | 4259 | FALSE | 1 |
| M | 215 | 5350 | 4259 | FALSE | 4 |
| M | 216 | 5350 | 4259 | FALSE | 2 |
| M | 217 | 5350 | 3531 | FALSE | 1 |
| M | 218 | 5890 | 0 | FALSE | 5 |
| M | 219 | 5890 | 0 | FALSE | 2 |
| M | 220 | 6817 | 0 | FALSE | 1 |
| M | 221 | 6817 | 0 | FALSE | 0 |
| M | 222 | 8065 | 0 | FALSE | 4 |
| N | 1 | 1964 | 3271 | FALSE | 3 |
| N | 2 | 2509 | 3271 | FALSE | 2 |
| N | 3 | 2509 | 3271 | FALSE | 3 |
| N | 4 | 2509 | 3271 | FALSE | 1 |
| N | 5 | 2509 | 3271 | FALSE | 5 |
| N | 6 | 2509 | 3271 | FALSE | 3 |
| N | 7 | 2509 | 3271 | FALSE | 4 |
| N | 8 | 2263 | 3271 | FALSE | 3 |

| | | | | | |
|---|---|---|---|---|---|
| N | 9 | 2263 | 3271 | FALSE | 5 |
| N | 10 | 2263 | 3271 | FALSE | 5 |
| N | 11 | 2203 | 4018 | FALSE | 1 |
| N | 12 | 2203 | 4018 | FALSE | 4 |
| N | 13 | 2044 | 6999 | FALSE | 4 |
| N | 14 | 2044 | 6999 | FALSE | 3 |
| N | 15 | 2525 | 0 | FALSE | 3 |
| N | 16 | 2525 | 0 | FALSE | 4 |
| N | 17 | 1621 | 0 | FALSE | 5 |
| N | 18 | 1619 | 1678 | FALSE | 7 |
| N | 19 | 1619 | 1678 | FALSE | 4 |
| N | 20 | 1807 | 1678 | FALSE | 3 |
| N | 21 | 1807 | 1678 | FALSE | 3 |
| N | 22 | 1807 | 1612 | FALSE | 3 |
| N | 23 | 1949 | 1612 | FALSE | 3 |
| N | 24 | 1949 | 1612 | FALSE | 6 |
| N | 25 | 2290 | 1612 | FALSE | 6 |
| N | 26 | 2370 | 1612 | FALSE | 2 |
| N | 27 | 2370 | 1612 | FALSE | 3 |
| N | 28 | 2370 | 1612 | FALSE | 3 |
| N | 29 | 2370 | 1612 | FALSE | 4 |
| N | 30 | 2370 | 1612 | FALSE | 9 |
| N | 31 | 3185 | 2523 | FALSE | 4 |
| N | 32 | 3185 | 2523 | FALSE | 4 |
| N | 33 | 3381 | 3791 | FALSE | 4 |
| N | 34 | 3381 | 3791 | FALSE | 2 |
| N | 35 | 3780 | 3791 | FALSE | 6 |
| N | 36 | 3780 | 3791 | FALSE | 4 |
| N | 37 | 3780 | 6165 | FALSE | 4 |
| N | 38 | 3780 | 6165 | FALSE | 1 |
| N | 39 | 3780 | 6165 | FALSE | 4 |
| N | 40 | 4815 | 6165 | FALSE | 4 |
| N | 41 | 4815 | 6165 | FALSE | 5 |
| N | 42 | 4815 | 6165 | FALSE | 3 |
| N | 43 | 4815 | 3903 | FALSE | 3 |
| N | 44 | 3941 | 4732 | FALSE | 5 |
| N | 45 | 3941 | 4732 | FALSE | 3 |
| N | 46 | 1319 | 4015 | FALSE | 6 |
| N | 47 | 1319 | 4015 | FALSE | 0 |
| N | 48 | 1319 | 4015 | FALSE | 3 |
| N | 49 | 1558 | 4015 | FALSE | 1 |
| N | 50 | 1765 | 4015 | FALSE | 1 |
| N | 51 | 1765 | 4015 | FALSE | 2 |
| N | 52 | 1765 | 4015 | FALSE | 6 |
| N | 53 | 1780 | 4015 | FALSE | 4 |
| N | 54 | 1816 | 4015 | FALSE | 4 |
| N | 55 | 1816 | 4015 | FALSE | 2 |
| N | 56 | 1816 | 4015 | FALSE | 4 |
| N | 57 | 1816 | 4015 | FALSE | 3 |
| N | 58 | 1704 | 4591 | FALSE | 4 |
| N | 59 | 1781 | 1980 | FALSE | 4 |
| N | 60 | 1781 | 3220 | FALSE | 4 |
| N | 61 | 1667 | 3220 | FALSE | 4 |
| N | 62 | 5241 | 3679 | FALSE | 3 |
| N | 63 | 5241 | 3679 | FALSE | 5 |
| N | 64 | 5733 | 3679 | FALSE | 3 |
| N | 65 | 6325 | 3313 | FALSE | 3 |
| N | 66 | 6325 | 3313 | FALSE | 3 |
| N | 67 | 6325 | 3313 | FALSE | 3 |
| N | 68 | 7225 | 3313 | FALSE | 5 |
| N | 69 | 8542 | 3313 | FALSE | 4 |
| N | 70 | 8542 | 3313 | FALSE | 3 |

| | | | | | |
|---|---|---|---|---|---|
| N | 71 | 7151 | 3656 | FALSE | 3 |
| N | 72 | 7151 | 3656 | FALSE | 4 |
| N | 73 | 6732 | 3438 | FALSE | 4 |
| N | 74 | 6732 | 3816 | FALSE | 3 |
| N | 75 | 6732 | 3439 | FALSE | 3 |
| N | 76 | 7859 | 3439 | FALSE | 3 |
| N | 77 | 2256 | 2820 | FALSE | 2 |
| N | 78 | 2256 | 2368 | FALSE | 4 |
| N | 79 | 2012 | 2290 | FALSE | 4 |
| N | 80 | 2012 | 2217 | FALSE | 3 |
| N | 81 | 1805 | 2049 | FALSE | 4 |
| N | 82 | 1805 | 2049 | FALSE | 3 |
| N | 83 | 1805 | 2049 | FALSE | 2 |
| N | 84 | 1805 | 2049 | FALSE | 1 |
| N | 85 | 1805 | 2049 | FALSE | 4 |
| N | 86 | 1842 | 1439 | FALSE | 2 |
| N | 87 | 1842 | 1437 | FALSE | 2 |
| N | 88 | 1317 | 1466 | FALSE | 2 |
| N | 89 | 1317 | 1466 | FALSE | 4 |
| N | 90 | 1317 | 1466 | FALSE | 4 |
| N | 91 | 1317 | 1466 | FALSE | 3 |
| N | 92 | 1204 | 1466 | FALSE | 3 |
| N | 93 | 1204 | 1354 | FALSE | 5 |
| N | 94 | 1290 | 1173 | FALSE | 1 |
| N | 95 | 1290 | 1235 | FALSE | 5 |
| N | 96 | 0 | 1426 | FALSE | 3 |
| N | 97 | 0 | 1426 | FALSE | 2 |
| N | 98 | 0 | 1368 | TRUE | 2 |
| N | 99 | 0 | 1282 | TRUE | 3 |
| N | 100 | 0 | 1282 | TRUE | 2 |
| N | 101 | 0 | 1282 | TRUE | 4 |
| N | 102 | 0 | 1209 | FALSE | 3 |
| N | 103 | 0 | 1209 | TRUE | 5 |
| N | 104 | 0 | 1209 | TRUE | 3 |
| N | 105 | 0 | 1209 | FALSE | 3 |
| N | 106 | 0 | 1209 | TRUE | 3 |
| N | 107 | 0 | 1209 | TRUE | 2 |
| N | 108 | 0 | 1209 | TRUE | 2 |
| N | 109 | 0 | 1209 | TRUE | 1 |
| N | 110 | 0 | 1209 | TRUE | 3 |
| N | 111 | 0 | 1209 | TRUE | 2 |
| N | 112 | 0 | 1209 | TRUE | 1 |
| N | 113 | 0 | 1108 | TRUE | 3 |
| N | 114 | 0 | 2753 | TRUE | 4 |
| N | 115 | 1788 | 2300 | TRUE | 4 |
| N | 116 | 1908 | 2936 | TRUE | 4 |
| N | 117 | 6273 | 2861 | TRUE | 3 |
| N | 118 | 8850 | 4345 | FALSE | 1 |
| N | 119 | 8850 | 4345 | FALSE | 3 |
| N | 120 | 8302 | 4949 | FALSE | 3 |
| N | 121 | 8302 | 4949 | FALSE | 4 |
| N | 122 | 8302 | 4949 | FALSE | 4 |
| N | 123 | 8302 | 4513 | FALSE | 2 |
| N | 124 | 8302 | 4513 | FALSE | 4 |
| N | 125 | 8302 | 4513 | FALSE | 4 |
| N | 126 | 8192 | 4513 | FALSE | 4 |
| N | 127 | 8192 | 4513 | FALSE | 3 |
| N | 128 | 8192 | 4513 | FALSE | 5 |
| N | 129 | 8192 | 4702 | FALSE | 6 |
| N | 130 | 10754 | 5175 | FALSE | 1 |
| N | 131 | 16571 | 5230 | FALSE | 3 |
| N | 132 | 11646 | 5764 | FALSE | 4 |

| | | | | | |
|---|---|---|---|---|---|
| N | 133 | 3226 | 4236 | FALSE | 3 |
| N | 134 | 3226 | 4236 | FALSE | 4 |
| N | 135 | 2303 | 1766 | FALSE | 4 |
| N | 136 | 3311 | 5800 | FALSE | 4 |
| N | 137 | 3311 | 5800 | FALSE | 5 |
| N | 138 | 3311 | 6994 | FALSE | 4 |
| N | 139 | 3311 | 6994 | FALSE | 4 |
| N | 140 | 3311 | 6994 | FALSE | 4 |
| N | 141 | 2817 | 6994 | FALSE | 2 |
| N | 142 | 2817 | 6994 | FALSE | 6 |
| N | 143 | 2817 | 6994 | FALSE | 2 |
| N | 144 | 2668 | 8490 | FALSE | 5 |
| N | 145 | 2668 | 8490 | FALSE | 2 |
| N | 146 | 2668 | 8490 | FALSE | 3 |
| N | 147 | 2668 | 8490 | FALSE | 4 |
| N | 148 | 2881 | 8490 | FALSE | 5 |
| N | 149 | 2881 | 8490 | FALSE | 4 |
| N | 150 | 4150 | 8490 | FALSE | 1 |
| N | 151 | 1626 | 0 | FALSE | 5 |
| N | 152 | 1626 | 0 | FALSE | 6 |
| N | 153 | 1626 | 0 | FALSE | 2 |
| N | 154 | 1626 | 0 | FALSE | 4 |
| N | 155 | 2528 | 0 | FALSE | 4 |
| N | 156 | 2233 | 0 | FALSE | 4 |
| N | 157 | 2025 | 0 | FALSE | 3 |
| N | 158 | 2025 | 0 | FALSE | 4 |
| N | 159 | 2059 | 0 | FALSE | 2 |
| N | 160 | 2038 | 0 | FALSE | 3 |
| N | 161 | 1983 | 0 | FALSE | 3 |
| N | 162 | 1983 | 0 | FALSE | 3 |
| N | 163 | 2024 | 0 | FALSE | 4 |
| N | 164 | 2024 | 0 | FALSE | 5 |
| N | 165 | 2024 | 0 | TRUE | 2 |
| N | 166 | 2024 | 0 | FALSE | 2 |
| N | 167 | 2024 | 0 | TRUE | 7 |
| N | 168 | 2024 | 0 | TRUE | 4 |
| N | 169 | 2024 | 0 | TRUE | 3 |
| N | 170 | 1751 | 0 | FALSE | 4 |
| N | 171 | 1818 | 0 | FALSE | 1 |
| N | 172 | 1922 | 0 | FALSE | 4 |
| N | 173 | 1922 | 0 | FALSE | 3 |
| N | 174 | 1822 | 12364 | TRUE | 4 |
| N | 175 | 1792 | 10319 | FALSE | 4 |
| N | 176 | 1890 | 8851 | FALSE | 1 |
| N | 177 | 12294 | 8851 | FALSE | 2 |
| N | 178 | 20105 | 8851 | FALSE | 4 |
| N | 179 | 19767 | 8851 | FALSE | 4 |
| N | 180 | 19767 | 8851 | FALSE | 5 |
| N | 181 | 19767 | 8851 | FALSE | 5 |
| N | 182 | 18513 | 8851 | FALSE | 4 |
| N | 183 | 18513 | 8851 | FALSE | 2 |
| N | 184 | 19017 | 7773 | FALSE | 3 |
| N | 185 | 18039 | 7773 | FALSE | 4 |
| N | 186 | 16566 | 8194 | FALSE | 4 |
| N | 187 | 15464 | 8544 | FALSE | 2 |
| N | 188 | 15464 | 8013 | FALSE | 2 |
| N | 189 | 15506 | 5527 | FALSE | 3 |
| N | 190 | 15506 | 6092 | FALSE | 5 |
| N | 191 | 14394 | 7008 | FALSE | 5 |
| N | 192 | 11961 | 7008 | FALSE | 2 |
| N | 193 | 9963 | 6786 | FALSE | 5 |
| N | 194 | 7935 | 6786 | FALSE | 4 |

| | | | | | |
|---|---|---|---|---|---|
| N | 195 | 7257 | 6786 | FALSE | 6 |
| N | 196 | 7257 | 6786 | FALSE | 4 |
| N | 197 | 6843 | 6786 | FALSE | 4 |
| N | 198 | 6843 | 6786 | FALSE | 3 |
| N | 199 | 2657 | 7881 | FALSE | 6 |
| N | 200 | 1665 | 7881 | FALSE | 2 |
| N | 201 | 1642 | 6677 | FALSE | 2 |
| N | 202 | 1358 | 4500 | FALSE | 4 |
| N | 203 | 1358 | 5394 | FALSE | 5 |
| N | 204 | 1243 | 4047 | FALSE | 5 |
| N | 205 | 1243 | 4047 | FALSE | 2 |
| N | 206 | 1292 | 4047 | FALSE | 2 |
| N | 207 | 1234 | 4047 | FALSE | 5 |
| N | 208 | 1234 | 3234 | FALSE | 6 |
| N | 209 | 1234 | 3234 | FALSE | 4 |
| N | 210 | 1273 | 3234 | FALSE | 6 |
| N | 211 | 1273 | 3234 | FALSE | 5 |
| N | 212 | 1273 | 3234 | FALSE | 6 |
| N | 213 | 1273 | 3234 | FALSE | 2 |
| N | 214 | 1311 | 3234 | FALSE | 5 |
| N | 215 | 0 | 3234 | FALSE | 4 |
| N | 216 | 0 | 1354 | FALSE | 4 |
| N | 217 | 0 | 1354 | FALSE | 6 |
| N | 218 | 0 | 1354 | FALSE | 2 |
| N | 228 | 3439 | 0 | FALSE | 6 |
| N | 229 | 3439 | 0 | FALSE | 3 |
| N | 230 | 3439 | 0 | FALSE | 3 |
| N | 231 | 3439 | 6342 | FALSE | 1 |
| N | 232 | 3439 | 6342 | FALSE | 4 |
| N | 233 | 3439 | 6342 | FALSE | 3 |
| N | 234 | 3439 | 6342 | FALSE | 3 |
| N | 235 | 3439 | 6342 | FALSE | 5 |
| N | 236 | 3439 | 6342 | FALSE | 5 |
| N | 237 | 3439 | 6342 | FALSE | 3 |
| N | 238 | 3439 | 6342 | FALSE | 5 |
| N | 239 | 3439 | 6342 | FALSE | 4 |
| N | 240 | 3439 | 6342 | FALSE | 4 |
| N | 241 | 3439 | 6342 | FALSE | 3 |
| N | 242 | 3722 | 6342 | FALSE | 4 |
| N | 243 | 3154 | 6342 | FALSE | 4 |
| N | 244 | 2924 | 6342 | FALSE | 4 |
| N | 245 | 2523 | 6342 | FALSE | 3 |
| N | 246 | 2523 | 0 | FALSE | 5 |
| N | 247 | 2523 | 0 | FALSE | 4 |
| N | 248 | 2395 | 0 | FALSE | 5 |
| N | 249 | 2395 | 0 | FALSE | 1 |
| N | 250 | 2395 | 0 | FALSE | 2 |
| N | 251 | 2395 | 0 | FALSE | 4 |
| N | 252 | 2395 | 0 | FALSE | 5 |
| N | 253 | 3495 | 14565 | FALSE | 8 |
| N | 254 | 3495 | 14565 | FALSE | 8 |
| N | 255 | 3495 | 8143 | FALSE | 7 |
| N | 256 | 3218 | 8143 | FALSE | 4 |
| N | 257 | 3105 | 8143 | FALSE | 1 |
| N | 258 | 3239 | 8143 | FALSE | 3 |
| N | 259 | 3394 | 8143 | FALSE | 2 |
| N | 260 | 3912 | 8143 | FALSE | 2 |
| N | 261 | 3912 | 8143 | FALSE | 1 |
| N | 262 | 3912 | 8143 | FALSE | 1 |
| N | 263 | 5684 | 8143 | FALSE | 2 |
| N | 264 | 5684 | 8143 | FALSE | 4 |
| N | 265 | 5684 | 8143 | FALSE | 3 |

| | | | | | |
|---|---|---|---|---|---|
| N | 266 | 5684 | 8143 | FALSE | 2 |
| N | 267 | 5684 | 8143 | FALSE | 3 |
| N | 268 | 1274 | 1720 | FALSE | 4 |
| N | 269 | 1274 | 1720 | FALSE | 2 |
| N | 270 | 1274 | 0 | FALSE | 3 |
| N | 272 | 3277 | 0 | FALSE | 4 |
| N | 273 | 3277 | 0 | FALSE | 3 |
| N | 274 | 3277 | 0 | FALSE | 3 |
| N | 275 | 2412 | 1161 | FALSE | 3 |
| N | 276 | 2412 | 1372 | FALSE | 2 |
| N | 277 | 2412 | 1592 | FALSE | 2 |
| N | 278 | 1993 | 1556 | FALSE | 3 |
| N | 279 | 4390 | 1549 | FALSE | 2 |
| N | 280 | 7457 | 2282 | FALSE | 2 |
| N | 281 | 7457 | 2282 | FALSE | 0 |
| N | 282 | 7457 | 2282 | FALSE | 3 |
| N | 283 | 7457 | 2191 | FALSE | 5 |
| N | 284 | 7082 | 2212 | FALSE | 4 |
| N | 285 | 7082 | 2108 | FALSE | 5 |
| N | 286 | 7082 | 2108 | FALSE | 2 |
| N | 287 | 7399 | 2108 | FALSE | 2 |
| N | 288 | 7399 | 2108 | FALSE | 2 |
| N | 289 | 8475 | 2108 | FALSE | 5 |
| N | 290 | 8400 | 2362 | FALSE | 6 |
| N | 291 | 8400 | 2440 | FALSE | 1 |
| N | 292 | 8400 | 2486 | FALSE | 2 |
| N | 293 | 10605 | 2615 | FALSE | 1 |
| N | 294 | 10411 | 2975 | FALSE | 4 |
| N | 295 | 8291 | 4140 | FALSE | 6 |
| N | 296 | 10143 | 4140 | FALSE | 3 |
| N | 297 | 10143 | 4140 | FALSE | 5 |
| N | 298 | 10143 | 4808 | FALSE | 2 |
| N | 299 | 11403 | 5612 | FALSE | 3 |
| N | 300 | 11403 | 7881 | FALSE | 3 |
| N | 301 | 11403 | 7881 | FALSE | 2 |
| N | 302 | 11403 | 7881 | FALSE | 6 |
| N | 303 | 11403 | 7881 | FALSE | 3 |
| N | 304 | 9406 | 7881 | FALSE | 2 |
| N | 305 | 21257 | 11808 | FALSE | 4 |
| N | 306 | 21257 | 11808 | FALSE | 1 |
| N | 307 | 21257 | 11808 | FALSE | 3 |
| N | 308 | 21257 | 11808 | FALSE | 4 |
| N | 309 | 21257 | 11808 | FALSE | 4 |
| N | 310 | 21257 | 0 | FALSE | 4 |
| N | 316 | 1621 | 6027 | FALSE | 5 |
| N | 317 | 1621 | 6027 | FALSE | 2 |
| N | 318 | 1621 | 6027 | FALSE | 5 |
| N | 319 | 1621 | 6027 | FALSE | 4 |
| N | 320 | 1621 | 6027 | FALSE | 0 |
| N | 321 | 1397 | 5409 | FALSE | 4 |
| N | 322 | 1397 | 5409 | FALSE | 3 |
| N | 323 | 6013 | 5043 | FALSE | 5 |
| N | 324 | 6013 | 5043 | FALSE | 1 |
| N | 325 | 6013 | 5043 | FALSE | 1 |
| N | 326 | 6013 | 5043 | FALSE | 5 |
| N | 327 | 6013 | 5043 | FALSE | 7 |
| N | 328 | 6013 | 5043 | FALSE | 3 |
| N | 329 | 6013 | 5043 | FALSE | 2 |
| N | 330 | 6013 | 5043 | FALSE | 3 |
| N | 331 | 8208 | 4550 | FALSE | 3 |
| N | 332 | 5978 | 4550 | FALSE | 3 |
| N | 333 | 5978 | 4550 | FALSE | 2 |

| | | | | | |
|---|---|---|---|---|---|
| N | 334 | 5978 | 4550 | FALSE | 3 |
| N | 335 | 5978 | 3637 | FALSE | 1 |
| N | 336 | 8381 | 3061 | FALSE | 5 |
| N | 337 | 8381 | 3061 | FALSE | 2 |
| N | 338 | 1720 | 1811 | FALSE | 1 |
| N | 339 | 1585 | 1811 | FALSE | 1 |
| N | 340 | 1585 | 1811 | FALSE | 3 |
| N | 341 | 1585 | 1811 | FALSE | 4 |
| N | 342 | 1585 | 2247 | FALSE | 2 |
| N | 343 | 1585 | 2247 | FALSE | 2 |
| N | 344 | 1585 | 2247 | FALSE | 3 |
| N | 345 | 1421 | 2247 | FALSE | 2 |
| N | 346 | 1421 | 2247 | FALSE | 3 |
| N | 347 | 1405 | 2247 | FALSE | 2 |
| N | 348 | 1405 | 2247 | FALSE | 4 |
| N | 349 | 1405 | 2247 | FALSE | 4 |
| N | 350 | 1405 | 2684 | FALSE | 3 |
| N | 351 | 1405 | 2684 | FALSE | 2 |
| N | 352 | 1405 | 2684 | FALSE | 5 |
| N | 353 | 1023 | 2684 | FALSE | 3 |
| N | 354 | 1920 | 2684 | FALSE | 5 |
| N | 355 | 1920 | 2684 | FALSE | 3 |
| N | 356 | 1920 | 2684 | FALSE | 2 |
| N | 357 | 1920 | 0 | FALSE | 3 |
| N | 358 | 1753 | 0 | FALSE | 3 |
| N | 359 | 1753 | 0 | FALSE | 4 |
| N | 360 | 3462 | 0 | FALSE | 3 |
| N | 361 | 2930 | 0 | FALSE | 6 |
| N | 362 | 3424 | 0 | FALSE | 3 |
| N | 363 | 3424 | 0 | FALSE | 2 |
| N | 364 | 3424 | 1159 | FALSE | 4 |
| N | 365 | 3734 | 1159 | FALSE | 4 |
| N | 366 | 3464 | 1159 | FALSE | 2 |
| N | 367 | 3234 | 1881 | FALSE | 4 |
| N | 368 | 3234 | 1881 | FALSE | 4 |
| N | 369 | 3336 | 1881 | FALSE | 3 |
| N | 370 | 3336 | 1881 | FALSE | 1 |
| N | 371 | 3336 | 1881 | FALSE | 4 |
| N | 372 | 3336 | 1881 | FALSE | 0 |
| N | 373 | 3657 | 1881 | FALSE | 5 |
| N | 374 | 3657 | 1881 | FALSE | 3 |
| N | 375 | 3082 | 1881 | FALSE | 2 |
| N | 376 | 3179 | 1881 | FALSE | 5 |
| N | 377 | 7651 | 1668 | FALSE | 1 |
| N | 378 | 8031 | 1878 | FALSE | 4 |
| N | 379 | 8031 | 2118 | FALSE | 4 |
| N | 380 | 8386 | 2118 | FALSE | 3 |
| N | 381 | 9801 | 2118 | FALSE | 4 |
| N | 382 | 11961 | 1876 | FALSE | 1 |
| N | 383 | 11961 | 1876 | FALSE | 5 |
| N | 384 | 11961 | 1876 | FALSE | 3 |
| N | 385 | 11961 | 1876 | FALSE | 3 |
| N | 386 | 11961 | 1876 | FALSE | 4 |
| N | 387 | 11961 | 1876 | FALSE | 4 |
| N | 388 | 11961 | 1773 | FALSE | 2 |
| N | 389 | 11961 | 1831 | FALSE | 5 |
| N | 390 | 11961 | 1731 | FALSE | 5 |
| N | 391 | 12284 | 1708 | FALSE | 4 |
| N | 392 | 6794 | 1679 | FALSE | 3 |
| N | 393 | 1883 | 1513 | FALSE | 5 |
| N | 394 | 1883 | 1513 | FALSE | 3 |
| N | 395 | 1883 | 1520 | FALSE | 2 |

| | | | | | |
|---|---|---|---|---|---|
| N | 396 | 1716 | 1604 | FALSE | 4 |
| N | 397 | 1716 | 1584 | FALSE | 4 |
| N | 398 | 1716 | 1584 | FALSE | 4 |
| N | 399 | 1716 | 1584 | FALSE | 5 |
| N | 400 | 1716 | 1584 | FALSE | 3 |
| N | 401 | 1716 | 1584 | FALSE | 2 |
| N | 402 | 1716 | 1584 | FALSE | 3 |
| N | 403 | 1716 | 1587 | FALSE | 2 |
| N | 404 | 1716 | 1517 | FALSE | 5 |
| N | 405 | 1716 | 1554 | FALSE | 0 |
| N | 406 | 1480 | 1545 | FALSE | 5 |
| N | 407 | 1480 | 1703 | FALSE | 3 |
| N | 408 | 1215 | 1703 | FALSE | 1 |
| N | 409 | 1215 | 1703 | FALSE | 3 |
| N | 410 | 1215 | 1778 | FALSE | 3 |
| N | 411 | 0 | 1447 | FALSE | 1 |

### Table S10

| Labeling of mutation (ORF:Old AA/Residue/New AA) | SARS-CoV-2 Lineage | ORF | Is in epitope site (HDPA) | Is in cross-reactive epitope site (HDPA) | ORF protein site (AA) | Is in differential epitope site (HDPA) |
|---|---|---|---|---|---|---|
| N:D3L | Alpha (B.1.1.7) | N | TRUE | FALSE | 3 | FALSE |
| N:R203K | Alpha (B.1.1.7) | N | TRUE | FALSE | 203 | FALSE |
| N:G204R | Alpha (B.1.1.7) | N | TRUE | FALSE | 204 | FALSE |
| N:S235F | Alpha (B.1.1.7) | N | TRUE | FALSE | 235 | FALSE |
| ORF1a:T1001I | Alpha (B.1.1.7) | ORF1A | TRUE | FALSE | 1001 | FALSE |
| ORF1a:A1708D | Alpha (B.1.1.7) | ORF1A | TRUE | FALSE | 1708 | FALSE |
| ORF1a:I2230T | Alpha (B.1.1.7) | ORF1A | TRUE | FALSE | 2230 | FALSE |
| ORF1b:P314L | Alpha (B.1.1.7) | ORF1B | FALSE | FALSE | 314 | FALSE |
| ORF8:R52I | Alpha (B.1.1.7) | ORF8 | FALSE | FALSE | 52 | FALSE |
| ORF8:Y73C | Alpha (B.1.1.7) | ORF8 | FALSE | FALSE | 73 | FALSE |
| S:N501Y | Alpha (B.1.1.7) | S | TRUE | FALSE | 501 | FALSE |
| S:A570D | Alpha (B.1.1.7) | S | TRUE | FALSE | 570 | TRUE |
| S:D614G | Alpha (B.1.1.7) | S | TRUE | FALSE | 614 | FALSE |
| S:P681H | Alpha (B.1.1.7) | S | TRUE | FALSE | 681 | FALSE |
| S:T716I | Alpha (B.1.1.7) | S | TRUE | FALSE | 716 | FALSE |
| S:S982A | Alpha (B.1.1.7) | S | TRUE | TRUE | 982 | FALSE |
| S:D1118H | Alpha (B.1.1.7) | S | TRUE | FALSE | 1118 | TRUE |
| E:P71L | Beta (B.1.351) | E | FALSE | FALSE | 71 | FALSE |
| N:T205I | Beta (B.1.351) | N | TRUE | FALSE | 205 | FALSE |
| ORF1a:T265I | Beta (B.1.351) | ORF1A | TRUE | FALSE | 265 | FALSE |
| ORF1a:K1655N | Beta (B.1.351) | ORF1A | FALSE | FALSE | 1655 | FALSE |
| ORF1a:K3353R | Beta (B.1.351) | ORF1A | TRUE | FALSE | 3353 | FALSE |
| ORF3a:Q57H | Beta (B.1.351) | ORF3A | FALSE | FALSE | 57 | FALSE |
| ORF3a:S171L | Beta (B.1.351) | ORF3A | FALSE | FALSE | 171 | FALSE |
| S:D80A | Beta (B.1.351) | S | TRUE | FALSE | 80 | TRUE |
| S:D215G | Beta (B.1.351) | S | TRUE | FALSE | 215 | FALSE |
| S:L242H | Beta (B.1.351) | S | TRUE | FALSE | 242 | FALSE |
| S:K417N | Beta (B.1.351) | S | TRUE | FALSE | 417 | TRUE |
| S:D614G | Beta (B.1.351) | S | TRUE | FALSE | 614 | FALSE |
| S:A701V | Beta (B.1.351) | S | TRUE | FALSE | 701 | TRUE |
| N:P80R | Gamma (P.1) | N | TRUE | FALSE | 80 | FALSE |
| N:R203K | Gamma (P.1) | N | TRUE | FALSE | 203 | FALSE |
| N:G204R | Gamma (P.1) | N | TRUE | FALSE | 204 | FALSE |
| ORF1a:S1188L | Gamma (P.1) | ORF1A | FALSE | FALSE | 1188 | FALSE |
| ORF1a:K1795Q | Gamma (P.1) | ORF1A | TRUE | FALSE | 1795 | FALSE |
| ORF1b:P314L | Gamma (P.1) | ORF1B | FALSE | FALSE | 314 | FALSE |
| ORF1b:E1264D | Gamma (P.1) | ORF1B | TRUE | TRUE | 1264 | FALSE |
| ORF3a:S253P | Gamma (P.1) | ORF3A | FALSE | FALSE | 253 | FALSE |
| ORF8:E92K | Gamma (P.1) | ORF8 | FALSE | FALSE | 92 | FALSE |
| S:L18F | Gamma (P.1) | S | TRUE | FALSE | 18 | FALSE |
| S:T20N | Gamma (P.1) | S | TRUE | FALSE | 20 | FALSE |
| S:P26S | Gamma (P.1) | S | TRUE | FALSE | 26 | FALSE |
| S:D138Y | Gamma (P.1) | S | TRUE | FALSE | 138 | TRUE |
| S:R190S | Gamma (P.1) | S | TRUE | FALSE | 190 | FALSE |
| S:K417T | Gamma (P.1) | S | TRUE | FALSE | 417 | TRUE |
| S:E484K | Gamma (P.1) | S | TRUE | FALSE | 484 | FALSE |
| S:N501Y | Gamma (P.1) | S | TRUE | FALSE | 501 | FALSE |
| S:D614G | Gamma (P.1) | S | TRUE | FALSE | 614 | FALSE |
| S:H655Y | Gamma (P.1) | S | TRUE | FALSE | 655 | FALSE |
| S:T1027I | Gamma (P.1) | S | TRUE | TRUE | 1027 | FALSE |

| | | | | | | |
|---|---|---|---|---|---|---|
| S:V1176F | Gamma (P.1) | S | TRUE | FALSE | 1176 | FALSE |
| N:T205I | Epsilon (B.1.427+B.1.429) | N | TRUE | FALSE | 205 | FALSE |
| ORF1a:T265I | Epsilon (B.1.427+B.1.429) | ORF1A | TRUE | FALSE | 265 | FALSE |
| ORF1a:I4205V | Epsilon (B.1.427+B.1.429) | ORF1A | TRUE | TRUE | 4205 | FALSE |
| ORF1a:S3158T | Epsilon (B.1.427+B.1.429) | ORF1A | FALSE | FALSE | 3158 | FALSE |
| ORF1b:P314L | Epsilon (B.1.427+B.1.429) | ORF1B | FALSE | FALSE | 314 | FALSE |
| ORF1b:P976L | Epsilon (B.1.427+B.1.429) | ORF1B | TRUE | TRUE | 976 | FALSE |
| ORF1b:D1183Y | Epsilon (B.1.427+B.1.429) | ORF1B | TRUE | TRUE | 1183 | FALSE |
| ORF3a:Q57H | Epsilon (B.1.427+B.1.429) | ORF3A | FALSE | FALSE | 57 | FALSE |
| S:S13I | Epsilon (B.1.427+B.1.429) | S | TRUE | FALSE | 13 | FALSE |
| S:W152C | Epsilon (B.1.427+B.1.429) | S | TRUE | FALSE | 152 | TRUE |
| S:L452R | Epsilon (B.1.427+B.1.429) | S | TRUE | FALSE | 452 | FALSE |
| S:D614G | Epsilon (B.1.427+B.1.429) | S | TRUE | FALSE | 614 | FALSE |
| M:I82T | Kappa+Delta (B.1.617.X) | M | TRUE | FALSE | 82 | FALSE |
| N:D63G | Kappa+Delta (B.1.617.X) | N | TRUE | FALSE | 63 | FALSE |
| N:R203M | Kappa+Delta (B.1.617.X) | N | TRUE | FALSE | 203 | FALSE |
| N:D377Y | Kappa+Delta (B.1.617.X) | N | TRUE | FALSE | 377 | TRUE |
| ORF1b:P314L | Kappa+Delta (B.1.617.X) | ORF1B | FALSE | FALSE | 314 | FALSE |
| ORF3a:S26L | Kappa+Delta (B.1.617.X) | ORF3A | FALSE | FALSE | 26 | FALSE |
| ORF7a:V82A | Kappa+Delta (B.1.617.X) | ORF7A | FALSE | FALSE | 82 | FALSE |
| S:T19R | Kappa+Delta (B.1.617.X) | S | TRUE | FALSE | 19 | FALSE |
| S:L452R | Kappa+Delta (B.1.617.X) | S | TRUE | FALSE | 452 | FALSE |
| S:T478K | Kappa+Delta (B.1.617.X) | S | TRUE | FALSE | 478 | TRUE |
| S:D614G | Kappa+Delta (B.1.617.X) | S | TRUE | FALSE | 614 | FALSE |
| S:P681R | Kappa+Delta (B.1.617.X) | S | TRUE | FALSE | 681 | FALSE |
| S:D950N | Kappa+Delta (B.1.617.X) | S | TRUE | FALSE | 950 | FALSE |
| M:I82T | Eta (B.1.525) | M | TRUE | FALSE | 82 | FALSE |
| N:S2Y | Eta (B.1.525) | N | TRUE | FALSE | 2 | FALSE |
| N:A12G | Eta (B.1.525) | N | TRUE | FALSE | 12 | FALSE |
| N:T205I | Eta (B.1.525) | N | TRUE | FALSE | 205 | FALSE |
| ORF1a:T2007I | Eta (B.1.525) | ORF1A | TRUE | FALSE | 2007 | FALSE |
| ORF1b:P314F | Eta (B.1.525) | ORF1B | FALSE | FALSE | 314 | FALSE |
| S:A67V | Eta (B.1.525) | S | FALSE | FALSE | 67 | FALSE |
| S:E484K | Eta (B.1.525) | S | TRUE | FALSE | 484 | FALSE |
| S:D614G | Eta (B.1.525) | S | TRUE | FALSE | 614 | FALSE |
| S:Q677H | Eta (B.1.525) | S | TRUE | FALSE | 677 | FALSE |
| S:F888L | Eta (B.1.525) | S | TRUE | FALSE | 888 | FALSE |
| ORF1a:T265I | Iota (B.1.526) | ORF1A | TRUE | FALSE | 265 | FALSE |
| ORF1a:L3201P | Iota (B.1.526) | ORF1A | TRUE | FALSE | 3201 | FALSE |
| ORF1b:P314L | Iota (B.1.526) | ORF1B | FALSE | FALSE | 314 | FALSE |
| ORF3a:P42L | Iota (B.1.526) | ORF3A | FALSE | FALSE | 42 | FALSE |
| ORF3a:Q57H | Iota (B.1.526) | ORF3A | FALSE | FALSE | 57 | FALSE |
| ORF8:T11I | Iota (B.1.526) | ORF8 | FALSE | FALSE | 11 | FALSE |
| S:D614G | Iota (B.1.526) | S | TRUE | FALSE | 614 | FALSE |
| N:P13L | Lambda (C.37) | N | TRUE | FALSE | 13 | FALSE |
| N:R203K | Lambda (C.37) | N | TRUE | FALSE | 203 | FALSE |
| N:G204R | Lambda (C.37) | N | TRUE | FALSE | 204 | FALSE |
| N:G214C | Lambda (C.37) | N | TRUE | FALSE | 214 | FALSE |
| ORF1a:T1246I | Lambda (C.37) | ORF1A | TRUE | FALSE | 1246 | FALSE |
| ORF1a:P2287S | Lambda (C.37) | ORF1A | TRUE | FALSE | 2287 | FALSE |
| ORF1a:F2387V | Lambda (C.37) | ORF1A | TRUE | FALSE | 2387 | FALSE |

| | | | | | | |
|---|---|---|---|---|---|---|
| ORF1a:L3201P | Lambda (C.37) | ORF1A | TRUE | FALSE | 3201 | FALSE |
| ORF1a:T3255I | Lambda (C.37) | ORF1A | TRUE | TRUE | 3255 | FALSE |
| ORF1a:G3278S | Lambda (C.37) | ORF1A | TRUE | TRUE | 3278 | FALSE |
| ORF1b:P314L | Lambda (C.37) | ORF1B | FALSE | FALSE | 314 | FALSE |
| S:G75V | Lambda (C.37) | S | TRUE | FALSE | 75 | FALSE |
| S:T76I | Lambda (C.37) | S | TRUE | FALSE | 76 | FALSE |
| S:L452Q | Lambda (C.37) | S | TRUE | FALSE | 452 | FALSE |
| S:F490S | Lambda (C.37) | S | TRUE | FALSE | 490 | FALSE |
| S:D614G | Lambda (C.37) | S | TRUE | FALSE | 614 | FALSE |
| S:T859N | Lambda (C.37) | S | TRUE | TRUE | 859 | FALSE |
| S:Y505H | Omicron (B.1.1.529) | S | TRUE | FALSE | 505 | FALSE |
| N:G204R | Omicron (B.1.1.529) | N | TRUE | FALSE | 204 | FALSE |
| N:R203K | Omicron (B.1.1.529) | N | TRUE | FALSE | 203 | FALSE |
| ORF1a:A2710T | Omicron (B.1.1.529) | ORF1A | TRUE | FALSE | 2710 | FALSE |
| ORF1a:I3758V | Omicron (B.1.1.529) | ORF1A | TRUE | FALSE | 3758 | FALSE |
| ORF1a:K856R | Omicron (B.1.1.529) | ORF1A | TRUE | FALSE | 856 | FALSE |
| ORF1a:T3255I | Omicron (B.1.1.529) | ORF1A | TRUE | TRUE | 3255 | FALSE |
| ORF1b:I1566V | Omicron (B.1.1.529) | ORF1B | TRUE | FALSE | 1566 | FALSE |
| ORF1b:P314L | Omicron (B.1.1.529) | ORF1B | FALSE | FALSE | 314 | FALSE |
| S:D614G | Omicron (B.1.1.529) | S | TRUE | FALSE | 614 | FALSE |
| S:E484A | Omicron (B.1.1.529) | S | TRUE | FALSE | 484 | FALSE |
| S:G339D | Omicron (B.1.1.529) | S | TRUE | FALSE | 339 | FALSE |
| S:G446S | Omicron (B.1.1.529) | S | TRUE | FALSE | 446 | FALSE |
| S:G496S | Omicron (B.1.1.529) | S | TRUE | FALSE | 496 | FALSE |
| S:H655Y | Omicron (B.1.1.529) | S | TRUE | FALSE | 655 | FALSE |
| S:K417N | Omicron (B.1.1.529) | S | TRUE | FALSE | 417 | TRUE |
| S:N440K | Omicron (B.1.1.529) | S | TRUE | FALSE | 440 | FALSE |
| S:N501Y | Omicron (B.1.1.529) | S | TRUE | FALSE | 501 | FALSE |
| S:N679K | Omicron (B.1.1.529) | S | TRUE | FALSE | 679 | FALSE |
| S:P681H | Omicron (B.1.1.529) | S | TRUE | FALSE | 681 | FALSE |
| S:Q493R | Omicron (B.1.1.529) | S | TRUE | FALSE | 493 | FALSE |
| S:Q498R | Omicron (B.1.1.529) | S | TRUE | FALSE | 498 | FALSE |
| S:S371L | Omicron (B.1.1.529) | S | TRUE | FALSE | 371 | FALSE |
| S:S373P | Omicron (B.1.1.529) | S | TRUE | FALSE | 373 | FALSE |
| S:S375F | Omicron (B.1.1.529) | S | TRUE | FALSE | 375 | FALSE |
| S:S477N | Omicron (B.1.1.529) | S | TRUE | FALSE | 477 | TRUE |
| S:T478K | Omicron (B.1.1.529) | S | TRUE | FALSE | 478 | TRUE |
| S:T547K | Omicron (B.1.1.529) | S | TRUE | FALSE | 547 | FALSE |
| ORF1a:L2084I | Omicron (B.1.1.529) | ORF1A | TRUE | FALSE | 2084 | FALSE |
| S:A67V | Omicron (B.1.1.529) | S | FALSE | FALSE | 67 | FALSE |
| S:Y145D | Omicron (B.1.1.529) | S | TRUE | FALSE | 145 | TRUE |
| E:T9I | Omicron (B.1.1.529) | E | TRUE | FALSE | 9 | FALSE |
| M:A63T | Omicron (B.1.1.529) | M | FALSE | FALSE | 63 | FALSE |
| M:D3G | Omicron (B.1.1.529) | M | TRUE | FALSE | 3 | FALSE |
| M:Q19E | Omicron (B.1.1.529) | M | TRUE | FALSE | 19 | TRUE |
| S:L212I | Omicron (B.1.1.529) | S | TRUE | FALSE | 212 | FALSE |
| S:T95I | Omicron (B.1.1.529) | S | TRUE | FALSE | 95 | FALSE |
| ORF1a:P3395H | Omicron (B.1.1.529) | ORF1A | FALSE | FALSE | 3395 | FALSE |
| N:R203K | Theta (P.3) | N | TRUE | FALSE | 203 | FALSE |
| N:G204R | Theta (P.3) | N | TRUE | FALSE | 204 | FALSE |
| ORF1a:D1554G | Theta (P.3) | ORF1A | TRUE | FALSE | 1554 | FALSE |

| | | | | | | |
|---|---|---|---|---|---|---|
| ORF1a:S2625F | Theta (P.3) | ORF1A | TRUE | FALSE | 2625 | FALSE |
| ORF1a:D2980N | Theta (P.3) | ORF1A | TRUE | TRUE | 2980 | FALSE |
| ORF1a:L3201P | Theta (P.3) | ORF1A | TRUE | FALSE | 3201 | FALSE |
| ORF1a:D3681E | Theta (P.3) | ORF1A | FALSE | FALSE | 3681 | FALSE |
| ORF1a:L3930F | Theta (P.3) | ORF1A | TRUE | FALSE | 3930 | FALSE |
| ORF1b:P314L | Theta (P.3) | ORF1B | FALSE | FALSE | 314 | FALSE |
| ORF1b:A1291V | Theta (P.3) | ORF1B | TRUE | TRUE | 1291 | FALSE |
| ORF8:K2Q | Theta (P.3) | ORF8 | FALSE | FALSE | 2 | FALSE |
| S:E484K | Theta (P.3) | S | TRUE | FALSE | 484 | FALSE |
| S:N501Y | Theta (P.3) | S | TRUE | FALSE | 501 | FALSE |
| S:D614G | Theta (P.3) | S | TRUE | FALSE | 614 | FALSE |
| S:P681H | Theta (P.3) | S | TRUE | FALSE | 681 | FALSE |
| S:E1092K | Theta (P.3) | S | TRUE | FALSE | 1092 | FALSE |
| S:H1101Y | Theta (P.3) | S | TRUE | FALSE | 1101 | FALSE |
| S:V1176F | Theta (P.3) | S | TRUE | FALSE | 1176 | FALSE |
| N:A119S | Zeta (P.2) | N | TRUE | FALSE | 119 | TRUE |
| N:R203K | Zeta (P.2) | N | TRUE | FALSE | 203 | FALSE |
| N:G204R | Zeta (P.2) | N | TRUE | FALSE | 204 | FALSE |
| N:M234I | Zeta (P.2) | N | TRUE | FALSE | 234 | FALSE |
| ORF1a:L3468V | Zeta (P.2) | ORF1A | TRUE | FALSE | 3468 | FALSE |
| ORF1a:L3930F | Zeta (P.2) | ORF1A | TRUE | FALSE | 3930 | FALSE |
| ORF1b:P314L | Zeta (P.2) | ORF1B | FALSE | FALSE | 314 | FALSE |
| S:E484K | Zeta (P.2) | S | TRUE | FALSE | 484 | FALSE |
| S:D614G | Zeta (P.2) | S | TRUE | FALSE | 614 | FALSE |
| S:V1176F | Zeta (P.2) | S | TRUE | FALSE | 1176 | FALSE |
| N:S197L | A.2.5 | N | TRUE | FALSE | 197 | FALSE |
| N:M234I | A.2.5 | N | TRUE | FALSE | 234 | FALSE |
| N:P365S | A.2.5 | N | TRUE | FALSE | 365 | TRUE |
| N:P383L | A.2.5 | N | TRUE | FALSE | 383 | TRUE |
| ORF1a:L4F | A.2.5 | ORF1A | TRUE | FALSE | 4 | FALSE |
| ORF1a:K1657E | A.2.5 | ORF1A | FALSE | FALSE | 1657 | FALSE |
| ORF1a:F3071Y | A.2.5 | ORF1A | FALSE | FALSE | 3071 | FALSE |
| ORF1a:T3255I | A.2.5 | ORF1A | TRUE | TRUE | 3255 | FALSE |
| ORF1a:H3580Q | A.2.5 | ORF1A | FALSE | FALSE | 3580 | FALSE |
| ORF1b:P1000L | A.2.5 | ORF1B | FALSE | FALSE | 1000 | FALSE |
| ORF3a:S74F | A.2.5 | ORF3A | FALSE | FALSE | 74 | FALSE |
| ORF3a:G196V | A.2.5 | ORF3A | FALSE | FALSE | 196 | FALSE |
| ORF8:L84S | A.2.5 | ORF8 | FALSE | FALSE | 84 | FALSE |
| S:L452R | A.2.5 | S | TRUE | FALSE | 452 | FALSE |
| S:D614G | A.2.5 | S | TRUE | FALSE | 614 | FALSE |
| M:I82T | B.1.1.318 | M | TRUE | FALSE | 82 | FALSE |
| N:R203K | B.1.1.318 | N | TRUE | FALSE | 203 | FALSE |
| N:G204R | B.1.1.318 | N | TRUE | FALSE | 204 | FALSE |
| N:A208G | B.1.1.318 | N | TRUE | FALSE | 208 | FALSE |
| ORF1a:E1196V | B.1.1.318 | ORF1A | TRUE | FALSE | 1196 | FALSE |
| ORF1a:K2511N | B.1.1.318 | ORF1A | FALSE | FALSE | 2511 | FALSE |
| ORF1a:T2936I | B.1.1.318 | ORF1A | FALSE | FALSE | 2936 | FALSE |
| ORF1a:A3209V | B.1.1.318 | ORF1A | FALSE | FALSE | 3209 | FALSE |
| ORF1a:T3284I | B.1.1.318 | ORF1A | TRUE | TRUE | 3284 | FALSE |
| ORF1b:P314L | B.1.1.318 | ORF1B | FALSE | FALSE | 314 | FALSE |
| ORF1b:V2371M | B.1.1.318 | ORF1B | FALSE | FALSE | 2371 | FALSE |

| | | | | | | |
|---|---|---|---|---|---|---|
| S:T95I | B.1.1.318 | S | TRUE | FALSE | 95 | FALSE |
| S:E484K | B.1.1.318 | S | TRUE | FALSE | 484 | FALSE |
| S:D614G | B.1.1.318 | S | TRUE | FALSE | 614 | FALSE |
| S:P681H | B.1.1.318 | S | TRUE | FALSE | 681 | FALSE |
| S:D796H | B.1.1.318 | S | FALSE | FALSE | 796 | FALSE |
| N:R203K | B.1.1.519 | N | TRUE | FALSE | 203 | FALSE |
| N:G204R | B.1.1.519 | N | TRUE | FALSE | 204 | FALSE |
| ORF1a:P959S | B.1.1.519 | ORF1A | TRUE | FALSE | 959 | FALSE |
| ORF1a:T3255I | B.1.1.519 | ORF1A | TRUE | TRUE | 3255 | FALSE |
| ORF1a:I3618V | B.1.1.519 | ORF1A | TRUE | FALSE | 3618 | FALSE |
| ORF1a:T4175I | B.1.1.519 | ORF1A | TRUE | FALSE | 4175 | FALSE |
| ORF1b:P314L | B.1.1.519 | ORF1B | FALSE | FALSE | 314 | FALSE |
| S:T478K | B.1.1.519 | S | TRUE | FALSE | 478 | TRUE |
| S:D614G | B.1.1.519 | S | TRUE | FALSE | 614 | FALSE |
| S:P681H | B.1.1.519 | S | TRUE | FALSE | 681 | FALSE |
| S:T732A | B.1.1.519 | S | TRUE | TRUE | 732 | FALSE |
| N:T205I | B.1.466.2 | N | TRUE | FALSE | 205 | FALSE |
| ORF1a:T1168I | B.1.466.2 | ORF1A | TRUE | FALSE | 1168 | FALSE |
| ORF1a:P1640L | B.1.466.2 | ORF1A | TRUE | FALSE | 1640 | FALSE |
| ORF1b:P314L | B.1.466.2 | ORF1B | FALSE | FALSE | 314 | FALSE |
| ORF3a:Q57H | B.1.466.2 | ORF3A | FALSE | FALSE | 57 | FALSE |
| S:N439K | B.1.466.2 | S | TRUE | FALSE | 439 | FALSE |
| S:D614G | B.1.466.2 | S | TRUE | FALSE | 614 | FALSE |
| N:T205I | B.1.621 | N | TRUE | FALSE | 205 | FALSE |
| ORF1a:T1055A | B.1.621 | ORF1A | TRUE | FALSE | 1055 | FALSE |
| ORF1a:T1538I | B.1.621 | ORF1A | TRUE | FALSE | 1538 | FALSE |
| ORF1a:T3255I | B.1.621 | ORF1A | TRUE | TRUE | 3255 | FALSE |
| ORF1a:Q3729R | B.1.621 | ORF1A | FALSE | FALSE | 3729 | FALSE |
| ORF1b:P314L | B.1.621 | ORF1B | FALSE | FALSE | 314 | FALSE |
| ORF1b:P1342S | B.1.621 | ORF1B | TRUE | TRUE | 1342 | FALSE |
| ORF3a:Q57H | B.1.621 | ORF3A | FALSE | FALSE | 57 | FALSE |
| ORF8:T11K | B.1.621 | ORF8 | FALSE | FALSE | 11 | FALSE |
| ORF8:P38S | B.1.621 | ORF8 | FALSE | FALSE | 38 | FALSE |
| S:T95I | B.1.621 | S | TRUE | FALSE | 95 | FALSE |
| S:R346K | B.1.621 | S | TRUE | FALSE | 346 | TRUE |
| S:E484K | B.1.621 | S | TRUE | FALSE | 484 | FALSE |
| S:N501Y | B.1.621 | S | TRUE | FALSE | 501 | FALSE |
| S:D614G | B.1.621 | S | TRUE | FALSE | 614 | FALSE |
| S:P681H | B.1.621 | S | TRUE | FALSE | 681 | FALSE |
| S:D950N | B.1.621 | S | TRUE | FALSE | 950 | FALSE |
| N:T205I | B.1.214.2 | N | TRUE | FALSE | 205 | FALSE |
| ORF1a:I1398V | B.1.214.2 | ORF1A | TRUE | FALSE | 1398 | FALSE |
| ORF1a:T1881I | B.1.214.2 | ORF1A | TRUE | FALSE | 1881 | FALSE |
| ORF1a:A4016V | B.1.214.2 | ORF1A | TRUE | TRUE | 4016 | FALSE |
| ORF1b:P314L | B.1.214.2 | ORF1B | FALSE | FALSE | 314 | FALSE |
| S:Q414K | B.1.214.2 | S | TRUE | FALSE | 414 | TRUE |
| S:D614G | B.1.214.2 | S | TRUE | FALSE | 614 | FALSE |
| S:T716I | B.1.214.2 | S | TRUE | FALSE | 716 | FALSE |
| M:A63T | AV.1 | M | FALSE | FALSE | 63 | FALSE |
| M:H125Y | AV.1 | M | TRUE | TRUE | 125 | TRUE |
| N:I157V | AV.1 | N | TRUE | FALSE | 157 | FALSE |

| | | | | | | |
|---|---|---|---|---|---|---|
| N:R203K | AV.1 | N | TRUE | FALSE | 203 | FALSE |
| N:G204R | AV.1 | N | TRUE | FALSE | 204 | FALSE |
| ORF1a:D75G | AV.1 | ORF1A | TRUE | FALSE | 75 | FALSE |
| ORF1a:G519S | AV.1 | ORF1A | FALSE | FALSE | 519 | FALSE |
| ORF1a:A591V | AV.1 | ORF1A | TRUE | FALSE | 591 | FALSE |
| ORF1a:H1160Y | AV.1 | ORF1A | TRUE | FALSE | 1160 | FALSE |
| ORF1a:P1640L | AV.1 | ORF1A | TRUE | FALSE | 1640 | FALSE |
| ORF1a:K1745I | AV.1 | ORF1A | TRUE | FALSE | 1745 | FALSE |
| ORF1a:N2405S | AV.1 | ORF1A | FALSE | FALSE | 2405 | FALSE |
| ORF1a:A3209V | AV.1 | ORF1A | FALSE | FALSE | 3209 | FALSE |
| ORF1b:P314L | AV.1 | ORF1B | FALSE | FALSE | 314 | FALSE |
| ORF1b:S425A | AV.1 | ORF1B | TRUE | TRUE | 425 | FALSE |
| ORF1b:T1404M | AV.1 | ORF1B | TRUE | TRUE | 1404 | FALSE |
| ORF1b:A1643V | AV.1 | ORF1B | TRUE | TRUE | 1643 | FALSE |
| S:D80G | AV.1 | S | TRUE | FALSE | 80 | TRUE |
| S:T95I | AV.1 | S | TRUE | FALSE | 95 | FALSE |
| S:G142D | AV.1 | S | TRUE | FALSE | 142 | TRUE |
| S:E484K | AV.1 | S | TRUE | FALSE | 484 | FALSE |
| S:D614G | AV.1 | S | TRUE | FALSE | 614 | FALSE |
| S:P681H | AV.1 | S | TRUE | FALSE | 681 | FALSE |
| S:I1130V | AV.1 | S | TRUE | FALSE | 1130 | TRUE |
| S:D1139H | AV.1 | S | TRUE | TRUE | 1139 | FALSE |
| M:L16I | AT.1 | M | TRUE | FALSE | 16 | FALSE |
| N:R203K | AT.1 | N | TRUE | FALSE | 203 | FALSE |
| N:G204R | AT.1 | N | TRUE | FALSE | 204 | FALSE |
| ORF1a:S376L | AT.1 | ORF1A | TRUE | FALSE | 376 | FALSE |
| ORF1a:V1006F | AT.1 | ORF1A | TRUE | FALSE | 1006 | FALSE |
| ORF1a:T2247N | AT.1 | ORF1A | TRUE | FALSE | 2247 | FALSE |
| ORF1a:T3255I | AT.1 | ORF1A | TRUE | TRUE | 3255 | FALSE |
| ORF1a:Q3729R | AT.1 | ORF1A | FALSE | FALSE | 3729 | FALSE |
| ORF1a:S4119T | AT.1 | ORF1A | TRUE | FALSE | 4119 | FALSE |
| ORF1b:P314L | AT.1 | ORF1B | FALSE | FALSE | 314 | FALSE |
| ORF1b:T1173N | AT.1 | ORF1B | TRUE | FALSE | 1173 | FALSE |
| ORF1b:V1905L | AT.1 | ORF1B | TRUE | TRUE | 1905 | FALSE |
| ORF1b:A2431V | AT.1 | ORF1B | TRUE | TRUE | 2431 | FALSE |
| ORF3a:L95M | AT.1 | ORF3A | FALSE | FALSE | 95 | FALSE |
| S:P9L | AT.1 | S | TRUE | FALSE | 9 | FALSE |
| S:C136Y | AT.1 | S | TRUE | FALSE | 136 | FALSE |
| S:D215G | AT.1 | S | TRUE | FALSE | 215 | FALSE |
| S:H245P | AT.1 | S | TRUE | FALSE | 245 | FALSE |
| S:E484K | AT.1 | S | TRUE | FALSE | 484 | FALSE |
| S:D614G | AT.1 | S | TRUE | FALSE | 614 | FALSE |
| S:E780K | AT.1 | S | TRUE | FALSE | 780 | FALSE |
| M:I82T | C.36.3 | M | TRUE | FALSE | 82 | FALSE |
| N:R203K | C.36.3 | N | TRUE | FALSE | 203 | FALSE |
| N:G204R | C.36.3 | N | TRUE | FALSE | 204 | FALSE |
| N:G212V | C.36.3 | N | TRUE | FALSE | 212 | FALSE |
| ORF1a:E102K | C.36.3 | ORF1A | TRUE | FALSE | 102 | FALSE |
| ORF1a:A859V | C.36.3 | ORF1A | TRUE | FALSE | 859 | FALSE |
| ORF1a:T1246I | C.36.3 | ORF1A | TRUE | FALSE | 1246 | FALSE |
| ORF1a:D1639N | C.36.3 | ORF1A | TRUE | FALSE | 1639 | FALSE |

| | | | | | | |
|---|---|---|---|---|---|---|
| ORF1a:P2287S | C.36.3 | ORF1A | TRUE | FALSE | 2287 | FALSE |
| ORF1a:D2980N | C.36.3 | ORF1A | TRUE | TRUE | 2980 | FALSE |
| ORF1a:D3222N | C.36.3 | ORF1A | TRUE | TRUE | 3222 | FALSE |
| ORF1a:G3278S | C.36.3 | ORF1A | TRUE | TRUE | 3278 | FALSE |
| ORF1a:S3687L | C.36.3 | ORF1A | FALSE | FALSE | 3687 | FALSE |
| ORF1a:L3691S | C.36.3 | ORF1A | FALSE | FALSE | 3691 | FALSE |
| ORF1a:T4090I | C.36.3 | ORF1A | TRUE | FALSE | 4090 | FALSE |
| ORF1b:P314L | C.36.3 | ORF1B | FALSE | FALSE | 314 | FALSE |
| ORF1b:D1028Y | C.36.3 | ORF1B | TRUE | TRUE | 1028 | FALSE |
| ORF7b:A43S | C.36.3 | ORF7B | FALSE | FALSE | 43 | FALSE |
| S:S12F | C.36.3 | S | TRUE | FALSE | 12 | FALSE |
| S:W152R | C.36.3 | S | TRUE | FALSE | 152 | TRUE |
| S:R346S | C.36.3 | S | TRUE | FALSE | 346 | TRUE |
| S:L452R | C.36.3 | S | TRUE | FALSE | 452 | FALSE |
| S:D614G | C.36.3 | S | TRUE | FALSE | 614 | FALSE |
| S:Q677H | C.36.3 | S | TRUE | FALSE | 677 | FALSE |
| S:A899S | C.36.3 | S | TRUE | TRUE | 899 | FALSE |
| M:F28L | R.1 | M | TRUE | FALSE | 28 | TRUE |
| N:S187L | R.1 | N | TRUE | FALSE | 187 | FALSE |
| N:R203K | R.1 | N | TRUE | FALSE | 203 | FALSE |
| N:G204R | R.1 | N | TRUE | FALSE | 204 | FALSE |
| N:Q418H | R.1 | N | FALSE | FALSE | 418 | FALSE |
| ORF1b:P314L | R.1 | ORF1B | FALSE | FALSE | 314 | FALSE |
| ORF1b:G1362R | R.1 | ORF1B | TRUE | TRUE | 1362 | FALSE |
| ORF1b:P1936H | R.1 | ORF1B | FALSE | FALSE | 1936 | FALSE |
| S:W152L | R.1 | S | TRUE | FALSE | 152 | TRUE |
| S:E484K | R.1 | S | TRUE | FALSE | 484 | FALSE |
| S:D614G | R.1 | S | TRUE | FALSE | 614 | FALSE |
| S:G769V | R.1 | S | TRUE | FALSE | 769 | FALSE |
| N:R203K | R.2 | N | TRUE | FALSE | 203 | FALSE |
| N:G204R | R.2 | N | TRUE | FALSE | 204 | FALSE |
| ORF1a:A1049V | R.2 | ORF1A | TRUE | FALSE | 1049 | FALSE |
| ORF1a:K1202N | R.2 | ORF1A | TRUE | FALSE | 1202 | FALSE |
| ORF1a:Q1592R | R.2 | ORF1A | TRUE | FALSE | 1592 | FALSE |
| ORF1a:D4085G | R.2 | ORF1A | TRUE | FALSE | 4085 | FALSE |
| ORF1b:P314L | R.2 | ORF1B | FALSE | FALSE | 314 | FALSE |
| S:D614G | R.2 | S | TRUE | FALSE | 614 | FALSE |
| S:Q677H | R.2 | S | TRUE | FALSE | 677 | FALSE |
| S:T732S | R.2 | S | TRUE | TRUE | 732 | FALSE |
| ORF9c:G50R | Alpha (B.1.1.7) | ORF9c | TRUE | FALSE | 50 | FALSE |
| ORF9c:L52F | Beta (B.1.351) | ORF9c | TRUE | FALSE | 52 | FALSE |
| ORF9c:G50R | Gamma (P.1) | ORF9c | TRUE | FALSE | 50 | FALSE |
| ORF9c:L52F | Epsilon (B.1.427+B.1.429) | ORF9c | TRUE | FALSE | 52 | FALSE |
| ORF9c:G50W | Kappa+Delta (B.1.617.X) | ORF9c | TRUE | FALSE | 50 | FALSE |
| ORF9c:L52F | Eta (B.1.525) | ORF9c | TRUE | FALSE | 52 | FALSE |
| ORF9c:G50R | Lambda (C.37) | ORF9c | TRUE | FALSE | 50 | FALSE |
| ORF9c:M60I | Lambda (C.37) | ORF9c | TRUE | FALSE | 60 | FALSE |
| ORF9c:G50R | Omicron (B.1.1.529) | ORF9c | TRUE | FALSE | 50 | FALSE |
| ORF9c:G50R | Theta (P.3) | ORF9c | TRUE | FALSE | 50 | FALSE |
| ORF9c:G50R | Zeta (P.2) | ORF9c | TRUE | FALSE | 50 | FALSE |
| ORF9c:Q44Stop | A.2.5 | ORF9c | TRUE | FALSE | 44Sto | FALSE |

| | | | | | | |
|---|---|---|---|---|---|---|
| ORF9c:G50R | B.1.1.318 | ORF9c | TRUE | FALSE | 50 | FALSE |
| ORF9c:L55V | B.1.1.318 | ORF9c | TRUE | FALSE | 55 | FALSE |
| ORF9c:G50R | B.1.1.519 | ORF9c | TRUE | FALSE | 50 | FALSE |
| ORF9c:L52F | B.1.466.2 | ORF9c | TRUE | FALSE | 52 | FALSE |
| ORF9c:L52F | B.1.621 | ORF9c | TRUE | FALSE | 52 | FALSE |
| ORF9c:L52F | B.1.214.2 | ORF9c | TRUE | FALSE | 52 | FALSE |
| ORF9c:G50R | AV.1 | ORF9c | TRUE | FALSE | 50 | FALSE |
| ORF9c:G50R | AT.1 | ORF9c | TRUE | FALSE | 50 | FALSE |
| ORF9c:G50R | C.36.3 | ORF9c | TRUE | FALSE | 50 | FALSE |
| ORF9c:A59S | C.36.3 | ORF9c | TRUE | FALSE | 59 | FALSE |
| ORF9c:H34Y | R.1 | ORF9c | TRUE | FALSE | 34 | FALSE |
| ORF9c:G50R | R.1 | ORF9c | TRUE | FALSE | 50 | FALSE |
| ORF9c:G50R | R.2 | ORF9c | TRUE | FALSE | 50 | FALSE |

| Name | Genus | Subgenus | Taxonomy ID | ORF1AB | Spike | Envelope | Membrane | Nucleocapsid |
|---|---|---|---|---|---|---|---|---|
| SARS-CoV-2 | Betacoronavirus | Sarbecovirus | NCBI:txid2697049 | YP_009724389.1 | YP_009724390.1 | YP_009724392.1 | YP_009724393.1 | YP_009724397.2 |
| 229E | Alphacoronavirus | Duvinacovirus | NCBI:txid11137 | ARU07599.1 | ARU07601.1 | ARU07603.1 | ARU07604.1 | ARU07605.1 |
| NL63 | Alphacoronavirus | Setracovirus | NCBI:txid277944 | YP_003766.2 | YP_003767.1 | YP_003769.1 | YP_003770.1 | YP_003771.1 |
| OC43 | Betacoronavirus | Embecovirus | NCBI:txid31631 | YP_009555238.1 | YP_009555241.1 | YP_009555243.1 | YP_009555244.1 | YP_009555245.1 |
| HKU1 | Betacoronavirus | Embecovirus | NCBI:txid290028 | YP_173236.1 | YP_173238.1 | YP_173240.1 | YP_173241.1 | YP_173242.1 |

Table S12
